# PIME: a package for discovery of novel differences among microbial communities

**DOI:** 10.1101/632182

**Authors:** Luiz Fernando W. Roesch, Priscila Thiago Dobbler, Victor Satler Pylro, Bryan Kolaczkowski, Jennifer C. Drew, Eric W. Triplett

## Abstract

Massive sequencing of genetic markers, such as the 16S rRNA gene for prokaryotes, allows the comparative analysis of diversity and abundance of whole microbial communities. However, the data used for profiling microbial communities is usually low in signal and high in noise preventing the identification of real differences among treatments. PIME (Prevalence Interval for Microbiome Evaluation) fills this gap by removing those taxa that may be high in relative abundance in just a few samples but have a low prevalence overall. The reliability and robustness of PIME were compare against the existing methods and verified by a number of approaches using 16S rRNA independent datasets. To remove the noise, PIME filters microbial taxa not shared in a per treatment prevalence interval starting at 5% with increments of 5% at each filtering step. For each prevalence interval, hundreds of decision trees are calculated to predict the likelihood of detecting differences in treatments. The best prevalence-filtered dataset is user-selected by choosing the prevalence interval that keeps the majority of the 16S rRNA reads in the dataset and shows the lowest error rate. To obtain the likelihood of introducing bias while building prevalence-filtered datasets, an error detection step based in random permutations is also included. A reanalysis of previews published datasets with PIME uncovered previously missed microbial associations improving the ability to detect important organisms, which may be masked when only relative abundance is considered.

## INTRODUCTION

Sequencing of amplified genetic markers (metataxonomics), *e.g.* the 16S rRNA gene, is traditionally used for testing hypotheses based on microbial community composition. Taxonomic differences among treatments or outcomes in microbiome surveys have changed our understanding of the role played by microorganisms in the environment, plant and animal hosts, including humans. The major challenge for using this information is their interpretation for the discovery of the drivers of microbial diversity, the main taxa related to a given factor and the reduction in false discovery rates. Generally, as microbiome studies present a large number of taxa that are low in prevalence in many of the samples (1), these approaches frequently include a variety of pre-filtering steps. Those steps include, but are not limited, to the exclusion of sequences and/or taxonomic unities with low abundance, low variation, or low presence across all samples. Moreover, removing arguably uninformative information, pre-filtering is also advantageous because low abundance features in metataxonomic surveys might be also due to sequencing errors or low level of contaminants from commercial kits (2, 3).

Besides filtering low abundance sequences, a frequent approach involves the exclusion of microbial taxa under low prevalence across all samples. The prevalence of microbes in the human microbiome is characterized by variable distribution patterns (4) with prominent abundance of some strains in some subjects and nearly absence in others. While this unusual distribution might be focus of research for future experimental study (4), identify microbial taxonomic unities present in the majority of the subjects, also known as microbial core, has been one of the primary goals of the Human Microbiome Project (5, 6). The central objective of obtaining a healthy core microbiome is to use it to identify significant deviations from normality that might be associated with disease states, for example.

Many tools such as DADA2 (7), Phyloseq (8), Qiime (9), UPARSE (10), MG-RAST (11), mothur (12), MicrobiomeAnalyst (13) among others, have been developed to contrast experimental factors in microbiome studies. The choice of a given analyses package is usually based on the user’s level of experience in bioinformatics and on the available resources at the user’s host institution (14), but unfortunately, the most used approaches embedded in these packages rarely consider microbial prevalence.

Based on the microbial core concept, here we propose a new workflow designed to identify and remove the within group variation found in metataxonomic surveys (16S rRNA datasets) by capturing only biological differences at high sample prevalence levels. That means in an experiment comparing two treatments (e.g. health against diseased subjects) one core for each treatment will be calculated and relevant microbial taxa responsible for differences within microbial cores will be detected. To implement this concept, we developed an R package called PIME (Prevalence Interval for Microbiome Evaluation). PIME is a tool specifically designed to work with datasets presenting high variations among samples. It removes per group microbial taxa to keep only those taxa that are shared at some level of prevalence, using a machine learning algorithm. For each prevalence level a list with the most relevant taxa responsible for differences between or among groups is provided. To obtain the likelihood of introducing bias while building prevalence-filtered datasets, an error detection step based on randomizations is also included.

## PROGRAM DESCRIPTION AND METHODS

### Bioinformatics Workflow

The bioinformatics workflow described here is embedded into an R package called PIME (Prevalence Intervals for Microbiome Evaluation) available at: https://github.com/microEcology/PIME. PIME identifies statistically significant bacterial community differences taking into account the proportion of samples hosting a specific microbial community in a given time period. For the purpose of this work, prevalence was defined as the proportion of individuals in a specific group who share taxa, irrespective of the abundance, at the time of sampling. That is, a prevalence cutoff of 50% means that the taxa selected at this prevalence interval are found in 50% of subjects. PIME’s strategy is based on four fundamental steps depicted in Figure 1 and explained below:

**FIGURE 1.**
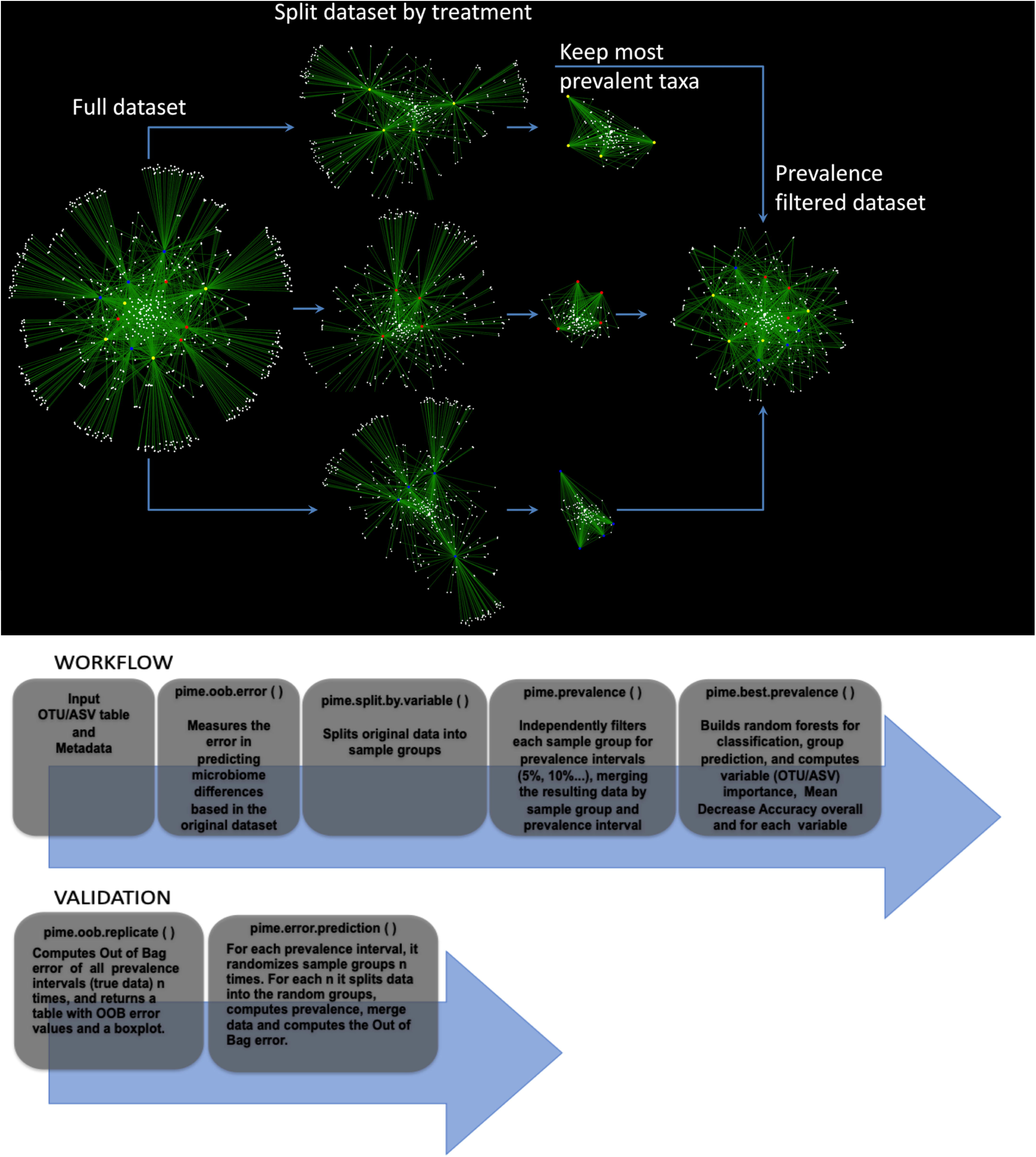
Empirical representation of steps used in PIME. Top panel. Simplified schema illustrating PIME method with a subset of 12 saliva’s microbiome samples. Each sample (red, yellow and blue circles) is connected to an ASV (white circles) through edges (green). ASVs observed in more than one sample are connected by at least two edges and are displayed at the center of the network. ASVs present in only one sample are connected by a single edge and are displayed at the border of the network. The first step applied by PIME is to split the full dataset according to the treatments defined by the user. Within this example red, yellow and blue circles depict three different treatments. At each of the three new groups the low prevalent ASVs are removed. Finally, the subsets are merged to compose a new filtered dataset used in the downstream analysis. Bottom panel. Step-by-step representation of PIME workflow and validation.

#### I) Prediction of differences on full dataset

PIME takes a phyloseq object (8) as input, builds hundreds of decision trees using a supervised machine learning algorithm and combines them into a single model to predict the likelihood of detecting any user predefined factor (e.g. difference between treatments) as source of sample variation (15). The model performance is indicated by the out-of-bag (OOB) estimate of the error rate calculated by training the algorithm on a subset of samples and tested on the remaining samples. Values can vary from 0 to 1, where zero indicates the model has 100% accuracy and 1 that the model has zero accuracy. This overall measurement of accuracy can be interpreted as an estimate of error obtained when the model is applied to new observations. Higher OOB error indicates low accuracy of the model in predicting differences among the variables tested. In this case, PIME might be used as an alternative to reduce noise by removing microbial taxa with low prevalence among samples. This might help to improve the model accuracy. This first step using the full dataset is implemented in a function called *pime.oob.error*. This function is run using the dataset without any filtering proposed by PIME. After obtaining the OOB error rate, the user should decide whether or not running PIME is adequate to the dataset. For instance, an OOB error close to zero indicates the prevalence filtering with PIME is not necessary, as the model accuracy is already reasonably good. On the other hand, if OOB error rate is greater than zero, filtering the dataset using PIME might improve the model accuracy. The user might then run the next function called *pime.split.by.variable*, which is described below.

#### II) Split the dataset by predictor variable and compute prevalence intervals

The full dataset is split according to treatments (or variables) defined by the user in the metadata file. PIME can deal with two or more variables. Each variable will be used to define data subsets. Those per variable subsets will be filtered using different prevalence levels from 5% up to 95% with increments of 5% for each level (see Figure 1 for a simplified schema illustrating this filtering step). Prevalence levels (usually high prevalence levels – e.g. 90%) where samples have zero counts are not calculated. After removal of taxa that did not match the prevalence criteria, the subsets are merged to compose a new filtered dataset (one per prevalence interval) used in the downstream analysis. This step is implemented in two functions called: *pime.split.by.variable* and *pime.prevalence*. The *pime.split.by.variable* function uses the original dataset as input and its output is used as input to the *pime.prevalence.* The function *pime.prevalence* keeps, for each treatment group, every OTU/ASV according to the following logical equation:

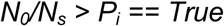

*Where: N*_*0*_ is the number of OTUs/ASVs with *Sum*>*0, Ns* is the number of samples and *P*_*i*_ is the prevalence interval *P*_i_=*0.05, …, Pmax.*

#### III) Computation of OOB error on each prevalence interval and importance of each taxa to differentiate microbial communities

At this step Random Forests analysis (15) is used to determine the level of prevalence that provides the best model to predict differences in the communities while still including as many taxa as possible in the analysis. The approach uses multiple learning algorithms to run classifications based on decision trees. After prevalence filtering, for each prevalence interval the OOB error rate, the number of remaining taxa and sequences is calculated. The results are provided in a table that can be used to decide the best prevalence interval that provides a classification model with reasonably good accuracy. This step is implemented in a function called: *pime.best.prevalence.* Within the same function, the contribution of each taxa to the mean decrease in classification accuracy is calculated. High values of mean decrease accuracy indicate the importance of taxa to differentiate two or more microbial communities. The user can access the importance of taxa in each of the prevalence interval calculated.

#### IV) Validation

To obtain the likelihood of introducing bias while building prevalence-filtered datasets, an error detection step is also included under the following rationale: Consider the scenario in which the null hypothesis of “no difference between groups” is false. If we randomly shuffle the labels that identify the sample groups and run the test again the expected outcome is that the randomized dataset will have a small chance to present distinct groups. Running the test multiple times with the random dataset would produce a high OOB error rate in most cases. This error detection test is implemented in two functions called pime.error.prediction and pime.oob.replicate. The first function randomizes the samples labels into arbitrary groupings using 100 random permutations. For each randomized prevalence filtered dataset, the OOB error rate is calculated to determine whether differences in the original groups occur by chance. The second function performs the Random Forest analyses and computes the OOB error for 100 replications in each prevalence interval without randomizing the sample labels. The biological difference among samples is expected to be greater than the differences generated randomly. Thus, the greatest fraction of randomizations should generate high error rates. On the other hand, no improvement in accuracy is expected within the randomized dataset.

### Empirical Validation

The PIME workflow was compared against other existing filtering methods and by using empirical tests with 16S rDNA datasets. The performance of PIME was compared against filtering methods based on overall prevalence, low abundance and low variance. Also, four 16S rDNA datasets were analyzed using PIME to illustrate its usefulness. These include an assessment of: a) the association between diet and saliva microbiome composition (unpublished original research); b) the gut microbiome in subjects at high genetic risk for type 1 diabetes (16); c) the vaginal microbiome in pregnant women randomized to receive milk with or without probiotic bacterial strains (17); and d) the saliva microbome compared the left antecubital fossa of healthy individuals (Human Microbiome Consortium, 2012).

The 16S rRNA gene sequences generated in this work have been deposited in NCBI’s Short Raw Archive and are accessible through BioProject ID PRJNA504439.

#### Comparison with other existing filtering methods

Comparisons were performed using a dataset composed by 16S rRNA sequences from microbes extracted from saliva of 125 undergraduate and graduate students from the University of Florida (accessible through BioProject ID PRJNA504439). The following filtering tests were performed: a) filtering the dataset by overall prevalence. To be kept, taxa must be present in at least 20% of the subjects; b) filtering the dataset by abundance. To be kept, taxa must have at least 5 sequences; c) filtering by low variance. To be kept taxa must have variance higher than 20%. Filtered datasets were compared against the prevalence interval of 65% as calculated by PIME as the best prevalence interval where the OOB error was zero. A record of this analysis containing a step-by-step R-code and results is provided in the Supplementary File S1.

### Performance evaluation with 16S rRNA datasets

A novel and three published datasets were analyzed with PIME. The novel dataset used in this work comprised of 16S rRNA gene sequences from saliva samples obtained from 125 undergraduate and graduate students from the University of Florida. The study assessed the subject’s diet as a factor influencing the saliva microbiome. This study was approved by the University of Florida’s Institutional Review Board and assigned number IRB201602134. Approximately 224 undergraduate and graduate students taking three courses were invited to anonymously participate in this study as volunteers. A study coordinator was chosen to collect samples and code the samples so that those who did the analysis were unaware of the identity of the volunteers. To assess the diet, the subjects also completed the KIDMED survey (18). The sampling collection, DNA extraction and library preparation are described below.

#### Sampling collection, DNA extraction and library preparation

Of the 224 students invited, 125 volunteers obtained the saliva sample collection and provided 2 ml of saliva. The samples were taken from each subject using the GeneFiX™ Saliva DNA Collection device. The collection kit allows immediate stabilization of the DNA. Total DNA was extracted using the GeneFix™ Saliva-prep-2 kit (Cell Projects Ltd, Harrietsham, UK) following the manufacturer’s protocol. DNA samples were stored at −20 °C until use.

To assess the diet, the subjects also completed the KIDMED survey (18). The KIDMED Index is based on a series of 16 questions, which measures the degree to which a subject adheres to the Mediterranean diet. The KIDMED index has been validated with nutritional data (19) and was much simpler to implement than a diet diary or a serum-based nutrition analysis. Participant’s age and gender were also obtained.

The 16S rRNA library preparation was performed as described previously (16) and sequenced with Illumina MiSeq: 2×300 cycles run. The raw fastq files were used to build a table of exact amplicon sequence variants (ASVs) with DADA2 version 1.8 (7). Taxonomy was assigned to each ASV using the SILVA ribosomal RNA gene database version v132 (20). A detailed R script containing the code used to generate the ASV table is provided in the Supplementary File S2. Downstream analyses were carried out using a rarefied dataset of 24,900 sequences as previously recommended by Lemos et al. (21).

#### Description of the previously published datasets

The first previously published dataset used here was described by Davis-Richardson et al. (16) and comprised of partial 16S rDNA sequences from fecal samples of 76 subjects born between 1996 and 2007 at the Turku University Hospital in southwestern Finland. All subjects were at high genetic risk for type 1 diabetes. The cohort was retroactively selected to create an age-matched genotype-controlled set of subjects for the investigation of the microbiome as an environmental factor influencing the development of Type-1 diabetes. The raw Fastq files were obtained and sequences were processed using DADA2 version 1.8 (7), as described above. Cases were defined as subjects who developed at least two persistent islet cell autoantibody (ICA), IAA, GADA, or IA-2A. Controls were defined as subjects with no detectable islet autoantibodies. Samples from subjects older than one year and post seroconversion were removed.

The second published dataset used here was previously described by Avershina et al. (17). The dataset comprised of amplified and sequenced 16S rRNA genes from vaginal swab samples collected from a cohort of 256 pregnant women. These subjects were randomized to receive a daily dose of fermented milk containing probiotic bacterial strains, or milk without probiotics. An OTU table with 3,000 reads per sample and the accompanying metadata were kindly provided by the corresponding author. This table was used in all downstream bioinformatics and statistical analysis. Only those samples collected at the 36^th^ week of gestation were used in these analyses.

The third previously published dataset comprised of 16S rRNA gene sequences from the V1-V3 hypervariable region downloaded from the NIH Human Microbiome Project (https://www.hmpdacc.org/HMQCP/#data). The final OTU table processed by Qiime (9) using an OTU-clustering strategy and accompanying metadata were obtained and loaded into the R environment. After removing singletons, only saliva and left antecubital fossa samples were kept. The final dataset comprised of 113 saliva samples and 59 left antecubital fossa samples all rarefied at 2,000 sequences pre sample. A record of all statistical analyses comparing the datasets with and without using PIME including the R-code and results is included in Supplementary File S3.

### Performance of PIME compared against other filtering methods

The results comparing the performance of PIME with other filtering methods are presented in Figure 2. After quality filtering the saliva dataset, a total of 4,981,638 high-quality paired sequences, 400 bp long, were obtained from all subjects. An average 44,258 reads per sample were obtained. The dataset was rarefied to 24,900 reads per sample in all analyses commensurate with the lowest number of reads found in any one sample. This number of reads was sufficient to accurately reflect the microbial diversity in these samples given the low complexity of saliva samples. The best prevalence interval calculated by PIME was at 65%. This prevalence interval was used to compare the performance of PIME against the other filtering methods. The original dataset, without any filtering, presented 4,555 taxa and a total of 3,112,500 sequences after rarefaction. Both prevalence overall and PIME excluded the highest proportion of ASVs and sequences while filtering by abundance or variance excluded only 78% of ASVs and kept 99.9% of the reads. Nevertheless, the overall prevalence kept 84% of the sequences while PIME kept 68% of the total number of sequences. Without using the PIME filtering the OOB error obtained while attempting to classify the salivary microbe according to the three diet categories was 44% indicating the model had low accuracy in predicting diet according to the microbiota. Overall prevalence, abundance and variance filtering also presented low accuracy in classifying diet according to the microbiota however, after PIME filtering the accuracy of the model increased to 100%.

**FIGURE 2.**
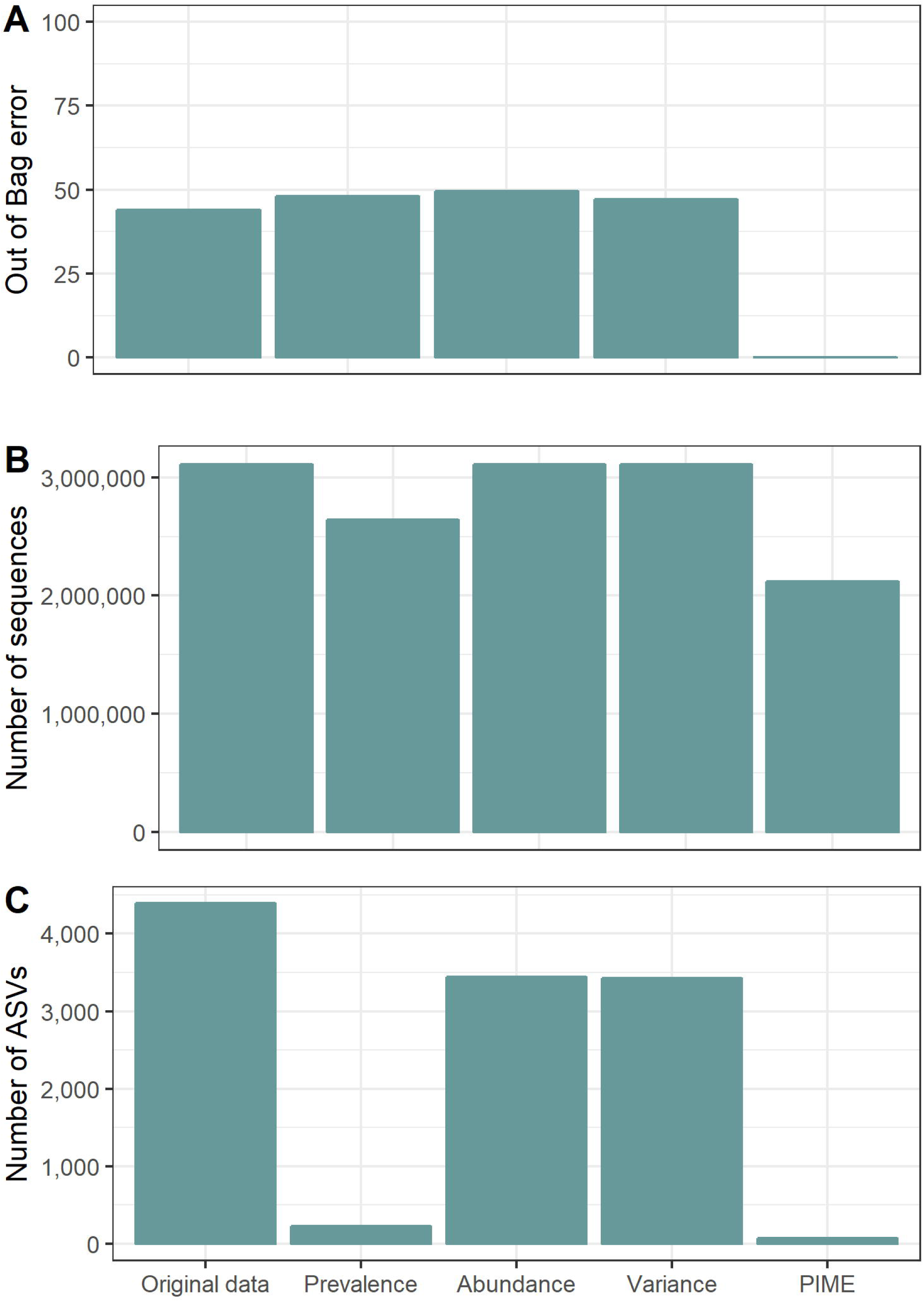
Performance of PIME compared to other filtering methods. A) Out of Bag error rate (OOB error rate); B) total number of sequences; C) Total number of ASVs. Prevalence = filter by overall taxa prevalence in at least 20% of the subjects; Abundance = filter by abundance of at least 5 sequences; Variance = filter by variance higher than 20%. PIME = filter by prevalence interval of 65%.

### PIME application and effectiveness

Different datasets were used to validate the PIME workflow. The computations of the OOB error rate from random forests, the number of taxa and the number of remaining sequences for each prevalence interval from the diet-saliva dataset are presented in Table 1. Stringent criteria for definition of prevalence lead to greater improvement in accuracy for predicting diet based on the salivary microbiota. The prevalence interval of 65% provided the best separation of microbial communities (OOB error = zero) while still including the majority of the sequences in the analysis. This prevalence interval was chosen for further analysis, but other intervals of prevalence can also be tested. For instance, the prevalence interval of 25% had OOB error of 7.2%. This indicates that the model is 92.8% accurate, which is a reasonably good model and keep 88% of sequences. The importance of each ASV in finding microbiome differences among diet categories (high, medium, or low diet categories) for the prevalence interval of 65% is presented in Table 2. The table indicates the ability of each variable to classify the microbes according to the three diet categories. The ASVs are ordered as most- to least-important. The more the accuracy of the random forest decreases due to the exclusion of a single ASV, the more important that variable is, and therefore variables with a large mean decrease in accuracy are more important for classification of the ASVs according to diet. The mean decrease accuracy of the unfiltered dataset presented negative values, which are a clear warning sign the model might be overfitting noise (Table 2). On the other hand, after PIME filtering, the mean decrease accuracy values were all positive indicating a true contribution of each ASV to classify diet according to the microbiota. Altogether, the results indicated that after PIME filtering differences in the saliva microbiome was partially explained by diet rather than by random distribution patterns. The traditional approach, not accounting for microbial prevalence, was unable to distinguish these differences.

**Table 1.** Computations of the out-of-bag error rate from random forests, number of taxa and number of remaining sequences for each prevalence interval from the diet-saliva dataset.

| Prevalence Interval | OOB error rate (%) | Number of OTUs | Number of seqs. |
| --- | --- | --- | --- |
| 0.05 | 44.8 | 1,158 | 2,915,670 |
| 0.10 | 34.4 | 627 | 2,797,858 |
| 0.15 | 24 | 448 | 2,715,217 |
| 0.20 | 16.8 | 327 | 2,653,319 |
| 0.25 | 7.2 | 235 | 2,587,433 |
| 0.30 | 7.2 | 196 | 2,547,536 |
| 0.35 | 2.4 | 169 | 2,479,055 |
| 0.40 | 0.8 | 151 | 2,438,026 |
| 0.45 | 1.6 | 130 | 2,383,250 |
| 0.50 | 1.6 | 104 | 2,302,147 |
| 0.55 | 1.6 | 91 | 2,264,871 |
| 0.60 | 0.8 | 84 | 2,222,431 |
| 0.65 | 0 | 76 | 2,120,215 |
| 0.70 | 0 | 66 | 1,972,958 |
| 0.75 | 0 | 53 | 1,824,764 |
| 0.80 | 0.8 | 43 | 1,747,231 |
| 0.85 | 0 | 34 | 1,612,451 |
| 0.90 | 0 | 26 | 1,351,690 |
| 0.95 | 0 | 17 | 1,078,200 |

**Table 2.**
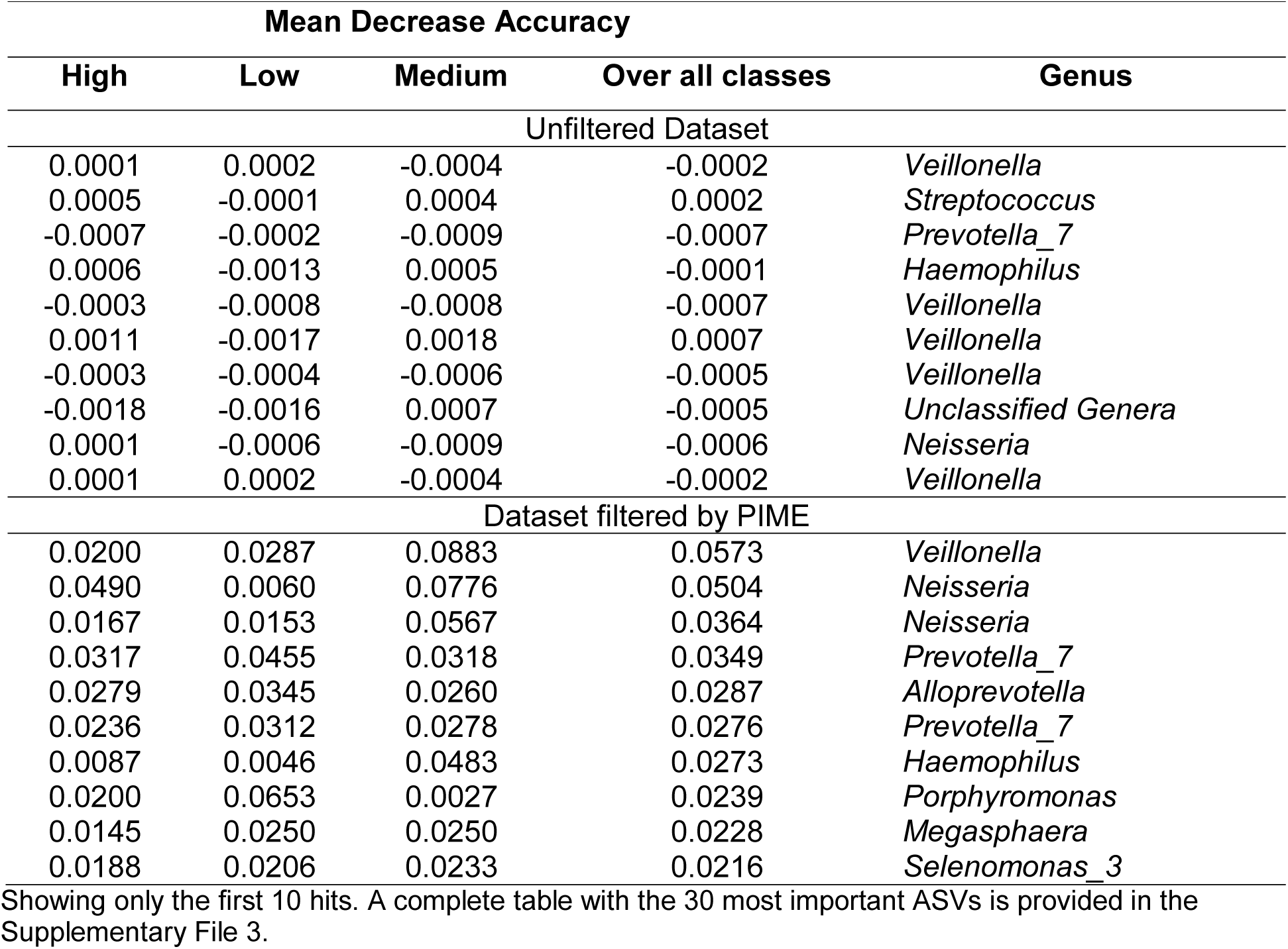
Importance of ASVs measured by mean decrease accuracy to differentiate the three diet categories (High, Low and Medium) from the diet-saliva dataset.

Following this first test, 16S rDNA data from stool of 76 children at high genetic risk for type 1 diabetes (16) were tested for prevalence differences in those samples from children who remained healthy versus those that became autoimmune. The computations of the OOB error rate from random forests, the number of taxa and the number of remaining sequences for each prevalence interval from the dataset described by Davis-Richardson *et al.* (2014) are presented in Table 3. PIME was able to calculate prevalence interval up to 70%. At prevalence intervals higher than 70% samples had zero counts and prevalence was not calculated. As expected, the OOB error rate decreased with higher prevalence intervals. At 60% prevalence interval the OOB error was zero and the number of remaining sequences was 1,165,304. The importance of each ASV in finding microbiome differences among cases and controls subjects under risk for T1 diabetes for the prevalence interval of 60% is presented in Table 4. Comparing the results obtained by the unfiltered dataset with the PIME filtered dataset we observe an improvement in accuracy. Previously, Davis-Richardson et al. (2014) discovered that the relative abundance of *Bacteroides* was significantly higher in autoimmune vs. control subjects. The higher abundance of *Bacteroides* was confirmed by PIME and other Amplicon Sequence Variants (ASVs) belonging to *Bifidobacterium* genus were also found associated with autoimmune subjects.

**Table 3.** Computations of the out-of-bag error rate from random forests, number of taxa and number of remaining sequences for each prevalence interval from the dataset described by Davis-Richardson *et al.* (2014)

| Prevalence Interval | OOB error rate (%) | Number of OTUs | Number of seqs. |
| --- | --- | --- | --- |
| 0.05 | 13.17 | 979 | 3,152,779 |
| 0.10 | 10.64 | 556 | 2,975,527 |
| 0.15 | 6.16 | 431 | 2,891,465 |
| 0.20 | 3.64 | 335 | 2,773,432 |
| 0.25 | 3.36 | 258 | 2,556,564 |
| 0.30 | 2.8 | 219 | 2,457,867 |
| 0.35 | 2.24 | 175 | 2,339,966 |
| 0.40 | 2.24 | 143 | 2,119,062 |
| 0.45 | 1.4 | 115 | 1,922,701 |
| 0.50 | 2.52 | 99 | 1,733,801 |
| 0.55 | 1.12 | 74 | 1,357,557 |
| 0.60 | 0 | 62 | 1,165,304 |
| 0.65 | 0.28 | 49 | 1,026,879 |
| 0.70 | 0.56 | 39 | 897,367 |

**Table 4.** Importance of ASVs measured by mean decrease accuracy to differentiate the cases and controls at high genetic risk for type 1 diabetes from the dataset described by Davis-Richardson *et al.* (2014).

| Mean Decrease Accuracy |  |  |  |
| --- | --- | --- | --- |
| Controls | Cases | Over all classes | Genus |
| Unfiltered Dataset |  |  |  |
| 0.0028 | 0.0061 | 0.0040 | *Bacteroides* |
| 0.0031 | 0.0033 | 0.0031 | *Bacteroides* |
| 0.0011 | 0.0036 | 0.0020 | *Bacteroides* |
| 0.0037 | 0.0017 | 0.0030 | *Bacteroides* |
| 0.0006 | 0.0012 | 0.0008 | *Bacteroides* |
| 0.0012 | 0.0003 | 0.0008 | *Bacteroides* |
| 0.0025 | 0.0020 | 0.0023 | *Bacteroides* |
| 0.0021 | 0.0015 | 0.0019 | *Bacteroides* |
| 0.0028 | 0.0047 | 0.0035 | *Bacteroides* |
| 0.0005 | 0.0008 | 0.0006 | *Bifidobacterium* |
| Dataset filtered by PIME |  |  |  |
| 0.0588 | 0.0148 | 0.0422 | *Bacteroides* |
| 0.0470 | 0.0119 | 0.0339 | *Bacteroides* |
| 0.0386 | 0.0118 | 0.0284 | *Bifidobacterium* |
| 0.0372 | 0.0073 | 0.0260 | *Bifidobacterium* |
| 0.0375 | 0.0063 | 0.0257 | *Bacteroides* |
| 0.0366 | 0.0070 | 0.0255 | *Bacteroides* |
| 0.0356 | 0.0074 | 0.0251 | *Bacteroides* |
| 0.0372 | 0.0045 | 0.0251 | *Bacteroides* |
| 0.0366 | 0.0058 | 0.0250 | *Bacteroides* |
| 0.0346 | 0.0076 | 0.0245 | *Bacteroides* |

Showing only the first 10 hits. A complete table with the 30 most important ASVs is provided in the Supplementary File 3.

In the third dataset tested, taxa were equally likely to be detected in the probiotic and placebo groups (17). Prevalence testing by PIME also does not capture any difference between treatments (Table 5). As the vaginal environment is dominated by *Lactobacillus*, a severe drop in the number of sequences at 5% prevalence interval was observed, The OOB error rate of the overall model obtained by Random Forest analysis suggests that irrespective of the prevalence interval no distinction between probiotic consumption and placebo exists (Supplementary File 3). Those results confirm the author’s previous findings and demonstrate our approach is not prone to type I errors (finding false positive results).

**Table 5.** Computations of the out-of-bag error rate from random forests, number of taxa and number of remaining sequences for each prevalence interval from the dataset described by Avershina *et al.*, (2017).

| Prevalence Interval | OOB error rate (%) | Number of OTUs | Number of seqs. |
| --- | --- | --- | --- |
| 0.05 | 28.06 | 123 | 988,309 |
| 0.10 | 14.93 | 68 | 967,461 |
| 0.15 | 20.90 | 40 | 954,379 |
| 0.20 | 21.19 | 26 | 931,177 |
| 0.25 | 27.46 | 22 | 929,563 |
| 0.30 | 25.67 | 17 | 903,330 |
| 0.35 | 33.73 | 14 | 899,992 |
| 0.40 | 17.91 | 14 | 898,656 |
| 0.45 | 20.00 | 10 | 889,244 |
| 0.50 | 42.69 | 7 | 881,104 |
| 0.55 | 14.63 | 7 | 853,874 |
| 0.60 | 17.01 | 6 | 853,217 |
| 0.65 | 18.21 | 5 | 797,394 |
| 0.70 | 53.73 | 3 | 745,832 |
| 0.75 | 47.16 | 2 | 663,347 |
| 0.80 | 46.57 | 2 | 663,347 |
| 0.85 | 46.87 | 2 | 663,347 |
| 0.90 | 47.16 | 2 | 663,347 |

Finally, PIME was tested using the association between saliva microbiome and the left antecubital fossa, a dataset from the Human Microbiome Project (22). These two distinctive human microbial habitats were selected as they are expected to harbor very different communities. As predicted, PIME showed that the microbial habitats tested are very distinct. The OOB error rate was 0.005 within the original dataset and zero at all prevalence intervals applied (Table 6) indicating the prevalence filtering does not increase the differentiation between these very different microbial habitats.

**Table 6.** Computations of the out-of-bag error rate from random forests, number of taxa and number of remaining sequences for each prevalence interval from the Human Microbiome Project dataset.

| Prevalence Interval | OOB error rate (%) | Number of OTUs | Number of seqs. |
| --- | --- | --- | --- |
| 0.05 | 0 | 509 | 190,455 |
| 0.10 | 0 | 509 | 189,296 |
| 0.15 | 0 | 509 | 188,555 |
| 0.20 | 0 | 506 | 188,189 |
| 0.25 | 0 | 473 | 183,574 |
| 0.30 | 0 | 421 | 176,217 |
| 0.35 | 0 | 328 | 161,346 |
| 0.40 | 0 | 274 | 151,525 |
| 0.45 | 0 | 234 | 142,112 |
| 0.50 | 0 | 185 | 129,369 |
| 0.55 | 0 | 143 | 112,978 |
| 0.60 | 0 | 118 | 104,474 |
| 0.65 | 0 | 74 | 82,052 |
| 0.70 | 0 | 48 | 62,817 |
| 0.75 | 0 | 34 | 56,457 |
| 0.80 | 0 | 23 | 46,475 |
| 0.85 | 0 | 13 | 34,459 |

### Likelihood of introducing bias while building prevalence-filtered datasets

The results obtained by the PIME error detection step are presented in Figure 3. The biological difference among samples is expected to be greater than the differences generated randomly. This way, as the prevalence interval increases the OOB error might decrease. As expected, the OOB error rate of samples with true biological relevant differences (Figures 3A, 3B and 3D) decreased (or remained constant in low noise datasets – Figure 3D) with the increase in the prevalence interval definition. On the other hand, random sampling produced OOB error rate always higher than those obtained based on the original dataset. In datasets with no expected biological relevant differences (Figure 3C), the OOB error did not decrease with higher prevalence interval definitions and the randomized dataset produced higher OOB error rates. Thus, the signal to noise ratio increases with the prevalence intervals generating low OOB error rate values while no improvements in accuracy are observed within the randomized datasets. This error detection analysis showed that no bias was introduced while building prevalence-filtered datasets confirming this workflow is not prone to type I errors.

**FIGURE 3.**
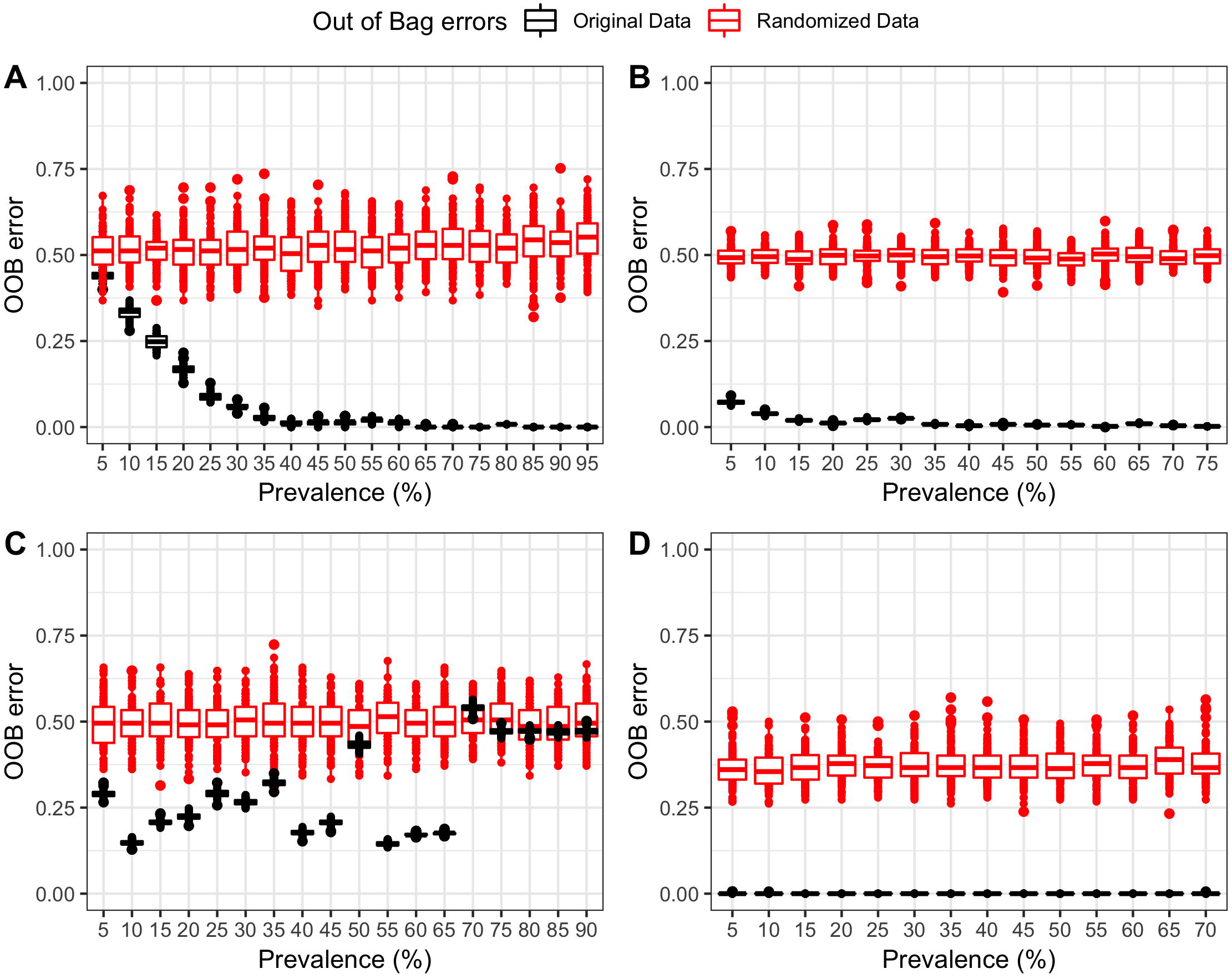
Boxplot depicting the PIME error detection step. Red boxes represent the OOB error rate obtained by randomly shuffling the labels into arbitrary groupings using 100 random permutations and running pime.error.prediction function at each randomization for each prevalence interval. Black boxes represent the OOB error rate against the 100 replications in each prevalence interval against the original sampling labels obtained by running pime.oob.replicate function. (A) Original dataset from salivary microbiome samples. (B) Data from the gut microbial of 76 children at high genetic risk for type 1 diabetes. (C) Data from the vaginal microbiome of pregnant women randomized to receive milk with or without probiotic bacterial strains. (D) Data from the microbiome of saliva and left antecubital fossa of healthy individuals. Boxes span the first to third quartiles; the horizontal line inside the boxes represents the median. Whiskers extending vertically from the boxes indicate variability outside the upper and lower quartiles, and the circles indicate outliers.

## CONCLUSIONS

Prevalence is a key epidemiological concept involving the counting of the number of people affected by a disease (23, 24). PIME was designed based on this concept. Here we argue the importance of a microbial community found in a single sample is smaller than if the same community is present in the majority of samples. Under such rationale, we designed a workflow capable of improving the ability to detect important organisms as it considers the extent to which an organism is present across a given population, which may be masked when only relative abundance is considered. Challenges in microbiome data include sparcity (presence of many zeros) and large variance in distribution patters (also known as over-dispersion) with prominent abundance of some microbes in some subjects/samples and nearly absence in others (4, 25). The current major challenge for using this information is indubitably how to convert it into rational biological conclusions providing control for error rates of false discoveries. Many tools have been successfully developed aiming to contrast experimental factors but they usually only take into account the microbial abundance and/or presence/absence. Thus, PIME is expected surpass those challenges including the concept of a per treatment microbial prevalence in the analysis. This approach greatly improves the results by removing interpersonal variation within groups (unique microbes found in a single subject/sample) and keeping only microbes found in most of the subjects of a population (likely to be associated with a experimental variable). This approach reveals microbes important in a disease that may be overlooked by traditional methods leading to a greater understanding of pathogenesis and the identification of potential probiotic treatment and prevention strategies.

Several tools designed to support microbiome statistical data analysis include data filtering as one of the first steps. The most commonly used filtering includes the exclusion of low count features (low abundance) using a minimum, yet arbitrary, cutoff, low variance (assuming that features under low variance are very unlikely to be significant in the comparative analysis) and low overall prevalence. Arguably filtering those uninformative taxa can improve the data sparsity issue, improving statistical power. Here we compare the performance of PIME with these other filtering methods. PIME outperformed all of those other approaches reducing the error rate and detecting microbial community differences where none were seen by other methods. To illustrate the application and the value of PIME, it was also implemented in a variety of 16S rRNA datasets. Within all of our tests we confirmed previews findings and improved the results.

During the course of analysis and tests we also detected some potential limitations of PIME. As PIME relies strongly on group prevalence, it is sensitive to the quality of sample groups. Poorly categorized groups made up of subjects/samples with very different microbial composition might affect the prevalence computations and therefore PIME might not be as effective in suggesting a good prevalence interval for filtering. For datasets with very large number of samples, PIME might not find a clear prevalence interval for data filtering. With increased number of samples, the chance of sampling different “cores” or subpopulations is also increased. In addition, when there is large heterogeneity within sample groups, coupled with high data sparsity, prevalence computation might not be successful. Another possible limitation of PIME rises from random forests method, wrapped in pime.oob.error and pime.best.prevalence functions. Random forests models are sensitive to multicollinear variables when informing variable importance, though it doesn’t affect prediction errors. Colinear variables might have inaccurate importance values as the difference is explained by the, randomly, first chosen variable and little information is added to the model after this.

## Supporting information

Comparison of PIME with other existing filtering methods.

The pipeline used to assign 16S rRNA sequences to ASVs.

Detailed and reproducible description of PIME data analysis.

## Acknowledgements

L.F.W.R. and P.C.T.D. were supported by Conselho Nacional de Desenvolvimento Científico e Tecnológico (CNPq) and Coordenação de Aperfeiçoamento de Pessoal de Nível Superior (CAPES), respectively. The work was supported by grants to E.W.T. from NIH (5R21AI120195-02) and JDRF (1-INO-2018-637-A-N).

## Availability and implementation

The R package, installation instructions and a step-by-step example on how to use PIME are freely available at: https://github.com/microEcology/PIME.

## Supplementary information

**Supplementary File S1.** Comparison of PIME with other existing filtering methods.

**Supplementary File S2.** The pipeline used to assign 16S rRNA sequences to ASVs.

**Supplementary File S3.** Detailed and reproducible description of PIME data analysis.

