## Supplementary material for "PIME: a package for discovery of novel differences among microbial communities": Comparison of PIME with other existing filtering methods.

### Comparison with other existing filtering methods

*Roesch et al.*

*4/11/2019*

##### Load objects generated by dada2

##### Make a phyloseq object

```
library(phyloseq)
library(ggplot2)
library(microbiome)
library(vegan)
library(knitr)
library(ggpubr)
library(pime)
seqtab = readRDS("/Users/luizroesch/Desktop/Saliva/seqs/dada2/fastq/seqtab_final.rds")
tax= readRDS("/Users/luizroesch/Desktop/Saliva/seqs/dada2/fastq/tax_final.rds")
taxa = readRDS("/Users/luizroesch/Desktop/Saliva/seqs/dada2/fastq/taxa_sp_final.rds")
map <- "/Users/luizroesch/Desktop/Saliva/seqs/dada2/map.txt"
ps <- phyloseq(otu_table(seqtab, taxa_are_rows=FALSE),
               tax_table(taxa))
sample_metadata = import_qiime_sample_data(map)
saliva =merge_phyloseq(ps, sample_metadata)
summarize_phyloseq(saliva)
```

```
## Compositional = NO
## 1] Min. number of reads = 24956
## 2] Max. number of reads = 223138
## 3] Total number of reads = 5532280
## 4] Average number of reads = 44258.24
## 5] Median number of reads = 39432
```

```
## 7] Sparsity = 0.95352096597146
## 6] Any OTU sum to 1 or less? NO
## 8] Number of singletons = 0
## 9] Percent of OTUs that are singletons 0
## 10] Number of sample variables are: 21
## X.NAME
## KIDMED_index
## Gender
## Age
## Fruit.daily.
## Second.fruit.daily.
## Vegetable.daily.
## Second.vegetable.daily.
## Fish.regularly.
## Pulses..1.weekly
## Pasta.rice..5.weekly
## Cereals.grains.breakfast.
## Nut.2.3.weekly
## Olive.oil.used.
## Dairy.at.breakfast
## yogurt.cheese.3.weekly.
## Fast.food..1.weekly
## Skip.breakfast..3.weekly.
## Baked.goods.pastires.breakfast.
## Sweets.3.daily.
## Total
```

```
rank_names(saliva)
```

```
## [1] "Kingdom" "Phylum"  "Class"   "Order"   "Family"  "Genus"   "Species"
```

```
saliva
```

```
## phyloseq-class experiment-level object
## otu_table()   OTU Table:         [ 4555 taxa and 125 samples ]
## sample_data() Sample Data:       [ 125 samples by 21 sample variables ]
## tax_table()   Taxonomy Table:    [ 4555 taxa by 7 taxonomic ranks ]
```

```
set.seed(2125)
salivaR = rarefy_even_depth(saliva, sample.size = 24900)
```

### To test the performance of PIME: comparison with other existing filtering methods

#### Filtering by overall prevalence 20%

```
saliva.prev20 = filter_taxa(salivaR, function(x) sum(x > 0) > (0.2*length(x)), TRUE)
saliva.prev20
```

```
## phyloseq-class experiment-level object
## otu_table()   OTU Table:         [ 225 taxa and 125 samples ]
## sample_data() Sample Data:       [ 125 samples by 21 sample variables ]
## tax_table()   Taxonomy Table:    [ 225 taxa by 7 taxonomic ranks ]
```

#### Randon Forest approach to obtain the OOB error rate

```r
library(randomForest)
train= otu_table(saliva.prev20)
# Make one column for our outcome/response variable
KIDMED_index <- as.factor(sample_data(saliva.prev20)$KIDMED_index)
# Combine them into 1 data frame
training.set <- data.frame(KIDMED_index, train)
train.model = randomForest(KIDMED_index ~ ., data = training.set, importance = TRUE)
print(train.model)
```

```
##
## Call:
##  randomForest(formula = KIDMED_index ~ ., data = training.set,      importance = TRUE)
##                Type of random forest: classification
##                      Number of trees: 500
## No. of variables tried at each split: 15
##
##         OOB estimate of  error rate: 48%
## Confusion matrix:
##        High Low Medium class.error
## High      0   2     22   1.0000000
## Low       1   8     28   0.7837838
## Medium    0   7     57   0.1093750
```

#### Filtering by abundance

```r
saliva.ab5 = filter_taxa(salivaR, function(x) sum(x) > 5, TRUE)
saliva.ab5
```

```
## phyloseq-class experiment-level object
## otu_table()   OTU Table:         [ 3447 taxa and 125 samples ]
## sample_data() Sample Data:       [ 125 samples by 21 sample variables ]
## tax_table()   Taxonomy Table:    [ 3447 taxa by 7 taxonomic ranks ]
```

#### Randon Forest approach to obtain the OOB error rate

```r
library(randomForest)
train= otu_table(saliva.ab5)
# Make one column for our outcome/response variable
KIDMED_index <- as.factor(sample_data(saliva.ab5)$KIDMED_index)
# Combine them into 1 data frame
training.set <- data.frame(KIDMED_index, train)
train.model = randomForest(KIDMED_index ~ ., data = training.set, importance = TRUE)
print(train.model)
```

```
##
## Call:
##  randomForest(formula = KIDMED_index ~ ., data = training.set,      importance = TRUE)
##                Type of random forest: classification
##                      Number of trees: 500
## No. of variables tried at each split: 58
```

```
##
##         OOB estimate of  error rate: 49.6%
## Confusion matrix:
##        High Low Medium class.error
## High      0   2     22   1.0000000
## Low       0   4     33   0.8918919
## Medium    0   5     59   0.0781250
```

##### Filtering by low variance

```
saliva.var20 = filter_taxa(salivaR, function(x) var(x) > 0.2, TRUE)
saliva.var20
```

```
## phyloseq-class experiment-level object
## otu_table()   OTU Table:         [ 3427 taxa and 125 samples ]
## sample_data() Sample Data:       [ 125 samples by 21 sample variables ]
## tax_table()   Taxonomy Table:    [ 3427 taxa by 7 taxonomic ranks ]
```

##### Randon Forest approach to obtain the OOB error rate

```
library(randomForest)
train= otu_table(saliva.var20)
# Make one column for our outcome/response variable
KIDMED_index <- as.factor(sample_data(saliva.var20)$KIDMED_index)
# Combine them into 1 data frame
training.set <- data.frame(KIDMED_index, train)
train.model = randomForest(KIDMED_index ~ ., data = training.set, importance = TRUE)
print(train.model)
```

```
##
## Call:
##  randomForest(formula = KIDMED_index ~ ., data = training.set,      importance = TRUE)
##                Type of random forest: classification
##                      Number of trees: 500
## No. of variables tried at each split: 58
##
##         OOB estimate of  error rate: 47.2%
## Confusion matrix:
##        High Low Medium class.error
## High      0   2     22   1.0000000
## Low       0   6     31   0.8378378
## Medium    0   4     60   0.0625000
```

#### Running PIME

##### Prediction using random forests on full dataset

```
OOB_error_full=pime.oob.error(salivaR, "KIDMED_index")
OOB_error_full
```

```
## [1] 0.44
```

The OOB error rate <=0.1, indicated the dataset present large differences, and pime might not remove much of the noise. Higher OOB error rate indicates that the next functions should be run to find the best prevalence interval for the dataset.

##### Split the dataset by predictor variable

```r
per_variable_obj= pime.split.by.variable(salivaR, "KIDMED_index")
per_variable_obj
```

```
## $High
## phyloseq-class experiment-level object
## otu_table()   OTU Table:         [ 1749 taxa and 24 samples ]
## sample_data() Sample Data:       [ 24 samples by 21 sample variables ]
## tax_table()   Taxonomy Table:    [ 1749 taxa by 7 taxonomic ranks ]
##
## $Low
## phyloseq-class experiment-level object
## otu_table()   OTU Table:         [ 2132 taxa and 37 samples ]
## sample_data() Sample Data:       [ 37 samples by 21 sample variables ]
## tax_table()   Taxonomy Table:    [ 2132 taxa by 7 taxonomic ranks ]
##
## $Medium
## phyloseq-class experiment-level object
## otu_table()   OTU Table:         [ 3048 taxa and 64 samples ]
## sample_data() Sample Data:       [ 64 samples by 21 sample variables ]
## tax_table()   Taxonomy Table:    [ 3048 taxa by 7 taxonomic ranks ]
```

##### Calculate the highest possible prevalence intervals

This function calculates prevalence for different intervals by increments of 5. The input file is the output from the pime.split.by.variable(per_variable_obj)

```r
prevalences=pime.prevalence(per_variable_obj)
head(prevalences)
```

```
## $`5`
## phyloseq-class experiment-level object
## otu_table()   OTU Table:         [ 1158 taxa and 125 samples ]
## sample_data() Sample Data:       [ 125 samples by 21 sample variables ]
## tax_table()   Taxonomy Table:    [ 1158 taxa by 7 taxonomic ranks ]
##
## $`10`
## phyloseq-class experiment-level object
## otu_table()   OTU Table:         [ 627 taxa and 125 samples ]
## sample_data() Sample Data:       [ 125 samples by 21 sample variables ]
## tax_table()   Taxonomy Table:    [ 627 taxa by 7 taxonomic ranks ]
##
## $`15`
## phyloseq-class experiment-level object
## otu_table()   OTU Table:         [ 448 taxa and 125 samples ]
## sample_data() Sample Data:       [ 125 samples by 21 sample variables ]
```

```
## tax_table()   Taxonomy Table:    [ 448 taxa by 7 taxonomic ranks ]
##
## $`20`
## phyloseq-class experiment-level object
## otu_table()   OTU Table:         [ 327 taxa and 125 samples ]
## sample_data() Sample Data:       [ 125 samples by 21 sample variables ]
## tax_table()   Taxonomy Table:    [ 327 taxa by 7 taxonomic ranks ]
##
## $`25`
## phyloseq-class experiment-level object
## otu_table()   OTU Table:         [ 235 taxa and 125 samples ]
## sample_data() Sample Data:       [ 125 samples by 21 sample variables ]
## tax_table()   Taxonomy Table:    [ 235 taxa by 7 taxonomic ranks ]
##
## $`30`
## phyloseq-class experiment-level object
## otu_table()   OTU Table:         [ 196 taxa and 125 samples ]
## sample_data() Sample Data:       [ 125 samples by 21 sample variables ]
## tax_table()   Taxonomy Table:    [ 196 taxa by 7 taxonomic ranks ]
```

#### Calculate the best prevalence interval for the dataset

This function will return a table with Out of Bag error from random forests for each prevalence interval. The number of taxa and the number of remaining sequences for each prevalence interval are also computed. The best prevalence interval value provides the clearest separation of communities while still including a majority of the taxa in the analysis. If true differences are present. It will be represented by the first interval in which the OOB error rate is zero or close to zero. The input file is the list of prevalences generated by the pime.prevalence (prevalences) and the predictor variable ("KIDMED_index").

```r
set.seed(2125)
best.prev=pime.best.prevalence(prevalences, "KIDMED_index")
```

```
##        Interval OOB error rate (%) OTUs   Nseqs
##   Prevalence 5%               44.8 1158 2915670
##  Prevalence 10%               34.4  627 2797858
##  Prevalence 15%                 24  448 2715217
##  Prevalence 20%               16.8  327 2653319
##  Prevalence 25%                7.2  235 2587433
##  Prevalence 30%                7.2  196 2547536
##  Prevalence 35%                2.4  169 2479055
##  Prevalence 40%                0.8  151 2438026
##  Prevalence 45%                1.6  130 2383250
##  Prevalence 50%                1.6  104 2302147
##  Prevalence 55%                1.6   91 2264871
##  Prevalence 60%                0.8   84 2222431
##  Prevalence 65%                  0   76 2120215
##  Prevalence 70%                  0   66 1972958
##  Prevalence 75%                  0   53 1824764
##  Prevalence 80%                0.8   43 1747231
##  Prevalence 85%                  0   34 1612451
##  Prevalence 90%                  0   26 1351690
##  Prevalence 95%                  0   17 1078200
```
