## Supplementary material for "PIME: a package for discovery of novel differences among microbial communities": The pipeline used to assign 16S rRNA sequences to ASVs.

Roesch\_pipeline\_dada2.R


### Roesch\_pipeline\_dada2.R

###### *luizroesch*

###### *Tue Aug 14 13:36:27 2018*

```
#sequences were first joined and demultiplexed by using Qiime
#no quality filtering was applied at this time
#commands performed in Qiime to join de ends and demultiplex the samples
#join_paired_ends.py -f /Users/luizroesch/uclust8/f_seqs.fastq -r /Users/luizroesch/uclust8/r_seqs.fastq -b /Users/luizroesch/uclust8/barcodes.fastq -o joined
#split_libraries_fastq.py -i /Users/luizroesch/uclust8/Analysis/joined/fastqjoin.join.fastq -b /Users/luizroesch/uclust8/Analysis/joined/fastqjoin.join_barcodes.fastq   -o splitout -m /Users/luizroesch/uclust8/map.txt --store_demultiplexed_fastq -r 0 -q 0 -n 100 --barcode_type 11 --rev_comp_barcode
#split_sequence_file_on_sample_ids.py -i /Users/luizroesch/Desktop/Saliva/seqs/Analysis/splitout/seqs.fastq --file_type fastq -o out_fastq/

# Assign ASVs to sequences ------------------------------------------------

library(dada2); packageVersion("dada2")
```

```
## Loading required package: Rcpp
```

```
## [1] '1.9.0'
```

```
#directory containing the fastq files after unzipping
path = "/Users/luizroesch/Desktop/Saliva/seqs/dada2/fastq"
list.files(path)
```

```
##   [1] "9010_.fastq"       "9011_.fastq"       "9012_.fastq"      
##   [4] "9013_.fastq"       "9014_.fastq"       "9015_.fastq"      
##   [7] "9016_.fastq"       "9018_.fastq"       "9019_.fastq"      
##  [10] "9020_.fastq"       "9021_.fastq"       "9022_.fastq"      
##  [13] "9023_.fastq"       "9024_.fastq"       "9025_.fastq"      
##  [16] "9026_.fastq"       "9027_.fastq"       "9028_.fastq"      
##  [19] "9029_.fastq"       "9030_.fastq"       "9031_.fastq"      
##  [22] "9032_.fastq"       "9033_.fastq"       "9034_.fastq"      
##  [25] "9035_.fastq"       "9036_.fastq"       "9037_.fastq"      
##  [28] "9038_.fastq"       "9039_.fastq"       "9041_.fastq"      
##  [31] "9042_.fastq"       "9043_.fastq"       "9044_.fastq"      
##  [34] "9045_.fastq"       "9048_.fastq"       "9049_.fastq"      
##  [37] "9050_.fastq"       "9051_.fastq"       "9052_.fastq"      
##  [40] "9053_.fastq"       "9054_.fastq"       "9055_.fastq"      
##  [43] "9056_.fastq"       "9057_.fastq"       "9058_.fastq"      
##  [46] "9059_.fastq"       "9061_.fastq"       "9062_.fastq"      
##  [49] "9063_.fastq"       "9064_.fastq"       "9065_.fastq"      
##  [52] "9066_.fastq"       "9067_.fastq"       "9068_.fastq"      
##  [55] "9069_.fastq"       "9070_.fastq"       "9071_.fastq"      
##  [58] "9073_.fastq"       "9074_.fastq"       "9075_.fastq"      
##  [61] "9076_.fastq"       "9077_.fastq"       "9079_.fastq"      
##  [64] "9080_.fastq"       "9081_.fastq"       "9083_.fastq"      
##  [67] "9084_.fastq"       "9085_.fastq"       "9086_.fastq"      
##  [70] "9087_.fastq"       "9088_.fastq"       "9089_.fastq"      
##  [73] "9090_.fastq"       "9091_.fastq"       "9093_.fastq"      
##  [76] "9094_.fastq"       "9095_.fastq"       "9096_.fastq"      
##  [79] "9097_.fastq"       "9098_.fastq"       "9099_.fastq"      
##  [82] "9110_.fastq"       "9111_.fastq"       "9112_.fastq"      
##  [85] "9113_.fastq"       "9114_.fastq"       "9120_.fastq"      
##  [88] "9121_.fastq"       "9122_.fastq"       "9123_.fastq"      
##  [91] "9124_.fastq"       "9130_.fastq"       "9131_.fastq"      
##  [94] "9132_.fastq"       "9133_.fastq"       "9134_.fastq"      
##  [97] "9140_.fastq"       "9141_.fastq"       "9142_.fastq"      
## [100] "9143_.fastq"       "9144_.fastq"       "9150_.fastq"      
## [103] "9151_.fastq"       "9152_.fastq"       "9153_.fastq"      
## [106] "9154_.fastq"       "9160_.fastq"       "9161_.fastq"      
## [109] "9162_.fastq"       "9164_.fastq"       "9170_.fastq"      
## [112] "9171_.fastq"       "9172_.fastq"       "9173_.fastq"      
## [115] "9174_.fastq"       "9180_.fastq"       "9181_.fastq"      
## [118] "9182_.fastq"       "9183_.fastq"       "9184_.fastq"      
## [121] "9190_.fastq"       "9191_.fastq"       "9192_.fastq"      
## [124] "9193_.fastq"       "9194_.fastq"       "filtered"         
## [127] "seqtab_final.rds"  "seqtab.rds"        "tax_final.rds"    
## [130] "taxa_sp_final.rds"
```

```
#read in the names of the fastq files, 
#and perform some string manipulation to get lists of 
#the fastq files in matched order
fnFs <- sort(list.files(path, pattern="_.fastq", full.names = TRUE))
sample.names <- sapply(strsplit(basename(fnFs), "_"), `[`, 1)
sample.names
```

```
##   [1] "9010" "9011" "9012" "9013" "9014" "9015" "9016" "9018" "9019" "9020"
##  [11] "9021" "9022" "9023" "9024" "9025" "9026" "9027" "9028" "9029" "9030"
##  [21] "9031" "9032" "9033" "9034" "9035" "9036" "9037" "9038" "9039" "9041"
##  [31] "9042" "9043" "9044" "9045" "9048" "9049" "9050" "9051" "9052" "9053"
##  [41] "9054" "9055" "9056" "9057" "9058" "9059" "9061" "9062" "9063" "9064"
##  [51] "9065" "9066" "9067" "9068" "9069" "9070" "9071" "9073" "9074" "9075"
##  [61] "9076" "9077" "9079" "9080" "9081" "9083" "9084" "9085" "9086" "9087"
##  [71] "9088" "9089" "9090" "9091" "9093" "9094" "9095" "9096" "9097" "9098"
##  [81] "9099" "9110" "9111" "9112" "9113" "9114" "9120" "9121" "9122" "9123"
##  [91] "9124" "9130" "9131" "9132" "9133" "9134" "9140" "9141" "9142" "9143"
## [101] "9144" "9150" "9151" "9152" "9153" "9154" "9160" "9161" "9162" "9164"
## [111] "9170" "9171" "9172" "9173" "9174" "9180" "9181" "9182" "9183" "9184"
## [121] "9190" "9191" "9192" "9193" "9194"
```

```
#visualizing the quality profiles of the reads
plotQualityProfile(fnFs[1:4])
```

```
## Scale for 'y' is already present. Adding another scale for 'y', which
## will replace the existing scale.
```

```
#Assign the filenames for the filtered fastq.gz files
filtFs <- file.path(path, "filtered", paste0(sample.names, "_filt.fastq.gz"))


#Filter and trim
out <- filterAndTrim(fnFs, filtFs, truncLen=c(400), trimLeft=c(5), maxN=0, maxEE=c(1), truncQ=11, rm.phix=TRUE, compress=TRUE, multithread=TRUE)
head(out)
```

```
##             reads.in reads.out
## 9010_.fastq    95312     46578
## 9011_.fastq   106219     55553
## 9012_.fastq   114012     57769
## 9013_.fastq    98657     50539
## 9014_.fastq   105345     54613
## 9015_.fastq    99007     48696
```

```
filtpath <- "/Users/luizroesch/Desktop/Saliva/seqs/dada2/fastq/filtered"
filts <- list.files(filtpath, pattern="fastq.gz", full.names=TRUE)
plotQualityProfile(filts[1:4])
```

```
sample.names <- sapply(strsplit(basename(filts), "_"), `[`, 1)
names(filts) <- sample.names

# Learn error rates
set.seed(2125)
err <- learnErrors(filts, nbases = 1e8, multithread=TRUE, randomize=TRUE)
```

```
## 112513775 total bases in 284845 reads from 6 samples will be used for learning the error rates.
## Initializing error rates to maximum possible estimate.
## selfConsist step 1 ......
##    selfConsist step 2
##    selfConsist step 3
##    selfConsist step 4
##    selfConsist step 5
##    selfConsist step 6
## Convergence after  6  rounds.
```

```
# Infer sequence variants
dds <- vector("list", length(sample.names))
names(dds) <- sample.names
for(sam in sample.names) {
  cat("Processing:", sam, "\n")
  derep <- derepFastq(filts[[sam]])
  dds[[sam]] <- dada(derep, err=err, multithread=TRUE)
}
```

```
## Processing: 9010 
## Sample 1 - 46578 reads in 15769 unique sequences.
## Processing: 9011 
## Sample 1 - 55553 reads in 18033 unique sequences.
## Processing: 9012 
## Sample 1 - 57769 reads in 16228 unique sequences.
## Processing: 9013 
## Sample 1 - 50539 reads in 16715 unique sequences.
## Processing: 9014 
## Sample 1 - 54613 reads in 15349 unique sequences.
## Processing: 9015 
## Sample 1 - 48696 reads in 18313 unique sequences.
## Processing: 9016 
## Sample 1 - 39436 reads in 13565 unique sequences.
## Processing: 9018 
## Sample 1 - 43137 reads in 14101 unique sequences.
## Processing: 9019 
## Sample 1 - 49171 reads in 15504 unique sequences.
## Processing: 9020 
## Sample 1 - 33998 reads in 10980 unique sequences.
## Processing: 9021 
## Sample 1 - 49282 reads in 14565 unique sequences.
## Processing: 9022 
## Sample 1 - 50095 reads in 16838 unique sequences.
## Processing: 9023 
## Sample 1 - 50338 reads in 16589 unique sequences.
## Processing: 9024 
## Sample 1 - 48141 reads in 14755 unique sequences.
## Processing: 9025 
## Sample 1 - 40292 reads in 12122 unique sequences.
## Processing: 9026 
## Sample 1 - 51643 reads in 14600 unique sequences.
## Processing: 9027 
## Sample 1 - 43945 reads in 10961 unique sequences.
## Processing: 9028 
## Sample 1 - 95647 reads in 28977 unique sequences.
## Processing: 9029 
## Sample 1 - 229364 reads in 64601 unique sequences.
## Processing: 9030 
## Sample 1 - 44799 reads in 10067 unique sequences.
## Processing: 9031 
## Sample 1 - 48013 reads in 14838 unique sequences.
## Processing: 9032 
## Sample 1 - 47005 reads in 15765 unique sequences.
## Processing: 9033 
## Sample 1 - 47576 reads in 15067 unique sequences.
## Processing: 9034 
## Sample 1 - 48428 reads in 18078 unique sequences.
## Processing: 9035 
## Sample 1 - 51841 reads in 19547 unique sequences.
## Processing: 9036 
## Sample 1 - 45331 reads in 14720 unique sequences.
## Processing: 9037 
## Sample 1 - 52125 reads in 20772 unique sequences.
## Processing: 9038 
## Sample 1 - 51521 reads in 17687 unique sequences.
## Processing: 9039 
## Sample 1 - 55319 reads in 16737 unique sequences.
## Processing: 9041 
## Sample 1 - 43658 reads in 11690 unique sequences.
## Processing: 9042 
## Sample 1 - 47795 reads in 13012 unique sequences.
## Processing: 9043 
## Sample 1 - 50003 reads in 16406 unique sequences.
## Processing: 9044 
## Sample 1 - 40547 reads in 14185 unique sequences.
## Processing: 9045 
## Sample 1 - 56254 reads in 22314 unique sequences.
## Processing: 9048 
## Sample 1 - 39176 reads in 12881 unique sequences.
## Processing: 9049 
## Sample 1 - 51732 reads in 16012 unique sequences.
## Processing: 9050 
## Sample 1 - 49470 reads in 15647 unique sequences.
## Processing: 9051 
## Sample 1 - 47233 reads in 15680 unique sequences.
## Processing: 9052 
## Sample 1 - 43753 reads in 15105 unique sequences.
## Processing: 9053 
## Sample 1 - 40147 reads in 12193 unique sequences.
## Processing: 9054 
## Sample 1 - 43228 reads in 12570 unique sequences.
## Processing: 9055 
## Sample 1 - 38018 reads in 11581 unique sequences.
## Processing: 9056 
## Sample 1 - 46792 reads in 14616 unique sequences.
## Processing: 9057 
## Sample 1 - 262863 reads in 65431 unique sequences.
## Processing: 9058 
## Sample 1 - 39318 reads in 14606 unique sequences.
## Processing: 9059 
## Sample 1 - 45676 reads in 11951 unique sequences.
## Processing: 9061 
## Sample 1 - 46038 reads in 15093 unique sequences.
## Processing: 9062 
## Sample 1 - 45081 reads in 11963 unique sequences.
## Processing: 9063 
## Sample 1 - 43780 reads in 14132 unique sequences.
## Processing: 9064 
## Sample 1 - 48518 reads in 12610 unique sequences.
## Processing: 9065 
## Sample 1 - 46992 reads in 15326 unique sequences.
## Processing: 9066 
## Sample 1 - 42961 reads in 14025 unique sequences.
## Processing: 9067 
## Sample 1 - 50242 reads in 18058 unique sequences.
## Processing: 9068 
## Sample 1 - 261357 reads in 55238 unique sequences.
## Processing: 9069 
## Sample 1 - 44351 reads in 14625 unique sequences.
## Processing: 9070 
## Sample 1 - 38678 reads in 12391 unique sequences.
## Processing: 9071 
## Sample 1 - 42482 reads in 13136 unique sequences.
## Processing: 9073 
## Sample 1 - 48318 reads in 17582 unique sequences.
## Processing: 9074 
## Sample 1 - 52042 reads in 13589 unique sequences.
## Processing: 9075 
## Sample 1 - 27249 reads in 8910 unique sequences.
## Processing: 9076 
## Sample 1 - 40103 reads in 13200 unique sequences.
## Processing: 9077 
## Sample 1 - 46050 reads in 15965 unique sequences.
## Processing: 9079 
## Sample 1 - 42801 reads in 14749 unique sequences.
## Processing: 9080 
## Sample 1 - 43438 reads in 15434 unique sequences.
## Processing: 9081 
## Sample 1 - 40441 reads in 12378 unique sequences.
## Processing: 9083 
## Sample 1 - 40784 reads in 11790 unique sequences.
## Processing: 9084 
## Sample 1 - 39411 reads in 14798 unique sequences.
## Processing: 9085 
## Sample 1 - 42592 reads in 14727 unique sequences.
## Processing: 9086 
## Sample 1 - 44106 reads in 14386 unique sequences.
## Processing: 9087 
## Sample 1 - 49672 reads in 20317 unique sequences.
## Processing: 9088 
## Sample 1 - 52236 reads in 15866 unique sequences.
## Processing: 9089 
## Sample 1 - 52870 reads in 16640 unique sequences.
## Processing: 9090 
## Sample 1 - 43088 reads in 14188 unique sequences.
## Processing: 9091 
## Sample 1 - 50989 reads in 15937 unique sequences.
## Processing: 9093 
## Sample 1 - 45912 reads in 15609 unique sequences.
## Processing: 9094 
## Sample 1 - 45906 reads in 17375 unique sequences.
## Processing: 9095 
## Sample 1 - 61969 reads in 17092 unique sequences.
## Processing: 9096 
## Sample 1 - 44936 reads in 14316 unique sequences.
## Processing: 9097 
## Sample 1 - 43434 reads in 12917 unique sequences.
## Processing: 9098 
## Sample 1 - 56326 reads in 19253 unique sequences.
## Processing: 9099 
## Sample 1 - 47870 reads in 16125 unique sequences.
## Processing: 9110 
## Sample 1 - 42763 reads in 15310 unique sequences.
## Processing: 9111 
## Sample 1 - 44070 reads in 13146 unique sequences.
## Processing: 9112 
## Sample 1 - 41518 reads in 13504 unique sequences.
## Processing: 9113 
## Sample 1 - 53612 reads in 11474 unique sequences.
## Processing: 9114 
## Sample 1 - 48348 reads in 17379 unique sequences.
## Processing: 9120 
## Sample 1 - 49683 reads in 15431 unique sequences.
## Processing: 9121 
## Sample 1 - 40946 reads in 14179 unique sequences.
## Processing: 9122 
## Sample 1 - 45331 reads in 14462 unique sequences.
## Processing: 9123 
## Sample 1 - 41819 reads in 14703 unique sequences.
## Processing: 9124 
## Sample 1 - 44198 reads in 13221 unique sequences.
## Processing: 9130 
## Sample 1 - 38492 reads in 13745 unique sequences.
## Processing: 9131 
## Sample 1 - 45703 reads in 18543 unique sequences.
## Processing: 9132 
## Sample 1 - 34585 reads in 11842 unique sequences.
## Processing: 9133 
## Sample 1 - 42105 reads in 16122 unique sequences.
## Processing: 9134 
## Sample 1 - 46576 reads in 14598 unique sequences.
## Processing: 9140 
## Sample 1 - 44194 reads in 13778 unique sequences.
## Processing: 9141 
## Sample 1 - 51611 reads in 17322 unique sequences.
## Processing: 9142 
## Sample 1 - 48490 reads in 13369 unique sequences.
## Processing: 9143 
## Sample 1 - 46288 reads in 13602 unique sequences.
## Processing: 9144 
## Sample 1 - 51707 reads in 14380 unique sequences.
## Processing: 9150 
## Sample 1 - 42730 reads in 14686 unique sequences.
## Processing: 9151 
## Sample 1 - 42874 reads in 13413 unique sequences.
## Processing: 9152 
## Sample 1 - 44582 reads in 12667 unique sequences.
## Processing: 9153 
## Sample 1 - 50688 reads in 15233 unique sequences.
## Processing: 9154 
## Sample 1 - 51432 reads in 14048 unique sequences.
## Processing: 9160 
## Sample 1 - 42278 reads in 15088 unique sequences.
## Processing: 9161 
## Sample 1 - 36945 reads in 10877 unique sequences.
## Processing: 9162 
## Sample 1 - 40107 reads in 11882 unique sequences.
## Processing: 9164 
## Sample 1 - 218663 reads in 58594 unique sequences.
## Processing: 9170 
## Sample 1 - 49328 reads in 15323 unique sequences.
## Processing: 9171 
## Sample 1 - 47284 reads in 14914 unique sequences.
## Processing: 9172 
## Sample 1 - 47389 reads in 14768 unique sequences.
## Processing: 9173 
## Sample 1 - 54511 reads in 20168 unique sequences.
## Processing: 9174 
## Sample 1 - 39870 reads in 13528 unique sequences.
## Processing: 9180 
## Sample 1 - 36210 reads in 13513 unique sequences.
## Processing: 9181 
## Sample 1 - 49903 reads in 16753 unique sequences.
## Processing: 9182 
## Sample 1 - 43460 reads in 14556 unique sequences.
## Processing: 9183 
## Sample 1 - 43198 reads in 13670 unique sequences.
## Processing: 9184 
## Sample 1 - 48523 reads in 18138 unique sequences.
## Processing: 9190 
## Sample 1 - 37177 reads in 12427 unique sequences.
## Processing: 9191 
## Sample 1 - 35330 reads in 11469 unique sequences.
## Processing: 9192 
## Sample 1 - 39690 reads in 14189 unique sequences.
## Processing: 9193 
## Sample 1 - 42952 reads in 14079 unique sequences.
## Processing: 9194 
## Sample 1 - 49241 reads in 17570 unique sequences.
```

```
# Construct sequence table and write to disk
seqtab <- makeSequenceTable(dds)

saveRDS(seqtab, "/Users/luizroesch/Desktop/Saliva/seqs/dada2/fastq/seqtab.rds")
st.all <- readRDS("/Users/luizroesch/Desktop/Saliva/seqs/dada2/fastq/seqtab.rds")
# Remove chimeras
seqtab <- removeBimeraDenovo(st.all, method="consensus", multithread=TRUE)
dim(seqtab)
```

```
## [1]  125 4555
```

```
tax <- assignTaxonomy(seqtab, "/Users/luizroesch/uclust8/silva_nr_v132_train_set.fa.gz", multithread=TRUE)
unname(tax)
```

```
##         [,1]       [,2]                  [,3]                         
##    [1,] "Bacteria" "Firmicutes"          "Negativicutes"              
##    [2,] "Bacteria" "Firmicutes"          "Bacilli"                    
##    [3,] "Bacteria" "Bacteroidetes"       "Bacteroidia"                
##    [4,] "Bacteria" "Proteobacteria"      "Gammaproteobacteria"        
##    [5,] "Bacteria" "Firmicutes"          "Negativicutes"              
##    [6,] "Bacteria" "Firmicutes"          "Negativicutes"              
##    [7,] "Bacteria" "Firmicutes"          "Negativicutes"              
##    [8,] "Bacteria" "Patescibacteria"     "Saccharimonadia"            
##    [9,] "Bacteria" "Proteobacteria"      "Gammaproteobacteria"        
##   [10,] "Bacteria" "Firmicutes"          "Negativicutes"              
##   [11,] "Bacteria" "Proteobacteria"      "Gammaproteobacteria"        
##   [12,] "Bacteria" "Firmicutes"          "Negativicutes"              
##   [13,] "Bacteria" "Fusobacteria"        "Fusobacteriia"              
##   [14,] "Bacteria" "Firmicutes"          "Negativicutes"              
##   [15,] "Bacteria" "Bacteroidetes"       "Bacteroidia"                
##   [16,] "Bacteria" "Epsilonbacteraeota"  "Campylobacteria"            
##   [17,] "Bacteria" "Proteobacteria"      "Gammaproteobacteria"        
##   [18,] "Bacteria" "Firmicutes"          "Clostridia"                 
##   [19,] "Bacteria" "Firmicutes"          "Negativicutes"              
##   [20,] "Bacteria" "Bacteroidetes"       "Bacteroidia"                
##   [21,] "Bacteria" "Firmicutes"          "Bacilli"                    
##   [22,] "Bacteria" "Bacteroidetes"       "Bacteroidia"                
##   [23,] "Bacteria" "Bacteroidetes"       "Bacteroidia"                
##   [24,] "Bacteria" "Firmicutes"          "Negativicutes"              
##   [25,] "Bacteria" "Proteobacteria"      "Gammaproteobacteria"        
##   [26,] "Bacteria" "Bacteroidetes"       "Bacteroidia"                
##   [27,] "Bacteria" "Bacteroidetes"       "Bacteroidia"                
##   [28,] "Bacteria" "Firmicutes"          "Negativicutes"              
##   [29,] "Bacteria" "Firmicutes"          "Bacilli"                    
##   [30,] "Bacteria" "Firmicutes"          "Bacilli"                    
##   [31,] "Bacteria" "Bacteroidetes"       "Bacteroidia"                
##   [32,] "Bacteria" "Firmicutes"          "Bacilli"                    
##   [33,] "Bacteria" "Bacteroidetes"       "Bacteroidia"                
##   [34,] "Bacteria" "Bacteroidetes"       "Bacteroidia"                
##   [35,] "Bacteria" "Firmicutes"          "Clostridia"                 
##   [36,] "Bacteria" "Firmicutes"          "Negativicutes"              
##   [37,] "Bacteria" "Bacteroidetes"       "Bacteroidia"                
##   [38,] "Bacteria" "Firmicutes"          "Negativicutes"              
##   [39,] "Bacteria" "Firmicutes"          "Clostridia"                 
##   [40,] "Bacteria" "Firmicutes"          "Bacilli"                    
##   [41,] "Bacteria" "Bacteroidetes"       "Bacteroidia"                
##   [42,] "Bacteria" "Firmicutes"          "Bacilli"                    
##   [43,] "Bacteria" "Firmicutes"          "Bacilli"                    
##   [44,] "Bacteria" "Bacteroidetes"       "Bacteroidia"                
##   [45,] "Bacteria" "Proteobacteria"      "Gammaproteobacteria"        
##   [46,] "Bacteria" "Bacteroidetes"       "Bacteroidia"                
##   [47,] "Bacteria" "Firmicutes"          "Negativicutes"              
##   [48,] "Bacteria" "Firmicutes"          "Negativicutes"              
##   [49,] "Bacteria" "Firmicutes"          "Clostridia"                 
##   [50,] "Bacteria" "Fusobacteria"        "Fusobacteriia"              
##   [51,] "Bacteria" "Firmicutes"          "Bacilli"                    
##   [52,] "Bacteria" "Proteobacteria"      "Gammaproteobacteria"        
##   [53,] "Bacteria" "Firmicutes"          "Bacilli"                    
##   [54,] "Bacteria" "Fusobacteria"        "Fusobacteriia"              
##   [55,] "Bacteria" "Bacteroidetes"       "Bacteroidia"                
##   [56,] "Bacteria" "Bacteroidetes"       "Bacteroidia"                
##   [57,] "Bacteria" "Firmicutes"          "Negativicutes"              
##   [58,] "Bacteria" "Firmicutes"          "Clostridia"                 
##   [59,] "Bacteria" "Actinobacteria"      "Coriobacteriia"             
##   [60,] "Bacteria" "Proteobacteria"      "Gammaproteobacteria"        
##   [61,] "Bacteria" "Proteobacteria"      "Gammaproteobacteria"        
##   [62,] "Bacteria" "Bacteroidetes"       "Bacteroidia"                
##   [63,] "Bacteria" "Firmicutes"          "Clostridia"                 
##   [64,] "Bacteria" "Firmicutes"          "Negativicutes"              
##   [65,] "Bacteria" "Proteobacteria"      "Gammaproteobacteria"        
##   [66,] "Bacteria" "Firmicutes"          "Negativicutes"              
##   [67,] "Bacteria" "Patescibacteria"     "Saccharimonadia"            
##   [68,] "Bacteria" "Actinobacteria"      "Actinobacteria"             
##   [69,] "Bacteria" "Proteobacteria"      "Gammaproteobacteria"        
##   [70,] "Bacteria" "Firmicutes"          "Clostridia"                 
##   [71,] "Bacteria" "Bacteroidetes"       "Bacteroidia"                
##   [72,] "Bacteria" "Actinobacteria"      "Actinobacteria"             
##   [73,] "Bacteria" "Firmicutes"          "Clostridia"                 
##   [74,] "Bacteria" "Firmicutes"          "Negativicutes"              
##   [75,] "Bacteria" "Actinobacteria"      "Actinobacteria"             
##   [76,] "Bacteria" "Proteobacteria"      "Gammaproteobacteria"        
##   [77,] "Bacteria" "Firmicutes"          "Clostridia"                 
##   [78,] "Bacteria" "Bacteroidetes"       "Bacteroidia"                
##   [79,] "Bacteria" "Firmicutes"          "Negativicutes"              
##   [80,] "Bacteria" "Firmicutes"          "Clostridia"                 
##   [81,] "Bacteria" "Proteobacteria"      "Gammaproteobacteria"        
##   [82,] "Bacteria" "Firmicutes"          "Negativicutes"              
##   [83,] "Bacteria" "Bacteroidetes"       "Bacteroidia"                
##   [84,] "Bacteria" "Bacteroidetes"       "Bacteroidia"                
##   [85,] "Bacteria" "Firmicutes"          "Negativicutes"              
##   [86,] "Bacteria" "Bacteroidetes"       "Bacteroidia"                
##   [87,] "Bacteria" "Fusobacteria"        "Fusobacteriia"              
##   [88,] "Bacteria" "Firmicutes"          "Clostridia"                 
##   [89,] "Bacteria" "Bacteroidetes"       "Bacteroidia"                
##   [90,] "Bacteria" "Firmicutes"          "Clostridia"                 
##   [91,] "Bacteria" "Actinobacteria"      "Actinobacteria"             
##   [92,] "Bacteria" "Proteobacteria"      "Gammaproteobacteria"        
##   [93,] "Bacteria" "Actinobacteria"      "Actinobacteria"             
##   [94,] "Bacteria" "Firmicutes"          "Bacilli"                    
##   [95,] "Bacteria" "Bacteroidetes"       "Bacteroidia"                
##   [96,] "Bacteria" "Proteobacteria"      "Gammaproteobacteria"        
##   [97,] "Bacteria" "Fusobacteria"        "Fusobacteriia"              
##   [98,] "Bacteria" "Firmicutes"          "Negativicutes"              
##   [99,] "Bacteria" "Fusobacteria"        "Fusobacteriia"              
##  [100,] "Bacteria" "Bacteroidetes"       "Bacteroidia"                
##  [101,] "Bacteria" "Bacteroidetes"       "Bacteroidia"                
##  [102,] "Bacteria" "Proteobacteria"      "Gammaproteobacteria"        
##  [103,] "Bacteria" "Proteobacteria"      "Gammaproteobacteria"        
##  [104,] "Bacteria" "Bacteroidetes"       "Bacteroidia"                
##  [105,] "Bacteria" "Firmicutes"          "Bacilli"                    
##  [106,] "Bacteria" "Firmicutes"          "Bacilli"                    
##  [107,] "Bacteria" "Firmicutes"          "Negativicutes"              
##  [108,] "Bacteria" "Firmicutes"          "Negativicutes"              
##  [109,] "Bacteria" "Actinobacteria"      "Actinobacteria"             
##  [110,] "Bacteria" "Proteobacteria"      "Gammaproteobacteria"        
##  [111,] "Bacteria" "Epsilonbacteraeota"  "Campylobacteria"            
##  [112,] "Bacteria" "Bacteroidetes"       "Bacteroidia"                
##  [113,] "Bacteria" "Proteobacteria"      "Gammaproteobacteria"        
##  [114,] "Bacteria" "Proteobacteria"      "Gammaproteobacteria"        
##  [115,] "Bacteria" "Patescibacteria"     "Saccharimonadia"            
##  [116,] "Bacteria" "Bacteroidetes"       "Bacteroidia"                
##  [117,] "Bacteria" "Proteobacteria"      "Gammaproteobacteria"        
##  [118,] "Bacteria" "Proteobacteria"      "Gammaproteobacteria"        
##  [119,] "Bacteria" "Proteobacteria"      "Gammaproteobacteria"        
##  [120,] "Bacteria" "Proteobacteria"      "Gammaproteobacteria"        
##  [121,] "Bacteria" "Proteobacteria"      "Gammaproteobacteria"        
##  [122,] "Bacteria" "Bacteroidetes"       "Bacteroidia"                
##  [123,] "Bacteria" "Firmicutes"          "Bacilli"                    
##  [124,] "Bacteria" "Bacteroidetes"       "Bacteroidia"                
##  [125,] "Bacteria" "Proteobacteria"      "Gammaproteobacteria"        
##  [126,] "Bacteria" "Bacteroidetes"       "Bacteroidia"                
##  [127,] "Bacteria" "Firmicutes"          "Negativicutes"              
##  [128,] "Bacteria" "Proteobacteria"      "Gammaproteobacteria"        
##  [129,] "Bacteria" "Proteobacteria"      "Gammaproteobacteria"        
##  [130,] "Bacteria" "Proteobacteria"      "Gammaproteobacteria"        
##  [131,] "Bacteria" "Proteobacteria"      "Gammaproteobacteria"        
##  [132,] "Bacteria" "Bacteroidetes"       "Bacteroidia"                
##  [133,] "Bacteria" "Patescibacteria"     "Saccharimonadia"            
##  [134,] "Bacteria" "Bacteroidetes"       "Bacteroidia"                
##  [135,] "Bacteria" "Firmicutes"          "Negativicutes"              
##  [136,] "Bacteria" "Firmicutes"          "Negativicutes"              
##  [137,] "Bacteria" "Bacteroidetes"       "Bacteroidia"                
##  [138,] "Bacteria" "Bacteroidetes"       "Bacteroidia"                
##  [139,] "Bacteria" "Bacteroidetes"       "Bacteroidia"                
##  [140,] "Bacteria" "Firmicutes"          "Bacilli"                    
##  [141,] "Bacteria" "Firmicutes"          "Negativicutes"              
##  [142,] "Bacteria" "Firmicutes"          "Negativicutes"              
##  [143,] "Bacteria" "Fusobacteria"        "Fusobacteriia"              
##  [144,] "Bacteria" "Firmicutes"          "Negativicutes"              
##  [145,] "Bacteria" "Firmicutes"          "Bacilli"                    
##  [146,] "Bacteria" "Bacteroidetes"       "Bacteroidia"                
##  [147,] "Bacteria" "Proteobacteria"      "Gammaproteobacteria"        
##  [148,] "Bacteria" "Firmicutes"          "Negativicutes"              
##  [149,] "Bacteria" "Firmicutes"          "Negativicutes"              
##  [150,] "Bacteria" "Firmicutes"          "Negativicutes"              
##  [151,] "Bacteria" "Proteobacteria"      "Gammaproteobacteria"        
##  [152,] "Bacteria" "Bacteroidetes"       "Bacteroidia"                
##  [153,] "Bacteria" "Firmicutes"          "Negativicutes"              
##  [154,] "Bacteria" "Bacteroidetes"       "Bacteroidia"                
##  [155,] "Bacteria" "Firmicutes"          "Clostridia"                 
##  [156,] "Bacteria" "Proteobacteria"      "Gammaproteobacteria"        
##  [157,] "Bacteria" "Firmicutes"          "Negativicutes"              
##  [158,] "Bacteria" "Bacteroidetes"       "Bacteroidia"                
##  [159,] "Bacteria" "Firmicutes"          "Negativicutes"              
##  [160,] "Bacteria" "Firmicutes"          "Clostridia"                 
##  [161,] "Bacteria" "Firmicutes"          "Bacilli"                    
##  [162,] "Bacteria" "Actinobacteria"      "Actinobacteria"             
##  [163,] "Bacteria" "Bacteroidetes"       "Bacteroidia"                
##  [164,] "Bacteria" "Firmicutes"          "Negativicutes"              
##  [165,] "Bacteria" "Epsilonbacteraeota"  "Campylobacteria"            
##  [166,] "Bacteria" "Bacteroidetes"       "Bacteroidia"                
##  [167,] "Bacteria" "Bacteroidetes"       "Bacteroidia"                
##  [168,] "Bacteria" "Firmicutes"          "Bacilli"                    
##  [169,] "Bacteria" "Bacteroidetes"       "Bacteroidia"                
##  [170,] "Bacteria" "Proteobacteria"      "Gammaproteobacteria"        
##  [171,] "Bacteria" "Firmicutes"          "Bacilli"                    
##  [172,] "Bacteria" "Firmicutes"          "Bacilli"                    
##  [173,] "Bacteria" "Patescibacteria"     "Saccharimonadia"            
##  [174,] "Bacteria" "Bacteroidetes"       "Bacteroidia"                
##  [175,] "Bacteria" "Bacteroidetes"       "Bacteroidia"                
##  [176,] "Bacteria" "Firmicutes"          "Clostridia"                 
##  [177,] "Bacteria" "Firmicutes"          "Negativicutes"              
##  [178,] "Bacteria" "Firmicutes"          "Negativicutes"              
##  [179,] "Bacteria" "Bacteroidetes"       "Bacteroidia"                
##  [180,] "Bacteria" "Proteobacteria"      "Gammaproteobacteria"        
##  [181,] "Bacteria" "Patescibacteria"     "Gracilibacteria"            
##  [182,] "Bacteria" "Firmicutes"          "Negativicutes"              
##  [183,] "Bacteria" "Proteobacteria"      "Gammaproteobacteria"        
##  [184,] "Bacteria" "Firmicutes"          "Negativicutes"              
##  [185,] "Bacteria" "Proteobacteria"      "Gammaproteobacteria"        
##  [186,] "Bacteria" "Firmicutes"          "Negativicutes"              
##  [187,] "Bacteria" "Firmicutes"          "Negativicutes"              
##  [188,] "Bacteria" "Bacteroidetes"       "Bacteroidia"                
##  [189,] "Bacteria" "Bacteroidetes"       "Bacteroidia"                
##  [190,] "Bacteria" "Proteobacteria"      "Gammaproteobacteria"        
##  [191,] "Bacteria" "Bacteroidetes"       "Bacteroidia"                
##  [192,] "Bacteria" "Bacteroidetes"       "Bacteroidia"                
##  [193,] "Bacteria" "Proteobacteria"      "Gammaproteobacteria"        
##  [194,] "Bacteria" "Bacteroidetes"       "Bacteroidia"                
##  [195,] "Bacteria" "Firmicutes"          "Negativicutes"              
##  [196,] "Bacteria" "Firmicutes"          "Negativicutes"              
##  [197,] "Bacteria" "Fusobacteria"        "Fusobacteriia"              
##  [198,] "Bacteria" "Firmicutes"          "Negativicutes"              
##  [199,] "Bacteria" "Bacteroidetes"       "Bacteroidia"                
##  [200,] "Bacteria" "Firmicutes"          "Negativicutes"              
##  [201,] "Bacteria" "Bacteroidetes"       "Bacteroidia"                
##  [202,] "Bacteria" "Firmicutes"          "Clostridia"                 
##  [203,] "Bacteria" "Epsilonbacteraeota"  "Campylobacteria"            
##  [204,] "Bacteria" "Bacteroidetes"       "Bacteroidia"                
##  [205,] "Bacteria" "Firmicutes"          "Negativicutes"              
##  [206,] "Bacteria" "Firmicutes"          "Negativicutes"              
##  [207,] "Bacteria" "Bacteroidetes"       "Bacteroidia"                
##  [208,] "Bacteria" "Proteobacteria"      "Gammaproteobacteria"        
##  [209,] "Bacteria" "Proteobacteria"      "Gammaproteobacteria"        
##  [210,] "Bacteria" "Firmicutes"          "Bacilli"                    
##  [211,] "Bacteria" "Firmicutes"          "Clostridia"                 
##  [212,] "Bacteria" "Firmicutes"          "Negativicutes"              
##  [213,] "Bacteria" "Proteobacteria"      "Gammaproteobacteria"        
##  [214,] "Bacteria" "Proteobacteria"      "Gammaproteobacteria"        
##  [215,] "Bacteria" "Firmicutes"          "Negativicutes"              
##  [216,] "Bacteria" "Bacteroidetes"       "Bacteroidia"                
##  [217,] "Bacteria" "Firmicutes"          "Bacilli"                    
##  [218,] "Bacteria" "Proteobacteria"      "Gammaproteobacteria"        
##  [219,] "Bacteria" "Firmicutes"          "Negativicutes"              
##  [220,] "Bacteria" "Firmicutes"          "Bacilli"                    
##  [221,] "Bacteria" "Firmicutes"          "Clostridia"                 
##  [222,] "Bacteria" "Firmicutes"          "Bacilli"                    
##  [223,] "Bacteria" "Firmicutes"          "Clostridia"                 
##  [224,] "Bacteria" "Bacteroidetes"       "Bacteroidia"                
##  [225,] "Bacteria" "Fusobacteria"        "Fusobacteriia"              
##  [226,] "Bacteria" "Firmicutes"          "Negativicutes"              
##  [227,] "Bacteria" "Bacteroidetes"       "Bacteroidia"                
##  [228,] "Bacteria" "Firmicutes"          "Bacilli"                    
##  [229,] "Bacteria" "Bacteroidetes"       "Bacteroidia"                
##  [230,] "Bacteria" "Bacteroidetes"       "Bacteroidia"                
##  [231,] "Bacteria" "Fusobacteria"        "Fusobacteriia"              
##  [232,] "Bacteria" "Fusobacteria"        "Fusobacteriia"              
##  [233,] "Bacteria" "Proteobacteria"      "Gammaproteobacteria"        
##  [234,] "Bacteria" "Fusobacteria"        "Fusobacteriia"              
##  [235,] "Bacteria" "Firmicutes"          "Negativicutes"              
##  [236,] "Bacteria" "Patescibacteria"     "Saccharimonadia"            
##  [237,] "Bacteria" "Bacteroidetes"       "Bacteroidia"                
##  [238,] "Bacteria" "Firmicutes"          "Negativicutes"              
##  [239,] "Bacteria" "Bacteroidetes"       "Bacteroidia"                
##  [240,] "Bacteria" "Firmicutes"          "Negativicutes"              
##  [241,] "Bacteria" "Fusobacteria"        "Fusobacteriia"              
##  [242,] "Bacteria" "Bacteroidetes"       "Bacteroidia"                
##  [243,] "Bacteria" "Firmicutes"          "Negativicutes"              
##  [244,] "Bacteria" "Cyanobacteria"       "Oxyphotobacteria"           
##  [245,] "Bacteria" "Bacteroidetes"       "Bacteroidia"                
##  [246,] "Bacteria" "Proteobacteria"      "Gammaproteobacteria"        
##  [247,] "Bacteria" "Proteobacteria"      "Gammaproteobacteria"        
##  [248,] "Bacteria" "Cyanobacteria"       "Oxyphotobacteria"           
##  [249,] "Bacteria" "Patescibacteria"     "Gracilibacteria"            
##  [250,] "Bacteria" "Bacteroidetes"       "Bacteroidia"                
##  [251,] "Bacteria" "Bacteroidetes"       "Bacteroidia"                
##  [252,] "Bacteria" "Bacteroidetes"       "Bacteroidia"                
##  [253,] "Bacteria" "Firmicutes"          "Clostridia"                 
##  [254,] "Bacteria" "Proteobacteria"      "Gammaproteobacteria"        
##  [255,] "Bacteria" "Proteobacteria"      "Gammaproteobacteria"        
##  [256,] "Bacteria" "Firmicutes"          "Negativicutes"              
##  [257,] "Bacteria" "Firmicutes"          "Negativicutes"              
##  [258,] "Bacteria" "Proteobacteria"      "Gammaproteobacteria"        
##  [259,] "Bacteria" "Patescibacteria"     "Saccharimonadia"            
##  [260,] "Bacteria" "Actinobacteria"      "Actinobacteria"             
##  [261,] "Bacteria" "Firmicutes"          "Clostridia"                 
##  [262,] "Bacteria" "Bacteroidetes"       "Bacteroidia"                
##  [263,] "Bacteria" "Bacteroidetes"       "Bacteroidia"                
##  [264,] "Bacteria" "Proteobacteria"      "Gammaproteobacteria"        
##  [265,] "Bacteria" "Patescibacteria"     "Saccharimonadia"            
##  [266,] "Bacteria" "Proteobacteria"      "Gammaproteobacteria"        
##  [267,] "Bacteria" "Firmicutes"          "Erysipelotrichia"           
##  [268,] "Bacteria" "Bacteroidetes"       "Bacteroidia"                
##  [269,] "Bacteria" "Firmicutes"          "Clostridia"                 
##  [270,] "Bacteria" "Proteobacteria"      "Gammaproteobacteria"        
##  [271,] "Bacteria" "Firmicutes"          "Negativicutes"              
##  [272,] "Bacteria" "Proteobacteria"      "Gammaproteobacteria"        
##  [273,] "Bacteria" "Proteobacteria"      "Gammaproteobacteria"        
##  [274,] "Bacteria" "Firmicutes"          "Negativicutes"              
##  [275,] "Bacteria" "Firmicutes"          "Bacilli"                    
##  [276,] "Bacteria" "Firmicutes"          "Negativicutes"              
##  [277,] "Bacteria" "Bacteroidetes"       "Bacteroidia"                
##  [278,] "Bacteria" "Bacteroidetes"       "Bacteroidia"                
##  [279,] "Bacteria" "Proteobacteria"      "Gammaproteobacteria"        
##  [280,] "Bacteria" "Bacteroidetes"       "Bacteroidia"                
##  [281,] "Bacteria" "Fusobacteria"        "Fusobacteriia"              
##  [282,] "Bacteria" "Firmicutes"          NA                           
##  [283,] "Bacteria" "Fusobacteria"        "Fusobacteriia"              
##  [284,] "Bacteria" "Bacteroidetes"       "Bacteroidia"                
##  [285,] "Bacteria" "Bacteroidetes"       "Bacteroidia"                
##  [286,] "Bacteria" "Patescibacteria"     "Gracilibacteria"            
##  [287,] "Bacteria" "Bacteroidetes"       "Bacteroidia"                
##  [288,] "Bacteria" "Firmicutes"          "Negativicutes"              
##  [289,] "Bacteria" "Bacteroidetes"       "Bacteroidia"                
##  [290,] "Bacteria" "Firmicutes"          "Clostridia"                 
##  [291,] "Bacteria" "Bacteroidetes"       "Bacteroidia"                
##  [292,] "Bacteria" "Proteobacteria"      "Gammaproteobacteria"        
##  [293,] "Bacteria" "Firmicutes"          "Negativicutes"              
##  [294,] "Bacteria" "Patescibacteria"     "Saccharimonadia"            
##  [295,] "Bacteria" "Firmicutes"          "Bacilli"                    
##  [296,] "Bacteria" "Firmicutes"          "Negativicutes"              
##  [297,] "Bacteria" "Firmicutes"          NA                           
##  [298,] "Bacteria" "Firmicutes"          "Negativicutes"              
##  [299,] "Bacteria" "Proteobacteria"      "Gammaproteobacteria"        
##  [300,] "Bacteria" "Bacteroidetes"       "Bacteroidia"                
##  [301,] "Bacteria" "Bacteroidetes"       "Bacteroidia"                
##  [302,] "Bacteria" "Fusobacteria"        "Fusobacteriia"              
##  [303,] "Bacteria" "Bacteroidetes"       "Bacteroidia"                
##  [304,] "Bacteria" "Fusobacteria"        "Fusobacteriia"              
##  [305,] "Bacteria" "Fusobacteria"        "Fusobacteriia"              
##  [306,] "Bacteria" "Bacteroidetes"       "Bacteroidia"                
##  [307,] "Bacteria" "Firmicutes"          "Negativicutes"              
##  [308,] "Bacteria" "Fusobacteria"        "Fusobacteriia"              
##  [309,] "Bacteria" "Firmicutes"          "Clostridia"                 
##  [310,] "Bacteria" "Proteobacteria"      "Gammaproteobacteria"        
##  [311,] "Bacteria" "Firmicutes"          "Negativicutes"              
##  [312,] "Bacteria" "Fusobacteria"        "Fusobacteriia"              
##  [313,] "Bacteria" "Patescibacteria"     "Saccharimonadia"            
##  [314,] "Bacteria" "Epsilonbacteraeota"  "Campylobacteria"            
##  [315,] "Bacteria" "Firmicutes"          "Negativicutes"              
##  [316,] "Bacteria" "Bacteroidetes"       "Bacteroidia"                
##  [317,] "Bacteria" "Firmicutes"          "Negativicutes"              
##  [318,] "Bacteria" "Firmicutes"          "Clostridia"                 
##  [319,] "Bacteria" "Firmicutes"          "Clostridia"                 
##  [320,] "Bacteria" "Bacteroidetes"       "Bacteroidia"                
##  [321,] "Bacteria" "Actinobacteria"      "Coriobacteriia"             
##  [322,] "Bacteria" "Firmicutes"          "Bacilli"                    
##  [323,] "Bacteria" "Patescibacteria"     "Saccharimonadia"            
##  [324,] "Bacteria" "Firmicutes"          "Negativicutes"              
##  [325,] "Bacteria" "Proteobacteria"      "Gammaproteobacteria"        
##  [326,] "Bacteria" "Patescibacteria"     "Saccharimonadia"            
##  [327,] "Bacteria" "Bacteroidetes"       "Bacteroidia"                
##  [328,] "Bacteria" "Actinobacteria"      "Coriobacteriia"             
##  [329,] "Bacteria" "Firmicutes"          "Negativicutes"              
##  [330,] "Bacteria" "Bacteroidetes"       "Bacteroidia"                
##  [331,] "Bacteria" "Bacteroidetes"       "Bacteroidia"                
##  [332,] "Bacteria" "Patescibacteria"     "Saccharimonadia"            
##  [333,] "Bacteria" "Firmicutes"          "Clostridia"                 
##  [334,] "Bacteria" "Firmicutes"          "Negativicutes"              
##  [335,] "Bacteria" "Firmicutes"          "Clostridia"                 
##  [336,] "Bacteria" "Fusobacteria"        "Fusobacteriia"              
##  [337,] "Bacteria" "Firmicutes"          "Bacilli"                    
##  [338,] "Bacteria" "Fusobacteria"        "Fusobacteriia"              
##  [339,] "Bacteria" "Actinobacteria"      "Actinobacteria"             
##  [340,] "Bacteria" "Fusobacteria"        "Fusobacteriia"              
##  [341,] "Bacteria" "Fusobacteria"        "Fusobacteriia"              
##  [342,] "Bacteria" "Bacteroidetes"       "Bacteroidia"                
##  [343,] "Bacteria" "Proteobacteria"      "Gammaproteobacteria"        
##  [344,] "Bacteria" "Bacteroidetes"       "Bacteroidia"                
##  [345,] "Bacteria" "Firmicutes"          "Clostridia"                 
##  [346,] "Bacteria" "Proteobacteria"      "Gammaproteobacteria"        
##  [347,] "Bacteria" "Firmicutes"          NA                           
##  [348,] "Bacteria" "Proteobacteria"      "Gammaproteobacteria"        
##  [349,] "Bacteria" "Proteobacteria"      "Gammaproteobacteria"        
##  [350,] "Bacteria" "Bacteroidetes"       "Bacteroidia"                
##  [351,] "Bacteria" "Firmicutes"          "Clostridia"                 
##  [352,] "Bacteria" "Firmicutes"          "Negativicutes"              
##  [353,] "Bacteria" "Bacteroidetes"       "Bacteroidia"                
##  [354,] "Bacteria" "Proteobacteria"      "Gammaproteobacteria"        
##  [355,] "Bacteria" "Fusobacteria"        "Fusobacteriia"              
##  [356,] "Bacteria" "Firmicutes"          "Negativicutes"              
##  [357,] "Bacteria" "Fusobacteria"        "Fusobacteriia"              
##  [358,] "Bacteria" "Bacteroidetes"       "Bacteroidia"                
##  [359,] "Bacteria" "Firmicutes"          "Negativicutes"              
##  [360,] "Bacteria" "Firmicutes"          "Clostridia"                 
##  [361,] "Bacteria" "Firmicutes"          "Clostridia"                 
##  [362,] "Bacteria" "Proteobacteria"      "Gammaproteobacteria"        
##  [363,] "Bacteria" "Bacteroidetes"       "Bacteroidia"                
##  [364,] "Bacteria" "Firmicutes"          "Clostridia"                 
##  [365,] "Bacteria" "Fusobacteria"        "Fusobacteriia"              
##  [366,] "Bacteria" "Patescibacteria"     "Saccharimonadia"            
##  [367,] "Bacteria" "Firmicutes"          "Clostridia"                 
##  [368,] "Bacteria" "Firmicutes"          "Clostridia"                 
##  [369,] "Bacteria" "Actinobacteria"      "Actinobacteria"             
##  [370,] "Bacteria" "Bacteroidetes"       "Bacteroidia"                
##  [371,] "Bacteria" "Patescibacteria"     "Saccharimonadia"            
##  [372,] "Bacteria" "Firmicutes"          "Bacilli"                    
##  [373,] "Bacteria" "Firmicutes"          "Bacilli"                    
##  [374,] "Bacteria" "Firmicutes"          "Negativicutes"              
##  [375,] "Bacteria" "Patescibacteria"     "Saccharimonadia"            
##  [376,] "Bacteria" "Firmicutes"          "Negativicutes"              
##  [377,] "Bacteria" "Fusobacteria"        "Fusobacteriia"              
##  [378,] "Bacteria" "Firmicutes"          "Bacilli"                    
##  [379,] "Bacteria" "Firmicutes"          "Negativicutes"              
##  [380,] "Bacteria" "Fusobacteria"        "Fusobacteriia"              
##  [381,] "Bacteria" "Fusobacteria"        "Fusobacteriia"              
##  [382,] "Bacteria" "Firmicutes"          "Clostridia"                 
##  [383,] "Bacteria" "Bacteroidetes"       "Bacteroidia"                
##  [384,] "Bacteria" "Bacteroidetes"       "Bacteroidia"                
##  [385,] "Bacteria" "Firmicutes"          "Bacilli"                    
##  [386,] "Bacteria" "Proteobacteria"      "Gammaproteobacteria"        
##  [387,] "Bacteria" "Fusobacteria"        "Fusobacteriia"              
##  [388,] "Bacteria" "Actinobacteria"      "Actinobacteria"             
##  [389,] "Bacteria" "Fusobacteria"        "Fusobacteriia"              
##  [390,] "Bacteria" "Bacteroidetes"       "Bacteroidia"                
##  [391,] "Bacteria" "Actinobacteria"      "Actinobacteria"             
##  [392,] "Bacteria" "Fusobacteria"        "Fusobacteriia"              
##  [393,] "Bacteria" "Proteobacteria"      "Gammaproteobacteria"        
##  [394,] "Bacteria" "Proteobacteria"      "Gammaproteobacteria"        
##  [395,] "Bacteria" "Firmicutes"          "Negativicutes"              
##  [396,] "Bacteria" "Bacteroidetes"       "Bacteroidia"                
##  [397,] "Bacteria" "Bacteroidetes"       "Bacteroidia"                
##  [398,] "Bacteria" "Proteobacteria"      "Gammaproteobacteria"        
##  [399,] "Bacteria" "Fusobacteria"        "Fusobacteriia"              
##  [400,] "Bacteria" "Proteobacteria"      "Gammaproteobacteria"        
##  [401,] "Bacteria" "Firmicutes"          "Bacilli"                    
##  [402,] "Bacteria" "Bacteroidetes"       "Bacteroidia"                
##  [403,] "Bacteria" "Bacteroidetes"       "Bacteroidia"                
##  [404,] "Bacteria" "Bacteroidetes"       "Bacteroidia"                
##  [405,] "Bacteria" "Firmicutes"          "Clostridia"                 
##  [406,] "Bacteria" "Firmicutes"          "Negativicutes"              
##  [407,] "Bacteria" "Bacteroidetes"       "Bacteroidia"                
##  [408,] "Bacteria" "Firmicutes"          "Bacilli"                    
##  [409,] "Bacteria" "Firmicutes"          "Negativicutes"              
##  [410,] "Bacteria" "Bacteroidetes"       "Bacteroidia"                
##  [411,] "Bacteria" "Bacteroidetes"       "Bacteroidia"                
##  [412,] "Bacteria" "Bacteroidetes"       "Bacteroidia"                
##  [413,] "Bacteria" "Proteobacteria"      "Gammaproteobacteria"        
##  [414,] "Bacteria" "Firmicutes"          "Clostridia"                 
##  [415,] "Bacteria" "Bacteroidetes"       "Bacteroidia"                
##  [416,] "Bacteria" "Proteobacteria"      "Gammaproteobacteria"        
##  [417,] "Bacteria" "Bacteroidetes"       "Bacteroidia"                
##  [418,] "Bacteria" "Bacteroidetes"       "Bacteroidia"                
##  [419,] "Bacteria" "Fusobacteria"        "Fusobacteriia"              
##  [420,] "Bacteria" "Bacteroidetes"       "Bacteroidia"                
##  [421,] "Bacteria" "Firmicutes"          "Bacilli"                    
##  [422,] "Bacteria" "Firmicutes"          "Negativicutes"              
##  [423,] "Bacteria" "Firmicutes"          "Negativicutes"              
##  [424,] "Bacteria" "Firmicutes"          "Negativicutes"              
##  [425,] "Bacteria" "Proteobacteria"      "Gammaproteobacteria"        
##  [426,] "Bacteria" "Bacteroidetes"       "Bacteroidia"                
##  [427,] "Bacteria" "Bacteroidetes"       "Bacteroidia"                
##  [428,] "Bacteria" "Bacteroidetes"       "Bacteroidia"                
##  [429,] "Bacteria" "Firmicutes"          "Negativicutes"              
##  [430,] "Bacteria" "Firmicutes"          "Negativicutes"              
##  [431,] "Bacteria" "Firmicutes"          "Negativicutes"              
##  [432,] "Bacteria" "Bacteroidetes"       "Bacteroidia"                
##  [433,] "Bacteria" "Firmicutes"          "Negativicutes"              
##  [434,] "Bacteria" "Bacteroidetes"       "Bacteroidia"                
##  [435,] "Bacteria" "Firmicutes"          "Clostridia"                 
##  [436,] "Bacteria" "Firmicutes"          "Negativicutes"              
##  [437,] "Bacteria" "Bacteroidetes"       "Bacteroidia"                
##  [438,] "Bacteria" "Firmicutes"          "Bacilli"                    
##  [439,] "Bacteria" "Bacteroidetes"       "Bacteroidia"                
##  [440,] "Bacteria" "Proteobacteria"      "Gammaproteobacteria"        
##  [441,] "Bacteria" "Bacteroidetes"       "Bacteroidia"                
##  [442,] "Bacteria" "Fusobacteria"        "Fusobacteriia"              
##  [443,] "Bacteria" "Firmicutes"          "Negativicutes"              
##  [444,] "Bacteria" "Firmicutes"          "Negativicutes"              
##  [445,] "Bacteria" "Firmicutes"          "Negativicutes"              
##  [446,] "Bacteria" "Firmicutes"          "Negativicutes"              
##  [447,] "Bacteria" "Firmicutes"          "Bacilli"                    
##  [448,] "Bacteria" "Fusobacteria"        "Fusobacteriia"              
##  [449,] "Bacteria" "Bacteroidetes"       "Bacteroidia"                
##  [450,] "Bacteria" "Fusobacteria"        "Fusobacteriia"              
##  [451,] "Bacteria" "Bacteroidetes"       "Bacteroidia"                
##  [452,] "Bacteria" "Epsilonbacteraeota"  "Campylobacteria"            
##  [453,] "Bacteria" "Firmicutes"          "Clostridia"                 
##  [454,] "Bacteria" "Bacteroidetes"       "Bacteroidia"                
##  [455,] "Bacteria" "Firmicutes"          "Negativicutes"              
##  [456,] "Bacteria" "Firmicutes"          "Clostridia"                 
##  [457,] "Bacteria" "Proteobacteria"      "Gammaproteobacteria"        
##  [458,] "Bacteria" "Actinobacteria"      "Actinobacteria"             
##  [459,] "Bacteria" "Fusobacteria"        "Fusobacteriia"              
##  [460,] "Bacteria" "Firmicutes"          "Negativicutes"              
##  [461,] "Bacteria" "Patescibacteria"     "Saccharimonadia"            
##  [462,] "Bacteria" "Fusobacteria"        "Fusobacteriia"              
##  [463,] "Bacteria" "Bacteroidetes"       "Bacteroidia"                
##  [464,] "Bacteria" "Firmicutes"          "Bacilli"                    
##  [465,] "Bacteria" "Bacteroidetes"       "Bacteroidia"                
##  [466,] "Bacteria" "Firmicutes"          "Negativicutes"              
##  [467,] "Bacteria" "Bacteroidetes"       "Bacteroidia"                
##  [468,] "Bacteria" "Proteobacteria"      "Gammaproteobacteria"        
##  [469,] "Bacteria" "Patescibacteria"     "Saccharimonadia"            
##  [470,] "Bacteria" "Proteobacteria"      "Gammaproteobacteria"        
##  [471,] "Bacteria" "Proteobacteria"      "Gammaproteobacteria"        
##  [472,] "Bacteria" "Bacteroidetes"       "Bacteroidia"                
##  [473,] "Bacteria" "Bacteroidetes"       "Bacteroidia"                
##  [474,] "Bacteria" "Firmicutes"          "Negativicutes"              
##  [475,] "Bacteria" "Bacteroidetes"       "Bacteroidia"                
##  [476,] "Bacteria" "Proteobacteria"      "Gammaproteobacteria"        
##  [477,] "Bacteria" "Bacteroidetes"       "Bacteroidia"                
##  [478,] "Bacteria" "Proteobacteria"      "Gammaproteobacteria"        
##  [479,] "Bacteria" "Proteobacteria"      "Gammaproteobacteria"        
##  [480,] "Bacteria" "Fusobacteria"        "Fusobacteriia"              
##  [481,] "Bacteria" "Bacteroidetes"       "Bacteroidia"                
##  [482,] "Bacteria" "Bacteroidetes"       "Bacteroidia"                
##  [483,] "Bacteria" "Fusobacteria"        "Fusobacteriia"              
##  [484,] "Bacteria" "Firmicutes"          "Negativicutes"              
##  [485,] "Bacteria" "Firmicutes"          "Bacilli"                    
##  [486,] "Bacteria" "Fusobacteria"        "Fusobacteriia"              
##  [487,] "Bacteria" "Bacteroidetes"       "Bacteroidia"                
##  [488,] "Bacteria" "Fusobacteria"        "Fusobacteriia"              
##  [489,] "Bacteria" "Bacteroidetes"       "Bacteroidia"                
##  [490,] "Bacteria" "Epsilonbacteraeota"  "Campylobacteria"            
##  [491,] "Bacteria" "Firmicutes"          "Negativicutes"              
##  [492,] "Bacteria" "Epsilonbacteraeota"  "Campylobacteria"            
##  [493,] "Bacteria" "Bacteroidetes"       "Bacteroidia"                
##  [494,] "Bacteria" "Fusobacteria"        "Fusobacteriia"              
##  [495,] "Bacteria" "Bacteroidetes"       "Bacteroidia"                
##  [496,] "Bacteria" "Firmicutes"          "Negativicutes"              
##  [497,] "Bacteria" "Proteobacteria"      "Gammaproteobacteria"        
##  [498,] "Bacteria" "Bacteroidetes"       "Bacteroidia"                
##  [499,] "Bacteria" "Firmicutes"          "Clostridia"                 
##  [500,] "Bacteria" "Firmicutes"          "Negativicutes"              
##  [501,] "Bacteria" "Firmicutes"          "Negativicutes"              
##  [502,] "Bacteria" "Actinobacteria"      "Actinobacteria"             
##  [503,] "Bacteria" "Proteobacteria"      "Gammaproteobacteria"        
##  [504,] "Bacteria" "Firmicutes"          "Negativicutes"              
##  [505,] "Bacteria" "Proteobacteria"      "Gammaproteobacteria"        
##  [506,] "Bacteria" "Fusobacteria"        "Fusobacteriia"              
##  [507,] "Bacteria" "Actinobacteria"      "Actinobacteria"             
##  [508,] "Bacteria" "Bacteroidetes"       "Bacteroidia"                
##  [509,] "Bacteria" "Bacteroidetes"       "Bacteroidia"                
##  [510,] "Bacteria" "Proteobacteria"      "Gammaproteobacteria"        
##  [511,] "Bacteria" "Firmicutes"          "Negativicutes"              
##  [512,] "Bacteria" "Firmicutes"          "Negativicutes"              
##  [513,] "Bacteria" "Proteobacteria"      "Gammaproteobacteria"        
##  [514,] "Bacteria" "Firmicutes"          "Negativicutes"              
##  [515,] "Bacteria" "Firmicutes"          "Negativicutes"              
##  [516,] "Bacteria" "Firmicutes"          "Negativicutes"              
##  [517,] "Bacteria" "Bacteroidetes"       "Bacteroidia"                
##  [518,] "Bacteria" "Proteobacteria"      "Gammaproteobacteria"        
##  [519,] "Bacteria" "Firmicutes"          "Negativicutes"              
##  [520,] "Bacteria" "Actinobacteria"      "Actinobacteria"             
##  [521,] "Bacteria" "Firmicutes"          "Bacilli"                    
##  [522,] "Bacteria" "Actinobacteria"      "Actinobacteria"             
##  [523,] "Bacteria" "Firmicutes"          "Clostridia"                 
##  [524,] "Bacteria" "Firmicutes"          "Negativicutes"              
##  [525,] "Bacteria" "Fusobacteria"        "Fusobacteriia"              
##  [526,] "Bacteria" "Bacteroidetes"       "Bacteroidia"                
##  [527,] "Bacteria" "Bacteroidetes"       "Bacteroidia"                
##  [528,] "Bacteria" "Proteobacteria"      "Gammaproteobacteria"        
##  [529,] "Bacteria" "Epsilonbacteraeota"  "Campylobacteria"            
##  [530,] "Bacteria" "Firmicutes"          "Bacilli"                    
##  [531,] "Bacteria" "Proteobacteria"      "Gammaproteobacteria"        
##  [532,] "Bacteria" "Bacteroidetes"       "Bacteroidia"                
##  [533,] "Bacteria" "Patescibacteria"     "Saccharimonadia"            
##  [534,] "Bacteria" "Firmicutes"          "Negativicutes"              
##  [535,] "Bacteria" "Bacteroidetes"       "Bacteroidia"                
##  [536,] "Bacteria" "Bacteroidetes"       "Bacteroidia"                
##  [537,] "Bacteria" "Firmicutes"          "Negativicutes"              
##  [538,] "Bacteria" "Proteobacteria"      "Gammaproteobacteria"        
##  [539,] "Bacteria" "Bacteroidetes"       "Bacteroidia"                
##  [540,] "Bacteria" "Proteobacteria"      "Gammaproteobacteria"        
##  [541,] "Bacteria" "Fusobacteria"        "Fusobacteriia"              
##  [542,] "Bacteria" "Firmicutes"          "Bacilli"                    
##  [543,] "Bacteria" "Bacteroidetes"       "Bacteroidia"                
##  [544,] "Bacteria" "Bacteroidetes"       "Bacteroidia"                
##  [545,] "Bacteria" "Firmicutes"          "Clostridia"                 
##  [546,] "Bacteria" "Bacteroidetes"       "Bacteroidia"                
##  [547,] "Bacteria" "Proteobacteria"      "Gammaproteobacteria"        
##  [548,] "Bacteria" NA                    NA                           
##  [549,] "Bacteria" "Firmicutes"          "Negativicutes"              
##  [550,] "Bacteria" "Bacteroidetes"       "Bacteroidia"                
##  [551,] "Bacteria" "Firmicutes"          "Bacilli"                    
##  [552,] "Bacteria" "Proteobacteria"      "Gammaproteobacteria"        
##  [553,] "Bacteria" "Bacteroidetes"       "Bacteroidia"                
##  [554,] "Bacteria" "Bacteroidetes"       "Bacteroidia"                
##  [555,] "Bacteria" "Bacteroidetes"       "Bacteroidia"                
##  [556,] "Bacteria" "Firmicutes"          "Negativicutes"              
##  [557,] "Bacteria" "Firmicutes"          "Negativicutes"              
##  [558,] "Bacteria" "Firmicutes"          "Bacilli"                    
##  [559,] "Bacteria" "Bacteroidetes"       "Bacteroidia"                
##  [560,] "Bacteria" "Proteobacteria"      "Gammaproteobacteria"        
##  [561,] "Bacteria" "Bacteroidetes"       "Bacteroidia"                
##  [562,] "Bacteria" "Patescibacteria"     "Saccharimonadia"            
##  [563,] "Bacteria" "Proteobacteria"      "Gammaproteobacteria"        
##  [564,] "Bacteria" "Firmicutes"          "Bacilli"                    
##  [565,] "Bacteria" "Firmicutes"          "Negativicutes"              
##  [566,] "Bacteria" "Bacteroidetes"       "Bacteroidia"                
##  [567,] "Bacteria" "Firmicutes"          "Bacilli"                    
##  [568,] "Bacteria" "Bacteroidetes"       "Bacteroidia"                
##  [569,] "Bacteria" "Bacteroidetes"       "Bacteroidia"                
##  [570,] "Bacteria" "Fusobacteria"        "Fusobacteriia"              
##  [571,] "Bacteria" "Bacteroidetes"       "Bacteroidia"                
##  [572,] "Bacteria" "Bacteroidetes"       "Bacteroidia"                
##  [573,] "Bacteria" "Proteobacteria"      "Gammaproteobacteria"        
##  [574,] "Bacteria" "Bacteroidetes"       "Bacteroidia"                
##  [575,] "Bacteria" "Bacteroidetes"       "Bacteroidia"                
##  [576,] "Bacteria" "Bacteroidetes"       "Bacteroidia"                
##  [577,] "Bacteria" "Patescibacteria"     "Saccharimonadia"            
##  [578,] "Bacteria" "Firmicutes"          "Negativicutes"              
##  [579,] "Bacteria" "Firmicutes"          "Negativicutes"              
##  [580,] "Bacteria" "Firmicutes"          "Negativicutes"              
##  [581,] "Bacteria" "Fusobacteria"        "Fusobacteriia"              
##  [582,] "Bacteria" "Proteobacteria"      "Gammaproteobacteria"        
##  [583,] "Bacteria" "Fusobacteria"        "Fusobacteriia"              
##  [584,] "Bacteria" "Bacteroidetes"       "Bacteroidia"                
##  [585,] "Bacteria" "Proteobacteria"      "Gammaproteobacteria"        
##  [586,] "Bacteria" "Fusobacteria"        "Fusobacteriia"              
##  [587,] "Bacteria" "Fusobacteria"        "Fusobacteriia"              
##  [588,] "Bacteria" "Bacteroidetes"       "Bacteroidia"                
##  [589,] "Bacteria" "Fusobacteria"        "Fusobacteriia"              
##  [590,] "Bacteria" "Bacteroidetes"       "Bacteroidia"                
##  [591,] "Bacteria" "Proteobacteria"      "Gammaproteobacteria"        
##  [592,] "Bacteria" "Bacteroidetes"       "Bacteroidia"                
##  [593,] "Bacteria" "Bacteroidetes"       "Bacteroidia"                
##  [594,] "Bacteria" "Bacteroidetes"       "Bacteroidia"                
##  [595,] "Bacteria" "Proteobacteria"      "Gammaproteobacteria"        
##  [596,] "Bacteria" "Bacteroidetes"       "Bacteroidia"                
##  [597,] "Bacteria" "Firmicutes"          "Negativicutes"              
##  [598,] "Bacteria" "Proteobacteria"      "Gammaproteobacteria"        
##  [599,] "Bacteria" "Firmicutes"          "Clostridia"                 
##  [600,] "Bacteria" "Firmicutes"          "Negativicutes"              
##  [601,] "Bacteria" "Proteobacteria"      "Gammaproteobacteria"        
##  [602,] "Bacteria" "Bacteroidetes"       "Bacteroidia"                
##  [603,] "Bacteria" "Bacteroidetes"       "Bacteroidia"                
##  [604,] "Bacteria" "Firmicutes"          "Negativicutes"              
##  [605,] "Bacteria" "Proteobacteria"      "Gammaproteobacteria"        
##  [606,] "Bacteria" "Fusobacteria"        "Fusobacteriia"              
##  [607,] "Bacteria" "Bacteroidetes"       "Bacteroidia"                
##  [608,] "Bacteria" "Patescibacteria"     "Saccharimonadia"            
##  [609,] "Bacteria" "Bacteroidetes"       "Bacteroidia"                
##  [610,] "Bacteria" "Patescibacteria"     "Saccharimonadia"            
##  [611,] "Bacteria" "Proteobacteria"      "Gammaproteobacteria"        
##  [612,] "Bacteria" "Firmicutes"          "Negativicutes"              
##  [613,] "Bacteria" "Fusobacteria"        "Fusobacteriia"              
##  [614,] "Bacteria" "Fusobacteria"        "Fusobacteriia"              
##  [615,] "Bacteria" "Bacteroidetes"       "Bacteroidia"                
##  [616,] "Bacteria" "Firmicutes"          "Negativicutes"              
##  [617,] "Bacteria" "Bacteroidetes"       "Bacteroidia"                
##  [618,] "Bacteria" "Firmicutes"          "Negativicutes"              
##  [619,] "Bacteria" "Firmicutes"          "Bacilli"                    
##  [620,] "Bacteria" "Bacteroidetes"       "Bacteroidia"                
##  [621,] "Bacteria" "Firmicutes"          "Negativicutes"              
##  [622,] "Bacteria" "Bacteroidetes"       "Bacteroidia"                
##  [623,] "Bacteria" "Firmicutes"          "Negativicutes"              
##  [624,] "Bacteria" "Bacteroidetes"       "Bacteroidia"                
##  [625,] "Bacteria" "Bacteroidetes"       "Bacteroidia"                
##  [626,] "Bacteria" "Bacteroidetes"       "Bacteroidia"                
##  [627,] "Bacteria" "Bacteroidetes"       "Bacteroidia"                
##  [628,] "Bacteria" "Fusobacteria"        "Fusobacteriia"              
##  [629,] "Bacteria" "Bacteroidetes"       "Bacteroidia"                
##  [630,] "Bacteria" "Actinobacteria"      "Actinobacteria"             
##  [631,] "Bacteria" "Firmicutes"          "Negativicutes"              
##  [632,] "Bacteria" "Proteobacteria"      "Gammaproteobacteria"        
##  [633,] "Bacteria" "Actinobacteria"      "Coriobacteriia"             
##  [634,] "Bacteria" "Firmicutes"          "Negativicutes"              
##  [635,] "Bacteria" "Patescibacteria"     "Saccharimonadia"            
##  [636,] "Bacteria" "Actinobacteria"      "Actinobacteria"             
##  [637,] "Bacteria" "Bacteroidetes"       "Bacteroidia"                
##  [638,] "Bacteria" "Proteobacteria"      "Gammaproteobacteria"        
##  [639,] "Bacteria" "Firmicutes"          "Bacilli"                    
##  [640,] "Bacteria" "Patescibacteria"     "Gracilibacteria"            
##  [641,] "Bacteria" "Firmicutes"          "Negativicutes"              
##  [642,] "Bacteria" "Actinobacteria"      "Actinobacteria"             
##  [643,] "Bacteria" "Proteobacteria"      "Gammaproteobacteria"        
##  [644,] "Bacteria" "Bacteroidetes"       "Bacteroidia"                
##  [645,] "Bacteria" "Bacteroidetes"       "Bacteroidia"                
##  [646,] "Bacteria" "Bacteroidetes"       "Bacteroidia"                
##  [647,] "Bacteria" "Actinobacteria"      "Actinobacteria"             
##  [648,] "Bacteria" "Firmicutes"          "Bacilli"                    
##  [649,] "Bacteria" "Bacteroidetes"       "Bacteroidia"                
##  [650,] "Bacteria" "Firmicutes"          "Clostridia"                 
##  [651,] "Bacteria" "Firmicutes"          "Negativicutes"              
##  [652,] "Bacteria" "Bacteroidetes"       "Bacteroidia"                
##  [653,] "Bacteria" "Firmicutes"          "Negativicutes"              
##  [654,] "Bacteria" "Patescibacteria"     "Saccharimonadia"            
##  [655,] "Bacteria" "Bacteroidetes"       "Bacteroidia"                
##  [656,] "Bacteria" "Firmicutes"          "Negativicutes"              
##  [657,] "Bacteria" "Fusobacteria"        "Fusobacteriia"              
##  [658,] "Bacteria" "Bacteroidetes"       "Bacteroidia"                
##  [659,] "Bacteria" "Firmicutes"          "Negativicutes"              
##  [660,] "Bacteria" "Fusobacteria"        "Fusobacteriia"              
##  [661,] "Bacteria" "Actinobacteria"      "Actinobacteria"             
##  [662,] "Bacteria" "Fusobacteria"        "Fusobacteriia"              
##  [663,] "Bacteria" "Fusobacteria"        "Fusobacteriia"              
##  [664,] "Bacteria" "Proteobacteria"      "Gammaproteobacteria"        
##  [665,] "Bacteria" "Fusobacteria"        "Fusobacteriia"              
##  [666,] "Bacteria" "Firmicutes"          "Negativicutes"              
##  [667,] "Bacteria" "Cyanobacteria"       "Oxyphotobacteria"           
##  [668,] "Bacteria" "Firmicutes"          "Bacilli"                    
##  [669,] "Bacteria" "Bacteroidetes"       "Bacteroidia"                
##  [670,] "Bacteria" "Bacteroidetes"       "Bacteroidia"                
##  [671,] "Bacteria" "Actinobacteria"      "Actinobacteria"             
##  [672,] "Bacteria" "Actinobacteria"      "Actinobacteria"             
##  [673,] "Bacteria" "Patescibacteria"     "Saccharimonadia"            
##  [674,] "Bacteria" "Bacteroidetes"       "Bacteroidia"                
##  [675,] "Bacteria" "Firmicutes"          "Bacilli"                    
##  [676,] "Bacteria" "Proteobacteria"      "Gammaproteobacteria"        
##  [677,] "Bacteria" "Bacteroidetes"       "Bacteroidia"                
##  [678,] "Bacteria" "Firmicutes"          "Negativicutes"              
##  [679,] "Bacteria" "Bacteroidetes"       "Bacteroidia"                
##  [680,] "Bacteria" "Firmicutes"          NA                           
##  [681,] "Bacteria" "Bacteroidetes"       "Bacteroidia"                
##  [682,] "Bacteria" "Proteobacteria"      "Gammaproteobacteria"        
##  [683,] "Bacteria" "Firmicutes"          "Negativicutes"              
##  [684,] "Bacteria" "Bacteroidetes"       "Bacteroidia"                
##  [685,] "Bacteria" "Bacteroidetes"       "Bacteroidia"                
##  [686,] "Bacteria" "Bacteroidetes"       "Bacteroidia"                
##  [687,] "Bacteria" "Patescibacteria"     "Saccharimonadia"            
##  [688,] "Bacteria" "Firmicutes"          "Bacilli"                    
##  [689,] "Bacteria" "Firmicutes"          "Negativicutes"              
##  [690,] "Bacteria" "Firmicutes"          "Negativicutes"              
##  [691,] "Bacteria" "Bacteroidetes"       "Bacteroidia"                
##  [692,] "Bacteria" "Patescibacteria"     "Saccharimonadia"            
##  [693,] "Bacteria" "Patescibacteria"     "Gracilibacteria"            
##  [694,] "Bacteria" "Bacteroidetes"       "Bacteroidia"                
##  [695,] "Bacteria" "Firmicutes"          "Negativicutes"              
##  [696,] "Bacteria" "Firmicutes"          "Bacilli"                    
##  [697,] "Bacteria" "Firmicutes"          "Negativicutes"              
##  [698,] "Bacteria" "Bacteroidetes"       "Bacteroidia"                
##  [699,] "Bacteria" "Bacteroidetes"       "Bacteroidia"                
##  [700,] "Bacteria" "Spirochaetes"        "Spirochaetia"               
##  [701,] "Bacteria" "Bacteroidetes"       "Bacteroidia"                
##  [702,] "Bacteria" "Firmicutes"          "Negativicutes"              
##  [703,] "Bacteria" "Fusobacteria"        "Fusobacteriia"              
##  [704,] "Bacteria" "Epsilonbacteraeota"  "Campylobacteria"            
##  [705,] "Bacteria" "Bacteroidetes"       "Bacteroidia"                
##  [706,] "Bacteria" "Patescibacteria"     "Saccharimonadia"            
##  [707,] "Bacteria" "Actinobacteria"      "Actinobacteria"             
##  [708,] "Bacteria" "Bacteroidetes"       "Bacteroidia"                
##  [709,] "Bacteria" "Bacteroidetes"       "Bacteroidia"                
##  [710,] "Bacteria" "Proteobacteria"      "Gammaproteobacteria"        
##  [711,] "Bacteria" "Bacteroidetes"       "Bacteroidia"                
##  [712,] "Bacteria" "Firmicutes"          "Negativicutes"              
##  [713,] "Bacteria" "Bacteroidetes"       "Bacteroidia"                
##  [714,] "Bacteria" "Bacteroidetes"       "Bacteroidia"                
##  [715,] "Bacteria" "Proteobacteria"      "Gammaproteobacteria"        
##  [716,] "Bacteria" "Patescibacteria"     "Saccharimonadia"            
##  [717,] "Bacteria" "Firmicutes"          "Bacilli"                    
##  [718,] "Bacteria" "Spirochaetes"        "Spirochaetia"               
##  [719,] "Bacteria" "Bacteroidetes"       "Bacteroidia"                
##  [720,] "Bacteria" "Proteobacteria"      "Gammaproteobacteria"        
##  [721,] "Bacteria" "Bacteroidetes"       "Bacteroidia"                
##  [722,] "Bacteria" "Bacteroidetes"       "Bacteroidia"                
##  [723,] "Bacteria" "Bacteroidetes"       "Bacteroidia"                
##  [724,] "Bacteria" "Proteobacteria"      "Gammaproteobacteria"        
##  [725,] "Bacteria" "Proteobacteria"      "Gammaproteobacteria"        
##  [726,] "Bacteria" "Proteobacteria"      "Gammaproteobacteria"        
##  [727,] "Bacteria" "Proteobacteria"      "Gammaproteobacteria"        
##  [728,] "Bacteria" "Firmicutes"          "Negativicutes"              
##  [729,] "Bacteria" "Firmicutes"          "Negativicutes"              
##  [730,] "Bacteria" "Bacteroidetes"       "Bacteroidia"                
##  [731,] "Bacteria" "Proteobacteria"      "Gammaproteobacteria"        
##  [732,] "Bacteria" "Fusobacteria"        "Fusobacteriia"              
##  [733,] "Bacteria" "Proteobacteria"      "Gammaproteobacteria"        
##  [734,] "Bacteria" "Firmicutes"          "Negativicutes"              
##  [735,] "Bacteria" "Proteobacteria"      "Gammaproteobacteria"        
##  [736,] "Bacteria" "Bacteroidetes"       "Bacteroidia"                
##  [737,] "Bacteria" "Proteobacteria"      "Gammaproteobacteria"        
##  [738,] "Bacteria" "Actinobacteria"      "Actinobacteria"             
##  [739,] "Bacteria" "Proteobacteria"      "Gammaproteobacteria"        
##  [740,] "Bacteria" "Bacteroidetes"       "Bacteroidia"                
##  [741,] "Bacteria" "Fusobacteria"        "Fusobacteriia"              
##  [742,] "Bacteria" "Firmicutes"          "Bacilli"                    
##  [743,] "Bacteria" "Actinobacteria"      "Actinobacteria"             
##  [744,] "Bacteria" "Firmicutes"          "Bacilli"                    
##  [745,] "Bacteria" "Firmicutes"          "Negativicutes"              
##  [746,] "Bacteria" "Firmicutes"          "Negativicutes"              
##  [747,] "Bacteria" "Bacteroidetes"       "Bacteroidia"                
##  [748,] "Bacteria" "Patescibacteria"     "Saccharimonadia"            
##  [749,] "Bacteria" "Fusobacteria"        "Fusobacteriia"              
##  [750,] "Bacteria" "Proteobacteria"      "Gammaproteobacteria"        
##  [751,] "Bacteria" "Bacteroidetes"       "Bacteroidia"                
##  [752,] "Bacteria" "Bacteroidetes"       "Bacteroidia"                
##  [753,] "Bacteria" "Bacteroidetes"       "Bacteroidia"                
##  [754,] "Bacteria" "Bacteroidetes"       "Bacteroidia"                
##  [755,] "Bacteria" "Firmicutes"          "Negativicutes"              
##  [756,] "Bacteria" "Firmicutes"          "Negativicutes"              
##  [757,] "Bacteria" "Bacteroidetes"       "Bacteroidia"                
##  [758,] "Bacteria" "Fusobacteria"        "Fusobacteriia"              
##  [759,] "Bacteria" "Firmicutes"          "Negativicutes"              
##  [760,] "Bacteria" "Bacteroidetes"       "Bacteroidia"                
##  [761,] "Bacteria" "Firmicutes"          "Negativicutes"              
##  [762,] "Bacteria" "Bacteroidetes"       "Bacteroidia"                
##  [763,] "Bacteria" "Bacteroidetes"       "Bacteroidia"                
##  [764,] "Bacteria" "Firmicutes"          "Negativicutes"              
##  [765,] "Bacteria" "Fusobacteria"        "Fusobacteriia"              
##  [766,] "Bacteria" "Firmicutes"          "Negativicutes"              
##  [767,] "Bacteria" "Fusobacteria"        "Fusobacteriia"              
##  [768,] "Bacteria" "Fusobacteria"        "Fusobacteriia"              
##  [769,] "Bacteria" "Bacteroidetes"       "Bacteroidia"                
##  [770,] "Bacteria" "Bacteroidetes"       "Bacteroidia"                
##  [771,] "Bacteria" "Firmicutes"          "Bacilli"                    
##  [772,] "Bacteria" "Fusobacteria"        "Fusobacteriia"              
##  [773,] "Bacteria" "Bacteroidetes"       "Bacteroidia"                
##  [774,] "Bacteria" "Firmicutes"          "Bacilli"                    
##  [775,] "Bacteria" "Actinobacteria"      "Actinobacteria"             
##  [776,] "Bacteria" "Firmicutes"          "Negativicutes"              
##  [777,] "Bacteria" "Firmicutes"          "Negativicutes"              
##  [778,] "Bacteria" "Bacteroidetes"       "Bacteroidia"                
##  [779,] "Bacteria" "Proteobacteria"      "Gammaproteobacteria"        
##  [780,] "Bacteria" "Firmicutes"          "Clostridia"                 
##  [781,] "Bacteria" "Bacteroidetes"       "Bacteroidia"                
##  [782,] "Bacteria" "Proteobacteria"      "Gammaproteobacteria"        
##  [783,] "Bacteria" "Bacteroidetes"       "Bacteroidia"                
##  [784,] "Bacteria" "Proteobacteria"      "Gammaproteobacteria"        
##  [785,] "Bacteria" "Firmicutes"          "Clostridia"                 
##  [786,] "Bacteria" "Firmicutes"          "Bacilli"                    
##  [787,] "Bacteria" "Bacteroidetes"       "Bacteroidia"                
##  [788,] "Bacteria" "Bacteroidetes"       "Bacteroidia"                
##  [789,] "Bacteria" "Proteobacteria"      "Gammaproteobacteria"        
##  [790,] "Bacteria" "Firmicutes"          "Clostridia"                 
##  [791,] "Bacteria" "Firmicutes"          "Negativicutes"              
##  [792,] "Bacteria" "Firmicutes"          "Clostridia"                 
##  [793,] "Bacteria" "Firmicutes"          "Negativicutes"              
##  [794,] "Bacteria" "Fusobacteria"        "Fusobacteriia"              
##  [795,] "Bacteria" "Epsilonbacteraeota"  "Campylobacteria"            
##  [796,] "Bacteria" "Bacteroidetes"       "Bacteroidia"                
##  [797,] "Bacteria" "Firmicutes"          NA                           
##  [798,] "Bacteria" "Firmicutes"          "Negativicutes"              
##  [799,] "Bacteria" "Firmicutes"          "Negativicutes"              
##  [800,] "Bacteria" "Proteobacteria"      "Gammaproteobacteria"        
##  [801,] "Bacteria" "Bacteroidetes"       "Bacteroidia"                
##  [802,] "Bacteria" "Fusobacteria"        "Fusobacteriia"              
##  [803,] "Bacteria" "Bacteroidetes"       "Bacteroidia"                
##  [804,] "Bacteria" "Bacteroidetes"       "Bacteroidia"                
##  [805,] "Bacteria" "Actinobacteria"      "Actinobacteria"             
##  [806,] "Bacteria" "Firmicutes"          "Clostridia"                 
##  [807,] "Bacteria" "Proteobacteria"      "Gammaproteobacteria"        
##  [808,] "Bacteria" "Firmicutes"          "Bacilli"                    
##  [809,] "Bacteria" "Bacteroidetes"       "Bacteroidia"                
##  [810,] "Bacteria" "Patescibacteria"     "Saccharimonadia"            
##  [811,] "Bacteria" "Bacteroidetes"       "Bacteroidia"                
##  [812,] "Bacteria" "Actinobacteria"      "Actinobacteria"             
##  [813,] "Bacteria" "Fusobacteria"        "Fusobacteriia"              
##  [814,] "Bacteria" "Firmicutes"          "Negativicutes"              
##  [815,] "Bacteria" "Bacteroidetes"       "Bacteroidia"                
##  [816,] "Bacteria" "Proteobacteria"      "Gammaproteobacteria"        
##  [817,] "Bacteria" "Bacteroidetes"       "Bacteroidia"                
##  [818,] "Bacteria" "Firmicutes"          "Negativicutes"              
##  [819,] "Bacteria" "Bacteroidetes"       "Bacteroidia"                
##  [820,] "Bacteria" "Patescibacteria"     "Saccharimonadia"            
##  [821,] "Bacteria" "Proteobacteria"      "Gammaproteobacteria"        
##  [822,] "Bacteria" "Fusobacteria"        "Fusobacteriia"              
##  [823,] "Bacteria" "Firmicutes"          "Bacilli"                    
##  [824,] "Bacteria" "Bacteroidetes"       "Bacteroidia"                
##  [825,] "Bacteria" "Fusobacteria"        "Fusobacteriia"              
##  [826,] "Bacteria" "Bacteroidetes"       "Bacteroidia"                
##  [827,] "Bacteria" "Bacteroidetes"       "Bacteroidia"                
##  [828,] "Bacteria" "Proteobacteria"      "Gammaproteobacteria"        
##  [829,] "Bacteria" "Bacteroidetes"       "Bacteroidia"                
##  [830,] "Bacteria" "Proteobacteria"      "Gammaproteobacteria"        
##  [831,] "Bacteria" "Firmicutes"          "Negativicutes"              
##  [832,] "Bacteria" "Bacteroidetes"       "Bacteroidia"                
##  [833,] "Bacteria" "Firmicutes"          "Clostridia"                 
##  [834,] "Bacteria" "Actinobacteria"      "Actinobacteria"             
##  [835,] "Bacteria" "Firmicutes"          "Negativicutes"              
##  [836,] "Bacteria" "Firmicutes"          "Negativicutes"              
##  [837,] "Bacteria" "Firmicutes"          "Bacilli"                    
##  [838,] "Bacteria" "Bacteroidetes"       "Bacteroidia"                
##  [839,] "Bacteria" "Epsilonbacteraeota"  "Campylobacteria"            
##  [840,] "Bacteria" "Patescibacteria"     "Saccharimonadia"            
##  [841,] "Bacteria" "Firmicutes"          "Negativicutes"              
##  [842,] "Bacteria" "Bacteroidetes"       "Bacteroidia"                
##  [843,] "Bacteria" "Firmicutes"          "Clostridia"                 
##  [844,] "Bacteria" "Bacteroidetes"       "Bacteroidia"                
##  [845,] "Bacteria" "Firmicutes"          "Bacilli"                    
##  [846,] "Bacteria" "Actinobacteria"      "Actinobacteria"             
##  [847,] "Bacteria" "Firmicutes"          "Negativicutes"              
##  [848,] "Bacteria" "Actinobacteria"      "Actinobacteria"             
##  [849,] "Bacteria" "Bacteroidetes"       "Bacteroidia"                
##  [850,] "Bacteria" "Proteobacteria"      "Gammaproteobacteria"        
##  [851,] "Bacteria" "Bacteroidetes"       "Bacteroidia"                
##  [852,] "Bacteria" "Firmicutes"          "Bacilli"                    
##  [853,] "Bacteria" "Bacteroidetes"       "Bacteroidia"                
##  [854,] "Bacteria" "Firmicutes"          "Negativicutes"              
##  [855,] "Bacteria" "Bacteroidetes"       "Bacteroidia"                
##  [856,] "Bacteria" "Firmicutes"          "Clostridia"                 
##  [857,] "Bacteria" "Bacteroidetes"       "Bacteroidia"                
##  [858,] "Bacteria" "Firmicutes"          "Bacilli"                    
##  [859,] "Bacteria" "Bacteroidetes"       "Bacteroidia"                
##  [860,] "Bacteria" "Patescibacteria"     "Saccharimonadia"            
##  [861,] "Bacteria" "Firmicutes"          "Clostridia"                 
##  [862,] "Bacteria" "Firmicutes"          "Clostridia"                 
##  [863,] "Bacteria" "Bacteroidetes"       "Bacteroidia"                
##  [864,] "Bacteria" "Bacteroidetes"       "Bacteroidia"                
##  [865,] "Bacteria" "Bacteroidetes"       "Bacteroidia"                
##  [866,] "Bacteria" "Bacteroidetes"       "Bacteroidia"                
##  [867,] "Bacteria" "Actinobacteria"      "Actinobacteria"             
##  [868,] "Bacteria" "Bacteroidetes"       "Bacteroidia"                
##  [869,] "Bacteria" "Firmicutes"          "Negativicutes"              
##  [870,] "Bacteria" "Actinobacteria"      "Actinobacteria"             
##  [871,] "Bacteria" "Actinobacteria"      "Actinobacteria"             
##  [872,] "Bacteria" "Firmicutes"          "Negativicutes"              
##  [873,] "Bacteria" "Bacteroidetes"       "Bacteroidia"                
##  [874,] "Bacteria" "Bacteroidetes"       "Bacteroidia"                
##  [875,] "Bacteria" "Actinobacteria"      "Actinobacteria"             
##  [876,] "Bacteria" "Fusobacteria"        "Fusobacteriia"              
##  [877,] "Bacteria" "Patescibacteria"     "Saccharimonadia"            
##  [878,] "Bacteria" "Firmicutes"          "Negativicutes"              
##  [879,] "Bacteria" "Actinobacteria"      "Actinobacteria"             
##  [880,] "Bacteria" "Firmicutes"          "Negativicutes"              
##  [881,] "Bacteria" "Proteobacteria"      "Gammaproteobacteria"        
##  [882,] "Bacteria" "Firmicutes"          "Clostridia"                 
##  [883,] "Bacteria" "Firmicutes"          "Negativicutes"              
##  [884,] "Bacteria" "Bacteroidetes"       "Bacteroidia"                
##  [885,] "Bacteria" "Bacteroidetes"       "Bacteroidia"                
##  [886,] "Bacteria" "Bacteroidetes"       "Bacteroidia"                
##  [887,] "Bacteria" "Firmicutes"          "Negativicutes"              
##  [888,] "Bacteria" "Actinobacteria"      "Actinobacteria"             
##  [889,] "Bacteria" "Actinobacteria"      "Actinobacteria"             
##  [890,] "Bacteria" "Firmicutes"          "Clostridia"                 
##  [891,] "Bacteria" "Proteobacteria"      "Gammaproteobacteria"        
##  [892,] "Bacteria" "Firmicutes"          "Bacilli"                    
##  [893,] "Bacteria" "Patescibacteria"     "Saccharimonadia"            
##  [894,] "Bacteria" "Bacteroidetes"       "Bacteroidia"                
##  [895,] "Bacteria" "Actinobacteria"      "Actinobacteria"             
##  [896,] "Bacteria" "Bacteroidetes"       "Bacteroidia"                
##  [897,] "Bacteria" "Bacteroidetes"       "Bacteroidia"                
##  [898,] "Bacteria" "Bacteroidetes"       "Bacteroidia"                
##  [899,] "Bacteria" "Patescibacteria"     "Saccharimonadia"            
##  [900,] "Bacteria" "Firmicutes"          NA                           
##  [901,] "Bacteria" "Fusobacteria"        "Fusobacteriia"              
##  [902,] "Bacteria" "Proteobacteria"      "Gammaproteobacteria"        
##  [903,] "Bacteria" "Firmicutes"          "Bacilli"                    
##  [904,] "Bacteria" "Firmicutes"          "Clostridia"                 
##  [905,] "Bacteria" "Actinobacteria"      "Actinobacteria"             
##  [906,] "Bacteria" "Bacteroidetes"       "Bacteroidia"                
##  [907,] "Bacteria" "Firmicutes"          "Bacilli"                    
##  [908,] "Bacteria" "Proteobacteria"      "Gammaproteobacteria"        
##  [909,] "Bacteria" "Firmicutes"          "Negativicutes"              
##  [910,] "Bacteria" "Proteobacteria"      "Gammaproteobacteria"        
##  [911,] "Bacteria" "Firmicutes"          "Negativicutes"              
##  [912,] "Bacteria" "Patescibacteria"     "Saccharimonadia"            
##  [913,] "Bacteria" "Proteobacteria"      "Gammaproteobacteria"        
##  [914,] "Bacteria" "Proteobacteria"      "Gammaproteobacteria"        
##  [915,] "Bacteria" "Actinobacteria"      "Actinobacteria"             
##  [916,] "Bacteria" "Firmicutes"          "Negativicutes"              
##  [917,] "Bacteria" "Bacteroidetes"       "Bacteroidia"                
##  [918,] "Bacteria" "Firmicutes"          "Clostridia"                 
##  [919,] "Bacteria" "Actinobacteria"      "Actinobacteria"             
##  [920,] "Bacteria" "Proteobacteria"      "Gammaproteobacteria"        
##  [921,] "Bacteria" "Epsilonbacteraeota"  "Campylobacteria"            
##  [922,] "Bacteria" "Firmicutes"          "Negativicutes"              
##  [923,] "Bacteria" "Firmicutes"          "Negativicutes"              
##  [924,] "Bacteria" "Actinobacteria"      "Actinobacteria"             
##  [925,] "Bacteria" "Fusobacteria"        "Fusobacteriia"              
##  [926,] "Bacteria" "Proteobacteria"      "Gammaproteobacteria"        
##  [927,] "Bacteria" "Fusobacteria"        "Fusobacteriia"              
##  [928,] "Bacteria" "Fusobacteria"        "Fusobacteriia"              
##  [929,] "Bacteria" "Firmicutes"          "Negativicutes"              
##  [930,] "Bacteria" "Bacteroidetes"       "Bacteroidia"                
##  [931,] "Bacteria" "Fusobacteria"        "Fusobacteriia"              
##  [932,] "Bacteria" "Firmicutes"          "Bacilli"                    
##  [933,] "Bacteria" "Firmicutes"          NA                           
##  [934,] "Bacteria" "Firmicutes"          "Negativicutes"              
##  [935,] "Bacteria" "Bacteroidetes"       "Bacteroidia"                
##  [936,] "Bacteria" "Firmicutes"          "Clostridia"                 
##  [937,] "Bacteria" "Fusobacteria"        "Fusobacteriia"              
##  [938,] "Bacteria" "Firmicutes"          "Erysipelotrichia"           
##  [939,] "Bacteria" "Firmicutes"          "Bacilli"                    
##  [940,] "Bacteria" "Bacteroidetes"       "Bacteroidia"                
##  [941,] "Bacteria" "Firmicutes"          "Bacilli"                    
##  [942,] "Bacteria" "Actinobacteria"      "Actinobacteria"             
##  [943,] "Bacteria" "Epsilonbacteraeota"  "Campylobacteria"            
##  [944,] "Bacteria" "Spirochaetes"        "Spirochaetia"               
##  [945,] "Bacteria" "Firmicutes"          "Bacilli"                    
##  [946,] "Bacteria" "Actinobacteria"      "Actinobacteria"             
##  [947,] "Bacteria" "Firmicutes"          "Bacilli"                    
##  [948,] "Bacteria" "Fusobacteria"        "Fusobacteriia"              
##  [949,] "Bacteria" "Fusobacteria"        "Fusobacteriia"              
##  [950,] "Bacteria" "Firmicutes"          "Clostridia"                 
##  [951,] "Bacteria" "Fusobacteria"        "Fusobacteriia"              
##  [952,] "Bacteria" "Firmicutes"          "Negativicutes"              
##  [953,] "Bacteria" "Firmicutes"          NA                           
##  [954,] "Bacteria" "Spirochaetes"        "Spirochaetia"               
##  [955,] "Bacteria" "Bacteroidetes"       "Bacteroidia"                
##  [956,] "Bacteria" "Actinobacteria"      "Actinobacteria"             
##  [957,] "Bacteria" "Firmicutes"          "Negativicutes"              
##  [958,] "Bacteria" "Patescibacteria"     "Saccharimonadia"            
##  [959,] "Bacteria" "Firmicutes"          "Bacilli"                    
##  [960,] "Bacteria" "Bacteroidetes"       "Bacteroidia"                
##  [961,] "Bacteria" "Firmicutes"          "Negativicutes"              
##  [962,] "Bacteria" "Firmicutes"          "Bacilli"                    
##  [963,] "Bacteria" "Firmicutes"          "Negativicutes"              
##  [964,] "Bacteria" "Proteobacteria"      "Gammaproteobacteria"        
##  [965,] "Bacteria" "Fusobacteria"        "Fusobacteriia"              
##  [966,] "Bacteria" "Firmicutes"          "Clostridia"                 
##  [967,] "Bacteria" "Bacteroidetes"       "Bacteroidia"                
##  [968,] "Bacteria" "Fusobacteria"        "Fusobacteriia"              
##  [969,] "Bacteria" "Firmicutes"          "Clostridia"                 
##  [970,] "Bacteria" "Bacteroidetes"       "Bacteroidia"                
##  [971,] "Bacteria" "Fusobacteria"        "Fusobacteriia"              
##  [972,] "Bacteria" "Firmicutes"          "Negativicutes"              
##  [973,] "Bacteria" "Patescibacteria"     "Saccharimonadia"            
##  [974,] "Bacteria" "Firmicutes"          "Negativicutes"              
##  [975,] "Bacteria" "Bacteroidetes"       "Bacteroidia"                
##  [976,] "Bacteria" "Firmicutes"          "Clostridia"                 
##  [977,] "Bacteria" "Firmicutes"          "Clostridia"                 
##  [978,] "Bacteria" "Firmicutes"          "Negativicutes"              
##  [979,] "Bacteria" "Actinobacteria"      "Actinobacteria"             
##  [980,] "Bacteria" "Actinobacteria"      "Actinobacteria"             
##  [981,] "Bacteria" "Fusobacteria"        "Fusobacteriia"              
##  [982,] "Bacteria" "Bacteroidetes"       "Bacteroidia"                
##  [983,] "Bacteria" "Fusobacteria"        "Fusobacteriia"              
##  [984,] "Bacteria" "Actinobacteria"      "Actinobacteria"             
##  [985,] "Bacteria" "Epsilonbacteraeota"  "Campylobacteria"            
##  [986,] "Bacteria" "Firmicutes"          "Negativicutes"              
##  [987,] "Bacteria" "Actinobacteria"      "Actinobacteria"             
##  [988,] "Bacteria" "Proteobacteria"      "Gammaproteobacteria"        
##  [989,] "Bacteria" "Proteobacteria"      "Gammaproteobacteria"        
##  [990,] "Bacteria" "Firmicutes"          "Negativicutes"              
##  [991,] "Bacteria" "Bacteroidetes"       "Bacteroidia"                
##  [992,] "Bacteria" "Actinobacteria"      "Actinobacteria"             
##  [993,] "Bacteria" "Actinobacteria"      "Actinobacteria"             
##  [994,] "Bacteria" "Firmicutes"          "Negativicutes"              
##  [995,] "Bacteria" "Firmicutes"          "Negativicutes"              
##  [996,] "Bacteria" "Fusobacteria"        "Fusobacteriia"              
##  [997,] "Bacteria" "Fusobacteria"        "Fusobacteriia"              
##  [998,] "Bacteria" "Patescibacteria"     "Saccharimonadia"            
##  [999,] "Bacteria" "Bacteroidetes"       "Bacteroidia"                
## [1000,] "Bacteria" "Firmicutes"          "Negativicutes"              
## [1001,] "Bacteria" "Firmicutes"          "Negativicutes"              
## [1002,] "Bacteria" "Firmicutes"          "Negativicutes"              
## [1003,] "Bacteria" "Proteobacteria"      "Gammaproteobacteria"        
## [1004,] "Bacteria" "Bacteroidetes"       "Bacteroidia"                
## [1005,] "Bacteria" "Bacteroidetes"       "Bacteroidia"                
## [1006,] "Bacteria" "Bacteroidetes"       "Bacteroidia"                
## [1007,] "Bacteria" "Firmicutes"          "Clostridia"                 
## [1008,] "Bacteria" "Bacteroidetes"       "Bacteroidia"                
## [1009,] "Bacteria" "Bacteroidetes"       "Bacteroidia"                
## [1010,] "Bacteria" "Firmicutes"          "Clostridia"                 
## [1011,] "Bacteria" "Firmicutes"          "Negativicutes"              
## [1012,] "Bacteria" "Patescibacteria"     "Saccharimonadia"            
## [1013,] "Bacteria" "Proteobacteria"      "Gammaproteobacteria"        
## [1014,] "Bacteria" "Patescibacteria"     "Saccharimonadia"            
## [1015,] "Bacteria" "Firmicutes"          "Clostridia"                 
## [1016,] "Bacteria" "Proteobacteria"      "Gammaproteobacteria"        
## [1017,] "Bacteria" "Patescibacteria"     "Saccharimonadia"            
## [1018,] "Bacteria" "Bacteroidetes"       "Bacteroidia"                
## [1019,] "Bacteria" "Fusobacteria"        "Fusobacteriia"              
## [1020,] "Bacteria" "Proteobacteria"      "Gammaproteobacteria"        
## [1021,] "Bacteria" "Bacteroidetes"       "Bacteroidia"                
## [1022,] "Bacteria" "Fusobacteria"        "Fusobacteriia"              
## [1023,] "Bacteria" "Spirochaetes"        "Spirochaetia"               
## [1024,] "Bacteria" "Firmicutes"          "Bacilli"                    
## [1025,] "Bacteria" "Firmicutes"          "Bacilli"                    
## [1026,] "Bacteria" "Actinobacteria"      "Actinobacteria"             
## [1027,] "Bacteria" "Fusobacteria"        "Fusobacteriia"              
## [1028,] "Bacteria" "Bacteroidetes"       "Bacteroidia"                
## [1029,] "Bacteria" "Bacteroidetes"       "Bacteroidia"                
## [1030,] "Bacteria" "Proteobacteria"      "Gammaproteobacteria"        
## [1031,] "Bacteria" "Fusobacteria"        "Fusobacteriia"              
## [1032,] "Bacteria" "Fusobacteria"        "Fusobacteriia"              
## [1033,] "Bacteria" "Fusobacteria"        "Fusobacteriia"              
## [1034,] "Bacteria" "Firmicutes"          "Negativicutes"              
## [1035,] "Bacteria" "Bacteroidetes"       "Bacteroidia"                
## [1036,] "Bacteria" "Firmicutes"          "Negativicutes"              
## [1037,] "Bacteria" "Firmicutes"          "Negativicutes"              
## [1038,] "Bacteria" "Firmicutes"          "Clostridia"                 
## [1039,] "Bacteria" "Bacteroidetes"       "Bacteroidia"                
## [1040,] "Bacteria" "Firmicutes"          "Negativicutes"              
## [1041,] "Bacteria" "Bacteroidetes"       "Bacteroidia"                
## [1042,] "Bacteria" "Firmicutes"          "Clostridia"                 
## [1043,] "Bacteria" "Fusobacteria"        "Fusobacteriia"              
## [1044,] "Bacteria" "Firmicutes"          "Negativicutes"              
## [1045,] "Bacteria" "Actinobacteria"      "Actinobacteria"             
## [1046,] "Bacteria" "Bacteroidetes"       "Bacteroidia"                
## [1047,] "Bacteria" "Firmicutes"          "Negativicutes"              
## [1048,] "Bacteria" "Bacteroidetes"       "Bacteroidia"                
## [1049,] "Bacteria" "Firmicutes"          "Negativicutes"              
## [1050,] "Bacteria" "Fusobacteria"        "Fusobacteriia"              
## [1051,] "Bacteria" "Firmicutes"          "Clostridia"                 
## [1052,] "Bacteria" "Firmicutes"          "Bacilli"                    
## [1053,] "Bacteria" "Bacteroidetes"       "Bacteroidia"                
## [1054,] "Bacteria" "Proteobacteria"      "Gammaproteobacteria"        
## [1055,] "Bacteria" "Bacteroidetes"       "Bacteroidia"                
## [1056,] "Bacteria" "Firmicutes"          "Bacilli"                    
## [1057,] "Bacteria" "Proteobacteria"      "Gammaproteobacteria"        
## [1058,] "Bacteria" "Bacteroidetes"       "Bacteroidia"                
## [1059,] "Bacteria" "Firmicutes"          "Clostridia"                 
## [1060,] "Bacteria" "Bacteroidetes"       "Bacteroidia"                
## [1061,] "Bacteria" "Firmicutes"          "Negativicutes"              
## [1062,] "Bacteria" "Firmicutes"          "Bacilli"                    
## [1063,] "Bacteria" "Bacteroidetes"       "Bacteroidia"                
## [1064,] "Bacteria" "Fusobacteria"        "Fusobacteriia"              
## [1065,] "Bacteria" "Firmicutes"          "Negativicutes"              
## [1066,] "Bacteria" "Fusobacteria"        "Fusobacteriia"              
## [1067,] "Bacteria" "Firmicutes"          "Negativicutes"              
## [1068,] "Bacteria" "Proteobacteria"      "Gammaproteobacteria"        
## [1069,] "Bacteria" "Fusobacteria"        "Fusobacteriia"              
## [1070,] "Bacteria" "Firmicutes"          "Negativicutes"              
## [1071,] "Bacteria" "Proteobacteria"      "Gammaproteobacteria"        
## [1072,] "Bacteria" "Bacteroidetes"       "Bacteroidia"                
## [1073,] "Bacteria" "Bacteroidetes"       "Bacteroidia"                
## [1074,] "Bacteria" "Actinobacteria"      "Coriobacteriia"             
## [1075,] "Bacteria" "Bacteroidetes"       "Bacteroidia"                
## [1076,] "Bacteria" "Proteobacteria"      "Gammaproteobacteria"        
## [1077,] "Bacteria" NA                    NA                           
## [1078,] "Bacteria" "Bacteroidetes"       "Bacteroidia"                
## [1079,] "Bacteria" "Firmicutes"          "Negativicutes"              
## [1080,] "Bacteria" "Firmicutes"          "Clostridia"                 
## [1081,] "Bacteria" "Bacteroidetes"       "Bacteroidia"                
## [1082,] "Bacteria" "Patescibacteria"     "Saccharimonadia"            
## [1083,] "Bacteria" "Bacteroidetes"       "Bacteroidia"                
## [1084,] "Bacteria" "Firmicutes"          "Negativicutes"              
## [1085,] "Bacteria" "Firmicutes"          "Negativicutes"              
## [1086,] "Bacteria" "Firmicutes"          "Bacilli"                    
## [1087,] "Bacteria" "Firmicutes"          "Bacilli"                    
## [1088,] "Bacteria" "Bacteroidetes"       "Bacteroidia"                
## [1089,] "Bacteria" "Firmicutes"          "Negativicutes"              
## [1090,] "Bacteria" "Bacteroidetes"       "Bacteroidia"                
## [1091,] "Bacteria" "Actinobacteria"      "Coriobacteriia"             
## [1092,] "Bacteria" "Bacteroidetes"       "Bacteroidia"                
## [1093,] "Bacteria" "Bacteroidetes"       "Bacteroidia"                
## [1094,] "Bacteria" "Bacteroidetes"       "Bacteroidia"                
## [1095,] "Bacteria" "Bacteroidetes"       "Bacteroidia"                
## [1096,] "Bacteria" "Firmicutes"          "Negativicutes"              
## [1097,] "Bacteria" "Bacteroidetes"       "Bacteroidia"                
## [1098,] "Bacteria" "Firmicutes"          "Bacilli"                    
## [1099,] "Bacteria" "Actinobacteria"      "Actinobacteria"             
## [1100,] "Bacteria" "Bacteroidetes"       "Bacteroidia"                
## [1101,] "Bacteria" "Patescibacteria"     "Saccharimonadia"            
## [1102,] "Bacteria" "Firmicutes"          "Negativicutes"              
## [1103,] "Bacteria" "Firmicutes"          "Negativicutes"              
## [1104,] "Bacteria" "Bacteroidetes"       "Bacteroidia"                
## [1105,] "Bacteria" "Bacteroidetes"       "Bacteroidia"                
## [1106,] "Bacteria" "Firmicutes"          "Negativicutes"              
## [1107,] "Bacteria" "Bacteroidetes"       "Bacteroidia"                
## [1108,] "Bacteria" "Fusobacteria"        "Fusobacteriia"              
## [1109,] "Bacteria" "Bacteroidetes"       "Bacteroidia"                
## [1110,] "Bacteria" "Firmicutes"          "Negativicutes"              
## [1111,] "Bacteria" "Bacteroidetes"       "Bacteroidia"                
## [1112,] "Bacteria" "Fusobacteria"        "Fusobacteriia"              
## [1113,] "Bacteria" "Bacteroidetes"       "Bacteroidia"                
## [1114,] "Bacteria" "Firmicutes"          "Clostridia"                 
## [1115,] "Bacteria" "Patescibacteria"     "Saccharimonadia"            
## [1116,] "Bacteria" "Patescibacteria"     "Saccharimonadia"            
## [1117,] "Bacteria" "Bacteroidetes"       "Bacteroidia"                
## [1118,] "Bacteria" "Bacteroidetes"       "Bacteroidia"                
## [1119,] "Bacteria" "Actinobacteria"      "Actinobacteria"             
## [1120,] "Bacteria" "Patescibacteria"     "Saccharimonadia"            
## [1121,] "Bacteria" "Firmicutes"          "Negativicutes"              
## [1122,] "Bacteria" "Firmicutes"          "Negativicutes"              
## [1123,] "Bacteria" "Bacteroidetes"       "Bacteroidia"                
## [1124,] "Bacteria" "Epsilonbacteraeota"  "Campylobacteria"            
## [1125,] "Bacteria" "Actinobacteria"      "Actinobacteria"             
## [1126,] "Bacteria" "Cyanobacteria"       "Oxyphotobacteria"           
## [1127,] "Bacteria" "Bacteroidetes"       "Bacteroidia"                
## [1128,] "Bacteria" "Firmicutes"          NA                           
## [1129,] "Bacteria" "Fusobacteria"        "Fusobacteriia"              
## [1130,] "Bacteria" "Proteobacteria"      "Gammaproteobacteria"        
## [1131,] "Bacteria" "Firmicutes"          "Clostridia"                 
## [1132,] "Bacteria" "Bacteroidetes"       "Bacteroidia"                
## [1133,] "Bacteria" "Bacteroidetes"       "Bacteroidia"                
## [1134,] "Bacteria" "Firmicutes"          "Negativicutes"              
## [1135,] "Bacteria" "Proteobacteria"      "Gammaproteobacteria"        
## [1136,] "Bacteria" "Proteobacteria"      "Gammaproteobacteria"        
## [1137,] "Bacteria" "Bacteroidetes"       "Bacteroidia"                
## [1138,] "Bacteria" "Bacteroidetes"       "Bacteroidia"                
## [1139,] "Bacteria" "Proteobacteria"      "Gammaproteobacteria"        
## [1140,] "Bacteria" "Firmicutes"          "Bacilli"                    
## [1141,] "Bacteria" "Proteobacteria"      "Gammaproteobacteria"        
## [1142,] "Bacteria" "Bacteroidetes"       "Bacteroidia"                
## [1143,] "Bacteria" "Firmicutes"          "Clostridia"                 
## [1144,] "Bacteria" "Firmicutes"          "Bacilli"                    
## [1145,] "Bacteria" "Actinobacteria"      "Actinobacteria"             
## [1146,] "Bacteria" "Firmicutes"          "Bacilli"                    
## [1147,] "Bacteria" "Bacteroidetes"       "Bacteroidia"                
## [1148,] "Bacteria" "Firmicutes"          "Negativicutes"              
## [1149,] "Bacteria" "Firmicutes"          "Clostridia"                 
## [1150,] "Bacteria" "Actinobacteria"      "Actinobacteria"             
## [1151,] "Bacteria" "Fusobacteria"        "Fusobacteriia"              
## [1152,] "Bacteria" "Proteobacteria"      "Gammaproteobacteria"        
## [1153,] "Bacteria" "Firmicutes"          "Bacilli"                    
## [1154,] "Bacteria" "Proteobacteria"      "Gammaproteobacteria"        
## [1155,] "Bacteria" "Bacteroidetes"       "Bacteroidia"                
## [1156,] "Bacteria" "Proteobacteria"      "Gammaproteobacteria"        
## [1157,] "Bacteria" "Firmicutes"          "Bacilli"                    
## [1158,] "Bacteria" "Fusobacteria"        "Fusobacteriia"              
## [1159,] "Bacteria" "Firmicutes"          "Clostridia"                 
## [1160,] "Bacteria" "Bacteroidetes"       "Bacteroidia"                
## [1161,] "Bacteria" "Bacteroidetes"       "Bacteroidia"                
## [1162,] "Bacteria" "Firmicutes"          "Bacilli"                    
## [1163,] "Bacteria" "Tenericutes"         "Mollicutes"                 
## [1164,] "Bacteria" "Proteobacteria"      "Gammaproteobacteria"        
## [1165,] "Bacteria" "Patescibacteria"     "Gracilibacteria"            
## [1166,] "Bacteria" "Firmicutes"          "Negativicutes"              
## [1167,] "Bacteria" "Proteobacteria"      "Gammaproteobacteria"        
## [1168,] "Bacteria" "Firmicutes"          "Negativicutes"              
## [1169,] "Bacteria" "Firmicutes"          "Negativicutes"              
## [1170,] "Bacteria" "Firmicutes"          "Bacilli"                    
## [1171,] "Bacteria" "Bacteroidetes"       "Bacteroidia"                
## [1172,] "Bacteria" "Patescibacteria"     "Saccharimonadia"            
## [1173,] "Bacteria" "Bacteroidetes"       "Bacteroidia"                
## [1174,] "Bacteria" "Firmicutes"          "Negativicutes"              
## [1175,] "Bacteria" "Firmicutes"          "Negativicutes"              
## [1176,] "Bacteria" "Firmicutes"          "Negativicutes"              
## [1177,] "Bacteria" "Bacteroidetes"       "Bacteroidia"                
## [1178,] "Bacteria" "Fusobacteria"        "Fusobacteriia"              
## [1179,] "Bacteria" "Proteobacteria"      "Gammaproteobacteria"        
## [1180,] "Bacteria" "Bacteroidetes"       "Bacteroidia"                
## [1181,] "Bacteria" "Firmicutes"          "Clostridia"                 
## [1182,] "Bacteria" "Firmicutes"          "Clostridia"                 
## [1183,] "Bacteria" "Firmicutes"          "Bacilli"                    
## [1184,] "Bacteria" "Bacteroidetes"       "Bacteroidia"                
## [1185,] "Bacteria" "Proteobacteria"      "Gammaproteobacteria"        
## [1186,] "Bacteria" "Proteobacteria"      "Gammaproteobacteria"        
## [1187,] "Bacteria" "Bacteroidetes"       "Bacteroidia"                
## [1188,] "Bacteria" "Firmicutes"          "Negativicutes"              
## [1189,] "Bacteria" "Bacteroidetes"       "Bacteroidia"                
## [1190,] "Bacteria" "Bacteroidetes"       "Bacteroidia"                
## [1191,] "Bacteria" "Actinobacteria"      "Actinobacteria"             
## [1192,] "Bacteria" "Fusobacteria"        "Fusobacteriia"              
## [1193,] "Bacteria" "Actinobacteria"      "Actinobacteria"             
## [1194,] "Bacteria" "Proteobacteria"      "Gammaproteobacteria"        
## [1195,] "Bacteria" "Bacteroidetes"       "Bacteroidia"                
## [1196,] "Bacteria" "Bacteroidetes"       "Bacteroidia"                
## [1197,] "Bacteria" "Firmicutes"          "Negativicutes"              
## [1198,] "Bacteria" "Firmicutes"          "Clostridia"                 
## [1199,] "Bacteria" "Firmicutes"          "Bacilli"                    
## [1200,] "Bacteria" "Bacteroidetes"       "Bacteroidia"                
## [1201,] "Bacteria" "Firmicutes"          "Negativicutes"              
## [1202,] "Bacteria" "Fusobacteria"        "Fusobacteriia"              
## [1203,] "Bacteria" "Actinobacteria"      "Actinobacteria"             
## [1204,] "Bacteria" "Bacteroidetes"       "Bacteroidia"                
## [1205,] "Bacteria" "Synergistetes"       "Synergistia"                
## [1206,] "Bacteria" "Epsilonbacteraeota"  "Campylobacteria"            
## [1207,] "Bacteria" "Proteobacteria"      "Gammaproteobacteria"        
## [1208,] "Bacteria" "Firmicutes"          "Negativicutes"              
## [1209,] "Bacteria" "Proteobacteria"      "Alphaproteobacteria"        
## [1210,] "Bacteria" "Firmicutes"          "Negativicutes"              
## [1211,] "Bacteria" "Fusobacteria"        "Fusobacteriia"              
## [1212,] "Bacteria" "Patescibacteria"     "Saccharimonadia"            
## [1213,] "Bacteria" "Firmicutes"          "Negativicutes"              
## [1214,] "Bacteria" "Fusobacteria"        "Fusobacteriia"              
## [1215,] "Bacteria" "Patescibacteria"     "Saccharimonadia"            
## [1216,] "Bacteria" "Firmicutes"          "Clostridia"                 
## [1217,] "Bacteria" "Spirochaetes"        "Spirochaetia"               
## [1218,] "Bacteria" "Bacteroidetes"       "Bacteroidia"                
## [1219,] "Bacteria" "Firmicutes"          "Clostridia"                 
## [1220,] "Bacteria" "Firmicutes"          "Clostridia"                 
## [1221,] "Bacteria" "Firmicutes"          "Negativicutes"              
## [1222,] "Bacteria" "Actinobacteria"      "Actinobacteria"             
## [1223,] "Bacteria" "Bacteroidetes"       "Bacteroidia"                
## [1224,] "Bacteria" "Firmicutes"          NA                           
## [1225,] "Bacteria" "Firmicutes"          "Negativicutes"              
## [1226,] "Bacteria" "Firmicutes"          NA                           
## [1227,] "Bacteria" "Firmicutes"          "Negativicutes"              
## [1228,] "Bacteria" "Proteobacteria"      "Gammaproteobacteria"        
## [1229,] "Bacteria" "Bacteroidetes"       "Bacteroidia"                
## [1230,] "Bacteria" "Firmicutes"          "Negativicutes"              
## [1231,] "Bacteria" "Bacteroidetes"       "Bacteroidia"                
## [1232,] "Bacteria" "Bacteroidetes"       "Bacteroidia"                
## [1233,] "Bacteria" "Bacteroidetes"       "Bacteroidia"                
## [1234,] "Bacteria" "Firmicutes"          "Negativicutes"              
## [1235,] "Bacteria" "Bacteroidetes"       "Bacteroidia"                
## [1236,] "Bacteria" "Firmicutes"          "Clostridia"                 
## [1237,] "Bacteria" "Fusobacteria"        "Fusobacteriia"              
## [1238,] "Bacteria" "Actinobacteria"      "Actinobacteria"             
## [1239,] "Bacteria" "Firmicutes"          "Negativicutes"              
## [1240,] "Bacteria" "Patescibacteria"     "Saccharimonadia"            
## [1241,] "Bacteria" "Bacteroidetes"       "Bacteroidia"                
## [1242,] "Bacteria" "Bacteroidetes"       "Bacteroidia"                
## [1243,] "Bacteria" "Firmicutes"          "Clostridia"                 
## [1244,] "Bacteria" "Firmicutes"          "Negativicutes"              
## [1245,] "Bacteria" "Firmicutes"          "Negativicutes"              
## [1246,] "Bacteria" "Fusobacteria"        "Fusobacteriia"              
## [1247,] "Bacteria" "Firmicutes"          NA                           
## [1248,] "Bacteria" "Bacteroidetes"       "Bacteroidia"                
## [1249,] "Bacteria" "Firmicutes"          "Negativicutes"              
## [1250,] "Bacteria" "Actinobacteria"      "Actinobacteria"             
## [1251,] "Bacteria" "Fusobacteria"        "Fusobacteriia"              
## [1252,] "Bacteria" "Bacteroidetes"       "Bacteroidia"                
## [1253,] "Bacteria" "Bacteroidetes"       "Bacteroidia"                
## [1254,] "Bacteria" "Actinobacteria"      "Actinobacteria"             
## [1255,] "Bacteria" "Bacteroidetes"       "Bacteroidia"                
## [1256,] "Bacteria" "Bacteroidetes"       "Bacteroidia"                
## [1257,] "Bacteria" "Firmicutes"          "Negativicutes"              
## [1258,] "Bacteria" "Bacteroidetes"       "Bacteroidia"                
## [1259,] "Bacteria" "Firmicutes"          "Bacilli"                    
## [1260,] "Bacteria" "Bacteroidetes"       "Bacteroidia"                
## [1261,] "Bacteria" "Proteobacteria"      "Gammaproteobacteria"        
## [1262,] "Bacteria" "Firmicutes"          "Negativicutes"              
## [1263,] "Bacteria" "Patescibacteria"     "Saccharimonadia"            
## [1264,] "Bacteria" "Actinobacteria"      "Actinobacteria"             
## [1265,] "Bacteria" "Actinobacteria"      "Actinobacteria"             
## [1266,] "Bacteria" "Firmicutes"          "Negativicutes"              
## [1267,] "Bacteria" "Actinobacteria"      "Coriobacteriia"             
## [1268,] "Bacteria" "Bacteroidetes"       "Bacteroidia"                
## [1269,] "Bacteria" "Firmicutes"          "Clostridia"                 
## [1270,] "Bacteria" "Bacteroidetes"       "Bacteroidia"                
## [1271,] "Bacteria" "Firmicutes"          "Negativicutes"              
## [1272,] "Bacteria" "Firmicutes"          "Negativicutes"              
## [1273,] "Bacteria" "Fusobacteria"        "Fusobacteriia"              
## [1274,] "Bacteria" "Bacteroidetes"       "Bacteroidia"                
## [1275,] "Bacteria" "Spirochaetes"        "Spirochaetia"               
## [1276,] "Bacteria" "Bacteroidetes"       "Bacteroidia"                
## [1277,] "Bacteria" "Firmicutes"          "Negativicutes"              
## [1278,] "Bacteria" "Firmicutes"          "Negativicutes"              
## [1279,] "Bacteria" "Firmicutes"          "Negativicutes"              
## [1280,] "Bacteria" "Proteobacteria"      "Gammaproteobacteria"        
## [1281,] "Bacteria" "Proteobacteria"      "Gammaproteobacteria"        
## [1282,] "Bacteria" "Bacteroidetes"       "Bacteroidia"                
## [1283,] "Bacteria" "Patescibacteria"     "Saccharimonadia"            
## [1284,] "Bacteria" "Firmicutes"          "Negativicutes"              
## [1285,] "Bacteria" "Fusobacteria"        "Fusobacteriia"              
## [1286,] "Bacteria" "Actinobacteria"      "Actinobacteria"             
## [1287,] "Bacteria" "Firmicutes"          "Negativicutes"              
## [1288,] "Bacteria" "Firmicutes"          "Clostridia"                 
## [1289,] "Bacteria" "Tenericutes"         "Mollicutes"                 
## [1290,] "Bacteria" "Fusobacteria"        "Fusobacteriia"              
## [1291,] "Bacteria" "Bacteroidetes"       "Bacteroidia"                
## [1292,] "Bacteria" "Firmicutes"          "Clostridia"                 
## [1293,] "Bacteria" "Firmicutes"          "Clostridia"                 
## [1294,] "Bacteria" "Fusobacteria"        "Fusobacteriia"              
## [1295,] "Bacteria" "Proteobacteria"      "Gammaproteobacteria"        
## [1296,] "Bacteria" "Bacteroidetes"       "Bacteroidia"                
## [1297,] "Bacteria" "Firmicutes"          "Clostridia"                 
## [1298,] "Bacteria" "Bacteroidetes"       "Bacteroidia"                
## [1299,] "Bacteria" "Bacteroidetes"       "Bacteroidia"                
## [1300,] "Bacteria" "Bacteroidetes"       "Bacteroidia"                
## [1301,] "Bacteria" "Firmicutes"          "Clostridia"                 
## [1302,] "Bacteria" "Bacteroidetes"       "Bacteroidia"                
## [1303,] "Bacteria" "Firmicutes"          "Negativicutes"              
## [1304,] "Bacteria" "Bacteroidetes"       "Bacteroidia"                
## [1305,] "Bacteria" "Actinobacteria"      "Actinobacteria"             
## [1306,] "Bacteria" "Firmicutes"          "Bacilli"                    
## [1307,] "Bacteria" "Proteobacteria"      "Gammaproteobacteria"        
## [1308,] "Bacteria" "Proteobacteria"      "Gammaproteobacteria"        
## [1309,] "Bacteria" "Spirochaetes"        "Spirochaetia"               
## [1310,] "Bacteria" "Proteobacteria"      "Gammaproteobacteria"        
## [1311,] "Bacteria" "Proteobacteria"      "Gammaproteobacteria"        
## [1312,] "Bacteria" "Firmicutes"          "Clostridia"                 
## [1313,] "Bacteria" "Bacteroidetes"       "Bacteroidia"                
## [1314,] "Bacteria" "Firmicutes"          "Clostridia"                 
## [1315,] "Bacteria" "Bacteroidetes"       "Bacteroidia"                
## [1316,] "Bacteria" "Bacteroidetes"       "Bacteroidia"                
## [1317,] "Bacteria" "Epsilonbacteraeota"  "Campylobacteria"            
## [1318,] "Bacteria" "Firmicutes"          "Negativicutes"              
## [1319,] "Bacteria" "Firmicutes"          "Negativicutes"              
## [1320,] "Bacteria" "Firmicutes"          "Bacilli"                    
## [1321,] "Bacteria" "Firmicutes"          "Negativicutes"              
## [1322,] "Bacteria" "Epsilonbacteraeota"  "Campylobacteria"            
## [1323,] "Bacteria" "Firmicutes"          "Negativicutes"              
## [1324,] "Bacteria" "Firmicutes"          "Bacilli"                    
## [1325,] "Bacteria" "Bacteroidetes"       "Bacteroidia"                
## [1326,] "Bacteria" "Proteobacteria"      "Gammaproteobacteria"        
## [1327,] "Bacteria" "Firmicutes"          "Bacilli"                    
## [1328,] "Bacteria" "Firmicutes"          "Negativicutes"              
## [1329,] "Bacteria" "Bacteroidetes"       "Bacteroidia"                
## [1330,] "Bacteria" "Bacteroidetes"       "Bacteroidia"                
## [1331,] "Bacteria" "Firmicutes"          "Clostridia"                 
## [1332,] "Bacteria" "Bacteroidetes"       "Bacteroidia"                
## [1333,] "Bacteria" "Firmicutes"          "Negativicutes"              
## [1334,] "Bacteria" "Bacteroidetes"       "Bacteroidia"                
## [1335,] "Bacteria" "Fusobacteria"        "Fusobacteriia"              
## [1336,] "Bacteria" "Fusobacteria"        "Fusobacteriia"              
## [1337,] "Bacteria" "Firmicutes"          "Negativicutes"              
## [1338,] "Bacteria" "Firmicutes"          "Clostridia"                 
## [1339,] "Bacteria" "Proteobacteria"      "Gammaproteobacteria"        
## [1340,] "Bacteria" "Firmicutes"          "Negativicutes"              
## [1341,] "Bacteria" "Firmicutes"          NA                           
## [1342,] "Bacteria" "Fusobacteria"        "Fusobacteriia"              
## [1343,] "Bacteria" "Bacteroidetes"       "Bacteroidia"                
## [1344,] "Bacteria" "Patescibacteria"     "Saccharimonadia"            
## [1345,] "Bacteria" "Bacteroidetes"       "Bacteroidia"                
## [1346,] "Bacteria" "Patescibacteria"     "Saccharimonadia"            
## [1347,] "Bacteria" "Bacteroidetes"       "Bacteroidia"                
## [1348,] "Bacteria" "Bacteroidetes"       "Bacteroidia"                
## [1349,] "Bacteria" "Proteobacteria"      "Gammaproteobacteria"        
## [1350,] "Bacteria" "Actinobacteria"      "Actinobacteria"             
## [1351,] "Bacteria" "Actinobacteria"      "Actinobacteria"             
## [1352,] "Bacteria" "Actinobacteria"      "Actinobacteria"             
## [1353,] "Bacteria" "Bacteroidetes"       "Bacteroidia"                
## [1354,] "Bacteria" "Firmicutes"          "Negativicutes"              
## [1355,] "Bacteria" "Epsilonbacteraeota"  "Campylobacteria"            
## [1356,] "Bacteria" "Firmicutes"          NA                           
## [1357,] "Bacteria" "Bacteroidetes"       "Bacteroidia"                
## [1358,] "Bacteria" "Tenericutes"         "Mollicutes"                 
## [1359,] "Bacteria" "Firmicutes"          "Negativicutes"              
## [1360,] "Bacteria" "Bacteroidetes"       "Bacteroidia"                
## [1361,] "Bacteria" "Proteobacteria"      "Gammaproteobacteria"        
## [1362,] "Bacteria" "Firmicutes"          NA                           
## [1363,] "Bacteria" "Firmicutes"          NA                           
## [1364,] "Bacteria" "Actinobacteria"      "Actinobacteria"             
## [1365,] "Bacteria" "Bacteroidetes"       "Bacteroidia"                
## [1366,] "Bacteria" "Bacteroidetes"       "Bacteroidia"                
## [1367,] "Bacteria" "Bacteroidetes"       "Bacteroidia"                
## [1368,] "Bacteria" "Proteobacteria"      "Gammaproteobacteria"        
## [1369,] "Bacteria" "Fusobacteria"        "Fusobacteriia"              
## [1370,] "Bacteria" "Firmicutes"          "Clostridia"                 
## [1371,] "Bacteria" "Bacteroidetes"       "Bacteroidia"                
## [1372,] "Bacteria" "Fusobacteria"        "Fusobacteriia"              
## [1373,] "Bacteria" "Firmicutes"          "Bacilli"                    
## [1374,] "Bacteria" "Proteobacteria"      "Gammaproteobacteria"        
## [1375,] "Bacteria" "Fusobacteria"        "Fusobacteriia"              
## [1376,] "Bacteria" "Proteobacteria"      "Gammaproteobacteria"        
## [1377,] "Bacteria" "Firmicutes"          "Negativicutes"              
## [1378,] "Bacteria" "Bacteroidetes"       "Bacteroidia"                
## [1379,] "Bacteria" "Bacteroidetes"       "Bacteroidia"                
## [1380,] "Bacteria" "Patescibacteria"     "Gracilibacteria"            
## [1381,] "Bacteria" "Firmicutes"          "Clostridia"                 
## [1382,] "Bacteria" "Bacteroidetes"       "Bacteroidia"                
## [1383,] "Bacteria" "Firmicutes"          "Negativicutes"              
## [1384,] "Bacteria" "Firmicutes"          "Bacilli"                    
## [1385,] "Bacteria" "Firmicutes"          "Negativicutes"              
## [1386,] "Bacteria" "Firmicutes"          "Bacilli"                    
## [1387,] "Bacteria" "Fusobacteria"        "Fusobacteriia"              
## [1388,] "Bacteria" "Bacteroidetes"       "Bacteroidia"                
## [1389,] "Bacteria" "Firmicutes"          "Negativicutes"              
## [1390,] "Bacteria" "Bacteroidetes"       "Bacteroidia"                
## [1391,] "Bacteria" "Patescibacteria"     "Saccharimonadia"            
## [1392,] "Bacteria" "Bacteroidetes"       "Bacteroidia"                
## [1393,] "Bacteria" "Fusobacteria"        "Fusobacteriia"              
## [1394,] "Bacteria" "Bacteroidetes"       "Bacteroidia"                
## [1395,] "Bacteria" "Fusobacteria"        "Fusobacteriia"              
## [1396,] "Bacteria" "Firmicutes"          NA                           
## [1397,] "Bacteria" "Bacteroidetes"       "Bacteroidia"                
## [1398,] "Bacteria" "Bacteroidetes"       "Bacteroidia"                
## [1399,] "Bacteria" "Firmicutes"          "Bacilli"                    
## [1400,] "Bacteria" "Bacteroidetes"       "Bacteroidia"                
## [1401,] "Bacteria" "Fusobacteria"        "Fusobacteriia"              
## [1402,] "Bacteria" "Bacteroidetes"       "Bacteroidia"                
## [1403,] "Bacteria" "Fusobacteria"        "Fusobacteriia"              
## [1404,] "Bacteria" "Bacteroidetes"       "Bacteroidia"                
## [1405,] "Bacteria" "Epsilonbacteraeota"  "Campylobacteria"            
## [1406,] "Bacteria" "Proteobacteria"      "Gammaproteobacteria"        
## [1407,] "Bacteria" "Fusobacteria"        "Fusobacteriia"              
## [1408,] "Bacteria" "Firmicutes"          "Negativicutes"              
## [1409,] "Bacteria" "Proteobacteria"      "Gammaproteobacteria"        
## [1410,] "Bacteria" "Fusobacteria"        "Fusobacteriia"              
## [1411,] "Bacteria" "Firmicutes"          "Clostridia"                 
## [1412,] "Bacteria" "Firmicutes"          "Bacilli"                    
## [1413,] "Bacteria" "Proteobacteria"      "Gammaproteobacteria"        
## [1414,] "Bacteria" "Proteobacteria"      "Gammaproteobacteria"        
## [1415,] "Bacteria" "Bacteroidetes"       "Bacteroidia"                
## [1416,] "Bacteria" "Actinobacteria"      "Actinobacteria"             
## [1417,] "Bacteria" "Spirochaetes"        "Spirochaetia"               
## [1418,] "Bacteria" "Firmicutes"          "Negativicutes"              
## [1419,] "Bacteria" "Fusobacteria"        "Fusobacteriia"              
## [1420,] "Bacteria" "Fusobacteria"        "Fusobacteriia"              
## [1421,] "Bacteria" "Epsilonbacteraeota"  "Campylobacteria"            
## [1422,] "Bacteria" "Firmicutes"          "Negativicutes"              
## [1423,] "Bacteria" "Fusobacteria"        "Fusobacteriia"              
## [1424,] "Bacteria" "Bacteroidetes"       "Bacteroidia"                
## [1425,] "Bacteria" "Firmicutes"          "Negativicutes"              
## [1426,] "Bacteria" "Epsilonbacteraeota"  "Campylobacteria"            
## [1427,] "Bacteria" "Bacteroidetes"       "Bacteroidia"                
## [1428,] "Bacteria" "Actinobacteria"      "Actinobacteria"             
## [1429,] "Bacteria" "Bacteroidetes"       "Bacteroidia"                
## [1430,] "Bacteria" "Bacteroidetes"       "Bacteroidia"                
## [1431,] "Bacteria" "Firmicutes"          "Negativicutes"              
## [1432,] "Bacteria" "Spirochaetes"        "Spirochaetia"               
## [1433,] "Bacteria" "Bacteroidetes"       "Bacteroidia"                
## [1434,] "Bacteria" "Firmicutes"          "Bacilli"                    
## [1435,] "Bacteria" "Proteobacteria"      "Gammaproteobacteria"        
## [1436,] "Bacteria" "Patescibacteria"     "Saccharimonadia"            
## [1437,] "Bacteria" "Firmicutes"          "Clostridia"                 
## [1438,] "Bacteria" "Firmicutes"          "Negativicutes"              
## [1439,] "Bacteria" "Proteobacteria"      "Gammaproteobacteria"        
## [1440,] "Bacteria" "Actinobacteria"      "Actinobacteria"             
## [1441,] "Bacteria" "Firmicutes"          "Negativicutes"              
## [1442,] "Bacteria" "Firmicutes"          "Negativicutes"              
## [1443,] "Bacteria" "Bacteroidetes"       "Bacteroidia"                
## [1444,] "Bacteria" "Bacteroidetes"       "Bacteroidia"                
## [1445,] "Bacteria" "Firmicutes"          "Clostridia"                 
## [1446,] "Bacteria" "Proteobacteria"      "Gammaproteobacteria"        
## [1447,] "Bacteria" NA                    NA                           
## [1448,] "Bacteria" "Proteobacteria"      "Gammaproteobacteria"        
## [1449,] "Bacteria" "Bacteroidetes"       "Bacteroidia"                
## [1450,] "Bacteria" "Firmicutes"          "Bacilli"                    
## [1451,] "Bacteria" "Firmicutes"          "Bacilli"                    
## [1452,] "Bacteria" "Actinobacteria"      "Actinobacteria"             
## [1453,] "Bacteria" "Bacteroidetes"       "Bacteroidia"                
## [1454,] "Bacteria" "Firmicutes"          "Negativicutes"              
## [1455,] "Bacteria" "Firmicutes"          "Negativicutes"              
## [1456,] "Bacteria" "Proteobacteria"      "Gammaproteobacteria"        
## [1457,] "Bacteria" "Bacteroidetes"       "Bacteroidia"                
## [1458,] "Bacteria" "Patescibacteria"     "Saccharimonadia"            
## [1459,] "Bacteria" "Firmicutes"          "Negativicutes"              
## [1460,] "Bacteria" "Fusobacteria"        "Fusobacteriia"              
## [1461,] "Bacteria" "Bacteroidetes"       "Bacteroidia"                
## [1462,] "Bacteria" "Bacteroidetes"       "Bacteroidia"                
## [1463,] "Bacteria" "Spirochaetes"        "Spirochaetia"               
## [1464,] "Bacteria" "Firmicutes"          "Clostridia"                 
## [1465,] "Bacteria" "Fusobacteria"        "Fusobacteriia"              
## [1466,] "Bacteria" "Actinobacteria"      "Actinobacteria"             
## [1467,] "Bacteria" "Bacteroidetes"       "Bacteroidia"                
## [1468,] "Bacteria" "Bacteroidetes"       "Bacteroidia"                
## [1469,] "Bacteria" "Firmicutes"          "Bacilli"                    
## [1470,] "Bacteria" "Bacteroidetes"       "Bacteroidia"                
## [1471,] "Bacteria" NA                    NA                           
## [1472,] "Bacteria" "Fusobacteria"        "Fusobacteriia"              
## [1473,] "Bacteria" "Bacteroidetes"       "Bacteroidia"                
## [1474,] "Bacteria" "Patescibacteria"     "Saccharimonadia"            
## [1475,] "Bacteria" "Bacteroidetes"       "Bacteroidia"                
## [1476,] "Bacteria" "Firmicutes"          "Negativicutes"              
## [1477,] "Bacteria" "Fusobacteria"        "Fusobacteriia"              
## [1478,] "Bacteria" "Firmicutes"          "Clostridia"                 
## [1479,] "Bacteria" "Proteobacteria"      "Gammaproteobacteria"        
## [1480,] "Bacteria" "Firmicutes"          "Negativicutes"              
## [1481,] "Bacteria" "Bacteroidetes"       "Bacteroidia"                
## [1482,] "Bacteria" "Firmicutes"          "Negativicutes"              
## [1483,] "Bacteria" "Bacteroidetes"       "Bacteroidia"                
## [1484,] "Bacteria" "Bacteroidetes"       "Bacteroidia"                
## [1485,] "Bacteria" "Bacteroidetes"       "Bacteroidia"                
## [1486,] "Bacteria" "Firmicutes"          "Negativicutes"              
## [1487,] "Bacteria" "Bacteroidetes"       "Bacteroidia"                
## [1488,] "Bacteria" "Firmicutes"          "Negativicutes"              
## [1489,] "Bacteria" "Proteobacteria"      "Gammaproteobacteria"        
## [1490,] "Bacteria" "Actinobacteria"      "Actinobacteria"             
## [1491,] "Bacteria" "Fusobacteria"        "Fusobacteriia"              
## [1492,] "Bacteria" "Firmicutes"          "Clostridia"                 
## [1493,] "Bacteria" "Firmicutes"          "Clostridia"                 
## [1494,] "Bacteria" "Firmicutes"          "Negativicutes"              
## [1495,] "Bacteria" "Proteobacteria"      "Gammaproteobacteria"        
## [1496,] "Bacteria" "Proteobacteria"      "Gammaproteobacteria"        
## [1497,] "Bacteria" "Bacteroidetes"       "Bacteroidia"                
## [1498,] "Bacteria" "Proteobacteria"      "Gammaproteobacteria"        
## [1499,] "Bacteria" "Patescibacteria"     "Saccharimonadia"            
## [1500,] "Bacteria" "Firmicutes"          "Bacilli"                    
## [1501,] "Bacteria" "Actinobacteria"      "Actinobacteria"             
## [1502,] "Bacteria" "Epsilonbacteraeota"  "Campylobacteria"            
## [1503,] "Bacteria" "Bacteroidetes"       "Bacteroidia"                
## [1504,] "Bacteria" "Firmicutes"          "Negativicutes"              
## [1505,] "Bacteria" "Proteobacteria"      "Gammaproteobacteria"        
## [1506,] "Bacteria" "Patescibacteria"     "Saccharimonadia"            
## [1507,] "Bacteria" "Bacteroidetes"       "Bacteroidia"                
## [1508,] "Bacteria" "Bacteroidetes"       "Bacteroidia"                
## [1509,] "Bacteria" "Proteobacteria"      "Gammaproteobacteria"        
## [1510,] "Bacteria" "Fusobacteria"        "Fusobacteriia"              
## [1511,] "Bacteria" "Proteobacteria"      "Gammaproteobacteria"        
## [1512,] "Bacteria" "Bacteroidetes"       "Bacteroidia"                
## [1513,] "Bacteria" "Bacteroidetes"       "Bacteroidia"                
## [1514,] "Bacteria" "Firmicutes"          "Bacilli"                    
## [1515,] "Bacteria" "Actinobacteria"      "Actinobacteria"             
## [1516,] "Bacteria" "Bacteroidetes"       "Bacteroidia"                
## [1517,] "Bacteria" "Proteobacteria"      "Gammaproteobacteria"        
## [1518,] "Bacteria" "Firmicutes"          "Clostridia"                 
## [1519,] "Bacteria" "Firmicutes"          "Bacilli"                    
## [1520,] "Bacteria" "Bacteroidetes"       "Bacteroidia"                
## [1521,] "Bacteria" "Fusobacteria"        "Fusobacteriia"              
## [1522,] "Bacteria" "Bacteroidetes"       "Bacteroidia"                
## [1523,] "Bacteria" "Fusobacteria"        "Fusobacteriia"              
## [1524,] "Bacteria" "Proteobacteria"      "Gammaproteobacteria"        
## [1525,] "Bacteria" "Actinobacteria"      "Coriobacteriia"             
## [1526,] "Bacteria" "Firmicutes"          "Negativicutes"              
## [1527,] "Bacteria" "Bacteroidetes"       "Bacteroidia"                
## [1528,] "Bacteria" "Firmicutes"          NA                           
## [1529,] "Bacteria" "Bacteroidetes"       "Bacteroidia"                
## [1530,] "Bacteria" "Firmicutes"          "Clostridia"                 
## [1531,] "Bacteria" "Epsilonbacteraeota"  "Campylobacteria"            
## [1532,] "Bacteria" "Patescibacteria"     "Saccharimonadia"            
## [1533,] "Bacteria" "Fusobacteria"        "Fusobacteriia"              
## [1534,] "Bacteria" "Firmicutes"          "Clostridia"                 
## [1535,] "Bacteria" "Firmicutes"          NA                           
## [1536,] "Bacteria" "Bacteroidetes"       "Bacteroidia"                
## [1537,] "Bacteria" "Bacteroidetes"       "Bacteroidia"                
## [1538,] "Bacteria" "Patescibacteria"     "Gracilibacteria"            
## [1539,] "Bacteria" "Actinobacteria"      "Actinobacteria"             
## [1540,] "Bacteria" "Firmicutes"          "Negativicutes"              
## [1541,] "Bacteria" "Firmicutes"          "Negativicutes"              
## [1542,] "Bacteria" "Bacteroidetes"       "Bacteroidia"                
## [1543,] "Bacteria" "Firmicutes"          "Bacilli"                    
## [1544,] "Bacteria" "Spirochaetes"        "Spirochaetia"               
## [1545,] "Bacteria" "Firmicutes"          "Clostridia"                 
## [1546,] "Bacteria" "Firmicutes"          "Negativicutes"              
## [1547,] "Bacteria" "Bacteroidetes"       "Bacteroidia"                
## [1548,] "Bacteria" "Actinobacteria"      "Actinobacteria"             
## [1549,] "Bacteria" "Fusobacteria"        "Fusobacteriia"              
## [1550,] "Bacteria" "Firmicutes"          "Negativicutes"              
## [1551,] "Bacteria" "Firmicutes"          "Clostridia"                 
## [1552,] "Bacteria" "Patescibacteria"     "Gracilibacteria"            
## [1553,] "Bacteria" "Fusobacteria"        "Fusobacteriia"              
## [1554,] "Bacteria" "Firmicutes"          "Negativicutes"              
## [1555,] "Bacteria" "Bacteroidetes"       "Bacteroidia"                
## [1556,] "Bacteria" "Actinobacteria"      "Actinobacteria"             
## [1557,] "Bacteria" "Bacteroidetes"       "Bacteroidia"                
## [1558,] "Bacteria" "Firmicutes"          "Negativicutes"              
## [1559,] "Bacteria" "Fusobacteria"        "Fusobacteriia"              
## [1560,] "Bacteria" "Cyanobacteria"       "Oxyphotobacteria"           
## [1561,] "Bacteria" "Bacteroidetes"       "Bacteroidia"                
## [1562,] "Bacteria" "Bacteroidetes"       "Bacteroidia"                
## [1563,] "Bacteria" "Firmicutes"          "Negativicutes"              
## [1564,] "Bacteria" "Fusobacteria"        "Fusobacteriia"              
## [1565,] "Bacteria" "Bacteroidetes"       "Bacteroidia"                
## [1566,] "Bacteria" "Firmicutes"          "Clostridia"                 
## [1567,] "Bacteria" "Firmicutes"          "Negativicutes"              
## [1568,] "Bacteria" "Firmicutes"          "Bacilli"                    
## [1569,] "Bacteria" "Epsilonbacteraeota"  "Campylobacteria"            
## [1570,] "Bacteria" "Bacteroidetes"       "Bacteroidia"                
## [1571,] "Bacteria" "Bacteroidetes"       "Bacteroidia"                
## [1572,] "Bacteria" "Bacteroidetes"       "Bacteroidia"                
## [1573,] "Bacteria" "Firmicutes"          "Negativicutes"              
## [1574,] "Bacteria" "Fusobacteria"        "Fusobacteriia"              
## [1575,] "Bacteria" "Bacteroidetes"       "Bacteroidia"                
## [1576,] "Bacteria" "Bacteroidetes"       "Bacteroidia"                
## [1577,] "Bacteria" "Fusobacteria"        "Fusobacteriia"              
## [1578,] "Bacteria" "Bacteroidetes"       "Bacteroidia"                
## [1579,] "Bacteria" "Bacteroidetes"       "Bacteroidia"                
## [1580,] "Bacteria" "Bacteroidetes"       "Bacteroidia"                
## [1581,] "Bacteria" "Bacteroidetes"       "Bacteroidia"                
## [1582,] "Bacteria" "Firmicutes"          "Negativicutes"              
## [1583,] "Bacteria" "Firmicutes"          "Bacilli"                    
## [1584,] "Bacteria" "Spirochaetes"        "Spirochaetia"               
## [1585,] "Bacteria" "Epsilonbacteraeota"  "Campylobacteria"            
## [1586,] "Bacteria" "Bacteroidetes"       "Bacteroidia"                
## [1587,] "Bacteria" "Firmicutes"          "Negativicutes"              
## [1588,] "Bacteria" "Firmicutes"          "Negativicutes"              
## [1589,] "Bacteria" "Bacteroidetes"       "Bacteroidia"                
## [1590,] "Bacteria" "Epsilonbacteraeota"  "Campylobacteria"            
## [1591,] "Bacteria" "Bacteroidetes"       "Bacteroidia"                
## [1592,] "Bacteria" "Firmicutes"          "Bacilli"                    
## [1593,] "Bacteria" "Proteobacteria"      "Gammaproteobacteria"        
## [1594,] "Bacteria" "Bacteroidetes"       "Bacteroidia"                
## [1595,] "Bacteria" "Proteobacteria"      "Gammaproteobacteria"        
## [1596,] "Bacteria" "Firmicutes"          "Negativicutes"              
## [1597,] "Bacteria" "Bacteroidetes"       "Bacteroidia"                
## [1598,] "Bacteria" "Bacteroidetes"       "Bacteroidia"                
## [1599,] "Bacteria" "Bacteroidetes"       "Bacteroidia"                
## [1600,] "Bacteria" "Bacteroidetes"       "Bacteroidia"                
## [1601,] "Bacteria" "Firmicutes"          "Negativicutes"              
## [1602,] "Bacteria" "Bacteroidetes"       "Bacteroidia"                
## [1603,] "Bacteria" "Bacteroidetes"       "Bacteroidia"                
## [1604,] "Bacteria" "Patescibacteria"     "Saccharimonadia"            
## [1605,] "Bacteria" "Patescibacteria"     "Saccharimonadia"            
## [1606,] "Bacteria" "Firmicutes"          "Bacilli"                    
## [1607,] "Bacteria" "Bacteroidetes"       "Bacteroidia"                
## [1608,] "Bacteria" "Firmicutes"          "Negativicutes"              
## [1609,] "Bacteria" "Proteobacteria"      "Gammaproteobacteria"        
## [1610,] "Bacteria" "Bacteroidetes"       "Bacteroidia"                
## [1611,] "Bacteria" "Fusobacteria"        "Fusobacteriia"              
## [1612,] "Bacteria" "Fusobacteria"        "Fusobacteriia"              
## [1613,] "Bacteria" "Proteobacteria"      "Gammaproteobacteria"        
## [1614,] "Bacteria" "Bacteroidetes"       "Bacteroidia"                
## [1615,] "Bacteria" "Proteobacteria"      "Gammaproteobacteria"        
## [1616,] "Bacteria" "Cyanobacteria"       "Oxyphotobacteria"           
## [1617,] "Bacteria" "Actinobacteria"      "Actinobacteria"             
## [1618,] "Bacteria" "Firmicutes"          "Negativicutes"              
## [1619,] "Bacteria" "Firmicutes"          "Negativicutes"              
## [1620,] "Bacteria" "Fusobacteria"        "Fusobacteriia"              
## [1621,] "Bacteria" "Actinobacteria"      "Actinobacteria"             
## [1622,] "Bacteria" "Proteobacteria"      "Gammaproteobacteria"        
## [1623,] "Bacteria" "Fusobacteria"        "Fusobacteriia"              
## [1624,] "Bacteria" "Bacteroidetes"       "Bacteroidia"                
## [1625,] "Bacteria" "Fusobacteria"        "Fusobacteriia"              
## [1626,] "Bacteria" "Firmicutes"          "Bacilli"                    
## [1627,] "Bacteria" "Bacteroidetes"       "Bacteroidia"                
## [1628,] "Bacteria" "Firmicutes"          "Clostridia"                 
## [1629,] "Bacteria" "Firmicutes"          "Clostridia"                 
## [1630,] "Bacteria" "Bacteroidetes"       "Bacteroidia"                
## [1631,] "Bacteria" "Bacteroidetes"       "Bacteroidia"                
## [1632,] "Bacteria" "Firmicutes"          "Negativicutes"              
## [1633,] "Bacteria" "Fusobacteria"        "Fusobacteriia"              
## [1634,] "Bacteria" "Firmicutes"          "Negativicutes"              
## [1635,] "Bacteria" "Bacteroidetes"       "Bacteroidia"                
## [1636,] "Bacteria" "Proteobacteria"      "Gammaproteobacteria"        
## [1637,] "Bacteria" "Firmicutes"          "Negativicutes"              
## [1638,] "Bacteria" "Bacteroidetes"       "Bacteroidia"                
## [1639,] "Bacteria" "Bacteroidetes"       "Bacteroidia"                
## [1640,] "Bacteria" "Bacteroidetes"       "Bacteroidia"                
## [1641,] "Bacteria" "Actinobacteria"      "Actinobacteria"             
## [1642,] "Bacteria" "Bacteroidetes"       "Bacteroidia"                
## [1643,] "Bacteria" "Firmicutes"          "Clostridia"                 
## [1644,] "Bacteria" "Patescibacteria"     "Gracilibacteria"            
## [1645,] "Bacteria" "Bacteroidetes"       "Bacteroidia"                
## [1646,] "Bacteria" "Fusobacteria"        "Fusobacteriia"              
## [1647,] "Bacteria" "Firmicutes"          "Clostridia"                 
## [1648,] "Bacteria" "Firmicutes"          "Clostridia"                 
## [1649,] "Bacteria" "Fusobacteria"        "Fusobacteriia"              
## [1650,] "Bacteria" "Fusobacteria"        "Fusobacteriia"              
## [1651,] "Bacteria" "Firmicutes"          "Clostridia"                 
## [1652,] "Bacteria" "Bacteroidetes"       "Bacteroidia"                
## [1653,] "Bacteria" "Firmicutes"          "Negativicutes"              
## [1654,] "Bacteria" "Epsilonbacteraeota"  "Campylobacteria"            
## [1655,] "Bacteria" "Patescibacteria"     "Saccharimonadia"            
## [1656,] "Bacteria" "Actinobacteria"      "Coriobacteriia"             
## [1657,] "Bacteria" "Bacteroidetes"       "Bacteroidia"                
## [1658,] "Bacteria" "Spirochaetes"        "Spirochaetia"               
## [1659,] "Bacteria" "Proteobacteria"      "Gammaproteobacteria"        
## [1660,] "Bacteria" "Proteobacteria"      "Gammaproteobacteria"        
## [1661,] "Bacteria" "Bacteroidetes"       "Bacteroidia"                
## [1662,] "Bacteria" "Bacteroidetes"       "Bacteroidia"                
## [1663,] "Bacteria" "Firmicutes"          "Negativicutes"              
## [1664,] "Bacteria" "Bacteroidetes"       "Bacteroidia"                
## [1665,] "Bacteria" "Bacteroidetes"       "Bacteroidia"                
## [1666,] "Bacteria" "Fusobacteria"        "Fusobacteriia"              
## [1667,] "Bacteria" "Fusobacteria"        "Fusobacteriia"              
## [1668,] "Bacteria" "Firmicutes"          "Clostridia"                 
## [1669,] "Bacteria" "Spirochaetes"        "Spirochaetia"               
## [1670,] "Bacteria" "Fusobacteria"        "Fusobacteriia"              
## [1671,] "Bacteria" "Firmicutes"          NA                           
## [1672,] "Bacteria" "Bacteroidetes"       "Bacteroidia"                
## [1673,] "Bacteria" "Proteobacteria"      "Gammaproteobacteria"        
## [1674,] "Bacteria" "Proteobacteria"      "Gammaproteobacteria"        
## [1675,] "Bacteria" "Bacteroidetes"       "Bacteroidia"                
## [1676,] "Bacteria" "Firmicutes"          "Clostridia"                 
## [1677,] "Bacteria" "Firmicutes"          "Bacilli"                    
## [1678,] "Bacteria" "Firmicutes"          NA                           
## [1679,] "Bacteria" "Proteobacteria"      "Gammaproteobacteria"        
## [1680,] "Bacteria" "Patescibacteria"     "Saccharimonadia"            
## [1681,] "Bacteria" "Bacteroidetes"       "Bacteroidia"                
## [1682,] "Bacteria" "Bacteroidetes"       "Bacteroidia"                
## [1683,] "Bacteria" "Firmicutes"          "Bacilli"                    
## [1684,] "Bacteria" "Proteobacteria"      "Gammaproteobacteria"        
## [1685,] "Bacteria" "Firmicutes"          "Negativicutes"              
## [1686,] "Bacteria" "Firmicutes"          "Bacilli"                    
## [1687,] "Bacteria" "Bacteroidetes"       "Bacteroidia"                
## [1688,] "Bacteria" "Firmicutes"          "Negativicutes"              
## [1689,] "Bacteria" "Fusobacteria"        "Fusobacteriia"              
## [1690,] "Bacteria" "Proteobacteria"      "Gammaproteobacteria"        
## [1691,] "Bacteria" "Fusobacteria"        "Fusobacteriia"              
## [1692,] "Bacteria" "Bacteroidetes"       "Bacteroidia"                
## [1693,] "Bacteria" "Fusobacteria"        "Fusobacteriia"              
## [1694,] "Bacteria" "Bacteroidetes"       "Bacteroidia"                
## [1695,] "Bacteria" "Bacteroidetes"       "Bacteroidia"                
## [1696,] "Bacteria" "Actinobacteria"      "Actinobacteria"             
## [1697,] "Bacteria" "Bacteroidetes"       "Bacteroidia"                
## [1698,] "Bacteria" "Epsilonbacteraeota"  "Campylobacteria"            
## [1699,] "Bacteria" "Proteobacteria"      "Gammaproteobacteria"        
## [1700,] "Bacteria" "Actinobacteria"      "Actinobacteria"             
## [1701,] "Bacteria" "Patescibacteria"     "Saccharimonadia"            
## [1702,] "Bacteria" "Fusobacteria"        "Fusobacteriia"              
## [1703,] "Bacteria" "Firmicutes"          "Bacilli"                    
## [1704,] "Bacteria" "Proteobacteria"      "Gammaproteobacteria"        
## [1705,] "Bacteria" "Bacteroidetes"       "Bacteroidia"                
## [1706,] "Bacteria" "Proteobacteria"      "Gammaproteobacteria"        
## [1707,] "Bacteria" "Firmicutes"          "Bacilli"                    
## [1708,] "Bacteria" "Firmicutes"          "Negativicutes"              
## [1709,] "Bacteria" "Firmicutes"          "Bacilli"                    
## [1710,] "Bacteria" "Bacteroidetes"       "Bacteroidia"                
## [1711,] "Bacteria" "Firmicutes"          "Negativicutes"              
## [1712,] "Bacteria" "Firmicutes"          "Negativicutes"              
## [1713,] "Bacteria" "Firmicutes"          "Clostridia"                 
## [1714,] "Bacteria" "Firmicutes"          "Negativicutes"              
## [1715,] "Bacteria" "Bacteroidetes"       "Bacteroidia"                
## [1716,] "Bacteria" "Fusobacteria"        "Fusobacteriia"              
## [1717,] "Bacteria" "Proteobacteria"      "Gammaproteobacteria"        
## [1718,] "Bacteria" "Bacteroidetes"       "Bacteroidia"                
## [1719,] "Bacteria" "Firmicutes"          "Clostridia"                 
## [1720,] "Bacteria" "Proteobacteria"      "Gammaproteobacteria"        
## [1721,] "Bacteria" "Bacteroidetes"       "Bacteroidia"                
## [1722,] "Bacteria" "Bacteroidetes"       "Bacteroidia"                
## [1723,] "Bacteria" "Firmicutes"          "Negativicutes"              
## [1724,] "Bacteria" "Bacteroidetes"       "Bacteroidia"                
## [1725,] "Bacteria" "Proteobacteria"      "Gammaproteobacteria"        
## [1726,] "Bacteria" "Bacteroidetes"       "Bacteroidia"                
## [1727,] "Bacteria" "Actinobacteria"      "Coriobacteriia"             
## [1728,] "Bacteria" "Bacteroidetes"       "Bacteroidia"                
## [1729,] "Bacteria" "Bacteroidetes"       "Bacteroidia"                
## [1730,] "Bacteria" "Fusobacteria"        "Fusobacteriia"              
## [1731,] "Bacteria" "Firmicutes"          "Negativicutes"              
## [1732,] "Bacteria" "Actinobacteria"      "Actinobacteria"             
## [1733,] "Bacteria" "Bacteroidetes"       "Bacteroidia"                
## [1734,] "Bacteria" "Bacteroidetes"       "Bacteroidia"                
## [1735,] "Bacteria" "Bacteroidetes"       "Bacteroidia"                
## [1736,] "Bacteria" "Bacteroidetes"       "Bacteroidia"                
## [1737,] "Bacteria" "Firmicutes"          "Clostridia"                 
## [1738,] "Bacteria" "Bacteroidetes"       "Bacteroidia"                
## [1739,] "Bacteria" "Bacteroidetes"       "Bacteroidia"                
## [1740,] "Bacteria" "Bacteroidetes"       "Bacteroidia"                
## [1741,] "Bacteria" "Firmicutes"          "Negativicutes"              
## [1742,] "Bacteria" "Bacteroidetes"       "Bacteroidia"                
## [1743,] "Bacteria" "Firmicutes"          "Bacilli"                    
## [1744,] "Bacteria" "Bacteroidetes"       "Bacteroidia"                
## [1745,] "Bacteria" "Proteobacteria"      "Gammaproteobacteria"        
## [1746,] "Bacteria" "Bacteroidetes"       "Bacteroidia"                
## [1747,] "Bacteria" "Bacteroidetes"       "Bacteroidia"                
## [1748,] "Bacteria" "Fusobacteria"        "Fusobacteriia"              
## [1749,] "Bacteria" "Bacteroidetes"       "Bacteroidia"                
## [1750,] "Bacteria" "Bacteroidetes"       "Bacteroidia"                
## [1751,] "Bacteria" "Firmicutes"          "Bacilli"                    
## [1752,] "Bacteria" "Firmicutes"          "Negativicutes"              
## [1753,] "Bacteria" "Bacteroidetes"       "Bacteroidia"                
## [1754,] "Bacteria" "Bacteroidetes"       "Bacteroidia"                
## [1755,] "Bacteria" "Firmicutes"          "Negativicutes"              
## [1756,] "Bacteria" "Firmicutes"          "Bacilli"                    
## [1757,] "Bacteria" "Actinobacteria"      "Actinobacteria"             
## [1758,] "Bacteria" "Bacteroidetes"       "Bacteroidia"                
## [1759,] "Bacteria" "Bacteroidetes"       "Bacteroidia"                
## [1760,] "Bacteria" "Patescibacteria"     "Saccharimonadia"            
## [1761,] "Bacteria" "Bacteroidetes"       "Bacteroidia"                
## [1762,] "Bacteria" "Firmicutes"          "Bacilli"                    
## [1763,] "Bacteria" "Firmicutes"          "Bacilli"                    
## [1764,] "Bacteria" "Firmicutes"          "Negativicutes"              
## [1765,] "Bacteria" "Firmicutes"          "Clostridia"                 
## [1766,] "Bacteria" "Bacteroidetes"       "Bacteroidia"                
## [1767,] "Bacteria" "Firmicutes"          "Clostridia"                 
## [1768,] "Bacteria" "Bacteroidetes"       "Bacteroidia"                
## [1769,] "Bacteria" "Firmicutes"          "Negativicutes"              
## [1770,] "Bacteria" "Bacteroidetes"       "Bacteroidia"                
## [1771,] "Bacteria" "Firmicutes"          "Negativicutes"              
## [1772,] "Bacteria" "Proteobacteria"      "Gammaproteobacteria"        
## [1773,] "Bacteria" "Firmicutes"          "Clostridia"                 
## [1774,] "Bacteria" "Actinobacteria"      "Actinobacteria"             
## [1775,] "Bacteria" "Firmicutes"          "Clostridia"                 
## [1776,] "Bacteria" "Firmicutes"          "Bacilli"                    
## [1777,] "Bacteria" "Proteobacteria"      "Gammaproteobacteria"        
## [1778,] "Bacteria" "Proteobacteria"      "Gammaproteobacteria"        
## [1779,] "Bacteria" "Fusobacteria"        "Fusobacteriia"              
## [1780,] "Bacteria" "Bacteroidetes"       "Bacteroidia"                
## [1781,] "Bacteria" "Firmicutes"          "Clostridia"                 
## [1782,] "Bacteria" "Proteobacteria"      "Gammaproteobacteria"        
## [1783,] "Bacteria" "Actinobacteria"      "Actinobacteria"             
## [1784,] "Bacteria" "Bacteroidetes"       "Bacteroidia"                
## [1785,] "Bacteria" "Bacteroidetes"       "Bacteroidia"                
## [1786,] "Bacteria" "Bacteroidetes"       "Bacteroidia"                
## [1787,] "Bacteria" "Firmicutes"          "Negativicutes"              
## [1788,] "Bacteria" "Bacteroidetes"       "Bacteroidia"                
## [1789,] "Bacteria" "Proteobacteria"      "Gammaproteobacteria"        
## [1790,] "Bacteria" "Fusobacteria"        "Fusobacteriia"              
## [1791,] "Bacteria" "Proteobacteria"      "Gammaproteobacteria"        
## [1792,] "Bacteria" "Firmicutes"          NA                           
## [1793,] "Bacteria" "Proteobacteria"      "Gammaproteobacteria"        
## [1794,] "Bacteria" "Firmicutes"          "Clostridia"                 
## [1795,] "Bacteria" "Firmicutes"          NA                           
## [1796,] "Bacteria" "Bacteroidetes"       "Bacteroidia"                
## [1797,] "Bacteria" "Bacteroidetes"       "Bacteroidia"                
## [1798,] "Bacteria" "Firmicutes"          "Clostridia"                 
## [1799,] "Bacteria" "Firmicutes"          "Bacilli"                    
## [1800,] "Bacteria" "Firmicutes"          "Clostridia"                 
## [1801,] "Bacteria" "Firmicutes"          "Clostridia"                 
## [1802,] "Bacteria" "Firmicutes"          "Negativicutes"              
## [1803,] "Bacteria" "Fusobacteria"        "Fusobacteriia"              
## [1804,] "Bacteria" "Firmicutes"          "Clostridia"                 
## [1805,] "Bacteria" "Fusobacteria"        "Fusobacteriia"              
## [1806,] "Bacteria" "Bacteroidetes"       "Bacteroidia"                
## [1807,] "Bacteria" "Fusobacteria"        "Fusobacteriia"              
## [1808,] "Bacteria" "Firmicutes"          "Bacilli"                    
## [1809,] "Bacteria" "Actinobacteria"      "Actinobacteria"             
## [1810,] "Bacteria" "Actinobacteria"      "Actinobacteria"             
## [1811,] "Bacteria" "Firmicutes"          "Negativicutes"              
## [1812,] "Bacteria" "Firmicutes"          "Negativicutes"              
## [1813,] "Bacteria" "Firmicutes"          "Negativicutes"              
## [1814,] "Bacteria" "Fusobacteria"        "Fusobacteriia"              
## [1815,] "Bacteria" "Proteobacteria"      "Gammaproteobacteria"        
## [1816,] "Bacteria" "Proteobacteria"      "Gammaproteobacteria"        
## [1817,] "Bacteria" "Proteobacteria"      "Gammaproteobacteria"        
## [1818,] "Bacteria" "Bacteroidetes"       "Bacteroidia"                
## [1819,] "Bacteria" "Bacteroidetes"       "Bacteroidia"                
## [1820,] "Bacteria" "Bacteroidetes"       "Bacteroidia"                
## [1821,] "Bacteria" "Proteobacteria"      "Gammaproteobacteria"        
## [1822,] "Bacteria" "Bacteroidetes"       "Bacteroidia"                
## [1823,] "Bacteria" "Firmicutes"          "Negativicutes"              
## [1824,] "Bacteria" "Firmicutes"          "Negativicutes"              
## [1825,] "Bacteria" "Firmicutes"          "Bacilli"                    
## [1826,] "Bacteria" "Firmicutes"          "Negativicutes"              
## [1827,] "Bacteria" "Bacteroidetes"       "Bacteroidia"                
## [1828,] "Bacteria" "Proteobacteria"      "Gammaproteobacteria"        
## [1829,] "Bacteria" "Firmicutes"          "Clostridia"                 
## [1830,] "Bacteria" "Fusobacteria"        "Fusobacteriia"              
## [1831,] "Bacteria" "Fusobacteria"        "Fusobacteriia"              
## [1832,] "Bacteria" "Bacteroidetes"       "Bacteroidia"                
## [1833,] "Bacteria" "Actinobacteria"      "Actinobacteria"             
## [1834,] "Bacteria" "Patescibacteria"     "Saccharimonadia"            
## [1835,] "Bacteria" "Bacteroidetes"       "Bacteroidia"                
## [1836,] "Bacteria" "Spirochaetes"        "Spirochaetia"               
## [1837,] "Bacteria" NA                    NA                           
## [1838,] "Bacteria" "Bacteroidetes"       "Bacteroidia"                
## [1839,] "Bacteria" "Fusobacteria"        "Fusobacteriia"              
## [1840,] "Bacteria" "Spirochaetes"        "Spirochaetia"               
## [1841,] "Bacteria" "Firmicutes"          "Negativicutes"              
## [1842,] "Bacteria" "Firmicutes"          "Clostridia"                 
## [1843,] "Bacteria" "Bacteroidetes"       "Bacteroidia"                
## [1844,] "Bacteria" "Bacteroidetes"       "Bacteroidia"                
## [1845,] "Bacteria" "Firmicutes"          "Clostridia"                 
## [1846,] "Bacteria" "Proteobacteria"      "Gammaproteobacteria"        
## [1847,] "Bacteria" "Bacteroidetes"       "Bacteroidia"                
## [1848,] "Bacteria" "Bacteroidetes"       "Bacteroidia"                
## [1849,] "Bacteria" "Bacteroidetes"       "Bacteroidia"                
## [1850,] "Bacteria" "Bacteroidetes"       "Bacteroidia"                
## [1851,] "Bacteria" "Synergistetes"       "Synergistia"                
## [1852,] "Bacteria" "Bacteroidetes"       "Bacteroidia"                
## [1853,] "Bacteria" "Bacteroidetes"       "Bacteroidia"                
## [1854,] "Bacteria" "Proteobacteria"      "Gammaproteobacteria"        
## [1855,] "Bacteria" "Bacteroidetes"       "Bacteroidia"                
## [1856,] "Bacteria" "Proteobacteria"      "Gammaproteobacteria"        
## [1857,] "Bacteria" "Bacteroidetes"       "Bacteroidia"                
## [1858,] "Bacteria" "Bacteroidetes"       "Bacteroidia"                
## [1859,] "Bacteria" "Proteobacteria"      "Gammaproteobacteria"        
## [1860,] "Bacteria" "Firmicutes"          "Clostridia"                 
## [1861,] "Bacteria" "Proteobacteria"      "Gammaproteobacteria"        
## [1862,] "Bacteria" "Bacteroidetes"       "Bacteroidia"                
## [1863,] "Bacteria" "Proteobacteria"      "Gammaproteobacteria"        
## [1864,] "Bacteria" "Fusobacteria"        "Fusobacteriia"              
## [1865,] "Bacteria" "Proteobacteria"      "Gammaproteobacteria"        
## [1866,] "Bacteria" "Bacteroidetes"       "Bacteroidia"                
## [1867,] "Bacteria" "Fusobacteria"        "Fusobacteriia"              
## [1868,] "Bacteria" "Firmicutes"          "Clostridia"                 
## [1869,] "Bacteria" "Proteobacteria"      "Gammaproteobacteria"        
## [1870,] "Bacteria" "Firmicutes"          "Negativicutes"              
## [1871,] "Bacteria" "Proteobacteria"      "Gammaproteobacteria"        
## [1872,] "Bacteria" "Firmicutes"          "Bacilli"                    
## [1873,] "Bacteria" "Proteobacteria"      "Gammaproteobacteria"        
## [1874,] "Bacteria" "Firmicutes"          "Negativicutes"              
## [1875,] "Bacteria" "Fusobacteria"        "Fusobacteriia"              
## [1876,] "Bacteria" "Firmicutes"          "Negativicutes"              
## [1877,] "Bacteria" "Actinobacteria"      "Actinobacteria"             
## [1878,] "Bacteria" "Bacteroidetes"       "Bacteroidia"                
## [1879,] "Bacteria" "Proteobacteria"      "Gammaproteobacteria"        
## [1880,] "Bacteria" "Firmicutes"          "Clostridia"                 
## [1881,] "Bacteria" "Bacteroidetes"       "Bacteroidia"                
## [1882,] "Bacteria" "Firmicutes"          "Negativicutes"              
## [1883,] "Bacteria" "Fusobacteria"        "Fusobacteriia"              
## [1884,] "Bacteria" "Bacteroidetes"       "Bacteroidia"                
## [1885,] "Bacteria" "Actinobacteria"      "Actinobacteria"             
## [1886,] "Bacteria" "Bacteroidetes"       "Bacteroidia"                
## [1887,] "Bacteria" "Firmicutes"          "Negativicutes"              
## [1888,] "Bacteria" "Actinobacteria"      "Actinobacteria"             
## [1889,] "Bacteria" "Bacteroidetes"       "Bacteroidia"                
## [1890,] "Bacteria" "Bacteroidetes"       "Bacteroidia"                
## [1891,] "Bacteria" "Proteobacteria"      "Gammaproteobacteria"        
## [1892,] "Bacteria" "Proteobacteria"      "Gammaproteobacteria"        
## [1893,] "Bacteria" "Firmicutes"          "Negativicutes"              
## [1894,] "Bacteria" "Fusobacteria"        "Fusobacteriia"              
## [1895,] "Bacteria" "Firmicutes"          "Negativicutes"              
## [1896,] "Bacteria" "Firmicutes"          "Clostridia"                 
## [1897,] "Bacteria" "Bacteroidetes"       "Bacteroidia"                
## [1898,] "Bacteria" "Firmicutes"          "Bacilli"                    
## [1899,] "Bacteria" "Proteobacteria"      "Gammaproteobacteria"        
## [1900,] "Bacteria" "Firmicutes"          "Negativicutes"              
## [1901,] "Bacteria" "Firmicutes"          "Clostridia"                 
## [1902,] "Bacteria" "Firmicutes"          "Negativicutes"              
## [1903,] "Bacteria" "Bacteroidetes"       "Bacteroidia"                
## [1904,] "Bacteria" "Bacteroidetes"       "Bacteroidia"                
## [1905,] "Bacteria" "Fusobacteria"        "Fusobacteriia"              
## [1906,] "Bacteria" "Patescibacteria"     "Saccharimonadia"            
## [1907,] "Bacteria" "Firmicutes"          "Clostridia"                 
## [1908,] "Bacteria" "Bacteroidetes"       "Bacteroidia"                
## [1909,] "Bacteria" "Firmicutes"          "Clostridia"                 
## [1910,] "Bacteria" "Fusobacteria"        "Fusobacteriia"              
## [1911,] "Bacteria" "Proteobacteria"      "Gammaproteobacteria"        
## [1912,] "Bacteria" "Bacteroidetes"       "Bacteroidia"                
## [1913,] "Bacteria" "Firmicutes"          "Negativicutes"              
## [1914,] "Bacteria" "Bacteroidetes"       "Bacteroidia"                
## [1915,] "Bacteria" "Bacteroidetes"       "Bacteroidia"                
## [1916,] "Bacteria" "Spirochaetes"        "Spirochaetia"               
## [1917,] "Bacteria" "Bacteroidetes"       "Bacteroidia"                
## [1918,] "Bacteria" "Bacteroidetes"       "Bacteroidia"                
## [1919,] "Bacteria" "Actinobacteria"      "Actinobacteria"             
## [1920,] "Bacteria" "Firmicutes"          "Clostridia"                 
## [1921,] "Bacteria" "Bacteroidetes"       "Bacteroidia"                
## [1922,] "Bacteria" "Firmicutes"          "Clostridia"                 
## [1923,] "Bacteria" "Proteobacteria"      "Gammaproteobacteria"        
## [1924,] "Bacteria" "Proteobacteria"      "Gammaproteobacteria"        
## [1925,] "Bacteria" "Bacteroidetes"       "Bacteroidia"                
## [1926,] "Bacteria" "Bacteroidetes"       "Bacteroidia"                
## [1927,] "Bacteria" "Epsilonbacteraeota"  "Campylobacteria"            
## [1928,] "Bacteria" "Proteobacteria"      "Gammaproteobacteria"        
## [1929,] "Bacteria" "Firmicutes"          "Negativicutes"              
## [1930,] "Bacteria" "Bacteroidetes"       "Bacteroidia"                
## [1931,] "Bacteria" "Fusobacteria"        "Fusobacteriia"              
## [1932,] "Bacteria" "Bacteroidetes"       "Bacteroidia"                
## [1933,] "Bacteria" "Patescibacteria"     "Saccharimonadia"            
## [1934,] "Bacteria" "Bacteroidetes"       "Bacteroidia"                
## [1935,] "Bacteria" "Actinobacteria"      "Actinobacteria"             
## [1936,] "Bacteria" "Actinobacteria"      "Actinobacteria"             
## [1937,] "Bacteria" "Proteobacteria"      "Gammaproteobacteria"        
## [1938,] "Bacteria" "Bacteroidetes"       "Bacteroidia"                
## [1939,] "Bacteria" "Bacteroidetes"       "Bacteroidia"                
## [1940,] "Bacteria" "Fusobacteria"        "Fusobacteriia"              
## [1941,] "Bacteria" "Firmicutes"          "Bacilli"                    
## [1942,] "Bacteria" "Actinobacteria"      "Actinobacteria"             
## [1943,] "Bacteria" "Bacteroidetes"       "Bacteroidia"                
## [1944,] "Bacteria" "Proteobacteria"      "Gammaproteobacteria"        
## [1945,] "Bacteria" "Firmicutes"          "Clostridia"                 
## [1946,] "Bacteria" "Spirochaetes"        "Spirochaetia"               
## [1947,] "Bacteria" "Proteobacteria"      "Gammaproteobacteria"        
## [1948,] "Bacteria" "Fusobacteria"        "Fusobacteriia"              
## [1949,] "Bacteria" "Bacteroidetes"       "Bacteroidia"                
## [1950,] "Bacteria" "Firmicutes"          "Negativicutes"              
## [1951,] "Bacteria" "Bacteroidetes"       "Bacteroidia"                
## [1952,] "Bacteria" "Firmicutes"          "Clostridia"                 
## [1953,] "Bacteria" "Firmicutes"          "Negativicutes"              
## [1954,] "Bacteria" "Proteobacteria"      "Gammaproteobacteria"        
## [1955,] "Bacteria" "Bacteroidetes"       "Bacteroidia"                
## [1956,] "Bacteria" "Bacteroidetes"       "Bacteroidia"                
## [1957,] "Bacteria" "Firmicutes"          "Clostridia"                 
## [1958,] "Bacteria" "Bacteroidetes"       "Bacteroidia"                
## [1959,] "Bacteria" "Bacteroidetes"       "Bacteroidia"                
## [1960,] "Bacteria" "Bacteroidetes"       "Bacteroidia"                
## [1961,] "Bacteria" "Bacteroidetes"       "Bacteroidia"                
## [1962,] "Bacteria" "Actinobacteria"      "Coriobacteriia"             
## [1963,] "Bacteria" "Bacteroidetes"       "Bacteroidia"                
## [1964,] "Bacteria" "Actinobacteria"      "Actinobacteria"             
## [1965,] "Bacteria" "Fusobacteria"        "Fusobacteriia"              
## [1966,] "Bacteria" "Firmicutes"          "Negativicutes"              
## [1967,] "Bacteria" "Bacteroidetes"       "Bacteroidia"                
## [1968,] "Bacteria" "Firmicutes"          "Negativicutes"              
## [1969,] "Bacteria" "Fusobacteria"        "Fusobacteriia"              
## [1970,] "Bacteria" "Proteobacteria"      "Gammaproteobacteria"        
## [1971,] "Bacteria" "Fusobacteria"        "Fusobacteriia"              
## [1972,] "Bacteria" "Firmicutes"          "Clostridia"                 
## [1973,] "Bacteria" "Bacteroidetes"       "Bacteroidia"                
## [1974,] "Bacteria" "Firmicutes"          "Clostridia"                 
## [1975,] "Bacteria" "Firmicutes"          "Clostridia"                 
## [1976,] "Bacteria" "Bacteroidetes"       "Bacteroidia"                
## [1977,] "Bacteria" "Firmicutes"          "Clostridia"                 
## [1978,] "Bacteria" "Bacteroidetes"       "Bacteroidia"                
## [1979,] "Bacteria" "Bacteroidetes"       "Bacteroidia"                
## [1980,] "Bacteria" "Bacteroidetes"       "Bacteroidia"                
## [1981,] "Bacteria" "Bacteroidetes"       "Bacteroidia"                
## [1982,] "Bacteria" "Fusobacteria"        "Fusobacteriia"              
## [1983,] "Bacteria" "Firmicutes"          "Negativicutes"              
## [1984,] "Bacteria" "Proteobacteria"      "Gammaproteobacteria"        
## [1985,] "Bacteria" "Bacteroidetes"       "Bacteroidia"                
## [1986,] "Bacteria" "Firmicutes"          "Negativicutes"              
## [1987,] "Bacteria" "Proteobacteria"      "Gammaproteobacteria"        
## [1988,] "Bacteria" "Actinobacteria"      "Actinobacteria"             
## [1989,] "Bacteria" "Fusobacteria"        "Fusobacteriia"              
## [1990,] "Bacteria" "Firmicutes"          "Bacilli"                    
## [1991,] "Bacteria" "Bacteroidetes"       "Bacteroidia"                
## [1992,] "Bacteria" "Bacteroidetes"       "Bacteroidia"                
## [1993,] "Bacteria" "Bacteroidetes"       "Bacteroidia"                
## [1994,] "Bacteria" "Proteobacteria"      "Gammaproteobacteria"        
## [1995,] "Bacteria" "Firmicutes"          "Negativicutes"              
## [1996,] "Bacteria" "Firmicutes"          "Bacilli"                    
## [1997,] "Bacteria" "Bacteroidetes"       "Bacteroidia"                
## [1998,] "Bacteria" "Firmicutes"          "Negativicutes"              
## [1999,] "Bacteria" "Firmicutes"          "Clostridia"                 
## [2000,] "Bacteria" "Proteobacteria"      "Gammaproteobacteria"        
## [2001,] "Bacteria" "Fusobacteria"        "Fusobacteriia"              
## [2002,] "Bacteria" "Firmicutes"          "Clostridia"                 
## [2003,] "Bacteria" "Fusobacteria"        "Fusobacteriia"              
## [2004,] "Bacteria" "Bacteroidetes"       "Bacteroidia"                
## [2005,] "Bacteria" "Bacteroidetes"       "Bacteroidia"                
## [2006,] "Bacteria" "Firmicutes"          "Clostridia"                 
## [2007,] "Bacteria" "Bacteroidetes"       "Bacteroidia"                
## [2008,] "Bacteria" "Firmicutes"          "Negativicutes"              
## [2009,] "Bacteria" "Bacteroidetes"       "Bacteroidia"                
## [2010,] "Bacteria" "Proteobacteria"      "Gammaproteobacteria"        
## [2011,] "Bacteria" "Bacteroidetes"       "Bacteroidia"                
## [2012,] "Bacteria" "Bacteroidetes"       "Bacteroidia"                
## [2013,] "Bacteria" "Patescibacteria"     "Saccharimonadia"            
## [2014,] "Bacteria" "Bacteroidetes"       "Bacteroidia"                
## [2015,] "Bacteria" "Bacteroidetes"       "Bacteroidia"                
## [2016,] "Bacteria" "Firmicutes"          "Negativicutes"              
## [2017,] "Bacteria" "Bacteroidetes"       "Bacteroidia"                
## [2018,] "Bacteria" "Patescibacteria"     "Saccharimonadia"            
## [2019,] "Bacteria" "Firmicutes"          "Negativicutes"              
## [2020,] "Bacteria" "Fusobacteria"        "Fusobacteriia"              
## [2021,] "Bacteria" "Bacteroidetes"       "Bacteroidia"                
## [2022,] "Bacteria" "Actinobacteria"      "Actinobacteria"             
## [2023,] "Bacteria" "Proteobacteria"      "Gammaproteobacteria"        
## [2024,] "Bacteria" "Proteobacteria"      "Gammaproteobacteria"        
## [2025,] "Bacteria" "Proteobacteria"      "Gammaproteobacteria"        
## [2026,] "Bacteria" "Bacteroidetes"       "Bacteroidia"                
## [2027,] "Bacteria" NA                    NA                           
## [2028,] "Bacteria" "Bacteroidetes"       "Bacteroidia"                
## [2029,] "Bacteria" "Fusobacteria"        "Fusobacteriia"              
## [2030,] "Bacteria" "Fusobacteria"        "Fusobacteriia"              
## [2031,] "Bacteria" "Fusobacteria"        "Fusobacteriia"              
## [2032,] "Bacteria" "Patescibacteria"     "Saccharimonadia"            
## [2033,] "Bacteria" "Fusobacteria"        "Fusobacteriia"              
## [2034,] "Bacteria" "Firmicutes"          "Bacilli"                    
## [2035,] "Bacteria" "Firmicutes"          "Bacilli"                    
## [2036,] "Bacteria" "Firmicutes"          "Bacilli"                    
## [2037,] "Bacteria" "Synergistetes"       "Synergistia"                
## [2038,] "Bacteria" "Bacteroidetes"       "Bacteroidia"                
## [2039,] "Bacteria" "Fusobacteria"        "Fusobacteriia"              
## [2040,] "Bacteria" "Firmicutes"          "Negativicutes"              
## [2041,] "Bacteria" "Firmicutes"          "Negativicutes"              
## [2042,] "Bacteria" "Bacteroidetes"       "Bacteroidia"                
## [2043,] "Bacteria" "Proteobacteria"      "Gammaproteobacteria"        
## [2044,] "Bacteria" "Proteobacteria"      "Gammaproteobacteria"        
## [2045,] "Bacteria" "Patescibacteria"     "Gracilibacteria"            
## [2046,] "Bacteria" "Fusobacteria"        "Fusobacteriia"              
## [2047,] "Bacteria" "Firmicutes"          "Clostridia"                 
## [2048,] "Bacteria" "Firmicutes"          "Negativicutes"              
## [2049,] "Bacteria" "Epsilonbacteraeota"  "Campylobacteria"            
## [2050,] "Bacteria" "Bacteroidetes"       "Bacteroidia"                
## [2051,] "Bacteria" "Bacteroidetes"       "Bacteroidia"                
## [2052,] "Bacteria" "Firmicutes"          "Negativicutes"              
## [2053,] "Bacteria" "Bacteroidetes"       "Bacteroidia"                
## [2054,] "Bacteria" "Actinobacteria"      "Actinobacteria"             
## [2055,] "Bacteria" "Bacteroidetes"       "Bacteroidia"                
## [2056,] "Bacteria" "Firmicutes"          "Negativicutes"              
## [2057,] "Bacteria" "Patescibacteria"     "Saccharimonadia"            
## [2058,] "Bacteria" "Firmicutes"          "Negativicutes"              
## [2059,] "Bacteria" "Bacteroidetes"       "Bacteroidia"                
## [2060,] "Bacteria" "Firmicutes"          "Clostridia"                 
## [2061,] "Bacteria" "Bacteroidetes"       "Bacteroidia"                
## [2062,] "Bacteria" "Firmicutes"          "Negativicutes"              
## [2063,] "Bacteria" "Cyanobacteria"       "Oxyphotobacteria"           
## [2064,] "Bacteria" "Bacteroidetes"       "Bacteroidia"                
## [2065,] "Bacteria" "Firmicutes"          "Negativicutes"              
## [2066,] "Bacteria" "Proteobacteria"      "Gammaproteobacteria"        
## [2067,] "Bacteria" "Bacteroidetes"       "Bacteroidia"                
## [2068,] "Bacteria" "Bacteroidetes"       "Bacteroidia"                
## [2069,] "Bacteria" "Fusobacteria"        "Fusobacteriia"              
## [2070,] "Bacteria" "Patescibacteria"     "Saccharimonadia"            
## [2071,] "Bacteria" "Bacteroidetes"       "Bacteroidia"                
## [2072,] "Bacteria" "Epsilonbacteraeota"  "Campylobacteria"            
## [2073,] "Bacteria" "Firmicutes"          "Clostridia"                 
## [2074,] "Bacteria" "Proteobacteria"      "Gammaproteobacteria"        
## [2075,] "Bacteria" "Fusobacteria"        "Fusobacteriia"              
## [2076,] "Bacteria" "Firmicutes"          "Negativicutes"              
## [2077,] "Bacteria" "Spirochaetes"        "Spirochaetia"               
## [2078,] "Bacteria" "Fusobacteria"        "Fusobacteriia"              
## [2079,] "Bacteria" "Patescibacteria"     "Saccharimonadia"            
## [2080,] "Bacteria" "Bacteroidetes"       "Bacteroidia"                
## [2081,] "Bacteria" "Bacteroidetes"       "Bacteroidia"                
## [2082,] "Bacteria" "Bacteroidetes"       "Bacteroidia"                
## [2083,] "Bacteria" "Fusobacteria"        "Fusobacteriia"              
## [2084,] "Bacteria" NA                    NA                           
## [2085,] "Bacteria" "Proteobacteria"      "Gammaproteobacteria"        
## [2086,] "Bacteria" "Fusobacteria"        "Fusobacteriia"              
## [2087,] "Bacteria" "Firmicutes"          "Clostridia"                 
## [2088,] "Bacteria" "Bacteroidetes"       "Bacteroidia"                
## [2089,] "Bacteria" "Firmicutes"          "Negativicutes"              
## [2090,] "Bacteria" "Bacteroidetes"       "Bacteroidia"                
## [2091,] "Bacteria" "Spirochaetes"        "Spirochaetia"               
## [2092,] "Bacteria" "Actinobacteria"      "Actinobacteria"             
## [2093,] "Bacteria" "Actinobacteria"      "Actinobacteria"             
## [2094,] "Bacteria" "Bacteroidetes"       "Bacteroidia"                
## [2095,] "Bacteria" "Bacteroidetes"       "Bacteroidia"                
## [2096,] "Bacteria" "Fusobacteria"        "Fusobacteriia"              
## [2097,] "Bacteria" "Bacteroidetes"       "Bacteroidia"                
## [2098,] "Bacteria" "Spirochaetes"        "Spirochaetia"               
## [2099,] "Bacteria" "Fusobacteria"        "Fusobacteriia"              
## [2100,] "Bacteria" "Bacteroidetes"       "Bacteroidia"                
## [2101,] "Bacteria" "Bacteroidetes"       "Bacteroidia"                
## [2102,] "Bacteria" "Fusobacteria"        "Fusobacteriia"              
## [2103,] "Bacteria" "Firmicutes"          "Clostridia"                 
## [2104,] "Bacteria" "Bacteroidetes"       "Bacteroidia"                
## [2105,] "Bacteria" "Fusobacteria"        "Fusobacteriia"              
## [2106,] "Bacteria" "Proteobacteria"      "Gammaproteobacteria"        
## [2107,] "Bacteria" "Bacteroidetes"       "Bacteroidia"                
## [2108,] "Bacteria" "Firmicutes"          "Negativicutes"              
## [2109,] "Bacteria" "Bacteroidetes"       "Bacteroidia"                
## [2110,] "Bacteria" "Bacteroidetes"       "Bacteroidia"                
## [2111,] "Bacteria" "Bacteroidetes"       "Bacteroidia"                
## [2112,] "Bacteria" "Firmicutes"          "Clostridia"                 
## [2113,] "Bacteria" "Firmicutes"          "Negativicutes"              
## [2114,] "Bacteria" "Proteobacteria"      "Gammaproteobacteria"        
## [2115,] "Bacteria" "Bacteroidetes"       "Bacteroidia"                
## [2116,] "Bacteria" "Firmicutes"          NA                           
## [2117,] "Bacteria" "Firmicutes"          "Bacilli"                    
## [2118,] "Bacteria" "Bacteroidetes"       "Bacteroidia"                
## [2119,] "Bacteria" "Firmicutes"          "Negativicutes"              
## [2120,] "Bacteria" "Proteobacteria"      "Gammaproteobacteria"        
## [2121,] "Bacteria" "Bacteroidetes"       "Bacteroidia"                
## [2122,] "Bacteria" "Bacteroidetes"       "Bacteroidia"                
## [2123,] "Bacteria" "Patescibacteria"     "Gracilibacteria"            
## [2124,] "Bacteria" "Firmicutes"          "Negativicutes"              
## [2125,] "Bacteria" "Bacteroidetes"       "Bacteroidia"                
## [2126,] "Bacteria" "Firmicutes"          "Bacilli"                    
## [2127,] "Bacteria" "Bacteroidetes"       "Bacteroidia"                
## [2128,] "Bacteria" "Firmicutes"          "Negativicutes"              
## [2129,] "Bacteria" "Firmicutes"          "Bacilli"                    
## [2130,] "Bacteria" "Proteobacteria"      "Gammaproteobacteria"        
## [2131,] "Bacteria" "Bacteroidetes"       "Bacteroidia"                
## [2132,] "Bacteria" "Epsilonbacteraeota"  "Campylobacteria"            
## [2133,] "Bacteria" "Actinobacteria"      "Actinobacteria"             
## [2134,] "Bacteria" "Fusobacteria"        "Fusobacteriia"              
## [2135,] "Bacteria" "Bacteroidetes"       "Bacteroidia"                
## [2136,] "Bacteria" "Firmicutes"          "Negativicutes"              
## [2137,] "Bacteria" "Actinobacteria"      "Actinobacteria"             
## [2138,] "Bacteria" "Bacteroidetes"       "Bacteroidia"                
## [2139,] "Bacteria" "Bacteroidetes"       "Bacteroidia"                
## [2140,] "Bacteria" "Firmicutes"          "Clostridia"                 
## [2141,] "Bacteria" "Spirochaetes"        "Spirochaetia"               
## [2142,] "Bacteria" "Epsilonbacteraeota"  "Campylobacteria"            
## [2143,] "Bacteria" "Proteobacteria"      "Gammaproteobacteria"        
## [2144,] "Bacteria" "Firmicutes"          "Negativicutes"              
## [2145,] "Bacteria" "Bacteroidetes"       "Bacteroidia"                
## [2146,] "Bacteria" "Bacteroidetes"       "Bacteroidia"                
## [2147,] "Bacteria" "Bacteroidetes"       "Bacteroidia"                
## [2148,] "Bacteria" "Bacteroidetes"       "Bacteroidia"                
## [2149,] "Bacteria" "Fusobacteria"        "Fusobacteriia"              
## [2150,] "Bacteria" "Proteobacteria"      "Gammaproteobacteria"        
## [2151,] "Bacteria" "Patescibacteria"     "Saccharimonadia"            
## [2152,] "Bacteria" "Proteobacteria"      "Gammaproteobacteria"        
## [2153,] "Bacteria" "Epsilonbacteraeota"  "Campylobacteria"            
## [2154,] "Bacteria" "Bacteroidetes"       "Bacteroidia"                
## [2155,] "Bacteria" "Bacteroidetes"       "Bacteroidia"                
## [2156,] "Bacteria" "Bacteroidetes"       "Bacteroidia"                
## [2157,] "Bacteria" "Proteobacteria"      "Gammaproteobacteria"        
## [2158,] "Bacteria" NA                    NA                           
## [2159,] "Bacteria" "Bacteroidetes"       "Bacteroidia"                
## [2160,] "Bacteria" "Firmicutes"          "Bacilli"                    
## [2161,] "Bacteria" "Proteobacteria"      "Gammaproteobacteria"        
## [2162,] "Bacteria" "Actinobacteria"      "Actinobacteria"             
## [2163,] "Bacteria" "Fusobacteria"        "Fusobacteriia"              
## [2164,] "Bacteria" "Bacteroidetes"       "Bacteroidia"                
## [2165,] "Bacteria" "Firmicutes"          "Negativicutes"              
## [2166,] "Bacteria" "Bacteroidetes"       "Bacteroidia"                
## [2167,] "Bacteria" "Bacteroidetes"       "Bacteroidia"                
## [2168,] "Bacteria" "Spirochaetes"        "Spirochaetia"               
## [2169,] "Bacteria" "Bacteroidetes"       "Bacteroidia"                
## [2170,] "Bacteria" "Firmicutes"          "Clostridia"                 
## [2171,] "Bacteria" "Fusobacteria"        "Fusobacteriia"              
## [2172,] "Bacteria" "Firmicutes"          "Clostridia"                 
## [2173,] "Bacteria" "Actinobacteria"      "Actinobacteria"             
## [2174,] "Bacteria" "Bacteroidetes"       "Bacteroidia"                
## [2175,] "Bacteria" "Bacteroidetes"       "Bacteroidia"                
## [2176,] "Bacteria" "Fusobacteria"        "Fusobacteriia"              
## [2177,] "Bacteria" "Bacteroidetes"       "Bacteroidia"                
## [2178,] "Bacteria" "Firmicutes"          "Clostridia"                 
## [2179,] "Bacteria" "Firmicutes"          "Negativicutes"              
## [2180,] "Bacteria" "Firmicutes"          "Negativicutes"              
## [2181,] "Bacteria" "Fusobacteria"        "Fusobacteriia"              
## [2182,] "Bacteria" "Firmicutes"          "Clostridia"                 
## [2183,] "Bacteria" "Bacteroidetes"       "Bacteroidia"                
## [2184,] "Bacteria" "Bacteroidetes"       "Bacteroidia"                
## [2185,] "Bacteria" "Bacteroidetes"       "Bacteroidia"                
## [2186,] "Bacteria" "Fusobacteria"        "Fusobacteriia"              
## [2187,] "Bacteria" "Firmicutes"          "Negativicutes"              
## [2188,] "Bacteria" "Firmicutes"          "Negativicutes"              
## [2189,] "Bacteria" "Fusobacteria"        "Fusobacteriia"              
## [2190,] "Bacteria" "Firmicutes"          "Negativicutes"              
## [2191,] "Bacteria" "Bacteroidetes"       "Bacteroidia"                
## [2192,] "Bacteria" "Proteobacteria"      "Gammaproteobacteria"        
## [2193,] "Bacteria" "Bacteroidetes"       "Bacteroidia"                
## [2194,] "Bacteria" "Firmicutes"          "Negativicutes"              
## [2195,] "Bacteria" "Proteobacteria"      "Gammaproteobacteria"        
## [2196,] "Bacteria" "Patescibacteria"     "Saccharimonadia"            
## [2197,] "Bacteria" "Bacteroidetes"       "Bacteroidia"                
## [2198,] "Bacteria" "Fusobacteria"        "Fusobacteriia"              
## [2199,] "Bacteria" "Bacteroidetes"       "Bacteroidia"                
## [2200,] "Bacteria" "Firmicutes"          "Negativicutes"              
## [2201,] "Bacteria" "Actinobacteria"      "Actinobacteria"             
## [2202,] "Bacteria" "Bacteroidetes"       "Bacteroidia"                
## [2203,] "Bacteria" "Firmicutes"          "Negativicutes"              
## [2204,] "Bacteria" "Fusobacteria"        "Fusobacteriia"              
## [2205,] "Bacteria" "Firmicutes"          "Negativicutes"              
## [2206,] "Bacteria" "Proteobacteria"      "Gammaproteobacteria"        
## [2207,] "Bacteria" "Bacteroidetes"       "Bacteroidia"                
## [2208,] "Bacteria" "Proteobacteria"      "Gammaproteobacteria"        
## [2209,] "Bacteria" "Patescibacteria"     "Saccharimonadia"            
## [2210,] "Bacteria" "Spirochaetes"        "Spirochaetia"               
## [2211,] "Bacteria" "Firmicutes"          "Clostridia"                 
## [2212,] "Bacteria" "Bacteroidetes"       "Bacteroidia"                
## [2213,] "Bacteria" "Bacteroidetes"       "Bacteroidia"                
## [2214,] "Bacteria" "Firmicutes"          "Negativicutes"              
## [2215,] "Bacteria" "Bacteroidetes"       "Bacteroidia"                
## [2216,] "Bacteria" "Bacteroidetes"       "Bacteroidia"                
## [2217,] "Bacteria" "Bacteroidetes"       "Bacteroidia"                
## [2218,] "Bacteria" "Firmicutes"          "Negativicutes"              
## [2219,] "Bacteria" "Bacteroidetes"       "Bacteroidia"                
## [2220,] "Bacteria" "Bacteroidetes"       "Bacteroidia"                
## [2221,] "Bacteria" "Patescibacteria"     "Saccharimonadia"            
## [2222,] "Bacteria" "Bacteroidetes"       "Bacteroidia"                
## [2223,] "Bacteria" "Cyanobacteria"       "Oxyphotobacteria"           
## [2224,] "Bacteria" "Bacteroidetes"       "Bacteroidia"                
## [2225,] "Bacteria" "Firmicutes"          "Negativicutes"              
## [2226,] "Bacteria" "Patescibacteria"     "Saccharimonadia"            
## [2227,] "Bacteria" "Firmicutes"          "Bacilli"                    
## [2228,] "Bacteria" "Bacteroidetes"       "Bacteroidia"                
## [2229,] "Bacteria" "Firmicutes"          "Clostridia"                 
## [2230,] "Bacteria" NA                    NA                           
## [2231,] "Bacteria" "Actinobacteria"      "Actinobacteria"             
## [2232,] "Bacteria" "Firmicutes"          "Negativicutes"              
## [2233,] "Bacteria" "Bacteroidetes"       "Bacteroidia"                
## [2234,] "Bacteria" "Firmicutes"          "Negativicutes"              
## [2235,] "Bacteria" "Firmicutes"          "Clostridia"                 
## [2236,] "Bacteria" "Firmicutes"          "Negativicutes"              
## [2237,] "Bacteria" "Bacteroidetes"       "Bacteroidia"                
## [2238,] "Bacteria" "Fusobacteria"        "Fusobacteriia"              
## [2239,] "Bacteria" "Firmicutes"          "Negativicutes"              
## [2240,] "Bacteria" "Actinobacteria"      "Actinobacteria"             
## [2241,] "Bacteria" "Bacteroidetes"       "Bacteroidia"                
## [2242,] "Bacteria" "Cyanobacteria"       "Oxyphotobacteria"           
## [2243,] "Bacteria" "Bacteroidetes"       "Bacteroidia"                
## [2244,] "Bacteria" "Bacteroidetes"       "Bacteroidia"                
## [2245,] "Bacteria" "Proteobacteria"      "Gammaproteobacteria"        
## [2246,] "Bacteria" "Bacteroidetes"       "Bacteroidia"                
## [2247,] "Bacteria" "Bacteroidetes"       "Bacteroidia"                
## [2248,] "Bacteria" "Bacteroidetes"       "Bacteroidia"                
## [2249,] "Bacteria" NA                    NA                           
## [2250,] "Bacteria" "Patescibacteria"     "Saccharimonadia"            
## [2251,] "Bacteria" "Actinobacteria"      "Coriobacteriia"             
## [2252,] "Bacteria" "Bacteroidetes"       "Bacteroidia"                
## [2253,] "Bacteria" "Firmicutes"          "Negativicutes"              
## [2254,] "Bacteria" "Firmicutes"          "Negativicutes"              
## [2255,] "Bacteria" "Synergistetes"       "Synergistia"                
## [2256,] "Bacteria" "Bacteroidetes"       "Bacteroidia"                
## [2257,] "Bacteria" "Bacteroidetes"       "Bacteroidia"                
## [2258,] "Bacteria" "Firmicutes"          "Clostridia"                 
## [2259,] "Bacteria" "Actinobacteria"      "Actinobacteria"             
## [2260,] "Bacteria" "Firmicutes"          "Bacilli"                    
## [2261,] "Bacteria" "Firmicutes"          "Negativicutes"              
## [2262,] "Bacteria" "Proteobacteria"      "Gammaproteobacteria"        
## [2263,] "Bacteria" "Proteobacteria"      "Gammaproteobacteria"        
## [2264,] "Bacteria" "Fusobacteria"        "Fusobacteriia"              
## [2265,] "Bacteria" "Bacteroidetes"       "Bacteroidia"                
## [2266,] "Bacteria" "Fusobacteria"        "Fusobacteriia"              
## [2267,] "Bacteria" "Firmicutes"          "Negativicutes"              
## [2268,] "Bacteria" "Proteobacteria"      "Gammaproteobacteria"        
## [2269,] "Bacteria" "Bacteroidetes"       "Bacteroidia"                
## [2270,] "Bacteria" "Bacteroidetes"       "Bacteroidia"                
## [2271,] "Bacteria" "Fusobacteria"        "Fusobacteriia"              
## [2272,] "Bacteria" "Patescibacteria"     "Gracilibacteria"            
## [2273,] "Bacteria" "Fusobacteria"        "Fusobacteriia"              
## [2274,] "Bacteria" "Firmicutes"          "Clostridia"                 
## [2275,] "Bacteria" "Bacteroidetes"       "Bacteroidia"                
## [2276,] "Bacteria" "Bacteroidetes"       "Bacteroidia"                
## [2277,] "Bacteria" "Firmicutes"          "Clostridia"                 
## [2278,] "Bacteria" "Firmicutes"          "Negativicutes"              
## [2279,] "Bacteria" "Proteobacteria"      "Gammaproteobacteria"        
## [2280,] "Bacteria" "Firmicutes"          NA                           
## [2281,] "Bacteria" "Proteobacteria"      "Gammaproteobacteria"        
## [2282,] "Bacteria" "Firmicutes"          "Clostridia"                 
## [2283,] "Bacteria" "Bacteroidetes"       "Bacteroidia"                
## [2284,] "Bacteria" "Bacteroidetes"       "Bacteroidia"                
## [2285,] "Bacteria" "Bacteroidetes"       "Bacteroidia"                
## [2286,] "Bacteria" "Firmicutes"          "Clostridia"                 
## [2287,] "Bacteria" "Bacteroidetes"       "Bacteroidia"                
## [2288,] "Bacteria" "Bacteroidetes"       "Bacteroidia"                
## [2289,] "Bacteria" "Patescibacteria"     "Gracilibacteria"            
## [2290,] "Bacteria" "Fusobacteria"        "Fusobacteriia"              
## [2291,] "Bacteria" "Firmicutes"          "Negativicutes"              
## [2292,] "Bacteria" "Proteobacteria"      "Gammaproteobacteria"        
## [2293,] "Bacteria" "Bacteroidetes"       "Bacteroidia"                
## [2294,] "Bacteria" "Bacteroidetes"       "Bacteroidia"                
## [2295,] "Bacteria" "Bacteroidetes"       "Bacteroidia"                
## [2296,] "Bacteria" "Proteobacteria"      "Gammaproteobacteria"        
## [2297,] "Bacteria" "Bacteroidetes"       "Bacteroidia"                
## [2298,] "Bacteria" "Bacteroidetes"       "Bacteroidia"                
## [2299,] "Bacteria" "Bacteroidetes"       "Bacteroidia"                
## [2300,] "Bacteria" "Bacteroidetes"       "Bacteroidia"                
## [2301,] "Bacteria" "Firmicutes"          "Negativicutes"              
## [2302,] "Bacteria" "Firmicutes"          "Clostridia"                 
## [2303,] "Bacteria" "Actinobacteria"      "Actinobacteria"             
## [2304,] "Bacteria" "Firmicutes"          "Clostridia"                 
## [2305,] "Bacteria" "Spirochaetes"        "Spirochaetia"               
## [2306,] "Bacteria" "Bacteroidetes"       "Bacteroidia"                
## [2307,] "Bacteria" "Patescibacteria"     "Saccharimonadia"            
## [2308,] "Bacteria" "Bacteroidetes"       "Bacteroidia"                
## [2309,] "Bacteria" "Firmicutes"          "Clostridia"                 
## [2310,] "Bacteria" "Bacteroidetes"       "Bacteroidia"                
## [2311,] "Bacteria" "Firmicutes"          "Clostridia"                 
## [2312,] "Bacteria" "Firmicutes"          "Negativicutes"              
## [2313,] "Bacteria" "Actinobacteria"      "Actinobacteria"             
## [2314,] "Bacteria" "Bacteroidetes"       "Bacteroidia"                
## [2315,] "Bacteria" "Fusobacteria"        "Fusobacteriia"              
## [2316,] "Bacteria" "Actinobacteria"      "Actinobacteria"             
## [2317,] "Bacteria" "Firmicutes"          NA                           
## [2318,] "Bacteria" "Bacteroidetes"       "Bacteroidia"                
## [2319,] "Bacteria" "Firmicutes"          "Clostridia"                 
## [2320,] "Bacteria" "Patescibacteria"     "Saccharimonadia"            
## [2321,] "Bacteria" "Bacteroidetes"       "Bacteroidia"                
## [2322,] "Bacteria" "Bacteroidetes"       "Bacteroidia"                
## [2323,] "Bacteria" "Bacteroidetes"       "Bacteroidia"                
## [2324,] "Bacteria" "Bacteroidetes"       "Bacteroidia"                
## [2325,] "Bacteria" "Bacteroidetes"       "Bacteroidia"                
## [2326,] "Bacteria" "Firmicutes"          NA                           
## [2327,] "Bacteria" "Fusobacteria"        "Fusobacteriia"              
## [2328,] "Bacteria" "Bacteroidetes"       "Bacteroidia"                
## [2329,] "Bacteria" "Firmicutes"          "Clostridia"                 
## [2330,] "Bacteria" "Bacteroidetes"       "Bacteroidia"                
## [2331,] "Bacteria" "Bacteroidetes"       "Bacteroidia"                
## [2332,] "Bacteria" "Bacteroidetes"       "Bacteroidia"                
## [2333,] "Bacteria" "Bacteroidetes"       "Bacteroidia"                
## [2334,] "Bacteria" "Bacteroidetes"       "Bacteroidia"                
## [2335,] "Bacteria" "Proteobacteria"      "Gammaproteobacteria"        
## [2336,] "Bacteria" "Proteobacteria"      "Gammaproteobacteria"        
## [2337,] "Bacteria" "Bacteroidetes"       "Bacteroidia"                
## [2338,] "Bacteria" "Bacteroidetes"       "Bacteroidia"                
## [2339,] "Bacteria" "Firmicutes"          "Bacilli"                    
## [2340,] "Bacteria" "Bacteroidetes"       "Bacteroidia"                
## [2341,] "Bacteria" "Patescibacteria"     "Saccharimonadia"            
## [2342,] "Bacteria" "Firmicutes"          "Clostridia"                 
## [2343,] "Bacteria" "Proteobacteria"      "Gammaproteobacteria"        
## [2344,] "Bacteria" "Bacteroidetes"       "Bacteroidia"                
## [2345,] "Bacteria" "Bacteroidetes"       "Bacteroidia"                
## [2346,] "Bacteria" "Actinobacteria"      "Actinobacteria"             
## [2347,] "Bacteria" "Firmicutes"          "Negativicutes"              
## [2348,] "Bacteria" "Actinobacteria"      "Actinobacteria"             
## [2349,] "Bacteria" "Synergistetes"       "Synergistia"                
## [2350,] "Bacteria" "Spirochaetes"        "Spirochaetia"               
## [2351,] "Bacteria" "Bacteroidetes"       "Bacteroidia"                
## [2352,] "Bacteria" "Bacteroidetes"       "Bacteroidia"                
## [2353,] "Bacteria" "Actinobacteria"      "Actinobacteria"             
## [2354,] "Bacteria" "Actinobacteria"      "Actinobacteria"             
## [2355,] "Bacteria" "Firmicutes"          "Clostridia"                 
## [2356,] "Bacteria" "Bacteroidetes"       "Bacteroidia"                
## [2357,] "Bacteria" "Firmicutes"          "Negativicutes"              
## [2358,] "Bacteria" "Bacteroidetes"       "Bacteroidia"                
## [2359,] "Bacteria" "Bacteroidetes"       "Bacteroidia"                
## [2360,] "Bacteria" "Firmicutes"          "Negativicutes"              
## [2361,] "Bacteria" "Proteobacteria"      "Gammaproteobacteria"        
## [2362,] "Bacteria" "Firmicutes"          "Negativicutes"              
## [2363,] "Bacteria" "Bacteroidetes"       "Bacteroidia"                
## [2364,] "Bacteria" "Firmicutes"          "Negativicutes"              
## [2365,] "Bacteria" "Firmicutes"          "Negativicutes"              
## [2366,] "Bacteria" "Fusobacteria"        "Fusobacteriia"              
## [2367,] "Bacteria" "Actinobacteria"      "Actinobacteria"             
## [2368,] "Bacteria" "Bacteroidetes"       "Bacteroidia"                
## [2369,] "Bacteria" "Proteobacteria"      "Gammaproteobacteria"        
## [2370,] "Bacteria" "Fusobacteria"        "Fusobacteriia"              
## [2371,] "Bacteria" "Bacteroidetes"       "Bacteroidia"                
## [2372,] "Bacteria" "Firmicutes"          "Negativicutes"              
## [2373,] "Bacteria" "Bacteroidetes"       "Bacteroidia"                
## [2374,] "Bacteria" "Bacteroidetes"       "Bacteroidia"                
## [2375,] "Bacteria" "Firmicutes"          "Negativicutes"              
## [2376,] "Bacteria" "Firmicutes"          "Clostridia"                 
## [2377,] "Bacteria" "Proteobacteria"      "Gammaproteobacteria"        
## [2378,] "Bacteria" "Spirochaetes"        "Spirochaetia"               
## [2379,] "Bacteria" "Bacteroidetes"       "Bacteroidia"                
## [2380,] "Bacteria" "Fusobacteria"        "Fusobacteriia"              
## [2381,] "Bacteria" "Bacteroidetes"       "Bacteroidia"                
## [2382,] "Bacteria" "Fusobacteria"        "Fusobacteriia"              
## [2383,] "Bacteria" "Firmicutes"          "Clostridia"                 
## [2384,] "Bacteria" "Bacteroidetes"       "Bacteroidia"                
## [2385,] "Bacteria" "Proteobacteria"      "Gammaproteobacteria"        
## [2386,] "Bacteria" "Bacteroidetes"       "Bacteroidia"                
## [2387,] "Bacteria" "Firmicutes"          "Negativicutes"              
## [2388,] "Bacteria" "Actinobacteria"      "Actinobacteria"             
## [2389,] "Bacteria" "Bacteroidetes"       "Bacteroidia"                
## [2390,] "Bacteria" "Firmicutes"          "Negativicutes"              
## [2391,] "Bacteria" "Fusobacteria"        "Fusobacteriia"              
## [2392,] "Bacteria" "Firmicutes"          "Negativicutes"              
## [2393,] "Bacteria" "Fusobacteria"        "Fusobacteriia"              
## [2394,] "Bacteria" "Patescibacteria"     "Saccharimonadia"            
## [2395,] "Bacteria" "Firmicutes"          "Negativicutes"              
## [2396,] "Bacteria" "Firmicutes"          "Negativicutes"              
## [2397,] "Bacteria" "Firmicutes"          "Negativicutes"              
## [2398,] "Bacteria" "Actinobacteria"      "Actinobacteria"             
## [2399,] "Bacteria" "Actinobacteria"      "Actinobacteria"             
## [2400,] "Bacteria" "Bacteroidetes"       "Bacteroidia"                
## [2401,] "Bacteria" "Actinobacteria"      "Coriobacteriia"             
## [2402,] "Bacteria" "Spirochaetes"        "Spirochaetia"               
## [2403,] "Bacteria" "Bacteroidetes"       "Bacteroidia"                
## [2404,] "Bacteria" "Fusobacteria"        "Fusobacteriia"              
## [2405,] "Bacteria" "Firmicutes"          "Clostridia"                 
## [2406,] "Bacteria" "Firmicutes"          "Negativicutes"              
## [2407,] "Bacteria" "Bacteroidetes"       "Bacteroidia"                
## [2408,] "Bacteria" "Bacteroidetes"       "Bacteroidia"                
## [2409,] "Bacteria" "Fusobacteria"        "Fusobacteriia"              
## [2410,] "Bacteria" "Bacteroidetes"       "Bacteroidia"                
## [2411,] "Bacteria" "Firmicutes"          "Negativicutes"              
## [2412,] "Bacteria" "Firmicutes"          NA                           
## [2413,] "Bacteria" "Patescibacteria"     "Saccharimonadia"            
## [2414,] "Bacteria" "Bacteroidetes"       "Bacteroidia"                
## [2415,] "Bacteria" "Firmicutes"          "Negativicutes"              
## [2416,] "Bacteria" "Firmicutes"          "Negativicutes"              
## [2417,] "Bacteria" "Firmicutes"          "Negativicutes"              
## [2418,] "Bacteria" "Bacteroidetes"       "Bacteroidia"                
## [2419,] "Bacteria" "Bacteroidetes"       "Bacteroidia"                
## [2420,] "Bacteria" "Firmicutes"          "Clostridia"                 
## [2421,] "Bacteria" "Proteobacteria"      "Gammaproteobacteria"        
## [2422,] "Bacteria" "Actinobacteria"      "Actinobacteria"             
## [2423,] "Bacteria" "Actinobacteria"      "Actinobacteria"             
## [2424,] "Bacteria" "Firmicutes"          NA                           
## [2425,] "Bacteria" "Bacteroidetes"       "Bacteroidia"                
## [2426,] "Bacteria" "Cyanobacteria"       "Oxyphotobacteria"           
## [2427,] "Bacteria" "Bacteroidetes"       "Bacteroidia"                
## [2428,] "Bacteria" "Bacteroidetes"       "Bacteroidia"                
## [2429,] "Bacteria" "Bacteroidetes"       "Bacteroidia"                
## [2430,] "Bacteria" "Bacteroidetes"       "Bacteroidia"                
## [2431,] "Bacteria" "Bacteroidetes"       "Bacteroidia"                
## [2432,] "Bacteria" "Firmicutes"          "Clostridia"                 
## [2433,] "Bacteria" "Firmicutes"          "Negativicutes"              
## [2434,] "Bacteria" "Bacteroidetes"       "Bacteroidia"                
## [2435,] "Bacteria" "Actinobacteria"      "Actinobacteria"             
## [2436,] "Bacteria" "Bacteroidetes"       "Bacteroidia"                
## [2437,] "Bacteria" "Spirochaetes"        "Spirochaetia"               
## [2438,] "Bacteria" "Firmicutes"          NA                           
## [2439,] "Bacteria" "Bacteroidetes"       "Bacteroidia"                
## [2440,] "Bacteria" "Cyanobacteria"       "Oxyphotobacteria"           
## [2441,] "Bacteria" "Actinobacteria"      "Actinobacteria"             
## [2442,] "Bacteria" "Bacteroidetes"       "Bacteroidia"                
## [2443,] "Bacteria" "Proteobacteria"      "Gammaproteobacteria"        
## [2444,] "Bacteria" "Proteobacteria"      "Gammaproteobacteria"        
## [2445,] "Bacteria" "Firmicutes"          "Clostridia"                 
## [2446,] "Bacteria" "Bacteroidetes"       "Bacteroidia"                
## [2447,] "Bacteria" "Bacteroidetes"       "Bacteroidia"                
## [2448,] "Bacteria" "Firmicutes"          "Bacilli"                    
## [2449,] "Bacteria" "Patescibacteria"     "Saccharimonadia"            
## [2450,] "Bacteria" "Bacteroidetes"       "Bacteroidia"                
## [2451,] "Bacteria" "Firmicutes"          NA                           
## [2452,] "Bacteria" "Bacteroidetes"       "Bacteroidia"                
## [2453,] "Bacteria" "Proteobacteria"      "Gammaproteobacteria"        
## [2454,] "Bacteria" "Bacteroidetes"       "Bacteroidia"                
## [2455,] "Bacteria" "Firmicutes"          "Clostridia"                 
## [2456,] "Bacteria" "Actinobacteria"      "Actinobacteria"             
## [2457,] "Bacteria" "Proteobacteria"      "Gammaproteobacteria"        
## [2458,] "Bacteria" "Bacteroidetes"       "Bacteroidia"                
## [2459,] "Bacteria" "Firmicutes"          "Clostridia"                 
## [2460,] "Bacteria" "Spirochaetes"        "Spirochaetia"               
## [2461,] "Bacteria" "Firmicutes"          "Clostridia"                 
## [2462,] "Bacteria" "Firmicutes"          "Negativicutes"              
## [2463,] "Bacteria" "Bacteroidetes"       "Bacteroidia"                
## [2464,] "Bacteria" "Firmicutes"          "Clostridia"                 
## [2465,] "Bacteria" "Bacteroidetes"       "Bacteroidia"                
## [2466,] "Bacteria" "Bacteroidetes"       "Bacteroidia"                
## [2467,] "Bacteria" "Proteobacteria"      "Gammaproteobacteria"        
## [2468,] "Bacteria" "Bacteroidetes"       "Bacteroidia"                
## [2469,] "Bacteria" "Bacteroidetes"       "Bacteroidia"                
## [2470,] "Bacteria" "Bacteroidetes"       "Bacteroidia"                
## [2471,] "Bacteria" "Proteobacteria"      "Gammaproteobacteria"        
## [2472,] "Bacteria" "Proteobacteria"      "Gammaproteobacteria"        
## [2473,] "Bacteria" "Firmicutes"          "Negativicutes"              
## [2474,] "Bacteria" "Firmicutes"          "Clostridia"                 
## [2475,] "Bacteria" "Bacteroidetes"       "Bacteroidia"                
## [2476,] "Bacteria" "Bacteroidetes"       "Bacteroidia"                
## [2477,] "Bacteria" "Patescibacteria"     "Saccharimonadia"            
## [2478,] "Bacteria" "Firmicutes"          "Negativicutes"              
## [2479,] "Bacteria" "Firmicutes"          "Negativicutes"              
## [2480,] "Bacteria" "Actinobacteria"      "Actinobacteria"             
## [2481,] "Bacteria" "Actinobacteria"      "Actinobacteria"             
## [2482,] "Bacteria" "Firmicutes"          "Clostridia"                 
## [2483,] "Bacteria" "Bacteroidetes"       "Bacteroidia"                
## [2484,] "Bacteria" "Spirochaetes"        "Spirochaetia"               
## [2485,] "Bacteria" "Proteobacteria"      "Gammaproteobacteria"        
## [2486,] "Bacteria" "Firmicutes"          "Clostridia"                 
## [2487,] "Bacteria" "Bacteroidetes"       "Bacteroidia"                
## [2488,] "Bacteria" "Bacteroidetes"       "Bacteroidia"                
## [2489,] "Bacteria" "Proteobacteria"      "Gammaproteobacteria"        
## [2490,] "Bacteria" "Bacteroidetes"       "Bacteroidia"                
## [2491,] "Bacteria" "Fusobacteria"        "Fusobacteriia"              
## [2492,] "Bacteria" "Firmicutes"          "Negativicutes"              
## [2493,] "Bacteria" "Bacteroidetes"       "Bacteroidia"                
## [2494,] "Bacteria" "Bacteroidetes"       "Bacteroidia"                
## [2495,] "Bacteria" "Bacteroidetes"       "Bacteroidia"                
## [2496,] "Bacteria" "Fusobacteria"        "Fusobacteriia"              
## [2497,] "Bacteria" "Actinobacteria"      "Actinobacteria"             
## [2498,] "Bacteria" "Proteobacteria"      "Gammaproteobacteria"        
## [2499,] "Bacteria" "Bacteroidetes"       "Bacteroidia"                
## [2500,] "Bacteria" "Actinobacteria"      "Actinobacteria"             
## [2501,] "Bacteria" "Firmicutes"          "Negativicutes"              
## [2502,] "Bacteria" "Bacteroidetes"       "Bacteroidia"                
## [2503,] "Bacteria" "Bacteroidetes"       "Bacteroidia"                
## [2504,] "Bacteria" "Proteobacteria"      "Gammaproteobacteria"        
## [2505,] "Bacteria" "Fusobacteria"        "Fusobacteriia"              
## [2506,] "Bacteria" "Bacteroidetes"       "Bacteroidia"                
## [2507,] "Bacteria" "Bacteroidetes"       "Bacteroidia"                
## [2508,] "Bacteria" "Bacteroidetes"       "Bacteroidia"                
## [2509,] "Bacteria" "Firmicutes"          "Bacilli"                    
## [2510,] "Bacteria" "Patescibacteria"     "Saccharimonadia"            
## [2511,] "Bacteria" "Bacteroidetes"       "Bacteroidia"                
## [2512,] "Bacteria" "Proteobacteria"      "Gammaproteobacteria"        
## [2513,] "Bacteria" "Firmicutes"          "Clostridia"                 
## [2514,] "Bacteria" "Firmicutes"          "Clostridia"                 
## [2515,] "Bacteria" "Bacteroidetes"       "Bacteroidia"                
## [2516,] "Bacteria" "Proteobacteria"      "Gammaproteobacteria"        
## [2517,] "Bacteria" "Actinobacteria"      "Actinobacteria"             
## [2518,] "Bacteria" "Fusobacteria"        "Fusobacteriia"              
## [2519,] "Bacteria" "Actinobacteria"      "Actinobacteria"             
## [2520,] "Bacteria" "Actinobacteria"      "Actinobacteria"             
## [2521,] "Bacteria" "Proteobacteria"      "Gammaproteobacteria"        
## [2522,] "Bacteria" "Fusobacteria"        "Fusobacteriia"              
## [2523,] "Bacteria" "Bacteroidetes"       "Bacteroidia"                
## [2524,] "Bacteria" "Firmicutes"          "Negativicutes"              
## [2525,] "Bacteria" "Proteobacteria"      "Gammaproteobacteria"        
## [2526,] "Bacteria" "Actinobacteria"      "Actinobacteria"             
## [2527,] "Bacteria" "Bacteroidetes"       "Bacteroidia"                
## [2528,] "Bacteria" "Firmicutes"          "Bacilli"                    
## [2529,] "Bacteria" "Fusobacteria"        "Fusobacteriia"              
## [2530,] "Bacteria" "Bacteroidetes"       "Bacteroidia"                
## [2531,] "Bacteria" "Firmicutes"          "Negativicutes"              
## [2532,] "Bacteria" "Firmicutes"          "Negativicutes"              
## [2533,] "Bacteria" "Bacteroidetes"       "Bacteroidia"                
## [2534,] "Bacteria" "Bacteroidetes"       "Bacteroidia"                
## [2535,] "Bacteria" "Bacteroidetes"       "Bacteroidia"                
## [2536,] "Bacteria" "Proteobacteria"      "Gammaproteobacteria"        
## [2537,] "Bacteria" "Firmicutes"          "Bacilli"                    
## [2538,] "Bacteria" "Bacteroidetes"       "Bacteroidia"                
## [2539,] "Bacteria" "Actinobacteria"      "Actinobacteria"             
## [2540,] "Bacteria" "Proteobacteria"      "Gammaproteobacteria"        
## [2541,] "Bacteria" "Bacteroidetes"       "Bacteroidia"                
## [2542,] "Bacteria" "Firmicutes"          "Clostridia"                 
## [2543,] "Bacteria" "Fusobacteria"        "Fusobacteriia"              
## [2544,] "Bacteria" "Bacteroidetes"       "Bacteroidia"                
## [2545,] "Bacteria" "Tenericutes"         "Mollicutes"                 
## [2546,] "Bacteria" "Fusobacteria"        "Fusobacteriia"              
## [2547,] "Bacteria" "Fusobacteria"        "Fusobacteriia"              
## [2548,] "Bacteria" "Actinobacteria"      "Actinobacteria"             
## [2549,] "Bacteria" "Firmicutes"          "Negativicutes"              
## [2550,] "Bacteria" "Firmicutes"          "Clostridia"                 
## [2551,] "Bacteria" "Firmicutes"          "Negativicutes"              
## [2552,] "Bacteria" "Bacteroidetes"       "Bacteroidia"                
## [2553,] "Bacteria" "Fusobacteria"        "Fusobacteriia"              
## [2554,] "Bacteria" "Fusobacteria"        "Fusobacteriia"              
## [2555,] "Bacteria" "Fusobacteria"        "Fusobacteriia"              
## [2556,] "Bacteria" "Bacteroidetes"       "Bacteroidia"                
## [2557,] "Bacteria" "Proteobacteria"      "Gammaproteobacteria"        
## [2558,] "Bacteria" "Spirochaetes"        "Spirochaetia"               
## [2559,] "Bacteria" "Bacteroidetes"       "Bacteroidia"                
## [2560,] "Bacteria" "Firmicutes"          "Negativicutes"              
## [2561,] "Bacteria" "Firmicutes"          "Negativicutes"              
## [2562,] "Bacteria" "Spirochaetes"        "Spirochaetia"               
## [2563,] "Bacteria" "Spirochaetes"        "Spirochaetia"               
## [2564,] "Bacteria" "Proteobacteria"      "Gammaproteobacteria"        
## [2565,] "Bacteria" "Firmicutes"          "Clostridia"                 
## [2566,] "Bacteria" "Firmicutes"          "Bacilli"                    
## [2567,] "Bacteria" "Proteobacteria"      "Gammaproteobacteria"        
## [2568,] "Bacteria" "Firmicutes"          "Bacilli"                    
## [2569,] "Bacteria" "Bacteroidetes"       "Bacteroidia"                
## [2570,] "Bacteria" "Firmicutes"          "Negativicutes"              
## [2571,] "Bacteria" "Bacteroidetes"       "Bacteroidia"                
## [2572,] "Bacteria" "Bacteroidetes"       "Bacteroidia"                
## [2573,] "Bacteria" "Proteobacteria"      "Gammaproteobacteria"        
## [2574,] "Bacteria" "Fusobacteria"        "Fusobacteriia"              
## [2575,] "Bacteria" "Patescibacteria"     "Saccharimonadia"            
## [2576,] "Bacteria" "Bacteroidetes"       "Bacteroidia"                
## [2577,] "Bacteria" "Actinobacteria"      "Actinobacteria"             
## [2578,] "Bacteria" "Fusobacteria"        "Fusobacteriia"              
## [2579,] "Bacteria" "Bacteroidetes"       "Bacteroidia"                
## [2580,] "Bacteria" "Bacteroidetes"       "Bacteroidia"                
## [2581,] "Bacteria" "Bacteroidetes"       "Bacteroidia"                
## [2582,] "Bacteria" "Fusobacteria"        "Fusobacteriia"              
## [2583,] "Bacteria" "Bacteroidetes"       "Bacteroidia"                
## [2584,] "Bacteria" "Firmicutes"          "Negativicutes"              
## [2585,] "Bacteria" "Bacteroidetes"       "Bacteroidia"                
## [2586,] "Bacteria" "Patescibacteria"     "Saccharimonadia"            
## [2587,] "Bacteria" "Bacteroidetes"       "Bacteroidia"                
## [2588,] "Bacteria" "Firmicutes"          "Clostridia"                 
## [2589,] "Bacteria" "Firmicutes"          "Clostridia"                 
## [2590,] "Bacteria" "Bacteroidetes"       "Bacteroidia"                
## [2591,] "Bacteria" "Epsilonbacteraeota"  "Campylobacteria"            
## [2592,] "Bacteria" "Firmicutes"          "Negativicutes"              
## [2593,] "Bacteria" "Firmicutes"          "Bacilli"                    
## [2594,] "Bacteria" "Proteobacteria"      "Gammaproteobacteria"        
## [2595,] "Bacteria" "Bacteroidetes"       "Bacteroidia"                
## [2596,] "Bacteria" "Firmicutes"          "Bacilli"                    
## [2597,] "Bacteria" "Bacteroidetes"       "Bacteroidia"                
## [2598,] "Bacteria" "Firmicutes"          NA                           
## [2599,] "Bacteria" "Bacteroidetes"       "Bacteroidia"                
## [2600,] "Bacteria" "Firmicutes"          "Negativicutes"              
## [2601,] "Bacteria" "Firmicutes"          "Negativicutes"              
## [2602,] "Bacteria" "Proteobacteria"      "Gammaproteobacteria"        
## [2603,] "Bacteria" "Bacteroidetes"       "Bacteroidia"                
## [2604,] "Bacteria" "Firmicutes"          "Negativicutes"              
## [2605,] "Bacteria" "Bacteroidetes"       "Bacteroidia"                
## [2606,] "Bacteria" "Proteobacteria"      "Gammaproteobacteria"        
## [2607,] "Bacteria" "Actinobacteria"      "Actinobacteria"             
## [2608,] "Bacteria" NA                    NA                           
## [2609,] "Bacteria" "Firmicutes"          NA                           
## [2610,] "Bacteria" "Actinobacteria"      "Actinobacteria"             
## [2611,] "Bacteria" "Firmicutes"          "Clostridia"                 
## [2612,] "Bacteria" "Bacteroidetes"       "Bacteroidia"                
## [2613,] "Bacteria" "Actinobacteria"      "Actinobacteria"             
## [2614,] "Bacteria" "Bacteroidetes"       "Bacteroidia"                
## [2615,] "Bacteria" "Firmicutes"          NA                           
## [2616,] "Bacteria" "Bacteroidetes"       "Bacteroidia"                
## [2617,] "Bacteria" "Proteobacteria"      "Gammaproteobacteria"        
## [2618,] "Bacteria" "Firmicutes"          "Negativicutes"              
## [2619,] "Bacteria" "Proteobacteria"      "Gammaproteobacteria"        
## [2620,] "Bacteria" "Fusobacteria"        "Fusobacteriia"              
## [2621,] "Bacteria" "Fusobacteria"        "Fusobacteriia"              
## [2622,] "Bacteria" "Bacteroidetes"       "Bacteroidia"                
## [2623,] "Bacteria" "Bacteroidetes"       "Bacteroidia"                
## [2624,] "Bacteria" "Proteobacteria"      "Gammaproteobacteria"        
## [2625,] "Bacteria" "Firmicutes"          "Clostridia"                 
## [2626,] "Bacteria" "Patescibacteria"     "Gracilibacteria"            
## [2627,] "Bacteria" "Firmicutes"          "Negativicutes"              
## [2628,] "Bacteria" "Firmicutes"          "Bacilli"                    
## [2629,] "Bacteria" "Actinobacteria"      "Actinobacteria"             
## [2630,] "Bacteria" "Firmicutes"          "Clostridia"                 
## [2631,] "Bacteria" "Firmicutes"          NA                           
## [2632,] "Bacteria" "Spirochaetes"        "Spirochaetia"               
## [2633,] "Bacteria" "Firmicutes"          "Bacilli"                    
## [2634,] "Bacteria" "Bacteroidetes"       "Bacteroidia"                
## [2635,] "Bacteria" "Epsilonbacteraeota"  "Campylobacteria"            
## [2636,] "Bacteria" "Firmicutes"          "Negativicutes"              
## [2637,] "Bacteria" "Epsilonbacteraeota"  "Campylobacteria"            
## [2638,] "Bacteria" "Epsilonbacteraeota"  "Campylobacteria"            
## [2639,] "Bacteria" "Bacteroidetes"       "Bacteroidia"                
## [2640,] "Bacteria" "Fusobacteria"        "Fusobacteriia"              
## [2641,] "Bacteria" "Epsilonbacteraeota"  "Campylobacteria"            
## [2642,] "Bacteria" "Firmicutes"          "Negativicutes"              
## [2643,] "Bacteria" "Bacteroidetes"       "Bacteroidia"                
## [2644,] "Bacteria" "Firmicutes"          "Bacilli"                    
## [2645,] "Bacteria" "Proteobacteria"      "Gammaproteobacteria"        
## [2646,] "Bacteria" "Firmicutes"          "Clostridia"                 
## [2647,] "Bacteria" "Bacteroidetes"       "Bacteroidia"                
## [2648,] "Bacteria" "Firmicutes"          "Clostridia"                 
## [2649,] "Bacteria" "Bacteroidetes"       "Bacteroidia"                
## [2650,] "Bacteria" "Firmicutes"          "Bacilli"                    
## [2651,] "Bacteria" "Spirochaetes"        "Spirochaetia"               
## [2652,] "Bacteria" "Proteobacteria"      "Gammaproteobacteria"        
## [2653,] "Bacteria" "Fusobacteria"        "Fusobacteriia"              
## [2654,] "Bacteria" "Bacteroidetes"       "Bacteroidia"                
## [2655,] "Bacteria" "Firmicutes"          "Negativicutes"              
## [2656,] "Bacteria" "Bacteroidetes"       "Bacteroidia"                
## [2657,] "Bacteria" "Bacteroidetes"       "Bacteroidia"                
## [2658,] "Bacteria" "Firmicutes"          "Clostridia"                 
## [2659,] "Bacteria" "Bacteroidetes"       "Bacteroidia"                
## [2660,] "Bacteria" "Bacteroidetes"       "Bacteroidia"                
## [2661,] "Bacteria" "Bacteroidetes"       "Bacteroidia"                
## [2662,] "Bacteria" "Firmicutes"          "Negativicutes"              
## [2663,] "Bacteria" "Actinobacteria"      "Actinobacteria"             
## [2664,] "Bacteria" "Actinobacteria"      "Actinobacteria"             
## [2665,] "Bacteria" "Fusobacteria"        "Fusobacteriia"              
## [2666,] "Bacteria" "Firmicutes"          "Negativicutes"              
## [2667,] "Bacteria" NA                    NA                           
## [2668,] "Bacteria" "Firmicutes"          "Negativicutes"              
## [2669,] "Bacteria" "Firmicutes"          "Clostridia"                 
## [2670,] "Bacteria" "Bacteroidetes"       "Bacteroidia"                
## [2671,] "Bacteria" "Firmicutes"          "Negativicutes"              
## [2672,] "Bacteria" "Proteobacteria"      "Gammaproteobacteria"        
## [2673,] "Bacteria" "Bacteroidetes"       "Bacteroidia"                
## [2674,] "Bacteria" "Bacteroidetes"       "Bacteroidia"                
## [2675,] "Bacteria" "Spirochaetes"        "Spirochaetia"               
## [2676,] "Bacteria" "Bacteroidetes"       "Bacteroidia"                
## [2677,] "Bacteria" "Actinobacteria"      "Actinobacteria"             
## [2678,] "Bacteria" "Tenericutes"         "Mollicutes"                 
## [2679,] "Bacteria" "Proteobacteria"      "Gammaproteobacteria"        
## [2680,] "Bacteria" "Bacteroidetes"       "Bacteroidia"                
## [2681,] "Bacteria" "Proteobacteria"      "Gammaproteobacteria"        
## [2682,] "Bacteria" "Bacteroidetes"       "Bacteroidia"                
## [2683,] "Bacteria" "Actinobacteria"      "Actinobacteria"             
## [2684,] "Bacteria" "Bacteroidetes"       "Bacteroidia"                
## [2685,] "Bacteria" "Bacteroidetes"       "Bacteroidia"                
## [2686,] "Bacteria" "Fusobacteria"        "Fusobacteriia"              
## [2687,] "Bacteria" "Actinobacteria"      "Coriobacteriia"             
## [2688,] "Bacteria" "Patescibacteria"     "Saccharimonadia"            
## [2689,] "Bacteria" "Actinobacteria"      "Actinobacteria"             
## [2690,] "Bacteria" NA                    NA                           
## [2691,] "Bacteria" "Firmicutes"          "Clostridia"                 
## [2692,] "Bacteria" "Fusobacteria"        "Fusobacteriia"              
## [2693,] "Bacteria" "Firmicutes"          "Clostridia"                 
## [2694,] "Bacteria" "Bacteroidetes"       "Bacteroidia"                
## [2695,] "Bacteria" "Bacteroidetes"       "Bacteroidia"                
## [2696,] "Bacteria" "Firmicutes"          NA                           
## [2697,] "Bacteria" "Firmicutes"          "Negativicutes"              
## [2698,] "Bacteria" "Firmicutes"          "Negativicutes"              
## [2699,] "Bacteria" "Firmicutes"          "Negativicutes"              
## [2700,] "Bacteria" "Bacteroidetes"       "Bacteroidia"                
## [2701,] "Bacteria" NA                    NA                           
## [2702,] "Bacteria" "Firmicutes"          "Negativicutes"              
## [2703,] "Bacteria" "Bacteroidetes"       "Bacteroidia"                
## [2704,] "Bacteria" "Bacteroidetes"       "Bacteroidia"                
## [2705,] "Bacteria" "Proteobacteria"      "Gammaproteobacteria"        
## [2706,] "Bacteria" "Fusobacteria"        "Fusobacteriia"              
## [2707,] "Bacteria" "Firmicutes"          "Negativicutes"              
## [2708,] "Bacteria" "Fusobacteria"        "Fusobacteriia"              
## [2709,] "Bacteria" "Firmicutes"          "Negativicutes"              
## [2710,] "Bacteria" "Bacteroidetes"       "Bacteroidia"                
## [2711,] "Bacteria" "Bacteroidetes"       "Bacteroidia"                
## [2712,] "Bacteria" "Fusobacteria"        "Fusobacteriia"              
## [2713,] "Bacteria" "Firmicutes"          "Negativicutes"              
## [2714,] "Bacteria" "Fusobacteria"        "Fusobacteriia"              
## [2715,] "Bacteria" "Bacteroidetes"       "Bacteroidia"                
## [2716,] "Bacteria" "Fusobacteria"        "Fusobacteriia"              
## [2717,] "Bacteria" "Fusobacteria"        "Fusobacteriia"              
## [2718,] "Bacteria" "Bacteroidetes"       "Bacteroidia"                
## [2719,] "Bacteria" "Proteobacteria"      "Gammaproteobacteria"        
## [2720,] "Bacteria" "Firmicutes"          "Clostridia"                 
## [2721,] "Bacteria" "Spirochaetes"        "Spirochaetia"               
## [2722,] "Bacteria" "Fusobacteria"        "Fusobacteriia"              
## [2723,] "Bacteria" "Firmicutes"          "Clostridia"                 
## [2724,] "Bacteria" "Bacteroidetes"       "Bacteroidia"                
## [2725,] "Bacteria" "Bacteroidetes"       "Bacteroidia"                
## [2726,] "Bacteria" "Proteobacteria"      "Gammaproteobacteria"        
## [2727,] "Bacteria" "Bacteroidetes"       "Bacteroidia"                
## [2728,] "Bacteria" "Actinobacteria"      "Actinobacteria"             
## [2729,] "Bacteria" "Bacteroidetes"       "Bacteroidia"                
## [2730,] "Bacteria" "Epsilonbacteraeota"  "Campylobacteria"            
## [2731,] "Bacteria" "Firmicutes"          "Clostridia"                 
## [2732,] "Bacteria" "Proteobacteria"      "Gammaproteobacteria"        
## [2733,] "Bacteria" "Bacteroidetes"       "Bacteroidia"                
## [2734,] "Bacteria" "Fusobacteria"        "Fusobacteriia"              
## [2735,] "Bacteria" "Fusobacteria"        "Fusobacteriia"              
## [2736,] "Bacteria" "Fusobacteria"        "Fusobacteriia"              
## [2737,] "Bacteria" "Proteobacteria"      "Gammaproteobacteria"        
## [2738,] "Bacteria" "Bacteroidetes"       "Bacteroidia"                
## [2739,] "Bacteria" "Actinobacteria"      "Actinobacteria"             
## [2740,] "Bacteria" NA                    NA                           
## [2741,] "Bacteria" "Firmicutes"          NA                           
## [2742,] "Bacteria" "Bacteroidetes"       "Bacteroidia"                
## [2743,] "Bacteria" "Firmicutes"          "Erysipelotrichia"           
## [2744,] "Bacteria" "Firmicutes"          "Negativicutes"              
## [2745,] "Bacteria" "Proteobacteria"      "Gammaproteobacteria"        
## [2746,] "Bacteria" "Proteobacteria"      "Gammaproteobacteria"        
## [2747,] "Bacteria" "Firmicutes"          "Negativicutes"              
## [2748,] "Bacteria" "Bacteroidetes"       "Bacteroidia"                
## [2749,] "Bacteria" "Bacteroidetes"       "Bacteroidia"                
## [2750,] "Bacteria" "Bacteroidetes"       "Bacteroidia"                
## [2751,] "Bacteria" "Patescibacteria"     "Saccharimonadia"            
## [2752,] "Bacteria" "Bacteroidetes"       "Bacteroidia"                
## [2753,] "Bacteria" "Bacteroidetes"       "Bacteroidia"                
## [2754,] "Bacteria" "Bacteroidetes"       "Bacteroidia"                
## [2755,] "Bacteria" "Proteobacteria"      "Gammaproteobacteria"        
## [2756,] "Bacteria" NA                    NA                           
## [2757,] "Bacteria" "Bacteroidetes"       "Bacteroidia"                
## [2758,] "Bacteria" "Fusobacteria"        "Fusobacteriia"              
## [2759,] "Bacteria" "Bacteroidetes"       "Bacteroidia"                
## [2760,] "Bacteria" "Actinobacteria"      "Coriobacteriia"             
## [2761,] "Bacteria" "Bacteroidetes"       "Bacteroidia"                
## [2762,] "Bacteria" "Bacteroidetes"       "Bacteroidia"                
## [2763,] "Bacteria" "Bacteroidetes"       "Bacteroidia"                
## [2764,] "Bacteria" "Actinobacteria"      "Actinobacteria"             
## [2765,] "Bacteria" "Actinobacteria"      "Actinobacteria"             
## [2766,] "Bacteria" "Bacteroidetes"       "Bacteroidia"                
## [2767,] "Bacteria" "Firmicutes"          "Bacilli"                    
## [2768,] "Bacteria" "Firmicutes"          "Bacilli"                    
## [2769,] "Bacteria" "Firmicutes"          "Clostridia"                 
## [2770,] "Bacteria" "Firmicutes"          "Negativicutes"              
## [2771,] "Bacteria" "Patescibacteria"     "Saccharimonadia"            
## [2772,] "Bacteria" "Firmicutes"          "Bacilli"                    
## [2773,] "Bacteria" "Bacteroidetes"       "Bacteroidia"                
## [2774,] "Bacteria" "Firmicutes"          "Negativicutes"              
## [2775,] "Bacteria" "Epsilonbacteraeota"  "Campylobacteria"            
## [2776,] "Bacteria" "Actinobacteria"      "Actinobacteria"             
## [2777,] "Bacteria" "Proteobacteria"      "Gammaproteobacteria"        
## [2778,] "Bacteria" "Actinobacteria"      "Actinobacteria"             
## [2779,] "Bacteria" "Bacteroidetes"       "Bacteroidia"                
## [2780,] "Bacteria" "Bacteroidetes"       "Bacteroidia"                
## [2781,] "Bacteria" "Firmicutes"          NA                           
## [2782,] "Bacteria" "Bacteroidetes"       "Bacteroidia"                
## [2783,] "Bacteria" "Firmicutes"          "Clostridia"                 
## [2784,] "Bacteria" "Firmicutes"          "Clostridia"                 
## [2785,] "Bacteria" "Actinobacteria"      "Actinobacteria"             
## [2786,] "Bacteria" "Actinobacteria"      "Actinobacteria"             
## [2787,] "Bacteria" "Bacteroidetes"       "Bacteroidia"                
## [2788,] "Bacteria" "Firmicutes"          "Negativicutes"              
## [2789,] "Bacteria" NA                    NA                           
## [2790,] "Bacteria" "Bacteroidetes"       "Bacteroidia"                
## [2791,] "Bacteria" "Bacteroidetes"       "Bacteroidia"                
## [2792,] "Bacteria" "Bacteroidetes"       "Bacteroidia"                
## [2793,] "Bacteria" "Firmicutes"          "Negativicutes"              
## [2794,] "Bacteria" "Firmicutes"          "Clostridia"                 
## [2795,] "Bacteria" "Proteobacteria"      "Gammaproteobacteria"        
## [2796,] "Bacteria" "Bacteroidetes"       "Bacteroidia"                
## [2797,] "Bacteria" "Bacteroidetes"       "Bacteroidia"                
## [2798,] "Bacteria" "Bacteroidetes"       "Bacteroidia"                
## [2799,] "Bacteria" "Bacteroidetes"       "Bacteroidia"                
## [2800,] "Bacteria" "Proteobacteria"      "Gammaproteobacteria"        
## [2801,] "Bacteria" "Proteobacteria"      "Gammaproteobacteria"        
## [2802,] "Bacteria" "Bacteroidetes"       "Bacteroidia"                
## [2803,] "Bacteria" "Bacteroidetes"       "Bacteroidia"                
## [2804,] "Bacteria" "Firmicutes"          "Negativicutes"              
## [2805,] "Bacteria" "Firmicutes"          "Clostridia"                 
## [2806,] "Bacteria" "Bacteroidetes"       "Bacteroidia"                
## [2807,] "Bacteria" "Bacteroidetes"       "Bacteroidia"                
## [2808,] "Bacteria" "Firmicutes"          "Negativicutes"              
## [2809,] "Bacteria" "Fusobacteria"        "Fusobacteriia"              
## [2810,] "Bacteria" "Bacteroidetes"       "Bacteroidia"                
## [2811,] "Bacteria" "Bacteroidetes"       "Bacteroidia"                
## [2812,] "Bacteria" "Fusobacteria"        "Fusobacteriia"              
## [2813,] "Bacteria" "Proteobacteria"      "Gammaproteobacteria"        
## [2814,] "Bacteria" "Fusobacteria"        "Fusobacteriia"              
## [2815,] "Bacteria" "Bacteroidetes"       "Bacteroidia"                
## [2816,] "Bacteria" "Bacteroidetes"       "Bacteroidia"                
## [2817,] "Bacteria" "Bacteroidetes"       "Bacteroidia"                
## [2818,] "Bacteria" "Firmicutes"          "Negativicutes"              
## [2819,] "Bacteria" "Firmicutes"          "Clostridia"                 
## [2820,] "Bacteria" "Bacteroidetes"       "Bacteroidia"                
## [2821,] "Bacteria" "Firmicutes"          "Negativicutes"              
## [2822,] "Bacteria" "Bacteroidetes"       "Bacteroidia"                
## [2823,] "Bacteria" "Bacteroidetes"       "Bacteroidia"                
## [2824,] "Bacteria" "Bacteroidetes"       "Bacteroidia"                
## [2825,] "Bacteria" "Bacteroidetes"       "Bacteroidia"                
## [2826,] "Bacteria" "Bacteroidetes"       "Bacteroidia"                
## [2827,] "Bacteria" "Firmicutes"          "Negativicutes"              
## [2828,] "Bacteria" "Firmicutes"          "Negativicutes"              
## [2829,] "Bacteria" "Patescibacteria"     "Saccharimonadia"            
## [2830,] "Bacteria" "Firmicutes"          "Clostridia"                 
## [2831,] "Bacteria" "Patescibacteria"     "Saccharimonadia"            
## [2832,] "Bacteria" "Proteobacteria"      "Gammaproteobacteria"        
## [2833,] "Bacteria" "Proteobacteria"      "Gammaproteobacteria"        
## [2834,] "Bacteria" "Spirochaetes"        "Spirochaetia"               
## [2835,] "Bacteria" "Firmicutes"          "Negativicutes"              
## [2836,] "Bacteria" "Bacteroidetes"       "Bacteroidia"                
## [2837,] "Bacteria" "Firmicutes"          "Negativicutes"              
## [2838,] "Bacteria" "Fusobacteria"        "Fusobacteriia"              
## [2839,] "Bacteria" "Bacteroidetes"       "Bacteroidia"                
## [2840,] "Bacteria" "Firmicutes"          "Clostridia"                 
## [2841,] "Bacteria" "Firmicutes"          "Clostridia"                 
## [2842,] "Bacteria" "Firmicutes"          "Negativicutes"              
## [2843,] "Bacteria" "Proteobacteria"      "Gammaproteobacteria"        
## [2844,] "Bacteria" "Firmicutes"          "Negativicutes"              
## [2845,] "Bacteria" "Bacteroidetes"       "Bacteroidia"                
## [2846,] "Bacteria" "Firmicutes"          "Negativicutes"              
## [2847,] "Bacteria" "Proteobacteria"      "Gammaproteobacteria"        
## [2848,] "Bacteria" "Firmicutes"          "Bacilli"                    
## [2849,] "Bacteria" "Firmicutes"          "Negativicutes"              
## [2850,] "Bacteria" "Bacteroidetes"       "Bacteroidia"                
## [2851,] "Bacteria" "Proteobacteria"      "Gammaproteobacteria"        
## [2852,] "Bacteria" "Fusobacteria"        "Fusobacteriia"              
## [2853,] "Bacteria" "Firmicutes"          "Negativicutes"              
## [2854,] "Bacteria" "Bacteroidetes"       "Bacteroidia"                
## [2855,] "Bacteria" "Proteobacteria"      "Gammaproteobacteria"        
## [2856,] "Bacteria" "Actinobacteria"      "Actinobacteria"             
## [2857,] "Bacteria" "Proteobacteria"      "Gammaproteobacteria"        
## [2858,] "Bacteria" "Bacteroidetes"       "Bacteroidia"                
## [2859,] "Bacteria" "Bacteroidetes"       "Bacteroidia"                
## [2860,] "Bacteria" "Proteobacteria"      "Gammaproteobacteria"        
## [2861,] "Bacteria" "Firmicutes"          "Negativicutes"              
## [2862,] "Bacteria" "Firmicutes"          "Negativicutes"              
## [2863,] "Bacteria" "Bacteroidetes"       "Bacteroidia"                
## [2864,] "Bacteria" "Actinobacteria"      "Actinobacteria"             
## [2865,] "Bacteria" "Bacteroidetes"       "Bacteroidia"                
## [2866,] "Bacteria" "Tenericutes"         "Mollicutes"                 
## [2867,] "Bacteria" "Bacteroidetes"       "Bacteroidia"                
## [2868,] "Bacteria" "Firmicutes"          "Clostridia"                 
## [2869,] "Bacteria" "Actinobacteria"      "Actinobacteria"             
## [2870,] "Bacteria" "Firmicutes"          "Clostridia"                 
## [2871,] "Bacteria" "Bacteroidetes"       "Bacteroidia"                
## [2872,] "Bacteria" "Firmicutes"          NA                           
## [2873,] "Bacteria" "Proteobacteria"      "Gammaproteobacteria"        
## [2874,] "Bacteria" "Bacteroidetes"       "Bacteroidia"                
## [2875,] "Bacteria" "Bacteroidetes"       "Bacteroidia"                
## [2876,] "Bacteria" "Bacteroidetes"       "Bacteroidia"                
## [2877,] "Bacteria" "Proteobacteria"      "Gammaproteobacteria"        
## [2878,] "Bacteria" "Bacteroidetes"       "Bacteroidia"                
## [2879,] "Bacteria" "Firmicutes"          "Bacilli"                    
## [2880,] "Bacteria" "Firmicutes"          NA                           
## [2881,] "Bacteria" "Bacteroidetes"       "Bacteroidia"                
## [2882,] "Bacteria" "Proteobacteria"      "Gammaproteobacteria"        
## [2883,] "Bacteria" "Firmicutes"          "Negativicutes"              
## [2884,] "Bacteria" "Firmicutes"          "Clostridia"                 
## [2885,] "Bacteria" "Bacteroidetes"       "Bacteroidia"                
## [2886,] "Bacteria" "Firmicutes"          "Clostridia"                 
## [2887,] "Bacteria" "Firmicutes"          "Clostridia"                 
## [2888,] "Bacteria" "Fusobacteria"        "Fusobacteriia"              
## [2889,] "Bacteria" "Bacteroidetes"       "Bacteroidia"                
## [2890,] "Bacteria" "Bacteroidetes"       "Bacteroidia"                
## [2891,] "Bacteria" "Proteobacteria"      "Gammaproteobacteria"        
## [2892,] "Bacteria" "Bacteroidetes"       "Bacteroidia"                
## [2893,] "Bacteria" "Bacteroidetes"       "Bacteroidia"                
## [2894,] "Bacteria" "Proteobacteria"      "Gammaproteobacteria"        
## [2895,] "Bacteria" "Firmicutes"          NA                           
## [2896,] "Bacteria" "Fusobacteria"        "Fusobacteriia"              
## [2897,] "Bacteria" "Bacteroidetes"       "Bacteroidia"                
## [2898,] "Bacteria" "Bacteroidetes"       "Bacteroidia"                
## [2899,] "Bacteria" "Patescibacteria"     "Saccharimonadia"            
## [2900,] "Bacteria" "Fusobacteria"        "Fusobacteriia"              
## [2901,] "Bacteria" "Bacteroidetes"       "Bacteroidia"                
## [2902,] "Bacteria" "Firmicutes"          "Negativicutes"              
## [2903,] "Bacteria" "Actinobacteria"      "Actinobacteria"             
## [2904,] "Bacteria" "Bacteroidetes"       "Bacteroidia"                
## [2905,] "Bacteria" "Spirochaetes"        "Spirochaetia"               
## [2906,] "Bacteria" "Proteobacteria"      "Gammaproteobacteria"        
## [2907,] "Bacteria" "Spirochaetes"        "Spirochaetia"               
## [2908,] "Bacteria" "Epsilonbacteraeota"  "Campylobacteria"            
## [2909,] "Bacteria" "Firmicutes"          "Clostridia"                 
## [2910,] "Bacteria" "Proteobacteria"      "Gammaproteobacteria"        
## [2911,] "Bacteria" "Firmicutes"          "Negativicutes"              
## [2912,] "Bacteria" "Tenericutes"         "Mollicutes"                 
## [2913,] "Bacteria" "Bacteroidetes"       "Bacteroidia"                
## [2914,] "Bacteria" NA                    NA                           
## [2915,] "Bacteria" "Patescibacteria"     "Saccharimonadia"            
## [2916,] "Bacteria" "Bacteroidetes"       "Bacteroidia"                
## [2917,] "Bacteria" "Patescibacteria"     "Saccharimonadia"            
## [2918,] "Bacteria" "Firmicutes"          "Negativicutes"              
## [2919,] "Bacteria" "Fusobacteria"        "Fusobacteriia"              
## [2920,] "Bacteria" "Firmicutes"          "Negativicutes"              
## [2921,] "Bacteria" "Bacteroidetes"       "Bacteroidia"                
## [2922,] "Bacteria" "Actinobacteria"      "Actinobacteria"             
## [2923,] "Bacteria" "Proteobacteria"      "Gammaproteobacteria"        
## [2924,] "Bacteria" "Firmicutes"          "Bacilli"                    
## [2925,] "Bacteria" "Firmicutes"          "Clostridia"                 
## [2926,] "Bacteria" "Fusobacteria"        "Fusobacteriia"              
## [2927,] "Bacteria" "Proteobacteria"      "Gammaproteobacteria"        
## [2928,] "Bacteria" "Firmicutes"          "Bacilli"                    
## [2929,] "Bacteria" "Actinobacteria"      "Actinobacteria"             
## [2930,] "Bacteria" "Actinobacteria"      "Actinobacteria"             
## [2931,] "Bacteria" "Firmicutes"          "Clostridia"                 
## [2932,] "Bacteria" "Proteobacteria"      "Gammaproteobacteria"        
## [2933,] "Bacteria" "Firmicutes"          "Bacilli"                    
## [2934,] "Bacteria" "Actinobacteria"      "Coriobacteriia"             
## [2935,] "Bacteria" "Actinobacteria"      "Actinobacteria"             
## [2936,] "Bacteria" "Firmicutes"          "Negativicutes"              
## [2937,] "Bacteria" "Firmicutes"          NA                           
## [2938,] "Bacteria" "Fusobacteria"        "Fusobacteriia"              
## [2939,] "Bacteria" "Bacteroidetes"       "Bacteroidia"                
## [2940,] "Bacteria" "Bacteroidetes"       "Bacteroidia"                
## [2941,] "Bacteria" "Actinobacteria"      "Actinobacteria"             
## [2942,] "Bacteria" "Patescibacteria"     "Saccharimonadia"            
## [2943,] "Bacteria" "Bacteroidetes"       "Bacteroidia"                
## [2944,] "Bacteria" "Firmicutes"          "Bacilli"                    
## [2945,] "Bacteria" "Bacteroidetes"       "Bacteroidia"                
## [2946,] "Bacteria" "Bacteroidetes"       "Bacteroidia"                
## [2947,] "Bacteria" "Bacteroidetes"       "Bacteroidia"                
## [2948,] "Bacteria" "Firmicutes"          NA                           
## [2949,] "Bacteria" "Firmicutes"          "Clostridia"                 
## [2950,] "Bacteria" "Firmicutes"          "Negativicutes"              
## [2951,] "Bacteria" "Firmicutes"          "Negativicutes"              
## [2952,] "Bacteria" "Bacteroidetes"       "Bacteroidia"                
## [2953,] "Bacteria" "Bacteroidetes"       "Bacteroidia"                
## [2954,] "Bacteria" "Bacteroidetes"       "Bacteroidia"                
## [2955,] "Bacteria" "Firmicutes"          "Clostridia"                 
## [2956,] "Bacteria" "Cyanobacteria"       "Oxyphotobacteria"           
## [2957,] "Bacteria" "Proteobacteria"      "Gammaproteobacteria"        
## [2958,] "Bacteria" "Fusobacteria"        "Fusobacteriia"              
## [2959,] "Bacteria" "Firmicutes"          "Negativicutes"              
## [2960,] "Bacteria" "Firmicutes"          "Negativicutes"              
## [2961,] "Bacteria" "Bacteroidetes"       "Bacteroidia"                
## [2962,] "Bacteria" "Proteobacteria"      "Gammaproteobacteria"        
## [2963,] "Bacteria" "Firmicutes"          "Negativicutes"              
## [2964,] "Bacteria" "Firmicutes"          "Negativicutes"              
## [2965,] "Bacteria" "Proteobacteria"      "Gammaproteobacteria"        
## [2966,] "Bacteria" "Firmicutes"          "Negativicutes"              
## [2967,] "Bacteria" "Firmicutes"          "Negativicutes"              
## [2968,] "Bacteria" "Firmicutes"          NA                           
## [2969,] "Bacteria" "Bacteroidetes"       "Bacteroidia"                
## [2970,] "Bacteria" "Fusobacteria"        "Fusobacteriia"              
## [2971,] "Bacteria" "Proteobacteria"      "Gammaproteobacteria"        
## [2972,] "Bacteria" "Patescibacteria"     "Gracilibacteria"            
## [2973,] "Bacteria" "Bacteroidetes"       "Bacteroidia"                
## [2974,] "Bacteria" "Bacteroidetes"       "Bacteroidia"                
## [2975,] "Bacteria" "Firmicutes"          "Bacilli"                    
## [2976,] "Bacteria" "Spirochaetes"        "Spirochaetia"               
## [2977,] "Bacteria" "Firmicutes"          "Clostridia"                 
## [2978,] "Bacteria" "Tenericutes"         "Mollicutes"                 
## [2979,] "Bacteria" "Firmicutes"          "Negativicutes"              
## [2980,] "Bacteria" "Bacteroidetes"       "Bacteroidia"                
## [2981,] "Bacteria" "Proteobacteria"      "Gammaproteobacteria"        
## [2982,] "Bacteria" "Spirochaetes"        "Spirochaetia"               
## [2983,] "Bacteria" "Epsilonbacteraeota"  "Campylobacteria"            
## [2984,] "Bacteria" "Bacteroidetes"       "Bacteroidia"                
## [2985,] "Bacteria" "Firmicutes"          "Clostridia"                 
## [2986,] "Bacteria" "Proteobacteria"      "Gammaproteobacteria"        
## [2987,] "Bacteria" "Actinobacteria"      "Actinobacteria"             
## [2988,] "Bacteria" "Bacteroidetes"       "Bacteroidia"                
## [2989,] "Bacteria" "Bacteroidetes"       "Bacteroidia"                
## [2990,] "Bacteria" "Proteobacteria"      "Gammaproteobacteria"        
## [2991,] "Bacteria" "Firmicutes"          "Negativicutes"              
## [2992,] "Bacteria" "Synergistetes"       "Synergistia"                
## [2993,] "Bacteria" "Firmicutes"          "Bacilli"                    
## [2994,] "Bacteria" "Actinobacteria"      "Actinobacteria"             
## [2995,] "Bacteria" "Bacteroidetes"       "Bacteroidia"                
## [2996,] "Bacteria" "Firmicutes"          "Clostridia"                 
## [2997,] "Bacteria" "Firmicutes"          "Negativicutes"              
## [2998,] "Bacteria" "Patescibacteria"     "Saccharimonadia"            
## [2999,] "Bacteria" "Proteobacteria"      "Gammaproteobacteria"        
## [3000,] "Bacteria" "Firmicutes"          "Negativicutes"              
## [3001,] "Bacteria" "Bacteroidetes"       "Bacteroidia"                
## [3002,] "Bacteria" "Fusobacteria"        "Fusobacteriia"              
## [3003,] "Bacteria" "Firmicutes"          NA                           
## [3004,] "Bacteria" "Bacteroidetes"       "Bacteroidia"                
## [3005,] "Bacteria" "Proteobacteria"      "Gammaproteobacteria"        
## [3006,] "Bacteria" "Proteobacteria"      "Gammaproteobacteria"        
## [3007,] "Bacteria" "Firmicutes"          "Negativicutes"              
## [3008,] "Bacteria" "Actinobacteria"      "Actinobacteria"             
## [3009,] "Bacteria" "Firmicutes"          "Clostridia"                 
## [3010,] "Bacteria" "Firmicutes"          "Negativicutes"              
## [3011,] "Bacteria" "Fusobacteria"        "Fusobacteriia"              
## [3012,] "Bacteria" "Firmicutes"          "Negativicutes"              
## [3013,] "Bacteria" "Firmicutes"          NA                           
## [3014,] "Bacteria" "Bacteroidetes"       "Bacteroidia"                
## [3015,] "Bacteria" "Spirochaetes"        "Spirochaetia"               
## [3016,] "Bacteria" NA                    NA                           
## [3017,] "Bacteria" "Proteobacteria"      "Gammaproteobacteria"        
## [3018,] "Bacteria" "Firmicutes"          "Negativicutes"              
## [3019,] "Bacteria" "Bacteroidetes"       "Bacteroidia"                
## [3020,] "Bacteria" NA                    NA                           
## [3021,] "Bacteria" "Bacteroidetes"       "Bacteroidia"                
## [3022,] "Bacteria" "Firmicutes"          "Clostridia"                 
## [3023,] "Bacteria" "Bacteroidetes"       "Bacteroidia"                
## [3024,] "Bacteria" "Bacteroidetes"       "Bacteroidia"                
## [3025,] "Bacteria" "Bacteroidetes"       "Bacteroidia"                
## [3026,] "Bacteria" "Patescibacteria"     "Gracilibacteria"            
## [3027,] "Bacteria" "Bacteroidetes"       "Bacteroidia"                
## [3028,] "Bacteria" "Patescibacteria"     "Gracilibacteria"            
## [3029,] "Bacteria" "Fusobacteria"        "Fusobacteriia"              
## [3030,] "Bacteria" "Bacteroidetes"       "Bacteroidia"                
## [3031,] "Bacteria" "Bacteroidetes"       "Bacteroidia"                
## [3032,] "Bacteria" "Proteobacteria"      "Gammaproteobacteria"        
## [3033,] "Bacteria" "Bacteroidetes"       "Bacteroidia"                
## [3034,] "Bacteria" NA                    NA                           
## [3035,] "Bacteria" "Firmicutes"          "Bacilli"                    
## [3036,] "Bacteria" "Firmicutes"          "Clostridia"                 
## [3037,] "Bacteria" "Firmicutes"          "Clostridia"                 
## [3038,] "Bacteria" "Bacteroidetes"       "Bacteroidia"                
## [3039,] "Bacteria" "Actinobacteria"      "Actinobacteria"             
## [3040,] "Bacteria" "Firmicutes"          "Bacilli"                    
## [3041,] "Bacteria" "Epsilonbacteraeota"  "Campylobacteria"            
## [3042,] "Bacteria" "Bacteroidetes"       "Bacteroidia"                
## [3043,] "Bacteria" "Firmicutes"          "Negativicutes"              
## [3044,] "Bacteria" "Firmicutes"          "Negativicutes"              
## [3045,] "Bacteria" "Bacteroidetes"       "Bacteroidia"                
## [3046,] "Bacteria" "Proteobacteria"      "Gammaproteobacteria"        
## [3047,] "Bacteria" "Actinobacteria"      "Actinobacteria"             
## [3048,] "Bacteria" "Proteobacteria"      "Gammaproteobacteria"        
## [3049,] "Bacteria" "Patescibacteria"     "Gracilibacteria"            
## [3050,] "Bacteria" "Firmicutes"          "Clostridia"                 
## [3051,] "Bacteria" "Bacteroidetes"       "Bacteroidia"                
## [3052,] "Bacteria" "Proteobacteria"      "Gammaproteobacteria"        
## [3053,] "Bacteria" "Fusobacteria"        "Fusobacteriia"              
## [3054,] "Bacteria" "Fusobacteria"        "Fusobacteriia"              
## [3055,] "Bacteria" "Firmicutes"          "Clostridia"                 
## [3056,] "Bacteria" "Bacteroidetes"       "Bacteroidia"                
## [3057,] "Bacteria" "Bacteroidetes"       "Bacteroidia"                
## [3058,] "Bacteria" "Fusobacteria"        "Fusobacteriia"              
## [3059,] "Bacteria" "Bacteroidetes"       "Bacteroidia"                
## [3060,] "Bacteria" "Bacteroidetes"       "Bacteroidia"                
## [3061,] "Bacteria" "Bacteroidetes"       "Bacteroidia"                
## [3062,] "Bacteria" "Fusobacteria"        "Fusobacteriia"              
## [3063,] "Bacteria" "Firmicutes"          "Negativicutes"              
## [3064,] "Bacteria" "Firmicutes"          "Clostridia"                 
## [3065,] "Bacteria" "Actinobacteria"      "Actinobacteria"             
## [3066,] "Bacteria" "Firmicutes"          "Clostridia"                 
## [3067,] "Bacteria" "Firmicutes"          "Negativicutes"              
## [3068,] "Bacteria" "Firmicutes"          "Negativicutes"              
## [3069,] "Bacteria" "Patescibacteria"     "Saccharimonadia"            
## [3070,] "Bacteria" "Actinobacteria"      "Actinobacteria"             
## [3071,] "Bacteria" "Firmicutes"          "Clostridia"                 
## [3072,] "Bacteria" "Firmicutes"          "Negativicutes"              
## [3073,] "Bacteria" "Bacteroidetes"       "Bacteroidia"                
## [3074,] "Bacteria" "Fusobacteria"        "Fusobacteriia"              
## [3075,] "Bacteria" "Bacteroidetes"       "Bacteroidia"                
## [3076,] "Bacteria" "Proteobacteria"      "Gammaproteobacteria"        
## [3077,] "Bacteria" "Firmicutes"          NA                           
## [3078,] "Bacteria" "Fusobacteria"        "Fusobacteriia"              
## [3079,] "Bacteria" "Proteobacteria"      "Gammaproteobacteria"        
## [3080,] "Bacteria" "Bacteroidetes"       "Bacteroidia"                
## [3081,] "Bacteria" "Spirochaetes"        "Spirochaetia"               
## [3082,] "Bacteria" "Actinobacteria"      "Actinobacteria"             
## [3083,] "Bacteria" "Actinobacteria"      "Actinobacteria"             
## [3084,] "Bacteria" "Bacteroidetes"       "Bacteroidia"                
## [3085,] "Bacteria" "Fusobacteria"        "Fusobacteriia"              
## [3086,] "Bacteria" "Firmicutes"          NA                           
## [3087,] "Bacteria" "Bacteroidetes"       "Bacteroidia"                
## [3088,] "Bacteria" "Proteobacteria"      "Gammaproteobacteria"        
## [3089,] "Bacteria" "Patescibacteria"     "Saccharimonadia"            
## [3090,] "Bacteria" "Firmicutes"          "Negativicutes"              
## [3091,] "Bacteria" "Firmicutes"          "Negativicutes"              
## [3092,] "Bacteria" "Actinobacteria"      "Actinobacteria"             
## [3093,] "Bacteria" "Fusobacteria"        "Fusobacteriia"              
## [3094,] "Bacteria" "Actinobacteria"      "Actinobacteria"             
## [3095,] "Bacteria" "Proteobacteria"      "Gammaproteobacteria"        
## [3096,] "Bacteria" "Bacteroidetes"       "Bacteroidia"                
## [3097,] "Bacteria" "Firmicutes"          "Negativicutes"              
## [3098,] "Bacteria" "Firmicutes"          "Negativicutes"              
## [3099,] "Bacteria" "Firmicutes"          NA                           
## [3100,] "Bacteria" "Bacteroidetes"       "Bacteroidia"                
## [3101,] "Bacteria" "Bacteroidetes"       "Bacteroidia"                
## [3102,] "Bacteria" "Spirochaetes"        "Spirochaetia"               
## [3103,] "Bacteria" NA                    NA                           
## [3104,] "Bacteria" "Bacteroidetes"       "Bacteroidia"                
## [3105,] "Bacteria" "Firmicutes"          NA                           
## [3106,] "Bacteria" "Actinobacteria"      "Actinobacteria"             
## [3107,] "Bacteria" "Bacteroidetes"       "Bacteroidia"                
## [3108,] "Bacteria" "Bacteroidetes"       "Bacteroidia"                
## [3109,] "Bacteria" "Proteobacteria"      "Gammaproteobacteria"        
## [3110,] "Bacteria" "Actinobacteria"      "Actinobacteria"             
## [3111,] "Bacteria" "Firmicutes"          NA                           
## [3112,] "Bacteria" "Fusobacteria"        "Fusobacteriia"              
## [3113,] "Bacteria" "Firmicutes"          "Negativicutes"              
## [3114,] "Bacteria" "Firmicutes"          "Clostridia"                 
## [3115,] "Bacteria" "Proteobacteria"      "Gammaproteobacteria"        
## [3116,] "Bacteria" "Firmicutes"          "Clostridia"                 
## [3117,] "Bacteria" "Firmicutes"          "Bacilli"                    
## [3118,] "Bacteria" "Bacteroidetes"       "Bacteroidia"                
## [3119,] "Bacteria" "Fusobacteria"        "Fusobacteriia"              
## [3120,] "Bacteria" "Bacteroidetes"       "Bacteroidia"                
## [3121,] "Bacteria" "Bacteroidetes"       "Bacteroidia"                
## [3122,] "Bacteria" "Actinobacteria"      "Actinobacteria"             
## [3123,] "Bacteria" "Actinobacteria"      "Actinobacteria"             
## [3124,] "Bacteria" "Bacteroidetes"       "Bacteroidia"                
## [3125,] "Bacteria" "Bacteroidetes"       "Bacteroidia"                
## [3126,] "Bacteria" "Firmicutes"          "Clostridia"                 
## [3127,] "Bacteria" "Firmicutes"          "Bacilli"                    
## [3128,] "Bacteria" "Bacteroidetes"       "Bacteroidia"                
## [3129,] "Bacteria" "Actinobacteria"      "Actinobacteria"             
## [3130,] "Bacteria" "Actinobacteria"      "Actinobacteria"             
## [3131,] "Bacteria" "Firmicutes"          "Negativicutes"              
## [3132,] "Bacteria" "Bacteroidetes"       "Bacteroidia"                
## [3133,] "Bacteria" "Bacteroidetes"       "Bacteroidia"                
## [3134,] "Bacteria" "Proteobacteria"      "Gammaproteobacteria"        
## [3135,] "Bacteria" "Bacteroidetes"       "Bacteroidia"                
## [3136,] "Bacteria" "Actinobacteria"      "Actinobacteria"             
## [3137,] "Bacteria" "Epsilonbacteraeota"  "Campylobacteria"            
## [3138,] "Bacteria" "Bacteroidetes"       "Bacteroidia"                
## [3139,] "Bacteria" "Bacteroidetes"       "Bacteroidia"                
## [3140,] "Bacteria" "Bacteroidetes"       "Bacteroidia"                
## [3141,] "Bacteria" "Bacteroidetes"       "Bacteroidia"                
## [3142,] "Bacteria" "Bacteroidetes"       "Bacteroidia"                
## [3143,] "Bacteria" "Bacteroidetes"       "Bacteroidia"                
## [3144,] "Bacteria" "Proteobacteria"      "Gammaproteobacteria"        
## [3145,] "Bacteria" "Patescibacteria"     "Saccharimonadia"            
## [3146,] "Bacteria" "Fusobacteria"        "Fusobacteriia"              
## [3147,] "Bacteria" "Fusobacteria"        "Fusobacteriia"              
## [3148,] "Bacteria" "Actinobacteria"      "Actinobacteria"             
## [3149,] "Bacteria" "Firmicutes"          NA                           
## [3150,] "Bacteria" "Spirochaetes"        "Spirochaetia"               
## [3151,] "Bacteria" "Firmicutes"          "Clostridia"                 
## [3152,] "Bacteria" "Proteobacteria"      "Alphaproteobacteria"        
## [3153,] "Bacteria" "Spirochaetes"        "Spirochaetia"               
## [3154,] "Bacteria" "Bacteroidetes"       "Bacteroidia"                
## [3155,] "Bacteria" "Bacteroidetes"       "Bacteroidia"                
## [3156,] "Bacteria" "Bacteroidetes"       "Bacteroidia"                
## [3157,] "Bacteria" "Firmicutes"          "Negativicutes"              
## [3158,] "Bacteria" "Firmicutes"          "Clostridia"                 
## [3159,] "Bacteria" "Proteobacteria"      "Gammaproteobacteria"        
## [3160,] "Bacteria" "Bacteroidetes"       "Bacteroidia"                
## [3161,] "Bacteria" "Epsilonbacteraeota"  "Campylobacteria"            
## [3162,] "Bacteria" "Proteobacteria"      "Gammaproteobacteria"        
## [3163,] "Bacteria" "Bacteroidetes"       "Bacteroidia"                
## [3164,] "Bacteria" "Bacteroidetes"       "Bacteroidia"                
## [3165,] "Bacteria" "Firmicutes"          "Clostridia"                 
## [3166,] "Bacteria" "Proteobacteria"      "Gammaproteobacteria"        
## [3167,] "Bacteria" "Bacteroidetes"       "Bacteroidia"                
## [3168,] "Bacteria" "Spirochaetes"        "Spirochaetia"               
## [3169,] "Bacteria" "Spirochaetes"        "Spirochaetia"               
## [3170,] "Bacteria" "Fusobacteria"        "Fusobacteriia"              
## [3171,] "Bacteria" "Epsilonbacteraeota"  "Campylobacteria"            
## [3172,] "Bacteria" "Tenericutes"         "Mollicutes"                 
## [3173,] "Bacteria" "Actinobacteria"      "Actinobacteria"             
## [3174,] "Bacteria" "Fusobacteria"        "Fusobacteriia"              
## [3175,] "Bacteria" "Cyanobacteria"       "Oxyphotobacteria"           
## [3176,] "Bacteria" "Epsilonbacteraeota"  "Campylobacteria"            
## [3177,] "Bacteria" "Bacteroidetes"       "Bacteroidia"                
## [3178,] "Bacteria" "Actinobacteria"      "Actinobacteria"             
## [3179,] "Bacteria" "Fusobacteria"        "Fusobacteriia"              
## [3180,] "Bacteria" "Proteobacteria"      "Gammaproteobacteria"        
## [3181,] "Bacteria" "Bacteroidetes"       "Bacteroidia"                
## [3182,] "Bacteria" "Spirochaetes"        "Spirochaetia"               
## [3183,] "Bacteria" "Fusobacteria"        "Fusobacteriia"              
## [3184,] "Bacteria" "Bacteroidetes"       "Bacteroidia"                
## [3185,] "Bacteria" "Fusobacteria"        "Fusobacteriia"              
## [3186,] "Bacteria" "Bacteroidetes"       "Bacteroidia"                
## [3187,] "Bacteria" "Proteobacteria"      "Gammaproteobacteria"        
## [3188,] "Bacteria" "Patescibacteria"     "Saccharimonadia"            
## [3189,] "Bacteria" "Bacteroidetes"       "Bacteroidia"                
## [3190,] "Bacteria" "Bacteroidetes"       "Bacteroidia"                
## [3191,] "Bacteria" "Spirochaetes"        "Spirochaetia"               
## [3192,] "Bacteria" "Proteobacteria"      "Gammaproteobacteria"        
## [3193,] "Bacteria" "Bacteroidetes"       "Bacteroidia"                
## [3194,] "Bacteria" "Fusobacteria"        "Fusobacteriia"              
## [3195,] "Bacteria" "Proteobacteria"      "Gammaproteobacteria"        
## [3196,] "Bacteria" "Fusobacteria"        "Fusobacteriia"              
## [3197,] "Bacteria" "Bacteroidetes"       "Bacteroidia"                
## [3198,] "Bacteria" "Bacteroidetes"       "Bacteroidia"                
## [3199,] "Bacteria" "Bacteroidetes"       "Bacteroidia"                
## [3200,] "Bacteria" "Epsilonbacteraeota"  "Campylobacteria"            
## [3201,] "Bacteria" "Proteobacteria"      "Gammaproteobacteria"        
## [3202,] "Bacteria" "Firmicutes"          "Clostridia"                 
## [3203,] "Bacteria" "Epsilonbacteraeota"  "Campylobacteria"            
## [3204,] "Bacteria" "Patescibacteria"     "Saccharimonadia"            
## [3205,] "Bacteria" "Actinobacteria"      "Actinobacteria"             
## [3206,] "Bacteria" "Bacteroidetes"       "Bacteroidia"                
## [3207,] "Bacteria" "Bacteroidetes"       "Bacteroidia"                
## [3208,] "Bacteria" "Bacteroidetes"       "Bacteroidia"                
## [3209,] "Bacteria" NA                    NA                           
## [3210,] "Bacteria" "Bacteroidetes"       "Bacteroidia"                
## [3211,] "Bacteria" "Actinobacteria"      "Actinobacteria"             
## [3212,] "Bacteria" "Bacteroidetes"       "Bacteroidia"                
## [3213,] "Bacteria" "Firmicutes"          "Clostridia"                 
## [3214,] "Bacteria" "Firmicutes"          "Clostridia"                 
## [3215,] "Bacteria" "Bacteroidetes"       "Bacteroidia"                
## [3216,] "Bacteria" "Bacteroidetes"       "Bacteroidia"                
## [3217,] "Bacteria" "Fusobacteria"        "Fusobacteriia"              
## [3218,] "Bacteria" "Bacteroidetes"       "Bacteroidia"                
## [3219,] "Bacteria" "Bacteroidetes"       "Bacteroidia"                
## [3220,] "Bacteria" "Firmicutes"          "Negativicutes"              
## [3221,] "Bacteria" "Firmicutes"          "Clostridia"                 
## [3222,] "Bacteria" "Firmicutes"          "Clostridia"                 
## [3223,] "Bacteria" "Firmicutes"          "Bacilli"                    
## [3224,] "Bacteria" "Firmicutes"          "Clostridia"                 
## [3225,] "Bacteria" "Firmicutes"          "Clostridia"                 
## [3226,] "Bacteria" "Actinobacteria"      "Actinobacteria"             
## [3227,] "Bacteria" "Fusobacteria"        "Fusobacteriia"              
## [3228,] "Bacteria" "Fusobacteria"        "Fusobacteriia"              
## [3229,] "Bacteria" "Firmicutes"          "Negativicutes"              
## [3230,] "Bacteria" "Fusobacteria"        "Fusobacteriia"              
## [3231,] "Bacteria" "Firmicutes"          NA                           
## [3232,] "Bacteria" "Bacteroidetes"       "Bacteroidia"                
## [3233,] "Bacteria" "Fusobacteria"        "Fusobacteriia"              
## [3234,] "Bacteria" "Proteobacteria"      "Gammaproteobacteria"        
## [3235,] "Bacteria" "Bacteroidetes"       "Bacteroidia"                
## [3236,] "Bacteria" NA                    NA                           
## [3237,] "Bacteria" "Cyanobacteria"       "Oxyphotobacteria"           
## [3238,] "Bacteria" "Firmicutes"          "Negativicutes"              
## [3239,] "Bacteria" "Firmicutes"          "Bacilli"                    
## [3240,] "Bacteria" "Bacteroidetes"       "Bacteroidia"                
## [3241,] "Bacteria" "Bacteroidetes"       "Bacteroidia"                
## [3242,] "Bacteria" "Firmicutes"          "Negativicutes"              
## [3243,] "Bacteria" "Bacteroidetes"       "Bacteroidia"                
## [3244,] "Bacteria" "Firmicutes"          "Clostridia"                 
## [3245,] "Bacteria" "Actinobacteria"      "Actinobacteria"             
## [3246,] "Bacteria" "Firmicutes"          "Clostridia"                 
## [3247,] "Bacteria" "Epsilonbacteraeota"  "Campylobacteria"            
## [3248,] "Bacteria" "Actinobacteria"      "Coriobacteriia"             
## [3249,] "Bacteria" "Bacteroidetes"       "Bacteroidia"                
## [3250,] "Bacteria" "Patescibacteria"     "Saccharimonadia"            
## [3251,] "Bacteria" "Actinobacteria"      "Coriobacteriia"             
## [3252,] "Bacteria" "Actinobacteria"      "Actinobacteria"             
## [3253,] "Bacteria" "Bacteroidetes"       "Bacteroidia"                
## [3254,] "Bacteria" "Bacteroidetes"       "Bacteroidia"                
## [3255,] "Bacteria" "Proteobacteria"      "Gammaproteobacteria"        
## [3256,] "Bacteria" "Bacteroidetes"       "Bacteroidia"                
## [3257,] "Bacteria" "Patescibacteria"     "Saccharimonadia"            
## [3258,] "Bacteria" "Fusobacteria"        "Fusobacteriia"              
## [3259,] "Bacteria" "Firmicutes"          "Bacilli"                    
## [3260,] "Bacteria" "Epsilonbacteraeota"  "Campylobacteria"            
## [3261,] "Bacteria" "Epsilonbacteraeota"  "Campylobacteria"            
## [3262,] "Bacteria" "Actinobacteria"      "Actinobacteria"             
## [3263,] "Bacteria" "Patescibacteria"     "Saccharimonadia"            
## [3264,] "Bacteria" "Fusobacteria"        "Fusobacteriia"              
## [3265,] "Bacteria" "Actinobacteria"      "Actinobacteria"             
## [3266,] "Bacteria" "Bacteroidetes"       "Bacteroidia"                
## [3267,] "Bacteria" "Bacteroidetes"       "Bacteroidia"                
## [3268,] "Bacteria" "Firmicutes"          "Bacilli"                    
## [3269,] "Bacteria" "Bacteroidetes"       "Bacteroidia"                
## [3270,] "Bacteria" "Fusobacteria"        "Fusobacteriia"              
## [3271,] "Bacteria" "Bacteroidetes"       "Bacteroidia"                
## [3272,] "Bacteria" "Bacteroidetes"       "Bacteroidia"                
## [3273,] "Bacteria" "Bacteroidetes"       "Bacteroidia"                
## [3274,] "Bacteria" "Patescibacteria"     "Saccharimonadia"            
## [3275,] "Bacteria" "Bacteroidetes"       "Bacteroidia"                
## [3276,] "Bacteria" "Actinobacteria"      "Actinobacteria"             
## [3277,] "Bacteria" NA                    NA                           
## [3278,] "Bacteria" "Firmicutes"          "Bacilli"                    
## [3279,] "Bacteria" "Spirochaetes"        "Spirochaetia"               
## [3280,] "Bacteria" "Bacteroidetes"       "Bacteroidia"                
## [3281,] "Bacteria" "Firmicutes"          "Negativicutes"              
## [3282,] "Bacteria" "Proteobacteria"      "Gammaproteobacteria"        
## [3283,] "Bacteria" "Bacteroidetes"       "Bacteroidia"                
## [3284,] "Bacteria" "Bacteroidetes"       "Bacteroidia"                
## [3285,] "Bacteria" "Proteobacteria"      "Gammaproteobacteria"        
## [3286,] "Bacteria" "Proteobacteria"      "Gammaproteobacteria"        
## [3287,] "Bacteria" "Proteobacteria"      "Gammaproteobacteria"        
## [3288,] "Bacteria" "Spirochaetes"        "Spirochaetia"               
## [3289,] "Bacteria" "Firmicutes"          "Clostridia"                 
## [3290,] "Bacteria" "Spirochaetes"        "Spirochaetia"               
## [3291,] "Bacteria" "Bacteroidetes"       "Bacteroidia"                
## [3292,] "Bacteria" "Proteobacteria"      "Gammaproteobacteria"        
## [3293,] "Bacteria" "Firmicutes"          "Negativicutes"              
## [3294,] "Bacteria" "Bacteroidetes"       "Bacteroidia"                
## [3295,] "Bacteria" "Bacteroidetes"       "Bacteroidia"                
## [3296,] "Bacteria" "Bacteroidetes"       "Bacteroidia"                
## [3297,] "Bacteria" "Actinobacteria"      "Actinobacteria"             
## [3298,] "Bacteria" "Bacteroidetes"       "Bacteroidia"                
## [3299,] "Bacteria" "Firmicutes"          "Bacilli"                    
## [3300,] "Bacteria" "Bacteroidetes"       "Bacteroidia"                
## [3301,] "Bacteria" "Bacteroidetes"       "Bacteroidia"                
## [3302,] "Bacteria" "Bacteroidetes"       "Bacteroidia"                
## [3303,] "Bacteria" "Bacteroidetes"       "Bacteroidia"                
## [3304,] "Bacteria" "Proteobacteria"      "Gammaproteobacteria"        
## [3305,] "Bacteria" "Proteobacteria"      "Gammaproteobacteria"        
## [3306,] "Bacteria" "Bacteroidetes"       "Bacteroidia"                
## [3307,] "Bacteria" "Bacteroidetes"       "Bacteroidia"                
## [3308,] "Bacteria" "Patescibacteria"     "Saccharimonadia"            
## [3309,] "Bacteria" "Proteobacteria"      "Gammaproteobacteria"        
## [3310,] "Bacteria" "Bacteroidetes"       "Bacteroidia"                
## [3311,] "Bacteria" "Bacteroidetes"       "Bacteroidia"                
## [3312,] "Bacteria" "Fusobacteria"        "Fusobacteriia"              
## [3313,] "Bacteria" "Bacteroidetes"       "Bacteroidia"                
## [3314,] "Bacteria" "Spirochaetes"        "Spirochaetia"               
## [3315,] "Bacteria" "Bacteroidetes"       "Bacteroidia"                
## [3316,] "Bacteria" "Patescibacteria"     "Saccharimonadia"            
## [3317,] "Bacteria" "Firmicutes"          "Clostridia"                 
## [3318,] "Bacteria" "Fusobacteria"        "Fusobacteriia"              
## [3319,] "Bacteria" "Proteobacteria"      "Gammaproteobacteria"        
## [3320,] "Bacteria" "Bacteroidetes"       "Bacteroidia"                
## [3321,] "Bacteria" "Bacteroidetes"       "Bacteroidia"                
## [3322,] "Bacteria" "Fusobacteria"        "Fusobacteriia"              
## [3323,] "Bacteria" "Bacteroidetes"       "Bacteroidia"                
## [3324,] "Bacteria" "Bacteroidetes"       "Bacteroidia"                
## [3325,] "Bacteria" "Bacteroidetes"       "Bacteroidia"                
## [3326,] "Bacteria" NA                    NA                           
## [3327,] "Bacteria" "Actinobacteria"      "Actinobacteria"             
## [3328,] "Bacteria" "Bacteroidetes"       "Bacteroidia"                
## [3329,] "Bacteria" "Bacteroidetes"       "Bacteroidia"                
## [3330,] "Bacteria" "Spirochaetes"        "Spirochaetia"               
## [3331,] "Bacteria" "Bacteroidetes"       "Bacteroidia"                
## [3332,] "Bacteria" "Bacteroidetes"       "Bacteroidia"                
## [3333,] "Bacteria" "Firmicutes"          "Negativicutes"              
## [3334,] "Bacteria" "Actinobacteria"      "Actinobacteria"             
## [3335,] "Bacteria" "Firmicutes"          "Bacilli"                    
## [3336,] "Bacteria" "Proteobacteria"      "Gammaproteobacteria"        
## [3337,] "Bacteria" "Bacteroidetes"       "Bacteroidia"                
## [3338,] "Bacteria" "Fusobacteria"        "Fusobacteriia"              
## [3339,] "Bacteria" "Patescibacteria"     "Saccharimonadia"            
## [3340,] "Bacteria" "Patescibacteria"     "Saccharimonadia"            
## [3341,] "Bacteria" "Firmicutes"          "Clostridia"                 
## [3342,] "Bacteria" "Fusobacteria"        "Fusobacteriia"              
## [3343,] "Bacteria" "Firmicutes"          NA                           
## [3344,] "Bacteria" "Firmicutes"          "Clostridia"                 
## [3345,] "Bacteria" "Patescibacteria"     "Saccharimonadia"            
## [3346,] "Bacteria" "Epsilonbacteraeota"  "Campylobacteria"            
## [3347,] "Bacteria" "Bacteroidetes"       "Bacteroidia"                
## [3348,] "Bacteria" "Proteobacteria"      "Gammaproteobacteria"        
## [3349,] "Bacteria" "Bacteroidetes"       "Bacteroidia"                
## [3350,] "Bacteria" "Fusobacteria"        "Fusobacteriia"              
## [3351,] "Bacteria" "Proteobacteria"      "Gammaproteobacteria"        
## [3352,] "Bacteria" "Proteobacteria"      "Gammaproteobacteria"        
## [3353,] "Bacteria" "Bacteroidetes"       "Bacteroidia"                
## [3354,] "Bacteria" "Proteobacteria"      "Gammaproteobacteria"        
## [3355,] "Bacteria" "Proteobacteria"      "Gammaproteobacteria"        
## [3356,] "Bacteria" "Firmicutes"          "Negativicutes"              
## [3357,] "Bacteria" "Actinobacteria"      "Coriobacteriia"             
## [3358,] "Bacteria" "Fusobacteria"        "Fusobacteriia"              
## [3359,] "Bacteria" "Spirochaetes"        "Spirochaetia"               
## [3360,] "Bacteria" "Firmicutes"          "Negativicutes"              
## [3361,] "Bacteria" "Actinobacteria"      "Actinobacteria"             
## [3362,] "Bacteria" "Bacteroidetes"       "Bacteroidia"                
## [3363,] "Bacteria" "Firmicutes"          "Negativicutes"              
## [3364,] "Bacteria" "Firmicutes"          "Clostridia"                 
## [3365,] "Bacteria" "Bacteroidetes"       "Bacteroidia"                
## [3366,] "Bacteria" "Bacteroidetes"       "Bacteroidia"                
## [3367,] "Bacteria" "Proteobacteria"      "Gammaproteobacteria"        
## [3368,] "Bacteria" "Actinobacteria"      "Actinobacteria"             
## [3369,] "Bacteria" "Bacteroidetes"       "Bacteroidia"                
## [3370,] "Bacteria" "Firmicutes"          "Negativicutes"              
## [3371,] "Bacteria" "Bacteroidetes"       "Bacteroidia"                
## [3372,] "Bacteria" "Fusobacteria"        "Fusobacteriia"              
## [3373,] "Bacteria" "Fusobacteria"        "Fusobacteriia"              
## [3374,] "Bacteria" "Spirochaetes"        "Spirochaetia"               
## [3375,] "Bacteria" "Bacteroidetes"       "Bacteroidia"                
## [3376,] "Bacteria" "Bacteroidetes"       "Bacteroidia"                
## [3377,] "Bacteria" "Bacteroidetes"       "Bacteroidia"                
## [3378,] "Bacteria" "Fusobacteria"        "Fusobacteriia"              
## [3379,] "Bacteria" "Bacteroidetes"       "Bacteroidia"                
## [3380,] "Bacteria" "Bacteroidetes"       "Bacteroidia"                
## [3381,] "Bacteria" "Bacteroidetes"       "Bacteroidia"                
## [3382,] "Bacteria" "Bacteroidetes"       "Bacteroidia"                
## [3383,] "Bacteria" "Firmicutes"          "Clostridia"                 
## [3384,] "Bacteria" "Bacteroidetes"       "Bacteroidia"                
## [3385,] "Bacteria" "Spirochaetes"        "Spirochaetia"               
## [3386,] "Bacteria" "Bacteroidetes"       "Bacteroidia"                
## [3387,] "Bacteria" "Firmicutes"          "Bacilli"                    
## [3388,] "Bacteria" "Firmicutes"          "Clostridia"                 
## [3389,] "Bacteria" "Bacteroidetes"       "Bacteroidia"                
## [3390,] "Bacteria" "Bacteroidetes"       "Bacteroidia"                
## [3391,] "Bacteria" "Firmicutes"          "Bacilli"                    
## [3392,] "Bacteria" "Bacteroidetes"       "Bacteroidia"                
## [3393,] "Bacteria" "Patescibacteria"     "Saccharimonadia"            
## [3394,] "Bacteria" NA                    NA                           
## [3395,] "Bacteria" "Fusobacteria"        "Fusobacteriia"              
## [3396,] "Bacteria" "Actinobacteria"      "Actinobacteria"             
## [3397,] "Bacteria" "Bacteroidetes"       "Bacteroidia"                
## [3398,] "Bacteria" "Actinobacteria"      "Actinobacteria"             
## [3399,] "Bacteria" "Firmicutes"          "Clostridia"                 
## [3400,] "Bacteria" "Bacteroidetes"       "Bacteroidia"                
## [3401,] "Bacteria" "Proteobacteria"      "Gammaproteobacteria"        
## [3402,] "Bacteria" "Proteobacteria"      "Gammaproteobacteria"        
## [3403,] "Bacteria" "Firmicutes"          "Clostridia"                 
## [3404,] "Bacteria" "Bacteroidetes"       "Bacteroidia"                
## [3405,] "Bacteria" "Bacteroidetes"       "Bacteroidia"                
## [3406,] "Bacteria" "Bacteroidetes"       "Bacteroidia"                
## [3407,] "Bacteria" "Bacteroidetes"       "Bacteroidia"                
## [3408,] "Bacteria" "Firmicutes"          "Bacilli"                    
## [3409,] "Bacteria" "Bacteroidetes"       "Bacteroidia"                
## [3410,] "Bacteria" "Proteobacteria"      "Gammaproteobacteria"        
## [3411,] "Bacteria" "Actinobacteria"      "Coriobacteriia"             
## [3412,] "Bacteria" "Bacteroidetes"       "Bacteroidia"                
## [3413,] "Bacteria" "Spirochaetes"        "Spirochaetia"               
## [3414,] "Bacteria" "Bacteroidetes"       "Bacteroidia"                
## [3415,] "Bacteria" "Bacteroidetes"       "Bacteroidia"                
## [3416,] "Bacteria" "Bacteroidetes"       "Bacteroidia"                
## [3417,] "Bacteria" "Fusobacteria"        "Fusobacteriia"              
## [3418,] "Bacteria" "Bacteroidetes"       "Bacteroidia"                
## [3419,] "Bacteria" "Firmicutes"          "Negativicutes"              
## [3420,] "Bacteria" "Firmicutes"          NA                           
## [3421,] "Bacteria" "Bacteroidetes"       "Bacteroidia"                
## [3422,] "Bacteria" "Firmicutes"          "Clostridia"                 
## [3423,] "Bacteria" "Proteobacteria"      "Gammaproteobacteria"        
## [3424,] "Bacteria" "Bacteroidetes"       "Bacteroidia"                
## [3425,] "Bacteria" "Proteobacteria"      "Gammaproteobacteria"        
## [3426,] "Bacteria" "Bacteroidetes"       "Bacteroidia"                
## [3427,] "Bacteria" "Bacteroidetes"       "Bacteroidia"                
## [3428,] "Bacteria" "Bacteroidetes"       "Bacteroidia"                
## [3429,] "Bacteria" "Proteobacteria"      "Gammaproteobacteria"        
## [3430,] "Bacteria" "Firmicutes"          "Bacilli"                    
## [3431,] "Bacteria" "Bacteroidetes"       "Bacteroidia"                
## [3432,] "Bacteria" "Bacteroidetes"       "Bacteroidia"                
## [3433,] "Bacteria" "Bacteroidetes"       "Bacteroidia"                
## [3434,] "Bacteria" "Bacteroidetes"       "Bacteroidia"                
## [3435,] "Bacteria" "Firmicutes"          NA                           
## [3436,] "Bacteria" "Proteobacteria"      "Gammaproteobacteria"        
## [3437,] "Bacteria" "Proteobacteria"      "Gammaproteobacteria"        
## [3438,] "Bacteria" "Firmicutes"          "Negativicutes"              
## [3439,] "Bacteria" "Proteobacteria"      "Gammaproteobacteria"        
## [3440,] "Bacteria" "Firmicutes"          "Bacilli"                    
## [3441,] "Bacteria" "Bacteroidetes"       "Bacteroidia"                
## [3442,] "Bacteria" "Actinobacteria"      "Coriobacteriia"             
## [3443,] "Bacteria" "Firmicutes"          "Clostridia"                 
## [3444,] "Bacteria" "Firmicutes"          "Negativicutes"              
## [3445,] "Bacteria" "Bacteroidetes"       "Bacteroidia"                
## [3446,] "Bacteria" "Firmicutes"          "Clostridia"                 
## [3447,] "Bacteria" "Epsilonbacteraeota"  "Campylobacteria"            
## [3448,] "Bacteria" "Bacteroidetes"       "Bacteroidia"                
## [3449,] "Bacteria" "Bacteroidetes"       "Bacteroidia"                
## [3450,] "Bacteria" "Fusobacteria"        "Fusobacteriia"              
## [3451,] "Bacteria" "Bacteroidetes"       "Bacteroidia"                
## [3452,] "Bacteria" "Fusobacteria"        "Fusobacteriia"              
## [3453,] "Bacteria" "Bacteroidetes"       "Bacteroidia"                
## [3454,] "Bacteria" "Bacteroidetes"       "Bacteroidia"                
## [3455,] "Bacteria" "Firmicutes"          "Negativicutes"              
## [3456,] "Bacteria" "Bacteroidetes"       "Bacteroidia"                
## [3457,] "Bacteria" "Fusobacteria"        "Fusobacteriia"              
## [3458,] "Bacteria" "Firmicutes"          "Negativicutes"              
## [3459,] "Bacteria" "Proteobacteria"      "Gammaproteobacteria"        
## [3460,] "Bacteria" "Actinobacteria"      "Actinobacteria"             
## [3461,] "Bacteria" "Firmicutes"          "Negativicutes"              
## [3462,] "Bacteria" "Proteobacteria"      "Gammaproteobacteria"        
## [3463,] "Bacteria" "Firmicutes"          "Clostridia"                 
## [3464,] "Bacteria" "Bacteroidetes"       "Bacteroidia"                
## [3465,] "Bacteria" "Bacteroidetes"       "Bacteroidia"                
## [3466,] "Bacteria" "Bacteroidetes"       "Bacteroidia"                
## [3467,] "Bacteria" "Firmicutes"          "Negativicutes"              
## [3468,] "Bacteria" "Firmicutes"          "Bacilli"                    
## [3469,] "Bacteria" "Bacteroidetes"       "Bacteroidia"                
## [3470,] "Bacteria" "Bacteroidetes"       "Bacteroidia"                
## [3471,] "Bacteria" NA                    NA                           
## [3472,] "Bacteria" "Proteobacteria"      "Gammaproteobacteria"        
## [3473,] "Bacteria" "Firmicutes"          "Clostridia"                 
## [3474,] "Bacteria" "Actinobacteria"      "Actinobacteria"             
## [3475,] "Bacteria" "Bacteroidetes"       "Bacteroidia"                
## [3476,] "Bacteria" "Bacteroidetes"       "Bacteroidia"                
## [3477,] "Bacteria" "Bacteroidetes"       "Bacteroidia"                
## [3478,] "Bacteria" "Proteobacteria"      "Gammaproteobacteria"        
## [3479,] "Bacteria" "Bacteroidetes"       "Bacteroidia"                
## [3480,] "Bacteria" "Firmicutes"          "Negativicutes"              
## [3481,] "Bacteria" "Proteobacteria"      "Gammaproteobacteria"        
## [3482,] "Bacteria" "Bacteroidetes"       "Bacteroidia"                
## [3483,] "Bacteria" "Firmicutes"          "Bacilli"                    
## [3484,] "Bacteria" "Firmicutes"          "Clostridia"                 
## [3485,] "Bacteria" "Bacteroidetes"       "Bacteroidia"                
## [3486,] "Bacteria" "Bacteroidetes"       "Bacteroidia"                
## [3487,] "Bacteria" "Bacteroidetes"       "Bacteroidia"                
## [3488,] "Bacteria" "Firmicutes"          "Negativicutes"              
## [3489,] "Bacteria" "Firmicutes"          "Negativicutes"              
## [3490,] "Bacteria" "Patescibacteria"     "Saccharimonadia"            
## [3491,] "Bacteria" "Bacteroidetes"       "Bacteroidia"                
## [3492,] "Bacteria" "Firmicutes"          "Negativicutes"              
## [3493,] "Bacteria" "Firmicutes"          "Negativicutes"              
## [3494,] "Bacteria" "Actinobacteria"      "Actinobacteria"             
## [3495,] "Bacteria" "Firmicutes"          NA                           
## [3496,] "Bacteria" "Proteobacteria"      "Gammaproteobacteria"        
## [3497,] "Bacteria" "Bacteroidetes"       "Bacteroidia"                
## [3498,] "Bacteria" "Cyanobacteria"       "Oxyphotobacteria"           
## [3499,] "Bacteria" "Firmicutes"          "Bacilli"                    
## [3500,] "Bacteria" "Bacteroidetes"       "Bacteroidia"                
## [3501,] "Bacteria" "Bacteroidetes"       "Bacteroidia"                
## [3502,] "Bacteria" "Firmicutes"          "Bacilli"                    
## [3503,] "Bacteria" "Proteobacteria"      "Gammaproteobacteria"        
## [3504,] "Bacteria" "Actinobacteria"      "Actinobacteria"             
## [3505,] "Bacteria" "Bacteroidetes"       "Bacteroidia"                
## [3506,] "Bacteria" "Bacteroidetes"       "Bacteroidia"                
## [3507,] "Bacteria" "Fusobacteria"        "Fusobacteriia"              
## [3508,] "Bacteria" "Fusobacteria"        "Fusobacteriia"              
## [3509,] "Bacteria" "Fusobacteria"        "Fusobacteriia"              
## [3510,] "Bacteria" "Patescibacteria"     "Saccharimonadia"            
## [3511,] "Bacteria" "Bacteroidetes"       "Bacteroidia"                
## [3512,] "Bacteria" "Proteobacteria"      "Gammaproteobacteria"        
## [3513,] "Bacteria" "Firmicutes"          "Clostridia"                 
## [3514,] "Bacteria" "Bacteroidetes"       "Bacteroidia"                
## [3515,] "Bacteria" "Firmicutes"          "Clostridia"                 
## [3516,] "Bacteria" "Firmicutes"          "Negativicutes"              
## [3517,] "Bacteria" "Firmicutes"          "Negativicutes"              
## [3518,] "Bacteria" "Proteobacteria"      "Alphaproteobacteria"        
## [3519,] "Bacteria" "Firmicutes"          "Negativicutes"              
## [3520,] "Bacteria" "Proteobacteria"      "Gammaproteobacteria"        
## [3521,] "Bacteria" "Proteobacteria"      "Gammaproteobacteria"        
## [3522,] "Bacteria" "Bacteroidetes"       "Bacteroidia"                
## [3523,] "Bacteria" "Firmicutes"          "Negativicutes"              
## [3524,] "Bacteria" "Proteobacteria"      "Gammaproteobacteria"        
## [3525,] "Bacteria" "Bacteroidetes"       "Bacteroidia"                
## [3526,] "Bacteria" "Firmicutes"          "Bacilli"                    
## [3527,] "Bacteria" "Proteobacteria"      "Gammaproteobacteria"        
## [3528,] "Bacteria" "Proteobacteria"      "Gammaproteobacteria"        
## [3529,] "Bacteria" "Bacteroidetes"       "Bacteroidia"                
## [3530,] "Bacteria" "Firmicutes"          "Bacilli"                    
## [3531,] "Bacteria" "Bacteroidetes"       "Bacteroidia"                
## [3532,] "Bacteria" "Bacteroidetes"       "Bacteroidia"                
## [3533,] "Bacteria" NA                    NA                           
## [3534,] "Bacteria" "Fusobacteria"        "Fusobacteriia"              
## [3535,] "Bacteria" "Firmicutes"          NA                           
## [3536,] "Bacteria" "Patescibacteria"     "Gracilibacteria"            
## [3537,] "Bacteria" "Patescibacteria"     "Saccharimonadia"            
## [3538,] "Bacteria" "Firmicutes"          "Negativicutes"              
## [3539,] "Bacteria" "Fusobacteria"        "Fusobacteriia"              
## [3540,] "Bacteria" "Bacteroidetes"       "Bacteroidia"                
## [3541,] "Bacteria" "Bacteroidetes"       "Bacteroidia"                
## [3542,] "Bacteria" "Fusobacteria"        "Fusobacteriia"              
## [3543,] "Bacteria" "Fusobacteria"        "Fusobacteriia"              
## [3544,] "Bacteria" "Proteobacteria"      "Gammaproteobacteria"        
## [3545,] "Bacteria" "Fusobacteria"        "Fusobacteriia"              
## [3546,] "Bacteria" "Firmicutes"          "Clostridia"                 
## [3547,] "Bacteria" "Proteobacteria"      "Gammaproteobacteria"        
## [3548,] "Bacteria" "Bacteroidetes"       "Bacteroidia"                
## [3549,] "Bacteria" "Actinobacteria"      "Actinobacteria"             
## [3550,] "Bacteria" NA                    NA                           
## [3551,] "Bacteria" "Firmicutes"          "Clostridia"                 
## [3552,] "Bacteria" "Firmicutes"          "Clostridia"                 
## [3553,] "Bacteria" "Proteobacteria"      "Gammaproteobacteria"        
## [3554,] "Bacteria" "Proteobacteria"      "Gammaproteobacteria"        
## [3555,] "Bacteria" "Spirochaetes"        "Spirochaetia"               
## [3556,] "Bacteria" "Firmicutes"          "Bacilli"                    
## [3557,] "Bacteria" "Firmicutes"          "Bacilli"                    
## [3558,] "Bacteria" "Actinobacteria"      "Actinobacteria"             
## [3559,] "Bacteria" "Actinobacteria"      "Actinobacteria"             
## [3560,] "Bacteria" "Firmicutes"          "Bacilli"                    
## [3561,] "Bacteria" "Spirochaetes"        "Spirochaetia"               
## [3562,] "Bacteria" "Firmicutes"          "Clostridia"                 
## [3563,] "Bacteria" "Epsilonbacteraeota"  "Campylobacteria"            
## [3564,] "Bacteria" "Firmicutes"          "Clostridia"                 
## [3565,] "Bacteria" "Bacteroidetes"       "Bacteroidia"                
## [3566,] "Bacteria" "Firmicutes"          "Negativicutes"              
## [3567,] "Bacteria" "Proteobacteria"      "Gammaproteobacteria"        
## [3568,] "Bacteria" "Firmicutes"          "Bacilli"                    
## [3569,] "Bacteria" "Firmicutes"          "Negativicutes"              
## [3570,] "Bacteria" "Bacteroidetes"       "Bacteroidia"                
## [3571,] "Bacteria" "Bacteroidetes"       "Bacteroidia"                
## [3572,] "Bacteria" "Epsilonbacteraeota"  "Campylobacteria"            
## [3573,] "Bacteria" "Fusobacteria"        "Fusobacteriia"              
## [3574,] "Bacteria" "Bacteroidetes"       "Bacteroidia"                
## [3575,] "Bacteria" "Firmicutes"          "Negativicutes"              
## [3576,] "Bacteria" "Firmicutes"          "Clostridia"                 
## [3577,] "Bacteria" "Firmicutes"          "Bacilli"                    
## [3578,] "Bacteria" "Firmicutes"          "Clostridia"                 
## [3579,] "Bacteria" "Bacteroidetes"       "Bacteroidia"                
## [3580,] "Bacteria" "Firmicutes"          "Clostridia"                 
## [3581,] "Bacteria" "Bacteroidetes"       "Bacteroidia"                
## [3582,] "Bacteria" "Spirochaetes"        "Spirochaetia"               
## [3583,] "Bacteria" "Firmicutes"          "Clostridia"                 
## [3584,] "Bacteria" "Actinobacteria"      "Actinobacteria"             
## [3585,] "Bacteria" "Bacteroidetes"       "Bacteroidia"                
## [3586,] "Bacteria" "Proteobacteria"      "Gammaproteobacteria"        
## [3587,] "Bacteria" "Bacteroidetes"       "Bacteroidia"                
## [3588,] "Bacteria" "Bacteroidetes"       "Bacteroidia"                
## [3589,] "Bacteria" "Proteobacteria"      NA                           
## [3590,] "Bacteria" "Epsilonbacteraeota"  "Campylobacteria"            
## [3591,] "Bacteria" "Firmicutes"          "Bacilli"                    
## [3592,] "Bacteria" "Bacteroidetes"       "Bacteroidia"                
## [3593,] "Bacteria" "Firmicutes"          "Bacilli"                    
## [3594,] "Bacteria" "Bacteroidetes"       "Bacteroidia"                
## [3595,] "Bacteria" NA                    NA                           
## [3596,] "Bacteria" "Fusobacteria"        "Fusobacteriia"              
## [3597,] "Bacteria" "Bacteroidetes"       "Bacteroidia"                
## [3598,] "Bacteria" "Bacteroidetes"       "Bacteroidia"                
## [3599,] "Bacteria" NA                    NA                           
## [3600,] "Bacteria" "Epsilonbacteraeota"  "Campylobacteria"            
## [3601,] "Bacteria" "Bacteroidetes"       "Bacteroidia"                
## [3602,] "Bacteria" "Proteobacteria"      "Gammaproteobacteria"        
## [3603,] "Bacteria" "Firmicutes"          "Negativicutes"              
## [3604,] "Bacteria" "Proteobacteria"      "Gammaproteobacteria"        
## [3605,] "Bacteria" "Actinobacteria"      "Actinobacteria"             
## [3606,] "Bacteria" "Bacteroidetes"       "Bacteroidia"                
## [3607,] "Bacteria" "Firmicutes"          "Bacilli"                    
## [3608,] "Bacteria" "Proteobacteria"      "Gammaproteobacteria"        
## [3609,] "Bacteria" "Fusobacteria"        "Fusobacteriia"              
## [3610,] "Bacteria" "Actinobacteria"      "Actinobacteria"             
## [3611,] "Bacteria" "Bacteroidetes"       "Bacteroidia"                
## [3612,] "Bacteria" "Firmicutes"          "Clostridia"                 
## [3613,] "Bacteria" "Fusobacteria"        "Fusobacteriia"              
## [3614,] "Bacteria" "Firmicutes"          "Negativicutes"              
## [3615,] "Bacteria" "Fusobacteria"        "Fusobacteriia"              
## [3616,] "Bacteria" "Firmicutes"          "Clostridia"                 
## [3617,] "Bacteria" "Fusobacteria"        "Fusobacteriia"              
## [3618,] "Bacteria" "Firmicutes"          "Clostridia"                 
## [3619,] "Bacteria" "Proteobacteria"      "Gammaproteobacteria"        
## [3620,] "Bacteria" "Actinobacteria"      "Coriobacteriia"             
## [3621,] "Bacteria" "Actinobacteria"      "Actinobacteria"             
## [3622,] "Bacteria" "Bacteroidetes"       "Bacteroidia"                
## [3623,] "Bacteria" "Proteobacteria"      "Gammaproteobacteria"        
## [3624,] "Bacteria" "Bacteroidetes"       "Bacteroidia"                
## [3625,] "Bacteria" "Bacteroidetes"       "Bacteroidia"                
## [3626,] "Bacteria" "Bacteroidetes"       "Bacteroidia"                
## [3627,] "Bacteria" "Proteobacteria"      "Gammaproteobacteria"        
## [3628,] "Bacteria" "Firmicutes"          "Clostridia"                 
## [3629,] "Bacteria" "Bacteroidetes"       "Bacteroidia"                
## [3630,] "Bacteria" "Patescibacteria"     "Saccharimonadia"            
## [3631,] "Bacteria" "Bacteroidetes"       "Bacteroidia"                
## [3632,] "Bacteria" "Bacteroidetes"       "Bacteroidia"                
## [3633,] "Bacteria" "Proteobacteria"      "Gammaproteobacteria"        
## [3634,] "Bacteria" "Firmicutes"          "Bacilli"                    
## [3635,] "Bacteria" "Bacteroidetes"       "Bacteroidia"                
## [3636,] "Bacteria" "Patescibacteria"     "Saccharimonadia"            
## [3637,] "Bacteria" "Bacteroidetes"       "Bacteroidia"                
## [3638,] "Bacteria" "Firmicutes"          "Clostridia"                 
## [3639,] "Bacteria" "Firmicutes"          "Negativicutes"              
## [3640,] "Bacteria" "Firmicutes"          "Negativicutes"              
## [3641,] "Bacteria" "Firmicutes"          "Negativicutes"              
## [3642,] "Bacteria" "Proteobacteria"      "Alphaproteobacteria"        
## [3643,] "Bacteria" "Proteobacteria"      "Gammaproteobacteria"        
## [3644,] "Bacteria" "Bacteroidetes"       "Bacteroidia"                
## [3645,] "Bacteria" "Proteobacteria"      "Gammaproteobacteria"        
## [3646,] "Bacteria" "Fusobacteria"        "Fusobacteriia"              
## [3647,] "Bacteria" "Actinobacteria"      "Actinobacteria"             
## [3648,] "Bacteria" "Firmicutes"          NA                           
## [3649,] "Bacteria" "Firmicutes"          "Clostridia"                 
## [3650,] "Bacteria" "Spirochaetes"        "Spirochaetia"               
## [3651,] "Bacteria" "Firmicutes"          "Clostridia"                 
## [3652,] "Bacteria" "Bacteroidetes"       "Bacteroidia"                
## [3653,] "Bacteria" NA                    NA                           
## [3654,] "Bacteria" "Bacteroidetes"       "Bacteroidia"                
## [3655,] "Bacteria" "Bacteroidetes"       "Bacteroidia"                
## [3656,] "Bacteria" "Proteobacteria"      "Gammaproteobacteria"        
## [3657,] "Bacteria" NA                    NA                           
## [3658,] "Bacteria" "Bacteroidetes"       "Bacteroidia"                
## [3659,] "Bacteria" "Firmicutes"          "Negativicutes"              
## [3660,] "Bacteria" "Epsilonbacteraeota"  "Campylobacteria"            
## [3661,] "Bacteria" "Firmicutes"          "Clostridia"                 
## [3662,] "Bacteria" "Actinobacteria"      "Actinobacteria"             
## [3663,] "Bacteria" "Bacteroidetes"       "Bacteroidia"                
## [3664,] "Bacteria" "Bacteroidetes"       "Bacteroidia"                
## [3665,] "Bacteria" "Bacteroidetes"       "Bacteroidia"                
## [3666,] "Bacteria" "Firmicutes"          "Negativicutes"              
## [3667,] "Bacteria" "Proteobacteria"      "Gammaproteobacteria"        
## [3668,] "Bacteria" "Proteobacteria"      "Gammaproteobacteria"        
## [3669,] "Bacteria" NA                    NA                           
## [3670,] "Bacteria" "Actinobacteria"      "Coriobacteriia"             
## [3671,] "Bacteria" "Firmicutes"          NA                           
## [3672,] "Bacteria" "Firmicutes"          NA                           
## [3673,] "Bacteria" "Bacteroidetes"       "Bacteroidia"                
## [3674,] "Bacteria" "Bacteroidetes"       "Bacteroidia"                
## [3675,] "Bacteria" "Firmicutes"          "Clostridia"                 
## [3676,] "Bacteria" "Bacteroidetes"       "Bacteroidia"                
## [3677,] "Bacteria" "Spirochaetes"        "Spirochaetia"               
## [3678,] "Bacteria" "Bacteroidetes"       "Bacteroidia"                
## [3679,] "Bacteria" "Bacteroidetes"       "Bacteroidia"                
## [3680,] "Bacteria" "Bacteroidetes"       "Bacteroidia"                
## [3681,] "Bacteria" "Firmicutes"          "Negativicutes"              
## [3682,] "Bacteria" "Bacteroidetes"       "Bacteroidia"                
## [3683,] "Bacteria" "Proteobacteria"      "Gammaproteobacteria"        
## [3684,] "Bacteria" "Proteobacteria"      "Gammaproteobacteria"        
## [3685,] "Bacteria" "Firmicutes"          "Bacilli"                    
## [3686,] "Bacteria" "Firmicutes"          "Clostridia"                 
## [3687,] "Bacteria" "Firmicutes"          NA                           
## [3688,] "Bacteria" "Bacteroidetes"       "Bacteroidia"                
## [3689,] "Bacteria" "Bacteroidetes"       "Bacteroidia"                
## [3690,] "Bacteria" "Firmicutes"          "Clostridia"                 
## [3691,] "Bacteria" "Fusobacteria"        "Fusobacteriia"              
## [3692,] "Bacteria" "Fusobacteria"        "Fusobacteriia"              
## [3693,] "Bacteria" "Bacteroidetes"       "Bacteroidia"                
## [3694,] "Bacteria" "Spirochaetes"        "Spirochaetia"               
## [3695,] "Bacteria" "Firmicutes"          "Clostridia"                 
## [3696,] "Bacteria" "Firmicutes"          "Negativicutes"              
## [3697,] "Bacteria" "Firmicutes"          "Clostridia"                 
## [3698,] "Bacteria" "Proteobacteria"      "Gammaproteobacteria"        
## [3699,] "Bacteria" "Fusobacteria"        "Fusobacteriia"              
## [3700,] "Bacteria" "Bacteroidetes"       "Bacteroidia"                
## [3701,] "Bacteria" "Firmicutes"          "Negativicutes"              
## [3702,] "Bacteria" "Epsilonbacteraeota"  "Campylobacteria"            
## [3703,] "Bacteria" NA                    NA                           
## [3704,] "Bacteria" "Fusobacteria"        "Fusobacteriia"              
## [3705,] "Bacteria" "Proteobacteria"      "Gammaproteobacteria"        
## [3706,] "Bacteria" "Firmicutes"          NA                           
## [3707,] "Bacteria" "Bacteroidetes"       "Bacteroidia"                
## [3708,] "Bacteria" "Proteobacteria"      "Gammaproteobacteria"        
## [3709,] "Bacteria" "Bacteroidetes"       "Bacteroidia"                
## [3710,] "Bacteria" "Bacteroidetes"       "Bacteroidia"                
## [3711,] "Bacteria" "Firmicutes"          "Clostridia"                 
## [3712,] "Bacteria" "Actinobacteria"      "Coriobacteriia"             
## [3713,] "Bacteria" "Firmicutes"          "Bacilli"                    
## [3714,] "Bacteria" "Firmicutes"          "Clostridia"                 
## [3715,] "Bacteria" "Firmicutes"          "Clostridia"                 
## [3716,] "Bacteria" "Proteobacteria"      "Gammaproteobacteria"        
## [3717,] "Bacteria" "Firmicutes"          "Bacilli"                    
## [3718,] "Bacteria" "Fusobacteria"        "Fusobacteriia"              
## [3719,] "Bacteria" "Patescibacteria"     "Saccharimonadia"            
## [3720,] "Bacteria" "Firmicutes"          "Bacilli"                    
## [3721,] "Bacteria" "Firmicutes"          "Negativicutes"              
## [3722,] "Bacteria" "Firmicutes"          "Bacilli"                    
## [3723,] "Bacteria" "Proteobacteria"      "Gammaproteobacteria"        
## [3724,] "Bacteria" "Firmicutes"          "Clostridia"                 
## [3725,] "Bacteria" "Firmicutes"          "Clostridia"                 
## [3726,] "Bacteria" "Epsilonbacteraeota"  "Campylobacteria"            
## [3727,] "Bacteria" "Epsilonbacteraeota"  "Campylobacteria"            
## [3728,] "Bacteria" "Firmicutes"          "Clostridia"                 
## [3729,] "Bacteria" "Firmicutes"          "Clostridia"                 
## [3730,] "Bacteria" "Firmicutes"          "Negativicutes"              
## [3731,] "Bacteria" "Firmicutes"          "Bacilli"                    
## [3732,] "Bacteria" "Spirochaetes"        "Spirochaetia"               
## [3733,] "Bacteria" "Firmicutes"          "Negativicutes"              
## [3734,] "Bacteria" "Bacteroidetes"       "Bacteroidia"                
## [3735,] "Bacteria" "Fusobacteria"        "Fusobacteriia"              
## [3736,] "Bacteria" "Firmicutes"          "Bacilli"                    
## [3737,] "Bacteria" "Bacteroidetes"       "Bacteroidia"                
## [3738,] "Bacteria" "Firmicutes"          "Clostridia"                 
## [3739,] "Bacteria" "Bacteroidetes"       "Bacteroidia"                
## [3740,] "Bacteria" "Bacteroidetes"       "Bacteroidia"                
## [3741,] "Bacteria" "Bacteroidetes"       "Bacteroidia"                
## [3742,] "Bacteria" "Proteobacteria"      "Gammaproteobacteria"        
## [3743,] "Bacteria" "Epsilonbacteraeota"  "Campylobacteria"            
## [3744,] "Bacteria" "Firmicutes"          "Bacilli"                    
## [3745,] "Bacteria" "Patescibacteria"     "Saccharimonadia"            
## [3746,] "Bacteria" "Bacteroidetes"       "Bacteroidia"                
## [3747,] "Bacteria" "Bacteroidetes"       "Bacteroidia"                
## [3748,] "Bacteria" "Bacteroidetes"       "Bacteroidia"                
## [3749,] "Bacteria" "Bacteroidetes"       "Bacteroidia"                
## [3750,] "Bacteria" "Epsilonbacteraeota"  "Campylobacteria"            
## [3751,] "Bacteria" "Firmicutes"          "Clostridia"                 
## [3752,] "Bacteria" "Firmicutes"          "Bacilli"                    
## [3753,] "Bacteria" "Firmicutes"          "Negativicutes"              
## [3754,] "Bacteria" "Bacteroidetes"       "Bacteroidia"                
## [3755,] "Bacteria" "Firmicutes"          "Clostridia"                 
## [3756,] "Bacteria" "Firmicutes"          "Clostridia"                 
## [3757,] "Bacteria" "Bacteroidetes"       "Bacteroidia"                
## [3758,] "Bacteria" "Firmicutes"          "Bacilli"                    
## [3759,] "Bacteria" "Firmicutes"          "Negativicutes"              
## [3760,] "Bacteria" "Patescibacteria"     "Saccharimonadia"            
## [3761,] "Bacteria" "Bacteroidetes"       "Bacteroidia"                
## [3762,] "Bacteria" "Firmicutes"          "Bacilli"                    
## [3763,] "Bacteria" "Spirochaetes"        "Spirochaetia"               
## [3764,] "Bacteria" "Bacteroidetes"       "Bacteroidia"                
## [3765,] "Bacteria" "Bacteroidetes"       "Bacteroidia"                
## [3766,] "Bacteria" "Bacteroidetes"       "Bacteroidia"                
## [3767,] "Bacteria" "Proteobacteria"      "Gammaproteobacteria"        
## [3768,] "Bacteria" "Firmicutes"          "Negativicutes"              
## [3769,] "Bacteria" "Actinobacteria"      "Actinobacteria"             
## [3770,] "Bacteria" "Proteobacteria"      "Gammaproteobacteria"        
## [3771,] "Bacteria" "Bacteroidetes"       "Bacteroidia"                
## [3772,] "Bacteria" "Patescibacteria"     "Saccharimonadia"            
## [3773,] "Bacteria" "Firmicutes"          "Clostridia"                 
## [3774,] "Bacteria" "Firmicutes"          "Clostridia"                 
## [3775,] "Bacteria" "Firmicutes"          "Negativicutes"              
## [3776,] "Bacteria" "Bacteroidetes"       "Bacteroidia"                
## [3777,] "Bacteria" "Bacteroidetes"       "Bacteroidia"                
## [3778,] "Bacteria" "Bacteroidetes"       "Bacteroidia"                
## [3779,] "Bacteria" "Bacteroidetes"       "Bacteroidia"                
## [3780,] "Bacteria" "Firmicutes"          "Clostridia"                 
## [3781,] "Bacteria" "Bacteroidetes"       "Bacteroidia"                
## [3782,] "Bacteria" "Bacteroidetes"       "Bacteroidia"                
## [3783,] "Bacteria" "Proteobacteria"      "Gammaproteobacteria"        
## [3784,] "Bacteria" "Bacteroidetes"       "Bacteroidia"                
## [3785,] "Bacteria" "Fusobacteria"        "Fusobacteriia"              
## [3786,] "Bacteria" "Bacteroidetes"       "Bacteroidia"                
## [3787,] "Bacteria" NA                    NA                           
## [3788,] "Bacteria" "Patescibacteria"     "Saccharimonadia"            
## [3789,] "Bacteria" "Proteobacteria"      "Gammaproteobacteria"        
## [3790,] "Bacteria" "Firmicutes"          "Negativicutes"              
## [3791,] "Bacteria" "Spirochaetes"        "Spirochaetia"               
## [3792,] "Bacteria" "Patescibacteria"     "Saccharimonadia"            
## [3793,] "Bacteria" "Bacteroidetes"       "Bacteroidia"                
## [3794,] "Bacteria" "Bacteroidetes"       "Bacteroidia"                
## [3795,] "Bacteria" "Firmicutes"          "Negativicutes"              
## [3796,] "Bacteria" "Patescibacteria"     "Saccharimonadia"            
## [3797,] "Bacteria" "Patescibacteria"     "Gracilibacteria"            
## [3798,] "Bacteria" "Firmicutes"          "Bacilli"                    
## [3799,] "Bacteria" "Bacteroidetes"       "Bacteroidia"                
## [3800,] "Bacteria" "Proteobacteria"      "Gammaproteobacteria"        
## [3801,] "Bacteria" "Firmicutes"          "Negativicutes"              
## [3802,] "Bacteria" "Bacteroidetes"       "Bacteroidia"                
## [3803,] "Bacteria" "Proteobacteria"      "Gammaproteobacteria"        
## [3804,] "Bacteria" "Firmicutes"          "Clostridia"                 
## [3805,] "Bacteria" "Firmicutes"          "Bacilli"                    
## [3806,] "Bacteria" "Patescibacteria"     "Saccharimonadia"            
## [3807,] "Bacteria" "Firmicutes"          "Clostridia"                 
## [3808,] "Bacteria" "Bacteroidetes"       "Bacteroidia"                
## [3809,] "Bacteria" "Actinobacteria"      "Actinobacteria"             
## [3810,] "Bacteria" "Actinobacteria"      "Actinobacteria"             
## [3811,] "Bacteria" "Firmicutes"          "Negativicutes"              
## [3812,] "Bacteria" "Firmicutes"          "Negativicutes"              
## [3813,] "Bacteria" "Firmicutes"          "Bacilli"                    
## [3814,] "Bacteria" "Firmicutes"          "Bacilli"                    
## [3815,] "Bacteria" "Bacteroidetes"       "Bacteroidia"                
## [3816,] "Bacteria" "Bacteroidetes"       "Bacteroidia"                
## [3817,] "Bacteria" "Firmicutes"          "Negativicutes"              
## [3818,] "Bacteria" "Firmicutes"          "Negativicutes"              
## [3819,] "Bacteria" "Bacteroidetes"       "Bacteroidia"                
## [3820,] "Bacteria" "Bacteroidetes"       "Bacteroidia"                
## [3821,] "Bacteria" "Patescibacteria"     "Gracilibacteria"            
## [3822,] "Bacteria" "Spirochaetes"        "Spirochaetia"               
## [3823,] "Bacteria" "Bacteroidetes"       "Bacteroidia"                
## [3824,] "Bacteria" "Firmicutes"          "Negativicutes"              
## [3825,] "Bacteria" "Spirochaetes"        "Spirochaetia"               
## [3826,] "Bacteria" "Firmicutes"          "Clostridia"                 
## [3827,] "Bacteria" "Patescibacteria"     "Saccharimonadia"            
## [3828,] "Bacteria" "Proteobacteria"      "Gammaproteobacteria"        
## [3829,] "Bacteria" "Firmicutes"          "Bacilli"                    
## [3830,] "Bacteria" "Patescibacteria"     "Saccharimonadia"            
## [3831,] "Bacteria" "Bacteroidetes"       "Bacteroidia"                
## [3832,] "Bacteria" "Proteobacteria"      "Gammaproteobacteria"        
## [3833,] "Bacteria" "Fusobacteria"        "Fusobacteriia"              
## [3834,] "Bacteria" "Actinobacteria"      "Actinobacteria"             
## [3835,] "Bacteria" "Bacteroidetes"       "Bacteroidia"                
## [3836,] "Bacteria" "Bacteroidetes"       "Bacteroidia"                
## [3837,] "Bacteria" "Firmicutes"          "Bacilli"                    
## [3838,] "Bacteria" "Bacteroidetes"       "Bacteroidia"                
## [3839,] "Bacteria" "Bacteroidetes"       "Bacteroidia"                
## [3840,] "Bacteria" "Firmicutes"          "Clostridia"                 
## [3841,] "Bacteria" "Bacteroidetes"       "Bacteroidia"                
## [3842,] "Bacteria" "Bacteroidetes"       "Bacteroidia"                
## [3843,] "Bacteria" "Bacteroidetes"       "Bacteroidia"                
## [3844,] "Bacteria" "Bacteroidetes"       "Bacteroidia"                
## [3845,] "Bacteria" "Firmicutes"          NA                           
## [3846,] "Bacteria" "Epsilonbacteraeota"  "Campylobacteria"            
## [3847,] "Bacteria" "Bacteroidetes"       "Bacteroidia"                
## [3848,] "Bacteria" "Firmicutes"          "Clostridia"                 
## [3849,] "Bacteria" "Spirochaetes"        "Spirochaetia"               
## [3850,] "Bacteria" "Firmicutes"          "Clostridia"                 
## [3851,] "Bacteria" "Proteobacteria"      "Gammaproteobacteria"        
## [3852,] "Bacteria" "Proteobacteria"      "Gammaproteobacteria"        
## [3853,] "Bacteria" "Bacteroidetes"       "Bacteroidia"                
## [3854,] "Bacteria" "Firmicutes"          "Bacilli"                    
## [3855,] "Bacteria" "Epsilonbacteraeota"  "Campylobacteria"            
## [3856,] "Bacteria" "Firmicutes"          "Bacilli"                    
## [3857,] "Bacteria" "Fusobacteria"        "Fusobacteriia"              
## [3858,] "Bacteria" "Bacteroidetes"       "Bacteroidia"                
## [3859,] "Bacteria" "Bacteroidetes"       "Bacteroidia"                
## [3860,] "Bacteria" "Bacteroidetes"       "Bacteroidia"                
## [3861,] "Bacteria" "Firmicutes"          "Bacilli"                    
## [3862,] "Bacteria" "Firmicutes"          "Clostridia"                 
## [3863,] "Bacteria" "Firmicutes"          "Clostridia"                 
## [3864,] "Bacteria" "Bacteroidetes"       "Bacteroidia"                
## [3865,] "Bacteria" "Fusobacteria"        "Fusobacteriia"              
## [3866,] "Bacteria" "Firmicutes"          "Clostridia"                 
## [3867,] "Bacteria" "Fusobacteria"        "Fusobacteriia"              
## [3868,] "Bacteria" "Bacteroidetes"       "Bacteroidia"                
## [3869,] "Bacteria" "Patescibacteria"     "Saccharimonadia"            
## [3870,] "Bacteria" "Bacteroidetes"       "Bacteroidia"                
## [3871,] "Bacteria" "Proteobacteria"      "Gammaproteobacteria"        
## [3872,] "Bacteria" "Proteobacteria"      "Gammaproteobacteria"        
## [3873,] "Bacteria" "Firmicutes"          "Bacilli"                    
## [3874,] "Bacteria" "Bacteroidetes"       "Bacteroidia"                
## [3875,] "Bacteria" "Firmicutes"          NA                           
## [3876,] "Bacteria" "Proteobacteria"      "Gammaproteobacteria"        
## [3877,] "Bacteria" "Bacteroidetes"       "Bacteroidia"                
## [3878,] "Bacteria" "Epsilonbacteraeota"  "Campylobacteria"            
## [3879,] "Bacteria" "Bacteroidetes"       "Bacteroidia"                
## [3880,] "Bacteria" "Firmicutes"          "Clostridia"                 
## [3881,] "Bacteria" "Spirochaetes"        "Spirochaetia"               
## [3882,] "Bacteria" "Bacteroidetes"       "Bacteroidia"                
## [3883,] "Bacteria" "Firmicutes"          "Bacilli"                    
## [3884,] "Bacteria" "Firmicutes"          "Negativicutes"              
## [3885,] "Bacteria" "Proteobacteria"      "Gammaproteobacteria"        
## [3886,] "Bacteria" "Firmicutes"          "Clostridia"                 
## [3887,] "Bacteria" "Firmicutes"          "Clostridia"                 
## [3888,] "Bacteria" "Firmicutes"          "Clostridia"                 
## [3889,] "Bacteria" "Firmicutes"          "Bacilli"                    
## [3890,] "Bacteria" "Firmicutes"          "Clostridia"                 
## [3891,] "Bacteria" "Patescibacteria"     "Saccharimonadia"            
## [3892,] "Bacteria" "Firmicutes"          NA                           
## [3893,] "Bacteria" "Firmicutes"          "Clostridia"                 
## [3894,] "Bacteria" "Firmicutes"          "Clostridia"                 
## [3895,] "Bacteria" "Actinobacteria"      "Actinobacteria"             
## [3896,] "Bacteria" "Bacteroidetes"       "Bacteroidia"                
## [3897,] "Bacteria" "Epsilonbacteraeota"  "Campylobacteria"            
## [3898,] "Bacteria" "Proteobacteria"      "Gammaproteobacteria"        
## [3899,] "Bacteria" "Proteobacteria"      "Gammaproteobacteria"        
## [3900,] "Bacteria" "Firmicutes"          "Bacilli"                    
## [3901,] "Bacteria" "Firmicutes"          "Bacilli"                    
## [3902,] "Bacteria" "Firmicutes"          "Negativicutes"              
## [3903,] "Bacteria" "Bacteroidetes"       "Bacteroidia"                
## [3904,] "Bacteria" "Bacteroidetes"       "Bacteroidia"                
## [3905,] "Bacteria" "Spirochaetes"        "Spirochaetia"               
## [3906,] "Bacteria" "Firmicutes"          "Clostridia"                 
## [3907,] "Bacteria" "Bacteroidetes"       "Bacteroidia"                
## [3908,] "Bacteria" "Proteobacteria"      "Gammaproteobacteria"        
## [3909,] "Bacteria" "Patescibacteria"     "Gracilibacteria"            
## [3910,] "Bacteria" "Bacteroidetes"       "Bacteroidia"                
## [3911,] "Bacteria" "Bacteroidetes"       "Bacteroidia"                
## [3912,] "Bacteria" "Epsilonbacteraeota"  "Campylobacteria"            
## [3913,] "Bacteria" "Firmicutes"          "Negativicutes"              
## [3914,] "Bacteria" "Firmicutes"          "Erysipelotrichia"           
## [3915,] "Bacteria" "Firmicutes"          "Bacilli"                    
## [3916,] "Bacteria" "Firmicutes"          "Bacilli"                    
## [3917,] "Bacteria" "Bacteroidetes"       "Bacteroidia"                
## [3918,] "Bacteria" "Fusobacteria"        "Fusobacteriia"              
## [3919,] "Bacteria" "Proteobacteria"      "Gammaproteobacteria"        
## [3920,] "Bacteria" "Firmicutes"          "Negativicutes"              
## [3921,] "Bacteria" "Actinobacteria"      "Actinobacteria"             
## [3922,] "Bacteria" "Bacteroidetes"       "Bacteroidia"                
## [3923,] "Bacteria" "Firmicutes"          NA                           
## [3924,] "Bacteria" "Proteobacteria"      "Gammaproteobacteria"        
## [3925,] "Bacteria" "Actinobacteria"      "Actinobacteria"             
## [3926,] "Bacteria" "Firmicutes"          "Negativicutes"              
## [3927,] "Bacteria" "Fusobacteria"        "Fusobacteriia"              
## [3928,] "Bacteria" "Proteobacteria"      "Gammaproteobacteria"        
## [3929,] "Bacteria" "Deinococcus-Thermus" "Deinococci"                 
## [3930,] "Bacteria" "Epsilonbacteraeota"  "Campylobacteria"            
## [3931,] "Bacteria" "Firmicutes"          "Clostridia"                 
## [3932,] "Bacteria" "Firmicutes"          "Clostridia"                 
## [3933,] "Bacteria" NA                    NA                           
## [3934,] "Bacteria" "Firmicutes"          "Clostridia"                 
## [3935,] "Bacteria" "Proteobacteria"      "Gammaproteobacteria"        
## [3936,] "Bacteria" "Spirochaetes"        "Spirochaetia"               
## [3937,] "Bacteria" "Firmicutes"          "Bacilli"                    
## [3938,] "Bacteria" "Proteobacteria"      "Gammaproteobacteria"        
## [3939,] "Bacteria" "Firmicutes"          "Bacilli"                    
## [3940,] "Bacteria" "Firmicutes"          "Clostridia"                 
## [3941,] "Bacteria" "Bacteroidetes"       "Bacteroidia"                
## [3942,] "Bacteria" "Bacteroidetes"       "Bacteroidia"                
## [3943,] "Bacteria" "Proteobacteria"      "Gammaproteobacteria"        
## [3944,] "Bacteria" "Firmicutes"          "Clostridia"                 
## [3945,] "Bacteria" "Bacteroidetes"       "Bacteroidia"                
## [3946,] "Bacteria" "Firmicutes"          "Bacilli"                    
## [3947,] "Bacteria" "Firmicutes"          "Negativicutes"              
## [3948,] "Bacteria" "Firmicutes"          "Bacilli"                    
## [3949,] "Bacteria" "Bacteroidetes"       "Bacteroidia"                
## [3950,] "Bacteria" "Firmicutes"          "Bacilli"                    
## [3951,] "Bacteria" "Firmicutes"          "Bacilli"                    
## [3952,] "Bacteria" "Bacteroidetes"       "Bacteroidia"                
## [3953,] "Bacteria" "Fusobacteria"        "Fusobacteriia"              
## [3954,] "Bacteria" "Proteobacteria"      "Gammaproteobacteria"        
## [3955,] "Bacteria" "Epsilonbacteraeota"  "Campylobacteria"            
## [3956,] "Bacteria" NA                    NA                           
## [3957,] "Bacteria" "Epsilonbacteraeota"  "Campylobacteria"            
## [3958,] "Bacteria" "Patescibacteria"     "Saccharimonadia"            
## [3959,] "Bacteria" "Bacteroidetes"       "Bacteroidia"                
## [3960,] "Bacteria" "Firmicutes"          "Negativicutes"              
## [3961,] "Bacteria" "Fusobacteria"        "Fusobacteriia"              
## [3962,] "Bacteria" "Patescibacteria"     "Saccharimonadia"            
## [3963,] "Bacteria" "Bacteroidetes"       "Bacteroidia"                
## [3964,] "Bacteria" "Bacteroidetes"       "Bacteroidia"                
## [3965,] "Bacteria" "Bacteroidetes"       "Bacteroidia"                
## [3966,] "Bacteria" "Firmicutes"          "Bacilli"                    
## [3967,] "Bacteria" NA                    NA                           
## [3968,] "Bacteria" "Fusobacteria"        "Fusobacteriia"              
## [3969,] "Bacteria" "Fusobacteria"        "Fusobacteriia"              
## [3970,] "Bacteria" "Epsilonbacteraeota"  "Campylobacteria"            
## [3971,] "Bacteria" "Firmicutes"          NA                           
## [3972,] "Bacteria" "Tenericutes"         "Mollicutes"                 
## [3973,] "Bacteria" "Tenericutes"         "Mollicutes"                 
## [3974,] "Bacteria" "Bacteroidetes"       "Bacteroidia"                
## [3975,] "Bacteria" "Proteobacteria"      "Deltaproteobacteria"        
## [3976,] "Bacteria" "Patescibacteria"     "Gracilibacteria"            
## [3977,] "Bacteria" "Bacteroidetes"       "Bacteroidia"                
## [3978,] "Bacteria" "Bacteroidetes"       "Bacteroidia"                
## [3979,] "Bacteria" "Firmicutes"          "Bacilli"                    
## [3980,] "Bacteria" "Fusobacteria"        "Fusobacteriia"              
## [3981,] "Bacteria" "Firmicutes"          "Clostridia"                 
## [3982,] "Bacteria" "Firmicutes"          "Bacilli"                    
## [3983,] "Bacteria" "Bacteroidetes"       "Bacteroidia"                
## [3984,] "Bacteria" "Firmicutes"          "Negativicutes"              
## [3985,] "Bacteria" "Firmicutes"          "Clostridia"                 
## [3986,] "Bacteria" "Bacteroidetes"       "Bacteroidia"                
## [3987,] "Bacteria" "Bacteroidetes"       "Bacteroidia"                
## [3988,] "Bacteria" "Bacteroidetes"       "Bacteroidia"                
## [3989,] "Bacteria" "Actinobacteria"      "Coriobacteriia"             
## [3990,] "Bacteria" "Firmicutes"          "Negativicutes"              
## [3991,] "Bacteria" "Fusobacteria"        "Fusobacteriia"              
## [3992,] "Bacteria" "Epsilonbacteraeota"  "Campylobacteria"            
## [3993,] "Bacteria" "Bacteroidetes"       "Bacteroidia"                
## [3994,] "Bacteria" "Firmicutes"          "Bacilli"                    
## [3995,] "Bacteria" "Spirochaetes"        "Spirochaetia"               
## [3996,] "Bacteria" "Bacteroidetes"       "Bacteroidia"                
## [3997,] "Bacteria" "Firmicutes"          "Clostridia"                 
## [3998,] "Bacteria" "Bacteroidetes"       "Bacteroidia"                
## [3999,] "Bacteria" "Patescibacteria"     "Saccharimonadia"            
## [4000,] "Bacteria" "Firmicutes"          NA                           
## [4001,] "Bacteria" "Firmicutes"          "Bacilli"                    
## [4002,] "Bacteria" "Spirochaetes"        "Spirochaetia"               
## [4003,] "Bacteria" "Firmicutes"          "Bacilli"                    
## [4004,] "Bacteria" "Bacteroidetes"       "Bacteroidia"                
## [4005,] "Bacteria" "Fusobacteria"        "Fusobacteriia"              
## [4006,] "Bacteria" "Bacteroidetes"       "Bacteroidia"                
## [4007,] "Bacteria" "Firmicutes"          "Negativicutes"              
## [4008,] "Bacteria" "Bacteroidetes"       "Bacteroidia"                
## [4009,] "Bacteria" "Bacteroidetes"       "Bacteroidia"                
## [4010,] "Bacteria" "Proteobacteria"      "Gammaproteobacteria"        
## [4011,] "Bacteria" "Firmicutes"          "Negativicutes"              
## [4012,] "Bacteria" "Bacteroidetes"       "Bacteroidia"                
## [4013,] "Bacteria" "Firmicutes"          "Clostridia"                 
## [4014,] "Bacteria" "Firmicutes"          "Negativicutes"              
## [4015,] "Bacteria" "Actinobacteria"      "Actinobacteria"             
## [4016,] "Bacteria" "Firmicutes"          "Bacilli"                    
## [4017,] "Bacteria" "Bacteroidetes"       "Bacteroidia"                
## [4018,] "Bacteria" "Firmicutes"          "Clostridia"                 
## [4019,] "Bacteria" "Firmicutes"          "Clostridia"                 
## [4020,] "Bacteria" "Actinobacteria"      "Actinobacteria"             
## [4021,] "Bacteria" "Bacteroidetes"       "Bacteroidia"                
## [4022,] "Bacteria" "Actinobacteria"      "Actinobacteria"             
## [4023,] "Bacteria" "Firmicutes"          "Negativicutes"              
## [4024,] "Bacteria" "Patescibacteria"     "Saccharimonadia"            
## [4025,] "Bacteria" "Bacteroidetes"       "Bacteroidia"                
## [4026,] "Bacteria" "Bacteroidetes"       "Bacteroidia"                
## [4027,] "Bacteria" "Firmicutes"          "Bacilli"                    
## [4028,] "Bacteria" "Firmicutes"          "Negativicutes"              
## [4029,] "Bacteria" "Firmicutes"          "Bacilli"                    
## [4030,] "Bacteria" "Patescibacteria"     "Saccharimonadia"            
## [4031,] "Bacteria" "Proteobacteria"      "Gammaproteobacteria"        
## [4032,] "Bacteria" "Actinobacteria"      "Actinobacteria"             
## [4033,] "Bacteria" "Spirochaetes"        "Spirochaetia"               
## [4034,] "Bacteria" "Firmicutes"          "Bacilli"                    
## [4035,] "Bacteria" "Fusobacteria"        "Fusobacteriia"              
## [4036,] "Bacteria" "Firmicutes"          "Clostridia"                 
## [4037,] "Bacteria" "Patescibacteria"     "Gracilibacteria"            
## [4038,] "Bacteria" "Firmicutes"          "Negativicutes"              
## [4039,] "Bacteria" "Proteobacteria"      "Gammaproteobacteria"        
## [4040,] "Bacteria" "Firmicutes"          "Bacilli"                    
## [4041,] "Bacteria" "Bacteroidetes"       "Bacteroidia"                
## [4042,] "Bacteria" "Fusobacteria"        "Fusobacteriia"              
## [4043,] "Bacteria" "Bacteroidetes"       "Bacteroidia"                
## [4044,] "Bacteria" "Bacteroidetes"       "Bacteroidia"                
## [4045,] "Bacteria" "Firmicutes"          "Clostridia"                 
## [4046,] "Bacteria" "Patescibacteria"     "Saccharimonadia"            
## [4047,] "Bacteria" "Bacteroidetes"       "Bacteroidia"                
## [4048,] "Bacteria" NA                    NA                           
## [4049,] "Bacteria" "Fusobacteria"        "Fusobacteriia"              
## [4050,] "Bacteria" "Bacteroidetes"       "Bacteroidia"                
## [4051,] "Bacteria" "Bacteroidetes"       "Bacteroidia"                
## [4052,] "Bacteria" "Bacteroidetes"       "Bacteroidia"                
## [4053,] "Bacteria" "Firmicutes"          "Bacilli"                    
## [4054,] "Bacteria" "Bacteroidetes"       "Bacteroidia"                
## [4055,] "Bacteria" "Actinobacteria"      "Actinobacteria"             
## [4056,] "Bacteria" "Fusobacteria"        "Fusobacteriia"              
## [4057,] "Bacteria" "Proteobacteria"      "Gammaproteobacteria"        
## [4058,] "Bacteria" "Bacteroidetes"       "Bacteroidia"                
## [4059,] "Bacteria" "Firmicutes"          "Bacilli"                    
## [4060,] "Bacteria" "Fusobacteria"        "Fusobacteriia"              
## [4061,] "Bacteria" "Cyanobacteria"       "Oxyphotobacteria"           
## [4062,] "Bacteria" "Actinobacteria"      "Actinobacteria"             
## [4063,] "Bacteria" "Patescibacteria"     "Saccharimonadia"            
## [4064,] "Bacteria" "Fusobacteria"        "Fusobacteriia"              
## [4065,] "Bacteria" "Firmicutes"          NA                           
## [4066,] "Bacteria" "Bacteroidetes"       "Bacteroidia"                
## [4067,] "Bacteria" "Firmicutes"          "Clostridia"                 
## [4068,] "Bacteria" "Firmicutes"          "Clostridia"                 
## [4069,] "Bacteria" "Bacteroidetes"       "Bacteroidia"                
## [4070,] "Bacteria" "Actinobacteria"      "Actinobacteria"             
## [4071,] "Bacteria" "Firmicutes"          "Bacilli"                    
## [4072,] "Bacteria" "Patescibacteria"     "Gracilibacteria"            
## [4073,] "Bacteria" "Firmicutes"          "Negativicutes"              
## [4074,] "Bacteria" "Firmicutes"          "Negativicutes"              
## [4075,] "Bacteria" "Firmicutes"          "Negativicutes"              
## [4076,] "Bacteria" "Firmicutes"          "Clostridia"                 
## [4077,] "Bacteria" "Bacteroidetes"       "Bacteroidia"                
## [4078,] "Bacteria" "Proteobacteria"      "Gammaproteobacteria"        
## [4079,] "Bacteria" "Bacteroidetes"       "Bacteroidia"                
## [4080,] "Bacteria" "Bacteroidetes"       "Bacteroidia"                
## [4081,] "Bacteria" "Firmicutes"          "Negativicutes"              
## [4082,] "Bacteria" "Firmicutes"          "Negativicutes"              
## [4083,] "Bacteria" "Firmicutes"          "Clostridia"                 
## [4084,] "Bacteria" "Epsilonbacteraeota"  "Campylobacteria"            
## [4085,] "Bacteria" "Cyanobacteria"       "Oxyphotobacteria"           
## [4086,] "Bacteria" "Firmicutes"          "Bacilli"                    
## [4087,] "Bacteria" "Bacteroidetes"       "Bacteroidia"                
## [4088,] "Bacteria" "Proteobacteria"      "Gammaproteobacteria"        
## [4089,] "Bacteria" "Actinobacteria"      "Actinobacteria"             
## [4090,] "Bacteria" NA                    NA                           
## [4091,] "Bacteria" "Actinobacteria"      "Actinobacteria"             
## [4092,] "Bacteria" "Firmicutes"          "Bacilli"                    
## [4093,] "Bacteria" "Firmicutes"          "Clostridia"                 
## [4094,] "Bacteria" "Firmicutes"          "Clostridia"                 
## [4095,] "Bacteria" "Proteobacteria"      "Gammaproteobacteria"        
## [4096,] "Bacteria" "Firmicutes"          "Clostridia"                 
## [4097,] "Bacteria" "Patescibacteria"     "Saccharimonadia"            
## [4098,] "Bacteria" "Proteobacteria"      "Gammaproteobacteria"        
## [4099,] "Bacteria" "Bacteroidetes"       "Bacteroidia"                
## [4100,] "Bacteria" "Firmicutes"          "Clostridia"                 
## [4101,] "Bacteria" "Actinobacteria"      "Actinobacteria"             
## [4102,] "Bacteria" "Bacteroidetes"       "Bacteroidia"                
## [4103,] "Bacteria" "Firmicutes"          "Clostridia"                 
## [4104,] "Bacteria" "Firmicutes"          "Clostridia"                 
## [4105,] "Bacteria" "Bacteroidetes"       "Bacteroidia"                
## [4106,] "Bacteria" "Bacteroidetes"       "Bacteroidia"                
## [4107,] "Bacteria" "Patescibacteria"     "Saccharimonadia"            
## [4108,] "Bacteria" "Patescibacteria"     "Saccharimonadia"            
## [4109,] "Bacteria" "Bacteroidetes"       "Bacteroidia"                
## [4110,] "Bacteria" "Firmicutes"          "Negativicutes"              
## [4111,] "Bacteria" "Firmicutes"          "Bacilli"                    
## [4112,] "Bacteria" "Firmicutes"          "Negativicutes"              
## [4113,] "Bacteria" "Fusobacteria"        "Fusobacteriia"              
## [4114,] "Bacteria" "Actinobacteria"      "Actinobacteria"             
## [4115,] "Bacteria" "Actinobacteria"      "Actinobacteria"             
## [4116,] "Bacteria" "Firmicutes"          "Clostridia"                 
## [4117,] "Bacteria" "Actinobacteria"      "Coriobacteriia"             
## [4118,] "Bacteria" "Bacteroidetes"       "Bacteroidia"                
## [4119,] "Bacteria" "Actinobacteria"      "Coriobacteriia"             
## [4120,] "Bacteria" "Bacteroidetes"       "Bacteroidia"                
## [4121,] "Bacteria" "Elusimicrobia"       "Elusimicrobia"              
## [4122,] "Bacteria" "Epsilonbacteraeota"  "Campylobacteria"            
## [4123,] "Bacteria" "Proteobacteria"      NA                           
## [4124,] "Bacteria" "Epsilonbacteraeota"  "Campylobacteria"            
## [4125,] "Bacteria" "Bacteroidetes"       "Bacteroidia"                
## [4126,] "Bacteria" "Patescibacteria"     "Saccharimonadia"            
## [4127,] "Bacteria" "Firmicutes"          "Bacilli"                    
## [4128,] "Bacteria" "Proteobacteria"      "Gammaproteobacteria"        
## [4129,] "Bacteria" "Bacteroidetes"       "Bacteroidia"                
## [4130,] "Bacteria" "Firmicutes"          "Clostridia"                 
## [4131,] "Bacteria" "Firmicutes"          "Bacilli"                    
## [4132,] "Bacteria" "Firmicutes"          "Clostridia"                 
## [4133,] "Bacteria" "Bacteroidetes"       "Bacteroidia"                
## [4134,] "Bacteria" "Bacteroidetes"       "Bacteroidia"                
## [4135,] "Bacteria" "Patescibacteria"     "Saccharimonadia"            
## [4136,] "Bacteria" "Patescibacteria"     "Saccharimonadia"            
## [4137,] "Bacteria" "Proteobacteria"      "Gammaproteobacteria"        
## [4138,] "Bacteria" "Patescibacteria"     "Saccharimonadia"            
## [4139,] "Bacteria" "Proteobacteria"      "Gammaproteobacteria"        
## [4140,] "Bacteria" "Firmicutes"          "Clostridia"                 
## [4141,] "Bacteria" "Proteobacteria"      "Gammaproteobacteria"        
## [4142,] "Bacteria" "Firmicutes"          "Clostridia"                 
## [4143,] "Bacteria" "Firmicutes"          "Clostridia"                 
## [4144,] "Bacteria" "Bacteroidetes"       "Bacteroidia"                
## [4145,] "Bacteria" "Firmicutes"          "Bacilli"                    
## [4146,] "Bacteria" "Bacteroidetes"       "Bacteroidia"                
## [4147,] "Bacteria" "Fusobacteria"        "Fusobacteriia"              
## [4148,] "Bacteria" "Firmicutes"          "Negativicutes"              
## [4149,] "Bacteria" "Bacteroidetes"       "Bacteroidia"                
## [4150,] "Bacteria" "Firmicutes"          "Clostridia"                 
## [4151,] "Bacteria" "Firmicutes"          "Clostridia"                 
## [4152,] "Bacteria" "Spirochaetes"        "Spirochaetia"               
## [4153,] "Bacteria" "Proteobacteria"      "Gammaproteobacteria"        
## [4154,] "Bacteria" "Firmicutes"          "Negativicutes"              
## [4155,] "Bacteria" "Bacteroidetes"       "Bacteroidia"                
## [4156,] "Bacteria" "Actinobacteria"      "Actinobacteria"             
## [4157,] "Bacteria" "Firmicutes"          "Negativicutes"              
## [4158,] "Bacteria" "Firmicutes"          "Bacilli"                    
## [4159,] "Bacteria" "Proteobacteria"      "Gammaproteobacteria"        
## [4160,] "Bacteria" "Bacteroidetes"       "Bacteroidia"                
## [4161,] "Bacteria" "Firmicutes"          "Clostridia"                 
## [4162,] "Bacteria" "Firmicutes"          "Negativicutes"              
## [4163,] "Bacteria" "Firmicutes"          "Clostridia"                 
## [4164,] "Bacteria" "Patescibacteria"     "Saccharimonadia"            
## [4165,] "Bacteria" "Fusobacteria"        "Fusobacteriia"              
## [4166,] "Bacteria" "Proteobacteria"      "Gammaproteobacteria"        
## [4167,] "Bacteria" "Firmicutes"          "Bacilli"                    
## [4168,] "Bacteria" "Fusobacteria"        "Fusobacteriia"              
## [4169,] "Bacteria" "Tenericutes"         "Mollicutes"                 
## [4170,] "Bacteria" "Firmicutes"          "Negativicutes"              
## [4171,] "Bacteria" "Bacteroidetes"       "Bacteroidia"                
## [4172,] "Bacteria" "Cyanobacteria"       "Oxyphotobacteria"           
## [4173,] "Bacteria" "Bacteroidetes"       "Bacteroidia"                
## [4174,] "Bacteria" "Firmicutes"          "Clostridia"                 
## [4175,] "Bacteria" "Spirochaetes"        "Spirochaetia"               
## [4176,] "Bacteria" "Proteobacteria"      "Gammaproteobacteria"        
## [4177,] "Bacteria" "Firmicutes"          "Clostridia"                 
## [4178,] "Bacteria" "Proteobacteria"      "Gammaproteobacteria"        
## [4179,] "Bacteria" "Bacteroidetes"       "Bacteroidia"                
## [4180,] "Bacteria" "Patescibacteria"     "Saccharimonadia"            
## [4181,] "Bacteria" "Firmicutes"          "Bacilli"                    
## [4182,] "Bacteria" "Proteobacteria"      "Gammaproteobacteria"        
## [4183,] "Bacteria" "Tenericutes"         "Mollicutes"                 
## [4184,] "Bacteria" "Bacteroidetes"       "Bacteroidia"                
## [4185,] "Bacteria" "Patescibacteria"     "Gracilibacteria"            
## [4186,] "Bacteria" "Firmicutes"          "Clostridia"                 
## [4187,] "Bacteria" "Firmicutes"          NA                           
## [4188,] "Bacteria" "Firmicutes"          "Negativicutes"              
## [4189,] "Bacteria" NA                    NA                           
## [4190,] "Bacteria" "Spirochaetes"        "Spirochaetia"               
## [4191,] "Bacteria" "Proteobacteria"      "Gammaproteobacteria"        
## [4192,] "Bacteria" "Proteobacteria"      "Gammaproteobacteria"        
## [4193,] "Bacteria" "Firmicutes"          "Bacilli"                    
## [4194,] "Bacteria" "Epsilonbacteraeota"  "Campylobacteria"            
## [4195,] "Bacteria" "Bacteroidetes"       "Bacteroidia"                
## [4196,] "Bacteria" "Firmicutes"          "Clostridia"                 
## [4197,] "Bacteria" "Firmicutes"          "Clostridia"                 
## [4198,] "Bacteria" "Firmicutes"          "Negativicutes"              
## [4199,] "Bacteria" "Actinobacteria"      "Actinobacteria"             
## [4200,] "Bacteria" "Proteobacteria"      "Gammaproteobacteria"        
## [4201,] "Bacteria" "Firmicutes"          "Bacilli"                    
## [4202,] "Bacteria" "Fusobacteria"        "Fusobacteriia"              
## [4203,] "Bacteria" "Firmicutes"          "Clostridia"                 
## [4204,] "Bacteria" "Bacteroidetes"       "Bacteroidia"                
## [4205,] "Bacteria" "Firmicutes"          "Negativicutes"              
## [4206,] "Bacteria" "Bacteroidetes"       "Bacteroidia"                
## [4207,] "Bacteria" "Actinobacteria"      "Actinobacteria"             
## [4208,] "Bacteria" "Firmicutes"          "Clostridia"                 
## [4209,] "Bacteria" "Fusobacteria"        "Fusobacteriia"              
## [4210,] "Bacteria" "Spirochaetes"        "Spirochaetia"               
## [4211,] "Bacteria" "Patescibacteria"     "Saccharimonadia"            
## [4212,] "Bacteria" "Firmicutes"          NA                           
## [4213,] "Bacteria" "Epsilonbacteraeota"  "Campylobacteria"            
## [4214,] "Bacteria" "Bacteroidetes"       "Bacteroidia"                
## [4215,] "Bacteria" "Bacteroidetes"       "Bacteroidia"                
## [4216,] "Bacteria" "Firmicutes"          NA                           
## [4217,] "Bacteria" "Patescibacteria"     "Saccharimonadia"            
## [4218,] "Bacteria" "Bacteroidetes"       "Bacteroidia"                
## [4219,] "Bacteria" "Firmicutes"          "Negativicutes"              
## [4220,] "Bacteria" "Epsilonbacteraeota"  "Campylobacteria"            
## [4221,] "Bacteria" "Proteobacteria"      "Gammaproteobacteria"        
## [4222,] "Bacteria" "Fusobacteria"        "Fusobacteriia"              
## [4223,] "Bacteria" "Bacteroidetes"       "Bacteroidia"                
## [4224,] "Bacteria" "Bacteroidetes"       "Bacteroidia"                
## [4225,] "Bacteria" "Firmicutes"          "Clostridia"                 
## [4226,] "Bacteria" "Fusobacteria"        "Fusobacteriia"              
## [4227,] "Bacteria" "Bacteroidetes"       "Bacteroidia"                
## [4228,] "Bacteria" "Firmicutes"          "Negativicutes"              
## [4229,] "Bacteria" "Proteobacteria"      "Gammaproteobacteria"        
## [4230,] "Bacteria" "Firmicutes"          "Bacilli"                    
## [4231,] "Bacteria" "Bacteroidetes"       "Bacteroidia"                
## [4232,] "Bacteria" "Proteobacteria"      "Gammaproteobacteria"        
## [4233,] "Bacteria" "Patescibacteria"     "Gracilibacteria"            
## [4234,] "Bacteria" "Bacteroidetes"       "Bacteroidia"                
## [4235,] "Bacteria" "Firmicutes"          "Bacilli"                    
## [4236,] "Bacteria" "Patescibacteria"     "Saccharimonadia"            
## [4237,] "Bacteria" "Firmicutes"          "Bacilli"                    
## [4238,] "Bacteria" "Spirochaetes"        "Spirochaetia"               
## [4239,] "Bacteria" "Actinobacteria"      "Actinobacteria"             
## [4240,] "Bacteria" "Proteobacteria"      "Gammaproteobacteria"        
## [4241,] "Bacteria" "Epsilonbacteraeota"  "Campylobacteria"            
## [4242,] "Bacteria" "Cyanobacteria"       "Oxyphotobacteria"           
## [4243,] "Bacteria" "Patescibacteria"     "Saccharimonadia"            
## [4244,] "Bacteria" "Firmicutes"          "Bacilli"                    
## [4245,] "Bacteria" "Bacteroidetes"       "Bacteroidia"                
## [4246,] "Bacteria" "Actinobacteria"      "Actinobacteria"             
## [4247,] "Bacteria" "Spirochaetes"        "Spirochaetia"               
## [4248,] "Bacteria" "Firmicutes"          "Bacilli"                    
## [4249,] "Bacteria" "Bacteroidetes"       "Bacteroidia"                
## [4250,] "Bacteria" "Spirochaetes"        "Spirochaetia"               
## [4251,] "Bacteria" "Firmicutes"          "Clostridia"                 
## [4252,] "Bacteria" "Firmicutes"          NA                           
## [4253,] "Bacteria" "Proteobacteria"      "Deltaproteobacteria"        
## [4254,] "Bacteria" "Tenericutes"         "Mollicutes"                 
## [4255,] "Bacteria" "Firmicutes"          "Bacilli"                    
## [4256,] "Bacteria" "Bacteroidetes"       "Bacteroidia"                
## [4257,] "Bacteria" "Actinobacteria"      "Actinobacteria"             
## [4258,] "Bacteria" "Epsilonbacteraeota"  "Campylobacteria"            
## [4259,] "Bacteria" "Bacteroidetes"       "Bacteroidia"                
## [4260,] "Bacteria" "Bacteroidetes"       "Bacteroidia"                
## [4261,] "Bacteria" "Patescibacteria"     "Saccharimonadia"            
## [4262,] "Bacteria" "Firmicutes"          "Bacilli"                    
## [4263,] "Bacteria" NA                    NA                           
## [4264,] "Bacteria" "Bacteroidetes"       "Bacteroidia"                
## [4265,] "Bacteria" "Bacteroidetes"       "Bacteroidia"                
## [4266,] "Bacteria" "Patescibacteria"     "Saccharimonadia"            
## [4267,] "Bacteria" "Actinobacteria"      "Actinobacteria"             
## [4268,] "Bacteria" "Actinobacteria"      "Coriobacteriia"             
## [4269,] "Bacteria" "Bacteroidetes"       "Bacteroidia"                
## [4270,] "Bacteria" "Firmicutes"          "Bacilli"                    
## [4271,] "Bacteria" "Firmicutes"          "Clostridia"                 
## [4272,] "Bacteria" "Firmicutes"          "Clostridia"                 
## [4273,] "Bacteria" "Actinobacteria"      "Actinobacteria"             
## [4274,] "Bacteria" "Proteobacteria"      "Gammaproteobacteria"        
## [4275,] "Bacteria" "Bacteroidetes"       "Bacteroidia"                
## [4276,] "Bacteria" "Firmicutes"          "Clostridia"                 
## [4277,] "Bacteria" "Firmicutes"          "Clostridia"                 
## [4278,] "Bacteria" "Firmicutes"          "Bacilli"                    
## [4279,] "Bacteria" "Firmicutes"          "Bacilli"                    
## [4280,] "Bacteria" "Bacteroidetes"       "Bacteroidia"                
## [4281,] "Bacteria" "Proteobacteria"      "Gammaproteobacteria"        
## [4282,] "Bacteria" "Fusobacteria"        "Fusobacteriia"              
## [4283,] "Bacteria" "Fusobacteria"        "Fusobacteriia"              
## [4284,] "Bacteria" "Proteobacteria"      "Gammaproteobacteria"        
## [4285,] "Bacteria" "Firmicutes"          "Erysipelotrichia"           
## [4286,] "Bacteria" "Firmicutes"          "Clostridia"                 
## [4287,] "Bacteria" "Bacteroidetes"       "Bacteroidia"                
## [4288,] "Bacteria" "Firmicutes"          "Bacilli"                    
## [4289,] "Bacteria" "Firmicutes"          NA                           
## [4290,] "Bacteria" "Firmicutes"          "Clostridia"                 
## [4291,] "Bacteria" "Firmicutes"          "Negativicutes"              
## [4292,] "Bacteria" "Bacteroidetes"       "Bacteroidia"                
## [4293,] "Bacteria" "Proteobacteria"      "Gammaproteobacteria"        
## [4294,] "Bacteria" "Bacteroidetes"       "Bacteroidia"                
## [4295,] "Bacteria" "Actinobacteria"      "Coriobacteriia"             
## [4296,] "Bacteria" "Bacteroidetes"       "Bacteroidia"                
## [4297,] "Bacteria" "Actinobacteria"      "Actinobacteria"             
## [4298,] "Bacteria" NA                    NA                           
## [4299,] "Bacteria" "Proteobacteria"      "Gammaproteobacteria"        
## [4300,] "Bacteria" "Firmicutes"          "Bacilli"                    
## [4301,] "Bacteria" "Firmicutes"          "Bacilli"                    
## [4302,] "Bacteria" "Spirochaetes"        "Spirochaetia"               
## [4303,] "Bacteria" "Bacteroidetes"       "Bacteroidia"                
## [4304,] "Bacteria" "Firmicutes"          "Negativicutes"              
## [4305,] "Bacteria" "Proteobacteria"      "Alphaproteobacteria"        
## [4306,] "Bacteria" "Firmicutes"          "Clostridia"                 
## [4307,] "Bacteria" "Spirochaetes"        "Spirochaetia"               
## [4308,] "Bacteria" "Firmicutes"          "Bacilli"                    
## [4309,] "Bacteria" "Proteobacteria"      "Gammaproteobacteria"        
## [4310,] "Bacteria" "Firmicutes"          "Negativicutes"              
## [4311,] "Bacteria" "Proteobacteria"      "Gammaproteobacteria"        
## [4312,] "Bacteria" "Firmicutes"          "Clostridia"                 
## [4313,] "Bacteria" "Patescibacteria"     "Saccharimonadia"            
## [4314,] "Bacteria" "Proteobacteria"      "Gammaproteobacteria"        
## [4315,] "Bacteria" "Bacteroidetes"       "Bacteroidia"                
## [4316,] "Bacteria" "Firmicutes"          "Clostridia"                 
## [4317,] "Bacteria" "Firmicutes"          NA                           
## [4318,] "Bacteria" "Actinobacteria"      "Actinobacteria"             
## [4319,] "Bacteria" "Spirochaetes"        "Spirochaetia"               
## [4320,] "Bacteria" "Firmicutes"          "Clostridia"                 
## [4321,] "Bacteria" "Firmicutes"          "Clostridia"                 
## [4322,] "Bacteria" "Proteobacteria"      "Gammaproteobacteria"        
## [4323,] "Bacteria" "Firmicutes"          "Clostridia"                 
## [4324,] "Bacteria" "Epsilonbacteraeota"  "Campylobacteria"            
## [4325,] "Bacteria" "Patescibacteria"     "Saccharimonadia"            
## [4326,] "Bacteria" "Proteobacteria"      "Gammaproteobacteria"        
## [4327,] "Bacteria" "Bacteroidetes"       "Bacteroidia"                
## [4328,] "Bacteria" "Firmicutes"          "Erysipelotrichia"           
## [4329,] "Bacteria" "Actinobacteria"      "Actinobacteria"             
## [4330,] "Bacteria" "Bacteroidetes"       "Bacteroidia"                
## [4331,] "Bacteria" "Firmicutes"          "Negativicutes"              
## [4332,] "Bacteria" "Proteobacteria"      "Gammaproteobacteria"        
## [4333,] "Bacteria" "Proteobacteria"      "Gammaproteobacteria"        
## [4334,] "Bacteria" "Bacteroidetes"       "Bacteroidia"                
## [4335,] "Bacteria" "Firmicutes"          "Bacilli"                    
## [4336,] "Bacteria" "Epsilonbacteraeota"  "Campylobacteria"            
## [4337,] "Bacteria" "Spirochaetes"        "Spirochaetia"               
## [4338,] "Bacteria" "Firmicutes"          "Clostridia"                 
## [4339,] "Bacteria" "Proteobacteria"      "Alphaproteobacteria"        
## [4340,] "Bacteria" "Proteobacteria"      "Gammaproteobacteria"        
## [4341,] "Bacteria" "Proteobacteria"      "Gammaproteobacteria"        
## [4342,] "Bacteria" "Firmicutes"          "Bacilli"                    
## [4343,] "Bacteria" "Bacteroidetes"       "Bacteroidia"                
## [4344,] "Bacteria" "Fusobacteria"        "Fusobacteriia"              
## [4345,] "Bacteria" "Bacteroidetes"       "Bacteroidia"                
## [4346,] "Bacteria" "Bacteroidetes"       "Bacteroidia"                
## [4347,] "Bacteria" "Firmicutes"          "Clostridia"                 
## [4348,] "Bacteria" "Bacteroidetes"       "Bacteroidia"                
## [4349,] "Bacteria" "Proteobacteria"      "Gammaproteobacteria"        
## [4350,] "Bacteria" "Bacteroidetes"       "Bacteroidia"                
## [4351,] "Bacteria" "Proteobacteria"      "Alphaproteobacteria"        
## [4352,] "Bacteria" "Firmicutes"          "Clostridia"                 
## [4353,] "Bacteria" "Bacteroidetes"       "Bacteroidia"                
## [4354,] "Bacteria" "Firmicutes"          "Bacilli"                    
## [4355,] "Bacteria" "Bacteroidetes"       "Bacteroidia"                
## [4356,] "Bacteria" "Firmicutes"          "Bacilli"                    
## [4357,] "Bacteria" "Proteobacteria"      "Gammaproteobacteria"        
## [4358,] "Bacteria" "Fusobacteria"        "Fusobacteriia"              
## [4359,] "Bacteria" "Proteobacteria"      "Gammaproteobacteria"        
## [4360,] "Bacteria" "Firmicutes"          "Clostridia"                 
## [4361,] "Bacteria" "Bacteroidetes"       "Bacteroidia"                
## [4362,] "Bacteria" "Fusobacteria"        "Fusobacteriia"              
## [4363,] "Bacteria" "Bacteroidetes"       "Bacteroidia"                
## [4364,] "Bacteria" "Firmicutes"          "Clostridia"                 
## [4365,] "Bacteria" "Proteobacteria"      "Gammaproteobacteria"        
## [4366,] "Bacteria" "Proteobacteria"      "Gammaproteobacteria"        
## [4367,] "Bacteria" "Firmicutes"          NA                           
## [4368,] "Bacteria" "Epsilonbacteraeota"  "Campylobacteria"            
## [4369,] "Bacteria" "Firmicutes"          "Clostridia"                 
## [4370,] "Bacteria" "Fusobacteria"        "Fusobacteriia"              
## [4371,] "Bacteria" "Proteobacteria"      "Gammaproteobacteria"        
## [4372,] "Bacteria" "Proteobacteria"      "Gammaproteobacteria"        
## [4373,] "Bacteria" "Firmicutes"          "Negativicutes"              
## [4374,] "Bacteria" "Fusobacteria"        "Fusobacteriia"              
## [4375,] "Bacteria" "Firmicutes"          "Negativicutes"              
## [4376,] "Bacteria" "Spirochaetes"        "Spirochaetia"               
## [4377,] "Bacteria" "Bacteroidetes"       "Bacteroidia"                
## [4378,] "Bacteria" "Firmicutes"          "Bacilli"                    
## [4379,] "Bacteria" "Firmicutes"          "Clostridia"                 
## [4380,] "Bacteria" "Proteobacteria"      "Gammaproteobacteria"        
## [4381,] "Bacteria" "Firmicutes"          NA                           
## [4382,] "Bacteria" "Bacteroidetes"       "Bacteroidia"                
## [4383,] "Bacteria" "Firmicutes"          "Bacilli"                    
## [4384,] "Bacteria" "Actinobacteria"      "Coriobacteriia"             
## [4385,] "Bacteria" "Firmicutes"          "Clostridia"                 
## [4386,] "Bacteria" "Bacteroidetes"       "Bacteroidia"                
## [4387,] "Bacteria" "Firmicutes"          "Negativicutes"              
## [4388,] "Bacteria" "Firmicutes"          "Clostridia"                 
## [4389,] "Bacteria" "Actinobacteria"      "Actinobacteria"             
## [4390,] "Bacteria" "Firmicutes"          "Negativicutes"              
## [4391,] "Bacteria" "Epsilonbacteraeota"  "Campylobacteria"            
## [4392,] "Bacteria" "Epsilonbacteraeota"  "Campylobacteria"            
## [4393,] "Bacteria" "Fusobacteria"        "Fusobacteriia"              
## [4394,] "Bacteria" "Firmicutes"          "Negativicutes"              
## [4395,] "Bacteria" "Firmicutes"          "Erysipelotrichia"           
## [4396,] "Bacteria" "Firmicutes"          "Negativicutes"              
## [4397,] "Bacteria" "Firmicutes"          "Negativicutes"              
## [4398,] "Bacteria" "Firmicutes"          "Negativicutes"              
## [4399,] "Bacteria" "Firmicutes"          "Bacilli"                    
## [4400,] "Bacteria" "Patescibacteria"     "Saccharimonadia"            
## [4401,] "Bacteria" "Firmicutes"          "Negativicutes"              
## [4402,] "Bacteria" "Fusobacteria"        "Fusobacteriia"              
## [4403,] "Bacteria" "Proteobacteria"      "Gammaproteobacteria"        
## [4404,] "Bacteria" "Patescibacteria"     "Saccharimonadia"            
## [4405,] "Bacteria" "Actinobacteria"      "Coriobacteriia"             
## [4406,] "Bacteria" "Bacteroidetes"       "Bacteroidia"                
## [4407,] "Bacteria" "Bacteroidetes"       "Bacteroidia"                
## [4408,] "Bacteria" "Proteobacteria"      "Alphaproteobacteria"        
## [4409,] "Bacteria" "Firmicutes"          "Bacilli"                    
## [4410,] "Bacteria" "Bacteroidetes"       "Bacteroidia"                
## [4411,] "Bacteria" "Firmicutes"          "Clostridia"                 
## [4412,] "Bacteria" "Firmicutes"          "Bacilli"                    
## [4413,] "Bacteria" "Actinobacteria"      "Coriobacteriia"             
## [4414,] "Bacteria" "Spirochaetes"        "Spirochaetia"               
## [4415,] "Bacteria" "Bacteroidetes"       "Bacteroidia"                
## [4416,] "Bacteria" "Epsilonbacteraeota"  "Campylobacteria"            
## [4417,] "Bacteria" "Synergistetes"       "Synergistia"                
## [4418,] "Bacteria" "Actinobacteria"      "Actinobacteria"             
## [4419,] "Bacteria" "Actinobacteria"      "Actinobacteria"             
## [4420,] "Bacteria" "Actinobacteria"      "Coriobacteriia"             
## [4421,] "Bacteria" "Firmicutes"          "Clostridia"                 
## [4422,] "Bacteria" "Tenericutes"         "Mollicutes"                 
## [4423,] "Bacteria" "Proteobacteria"      "Gammaproteobacteria"        
## [4424,] "Bacteria" "Firmicutes"          "Bacilli"                    
## [4425,] "Bacteria" "Epsilonbacteraeota"  "Campylobacteria"            
## [4426,] "Bacteria" "Actinobacteria"      "Actinobacteria"             
## [4427,] "Bacteria" "Firmicutes"          "Clostridia"                 
## [4428,] "Bacteria" "Firmicutes"          "Negativicutes"              
## [4429,] "Bacteria" "Bacteroidetes"       "Bacteroidia"                
## [4430,] "Bacteria" "Actinobacteria"      "Coriobacteriia"             
## [4431,] "Bacteria" "Bacteroidetes"       "Bacteroidia"                
## [4432,] "Bacteria" "Bacteroidetes"       "Bacteroidia"                
## [4433,] "Bacteria" "Patescibacteria"     "Saccharimonadia"            
## [4434,] "Bacteria" "Firmicutes"          "Clostridia"                 
## [4435,] "Bacteria" NA                    NA                           
## [4436,] "Bacteria" "Actinobacteria"      "Actinobacteria"             
## [4437,] "Bacteria" "Firmicutes"          NA                           
## [4438,] "Bacteria" "Patescibacteria"     "Saccharimonadia"            
## [4439,] "Bacteria" "Proteobacteria"      "Gammaproteobacteria"        
## [4440,] "Bacteria" "Firmicutes"          "Negativicutes"              
## [4441,] "Bacteria" "Firmicutes"          "Negativicutes"              
## [4442,] "Bacteria" "Proteobacteria"      "Gammaproteobacteria"        
## [4443,] "Bacteria" "Firmicutes"          "Erysipelotrichia"           
## [4444,] "Bacteria" "Proteobacteria"      "Gammaproteobacteria"        
## [4445,] "Bacteria" "Bacteroidetes"       "Bacteroidia"                
## [4446,] "Bacteria" "Spirochaetes"        "Spirochaetia"               
## [4447,] "Bacteria" "Bacteroidetes"       "Bacteroidia"                
## [4448,] "Bacteria" "Firmicutes"          "Bacilli"                    
## [4449,] "Bacteria" "Bacteroidetes"       "Bacteroidia"                
## [4450,] "Bacteria" "Bacteroidetes"       "Bacteroidia"                
## [4451,] "Bacteria" NA                    NA                           
## [4452,] "Bacteria" "Bacteroidetes"       "Bacteroidia"                
## [4453,] "Bacteria" "Actinobacteria"      "Actinobacteria"             
## [4454,] "Bacteria" "Proteobacteria"      "Gammaproteobacteria"        
## [4455,] "Bacteria" "Bacteroidetes"       "Bacteroidia"                
## [4456,] "Bacteria" "Fusobacteria"        "Fusobacteriia"              
## [4457,] "Bacteria" "Proteobacteria"      "Gammaproteobacteria"        
## [4458,] "Bacteria" "Firmicutes"          "Erysipelotrichia"           
## [4459,] "Bacteria" "Bacteroidetes"       "Bacteroidia"                
## [4460,] "Bacteria" "Proteobacteria"      "Gammaproteobacteria"        
## [4461,] "Bacteria" "Epsilonbacteraeota"  "Campylobacteria"            
## [4462,] "Bacteria" "Fusobacteria"        "Fusobacteriia"              
## [4463,] "Bacteria" "Firmicutes"          "Bacilli"                    
## [4464,] "Bacteria" "Bacteroidetes"       "Bacteroidia"                
## [4465,] "Bacteria" "Proteobacteria"      "Gammaproteobacteria"        
## [4466,] "Bacteria" "Actinobacteria"      "Coriobacteriia"             
## [4467,] "Bacteria" "Proteobacteria"      "Gammaproteobacteria"        
## [4468,] "Bacteria" "Firmicutes"          "Bacilli"                    
## [4469,] "Bacteria" NA                    NA                           
## [4470,] "Bacteria" "Bacteroidetes"       "Bacteroidia"                
## [4471,] "Bacteria" "Firmicutes"          "Clostridia"                 
## [4472,] "Bacteria" "Firmicutes"          "Clostridia"                 
## [4473,] "Bacteria" "Bacteroidetes"       "Bacteroidia"                
## [4474,] "Bacteria" "Bacteroidetes"       "Bacteroidia"                
## [4475,] "Bacteria" "Firmicutes"          "Clostridia"                 
## [4476,] "Bacteria" "Bacteroidetes"       "Bacteroidia"                
## [4477,] "Bacteria" "Firmicutes"          "Negativicutes"              
## [4478,] "Bacteria" "Bacteroidetes"       "Bacteroidia"                
## [4479,] "Bacteria" "Bacteroidetes"       "Bacteroidia"                
## [4480,] "Bacteria" "Firmicutes"          "Bacilli"                    
## [4481,] "Bacteria" "Proteobacteria"      "Gammaproteobacteria"        
## [4482,] "Bacteria" "Firmicutes"          "Clostridia"                 
## [4483,] "Bacteria" "Firmicutes"          "Bacilli"                    
## [4484,] "Bacteria" "Bacteroidetes"       "Bacteroidia"                
## [4485,] "Bacteria" "Firmicutes"          "Negativicutes"              
## [4486,] "Bacteria" "Firmicutes"          "Erysipelotrichia"           
## [4487,] "Bacteria" "Patescibacteria"     "Saccharimonadia"            
## [4488,] "Bacteria" "Patescibacteria"     "Saccharimonadia"            
## [4489,] "Bacteria" "Spirochaetes"        "Spirochaetia"               
## [4490,] "Bacteria" "Firmicutes"          "Clostridia"                 
## [4491,] "Bacteria" "Firmicutes"          "Clostridia"                 
## [4492,] "Bacteria" "Proteobacteria"      "Gammaproteobacteria"        
## [4493,] "Bacteria" "Firmicutes"          "Clostridia"                 
## [4494,] "Bacteria" "Proteobacteria"      "Gammaproteobacteria"        
## [4495,] "Bacteria" "Bacteroidetes"       "Bacteroidia"                
## [4496,] "Bacteria" "Proteobacteria"      "Gammaproteobacteria"        
## [4497,] "Bacteria" "Proteobacteria"      "Alphaproteobacteria"        
## [4498,] "Bacteria" "Firmicutes"          "Bacilli"                    
## [4499,] "Bacteria" "Bacteroidetes"       "Bacteroidia"                
## [4500,] "Bacteria" "Firmicutes"          "Negativicutes"              
## [4501,] "Bacteria" "Acidobacteria"       "Blastocatellia_(Subgroup_4)"
## [4502,] "Bacteria" "Firmicutes"          "Bacilli"                    
## [4503,] "Bacteria" "Proteobacteria"      "Gammaproteobacteria"        
## [4504,] "Bacteria" "Firmicutes"          "Clostridia"                 
## [4505,] "Bacteria" "Fusobacteria"        "Fusobacteriia"              
## [4506,] "Bacteria" "Bacteroidetes"       "Bacteroidia"                
## [4507,] "Bacteria" "Bacteroidetes"       "Bacteroidia"                
## [4508,] "Bacteria" "Firmicutes"          "Clostridia"                 
## [4509,] "Bacteria" "Patescibacteria"     "Saccharimonadia"            
## [4510,] "Bacteria" "Proteobacteria"      "Gammaproteobacteria"        
## [4511,] "Bacteria" "Bacteroidetes"       "Bacteroidia"                
## [4512,] "Bacteria" "Bacteroidetes"       "Bacteroidia"                
## [4513,] "Bacteria" "Actinobacteria"      "Actinobacteria"             
## [4514,] "Bacteria" "Firmicutes"          "Bacilli"                    
## [4515,] "Bacteria" "Fusobacteria"        "Fusobacteriia"              
## [4516,] "Bacteria" "Patescibacteria"     "Saccharimonadia"            
## [4517,] "Bacteria" "Fusobacteria"        "Fusobacteriia"              
## [4518,] "Bacteria" "Firmicutes"          "Negativicutes"              
## [4519,] "Bacteria" "Bacteroidetes"       "Bacteroidia"                
## [4520,] "Bacteria" "Firmicutes"          "Bacilli"                    
## [4521,] "Bacteria" "Proteobacteria"      "Gammaproteobacteria"        
## [4522,] "Bacteria" "Bacteroidetes"       "Bacteroidia"                
## [4523,] "Bacteria" "Bacteroidetes"       "Bacteroidia"                
## [4524,] "Bacteria" "Proteobacteria"      "Gammaproteobacteria"        
## [4525,] "Bacteria" "Firmicutes"          "Clostridia"                 
## [4526,] "Bacteria" "Bacteroidetes"       "Bacteroidia"                
## [4527,] "Bacteria" "Patescibacteria"     "Saccharimonadia"            
## [4528,] "Bacteria" "Bacteroidetes"       "Bacteroidia"                
## [4529,] "Bacteria" "Firmicutes"          "Negativicutes"              
## [4530,] "Bacteria" "Bacteroidetes"       "Bacteroidia"                
## [4531,] "Bacteria" "Bacteroidetes"       "Bacteroidia"                
## [4532,] "Bacteria" "Bacteroidetes"       "Bacteroidia"                
## [4533,] "Bacteria" NA                    NA                           
## [4534,] "Bacteria" "Proteobacteria"      "Gammaproteobacteria"        
## [4535,] "Bacteria" "Bacteroidetes"       "Bacteroidia"                
## [4536,] "Bacteria" "Actinobacteria"      "Actinobacteria"             
## [4537,] "Bacteria" "Actinobacteria"      "Coriobacteriia"             
## [4538,] "Bacteria" "Firmicutes"          "Bacilli"                    
## [4539,] "Bacteria" "Bacteroidetes"       "Bacteroidia"                
## [4540,] "Bacteria" "Bacteroidetes"       "Bacteroidia"                
## [4541,] "Bacteria" "Firmicutes"          "Clostridia"                 
## [4542,] "Bacteria" "Firmicutes"          "Clostridia"                 
## [4543,] "Bacteria" NA                    NA                           
## [4544,] "Bacteria" "Bacteroidetes"       "Bacteroidia"                
## [4545,] "Bacteria" "Epsilonbacteraeota"  "Campylobacteria"            
## [4546,] "Bacteria" "Spirochaetes"        "Spirochaetia"               
## [4547,] "Bacteria" "Proteobacteria"      "Gammaproteobacteria"        
## [4548,] "Bacteria" "Bacteroidetes"       "Bacteroidia"                
## [4549,] "Bacteria" "Firmicutes"          "Negativicutes"              
## [4550,] "Bacteria" "Firmicutes"          "Bacilli"                    
## [4551,] "Bacteria" "Fusobacteria"        "Fusobacteriia"              
## [4552,] "Bacteria" "Firmicutes"          "Bacilli"                    
## [4553,] "Bacteria" "Spirochaetes"        "Spirochaetia"               
## [4554,] "Bacteria" "Bacteroidetes"       "Bacteroidia"                
## [4555,] "Bacteria" "Firmicutes"          "Clostridia"                 
##         [,4]                          [,5]                           
##    [1,] "Selenomonadales"             "Veillonellaceae"              
##    [2,] "Lactobacillales"             "Streptococcaceae"             
##    [3,] "Bacteroidales"               "Prevotellaceae"               
##    [4,] "Pasteurellales"              "Pasteurellaceae"              
##    [5,] "Selenomonadales"             "Veillonellaceae"              
##    [6,] "Selenomonadales"             "Veillonellaceae"              
##    [7,] "Selenomonadales"             "Veillonellaceae"              
##    [8,] "Saccharimonadales"           "Saccharimonadaceae"           
##    [9,] "Betaproteobacteriales"       "Neisseriaceae"                
##   [10,] "Selenomonadales"             "Veillonellaceae"              
##   [11,] "Betaproteobacteriales"       "Neisseriaceae"                
##   [12,] "Selenomonadales"             "Veillonellaceae"              
##   [13,] "Fusobacteriales"             "Fusobacteriaceae"             
##   [14,] "Selenomonadales"             "Veillonellaceae"              
##   [15,] "Bacteroidales"               "Prevotellaceae"               
##   [16,] "Campylobacterales"           "Campylobacteraceae"           
##   [17,] "Pasteurellales"              "Pasteurellaceae"              
##   [18,] "Clostridiales"               "Lachnospiraceae"              
##   [19,] "Selenomonadales"             "Veillonellaceae"              
##   [20,] "Bacteroidales"               "Prevotellaceae"               
##   [21,] "Lactobacillales"             "Carnobacteriaceae"            
##   [22,] "Bacteroidales"               "Prevotellaceae"               
##   [23,] "Bacteroidales"               "Prevotellaceae"               
##   [24,] "Selenomonadales"             "Veillonellaceae"              
##   [25,] "Pasteurellales"              "Pasteurellaceae"              
##   [26,] "Bacteroidales"               "Prevotellaceae"               
##   [27,] "Bacteroidales"               "Porphyromonadaceae"           
##   [28,] "Selenomonadales"             "Veillonellaceae"              
##   [29,] "Lactobacillales"             "Streptococcaceae"             
##   [30,] "Lactobacillales"             "Streptococcaceae"             
##   [31,] "Bacteroidales"               "Porphyromonadaceae"           
##   [32,] "Lactobacillales"             "Streptococcaceae"             
##   [33,] "Bacteroidales"               "Prevotellaceae"               
##   [34,] "Bacteroidales"               "Prevotellaceae"               
##   [35,] "Clostridiales"               "Peptostreptococcaceae"        
##   [36,] "Selenomonadales"             "Veillonellaceae"              
##   [37,] "Bacteroidales"               "Porphyromonadaceae"           
##   [38,] "Selenomonadales"             "Veillonellaceae"              
##   [39,] "Clostridiales"               "Lachnospiraceae"              
##   [40,] "Lactobacillales"             "Streptococcaceae"             
##   [41,] "Bacteroidales"               "Prevotellaceae"               
##   [42,] "Lactobacillales"             "Streptococcaceae"             
##   [43,] "Lactobacillales"             "Streptococcaceae"             
##   [44,] "Bacteroidales"               "Prevotellaceae"               
##   [45,] "Betaproteobacteriales"       "Neisseriaceae"                
##   [46,] "Bacteroidales"               "Prevotellaceae"               
##   [47,] "Selenomonadales"             "Veillonellaceae"              
##   [48,] "Selenomonadales"             "Veillonellaceae"              
##   [49,] "Clostridiales"               "Lachnospiraceae"              
##   [50,] "Fusobacteriales"             "Leptotrichiaceae"             
##   [51,] "Lactobacillales"             "Streptococcaceae"             
##   [52,] "Pasteurellales"              "Pasteurellaceae"              
##   [53,] "Lactobacillales"             "Streptococcaceae"             
##   [54,] "Fusobacteriales"             "Leptotrichiaceae"             
##   [55,] "Bacteroidales"               "Prevotellaceae"               
##   [56,] "Bacteroidales"               "Prevotellaceae"               
##   [57,] "Selenomonadales"             "Veillonellaceae"              
##   [58,] "Clostridiales"               "Lachnospiraceae"              
##   [59,] "Coriobacteriales"            "Atopobiaceae"                 
##   [60,] "Pasteurellales"              "Pasteurellaceae"              
##   [61,] "Pasteurellales"              "Pasteurellaceae"              
##   [62,] "Bacteroidales"               "Prevotellaceae"               
##   [63,] "Clostridiales"               "Lachnospiraceae"              
##   [64,] "Selenomonadales"             "Veillonellaceae"              
##   [65,] "Pasteurellales"              "Pasteurellaceae"              
##   [66,] "Selenomonadales"             "Veillonellaceae"              
##   [67,] "Saccharimonadales"           "Saccharimonadaceae"           
##   [68,] "Micrococcales"               "Micrococcaceae"               
##   [69,] "Betaproteobacteriales"       "Neisseriaceae"                
##   [70,] "Clostridiales"               "Family_XI"                    
##   [71,] "Bacteroidales"               "Prevotellaceae"               
##   [72,] "Micrococcales"               "Micrococcaceae"               
##   [73,] "Clostridiales"               "Lachnospiraceae"              
##   [74,] "Selenomonadales"             "Veillonellaceae"              
##   [75,] "Micrococcales"               "Micrococcaceae"               
##   [76,] "Betaproteobacteriales"       "Neisseriaceae"                
##   [77,] "Clostridiales"               "Family_XIII"                  
##   [78,] "Bacteroidales"               "Porphyromonadaceae"           
##   [79,] "Selenomonadales"             "Veillonellaceae"              
##   [80,] "Clostridiales"               "Lachnospiraceae"              
##   [81,] "Pasteurellales"              "Pasteurellaceae"              
##   [82,] "Selenomonadales"             "Veillonellaceae"              
##   [83,] "Bacteroidales"               "Prevotellaceae"               
##   [84,] "Bacteroidales"               "Prevotellaceae"               
##   [85,] "Selenomonadales"             "Veillonellaceae"              
##   [86,] "Bacteroidales"               "Prevotellaceae"               
##   [87,] "Fusobacteriales"             "Leptotrichiaceae"             
##   [88,] "Clostridiales"               "Ruminococcaceae"              
##   [89,] "Bacteroidales"               "Prevotellaceae"               
##   [90,] "Clostridiales"               "Ruminococcaceae"              
##   [91,] "Micrococcales"               "Micrococcaceae"               
##   [92,] "Betaproteobacteriales"       "Neisseriaceae"                
##   [93,] "Micrococcales"               "Micrococcaceae"               
##   [94,] "Bacillales"                  "Family_XI"                    
##   [95,] "Bacteroidales"               "Prevotellaceae"               
##   [96,] "Betaproteobacteriales"       "Neisseriaceae"                
##   [97,] "Fusobacteriales"             "Fusobacteriaceae"             
##   [98,] "Selenomonadales"             "Veillonellaceae"              
##   [99,] "Fusobacteriales"             "Fusobacteriaceae"             
##  [100,] "Bacteroidales"               "Prevotellaceae"               
##  [101,] "Bacteroidales"               "Prevotellaceae"               
##  [102,] "Pasteurellales"              "Pasteurellaceae"              
##  [103,] "Pasteurellales"              "Pasteurellaceae"              
##  [104,] "Bacteroidales"               "Prevotellaceae"               
##  [105,] "Lactobacillales"             "Carnobacteriaceae"            
##  [106,] "Bacillales"                  "Family_XI"                    
##  [107,] "Selenomonadales"             "Veillonellaceae"              
##  [108,] "Selenomonadales"             "Veillonellaceae"              
##  [109,] "Micrococcales"               "Micrococcaceae"               
##  [110,] "Pasteurellales"              "Pasteurellaceae"              
##  [111,] "Campylobacterales"           "Campylobacteraceae"           
##  [112,] "Bacteroidales"               "Prevotellaceae"               
##  [113,] "Betaproteobacteriales"       "Neisseriaceae"                
##  [114,] "Betaproteobacteriales"       "Neisseriaceae"                
##  [115,] "Saccharimonadales"           NA                             
##  [116,] "Bacteroidales"               "Prevotellaceae"               
##  [117,] "Betaproteobacteriales"       "Neisseriaceae"                
##  [118,] "Pasteurellales"              "Pasteurellaceae"              
##  [119,] "Pasteurellales"              "Pasteurellaceae"              
##  [120,] "Pasteurellales"              "Pasteurellaceae"              
##  [121,] "Pasteurellales"              "Pasteurellaceae"              
##  [122,] "Bacteroidales"               "Prevotellaceae"               
##  [123,] "Lactobacillales"             "Streptococcaceae"             
##  [124,] "Bacteroidales"               "Prevotellaceae"               
##  [125,] "Betaproteobacteriales"       "Neisseriaceae"                
##  [126,] "Bacteroidales"               "Prevotellaceae"               
##  [127,] "Selenomonadales"             "Veillonellaceae"              
##  [128,] "Pasteurellales"              "Pasteurellaceae"              
##  [129,] "Pasteurellales"              "Pasteurellaceae"              
##  [130,] "Pasteurellales"              "Pasteurellaceae"              
##  [131,] "Betaproteobacteriales"       "Burkholderiaceae"             
##  [132,] "Bacteroidales"               "Prevotellaceae"               
##  [133,] "Saccharimonadales"           "Saccharimonadaceae"           
##  [134,] "Bacteroidales"               "Prevotellaceae"               
##  [135,] "Selenomonadales"             "Veillonellaceae"              
##  [136,] "Selenomonadales"             "Veillonellaceae"              
##  [137,] "Bacteroidales"               "Prevotellaceae"               
##  [138,] "Flavobacteriales"            "Flavobacteriaceae"            
##  [139,] "Bacteroidales"               "Prevotellaceae"               
##  [140,] "Lactobacillales"             "Streptococcaceae"             
##  [141,] "Selenomonadales"             "Veillonellaceae"              
##  [142,] "Selenomonadales"             "Veillonellaceae"              
##  [143,] "Fusobacteriales"             "Leptotrichiaceae"             
##  [144,] "Selenomonadales"             "Veillonellaceae"              
##  [145,] "Lactobacillales"             "Streptococcaceae"             
##  [146,] "Flavobacteriales"            "Weeksellaceae"                
##  [147,] "Betaproteobacteriales"       "Neisseriaceae"                
##  [148,] "Selenomonadales"             "Veillonellaceae"              
##  [149,] "Selenomonadales"             "Veillonellaceae"              
##  [150,] "Selenomonadales"             "Veillonellaceae"              
##  [151,] "Betaproteobacteriales"       "Neisseriaceae"                
##  [152,] "Bacteroidales"               "Prevotellaceae"               
##  [153,] "Selenomonadales"             "Veillonellaceae"              
##  [154,] "Bacteroidales"               "Prevotellaceae"               
##  [155,] "Clostridiales"               "Lachnospiraceae"              
##  [156,] "Enterobacteriales"           "Enterobacteriaceae"           
##  [157,] "Selenomonadales"             "Veillonellaceae"              
##  [158,] "Bacteroidales"               "Prevotellaceae"               
##  [159,] "Selenomonadales"             "Veillonellaceae"              
##  [160,] "Clostridiales"               "Lachnospiraceae"              
##  [161,] "Lactobacillales"             "Streptococcaceae"             
##  [162,] "Micrococcales"               "Micrococcaceae"               
##  [163,] "Bacteroidales"               "Prevotellaceae"               
##  [164,] "Selenomonadales"             "Veillonellaceae"              
##  [165,] "Campylobacterales"           "Campylobacteraceae"           
##  [166,] "Bacteroidales"               "Prevotellaceae"               
##  [167,] "Bacteroidales"               "Porphyromonadaceae"           
##  [168,] "Lactobacillales"             "Streptococcaceae"             
##  [169,] "Bacteroidales"               "Prevotellaceae"               
##  [170,] "Betaproteobacteriales"       "Neisseriaceae"                
##  [171,] "Lactobacillales"             "Streptococcaceae"             
##  [172,] "Lactobacillales"             "Streptococcaceae"             
##  [173,] "Saccharimonadales"           "Saccharimonadaceae"           
##  [174,] "Bacteroidales"               "Prevotellaceae"               
##  [175,] "Bacteroidales"               "Prevotellaceae"               
##  [176,] "Clostridiales"               "Lachnospiraceae"              
##  [177,] "Selenomonadales"             "Veillonellaceae"              
##  [178,] "Selenomonadales"             "Veillonellaceae"              
##  [179,] "Bacteroidales"               "Prevotellaceae"               
##  [180,] "Betaproteobacteriales"       "Neisseriaceae"                
##  [181,] "Absconditabacteriales_(SR1)" NA                             
##  [182,] "Selenomonadales"             "Veillonellaceae"              
##  [183,] "Betaproteobacteriales"       "Neisseriaceae"                
##  [184,] "Selenomonadales"             "Veillonellaceae"              
##  [185,] "Pasteurellales"              "Pasteurellaceae"              
##  [186,] "Selenomonadales"             "Veillonellaceae"              
##  [187,] "Selenomonadales"             "Veillonellaceae"              
##  [188,] "Bacteroidales"               "Prevotellaceae"               
##  [189,] "Bacteroidales"               "Prevotellaceae"               
##  [190,] "Pasteurellales"              "Pasteurellaceae"              
##  [191,] "Bacteroidales"               "Prevotellaceae"               
##  [192,] "Bacteroidales"               "Porphyromonadaceae"           
##  [193,] "Betaproteobacteriales"       "Neisseriaceae"                
##  [194,] "Bacteroidales"               "Prevotellaceae"               
##  [195,] "Selenomonadales"             "Veillonellaceae"              
##  [196,] "Selenomonadales"             "Veillonellaceae"              
##  [197,] "Fusobacteriales"             "Fusobacteriaceae"             
##  [198,] "Selenomonadales"             "Veillonellaceae"              
##  [199,] "Bacteroidales"               "Prevotellaceae"               
##  [200,] "Selenomonadales"             "Veillonellaceae"              
##  [201,] "Bacteroidales"               "Prevotellaceae"               
##  [202,] "Clostridiales"               "Lachnospiraceae"              
##  [203,] "Campylobacterales"           "Campylobacteraceae"           
##  [204,] "Bacteroidales"               "Prevotellaceae"               
##  [205,] "Selenomonadales"             "Veillonellaceae"              
##  [206,] "Selenomonadales"             "Veillonellaceae"              
##  [207,] "Flavobacteriales"            "Flavobacteriaceae"            
##  [208,] "Betaproteobacteriales"       "Neisseriaceae"                
##  [209,] "Betaproteobacteriales"       "Neisseriaceae"                
##  [210,] "Lactobacillales"             "Streptococcaceae"             
##  [211,] "Clostridiales"               "Family_XIII"                  
##  [212,] "Selenomonadales"             "Veillonellaceae"              
##  [213,] "Pasteurellales"              "Pasteurellaceae"              
##  [214,] "Betaproteobacteriales"       "Neisseriaceae"                
##  [215,] "Selenomonadales"             "Veillonellaceae"              
##  [216,] "Bacteroidales"               "Prevotellaceae"               
##  [217,] "Lactobacillales"             "Streptococcaceae"             
##  [218,] "Betaproteobacteriales"       "Neisseriaceae"                
##  [219,] "Selenomonadales"             "Veillonellaceae"              
##  [220,] "Lactobacillales"             "Streptococcaceae"             
##  [221,] "Clostridiales"               "Lachnospiraceae"              
##  [222,] "Lactobacillales"             "Streptococcaceae"             
##  [223,] "Clostridiales"               "Lachnospiraceae"              
##  [224,] "Bacteroidales"               "Prevotellaceae"               
##  [225,] "Fusobacteriales"             "Leptotrichiaceae"             
##  [226,] "Selenomonadales"             "Veillonellaceae"              
##  [227,] "Bacteroidales"               "Prevotellaceae"               
##  [228,] "Lactobacillales"             "Streptococcaceae"             
##  [229,] "Flavobacteriales"            "Flavobacteriaceae"            
##  [230,] "Flavobacteriales"            "Flavobacteriaceae"            
##  [231,] "Fusobacteriales"             "Leptotrichiaceae"             
##  [232,] "Fusobacteriales"             "Fusobacteriaceae"             
##  [233,] "Pasteurellales"              "Pasteurellaceae"              
##  [234,] "Fusobacteriales"             "Fusobacteriaceae"             
##  [235,] "Selenomonadales"             "Veillonellaceae"              
##  [236,] "Saccharimonadales"           NA                             
##  [237,] "Bacteroidales"               "Prevotellaceae"               
##  [238,] "Selenomonadales"             "Veillonellaceae"              
##  [239,] "Bacteroidales"               "Prevotellaceae"               
##  [240,] "Selenomonadales"             "Veillonellaceae"              
##  [241,] "Fusobacteriales"             "Fusobacteriaceae"             
##  [242,] "Bacteroidales"               "Porphyromonadaceae"           
##  [243,] "Selenomonadales"             "Veillonellaceae"              
##  [244,] "Chloroplast"                 NA                             
##  [245,] "Bacteroidales"               "Prevotellaceae"               
##  [246,] "Pasteurellales"              "Pasteurellaceae"              
##  [247,] "Betaproteobacteriales"       "Neisseriaceae"                
##  [248,] "Chloroplast"                 NA                             
##  [249,] "Absconditabacteriales_(SR1)" NA                             
##  [250,] "Bacteroidales"               "Prevotellaceae"               
##  [251,] "Bacteroidales"               "Prevotellaceae"               
##  [252,] "Bacteroidales"               "Prevotellaceae"               
##  [253,] "Clostridiales"               "Lachnospiraceae"              
##  [254,] "Pasteurellales"              "Pasteurellaceae"              
##  [255,] "Enterobacteriales"           "Enterobacteriaceae"           
##  [256,] "Selenomonadales"             "Veillonellaceae"              
##  [257,] "Selenomonadales"             "Veillonellaceae"              
##  [258,] "Pasteurellales"              "Pasteurellaceae"              
##  [259,] "Saccharimonadales"           "Saccharimonadaceae"           
##  [260,] "Actinomycetales"             "Actinomycetaceae"             
##  [261,] "Clostridiales"               "Lachnospiraceae"              
##  [262,] "Bacteroidales"               "Prevotellaceae"               
##  [263,] "Bacteroidales"               "Prevotellaceae"               
##  [264,] "Betaproteobacteriales"       "Neisseriaceae"                
##  [265,] "Saccharimonadales"           NA                             
##  [266,] "Betaproteobacteriales"       "Neisseriaceae"                
##  [267,] "Erysipelotrichales"          "Erysipelotrichaceae"          
##  [268,] "Bacteroidales"               "Prevotellaceae"               
##  [269,] "Clostridiales"               "Family_XI"                    
##  [270,] "Betaproteobacteriales"       "Neisseriaceae"                
##  [271,] "Selenomonadales"             "Veillonellaceae"              
##  [272,] "Betaproteobacteriales"       "Neisseriaceae"                
##  [273,] "Betaproteobacteriales"       "Neisseriaceae"                
##  [274,] "Selenomonadales"             "Veillonellaceae"              
##  [275,] "Lactobacillales"             "Streptococcaceae"             
##  [276,] "Selenomonadales"             "Veillonellaceae"              
##  [277,] "Bacteroidales"               "Prevotellaceae"               
##  [278,] "Flavobacteriales"            "Flavobacteriaceae"            
##  [279,] "Pasteurellales"              "Pasteurellaceae"              
##  [280,] "Bacteroidales"               "Prevotellaceae"               
##  [281,] "Fusobacteriales"             "Fusobacteriaceae"             
##  [282,] NA                            NA                             
##  [283,] "Fusobacteriales"             "Fusobacteriaceae"             
##  [284,] "Bacteroidales"               "Porphyromonadaceae"           
##  [285,] "Bacteroidales"               "Prevotellaceae"               
##  [286,] "Absconditabacteriales_(SR1)" NA                             
##  [287,] "Bacteroidales"               "Prevotellaceae"               
##  [288,] "Selenomonadales"             "Veillonellaceae"              
##  [289,] "Bacteroidales"               "Prevotellaceae"               
##  [290,] "Clostridiales"               "Lachnospiraceae"              
##  [291,] "Bacteroidales"               "Prevotellaceae"               
##  [292,] "Pasteurellales"              "Pasteurellaceae"              
##  [293,] "Selenomonadales"             "Veillonellaceae"              
##  [294,] "Saccharimonadales"           "Saccharimonadaceae"           
##  [295,] "Lactobacillales"             "Streptococcaceae"             
##  [296,] "Selenomonadales"             "Veillonellaceae"              
##  [297,] NA                            NA                             
##  [298,] "Selenomonadales"             "Veillonellaceae"              
##  [299,] "Betaproteobacteriales"       "Neisseriaceae"                
##  [300,] "Bacteroidales"               "Prevotellaceae"               
##  [301,] "Bacteroidales"               "Prevotellaceae"               
##  [302,] "Fusobacteriales"             "Leptotrichiaceae"             
##  [303,] "Bacteroidales"               "Prevotellaceae"               
##  [304,] "Fusobacteriales"             "Fusobacteriaceae"             
##  [305,] "Fusobacteriales"             "Fusobacteriaceae"             
##  [306,] "Bacteroidales"               "Prevotellaceae"               
##  [307,] "Selenomonadales"             "Veillonellaceae"              
##  [308,] "Fusobacteriales"             "Fusobacteriaceae"             
##  [309,] "Clostridiales"               "Peptostreptococcaceae"        
##  [310,] "Pasteurellales"              "Pasteurellaceae"              
##  [311,] "Selenomonadales"             "Veillonellaceae"              
##  [312,] "Fusobacteriales"             "Fusobacteriaceae"             
##  [313,] "Saccharimonadales"           NA                             
##  [314,] "Campylobacterales"           "Campylobacteraceae"           
##  [315,] "Selenomonadales"             "Veillonellaceae"              
##  [316,] "Bacteroidales"               "Prevotellaceae"               
##  [317,] "Selenomonadales"             "Veillonellaceae"              
##  [318,] "Clostridiales"               "Lachnospiraceae"              
##  [319,] "Clostridiales"               "Family_XIII"                  
##  [320,] "Bacteroidales"               "Prevotellaceae"               
##  [321,] "Coriobacteriales"            "Atopobiaceae"                 
##  [322,] "Lactobacillales"             "Streptococcaceae"             
##  [323,] "Saccharimonadales"           NA                             
##  [324,] "Selenomonadales"             "Veillonellaceae"              
##  [325,] "Pasteurellales"              "Pasteurellaceae"              
##  [326,] "Saccharimonadales"           "Saccharimonadaceae"           
##  [327,] "Bacteroidales"               "Porphyromonadaceae"           
##  [328,] "Coriobacteriales"            "Atopobiaceae"                 
##  [329,] "Selenomonadales"             "Veillonellaceae"              
##  [330,] "Bacteroidales"               "Prevotellaceae"               
##  [331,] "Bacteroidales"               "Prevotellaceae"               
##  [332,] "Saccharimonadales"           NA                             
##  [333,] "Clostridiales"               "Lachnospiraceae"              
##  [334,] "Selenomonadales"             "Veillonellaceae"              
##  [335,] "Clostridiales"               "Lachnospiraceae"              
##  [336,] "Fusobacteriales"             "Leptotrichiaceae"             
##  [337,] "Lactobacillales"             "Aerococcaceae"                
##  [338,] "Fusobacteriales"             "Leptotrichiaceae"             
##  [339,] "Micrococcales"               "Micrococcaceae"               
##  [340,] "Fusobacteriales"             "Fusobacteriaceae"             
##  [341,] "Fusobacteriales"             "Fusobacteriaceae"             
##  [342,] "Bacteroidales"               "Prevotellaceae"               
##  [343,] "Betaproteobacteriales"       "Neisseriaceae"                
##  [344,] "Bacteroidales"               "Prevotellaceae"               
##  [345,] "Clostridiales"               "Peptostreptococcaceae"        
##  [346,] "Betaproteobacteriales"       "Neisseriaceae"                
##  [347,] NA                            NA                             
##  [348,] "Betaproteobacteriales"       "Neisseriaceae"                
##  [349,] "Pasteurellales"              "Pasteurellaceae"              
##  [350,] "Bacteroidales"               "Prevotellaceae"               
##  [351,] "Clostridiales"               "Family_XIII"                  
##  [352,] "Selenomonadales"             "Veillonellaceae"              
##  [353,] "Bacteroidales"               "Prevotellaceae"               
##  [354,] "Pasteurellales"              "Pasteurellaceae"              
##  [355,] "Fusobacteriales"             "Fusobacteriaceae"             
##  [356,] "Selenomonadales"             "Veillonellaceae"              
##  [357,] "Fusobacteriales"             "Fusobacteriaceae"             
##  [358,] "Bacteroidales"               "Prevotellaceae"               
##  [359,] "Selenomonadales"             "Veillonellaceae"              
##  [360,] "Clostridiales"               "Peptococcaceae"               
##  [361,] "Clostridiales"               "Family_XIII"                  
##  [362,] "Pasteurellales"              "Pasteurellaceae"              
##  [363,] "Bacteroidales"               "Prevotellaceae"               
##  [364,] "Clostridiales"               "Peptostreptococcaceae"        
##  [365,] "Fusobacteriales"             NA                             
##  [366,] "Saccharimonadales"           NA                             
##  [367,] "Clostridiales"               "Lachnospiraceae"              
##  [368,] "Clostridiales"               "Lachnospiraceae"              
##  [369,] "Actinomycetales"             "Actinomycetaceae"             
##  [370,] "Bacteroidales"               "Prevotellaceae"               
##  [371,] "Saccharimonadales"           NA                             
##  [372,] "Bacillales"                  "Family_XI"                    
##  [373,] "Lactobacillales"             "Streptococcaceae"             
##  [374,] "Selenomonadales"             "Veillonellaceae"              
##  [375,] "Saccharimonadales"           NA                             
##  [376,] "Selenomonadales"             "Veillonellaceae"              
##  [377,] "Fusobacteriales"             "Fusobacteriaceae"             
##  [378,] "Lactobacillales"             "Streptococcaceae"             
##  [379,] "Selenomonadales"             "Veillonellaceae"              
##  [380,] "Fusobacteriales"             "Leptotrichiaceae"             
##  [381,] "Fusobacteriales"             "Fusobacteriaceae"             
##  [382,] "Clostridiales"               "Family_XI"                    
##  [383,] "Bacteroidales"               "Prevotellaceae"               
##  [384,] "Bacteroidales"               "Prevotellaceae"               
##  [385,] "Lactobacillales"             "Streptococcaceae"             
##  [386,] "Enterobacteriales"           "Enterobacteriaceae"           
##  [387,] "Fusobacteriales"             "Leptotrichiaceae"             
##  [388,] "Actinomycetales"             "Actinomycetaceae"             
##  [389,] "Fusobacteriales"             "Leptotrichiaceae"             
##  [390,] "Bacteroidales"               "Prevotellaceae"               
##  [391,] "Actinomycetales"             "Actinomycetaceae"             
##  [392,] "Fusobacteriales"             "Fusobacteriaceae"             
##  [393,] "Pasteurellales"              "Pasteurellaceae"              
##  [394,] "Enterobacteriales"           "Enterobacteriaceae"           
##  [395,] "Selenomonadales"             "Veillonellaceae"              
##  [396,] "Bacteroidales"               "Prevotellaceae"               
##  [397,] "Bacteroidales"               "Prevotellaceae"               
##  [398,] "Pasteurellales"              "Pasteurellaceae"              
##  [399,] "Fusobacteriales"             "Leptotrichiaceae"             
##  [400,] "Pasteurellales"              "Pasteurellaceae"              
##  [401,] "Lactobacillales"             "Streptococcaceae"             
##  [402,] "Flavobacteriales"            "Flavobacteriaceae"            
##  [403,] "Bacteroidales"               "Prevotellaceae"               
##  [404,] "Bacteroidales"               "Prevotellaceae"               
##  [405,] "Clostridiales"               "Lachnospiraceae"              
##  [406,] "Selenomonadales"             "Veillonellaceae"              
##  [407,] "Bacteroidales"               "Prevotellaceae"               
##  [408,] "Bacillales"                  "Family_XI"                    
##  [409,] "Selenomonadales"             "Veillonellaceae"              
##  [410,] "Bacteroidales"               "Porphyromonadaceae"           
##  [411,] "Bacteroidales"               "Prevotellaceae"               
##  [412,] "Bacteroidales"               "Prevotellaceae"               
##  [413,] "Betaproteobacteriales"       "Neisseriaceae"                
##  [414,] "Clostridiales"               "Lachnospiraceae"              
##  [415,] "Bacteroidales"               "Prevotellaceae"               
##  [416,] "Pseudomonadales"             "Moraxellaceae"                
##  [417,] "Bacteroidales"               "Prevotellaceae"               
##  [418,] "Bacteroidales"               "Prevotellaceae"               
##  [419,] "Fusobacteriales"             "Leptotrichiaceae"             
##  [420,] "Bacteroidales"               "Prevotellaceae"               
##  [421,] "Bacillales"                  "Family_XI"                    
##  [422,] "Selenomonadales"             "Veillonellaceae"              
##  [423,] "Selenomonadales"             "Veillonellaceae"              
##  [424,] "Selenomonadales"             "Veillonellaceae"              
##  [425,] "Pasteurellales"              "Pasteurellaceae"              
##  [426,] "Bacteroidales"               "Prevotellaceae"               
##  [427,] "Bacteroidales"               "Prevotellaceae"               
##  [428,] "Bacteroidales"               "Prevotellaceae"               
##  [429,] "Selenomonadales"             "Veillonellaceae"              
##  [430,] "Selenomonadales"             "Veillonellaceae"              
##  [431,] "Selenomonadales"             "Veillonellaceae"              
##  [432,] "Bacteroidales"               "Prevotellaceae"               
##  [433,] "Selenomonadales"             "Veillonellaceae"              
##  [434,] "Bacteroidales"               "Prevotellaceae"               
##  [435,] "Clostridiales"               "Lachnospiraceae"              
##  [436,] "Selenomonadales"             "Veillonellaceae"              
##  [437,] "Bacteroidales"               "Prevotellaceae"               
##  [438,] "Lactobacillales"             "Streptococcaceae"             
##  [439,] "Bacteroidales"               "Prevotellaceae"               
##  [440,] "Betaproteobacteriales"       "Neisseriaceae"                
##  [441,] "Bacteroidales"               "Porphyromonadaceae"           
##  [442,] "Fusobacteriales"             "Fusobacteriaceae"             
##  [443,] "Selenomonadales"             "Veillonellaceae"              
##  [444,] "Selenomonadales"             "Veillonellaceae"              
##  [445,] "Selenomonadales"             "Veillonellaceae"              
##  [446,] "Selenomonadales"             "Veillonellaceae"              
##  [447,] "Lactobacillales"             "Streptococcaceae"             
##  [448,] "Fusobacteriales"             "Leptotrichiaceae"             
##  [449,] "Flavobacteriales"            "Flavobacteriaceae"            
##  [450,] "Fusobacteriales"             "Leptotrichiaceae"             
##  [451,] "Bacteroidales"               "Prevotellaceae"               
##  [452,] "Campylobacterales"           NA                             
##  [453,] "Clostridiales"               "Family_XIII"                  
##  [454,] "Flavobacteriales"            "Flavobacteriaceae"            
##  [455,] "Selenomonadales"             "Veillonellaceae"              
##  [456,] "Clostridiales"               "Ruminococcaceae"              
##  [457,] "Betaproteobacteriales"       "Neisseriaceae"                
##  [458,] "Actinomycetales"             "Actinomycetaceae"             
##  [459,] "Fusobacteriales"             "Leptotrichiaceae"             
##  [460,] "Selenomonadales"             "Veillonellaceae"              
##  [461,] "Saccharimonadales"           "Saccharimonadaceae"           
##  [462,] "Fusobacteriales"             "Leptotrichiaceae"             
##  [463,] "Bacteroidales"               "Prevotellaceae"               
##  [464,] "Lactobacillales"             "Streptococcaceae"             
##  [465,] "Bacteroidales"               "Prevotellaceae"               
##  [466,] "Selenomonadales"             "Veillonellaceae"              
##  [467,] "Bacteroidales"               "Prevotellaceae"               
##  [468,] "Pasteurellales"              "Pasteurellaceae"              
##  [469,] "Saccharimonadales"           NA                             
##  [470,] "Betaproteobacteriales"       "Neisseriaceae"                
##  [471,] "Pasteurellales"              "Pasteurellaceae"              
##  [472,] "Bacteroidales"               "Prevotellaceae"               
##  [473,] "Bacteroidales"               "Prevotellaceae"               
##  [474,] "Selenomonadales"             "Veillonellaceae"              
##  [475,] "Bacteroidales"               "Prevotellaceae"               
##  [476,] "Pasteurellales"              "Pasteurellaceae"              
##  [477,] "Bacteroidales"               "Prevotellaceae"               
##  [478,] "Pasteurellales"              "Pasteurellaceae"              
##  [479,] "Enterobacteriales"           "Enterobacteriaceae"           
##  [480,] "Fusobacteriales"             "Leptotrichiaceae"             
##  [481,] "Bacteroidales"               "Prevotellaceae"               
##  [482,] "Bacteroidales"               "Prevotellaceae"               
##  [483,] "Fusobacteriales"             "Fusobacteriaceae"             
##  [484,] "Selenomonadales"             "Veillonellaceae"              
##  [485,] "Lactobacillales"             "Streptococcaceae"             
##  [486,] "Fusobacteriales"             NA                             
##  [487,] "Bacteroidales"               "Prevotellaceae"               
##  [488,] "Fusobacteriales"             "Leptotrichiaceae"             
##  [489,] "Bacteroidales"               "Porphyromonadaceae"           
##  [490,] "Campylobacterales"           "Campylobacteraceae"           
##  [491,] "Selenomonadales"             "Veillonellaceae"              
##  [492,] "Campylobacterales"           NA                             
##  [493,] "Bacteroidales"               "Prevotellaceae"               
##  [494,] "Fusobacteriales"             "Fusobacteriaceae"             
##  [495,] "Bacteroidales"               "Prevotellaceae"               
##  [496,] "Selenomonadales"             "Veillonellaceae"              
##  [497,] "Enterobacteriales"           "Enterobacteriaceae"           
##  [498,] "Bacteroidales"               "Prevotellaceae"               
##  [499,] "Clostridiales"               "Peptostreptococcaceae"        
##  [500,] "Selenomonadales"             "Veillonellaceae"              
##  [501,] "Selenomonadales"             "Veillonellaceae"              
##  [502,] "Actinomycetales"             "Actinomycetaceae"             
##  [503,] "Enterobacteriales"           "Enterobacteriaceae"           
##  [504,] "Selenomonadales"             "Veillonellaceae"              
##  [505,] "Betaproteobacteriales"       "Neisseriaceae"                
##  [506,] "Fusobacteriales"             "Leptotrichiaceae"             
##  [507,] "Actinomycetales"             "Actinomycetaceae"             
##  [508,] "Bacteroidales"               "Prevotellaceae"               
##  [509,] "Bacteroidales"               "Prevotellaceae"               
##  [510,] "Pasteurellales"              "Pasteurellaceae"              
##  [511,] "Selenomonadales"             "Veillonellaceae"              
##  [512,] "Selenomonadales"             "Veillonellaceae"              
##  [513,] "Pasteurellales"              "Pasteurellaceae"              
##  [514,] "Selenomonadales"             "Veillonellaceae"              
##  [515,] "Selenomonadales"             "Veillonellaceae"              
##  [516,] "Selenomonadales"             "Veillonellaceae"              
##  [517,] "Bacteroidales"               "Prevotellaceae"               
##  [518,] "Pasteurellales"              "Pasteurellaceae"              
##  [519,] "Selenomonadales"             "Veillonellaceae"              
##  [520,] "Actinomycetales"             "Actinomycetaceae"             
##  [521,] "Lactobacillales"             "Streptococcaceae"             
##  [522,] "Actinomycetales"             "Actinomycetaceae"             
##  [523,] "Clostridiales"               "Family_XI"                    
##  [524,] "Selenomonadales"             "Veillonellaceae"              
##  [525,] "Fusobacteriales"             "Fusobacteriaceae"             
##  [526,] "Bacteroidales"               "Prevotellaceae"               
##  [527,] "Bacteroidales"               "Prevotellaceae"               
##  [528,] "Betaproteobacteriales"       "Neisseriaceae"                
##  [529,] "Campylobacterales"           "Campylobacteraceae"           
##  [530,] "Lactobacillales"             "Streptococcaceae"             
##  [531,] "Cardiobacteriales"           "Cardiobacteriaceae"           
##  [532,] "Bacteroidales"               "Prevotellaceae"               
##  [533,] "Saccharimonadales"           NA                             
##  [534,] "Selenomonadales"             "Veillonellaceae"              
##  [535,] "Flavobacteriales"            "Flavobacteriaceae"            
##  [536,] "Bacteroidales"               "Prevotellaceae"               
##  [537,] "Selenomonadales"             "Veillonellaceae"              
##  [538,] "Pasteurellales"              "Pasteurellaceae"              
##  [539,] "Bacteroidales"               "Paludibacteraceae"            
##  [540,] "Enterobacteriales"           "Enterobacteriaceae"           
##  [541,] "Fusobacteriales"             "Leptotrichiaceae"             
##  [542,] "Lactobacillales"             "Streptococcaceae"             
##  [543,] "Bacteroidales"               "Porphyromonadaceae"           
##  [544,] "Bacteroidales"               "Prevotellaceae"               
##  [545,] "Clostridiales"               "Family_XIII"                  
##  [546,] "Bacteroidales"               "Prevotellaceae"               
##  [547,] "Betaproteobacteriales"       "Neisseriaceae"                
##  [548,] NA                            NA                             
##  [549,] "Selenomonadales"             "Veillonellaceae"              
##  [550,] "Bacteroidales"               "Prevotellaceae"               
##  [551,] "Bacillales"                  "Family_XI"                    
##  [552,] "Betaproteobacteriales"       "Neisseriaceae"                
##  [553,] "Bacteroidales"               "Prevotellaceae"               
##  [554,] "Flavobacteriales"            "Flavobacteriaceae"            
##  [555,] "Bacteroidales"               "Prevotellaceae"               
##  [556,] "Selenomonadales"             "Veillonellaceae"              
##  [557,] "Selenomonadales"             "Veillonellaceae"              
##  [558,] "Lactobacillales"             "Streptococcaceae"             
##  [559,] "Bacteroidales"               "Prevotellaceae"               
##  [560,] "Pasteurellales"              "Pasteurellaceae"              
##  [561,] "Bacteroidales"               "Porphyromonadaceae"           
##  [562,] "Saccharimonadales"           "Saccharimonadaceae"           
##  [563,] "Pasteurellales"              "Pasteurellaceae"              
##  [564,] "Lactobacillales"             "Streptococcaceae"             
##  [565,] "Selenomonadales"             "Veillonellaceae"              
##  [566,] "Bacteroidales"               "Prevotellaceae"               
##  [567,] "Lactobacillales"             "Streptococcaceae"             
##  [568,] "Bacteroidales"               "Prevotellaceae"               
##  [569,] "Bacteroidales"               "Prevotellaceae"               
##  [570,] "Fusobacteriales"             "Leptotrichiaceae"             
##  [571,] "Bacteroidales"               "Prevotellaceae"               
##  [572,] "Bacteroidales"               "Tannerellaceae"               
##  [573,] "Betaproteobacteriales"       "Neisseriaceae"                
##  [574,] "Bacteroidales"               "Prevotellaceae"               
##  [575,] "Flavobacteriales"            "Flavobacteriaceae"            
##  [576,] "Bacteroidales"               "Prevotellaceae"               
##  [577,] "Saccharimonadales"           "Saccharimonadaceae"           
##  [578,] "Selenomonadales"             "Veillonellaceae"              
##  [579,] "Selenomonadales"             "Veillonellaceae"              
##  [580,] "Selenomonadales"             "Veillonellaceae"              
##  [581,] "Fusobacteriales"             "Leptotrichiaceae"             
##  [582,] "Pasteurellales"              "Pasteurellaceae"              
##  [583,] "Fusobacteriales"             "Fusobacteriaceae"             
##  [584,] "Bacteroidales"               "Porphyromonadaceae"           
##  [585,] "Betaproteobacteriales"       "Neisseriaceae"                
##  [586,] "Fusobacteriales"             "Fusobacteriaceae"             
##  [587,] "Fusobacteriales"             "Fusobacteriaceae"             
##  [588,] "Bacteroidales"               "Tannerellaceae"               
##  [589,] "Fusobacteriales"             "Fusobacteriaceae"             
##  [590,] "Bacteroidales"               "Prevotellaceae"               
##  [591,] "Betaproteobacteriales"       "Neisseriaceae"                
##  [592,] "Bacteroidales"               "Prevotellaceae"               
##  [593,] "Bacteroidales"               "Prevotellaceae"               
##  [594,] "Bacteroidales"               "Prevotellaceae"               
##  [595,] "Pasteurellales"              "Pasteurellaceae"              
##  [596,] "Bacteroidales"               "Prevotellaceae"               
##  [597,] "Selenomonadales"             "Veillonellaceae"              
##  [598,] "Betaproteobacteriales"       "Neisseriaceae"                
##  [599,] "Clostridiales"               "Peptostreptococcaceae"        
##  [600,] "Selenomonadales"             "Veillonellaceae"              
##  [601,] "Pasteurellales"              "Pasteurellaceae"              
##  [602,] "Bacteroidales"               "Prevotellaceae"               
##  [603,] "Bacteroidales"               "Prevotellaceae"               
##  [604,] "Selenomonadales"             "Veillonellaceae"              
##  [605,] "Betaproteobacteriales"       "Neisseriaceae"                
##  [606,] "Fusobacteriales"             "Fusobacteriaceae"             
##  [607,] "Bacteroidales"               "Prevotellaceae"               
##  [608,] "Saccharimonadales"           "Saccharimonadaceae"           
##  [609,] "Bacteroidales"               "Prevotellaceae"               
##  [610,] "Saccharimonadales"           "Saccharimonadaceae"           
##  [611,] "Pasteurellales"              "Pasteurellaceae"              
##  [612,] "Selenomonadales"             "Veillonellaceae"              
##  [613,] "Fusobacteriales"             "Fusobacteriaceae"             
##  [614,] "Fusobacteriales"             "Leptotrichiaceae"             
##  [615,] "Bacteroidales"               "Prevotellaceae"               
##  [616,] "Selenomonadales"             "Veillonellaceae"              
##  [617,] "Bacteroidales"               "Prevotellaceae"               
##  [618,] "Selenomonadales"             "Veillonellaceae"              
##  [619,] "Bacillales"                  "Family_XI"                    
##  [620,] "Bacteroidales"               "Porphyromonadaceae"           
##  [621,] "Selenomonadales"             "Veillonellaceae"              
##  [622,] "Bacteroidales"               "Paludibacteraceae"            
##  [623,] "Selenomonadales"             "Veillonellaceae"              
##  [624,] "Bacteroidales"               "Prevotellaceae"               
##  [625,] "Bacteroidales"               "Prevotellaceae"               
##  [626,] "Bacteroidales"               "Prevotellaceae"               
##  [627,] "Bacteroidales"               "Prevotellaceae"               
##  [628,] "Fusobacteriales"             "Fusobacteriaceae"             
##  [629,] "Bacteroidales"               "Prevotellaceae"               
##  [630,] "Micrococcales"               "Micrococcaceae"               
##  [631,] "Selenomonadales"             "Veillonellaceae"              
##  [632,] "Pasteurellales"              "Pasteurellaceae"              
##  [633,] "Coriobacteriales"            "Atopobiaceae"                 
##  [634,] "Selenomonadales"             "Veillonellaceae"              
##  [635,] "Saccharimonadales"           "Saccharimonadaceae"           
##  [636,] "Micrococcales"               NA                             
##  [637,] "Bacteroidales"               "Porphyromonadaceae"           
##  [638,] "Pasteurellales"              "Pasteurellaceae"              
##  [639,] "Lactobacillales"             "Streptococcaceae"             
##  [640,] "Absconditabacteriales_(SR1)" NA                             
##  [641,] "Selenomonadales"             "Veillonellaceae"              
##  [642,] NA                            NA                             
##  [643,] "Enterobacteriales"           "Enterobacteriaceae"           
##  [644,] "Bacteroidales"               "Prevotellaceae"               
##  [645,] "Bacteroidales"               "Prevotellaceae"               
##  [646,] "Bacteroidales"               "Prevotellaceae"               
##  [647,] "Micrococcales"               "Micrococcaceae"               
##  [648,] "Lactobacillales"             "Streptococcaceae"             
##  [649,] "Bacteroidales"               "Prevotellaceae"               
##  [650,] "Clostridiales"               "Peptostreptococcaceae"        
##  [651,] "Selenomonadales"             "Veillonellaceae"              
##  [652,] "Bacteroidales"               "Prevotellaceae"               
##  [653,] "Selenomonadales"             "Veillonellaceae"              
##  [654,] "Saccharimonadales"           NA                             
##  [655,] "Bacteroidales"               "Prevotellaceae"               
##  [656,] "Selenomonadales"             "Veillonellaceae"              
##  [657,] "Fusobacteriales"             "Leptotrichiaceae"             
##  [658,] "Bacteroidales"               "Porphyromonadaceae"           
##  [659,] "Selenomonadales"             "Veillonellaceae"              
##  [660,] "Fusobacteriales"             "Leptotrichiaceae"             
##  [661,] "Micrococcales"               NA                             
##  [662,] "Fusobacteriales"             "Fusobacteriaceae"             
##  [663,] "Fusobacteriales"             "Fusobacteriaceae"             
##  [664,] "Pasteurellales"              "Pasteurellaceae"              
##  [665,] "Fusobacteriales"             "Leptotrichiaceae"             
##  [666,] "Selenomonadales"             "Veillonellaceae"              
##  [667,] "Chloroplast"                 NA                             
##  [668,] "Lactobacillales"             NA                             
##  [669,] "Sphingobacteriales"          "Lentimicrobiaceae"            
##  [670,] "Bacteroidales"               "Tannerellaceae"               
##  [671,] "Actinomycetales"             "Actinomycetaceae"             
##  [672,] "Micrococcales"               "Micrococcaceae"               
##  [673,] "Saccharimonadales"           NA                             
##  [674,] "Bacteroidales"               "Prevotellaceae"               
##  [675,] "Lactobacillales"             "Streptococcaceae"             
##  [676,] "Pasteurellales"              "Pasteurellaceae"              
##  [677,] "Bacteroidales"               "Prevotellaceae"               
##  [678,] "Selenomonadales"             "Veillonellaceae"              
##  [679,] "Bacteroidales"               "Prevotellaceae"               
##  [680,] NA                            NA                             
##  [681,] "Flavobacteriales"            "Flavobacteriaceae"            
##  [682,] "Pasteurellales"              "Pasteurellaceae"              
##  [683,] "Selenomonadales"             "Veillonellaceae"              
##  [684,] "Bacteroidales"               "Porphyromonadaceae"           
##  [685,] "Bacteroidales"               "Prevotellaceae"               
##  [686,] "Bacteroidales"               "Prevotellaceae"               
##  [687,] "Saccharimonadales"           "Saccharimonadaceae"           
##  [688,] "Lactobacillales"             "Streptococcaceae"             
##  [689,] "Selenomonadales"             "Veillonellaceae"              
##  [690,] "Selenomonadales"             "Veillonellaceae"              
##  [691,] "Bacteroidales"               "Porphyromonadaceae"           
##  [692,] "Saccharimonadales"           "Saccharimonadaceae"           
##  [693,] NA                            NA                             
##  [694,] "Bacteroidales"               "Prevotellaceae"               
##  [695,] "Selenomonadales"             "Veillonellaceae"              
##  [696,] "Lactobacillales"             "Streptococcaceae"             
##  [697,] "Selenomonadales"             "Veillonellaceae"              
##  [698,] "Bacteroidales"               "Prevotellaceae"               
##  [699,] "Bacteroidales"               "Prevotellaceae"               
##  [700,] "Spirochaetales"              "Spirochaetaceae"              
##  [701,] "Bacteroidales"               "Prevotellaceae"               
##  [702,] "Selenomonadales"             "Veillonellaceae"              
##  [703,] "Fusobacteriales"             "Leptotrichiaceae"             
##  [704,] "Campylobacterales"           "P6-b51"                       
##  [705,] "Bacteroidales"               "Prevotellaceae"               
##  [706,] "Saccharimonadales"           "Saccharimonadaceae"           
##  [707,] "Actinomycetales"             "Actinomycetaceae"             
##  [708,] "Bacteroidales"               "Prevotellaceae"               
##  [709,] "Bacteroidales"               "Prevotellaceae"               
##  [710,] "Pasteurellales"              "Pasteurellaceae"              
##  [711,] "Bacteroidales"               "Prevotellaceae"               
##  [712,] "Selenomonadales"             "Veillonellaceae"              
##  [713,] "Bacteroidales"               "Porphyromonadaceae"           
##  [714,] "Bacteroidales"               "Prevotellaceae"               
##  [715,] "Betaproteobacteriales"       "Neisseriaceae"                
##  [716,] "Saccharimonadales"           NA                             
##  [717,] "Lactobacillales"             "Streptococcaceae"             
##  [718,] "Spirochaetales"              "Spirochaetaceae"              
##  [719,] "Bacteroidales"               "Prevotellaceae"               
##  [720,] "Pasteurellales"              "Pasteurellaceae"              
##  [721,] "Bacteroidales"               "Prevotellaceae"               
##  [722,] "Bacteroidales"               "Prevotellaceae"               
##  [723,] "Flavobacteriales"            "Flavobacteriaceae"            
##  [724,] "Betaproteobacteriales"       "Burkholderiaceae"             
##  [725,] "Pasteurellales"              "Pasteurellaceae"              
##  [726,] "Betaproteobacteriales"       "Neisseriaceae"                
##  [727,] "Pasteurellales"              "Pasteurellaceae"              
##  [728,] "Selenomonadales"             "Veillonellaceae"              
##  [729,] "Selenomonadales"             "Veillonellaceae"              
##  [730,] "Flavobacteriales"            "Flavobacteriaceae"            
##  [731,] "Betaproteobacteriales"       "Neisseriaceae"                
##  [732,] "Fusobacteriales"             "Fusobacteriaceae"             
##  [733,] "Pasteurellales"              "Pasteurellaceae"              
##  [734,] "Selenomonadales"             "Veillonellaceae"              
##  [735,] "Betaproteobacteriales"       "Neisseriaceae"                
##  [736,] "Flavobacteriales"            "Flavobacteriaceae"            
##  [737,] "Betaproteobacteriales"       "Neisseriaceae"                
##  [738,] "Actinomycetales"             "Actinomycetaceae"             
##  [739,] "Betaproteobacteriales"       "Neisseriaceae"                
##  [740,] "Flavobacteriales"            "Weeksellaceae"                
##  [741,] "Fusobacteriales"             "Leptotrichiaceae"             
##  [742,] "Lactobacillales"             "Streptococcaceae"             
##  [743,] "Micrococcales"               "Micrococcaceae"               
##  [744,] "Lactobacillales"             "Streptococcaceae"             
##  [745,] "Selenomonadales"             "Veillonellaceae"              
##  [746,] "Selenomonadales"             "Veillonellaceae"              
##  [747,] "Bacteroidales"               "Prevotellaceae"               
##  [748,] "Saccharimonadales"           NA                             
##  [749,] "Fusobacteriales"             "Fusobacteriaceae"             
##  [750,] "Pasteurellales"              "Pasteurellaceae"              
##  [751,] "Bacteroidales"               "Prevotellaceae"               
##  [752,] "Bacteroidales"               "Prevotellaceae"               
##  [753,] "Bacteroidales"               "Prevotellaceae"               
##  [754,] "Flavobacteriales"            "Flavobacteriaceae"            
##  [755,] "Selenomonadales"             "Veillonellaceae"              
##  [756,] "Selenomonadales"             "Veillonellaceae"              
##  [757,] "Bacteroidales"               "Prevotellaceae"               
##  [758,] "Fusobacteriales"             "Leptotrichiaceae"             
##  [759,] "Selenomonadales"             "Veillonellaceae"              
##  [760,] "Bacteroidales"               "Porphyromonadaceae"           
##  [761,] "Selenomonadales"             "Veillonellaceae"              
##  [762,] "Bacteroidales"               "Porphyromonadaceae"           
##  [763,] "Bacteroidales"               "Prevotellaceae"               
##  [764,] "Selenomonadales"             "Veillonellaceae"              
##  [765,] "Fusobacteriales"             "Fusobacteriaceae"             
##  [766,] "Selenomonadales"             "Veillonellaceae"              
##  [767,] "Fusobacteriales"             "Fusobacteriaceae"             
##  [768,] "Fusobacteriales"             "Fusobacteriaceae"             
##  [769,] "Bacteroidales"               "Prevotellaceae"               
##  [770,] "Bacteroidales"               "Prevotellaceae"               
##  [771,] "Lactobacillales"             "Streptococcaceae"             
##  [772,] "Fusobacteriales"             "Leptotrichiaceae"             
##  [773,] "Bacteroidales"               "Prevotellaceae"               
##  [774,] "Lactobacillales"             "Streptococcaceae"             
##  [775,] "Actinomycetales"             "Actinomycetaceae"             
##  [776,] "Selenomonadales"             "Veillonellaceae"              
##  [777,] "Selenomonadales"             "Veillonellaceae"              
##  [778,] "Bacteroidales"               "Prevotellaceae"               
##  [779,] "Betaproteobacteriales"       "Neisseriaceae"                
##  [780,] "Clostridiales"               "Lachnospiraceae"              
##  [781,] "Bacteroidales"               "Prevotellaceae"               
##  [782,] "Pasteurellales"              "Pasteurellaceae"              
##  [783,] "Bacteroidales"               "Prevotellaceae"               
##  [784,] "Pasteurellales"              "Pasteurellaceae"              
##  [785,] "Clostridiales"               "Family_XI"                    
##  [786,] "Lactobacillales"             "Aerococcaceae"                
##  [787,] "Bacteroidales"               "Prevotellaceae"               
##  [788,] "Bacteroidales"               "Prevotellaceae"               
##  [789,] "Pasteurellales"              "Pasteurellaceae"              
##  [790,] "Clostridiales"               "Lachnospiraceae"              
##  [791,] "Selenomonadales"             "Veillonellaceae"              
##  [792,] "Clostridiales"               "Lachnospiraceae"              
##  [793,] "Selenomonadales"             "Veillonellaceae"              
##  [794,] "Fusobacteriales"             "Leptotrichiaceae"             
##  [795,] "Campylobacterales"           "Campylobacteraceae"           
##  [796,] "Bacteroidales"               "Prevotellaceae"               
##  [797,] NA                            NA                             
##  [798,] "Selenomonadales"             "Veillonellaceae"              
##  [799,] "Selenomonadales"             "Veillonellaceae"              
##  [800,] "Betaproteobacteriales"       "Neisseriaceae"                
##  [801,] "Bacteroidales"               "Prevotellaceae"               
##  [802,] "Fusobacteriales"             "Leptotrichiaceae"             
##  [803,] "Bacteroidales"               "Porphyromonadaceae"           
##  [804,] "Bacteroidales"               "Prevotellaceae"               
##  [805,] "Micrococcales"               NA                             
##  [806,] "Clostridiales"               "Lachnospiraceae"              
##  [807,] "Pasteurellales"              "Pasteurellaceae"              
##  [808,] "Lactobacillales"             "Streptococcaceae"             
##  [809,] "Bacteroidales"               "Prevotellaceae"               
##  [810,] "Saccharimonadales"           "Saccharimonadaceae"           
##  [811,] "Bacteroidales"               "Prevotellaceae"               
##  [812,] "Actinomycetales"             "Actinomycetaceae"             
##  [813,] "Fusobacteriales"             "Leptotrichiaceae"             
##  [814,] "Selenomonadales"             "Veillonellaceae"              
##  [815,] "Bacteroidales"               "Prevotellaceae"               
##  [816,] "Pasteurellales"              "Pasteurellaceae"              
##  [817,] "Bacteroidales"               "Porphyromonadaceae"           
##  [818,] "Selenomonadales"             "Veillonellaceae"              
##  [819,] "Bacteroidales"               "Prevotellaceae"               
##  [820,] "Saccharimonadales"           "Saccharimonadaceae"           
##  [821,] "Betaproteobacteriales"       "Neisseriaceae"                
##  [822,] "Fusobacteriales"             "Leptotrichiaceae"             
##  [823,] "Lactobacillales"             "Carnobacteriaceae"            
##  [824,] "Bacteroidales"               "Prevotellaceae"               
##  [825,] "Fusobacteriales"             "Leptotrichiaceae"             
##  [826,] "Bacteroidales"               "Prevotellaceae"               
##  [827,] "Bacteroidales"               "Prevotellaceae"               
##  [828,] "Betaproteobacteriales"       NA                             
##  [829,] "Bacteroidales"               "Prevotellaceae"               
##  [830,] "Pasteurellales"              "Pasteurellaceae"              
##  [831,] "Selenomonadales"             "Veillonellaceae"              
##  [832,] "Bacteroidales"               "Prevotellaceae"               
##  [833,] "Clostridiales"               "Peptostreptococcaceae"        
##  [834,] "Micrococcales"               NA                             
##  [835,] "Selenomonadales"             "Veillonellaceae"              
##  [836,] "Selenomonadales"             "Veillonellaceae"              
##  [837,] "Lactobacillales"             "Carnobacteriaceae"            
##  [838,] "Bacteroidales"               "Prevotellaceae"               
##  [839,] "Campylobacterales"           "Campylobacteraceae"           
##  [840,] "Saccharimonadales"           "Saccharimonadaceae"           
##  [841,] "Selenomonadales"             "Veillonellaceae"              
##  [842,] "Bacteroidales"               "Porphyromonadaceae"           
##  [843,] "Clostridiales"               "Family_XI"                    
##  [844,] "Bacteroidales"               "Porphyromonadaceae"           
##  [845,] NA                            NA                             
##  [846,] "Actinomycetales"             "Actinomycetaceae"             
##  [847,] "Selenomonadales"             "Veillonellaceae"              
##  [848,] "Micrococcales"               "Micrococcaceae"               
##  [849,] "Bacteroidales"               "Prevotellaceae"               
##  [850,] "Pasteurellales"              "Pasteurellaceae"              
##  [851,] "Bacteroidales"               "Prevotellaceae"               
##  [852,] "Lactobacillales"             "Streptococcaceae"             
##  [853,] "Bacteroidales"               "Prevotellaceae"               
##  [854,] "Selenomonadales"             "Veillonellaceae"              
##  [855,] "Bacteroidales"               "Prevotellaceae"               
##  [856,] "Clostridiales"               "Peptostreptococcaceae"        
##  [857,] "Bacteroidales"               "Prevotellaceae"               
##  [858,] "Lactobacillales"             "Streptococcaceae"             
##  [859,] "Bacteroidales"               "Prevotellaceae"               
##  [860,] "Saccharimonadales"           "Saccharimonadaceae"           
##  [861,] "Clostridiales"               "Defluviitaleaceae"            
##  [862,] "Clostridiales"               "Lachnospiraceae"              
##  [863,] "Bacteroidales"               "Prevotellaceae"               
##  [864,] "Bacteroidales"               "Prevotellaceae"               
##  [865,] "Bacteroidales"               "Prevotellaceae"               
##  [866,] "Bacteroidales"               "Porphyromonadaceae"           
##  [867,] "Actinomycetales"             "Actinomycetaceae"             
##  [868,] "Bacteroidales"               "Prevotellaceae"               
##  [869,] "Selenomonadales"             "Veillonellaceae"              
##  [870,] "Actinomycetales"             "Actinomycetaceae"             
##  [871,] "Actinomycetales"             "Actinomycetaceae"             
##  [872,] "Selenomonadales"             "Veillonellaceae"              
##  [873,] "Bacteroidales"               "Porphyromonadaceae"           
##  [874,] "Bacteroidales"               "Porphyromonadaceae"           
##  [875,] "Actinomycetales"             "Actinomycetaceae"             
##  [876,] "Fusobacteriales"             "Leptotrichiaceae"             
##  [877,] "Saccharimonadales"           "Saccharimonadaceae"           
##  [878,] "Selenomonadales"             "Veillonellaceae"              
##  [879,] "Actinomycetales"             "Actinomycetaceae"             
##  [880,] "Selenomonadales"             "Veillonellaceae"              
##  [881,] "Pasteurellales"              "Pasteurellaceae"              
##  [882,] "Clostridiales"               "Lachnospiraceae"              
##  [883,] "Selenomonadales"             "Veillonellaceae"              
##  [884,] "Bacteroidales"               "Prevotellaceae"               
##  [885,] "Bacteroidales"               "Prevotellaceae"               
##  [886,] "Bacteroidales"               "Porphyromonadaceae"           
##  [887,] "Selenomonadales"             "Veillonellaceae"              
##  [888,] "Actinomycetales"             "Actinomycetaceae"             
##  [889,] "Micrococcales"               NA                             
##  [890,] "Clostridiales"               "Lachnospiraceae"              
##  [891,] "Pasteurellales"              "Pasteurellaceae"              
##  [892,] "Lactobacillales"             "Streptococcaceae"             
##  [893,] "Saccharimonadales"           "Saccharimonadaceae"           
##  [894,] "Flavobacteriales"            "Weeksellaceae"                
##  [895,] "Actinomycetales"             "Actinomycetaceae"             
##  [896,] "Bacteroidales"               "Prevotellaceae"               
##  [897,] "Bacteroidales"               "Prevotellaceae"               
##  [898,] "Bacteroidales"               "Prevotellaceae"               
##  [899,] "Saccharimonadales"           "Saccharimonadaceae"           
##  [900,] NA                            NA                             
##  [901,] "Fusobacteriales"             "Fusobacteriaceae"             
##  [902,] "Pasteurellales"              "Pasteurellaceae"              
##  [903,] "Lactobacillales"             "Streptococcaceae"             
##  [904,] "Clostridiales"               "Ruminococcaceae"              
##  [905,] "Micrococcales"               NA                             
##  [906,] "Bacteroidales"               "Prevotellaceae"               
##  [907,] "Lactobacillales"             "Streptococcaceae"             
##  [908,] "Pasteurellales"              "Pasteurellaceae"              
##  [909,] "Selenomonadales"             "Veillonellaceae"              
##  [910,] "Pasteurellales"              "Pasteurellaceae"              
##  [911,] "Selenomonadales"             "Veillonellaceae"              
##  [912,] "Saccharimonadales"           "Saccharimonadaceae"           
##  [913,] "Betaproteobacteriales"       "Neisseriaceae"                
##  [914,] "Pasteurellales"              "Pasteurellaceae"              
##  [915,] "Bifidobacteriales"           "Bifidobacteriaceae"           
##  [916,] "Selenomonadales"             "Veillonellaceae"              
##  [917,] "Bacteroidales"               "Prevotellaceae"               
##  [918,] "Clostridiales"               "Lachnospiraceae"              
##  [919,] "Actinomycetales"             "Actinomycetaceae"             
##  [920,] "Betaproteobacteriales"       "Neisseriaceae"                
##  [921,] "Campylobacterales"           "Campylobacteraceae"           
##  [922,] "Selenomonadales"             "Veillonellaceae"              
##  [923,] "Selenomonadales"             "Veillonellaceae"              
##  [924,] NA                            NA                             
##  [925,] "Fusobacteriales"             "Leptotrichiaceae"             
##  [926,] "Betaproteobacteriales"       "Neisseriaceae"                
##  [927,] "Fusobacteriales"             "Fusobacteriaceae"             
##  [928,] "Fusobacteriales"             "Fusobacteriaceae"             
##  [929,] "Selenomonadales"             "Veillonellaceae"              
##  [930,] "Bacteroidales"               "Prevotellaceae"               
##  [931,] "Fusobacteriales"             "Leptotrichiaceae"             
##  [932,] "Lactobacillales"             "Streptococcaceae"             
##  [933,] NA                            NA                             
##  [934,] "Selenomonadales"             "Veillonellaceae"              
##  [935,] "Bacteroidales"               "Prevotellaceae"               
##  [936,] "Clostridiales"               "Family_XIII"                  
##  [937,] "Fusobacteriales"             "Leptotrichiaceae"             
##  [938,] "Erysipelotrichales"          "Erysipelotrichaceae"          
##  [939,] "Lactobacillales"             "Streptococcaceae"             
##  [940,] "Bacteroidales"               "Prevotellaceae"               
##  [941,] "Lactobacillales"             "Streptococcaceae"             
##  [942,] "Actinomycetales"             "Actinomycetaceae"             
##  [943,] "Campylobacterales"           NA                             
##  [944,] "Spirochaetales"              "Spirochaetaceae"              
##  [945,] "Lactobacillales"             "Streptococcaceae"             
##  [946,] "Actinomycetales"             "Actinomycetaceae"             
##  [947,] "Lactobacillales"             "Streptococcaceae"             
##  [948,] "Fusobacteriales"             "Leptotrichiaceae"             
##  [949,] "Fusobacteriales"             "Leptotrichiaceae"             
##  [950,] "Clostridiales"               "Lachnospiraceae"              
##  [951,] "Fusobacteriales"             "Fusobacteriaceae"             
##  [952,] "Selenomonadales"             "Veillonellaceae"              
##  [953,] NA                            NA                             
##  [954,] "Spirochaetales"              "Spirochaetaceae"              
##  [955,] "Bacteroidales"               "Prevotellaceae"               
##  [956,] "Micrococcales"               "Micrococcaceae"               
##  [957,] "Selenomonadales"             "Veillonellaceae"              
##  [958,] "Saccharimonadales"           "Saccharimonadaceae"           
##  [959,] "Lactobacillales"             "Streptococcaceae"             
##  [960,] "Bacteroidales"               "Porphyromonadaceae"           
##  [961,] "Selenomonadales"             "Veillonellaceae"              
##  [962,] "Lactobacillales"             "Streptococcaceae"             
##  [963,] "Selenomonadales"             "Veillonellaceae"              
##  [964,] "Enterobacteriales"           "Enterobacteriaceae"           
##  [965,] "Fusobacteriales"             "Fusobacteriaceae"             
##  [966,] "Clostridiales"               "Lachnospiraceae"              
##  [967,] "Bacteroidales"               "Prevotellaceae"               
##  [968,] "Fusobacteriales"             "Fusobacteriaceae"             
##  [969,] "Clostridiales"               "Ruminococcaceae"              
##  [970,] "Bacteroidales"               "Prevotellaceae"               
##  [971,] "Fusobacteriales"             "Fusobacteriaceae"             
##  [972,] "Selenomonadales"             "Veillonellaceae"              
##  [973,] "Saccharimonadales"           NA                             
##  [974,] "Selenomonadales"             "Veillonellaceae"              
##  [975,] "Bacteroidales"               "Prevotellaceae"               
##  [976,] "Clostridiales"               "Lachnospiraceae"              
##  [977,] "Clostridiales"               "Lachnospiraceae"              
##  [978,] "Selenomonadales"             "Veillonellaceae"              
##  [979,] "Actinomycetales"             "Actinomycetaceae"             
##  [980,] "Actinomycetales"             "Actinomycetaceae"             
##  [981,] "Fusobacteriales"             "Fusobacteriaceae"             
##  [982,] "Bacteroidales"               "Prevotellaceae"               
##  [983,] "Fusobacteriales"             "Leptotrichiaceae"             
##  [984,] "Actinomycetales"             "Actinomycetaceae"             
##  [985,] "Campylobacterales"           NA                             
##  [986,] "Selenomonadales"             "Veillonellaceae"              
##  [987,] "Actinomycetales"             "Actinomycetaceae"             
##  [988,] "Pasteurellales"              "Pasteurellaceae"              
##  [989,] "Pasteurellales"              "Pasteurellaceae"              
##  [990,] "Selenomonadales"             "Veillonellaceae"              
##  [991,] "Bacteroidales"               "Prevotellaceae"               
##  [992,] "Micrococcales"               NA                             
##  [993,] "Actinomycetales"             "Actinomycetaceae"             
##  [994,] "Selenomonadales"             "Veillonellaceae"              
##  [995,] "Selenomonadales"             "Veillonellaceae"              
##  [996,] "Fusobacteriales"             "Leptotrichiaceae"             
##  [997,] "Fusobacteriales"             "Leptotrichiaceae"             
##  [998,] "Saccharimonadales"           "Saccharimonadaceae"           
##  [999,] "Bacteroidales"               "Prevotellaceae"               
## [1000,] "Selenomonadales"             "Veillonellaceae"              
## [1001,] "Selenomonadales"             "Veillonellaceae"              
## [1002,] "Selenomonadales"             "Veillonellaceae"              
## [1003,] "Betaproteobacteriales"       "Neisseriaceae"                
## [1004,] "Bacteroidales"               "Porphyromonadaceae"           
## [1005,] "Bacteroidales"               "Prevotellaceae"               
## [1006,] "Bacteroidales"               "Prevotellaceae"               
## [1007,] "Clostridiales"               "Lachnospiraceae"              
## [1008,] "Bacteroidales"               "Prevotellaceae"               
## [1009,] "Bacteroidales"               "Prevotellaceae"               
## [1010,] "Clostridiales"               "Family_XI"                    
## [1011,] "Selenomonadales"             "Veillonellaceae"              
## [1012,] "Saccharimonadales"           "Saccharimonadaceae"           
## [1013,] "Betaproteobacteriales"       "Neisseriaceae"                
## [1014,] "Saccharimonadales"           NA                             
## [1015,] "Clostridiales"               "Lachnospiraceae"              
## [1016,] "Betaproteobacteriales"       "Neisseriaceae"                
## [1017,] "Saccharimonadales"           "Saccharimonadaceae"           
## [1018,] "Bacteroidales"               "Prevotellaceae"               
## [1019,] "Fusobacteriales"             "Fusobacteriaceae"             
## [1020,] "Enterobacteriales"           "Enterobacteriaceae"           
## [1021,] "Bacteroidales"               "Prevotellaceae"               
## [1022,] "Fusobacteriales"             "Leptotrichiaceae"             
## [1023,] "Spirochaetales"              "Spirochaetaceae"              
## [1024,] "Lactobacillales"             "Carnobacteriaceae"            
## [1025,] "Lactobacillales"             "Carnobacteriaceae"            
## [1026,] "Corynebacteriales"           "Corynebacteriaceae"           
## [1027,] "Fusobacteriales"             "Leptotrichiaceae"             
## [1028,] "Bacteroidales"               "Prevotellaceae"               
## [1029,] "Bacteroidales"               "Prevotellaceae"               
## [1030,] "Betaproteobacteriales"       "Neisseriaceae"                
## [1031,] "Fusobacteriales"             "Fusobacteriaceae"             
## [1032,] "Fusobacteriales"             "Leptotrichiaceae"             
## [1033,] "Fusobacteriales"             "Fusobacteriaceae"             
## [1034,] "Selenomonadales"             "Veillonellaceae"              
## [1035,] "Bacteroidales"               "Prevotellaceae"               
## [1036,] "Selenomonadales"             "Veillonellaceae"              
## [1037,] "Selenomonadales"             "Veillonellaceae"              
## [1038,] "Clostridiales"               "Lachnospiraceae"              
## [1039,] "Bacteroidales"               "Prevotellaceae"               
## [1040,] "Selenomonadales"             "Veillonellaceae"              
## [1041,] "Bacteroidales"               "Prevotellaceae"               
## [1042,] "Clostridiales"               "Lachnospiraceae"              
## [1043,] "Fusobacteriales"             "Leptotrichiaceae"             
## [1044,] "Selenomonadales"             "Veillonellaceae"              
## [1045,] "Actinomycetales"             "Actinomycetaceae"             
## [1046,] "Bacteroidales"               "Prevotellaceae"               
## [1047,] "Selenomonadales"             "Veillonellaceae"              
## [1048,] "Bacteroidales"               "Prevotellaceae"               
## [1049,] "Selenomonadales"             "Veillonellaceae"              
## [1050,] "Fusobacteriales"             NA                             
## [1051,] "Clostridiales"               "Lachnospiraceae"              
## [1052,] "Lactobacillales"             "Carnobacteriaceae"            
## [1053,] "Bacteroidales"               "Prevotellaceae"               
## [1054,] "Pasteurellales"              "Pasteurellaceae"              
## [1055,] "Bacteroidales"               "Prevotellaceae"               
## [1056,] "Lactobacillales"             "Streptococcaceae"             
## [1057,] "Pasteurellales"              "Pasteurellaceae"              
## [1058,] "Bacteroidales"               "Prevotellaceae"               
## [1059,] "Clostridiales"               "Lachnospiraceae"              
## [1060,] "Bacteroidales"               "Prevotellaceae"               
## [1061,] "Selenomonadales"             "Veillonellaceae"              
## [1062,] "Lactobacillales"             "Streptococcaceae"             
## [1063,] "Bacteroidales"               "Prevotellaceae"               
## [1064,] "Fusobacteriales"             "Fusobacteriaceae"             
## [1065,] "Selenomonadales"             "Veillonellaceae"              
## [1066,] "Fusobacteriales"             "Leptotrichiaceae"             
## [1067,] "Selenomonadales"             "Veillonellaceae"              
## [1068,] "Pasteurellales"              "Pasteurellaceae"              
## [1069,] "Fusobacteriales"             "Leptotrichiaceae"             
## [1070,] "Selenomonadales"             "Veillonellaceae"              
## [1071,] "Pasteurellales"              "Pasteurellaceae"              
## [1072,] "Bacteroidales"               "Porphyromonadaceae"           
## [1073,] "Bacteroidales"               "Prevotellaceae"               
## [1074,] "Coriobacteriales"            "Atopobiaceae"                 
## [1075,] "Flavobacteriales"            "Flavobacteriaceae"            
## [1076,] "Betaproteobacteriales"       "Neisseriaceae"                
## [1077,] NA                            NA                             
## [1078,] "Bacteroidales"               "Prevotellaceae"               
## [1079,] "Selenomonadales"             "Veillonellaceae"              
## [1080,] "Clostridiales"               "Lachnospiraceae"              
## [1081,] "Bacteroidales"               "Prevotellaceae"               
## [1082,] "Saccharimonadales"           "Saccharimonadaceae"           
## [1083,] "Flavobacteriales"            "Flavobacteriaceae"            
## [1084,] "Selenomonadales"             "Veillonellaceae"              
## [1085,] "Selenomonadales"             "Veillonellaceae"              
## [1086,] NA                            NA                             
## [1087,] "Lactobacillales"             "Streptococcaceae"             
## [1088,] "Bacteroidales"               "Prevotellaceae"               
## [1089,] "Selenomonadales"             "Veillonellaceae"              
## [1090,] "Bacteroidales"               "Prevotellaceae"               
## [1091,] "Coriobacteriales"            "Atopobiaceae"                 
## [1092,] "Bacteroidales"               "Prevotellaceae"               
## [1093,] "Bacteroidales"               "Prevotellaceae"               
## [1094,] "Bacteroidales"               "Prevotellaceae"               
## [1095,] "Flavobacteriales"            "Flavobacteriaceae"            
## [1096,] "Selenomonadales"             "Veillonellaceae"              
## [1097,] "Bacteroidales"               "Porphyromonadaceae"           
## [1098,] "Lactobacillales"             "Streptococcaceae"             
## [1099,] "Actinomycetales"             "Actinomycetaceae"             
## [1100,] "Bacteroidales"               "Prevotellaceae"               
## [1101,] "Saccharimonadales"           "Saccharimonadaceae"           
## [1102,] "Selenomonadales"             "Veillonellaceae"              
## [1103,] "Selenomonadales"             "Veillonellaceae"              
## [1104,] "Bacteroidales"               "Prevotellaceae"               
## [1105,] "Bacteroidales"               "Prevotellaceae"               
## [1106,] "Selenomonadales"             "Veillonellaceae"              
## [1107,] "Bacteroidales"               "Prevotellaceae"               
## [1108,] "Fusobacteriales"             "Fusobacteriaceae"             
## [1109,] "Flavobacteriales"            "Flavobacteriaceae"            
## [1110,] "Selenomonadales"             "Veillonellaceae"              
## [1111,] "Bacteroidales"               "Prevotellaceae"               
## [1112,] "Fusobacteriales"             "Leptotrichiaceae"             
## [1113,] "Bacteroidales"               "Prevotellaceae"               
## [1114,] "Clostridiales"               "Peptostreptococcaceae"        
## [1115,] "Saccharimonadales"           "Saccharimonadaceae"           
## [1116,] "Saccharimonadales"           "Saccharimonadaceae"           
## [1117,] "Flavobacteriales"            "Flavobacteriaceae"            
## [1118,] "Bacteroidales"               "Prevotellaceae"               
## [1119,] "Micrococcales"               "Micrococcaceae"               
## [1120,] "Saccharimonadales"           "Saccharimonadaceae"           
## [1121,] "Selenomonadales"             "Veillonellaceae"              
## [1122,] "Selenomonadales"             "Veillonellaceae"              
## [1123,] "Bacteroidales"               "Prevotellaceae"               
## [1124,] "Campylobacterales"           "Campylobacteraceae"           
## [1125,] "Actinomycetales"             "Actinomycetaceae"             
## [1126,] "Chloroplast"                 NA                             
## [1127,] "Bacteroidales"               "Porphyromonadaceae"           
## [1128,] NA                            NA                             
## [1129,] "Fusobacteriales"             "Fusobacteriaceae"             
## [1130,] "Pasteurellales"              "Pasteurellaceae"              
## [1131,] "Clostridiales"               "Lachnospiraceae"              
## [1132,] "Bacteroidales"               "Prevotellaceae"               
## [1133,] "Bacteroidales"               "Prevotellaceae"               
## [1134,] "Selenomonadales"             "Veillonellaceae"              
## [1135,] "Pasteurellales"              "Pasteurellaceae"              
## [1136,] "Pasteurellales"              "Pasteurellaceae"              
## [1137,] "Bacteroidales"               "Prevotellaceae"               
## [1138,] "Bacteroidales"               "Prevotellaceae"               
## [1139,] "Betaproteobacteriales"       "Neisseriaceae"                
## [1140,] "Lactobacillales"             NA                             
## [1141,] "Pseudomonadales"             "Moraxellaceae"                
## [1142,] "Bacteroidales"               "Prevotellaceae"               
## [1143,] "Clostridiales"               "Lachnospiraceae"              
## [1144,] "Lactobacillales"             "Streptococcaceae"             
## [1145,] "Actinomycetales"             "Actinomycetaceae"             
## [1146,] "Lactobacillales"             "Streptococcaceae"             
## [1147,] "Bacteroidales"               "Prevotellaceae"               
## [1148,] "Selenomonadales"             "Veillonellaceae"              
## [1149,] "Clostridiales"               "Lachnospiraceae"              
## [1150,] "Micrococcales"               NA                             
## [1151,] "Fusobacteriales"             "Leptotrichiaceae"             
## [1152,] "Pasteurellales"              "Pasteurellaceae"              
## [1153,] "Lactobacillales"             "Streptococcaceae"             
## [1154,] "Betaproteobacteriales"       "Neisseriaceae"                
## [1155,] "Bacteroidales"               "Prevotellaceae"               
## [1156,] "Pasteurellales"              "Pasteurellaceae"              
## [1157,] "Lactobacillales"             "Streptococcaceae"             
## [1158,] "Fusobacteriales"             "Fusobacteriaceae"             
## [1159,] "Clostridiales"               "Lachnospiraceae"              
## [1160,] "Bacteroidales"               "Prevotellaceae"               
## [1161,] "Bacteroidales"               "Prevotellaceae"               
## [1162,] "Lactobacillales"             "Streptococcaceae"             
## [1163,] "Mycoplasmatales"             "Mycoplasmataceae"             
## [1164,] "Pasteurellales"              "Pasteurellaceae"              
## [1165,] "Absconditabacteriales_(SR1)" NA                             
## [1166,] "Selenomonadales"             "Veillonellaceae"              
## [1167,] "Betaproteobacteriales"       "Neisseriaceae"                
## [1168,] "Selenomonadales"             "Veillonellaceae"              
## [1169,] "Selenomonadales"             "Veillonellaceae"              
## [1170,] "Lactobacillales"             "Streptococcaceae"             
## [1171,] "Bacteroidales"               "Prevotellaceae"               
## [1172,] "Saccharimonadales"           "Saccharimonadaceae"           
## [1173,] "Bacteroidales"               "Prevotellaceae"               
## [1174,] "Selenomonadales"             "Veillonellaceae"              
## [1175,] "Selenomonadales"             "Veillonellaceae"              
## [1176,] "Selenomonadales"             "Veillonellaceae"              
## [1177,] "Bacteroidales"               "Porphyromonadaceae"           
## [1178,] "Fusobacteriales"             "Leptotrichiaceae"             
## [1179,] "Betaproteobacteriales"       "Neisseriaceae"                
## [1180,] "Bacteroidales"               "Prevotellaceae"               
## [1181,] "Clostridiales"               "Lachnospiraceae"              
## [1182,] "Clostridiales"               "Peptostreptococcaceae"        
## [1183,] "Lactobacillales"             "Streptococcaceae"             
## [1184,] "Bacteroidales"               "Prevotellaceae"               
## [1185,] "Pasteurellales"              "Pasteurellaceae"              
## [1186,] "Pasteurellales"              "Pasteurellaceae"              
## [1187,] "Bacteroidales"               "Prevotellaceae"               
## [1188,] "Selenomonadales"             "Veillonellaceae"              
## [1189,] "Bacteroidales"               "Prevotellaceae"               
## [1190,] "Bacteroidales"               "Prevotellaceae"               
## [1191,] "Actinomycetales"             "Actinomycetaceae"             
## [1192,] "Fusobacteriales"             "Leptotrichiaceae"             
## [1193,] "Actinomycetales"             "Actinomycetaceae"             
## [1194,] "Pasteurellales"              "Pasteurellaceae"              
## [1195,] "Bacteroidales"               "Porphyromonadaceae"           
## [1196,] "Flavobacteriales"            "Flavobacteriaceae"            
## [1197,] "Selenomonadales"             "Veillonellaceae"              
## [1198,] "Clostridiales"               "Lachnospiraceae"              
## [1199,] "Lactobacillales"             "Streptococcaceae"             
## [1200,] "Bacteroidales"               "Prevotellaceae"               
## [1201,] "Selenomonadales"             "Veillonellaceae"              
## [1202,] "Fusobacteriales"             "Fusobacteriaceae"             
## [1203,] "Actinomycetales"             "Actinomycetaceae"             
## [1204,] "Bacteroidales"               "Prevotellaceae"               
## [1205,] "Synergistales"               "Synergistaceae"               
## [1206,] "Campylobacterales"           NA                             
## [1207,] "Betaproteobacteriales"       "Burkholderiaceae"             
## [1208,] "Selenomonadales"             "Veillonellaceae"              
## [1209,] "Rhizobiales"                 "Rhizobiaceae"                 
## [1210,] "Selenomonadales"             "Veillonellaceae"              
## [1211,] "Fusobacteriales"             "Fusobacteriaceae"             
## [1212,] "Saccharimonadales"           "Saccharimonadaceae"           
## [1213,] "Selenomonadales"             "Veillonellaceae"              
## [1214,] "Fusobacteriales"             "Fusobacteriaceae"             
## [1215,] "Saccharimonadales"           "Saccharimonadaceae"           
## [1216,] "Clostridiales"               "Lachnospiraceae"              
## [1217,] "Spirochaetales"              "Spirochaetaceae"              
## [1218,] "Bacteroidales"               "Prevotellaceae"               
## [1219,] "Clostridiales"               "Peptostreptococcaceae"        
## [1220,] "Clostridiales"               "Peptostreptococcaceae"        
## [1221,] "Selenomonadales"             "Veillonellaceae"              
## [1222,] "Actinomycetales"             "Actinomycetaceae"             
## [1223,] "Bacteroidales"               "Prevotellaceae"               
## [1224,] NA                            NA                             
## [1225,] "Selenomonadales"             "Veillonellaceae"              
## [1226,] NA                            NA                             
## [1227,] "Selenomonadales"             "Veillonellaceae"              
## [1228,] "Betaproteobacteriales"       "Burkholderiaceae"             
## [1229,] "Bacteroidales"               "Prevotellaceae"               
## [1230,] "Selenomonadales"             "Veillonellaceae"              
## [1231,] "Bacteroidales"               "Porphyromonadaceae"           
## [1232,] "Bacteroidales"               "Prevotellaceae"               
## [1233,] "Flavobacteriales"            "Weeksellaceae"                
## [1234,] "Selenomonadales"             "Veillonellaceae"              
## [1235,] "Bacteroidales"               "Porphyromonadaceae"           
## [1236,] "Clostridiales"               "Lachnospiraceae"              
## [1237,] "Fusobacteriales"             "Fusobacteriaceae"             
## [1238,] "Micrococcales"               "Micrococcaceae"               
## [1239,] "Selenomonadales"             "Veillonellaceae"              
## [1240,] "Saccharimonadales"           "Saccharimonadaceae"           
## [1241,] "Bacteroidales"               "Porphyromonadaceae"           
## [1242,] "Bacteroidales"               "Porphyromonadaceae"           
## [1243,] "Clostridiales"               "Peptostreptococcaceae"        
## [1244,] "Selenomonadales"             "Veillonellaceae"              
## [1245,] "Selenomonadales"             "Veillonellaceae"              
## [1246,] "Fusobacteriales"             "Leptotrichiaceae"             
## [1247,] NA                            NA                             
## [1248,] "Bacteroidales"               "Prevotellaceae"               
## [1249,] "Selenomonadales"             "Veillonellaceae"              
## [1250,] "Micrococcales"               NA                             
## [1251,] "Fusobacteriales"             NA                             
## [1252,] "Bacteroidales"               "Prevotellaceae"               
## [1253,] "Bacteroidales"               "Prevotellaceae"               
## [1254,] "Micrococcales"               NA                             
## [1255,] "Bacteroidales"               "Prevotellaceae"               
## [1256,] "Bacteroidales"               "Paludibacteraceae"            
## [1257,] "Selenomonadales"             "Veillonellaceae"              
## [1258,] "Bacteroidales"               "Prevotellaceae"               
## [1259,] "Lactobacillales"             "Streptococcaceae"             
## [1260,] "Bacteroidales"               "Prevotellaceae"               
## [1261,] "Betaproteobacteriales"       "Neisseriaceae"                
## [1262,] "Selenomonadales"             "Veillonellaceae"              
## [1263,] "Saccharimonadales"           "Saccharimonadaceae"           
## [1264,] NA                            NA                             
## [1265,] "Actinomycetales"             "Actinomycetaceae"             
## [1266,] "Selenomonadales"             "Veillonellaceae"              
## [1267,] "Coriobacteriales"            "Atopobiaceae"                 
## [1268,] "Bacteroidales"               "Prevotellaceae"               
## [1269,] "Clostridiales"               "Family_XIII"                  
## [1270,] "Bacteroidales"               "Prevotellaceae"               
## [1271,] "Selenomonadales"             "Veillonellaceae"              
## [1272,] "Selenomonadales"             "Veillonellaceae"              
## [1273,] "Fusobacteriales"             "Fusobacteriaceae"             
## [1274,] "Bacteroidales"               "Prevotellaceae"               
## [1275,] "Spirochaetales"              "Spirochaetaceae"              
## [1276,] "Bacteroidales"               "Prevotellaceae"               
## [1277,] "Selenomonadales"             "Veillonellaceae"              
## [1278,] "Selenomonadales"             "Veillonellaceae"              
## [1279,] "Selenomonadales"             "Veillonellaceae"              
## [1280,] "Pasteurellales"              "Pasteurellaceae"              
## [1281,] "Pasteurellales"              "Pasteurellaceae"              
## [1282,] "Bacteroidales"               "Prevotellaceae"               
## [1283,] "Saccharimonadales"           "Saccharimonadaceae"           
## [1284,] "Selenomonadales"             "Veillonellaceae"              
## [1285,] "Fusobacteriales"             "Fusobacteriaceae"             
## [1286,] "Corynebacteriales"           "Corynebacteriaceae"           
## [1287,] "Selenomonadales"             "Veillonellaceae"              
## [1288,] "Clostridiales"               "Lachnospiraceae"              
## [1289,] "Mycoplasmatales"             "Mycoplasmataceae"             
## [1290,] "Fusobacteriales"             "Fusobacteriaceae"             
## [1291,] "Bacteroidales"               "Prevotellaceae"               
## [1292,] "Clostridiales"               "Ruminococcaceae"              
## [1293,] "Clostridiales"               "Lachnospiraceae"              
## [1294,] "Fusobacteriales"             "Fusobacteriaceae"             
## [1295,] "Betaproteobacteriales"       "Burkholderiaceae"             
## [1296,] "Bacteroidales"               "Prevotellaceae"               
## [1297,] "Clostridiales"               "Lachnospiraceae"              
## [1298,] "Flavobacteriales"            "Weeksellaceae"                
## [1299,] "Bacteroidales"               "Prevotellaceae"               
## [1300,] "Bacteroidales"               "Porphyromonadaceae"           
## [1301,] "Clostridiales"               "Lachnospiraceae"              
## [1302,] "Bacteroidales"               "Prevotellaceae"               
## [1303,] "Selenomonadales"             "Veillonellaceae"              
## [1304,] "Bacteroidales"               "Prevotellaceae"               
## [1305,] "Actinomycetales"             "Actinomycetaceae"             
## [1306,] "Lactobacillales"             "Streptococcaceae"             
## [1307,] "Betaproteobacteriales"       "Neisseriaceae"                
## [1308,] "Betaproteobacteriales"       "Neisseriaceae"                
## [1309,] "Spirochaetales"              "Spirochaetaceae"              
## [1310,] "Pasteurellales"              "Pasteurellaceae"              
## [1311,] "Pasteurellales"              "Pasteurellaceae"              
## [1312,] "Clostridiales"               "Family_XIII"                  
## [1313,] "Bacteroidales"               "Prevotellaceae"               
## [1314,] "Clostridiales"               "Lachnospiraceae"              
## [1315,] "Bacteroidales"               "Prevotellaceae"               
## [1316,] "Sphingobacteriales"          "Lentimicrobiaceae"            
## [1317,] "Campylobacterales"           NA                             
## [1318,] "Selenomonadales"             "Veillonellaceae"              
## [1319,] "Selenomonadales"             "Veillonellaceae"              
## [1320,] "Lactobacillales"             "Streptococcaceae"             
## [1321,] "Selenomonadales"             "Veillonellaceae"              
## [1322,] "Campylobacterales"           "Campylobacteraceae"           
## [1323,] "Selenomonadales"             "Veillonellaceae"              
## [1324,] "Lactobacillales"             "Streptococcaceae"             
## [1325,] "Bacteroidales"               "Prevotellaceae"               
## [1326,] "Pasteurellales"              "Pasteurellaceae"              
## [1327,] "Lactobacillales"             "Streptococcaceae"             
## [1328,] "Selenomonadales"             "Veillonellaceae"              
## [1329,] "Bacteroidales"               "Prevotellaceae"               
## [1330,] "Bacteroidales"               "Porphyromonadaceae"           
## [1331,] "Clostridiales"               "Lachnospiraceae"              
## [1332,] "Bacteroidales"               "Prevotellaceae"               
## [1333,] "Selenomonadales"             "Veillonellaceae"              
## [1334,] "Bacteroidales"               "Prevotellaceae"               
## [1335,] "Fusobacteriales"             "Fusobacteriaceae"             
## [1336,] "Fusobacteriales"             "Leptotrichiaceae"             
## [1337,] "Selenomonadales"             "Veillonellaceae"              
## [1338,] "Clostridiales"               "Lachnospiraceae"              
## [1339,] "Pasteurellales"              "Pasteurellaceae"              
## [1340,] "Selenomonadales"             "Veillonellaceae"              
## [1341,] NA                            NA                             
## [1342,] "Fusobacteriales"             "Leptotrichiaceae"             
## [1343,] "Bacteroidales"               "Prevotellaceae"               
## [1344,] "Saccharimonadales"           "Saccharimonadaceae"           
## [1345,] "Flavobacteriales"            "Flavobacteriaceae"            
## [1346,] "Saccharimonadales"           NA                             
## [1347,] "Bacteroidales"               "Prevotellaceae"               
## [1348,] "Bacteroidales"               "Prevotellaceae"               
## [1349,] "Betaproteobacteriales"       "Neisseriaceae"                
## [1350,] "Micrococcales"               NA                             
## [1351,] "Actinomycetales"             "Actinomycetaceae"             
## [1352,] "Actinomycetales"             "Actinomycetaceae"             
## [1353,] "Bacteroidales"               "Porphyromonadaceae"           
## [1354,] "Selenomonadales"             "Veillonellaceae"              
## [1355,] "Campylobacterales"           NA                             
## [1356,] NA                            NA                             
## [1357,] "Bacteroidales"               "Prevotellaceae"               
## [1358,] "Acholeplasmatales"           "Acholeplasmataceae"           
## [1359,] "Selenomonadales"             "Veillonellaceae"              
## [1360,] "Bacteroidales"               "Prevotellaceae"               
## [1361,] "Betaproteobacteriales"       "Neisseriaceae"                
## [1362,] NA                            NA                             
## [1363,] NA                            NA                             
## [1364,] "Micrococcales"               NA                             
## [1365,] "Bacteroidales"               "Prevotellaceae"               
## [1366,] "Bacteroidales"               "Prevotellaceae"               
## [1367,] "Bacteroidales"               "Porphyromonadaceae"           
## [1368,] "Pasteurellales"              "Pasteurellaceae"              
## [1369,] "Fusobacteriales"             "Fusobacteriaceae"             
## [1370,] "Clostridiales"               "Ruminococcaceae"              
## [1371,] "Bacteroidales"               "Prevotellaceae"               
## [1372,] "Fusobacteriales"             "Leptotrichiaceae"             
## [1373,] "Bacillales"                  "Family_XI"                    
## [1374,] "Betaproteobacteriales"       "Neisseriaceae"                
## [1375,] "Fusobacteriales"             "Fusobacteriaceae"             
## [1376,] "Betaproteobacteriales"       "Neisseriaceae"                
## [1377,] "Selenomonadales"             "Veillonellaceae"              
## [1378,] "Bacteroidales"               "Prevotellaceae"               
## [1379,] "Bacteroidales"               "Prevotellaceae"               
## [1380,] NA                            NA                             
## [1381,] "Clostridiales"               "Lachnospiraceae"              
## [1382,] "Bacteroidales"               "Prevotellaceae"               
## [1383,] "Selenomonadales"             "Veillonellaceae"              
## [1384,] "Lactobacillales"             "Streptococcaceae"             
## [1385,] "Selenomonadales"             "Veillonellaceae"              
## [1386,] "Lactobacillales"             "Streptococcaceae"             
## [1387,] "Fusobacteriales"             "Fusobacteriaceae"             
## [1388,] "Bacteroidales"               "Prevotellaceae"               
## [1389,] "Selenomonadales"             "Veillonellaceae"              
## [1390,] "Bacteroidales"               "Prevotellaceae"               
## [1391,] "Saccharimonadales"           "Saccharimonadaceae"           
## [1392,] "Bacteroidales"               "Prevotellaceae"               
## [1393,] "Fusobacteriales"             "Fusobacteriaceae"             
## [1394,] "Bacteroidales"               "Prevotellaceae"               
## [1395,] "Fusobacteriales"             "Leptotrichiaceae"             
## [1396,] NA                            NA                             
## [1397,] "Bacteroidales"               "Prevotellaceae"               
## [1398,] "Bacteroidales"               "Prevotellaceae"               
## [1399,] "Lactobacillales"             "Carnobacteriaceae"            
## [1400,] "Flavobacteriales"            "Weeksellaceae"                
## [1401,] "Fusobacteriales"             "Leptotrichiaceae"             
## [1402,] "Bacteroidales"               "Prevotellaceae"               
## [1403,] "Fusobacteriales"             "Fusobacteriaceae"             
## [1404,] "Bacteroidales"               "Prevotellaceae"               
## [1405,] "Campylobacterales"           "Campylobacteraceae"           
## [1406,] "Pasteurellales"              "Pasteurellaceae"              
## [1407,] "Fusobacteriales"             "Leptotrichiaceae"             
## [1408,] "Selenomonadales"             "Veillonellaceae"              
## [1409,] "Pasteurellales"              "Pasteurellaceae"              
## [1410,] "Fusobacteriales"             "Fusobacteriaceae"             
## [1411,] "Clostridiales"               "Ruminococcaceae"              
## [1412,] "Bacillales"                  "Staphylococcaceae"            
## [1413,] "Betaproteobacteriales"       NA                             
## [1414,] "Betaproteobacteriales"       "Neisseriaceae"                
## [1415,] "Bacteroidales"               "Prevotellaceae"               
## [1416,] "Micrococcales"               NA                             
## [1417,] "Spirochaetales"              "Spirochaetaceae"              
## [1418,] "Selenomonadales"             "Veillonellaceae"              
## [1419,] "Fusobacteriales"             "Leptotrichiaceae"             
## [1420,] "Fusobacteriales"             "Leptotrichiaceae"             
## [1421,] "Campylobacterales"           "P6-b51"                       
## [1422,] "Selenomonadales"             "Veillonellaceae"              
## [1423,] "Fusobacteriales"             "Fusobacteriaceae"             
## [1424,] "Bacteroidales"               "Prevotellaceae"               
## [1425,] "Selenomonadales"             "Veillonellaceae"              
## [1426,] "Campylobacterales"           "Campylobacteraceae"           
## [1427,] "Bacteroidales"               "Prevotellaceae"               
## [1428,] "Actinomycetales"             "Actinomycetaceae"             
## [1429,] "Bacteroidales"               "Prevotellaceae"               
## [1430,] "Bacteroidales"               "Prevotellaceae"               
## [1431,] "Selenomonadales"             "Veillonellaceae"              
## [1432,] "Spirochaetales"              "Spirochaetaceae"              
## [1433,] "Bacteroidales"               "Prevotellaceae"               
## [1434,] "Lactobacillales"             "Streptococcaceae"             
## [1435,] "Pasteurellales"              "Pasteurellaceae"              
## [1436,] "Saccharimonadales"           "Saccharimonadaceae"           
## [1437,] "Clostridiales"               "Family_XI"                    
## [1438,] "Selenomonadales"             "Veillonellaceae"              
## [1439,] "Pasteurellales"              "Pasteurellaceae"              
## [1440,] "Corynebacteriales"           "Corynebacteriaceae"           
## [1441,] "Selenomonadales"             "Veillonellaceae"              
## [1442,] "Selenomonadales"             "Veillonellaceae"              
## [1443,] "Bacteroidales"               "Prevotellaceae"               
## [1444,] "Flavobacteriales"            "Weeksellaceae"                
## [1445,] "Clostridiales"               "Lachnospiraceae"              
## [1446,] "Pasteurellales"              "Pasteurellaceae"              
## [1447,] NA                            NA                             
## [1448,] "Pasteurellales"              "Pasteurellaceae"              
## [1449,] "Bacteroidales"               "Prevotellaceae"               
## [1450,] "Lactobacillales"             "Streptococcaceae"             
## [1451,] "Lactobacillales"             "Streptococcaceae"             
## [1452,] "Actinomycetales"             "Actinomycetaceae"             
## [1453,] "Bacteroidales"               "Prevotellaceae"               
## [1454,] "Selenomonadales"             "Veillonellaceae"              
## [1455,] "Selenomonadales"             "Veillonellaceae"              
## [1456,] "Enterobacteriales"           "Enterobacteriaceae"           
## [1457,] "Bacteroidales"               "Prevotellaceae"               
## [1458,] "Saccharimonadales"           "Saccharimonadaceae"           
## [1459,] "Selenomonadales"             "Veillonellaceae"              
## [1460,] "Fusobacteriales"             "Fusobacteriaceae"             
## [1461,] "Bacteroidales"               "Porphyromonadaceae"           
## [1462,] "Bacteroidales"               "Prevotellaceae"               
## [1463,] "Spirochaetales"              "Spirochaetaceae"              
## [1464,] "Clostridiales"               "Lachnospiraceae"              
## [1465,] "Fusobacteriales"             "Leptotrichiaceae"             
## [1466,] "Micrococcales"               NA                             
## [1467,] "Bacteroidales"               "Rikenellaceae"                
## [1468,] "Bacteroidales"               NA                             
## [1469,] "Lactobacillales"             "Streptococcaceae"             
## [1470,] "Bacteroidales"               "Prevotellaceae"               
## [1471,] NA                            NA                             
## [1472,] "Fusobacteriales"             "Fusobacteriaceae"             
## [1473,] "Bacteroidales"               "Prevotellaceae"               
## [1474,] "Saccharimonadales"           NA                             
## [1475,] "Bacteroidales"               "Prevotellaceae"               
## [1476,] "Selenomonadales"             "Veillonellaceae"              
## [1477,] "Fusobacteriales"             "Leptotrichiaceae"             
## [1478,] "Clostridiales"               "Lachnospiraceae"              
## [1479,] "Pasteurellales"              "Pasteurellaceae"              
## [1480,] "Selenomonadales"             "Veillonellaceae"              
## [1481,] "Bacteroidales"               "Prevotellaceae"               
## [1482,] "Selenomonadales"             "Veillonellaceae"              
## [1483,] "Bacteroidales"               "Prevotellaceae"               
## [1484,] "Bacteroidales"               "Prevotellaceae"               
## [1485,] "Bacteroidales"               "Prevotellaceae"               
## [1486,] "Selenomonadales"             "Veillonellaceae"              
## [1487,] "Bacteroidales"               "Prevotellaceae"               
## [1488,] "Selenomonadales"             "Veillonellaceae"              
## [1489,] "Betaproteobacteriales"       "Neisseriaceae"                
## [1490,] "Actinomycetales"             "Actinomycetaceae"             
## [1491,] "Fusobacteriales"             "Leptotrichiaceae"             
## [1492,] "Clostridiales"               "Lachnospiraceae"              
## [1493,] "Clostridiales"               "Lachnospiraceae"              
## [1494,] "Selenomonadales"             "Veillonellaceae"              
## [1495,] "Pseudomonadales"             "Moraxellaceae"                
## [1496,] "Pasteurellales"              "Pasteurellaceae"              
## [1497,] "Bacteroidales"               "Prevotellaceae"               
## [1498,] "Pasteurellales"              "Pasteurellaceae"              
## [1499,] "Saccharimonadales"           "Saccharimonadaceae"           
## [1500,] "Lactobacillales"             "Streptococcaceae"             
## [1501,] "Corynebacteriales"           "Corynebacteriaceae"           
## [1502,] "Campylobacterales"           "Campylobacteraceae"           
## [1503,] "Flavobacteriales"            "Weeksellaceae"                
## [1504,] "Selenomonadales"             "Veillonellaceae"              
## [1505,] "Betaproteobacteriales"       "Neisseriaceae"                
## [1506,] "Saccharimonadales"           "Saccharimonadaceae"           
## [1507,] "Bacteroidales"               "Prevotellaceae"               
## [1508,] "Bacteroidales"               "Prevotellaceae"               
## [1509,] "Pasteurellales"              "Pasteurellaceae"              
## [1510,] "Fusobacteriales"             "Fusobacteriaceae"             
## [1511,] "Betaproteobacteriales"       "Neisseriaceae"                
## [1512,] "Bacteroidales"               "Prevotellaceae"               
## [1513,] "Flavobacteriales"            "Flavobacteriaceae"            
## [1514,] "Lactobacillales"             "Carnobacteriaceae"            
## [1515,] "Micrococcales"               NA                             
## [1516,] "Bacteroidales"               "Prevotellaceae"               
## [1517,] "Pasteurellales"              "Pasteurellaceae"              
## [1518,] "Clostridiales"               "Lachnospiraceae"              
## [1519,] "Lactobacillales"             "Carnobacteriaceae"            
## [1520,] "Bacteroidales"               "Prevotellaceae"               
## [1521,] "Fusobacteriales"             "Leptotrichiaceae"             
## [1522,] "Bacteroidales"               "Prevotellaceae"               
## [1523,] "Fusobacteriales"             "Fusobacteriaceae"             
## [1524,] "Betaproteobacteriales"       "Neisseriaceae"                
## [1525,] "Coriobacteriales"            "Atopobiaceae"                 
## [1526,] "Selenomonadales"             "Veillonellaceae"              
## [1527,] "Bacteroidales"               "Prevotellaceae"               
## [1528,] NA                            NA                             
## [1529,] "Bacteroidales"               "Prevotellaceae"               
## [1530,] "Clostridiales"               "Lachnospiraceae"              
## [1531,] "Campylobacterales"           "Campylobacteraceae"           
## [1532,] "Saccharimonadales"           "Saccharimonadaceae"           
## [1533,] "Fusobacteriales"             "Fusobacteriaceae"             
## [1534,] "Clostridiales"               "Lachnospiraceae"              
## [1535,] NA                            NA                             
## [1536,] "Bacteroidales"               "Prevotellaceae"               
## [1537,] "Bacteroidales"               "Prevotellaceae"               
## [1538,] "Absconditabacteriales_(SR1)" NA                             
## [1539,] "Actinomycetales"             "Actinomycetaceae"             
## [1540,] "Selenomonadales"             "Veillonellaceae"              
## [1541,] "Selenomonadales"             "Veillonellaceae"              
## [1542,] "Bacteroidales"               "Prevotellaceae"               
## [1543,] "Lactobacillales"             "Streptococcaceae"             
## [1544,] "Spirochaetales"              "Spirochaetaceae"              
## [1545,] "Clostridiales"               "Ruminococcaceae"              
## [1546,] "Selenomonadales"             "Veillonellaceae"              
## [1547,] "Bacteroidales"               "Prevotellaceae"               
## [1548,] "Micrococcales"               NA                             
## [1549,] "Fusobacteriales"             "Fusobacteriaceae"             
## [1550,] "Selenomonadales"             "Veillonellaceae"              
## [1551,] "Clostridiales"               "Peptostreptococcaceae"        
## [1552,] "Absconditabacteriales_(SR1)" NA                             
## [1553,] "Fusobacteriales"             "Leptotrichiaceae"             
## [1554,] "Selenomonadales"             "Veillonellaceae"              
## [1555,] "Bacteroidales"               "Prevotellaceae"               
## [1556,] "Actinomycetales"             "Actinomycetaceae"             
## [1557,] "Bacteroidales"               "Prevotellaceae"               
## [1558,] "Selenomonadales"             "Veillonellaceae"              
## [1559,] "Fusobacteriales"             "Fusobacteriaceae"             
## [1560,] "Chloroplast"                 NA                             
## [1561,] "Bacteroidales"               "Prevotellaceae"               
## [1562,] "Bacteroidales"               "Prevotellaceae"               
## [1563,] "Selenomonadales"             "Veillonellaceae"              
## [1564,] "Fusobacteriales"             "Leptotrichiaceae"             
## [1565,] "Bacteroidales"               "Prevotellaceae"               
## [1566,] "Clostridiales"               "Peptococcaceae"               
## [1567,] "Selenomonadales"             "Veillonellaceae"              
## [1568,] "Lactobacillales"             "Carnobacteriaceae"            
## [1569,] "Campylobacterales"           NA                             
## [1570,] "Bacteroidales"               "Rikenellaceae"                
## [1571,] "Bacteroidales"               "Prevotellaceae"               
## [1572,] "Bacteroidales"               "Prevotellaceae"               
## [1573,] "Selenomonadales"             "Veillonellaceae"              
## [1574,] "Fusobacteriales"             "Fusobacteriaceae"             
## [1575,] "Bacteroidales"               "Prevotellaceae"               
## [1576,] "Bacteroidales"               "Porphyromonadaceae"           
## [1577,] "Fusobacteriales"             "Fusobacteriaceae"             
## [1578,] "Bacteroidales"               "Prevotellaceae"               
## [1579,] "Bacteroidales"               "Prevotellaceae"               
## [1580,] "Bacteroidales"               "Prevotellaceae"               
## [1581,] "Bacteroidales"               "Prevotellaceae"               
## [1582,] "Selenomonadales"             "Veillonellaceae"              
## [1583,] "Lactobacillales"             "Streptococcaceae"             
## [1584,] "Spirochaetales"              "Spirochaetaceae"              
## [1585,] "Campylobacterales"           NA                             
## [1586,] "Bacteroidales"               "Prevotellaceae"               
## [1587,] "Selenomonadales"             "Veillonellaceae"              
## [1588,] "Selenomonadales"             "Veillonellaceae"              
## [1589,] "Sphingobacteriales"          "Lentimicrobiaceae"            
## [1590,] "Campylobacterales"           NA                             
## [1591,] "Bacteroidales"               "Prevotellaceae"               
## [1592,] "Lactobacillales"             "Carnobacteriaceae"            
## [1593,] "Pasteurellales"              "Pasteurellaceae"              
## [1594,] "Flavobacteriales"            "Flavobacteriaceae"            
## [1595,] "Pasteurellales"              "Pasteurellaceae"              
## [1596,] "Selenomonadales"             "Veillonellaceae"              
## [1597,] "Bacteroidales"               "Prevotellaceae"               
## [1598,] "Bacteroidales"               "Prevotellaceae"               
## [1599,] "Bacteroidales"               "Prevotellaceae"               
## [1600,] "Bacteroidales"               "Prevotellaceae"               
## [1601,] "Selenomonadales"             "Veillonellaceae"              
## [1602,] "Bacteroidales"               "Porphyromonadaceae"           
## [1603,] "Bacteroidales"               "Prevotellaceae"               
## [1604,] "Saccharimonadales"           "Saccharimonadaceae"           
## [1605,] "Saccharimonadales"           "Saccharimonadaceae"           
## [1606,] "Lactobacillales"             "Streptococcaceae"             
## [1607,] "Bacteroidales"               "Prevotellaceae"               
## [1608,] "Selenomonadales"             "Veillonellaceae"              
## [1609,] "Betaproteobacteriales"       "Neisseriaceae"                
## [1610,] "Bacteroidales"               "Prevotellaceae"               
## [1611,] "Fusobacteriales"             "Fusobacteriaceae"             
## [1612,] "Fusobacteriales"             "Leptotrichiaceae"             
## [1613,] "Pasteurellales"              "Pasteurellaceae"              
## [1614,] "Bacteroidales"               "Prevotellaceae"               
## [1615,] "Betaproteobacteriales"       "Neisseriaceae"                
## [1616,] "Chloroplast"                 NA                             
## [1617,] "Actinomycetales"             "Actinomycetaceae"             
## [1618,] "Selenomonadales"             "Veillonellaceae"              
## [1619,] "Selenomonadales"             "Veillonellaceae"              
## [1620,] "Fusobacteriales"             "Leptotrichiaceae"             
## [1621,] "Actinomycetales"             "Actinomycetaceae"             
## [1622,] "Pasteurellales"              "Pasteurellaceae"              
## [1623,] "Fusobacteriales"             "Fusobacteriaceae"             
## [1624,] "Bacteroidales"               "Porphyromonadaceae"           
## [1625,] "Fusobacteriales"             "Leptotrichiaceae"             
## [1626,] "Lactobacillales"             "Streptococcaceae"             
## [1627,] "Bacteroidales"               "Prevotellaceae"               
## [1628,] "Clostridiales"               "Lachnospiraceae"              
## [1629,] "Clostridiales"               "Lachnospiraceae"              
## [1630,] "Bacteroidales"               "Prevotellaceae"               
## [1631,] "Bacteroidales"               "Prevotellaceae"               
## [1632,] "Selenomonadales"             "Veillonellaceae"              
## [1633,] "Fusobacteriales"             "Leptotrichiaceae"             
## [1634,] "Selenomonadales"             "Veillonellaceae"              
## [1635,] "Bacteroidales"               "Prevotellaceae"               
## [1636,] "Pasteurellales"              "Pasteurellaceae"              
## [1637,] "Selenomonadales"             "Veillonellaceae"              
## [1638,] "Bacteroidales"               "Prevotellaceae"               
## [1639,] "Bacteroidales"               "Prevotellaceae"               
## [1640,] "Bacteroidales"               "Porphyromonadaceae"           
## [1641,] "Actinomycetales"             "Actinomycetaceae"             
## [1642,] "Bacteroidales"               "Prevotellaceae"               
## [1643,] "Clostridiales"               "Family_XIII"                  
## [1644,] NA                            NA                             
## [1645,] "Bacteroidales"               "Prevotellaceae"               
## [1646,] "Fusobacteriales"             NA                             
## [1647,] "Clostridiales"               "Lachnospiraceae"              
## [1648,] "Clostridiales"               "Ruminococcaceae"              
## [1649,] "Fusobacteriales"             "Leptotrichiaceae"             
## [1650,] "Fusobacteriales"             "Leptotrichiaceae"             
## [1651,] "Clostridiales"               "Lachnospiraceae"              
## [1652,] "Bacteroidales"               "Prevotellaceae"               
## [1653,] "Selenomonadales"             "Veillonellaceae"              
## [1654,] "Campylobacterales"           "Campylobacteraceae"           
## [1655,] "Saccharimonadales"           "Saccharimonadaceae"           
## [1656,] "Coriobacteriales"            "Eggerthellaceae"              
## [1657,] "Bacteroidales"               "Prevotellaceae"               
## [1658,] "Spirochaetales"              "Spirochaetaceae"              
## [1659,] "Pasteurellales"              "Pasteurellaceae"              
## [1660,] "Enterobacteriales"           "Enterobacteriaceae"           
## [1661,] "Bacteroidales"               "Prevotellaceae"               
## [1662,] "Bacteroidales"               "Prevotellaceae"               
## [1663,] "Selenomonadales"             "Veillonellaceae"              
## [1664,] "Bacteroidales"               "Porphyromonadaceae"           
## [1665,] "Bacteroidales"               "Prevotellaceae"               
## [1666,] "Fusobacteriales"             "Leptotrichiaceae"             
## [1667,] "Fusobacteriales"             "Leptotrichiaceae"             
## [1668,] "Clostridiales"               "Lachnospiraceae"              
## [1669,] "Spirochaetales"              "Spirochaetaceae"              
## [1670,] "Fusobacteriales"             "Leptotrichiaceae"             
## [1671,] NA                            NA                             
## [1672,] "Sphingobacteriales"          "Lentimicrobiaceae"            
## [1673,] "Pseudomonadales"             "Moraxellaceae"                
## [1674,] "Pasteurellales"              "Pasteurellaceae"              
## [1675,] "Bacteroidales"               "Prevotellaceae"               
## [1676,] "Clostridiales"               "Lachnospiraceae"              
## [1677,] "Lactobacillales"             "Streptococcaceae"             
## [1678,] NA                            NA                             
## [1679,] "Pasteurellales"              "Pasteurellaceae"              
## [1680,] "Saccharimonadales"           "Saccharimonadaceae"           
## [1681,] "Bacteroidales"               "Prevotellaceae"               
## [1682,] "Bacteroidales"               "Porphyromonadaceae"           
## [1683,] "Lactobacillales"             NA                             
## [1684,] "Pasteurellales"              "Pasteurellaceae"              
## [1685,] "Selenomonadales"             "Veillonellaceae"              
## [1686,] "Lactobacillales"             "Streptococcaceae"             
## [1687,] "Bacteroidales"               "Prevotellaceae"               
## [1688,] "Selenomonadales"             "Veillonellaceae"              
## [1689,] "Fusobacteriales"             "Fusobacteriaceae"             
## [1690,] "Pasteurellales"              "Pasteurellaceae"              
## [1691,] "Fusobacteriales"             "Fusobacteriaceae"             
## [1692,] "Bacteroidales"               "Prevotellaceae"               
## [1693,] "Fusobacteriales"             "Leptotrichiaceae"             
## [1694,] "Bacteroidales"               "Prevotellaceae"               
## [1695,] "Bacteroidales"               "Prevotellaceae"               
## [1696,] "Actinomycetales"             "Actinomycetaceae"             
## [1697,] "Bacteroidales"               "Prevotellaceae"               
## [1698,] "Campylobacterales"           "Campylobacteraceae"           
## [1699,] "Pasteurellales"              "Pasteurellaceae"              
## [1700,] "Actinomycetales"             "Actinomycetaceae"             
## [1701,] "Saccharimonadales"           NA                             
## [1702,] "Fusobacteriales"             "Fusobacteriaceae"             
## [1703,] "Lactobacillales"             "Streptococcaceae"             
## [1704,] "Pasteurellales"              "Pasteurellaceae"              
## [1705,] "Bacteroidales"               "Prevotellaceae"               
## [1706,] "Betaproteobacteriales"       "Neisseriaceae"                
## [1707,] "Lactobacillales"             "Streptococcaceae"             
## [1708,] "Selenomonadales"             "Veillonellaceae"              
## [1709,] "Lactobacillales"             "Streptococcaceae"             
## [1710,] "Bacteroidales"               "Prevotellaceae"               
## [1711,] "Selenomonadales"             "Veillonellaceae"              
## [1712,] "Selenomonadales"             "Veillonellaceae"              
## [1713,] "Clostridiales"               "Lachnospiraceae"              
## [1714,] "Selenomonadales"             "Veillonellaceae"              
## [1715,] "Bacteroidales"               "Prevotellaceae"               
## [1716,] "Fusobacteriales"             "Leptotrichiaceae"             
## [1717,] "Pasteurellales"              "Pasteurellaceae"              
## [1718,] "Flavobacteriales"            "Flavobacteriaceae"            
## [1719,] "Clostridiales"               "Lachnospiraceae"              
## [1720,] "Betaproteobacteriales"       "Neisseriaceae"                
## [1721,] "Bacteroidales"               "Prevotellaceae"               
## [1722,] "Flavobacteriales"            "Weeksellaceae"                
## [1723,] "Selenomonadales"             "Veillonellaceae"              
## [1724,] "Bacteroidales"               "Prevotellaceae"               
## [1725,] "Pasteurellales"              "Pasteurellaceae"              
## [1726,] "Bacteroidales"               "Prevotellaceae"               
## [1727,] "Coriobacteriales"            "Atopobiaceae"                 
## [1728,] "Bacteroidales"               "Prevotellaceae"               
## [1729,] "Bacteroidales"               "Prevotellaceae"               
## [1730,] "Fusobacteriales"             "Fusobacteriaceae"             
## [1731,] "Selenomonadales"             "Veillonellaceae"              
## [1732,] "Corynebacteriales"           "Corynebacteriaceae"           
## [1733,] "Bacteroidales"               "Prevotellaceae"               
## [1734,] "Sphingobacteriales"          "Lentimicrobiaceae"            
## [1735,] "Bacteroidales"               "Prevotellaceae"               
## [1736,] "Bacteroidales"               "Prevotellaceae"               
## [1737,] "Clostridiales"               "Defluviitaleaceae"            
## [1738,] "Bacteroidales"               "Prevotellaceae"               
## [1739,] "Bacteroidales"               "Prevotellaceae"               
## [1740,] "Bacteroidales"               "Prevotellaceae"               
## [1741,] "Selenomonadales"             "Veillonellaceae"              
## [1742,] "Bacteroidales"               "Prevotellaceae"               
## [1743,] "Lactobacillales"             "Streptococcaceae"             
## [1744,] "Bacteroidales"               "Prevotellaceae"               
## [1745,] "Betaproteobacteriales"       "Neisseriaceae"                
## [1746,] "Bacteroidales"               "Porphyromonadaceae"           
## [1747,] "Bacteroidales"               "Porphyromonadaceae"           
## [1748,] "Fusobacteriales"             "Fusobacteriaceae"             
## [1749,] "Bacteroidales"               "Prevotellaceae"               
## [1750,] "Bacteroidales"               "Prevotellaceae"               
## [1751,] "Bacillales"                  "Family_XI"                    
## [1752,] "Selenomonadales"             "Veillonellaceae"              
## [1753,] "Bacteroidales"               "Prevotellaceae"               
## [1754,] "Bacteroidales"               "Prevotellaceae"               
## [1755,] "Selenomonadales"             "Veillonellaceae"              
## [1756,] "Lactobacillales"             "Carnobacteriaceae"            
## [1757,] "Micrococcales"               NA                             
## [1758,] "Bacteroidales"               "Prevotellaceae"               
## [1759,] "Bacteroidales"               "Prevotellaceae"               
## [1760,] "Saccharimonadales"           "Saccharimonadaceae"           
## [1761,] "Bacteroidales"               "Porphyromonadaceae"           
## [1762,] "Lactobacillales"             "Carnobacteriaceae"            
## [1763,] "Lactobacillales"             "Streptococcaceae"             
## [1764,] "Selenomonadales"             "Veillonellaceae"              
## [1765,] "Clostridiales"               "Lachnospiraceae"              
## [1766,] "Bacteroidales"               "Prevotellaceae"               
## [1767,] "Clostridiales"               "Ruminococcaceae"              
## [1768,] "Bacteroidales"               "Prevotellaceae"               
## [1769,] "Selenomonadales"             "Veillonellaceae"              
## [1770,] "Bacteroidales"               "Porphyromonadaceae"           
## [1771,] "Selenomonadales"             "Veillonellaceae"              
## [1772,] "Cardiobacteriales"           "Cardiobacteriaceae"           
## [1773,] "Clostridiales"               "Lachnospiraceae"              
## [1774,] "Actinomycetales"             "Actinomycetaceae"             
## [1775,] "Clostridiales"               "Ruminococcaceae"              
## [1776,] "Bacillales"                  "Family_XI"                    
## [1777,] "Pasteurellales"              "Pasteurellaceae"              
## [1778,] "Betaproteobacteriales"       "Neisseriaceae"                
## [1779,] "Fusobacteriales"             "Fusobacteriaceae"             
## [1780,] "Bacteroidales"               "Prevotellaceae"               
## [1781,] "Clostridiales"               "Lachnospiraceae"              
## [1782,] "Pasteurellales"              "Pasteurellaceae"              
## [1783,] "Micrococcales"               NA                             
## [1784,] "Bacteroidales"               "Prevotellaceae"               
## [1785,] "Bacteroidales"               "Prevotellaceae"               
## [1786,] "Bacteroidales"               "Prevotellaceae"               
## [1787,] "Selenomonadales"             "Veillonellaceae"              
## [1788,] "Bacteroidales"               "Prevotellaceae"               
## [1789,] "Pasteurellales"              "Pasteurellaceae"              
## [1790,] "Fusobacteriales"             "Leptotrichiaceae"             
## [1791,] "Pasteurellales"              "Pasteurellaceae"              
## [1792,] NA                            NA                             
## [1793,] NA                            NA                             
## [1794,] "Clostridiales"               "Peptostreptococcaceae"        
## [1795,] NA                            NA                             
## [1796,] "Bacteroidales"               "Porphyromonadaceae"           
## [1797,] "Bacteroidales"               "Prevotellaceae"               
## [1798,] "Clostridiales"               "Lachnospiraceae"              
## [1799,] "Lactobacillales"             "Streptococcaceae"             
## [1800,] "Clostridiales"               "Family_XIII"                  
## [1801,] "Clostridiales"               "Lachnospiraceae"              
## [1802,] "Selenomonadales"             "Veillonellaceae"              
## [1803,] "Fusobacteriales"             "Fusobacteriaceae"             
## [1804,] "Clostridiales"               "Lachnospiraceae"              
## [1805,] "Fusobacteriales"             "Fusobacteriaceae"             
## [1806,] "Flavobacteriales"            "Flavobacteriaceae"            
## [1807,] "Fusobacteriales"             "Fusobacteriaceae"             
## [1808,] "Lactobacillales"             "Carnobacteriaceae"            
## [1809,] "Micrococcales"               "Micrococcaceae"               
## [1810,] "Actinomycetales"             "Actinomycetaceae"             
## [1811,] "Selenomonadales"             "Veillonellaceae"              
## [1812,] "Selenomonadales"             "Veillonellaceae"              
## [1813,] "Selenomonadales"             "Veillonellaceae"              
## [1814,] "Fusobacteriales"             "Fusobacteriaceae"             
## [1815,] "Betaproteobacteriales"       "Neisseriaceae"                
## [1816,] "Pasteurellales"              "Pasteurellaceae"              
## [1817,] "Enterobacteriales"           "Enterobacteriaceae"           
## [1818,] "Bacteroidales"               "Prevotellaceae"               
## [1819,] "Bacteroidales"               "Porphyromonadaceae"           
## [1820,] "Bacteroidales"               "Prevotellaceae"               
## [1821,] "Betaproteobacteriales"       "Neisseriaceae"                
## [1822,] "Bacteroidales"               "Prevotellaceae"               
## [1823,] "Selenomonadales"             "Veillonellaceae"              
## [1824,] "Selenomonadales"             "Veillonellaceae"              
## [1825,] "Lactobacillales"             "Streptococcaceae"             
## [1826,] "Selenomonadales"             "Veillonellaceae"              
## [1827,] "Bacteroidales"               "Porphyromonadaceae"           
## [1828,] "Pasteurellales"              "Pasteurellaceae"              
## [1829,] "Clostridiales"               "Lachnospiraceae"              
## [1830,] "Fusobacteriales"             "Leptotrichiaceae"             
## [1831,] "Fusobacteriales"             "Leptotrichiaceae"             
## [1832,] "Bacteroidales"               "Prevotellaceae"               
## [1833,] "Corynebacteriales"           "Corynebacteriaceae"           
## [1834,] "Saccharimonadales"           "Saccharimonadaceae"           
## [1835,] "Flavobacteriales"            "Weeksellaceae"                
## [1836,] "Spirochaetales"              "Spirochaetaceae"              
## [1837,] NA                            NA                             
## [1838,] "Bacteroidales"               "Prevotellaceae"               
## [1839,] "Fusobacteriales"             "Leptotrichiaceae"             
## [1840,] "Spirochaetales"              "Spirochaetaceae"              
## [1841,] "Selenomonadales"             "Veillonellaceae"              
## [1842,] "Clostridiales"               "Family_XI"                    
## [1843,] "Bacteroidales"               "Prevotellaceae"               
## [1844,] "Bacteroidales"               "Porphyromonadaceae"           
## [1845,] "Clostridiales"               "Peptostreptococcaceae"        
## [1846,] "Betaproteobacteriales"       "Burkholderiaceae"             
## [1847,] "Bacteroidales"               "Prevotellaceae"               
## [1848,] "Bacteroidales"               "Prevotellaceae"               
## [1849,] "Bacteroidales"               "Prevotellaceae"               
## [1850,] "Bacteroidales"               "Prevotellaceae"               
## [1851,] "Synergistales"               "Synergistaceae"               
## [1852,] "Bacteroidales"               "Prevotellaceae"               
## [1853,] "Bacteroidales"               "Prevotellaceae"               
## [1854,] "Pasteurellales"              "Pasteurellaceae"              
## [1855,] "Bacteroidales"               "Prevotellaceae"               
## [1856,] "Pasteurellales"              "Pasteurellaceae"              
## [1857,] "Bacteroidales"               "Prevotellaceae"               
## [1858,] "Bacteroidales"               "Porphyromonadaceae"           
## [1859,] "Betaproteobacteriales"       "Neisseriaceae"                
## [1860,] "Clostridiales"               "Lachnospiraceae"              
## [1861,] "Pasteurellales"              "Pasteurellaceae"              
## [1862,] "Flavobacteriales"            "Flavobacteriaceae"            
## [1863,] "Betaproteobacteriales"       "Neisseriaceae"                
## [1864,] "Fusobacteriales"             "Fusobacteriaceae"             
## [1865,] "Pasteurellales"              "Pasteurellaceae"              
## [1866,] "Bacteroidales"               "Prevotellaceae"               
## [1867,] "Fusobacteriales"             "Leptotrichiaceae"             
## [1868,] "Clostridiales"               "Lachnospiraceae"              
## [1869,] "Pasteurellales"              "Pasteurellaceae"              
## [1870,] "Selenomonadales"             "Veillonellaceae"              
## [1871,] "Pasteurellales"              "Pasteurellaceae"              
## [1872,] "Lactobacillales"             "Streptococcaceae"             
## [1873,] "Pasteurellales"              "Pasteurellaceae"              
## [1874,] "Selenomonadales"             "Veillonellaceae"              
## [1875,] "Fusobacteriales"             "Fusobacteriaceae"             
## [1876,] "Selenomonadales"             "Veillonellaceae"              
## [1877,] NA                            NA                             
## [1878,] "Flavobacteriales"            "Flavobacteriaceae"            
## [1879,] "Enterobacteriales"           "Enterobacteriaceae"           
## [1880,] "Clostridiales"               "Ruminococcaceae"              
## [1881,] "Bacteroidales"               "Prevotellaceae"               
## [1882,] "Selenomonadales"             "Veillonellaceae"              
## [1883,] "Fusobacteriales"             "Fusobacteriaceae"             
## [1884,] "Bacteroidales"               "Prevotellaceae"               
## [1885,] "Actinomycetales"             "Actinomycetaceae"             
## [1886,] NA                            NA                             
## [1887,] "Selenomonadales"             "Veillonellaceae"              
## [1888,] "Micrococcales"               "Micrococcaceae"               
## [1889,] "Bacteroidales"               "Prevotellaceae"               
## [1890,] "Bacteroidales"               "Prevotellaceae"               
## [1891,] "Betaproteobacteriales"       "Neisseriaceae"                
## [1892,] "Betaproteobacteriales"       "Neisseriaceae"                
## [1893,] "Selenomonadales"             "Veillonellaceae"              
## [1894,] "Fusobacteriales"             "Fusobacteriaceae"             
## [1895,] "Selenomonadales"             "Veillonellaceae"              
## [1896,] "Clostridiales"               "Lachnospiraceae"              
## [1897,] "Bacteroidales"               "Prevotellaceae"               
## [1898,] "Lactobacillales"             "Streptococcaceae"             
## [1899,] "Pasteurellales"              "Pasteurellaceae"              
## [1900,] "Selenomonadales"             "Veillonellaceae"              
## [1901,] "Clostridiales"               "Lachnospiraceae"              
## [1902,] "Selenomonadales"             "Veillonellaceae"              
## [1903,] "Bacteroidales"               "Prevotellaceae"               
## [1904,] "Bacteroidales"               "Prevotellaceae"               
## [1905,] "Fusobacteriales"             "Fusobacteriaceae"             
## [1906,] "Saccharimonadales"           "Saccharimonadaceae"           
## [1907,] "Clostridiales"               "Peptostreptococcaceae"        
## [1908,] "Bacteroidales"               "Prevotellaceae"               
## [1909,] "Clostridiales"               "Lachnospiraceae"              
## [1910,] "Fusobacteriales"             "Fusobacteriaceae"             
## [1911,] "Pasteurellales"              "Pasteurellaceae"              
## [1912,] "Bacteroidales"               "Prevotellaceae"               
## [1913,] "Selenomonadales"             "Veillonellaceae"              
## [1914,] "Flavobacteriales"            "Flavobacteriaceae"            
## [1915,] "Bacteroidales"               "Prevotellaceae"               
## [1916,] "Spirochaetales"              "Spirochaetaceae"              
## [1917,] "Flavobacteriales"            "Flavobacteriaceae"            
## [1918,] "Flavobacteriales"            "Flavobacteriaceae"            
## [1919,] "Micrococcales"               NA                             
## [1920,] "Clostridiales"               "Lachnospiraceae"              
## [1921,] "Bacteroidales"               "Prevotellaceae"               
## [1922,] "Clostridiales"               "Lachnospiraceae"              
## [1923,] "Pseudomonadales"             "Pseudomonadaceae"             
## [1924,] "Enterobacteriales"           "Enterobacteriaceae"           
## [1925,] "Bacteroidales"               "Prevotellaceae"               
## [1926,] "Bacteroidales"               "Prevotellaceae"               
## [1927,] "Campylobacterales"           "Campylobacteraceae"           
## [1928,] "Pasteurellales"              "Pasteurellaceae"              
## [1929,] "Selenomonadales"             "Veillonellaceae"              
## [1930,] "Bacteroidales"               "Prevotellaceae"               
## [1931,] "Fusobacteriales"             "Leptotrichiaceae"             
## [1932,] "Bacteroidales"               "Prevotellaceae"               
## [1933,] "Saccharimonadales"           "Saccharimonadaceae"           
## [1934,] "Bacteroidales"               "Prevotellaceae"               
## [1935,] "Micrococcales"               NA                             
## [1936,] "Micrococcales"               NA                             
## [1937,] "Betaproteobacteriales"       "Neisseriaceae"                
## [1938,] "Bacteroidales"               "Prevotellaceae"               
## [1939,] "Bacteroidales"               "Prevotellaceae"               
## [1940,] "Fusobacteriales"             "Fusobacteriaceae"             
## [1941,] "Bacillales"                  "Family_XI"                    
## [1942,] "Micrococcales"               NA                             
## [1943,] "Bacteroidales"               "Prevotellaceae"               
## [1944,] "Betaproteobacteriales"       "Neisseriaceae"                
## [1945,] "Clostridiales"               "Lachnospiraceae"              
## [1946,] "Spirochaetales"              "Spirochaetaceae"              
## [1947,] "Betaproteobacteriales"       "Neisseriaceae"                
## [1948,] "Fusobacteriales"             "Fusobacteriaceae"             
## [1949,] "Flavobacteriales"            "Flavobacteriaceae"            
## [1950,] "Selenomonadales"             "Veillonellaceae"              
## [1951,] "Bacteroidales"               "Prevotellaceae"               
## [1952,] "Clostridiales"               "Lachnospiraceae"              
## [1953,] "Selenomonadales"             "Veillonellaceae"              
## [1954,] "Pasteurellales"              "Pasteurellaceae"              
## [1955,] "Bacteroidales"               "Prevotellaceae"               
## [1956,] "Bacteroidales"               "Prevotellaceae"               
## [1957,] "Clostridiales"               "Lachnospiraceae"              
## [1958,] "Bacteroidales"               "Porphyromonadaceae"           
## [1959,] "Bacteroidales"               "Prevotellaceae"               
## [1960,] "Bacteroidales"               "Prevotellaceae"               
## [1961,] "Flavobacteriales"            "Flavobacteriaceae"            
## [1962,] "Coriobacteriales"            "Atopobiaceae"                 
## [1963,] "Bacteroidales"               "Prevotellaceae"               
## [1964,] "Actinomycetales"             "Actinomycetaceae"             
## [1965,] "Fusobacteriales"             "Leptotrichiaceae"             
## [1966,] "Selenomonadales"             "Veillonellaceae"              
## [1967,] "Bacteroidales"               "Prevotellaceae"               
## [1968,] "Selenomonadales"             "Veillonellaceae"              
## [1969,] "Fusobacteriales"             "Fusobacteriaceae"             
## [1970,] "Betaproteobacteriales"       "Neisseriaceae"                
## [1971,] "Fusobacteriales"             NA                             
## [1972,] "Clostridiales"               "Peptostreptococcaceae"        
## [1973,] "Bacteroidales"               "Prevotellaceae"               
## [1974,] "Clostridiales"               "Lachnospiraceae"              
## [1975,] "Clostridiales"               "Family_XI"                    
## [1976,] "Bacteroidales"               "Prevotellaceae"               
## [1977,] "Clostridiales"               "Peptostreptococcaceae"        
## [1978,] "Bacteroidales"               "Prevotellaceae"               
## [1979,] "Bacteroidales"               "Porphyromonadaceae"           
## [1980,] "Bacteroidales"               "Prevotellaceae"               
## [1981,] "Bacteroidales"               "Prevotellaceae"               
## [1982,] "Fusobacteriales"             "Fusobacteriaceae"             
## [1983,] "Selenomonadales"             "Veillonellaceae"              
## [1984,] "Betaproteobacteriales"       "Neisseriaceae"                
## [1985,] "Bacteroidales"               "Porphyromonadaceae"           
## [1986,] "Selenomonadales"             "Veillonellaceae"              
## [1987,] "Pasteurellales"              "Pasteurellaceae"              
## [1988,] "Micrococcales"               NA                             
## [1989,] "Fusobacteriales"             "Leptotrichiaceae"             
## [1990,] "Lactobacillales"             "Streptococcaceae"             
## [1991,] "Bacteroidales"               "Prevotellaceae"               
## [1992,] "Bacteroidales"               "Rikenellaceae"                
## [1993,] "Flavobacteriales"            "Flavobacteriaceae"            
## [1994,] "Betaproteobacteriales"       "Neisseriaceae"                
## [1995,] "Selenomonadales"             "Veillonellaceae"              
## [1996,] "Bacillales"                  "Family_XI"                    
## [1997,] "Bacteroidales"               "Tannerellaceae"               
## [1998,] "Selenomonadales"             "Veillonellaceae"              
## [1999,] "Clostridiales"               "Lachnospiraceae"              
## [2000,] "Betaproteobacteriales"       "Neisseriaceae"                
## [2001,] "Fusobacteriales"             "Leptotrichiaceae"             
## [2002,] "Clostridiales"               "Peptostreptococcaceae"        
## [2003,] "Fusobacteriales"             "Fusobacteriaceae"             
## [2004,] "Bacteroidales"               "Paludibacteraceae"            
## [2005,] "Bacteroidales"               "Prevotellaceae"               
## [2006,] "Clostridiales"               "Lachnospiraceae"              
## [2007,] "Bacteroidales"               "Prevotellaceae"               
## [2008,] "Selenomonadales"             "Veillonellaceae"              
## [2009,] "Bacteroidales"               "Prevotellaceae"               
## [2010,] "Pasteurellales"              "Pasteurellaceae"              
## [2011,] "Bacteroidales"               "Prevotellaceae"               
## [2012,] "Bacteroidales"               "Porphyromonadaceae"           
## [2013,] "Saccharimonadales"           NA                             
## [2014,] "Bacteroidales"               "Prevotellaceae"               
## [2015,] "Bacteroidales"               "Prevotellaceae"               
## [2016,] "Selenomonadales"             "Veillonellaceae"              
## [2017,] "Flavobacteriales"            "Flavobacteriaceae"            
## [2018,] "Saccharimonadales"           "Saccharimonadaceae"           
## [2019,] "Selenomonadales"             "Veillonellaceae"              
## [2020,] "Fusobacteriales"             "Fusobacteriaceae"             
## [2021,] "Bacteroidales"               "Porphyromonadaceae"           
## [2022,] "Actinomycetales"             "Actinomycetaceae"             
## [2023,] "Pasteurellales"              "Pasteurellaceae"              
## [2024,] "Pasteurellales"              "Pasteurellaceae"              
## [2025,] "Betaproteobacteriales"       "Neisseriaceae"                
## [2026,] "Bacteroidales"               "Prevotellaceae"               
## [2027,] NA                            NA                             
## [2028,] "Bacteroidales"               "Prevotellaceae"               
## [2029,] "Fusobacteriales"             "Leptotrichiaceae"             
## [2030,] "Fusobacteriales"             NA                             
## [2031,] "Fusobacteriales"             "Leptotrichiaceae"             
## [2032,] "Saccharimonadales"           "Saccharimonadaceae"           
## [2033,] "Fusobacteriales"             "Fusobacteriaceae"             
## [2034,] "Lactobacillales"             "Streptococcaceae"             
## [2035,] "Lactobacillales"             "Streptococcaceae"             
## [2036,] "Lactobacillales"             "Carnobacteriaceae"            
## [2037,] "Synergistales"               "Synergistaceae"               
## [2038,] "Bacteroidales"               "Prevotellaceae"               
## [2039,] "Fusobacteriales"             "Fusobacteriaceae"             
## [2040,] "Selenomonadales"             "Veillonellaceae"              
## [2041,] "Selenomonadales"             "Veillonellaceae"              
## [2042,] "Bacteroidales"               "Prevotellaceae"               
## [2043,] "Pasteurellales"              "Pasteurellaceae"              
## [2044,] "Betaproteobacteriales"       "Neisseriaceae"                
## [2045,] "Absconditabacteriales_(SR1)" NA                             
## [2046,] "Fusobacteriales"             "Leptotrichiaceae"             
## [2047,] "Clostridiales"               "Ruminococcaceae"              
## [2048,] "Selenomonadales"             "Veillonellaceae"              
## [2049,] "Campylobacterales"           "Campylobacteraceae"           
## [2050,] "Bacteroidales"               "Prevotellaceae"               
## [2051,] "Bacteroidales"               "Prevotellaceae"               
## [2052,] "Selenomonadales"             "Veillonellaceae"              
## [2053,] "Bacteroidales"               "Prevotellaceae"               
## [2054,] "Actinomycetales"             "Actinomycetaceae"             
## [2055,] "Bacteroidales"               "Prevotellaceae"               
## [2056,] "Selenomonadales"             "Veillonellaceae"              
## [2057,] "Saccharimonadales"           "Saccharimonadaceae"           
## [2058,] "Selenomonadales"             "Veillonellaceae"              
## [2059,] "Bacteroidales"               "Prevotellaceae"               
## [2060,] "Clostridiales"               "Lachnospiraceae"              
## [2061,] "Bacteroidales"               "Prevotellaceae"               
## [2062,] "Selenomonadales"             "Veillonellaceae"              
## [2063,] "Chloroplast"                 NA                             
## [2064,] "Bacteroidales"               "Porphyromonadaceae"           
## [2065,] "Selenomonadales"             "Veillonellaceae"              
## [2066,] "Betaproteobacteriales"       "Neisseriaceae"                
## [2067,] "Bacteroidales"               "Prevotellaceae"               
## [2068,] "Bacteroidales"               "Prevotellaceae"               
## [2069,] "Fusobacteriales"             "Fusobacteriaceae"             
## [2070,] "Saccharimonadales"           "Saccharimonadaceae"           
## [2071,] "Bacteroidales"               "Prevotellaceae"               
## [2072,] "Campylobacterales"           NA                             
## [2073,] "Clostridiales"               "Lachnospiraceae"              
## [2074,] "Pasteurellales"              "Pasteurellaceae"              
## [2075,] "Fusobacteriales"             NA                             
## [2076,] "Selenomonadales"             "Veillonellaceae"              
## [2077,] "Spirochaetales"              "Spirochaetaceae"              
## [2078,] "Fusobacteriales"             "Fusobacteriaceae"             
## [2079,] "Saccharimonadales"           "Saccharimonadaceae"           
## [2080,] "Bacteroidales"               "Porphyromonadaceae"           
## [2081,] "Flavobacteriales"            "Flavobacteriaceae"            
## [2082,] "Bacteroidales"               "Prevotellaceae"               
## [2083,] "Fusobacteriales"             "Fusobacteriaceae"             
## [2084,] NA                            NA                             
## [2085,] "Betaproteobacteriales"       "Neisseriaceae"                
## [2086,] "Fusobacteriales"             "Leptotrichiaceae"             
## [2087,] "Clostridiales"               "Lachnospiraceae"              
## [2088,] "Bacteroidales"               "Prevotellaceae"               
## [2089,] "Selenomonadales"             "Veillonellaceae"              
## [2090,] "Bacteroidales"               "Prevotellaceae"               
## [2091,] "Spirochaetales"              "Spirochaetaceae"              
## [2092,] "Micrococcales"               NA                             
## [2093,] "Micrococcales"               NA                             
## [2094,] "Bacteroidales"               "Prevotellaceae"               
## [2095,] "Bacteroidales"               "Porphyromonadaceae"           
## [2096,] "Fusobacteriales"             "Fusobacteriaceae"             
## [2097,] "Bacteroidales"               "Porphyromonadaceae"           
## [2098,] "Spirochaetales"              "Spirochaetaceae"              
## [2099,] "Fusobacteriales"             "Leptotrichiaceae"             
## [2100,] "Bacteroidales"               "Prevotellaceae"               
## [2101,] "Bacteroidales"               "Prevotellaceae"               
## [2102,] "Fusobacteriales"             "Leptotrichiaceae"             
## [2103,] "Clostridiales"               "Lachnospiraceae"              
## [2104,] "Bacteroidales"               "Prevotellaceae"               
## [2105,] "Fusobacteriales"             "Leptotrichiaceae"             
## [2106,] "Pasteurellales"              "Pasteurellaceae"              
## [2107,] "Bacteroidales"               "Prevotellaceae"               
## [2108,] "Selenomonadales"             "Veillonellaceae"              
## [2109,] "Bacteroidales"               "Prevotellaceae"               
## [2110,] "Bacteroidales"               "Prevotellaceae"               
## [2111,] "Bacteroidales"               "Prevotellaceae"               
## [2112,] "Clostridiales"               "Lachnospiraceae"              
## [2113,] "Selenomonadales"             "Veillonellaceae"              
## [2114,] "Betaproteobacteriales"       "Neisseriaceae"                
## [2115,] "Bacteroidales"               "Prevotellaceae"               
## [2116,] NA                            NA                             
## [2117,] "Lactobacillales"             "Carnobacteriaceae"            
## [2118,] "Bacteroidales"               "Prevotellaceae"               
## [2119,] "Selenomonadales"             "Veillonellaceae"              
## [2120,] "Pasteurellales"              "Pasteurellaceae"              
## [2121,] "Flavobacteriales"            "Flavobacteriaceae"            
## [2122,] "Bacteroidales"               "Prevotellaceae"               
## [2123,] "JGI_0000069-P22"             NA                             
## [2124,] "Selenomonadales"             "Veillonellaceae"              
## [2125,] "Bacteroidales"               "Prevotellaceae"               
## [2126,] "Lactobacillales"             "Streptococcaceae"             
## [2127,] "Bacteroidales"               "Prevotellaceae"               
## [2128,] "Selenomonadales"             "Veillonellaceae"              
## [2129,] "Lactobacillales"             "Streptococcaceae"             
## [2130,] "Pasteurellales"              "Pasteurellaceae"              
## [2131,] "Bacteroidales"               "Prevotellaceae"               
## [2132,] "Campylobacterales"           NA                             
## [2133,] "Actinomycetales"             "Actinomycetaceae"             
## [2134,] "Fusobacteriales"             "Fusobacteriaceae"             
## [2135,] "Bacteroidales"               "Prevotellaceae"               
## [2136,] "Selenomonadales"             "Veillonellaceae"              
## [2137,] "Micrococcales"               "Micrococcaceae"               
## [2138,] "Bacteroidales"               "Prevotellaceae"               
## [2139,] "Bacteroidales"               "Prevotellaceae"               
## [2140,] "Clostridiales"               "Family_XIII"                  
## [2141,] "Spirochaetales"              "Spirochaetaceae"              
## [2142,] "Campylobacterales"           NA                             
## [2143,] "Betaproteobacteriales"       "Neisseriaceae"                
## [2144,] "Selenomonadales"             "Veillonellaceae"              
## [2145,] "Bacteroidales"               "Prevotellaceae"               
## [2146,] "Bacteroidales"               "Prevotellaceae"               
## [2147,] "Bacteroidales"               "Prevotellaceae"               
## [2148,] "Bacteroidales"               "Porphyromonadaceae"           
## [2149,] "Fusobacteriales"             "Leptotrichiaceae"             
## [2150,] "Betaproteobacteriales"       "Neisseriaceae"                
## [2151,] "Saccharimonadales"           "Saccharimonadaceae"           
## [2152,] "Cardiobacteriales"           "Cardiobacteriaceae"           
## [2153,] "Campylobacterales"           "Campylobacteraceae"           
## [2154,] "Bacteroidales"               "Prevotellaceae"               
## [2155,] "Bacteroidales"               "Prevotellaceae"               
## [2156,] "Bacteroidales"               "Prevotellaceae"               
## [2157,] "Betaproteobacteriales"       "Neisseriaceae"                
## [2158,] NA                            NA                             
## [2159,] "Bacteroidales"               "Porphyromonadaceae"           
## [2160,] "Lactobacillales"             "Carnobacteriaceae"            
## [2161,] "Pasteurellales"              "Pasteurellaceae"              
## [2162,] "Actinomycetales"             "Actinomycetaceae"             
## [2163,] "Fusobacteriales"             "Fusobacteriaceae"             
## [2164,] "Flavobacteriales"            "Flavobacteriaceae"            
## [2165,] "Selenomonadales"             "Veillonellaceae"              
## [2166,] "Bacteroidales"               "Porphyromonadaceae"           
## [2167,] "Bacteroidales"               "Prevotellaceae"               
## [2168,] "Spirochaetales"              "Spirochaetaceae"              
## [2169,] "Bacteroidales"               "Prevotellaceae"               
## [2170,] "Clostridiales"               "Lachnospiraceae"              
## [2171,] "Fusobacteriales"             "Fusobacteriaceae"             
## [2172,] "Clostridiales"               "Lachnospiraceae"              
## [2173,] "Actinomycetales"             "Actinomycetaceae"             
## [2174,] "Bacteroidales"               "Prevotellaceae"               
## [2175,] "Bacteroidales"               "Prevotellaceae"               
## [2176,] "Fusobacteriales"             "Leptotrichiaceae"             
## [2177,] "Flavobacteriales"            "Flavobacteriaceae"            
## [2178,] "Clostridiales"               "Lachnospiraceae"              
## [2179,] "Selenomonadales"             "Veillonellaceae"              
## [2180,] "Selenomonadales"             "Veillonellaceae"              
## [2181,] "Fusobacteriales"             "Fusobacteriaceae"             
## [2182,] "Clostridiales"               "Peptostreptococcaceae"        
## [2183,] "Bacteroidales"               "Porphyromonadaceae"           
## [2184,] "Bacteroidales"               "Prevotellaceae"               
## [2185,] "Bacteroidales"               "Prevotellaceae"               
## [2186,] "Fusobacteriales"             "Fusobacteriaceae"             
## [2187,] "Selenomonadales"             "Veillonellaceae"              
## [2188,] "Selenomonadales"             "Veillonellaceae"              
## [2189,] "Fusobacteriales"             "Fusobacteriaceae"             
## [2190,] "Selenomonadales"             "Veillonellaceae"              
## [2191,] "Bacteroidales"               "Prevotellaceae"               
## [2192,] "Betaproteobacteriales"       "Neisseriaceae"                
## [2193,] "Flavobacteriales"            "Weeksellaceae"                
## [2194,] "Selenomonadales"             "Veillonellaceae"              
## [2195,] "Pasteurellales"              "Pasteurellaceae"              
## [2196,] "Saccharimonadales"           NA                             
## [2197,] "Bacteroidales"               "Porphyromonadaceae"           
## [2198,] "Fusobacteriales"             "Fusobacteriaceae"             
## [2199,] "Bacteroidales"               "Prevotellaceae"               
## [2200,] "Selenomonadales"             "Veillonellaceae"              
## [2201,] "Actinomycetales"             "Actinomycetaceae"             
## [2202,] NA                            NA                             
## [2203,] "Selenomonadales"             "Veillonellaceae"              
## [2204,] "Fusobacteriales"             "Fusobacteriaceae"             
## [2205,] "Selenomonadales"             "Veillonellaceae"              
## [2206,] "Pasteurellales"              "Pasteurellaceae"              
## [2207,] "Bacteroidales"               "Prevotellaceae"               
## [2208,] "Pasteurellales"              "Pasteurellaceae"              
## [2209,] "Saccharimonadales"           "Saccharimonadaceae"           
## [2210,] "Spirochaetales"              "Spirochaetaceae"              
## [2211,] "Clostridiales"               "Peptostreptococcaceae"        
## [2212,] "Bacteroidales"               "Prevotellaceae"               
## [2213,] "Bacteroidales"               "Porphyromonadaceae"           
## [2214,] "Selenomonadales"             "Veillonellaceae"              
## [2215,] "Bacteroidales"               "Prevotellaceae"               
## [2216,] "Bacteroidales"               "Prevotellaceae"               
## [2217,] "Flavobacteriales"            "Flavobacteriaceae"            
## [2218,] "Selenomonadales"             "Veillonellaceae"              
## [2219,] "Bacteroidales"               "Prevotellaceae"               
## [2220,] "Bacteroidales"               "Prevotellaceae"               
## [2221,] "Saccharimonadales"           NA                             
## [2222,] "Flavobacteriales"            "Flavobacteriaceae"            
## [2223,] "Chloroplast"                 NA                             
## [2224,] "Flavobacteriales"            "Flavobacteriaceae"            
## [2225,] "Selenomonadales"             "Veillonellaceae"              
## [2226,] "Saccharimonadales"           "Saccharimonadaceae"           
## [2227,] "Lactobacillales"             "Streptococcaceae"             
## [2228,] "Bacteroidales"               "Prevotellaceae"               
## [2229,] "Clostridiales"               "Lachnospiraceae"              
## [2230,] NA                            NA                             
## [2231,] "Actinomycetales"             "Actinomycetaceae"             
## [2232,] "Selenomonadales"             "Veillonellaceae"              
## [2233,] "Bacteroidales"               "Prevotellaceae"               
## [2234,] "Selenomonadales"             "Veillonellaceae"              
## [2235,] "Clostridiales"               "Lachnospiraceae"              
## [2236,] "Selenomonadales"             "Veillonellaceae"              
## [2237,] "Bacteroidales"               "Prevotellaceae"               
## [2238,] "Fusobacteriales"             "Fusobacteriaceae"             
## [2239,] "Selenomonadales"             "Veillonellaceae"              
## [2240,] "Micrococcales"               NA                             
## [2241,] "Bacteroidales"               "Prevotellaceae"               
## [2242,] "Chloroplast"                 NA                             
## [2243,] "Bacteroidales"               "Porphyromonadaceae"           
## [2244,] "Bacteroidales"               "Prevotellaceae"               
## [2245,] "Pasteurellales"              "Pasteurellaceae"              
## [2246,] "Bacteroidales"               "Porphyromonadaceae"           
## [2247,] "Bacteroidales"               "Prevotellaceae"               
## [2248,] "Bacteroidales"               "Porphyromonadaceae"           
## [2249,] NA                            NA                             
## [2250,] "Saccharimonadales"           "Saccharimonadaceae"           
## [2251,] "Coriobacteriales"            "Atopobiaceae"                 
## [2252,] "Bacteroidales"               "Prevotellaceae"               
## [2253,] "Selenomonadales"             "Veillonellaceae"              
## [2254,] "Selenomonadales"             "Veillonellaceae"              
## [2255,] "Synergistales"               "Synergistaceae"               
## [2256,] "Bacteroidales"               "Prevotellaceae"               
## [2257,] "Bacteroidales"               "Prevotellaceae"               
## [2258,] "Clostridiales"               "Peptostreptococcaceae"        
## [2259,] "Corynebacteriales"           "Corynebacteriaceae"           
## [2260,] "Bacillales"                  "Family_XI"                    
## [2261,] "Selenomonadales"             "Veillonellaceae"              
## [2262,] "Pasteurellales"              "Pasteurellaceae"              
## [2263,] "Betaproteobacteriales"       "Neisseriaceae"                
## [2264,] "Fusobacteriales"             "Leptotrichiaceae"             
## [2265,] "Bacteroidales"               "Prevotellaceae"               
## [2266,] "Fusobacteriales"             "Leptotrichiaceae"             
## [2267,] "Selenomonadales"             "Veillonellaceae"              
## [2268,] "Betaproteobacteriales"       "Neisseriaceae"                
## [2269,] "Flavobacteriales"            "Flavobacteriaceae"            
## [2270,] "Bacteroidales"               "Prevotellaceae"               
## [2271,] "Fusobacteriales"             "Leptotrichiaceae"             
## [2272,] "Absconditabacteriales_(SR1)" NA                             
## [2273,] "Fusobacteriales"             NA                             
## [2274,] "Clostridiales"               "Lachnospiraceae"              
## [2275,] "Bacteroidales"               "Porphyromonadaceae"           
## [2276,] "Flavobacteriales"            "Flavobacteriaceae"            
## [2277,] "Clostridiales"               "Lachnospiraceae"              
## [2278,] "Selenomonadales"             "Veillonellaceae"              
## [2279,] "Pasteurellales"              "Pasteurellaceae"              
## [2280,] NA                            NA                             
## [2281,] "Betaproteobacteriales"       "Neisseriaceae"                
## [2282,] "Clostridiales"               "Lachnospiraceae"              
## [2283,] "Bacteroidales"               "Prevotellaceae"               
## [2284,] "Bacteroidales"               "Prevotellaceae"               
## [2285,] "Bacteroidales"               "Prevotellaceae"               
## [2286,] "Clostridiales"               "Lachnospiraceae"              
## [2287,] "Bacteroidales"               "Prevotellaceae"               
## [2288,] "Sphingobacteriales"          "Lentimicrobiaceae"            
## [2289,] "Absconditabacteriales_(SR1)" NA                             
## [2290,] "Fusobacteriales"             "Leptotrichiaceae"             
## [2291,] "Selenomonadales"             "Veillonellaceae"              
## [2292,] "Pasteurellales"              "Pasteurellaceae"              
## [2293,] "Bacteroidales"               "Prevotellaceae"               
## [2294,] "Bacteroidales"               "Prevotellaceae"               
## [2295,] "Bacteroidales"               "Porphyromonadaceae"           
## [2296,] "Betaproteobacteriales"       "Neisseriaceae"                
## [2297,] "Bacteroidales"               "Prevotellaceae"               
## [2298,] "Bacteroidales"               "Prevotellaceae"               
## [2299,] "Bacteroidales"               "Prevotellaceae"               
## [2300,] "Bacteroidales"               "Prevotellaceae"               
## [2301,] "Selenomonadales"             "Veillonellaceae"              
## [2302,] "Clostridiales"               "Peptostreptococcaceae"        
## [2303,] "Bifidobacteriales"           "Bifidobacteriaceae"           
## [2304,] "Clostridiales"               "Peptostreptococcaceae"        
## [2305,] "Spirochaetales"              "Spirochaetaceae"              
## [2306,] "Flavobacteriales"            "Weeksellaceae"                
## [2307,] "Saccharimonadales"           "Saccharimonadaceae"           
## [2308,] "Flavobacteriales"            "Flavobacteriaceae"            
## [2309,] "Clostridiales"               "Peptostreptococcaceae"        
## [2310,] "Bacteroidales"               "Prevotellaceae"               
## [2311,] "Clostridiales"               "Lachnospiraceae"              
## [2312,] "Selenomonadales"             "Veillonellaceae"              
## [2313,] "Micrococcales"               NA                             
## [2314,] "Bacteroidales"               "Prevotellaceae"               
## [2315,] "Fusobacteriales"             NA                             
## [2316,] "Micrococcales"               NA                             
## [2317,] NA                            NA                             
## [2318,] "Bacteroidales"               "Porphyromonadaceae"           
## [2319,] "Clostridiales"               "Lachnospiraceae"              
## [2320,] "Saccharimonadales"           NA                             
## [2321,] "Bacteroidales"               "Prevotellaceae"               
## [2322,] "Bacteroidales"               "Prevotellaceae"               
## [2323,] "Bacteroidales"               "Prevotellaceae"               
## [2324,] "Bacteroidales"               "Prevotellaceae"               
## [2325,] "Bacteroidales"               "Prevotellaceae"               
## [2326,] NA                            NA                             
## [2327,] "Fusobacteriales"             "Leptotrichiaceae"             
## [2328,] "Bacteroidales"               "Prevotellaceae"               
## [2329,] "Clostridiales"               "Ruminococcaceae"              
## [2330,] "Bacteroidales"               "Prevotellaceae"               
## [2331,] "Bacteroidales"               "Porphyromonadaceae"           
## [2332,] NA                            NA                             
## [2333,] "Bacteroidales"               "Prevotellaceae"               
## [2334,] "Bacteroidales"               "Prevotellaceae"               
## [2335,] "Betaproteobacteriales"       "Neisseriaceae"                
## [2336,] "Betaproteobacteriales"       "Neisseriaceae"                
## [2337,] "Bacteroidales"               "Prevotellaceae"               
## [2338,] "Bacteroidales"               "Prevotellaceae"               
## [2339,] "Lactobacillales"             "Streptococcaceae"             
## [2340,] "Bacteroidales"               "Prevotellaceae"               
## [2341,] "Saccharimonadales"           "Saccharimonadaceae"           
## [2342,] "Clostridiales"               "Lachnospiraceae"              
## [2343,] "Pasteurellales"              "Pasteurellaceae"              
## [2344,] "Bacteroidales"               "Porphyromonadaceae"           
## [2345,] "Bacteroidales"               "Porphyromonadaceae"           
## [2346,] "Micrococcales"               NA                             
## [2347,] "Selenomonadales"             "Veillonellaceae"              
## [2348,] "Corynebacteriales"           "Corynebacteriaceae"           
## [2349,] "Synergistales"               "Synergistaceae"               
## [2350,] "Spirochaetales"              "Spirochaetaceae"              
## [2351,] "Bacteroidales"               "Prevotellaceae"               
## [2352,] "Bacteroidales"               "Prevotellaceae"               
## [2353,] "Actinomycetales"             "Actinomycetaceae"             
## [2354,] "Micrococcales"               NA                             
## [2355,] "Clostridiales"               "Lachnospiraceae"              
## [2356,] "Bacteroidales"               "Prevotellaceae"               
## [2357,] "Selenomonadales"             "Veillonellaceae"              
## [2358,] "Bacteroidales"               "Prevotellaceae"               
## [2359,] "Bacteroidales"               "Prevotellaceae"               
## [2360,] "Selenomonadales"             "Veillonellaceae"              
## [2361,] "Enterobacteriales"           "Enterobacteriaceae"           
## [2362,] "Selenomonadales"             "Veillonellaceae"              
## [2363,] "Bacteroidales"               "Prevotellaceae"               
## [2364,] "Selenomonadales"             "Veillonellaceae"              
## [2365,] "Selenomonadales"             "Veillonellaceae"              
## [2366,] "Fusobacteriales"             "Fusobacteriaceae"             
## [2367,] "Actinomycetales"             "Actinomycetaceae"             
## [2368,] "Bacteroidales"               "Prevotellaceae"               
## [2369,] "Betaproteobacteriales"       "Neisseriaceae"                
## [2370,] "Fusobacteriales"             "Leptotrichiaceae"             
## [2371,] "Bacteroidales"               "Porphyromonadaceae"           
## [2372,] "Selenomonadales"             "Veillonellaceae"              
## [2373,] "Bacteroidales"               "Prevotellaceae"               
## [2374,] "Bacteroidales"               "Prevotellaceae"               
## [2375,] "Selenomonadales"             "Veillonellaceae"              
## [2376,] "Clostridiales"               "Peptostreptococcaceae"        
## [2377,] "Betaproteobacteriales"       "Neisseriaceae"                
## [2378,] "Spirochaetales"              "Spirochaetaceae"              
## [2379,] "Bacteroidales"               "Prevotellaceae"               
## [2380,] "Fusobacteriales"             "Fusobacteriaceae"             
## [2381,] "Bacteroidales"               "Prevotellaceae"               
## [2382,] "Fusobacteriales"             "Leptotrichiaceae"             
## [2383,] "Clostridiales"               "Family_XIII"                  
## [2384,] "Bacteroidales"               "Paludibacteraceae"            
## [2385,] "Pasteurellales"              "Pasteurellaceae"              
## [2386,] "Bacteroidales"               "Prevotellaceae"               
## [2387,] "Selenomonadales"             "Veillonellaceae"              
## [2388,] "Actinomycetales"             "Actinomycetaceae"             
## [2389,] "Bacteroidales"               "Prevotellaceae"               
## [2390,] "Selenomonadales"             "Veillonellaceae"              
## [2391,] "Fusobacteriales"             "Leptotrichiaceae"             
## [2392,] "Selenomonadales"             "Veillonellaceae"              
## [2393,] "Fusobacteriales"             "Leptotrichiaceae"             
## [2394,] "Saccharimonadales"           "Saccharimonadaceae"           
## [2395,] "Selenomonadales"             "Veillonellaceae"              
## [2396,] "Selenomonadales"             "Veillonellaceae"              
## [2397,] "Selenomonadales"             "Veillonellaceae"              
## [2398,] "Corynebacteriales"           "Corynebacteriaceae"           
## [2399,] "Actinomycetales"             "Actinomycetaceae"             
## [2400,] "Bacteroidales"               "Prevotellaceae"               
## [2401,] "Coriobacteriales"            "Atopobiaceae"                 
## [2402,] "Spirochaetales"              "Spirochaetaceae"              
## [2403,] "Bacteroidales"               "Prevotellaceae"               
## [2404,] "Fusobacteriales"             "Fusobacteriaceae"             
## [2405,] "Clostridiales"               "Lachnospiraceae"              
## [2406,] "Selenomonadales"             "Veillonellaceae"              
## [2407,] "Bacteroidales"               "Prevotellaceae"               
## [2408,] "Bacteroidales"               "Prevotellaceae"               
## [2409,] "Fusobacteriales"             "Leptotrichiaceae"             
## [2410,] "Bacteroidales"               "Prevotellaceae"               
## [2411,] "Selenomonadales"             "Veillonellaceae"              
## [2412,] NA                            NA                             
## [2413,] "Saccharimonadales"           "Saccharimonadaceae"           
## [2414,] "Bacteroidales"               "Prevotellaceae"               
## [2415,] "Selenomonadales"             "Veillonellaceae"              
## [2416,] "Selenomonadales"             "Veillonellaceae"              
## [2417,] "Selenomonadales"             "Veillonellaceae"              
## [2418,] "Bacteroidales"               "Prevotellaceae"               
## [2419,] "Bacteroidales"               "Prevotellaceae"               
## [2420,] "Clostridiales"               "Lachnospiraceae"              
## [2421,] "Pasteurellales"              "Pasteurellaceae"              
## [2422,] "Micrococcales"               NA                             
## [2423,] "Micrococcales"               NA                             
## [2424,] NA                            NA                             
## [2425,] "Bacteroidales"               "Prevotellaceae"               
## [2426,] "Chloroplast"                 NA                             
## [2427,] "Bacteroidales"               "Prevotellaceae"               
## [2428,] "Bacteroidales"               "Porphyromonadaceae"           
## [2429,] "Flavobacteriales"            "Flavobacteriaceae"            
## [2430,] "Bacteroidales"               "Prevotellaceae"               
## [2431,] "Flavobacteriales"            "Flavobacteriaceae"            
## [2432,] "Clostridiales"               "Lachnospiraceae"              
## [2433,] "Selenomonadales"             "Veillonellaceae"              
## [2434,] "Bacteroidales"               "Prevotellaceae"               
## [2435,] NA                            NA                             
## [2436,] "Bacteroidales"               "Prevotellaceae"               
## [2437,] "Spirochaetales"              "Spirochaetaceae"              
## [2438,] NA                            NA                             
## [2439,] "Bacteroidales"               "Prevotellaceae"               
## [2440,] "Chloroplast"                 NA                             
## [2441,] "Actinomycetales"             "Actinomycetaceae"             
## [2442,] "Bacteroidales"               "Porphyromonadaceae"           
## [2443,] "Betaproteobacteriales"       "Neisseriaceae"                
## [2444,] "Pasteurellales"              "Pasteurellaceae"              
## [2445,] "Clostridiales"               "Ruminococcaceae"              
## [2446,] "Bacteroidales"               "Prevotellaceae"               
## [2447,] "Bacteroidales"               "Prevotellaceae"               
## [2448,] "Lactobacillales"             "Streptococcaceae"             
## [2449,] "Saccharimonadales"           "Saccharimonadaceae"           
## [2450,] "Bacteroidales"               "Prevotellaceae"               
## [2451,] NA                            NA                             
## [2452,] "Bacteroidales"               "Prevotellaceae"               
## [2453,] "Betaproteobacteriales"       "Neisseriaceae"                
## [2454,] "Bacteroidales"               "Prevotellaceae"               
## [2455,] "Clostridiales"               "Lachnospiraceae"              
## [2456,] "Actinomycetales"             "Actinomycetaceae"             
## [2457,] "Betaproteobacteriales"       "Neisseriaceae"                
## [2458,] "Bacteroidales"               "Prevotellaceae"               
## [2459,] "Clostridiales"               "Lachnospiraceae"              
## [2460,] "Spirochaetales"              "Spirochaetaceae"              
## [2461,] "Clostridiales"               "Lachnospiraceae"              
## [2462,] "Selenomonadales"             "Veillonellaceae"              
## [2463,] "Bacteroidales"               "Prevotellaceae"               
## [2464,] "Clostridiales"               "Peptostreptococcaceae"        
## [2465,] "Bacteroidales"               "Prevotellaceae"               
## [2466,] "Bacteroidales"               "Prevotellaceae"               
## [2467,] "Betaproteobacteriales"       "Neisseriaceae"                
## [2468,] "Flavobacteriales"            "Weeksellaceae"                
## [2469,] "Bacteroidales"               "Prevotellaceae"               
## [2470,] "Bacteroidales"               "Prevotellaceae"               
## [2471,] "Betaproteobacteriales"       "Neisseriaceae"                
## [2472,] "Pasteurellales"              "Pasteurellaceae"              
## [2473,] "Selenomonadales"             "Veillonellaceae"              
## [2474,] "Clostridiales"               "Peptococcaceae"               
## [2475,] "Bacteroidales"               "Prevotellaceae"               
## [2476,] "Bacteroidales"               "Prevotellaceae"               
## [2477,] "Saccharimonadales"           NA                             
## [2478,] "Selenomonadales"             "Veillonellaceae"              
## [2479,] "Selenomonadales"             "Veillonellaceae"              
## [2480,] "Micrococcales"               NA                             
## [2481,] "Micrococcales"               NA                             
## [2482,] "Clostridiales"               "Lachnospiraceae"              
## [2483,] "Bacteroidales"               "Prevotellaceae"               
## [2484,] "Spirochaetales"              "Spirochaetaceae"              
## [2485,] "Betaproteobacteriales"       "Neisseriaceae"                
## [2486,] "Clostridiales"               "Family_XIII"                  
## [2487,] "Bacteroidales"               "Prevotellaceae"               
## [2488,] "Bacteroidales"               "Prevotellaceae"               
## [2489,] "Pasteurellales"              "Pasteurellaceae"              
## [2490,] "Bacteroidales"               "Prevotellaceae"               
## [2491,] "Fusobacteriales"             "Fusobacteriaceae"             
## [2492,] "Selenomonadales"             "Veillonellaceae"              
## [2493,] "Sphingobacteriales"          "Lentimicrobiaceae"            
## [2494,] "Bacteroidales"               "Prevotellaceae"               
## [2495,] "Flavobacteriales"            "Flavobacteriaceae"            
## [2496,] "Fusobacteriales"             "Fusobacteriaceae"             
## [2497,] "Actinomycetales"             "Actinomycetaceae"             
## [2498,] "Pasteurellales"              "Pasteurellaceae"              
## [2499,] "Bacteroidales"               NA                             
## [2500,] "Corynebacteriales"           "Corynebacteriaceae"           
## [2501,] "Selenomonadales"             "Veillonellaceae"              
## [2502,] "Bacteroidales"               "Prevotellaceae"               
## [2503,] "Bacteroidales"               "Prevotellaceae"               
## [2504,] "Enterobacteriales"           "Enterobacteriaceae"           
## [2505,] "Fusobacteriales"             "Fusobacteriaceae"             
## [2506,] "Bacteroidales"               "Prevotellaceae"               
## [2507,] "Flavobacteriales"            "Flavobacteriaceae"            
## [2508,] "Bacteroidales"               "Prevotellaceae"               
## [2509,] "Lactobacillales"             "Streptococcaceae"             
## [2510,] "Saccharimonadales"           "Saccharimonadaceae"           
## [2511,] "Flavobacteriales"            "Flavobacteriaceae"            
## [2512,] "Betaproteobacteriales"       "Neisseriaceae"                
## [2513,] "Clostridiales"               "Lachnospiraceae"              
## [2514,] "Clostridiales"               "Peptostreptococcaceae"        
## [2515,] "Bacteroidales"               "Prevotellaceae"               
## [2516,] "Betaproteobacteriales"       "Neisseriaceae"                
## [2517,] "Micrococcales"               "Micrococcaceae"               
## [2518,] "Fusobacteriales"             "Leptotrichiaceae"             
## [2519,] "Actinomycetales"             "Actinomycetaceae"             
## [2520,] "Actinomycetales"             "Actinomycetaceae"             
## [2521,] "Betaproteobacteriales"       "Neisseriaceae"                
## [2522,] "Fusobacteriales"             "Fusobacteriaceae"             
## [2523,] "Bacteroidales"               "Prevotellaceae"               
## [2524,] "Selenomonadales"             "Veillonellaceae"              
## [2525,] "Betaproteobacteriales"       "Neisseriaceae"                
## [2526,] "Micrococcales"               NA                             
## [2527,] "Bacteroidales"               "Prevotellaceae"               
## [2528,] "Lactobacillales"             "Streptococcaceae"             
## [2529,] "Fusobacteriales"             NA                             
## [2530,] "Bacteroidales"               "Tannerellaceae"               
## [2531,] "Selenomonadales"             "Veillonellaceae"              
## [2532,] "Selenomonadales"             "Veillonellaceae"              
## [2533,] "Bacteroidales"               "Prevotellaceae"               
## [2534,] "Flavobacteriales"            "Flavobacteriaceae"            
## [2535,] "Bacteroidales"               "Prevotellaceae"               
## [2536,] "Pasteurellales"              "Pasteurellaceae"              
## [2537,] "Lactobacillales"             "Streptococcaceae"             
## [2538,] "Sphingobacteriales"          "Lentimicrobiaceae"            
## [2539,] "Micrococcales"               NA                             
## [2540,] "Pasteurellales"              "Pasteurellaceae"              
## [2541,] "Bacteroidales"               "Prevotellaceae"               
## [2542,] "Clostridiales"               "Ruminococcaceae"              
## [2543,] "Fusobacteriales"             "Fusobacteriaceae"             
## [2544,] "Bacteroidales"               "Prevotellaceae"               
## [2545,] NA                            NA                             
## [2546,] "Fusobacteriales"             "Fusobacteriaceae"             
## [2547,] "Fusobacteriales"             "Leptotrichiaceae"             
## [2548,] "Micrococcales"               NA                             
## [2549,] "Selenomonadales"             "Veillonellaceae"              
## [2550,] "Clostridiales"               "Peptostreptococcaceae"        
## [2551,] "Selenomonadales"             "Veillonellaceae"              
## [2552,] "Bacteroidales"               "Prevotellaceae"               
## [2553,] "Fusobacteriales"             "Leptotrichiaceae"             
## [2554,] "Fusobacteriales"             "Leptotrichiaceae"             
## [2555,] "Fusobacteriales"             "Leptotrichiaceae"             
## [2556,] "Flavobacteriales"            "Flavobacteriaceae"            
## [2557,] "Pasteurellales"              "Pasteurellaceae"              
## [2558,] "Spirochaetales"              "Spirochaetaceae"              
## [2559,] "Bacteroidales"               NA                             
## [2560,] "Selenomonadales"             "Veillonellaceae"              
## [2561,] "Selenomonadales"             "Veillonellaceae"              
## [2562,] "Spirochaetales"              "Spirochaetaceae"              
## [2563,] "Spirochaetales"              "Spirochaetaceae"              
## [2564,] "Betaproteobacteriales"       "Neisseriaceae"                
## [2565,] "Clostridiales"               "Family_XI"                    
## [2566,] "Lactobacillales"             "Carnobacteriaceae"            
## [2567,] "Pasteurellales"              "Pasteurellaceae"              
## [2568,] "Lactobacillales"             "Streptococcaceae"             
## [2569,] "Bacteroidales"               "Prevotellaceae"               
## [2570,] "Selenomonadales"             "Veillonellaceae"              
## [2571,] "Bacteroidales"               "Prevotellaceae"               
## [2572,] "Bacteroidales"               "Prevotellaceae"               
## [2573,] "Pasteurellales"              "Pasteurellaceae"              
## [2574,] "Fusobacteriales"             "Leptotrichiaceae"             
## [2575,] "Saccharimonadales"           "Saccharimonadaceae"           
## [2576,] "Bacteroidales"               "Prevotellaceae"               
## [2577,] "Actinomycetales"             "Actinomycetaceae"             
## [2578,] "Fusobacteriales"             "Leptotrichiaceae"             
## [2579,] "Flavobacteriales"            "Flavobacteriaceae"            
## [2580,] "Bacteroidales"               "Porphyromonadaceae"           
## [2581,] "Bacteroidales"               "Porphyromonadaceae"           
## [2582,] "Fusobacteriales"             "Fusobacteriaceae"             
## [2583,] "Bacteroidales"               "Prevotellaceae"               
## [2584,] "Selenomonadales"             "Veillonellaceae"              
## [2585,] "Flavobacteriales"            "Flavobacteriaceae"            
## [2586,] "Saccharimonadales"           NA                             
## [2587,] "Bacteroidales"               "Rikenellaceae"                
## [2588,] "Clostridiales"               "Lachnospiraceae"              
## [2589,] "Clostridiales"               "Peptostreptococcaceae"        
## [2590,] "Bacteroidales"               "Prevotellaceae"               
## [2591,] "Campylobacterales"           NA                             
## [2592,] "Selenomonadales"             "Veillonellaceae"              
## [2593,] "Lactobacillales"             "Lactobacillaceae"             
## [2594,] "Betaproteobacteriales"       "Neisseriaceae"                
## [2595,] "Bacteroidales"               "Prevotellaceae"               
## [2596,] "Lactobacillales"             "Streptococcaceae"             
## [2597,] "Bacteroidales"               "Prevotellaceae"               
## [2598,] NA                            NA                             
## [2599,] "Sphingobacteriales"          "Lentimicrobiaceae"            
## [2600,] "Selenomonadales"             "Veillonellaceae"              
## [2601,] "Selenomonadales"             "Veillonellaceae"              
## [2602,] "Cardiobacteriales"           "Cardiobacteriaceae"           
## [2603,] "Bacteroidales"               "Prevotellaceae"               
## [2604,] "Selenomonadales"             "Veillonellaceae"              
## [2605,] "Bacteroidales"               "Prevotellaceae"               
## [2606,] "Betaproteobacteriales"       "Neisseriaceae"                
## [2607,] "Micrococcales"               NA                             
## [2608,] NA                            NA                             
## [2609,] NA                            NA                             
## [2610,] "Micrococcales"               NA                             
## [2611,] "Clostridiales"               "Peptostreptococcaceae"        
## [2612,] "Bacteroidales"               "Prevotellaceae"               
## [2613,] "Micrococcales"               NA                             
## [2614,] "Bacteroidales"               "Prevotellaceae"               
## [2615,] NA                            NA                             
## [2616,] "Bacteroidales"               "Prevotellaceae"               
## [2617,] "Pasteurellales"              "Pasteurellaceae"              
## [2618,] "Selenomonadales"             "Veillonellaceae"              
## [2619,] "Pasteurellales"              "Pasteurellaceae"              
## [2620,] "Fusobacteriales"             "Leptotrichiaceae"             
## [2621,] "Fusobacteriales"             "Fusobacteriaceae"             
## [2622,] "Bacteroidales"               "Prevotellaceae"               
## [2623,] "Flavobacteriales"            "Flavobacteriaceae"            
## [2624,] "Enterobacteriales"           "Enterobacteriaceae"           
## [2625,] "Clostridiales"               "Lachnospiraceae"              
## [2626,] "Absconditabacteriales_(SR1)" NA                             
## [2627,] "Selenomonadales"             "Veillonellaceae"              
## [2628,] "Lactobacillales"             "Carnobacteriaceae"            
## [2629,] "Corynebacteriales"           "Corynebacteriaceae"           
## [2630,] "Clostridiales"               "Lachnospiraceae"              
## [2631,] NA                            NA                             
## [2632,] "Spirochaetales"              "Spirochaetaceae"              
## [2633,] "Bacillales"                  "Family_XI"                    
## [2634,] "Bacteroidales"               "Prevotellaceae"               
## [2635,] "Campylobacterales"           NA                             
## [2636,] "Selenomonadales"             "Veillonellaceae"              
## [2637,] "Campylobacterales"           NA                             
## [2638,] "Campylobacterales"           NA                             
## [2639,] "Bacteroidales"               "Prevotellaceae"               
## [2640,] "Fusobacteriales"             "Leptotrichiaceae"             
## [2641,] "Campylobacterales"           "Campylobacteraceae"           
## [2642,] "Selenomonadales"             "Veillonellaceae"              
## [2643,] "Bacteroidales"               "Prevotellaceae"               
## [2644,] "Lactobacillales"             "Streptococcaceae"             
## [2645,] "Pasteurellales"              "Pasteurellaceae"              
## [2646,] "Clostridiales"               "Family_XI"                    
## [2647,] "Bacteroidales"               "Prevotellaceae"               
## [2648,] "Clostridiales"               "Ruminococcaceae"              
## [2649,] "Flavobacteriales"            "Flavobacteriaceae"            
## [2650,] "Lactobacillales"             "Streptococcaceae"             
## [2651,] "Spirochaetales"              "Spirochaetaceae"              
## [2652,] "Pasteurellales"              "Pasteurellaceae"              
## [2653,] "Fusobacteriales"             "Leptotrichiaceae"             
## [2654,] "Bacteroidales"               "Prevotellaceae"               
## [2655,] "Selenomonadales"             "Veillonellaceae"              
## [2656,] "Bacteroidales"               "Tannerellaceae"               
## [2657,] "Bacteroidales"               "Prevotellaceae"               
## [2658,] "Clostridiales"               "Peptostreptococcaceae"        
## [2659,] "Bacteroidales"               "Prevotellaceae"               
## [2660,] "Flavobacteriales"            "Flavobacteriaceae"            
## [2661,] "Flavobacteriales"            "Flavobacteriaceae"            
## [2662,] "Selenomonadales"             "Veillonellaceae"              
## [2663,] "Actinomycetales"             "Actinomycetaceae"             
## [2664,] "Actinomycetales"             "Actinomycetaceae"             
## [2665,] "Fusobacteriales"             "Leptotrichiaceae"             
## [2666,] "Selenomonadales"             "Veillonellaceae"              
## [2667,] NA                            NA                             
## [2668,] "Selenomonadales"             "Veillonellaceae"              
## [2669,] "Clostridiales"               "Family_XI"                    
## [2670,] "Flavobacteriales"            "Flavobacteriaceae"            
## [2671,] "Selenomonadales"             "Veillonellaceae"              
## [2672,] "Pasteurellales"              "Pasteurellaceae"              
## [2673,] "Bacteroidales"               "Prevotellaceae"               
## [2674,] "Bacteroidales"               "Prevotellaceae"               
## [2675,] "Spirochaetales"              "Spirochaetaceae"              
## [2676,] "Bacteroidales"               "Prevotellaceae"               
## [2677,] "Actinomycetales"             "Actinomycetaceae"             
## [2678,] "Mollicutes_RF39"             NA                             
## [2679,] "Enterobacteriales"           "Enterobacteriaceae"           
## [2680,] "Bacteroidales"               "Prevotellaceae"               
## [2681,] "Betaproteobacteriales"       "Burkholderiaceae"             
## [2682,] "Bacteroidales"               "Prevotellaceae"               
## [2683,] "Actinomycetales"             "Actinomycetaceae"             
## [2684,] "Flavobacteriales"            "Flavobacteriaceae"            
## [2685,] "Bacteroidales"               "Prevotellaceae"               
## [2686,] "Fusobacteriales"             "Leptotrichiaceae"             
## [2687,] "Coriobacteriales"            "Atopobiaceae"                 
## [2688,] "Saccharimonadales"           "Saccharimonadaceae"           
## [2689,] "Micrococcales"               NA                             
## [2690,] NA                            NA                             
## [2691,] "Clostridiales"               "Lachnospiraceae"              
## [2692,] "Fusobacteriales"             "Fusobacteriaceae"             
## [2693,] "Clostridiales"               "Clostridiales_vadinBB60_group"
## [2694,] "Bacteroidales"               "Prevotellaceae"               
## [2695,] "Bacteroidales"               "Prevotellaceae"               
## [2696,] NA                            NA                             
## [2697,] "Selenomonadales"             "Veillonellaceae"              
## [2698,] "Selenomonadales"             "Veillonellaceae"              
## [2699,] "Selenomonadales"             "Veillonellaceae"              
## [2700,] "Bacteroidales"               "Prevotellaceae"               
## [2701,] NA                            NA                             
## [2702,] "Selenomonadales"             "Veillonellaceae"              
## [2703,] "Sphingobacteriales"          "Lentimicrobiaceae"            
## [2704,] "Bacteroidales"               "Prevotellaceae"               
## [2705,] "Pasteurellales"              "Pasteurellaceae"              
## [2706,] "Fusobacteriales"             "Fusobacteriaceae"             
## [2707,] "Selenomonadales"             "Veillonellaceae"              
## [2708,] "Fusobacteriales"             "Fusobacteriaceae"             
## [2709,] "Selenomonadales"             "Veillonellaceae"              
## [2710,] "Bacteroidales"               "Prevotellaceae"               
## [2711,] "Flavobacteriales"            "Flavobacteriaceae"            
## [2712,] "Fusobacteriales"             "Fusobacteriaceae"             
## [2713,] "Selenomonadales"             "Veillonellaceae"              
## [2714,] "Fusobacteriales"             "Fusobacteriaceae"             
## [2715,] "Bacteroidales"               "Prevotellaceae"               
## [2716,] "Fusobacteriales"             "Leptotrichiaceae"             
## [2717,] "Fusobacteriales"             "Fusobacteriaceae"             
## [2718,] "Bacteroidales"               "Prevotellaceae"               
## [2719,] "Enterobacteriales"           "Enterobacteriaceae"           
## [2720,] "Clostridiales"               "Lachnospiraceae"              
## [2721,] "Spirochaetales"              "Spirochaetaceae"              
## [2722,] "Fusobacteriales"             "Leptotrichiaceae"             
## [2723,] "Clostridiales"               "Lachnospiraceae"              
## [2724,] "Bacteroidales"               "Prevotellaceae"               
## [2725,] "Bacteroidales"               "Prevotellaceae"               
## [2726,] "Betaproteobacteriales"       "Neisseriaceae"                
## [2727,] "Bacteroidales"               "Prevotellaceae"               
## [2728,] "Actinomycetales"             "Actinomycetaceae"             
## [2729,] "Bacteroidales"               "Prevotellaceae"               
## [2730,] "Campylobacterales"           NA                             
## [2731,] "Clostridiales"               "Lachnospiraceae"              
## [2732,] "Betaproteobacteriales"       "Neisseriaceae"                
## [2733,] "Bacteroidales"               "Prevotellaceae"               
## [2734,] "Fusobacteriales"             "Leptotrichiaceae"             
## [2735,] "Fusobacteriales"             "Fusobacteriaceae"             
## [2736,] "Fusobacteriales"             "Leptotrichiaceae"             
## [2737,] "Pasteurellales"              "Pasteurellaceae"              
## [2738,] "Bacteroidales"               "Prevotellaceae"               
## [2739,] "Micrococcales"               NA                             
## [2740,] NA                            NA                             
## [2741,] NA                            NA                             
## [2742,] "Bacteroidales"               "Prevotellaceae"               
## [2743,] "Erysipelotrichales"          "Erysipelotrichaceae"          
## [2744,] "Selenomonadales"             "Veillonellaceae"              
## [2745,] "Betaproteobacteriales"       "Neisseriaceae"                
## [2746,] "Betaproteobacteriales"       "Neisseriaceae"                
## [2747,] "Selenomonadales"             "Veillonellaceae"              
## [2748,] "Bacteroidales"               "Prevotellaceae"               
## [2749,] "Bacteroidales"               "Prevotellaceae"               
## [2750,] "Bacteroidales"               "Prevotellaceae"               
## [2751,] "Saccharimonadales"           "Saccharimonadaceae"           
## [2752,] "Bacteroidales"               "Prevotellaceae"               
## [2753,] "Bacteroidales"               "Tannerellaceae"               
## [2754,] "Bacteroidales"               "Prevotellaceae"               
## [2755,] "Betaproteobacteriales"       "Neisseriaceae"                
## [2756,] NA                            NA                             
## [2757,] "Flavobacteriales"            "Flavobacteriaceae"            
## [2758,] "Fusobacteriales"             "Fusobacteriaceae"             
## [2759,] "Bacteroidales"               "Prevotellaceae"               
## [2760,] "Coriobacteriales"            "Atopobiaceae"                 
## [2761,] "Bacteroidales"               "Prevotellaceae"               
## [2762,] "Bacteroidales"               "Porphyromonadaceae"           
## [2763,] "Bacteroidales"               "Prevotellaceae"               
## [2764,] "Micrococcales"               NA                             
## [2765,] NA                            NA                             
## [2766,] "Bacteroidales"               "Prevotellaceae"               
## [2767,] "Lactobacillales"             "Streptococcaceae"             
## [2768,] "Lactobacillales"             NA                             
## [2769,] "Clostridiales"               NA                             
## [2770,] "Selenomonadales"             "Veillonellaceae"              
## [2771,] "Saccharimonadales"           "Saccharimonadaceae"           
## [2772,] "Lactobacillales"             "Carnobacteriaceae"            
## [2773,] "Bacteroidales"               "Prevotellaceae"               
## [2774,] "Selenomonadales"             "Veillonellaceae"              
## [2775,] "Campylobacterales"           NA                             
## [2776,] "Corynebacteriales"           "Corynebacteriaceae"           
## [2777,] "Betaproteobacteriales"       "Neisseriaceae"                
## [2778,] "Corynebacteriales"           "Corynebacteriaceae"           
## [2779,] "Bacteroidales"               "Tannerellaceae"               
## [2780,] "Flavobacteriales"            "Weeksellaceae"                
## [2781,] NA                            NA                             
## [2782,] "Flavobacteriales"            "Flavobacteriaceae"            
## [2783,] "Clostridiales"               "Clostridiales_vadinBB60_group"
## [2784,] "Clostridiales"               "Family_XIII"                  
## [2785,] "Corynebacteriales"           "Corynebacteriaceae"           
## [2786,] "Actinomycetales"             "Actinomycetaceae"             
## [2787,] "Bacteroidales"               "Prevotellaceae"               
## [2788,] "Selenomonadales"             "Veillonellaceae"              
## [2789,] NA                            NA                             
## [2790,] "Bacteroidales"               "Prevotellaceae"               
## [2791,] "Bacteroidales"               "Prevotellaceae"               
## [2792,] "Flavobacteriales"            "Weeksellaceae"                
## [2793,] "Selenomonadales"             "Veillonellaceae"              
## [2794,] "Clostridiales"               NA                             
## [2795,] "Betaproteobacteriales"       "Neisseriaceae"                
## [2796,] "Bacteroidales"               "Prevotellaceae"               
## [2797,] "Bacteroidales"               "Prevotellaceae"               
## [2798,] "Flavobacteriales"            "Flavobacteriaceae"            
## [2799,] "Bacteroidales"               "Prevotellaceae"               
## [2800,] "Pasteurellales"              "Pasteurellaceae"              
## [2801,] "Pasteurellales"              "Pasteurellaceae"              
## [2802,] "Flavobacteriales"            "Flavobacteriaceae"            
## [2803,] "Bacteroidales"               "Prevotellaceae"               
## [2804,] "Selenomonadales"             "Veillonellaceae"              
## [2805,] "Clostridiales"               "Family_XIII"                  
## [2806,] "Bacteroidales"               "Prevotellaceae"               
## [2807,] "Bacteroidales"               "Prevotellaceae"               
## [2808,] "Selenomonadales"             "Veillonellaceae"              
## [2809,] "Fusobacteriales"             "Leptotrichiaceae"             
## [2810,] "Bacteroidales"               "Prevotellaceae"               
## [2811,] "Bacteroidales"               "Prevotellaceae"               
## [2812,] "Fusobacteriales"             "Fusobacteriaceae"             
## [2813,] "Pasteurellales"              "Pasteurellaceae"              
## [2814,] "Fusobacteriales"             "Leptotrichiaceae"             
## [2815,] "Bacteroidales"               "Prevotellaceae"               
## [2816,] "Bacteroidales"               "Prevotellaceae"               
## [2817,] "Bacteroidales"               "Prevotellaceae"               
## [2818,] "Selenomonadales"             "Veillonellaceae"              
## [2819,] "Clostridiales"               "Lachnospiraceae"              
## [2820,] "Bacteroidales"               "Prevotellaceae"               
## [2821,] "Selenomonadales"             "Veillonellaceae"              
## [2822,] "Bacteroidales"               "Prevotellaceae"               
## [2823,] "Bacteroidales"               "Prevotellaceae"               
## [2824,] "Bacteroidales"               "Prevotellaceae"               
## [2825,] "Flavobacteriales"            "Weeksellaceae"                
## [2826,] "Bacteroidales"               NA                             
## [2827,] "Selenomonadales"             "Veillonellaceae"              
## [2828,] "Selenomonadales"             "Veillonellaceae"              
## [2829,] "Saccharimonadales"           NA                             
## [2830,] "Clostridiales"               "Peptostreptococcaceae"        
## [2831,] "Saccharimonadales"           "Saccharimonadaceae"           
## [2832,] "Pasteurellales"              "Pasteurellaceae"              
## [2833,] "Pasteurellales"              "Pasteurellaceae"              
## [2834,] "Spirochaetales"              "Spirochaetaceae"              
## [2835,] "Selenomonadales"             "Veillonellaceae"              
## [2836,] "Bacteroidales"               "Prevotellaceae"               
## [2837,] "Selenomonadales"             "Veillonellaceae"              
## [2838,] "Fusobacteriales"             "Fusobacteriaceae"             
## [2839,] "Bacteroidales"               "Prevotellaceae"               
## [2840,] "Clostridiales"               "Family_XI"                    
## [2841,] "Clostridiales"               "Lachnospiraceae"              
## [2842,] "Selenomonadales"             "Veillonellaceae"              
## [2843,] "Betaproteobacteriales"       "Burkholderiaceae"             
## [2844,] "Selenomonadales"             "Veillonellaceae"              
## [2845,] "Bacteroidales"               "Prevotellaceae"               
## [2846,] "Selenomonadales"             "Veillonellaceae"              
## [2847,] "Pasteurellales"              "Pasteurellaceae"              
## [2848,] "Lactobacillales"             "Carnobacteriaceae"            
## [2849,] "Selenomonadales"             "Veillonellaceae"              
## [2850,] "Bacteroidales"               "Prevotellaceae"               
## [2851,] "Betaproteobacteriales"       "Neisseriaceae"                
## [2852,] "Fusobacteriales"             "Fusobacteriaceae"             
## [2853,] "Selenomonadales"             "Veillonellaceae"              
## [2854,] "Bacteroidales"               "Prevotellaceae"               
## [2855,] "Betaproteobacteriales"       "Neisseriaceae"                
## [2856,] "Actinomycetales"             "Actinomycetaceae"             
## [2857,] "Betaproteobacteriales"       "Neisseriaceae"                
## [2858,] "Bacteroidales"               "Porphyromonadaceae"           
## [2859,] "Bacteroidales"               "Prevotellaceae"               
## [2860,] "Betaproteobacteriales"       "Neisseriaceae"                
## [2861,] "Selenomonadales"             "Veillonellaceae"              
## [2862,] "Selenomonadales"             "Veillonellaceae"              
## [2863,] "Flavobacteriales"            "Weeksellaceae"                
## [2864,] "Micrococcales"               NA                             
## [2865,] "Bacteroidales"               "Tannerellaceae"               
## [2866,] "Mollicutes_RF39"             NA                             
## [2867,] "Sphingobacteriales"          "Lentimicrobiaceae"            
## [2868,] "Clostridiales"               "Lachnospiraceae"              
## [2869,] "Micrococcales"               NA                             
## [2870,] "Clostridiales"               "Lachnospiraceae"              
## [2871,] "Bacteroidales"               "Prevotellaceae"               
## [2872,] NA                            NA                             
## [2873,] "Pasteurellales"              "Pasteurellaceae"              
## [2874,] "Bacteroidales"               "Prevotellaceae"               
## [2875,] "Flavobacteriales"            "Flavobacteriaceae"            
## [2876,] "Bacteroidales"               "Prevotellaceae"               
## [2877,] "Enterobacteriales"           "Enterobacteriaceae"           
## [2878,] "Bacteroidales"               "Prevotellaceae"               
## [2879,] "Lactobacillales"             "Streptococcaceae"             
## [2880,] NA                            NA                             
## [2881,] "Sphingobacteriales"          "Lentimicrobiaceae"            
## [2882,] "Cardiobacteriales"           "Cardiobacteriaceae"           
## [2883,] "Selenomonadales"             "Veillonellaceae"              
## [2884,] "Clostridiales"               "Lachnospiraceae"              
## [2885,] "Bacteroidales"               "Prevotellaceae"               
## [2886,] "Clostridiales"               "Peptostreptococcaceae"        
## [2887,] "Clostridiales"               "Lachnospiraceae"              
## [2888,] "Fusobacteriales"             NA                             
## [2889,] "Flavobacteriales"            "Flavobacteriaceae"            
## [2890,] "Bacteroidales"               "Prevotellaceae"               
## [2891,] "Betaproteobacteriales"       "Neisseriaceae"                
## [2892,] "Bacteroidales"               "Prevotellaceae"               
## [2893,] "Bacteroidales"               "Prevotellaceae"               
## [2894,] "Pasteurellales"              "Pasteurellaceae"              
## [2895,] NA                            NA                             
## [2896,] "Fusobacteriales"             "Leptotrichiaceae"             
## [2897,] "Bacteroidales"               "Prevotellaceae"               
## [2898,] "Bacteroidales"               "Prevotellaceae"               
## [2899,] "Saccharimonadales"           NA                             
## [2900,] "Fusobacteriales"             "Fusobacteriaceae"             
## [2901,] "Bacteroidales"               "Prevotellaceae"               
## [2902,] "Selenomonadales"             "Veillonellaceae"              
## [2903,] "Actinomycetales"             "Actinomycetaceae"             
## [2904,] "Bacteroidales"               "Prevotellaceae"               
## [2905,] "Spirochaetales"              "Spirochaetaceae"              
## [2906,] "Pasteurellales"              "Pasteurellaceae"              
## [2907,] "Spirochaetales"              "Spirochaetaceae"              
## [2908,] "Campylobacterales"           "Campylobacteraceae"           
## [2909,] "Clostridiales"               "Lachnospiraceae"              
## [2910,] "Oceanospirillales"           "Halomonadaceae"               
## [2911,] "Selenomonadales"             "Veillonellaceae"              
## [2912,] "Mycoplasmatales"             "Mycoplasmataceae"             
## [2913,] "Bacteroidales"               "Prevotellaceae"               
## [2914,] NA                            NA                             
## [2915,] "Saccharimonadales"           "Saccharimonadaceae"           
## [2916,] "Flavobacteriales"            "Flavobacteriaceae"            
## [2917,] "Saccharimonadales"           "Saccharimonadaceae"           
## [2918,] "Selenomonadales"             "Veillonellaceae"              
## [2919,] "Fusobacteriales"             "Fusobacteriaceae"             
## [2920,] "Selenomonadales"             "Veillonellaceae"              
## [2921,] "Bacteroidales"               "Porphyromonadaceae"           
## [2922,] "Actinomycetales"             "Actinomycetaceae"             
## [2923,] "Betaproteobacteriales"       NA                             
## [2924,] "Lactobacillales"             "Streptococcaceae"             
## [2925,] "Clostridiales"               "Lachnospiraceae"              
## [2926,] "Fusobacteriales"             "Leptotrichiaceae"             
## [2927,] "Betaproteobacteriales"       "Neisseriaceae"                
## [2928,] "Lactobacillales"             "Streptococcaceae"             
## [2929,] "Actinomycetales"             "Actinomycetaceae"             
## [2930,] "Micrococcales"               NA                             
## [2931,] "Clostridiales"               "Ruminococcaceae"              
## [2932,] "Pasteurellales"              "Pasteurellaceae"              
## [2933,] "Lactobacillales"             "Carnobacteriaceae"            
## [2934,] "Coriobacteriales"            "Atopobiaceae"                 
## [2935,] "Actinomycetales"             "Actinomycetaceae"             
## [2936,] "Selenomonadales"             "Veillonellaceae"              
## [2937,] NA                            NA                             
## [2938,] "Fusobacteriales"             "Fusobacteriaceae"             
## [2939,] "Bacteroidales"               "Prevotellaceae"               
## [2940,] "Bacteroidales"               "Prevotellaceae"               
## [2941,] "Micrococcales"               NA                             
## [2942,] "Saccharimonadales"           "Saccharimonadaceae"           
## [2943,] "Bacteroidales"               "Prevotellaceae"               
## [2944,] "Bacillales"                  "Staphylococcaceae"            
## [2945,] "Flavobacteriales"            "Flavobacteriaceae"            
## [2946,] "Sphingobacteriales"          "Lentimicrobiaceae"            
## [2947,] "Bacteroidales"               "Tannerellaceae"               
## [2948,] NA                            NA                             
## [2949,] "Clostridiales"               "Lachnospiraceae"              
## [2950,] "Selenomonadales"             "Veillonellaceae"              
## [2951,] "Selenomonadales"             "Veillonellaceae"              
## [2952,] "Bacteroidales"               "Prevotellaceae"               
## [2953,] "Bacteroidales"               "Prevotellaceae"               
## [2954,] "Flavobacteriales"            "Flavobacteriaceae"            
## [2955,] "Clostridiales"               "Lachnospiraceae"              
## [2956,] "Chloroplast"                 NA                             
## [2957,] "Pasteurellales"              "Pasteurellaceae"              
## [2958,] "Fusobacteriales"             "Leptotrichiaceae"             
## [2959,] "Selenomonadales"             "Veillonellaceae"              
## [2960,] "Selenomonadales"             "Veillonellaceae"              
## [2961,] "Bacteroidales"               "Prevotellaceae"               
## [2962,] "Pasteurellales"              "Pasteurellaceae"              
## [2963,] "Selenomonadales"             "Veillonellaceae"              
## [2964,] "Selenomonadales"             "Veillonellaceae"              
## [2965,] "Betaproteobacteriales"       "Neisseriaceae"                
## [2966,] "Selenomonadales"             "Veillonellaceae"              
## [2967,] "Selenomonadales"             "Veillonellaceae"              
## [2968,] NA                            NA                             
## [2969,] "Bacteroidales"               "Prevotellaceae"               
## [2970,] "Fusobacteriales"             "Leptotrichiaceae"             
## [2971,] "Betaproteobacteriales"       "Neisseriaceae"                
## [2972,] "JGI_0000069-P22"             NA                             
## [2973,] "Bacteroidales"               "Prevotellaceae"               
## [2974,] "Bacteroidales"               "Rikenellaceae"                
## [2975,] "Bacillales"                  "Family_XI"                    
## [2976,] "Spirochaetales"              "Spirochaetaceae"              
## [2977,] "Clostridiales"               "Lachnospiraceae"              
## [2978,] "Mycoplasmatales"             "Mycoplasmataceae"             
## [2979,] "Selenomonadales"             "Veillonellaceae"              
## [2980,] "Bacteroidales"               "Prevotellaceae"               
## [2981,] "Enterobacteriales"           "Enterobacteriaceae"           
## [2982,] "Spirochaetales"              "Spirochaetaceae"              
## [2983,] "Campylobacterales"           "P6-b51"                       
## [2984,] "Bacteroidales"               "Porphyromonadaceae"           
## [2985,] "Clostridiales"               "Lachnospiraceae"              
## [2986,] "Pasteurellales"              "Pasteurellaceae"              
## [2987,] "Actinomycetales"             "Actinomycetaceae"             
## [2988,] "Bacteroidales"               "Prevotellaceae"               
## [2989,] "Flavobacteriales"            "Flavobacteriaceae"            
## [2990,] "Pasteurellales"              "Pasteurellaceae"              
## [2991,] "Selenomonadales"             "Veillonellaceae"              
## [2992,] "Synergistales"               "Synergistaceae"               
## [2993,] "Lactobacillales"             "Streptococcaceae"             
## [2994,] "Bifidobacteriales"           "Bifidobacteriaceae"           
## [2995,] "Bacteroidales"               "Prevotellaceae"               
## [2996,] "Clostridiales"               "Lachnospiraceae"              
## [2997,] "Selenomonadales"             "Veillonellaceae"              
## [2998,] "Saccharimonadales"           NA                             
## [2999,] "Enterobacteriales"           "Enterobacteriaceae"           
## [3000,] "Selenomonadales"             "Veillonellaceae"              
## [3001,] "Flavobacteriales"            "Flavobacteriaceae"            
## [3002,] "Fusobacteriales"             "Leptotrichiaceae"             
## [3003,] NA                            NA                             
## [3004,] "Bacteroidales"               "Prevotellaceae"               
## [3005,] "Betaproteobacteriales"       "Neisseriaceae"                
## [3006,] "Pasteurellales"              "Pasteurellaceae"              
## [3007,] "Selenomonadales"             "Veillonellaceae"              
## [3008,] "Corynebacteriales"           "Corynebacteriaceae"           
## [3009,] "Clostridiales"               "Lachnospiraceae"              
## [3010,] "Selenomonadales"             "Veillonellaceae"              
## [3011,] "Fusobacteriales"             "Fusobacteriaceae"             
## [3012,] "Selenomonadales"             "Veillonellaceae"              
## [3013,] NA                            NA                             
## [3014,] "Bacteroidales"               "Prevotellaceae"               
## [3015,] "Spirochaetales"              "Spirochaetaceae"              
## [3016,] NA                            NA                             
## [3017,] "Pasteurellales"              "Pasteurellaceae"              
## [3018,] "Selenomonadales"             "Veillonellaceae"              
## [3019,] "Bacteroidales"               "Porphyromonadaceae"           
## [3020,] NA                            NA                             
## [3021,] "Bacteroidales"               "Prevotellaceae"               
## [3022,] "Clostridiales"               "Peptostreptococcaceae"        
## [3023,] "Bacteroidales"               "Prevotellaceae"               
## [3024,] "Bacteroidales"               "Prevotellaceae"               
## [3025,] "Bacteroidales"               "Prevotellaceae"               
## [3026,] "Absconditabacteriales_(SR1)" NA                             
## [3027,] "Bacteroidales"               "Prevotellaceae"               
## [3028,] "Absconditabacteriales_(SR1)" NA                             
## [3029,] "Fusobacteriales"             "Fusobacteriaceae"             
## [3030,] "Flavobacteriales"            "Flavobacteriaceae"            
## [3031,] "Bacteroidales"               "Prevotellaceae"               
## [3032,] "Betaproteobacteriales"       "Neisseriaceae"                
## [3033,] "Bacteroidales"               "Prevotellaceae"               
## [3034,] NA                            NA                             
## [3035,] "Lactobacillales"             "Carnobacteriaceae"            
## [3036,] "Clostridiales"               "Lachnospiraceae"              
## [3037,] "Clostridiales"               "Lachnospiraceae"              
## [3038,] "Bacteroidales"               "Prevotellaceae"               
## [3039,] "Actinomycetales"             "Actinomycetaceae"             
## [3040,] "Lactobacillales"             "Streptococcaceae"             
## [3041,] "Campylobacterales"           NA                             
## [3042,] "Bacteroidales"               "Prevotellaceae"               
## [3043,] "Selenomonadales"             "Veillonellaceae"              
## [3044,] "Selenomonadales"             "Veillonellaceae"              
## [3045,] "Flavobacteriales"            "Flavobacteriaceae"            
## [3046,] "Betaproteobacteriales"       NA                             
## [3047,] "Actinomycetales"             "Actinomycetaceae"             
## [3048,] "Pasteurellales"              "Pasteurellaceae"              
## [3049,] "Absconditabacteriales_(SR1)" NA                             
## [3050,] "Clostridiales"               "Lachnospiraceae"              
## [3051,] "Bacteroidales"               "Prevotellaceae"               
## [3052,] "Pseudomonadales"             "Moraxellaceae"                
## [3053,] "Fusobacteriales"             "Fusobacteriaceae"             
## [3054,] "Fusobacteriales"             "Fusobacteriaceae"             
## [3055,] "Clostridiales"               "Peptostreptococcaceae"        
## [3056,] "Bacteroidales"               "Porphyromonadaceae"           
## [3057,] "Bacteroidales"               "Prevotellaceae"               
## [3058,] "Fusobacteriales"             "Leptotrichiaceae"             
## [3059,] "Bacteroidales"               "Prevotellaceae"               
## [3060,] "Flavobacteriales"            "Flavobacteriaceae"            
## [3061,] "Bacteroidales"               "Porphyromonadaceae"           
## [3062,] "Fusobacteriales"             "Leptotrichiaceae"             
## [3063,] "Selenomonadales"             "Veillonellaceae"              
## [3064,] "Clostridiales"               "Lachnospiraceae"              
## [3065,] "Bifidobacteriales"           "Bifidobacteriaceae"           
## [3066,] "Clostridiales"               "Lachnospiraceae"              
## [3067,] "Selenomonadales"             "Veillonellaceae"              
## [3068,] "Selenomonadales"             "Veillonellaceae"              
## [3069,] "Saccharimonadales"           "Saccharimonadaceae"           
## [3070,] "Micrococcales"               NA                             
## [3071,] "Clostridiales"               "Family_XIII"                  
## [3072,] "Selenomonadales"             "Veillonellaceae"              
## [3073,] "Flavobacteriales"            "Flavobacteriaceae"            
## [3074,] "Fusobacteriales"             "Leptotrichiaceae"             
## [3075,] "Bacteroidales"               "Prevotellaceae"               
## [3076,] "Betaproteobacteriales"       "Burkholderiaceae"             
## [3077,] NA                            NA                             
## [3078,] "Fusobacteriales"             "Leptotrichiaceae"             
## [3079,] "Betaproteobacteriales"       "Neisseriaceae"                
## [3080,] "Bacteroidales"               "Porphyromonadaceae"           
## [3081,] "Spirochaetales"              "Spirochaetaceae"              
## [3082,] "Actinomycetales"             "Actinomycetaceae"             
## [3083,] "Micrococcales"               "Micrococcaceae"               
## [3084,] "Bacteroidales"               "Prevotellaceae"               
## [3085,] "Fusobacteriales"             "Fusobacteriaceae"             
## [3086,] NA                            NA                             
## [3087,] "Bacteroidales"               "Prevotellaceae"               
## [3088,] "Betaproteobacteriales"       "Neisseriaceae"                
## [3089,] "Saccharimonadales"           "Saccharimonadaceae"           
## [3090,] "Selenomonadales"             "Veillonellaceae"              
## [3091,] "Selenomonadales"             "Veillonellaceae"              
## [3092,] "Corynebacteriales"           "Corynebacteriaceae"           
## [3093,] "Fusobacteriales"             "Fusobacteriaceae"             
## [3094,] "Actinomycetales"             "Actinomycetaceae"             
## [3095,] "Pasteurellales"              "Pasteurellaceae"              
## [3096,] "Bacteroidales"               "Prevotellaceae"               
## [3097,] "Selenomonadales"             "Veillonellaceae"              
## [3098,] "Selenomonadales"             "Veillonellaceae"              
## [3099,] NA                            NA                             
## [3100,] "Bacteroidales"               "Prevotellaceae"               
## [3101,] "Bacteroidales"               "Prevotellaceae"               
## [3102,] "Spirochaetales"              "Spirochaetaceae"              
## [3103,] NA                            NA                             
## [3104,] "Sphingobacteriales"          "Lentimicrobiaceae"            
## [3105,] NA                            NA                             
## [3106,] "Actinomycetales"             "Actinomycetaceae"             
## [3107,] "Bacteroidales"               "Prevotellaceae"               
## [3108,] "Bacteroidales"               "Prevotellaceae"               
## [3109,] "Pasteurellales"              "Pasteurellaceae"              
## [3110,] "Micrococcales"               NA                             
## [3111,] NA                            NA                             
## [3112,] "Fusobacteriales"             "Leptotrichiaceae"             
## [3113,] "Selenomonadales"             "Veillonellaceae"              
## [3114,] "Clostridiales"               "Lachnospiraceae"              
## [3115,] "Pasteurellales"              "Pasteurellaceae"              
## [3116,] "Clostridiales"               "Family_XIII"                  
## [3117,] "Lactobacillales"             "Streptococcaceae"             
## [3118,] "Flavobacteriales"            "Weeksellaceae"                
## [3119,] "Fusobacteriales"             "Leptotrichiaceae"             
## [3120,] "Bacteroidales"               "Prevotellaceae"               
## [3121,] "Bacteroidales"               "Prevotellaceae"               
## [3122,] "Micrococcales"               NA                             
## [3123,] "Actinomycetales"             "Actinomycetaceae"             
## [3124,] "Bacteroidales"               "Prevotellaceae"               
## [3125,] "Bacteroidales"               "Prevotellaceae"               
## [3126,] "Clostridiales"               "Defluviitaleaceae"            
## [3127,] "Lactobacillales"             "Carnobacteriaceae"            
## [3128,] "Bacteroidales"               "Prevotellaceae"               
## [3129,] "Micrococcales"               NA                             
## [3130,] "Actinomycetales"             "Actinomycetaceae"             
## [3131,] "Selenomonadales"             "Veillonellaceae"              
## [3132,] "Bacteroidales"               "Prevotellaceae"               
## [3133,] "Flavobacteriales"            "Flavobacteriaceae"            
## [3134,] "Pasteurellales"              "Pasteurellaceae"              
## [3135,] "Bacteroidales"               "Prevotellaceae"               
## [3136,] NA                            NA                             
## [3137,] "Campylobacterales"           NA                             
## [3138,] "Flavobacteriales"            "Flavobacteriaceae"            
## [3139,] "Flavobacteriales"            "Flavobacteriaceae"            
## [3140,] "Bacteroidales"               "Porphyromonadaceae"           
## [3141,] "Bacteroidales"               "Prevotellaceae"               
## [3142,] "Bacteroidales"               "Prevotellaceae"               
## [3143,] "Bacteroidales"               "Prevotellaceae"               
## [3144,] "Pasteurellales"              "Pasteurellaceae"              
## [3145,] "Saccharimonadales"           "Saccharimonadaceae"           
## [3146,] "Fusobacteriales"             "Leptotrichiaceae"             
## [3147,] "Fusobacteriales"             "Leptotrichiaceae"             
## [3148,] "Micrococcales"               NA                             
## [3149,] NA                            NA                             
## [3150,] "Spirochaetales"              "Spirochaetaceae"              
## [3151,] "Clostridiales"               "Ruminococcaceae"              
## [3152,] NA                            NA                             
## [3153,] "Spirochaetales"              "Spirochaetaceae"              
## [3154,] "Bacteroidales"               "Prevotellaceae"               
## [3155,] "Bacteroidales"               "Prevotellaceae"               
## [3156,] "Bacteroidales"               "Prevotellaceae"               
## [3157,] "Selenomonadales"             "Veillonellaceae"              
## [3158,] "Clostridiales"               "Lachnospiraceae"              
## [3159,] "Betaproteobacteriales"       "Neisseriaceae"                
## [3160,] "Bacteroidales"               "Tannerellaceae"               
## [3161,] "Campylobacterales"           NA                             
## [3162,] "Cardiobacteriales"           "Cardiobacteriaceae"           
## [3163,] "Bacteroidales"               "Porphyromonadaceae"           
## [3164,] "Bacteroidales"               "Prevotellaceae"               
## [3165,] "Clostridiales"               "Lachnospiraceae"              
## [3166,] "Pasteurellales"              "Pasteurellaceae"              
## [3167,] "Bacteroidales"               "Prevotellaceae"               
## [3168,] "Spirochaetales"              "Spirochaetaceae"              
## [3169,] "Spirochaetales"              "Spirochaetaceae"              
## [3170,] "Fusobacteriales"             "Leptotrichiaceae"             
## [3171,] "Campylobacterales"           "Campylobacteraceae"           
## [3172,] "Mollicutes_RF39"             NA                             
## [3173,] "Actinomycetales"             "Actinomycetaceae"             
## [3174,] "Fusobacteriales"             "Fusobacteriaceae"             
## [3175,] "Chloroplast"                 NA                             
## [3176,] "Campylobacterales"           NA                             
## [3177,] "Bacteroidales"               "Prevotellaceae"               
## [3178,] NA                            NA                             
## [3179,] "Fusobacteriales"             "Leptotrichiaceae"             
## [3180,] "Betaproteobacteriales"       "Neisseriaceae"                
## [3181,] "Flavobacteriales"            "Flavobacteriaceae"            
## [3182,] "Spirochaetales"              "Spirochaetaceae"              
## [3183,] "Fusobacteriales"             "Leptotrichiaceae"             
## [3184,] "Bacteroidales"               "Prevotellaceae"               
## [3185,] "Fusobacteriales"             "Fusobacteriaceae"             
## [3186,] "Bacteroidales"               "Rikenellaceae"                
## [3187,] "Betaproteobacteriales"       "Neisseriaceae"                
## [3188,] "Saccharimonadales"           NA                             
## [3189,] "Flavobacteriales"            "Flavobacteriaceae"            
## [3190,] "Bacteroidales"               "Prevotellaceae"               
## [3191,] "Spirochaetales"              "Spirochaetaceae"              
## [3192,] "Cardiobacteriales"           "Cardiobacteriaceae"           
## [3193,] "Bacteroidales"               "Prevotellaceae"               
## [3194,] "Fusobacteriales"             "Leptotrichiaceae"             
## [3195,] "Pasteurellales"              "Pasteurellaceae"              
## [3196,] "Fusobacteriales"             "Leptotrichiaceae"             
## [3197,] "Bacteroidales"               "Prevotellaceae"               
## [3198,] "Bacteroidales"               "Prevotellaceae"               
## [3199,] "Bacteroidales"               "Prevotellaceae"               
## [3200,] "Campylobacterales"           NA                             
## [3201,] "Pasteurellales"              "Pasteurellaceae"              
## [3202,] "Clostridiales"               "Family_XI"                    
## [3203,] "Campylobacterales"           "P6-b51"                       
## [3204,] "Saccharimonadales"           NA                             
## [3205,] "Micrococcales"               NA                             
## [3206,] "Bacteroidales"               "Prevotellaceae"               
## [3207,] "Flavobacteriales"            "Flavobacteriaceae"            
## [3208,] "Bacteroidales"               "Prevotellaceae"               
## [3209,] NA                            NA                             
## [3210,] "Bacteroidales"               "Prevotellaceae"               
## [3211,] "Corynebacteriales"           "Corynebacteriaceae"           
## [3212,] "Bacteroidales"               "Prevotellaceae"               
## [3213,] "Clostridiales"               "Lachnospiraceae"              
## [3214,] "Clostridiales"               "Peptostreptococcaceae"        
## [3215,] "Bacteroidales"               "Prevotellaceae"               
## [3216,] "Bacteroidales"               "Prevotellaceae"               
## [3217,] "Fusobacteriales"             "Leptotrichiaceae"             
## [3218,] "Bacteroidales"               "Prevotellaceae"               
## [3219,] "Bacteroidales"               "Bacteroidaceae"               
## [3220,] "Selenomonadales"             "Veillonellaceae"              
## [3221,] "Clostridiales"               "Ruminococcaceae"              
## [3222,] "Clostridiales"               "Lachnospiraceae"              
## [3223,] "Lactobacillales"             "Carnobacteriaceae"            
## [3224,] "Clostridiales"               "Lachnospiraceae"              
## [3225,] "Clostridiales"               "Ruminococcaceae"              
## [3226,] "Actinomycetales"             "Actinomycetaceae"             
## [3227,] "Fusobacteriales"             "Fusobacteriaceae"             
## [3228,] "Fusobacteriales"             "Leptotrichiaceae"             
## [3229,] "Selenomonadales"             "Veillonellaceae"              
## [3230,] "Fusobacteriales"             "Leptotrichiaceae"             
## [3231,] NA                            NA                             
## [3232,] "Bacteroidales"               "Prevotellaceae"               
## [3233,] "Fusobacteriales"             "Leptotrichiaceae"             
## [3234,] "Pasteurellales"              "Pasteurellaceae"              
## [3235,] "Bacteroidales"               "Prevotellaceae"               
## [3236,] NA                            NA                             
## [3237,] "Chloroplast"                 NA                             
## [3238,] "Selenomonadales"             "Veillonellaceae"              
## [3239,] "Lactobacillales"             "Streptococcaceae"             
## [3240,] "Bacteroidales"               "Prevotellaceae"               
## [3241,] "Bacteroidales"               "Prevotellaceae"               
## [3242,] "Selenomonadales"             "Veillonellaceae"              
## [3243,] "Bacteroidales"               NA                             
## [3244,] "Clostridiales"               "Lachnospiraceae"              
## [3245,] "Micrococcales"               "Micrococcaceae"               
## [3246,] "Clostridiales"               "Lachnospiraceae"              
## [3247,] "Campylobacterales"           NA                             
## [3248,] "Coriobacteriales"            "Coriobacteriaceae"            
## [3249,] "Bacteroidales"               "Prevotellaceae"               
## [3250,] "Saccharimonadales"           "Saccharimonadaceae"           
## [3251,] "Coriobacteriales"            "Atopobiaceae"                 
## [3252,] "Micrococcales"               NA                             
## [3253,] "Bacteroidales"               "Prevotellaceae"               
## [3254,] "Sphingobacteriales"          "Lentimicrobiaceae"            
## [3255,] "Betaproteobacteriales"       "Burkholderiaceae"             
## [3256,] "Bacteroidales"               "Porphyromonadaceae"           
## [3257,] "Saccharimonadales"           "Saccharimonadaceae"           
## [3258,] "Fusobacteriales"             "Leptotrichiaceae"             
## [3259,] "Lactobacillales"             "Streptococcaceae"             
## [3260,] "Campylobacterales"           "Campylobacteraceae"           
## [3261,] "Campylobacterales"           "Campylobacteraceae"           
## [3262,] "Micrococcales"               NA                             
## [3263,] "Saccharimonadales"           "Saccharimonadaceae"           
## [3264,] "Fusobacteriales"             "Leptotrichiaceae"             
## [3265,] "Micrococcales"               NA                             
## [3266,] "Bacteroidales"               "Prevotellaceae"               
## [3267,] "Bacteroidales"               "Prevotellaceae"               
## [3268,] "Lactobacillales"             "Carnobacteriaceae"            
## [3269,] "Bacteroidales"               "Prevotellaceae"               
## [3270,] "Fusobacteriales"             NA                             
## [3271,] "Bacteroidales"               "Prevotellaceae"               
## [3272,] "Bacteroidales"               "Prevotellaceae"               
## [3273,] "Bacteroidales"               "Prevotellaceae"               
## [3274,] "Saccharimonadales"           "Saccharimonadaceae"           
## [3275,] "Bacteroidales"               "Prevotellaceae"               
## [3276,] "Actinomycetales"             "Actinomycetaceae"             
## [3277,] NA                            NA                             
## [3278,] "Lactobacillales"             "Carnobacteriaceae"            
## [3279,] "Spirochaetales"              "Spirochaetaceae"              
## [3280,] "Bacteroidales"               "Prevotellaceae"               
## [3281,] "Selenomonadales"             "Veillonellaceae"              
## [3282,] "Cardiobacteriales"           "Cardiobacteriaceae"           
## [3283,] "Bacteroidales"               "Prevotellaceae"               
## [3284,] "Bacteroidales"               "Prevotellaceae"               
## [3285,] "Pasteurellales"              "Pasteurellaceae"              
## [3286,] "Pasteurellales"              "Pasteurellaceae"              
## [3287,] "Pasteurellales"              "Pasteurellaceae"              
## [3288,] "Spirochaetales"              "Spirochaetaceae"              
## [3289,] "Clostridiales"               "Lachnospiraceae"              
## [3290,] "Spirochaetales"              "Spirochaetaceae"              
## [3291,] "Bacteroidales"               "Prevotellaceae"               
## [3292,] "Pasteurellales"              "Pasteurellaceae"              
## [3293,] "Selenomonadales"             "Veillonellaceae"              
## [3294,] "Bacteroidales"               "Prevotellaceae"               
## [3295,] "Bacteroidales"               "Prevotellaceae"               
## [3296,] "Bacteroidales"               "Prevotellaceae"               
## [3297,] "Actinomycetales"             "Actinomycetaceae"             
## [3298,] "Bacteroidales"               "Prevotellaceae"               
## [3299,] "Lactobacillales"             "Streptococcaceae"             
## [3300,] "Bacteroidales"               "Prevotellaceae"               
## [3301,] "Bacteroidales"               "Prevotellaceae"               
## [3302,] "Flavobacteriales"            "Weeksellaceae"                
## [3303,] "Bacteroidales"               "Tannerellaceae"               
## [3304,] "Betaproteobacteriales"       "Neisseriaceae"                
## [3305,] "Betaproteobacteriales"       "Neisseriaceae"                
## [3306,] "Bacteroidales"               "Prevotellaceae"               
## [3307,] "Bacteroidales"               "Prevotellaceae"               
## [3308,] "Saccharimonadales"           "Saccharimonadaceae"           
## [3309,] "Pasteurellales"              "Pasteurellaceae"              
## [3310,] "Bacteroidales"               "Prevotellaceae"               
## [3311,] "Bacteroidales"               "Prevotellaceae"               
## [3312,] "Fusobacteriales"             "Leptotrichiaceae"             
## [3313,] "Flavobacteriales"            "Flavobacteriaceae"            
## [3314,] "Spirochaetales"              "Spirochaetaceae"              
## [3315,] "Bacteroidales"               "Prevotellaceae"               
## [3316,] "Saccharimonadales"           "Saccharimonadaceae"           
## [3317,] "Clostridiales"               "Lachnospiraceae"              
## [3318,] "Fusobacteriales"             "Leptotrichiaceae"             
## [3319,] "Betaproteobacteriales"       "Neisseriaceae"                
## [3320,] "Bacteroidales"               "Prevotellaceae"               
## [3321,] "Bacteroidales"               "Prevotellaceae"               
## [3322,] "Fusobacteriales"             "Leptotrichiaceae"             
## [3323,] "Bacteroidales"               "Prevotellaceae"               
## [3324,] "Bacteroidales"               "Prevotellaceae"               
## [3325,] "Bacteroidales"               "Prevotellaceae"               
## [3326,] NA                            NA                             
## [3327,] "Corynebacteriales"           "Corynebacteriaceae"           
## [3328,] "Flavobacteriales"            "Weeksellaceae"                
## [3329,] "Bacteroidales"               "Prevotellaceae"               
## [3330,] "Spirochaetales"              "Spirochaetaceae"              
## [3331,] "Bacteroidales"               "Porphyromonadaceae"           
## [3332,] "Bacteroidales"               "Prevotellaceae"               
## [3333,] "Selenomonadales"             "Veillonellaceae"              
## [3334,] "Micrococcales"               NA                             
## [3335,] "Bacillales"                  "Family_XI"                    
## [3336,] "Cardiobacteriales"           "Cardiobacteriaceae"           
## [3337,] "Bacteroidales"               "Prevotellaceae"               
## [3338,] "Fusobacteriales"             "Leptotrichiaceae"             
## [3339,] "Saccharimonadales"           "Saccharimonadaceae"           
## [3340,] "Saccharimonadales"           "Saccharimonadaceae"           
## [3341,] "Clostridiales"               "Family_XIII"                  
## [3342,] "Fusobacteriales"             "Fusobacteriaceae"             
## [3343,] NA                            NA                             
## [3344,] "Clostridiales"               "Lachnospiraceae"              
## [3345,] "Saccharimonadales"           "Saccharimonadaceae"           
## [3346,] "Campylobacterales"           NA                             
## [3347,] "Bacteroidales"               "Porphyromonadaceae"           
## [3348,] "Betaproteobacteriales"       "Neisseriaceae"                
## [3349,] "Flavobacteriales"            "Weeksellaceae"                
## [3350,] "Fusobacteriales"             "Leptotrichiaceae"             
## [3351,] "Pasteurellales"              "Pasteurellaceae"              
## [3352,] "Betaproteobacteriales"       "Neisseriaceae"                
## [3353,] "Bacteroidales"               "Prevotellaceae"               
## [3354,] "Betaproteobacteriales"       "Neisseriaceae"                
## [3355,] "Betaproteobacteriales"       "Neisseriaceae"                
## [3356,] "Selenomonadales"             "Veillonellaceae"              
## [3357,] "Coriobacteriales"            "Atopobiaceae"                 
## [3358,] "Fusobacteriales"             "Fusobacteriaceae"             
## [3359,] "Spirochaetales"              "Spirochaetaceae"              
## [3360,] "Selenomonadales"             "Veillonellaceae"              
## [3361,] NA                            NA                             
## [3362,] "Bacteroidales"               "Prevotellaceae"               
## [3363,] "Selenomonadales"             "Veillonellaceae"              
## [3364,] "Clostridiales"               "Family_XI"                    
## [3365,] "Bacteroidales"               "Prevotellaceae"               
## [3366,] "Bacteroidales"               "Prevotellaceae"               
## [3367,] "Pasteurellales"              "Pasteurellaceae"              
## [3368,] "Actinomycetales"             "Actinomycetaceae"             
## [3369,] "Flavobacteriales"            "Flavobacteriaceae"            
## [3370,] "Selenomonadales"             "Veillonellaceae"              
## [3371,] "Flavobacteriales"            "Flavobacteriaceae"            
## [3372,] "Fusobacteriales"             "Fusobacteriaceae"             
## [3373,] "Fusobacteriales"             "Leptotrichiaceae"             
## [3374,] "Spirochaetales"              "Spirochaetaceae"              
## [3375,] "Bacteroidales"               "Prevotellaceae"               
## [3376,] "Flavobacteriales"            "Flavobacteriaceae"            
## [3377,] "Bacteroidales"               "Prevotellaceae"               
## [3378,] "Fusobacteriales"             "Leptotrichiaceae"             
## [3379,] "Bacteroidales"               "Rikenellaceae"                
## [3380,] "Flavobacteriales"            "Flavobacteriaceae"            
## [3381,] "Flavobacteriales"            "Flavobacteriaceae"            
## [3382,] "Bacteroidales"               "Prevotellaceae"               
## [3383,] "Clostridiales"               "Lachnospiraceae"              
## [3384,] "Bacteroidales"               "Prevotellaceae"               
## [3385,] "Spirochaetales"              "Spirochaetaceae"              
## [3386,] "Bacteroidales"               "Porphyromonadaceae"           
## [3387,] "Lactobacillales"             "Streptococcaceae"             
## [3388,] "Clostridiales"               "Peptococcaceae"               
## [3389,] "Bacteroidales"               "Prevotellaceae"               
## [3390,] "Bacteroidales"               "Prevotellaceae"               
## [3391,] "Lactobacillales"             "Carnobacteriaceae"            
## [3392,] "Flavobacteriales"            "Flavobacteriaceae"            
## [3393,] "Saccharimonadales"           "Saccharimonadaceae"           
## [3394,] NA                            NA                             
## [3395,] "Fusobacteriales"             "Leptotrichiaceae"             
## [3396,] "Micrococcales"               "Micrococcaceae"               
## [3397,] "Bacteroidales"               "Porphyromonadaceae"           
## [3398,] "Micrococcales"               NA                             
## [3399,] "Clostridiales"               "Lachnospiraceae"              
## [3400,] "Bacteroidales"               "Porphyromonadaceae"           
## [3401,] "Enterobacteriales"           "Enterobacteriaceae"           
## [3402,] "Betaproteobacteriales"       "Neisseriaceae"                
## [3403,] "Clostridiales"               "Family_XI"                    
## [3404,] "Flavobacteriales"            "Flavobacteriaceae"            
## [3405,] "Bacteroidales"               "Prevotellaceae"               
## [3406,] "Bacteroidales"               "Prevotellaceae"               
## [3407,] "Bacteroidales"               "Prevotellaceae"               
## [3408,] "Lactobacillales"             "Aerococcaceae"                
## [3409,] "Flavobacteriales"            "Flavobacteriaceae"            
## [3410,] "Pasteurellales"              "Pasteurellaceae"              
## [3411,] "Coriobacteriales"            "Atopobiaceae"                 
## [3412,] "Bacteroidales"               "Prevotellaceae"               
## [3413,] "Spirochaetales"              "Spirochaetaceae"              
## [3414,] "Bacteroidales"               "Prevotellaceae"               
## [3415,] "Bacteroidales"               "Prevotellaceae"               
## [3416,] "Bacteroidales"               "Prevotellaceae"               
## [3417,] "Fusobacteriales"             "Leptotrichiaceae"             
## [3418,] "Bacteroidales"               "Porphyromonadaceae"           
## [3419,] "Selenomonadales"             "Veillonellaceae"              
## [3420,] NA                            NA                             
## [3421,] "Bacteroidales"               "Prevotellaceae"               
## [3422,] "Clostridiales"               "Lachnospiraceae"              
## [3423,] NA                            NA                             
## [3424,] "Bacteroidales"               "Prevotellaceae"               
## [3425,] "Pasteurellales"              "Pasteurellaceae"              
## [3426,] "Bacteroidales"               "Prevotellaceae"               
## [3427,] "Bacteroidales"               "Prevotellaceae"               
## [3428,] "Flavobacteriales"            "Flavobacteriaceae"            
## [3429,] "Betaproteobacteriales"       NA                             
## [3430,] NA                            NA                             
## [3431,] "Bacteroidales"               "Prevotellaceae"               
## [3432,] "Bacteroidales"               "Prevotellaceae"               
## [3433,] "Flavobacteriales"            "Weeksellaceae"                
## [3434,] "Bacteroidales"               "Prevotellaceae"               
## [3435,] NA                            NA                             
## [3436,] "Enterobacteriales"           "Enterobacteriaceae"           
## [3437,] "Pasteurellales"              "Pasteurellaceae"              
## [3438,] "Selenomonadales"             "Veillonellaceae"              
## [3439,] "Pseudomonadales"             "Pseudomonadaceae"             
## [3440,] "Lactobacillales"             "Aerococcaceae"                
## [3441,] "Flavobacteriales"            "Flavobacteriaceae"            
## [3442,] "Coriobacteriales"            "Atopobiaceae"                 
## [3443,] "Clostridiales"               "Peptostreptococcaceae"        
## [3444,] "Selenomonadales"             "Veillonellaceae"              
## [3445,] "Bacteroidales"               "Prevotellaceae"               
## [3446,] "Clostridiales"               "Lachnospiraceae"              
## [3447,] "Campylobacterales"           NA                             
## [3448,] "Bacteroidales"               "Prevotellaceae"               
## [3449,] "Bacteroidales"               "Prevotellaceae"               
## [3450,] "Fusobacteriales"             "Fusobacteriaceae"             
## [3451,] "Bacteroidales"               "Prevotellaceae"               
## [3452,] "Fusobacteriales"             "Leptotrichiaceae"             
## [3453,] "Bacteroidales"               "Prevotellaceae"               
## [3454,] "Bacteroidales"               "Prevotellaceae"               
## [3455,] "Selenomonadales"             "Veillonellaceae"              
## [3456,] "Bacteroidales"               "Porphyromonadaceae"           
## [3457,] "Fusobacteriales"             NA                             
## [3458,] "Selenomonadales"             "Veillonellaceae"              
## [3459,] "Pasteurellales"              "Pasteurellaceae"              
## [3460,] "Micrococcales"               NA                             
## [3461,] "Selenomonadales"             "Veillonellaceae"              
## [3462,] "Betaproteobacteriales"       "Neisseriaceae"                
## [3463,] "Clostridiales"               "Lachnospiraceae"              
## [3464,] "Bacteroidales"               "Prevotellaceae"               
## [3465,] "Bacteroidales"               "Porphyromonadaceae"           
## [3466,] "Bacteroidales"               NA                             
## [3467,] "Selenomonadales"             "Veillonellaceae"              
## [3468,] NA                            NA                             
## [3469,] "Bacteroidales"               "Prevotellaceae"               
## [3470,] "Bacteroidales"               "Prevotellaceae"               
## [3471,] NA                            NA                             
## [3472,] "Betaproteobacteriales"       NA                             
## [3473,] "Clostridiales"               "Lachnospiraceae"              
## [3474,] "Actinomycetales"             "Actinomycetaceae"             
## [3475,] "Bacteroidales"               "Porphyromonadaceae"           
## [3476,] "Bacteroidales"               "Prevotellaceae"               
## [3477,] "Bacteroidales"               "Porphyromonadaceae"           
## [3478,] "Betaproteobacteriales"       NA                             
## [3479,] "Bacteroidales"               NA                             
## [3480,] "Selenomonadales"             "Veillonellaceae"              
## [3481,] "Pasteurellales"              "Pasteurellaceae"              
## [3482,] "Flavobacteriales"            "Flavobacteriaceae"            
## [3483,] "Lactobacillales"             "Carnobacteriaceae"            
## [3484,] "Clostridiales"               "Lachnospiraceae"              
## [3485,] "Bacteroidales"               "Prevotellaceae"               
## [3486,] "Bacteroidales"               "Prevotellaceae"               
## [3487,] "Flavobacteriales"            "Flavobacteriaceae"            
## [3488,] "Selenomonadales"             "Veillonellaceae"              
## [3489,] "Selenomonadales"             "Veillonellaceae"              
## [3490,] "Saccharimonadales"           NA                             
## [3491,] "Bacteroidales"               "Prevotellaceae"               
## [3492,] "Selenomonadales"             "Veillonellaceae"              
## [3493,] "Selenomonadales"             "Veillonellaceae"              
## [3494,] "Micrococcales"               NA                             
## [3495,] NA                            NA                             
## [3496,] "Pasteurellales"              "Pasteurellaceae"              
## [3497,] "Bacteroidales"               "Prevotellaceae"               
## [3498,] "Chloroplast"                 NA                             
## [3499,] "Lactobacillales"             "Streptococcaceae"             
## [3500,] "Bacteroidales"               "Prevotellaceae"               
## [3501,] NA                            NA                             
## [3502,] "Lactobacillales"             "Streptococcaceae"             
## [3503,] "Pasteurellales"              "Pasteurellaceae"              
## [3504,] "Micrococcales"               NA                             
## [3505,] "Bacteroidales"               "Paludibacteraceae"            
## [3506,] "Bacteroidales"               "Prevotellaceae"               
## [3507,] "Fusobacteriales"             "Leptotrichiaceae"             
## [3508,] "Fusobacteriales"             "Fusobacteriaceae"             
## [3509,] "Fusobacteriales"             "Leptotrichiaceae"             
## [3510,] "Saccharimonadales"           "Saccharimonadaceae"           
## [3511,] "Flavobacteriales"            "Flavobacteriaceae"            
## [3512,] "Pasteurellales"              "Pasteurellaceae"              
## [3513,] "Clostridiales"               "Peptococcaceae"               
## [3514,] "Bacteroidales"               "Prevotellaceae"               
## [3515,] "Clostridiales"               "Lachnospiraceae"              
## [3516,] "Selenomonadales"             "Veillonellaceae"              
## [3517,] "Selenomonadales"             "Veillonellaceae"              
## [3518,] "Rickettsiales"               "Mitochondria"                 
## [3519,] "Selenomonadales"             "Veillonellaceae"              
## [3520,] "Pseudomonadales"             "Pseudomonadaceae"             
## [3521,] "Pasteurellales"              "Pasteurellaceae"              
## [3522,] "Bacteroidales"               "Prevotellaceae"               
## [3523,] "Selenomonadales"             "Veillonellaceae"              
## [3524,] "Pasteurellales"              "Pasteurellaceae"              
## [3525,] "Bacteroidales"               "Prevotellaceae"               
## [3526,] "Lactobacillales"             "Carnobacteriaceae"            
## [3527,] "Betaproteobacteriales"       "Neisseriaceae"                
## [3528,] "Betaproteobacteriales"       "Neisseriaceae"                
## [3529,] "Bacteroidales"               "Porphyromonadaceae"           
## [3530,] "Lactobacillales"             "Streptococcaceae"             
## [3531,] "Bacteroidales"               "Prevotellaceae"               
## [3532,] "Bacteroidales"               "Prevotellaceae"               
## [3533,] NA                            NA                             
## [3534,] "Fusobacteriales"             "Fusobacteriaceae"             
## [3535,] NA                            NA                             
## [3536,] NA                            NA                             
## [3537,] "Saccharimonadales"           "Saccharimonadaceae"           
## [3538,] "Selenomonadales"             "Veillonellaceae"              
## [3539,] "Fusobacteriales"             "Leptotrichiaceae"             
## [3540,] "Bacteroidales"               "Prevotellaceae"               
## [3541,] "Bacteroidales"               "Prevotellaceae"               
## [3542,] "Fusobacteriales"             "Leptotrichiaceae"             
## [3543,] "Fusobacteriales"             "Leptotrichiaceae"             
## [3544,] "Betaproteobacteriales"       "Neisseriaceae"                
## [3545,] "Fusobacteriales"             "Leptotrichiaceae"             
## [3546,] "Clostridiales"               "Family_XI"                    
## [3547,] NA                            NA                             
## [3548,] "Bacteroidales"               "Tannerellaceae"               
## [3549,] "Micrococcales"               "Micrococcaceae"               
## [3550,] NA                            NA                             
## [3551,] "Clostridiales"               "Lachnospiraceae"              
## [3552,] "Clostridiales"               "Lachnospiraceae"              
## [3553,] "Betaproteobacteriales"       "Neisseriaceae"                
## [3554,] "Pasteurellales"              "Pasteurellaceae"              
## [3555,] "Spirochaetales"              "Spirochaetaceae"              
## [3556,] "Lactobacillales"             "Streptococcaceae"             
## [3557,] "Lactobacillales"             "Streptococcaceae"             
## [3558,] "Micrococcales"               NA                             
## [3559,] "Micrococcales"               NA                             
## [3560,] "Lactobacillales"             "Lactobacillaceae"             
## [3561,] "Spirochaetales"              "Spirochaetaceae"              
## [3562,] "Clostridiales"               "Lachnospiraceae"              
## [3563,] "Campylobacterales"           NA                             
## [3564,] "Clostridiales"               "Peptostreptococcaceae"        
## [3565,] "Bacteroidales"               "Prevotellaceae"               
## [3566,] "Selenomonadales"             "Veillonellaceae"              
## [3567,] "Betaproteobacteriales"       "Neisseriaceae"                
## [3568,] "Lactobacillales"             "Streptococcaceae"             
## [3569,] "Selenomonadales"             "Veillonellaceae"              
## [3570,] "Bacteroidales"               "Prevotellaceae"               
## [3571,] "Flavobacteriales"            "Flavobacteriaceae"            
## [3572,] "Campylobacterales"           NA                             
## [3573,] "Fusobacteriales"             "Fusobacteriaceae"             
## [3574,] "Bacteroidales"               "Porphyromonadaceae"           
## [3575,] "Selenomonadales"             "Veillonellaceae"              
## [3576,] "Clostridiales"               "Lachnospiraceae"              
## [3577,] "Lactobacillales"             "Streptococcaceae"             
## [3578,] "Clostridiales"               "Lachnospiraceae"              
## [3579,] "Bacteroidales"               NA                             
## [3580,] "Clostridiales"               "Lachnospiraceae"              
## [3581,] "Bacteroidales"               "Prevotellaceae"               
## [3582,] "Spirochaetales"              "Spirochaetaceae"              
## [3583,] "Clostridiales"               "Peptococcaceae"               
## [3584,] "Micrococcales"               NA                             
## [3585,] "Bacteroidales"               "Prevotellaceae"               
## [3586,] "Betaproteobacteriales"       "Neisseriaceae"                
## [3587,] "Bacteroidales"               "Prevotellaceae"               
## [3588,] "Bacteroidales"               "Prevotellaceae"               
## [3589,] NA                            NA                             
## [3590,] "Campylobacterales"           NA                             
## [3591,] "Lactobacillales"             "Carnobacteriaceae"            
## [3592,] "Bacteroidales"               "Prevotellaceae"               
## [3593,] "Lactobacillales"             "Carnobacteriaceae"            
## [3594,] "Bacteroidales"               "Porphyromonadaceae"           
## [3595,] NA                            NA                             
## [3596,] "Fusobacteriales"             "Leptotrichiaceae"             
## [3597,] "Bacteroidales"               "Prevotellaceae"               
## [3598,] "Flavobacteriales"            "Flavobacteriaceae"            
## [3599,] NA                            NA                             
## [3600,] "Campylobacterales"           NA                             
## [3601,] "Bacteroidales"               "Prevotellaceae"               
## [3602,] "Betaproteobacteriales"       NA                             
## [3603,] "Selenomonadales"             "Veillonellaceae"              
## [3604,] "Betaproteobacteriales"       "Neisseriaceae"                
## [3605,] "Micrococcales"               NA                             
## [3606,] "Flavobacteriales"            "Weeksellaceae"                
## [3607,] "Lactobacillales"             "Streptococcaceae"             
## [3608,] "Betaproteobacteriales"       "Burkholderiaceae"             
## [3609,] "Fusobacteriales"             "Leptotrichiaceae"             
## [3610,] "Corynebacteriales"           "Corynebacteriaceae"           
## [3611,] "Bacteroidales"               "Porphyromonadaceae"           
## [3612,] "Clostridiales"               "Lachnospiraceae"              
## [3613,] "Fusobacteriales"             "Leptotrichiaceae"             
## [3614,] "Selenomonadales"             "Veillonellaceae"              
## [3615,] "Fusobacteriales"             "Fusobacteriaceae"             
## [3616,] "Clostridiales"               "Lachnospiraceae"              
## [3617,] "Fusobacteriales"             "Leptotrichiaceae"             
## [3618,] "Clostridiales"               "Peptostreptococcaceae"        
## [3619,] "Betaproteobacteriales"       "Neisseriaceae"                
## [3620,] "Coriobacteriales"            "Atopobiaceae"                 
## [3621,] "Micrococcales"               "Micrococcaceae"               
## [3622,] "Bacteroidales"               "Tannerellaceae"               
## [3623,] "Betaproteobacteriales"       "Neisseriaceae"                
## [3624,] "Bacteroidales"               "Prevotellaceae"               
## [3625,] "Bacteroidales"               "Rikenellaceae"                
## [3626,] "Bacteroidales"               "Prevotellaceae"               
## [3627,] "Pasteurellales"              "Pasteurellaceae"              
## [3628,] "Clostridiales"               "Lachnospiraceae"              
## [3629,] "Bacteroidales"               "Tannerellaceae"               
## [3630,] "Saccharimonadales"           NA                             
## [3631,] "Bacteroidales"               "Porphyromonadaceae"           
## [3632,] "Bacteroidales"               "Prevotellaceae"               
## [3633,] "Betaproteobacteriales"       "Neisseriaceae"                
## [3634,] "Lactobacillales"             NA                             
## [3635,] "Sphingobacteriales"          "Lentimicrobiaceae"            
## [3636,] "Saccharimonadales"           NA                             
## [3637,] "Flavobacteriales"            "Weeksellaceae"                
## [3638,] "Clostridiales"               "Ruminococcaceae"              
## [3639,] "Selenomonadales"             "Veillonellaceae"              
## [3640,] "Selenomonadales"             "Veillonellaceae"              
## [3641,] "Selenomonadales"             "Veillonellaceae"              
## [3642,] "Rhodospirillales"            NA                             
## [3643,] "Cardiobacteriales"           "Cardiobacteriaceae"           
## [3644,] "Bacteroidales"               NA                             
## [3645,] "Betaproteobacteriales"       NA                             
## [3646,] "Fusobacteriales"             "Fusobacteriaceae"             
## [3647,] NA                            NA                             
## [3648,] NA                            NA                             
## [3649,] "Clostridiales"               "Lachnospiraceae"              
## [3650,] "Spirochaetales"              "Spirochaetaceae"              
## [3651,] "Clostridiales"               "Ruminococcaceae"              
## [3652,] "Flavobacteriales"            "Flavobacteriaceae"            
## [3653,] NA                            NA                             
## [3654,] "Flavobacteriales"            "Flavobacteriaceae"            
## [3655,] "Bacteroidales"               "Prevotellaceae"               
## [3656,] "Pasteurellales"              "Pasteurellaceae"              
## [3657,] NA                            NA                             
## [3658,] NA                            NA                             
## [3659,] "Selenomonadales"             "Veillonellaceae"              
## [3660,] "Campylobacterales"           NA                             
## [3661,] "Clostridiales"               "Peptostreptococcaceae"        
## [3662,] "Micrococcales"               NA                             
## [3663,] "Bacteroidales"               "Prevotellaceae"               
## [3664,] "Bacteroidales"               "Prevotellaceae"               
## [3665,] "Bacteroidales"               "Prevotellaceae"               
## [3666,] "Selenomonadales"             "Veillonellaceae"              
## [3667,] "Pasteurellales"              "Pasteurellaceae"              
## [3668,] "Pasteurellales"              "Pasteurellaceae"              
## [3669,] NA                            NA                             
## [3670,] "Coriobacteriales"            "Atopobiaceae"                 
## [3671,] NA                            NA                             
## [3672,] NA                            NA                             
## [3673,] "Flavobacteriales"            "Flavobacteriaceae"            
## [3674,] "Bacteroidales"               "Prevotellaceae"               
## [3675,] "Clostridiales"               "Lachnospiraceae"              
## [3676,] "Bacteroidales"               "Prevotellaceae"               
## [3677,] "Spirochaetales"              "Spirochaetaceae"              
## [3678,] "Bacteroidales"               "Prevotellaceae"               
## [3679,] "Flavobacteriales"            "Weeksellaceae"                
## [3680,] "Bacteroidales"               "Prevotellaceae"               
## [3681,] "Selenomonadales"             "Veillonellaceae"              
## [3682,] "Bacteroidales"               "Prevotellaceae"               
## [3683,] "Betaproteobacteriales"       "Neisseriaceae"                
## [3684,] "Pasteurellales"              "Pasteurellaceae"              
## [3685,] "Lactobacillales"             "Aerococcaceae"                
## [3686,] "Clostridiales"               "Lachnospiraceae"              
## [3687,] NA                            NA                             
## [3688,] "Bacteroidales"               "Prevotellaceae"               
## [3689,] "Bacteroidales"               "Prevotellaceae"               
## [3690,] "Clostridiales"               "Lachnospiraceae"              
## [3691,] "Fusobacteriales"             "Leptotrichiaceae"             
## [3692,] "Fusobacteriales"             "Fusobacteriaceae"             
## [3693,] "Sphingobacteriales"          "Lentimicrobiaceae"            
## [3694,] "Spirochaetales"              "Spirochaetaceae"              
## [3695,] "Clostridiales"               "Lachnospiraceae"              
## [3696,] "Selenomonadales"             "Veillonellaceae"              
## [3697,] "Clostridiales"               "Ruminococcaceae"              
## [3698,] "Cardiobacteriales"           "Cardiobacteriaceae"           
## [3699,] "Fusobacteriales"             "Leptotrichiaceae"             
## [3700,] "Bacteroidales"               "Prevotellaceae"               
## [3701,] "Selenomonadales"             "Veillonellaceae"              
## [3702,] "Campylobacterales"           "Campylobacteraceae"           
## [3703,] NA                            NA                             
## [3704,] "Fusobacteriales"             "Leptotrichiaceae"             
## [3705,] "Betaproteobacteriales"       "Neisseriaceae"                
## [3706,] NA                            NA                             
## [3707,] "Bacteroidales"               "Prevotellaceae"               
## [3708,] "Betaproteobacteriales"       "Neisseriaceae"                
## [3709,] "Bacteroidales"               "Porphyromonadaceae"           
## [3710,] "Bacteroidales"               "Prevotellaceae"               
## [3711,] "Clostridiales"               "Lachnospiraceae"              
## [3712,] "Coriobacteriales"            "Atopobiaceae"                 
## [3713,] "Lactobacillales"             "Streptococcaceae"             
## [3714,] "Clostridiales"               "Family_XIII"                  
## [3715,] "Clostridiales"               "Lachnospiraceae"              
## [3716,] "Betaproteobacteriales"       "Burkholderiaceae"             
## [3717,] "Lactobacillales"             "Aerococcaceae"                
## [3718,] "Fusobacteriales"             "Fusobacteriaceae"             
## [3719,] "Saccharimonadales"           "Saccharimonadaceae"           
## [3720,] "Lactobacillales"             "Carnobacteriaceae"            
## [3721,] "Selenomonadales"             "Veillonellaceae"              
## [3722,] "Lactobacillales"             NA                             
## [3723,] "Pasteurellales"              "Pasteurellaceae"              
## [3724,] "Clostridiales"               "Lachnospiraceae"              
## [3725,] "Clostridiales"               "Lachnospiraceae"              
## [3726,] "Campylobacterales"           NA                             
## [3727,] "Campylobacterales"           NA                             
## [3728,] "Clostridiales"               "Peptostreptococcaceae"        
## [3729,] "Clostridiales"               "Lachnospiraceae"              
## [3730,] "Selenomonadales"             "Veillonellaceae"              
## [3731,] "Lactobacillales"             "Lactobacillaceae"             
## [3732,] "Spirochaetales"              "Spirochaetaceae"              
## [3733,] "Selenomonadales"             "Veillonellaceae"              
## [3734,] "Sphingobacteriales"          "Lentimicrobiaceae"            
## [3735,] "Fusobacteriales"             "Fusobacteriaceae"             
## [3736,] "Bacillales"                  "Family_XI"                    
## [3737,] "Bacteroidales"               "Prevotellaceae"               
## [3738,] "Clostridiales"               NA                             
## [3739,] "Bacteroidales"               "Porphyromonadaceae"           
## [3740,] "Bacteroidales"               "Prevotellaceae"               
## [3741,] "Bacteroidales"               "Prevotellaceae"               
## [3742,] "Betaproteobacteriales"       "Neisseriaceae"                
## [3743,] "Campylobacterales"           "Campylobacteraceae"           
## [3744,] "Lactobacillales"             "Streptococcaceae"             
## [3745,] "Saccharimonadales"           NA                             
## [3746,] "Flavobacteriales"            "Flavobacteriaceae"            
## [3747,] "Bacteroidales"               "Prevotellaceae"               
## [3748,] "Bacteroidales"               "Prevotellaceae"               
## [3749,] "Bacteroidales"               "Rikenellaceae"                
## [3750,] "Campylobacterales"           NA                             
## [3751,] "Clostridiales"               "Lachnospiraceae"              
## [3752,] "Lactobacillales"             "Streptococcaceae"             
## [3753,] "Selenomonadales"             "Veillonellaceae"              
## [3754,] "Bacteroidales"               "Prevotellaceae"               
## [3755,] "Clostridiales"               "Lachnospiraceae"              
## [3756,] "Clostridiales"               "Lachnospiraceae"              
## [3757,] "Bacteroidales"               "Prevotellaceae"               
## [3758,] "Lactobacillales"             "Streptococcaceae"             
## [3759,] "Selenomonadales"             "Veillonellaceae"              
## [3760,] "Saccharimonadales"           NA                             
## [3761,] "Bacteroidales"               "Prevotellaceae"               
## [3762,] "Lactobacillales"             "Streptococcaceae"             
## [3763,] "Spirochaetales"              "Spirochaetaceae"              
## [3764,] "Flavobacteriales"            "Weeksellaceae"                
## [3765,] "Bacteroidales"               "Prevotellaceae"               
## [3766,] "Bacteroidales"               "Prevotellaceae"               
## [3767,] "Betaproteobacteriales"       "Burkholderiaceae"             
## [3768,] "Selenomonadales"             "Veillonellaceae"              
## [3769,] "Corynebacteriales"           "Corynebacteriaceae"           
## [3770,] "Pasteurellales"              "Pasteurellaceae"              
## [3771,] "Bacteroidales"               "Prevotellaceae"               
## [3772,] "Saccharimonadales"           "Saccharimonadaceae"           
## [3773,] "Clostridiales"               "Peptostreptococcaceae"        
## [3774,] "Clostridiales"               "Lachnospiraceae"              
## [3775,] "Selenomonadales"             "Veillonellaceae"              
## [3776,] "Bacteroidales"               "Prevotellaceae"               
## [3777,] "Bacteroidales"               "Prevotellaceae"               
## [3778,] "Bacteroidales"               "Prevotellaceae"               
## [3779,] "Bacteroidales"               "Prevotellaceae"               
## [3780,] "Clostridiales"               "Family_XIII"                  
## [3781,] "Bacteroidales"               "Rikenellaceae"                
## [3782,] "Bacteroidales"               "Prevotellaceae"               
## [3783,] "Betaproteobacteriales"       "Neisseriaceae"                
## [3784,] "Bacteroidales"               "Prevotellaceae"               
## [3785,] "Fusobacteriales"             NA                             
## [3786,] "Bacteroidales"               "Tannerellaceae"               
## [3787,] NA                            NA                             
## [3788,] "Saccharimonadales"           "Saccharimonadaceae"           
## [3789,] "Betaproteobacteriales"       "Neisseriaceae"                
## [3790,] "Selenomonadales"             "Veillonellaceae"              
## [3791,] "Spirochaetales"              "Spirochaetaceae"              
## [3792,] "Saccharimonadales"           "Saccharimonadaceae"           
## [3793,] "Flavobacteriales"            "Flavobacteriaceae"            
## [3794,] "Bacteroidales"               "Prevotellaceae"               
## [3795,] "Selenomonadales"             "Veillonellaceae"              
## [3796,] "Saccharimonadales"           NA                             
## [3797,] "Absconditabacteriales_(SR1)" NA                             
## [3798,] "Lactobacillales"             "Carnobacteriaceae"            
## [3799,] "Bacteroidales"               "Prevotellaceae"               
## [3800,] "Pasteurellales"              "Pasteurellaceae"              
## [3801,] "Selenomonadales"             "Veillonellaceae"              
## [3802,] "Bacteroidales"               "Prevotellaceae"               
## [3803,] "Betaproteobacteriales"       "Neisseriaceae"                
## [3804,] "Clostridiales"               "Lachnospiraceae"              
## [3805,] "Lactobacillales"             "Streptococcaceae"             
## [3806,] "Saccharimonadales"           "Saccharimonadaceae"           
## [3807,] "Clostridiales"               "Lachnospiraceae"              
## [3808,] "Bacteroidales"               "Prevotellaceae"               
## [3809,] "Micrococcales"               NA                             
## [3810,] "Micrococcales"               NA                             
## [3811,] "Selenomonadales"             "Veillonellaceae"              
## [3812,] "Selenomonadales"             "Veillonellaceae"              
## [3813,] "Lactobacillales"             "Streptococcaceae"             
## [3814,] "Lactobacillales"             "Streptococcaceae"             
## [3815,] "Bacteroidales"               "Prevotellaceae"               
## [3816,] "Flavobacteriales"            "Weeksellaceae"                
## [3817,] "Selenomonadales"             "Veillonellaceae"              
## [3818,] "Selenomonadales"             "Veillonellaceae"              
## [3819,] "Flavobacteriales"            "Flavobacteriaceae"            
## [3820,] "Bacteroidales"               "Prevotellaceae"               
## [3821,] "Absconditabacteriales_(SR1)" NA                             
## [3822,] "Spirochaetales"              "Spirochaetaceae"              
## [3823,] "Bacteroidales"               "Prevotellaceae"               
## [3824,] "Selenomonadales"             "Veillonellaceae"              
## [3825,] "Spirochaetales"              "Spirochaetaceae"              
## [3826,] "Clostridiales"               "Lachnospiraceae"              
## [3827,] "Saccharimonadales"           NA                             
## [3828,] "Pasteurellales"              "Pasteurellaceae"              
## [3829,] "Lactobacillales"             "Lactobacillaceae"             
## [3830,] "Saccharimonadales"           "Saccharimonadaceae"           
## [3831,] "Bacteroidales"               "Prevotellaceae"               
## [3832,] "Betaproteobacteriales"       "Neisseriaceae"                
## [3833,] "Fusobacteriales"             "Leptotrichiaceae"             
## [3834,] "Micrococcales"               "Micrococcaceae"               
## [3835,] "Bacteroidales"               "Porphyromonadaceae"           
## [3836,] "Bacteroidales"               "Prevotellaceae"               
## [3837,] NA                            NA                             
## [3838,] "Bacteroidales"               "Prevotellaceae"               
## [3839,] "Bacteroidales"               "Prevotellaceae"               
## [3840,] "Clostridiales"               "Lachnospiraceae"              
## [3841,] "Bacteroidales"               "Prevotellaceae"               
## [3842,] "Flavobacteriales"            "Flavobacteriaceae"            
## [3843,] "Sphingobacteriales"          NA                             
## [3844,] "Bacteroidales"               "Bacteroidales_Incertae_Sedis" 
## [3845,] NA                            NA                             
## [3846,] "Campylobacterales"           NA                             
## [3847,] "Bacteroidales"               "Prevotellaceae"               
## [3848,] "Clostridiales"               "Lachnospiraceae"              
## [3849,] "Spirochaetales"              "Spirochaetaceae"              
## [3850,] "Clostridiales"               "Lachnospiraceae"              
## [3851,] "Betaproteobacteriales"       "Neisseriaceae"                
## [3852,] "Betaproteobacteriales"       "Neisseriaceae"                
## [3853,] "Bacteroidales"               NA                             
## [3854,] "Lactobacillales"             "Carnobacteriaceae"            
## [3855,] "Campylobacterales"           "Campylobacteraceae"           
## [3856,] "Lactobacillales"             NA                             
## [3857,] "Fusobacteriales"             "Leptotrichiaceae"             
## [3858,] "Bacteroidales"               "Prevotellaceae"               
## [3859,] "Bacteroidales"               "Prevotellaceae"               
## [3860,] "Flavobacteriales"            "Flavobacteriaceae"            
## [3861,] "Lactobacillales"             "Carnobacteriaceae"            
## [3862,] "Clostridiales"               "Lachnospiraceae"              
## [3863,] "Clostridiales"               "Lachnospiraceae"              
## [3864,] "Bacteroidales"               NA                             
## [3865,] "Fusobacteriales"             "Leptotrichiaceae"             
## [3866,] "Clostridiales"               "Lachnospiraceae"              
## [3867,] "Fusobacteriales"             NA                             
## [3868,] "Bacteroidales"               "Prevotellaceae"               
## [3869,] "Saccharimonadales"           "Saccharimonadaceae"           
## [3870,] "Bacteroidales"               "Porphyromonadaceae"           
## [3871,] "Pasteurellales"              "Pasteurellaceae"              
## [3872,] "Betaproteobacteriales"       "Burkholderiaceae"             
## [3873,] "Bacillales"                  NA                             
## [3874,] "Bacteroidales"               "Prevotellaceae"               
## [3875,] NA                            NA                             
## [3876,] "Pasteurellales"              "Pasteurellaceae"              
## [3877,] "Bacteroidales"               "Prevotellaceae"               
## [3878,] "Campylobacterales"           "Campylobacteraceae"           
## [3879,] "Flavobacteriales"            "Weeksellaceae"                
## [3880,] "Clostridiales"               "Family_XIII"                  
## [3881,] "Spirochaetales"              "Spirochaetaceae"              
## [3882,] "Bacteroidales"               "Prevotellaceae"               
## [3883,] "Lactobacillales"             "Carnobacteriaceae"            
## [3884,] "Selenomonadales"             "Veillonellaceae"              
## [3885,] "Pasteurellales"              "Pasteurellaceae"              
## [3886,] "Clostridiales"               "Lachnospiraceae"              
## [3887,] "Clostridiales"               "Family_XI"                    
## [3888,] "Clostridiales"               "Ruminococcaceae"              
## [3889,] "Lactobacillales"             "Streptococcaceae"             
## [3890,] "Clostridiales"               "Family_XIII"                  
## [3891,] "Saccharimonadales"           NA                             
## [3892,] NA                            NA                             
## [3893,] "Clostridiales"               "Lachnospiraceae"              
## [3894,] "Clostridiales"               "Lachnospiraceae"              
## [3895,] "Actinomycetales"             "Actinomycetaceae"             
## [3896,] "Bacteroidales"               "Prevotellaceae"               
## [3897,] "Campylobacterales"           NA                             
## [3898,] "Betaproteobacteriales"       NA                             
## [3899,] "Cardiobacteriales"           "Cardiobacteriaceae"           
## [3900,] "Lactobacillales"             "Carnobacteriaceae"            
## [3901,] "Bacillales"                  "Family_XI"                    
## [3902,] "Selenomonadales"             "Veillonellaceae"              
## [3903,] "Bacteroidales"               "Prevotellaceae"               
## [3904,] "Bacteroidales"               "Porphyromonadaceae"           
## [3905,] "Spirochaetales"              "Spirochaetaceae"              
## [3906,] "Clostridiales"               "Lachnospiraceae"              
## [3907,] "Bacteroidales"               "Prevotellaceae"               
## [3908,] "Pasteurellales"              "Pasteurellaceae"              
## [3909,] "JGI_0000069-P22"             NA                             
## [3910,] "Bacteroidales"               "Prevotellaceae"               
## [3911,] "Bacteroidales"               "Prevotellaceae"               
## [3912,] "Campylobacterales"           NA                             
## [3913,] "Selenomonadales"             "Veillonellaceae"              
## [3914,] "Erysipelotrichales"          "Erysipelotrichaceae"          
## [3915,] "Lactobacillales"             NA                             
## [3916,] "Bacillales"                  "Family_XI"                    
## [3917,] "Bacteroidales"               "Prevotellaceae"               
## [3918,] "Fusobacteriales"             "Fusobacteriaceae"             
## [3919,] "Pasteurellales"              "Pasteurellaceae"              
## [3920,] "Selenomonadales"             "Veillonellaceae"              
## [3921,] "Micrococcales"               NA                             
## [3922,] "Bacteroidales"               "Prevotellaceae"               
## [3923,] NA                            NA                             
## [3924,] "Betaproteobacteriales"       "Burkholderiaceae"             
## [3925,] "Micrococcales"               NA                             
## [3926,] "Selenomonadales"             "Veillonellaceae"              
## [3927,] "Fusobacteriales"             "Leptotrichiaceae"             
## [3928,] "Betaproteobacteriales"       "Burkholderiaceae"             
## [3929,] "Thermales"                   "Thermaceae"                   
## [3930,] "Campylobacterales"           NA                             
## [3931,] "Clostridiales"               "Lachnospiraceae"              
## [3932,] "Clostridiales"               "Family_XI"                    
## [3933,] NA                            NA                             
## [3934,] "Clostridiales"               NA                             
## [3935,] "Betaproteobacteriales"       "Burkholderiaceae"             
## [3936,] "Spirochaetales"              "Spirochaetaceae"              
## [3937,] "Lactobacillales"             "Lactobacillaceae"             
## [3938,] "Pseudomonadales"             "Moraxellaceae"                
## [3939,] "Lactobacillales"             "Lactobacillaceae"             
## [3940,] "Clostridiales"               "Family_XI"                    
## [3941,] "Bacteroidales"               "Prevotellaceae"               
## [3942,] "Bacteroidales"               "Prevotellaceae"               
## [3943,] "Pasteurellales"              "Pasteurellaceae"              
## [3944,] "Clostridiales"               "Lachnospiraceae"              
## [3945,] "Bacteroidales"               "Prevotellaceae"               
## [3946,] "Lactobacillales"             "Carnobacteriaceae"            
## [3947,] "Selenomonadales"             "Veillonellaceae"              
## [3948,] "Lactobacillales"             "Carnobacteriaceae"            
## [3949,] "Bacteroidales"               "Porphyromonadaceae"           
## [3950,] "Lactobacillales"             "Streptococcaceae"             
## [3951,] "Lactobacillales"             "Streptococcaceae"             
## [3952,] "Bacteroidales"               "Prevotellaceae"               
## [3953,] "Fusobacteriales"             "Leptotrichiaceae"             
## [3954,] "Pasteurellales"              "Pasteurellaceae"              
## [3955,] "Campylobacterales"           "Campylobacteraceae"           
## [3956,] NA                            NA                             
## [3957,] "Campylobacterales"           "P6-b51"                       
## [3958,] "Saccharimonadales"           "Saccharimonadaceae"           
## [3959,] "Bacteroidales"               "Prevotellaceae"               
## [3960,] "Selenomonadales"             "Veillonellaceae"              
## [3961,] "Fusobacteriales"             "Fusobacteriaceae"             
## [3962,] "Saccharimonadales"           NA                             
## [3963,] "Bacteroidales"               "Prevotellaceae"               
## [3964,] NA                            NA                             
## [3965,] "Bacteroidales"               "Prevotellaceae"               
## [3966,] "Lactobacillales"             "Carnobacteriaceae"            
## [3967,] NA                            NA                             
## [3968,] "Fusobacteriales"             "Leptotrichiaceae"             
## [3969,] "Fusobacteriales"             "Fusobacteriaceae"             
## [3970,] "Campylobacterales"           "Campylobacteraceae"           
## [3971,] NA                            NA                             
## [3972,] "Acholeplasmatales"           "Acholeplasmataceae"           
## [3973,] "Mycoplasmatales"             "Mycoplasmataceae"             
## [3974,] "Flavobacteriales"            "Weeksellaceae"                
## [3975,] "Desulfovibrionales"          "Desulfovibrionaceae"          
## [3976,] NA                            NA                             
## [3977,] "Bacteroidales"               "Prevotellaceae"               
## [3978,] "Bacteroidales"               "Porphyromonadaceae"           
## [3979,] "Lactobacillales"             "Streptococcaceae"             
## [3980,] "Fusobacteriales"             "Leptotrichiaceae"             
## [3981,] "Clostridiales"               "Lachnospiraceae"              
## [3982,] "Lactobacillales"             "Streptococcaceae"             
## [3983,] "Bacteroidales"               "Prevotellaceae"               
## [3984,] "Selenomonadales"             "Veillonellaceae"              
## [3985,] "Clostridiales"               "Lachnospiraceae"              
## [3986,] "Bacteroidales"               "Prevotellaceae"               
## [3987,] "Bacteroidales"               "Prevotellaceae"               
## [3988,] "Flavobacteriales"            "Flavobacteriaceae"            
## [3989,] "Coriobacteriales"            "Eggerthellaceae"              
## [3990,] "Selenomonadales"             "Veillonellaceae"              
## [3991,] "Fusobacteriales"             "Leptotrichiaceae"             
## [3992,] "Campylobacterales"           NA                             
## [3993,] "Bacteroidales"               "Prevotellaceae"               
## [3994,] "Lactobacillales"             "Streptococcaceae"             
## [3995,] "Spirochaetales"              "Spirochaetaceae"              
## [3996,] "Flavobacteriales"            "Flavobacteriaceae"            
## [3997,] "Clostridiales"               "Peptostreptococcaceae"        
## [3998,] "Sphingobacteriales"          "Lentimicrobiaceae"            
## [3999,] "Saccharimonadales"           NA                             
## [4000,] NA                            NA                             
## [4001,] "Lactobacillales"             "Streptococcaceae"             
## [4002,] "Spirochaetales"              "Spirochaetaceae"              
## [4003,] "Bacillales"                  "Family_XI"                    
## [4004,] "Flavobacteriales"            "Flavobacteriaceae"            
## [4005,] "Fusobacteriales"             "Leptotrichiaceae"             
## [4006,] "Bacteroidales"               "Prevotellaceae"               
## [4007,] "Selenomonadales"             "Veillonellaceae"              
## [4008,] "Bacteroidales"               "Prevotellaceae"               
## [4009,] "Flavobacteriales"            "Weeksellaceae"                
## [4010,] "Pasteurellales"              "Pasteurellaceae"              
## [4011,] "Selenomonadales"             "Veillonellaceae"              
## [4012,] "Bacteroidales"               "Tannerellaceae"               
## [4013,] "Clostridiales"               "Peptostreptococcaceae"        
## [4014,] "Selenomonadales"             "Veillonellaceae"              
## [4015,] "Actinomycetales"             "Actinomycetaceae"             
## [4016,] "Lactobacillales"             "Carnobacteriaceae"            
## [4017,] "Bacteroidales"               "Prevotellaceae"               
## [4018,] "Clostridiales"               "Clostridiales_vadinBB60_group"
## [4019,] "Clostridiales"               "Peptococcaceae"               
## [4020,] "Micrococcales"               "Micrococcaceae"               
## [4021,] "Bacteroidales"               "Prevotellaceae"               
## [4022,] "Actinomycetales"             "Actinomycetaceae"             
## [4023,] "Selenomonadales"             "Veillonellaceae"              
## [4024,] "Saccharimonadales"           NA                             
## [4025,] "Flavobacteriales"            "Flavobacteriaceae"            
## [4026,] "Bacteroidales"               "Prevotellaceae"               
## [4027,] "Lactobacillales"             "Streptococcaceae"             
## [4028,] "Selenomonadales"             "Veillonellaceae"              
## [4029,] "Lactobacillales"             NA                             
## [4030,] "Saccharimonadales"           NA                             
## [4031,] "Betaproteobacteriales"       "Neisseriaceae"                
## [4032,] "Actinomycetales"             "Actinomycetaceae"             
## [4033,] "Spirochaetales"              "Spirochaetaceae"              
## [4034,] "Lactobacillales"             "Carnobacteriaceae"            
## [4035,] "Fusobacteriales"             NA                             
## [4036,] "Clostridiales"               "Eubacteriaceae"               
## [4037,] "Absconditabacteriales_(SR1)" NA                             
## [4038,] "Selenomonadales"             "Veillonellaceae"              
## [4039,] "Betaproteobacteriales"       "Neisseriaceae"                
## [4040,] "Bacillales"                  NA                             
## [4041,] "Bacteroidales"               "Prevotellaceae"               
## [4042,] "Fusobacteriales"             "Fusobacteriaceae"             
## [4043,] "Bacteroidales"               "Prevotellaceae"               
## [4044,] "Flavobacteriales"            "Flavobacteriaceae"            
## [4045,] "Clostridiales"               "Lachnospiraceae"              
## [4046,] "Saccharimonadales"           "Saccharimonadaceae"           
## [4047,] "Bacteroidales"               "Prevotellaceae"               
## [4048,] NA                            NA                             
## [4049,] "Fusobacteriales"             NA                             
## [4050,] "Flavobacteriales"            "Weeksellaceae"                
## [4051,] "Bacteroidales"               "Prevotellaceae"               
## [4052,] "Bacteroidales"               "Prevotellaceae"               
## [4053,] "Lactobacillales"             NA                             
## [4054,] NA                            NA                             
## [4055,] NA                            NA                             
## [4056,] "Fusobacteriales"             "Fusobacteriaceae"             
## [4057,] NA                            NA                             
## [4058,] "Bacteroidales"               NA                             
## [4059,] "Lactobacillales"             "Streptococcaceae"             
## [4060,] "Fusobacteriales"             NA                             
## [4061,] "Chloroplast"                 NA                             
## [4062,] "Actinomycetales"             "Actinomycetaceae"             
## [4063,] "Saccharimonadales"           "Saccharimonadaceae"           
## [4064,] "Fusobacteriales"             "Leptotrichiaceae"             
## [4065,] NA                            NA                             
## [4066,] "Bacteroidales"               "Tannerellaceae"               
## [4067,] "Clostridiales"               "Lachnospiraceae"              
## [4068,] "Clostridiales"               "Lachnospiraceae"              
## [4069,] "Bacteroidales"               NA                             
## [4070,] "Corynebacteriales"           "Corynebacteriaceae"           
## [4071,] "Lactobacillales"             "Streptococcaceae"             
## [4072,] "Absconditabacteriales_(SR1)" NA                             
## [4073,] "Selenomonadales"             "Veillonellaceae"              
## [4074,] "Selenomonadales"             "Veillonellaceae"              
## [4075,] "Selenomonadales"             "Veillonellaceae"              
## [4076,] "Clostridiales"               "Lachnospiraceae"              
## [4077,] "Flavobacteriales"            "Flavobacteriaceae"            
## [4078,] "Pasteurellales"              "Pasteurellaceae"              
## [4079,] "Bacteroidales"               "Prevotellaceae"               
## [4080,] "Flavobacteriales"            "Flavobacteriaceae"            
## [4081,] "Selenomonadales"             "Veillonellaceae"              
## [4082,] "Selenomonadales"             "Veillonellaceae"              
## [4083,] "Clostridiales"               "Lachnospiraceae"              
## [4084,] "Campylobacterales"           NA                             
## [4085,] "Chloroplast"                 NA                             
## [4086,] "Lactobacillales"             NA                             
## [4087,] "Bacteroidales"               NA                             
## [4088,] "Betaproteobacteriales"       "Burkholderiaceae"             
## [4089,] "Corynebacteriales"           "Corynebacteriaceae"           
## [4090,] NA                            NA                             
## [4091,] NA                            NA                             
## [4092,] "Lactobacillales"             NA                             
## [4093,] "Clostridiales"               "Peptostreptococcaceae"        
## [4094,] "Clostridiales"               "Lachnospiraceae"              
## [4095,] "Betaproteobacteriales"       "Neisseriaceae"                
## [4096,] "Clostridiales"               "Ruminococcaceae"              
## [4097,] "Saccharimonadales"           "Saccharimonadaceae"           
## [4098,] "Pasteurellales"              "Pasteurellaceae"              
## [4099,] "Bacteroidales"               "Prevotellaceae"               
## [4100,] "Clostridiales"               "Peptococcaceae"               
## [4101,] "Micrococcales"               NA                             
## [4102,] "Flavobacteriales"            "Weeksellaceae"                
## [4103,] "Clostridiales"               "Family_XIII"                  
## [4104,] "Clostridiales"               "Family_XIII"                  
## [4105,] "Bacteroidales"               "Prevotellaceae"               
## [4106,] "Bacteroidales"               "Prevotellaceae"               
## [4107,] "Saccharimonadales"           NA                             
## [4108,] "Saccharimonadales"           "Saccharimonadaceae"           
## [4109,] "Bacteroidales"               "Porphyromonadaceae"           
## [4110,] "Selenomonadales"             "Veillonellaceae"              
## [4111,] "Lactobacillales"             "Carnobacteriaceae"            
## [4112,] "Selenomonadales"             "Veillonellaceae"              
## [4113,] "Fusobacteriales"             "Leptotrichiaceae"             
## [4114,] "Micrococcales"               NA                             
## [4115,] "Micrococcales"               NA                             
## [4116,] "Clostridiales"               "Peptostreptococcaceae"        
## [4117,] "Coriobacteriales"            "Atopobiaceae"                 
## [4118,] "Bacteroidales"               "Rikenellaceae"                
## [4119,] "Coriobacteriales"            "Atopobiaceae"                 
## [4120,] NA                            NA                             
## [4121,] "Lineage_IV"                  NA                             
## [4122,] "Campylobacterales"           "Rs-M59_termite_group"         
## [4123,] NA                            NA                             
## [4124,] "Campylobacterales"           NA                             
## [4125,] NA                            NA                             
## [4126,] "Saccharimonadales"           NA                             
## [4127,] "Lactobacillales"             "Streptococcaceae"             
## [4128,] "Pasteurellales"              "Pasteurellaceae"              
## [4129,] "Bacteroidales"               "Prevotellaceae"               
## [4130,] "Clostridiales"               "Lachnospiraceae"              
## [4131,] "Lactobacillales"             "Carnobacteriaceae"            
## [4132,] "Clostridiales"               "Peptostreptococcaceae"        
## [4133,] "Flavobacteriales"            "Flavobacteriaceae"            
## [4134,] "Bacteroidales"               "Prevotellaceae"               
## [4135,] "Saccharimonadales"           "Saccharimonadaceae"           
## [4136,] "Saccharimonadales"           "Saccharimonadaceae"           
## [4137,] "Betaproteobacteriales"       NA                             
## [4138,] "Saccharimonadales"           NA                             
## [4139,] "Cardiobacteriales"           "Cardiobacteriaceae"           
## [4140,] "Clostridiales"               "Ruminococcaceae"              
## [4141,] "Betaproteobacteriales"       "Neisseriaceae"                
## [4142,] "Clostridiales"               "Family_XI"                    
## [4143,] "Clostridiales"               "Lachnospiraceae"              
## [4144,] "Bacteroidales"               "Prevotellaceae"               
## [4145,] "Lactobacillales"             "Carnobacteriaceae"            
## [4146,] "Bacteroidales"               "Prevotellaceae"               
## [4147,] "Fusobacteriales"             "Leptotrichiaceae"             
## [4148,] "Selenomonadales"             "Veillonellaceae"              
## [4149,] "Bacteroidales"               "Prevotellaceae"               
## [4150,] "Clostridiales"               "Peptostreptococcaceae"        
## [4151,] "Clostridiales"               "Lachnospiraceae"              
## [4152,] "Spirochaetales"              "Spirochaetaceae"              
## [4153,] "Pasteurellales"              "Pasteurellaceae"              
## [4154,] "Selenomonadales"             "Veillonellaceae"              
## [4155,] "Bacteroidales"               "Prevotellaceae"               
## [4156,] "Corynebacteriales"           "Corynebacteriaceae"           
## [4157,] "Selenomonadales"             "Veillonellaceae"              
## [4158,] NA                            NA                             
## [4159,] "Betaproteobacteriales"       NA                             
## [4160,] "Bacteroidales"               "Prevotellaceae"               
## [4161,] "Clostridiales"               "Lachnospiraceae"              
## [4162,] "Selenomonadales"             "Veillonellaceae"              
## [4163,] "Clostridiales"               "Lachnospiraceae"              
## [4164,] "Saccharimonadales"           NA                             
## [4165,] "Fusobacteriales"             "Fusobacteriaceae"             
## [4166,] "Pasteurellales"              "Pasteurellaceae"              
## [4167,] "Bacillales"                  "Family_XI"                    
## [4168,] "Fusobacteriales"             "Fusobacteriaceae"             
## [4169,] NA                            NA                             
## [4170,] "Selenomonadales"             "Veillonellaceae"              
## [4171,] "Bacteroidales"               "Prevotellaceae"               
## [4172,] "Chloroplast"                 NA                             
## [4173,] "Bacteroidales"               "Prevotellaceae"               
## [4174,] "Clostridiales"               "Lachnospiraceae"              
## [4175,] "Spirochaetales"              "Spirochaetaceae"              
## [4176,] "Betaproteobacteriales"       "Burkholderiaceae"             
## [4177,] "Clostridiales"               "Family_XIII"                  
## [4178,] "Betaproteobacteriales"       "Neisseriaceae"                
## [4179,] "Bacteroidales"               "Prevotellaceae"               
## [4180,] "Saccharimonadales"           NA                             
## [4181,] "Lactobacillales"             "Carnobacteriaceae"            
## [4182,] "Betaproteobacteriales"       "Neisseriaceae"                
## [4183,] "Mycoplasmatales"             "Mycoplasmataceae"             
## [4184,] "Bacteroidales"               "Bacteroidaceae"               
## [4185,] "JGI_0000069-P22"             NA                             
## [4186,] "Clostridiales"               "Lachnospiraceae"              
## [4187,] NA                            NA                             
## [4188,] "Selenomonadales"             "Veillonellaceae"              
## [4189,] NA                            NA                             
## [4190,] "Spirochaetales"              "Spirochaetaceae"              
## [4191,] "Pasteurellales"              "Pasteurellaceae"              
## [4192,] "Betaproteobacteriales"       "Neisseriaceae"                
## [4193,] "Bacillales"                  "Family_XI"                    
## [4194,] "Campylobacterales"           NA                             
## [4195,] "Flavobacteriales"            "Flavobacteriaceae"            
## [4196,] "Clostridiales"               "Family_XIII"                  
## [4197,] "Clostridiales"               "Family_XI"                    
## [4198,] "Selenomonadales"             "Veillonellaceae"              
## [4199,] "Micrococcales"               NA                             
## [4200,] "Pseudomonadales"             "Moraxellaceae"                
## [4201,] "Lactobacillales"             "Streptococcaceae"             
## [4202,] "Fusobacteriales"             "Fusobacteriaceae"             
## [4203,] "Clostridiales"               "Lachnospiraceae"              
## [4204,] "Bacteroidales"               "Prevotellaceae"               
## [4205,] "Selenomonadales"             "Veillonellaceae"              
## [4206,] "Bacteroidales"               NA                             
## [4207,] "Corynebacteriales"           "Corynebacteriaceae"           
## [4208,] "Clostridiales"               "Lachnospiraceae"              
## [4209,] "Fusobacteriales"             "Leptotrichiaceae"             
## [4210,] "Spirochaetales"              "Spirochaetaceae"              
## [4211,] "Saccharimonadales"           "Saccharimonadaceae"           
## [4212,] NA                            NA                             
## [4213,] "Campylobacterales"           NA                             
## [4214,] "Bacteroidales"               "Prevotellaceae"               
## [4215,] "Bacteroidales"               "Prevotellaceae"               
## [4216,] NA                            NA                             
## [4217,] "Saccharimonadales"           "Saccharimonadaceae"           
## [4218,] "Bacteroidales"               "Prevotellaceae"               
## [4219,] "Selenomonadales"             "Veillonellaceae"              
## [4220,] "Campylobacterales"           NA                             
## [4221,] "Betaproteobacteriales"       NA                             
## [4222,] "Fusobacteriales"             NA                             
## [4223,] "Flavobacteriales"            "Weeksellaceae"                
## [4224,] "Flavobacteriales"            "Flavobacteriaceae"            
## [4225,] "Clostridiales"               "Lachnospiraceae"              
## [4226,] "Fusobacteriales"             "Leptotrichiaceae"             
## [4227,] "Bacteroidales"               "Prevotellaceae"               
## [4228,] "Selenomonadales"             "Veillonellaceae"              
## [4229,] "Pasteurellales"              "Pasteurellaceae"              
## [4230,] "Lactobacillales"             "Streptococcaceae"             
## [4231,] "Bacteroidales"               "Prevotellaceae"               
## [4232,] "Pasteurellales"              "Pasteurellaceae"              
## [4233,] "JGI_0000069-P22"             NA                             
## [4234,] "Bacteroidales"               "Prevotellaceae"               
## [4235,] "Lactobacillales"             "Carnobacteriaceae"            
## [4236,] "Saccharimonadales"           "Saccharimonadaceae"           
## [4237,] "Lactobacillales"             "Carnobacteriaceae"            
## [4238,] "Spirochaetales"              "Spirochaetaceae"              
## [4239,] "Corynebacteriales"           "Corynebacteriaceae"           
## [4240,] "Betaproteobacteriales"       "Neisseriaceae"                
## [4241,] "Campylobacterales"           NA                             
## [4242,] "Chloroplast"                 NA                             
## [4243,] "Saccharimonadales"           NA                             
## [4244,] "Lactobacillales"             "Carnobacteriaceae"            
## [4245,] "Bacteroidales"               "Porphyromonadaceae"           
## [4246,] "Actinomycetales"             "Actinomycetaceae"             
## [4247,] "Spirochaetales"              "Spirochaetaceae"              
## [4248,] "Bacillales"                  "Family_XI"                    
## [4249,] "Bacteroidales"               NA                             
## [4250,] "Spirochaetales"              "Spirochaetaceae"              
## [4251,] "Clostridiales"               "Family_XI"                    
## [4252,] NA                            NA                             
## [4253,] "Desulfobacterales"           "Desulfobulbaceae"             
## [4254,] "Izimaplasmatales"            NA                             
## [4255,] "Lactobacillales"             "Streptococcaceae"             
## [4256,] "Bacteroidales"               "Prevotellaceae"               
## [4257,] "Micrococcales"               NA                             
## [4258,] "Campylobacterales"           NA                             
## [4259,] "Flavobacteriales"            "Flavobacteriaceae"            
## [4260,] "Flavobacteriales"            "Weeksellaceae"                
## [4261,] "Saccharimonadales"           "Saccharimonadaceae"           
## [4262,] "Lactobacillales"             NA                             
## [4263,] NA                            NA                             
## [4264,] "Bacteroidales"               "Prevotellaceae"               
## [4265,] "Bacteroidales"               NA                             
## [4266,] "Saccharimonadales"           NA                             
## [4267,] "Micrococcales"               NA                             
## [4268,] "Coriobacteriales"            "Atopobiaceae"                 
## [4269,] "Bacteroidales"               "Prevotellaceae"               
## [4270,] "Lactobacillales"             "Streptococcaceae"             
## [4271,] "Clostridiales"               "Peptostreptococcaceae"        
## [4272,] "Clostridiales"               "Lachnospiraceae"              
## [4273,] NA                            NA                             
## [4274,] "Pasteurellales"              "Pasteurellaceae"              
## [4275,] "Bacteroidales"               "Prevotellaceae"               
## [4276,] "Clostridiales"               "Lachnospiraceae"              
## [4277,] "Clostridiales"               "Family_XIII"                  
## [4278,] "Lactobacillales"             "Streptococcaceae"             
## [4279,] "Lactobacillales"             "Streptococcaceae"             
## [4280,] "Bacteroidales"               "Porphyromonadaceae"           
## [4281,] "Betaproteobacteriales"       NA                             
## [4282,] "Fusobacteriales"             "Leptotrichiaceae"             
## [4283,] "Fusobacteriales"             "Fusobacteriaceae"             
## [4284,] "Betaproteobacteriales"       "Neisseriaceae"                
## [4285,] "Erysipelotrichales"          "Erysipelotrichaceae"          
## [4286,] "Clostridiales"               "Lachnospiraceae"              
## [4287,] "Bacteroidales"               "Bacteroidaceae"               
## [4288,] "Lactobacillales"             "Carnobacteriaceae"            
## [4289,] NA                            NA                             
## [4290,] "Clostridiales"               "Peptostreptococcaceae"        
## [4291,] "Selenomonadales"             "Veillonellaceae"              
## [4292,] "Bacteroidales"               NA                             
## [4293,] "Pasteurellales"              "Pasteurellaceae"              
## [4294,] NA                            NA                             
## [4295,] "Coriobacteriales"            "Atopobiaceae"                 
## [4296,] "Bacteroidales"               "Rikenellaceae"                
## [4297,] "Micrococcales"               "Micrococcaceae"               
## [4298,] NA                            NA                             
## [4299,] "Pasteurellales"              "Pasteurellaceae"              
## [4300,] "Lactobacillales"             "Carnobacteriaceae"            
## [4301,] "Lactobacillales"             NA                             
## [4302,] "Spirochaetales"              "Spirochaetaceae"              
## [4303,] "Flavobacteriales"            "Weeksellaceae"                
## [4304,] "Selenomonadales"             "Veillonellaceae"              
## [4305,] "Rickettsiales"               "Mitochondria"                 
## [4306,] "Clostridiales"               "Lachnospiraceae"              
## [4307,] "Spirochaetales"              "Spirochaetaceae"              
## [4308,] "Lactobacillales"             "Carnobacteriaceae"            
## [4309,] "Pasteurellales"              "Pasteurellaceae"              
## [4310,] "Selenomonadales"             "Veillonellaceae"              
## [4311,] "Betaproteobacteriales"       "Neisseriaceae"                
## [4312,] "Clostridiales"               "Peptostreptococcaceae"        
## [4313,] "Saccharimonadales"           NA                             
## [4314,] "Enterobacteriales"           "Enterobacteriaceae"           
## [4315,] "Bacteroidales"               NA                             
## [4316,] "Clostridiales"               "Lachnospiraceae"              
## [4317,] NA                            NA                             
## [4318,] "Micrococcales"               NA                             
## [4319,] "Spirochaetales"              "Spirochaetaceae"              
## [4320,] "Clostridiales"               "Peptostreptococcaceae"        
## [4321,] "Clostridiales"               "Ruminococcaceae"              
## [4322,] "Pasteurellales"              "Pasteurellaceae"              
## [4323,] "Clostridiales"               "Ruminococcaceae"              
## [4324,] "Campylobacterales"           NA                             
## [4325,] "Saccharimonadales"           "Saccharimonadaceae"           
## [4326,] "Cardiobacteriales"           "Cardiobacteriaceae"           
## [4327,] "Bacteroidales"               "Prevotellaceae"               
## [4328,] "Erysipelotrichales"          "Erysipelotrichaceae"          
## [4329,] "Bifidobacteriales"           "Bifidobacteriaceae"           
## [4330,] "Bacteroidales"               NA                             
## [4331,] "Selenomonadales"             "Veillonellaceae"              
## [4332,] "Betaproteobacteriales"       NA                             
## [4333,] "Betaproteobacteriales"       "Neisseriaceae"                
## [4334,] "Bacteroidales"               NA                             
## [4335,] "Lactobacillales"             "Streptococcaceae"             
## [4336,] "Campylobacterales"           "Helicobacteraceae"            
## [4337,] "Spirochaetales"              "Spirochaetaceae"              
## [4338,] "Clostridiales"               "Lachnospiraceae"              
## [4339,] "Sphingomonadales"            "Sphingomonadaceae"            
## [4340,] "Xanthomonadales"             "Xanthomonadaceae"             
## [4341,] "Betaproteobacteriales"       NA                             
## [4342,] "Bacillales"                  "Family_XI"                    
## [4343,] "Bacteroidales"               "Porphyromonadaceae"           
## [4344,] "Fusobacteriales"             "Leptotrichiaceae"             
## [4345,] "Bacteroidales"               "Rikenellaceae"                
## [4346,] "Bacteroidales"               NA                             
## [4347,] "Clostridiales"               "Family_XI"                    
## [4348,] "Bacteroidales"               "Prevotellaceae"               
## [4349,] "Betaproteobacteriales"       NA                             
## [4350,] "Bacteroidales"               "Tannerellaceae"               
## [4351,] "Rickettsiales"               "Mitochondria"                 
## [4352,] "Clostridiales"               "Defluviitaleaceae"            
## [4353,] "Bacteroidales"               "Prevotellaceae"               
## [4354,] "Bacillales"                  "Family_XI"                    
## [4355,] "Bacteroidales"               "Prevotellaceae"               
## [4356,] "Lactobacillales"             "Streptococcaceae"             
## [4357,] "Betaproteobacteriales"       "Burkholderiaceae"             
## [4358,] "Fusobacteriales"             "Fusobacteriaceae"             
## [4359,] "Pasteurellales"              "Pasteurellaceae"              
## [4360,] "Clostridiales"               "Lachnospiraceae"              
## [4361,] "Bacteroidales"               "Marinilabiliaceae"            
## [4362,] "Fusobacteriales"             "Leptotrichiaceae"             
## [4363,] "Flavobacteriales"            "Weeksellaceae"                
## [4364,] "Clostridiales"               "Peptostreptococcaceae"        
## [4365,] "Betaproteobacteriales"       NA                             
## [4366,] "Betaproteobacteriales"       NA                             
## [4367,] NA                            NA                             
## [4368,] "Campylobacterales"           NA                             
## [4369,] "Clostridiales"               "Family_XI"                    
## [4370,] "Fusobacteriales"             "Fusobacteriaceae"             
## [4371,] "Enterobacteriales"           "Enterobacteriaceae"           
## [4372,] "Cardiobacteriales"           "Cardiobacteriaceae"           
## [4373,] "Selenomonadales"             "Veillonellaceae"              
## [4374,] "Fusobacteriales"             "Leptotrichiaceae"             
## [4375,] "Selenomonadales"             "Veillonellaceae"              
## [4376,] "Spirochaetales"              "Spirochaetaceae"              
## [4377,] "Bacteroidales"               "Prevotellaceae"               
## [4378,] "Lactobacillales"             "Carnobacteriaceae"            
## [4379,] "Clostridiales"               "Lachnospiraceae"              
## [4380,] "Pasteurellales"              "Pasteurellaceae"              
## [4381,] NA                            NA                             
## [4382,] "Bacteroidales"               "Prevotellaceae"               
## [4383,] "Lactobacillales"             "Streptococcaceae"             
## [4384,] "Coriobacteriales"            "Eggerthellaceae"              
## [4385,] "Clostridiales"               "Lachnospiraceae"              
## [4386,] "Bacteroidales"               "Rikenellaceae"                
## [4387,] "Selenomonadales"             "Veillonellaceae"              
## [4388,] "Clostridiales"               "Lachnospiraceae"              
## [4389,] "Micrococcales"               NA                             
## [4390,] "Selenomonadales"             "Veillonellaceae"              
## [4391,] "Campylobacterales"           "P6-b51"                       
## [4392,] "Campylobacterales"           NA                             
## [4393,] "Fusobacteriales"             NA                             
## [4394,] "Selenomonadales"             "Veillonellaceae"              
## [4395,] "Erysipelotrichales"          "Erysipelotrichaceae"          
## [4396,] "Selenomonadales"             "Veillonellaceae"              
## [4397,] "Selenomonadales"             "Veillonellaceae"              
## [4398,] "Selenomonadales"             "Veillonellaceae"              
## [4399,] NA                            NA                             
## [4400,] "Saccharimonadales"           NA                             
## [4401,] "Selenomonadales"             "Veillonellaceae"              
## [4402,] "Fusobacteriales"             "Fusobacteriaceae"             
## [4403,] "Betaproteobacteriales"       "Neisseriaceae"                
## [4404,] "Saccharimonadales"           "Saccharimonadaceae"           
## [4405,] "Coriobacteriales"            "Atopobiaceae"                 
## [4406,] "Bacteroidales"               "Prevotellaceae"               
## [4407,] "Bacteroidales"               NA                             
## [4408,] "Rhizobiales"                 "Beijerinckiaceae"             
## [4409,] "Lactobacillales"             "Carnobacteriaceae"            
## [4410,] NA                            NA                             
## [4411,] "Clostridiales"               "Family_XIII"                  
## [4412,] "Lactobacillales"             "Lactobacillaceae"             
## [4413,] "Coriobacteriales"            "Atopobiaceae"                 
## [4414,] "Spirochaetales"              "Spirochaetaceae"              
## [4415,] "Flavobacteriales"            "Flavobacteriaceae"            
## [4416,] "Campylobacterales"           NA                             
## [4417,] "Synergistales"               "Synergistaceae"               
## [4418,] "Micrococcales"               NA                             
## [4419,] "Micrococcales"               NA                             
## [4420,] "Coriobacteriales"            "Atopobiaceae"                 
## [4421,] "Clostridiales"               "Lachnospiraceae"              
## [4422,] "Mycoplasmatales"             "Mycoplasmataceae"             
## [4423,] "Cardiobacteriales"           "Cardiobacteriaceae"           
## [4424,] "Bacillales"                  "Staphylococcaceae"            
## [4425,] "Campylobacterales"           NA                             
## [4426,] "Micrococcales"               NA                             
## [4427,] "Clostridiales"               "Lachnospiraceae"              
## [4428,] "Selenomonadales"             "Veillonellaceae"              
## [4429,] "Bacteroidales"               "Marinilabiliaceae"            
## [4430,] "Coriobacteriales"            "Atopobiaceae"                 
## [4431,] "Bacteroidales"               "Porphyromonadaceae"           
## [4432,] "Bacteroidales"               "Prevotellaceae"               
## [4433,] "Saccharimonadales"           NA                             
## [4434,] "Clostridiales"               "Peptococcaceae"               
## [4435,] NA                            NA                             
## [4436,] "Micrococcales"               NA                             
## [4437,] NA                            NA                             
## [4438,] "Saccharimonadales"           "Saccharimonadaceae"           
## [4439,] "Pasteurellales"              "Pasteurellaceae"              
## [4440,] "Selenomonadales"             "Veillonellaceae"              
## [4441,] "Selenomonadales"             "Veillonellaceae"              
## [4442,] "Pasteurellales"              "Pasteurellaceae"              
## [4443,] "Erysipelotrichales"          "Erysipelotrichaceae"          
## [4444,] "Cardiobacteriales"           "Cardiobacteriaceae"           
## [4445,] "Flavobacteriales"            "Flavobacteriaceae"            
## [4446,] "Spirochaetales"              "Spirochaetaceae"              
## [4447,] "Flavobacteriales"            "Flavobacteriaceae"            
## [4448,] "Bacillales"                  "Family_XI"                    
## [4449,] "Bacteroidales"               "Porphyromonadaceae"           
## [4450,] "Bacteroidales"               "Prevotellaceae"               
## [4451,] NA                            NA                             
## [4452,] "Bacteroidales"               "Prevotellaceae"               
## [4453,] "Actinomycetales"             "Actinomycetaceae"             
## [4454,] "Pasteurellales"              "Pasteurellaceae"              
## [4455,] "Flavobacteriales"            "Flavobacteriaceae"            
## [4456,] "Fusobacteriales"             "Fusobacteriaceae"             
## [4457,] "Betaproteobacteriales"       "Neisseriaceae"                
## [4458,] "Erysipelotrichales"          "Erysipelotrichaceae"          
## [4459,] "Bacteroidales"               "Prevotellaceae"               
## [4460,] "Pasteurellales"              "Pasteurellaceae"              
## [4461,] "Campylobacterales"           NA                             
## [4462,] "Fusobacteriales"             "Leptotrichiaceae"             
## [4463,] "Lactobacillales"             NA                             
## [4464,] "Bacteroidales"               NA                             
## [4465,] "Betaproteobacteriales"       "Neisseriaceae"                
## [4466,] "Coriobacteriales"            "Atopobiaceae"                 
## [4467,] "Betaproteobacteriales"       "Burkholderiaceae"             
## [4468,] "Bacillales"                  NA                             
## [4469,] NA                            NA                             
## [4470,] "Bacteroidales"               "Prevotellaceae"               
## [4471,] "Clostridiales"               "Lachnospiraceae"              
## [4472,] "Clostridiales"               "Lachnospiraceae"              
## [4473,] "Bacteroidales"               "Prevotellaceae"               
## [4474,] "Bacteroidales"               "Prevotellaceae"               
## [4475,] "Clostridiales"               "Lachnospiraceae"              
## [4476,] "Bacteroidales"               "Porphyromonadaceae"           
## [4477,] "Selenomonadales"             "Veillonellaceae"              
## [4478,] "Bacteroidales"               "Prevotellaceae"               
## [4479,] "Bacteroidales"               "Tannerellaceae"               
## [4480,] "Lactobacillales"             "Streptococcaceae"             
## [4481,] "Betaproteobacteriales"       NA                             
## [4482,] "Clostridiales"               "Peptococcaceae"               
## [4483,] "Lactobacillales"             "Streptococcaceae"             
## [4484,] "Bacteroidales"               "Prevotellaceae"               
## [4485,] "Selenomonadales"             "Veillonellaceae"              
## [4486,] "Erysipelotrichales"          "Erysipelotrichaceae"          
## [4487,] "Saccharimonadales"           NA                             
## [4488,] "Saccharimonadales"           NA                             
## [4489,] "Spirochaetales"              "Spirochaetaceae"              
## [4490,] "Clostridiales"               "Lachnospiraceae"              
## [4491,] "Clostridiales"               "Peptostreptococcaceae"        
## [4492,] "Betaproteobacteriales"       "Burkholderiaceae"             
## [4493,] "Clostridiales"               "Lachnospiraceae"              
## [4494,] "Pasteurellales"              "Pasteurellaceae"              
## [4495,] "Bacteroidales"               "Prevotellaceae"               
## [4496,] "Pseudomonadales"             "Pseudomonadaceae"             
## [4497,] "Rhizobiales"                 "Xanthobacteraceae"            
## [4498,] "Bacillales"                  "Family_XI"                    
## [4499,] "Bacteroidales"               "Prevotellaceae"               
## [4500,] "Selenomonadales"             "Veillonellaceae"              
## [4501,] "Blastocatellales"            "Blastocatellaceae"            
## [4502,] "Lactobacillales"             "Streptococcaceae"             
## [4503,] "Betaproteobacteriales"       "Neisseriaceae"                
## [4504,] "Clostridiales"               "Lachnospiraceae"              
## [4505,] "Fusobacteriales"             "Leptotrichiaceae"             
## [4506,] "Bacteroidales"               "Prevotellaceae"               
## [4507,] "Bacteroidales"               "Prevotellaceae"               
## [4508,] "Clostridiales"               "Lachnospiraceae"              
## [4509,] "Saccharimonadales"           "Saccharimonadaceae"           
## [4510,] "Pasteurellales"              "Pasteurellaceae"              
## [4511,] "Bacteroidales"               "Prevotellaceae"               
## [4512,] NA                            NA                             
## [4513,] "Micrococcales"               NA                             
## [4514,] "Lactobacillales"             "Carnobacteriaceae"            
## [4515,] "Fusobacteriales"             "Fusobacteriaceae"             
## [4516,] "Saccharimonadales"           "Saccharimonadaceae"           
## [4517,] "Fusobacteriales"             "Leptotrichiaceae"             
## [4518,] "Selenomonadales"             "Veillonellaceae"              
## [4519,] "Bacteroidales"               "Prevotellaceae"               
## [4520,] "Lactobacillales"             "Streptococcaceae"             
## [4521,] "Betaproteobacteriales"       "Neisseriaceae"                
## [4522,] "Bacteroidales"               "Prevotellaceae"               
## [4523,] "Bacteroidales"               "Prevotellaceae"               
## [4524,] "Betaproteobacteriales"       "Burkholderiaceae"             
## [4525,] "Clostridiales"               "Family_XI"                    
## [4526,] "Flavobacteriales"            "Flavobacteriaceae"            
## [4527,] "Saccharimonadales"           "Saccharimonadaceae"           
## [4528,] "Bacteroidales"               "Prevotellaceae"               
## [4529,] "Selenomonadales"             "Veillonellaceae"              
## [4530,] NA                            NA                             
## [4531,] "Bacteroidales"               "Porphyromonadaceae"           
## [4532,] "Bacteroidales"               "Porphyromonadaceae"           
## [4533,] NA                            NA                             
## [4534,] "Pasteurellales"              "Pasteurellaceae"              
## [4535,] "Bacteroidales"               "Porphyromonadaceae"           
## [4536,] "Micrococcales"               NA                             
## [4537,] "Coriobacteriales"            NA                             
## [4538,] "Lactobacillales"             "Streptococcaceae"             
## [4539,] "Bacteroidales"               "Bacteroidaceae"               
## [4540,] "Bacteroidales"               NA                             
## [4541,] "Clostridiales"               "Lachnospiraceae"              
## [4542,] "Clostridiales"               "Lachnospiraceae"              
## [4543,] NA                            NA                             
## [4544,] "Bacteroidales"               "Prevotellaceae"               
## [4545,] "Campylobacterales"           NA                             
## [4546,] "Spirochaetales"              "Spirochaetaceae"              
## [4547,] NA                            NA                             
## [4548,] "Bacteroidales"               "Rikenellaceae"                
## [4549,] "Selenomonadales"             "Veillonellaceae"              
## [4550,] "Lactobacillales"             "Streptococcaceae"             
## [4551,] "Fusobacteriales"             "Leptotrichiaceae"             
## [4552,] "Lactobacillales"             "Carnobacteriaceae"            
## [4553,] "Spirochaetales"              "Spirochaetaceae"              
## [4554,] "Bacteroidales"               "Prevotellaceae"               
## [4555,] "Clostridiales"               "Clostridiales_vadinBB60_group"
##         [,6]                             
##    [1,] "Veillonella"                    
##    [2,] "Streptococcus"                  
##    [3,] "Prevotella_7"                   
##    [4,] "Haemophilus"                    
##    [5,] "Veillonella"                    
##    [6,] "Veillonella"                    
##    [7,] "Veillonella"                    
##    [8,] NA                               
##    [9,] "Neisseria"                      
##   [10,] "Veillonella"                    
##   [11,] "Neisseria"                      
##   [12,] "Veillonella"                    
##   [13,] "Fusobacterium"                  
##   [14,] "Veillonella"                    
##   [15,] "Prevotella_7"                   
##   [16,] "Campylobacter"                  
##   [17,] "Haemophilus"                    
##   [18,] "Oribacterium"                   
##   [19,] "Veillonella"                    
##   [20,] "Alloprevotella"                 
##   [21,] "Granulicatella"                 
##   [22,] "Prevotella_7"                   
##   [23,] "Prevotella_7"                   
##   [24,] "Veillonella"                    
##   [25,] "Haemophilus"                    
##   [26,] "Prevotella_7"                   
##   [27,] "Porphyromonas"                  
##   [28,] "Veillonella"                    
##   [29,] "Streptococcus"                  
##   [30,] "Streptococcus"                  
##   [31,] "Porphyromonas"                  
##   [32,] "Streptococcus"                  
##   [33,] "Prevotella_6"                   
##   [34,] "Prevotella"                     
##   [35,] "Peptostreptococcus"             
##   [36,] "Megasphaera"                    
##   [37,] "Porphyromonas"                  
##   [38,] "Megasphaera"                    
##   [39,] "Lachnoanaerobaculum"            
##   [40,] "Streptococcus"                  
##   [41,] "Prevotella_7"                   
##   [42,] "Streptococcus"                  
##   [43,] "Streptococcus"                  
##   [44,] "Prevotella"                     
##   [45,] "Neisseria"                      
##   [46,] "Prevotella_7"                   
##   [47,] "Megasphaera"                    
##   [48,] "Veillonella"                    
##   [49,] "Lachnoanaerobaculum"            
##   [50,] "Leptotrichia"                   
##   [51,] "Streptococcus"                  
##   [52,] "Haemophilus"                    
##   [53,] "Streptococcus"                  
##   [54,] "Leptotrichia"                   
##   [55,] "Alloprevotella"                 
##   [56,] "Prevotella_7"                   
##   [57,] "Veillonella"                    
##   [58,] "Oribacterium"                   
##   [59,] "Atopobium"                      
##   [60,] "Haemophilus"                    
##   [61,] "Haemophilus"                    
##   [62,] "Prevotella_6"                   
##   [63,] "Oribacterium"                   
##   [64,] "Veillonella"                    
##   [65,] "Haemophilus"                    
##   [66,] "Selenomonas_3"                  
##   [67,] "Candidatus_Saccharimonas"       
##   [68,] "Rothia"                         
##   [69,] "Neisseria"                      
##   [70,] "Parvimonas"                     
##   [71,] "Alloprevotella"                 
##   [72,] "Rothia"                         
##   [73,] "Catonella"                      
##   [74,] "Veillonella"                    
##   [75,] "Rothia"                         
##   [76,] "Neisseria"                      
##   [77,] NA                               
##   [78,] "Porphyromonas"                  
##   [79,] "Veillonella"                    
##   [80,] "Stomatobaculum"                 
##   [81,] "Haemophilus"                    
##   [82,] "Veillonella"                    
##   [83,] "Prevotella_2"                   
##   [84,] "Prevotella"                     
##   [85,] "Megasphaera"                    
##   [86,] "Prevotella_7"                   
##   [87,] "Leptotrichia"                   
##   [88,] "Ruminococcaceae_UCG-014"        
##   [89,] "Alloprevotella"                 
##   [90,] "Ruminococcaceae_UCG-014"        
##   [91,] "Rothia"                         
##   [92,] "Neisseria"                      
##   [93,] "Rothia"                         
##   [94,] "Gemella"                        
##   [95,] "Prevotella_7"                   
##   [96,] "Neisseria"                      
##   [97,] "Fusobacterium"                  
##   [98,] "Veillonella"                    
##   [99,] "Fusobacterium"                  
##  [100,] "Prevotella_7"                   
##  [101,] "Prevotella_7"                   
##  [102,] "Haemophilus"                    
##  [103,] "Haemophilus"                    
##  [104,] "Prevotella"                     
##  [105,] "Granulicatella"                 
##  [106,] "Gemella"                        
##  [107,] "Veillonella"                    
##  [108,] "Veillonella"                    
##  [109,] "Rothia"                         
##  [110,] "Aggregatibacter"                
##  [111,] "Campylobacter"                  
##  [112,] "Prevotella"                     
##  [113,] "Neisseria"                      
##  [114,] "Neisseria"                      
##  [115,] NA                               
##  [116,] "Prevotella"                     
##  [117,] "Neisseria"                      
##  [118,] "Actinobacillus"                 
##  [119,] "Haemophilus"                    
##  [120,] "Haemophilus"                    
##  [121,] "Haemophilus"                    
##  [122,] "Prevotella"                     
##  [123,] "Streptococcus"                  
##  [124,] "Alloprevotella"                 
##  [125,] "Neisseria"                      
##  [126,] "Prevotella"                     
##  [127,] "Veillonella"                    
##  [128,] "Haemophilus"                    
##  [129,] "Aggregatibacter"                
##  [130,] "Haemophilus"                    
##  [131,] "Lautropia"                      
##  [132,] "Prevotella"                     
##  [133,] "Candidatus_Saccharimonas"       
##  [134,] "Prevotella_7"                   
##  [135,] "Veillonella"                    
##  [136,] "Veillonella"                    
##  [137,] "Prevotella_7"                   
##  [138,] "Capnocytophaga"                 
##  [139,] "Prevotella_7"                   
##  [140,] "Streptococcus"                  
##  [141,] "Veillonella"                    
##  [142,] "Megasphaera"                    
##  [143,] "Leptotrichia"                   
##  [144,] "Veillonella"                    
##  [145,] "Streptococcus"                  
##  [146,] "Bergeyella"                     
##  [147,] "Neisseria"                      
##  [148,] "Selenomonas_3"                  
##  [149,] "Veillonella"                    
##  [150,] "Veillonella"                    
##  [151,] "Neisseria"                      
##  [152,] "Prevotella_7"                   
##  [153,] "Veillonella"                    
##  [154,] "Alloprevotella"                 
##  [155,] "Butyrivibrio_2"                 
##  [156,] "Klebsiella"                     
##  [157,] NA                               
##  [158,] "Prevotella_7"                   
##  [159,] "Veillonella"                    
##  [160,] "Oribacterium"                   
##  [161,] "Streptococcus"                  
##  [162,] "Rothia"                         
##  [163,] "Alloprevotella"                 
##  [164,] "Veillonella"                    
##  [165,] "Campylobacter"                  
##  [166,] "Prevotella_7"                   
##  [167,] "Porphyromonas"                  
##  [168,] "Streptococcus"                  
##  [169,] "Alloprevotella"                 
##  [170,] "Neisseria"                      
##  [171,] "Streptococcus"                  
##  [172,] "Streptococcus"                  
##  [173,] "Candidatus_Saccharimonas"       
##  [174,] "Prevotella"                     
##  [175,] "Prevotella_7"                   
##  [176,] "Stomatobaculum"                 
##  [177,] "Veillonella"                    
##  [178,] "Veillonella"                    
##  [179,] "Prevotella_7"                   
##  [180,] "Neisseria"                      
##  [181,] NA                               
##  [182,] "Veillonella"                    
##  [183,] "Neisseria"                      
##  [184,] "Veillonella"                    
##  [185,] "Aggregatibacter"                
##  [186,] "Veillonella"                    
##  [187,] "Veillonella"                    
##  [188,] "Prevotella"                     
##  [189,] "Prevotella_7"                   
##  [190,] "Aggregatibacter"                
##  [191,] "Alloprevotella"                 
##  [192,] "Porphyromonas"                  
##  [193,] "Neisseria"                      
##  [194,] "Prevotella_7"                   
##  [195,] "Veillonella"                    
##  [196,] "Veillonella"                    
##  [197,] "Fusobacterium"                  
##  [198,] "Dialister"                      
##  [199,] "Prevotella"                     
##  [200,] "Selenomonas_3"                  
##  [201,] "Alloprevotella"                 
##  [202,] "Oribacterium"                   
##  [203,] "Campylobacter"                  
##  [204,] "Alloprevotella"                 
##  [205,] "Veillonella"                    
##  [206,] "Veillonella"                    
##  [207,] "Capnocytophaga"                 
##  [208,] "Neisseria"                      
##  [209,] "Kingella"                       
##  [210,] "Streptococcus"                  
##  [211,] NA                               
##  [212,] "Veillonella"                    
##  [213,] "Haemophilus"                    
##  [214,] "Neisseria"                      
##  [215,] "Megasphaera"                    
##  [216,] "Prevotella_7"                   
##  [217,] "Streptococcus"                  
##  [218,] "Neisseria"                      
##  [219,] "Megasphaera"                    
##  [220,] "Streptococcus"                  
##  [221,] "Oribacterium"                   
##  [222,] "Streptococcus"                  
##  [223,] "Johnsonella"                    
##  [224,] "Alloprevotella"                 
##  [225,] "Leptotrichia"                   
##  [226,] "Veillonella"                    
##  [227,] "Alloprevotella"                 
##  [228,] "Streptococcus"                  
##  [229,] "Capnocytophaga"                 
##  [230,] "Capnocytophaga"                 
##  [231,] "Leptotrichia"                   
##  [232,] "Fusobacterium"                  
##  [233,] "Haemophilus"                    
##  [234,] "Fusobacterium"                  
##  [235,] "Veillonella"                    
##  [236,] NA                               
##  [237,] "Alloprevotella"                 
##  [238,] "Veillonella"                    
##  [239,] "Prevotella_7"                   
##  [240,] "Veillonella"                    
##  [241,] "Fusobacterium"                  
##  [242,] "Porphyromonas"                  
##  [243,] NA                               
##  [244,] NA                               
##  [245,] "Prevotella_7"                   
##  [246,] "Aggregatibacter"                
##  [247,] "Neisseria"                      
##  [248,] NA                               
##  [249,] NA                               
##  [250,] "Prevotella_7"                   
##  [251,] "Alloprevotella"                 
##  [252,] "Prevotella_7"                   
##  [253,] "Johnsonella"                    
##  [254,] "Haemophilus"                    
##  [255,] "Enterobacter"                   
##  [256,] "Veillonella"                    
##  [257,] "Veillonella"                    
##  [258,] "Actinobacillus"                 
##  [259,] NA                               
##  [260,] "Actinomyces"                    
##  [261,] "Lachnoanaerobaculum"            
##  [262,] "Prevotella_7"                   
##  [263,] "Alloprevotella"                 
##  [264,] "Neisseria"                      
##  [265,] NA                               
##  [266,] "Bergeriella"                    
##  [267,] "Solobacterium"                  
##  [268,] "Prevotella"                     
##  [269,] "Parvimonas"                     
##  [270,] NA                               
##  [271,] "Veillonella"                    
##  [272,] "Neisseria"                      
##  [273,] "Neisseria"                      
##  [274,] "Veillonella"                    
##  [275,] "Streptococcus"                  
##  [276,] "Megasphaera"                    
##  [277,] "Prevotella"                     
##  [278,] "Capnocytophaga"                 
##  [279,] "Haemophilus"                    
##  [280,] "Alloprevotella"                 
##  [281,] "Fusobacterium"                  
##  [282,] NA                               
##  [283,] "Fusobacterium"                  
##  [284,] "Porphyromonas"                  
##  [285,] "Prevotella"                     
##  [286,] NA                               
##  [287,] "Alloprevotella"                 
##  [288,] "Veillonella"                    
##  [289,] "Alloprevotella"                 
##  [290,] "Oribacterium"                   
##  [291,] "Alloprevotella"                 
##  [292,] "Haemophilus"                    
##  [293,] "Veillonella"                    
##  [294,] "Candidatus_Saccharimonas"       
##  [295,] "Anthococcus"                    
##  [296,] "Veillonella"                    
##  [297,] NA                               
##  [298,] "Selenomonas_3"                  
##  [299,] "Neisseria"                      
##  [300,] "Alloprevotella"                 
##  [301,] "Prevotella_7"                   
##  [302,] "Leptotrichia"                   
##  [303,] "Prevotella"                     
##  [304,] NA                               
##  [305,] "Fusobacterium"                  
##  [306,] "Prevotella_7"                   
##  [307,] "Veillonella"                    
##  [308,] NA                               
##  [309,] NA                               
##  [310,] "Haemophilus"                    
##  [311,] "Veillonella"                    
##  [312,] "Fusobacterium"                  
##  [313,] NA                               
##  [314,] "Campylobacter"                  
##  [315,] "Veillonella"                    
##  [316,] "Prevotella_7"                   
##  [317,] "Veillonella"                    
##  [318,] "Catonella"                      
##  [319,] NA                               
##  [320,] "Prevotella"                     
##  [321,] "Atopobium"                      
##  [322,] "Streptococcus"                  
##  [323,] NA                               
##  [324,] "Veillonella"                    
##  [325,] "Haemophilus"                    
##  [326,] NA                               
##  [327,] "Porphyromonas"                  
##  [328,] "Atopobium"                      
##  [329,] "Veillonella"                    
##  [330,] "Prevotella"                     
##  [331,] "Alloprevotella"                 
##  [332,] NA                               
##  [333,] "Lachnoanaerobaculum"            
##  [334,] "Veillonella"                    
##  [335,] "Catonella"                      
##  [336,] "Leptotrichia"                   
##  [337,] "Abiotrophia"                    
##  [338,] "Leptotrichia"                   
##  [339,] "Rothia"                         
##  [340,] "Fusobacterium"                  
##  [341,] "Fusobacterium"                  
##  [342,] "Prevotella_6"                   
##  [343,] "Neisseria"                      
##  [344,] "Prevotella"                     
##  [345,] "Peptostreptococcus"             
##  [346,] "Kingella"                       
##  [347,] NA                               
##  [348,] "Neisseria"                      
##  [349,] "Haemophilus"                    
##  [350,] "Prevotella"                     
##  [351,] NA                               
##  [352,] "Veillonella"                    
##  [353,] "Alloprevotella"                 
##  [354,] "Actinobacillus"                 
##  [355,] "Fusobacterium"                  
##  [356,] "Veillonella"                    
##  [357,] "Fusobacterium"                  
##  [358,] "Prevotella_7"                   
##  [359,] "Veillonella"                    
##  [360,] "Peptococcus"                    
##  [361,] NA                               
##  [362,] NA                               
##  [363,] "Alloprevotella"                 
##  [364,] "Peptoanaerobacter"              
##  [365,] NA                               
##  [366,] NA                               
##  [367,] "Lachnoanaerobaculum"            
##  [368,] "Lachnoanaerobaculum"            
##  [369,] "Actinomyces"                    
##  [370,] "Prevotella_2"                   
##  [371,] NA                               
##  [372,] "Gemella"                        
##  [373,] "Streptococcus"                  
##  [374,] "Veillonella"                    
##  [375,] NA                               
##  [376,] "Veillonella"                    
##  [377,] "Fusobacterium"                  
##  [378,] NA                               
##  [379,] "Selenomonas"                    
##  [380,] "Leptotrichia"                   
##  [381,] "Fusobacterium"                  
##  [382,] "Parvimonas"                     
##  [383,] "Alloprevotella"                 
##  [384,] "Prevotella"                     
##  [385,] "Streptococcus"                  
##  [386,] "Enterobacter"                   
##  [387,] "Leptotrichia"                   
##  [388,] "Actinomyces"                    
##  [389,] "Leptotrichia"                   
##  [390,] "Prevotella_6"                   
##  [391,] "Actinomyces"                    
##  [392,] "Fusobacterium"                  
##  [393,] NA                               
##  [394,] "Enterobacter"                   
##  [395,] "Veillonella"                    
##  [396,] "Prevotella"                     
##  [397,] "Prevotella"                     
##  [398,] "Haemophilus"                    
##  [399,] "Leptotrichia"                   
##  [400,] "Haemophilus"                    
##  [401,] "Streptococcus"                  
##  [402,] "Capnocytophaga"                 
##  [403,] "Prevotella"                     
##  [404,] "Prevotella_7"                   
##  [405,] "Catonella"                      
##  [406,] "Veillonella"                    
##  [407,] "Alloprevotella"                 
##  [408,] "Gemella"                        
##  [409,] "Veillonella"                    
##  [410,] "Porphyromonas"                  
##  [411,] "Prevotella_7"                   
##  [412,] "Prevotella_7"                   
##  [413,] "Neisseria"                      
##  [414,] "Stomatobaculum"                 
##  [415,] "Prevotella_7"                   
##  [416,] "Moraxella"                      
##  [417,] "Alloprevotella"                 
##  [418,] "Prevotella_7"                   
##  [419,] "Leptotrichia"                   
##  [420,] "Prevotella_7"                   
##  [421,] "Gemella"                        
##  [422,] "Veillonella"                    
##  [423,] "Veillonella"                    
##  [424,] "Veillonella"                    
##  [425,] "Haemophilus"                    
##  [426,] "Alloprevotella"                 
##  [427,] "Prevotella"                     
##  [428,] "Prevotella"                     
##  [429,] "Veillonella"                    
##  [430,] "Veillonella"                    
##  [431,] "Selenomonas_3"                  
##  [432,] "Prevotella_7"                   
##  [433,] "Veillonella"                    
##  [434,] "Prevotella_7"                   
##  [435,] "Oribacterium"                   
##  [436,] "Veillonella"                    
##  [437,] "Prevotella_7"                   
##  [438,] "Streptococcus"                  
##  [439,] "Prevotella_7"                   
##  [440,] NA                               
##  [441,] "Porphyromonas"                  
##  [442,] "Fusobacterium"                  
##  [443,] "Veillonella"                    
##  [444,] "Veillonella"                    
##  [445,] "Veillonella"                    
##  [446,] NA                               
##  [447,] "Streptococcus"                  
##  [448,] "Leptotrichia"                   
##  [449,] "Capnocytophaga"                 
##  [450,] "Leptotrichia"                   
##  [451,] "Prevotella"                     
##  [452,] NA                               
##  [453,] "Mogibacterium"                  
##  [454,] "Capnocytophaga"                 
##  [455,] "Veillonella"                    
##  [456,] "Ruminococcaceae_UCG-014"        
##  [457,] NA                               
##  [458,] "Actinomyces"                    
##  [459,] "Leptotrichia"                   
##  [460,] "Veillonella"                    
##  [461,] NA                               
##  [462,] "Leptotrichia"                   
##  [463,] "Prevotella"                     
##  [464,] "Streptococcus"                  
##  [465,] "Prevotella_7"                   
##  [466,] "Dialister"                      
##  [467,] "Prevotella_7"                   
##  [468,] "Haemophilus"                    
##  [469,] NA                               
##  [470,] "Neisseria"                      
##  [471,] "Haemophilus"                    
##  [472,] "Alloprevotella"                 
##  [473,] "Prevotella_7"                   
##  [474,] "Veillonella"                    
##  [475,] "Prevotella_7"                   
##  [476,] "Haemophilus"                    
##  [477,] "Prevotella"                     
##  [478,] "Aggregatibacter"                
##  [479,] "Klebsiella"                     
##  [480,] "Leptotrichia"                   
##  [481,] "Alloprevotella"                 
##  [482,] "Prevotella_7"                   
##  [483,] "Fusobacterium"                  
##  [484,] "Veillonella"                    
##  [485,] "Streptococcus"                  
##  [486,] NA                               
##  [487,] "Prevotella_7"                   
##  [488,] "Leptotrichia"                   
##  [489,] "Porphyromonas"                  
##  [490,] "Campylobacter"                  
##  [491,] "Veillonella"                    
##  [492,] NA                               
##  [493,] "Prevotella_7"                   
##  [494,] NA                               
##  [495,] "Prevotella_7"                   
##  [496,] "Veillonella"                    
##  [497,] "Klebsiella"                     
##  [498,] "Prevotella_7"                   
##  [499,] "Filifactor"                     
##  [500,] "Veillonella"                    
##  [501,] "Veillonella"                    
##  [502,] "Actinomyces"                    
##  [503,] "Serratia"                       
##  [504,] "Veillonella"                    
##  [505,] "Neisseria"                      
##  [506,] "Leptotrichia"                   
##  [507,] "Actinomyces"                    
##  [508,] "Alloprevotella"                 
##  [509,] "Prevotella_7"                   
##  [510,] "Actinobacillus"                 
##  [511,] "Veillonella"                    
##  [512,] "Veillonella"                    
##  [513,] "Aggregatibacter"                
##  [514,] "Veillonella"                    
##  [515,] "Veillonella"                    
##  [516,] "Megasphaera"                    
##  [517,] "Prevotella_7"                   
##  [518,] "Haemophilus"                    
##  [519,] "Veillonella"                    
##  [520,] "Actinomyces"                    
##  [521,] "Anthococcus"                    
##  [522,] "Actinomyces"                    
##  [523,] "Parvimonas"                     
##  [524,] "Veillonella"                    
##  [525,] "Fusobacterium"                  
##  [526,] "Prevotella_7"                   
##  [527,] "Prevotella_7"                   
##  [528,] "Neisseria"                      
##  [529,] "Campylobacter"                  
##  [530,] "Streptococcus"                  
##  [531,] "Cardiobacterium"                
##  [532,] "Prevotella"                     
##  [533,] NA                               
##  [534,] "Veillonella"                    
##  [535,] "Capnocytophaga"                 
##  [536,] "Prevotella"                     
##  [537,] "Veillonella"                    
##  [538,] "Actinobacillus"                 
##  [539,] "F0058"                          
##  [540,] "Enterobacter"                   
##  [541,] "Leptotrichia"                   
##  [542,] NA                               
##  [543,] "Porphyromonas"                  
##  [544,] "Alloprevotella"                 
##  [545,] "Family_XIII_UCG-001"            
##  [546,] "Prevotella_7"                   
##  [547,] "Neisseria"                      
##  [548,] NA                               
##  [549,] "Veillonella"                    
##  [550,] "Alloprevotella"                 
##  [551,] "Gemella"                        
##  [552,] "Neisseria"                      
##  [553,] "Prevotella"                     
##  [554,] "Capnocytophaga"                 
##  [555,] "Prevotella"                     
##  [556,] "Veillonella"                    
##  [557,] NA                               
##  [558,] "Streptococcus"                  
##  [559,] "Prevotella"                     
##  [560,] "Cricetibacter"                  
##  [561,] "Porphyromonas"                  
##  [562,] NA                               
##  [563,] "Haemophilus"                    
##  [564,] "Lactococcus"                    
##  [565,] "Megasphaera"                    
##  [566,] "Prevotella_7"                   
##  [567,] "Streptococcus"                  
##  [568,] "Prevotella_7"                   
##  [569,] "Alloprevotella"                 
##  [570,] "Leptotrichia"                   
##  [571,] "Prevotella_7"                   
##  [572,] "Tannerella"                     
##  [573,] "Neisseria"                      
##  [574,] "Prevotella_7"                   
##  [575,] "Capnocytophaga"                 
##  [576,] "Prevotella"                     
##  [577,] NA                               
##  [578,] NA                               
##  [579,] "Veillonella"                    
##  [580,] "Veillonella"                    
##  [581,] "Leptotrichia"                   
##  [582,] "Haemophilus"                    
##  [583,] "Fusobacterium"                  
##  [584,] "Porphyromonas"                  
##  [585,] NA                               
##  [586,] "Fusobacterium"                  
##  [587,] "Fusobacterium"                  
##  [588,] "Tannerella"                     
##  [589,] "Fusobacterium"                  
##  [590,] "Prevotella"                     
##  [591,] "Neisseria"                      
##  [592,] "Prevotella_7"                   
##  [593,] "Prevotella_7"                   
##  [594,] "Prevotella_7"                   
##  [595,] "Actinobacillus"                 
##  [596,] "Prevotella_7"                   
##  [597,] "Veillonella"                    
##  [598,] "Neisseria"                      
##  [599,] "Peptostreptococcus"             
##  [600,] "Veillonella"                    
##  [601,] "Haemophilus"                    
##  [602,] "Prevotella_7"                   
##  [603,] "Prevotella_7"                   
##  [604,] "Selenomonas_3"                  
##  [605,] "Neisseria"                      
##  [606,] "Fusobacterium"                  
##  [607,] "Prevotella_7"                   
##  [608,] NA                               
##  [609,] "Alloprevotella"                 
##  [610,] NA                               
##  [611,] "Haemophilus"                    
##  [612,] "Veillonella"                    
##  [613,] "Fusobacterium"                  
##  [614,] "Leptotrichia"                   
##  [615,] "Prevotella"                     
##  [616,] "Veillonella"                    
##  [617,] "Alloprevotella"                 
##  [618,] "Veillonella"                    
##  [619,] "Gemella"                        
##  [620,] "Porphyromonas"                  
##  [621,] "Selenomonas_3"                  
##  [622,] "F0058"                          
##  [623,] "Veillonella"                    
##  [624,] "Prevotella_7"                   
##  [625,] "Prevotella_6"                   
##  [626,] "Prevotella"                     
##  [627,] "Prevotella"                     
##  [628,] "Fusobacterium"                  
##  [629,] "Prevotella"                     
##  [630,] "Rothia"                         
##  [631,] "Megasphaera"                    
##  [632,] NA                               
##  [633,] "Atopobium"                      
##  [634,] "Selenomonas"                    
##  [635,] NA                               
##  [636,] NA                               
##  [637,] "Porphyromonas"                  
##  [638,] "Haemophilus"                    
##  [639,] "Streptococcus"                  
##  [640,] NA                               
##  [641,] "Veillonella"                    
##  [642,] NA                               
##  [643,] "Serratia"                       
##  [644,] "Prevotella_7"                   
##  [645,] "Prevotella"                     
##  [646,] "Prevotella_7"                   
##  [647,] "Rothia"                         
##  [648,] "Streptococcus"                  
##  [649,] "Alloprevotella"                 
##  [650,] "Filifactor"                     
##  [651,] "Veillonella"                    
##  [652,] "Prevotella"                     
##  [653,] "Veillonella"                    
##  [654,] NA                               
##  [655,] "Alloprevotella"                 
##  [656,] "Veillonella"                    
##  [657,] "Leptotrichia"                   
##  [658,] "Porphyromonas"                  
##  [659,] "Veillonella"                    
##  [660,] "Leptotrichia"                   
##  [661,] NA                               
##  [662,] "Fusobacterium"                  
##  [663,] NA                               
##  [664,] "Haemophilus"                    
##  [665,] "Leptotrichia"                   
##  [666,] "Veillonella"                    
##  [667,] NA                               
##  [668,] NA                               
##  [669,] NA                               
##  [670,] "Tannerella"                     
##  [671,] "Actinomyces"                    
##  [672,] "Rothia"                         
##  [673,] NA                               
##  [674,] "Alloprevotella"                 
##  [675,] NA                               
##  [676,] "Haemophilus"                    
##  [677,] "Prevotella_7"                   
##  [678,] "Veillonella"                    
##  [679,] "Prevotella_7"                   
##  [680,] NA                               
##  [681,] "Capnocytophaga"                 
##  [682,] "Haemophilus"                    
##  [683,] "Selenomonas"                    
##  [684,] "Porphyromonas"                  
##  [685,] "Prevotella_7"                   
##  [686,] NA                               
##  [687,] NA                               
##  [688,] "Streptococcus"                  
##  [689,] "Megasphaera"                    
##  [690,] "Veillonella"                    
##  [691,] "Porphyromonas"                  
##  [692,] "Candidatus_Saccharimonas"       
##  [693,] NA                               
##  [694,] "Prevotella_7"                   
##  [695,] "Veillonella"                    
##  [696,] "Streptococcus"                  
##  [697,] "Megasphaera"                    
##  [698,] "Alloprevotella"                 
##  [699,] "Prevotellaceae_YAB2003_group"   
##  [700,] "Treponema_2"                    
##  [701,] "Prevotella_7"                   
##  [702,] "Veillonella"                    
##  [703,] "Leptotrichia"                   
##  [704,] NA                               
##  [705,] "Prevotella"                     
##  [706,] NA                               
##  [707,] "Actinomyces"                    
##  [708,] "Prevotella_7"                   
##  [709,] "Alloprevotella"                 
##  [710,] "Haemophilus"                    
##  [711,] "Prevotella_7"                   
##  [712,] "Veillonella"                    
##  [713,] "Porphyromonas"                  
##  [714,] "Prevotella"                     
##  [715,] "Neisseria"                      
##  [716,] NA                               
##  [717,] NA                               
##  [718,] "Treponema_2"                    
##  [719,] "Prevotella_2"                   
##  [720,] NA                               
##  [721,] "Prevotella"                     
##  [722,] "Prevotella"                     
##  [723,] "Capnocytophaga"                 
##  [724,] "Lautropia"                      
##  [725,] "Actinobacillus"                 
##  [726,] "Neisseria"                      
##  [727,] "Haemophilus"                    
##  [728,] "Veillonella"                    
##  [729,] "Veillonella"                    
##  [730,] "Capnocytophaga"                 
##  [731,] "Eikenella"                      
##  [732,] "Fusobacterium"                  
##  [733,] "Aggregatibacter"                
##  [734,] "Selenomonas_3"                  
##  [735,] NA                               
##  [736,] "Capnocytophaga"                 
##  [737,] "Eikenella"                      
##  [738,] "Actinomyces"                    
##  [739,] "Neisseria"                      
##  [740,] "Bergeyella"                     
##  [741,] "Leptotrichia"                   
##  [742,] "Streptococcus"                  
##  [743,] "Rothia"                         
##  [744,] "Streptococcus"                  
##  [745,] "Veillonella"                    
##  [746,] "Veillonella"                    
##  [747,] "Prevotella"                     
##  [748,] NA                               
##  [749,] "Fusobacterium"                  
##  [750,] NA                               
##  [751,] "Prevotella"                     
##  [752,] "Alloprevotella"                 
##  [753,] "Prevotellaceae_YAB2003_group"   
##  [754,] "Capnocytophaga"                 
##  [755,] NA                               
##  [756,] "Veillonella"                    
##  [757,] "Alloprevotella"                 
##  [758,] "Leptotrichia"                   
##  [759,] "Selenomonas"                    
##  [760,] "Porphyromonas"                  
##  [761,] "Veillonella"                    
##  [762,] "Porphyromonas"                  
##  [763,] "Prevotella"                     
##  [764,] "Selenomonas"                    
##  [765,] "Fusobacterium"                  
##  [766,] "Veillonella"                    
##  [767,] "Fusobacterium"                  
##  [768,] "Fusobacterium"                  
##  [769,] "Alloprevotella"                 
##  [770,] "Alloprevotella"                 
##  [771,] "Streptococcus"                  
##  [772,] "Leptotrichia"                   
##  [773,] "Alloprevotella"                 
##  [774,] "Streptococcus"                  
##  [775,] "Actinomyces"                    
##  [776,] "Veillonella"                    
##  [777,] "Veillonella"                    
##  [778,] "Alloprevotella"                 
##  [779,] "Neisseria"                      
##  [780,] "Oribacterium"                   
##  [781,] "Alloprevotella"                 
##  [782,] "Haemophilus"                    
##  [783,] "Prevotella_7"                   
##  [784,] "Aggregatibacter"                
##  [785,] "Parvimonas"                     
##  [786,] "Abiotrophia"                    
##  [787,] "Prevotella_2"                   
##  [788,] "Alloprevotella"                 
##  [789,] "Aggregatibacter"                
##  [790,] "Lachnoanaerobaculum"            
##  [791,] "Megasphaera"                    
##  [792,] "Oribacterium"                   
##  [793,] "Veillonella"                    
##  [794,] "Leptotrichia"                   
##  [795,] "Campylobacter"                  
##  [796,] "Prevotella"                     
##  [797,] NA                               
##  [798,] "Veillonella"                    
##  [799,] "Veillonella"                    
##  [800,] "Neisseria"                      
##  [801,] "Prevotella_7"                   
##  [802,] "Leptotrichia"                   
##  [803,] "Porphyromonas"                  
##  [804,] "Prevotellaceae_YAB2003_group"   
##  [805,] NA                               
##  [806,] NA                               
##  [807,] "Haemophilus"                    
##  [808,] NA                               
##  [809,] "Prevotella_7"                   
##  [810,] NA                               
##  [811,] "Prevotella_7"                   
##  [812,] "Actinomyces"                    
##  [813,] "Leptotrichia"                   
##  [814,] "Veillonella"                    
##  [815,] "Prevotella"                     
##  [816,] "Haemophilus"                    
##  [817,] "Porphyromonas"                  
##  [818,] "Selenomonas_3"                  
##  [819,] "Prevotella"                     
##  [820,] NA                               
##  [821,] "Bergeriella"                    
##  [822,] "Leptotrichia"                   
##  [823,] "Granulicatella"                 
##  [824,] "Alloprevotella"                 
##  [825,] "Leptotrichia"                   
##  [826,] "Prevotellaceae_YAB2003_group"   
##  [827,] "Alloprevotella"                 
##  [828,] NA                               
##  [829,] "Prevotella_7"                   
##  [830,] "Haemophilus"                    
##  [831,] "Veillonella"                    
##  [832,] "Prevotella_7"                   
##  [833,] NA                               
##  [834,] NA                               
##  [835,] "Megasphaera"                    
##  [836,] "Selenomonas_3"                  
##  [837,] "Granulicatella"                 
##  [838,] "Alloprevotella"                 
##  [839,] "Campylobacter"                  
##  [840,] NA                               
##  [841,] "Veillonella"                    
##  [842,] "Porphyromonas"                  
##  [843,] "Parvimonas"                     
##  [844,] "Porphyromonas"                  
##  [845,] NA                               
##  [846,] "Actinomyces"                    
##  [847,] "Veillonella"                    
##  [848,] "Rothia"                         
##  [849,] "Alloprevotella"                 
##  [850,] "Haemophilus"                    
##  [851,] "Prevotella_7"                   
##  [852,] "Streptococcus"                  
##  [853,] "Prevotella_7"                   
##  [854,] "Megasphaera"                    
##  [855,] "Prevotella_7"                   
##  [856,] "Peptostreptococcus"             
##  [857,] "Alloprevotella"                 
##  [858,] "Streptococcus"                  
##  [859,] "Alloprevotella"                 
##  [860,] NA                               
##  [861,] "Defluviitaleaceae_UCG-011"      
##  [862,] "Lachnoanaerobaculum"            
##  [863,] "Prevotella_7"                   
##  [864,] "Alloprevotella"                 
##  [865,] "Prevotella"                     
##  [866,] "Porphyromonas"                  
##  [867,] "Actinomyces"                    
##  [868,] "Prevotella_7"                   
##  [869,] "Selenomonas"                    
##  [870,] "Actinomyces"                    
##  [871,] "Actinomyces"                    
##  [872,] "Selenomonas_3"                  
##  [873,] "Porphyromonas"                  
##  [874,] "Porphyromonas"                  
##  [875,] "Actinomyces"                    
##  [876,] "Leptotrichia"                   
##  [877,] NA                               
##  [878,] NA                               
##  [879,] "Actinomyces"                    
##  [880,] "Veillonella"                    
##  [881,] "Aggregatibacter"                
##  [882,] "Oribacterium"                   
##  [883,] "Selenomonas_3"                  
##  [884,] "Prevotella"                     
##  [885,] "Alloprevotella"                 
##  [886,] "Porphyromonas"                  
##  [887,] NA                               
##  [888,] "Actinomyces"                    
##  [889,] NA                               
##  [890,] NA                               
##  [891,] "Haemophilus"                    
##  [892,] "Streptococcus"                  
##  [893,] NA                               
##  [894,] "Bergeyella"                     
##  [895,] "Actinomyces"                    
##  [896,] "Prevotella_7"                   
##  [897,] "Prevotella_2"                   
##  [898,] "Prevotella_2"                   
##  [899,] NA                               
##  [900,] NA                               
##  [901,] "Fusobacterium"                  
##  [902,] "Aggregatibacter"                
##  [903,] "Streptococcus"                  
##  [904,] "Ruminococcaceae_UCG-014"        
##  [905,] NA                               
##  [906,] "Prevotella_2"                   
##  [907,] "Streptococcus"                  
##  [908,] "Aggregatibacter"                
##  [909,] "Veillonella"                    
##  [910,] "Haemophilus"                    
##  [911,] "Veillonella"                    
##  [912,] NA                               
##  [913,] "Neisseria"                      
##  [914,] "Aggregatibacter"                
##  [915,] "Alloscardovia"                  
##  [916,] "Veillonella"                    
##  [917,] "Prevotella_6"                   
##  [918,] "Lachnoanaerobaculum"            
##  [919,] "Actinomyces"                    
##  [920,] "Neisseria"                      
##  [921,] "Campylobacter"                  
##  [922,] NA                               
##  [923,] "Veillonella"                    
##  [924,] NA                               
##  [925,] NA                               
##  [926,] "Neisseria"                      
##  [927,] NA                               
##  [928,] "Fusobacterium"                  
##  [929,] "Anaeroglobus"                   
##  [930,] "Alloprevotella"                 
##  [931,] "Leptotrichia"                   
##  [932,] "Streptococcus"                  
##  [933,] NA                               
##  [934,] "Veillonella"                    
##  [935,] "Alloprevotella"                 
##  [936,] NA                               
##  [937,] "Leptotrichia"                   
##  [938,] "Solobacterium"                  
##  [939,] "Streptococcus"                  
##  [940,] "Prevotella_7"                   
##  [941,] "Streptococcus"                  
##  [942,] "Actinomyces"                    
##  [943,] NA                               
##  [944,] "Treponema_2"                    
##  [945,] "Streptococcus"                  
##  [946,] "Actinomyces"                    
##  [947,] "Streptococcus"                  
##  [948,] "Sneathia"                       
##  [949,] "Leptotrichia"                   
##  [950,] "Lachnoanaerobaculum"            
##  [951,] NA                               
##  [952,] "Veillonella"                    
##  [953,] NA                               
##  [954,] "Treponema_2"                    
##  [955,] "Alloprevotella"                 
##  [956,] "Rothia"                         
##  [957,] "Megasphaera"                    
##  [958,] "Candidatus_Saccharimonas"       
##  [959,] "Streptococcus"                  
##  [960,] "Porphyromonas"                  
##  [961,] NA                               
##  [962,] "Streptococcus"                  
##  [963,] "Veillonella"                    
##  [964,] "Serratia"                       
##  [965,] "Fusobacterium"                  
##  [966,] "Lachnoanaerobaculum"            
##  [967,] "Prevotella"                     
##  [968,] "Fusobacterium"                  
##  [969,] "Ruminococcaceae_UCG-014"        
##  [970,] "Alloprevotella"                 
##  [971,] "Fusobacterium"                  
##  [972,] "Dialister"                      
##  [973,] NA                               
##  [974,] "Selenomonas_3"                  
##  [975,] "Prevotella_7"                   
##  [976,] "Lachnoanaerobaculum"            
##  [977,] "Oribacterium"                   
##  [978,] "Selenomonas_3"                  
##  [979,] "Actinomyces"                    
##  [980,] "Actinomyces"                    
##  [981,] "Fusobacterium"                  
##  [982,] "Prevotella"                     
##  [983,] "Leptotrichia"                   
##  [984,] "Actinomyces"                    
##  [985,] NA                               
##  [986,] "Veillonella"                    
##  [987,] "Actinomyces"                    
##  [988,] "Aggregatibacter"                
##  [989,] "Haemophilus"                    
##  [990,] "Veillonella"                    
##  [991,] "Prevotella"                     
##  [992,] NA                               
##  [993,] "Mobiluncus"                     
##  [994,] "Veillonella"                    
##  [995,] "Veillonella"                    
##  [996,] "Leptotrichia"                   
##  [997,] "Leptotrichia"                   
##  [998,] NA                               
##  [999,] "Alloprevotella"                 
## [1000,] "Veillonella"                    
## [1001,] "Veillonella"                    
## [1002,] "Veillonella"                    
## [1003,] "Simonsiella"                    
## [1004,] "Porphyromonas"                  
## [1005,] "Prevotella"                     
## [1006,] "Alloprevotella"                 
## [1007,] "Lachnoanaerobaculum"            
## [1008,] "Prevotella_7"                   
## [1009,] "Prevotella"                     
## [1010,] "Parvimonas"                     
## [1011,] "Veillonella"                    
## [1012,] NA                               
## [1013,] "Alysiella"                      
## [1014,] NA                               
## [1015,] "Oribacterium"                   
## [1016,] NA                               
## [1017,] NA                               
## [1018,] "Prevotella_2"                   
## [1019,] "Fusobacterium"                  
## [1020,] "Serratia"                       
## [1021,] "Prevotella_7"                   
## [1022,] "Leptotrichia"                   
## [1023,] "Treponema_2"                    
## [1024,] "Granulicatella"                 
## [1025,] "Granulicatella"                 
## [1026,] "Corynebacterium"                
## [1027,] "Streptobacillus"                
## [1028,] "Alloprevotella"                 
## [1029,] "Prevotella"                     
## [1030,] NA                               
## [1031,] NA                               
## [1032,] "Leptotrichia"                   
## [1033,] "Fusobacterium"                  
## [1034,] "Veillonella"                    
## [1035,] "Prevotella_7"                   
## [1036,] "Veillonella"                    
## [1037,] "Veillonella"                    
## [1038,] "Catonella"                      
## [1039,] "Alloprevotella"                 
## [1040,] "Veillonella"                    
## [1041,] "Prevotella_7"                   
## [1042,] "Johnsonella"                    
## [1043,] NA                               
## [1044,] "Veillonella"                    
## [1045,] "Actinomyces"                    
## [1046,] "Prevotella"                     
## [1047,] "Veillonella"                    
## [1048,] "Prevotella_7"                   
## [1049,] "Veillonella"                    
## [1050,] NA                               
## [1051,] NA                               
## [1052,] "Granulicatella"                 
## [1053,] "Prevotellaceae_YAB2003_group"   
## [1054,] "Aggregatibacter"                
## [1055,] "Prevotella_7"                   
## [1056,] "Streptococcus"                  
## [1057,] "Haemophilus"                    
## [1058,] "Prevotella"                     
## [1059,] "Catonella"                      
## [1060,] "Alloprevotella"                 
## [1061,] "Veillonella"                    
## [1062,] "Streptococcus"                  
## [1063,] "Prevotella"                     
## [1064,] "Fusobacterium"                  
## [1065,] "Selenomonas"                    
## [1066,] "Leptotrichia"                   
## [1067,] "Veillonella"                    
## [1068,] "Haemophilus"                    
## [1069,] "Leptotrichia"                   
## [1070,] "Veillonella"                    
## [1071,] "Haemophilus"                    
## [1072,] "Porphyromonas"                  
## [1073,] "Alloprevotella"                 
## [1074,] "Atopobium"                      
## [1075,] "Capnocytophaga"                 
## [1076,] NA                               
## [1077,] NA                               
## [1078,] "Alloprevotella"                 
## [1079,] "Veillonella"                    
## [1080,] "Oribacterium"                   
## [1081,] "Prevotella_7"                   
## [1082,] NA                               
## [1083,] "Capnocytophaga"                 
## [1084,] NA                               
## [1085,] "Veillonella"                    
## [1086,] NA                               
## [1087,] "Streptococcus"                  
## [1088,] "Prevotella_2"                   
## [1089,] "Selenomonas"                    
## [1090,] "Prevotella_7"                   
## [1091,] "Atopobium"                      
## [1092,] "Prevotella"                     
## [1093,] "Prevotella"                     
## [1094,] "Prevotella_6"                   
## [1095,] "Capnocytophaga"                 
## [1096,] "Veillonella"                    
## [1097,] "Porphyromonas"                  
## [1098,] NA                               
## [1099,] "Actinomyces"                    
## [1100,] "Prevotella"                     
## [1101,] NA                               
## [1102,] "Veillonella"                    
## [1103,] "Veillonella"                    
## [1104,] "Prevotella"                     
## [1105,] "Prevotella"                     
## [1106,] "Selenomonas_3"                  
## [1107,] "Prevotella_2"                   
## [1108,] "Fusobacterium"                  
## [1109,] "Capnocytophaga"                 
## [1110,] "Veillonella"                    
## [1111,] "Alloprevotella"                 
## [1112,] "Leptotrichia"                   
## [1113,] "Prevotella_2"                   
## [1114,] "Peptoanaerobacter"              
## [1115,] "Candidatus_Saccharimonas"       
## [1116,] NA                               
## [1117,] "Capnocytophaga"                 
## [1118,] "Prevotella_7"                   
## [1119,] "Rothia"                         
## [1120,] NA                               
## [1121,] "Veillonella"                    
## [1122,] NA                               
## [1123,] "Prevotella_7"                   
## [1124,] "Campylobacter"                  
## [1125,] "Actinomyces"                    
## [1126,] NA                               
## [1127,] "Porphyromonas"                  
## [1128,] NA                               
## [1129,] "Fusobacterium"                  
## [1130,] "Aggregatibacter"                
## [1131,] "Catonella"                      
## [1132,] "Prevotella_7"                   
## [1133,] "Prevotella_6"                   
## [1134,] "Anaeroglobus"                   
## [1135,] "Haemophilus"                    
## [1136,] "Haemophilus"                    
## [1137,] "Prevotella"                     
## [1138,] "Prevotella_7"                   
## [1139,] "Kingella"                       
## [1140,] NA                               
## [1141,] "Moraxella"                      
## [1142,] "Prevotella"                     
## [1143,] "Johnsonella"                    
## [1144,] "Streptococcus"                  
## [1145,] "Actinomyces"                    
## [1146,] NA                               
## [1147,] "Prevotella_2"                   
## [1148,] "Veillonella"                    
## [1149,] "Lachnoanaerobaculum"            
## [1150,] NA                               
## [1151,] "Leptotrichia"                   
## [1152,] "Aggregatibacter"                
## [1153,] "Streptococcus"                  
## [1154,] "Neisseria"                      
## [1155,] "Prevotella_7"                   
## [1156,] "Haemophilus"                    
## [1157,] "Streptococcus"                  
## [1158,] "Fusobacterium"                  
## [1159,] NA                               
## [1160,] "Prevotella_7"                   
## [1161,] "Prevotella"                     
## [1162,] "Streptococcus"                  
## [1163,] "Mycoplasma"                     
## [1164,] NA                               
## [1165,] NA                               
## [1166,] "Veillonella"                    
## [1167,] "Kingella"                       
## [1168,] "Megasphaera"                    
## [1169,] "Veillonella"                    
## [1170,] NA                               
## [1171,] "Prevotella"                     
## [1172,] NA                               
## [1173,] "Prevotella"                     
## [1174,] "Selenomonas"                    
## [1175,] "Veillonella"                    
## [1176,] "Veillonella"                    
## [1177,] "Porphyromonas"                  
## [1178,] "Leptotrichia"                   
## [1179,] "Eikenella"                      
## [1180,] "Alloprevotella"                 
## [1181,] "Stomatobaculum"                 
## [1182,] "Peptostreptococcus"             
## [1183,] "Streptococcus"                  
## [1184,] "Prevotella_7"                   
## [1185,] NA                               
## [1186,] "Cricetibacter"                  
## [1187,] "Prevotella_2"                   
## [1188,] "Veillonella"                    
## [1189,] "Prevotella_7"                   
## [1190,] "Prevotella_6"                   
## [1191,] "Actinomyces"                    
## [1192,] "Leptotrichia"                   
## [1193,] "Actinomyces"                    
## [1194,] "Haemophilus"                    
## [1195,] "Porphyromonas"                  
## [1196,] "Capnocytophaga"                 
## [1197,] "Veillonella"                    
## [1198,] "Lachnoanaerobaculum"            
## [1199,] "Streptococcus"                  
## [1200,] "Prevotella_7"                   
## [1201,] NA                               
## [1202,] "Fusobacterium"                  
## [1203,] "Actinomyces"                    
## [1204,] "Prevotella_7"                   
## [1205,] "Fretibacterium"                 
## [1206,] NA                               
## [1207,] "Lautropia"                      
## [1208,] "Selenomonas_3"                  
## [1209,] "Candidatus_Liberibacter"        
## [1210,] "Dialister"                      
## [1211,] "Fusobacterium"                  
## [1212,] NA                               
## [1213,] NA                               
## [1214,] "Fusobacterium"                  
## [1215,] NA                               
## [1216,] "Lachnospiraceae_AC2044_group"   
## [1217,] "Treponema_2"                    
## [1218,] "Alloprevotella"                 
## [1219,] "Peptostreptococcus"             
## [1220,] "Peptostreptococcus"             
## [1221,] "Selenomonas_3"                  
## [1222,] "Actinomyces"                    
## [1223,] "Alloprevotella"                 
## [1224,] NA                               
## [1225,] "Veillonella"                    
## [1226,] NA                               
## [1227,] "Selenomonas_3"                  
## [1228,] "Lautropia"                      
## [1229,] "Prevotella"                     
## [1230,] "Megasphaera"                    
## [1231,] "Porphyromonas"                  
## [1232,] "Prevotella_2"                   
## [1233,] "Bergeyella"                     
## [1234,] "Veillonella"                    
## [1235,] "Porphyromonas"                  
## [1236,] "Stomatobaculum"                 
## [1237,] "Fusobacterium"                  
## [1238,] "Rothia"                         
## [1239,] "Veillonella"                    
## [1240,] NA                               
## [1241,] "Porphyromonas"                  
## [1242,] "Porphyromonas"                  
## [1243,] NA                               
## [1244,] NA                               
## [1245,] "Veillonella"                    
## [1246,] "Leptotrichia"                   
## [1247,] NA                               
## [1248,] "Prevotella_7"                   
## [1249,] "Selenomonas_3"                  
## [1250,] NA                               
## [1251,] NA                               
## [1252,] "Prevotella"                     
## [1253,] "Prevotella_7"                   
## [1254,] NA                               
## [1255,] "Alloprevotella"                 
## [1256,] "F0058"                          
## [1257,] "Selenomonas_3"                  
## [1258,] "Alloprevotella"                 
## [1259,] "Anthococcus"                    
## [1260,] "Prevotella_7"                   
## [1261,] "Simonsiella"                    
## [1262,] "Veillonella"                    
## [1263,] NA                               
## [1264,] NA                               
## [1265,] "Actinomyces"                    
## [1266,] "Veillonella"                    
## [1267,] "Atopobium"                      
## [1268,] "Prevotella_7"                   
## [1269,] NA                               
## [1270,] "Prevotella_2"                   
## [1271,] NA                               
## [1272,] "Veillonella"                    
## [1273,] NA                               
## [1274,] "Prevotella_2"                   
## [1275,] "Treponema_2"                    
## [1276,] "Alloprevotella"                 
## [1277,] "Selenomonas_3"                  
## [1278,] "Veillonella"                    
## [1279,] "Veillonella"                    
## [1280,] "Aggregatibacter"                
## [1281,] "Actinobacillus"                 
## [1282,] "Prevotella"                     
## [1283,] "Candidatus_Saccharimonas"       
## [1284,] "Selenomonas_3"                  
## [1285,] "Fusobacterium"                  
## [1286,] "Corynebacterium"                
## [1287,] "Megasphaera"                    
## [1288,] "Lachnoanaerobaculum"            
## [1289,] "Mycoplasma"                     
## [1290,] "Fusobacterium"                  
## [1291,] "Prevotella_7"                   
## [1292,] "Ruminococcaceae_UCG-014"        
## [1293,] "Lachnoanaerobaculum"            
## [1294,] "Fusobacterium"                  
## [1295,] "Lautropia"                      
## [1296,] "Prevotellaceae_YAB2003_group"   
## [1297,] "Lachnoanaerobaculum"            
## [1298,] "Bergeyella"                     
## [1299,] "Prevotella_7"                   
## [1300,] "Porphyromonas"                  
## [1301,] "Stomatobaculum"                 
## [1302,] "Prevotella"                     
## [1303,] "Veillonella"                    
## [1304,] "Prevotella"                     
## [1305,] "Actinomyces"                    
## [1306,] "Streptococcus"                  
## [1307,] "Kingella"                       
## [1308,] "Simonsiella"                    
## [1309,] "Treponema_2"                    
## [1310,] "Actinobacillus"                 
## [1311,] NA                               
## [1312,] NA                               
## [1313,] "Prevotella_7"                   
## [1314,] "Catonella"                      
## [1315,] NA                               
## [1316,] NA                               
## [1317,] NA                               
## [1318,] "Veillonella"                    
## [1319,] "Selenomonas"                    
## [1320,] "Streptococcus"                  
## [1321,] NA                               
## [1322,] "Campylobacter"                  
## [1323,] "Veillonella"                    
## [1324,] "Streptococcus"                  
## [1325,] "Prevotella_7"                   
## [1326,] "Haemophilus"                    
## [1327,] NA                               
## [1328,] "Selenomonas_3"                  
## [1329,] "Prevotella_7"                   
## [1330,] "Porphyromonas"                  
## [1331,] "Catonella"                      
## [1332,] "Prevotella_2"                   
## [1333,] "Veillonella"                    
## [1334,] "Prevotella_7"                   
## [1335,] NA                               
## [1336,] NA                               
## [1337,] "Veillonella"                    
## [1338,] "Johnsonella"                    
## [1339,] NA                               
## [1340,] "Veillonella"                    
## [1341,] NA                               
## [1342,] "Leptotrichia"                   
## [1343,] "Alloprevotella"                 
## [1344,] "Candidatus_Saccharimonas"       
## [1345,] "Capnocytophaga"                 
## [1346,] NA                               
## [1347,] "Alloprevotella"                 
## [1348,] "Prevotella"                     
## [1349,] "Alysiella"                      
## [1350,] NA                               
## [1351,] "Actinomyces"                    
## [1352,] "Actinomyces"                    
## [1353,] "Porphyromonas"                  
## [1354,] "Selenomonas"                    
## [1355,] NA                               
## [1356,] NA                               
## [1357,] "Prevotella"                     
## [1358,] "Acholeplasma"                   
## [1359,] "Veillonella"                    
## [1360,] "Alloprevotella"                 
## [1361,] "Kingella"                       
## [1362,] NA                               
## [1363,] NA                               
## [1364,] NA                               
## [1365,] "Prevotella_7"                   
## [1366,] "Prevotella_7"                   
## [1367,] "Porphyromonas"                  
## [1368,] "Haemophilus"                    
## [1369,] "Fusobacterium"                  
## [1370,] "Ruminococcaceae_UCG-014"        
## [1371,] "Alloprevotella"                 
## [1372,] "Leptotrichia"                   
## [1373,] "Gemella"                        
## [1374,] "Neisseria"                      
## [1375,] NA                               
## [1376,] "Neisseria"                      
## [1377,] "Veillonella"                    
## [1378,] "Prevotella_7"                   
## [1379,] "Prevotella_7"                   
## [1380,] NA                               
## [1381,] "Butyrivibrio_2"                 
## [1382,] "Prevotella_7"                   
## [1383,] "Veillonella"                    
## [1384,] "Streptococcus"                  
## [1385,] NA                               
## [1386,] "Streptococcus"                  
## [1387,] "Fusobacterium"                  
## [1388,] "Prevotella_7"                   
## [1389,] NA                               
## [1390,] "Alloprevotella"                 
## [1391,] NA                               
## [1392,] "Alloprevotella"                 
## [1393,] "Fusobacterium"                  
## [1394,] "Prevotella_7"                   
## [1395,] "Leptotrichia"                   
## [1396,] NA                               
## [1397,] "Prevotella_7"                   
## [1398,] "Prevotella_7"                   
## [1399,] NA                               
## [1400,] "Bergeyella"                     
## [1401,] "Streptobacillus"                
## [1402,] "Alloprevotella"                 
## [1403,] "Fusobacterium"                  
## [1404,] "Alloprevotella"                 
## [1405,] "Campylobacter"                  
## [1406,] "Haemophilus"                    
## [1407,] "Leptotrichia"                   
## [1408,] "Veillonella"                    
## [1409,] "Haemophilus"                    
## [1410,] "Fusobacterium"                  
## [1411,] "Ruminococcaceae_UCG-014"        
## [1412,] "Staphylococcus"                 
## [1413,] NA                               
## [1414,] "Eikenella"                      
## [1415,] "Prevotella_2"                   
## [1416,] NA                               
## [1417,] "Treponema_2"                    
## [1418,] "Selenomonas"                    
## [1419,] NA                               
## [1420,] "Leptotrichia"                   
## [1421,] NA                               
## [1422,] "Megasphaera"                    
## [1423,] "Fusobacterium"                  
## [1424,] "Prevotella_7"                   
## [1425,] "Veillonella"                    
## [1426,] "Campylobacter"                  
## [1427,] "Prevotellaceae_YAB2003_group"   
## [1428,] "Actinomyces"                    
## [1429,] "Prevotella"                     
## [1430,] "Prevotella"                     
## [1431,] "Selenomonas"                    
## [1432,] "Treponema_2"                    
## [1433,] "Prevotella_2"                   
## [1434,] "Anthococcus"                    
## [1435,] "Actinobacillus"                 
## [1436,] NA                               
## [1437,] "Parvimonas"                     
## [1438,] "Veillonella"                    
## [1439,] "Haemophilus"                    
## [1440,] "Corynebacterium"                
## [1441,] NA                               
## [1442,] "Veillonella"                    
## [1443,] "Alloprevotella"                 
## [1444,] "Bergeyella"                     
## [1445,] "Shuttleworthia"                 
## [1446,] "Actinobacillus"                 
## [1447,] NA                               
## [1448,] "Chelonobacter"                  
## [1449,] "Prevotella"                     
## [1450,] "Streptococcus"                  
## [1451,] NA                               
## [1452,] "Actinomyces"                    
## [1453,] "Prevotella"                     
## [1454,] "Veillonella"                    
## [1455,] "Megasphaera"                    
## [1456,] "Serratia"                       
## [1457,] "Prevotella"                     
## [1458,] NA                               
## [1459,] "Veillonella"                    
## [1460,] "Fusobacterium"                  
## [1461,] "Porphyromonas"                  
## [1462,] "Alloprevotella"                 
## [1463,] "Treponema_2"                    
## [1464,] "Johnsonella"                    
## [1465,] "Leptotrichia"                   
## [1466,] NA                               
## [1467,] "Rikenellaceae_RC9_gut_group"    
## [1468,] NA                               
## [1469,] NA                               
## [1470,] "Prevotella"                     
## [1471,] NA                               
## [1472,] NA                               
## [1473,] "Prevotella"                     
## [1474,] NA                               
## [1475,] "Prevotella"                     
## [1476,] "Selenomonas_3"                  
## [1477,] "Leptotrichia"                   
## [1478,] "Lachnoanaerobaculum"            
## [1479,] "Aggregatibacter"                
## [1480,] "Selenomonas"                    
## [1481,] "Prevotella"                     
## [1482,] "Selenomonas_3"                  
## [1483,] "Prevotella_6"                   
## [1484,] "Alloprevotella"                 
## [1485,] "Prevotella"                     
## [1486,] "Selenomonas_3"                  
## [1487,] "Prevotella"                     
## [1488,] "Veillonella"                    
## [1489,] "Neisseria"                      
## [1490,] "Actinomyces"                    
## [1491,] "Leptotrichia"                   
## [1492,] "Stomatobaculum"                 
## [1493,] "Lachnoanaerobaculum"            
## [1494,] "Veillonella"                    
## [1495,] "Moraxella"                      
## [1496,] "Haemophilus"                    
## [1497,] "Prevotella"                     
## [1498,] "Actinobacillus"                 
## [1499,] NA                               
## [1500,] "Streptococcus"                  
## [1501,] "Corynebacterium"                
## [1502,] "Campylobacter"                  
## [1503,] "Bergeyella"                     
## [1504,] "Selenomonas_3"                  
## [1505,] "Neisseria"                      
## [1506,] NA                               
## [1507,] "Prevotella_7"                   
## [1508,] "Alloprevotella"                 
## [1509,] "Aggregatibacter"                
## [1510,] "Fusobacterium"                  
## [1511,] "Neisseria"                      
## [1512,] "Prevotella"                     
## [1513,] "Capnocytophaga"                 
## [1514,] "Granulicatella"                 
## [1515,] NA                               
## [1516,] "Alloprevotella"                 
## [1517,] NA                               
## [1518,] "Catonella"                      
## [1519,] "Granulicatella"                 
## [1520,] "Prevotella"                     
## [1521,] "Leptotrichia"                   
## [1522,] "Prevotella_6"                   
## [1523,] "Fusobacterium"                  
## [1524,] "Simonsiella"                    
## [1525,] "Atopobium"                      
## [1526,] "Veillonella"                    
## [1527,] "Prevotella_7"                   
## [1528,] NA                               
## [1529,] "Prevotella"                     
## [1530,] "Stomatobaculum"                 
## [1531,] "Campylobacter"                  
## [1532,] NA                               
## [1533,] NA                               
## [1534,] "Catonella"                      
## [1535,] NA                               
## [1536,] "Prevotella_7"                   
## [1537,] "Prevotella_2"                   
## [1538,] NA                               
## [1539,] "Actinomyces"                    
## [1540,] NA                               
## [1541,] "Selenomonas_3"                  
## [1542,] "Prevotella_6"                   
## [1543,] "Streptococcus"                  
## [1544,] "Treponema_2"                    
## [1545,] "Ruminococcaceae_UCG-014"        
## [1546,] "Selenomonas_3"                  
## [1547,] "Alloprevotella"                 
## [1548,] NA                               
## [1549,] "Fusobacterium"                  
## [1550,] "Selenomonas_3"                  
## [1551,] NA                               
## [1552,] NA                               
## [1553,] "Leptotrichia"                   
## [1554,] "Veillonella"                    
## [1555,] "Alloprevotella"                 
## [1556,] "Actinomyces"                    
## [1557,] "Prevotella_2"                   
## [1558,] "Megasphaera"                    
## [1559,] "Fusobacterium"                  
## [1560,] NA                               
## [1561,] "Prevotella_7"                   
## [1562,] "Prevotella_2"                   
## [1563,] "Selenomonas_3"                  
## [1564,] "Leptotrichia"                   
## [1565,] "Prevotella_7"                   
## [1566,] "Peptococcus"                    
## [1567,] "Megasphaera"                    
## [1568,] NA                               
## [1569,] NA                               
## [1570,] "Rikenellaceae_RC9_gut_group"    
## [1571,] "Prevotella_2"                   
## [1572,] "Prevotella_2"                   
## [1573,] "Veillonella"                    
## [1574,] "Fusobacterium"                  
## [1575,] "Prevotella"                     
## [1576,] "Porphyromonas"                  
## [1577,] "Fusobacterium"                  
## [1578,] "Prevotella_2"                   
## [1579,] "Alloprevotella"                 
## [1580,] "Prevotella"                     
## [1581,] "Prevotellaceae_YAB2003_group"   
## [1582,] "Veillonella"                    
## [1583,] "Streptococcus"                  
## [1584,] "Treponema_2"                    
## [1585,] NA                               
## [1586,] "Alloprevotella"                 
## [1587,] "Veillonella"                    
## [1588,] "Veillonella"                    
## [1589,] NA                               
## [1590,] NA                               
## [1591,] "Prevotella"                     
## [1592,] "Granulicatella"                 
## [1593,] "Haemophilus"                    
## [1594,] "Capnocytophaga"                 
## [1595,] "Actinobacillus"                 
## [1596,] "Veillonella"                    
## [1597,] "Prevotella_2"                   
## [1598,] "Prevotella_2"                   
## [1599,] "Prevotella_7"                   
## [1600,] "Prevotellaceae_YAB2003_group"   
## [1601,] "Veillonella"                    
## [1602,] "Porphyromonas"                  
## [1603,] "Prevotella_6"                   
## [1604,] NA                               
## [1605,] NA                               
## [1606,] "Streptococcus"                  
## [1607,] "Prevotella"                     
## [1608,] "Megasphaera"                    
## [1609,] "Neisseria"                      
## [1610,] "Prevotella_2"                   
## [1611,] "Fusobacterium"                  
## [1612,] "Leptotrichia"                   
## [1613,] "Haemophilus"                    
## [1614,] "Prevotella"                     
## [1615,] NA                               
## [1616,] NA                               
## [1617,] "Actinomyces"                    
## [1618,] "Veillonella"                    
## [1619,] "Selenomonas"                    
## [1620,] "Leptotrichia"                   
## [1621,] "Actinomyces"                    
## [1622,] "Haemophilus"                    
## [1623,] "Fusobacterium"                  
## [1624,] "Porphyromonas"                  
## [1625,] "Leptotrichia"                   
## [1626,] "Streptococcus"                  
## [1627,] "Prevotella_7"                   
## [1628,] "Lachnoanaerobaculum"            
## [1629,] "Oribacterium"                   
## [1630,] "Prevotella_7"                   
## [1631,] "Prevotella"                     
## [1632,] "Selenomonas"                    
## [1633,] "Leptotrichia"                   
## [1634,] NA                               
## [1635,] "Alloprevotella"                 
## [1636,] "Haemophilus"                    
## [1637,] "Selenomonas_3"                  
## [1638,] "Prevotella"                     
## [1639,] "Alloprevotella"                 
## [1640,] "Porphyromonas"                  
## [1641,] "Actinomyces"                    
## [1642,] "Prevotella"                     
## [1643,] NA                               
## [1644,] NA                               
## [1645,] "Prevotella_2"                   
## [1646,] NA                               
## [1647,] "Lachnoanaerobaculum"            
## [1648,] "Ruminococcaceae_UCG-014"        
## [1649,] "Leptotrichia"                   
## [1650,] "Leptotrichia"                   
## [1651,] "Stomatobaculum"                 
## [1652,] "Alloprevotella"                 
## [1653,] "Selenomonas"                    
## [1654,] "Campylobacter"                  
## [1655,] "Candidatus_Saccharimonas"       
## [1656,] "Cryptobacterium"                
## [1657,] "Prevotella_7"                   
## [1658,] "Treponema_2"                    
## [1659,] "Haemophilus"                    
## [1660,] "Rahnella"                       
## [1661,] "Alloprevotella"                 
## [1662,] "Alloprevotella"                 
## [1663,] "Veillonella"                    
## [1664,] "Porphyromonas"                  
## [1665,] "Alloprevotella"                 
## [1666,] "Leptotrichia"                   
## [1667,] "Leptotrichia"                   
## [1668,] "Lachnoanaerobaculum"            
## [1669,] "Treponema_2"                    
## [1670,] "Leptotrichia"                   
## [1671,] NA                               
## [1672,] NA                               
## [1673,] "Moraxella"                      
## [1674,] "Haemophilus"                    
## [1675,] "Prevotella_7"                   
## [1676,] "Lachnoanaerobaculum"            
## [1677,] "Anthococcus"                    
## [1678,] NA                               
## [1679,] "Cricetibacter"                  
## [1680,] NA                               
## [1681,] "Prevotella"                     
## [1682,] NA                               
## [1683,] NA                               
## [1684,] "Haemophilus"                    
## [1685,] "Selenomonas_3"                  
## [1686,] "Streptococcus"                  
## [1687,] "Prevotella"                     
## [1688,] "Veillonella"                    
## [1689,] "Fusobacterium"                  
## [1690,] "Actinobacillus"                 
## [1691,] "Fusobacterium"                  
## [1692,] "Prevotellaceae_YAB2003_group"   
## [1693,] "Streptobacillus"                
## [1694,] "Alloprevotella"                 
## [1695,] "Prevotella_2"                   
## [1696,] "Actinomyces"                    
## [1697,] "Prevotellaceae_YAB2003_group"   
## [1698,] "Campylobacter"                  
## [1699,] "Haemophilus"                    
## [1700,] "Actinomyces"                    
## [1701,] NA                               
## [1702,] NA                               
## [1703,] "Streptococcus"                  
## [1704,] "Haemophilus"                    
## [1705,] "Prevotella"                     
## [1706,] "Kingella"                       
## [1707,] "Streptococcus"                  
## [1708,] "Selenomonas"                    
## [1709,] "Streptococcus"                  
## [1710,] "Prevotella_7"                   
## [1711,] "Selenomonas_3"                  
## [1712,] NA                               
## [1713,] "Lachnoanaerobaculum"            
## [1714,] "Veillonella"                    
## [1715,] "Alloprevotella"                 
## [1716,] "Leptotrichia"                   
## [1717,] "Haemophilus"                    
## [1718,] "Capnocytophaga"                 
## [1719,] "Oribacterium"                   
## [1720,] "Neisseria"                      
## [1721,] "Prevotella"                     
## [1722,] "Bergeyella"                     
## [1723,] "Veillonella"                    
## [1724,] "Prevotella_7"                   
## [1725,] "Aggregatibacter"                
## [1726,] "Prevotella"                     
## [1727,] "Atopobium"                      
## [1728,] "Prevotella_7"                   
## [1729,] "Prevotella_6"                   
## [1730,] "Fusobacterium"                  
## [1731,] "Veillonella"                    
## [1732,] "Corynebacterium"                
## [1733,] "Prevotella"                     
## [1734,] NA                               
## [1735,] "Alloprevotella"                 
## [1736,] "Prevotella"                     
## [1737,] "Defluviitaleaceae_UCG-011"      
## [1738,] "Prevotella"                     
## [1739,] "Prevotella_7"                   
## [1740,] "Prevotella"                     
## [1741,] NA                               
## [1742,] "Alloprevotella"                 
## [1743,] "Streptococcus"                  
## [1744,] "Prevotella"                     
## [1745,] "Neisseria"                      
## [1746,] "Porphyromonas"                  
## [1747,] "Porphyromonas"                  
## [1748,] "Fusobacterium"                  
## [1749,] "Prevotella_7"                   
## [1750,] "Alloprevotella"                 
## [1751,] "Gemella"                        
## [1752,] "Veillonella"                    
## [1753,] "Prevotella_7"                   
## [1754,] "Prevotella_2"                   
## [1755,] NA                               
## [1756,] "Granulicatella"                 
## [1757,] NA                               
## [1758,] "Alloprevotella"                 
## [1759,] "Prevotella"                     
## [1760,] "Candidatus_Saccharimonas"       
## [1761,] "Porphyromonas"                  
## [1762,] "Granulicatella"                 
## [1763,] "Streptococcus"                  
## [1764,] "Selenomonas"                    
## [1765,] NA                               
## [1766,] "Prevotella_6"                   
## [1767,] "Ruminococcaceae_UCG-014"        
## [1768,] "Prevotella_7"                   
## [1769,] "Selenomonas_3"                  
## [1770,] "Porphyromonas"                  
## [1771,] "Selenomonas"                    
## [1772,] "Cardiobacterium"                
## [1773,] "Stomatobaculum"                 
## [1774,] "Actinomyces"                    
## [1775,] "Ruminococcaceae_UCG-014"        
## [1776,] "Gemella"                        
## [1777,] "Haemophilus"                    
## [1778,] NA                               
## [1779,] "Fusobacterium"                  
## [1780,] "Prevotella_7"                   
## [1781,] NA                               
## [1782,] "Necropsobacter"                 
## [1783,] NA                               
## [1784,] "Alloprevotella"                 
## [1785,] "Prevotella_7"                   
## [1786,] "Alloprevotella"                 
## [1787,] "Centipeda"                      
## [1788,] "Prevotella"                     
## [1789,] NA                               
## [1790,] "Leptotrichia"                   
## [1791,] "Haemophilus"                    
## [1792,] NA                               
## [1793,] NA                               
## [1794,] "Peptostreptococcus"             
## [1795,] NA                               
## [1796,] NA                               
## [1797,] "Alloprevotella"                 
## [1798,] "Catonella"                      
## [1799,] "Streptococcus"                  
## [1800,] NA                               
## [1801,] "Oribacterium"                   
## [1802,] "Veillonella"                    
## [1803,] "Fusobacterium"                  
## [1804,] "Johnsonella"                    
## [1805,] "Fusobacterium"                  
## [1806,] "Capnocytophaga"                 
## [1807,] NA                               
## [1808,] "Granulicatella"                 
## [1809,] "Rothia"                         
## [1810,] "Actinomyces"                    
## [1811,] "Veillonella"                    
## [1812,] "Veillonella"                    
## [1813,] "Veillonella"                    
## [1814,] "Fusobacterium"                  
## [1815,] NA                               
## [1816,] NA                               
## [1817,] "Atlantibacter"                  
## [1818,] "Prevotella_7"                   
## [1819,] "Porphyromonas"                  
## [1820,] "Alloprevotella"                 
## [1821,] "Alysiella"                      
## [1822,] "Prevotella"                     
## [1823,] "Selenomonas_4"                  
## [1824,] "Veillonella"                    
## [1825,] "Anthococcus"                    
## [1826,] "Anaeroglobus"                   
## [1827,] "Porphyromonas"                  
## [1828,] "Aggregatibacter"                
## [1829,] "Oribacterium"                   
## [1830,] "Leptotrichia"                   
## [1831,] NA                               
## [1832,] "Prevotella_2"                   
## [1833,] "Corynebacterium"                
## [1834,] NA                               
## [1835,] "Bergeyella"                     
## [1836,] "Treponema_2"                    
## [1837,] NA                               
## [1838,] "Prevotellaceae_YAB2003_group"   
## [1839,] "Leptotrichia"                   
## [1840,] "Treponema_2"                    
## [1841,] "Veillonella"                    
## [1842,] "Ezakiella"                      
## [1843,] "Prevotella_7"                   
## [1844,] "Porphyromonas"                  
## [1845,] "Peptostreptococcus"             
## [1846,] "Lautropia"                      
## [1847,] "Alloprevotella"                 
## [1848,] "Prevotella_7"                   
## [1849,] "Prevotella_2"                   
## [1850,] "Prevotella_2"                   
## [1851,] "Fretibacterium"                 
## [1852,] "Prevotella_7"                   
## [1853,] "Alloprevotella"                 
## [1854,] "Haemophilus"                    
## [1855,] "Prevotella_7"                   
## [1856,] "Haemophilus"                    
## [1857,] "Prevotella_7"                   
## [1858,] "Porphyromonas"                  
## [1859,] NA                               
## [1860,] "Lachnoanaerobaculum"            
## [1861,] "Actinobacillus"                 
## [1862,] "Capnocytophaga"                 
## [1863,] "Alysiella"                      
## [1864,] "Fusobacterium"                  
## [1865,] "Aggregatibacter"                
## [1866,] "Prevotella_2"                   
## [1867,] NA                               
## [1868,] "Oribacterium"                   
## [1869,] "Haemophilus"                    
## [1870,] NA                               
## [1871,] "Haemophilus"                    
## [1872,] "Streptococcus"                  
## [1873,] "Haemophilus"                    
## [1874,] "Selenomonas_4"                  
## [1875,] "Fusobacterium"                  
## [1876,] NA                               
## [1877,] NA                               
## [1878,] "Capnocytophaga"                 
## [1879,] NA                               
## [1880,] "Ruminococcaceae_UCG-014"        
## [1881,] "Alloprevotella"                 
## [1882,] "Selenomonas_3"                  
## [1883,] "Fusobacterium"                  
## [1884,] "Prevotella_7"                   
## [1885,] "Actinomyces"                    
## [1886,] NA                               
## [1887,] NA                               
## [1888,] "Rothia"                         
## [1889,] "Alloprevotella"                 
## [1890,] "Prevotella"                     
## [1891,] "Neisseria"                      
## [1892,] "Kingella"                       
## [1893,] "Veillonella"                    
## [1894,] NA                               
## [1895,] "Veillonella"                    
## [1896,] "Oribacterium"                   
## [1897,] "Prevotella_7"                   
## [1898,] "Streptococcus"                  
## [1899,] "Aggregatibacter"                
## [1900,] "Veillonella"                    
## [1901,] "Johnsonella"                    
## [1902,] "Veillonella"                    
## [1903,] "Alloprevotella"                 
## [1904,] "Alloprevotella"                 
## [1905,] "Fusobacterium"                  
## [1906,] NA                               
## [1907,] NA                               
## [1908,] "Alloprevotella"                 
## [1909,] "Catonella"                      
## [1910,] "Fusobacterium"                  
## [1911,] "Aggregatibacter"                
## [1912,] "Alloprevotella"                 
## [1913,] "Veillonella"                    
## [1914,] "Capnocytophaga"                 
## [1915,] "Prevotella_7"                   
## [1916,] "Treponema_2"                    
## [1917,] "Capnocytophaga"                 
## [1918,] "Capnocytophaga"                 
## [1919,] NA                               
## [1920,] "Lachnoanaerobaculum"            
## [1921,] "Prevotella_7"                   
## [1922,] "Stomatobaculum"                 
## [1923,] "Pseudomonas"                    
## [1924,] "Rahnella"                       
## [1925,] "Prevotella_6"                   
## [1926,] "Prevotella"                     
## [1927,] "Campylobacter"                  
## [1928,] "Actinobacillus"                 
## [1929,] NA                               
## [1930,] "Alloprevotella"                 
## [1931,] "Leptotrichia"                   
## [1932,] "Prevotella"                     
## [1933,] NA                               
## [1934,] "Prevotella"                     
## [1935,] NA                               
## [1936,] NA                               
## [1937,] NA                               
## [1938,] "Prevotella_7"                   
## [1939,] "Prevotella_2"                   
## [1940,] NA                               
## [1941,] "Gemella"                        
## [1942,] NA                               
## [1943,] "Prevotella"                     
## [1944,] "Bergeriella"                    
## [1945,] "Johnsonella"                    
## [1946,] "Treponema_2"                    
## [1947,] "Kingella"                       
## [1948,] NA                               
## [1949,] "Capnocytophaga"                 
## [1950,] NA                               
## [1951,] "Prevotella_7"                   
## [1952,] "Lachnoanaerobaculum"            
## [1953,] "Veillonella"                    
## [1954,] NA                               
## [1955,] NA                               
## [1956,] "Prevotella_2"                   
## [1957,] "Lachnoanaerobaculum"            
## [1958,] "Porphyromonas"                  
## [1959,] "Prevotella_7"                   
## [1960,] "Prevotella"                     
## [1961,] "Capnocytophaga"                 
## [1962,] "Atopobium"                      
## [1963,] "Alloprevotella"                 
## [1964,] "Actinomyces"                    
## [1965,] "Leptotrichia"                   
## [1966,] NA                               
## [1967,] "Prevotella_7"                   
## [1968,] "Veillonella"                    
## [1969,] "Fusobacterium"                  
## [1970,] "Neisseria"                      
## [1971,] NA                               
## [1972,] "Peptoanaerobacter"              
## [1973,] "Alloprevotella"                 
## [1974,] "Johnsonella"                    
## [1975,] "Peptoniphilus"                  
## [1976,] "Alloprevotella"                 
## [1977,] "Peptostreptococcus"             
## [1978,] "Alloprevotella"                 
## [1979,] "Porphyromonas"                  
## [1980,] "Prevotella_7"                   
## [1981,] "Prevotella_7"                   
## [1982,] "Fusobacterium"                  
## [1983,] "Veillonella"                    
## [1984,] "Neisseria"                      
## [1985,] "Porphyromonas"                  
## [1986,] "Veillonella"                    
## [1987,] "Haemophilus"                    
## [1988,] NA                               
## [1989,] "Leptotrichia"                   
## [1990,] NA                               
## [1991,] "Alloprevotella"                 
## [1992,] "Rikenellaceae_RC9_gut_group"    
## [1993,] "Capnocytophaga"                 
## [1994,] NA                               
## [1995,] "Centipeda"                      
## [1996,] "Gemella"                        
## [1997,] "Tannerella"                     
## [1998,] "Veillonella"                    
## [1999,] NA                               
## [2000,] "Neisseria"                      
## [2001,] "Leptotrichia"                   
## [2002,] "Peptostreptococcus"             
## [2003,] "Fusobacterium"                  
## [2004,] "F0058"                          
## [2005,] "Prevotella_7"                   
## [2006,] "Lachnoanaerobaculum"            
## [2007,] "Prevotella"                     
## [2008,] "Veillonella"                    
## [2009,] "Prevotella"                     
## [2010,] "Haemophilus"                    
## [2011,] "Prevotella"                     
## [2012,] "Porphyromonas"                  
## [2013,] NA                               
## [2014,] "Prevotella_7"                   
## [2015,] "Prevotella"                     
## [2016,] "Veillonella"                    
## [2017,] "Capnocytophaga"                 
## [2018,] NA                               
## [2019,] "Veillonella"                    
## [2020,] "Fusobacterium"                  
## [2021,] "Porphyromonas"                  
## [2022,] "Actinomyces"                    
## [2023,] "Actinobacillus"                 
## [2024,] "Haemophilus"                    
## [2025,] "Neisseria"                      
## [2026,] "Prevotella_7"                   
## [2027,] NA                               
## [2028,] "Prevotella"                     
## [2029,] "Leptotrichia"                   
## [2030,] NA                               
## [2031,] "Leptotrichia"                   
## [2032,] "Candidatus_Saccharimonas"       
## [2033,] NA                               
## [2034,] "Anthococcus"                    
## [2035,] "Anthococcus"                    
## [2036,] "Granulicatella"                 
## [2037,] "Fretibacterium"                 
## [2038,] "Prevotella_7"                   
## [2039,] "Fusobacterium"                  
## [2040,] "Selenomonas"                    
## [2041,] "Veillonella"                    
## [2042,] "Prevotella_7"                   
## [2043,] "Haemophilus"                    
## [2044,] "Bergeriella"                    
## [2045,] NA                               
## [2046,] "Leptotrichia"                   
## [2047,] "Ruminococcaceae_UCG-014"        
## [2048,] "Veillonella"                    
## [2049,] "Campylobacter"                  
## [2050,] "Prevotella_6"                   
## [2051,] "Prevotella_7"                   
## [2052,] "Dialister"                      
## [2053,] "Prevotella_7"                   
## [2054,] "Actinomyces"                    
## [2055,] "Prevotella_7"                   
## [2056,] "Selenomonas"                    
## [2057,] "Candidatus_Saccharimonas"       
## [2058,] NA                               
## [2059,] "Prevotella"                     
## [2060,] "Catonella"                      
## [2061,] "Alloprevotella"                 
## [2062,] "Veillonella"                    
## [2063,] NA                               
## [2064,] "Porphyromonas"                  
## [2065,] "Veillonella"                    
## [2066,] "Neisseria"                      
## [2067,] "Prevotella_7"                   
## [2068,] "Prevotella"                     
## [2069,] "Fusobacterium"                  
## [2070,] "Candidatus_Saccharimonas"       
## [2071,] "Prevotellaceae_YAB2003_group"   
## [2072,] NA                               
## [2073,] "Oribacterium"                   
## [2074,] "Haemophilus"                    
## [2075,] NA                               
## [2076,] "Veillonella"                    
## [2077,] "Treponema_2"                    
## [2078,] "Fusobacterium"                  
## [2079,] "Candidatus_Saccharimonas"       
## [2080,] "Porphyromonas"                  
## [2081,] "Capnocytophaga"                 
## [2082,] "Prevotella_7"                   
## [2083,] NA                               
## [2084,] NA                               
## [2085,] "Neisseria"                      
## [2086,] "Leptotrichia"                   
## [2087,] "Lachnoanaerobaculum"            
## [2088,] "Prevotella"                     
## [2089,] "Dialister"                      
## [2090,] "Prevotella_2"                   
## [2091,] "Treponema_2"                    
## [2092,] NA                               
## [2093,] NA                               
## [2094,] "Prevotella"                     
## [2095,] "Porphyromonas"                  
## [2096,] "Fusobacterium"                  
## [2097,] NA                               
## [2098,] "Treponema_2"                    
## [2099,] "Leptotrichia"                   
## [2100,] "Prevotella"                     
## [2101,] "Prevotella_7"                   
## [2102,] "Leptotrichia"                   
## [2103,] "Catonella"                      
## [2104,] "Prevotella"                     
## [2105,] "Leptotrichia"                   
## [2106,] "Aggregatibacter"                
## [2107,] "Prevotella_7"                   
## [2108,] "Veillonella"                    
## [2109,] "Prevotella_2"                   
## [2110,] "Prevotella_2"                   
## [2111,] "Alloprevotella"                 
## [2112,] NA                               
## [2113,] "Selenomonas_3"                  
## [2114,] "Neisseria"                      
## [2115,] "Prevotellaceae_YAB2003_group"   
## [2116,] NA                               
## [2117,] "Granulicatella"                 
## [2118,] "Prevotella_2"                   
## [2119,] "Selenomonas_3"                  
## [2120,] "Haemophilus"                    
## [2121,] "Capnocytophaga"                 
## [2122,] "Prevotellaceae_YAB2003_group"   
## [2123,] NA                               
## [2124,] "Selenomonas"                    
## [2125,] "Prevotella_7"                   
## [2126,] "Streptococcus"                  
## [2127,] "Alloprevotella"                 
## [2128,] "Veillonella"                    
## [2129,] "Streptococcus"                  
## [2130,] "Haemophilus"                    
## [2131,] "Alloprevotella"                 
## [2132,] NA                               
## [2133,] "Actinomyces"                    
## [2134,] "Fusobacterium"                  
## [2135,] "Prevotella_6"                   
## [2136,] "Veillonella"                    
## [2137,] "Rothia"                         
## [2138,] "Alloprevotella"                 
## [2139,] "Prevotella_7"                   
## [2140,] NA                               
## [2141,] "Treponema_2"                    
## [2142,] NA                               
## [2143,] NA                               
## [2144,] "Veillonella"                    
## [2145,] "Prevotella_7"                   
## [2146,] "Alloprevotella"                 
## [2147,] "Alloprevotella"                 
## [2148,] "Porphyromonas"                  
## [2149,] "Leptotrichia"                   
## [2150,] NA                               
## [2151,] NA                               
## [2152,] "Cardiobacterium"                
## [2153,] "Campylobacter"                  
## [2154,] "Prevotella_2"                   
## [2155,] "Prevotellaceae_YAB2003_group"   
## [2156,] "Prevotella_7"                   
## [2157,] "Neisseria"                      
## [2158,] NA                               
## [2159,] "Porphyromonas"                  
## [2160,] "Granulicatella"                 
## [2161,] "Haemophilus"                    
## [2162,] "Actinomyces"                    
## [2163,] NA                               
## [2164,] "Capnocytophaga"                 
## [2165,] "Selenomonas_3"                  
## [2166,] NA                               
## [2167,] "Prevotellaceae_YAB2003_group"   
## [2168,] "Treponema_2"                    
## [2169,] "Prevotella_2"                   
## [2170,] "Butyrivibrio_2"                 
## [2171,] "Fusobacterium"                  
## [2172,] "Oribacterium"                   
## [2173,] "Actinomyces"                    
## [2174,] "Alloprevotella"                 
## [2175,] "Alloprevotella"                 
## [2176,] "Leptotrichia"                   
## [2177,] "Capnocytophaga"                 
## [2178,] NA                               
## [2179,] "Veillonella"                    
## [2180,] NA                               
## [2181,] NA                               
## [2182,] "Peptostreptococcus"             
## [2183,] "Porphyromonas"                  
## [2184,] "Prevotella"                     
## [2185,] "Prevotella"                     
## [2186,] "Fusobacterium"                  
## [2187,] "Veillonella"                    
## [2188,] "Selenomonas_3"                  
## [2189,] "Fusobacterium"                  
## [2190,] NA                               
## [2191,] "Prevotella"                     
## [2192,] "Simonsiella"                    
## [2193,] "Bergeyella"                     
## [2194,] "Veillonella"                    
## [2195,] "Haemophilus"                    
## [2196,] NA                               
## [2197,] "Porphyromonas"                  
## [2198,] NA                               
## [2199,] "Alloprevotella"                 
## [2200,] NA                               
## [2201,] "F0332"                          
## [2202,] NA                               
## [2203,] "Selenomonas_3"                  
## [2204,] NA                               
## [2205,] NA                               
## [2206,] "Haemophilus"                    
## [2207,] "Prevotella_7"                   
## [2208,] "Aggregatibacter"                
## [2209,] "Candidatus_Saccharimonas"       
## [2210,] "Treponema_2"                    
## [2211,] NA                               
## [2212,] "Prevotella_7"                   
## [2213,] "Porphyromonas"                  
## [2214,] "Selenomonas_3"                  
## [2215,] NA                               
## [2216,] "Prevotella"                     
## [2217,] "Capnocytophaga"                 
## [2218,] "Selenomonas_3"                  
## [2219,] "Prevotella_7"                   
## [2220,] NA                               
## [2221,] NA                               
## [2222,] "Capnocytophaga"                 
## [2223,] NA                               
## [2224,] "Capnocytophaga"                 
## [2225,] "Veillonella"                    
## [2226,] NA                               
## [2227,] "Streptococcus"                  
## [2228,] "Prevotella_7"                   
## [2229,] "Lachnoanaerobaculum"            
## [2230,] NA                               
## [2231,] "Actinomyces"                    
## [2232,] "Selenomonas_3"                  
## [2233,] "Prevotella_2"                   
## [2234,] "Selenomonas"                    
## [2235,] "Lachnoanaerobaculum"            
## [2236,] "Veillonella"                    
## [2237,] "Prevotella_7"                   
## [2238,] NA                               
## [2239,] "Veillonella"                    
## [2240,] NA                               
## [2241,] "Prevotella_7"                   
## [2242,] NA                               
## [2243,] "Porphyromonas"                  
## [2244,] "Prevotellaceae_YAB2003_group"   
## [2245,] "Haemophilus"                    
## [2246,] NA                               
## [2247,] "Prevotellaceae_YAB2003_group"   
## [2248,] "Porphyromonas"                  
## [2249,] NA                               
## [2250,] "Candidatus_Saccharimonas"       
## [2251,] "Atopobium"                      
## [2252,] "Prevotella_2"                   
## [2253,] "Selenomonas"                    
## [2254,] "Dialister"                      
## [2255,] "Fretibacterium"                 
## [2256,] "Prevotella"                     
## [2257,] "Prevotella_7"                   
## [2258,] "Peptoanaerobacter"              
## [2259,] "Corynebacterium"                
## [2260,] "Gemella"                        
## [2261,] "Veillonella"                    
## [2262,] "Necropsobacter"                 
## [2263,] NA                               
## [2264,] "Leptotrichia"                   
## [2265,] "Prevotella_7"                   
## [2266,] "Leptotrichia"                   
## [2267,] "Anaeroglobus"                   
## [2268,] NA                               
## [2269,] "Capnocytophaga"                 
## [2270,] "Prevotella_2"                   
## [2271,] "Leptotrichia"                   
## [2272,] NA                               
## [2273,] NA                               
## [2274,] NA                               
## [2275,] "Porphyromonas"                  
## [2276,] "Capnocytophaga"                 
## [2277,] "Stomatobaculum"                 
## [2278,] "Selenomonas"                    
## [2279,] "Actinobacillus"                 
## [2280,] NA                               
## [2281,] NA                               
## [2282,] NA                               
## [2283,] "Prevotella_7"                   
## [2284,] "Prevotella_2"                   
## [2285,] "Prevotella"                     
## [2286,] "Lachnoanaerobaculum"            
## [2287,] "Alloprevotella"                 
## [2288,] NA                               
## [2289,] NA                               
## [2290,] "Leptotrichia"                   
## [2291,] "Veillonella"                    
## [2292,] "Aggregatibacter"                
## [2293,] "Prevotella"                     
## [2294,] "Alloprevotella"                 
## [2295,] "Porphyromonas"                  
## [2296,] "Neisseria"                      
## [2297,] "Prevotella_7"                   
## [2298,] "Prevotella"                     
## [2299,] "Prevotella_7"                   
## [2300,] "Prevotella_7"                   
## [2301,] NA                               
## [2302,] NA                               
## [2303,] "Scardovia"                      
## [2304,] NA                               
## [2305,] "Treponema_2"                    
## [2306,] "Bergeyella"                     
## [2307,] NA                               
## [2308,] "Capnocytophaga"                 
## [2309,] "Peptostreptococcus"             
## [2310,] "Prevotella_7"                   
## [2311,] NA                               
## [2312,] "Veillonella"                    
## [2313,] NA                               
## [2314,] "Prevotella"                     
## [2315,] NA                               
## [2316,] NA                               
## [2317,] NA                               
## [2318,] "Porphyromonas"                  
## [2319,] "Stomatobaculum"                 
## [2320,] NA                               
## [2321,] "Alloprevotella"                 
## [2322,] "Prevotella_7"                   
## [2323,] "Prevotella_6"                   
## [2324,] "Alloprevotella"                 
## [2325,] "Prevotella_6"                   
## [2326,] NA                               
## [2327,] NA                               
## [2328,] "Prevotella_7"                   
## [2329,] "Ruminococcaceae_UCG-014"        
## [2330,] "Alloprevotella"                 
## [2331,] "Porphyromonas"                  
## [2332,] NA                               
## [2333,] "Prevotella"                     
## [2334,] "Prevotellaceae_YAB2003_group"   
## [2335,] "Neisseria"                      
## [2336,] "Kingella"                       
## [2337,] "Prevotella"                     
## [2338,] "Prevotella"                     
## [2339,] "Streptococcus"                  
## [2340,] "Alloprevotella"                 
## [2341,] NA                               
## [2342,] "Lachnoanaerobaculum"            
## [2343,] "Haemophilus"                    
## [2344,] "Porphyromonas"                  
## [2345,] "Porphyromonas"                  
## [2346,] NA                               
## [2347,] NA                               
## [2348,] "Corynebacterium"                
## [2349,] "Fretibacterium"                 
## [2350,] "Treponema_2"                    
## [2351,] NA                               
## [2352,] "Prevotella"                     
## [2353,] "Actinomyces"                    
## [2354,] NA                               
## [2355,] "Lachnoanaerobaculum"            
## [2356,] "Prevotella"                     
## [2357,] "Veillonella"                    
## [2358,] "Prevotella"                     
## [2359,] "Prevotella_7"                   
## [2360,] "Selenomonas_3"                  
## [2361,] NA                               
## [2362,] "Selenomonas_3"                  
## [2363,] "Prevotella_7"                   
## [2364,] NA                               
## [2365,] "Selenomonas"                    
## [2366,] NA                               
## [2367,] "Actinomyces"                    
## [2368,] "Prevotella"                     
## [2369,] "Eikenella"                      
## [2370,] "Oceanivirga"                    
## [2371,] "Porphyromonas"                  
## [2372,] "Veillonella"                    
## [2373,] "Alloprevotella"                 
## [2374,] "Alloprevotella"                 
## [2375,] "Selenomonas_3"                  
## [2376,] "Peptoanaerobacter"              
## [2377,] "Alysiella"                      
## [2378,] "Treponema_2"                    
## [2379,] "Prevotella"                     
## [2380,] "Fusobacterium"                  
## [2381,] "Prevotella_7"                   
## [2382,] "Leptotrichia"                   
## [2383,] NA                               
## [2384,] "F0058"                          
## [2385,] "Haemophilus"                    
## [2386,] "Alloprevotella"                 
## [2387,] "Veillonella"                    
## [2388,] "Actinomyces"                    
## [2389,] "Alloprevotella"                 
## [2390,] "Veillonella"                    
## [2391,] "Leptotrichia"                   
## [2392,] "Selenomonas_3"                  
## [2393,] "Leptotrichia"                   
## [2394,] NA                               
## [2395,] "Selenomonas"                    
## [2396,] "Veillonella"                    
## [2397,] "Veillonella"                    
## [2398,] "Corynebacterium"                
## [2399,] "Actinomyces"                    
## [2400,] "Prevotella_7"                   
## [2401,] "Atopobium"                      
## [2402,] "Treponema_2"                    
## [2403,] "Prevotella_7"                   
## [2404,] NA                               
## [2405,] "Lachnoanaerobaculum"            
## [2406,] "Veillonella"                    
## [2407,] "Alloprevotella"                 
## [2408,] "Prevotella_7"                   
## [2409,] "Leptotrichia"                   
## [2410,] "Prevotella_7"                   
## [2411,] "Veillonella"                    
## [2412,] NA                               
## [2413,] "Candidatus_Saccharimonas"       
## [2414,] "Prevotellaceae_YAB2003_group"   
## [2415,] "Selenomonas_3"                  
## [2416,] "Selenomonas_3"                  
## [2417,] "Veillonella"                    
## [2418,] "Alloprevotella"                 
## [2419,] "Prevotella_7"                   
## [2420,] "Johnsonella"                    
## [2421,] "Haemophilus"                    
## [2422,] NA                               
## [2423,] NA                               
## [2424,] NA                               
## [2425,] "Alloprevotella"                 
## [2426,] NA                               
## [2427,] "Prevotella_6"                   
## [2428,] "Porphyromonas"                  
## [2429,] "Capnocytophaga"                 
## [2430,] "Alloprevotella"                 
## [2431,] "Capnocytophaga"                 
## [2432,] "Johnsonella"                    
## [2433,] "Selenomonas_3"                  
## [2434,] "Prevotella_2"                   
## [2435,] NA                               
## [2436,] "Prevotella_7"                   
## [2437,] "Treponema_2"                    
## [2438,] NA                               
## [2439,] "Prevotella_7"                   
## [2440,] NA                               
## [2441,] "Actinomyces"                    
## [2442,] "Porphyromonas"                  
## [2443,] NA                               
## [2444,] "Haemophilus"                    
## [2445,] "Ruminococcaceae_UCG-014"        
## [2446,] "Alloprevotella"                 
## [2447,] "Prevotella_6"                   
## [2448,] "Anthococcus"                    
## [2449,] NA                               
## [2450,] NA                               
## [2451,] NA                               
## [2452,] "Prevotella_7"                   
## [2453,] "Bergeriella"                    
## [2454,] "Prevotellaceae_YAB2003_group"   
## [2455,] "Johnsonella"                    
## [2456,] "Actinomyces"                    
## [2457,] "Kingella"                       
## [2458,] "Prevotella_2"                   
## [2459,] "Lachnoanaerobaculum"            
## [2460,] "Treponema_2"                    
## [2461,] "Stomatobaculum"                 
## [2462,] "Dialister"                      
## [2463,] "Alloprevotella"                 
## [2464,] "Peptostreptococcus"             
## [2465,] "Prevotella"                     
## [2466,] "Prevotella_6"                   
## [2467,] NA                               
## [2468,] "Bergeyella"                     
## [2469,] "Prevotella"                     
## [2470,] "Prevotella_2"                   
## [2471,] "Bergeriella"                    
## [2472,] "Aggregatibacter"                
## [2473,] NA                               
## [2474,] "Peptococcus"                    
## [2475,] "Prevotellaceae_YAB2003_group"   
## [2476,] NA                               
## [2477,] NA                               
## [2478,] "Selenomonas_3"                  
## [2479,] "Centipeda"                      
## [2480,] NA                               
## [2481,] NA                               
## [2482,] "Lachnoanaerobaculum"            
## [2483,] "Prevotella_7"                   
## [2484,] "Treponema_2"                    
## [2485,] "Neisseria"                      
## [2486,] "Family_XIII_UCG-001"            
## [2487,] "Prevotella_7"                   
## [2488,] "Prevotella_7"                   
## [2489,] "Haemophilus"                    
## [2490,] "Prevotella"                     
## [2491,] NA                               
## [2492,] "Veillonella"                    
## [2493,] NA                               
## [2494,] "Prevotella"                     
## [2495,] "Capnocytophaga"                 
## [2496,] "Fusobacterium"                  
## [2497,] "Actinomyces"                    
## [2498,] NA                               
## [2499,] NA                               
## [2500,] "Corynebacterium"                
## [2501,] "Selenomonas_3"                  
## [2502,] "Prevotella"                     
## [2503,] "Alloprevotella"                 
## [2504,] NA                               
## [2505,] "Fusobacterium"                  
## [2506,] "Alloprevotella"                 
## [2507,] "Capnocytophaga"                 
## [2508,] "Prevotella"                     
## [2509,] "Streptococcus"                  
## [2510,] NA                               
## [2511,] "Capnocytophaga"                 
## [2512,] "Neisseria"                      
## [2513,] "Lachnoanaerobaculum"            
## [2514,] NA                               
## [2515,] "Alloprevotella"                 
## [2516,] "Kingella"                       
## [2517,] "Rothia"                         
## [2518,] NA                               
## [2519,] "Actinomyces"                    
## [2520,] "Actinomyces"                    
## [2521,] "Bergeriella"                    
## [2522,] NA                               
## [2523,] "Alloprevotella"                 
## [2524,] "Selenomonas"                    
## [2525,] "Neisseria"                      
## [2526,] NA                               
## [2527,] "Prevotella"                     
## [2528,] NA                               
## [2529,] NA                               
## [2530,] "Tannerella"                     
## [2531,] "Selenomonas_3"                  
## [2532,] "Dialister"                      
## [2533,] "Alloprevotella"                 
## [2534,] "Capnocytophaga"                 
## [2535,] "Prevotella_7"                   
## [2536,] NA                               
## [2537,] "Anthococcus"                    
## [2538,] NA                               
## [2539,] NA                               
## [2540,] "Haemophilus"                    
## [2541,] "Prevotellaceae_YAB2003_group"   
## [2542,] "Ruminococcaceae_UCG-014"        
## [2543,] NA                               
## [2544,] "Prevotella_6"                   
## [2545,] NA                               
## [2546,] NA                               
## [2547,] "Leptotrichia"                   
## [2548,] NA                               
## [2549,] "Veillonella"                    
## [2550,] NA                               
## [2551,] "Megasphaera"                    
## [2552,] "Prevotella_7"                   
## [2553,] "Leptotrichia"                   
## [2554,] "Leptotrichia"                   
## [2555,] "Streptobacillus"                
## [2556,] "Capnocytophaga"                 
## [2557,] "Haemophilus"                    
## [2558,] "Treponema_2"                    
## [2559,] NA                               
## [2560,] "Dialister"                      
## [2561,] "Veillonella"                    
## [2562,] "Treponema_2"                    
## [2563,] "Treponema_2"                    
## [2564,] NA                               
## [2565,] "Parvimonas"                     
## [2566,] NA                               
## [2567,] "Haemophilus"                    
## [2568,] "Anthococcus"                    
## [2569,] "Prevotella_6"                   
## [2570,] "Veillonella"                    
## [2571,] "Prevotella_6"                   
## [2572,] "Prevotella_7"                   
## [2573,] "Aggregatibacter"                
## [2574,] "Leptotrichia"                   
## [2575,] NA                               
## [2576,] "Prevotella_7"                   
## [2577,] "Actinomyces"                    
## [2578,] "Leptotrichia"                   
## [2579,] "Capnocytophaga"                 
## [2580,] "Porphyromonas"                  
## [2581,] "Porphyromonas"                  
## [2582,] "Fusobacterium"                  
## [2583,] "Prevotella_7"                   
## [2584,] "Selenomonas_3"                  
## [2585,] "Capnocytophaga"                 
## [2586,] NA                               
## [2587,] "Rikenellaceae_RC9_gut_group"    
## [2588,] NA                               
## [2589,] "Peptostreptococcus"             
## [2590,] "Prevotella_2"                   
## [2591,] NA                               
## [2592,] "Veillonella"                    
## [2593,] "Lactobacillus"                  
## [2594,] "Neisseria"                      
## [2595,] "Prevotella"                     
## [2596,] "Streptococcus"                  
## [2597,] "Prevotella_7"                   
## [2598,] NA                               
## [2599,] NA                               
## [2600,] "Selenomonas_3"                  
## [2601,] "Selenomonas_3"                  
## [2602,] "Cardiobacterium"                
## [2603,] "Prevotella_7"                   
## [2604,] "Veillonella"                    
## [2605,] "Prevotella"                     
## [2606,] NA                               
## [2607,] NA                               
## [2608,] NA                               
## [2609,] NA                               
## [2610,] NA                               
## [2611,] "Filifactor"                     
## [2612,] "Prevotella_7"                   
## [2613,] NA                               
## [2614,] "Prevotella"                     
## [2615,] NA                               
## [2616,] "Prevotella"                     
## [2617,] "Haemophilus"                    
## [2618,] "Veillonella"                    
## [2619,] "Aggregatibacter"                
## [2620,] "Leptotrichia"                   
## [2621,] "Fusobacterium"                  
## [2622,] "Prevotella"                     
## [2623,] "Capnocytophaga"                 
## [2624,] "Atlantibacter"                  
## [2625,] "Lachnoanaerobaculum"            
## [2626,] NA                               
## [2627,] "Veillonella"                    
## [2628,] NA                               
## [2629,] "Corynebacterium"                
## [2630,] "Oribacterium"                   
## [2631,] NA                               
## [2632,] "Treponema_2"                    
## [2633,] "Gemella"                        
## [2634,] "Prevotella_7"                   
## [2635,] NA                               
## [2636,] "Dialister"                      
## [2637,] NA                               
## [2638,] NA                               
## [2639,] "Alloprevotella"                 
## [2640,] NA                               
## [2641,] "Campylobacter"                  
## [2642,] "Veillonella"                    
## [2643,] "Prevotellaceae_YAB2003_group"   
## [2644,] "Streptococcus"                  
## [2645,] NA                               
## [2646,] "Parvimonas"                     
## [2647,] "Prevotella"                     
## [2648,] "Ruminococcaceae_UCG-014"        
## [2649,] "Capnocytophaga"                 
## [2650,] "Streptococcus"                  
## [2651,] "Treponema_2"                    
## [2652,] "Actinobacillus"                 
## [2653,] "Leptotrichia"                   
## [2654,] "Prevotella_7"                   
## [2655,] "Selenomonas_3"                  
## [2656,] "Tannerella"                     
## [2657,] "Alloprevotella"                 
## [2658,] "Peptoanaerobacter"              
## [2659,] "Prevotellaceae_YAB2003_group"   
## [2660,] "Capnocytophaga"                 
## [2661,] "Capnocytophaga"                 
## [2662,] "Selenomonas_3"                  
## [2663,] "Actinomyces"                    
## [2664,] "Actinomyces"                    
## [2665,] "Leptotrichia"                   
## [2666,] "Selenomonas_3"                  
## [2667,] NA                               
## [2668,] "Selenomonas_3"                  
## [2669,] "Parvimonas"                     
## [2670,] "Capnocytophaga"                 
## [2671,] "Veillonella"                    
## [2672,] "Necropsobacter"                 
## [2673,] "Prevotella"                     
## [2674,] "Prevotella"                     
## [2675,] "Treponema_2"                    
## [2676,] "Prevotella"                     
## [2677,] "Actinomyces"                    
## [2678,] NA                               
## [2679,] "Raoultella"                     
## [2680,] NA                               
## [2681,] "Lautropia"                      
## [2682,] "Prevotella_7"                   
## [2683,] "Actinomyces"                    
## [2684,] "Capnocytophaga"                 
## [2685,] "Prevotella_7"                   
## [2686,] NA                               
## [2687,] "Atopobium"                      
## [2688,] NA                               
## [2689,] NA                               
## [2690,] NA                               
## [2691,] NA                               
## [2692,] "Fusobacterium"                  
## [2693,] NA                               
## [2694,] "Prevotella"                     
## [2695,] "Prevotella_2"                   
## [2696,] NA                               
## [2697,] "Veillonella"                    
## [2698,] "Veillonella"                    
## [2699,] "Veillonella"                    
## [2700,] "Prevotella_7"                   
## [2701,] NA                               
## [2702,] "Veillonella"                    
## [2703,] NA                               
## [2704,] "Prevotella"                     
## [2705,] "Chelonobacter"                  
## [2706,] "Fusobacterium"                  
## [2707,] "Selenomonas_3"                  
## [2708,] "Fusobacterium"                  
## [2709,] "Veillonella"                    
## [2710,] "Prevotella_7"                   
## [2711,] "Capnocytophaga"                 
## [2712,] NA                               
## [2713,] "Veillonella"                    
## [2714,] "Fusobacterium"                  
## [2715,] "Prevotella_7"                   
## [2716,] "Leptotrichia"                   
## [2717,] NA                               
## [2718,] "Prevotella_2"                   
## [2719,] "Atlantibacter"                  
## [2720,] NA                               
## [2721,] "Treponema_2"                    
## [2722,] NA                               
## [2723,] "Oribacterium"                   
## [2724,] "Prevotella_7"                   
## [2725,] "Prevotella"                     
## [2726,] "Kingella"                       
## [2727,] "Prevotella_7"                   
## [2728,] "Actinomyces"                    
## [2729,] "Prevotella_7"                   
## [2730,] NA                               
## [2731,] NA                               
## [2732,] "Kingella"                       
## [2733,] "Prevotella_2"                   
## [2734,] "Leptotrichia"                   
## [2735,] "Fusobacterium"                  
## [2736,] "Leptotrichia"                   
## [2737,] "Aggregatibacter"                
## [2738,] "Prevotella_7"                   
## [2739,] NA                               
## [2740,] NA                               
## [2741,] NA                               
## [2742,] NA                               
## [2743,] "Solobacterium"                  
## [2744,] "Selenomonas_4"                  
## [2745,] "Neisseria"                      
## [2746,] "Kingella"                       
## [2747,] "Centipeda"                      
## [2748,] "Prevotella_2"                   
## [2749,] "Prevotella_6"                   
## [2750,] "Prevotella_6"                   
## [2751,] NA                               
## [2752,] "Prevotella_7"                   
## [2753,] "Tannerella"                     
## [2754,] "Alloprevotella"                 
## [2755,] "Eikenella"                      
## [2756,] NA                               
## [2757,] "Capnocytophaga"                 
## [2758,] "Fusobacterium"                  
## [2759,] "Prevotella_7"                   
## [2760,] "Atopobium"                      
## [2761,] "Prevotella_7"                   
## [2762,] "Porphyromonas"                  
## [2763,] "Prevotella_2"                   
## [2764,] NA                               
## [2765,] NA                               
## [2766,] "Alloprevotella"                 
## [2767,] "Streptococcus"                  
## [2768,] NA                               
## [2769,] NA                               
## [2770,] "Selenomonas"                    
## [2771,] NA                               
## [2772,] NA                               
## [2773,] "Prevotellaceae_YAB2003_group"   
## [2774,] "Selenomonas"                    
## [2775,] NA                               
## [2776,] "Corynebacterium"                
## [2777,] "Eikenella"                      
## [2778,] "Corynebacterium"                
## [2779,] "Tannerella"                     
## [2780,] "Bergeyella"                     
## [2781,] NA                               
## [2782,] "Capnocytophaga"                 
## [2783,] NA                               
## [2784,] NA                               
## [2785,] "Corynebacterium"                
## [2786,] "Actinomyces"                    
## [2787,] "Prevotella_7"                   
## [2788,] "Veillonella"                    
## [2789,] NA                               
## [2790,] "Alloprevotella"                 
## [2791,] "Prevotella"                     
## [2792,] "Bergeyella"                     
## [2793,] NA                               
## [2794,] NA                               
## [2795,] "Neisseria"                      
## [2796,] "Prevotellaceae_YAB2003_group"   
## [2797,] "Prevotella_7"                   
## [2798,] "Capnocytophaga"                 
## [2799,] "Prevotella"                     
## [2800,] NA                               
## [2801,] "Actinobacillus"                 
## [2802,] "Capnocytophaga"                 
## [2803,] "Prevotella_7"                   
## [2804,] "Veillonella"                    
## [2805,] NA                               
## [2806,] "Prevotella_7"                   
## [2807,] "Alloprevotella"                 
## [2808,] NA                               
## [2809,] "Leptotrichia"                   
## [2810,] "Prevotella_7"                   
## [2811,] "Prevotella_7"                   
## [2812,] "Fusobacterium"                  
## [2813,] "Haemophilus"                    
## [2814,] NA                               
## [2815,] "Prevotella"                     
## [2816,] "Prevotella_7"                   
## [2817,] "Prevotella_7"                   
## [2818,] "Selenomonas_3"                  
## [2819,] "Stomatobaculum"                 
## [2820,] "Alloprevotella"                 
## [2821,] "Selenomonas_3"                  
## [2822,] "Prevotella"                     
## [2823,] "Alloprevotella"                 
## [2824,] "Alloprevotella"                 
## [2825,] "Bergeyella"                     
## [2826,] NA                               
## [2827,] "Megasphaera"                    
## [2828,] "Selenomonas"                    
## [2829,] NA                               
## [2830,] "Peptoanaerobacter"              
## [2831,] "Candidatus_Saccharimonas"       
## [2832,] "Aggregatibacter"                
## [2833,] "Haemophilus"                    
## [2834,] "Treponema_2"                    
## [2835,] NA                               
## [2836,] "Alloprevotella"                 
## [2837,] "Dialister"                      
## [2838,] NA                               
## [2839,] "Prevotella_7"                   
## [2840,] "Parvimonas"                     
## [2841,] "Lachnoanaerobaculum"            
## [2842,] "Selenomonas_3"                  
## [2843,] "Lautropia"                      
## [2844,] "Selenomonas_3"                  
## [2845,] "Prevotellaceae_YAB2003_group"   
## [2846,] "Selenomonas"                    
## [2847,] "Aggregatibacter"                
## [2848,] NA                               
## [2849,] "Selenomonas_3"                  
## [2850,] "Prevotella_7"                   
## [2851,] "Kingella"                       
## [2852,] "Fusobacterium"                  
## [2853,] "Selenomonas_3"                  
## [2854,] "Prevotella_7"                   
## [2855,] "Neisseria"                      
## [2856,] "Actinomyces"                    
## [2857,] "Simonsiella"                    
## [2858,] "Porphyromonas"                  
## [2859,] "Prevotella_2"                   
## [2860,] "Bergeriella"                    
## [2861,] "Selenomonas_3"                  
## [2862,] NA                               
## [2863,] "Bergeyella"                     
## [2864,] NA                               
## [2865,] "Tannerella"                     
## [2866,] NA                               
## [2867,] NA                               
## [2868,] NA                               
## [2869,] NA                               
## [2870,] "Lachnospiraceae_AC2044_group"   
## [2871,] "Prevotella"                     
## [2872,] NA                               
## [2873,] "Haemophilus"                    
## [2874,] "Prevotella_7"                   
## [2875,] "Capnocytophaga"                 
## [2876,] "Prevotella_7"                   
## [2877,] NA                               
## [2878,] "Prevotella_7"                   
## [2879,] "Anthococcus"                    
## [2880,] NA                               
## [2881,] NA                               
## [2882,] "Cardiobacterium"                
## [2883,] "Selenomonas"                    
## [2884,] NA                               
## [2885,] "Prevotella"                     
## [2886,] NA                               
## [2887,] "Stomatobaculum"                 
## [2888,] NA                               
## [2889,] "Capnocytophaga"                 
## [2890,] "Prevotella_7"                   
## [2891,] "Neisseria"                      
## [2892,] "Alloprevotella"                 
## [2893,] "Prevotella_7"                   
## [2894,] NA                               
## [2895,] NA                               
## [2896,] "Leptotrichia"                   
## [2897,] "Alloprevotella"                 
## [2898,] NA                               
## [2899,] NA                               
## [2900,] "Fusobacterium"                  
## [2901,] "Prevotella"                     
## [2902,] "Dialister"                      
## [2903,] "Actinomyces"                    
## [2904,] "Prevotella"                     
## [2905,] "Treponema_2"                    
## [2906,] "Haemophilus"                    
## [2907,] "Treponema_2"                    
## [2908,] "Campylobacter"                  
## [2909,] "Lachnoanaerobaculum"            
## [2910,] "Cobetia"                        
## [2911,] "Veillonella"                    
## [2912,] "Mycoplasma"                     
## [2913,] "Alloprevotella"                 
## [2914,] NA                               
## [2915,] NA                               
## [2916,] "Capnocytophaga"                 
## [2917,] "Candidatus_Saccharimonas"       
## [2918,] NA                               
## [2919,] "Fusobacterium"                  
## [2920,] "Veillonella"                    
## [2921,] "Porphyromonas"                  
## [2922,] "Actinomyces"                    
## [2923,] NA                               
## [2924,] "Anthococcus"                    
## [2925,] "Catonella"                      
## [2926,] "Leptotrichia"                   
## [2927,] NA                               
## [2928,] "Streptococcus"                  
## [2929,] "Arcanobacterium"                
## [2930,] NA                               
## [2931,] "Faecalibacterium"               
## [2932,] "Necropsobacter"                 
## [2933,] NA                               
## [2934,] "Atopobium"                      
## [2935,] "Actinomyces"                    
## [2936,] "Veillonella"                    
## [2937,] NA                               
## [2938,] NA                               
## [2939,] "Prevotella_7"                   
## [2940,] "Prevotella"                     
## [2941,] NA                               
## [2942,] NA                               
## [2943,] "Alloprevotella"                 
## [2944,] "Staphylococcus"                 
## [2945,] "Capnocytophaga"                 
## [2946,] NA                               
## [2947,] "Tannerella"                     
## [2948,] NA                               
## [2949,] "Catonella"                      
## [2950,] "Selenomonas"                    
## [2951,] NA                               
## [2952,] "Prevotella_7"                   
## [2953,] "Prevotella_7"                   
## [2954,] "Capnocytophaga"                 
## [2955,] "Oribacterium"                   
## [2956,] NA                               
## [2957,] "Haemophilus"                    
## [2958,] "Leptotrichia"                   
## [2959,] "Veillonella"                    
## [2960,] "Veillonella"                    
## [2961,] "Alloprevotella"                 
## [2962,] "Aggregatibacter"                
## [2963,] "Veillonella"                    
## [2964,] "Veillonella"                    
## [2965,] "Eikenella"                      
## [2966,] "Selenomonas_3"                  
## [2967,] "Veillonella"                    
## [2968,] NA                               
## [2969,] NA                               
## [2970,] "Leptotrichia"                   
## [2971,] "Neisseria"                      
## [2972,] NA                               
## [2973,] "Prevotella_7"                   
## [2974,] "Rikenellaceae_RC9_gut_group"    
## [2975,] "Gemella"                        
## [2976,] "Treponema_2"                    
## [2977,] "Lachnoanaerobaculum"            
## [2978,] "Mycoplasma"                     
## [2979,] "Veillonella"                    
## [2980,] "Prevotella"                     
## [2981,] "Citrobacter"                    
## [2982,] "Treponema_2"                    
## [2983,] NA                               
## [2984,] "Porphyromonas"                  
## [2985,] "Johnsonella"                    
## [2986,] NA                               
## [2987,] "F0332"                          
## [2988,] "Prevotella_2"                   
## [2989,] "Capnocytophaga"                 
## [2990,] NA                               
## [2991,] "Dialister"                      
## [2992,] "Fretibacterium"                 
## [2993,] "Streptococcus"                  
## [2994,] "Alloscardovia"                  
## [2995,] "Prevotella_6"                   
## [2996,] "Catonella"                      
## [2997,] "Dialister"                      
## [2998,] NA                               
## [2999,] "Rahnella"                       
## [3000,] "Selenomonas_3"                  
## [3001,] "Capnocytophaga"                 
## [3002,] "Leptotrichia"                   
## [3003,] NA                               
## [3004,] "Prevotella_7"                   
## [3005,] NA                               
## [3006,] "Cricetibacter"                  
## [3007,] "Veillonella"                    
## [3008,] "Lawsonella"                     
## [3009,] "Johnsonella"                    
## [3010,] "Selenomonas"                    
## [3011,] NA                               
## [3012,] "Selenomonas_3"                  
## [3013,] NA                               
## [3014,] "Prevotella"                     
## [3015,] "Treponema_2"                    
## [3016,] NA                               
## [3017,] NA                               
## [3018,] NA                               
## [3019,] "Porphyromonas"                  
## [3020,] NA                               
## [3021,] "Alloprevotella"                 
## [3022,] "Peptoanaerobacter"              
## [3023,] "Prevotella_7"                   
## [3024,] "Prevotella"                     
## [3025,] "Prevotella_7"                   
## [3026,] NA                               
## [3027,] "Prevotella_7"                   
## [3028,] NA                               
## [3029,] "Fusobacterium"                  
## [3030,] "Capnocytophaga"                 
## [3031,] "Prevotella"                     
## [3032,] NA                               
## [3033,] "Prevotella"                     
## [3034,] NA                               
## [3035,] NA                               
## [3036,] "Stomatobaculum"                 
## [3037,] "Lachnoanaerobaculum"            
## [3038,] "Alloprevotella"                 
## [3039,] "Actinomyces"                    
## [3040,] "Anthococcus"                    
## [3041,] NA                               
## [3042,] "Alloprevotella"                 
## [3043,] "Selenomonas_3"                  
## [3044,] "Selenomonas"                    
## [3045,] "Capnocytophaga"                 
## [3046,] NA                               
## [3047,] "Actinomyces"                    
## [3048,] "Cricetibacter"                  
## [3049,] NA                               
## [3050,] NA                               
## [3051,] NA                               
## [3052,] "Moraxella"                      
## [3053,] NA                               
## [3054,] NA                               
## [3055,] "Filifactor"                     
## [3056,] "Porphyromonas"                  
## [3057,] "Prevotella"                     
## [3058,] "Oceanivirga"                    
## [3059,] "Prevotella_7"                   
## [3060,] "Capnocytophaga"                 
## [3061,] "Porphyromonas"                  
## [3062,] "Leptotrichia"                   
## [3063,] "Veillonella"                    
## [3064,] "Lachnoanaerobaculum"            
## [3065,] "Alloscardovia"                  
## [3066,] "Lachnoanaerobaculum"            
## [3067,] "Selenomonas_3"                  
## [3068,] "Selenomonas_3"                  
## [3069,] "Candidatus_Saccharimonas"       
## [3070,] NA                               
## [3071,] NA                               
## [3072,] "Selenomonas_3"                  
## [3073,] "Capnocytophaga"                 
## [3074,] "Streptobacillus"                
## [3075,] "Alloprevotella"                 
## [3076,] "Lautropia"                      
## [3077,] NA                               
## [3078,] "Leptotrichia"                   
## [3079,] "Simonsiella"                    
## [3080,] "Porphyromonas"                  
## [3081,] "Treponema_2"                    
## [3082,] "Actinomyces"                    
## [3083,] "Rothia"                         
## [3084,] "Prevotella_7"                   
## [3085,] "Fusobacterium"                  
## [3086,] NA                               
## [3087,] "Prevotellaceae_YAB2003_group"   
## [3088,] NA                               
## [3089,] NA                               
## [3090,] "Selenomonas_3"                  
## [3091,] "Selenomonas_3"                  
## [3092,] "Corynebacterium"                
## [3093,] NA                               
## [3094,] "Actinomyces"                    
## [3095,] NA                               
## [3096,] NA                               
## [3097,] "Centipeda"                      
## [3098,] "Veillonella"                    
## [3099,] NA                               
## [3100,] "Prevotella_7"                   
## [3101,] "Prevotella"                     
## [3102,] "Treponema_2"                    
## [3103,] NA                               
## [3104,] NA                               
## [3105,] NA                               
## [3106,] "Actinomyces"                    
## [3107,] "Prevotella"                     
## [3108,] "Prevotella_7"                   
## [3109,] "Haemophilus"                    
## [3110,] NA                               
## [3111,] NA                               
## [3112,] NA                               
## [3113,] NA                               
## [3114,] "Lachnoanaerobaculum"            
## [3115,] NA                               
## [3116,] NA                               
## [3117,] "Anthococcus"                    
## [3118,] "Bergeyella"                     
## [3119,] NA                               
## [3120,] "Prevotella_2"                   
## [3121,] "Alloprevotella"                 
## [3122,] NA                               
## [3123,] "Actinomyces"                    
## [3124,] "Prevotellaceae_YAB2003_group"   
## [3125,] "Prevotella_7"                   
## [3126,] "Defluviitaleaceae_UCG-011"      
## [3127,] NA                               
## [3128,] "Prevotella_2"                   
## [3129,] NA                               
## [3130,] "Actinomyces"                    
## [3131,] "Selenomonas_3"                  
## [3132,] "Prevotella_7"                   
## [3133,] "Capnocytophaga"                 
## [3134,] NA                               
## [3135,] "Prevotella_7"                   
## [3136,] NA                               
## [3137,] NA                               
## [3138,] "Capnocytophaga"                 
## [3139,] "Capnocytophaga"                 
## [3140,] "Porphyromonas"                  
## [3141,] "Prevotella_7"                   
## [3142,] "Prevotella"                     
## [3143,] "Prevotella"                     
## [3144,] "Aggregatibacter"                
## [3145,] "Candidatus_Saccharimonas"       
## [3146,] "Leptotrichia"                   
## [3147,] "Leptotrichia"                   
## [3148,] NA                               
## [3149,] NA                               
## [3150,] "Treponema_2"                    
## [3151,] "Ruminococcaceae_UCG-014"        
## [3152,] NA                               
## [3153,] "Treponema_2"                    
## [3154,] "Prevotella"                     
## [3155,] "Prevotella"                     
## [3156,] "Prevotella_7"                   
## [3157,] "Veillonella"                    
## [3158,] "Johnsonella"                    
## [3159,] "Neisseria"                      
## [3160,] "Tannerella"                     
## [3161,] NA                               
## [3162,] "Cardiobacterium"                
## [3163,] "Porphyromonas"                  
## [3164,] "Prevotella_7"                   
## [3165,] "Stomatobaculum"                 
## [3166,] "Aggregatibacter"                
## [3167,] "Alloprevotella"                 
## [3168,] "Treponema_2"                    
## [3169,] "Treponema_2"                    
## [3170,] "Leptotrichia"                   
## [3171,] "Campylobacter"                  
## [3172,] NA                               
## [3173,] "Actinomyces"                    
## [3174,] NA                               
## [3175,] NA                               
## [3176,] NA                               
## [3177,] "Alloprevotella"                 
## [3178,] NA                               
## [3179,] NA                               
## [3180,] NA                               
## [3181,] "Capnocytophaga"                 
## [3182,] "Treponema_2"                    
## [3183,] "Leptotrichia"                   
## [3184,] "Prevotella_2"                   
## [3185,] "Fusobacterium"                  
## [3186,] "Blvii28_wastewater-sludge_group"
## [3187,] "Neisseria"                      
## [3188,] NA                               
## [3189,] "Capnocytophaga"                 
## [3190,] "Prevotella_7"                   
## [3191,] "Treponema_2"                    
## [3192,] "Cardiobacterium"                
## [3193,] "Prevotella_7"                   
## [3194,] NA                               
## [3195,] NA                               
## [3196,] NA                               
## [3197,] "Prevotella_7"                   
## [3198,] "Prevotella_7"                   
## [3199,] "Prevotella"                     
## [3200,] NA                               
## [3201,] NA                               
## [3202,] "Anaerococcus"                   
## [3203,] NA                               
## [3204,] NA                               
## [3205,] NA                               
## [3206,] "Prevotella"                     
## [3207,] "Capnocytophaga"                 
## [3208,] "Prevotellaceae_YAB2003_group"   
## [3209,] NA                               
## [3210,] "Alloprevotella"                 
## [3211,] "Corynebacterium"                
## [3212,] "Prevotella"                     
## [3213,] "Butyrivibrio_2"                 
## [3214,] NA                               
## [3215,] "Prevotella_7"                   
## [3216,] "Prevotella_7"                   
## [3217,] "Leptotrichia"                   
## [3218,] "Prevotellaceae_YAB2003_group"   
## [3219,] "Bacteroides"                    
## [3220,] "Anaeroglobus"                   
## [3221,] NA                               
## [3222,] "Johnsonella"                    
## [3223,] "Granulicatella"                 
## [3224,] NA                               
## [3225,] "Faecalibacterium"               
## [3226,] "Actinomyces"                    
## [3227,] NA                               
## [3228,] NA                               
## [3229,] "Selenomonas_3"                  
## [3230,] "Leptotrichia"                   
## [3231,] NA                               
## [3232,] "Prevotella_7"                   
## [3233,] "Leptotrichia"                   
## [3234,] "Necropsobacter"                 
## [3235,] "Prevotella_2"                   
## [3236,] NA                               
## [3237,] NA                               
## [3238,] "Selenomonas"                    
## [3239,] NA                               
## [3240,] "Prevotella"                     
## [3241,] "Prevotella_7"                   
## [3242,] "Selenomonas"                    
## [3243,] NA                               
## [3244,] "Lachnoanaerobaculum"            
## [3245,] "Rothia"                         
## [3246,] "Shuttleworthia"                 
## [3247,] NA                               
## [3248,] "Collinsella"                    
## [3249,] "Prevotella_7"                   
## [3250,] NA                               
## [3251,] "Atopobium"                      
## [3252,] NA                               
## [3253,] "Prevotella"                     
## [3254,] NA                               
## [3255,] "Lautropia"                      
## [3256,] "Porphyromonas"                  
## [3257,] NA                               
## [3258,] "Leptotrichia"                   
## [3259,] "Anthococcus"                    
## [3260,] "Campylobacter"                  
## [3261,] "Campylobacter"                  
## [3262,] NA                               
## [3263,] "Candidatus_Saccharimonas"       
## [3264,] NA                               
## [3265,] NA                               
## [3266,] NA                               
## [3267,] "Alloprevotella"                 
## [3268,] NA                               
## [3269,] "Alloprevotella"                 
## [3270,] NA                               
## [3271,] "Alloprevotella"                 
## [3272,] "Alloprevotella"                 
## [3273,] "Prevotella"                     
## [3274,] NA                               
## [3275,] "Prevotella"                     
## [3276,] "Actinomyces"                    
## [3277,] NA                               
## [3278,] NA                               
## [3279,] "Treponema_2"                    
## [3280,] "Prevotella_7"                   
## [3281,] "Selenomonas_3"                  
## [3282,] "Cardiobacterium"                
## [3283,] "Alloprevotella"                 
## [3284,] "Prevotella_2"                   
## [3285,] NA                               
## [3286,] "Necropsobacter"                 
## [3287,] "Cricetibacter"                  
## [3288,] "Treponema_2"                    
## [3289,] "Lachnoanaerobaculum"            
## [3290,] "Treponema_2"                    
## [3291,] "Prevotella_6"                   
## [3292,] "Necropsobacter"                 
## [3293,] NA                               
## [3294,] "Prevotella_7"                   
## [3295,] "Prevotella"                     
## [3296,] "Prevotella"                     
## [3297,] "Actinomyces"                    
## [3298,] "Prevotella"                     
## [3299,] NA                               
## [3300,] "Prevotella_7"                   
## [3301,] "Alloprevotella"                 
## [3302,] "Bergeyella"                     
## [3303,] "Tannerella"                     
## [3304,] "Kingella"                       
## [3305,] "Kingella"                       
## [3306,] "Prevotella_2"                   
## [3307,] "Prevotella_7"                   
## [3308,] NA                               
## [3309,] NA                               
## [3310,] "Prevotellaceae_YAB2003_group"   
## [3311,] "Prevotella"                     
## [3312,] "Leptotrichia"                   
## [3313,] "Capnocytophaga"                 
## [3314,] "Treponema_2"                    
## [3315,] NA                               
## [3316,] NA                               
## [3317,] "Catonella"                      
## [3318,] "Leptotrichia"                   
## [3319,] "Bergeriella"                    
## [3320,] "Alloprevotella"                 
## [3321,] "Prevotella"                     
## [3322,] "Leptotrichia"                   
## [3323,] "Prevotella_7"                   
## [3324,] "Prevotella"                     
## [3325,] "Prevotella"                     
## [3326,] NA                               
## [3327,] "Corynebacterium"                
## [3328,] "Bergeyella"                     
## [3329,] "Prevotellaceae_YAB2003_group"   
## [3330,] "Treponema_2"                    
## [3331,] "Porphyromonas"                  
## [3332,] "Prevotella"                     
## [3333,] NA                               
## [3334,] NA                               
## [3335,] "Gemella"                        
## [3336,] "Cardiobacterium"                
## [3337,] "Prevotella_7"                   
## [3338,] "Leptotrichia"                   
## [3339,] "Candidatus_Saccharimonas"       
## [3340,] NA                               
## [3341,] NA                               
## [3342,] NA                               
## [3343,] NA                               
## [3344,] NA                               
## [3345,] NA                               
## [3346,] NA                               
## [3347,] NA                               
## [3348,] "Bergeriella"                    
## [3349,] NA                               
## [3350,] "Leptotrichia"                   
## [3351,] NA                               
## [3352,] "Bergeriella"                    
## [3353,] "Prevotella"                     
## [3354,] NA                               
## [3355,] NA                               
## [3356,] "Veillonella"                    
## [3357,] "Atopobium"                      
## [3358,] NA                               
## [3359,] "Treponema_2"                    
## [3360,] "Veillonella"                    
## [3361,] NA                               
## [3362,] "Prevotella_2"                   
## [3363,] NA                               
## [3364,] "Peptoniphilus"                  
## [3365,] "Prevotella"                     
## [3366,] "Prevotella_7"                   
## [3367,] "Necropsobacter"                 
## [3368,] "Actinomyces"                    
## [3369,] "Capnocytophaga"                 
## [3370,] "Veillonella"                    
## [3371,] "Capnocytophaga"                 
## [3372,] "Fusobacterium"                  
## [3373,] "Leptotrichia"                   
## [3374,] "Treponema_2"                    
## [3375,] "Prevotella"                     
## [3376,] "Capnocytophaga"                 
## [3377,] "Prevotella_7"                   
## [3378,] "Leptotrichia"                   
## [3379,] "Rikenellaceae_RC9_gut_group"    
## [3380,] "Capnocytophaga"                 
## [3381,] "Capnocytophaga"                 
## [3382,] "Prevotella_7"                   
## [3383,] "Lachnoanaerobaculum"            
## [3384,] "Prevotella_7"                   
## [3385,] "Treponema_2"                    
## [3386,] "Porphyromonas"                  
## [3387,] "Streptococcus"                  
## [3388,] "Peptococcus"                    
## [3389,] "Prevotella_7"                   
## [3390,] "Prevotella_2"                   
## [3391,] "Granulicatella"                 
## [3392,] "Capnocytophaga"                 
## [3393,] NA                               
## [3394,] NA                               
## [3395,] "Leptotrichia"                   
## [3396,] "Rothia"                         
## [3397,] "Porphyromonas"                  
## [3398,] NA                               
## [3399,] NA                               
## [3400,] "Porphyromonas"                  
## [3401,] "Enterobacter"                   
## [3402,] NA                               
## [3403,] "Parvimonas"                     
## [3404,] "Capnocytophaga"                 
## [3405,] "Prevotella_7"                   
## [3406,] "Alloprevotella"                 
## [3407,] "Prevotella"                     
## [3408,] "Abiotrophia"                    
## [3409,] "Capnocytophaga"                 
## [3410,] "Cricetibacter"                  
## [3411,] "Atopobium"                      
## [3412,] "Alloprevotella"                 
## [3413,] "Treponema_2"                    
## [3414,] "Prevotella"                     
## [3415,] "Prevotella"                     
## [3416,] "Prevotella_7"                   
## [3417,] "Leptotrichia"                   
## [3418,] "Porphyromonas"                  
## [3419,] "Veillonella"                    
## [3420,] NA                               
## [3421,] NA                               
## [3422,] "Oribacterium"                   
## [3423,] NA                               
## [3424,] "Prevotella"                     
## [3425,] "Necropsobacter"                 
## [3426,] "Prevotellaceae_YAB2003_group"   
## [3427,] "Alloprevotella"                 
## [3428,] "Capnocytophaga"                 
## [3429,] NA                               
## [3430,] NA                               
## [3431,] NA                               
## [3432,] "Prevotella"                     
## [3433,] "Soonwooa"                       
## [3434,] "Prevotellaceae_YAB2003_group"   
## [3435,] NA                               
## [3436,] "Pantoea"                        
## [3437,] NA                               
## [3438,] NA                               
## [3439,] "Pseudomonas"                    
## [3440,] "Abiotrophia"                    
## [3441,] "Capnocytophaga"                 
## [3442,] NA                               
## [3443,] "Peptostreptococcus"             
## [3444,] NA                               
## [3445,] "Prevotella"                     
## [3446,] "Stomatobaculum"                 
## [3447,] NA                               
## [3448,] "Prevotella"                     
## [3449,] "Prevotella_2"                   
## [3450,] NA                               
## [3451,] "Prevotella_7"                   
## [3452,] "Leptotrichia"                   
## [3453,] "Prevotella_7"                   
## [3454,] "Alloprevotella"                 
## [3455,] "Selenomonas_3"                  
## [3456,] "Porphyromonas"                  
## [3457,] NA                               
## [3458,] NA                               
## [3459,] "Aggregatibacter"                
## [3460,] NA                               
## [3461,] "Selenomonas_4"                  
## [3462,] "Simonsiella"                    
## [3463,] "Johnsonella"                    
## [3464,] "Prevotella_2"                   
## [3465,] "Porphyromonas"                  
## [3466,] NA                               
## [3467,] "Centipeda"                      
## [3468,] NA                               
## [3469,] "Prevotella"                     
## [3470,] "Alloprevotella"                 
## [3471,] NA                               
## [3472,] NA                               
## [3473,] NA                               
## [3474,] "Actinomyces"                    
## [3475,] NA                               
## [3476,] "Prevotellaceae_YAB2003_group"   
## [3477,] "Porphyromonas"                  
## [3478,] NA                               
## [3479,] NA                               
## [3480,] "Selenomonas_3"                  
## [3481,] "Necropsobacter"                 
## [3482,] "Capnocytophaga"                 
## [3483,] NA                               
## [3484,] "Howardella"                     
## [3485,] "Prevotella_7"                   
## [3486,] "Alloprevotella"                 
## [3487,] "Capnocytophaga"                 
## [3488,] "Selenomonas_3"                  
## [3489,] NA                               
## [3490,] NA                               
## [3491,] "Prevotella_7"                   
## [3492,] "Selenomonas_3"                  
## [3493,] "Selenomonas_4"                  
## [3494,] NA                               
## [3495,] NA                               
## [3496,] NA                               
## [3497,] "Prevotella_7"                   
## [3498,] NA                               
## [3499,] "Anthococcus"                    
## [3500,] "Prevotella_6"                   
## [3501,] NA                               
## [3502,] NA                               
## [3503,] "Aggregatibacter"                
## [3504,] NA                               
## [3505,] "F0058"                          
## [3506,] "Alloprevotella"                 
## [3507,] NA                               
## [3508,] NA                               
## [3509,] "Leptotrichia"                   
## [3510,] NA                               
## [3511,] "Capnocytophaga"                 
## [3512,] "Actinobacillus"                 
## [3513,] "Peptococcus"                    
## [3514,] "Prevotella_6"                   
## [3515,] NA                               
## [3516,] "Selenomonas_3"                  
## [3517,] NA                               
## [3518,] NA                               
## [3519,] "Selenomonas_3"                  
## [3520,] "Pseudomonas"                    
## [3521,] "Necropsobacter"                 
## [3522,] NA                               
## [3523,] NA                               
## [3524,] NA                               
## [3525,] "Prevotella"                     
## [3526,] NA                               
## [3527,] NA                               
## [3528,] "Neisseria"                      
## [3529,] "Porphyromonas"                  
## [3530,] "Anthococcus"                    
## [3531,] "Prevotella"                     
## [3532,] "Prevotella"                     
## [3533,] NA                               
## [3534,] NA                               
## [3535,] NA                               
## [3536,] NA                               
## [3537,] NA                               
## [3538,] "Selenomonas"                    
## [3539,] "Leptotrichia"                   
## [3540,] NA                               
## [3541,] "Prevotella_7"                   
## [3542,] "Streptobacillus"                
## [3543,] "Leptotrichia"                   
## [3544,] "Neisseria"                      
## [3545,] "Leptotrichia"                   
## [3546,] "Parvimonas"                     
## [3547,] NA                               
## [3548,] "Tannerella"                     
## [3549,] NA                               
## [3550,] NA                               
## [3551,] NA                               
## [3552,] NA                               
## [3553,] "Neisseria"                      
## [3554,] "Cricetibacter"                  
## [3555,] NA                               
## [3556,] NA                               
## [3557,] "Anthococcus"                    
## [3558,] NA                               
## [3559,] NA                               
## [3560,] "Lactobacillus"                  
## [3561,] "Treponema_2"                    
## [3562,] "Johnsonella"                    
## [3563,] NA                               
## [3564,] "Peptostreptococcus"             
## [3565,] "Prevotella"                     
## [3566,] "Selenomonas_3"                  
## [3567,] "Neisseria"                      
## [3568,] NA                               
## [3569,] "Veillonella"                    
## [3570,] "Alloprevotella"                 
## [3571,] "Capnocytophaga"                 
## [3572,] NA                               
## [3573,] NA                               
## [3574,] NA                               
## [3575,] NA                               
## [3576,] "Oribacterium"                   
## [3577,] NA                               
## [3578,] "Lachnoanaerobaculum"            
## [3579,] NA                               
## [3580,] "Howardella"                     
## [3581,] "Prevotella_7"                   
## [3582,] "Treponema_2"                    
## [3583,] "Peptococcus"                    
## [3584,] NA                               
## [3585,] "Prevotella_7"                   
## [3586,] "Bergeriella"                    
## [3587,] "Prevotellaceae_YAB2003_group"   
## [3588,] "Prevotella_2"                   
## [3589,] NA                               
## [3590,] NA                               
## [3591,] NA                               
## [3592,] "Alloprevotella"                 
## [3593,] NA                               
## [3594,] "Porphyromonas"                  
## [3595,] NA                               
## [3596,] NA                               
## [3597,] "Alloprevotella"                 
## [3598,] "Capnocytophaga"                 
## [3599,] NA                               
## [3600,] NA                               
## [3601,] "Prevotella"                     
## [3602,] NA                               
## [3603,] NA                               
## [3604,] "Bergeriella"                    
## [3605,] NA                               
## [3606,] "Bergeyella"                     
## [3607,] NA                               
## [3608,] "Comamonas"                      
## [3609,] "Leptotrichia"                   
## [3610,] "Corynebacterium"                
## [3611,] "Porphyromonas"                  
## [3612,] "Johnsonella"                    
## [3613,] "Leptotrichia"                   
## [3614,] "Dialister"                      
## [3615,] NA                               
## [3616,] NA                               
## [3617,] "Leptotrichia"                   
## [3618,] "Peptostreptococcus"             
## [3619,] "Kingella"                       
## [3620,] "Atopobium"                      
## [3621,] "Rothia"                         
## [3622,] "Tannerella"                     
## [3623,] "Kingella"                       
## [3624,] "Prevotella"                     
## [3625,] "Rikenellaceae_RC9_gut_group"    
## [3626,] "Prevotella_7"                   
## [3627,] NA                               
## [3628,] NA                               
## [3629,] "Tannerella"                     
## [3630,] NA                               
## [3631,] "Porphyromonas"                  
## [3632,] "Prevotella"                     
## [3633,] "Bergeriella"                    
## [3634,] NA                               
## [3635,] NA                               
## [3636,] NA                               
## [3637,] "Bergeyella"                     
## [3638,] "Ruminococcaceae_UCG-014"        
## [3639,] "Veillonella"                    
## [3640,] "Veillonella"                    
## [3641,] "Veillonella"                    
## [3642,] NA                               
## [3643,] "Cardiobacterium"                
## [3644,] NA                               
## [3645,] NA                               
## [3646,] NA                               
## [3647,] NA                               
## [3648,] NA                               
## [3649,] "Shuttleworthia"                 
## [3650,] "Treponema_2"                    
## [3651,] "Ruminococcaceae_UCG-014"        
## [3652,] "Capnocytophaga"                 
## [3653,] NA                               
## [3654,] "Capnocytophaga"                 
## [3655,] "Alloprevotella"                 
## [3656,] "Necropsobacter"                 
## [3657,] NA                               
## [3658,] NA                               
## [3659,] NA                               
## [3660,] NA                               
## [3661,] "Peptostreptococcus"             
## [3662,] NA                               
## [3663,] "Prevotella_7"                   
## [3664,] "Prevotella"                     
## [3665,] "Prevotella_7"                   
## [3666,] "Veillonella"                    
## [3667,] "Rodentibacter"                  
## [3668,] NA                               
## [3669,] NA                               
## [3670,] "Atopobium"                      
## [3671,] NA                               
## [3672,] NA                               
## [3673,] "Capnocytophaga"                 
## [3674,] "Prevotella"                     
## [3675,] NA                               
## [3676,] "Prevotella_7"                   
## [3677,] "Treponema_2"                    
## [3678,] "Prevotella"                     
## [3679,] "Bergeyella"                     
## [3680,] "Prevotellaceae_YAB2003_group"   
## [3681,] "Selenomonas_3"                  
## [3682,] "Alloprevotella"                 
## [3683,] "Kingella"                       
## [3684,] "Haemophilus"                    
## [3685,] "Abiotrophia"                    
## [3686,] NA                               
## [3687,] NA                               
## [3688,] "Prevotella_2"                   
## [3689,] "Prevotella_7"                   
## [3690,] "Agathobacter"                   
## [3691,] NA                               
## [3692,] NA                               
## [3693,] NA                               
## [3694,] "Treponema_2"                    
## [3695,] "Butyrivibrio_2"                 
## [3696,] NA                               
## [3697,] "Faecalibacterium"               
## [3698,] "Cardiobacterium"                
## [3699,] "Leptotrichia"                   
## [3700,] "Prevotella_7"                   
## [3701,] "Selenomonas_3"                  
## [3702,] "Campylobacter"                  
## [3703,] NA                               
## [3704,] "Leptotrichia"                   
## [3705,] "Kingella"                       
## [3706,] NA                               
## [3707,] "Prevotella"                     
## [3708,] "Kingella"                       
## [3709,] NA                               
## [3710,] "Prevotella_7"                   
## [3711,] "Lachnoanaerobaculum"            
## [3712,] "Atopobium"                      
## [3713,] NA                               
## [3714,] NA                               
## [3715,] NA                               
## [3716,] "Comamonas"                      
## [3717,] "Abiotrophia"                    
## [3718,] "Fusobacterium"                  
## [3719,] NA                               
## [3720,] NA                               
## [3721,] "Selenomonas_3"                  
## [3722,] NA                               
## [3723,] "Aggregatibacter"                
## [3724,] "Lachnoanaerobaculum"            
## [3725,] "Shuttleworthia"                 
## [3726,] NA                               
## [3727,] NA                               
## [3728,] "Filifactor"                     
## [3729,] "Lachnoanaerobaculum"            
## [3730,] NA                               
## [3731,] "Lactobacillus"                  
## [3732,] "Treponema_2"                    
## [3733,] NA                               
## [3734,] NA                               
## [3735,] NA                               
## [3736,] "Gemella"                        
## [3737,] "Prevotella"                     
## [3738,] NA                               
## [3739,] "Porphyromonas"                  
## [3740,] "Prevotellaceae_YAB2003_group"   
## [3741,] "Prevotella_6"                   
## [3742,] "Simonsiella"                    
## [3743,] "Campylobacter"                  
## [3744,] NA                               
## [3745,] NA                               
## [3746,] "Capnocytophaga"                 
## [3747,] "Prevotella_7"                   
## [3748,] "Prevotella_7"                   
## [3749,] "Rikenellaceae_RC9_gut_group"    
## [3750,] NA                               
## [3751,] "Stomatobaculum"                 
## [3752,] NA                               
## [3753,] NA                               
## [3754,] "Prevotella_7"                   
## [3755,] "Stomatobaculum"                 
## [3756,] "Lachnoanaerobaculum"            
## [3757,] "Prevotellaceae_YAB2003_group"   
## [3758,] NA                               
## [3759,] "Selenomonas_3"                  
## [3760,] NA                               
## [3761,] "Alloprevotella"                 
## [3762,] NA                               
## [3763,] "Treponema_2"                    
## [3764,] "Bergeyella"                     
## [3765,] "Prevotella_7"                   
## [3766,] "Prevotella_7"                   
## [3767,] "Lautropia"                      
## [3768,] "Dialister"                      
## [3769,] "Corynebacterium"                
## [3770,] "Chelonobacter"                  
## [3771,] "Prevotellaceae_YAB2003_group"   
## [3772,] NA                               
## [3773,] "Peptostreptococcus"             
## [3774,] "Lachnoanaerobaculum"            
## [3775,] "Selenomonas_3"                  
## [3776,] "Alloprevotella"                 
## [3777,] "Prevotella_7"                   
## [3778,] "Alloprevotella"                 
## [3779,] "Prevotella_7"                   
## [3780,] NA                               
## [3781,] "Rikenellaceae_RC9_gut_group"    
## [3782,] "Prevotellaceae_YAB2003_group"   
## [3783,] "Simonsiella"                    
## [3784,] "Alloprevotella"                 
## [3785,] NA                               
## [3786,] "Tannerella"                     
## [3787,] NA                               
## [3788,] NA                               
## [3789,] NA                               
## [3790,] "Selenomonas"                    
## [3791,] "Treponema_2"                    
## [3792,] NA                               
## [3793,] "Capnocytophaga"                 
## [3794,] "Alloprevotella"                 
## [3795,] "Selenomonas"                    
## [3796,] NA                               
## [3797,] NA                               
## [3798,] NA                               
## [3799,] "Prevotella"                     
## [3800,] NA                               
## [3801,] NA                               
## [3802,] "Prevotellaceae_YAB2003_group"   
## [3803,] "Simonsiella"                    
## [3804,] "Lachnoanaerobaculum"            
## [3805,] "Anthococcus"                    
## [3806,] "Candidatus_Saccharimonas"       
## [3807,] "Johnsonella"                    
## [3808,] "Prevotella"                     
## [3809,] NA                               
## [3810,] NA                               
## [3811,] "Megasphaera"                    
## [3812,] "Veillonella"                    
## [3813,] "Anthococcus"                    
## [3814,] NA                               
## [3815,] "Prevotellaceae_YAB2003_group"   
## [3816,] "Bergeyella"                     
## [3817,] "Selenomonas_3"                  
## [3818,] "Selenomonas_3"                  
## [3819,] "Capnocytophaga"                 
## [3820,] "Prevotella"                     
## [3821,] NA                               
## [3822,] "Treponema_2"                    
## [3823,] "Prevotella_7"                   
## [3824,] "Veillonella"                    
## [3825,] "Treponema_2"                    
## [3826,] "Lachnoanaerobaculum"            
## [3827,] NA                               
## [3828,] NA                               
## [3829,] "Lactobacillus"                  
## [3830,] NA                               
## [3831,] "Prevotella_7"                   
## [3832,] "Simonsiella"                    
## [3833,] NA                               
## [3834,] "Rothia"                         
## [3835,] "Porphyromonas"                  
## [3836,] "Alloprevotella"                 
## [3837,] NA                               
## [3838,] "Prevotella_2"                   
## [3839,] "Prevotella_7"                   
## [3840,] "Catonella"                      
## [3841,] NA                               
## [3842,] "Capnocytophaga"                 
## [3843,] NA                               
## [3844,] "Phocaeicola"                    
## [3845,] NA                               
## [3846,] NA                               
## [3847,] "Prevotella_7"                   
## [3848,] "Cuneatibacter"                  
## [3849,] "Treponema_2"                    
## [3850,] "Lachnoanaerobaculum"            
## [3851,] "Bergeriella"                    
## [3852,] "Eikenella"                      
## [3853,] NA                               
## [3854,] NA                               
## [3855,] "Campylobacter"                  
## [3856,] NA                               
## [3857,] "Leptotrichia"                   
## [3858,] "Prevotella_7"                   
## [3859,] "Alloprevotella"                 
## [3860,] "Capnocytophaga"                 
## [3861,] NA                               
## [3862,] "Lachnoanaerobaculum"            
## [3863,] "Johnsonella"                    
## [3864,] NA                               
## [3865,] NA                               
## [3866,] "Lachnoanaerobaculum"            
## [3867,] NA                               
## [3868,] "Prevotella"                     
## [3869,] NA                               
## [3870,] "Porphyromonas"                  
## [3871,] "Haemophilus"                    
## [3872,] "Comamonas"                      
## [3873,] NA                               
## [3874,] "Prevotellaceae_YAB2003_group"   
## [3875,] NA                               
## [3876,] "Necropsobacter"                 
## [3877,] "Prevotella_7"                   
## [3878,] "Campylobacter"                  
## [3879,] "Bergeyella"                     
## [3880,] NA                               
## [3881,] "Treponema_2"                    
## [3882,] "Prevotella_7"                   
## [3883,] NA                               
## [3884,] "Dialister"                      
## [3885,] NA                               
## [3886,] NA                               
## [3887,] "Ezakiella"                      
## [3888,] "Ruminococcaceae_UCG-014"        
## [3889,] NA                               
## [3890,] "Mogibacterium"                  
## [3891,] NA                               
## [3892,] NA                               
## [3893,] "Lachnoanaerobaculum"            
## [3894,] NA                               
## [3895,] "Actinomyces"                    
## [3896,] "Prevotella"                     
## [3897,] NA                               
## [3898,] NA                               
## [3899,] "Cardiobacterium"                
## [3900,] NA                               
## [3901,] "Gemella"                        
## [3902,] "Selenomonas_4"                  
## [3903,] "Prevotella_2"                   
## [3904,] "Porphyromonas"                  
## [3905,] "Treponema_2"                    
## [3906,] "Catonella"                      
## [3907,] "Prevotella"                     
## [3908,] "Necropsobacter"                 
## [3909,] NA                               
## [3910,] "Prevotella_7"                   
## [3911,] "Prevotella_7"                   
## [3912,] NA                               
## [3913,] "Selenomonas_3"                  
## [3914,] "Solobacterium"                  
## [3915,] NA                               
## [3916,] "Gemella"                        
## [3917,] "Prevotella_7"                   
## [3918,] NA                               
## [3919,] "Aggregatibacter"                
## [3920,] NA                               
## [3921,] NA                               
## [3922,] "Prevotella"                     
## [3923,] NA                               
## [3924,] "Duganella"                      
## [3925,] NA                               
## [3926,] "Anaeroglobus"                   
## [3927,] NA                               
## [3928,] NA                               
## [3929,] "Thermus"                        
## [3930,] NA                               
## [3931,] NA                               
## [3932,] NA                               
## [3933,] NA                               
## [3934,] NA                               
## [3935,] "Comamonas"                      
## [3936,] "Treponema_2"                    
## [3937,] "Lactobacillus"                  
## [3938,] "Enhydrobacter"                  
## [3939,] "Lactobacillus"                  
## [3940,] "Anaerococcus"                   
## [3941,] "Prevotella_7"                   
## [3942,] "Alloprevotella"                 
## [3943,] NA                               
## [3944,] "Stomatobaculum"                 
## [3945,] "Prevotella_2"                   
## [3946,] NA                               
## [3947,] NA                               
## [3948,] NA                               
## [3949,] "Porphyromonas"                  
## [3950,] NA                               
## [3951,] "Anthococcus"                    
## [3952,] "Prevotella"                     
## [3953,] NA                               
## [3954,] NA                               
## [3955,] "Campylobacter"                  
## [3956,] NA                               
## [3957,] NA                               
## [3958,] NA                               
## [3959,] "Prevotella"                     
## [3960,] "Veillonella"                    
## [3961,] NA                               
## [3962,] NA                               
## [3963,] "Prevotella_6"                   
## [3964,] NA                               
## [3965,] "Prevotella_7"                   
## [3966,] NA                               
## [3967,] NA                               
## [3968,] "Leptotrichia"                   
## [3969,] NA                               
## [3970,] "Campylobacter"                  
## [3971,] NA                               
## [3972,] "Acholeplasma"                   
## [3973,] "Mycoplasma"                     
## [3974,] "Bergeyella"                     
## [3975,] "Desulfovibrio"                  
## [3976,] NA                               
## [3977,] "Prevotella"                     
## [3978,] NA                               
## [3979,] "Anthococcus"                    
## [3980,] NA                               
## [3981,] NA                               
## [3982,] "Anthococcus"                    
## [3983,] "Prevotella"                     
## [3984,] "Centipeda"                      
## [3985,] "Stomatobaculum"                 
## [3986,] "Alloprevotella"                 
## [3987,] "Prevotella"                     
## [3988,] "Capnocytophaga"                 
## [3989,] "Cryptobacterium"                
## [3990,] "Dialister"                      
## [3991,] "Leptotrichia"                   
## [3992,] NA                               
## [3993,] "Alloprevotella"                 
## [3994,] "Streptococcus"                  
## [3995,] "Treponema_2"                    
## [3996,] "Capnocytophaga"                 
## [3997,] "Peptoanaerobacter"              
## [3998,] NA                               
## [3999,] NA                               
## [4000,] NA                               
## [4001,] "Anthococcus"                    
## [4002,] "Treponema_2"                    
## [4003,] "Gemella"                        
## [4004,] "Capnocytophaga"                 
## [4005,] "Leptotrichia"                   
## [4006,] "Prevotella_7"                   
## [4007,] "Selenomonas_3"                  
## [4008,] "Alloprevotella"                 
## [4009,] "Bergeyella"                     
## [4010,] "Aggregatibacter"                
## [4011,] NA                               
## [4012,] "Tannerella"                     
## [4013,] "Peptostreptococcus"             
## [4014,] "Selenomonas_3"                  
## [4015,] "Actinomyces"                    
## [4016,] NA                               
## [4017,] "Prevotella"                     
## [4018,] NA                               
## [4019,] "Peptococcus"                    
## [4020,] "Rothia"                         
## [4021,] "Prevotella_7"                   
## [4022,] "Actinomyces"                    
## [4023,] "Veillonella"                    
## [4024,] NA                               
## [4025,] "Capnocytophaga"                 
## [4026,] "Prevotella"                     
## [4027,] "Anthococcus"                    
## [4028,] "Selenomonas_3"                  
## [4029,] NA                               
## [4030,] NA                               
## [4031,] "Neisseria"                      
## [4032,] "Actinomyces"                    
## [4033,] "Treponema_2"                    
## [4034,] "Isobaculum"                     
## [4035,] NA                               
## [4036,] "Pseudoramibacter"               
## [4037,] NA                               
## [4038,] "Veillonella"                    
## [4039,] "Neisseria"                      
## [4040,] NA                               
## [4041,] "Alloprevotella"                 
## [4042,] NA                               
## [4043,] "Prevotella_2"                   
## [4044,] "Capnocytophaga"                 
## [4045,] "Lachnoanaerobaculum"            
## [4046,] NA                               
## [4047,] "Prevotella_7"                   
## [4048,] NA                               
## [4049,] NA                               
## [4050,] "Bergeyella"                     
## [4051,] "Prevotella_7"                   
## [4052,] "Prevotella_7"                   
## [4053,] NA                               
## [4054,] NA                               
## [4055,] NA                               
## [4056,] NA                               
## [4057,] NA                               
## [4058,] NA                               
## [4059,] "Streptococcus"                  
## [4060,] NA                               
## [4061,] NA                               
## [4062,] "Actinomyces"                    
## [4063,] "Candidatus_Saccharimonas"       
## [4064,] NA                               
## [4065,] NA                               
## [4066,] "Tannerella"                     
## [4067,] "Stomatobaculum"                 
## [4068,] NA                               
## [4069,] NA                               
## [4070,] "Corynebacterium"                
## [4071,] "Anthococcus"                    
## [4072,] NA                               
## [4073,] "Selenomonas_4"                  
## [4074,] "Selenomonas_3"                  
## [4075,] NA                               
## [4076,] "Lachnoanaerobaculum"            
## [4077,] "Capnocytophaga"                 
## [4078,] NA                               
## [4079,] "Alloprevotella"                 
## [4080,] "Capnocytophaga"                 
## [4081,] NA                               
## [4082,] "Anaeroglobus"                   
## [4083,] NA                               
## [4084,] NA                               
## [4085,] NA                               
## [4086,] NA                               
## [4087,] NA                               
## [4088,] "Lautropia"                      
## [4089,] "Corynebacterium"                
## [4090,] NA                               
## [4091,] NA                               
## [4092,] NA                               
## [4093,] "Peptostreptococcus"             
## [4094,] "Stomatobaculum"                 
## [4095,] NA                               
## [4096,] "Ruminococcaceae_UCG-014"        
## [4097,] "Candidatus_Saccharimonas"       
## [4098,] NA                               
## [4099,] "Alloprevotella"                 
## [4100,] "Peptococcus"                    
## [4101,] NA                               
## [4102,] "Chryseobacterium"               
## [4103,] NA                               
## [4104,] "Mogibacterium"                  
## [4105,] "Alloprevotella"                 
## [4106,] "Prevotella_7"                   
## [4107,] NA                               
## [4108,] NA                               
## [4109,] "Porphyromonas"                  
## [4110,] "Anaeroglobus"                   
## [4111,] "Granulicatella"                 
## [4112,] "Veillonella"                    
## [4113,] NA                               
## [4114,] NA                               
## [4115,] NA                               
## [4116,] "Filifactor"                     
## [4117,] "Atopobium"                      
## [4118,] "Rikenellaceae_RC9_gut_group"    
## [4119,] "Atopobium"                      
## [4120,] NA                               
## [4121,] NA                               
## [4122,] NA                               
## [4123,] NA                               
## [4124,] NA                               
## [4125,] NA                               
## [4126,] NA                               
## [4127,] "Anthococcus"                    
## [4128,] "Necropsobacter"                 
## [4129,] "Prevotella"                     
## [4130,] NA                               
## [4131,] NA                               
## [4132,] NA                               
## [4133,] "Capnocytophaga"                 
## [4134,] "Prevotella"                     
## [4135,] NA                               
## [4136,] NA                               
## [4137,] NA                               
## [4138,] NA                               
## [4139,] "Cardiobacterium"                
## [4140,] "Ruminococcaceae_UCG-014"        
## [4141,] "Bergeriella"                    
## [4142,] "Parvimonas"                     
## [4143,] "Butyrivibrio_2"                 
## [4144,] "Prevotella_7"                   
## [4145,] NA                               
## [4146,] "Prevotella_2"                   
## [4147,] NA                               
## [4148,] "Veillonella"                    
## [4149,] "Prevotella"                     
## [4150,] "Filifactor"                     
## [4151,] NA                               
## [4152,] "Treponema_2"                    
## [4153,] NA                               
## [4154,] "Dialister"                      
## [4155,] "Prevotella_2"                   
## [4156,] "Corynebacterium"                
## [4157,] "Veillonella"                    
## [4158,] NA                               
## [4159,] NA                               
## [4160,] "Prevotella_7"                   
## [4161,] NA                               
## [4162,] NA                               
## [4163,] NA                               
## [4164,] NA                               
## [4165,] NA                               
## [4166,] "Necropsobacter"                 
## [4167,] "Gemella"                        
## [4168,] NA                               
## [4169,] NA                               
## [4170,] "Anaeroglobus"                   
## [4171,] "Prevotella_2"                   
## [4172,] NA                               
## [4173,] "Alloprevotella"                 
## [4174,] "Johnsonella"                    
## [4175,] "Treponema_2"                    
## [4176,] "Comamonas"                      
## [4177,] NA                               
## [4178,] "Neisseria"                      
## [4179,] "Prevotellaceae_YAB2003_group"   
## [4180,] NA                               
## [4181,] NA                               
## [4182,] "Bergeriella"                    
## [4183,] "Mycoplasma"                     
## [4184,] "Bacteroides"                    
## [4185,] NA                               
## [4186,] NA                               
## [4187,] NA                               
## [4188,] "Selenomonas"                    
## [4189,] NA                               
## [4190,] "Treponema_2"                    
## [4191,] NA                               
## [4192,] "Bergeriella"                    
## [4193,] "Gemella"                        
## [4194,] NA                               
## [4195,] "Capnocytophaga"                 
## [4196,] NA                               
## [4197,] "Parvimonas"                     
## [4198,] "Veillonella"                    
## [4199,] NA                               
## [4200,] "Acinetobacter"                  
## [4201,] "Anthococcus"                    
## [4202,] "Fusobacterium"                  
## [4203,] "Stomatobaculum"                 
## [4204,] "Prevotella_7"                   
## [4205,] "Veillonella"                    
## [4206,] NA                               
## [4207,] "Lawsonella"                     
## [4208,] "Lachnoanaerobaculum"            
## [4209,] "Streptobacillus"                
## [4210,] "Treponema_2"                    
## [4211,] NA                               
## [4212,] NA                               
## [4213,] NA                               
## [4214,] "Prevotella_7"                   
## [4215,] "Prevotella_2"                   
## [4216,] NA                               
## [4217,] "Candidatus_Saccharimonas"       
## [4218,] "Prevotella_7"                   
## [4219,] NA                               
## [4220,] NA                               
## [4221,] NA                               
## [4222,] NA                               
## [4223,] "Soonwooa"                       
## [4224,] "Capnocytophaga"                 
## [4225,] NA                               
## [4226,] "Leptotrichia"                   
## [4227,] "Prevotella_7"                   
## [4228,] NA                               
## [4229,] "Necropsobacter"                 
## [4230,] NA                               
## [4231,] "Prevotella_7"                   
## [4232,] "Actinobacillus"                 
## [4233,] NA                               
## [4234,] "Prevotella_7"                   
## [4235,] NA                               
## [4236,] "Candidatus_Saccharimonas"       
## [4237,] NA                               
## [4238,] "Treponema_2"                    
## [4239,] "Corynebacterium"                
## [4240,] "Bergeriella"                    
## [4241,] NA                               
## [4242,] NA                               
## [4243,] NA                               
## [4244,] NA                               
## [4245,] "Porphyromonas"                  
## [4246,] "Actinomyces"                    
## [4247,] "Treponema_2"                    
## [4248,] "Gemella"                        
## [4249,] NA                               
## [4250,] "Treponema_2"                    
## [4251,] "Finegoldia"                     
## [4252,] NA                               
## [4253,] "Desulfobulbus"                  
## [4254,] NA                               
## [4255,] NA                               
## [4256,] "Prevotella_7"                   
## [4257,] NA                               
## [4258,] NA                               
## [4259,] "Capnocytophaga"                 
## [4260,] "Bergeyella"                     
## [4261,] NA                               
## [4262,] NA                               
## [4263,] NA                               
## [4264,] "Prevotella"                     
## [4265,] NA                               
## [4266,] NA                               
## [4267,] NA                               
## [4268,] "Atopobium"                      
## [4269,] "Prevotella"                     
## [4270,] "Anthococcus"                    
## [4271,] "Peptostreptococcus"             
## [4272,] "Lachnoanaerobaculum"            
## [4273,] NA                               
## [4274,] NA                               
## [4275,] "Prevotella_2"                   
## [4276,] "Catonella"                      
## [4277,] NA                               
## [4278,] "Anthococcus"                    
## [4279,] "Anthococcus"                    
## [4280,] "Porphyromonas"                  
## [4281,] NA                               
## [4282,] NA                               
## [4283,] NA                               
## [4284,] NA                               
## [4285,] "Solobacterium"                  
## [4286,] NA                               
## [4287,] "Bacteroides"                    
## [4288,] NA                               
## [4289,] NA                               
## [4290,] "Peptostreptococcus"             
## [4291,] NA                               
## [4292,] NA                               
## [4293,] "Cricetibacter"                  
## [4294,] NA                               
## [4295,] "Atopobium"                      
## [4296,] "Rikenellaceae_RC9_gut_group"    
## [4297,] NA                               
## [4298,] NA                               
## [4299,] NA                               
## [4300,] NA                               
## [4301,] NA                               
## [4302,] "Treponema_2"                    
## [4303,] "Chryseobacterium"               
## [4304,] "Selenomonas_3"                  
## [4305,] NA                               
## [4306,] NA                               
## [4307,] "Treponema_2"                    
## [4308,] NA                               
## [4309,] NA                               
## [4310,] "Veillonella"                    
## [4311,] "Bergeriella"                    
## [4312,] "Peptostreptococcus"             
## [4313,] NA                               
## [4314,] NA                               
## [4315,] NA                               
## [4316,] NA                               
## [4317,] NA                               
## [4318,] NA                               
## [4319,] "Treponema_2"                    
## [4320,] "Filifactor"                     
## [4321,] "Ruminococcaceae_UCG-014"        
## [4322,] NA                               
## [4323,] "Subdoligranulum"                
## [4324,] NA                               
## [4325,] NA                               
## [4326,] "Cardiobacterium"                
## [4327,] NA                               
## [4328,] "Erysipelotrichaceae_UCG-006"    
## [4329,] "Bifidobacterium"                
## [4330,] NA                               
## [4331,] "Veillonella"                    
## [4332,] NA                               
## [4333,] "Bergeriella"                    
## [4334,] NA                               
## [4335,] "Anthococcus"                    
## [4336,] "Wolinella"                      
## [4337,] "Treponema_2"                    
## [4338,] NA                               
## [4339,] "Sphingobium"                    
## [4340,] "Xanthomonas"                    
## [4341,] NA                               
## [4342,] "Gemella"                        
## [4343,] NA                               
## [4344,] "Leptotrichia"                   
## [4345,] "Rikenellaceae_RC9_gut_group"    
## [4346,] NA                               
## [4347,] "Parvimonas"                     
## [4348,] "Prevotella"                     
## [4349,] NA                               
## [4350,] "Tannerella"                     
## [4351,] NA                               
## [4352,] "Defluviitaleaceae_UCG-011"      
## [4353,] "Prevotella_2"                   
## [4354,] "Gemella"                        
## [4355,] "Alloprevotella"                 
## [4356,] NA                               
## [4357,] NA                               
## [4358,] "Fusobacterium"                  
## [4359,] NA                               
## [4360,] NA                               
## [4361,] NA                               
## [4362,] "Leptotrichia"                   
## [4363,] "Bergeyella"                     
## [4364,] "Peptostreptococcus"             
## [4365,] NA                               
## [4366,] NA                               
## [4367,] NA                               
## [4368,] NA                               
## [4369,] "Parvimonas"                     
## [4370,] NA                               
## [4371,] "Lelliottia"                     
## [4372,] "Cardiobacterium"                
## [4373,] NA                               
## [4374,] "Leptotrichia"                   
## [4375,] "Dialister"                      
## [4376,] "Treponema_2"                    
## [4377,] "Prevotella"                     
## [4378,] NA                               
## [4379,] "Lachnoanaerobaculum"            
## [4380,] NA                               
## [4381,] NA                               
## [4382,] "Prevotella"                     
## [4383,] NA                               
## [4384,] "Cryptobacterium"                
## [4385,] NA                               
## [4386,] "Rikenellaceae_RC9_gut_group"    
## [4387,] "Veillonella"                    
## [4388,] "Lachnoanaerobaculum"            
## [4389,] NA                               
## [4390,] "Selenomonas_4"                  
## [4391,] NA                               
## [4392,] NA                               
## [4393,] NA                               
## [4394,] "Selenomonas_4"                  
## [4395,] "Solobacterium"                  
## [4396,] "Centipeda"                      
## [4397,] "Centipeda"                      
## [4398,] NA                               
## [4399,] NA                               
## [4400,] NA                               
## [4401,] "Anaeroglobus"                   
## [4402,] NA                               
## [4403,] "Kingella"                       
## [4404,] "Candidatus_Saccharimonas"       
## [4405,] "Atopobium"                      
## [4406,] "Prevotella"                     
## [4407,] NA                               
## [4408,] "Methylobacterium"               
## [4409,] NA                               
## [4410,] NA                               
## [4411,] "Family_XIII_UCG-001"            
## [4412,] "Lactobacillus"                  
## [4413,] "Atopobium"                      
## [4414,] "Treponema_2"                    
## [4415,] NA                               
## [4416,] NA                               
## [4417,] "Fretibacterium"                 
## [4418,] NA                               
## [4419,] NA                               
## [4420,] "Atopobium"                      
## [4421,] NA                               
## [4422,] "Mycoplasma"                     
## [4423,] "Cardiobacterium"                
## [4424,] "Staphylococcus"                 
## [4425,] NA                               
## [4426,] NA                               
## [4427,] "Johnsonella"                    
## [4428,] NA                               
## [4429,] NA                               
## [4430,] "Atopobium"                      
## [4431,] NA                               
## [4432,] "Prevotellaceae_YAB2003_group"   
## [4433,] NA                               
## [4434,] "Peptococcus"                    
## [4435,] NA                               
## [4436,] NA                               
## [4437,] NA                               
## [4438,] "Candidatus_Saccharimonas"       
## [4439,] NA                               
## [4440,] NA                               
## [4441,] "Megasphaera"                    
## [4442,] "Actinobacillus"                 
## [4443,] "Bulleidia"                      
## [4444,] "Suttonella"                     
## [4445,] "Capnocytophaga"                 
## [4446,] "Treponema_2"                    
## [4447,] "Capnocytophaga"                 
## [4448,] "Gemella"                        
## [4449,] "Porphyromonas"                  
## [4450,] "Prevotella"                     
## [4451,] NA                               
## [4452,] "Prevotella"                     
## [4453,] "Actinomyces"                    
## [4454,] "Chelonobacter"                  
## [4455,] NA                               
## [4456,] NA                               
## [4457,] "Bergeriella"                    
## [4458,] "Solobacterium"                  
## [4459,] "Alloprevotella"                 
## [4460,] NA                               
## [4461,] NA                               
## [4462,] NA                               
## [4463,] NA                               
## [4464,] NA                               
## [4465,] NA                               
## [4466,] "Atopobium"                      
## [4467,] "Lautropia"                      
## [4468,] NA                               
## [4469,] NA                               
## [4470,] "Prevotella_7"                   
## [4471,] "Anaerostipes"                   
## [4472,] "Stomatobaculum"                 
## [4473,] "Prevotella"                     
## [4474,] "Alloprevotella"                 
## [4475,] "Johnsonella"                    
## [4476,] "Porphyromonas"                  
## [4477,] "Selenomonas_4"                  
## [4478,] "Prevotella_7"                   
## [4479,] "Tannerella"                     
## [4480,] "Anthococcus"                    
## [4481,] NA                               
## [4482,] "Peptococcus"                    
## [4483,] NA                               
## [4484,] "Alloprevotella"                 
## [4485,] "Selenomonas"                    
## [4486,] "Erysipelotrichaceae_UCG-008"    
## [4487,] NA                               
## [4488,] NA                               
## [4489,] "Treponema_2"                    
## [4490,] "Catonella"                      
## [4491,] "Peptoanaerobacter"              
## [4492,] "Delftia"                        
## [4493,] NA                               
## [4494,] NA                               
## [4495,] "Prevotella"                     
## [4496,] "Pseudomonas"                    
## [4497,] "Afipia"                         
## [4498,] "Gemella"                        
## [4499,] "Prevotella_7"                   
## [4500,] NA                               
## [4501,] NA                               
## [4502,] "Anthococcus"                    
## [4503,] "Bergeriella"                    
## [4504,] "Johnsonella"                    
## [4505,] NA                               
## [4506,] "Prevotella_7"                   
## [4507,] "Prevotella"                     
## [4508,] "Lachnoanaerobaculum"            
## [4509,] NA                               
## [4510,] "Necropsobacter"                 
## [4511,] "Prevotella_7"                   
## [4512,] NA                               
## [4513,] NA                               
## [4514,] NA                               
## [4515,] NA                               
## [4516,] "Candidatus_Saccharimonas"       
## [4517,] "Leptotrichia"                   
## [4518,] "Dialister"                      
## [4519,] NA                               
## [4520,] NA                               
## [4521,] "Bergeriella"                    
## [4522,] "Prevotella_7"                   
## [4523,] "Prevotella_7"                   
## [4524,] "Brachymonas"                    
## [4525,] "W5053"                          
## [4526,] "Capnocytophaga"                 
## [4527,] "Candidatus_Saccharimonas"       
## [4528,] "Alloprevotella"                 
## [4529,] "Centipeda"                      
## [4530,] NA                               
## [4531,] "Porphyromonas"                  
## [4532,] NA                               
## [4533,] NA                               
## [4534,] NA                               
## [4535,] "Porphyromonas"                  
## [4536,] NA                               
## [4537,] NA                               
## [4538,] "Anthococcus"                    
## [4539,] "Bacteroides"                    
## [4540,] NA                               
## [4541,] NA                               
## [4542,] "Stomatobaculum"                 
## [4543,] NA                               
## [4544,] NA                               
## [4545,] NA                               
## [4546,] "Treponema_2"                    
## [4547,] NA                               
## [4548,] "Rikenellaceae_RC9_gut_group"    
## [4549,] "Centipeda"                      
## [4550,] "Streptococcus"                  
## [4551,] "Leptotrichia"                   
## [4552,] NA                               
## [4553,] "Treponema_2"                    
## [4554,] "Prevotella_2"                   
## [4555,] NA
```

```
#inspect the taxonomic assignments
taxa.print <- tax
rownames(taxa.print) <- NULL
head(taxa.print)
```

```
##      Kingdom    Phylum           Class                 Order            
## [1,] "Bacteria" "Firmicutes"     "Negativicutes"       "Selenomonadales"
## [2,] "Bacteria" "Firmicutes"     "Bacilli"             "Lactobacillales"
## [3,] "Bacteria" "Bacteroidetes"  "Bacteroidia"         "Bacteroidales"  
## [4,] "Bacteria" "Proteobacteria" "Gammaproteobacteria" "Pasteurellales" 
## [5,] "Bacteria" "Firmicutes"     "Negativicutes"       "Selenomonadales"
## [6,] "Bacteria" "Firmicutes"     "Negativicutes"       "Selenomonadales"
##      Family             Genus          
## [1,] "Veillonellaceae"  "Veillonella"  
## [2,] "Streptococcaceae" "Streptococcus"
## [3,] "Prevotellaceae"   "Prevotella_7" 
## [4,] "Pasteurellaceae"  "Haemophilus"  
## [5,] "Veillonellaceae"  "Veillonella"  
## [6,] "Veillonellaceae"  "Veillonella"
```

```
table(nchar(getSequences(seqtab)))
```

```
## 
##  395 
## 4555
```

```
#make species level assignments
taxa <- addSpecies(tax, "/Users/luizroesch/uclust8/silva_species_assignment_v132.fa.gz",  verbose=TRUE)
```

```
## 330 out of 4555 were assigned to the species level.
## Of which 321 had genera consistent with the input table.
```

```
unname(taxa)
```

```
##         [,1]       [,2]                  [,3]                         
##    [1,] "Bacteria" "Firmicutes"          "Negativicutes"              
##    [2,] "Bacteria" "Firmicutes"          "Bacilli"                    
##    [3,] "Bacteria" "Bacteroidetes"       "Bacteroidia"                
##    [4,] "Bacteria" "Proteobacteria"      "Gammaproteobacteria"        
##    [5,] "Bacteria" "Firmicutes"          "Negativicutes"              
##    [6,] "Bacteria" "Firmicutes"          "Negativicutes"              
##    [7,] "Bacteria" "Firmicutes"          "Negativicutes"              
##    [8,] "Bacteria" "Patescibacteria"     "Saccharimonadia"            
##    [9,] "Bacteria" "Proteobacteria"      "Gammaproteobacteria"        
##   [10,] "Bacteria" "Firmicutes"          "Negativicutes"              
##   [11,] "Bacteria" "Proteobacteria"      "Gammaproteobacteria"        
##   [12,] "Bacteria" "Firmicutes"          "Negativicutes"              
##   [13,] "Bacteria" "Fusobacteria"        "Fusobacteriia"              
##   [14,] "Bacteria" "Firmicutes"          "Negativicutes"              
##   [15,] "Bacteria" "Bacteroidetes"       "Bacteroidia"                
##   [16,] "Bacteria" "Epsilonbacteraeota"  "Campylobacteria"            
##   [17,] "Bacteria" "Proteobacteria"      "Gammaproteobacteria"        
##   [18,] "Bacteria" "Firmicutes"          "Clostridia"                 
##   [19,] "Bacteria" "Firmicutes"          "Negativicutes"              
##   [20,] "Bacteria" "Bacteroidetes"       "Bacteroidia"                
##   [21,] "Bacteria" "Firmicutes"          "Bacilli"                    
##   [22,] "Bacteria" "Bacteroidetes"       "Bacteroidia"                
##   [23,] "Bacteria" "Bacteroidetes"       "Bacteroidia"                
##   [24,] "Bacteria" "Firmicutes"          "Negativicutes"              
##   [25,] "Bacteria" "Proteobacteria"      "Gammaproteobacteria"        
##   [26,] "Bacteria" "Bacteroidetes"       "Bacteroidia"                
##   [27,] "Bacteria" "Bacteroidetes"       "Bacteroidia"                
##   [28,] "Bacteria" "Firmicutes"          "Negativicutes"              
##   [29,] "Bacteria" "Firmicutes"          "Bacilli"                    
##   [30,] "Bacteria" "Firmicutes"          "Bacilli"                    
##   [31,] "Bacteria" "Bacteroidetes"       "Bacteroidia"                
##   [32,] "Bacteria" "Firmicutes"          "Bacilli"                    
##   [33,] "Bacteria" "Bacteroidetes"       "Bacteroidia"                
##   [34,] "Bacteria" "Bacteroidetes"       "Bacteroidia"                
##   [35,] "Bacteria" "Firmicutes"          "Clostridia"                 
##   [36,] "Bacteria" "Firmicutes"          "Negativicutes"              
##   [37,] "Bacteria" "Bacteroidetes"       "Bacteroidia"                
##   [38,] "Bacteria" "Firmicutes"          "Negativicutes"              
##   [39,] "Bacteria" "Firmicutes"          "Clostridia"                 
##   [40,] "Bacteria" "Firmicutes"          "Bacilli"                    
##   [41,] "Bacteria" "Bacteroidetes"       "Bacteroidia"                
##   [42,] "Bacteria" "Firmicutes"          "Bacilli"                    
##   [43,] "Bacteria" "Firmicutes"          "Bacilli"                    
##   [44,] "Bacteria" "Bacteroidetes"       "Bacteroidia"                
##   [45,] "Bacteria" "Proteobacteria"      "Gammaproteobacteria"        
##   [46,] "Bacteria" "Bacteroidetes"       "Bacteroidia"                
##   [47,] "Bacteria" "Firmicutes"          "Negativicutes"              
##   [48,] "Bacteria" "Firmicutes"          "Negativicutes"              
##   [49,] "Bacteria" "Firmicutes"          "Clostridia"                 
##   [50,] "Bacteria" "Fusobacteria"        "Fusobacteriia"              
##   [51,] "Bacteria" "Firmicutes"          "Bacilli"                    
##   [52,] "Bacteria" "Proteobacteria"      "Gammaproteobacteria"        
##   [53,] "Bacteria" "Firmicutes"          "Bacilli"                    
##   [54,] "Bacteria" "Fusobacteria"        "Fusobacteriia"              
##   [55,] "Bacteria" "Bacteroidetes"       "Bacteroidia"                
##   [56,] "Bacteria" "Bacteroidetes"       "Bacteroidia"                
##   [57,] "Bacteria" "Firmicutes"          "Negativicutes"              
##   [58,] "Bacteria" "Firmicutes"          "Clostridia"                 
##   [59,] "Bacteria" "Actinobacteria"      "Coriobacteriia"             
##   [60,] "Bacteria" "Proteobacteria"      "Gammaproteobacteria"        
##   [61,] "Bacteria" "Proteobacteria"      "Gammaproteobacteria"        
##   [62,] "Bacteria" "Bacteroidetes"       "Bacteroidia"                
##   [63,] "Bacteria" "Firmicutes"          "Clostridia"                 
##   [64,] "Bacteria" "Firmicutes"          "Negativicutes"              
##   [65,] "Bacteria" "Proteobacteria"      "Gammaproteobacteria"        
##   [66,] "Bacteria" "Firmicutes"          "Negativicutes"              
##   [67,] "Bacteria" "Patescibacteria"     "Saccharimonadia"            
##   [68,] "Bacteria" "Actinobacteria"      "Actinobacteria"             
##   [69,] "Bacteria" "Proteobacteria"      "Gammaproteobacteria"        
##   [70,] "Bacteria" "Firmicutes"          "Clostridia"                 
##   [71,] "Bacteria" "Bacteroidetes"       "Bacteroidia"                
##   [72,] "Bacteria" "Actinobacteria"      "Actinobacteria"             
##   [73,] "Bacteria" "Firmicutes"          "Clostridia"                 
##   [74,] "Bacteria" "Firmicutes"          "Negativicutes"              
##   [75,] "Bacteria" "Actinobacteria"      "Actinobacteria"             
##   [76,] "Bacteria" "Proteobacteria"      "Gammaproteobacteria"        
##   [77,] "Bacteria" "Firmicutes"          "Clostridia"                 
##   [78,] "Bacteria" "Bacteroidetes"       "Bacteroidia"                
##   [79,] "Bacteria" "Firmicutes"          "Negativicutes"              
##   [80,] "Bacteria" "Firmicutes"          "Clostridia"                 
##   [81,] "Bacteria" "Proteobacteria"      "Gammaproteobacteria"        
##   [82,] "Bacteria" "Firmicutes"          "Negativicutes"              
##   [83,] "Bacteria" "Bacteroidetes"       "Bacteroidia"                
##   [84,] "Bacteria" "Bacteroidetes"       "Bacteroidia"                
##   [85,] "Bacteria" "Firmicutes"          "Negativicutes"              
##   [86,] "Bacteria" "Bacteroidetes"       "Bacteroidia"                
##   [87,] "Bacteria" "Fusobacteria"        "Fusobacteriia"              
##   [88,] "Bacteria" "Firmicutes"          "Clostridia"                 
##   [89,] "Bacteria" "Bacteroidetes"       "Bacteroidia"                
##   [90,] "Bacteria" "Firmicutes"          "Clostridia"                 
##   [91,] "Bacteria" "Actinobacteria"      "Actinobacteria"             
##   [92,] "Bacteria" "Proteobacteria"      "Gammaproteobacteria"        
##   [93,] "Bacteria" "Actinobacteria"      "Actinobacteria"             
##   [94,] "Bacteria" "Firmicutes"          "Bacilli"                    
##   [95,] "Bacteria" "Bacteroidetes"       "Bacteroidia"                
##   [96,] "Bacteria" "Proteobacteria"      "Gammaproteobacteria"        
##   [97,] "Bacteria" "Fusobacteria"        "Fusobacteriia"              
##   [98,] "Bacteria" "Firmicutes"          "Negativicutes"              
##   [99,] "Bacteria" "Fusobacteria"        "Fusobacteriia"              
##  [100,] "Bacteria" "Bacteroidetes"       "Bacteroidia"                
##  [101,] "Bacteria" "Bacteroidetes"       "Bacteroidia"                
##  [102,] "Bacteria" "Proteobacteria"      "Gammaproteobacteria"        
##  [103,] "Bacteria" "Proteobacteria"      "Gammaproteobacteria"        
##  [104,] "Bacteria" "Bacteroidetes"       "Bacteroidia"                
##  [105,] "Bacteria" "Firmicutes"          "Bacilli"                    
##  [106,] "Bacteria" "Firmicutes"          "Bacilli"                    
##  [107,] "Bacteria" "Firmicutes"          "Negativicutes"              
##  [108,] "Bacteria" "Firmicutes"          "Negativicutes"              
##  [109,] "Bacteria" "Actinobacteria"      "Actinobacteria"             
##  [110,] "Bacteria" "Proteobacteria"      "Gammaproteobacteria"        
##  [111,] "Bacteria" "Epsilonbacteraeota"  "Campylobacteria"            
##  [112,] "Bacteria" "Bacteroidetes"       "Bacteroidia"                
##  [113,] "Bacteria" "Proteobacteria"      "Gammaproteobacteria"        
##  [114,] "Bacteria" "Proteobacteria"      "Gammaproteobacteria"        
##  [115,] "Bacteria" "Patescibacteria"     "Saccharimonadia"            
##  [116,] "Bacteria" "Bacteroidetes"       "Bacteroidia"                
##  [117,] "Bacteria" "Proteobacteria"      "Gammaproteobacteria"        
##  [118,] "Bacteria" "Proteobacteria"      "Gammaproteobacteria"        
##  [119,] "Bacteria" "Proteobacteria"      "Gammaproteobacteria"        
##  [120,] "Bacteria" "Proteobacteria"      "Gammaproteobacteria"        
##  [121,] "Bacteria" "Proteobacteria"      "Gammaproteobacteria"        
##  [122,] "Bacteria" "Bacteroidetes"       "Bacteroidia"                
##  [123,] "Bacteria" "Firmicutes"          "Bacilli"                    
##  [124,] "Bacteria" "Bacteroidetes"       "Bacteroidia"                
##  [125,] "Bacteria" "Proteobacteria"      "Gammaproteobacteria"        
##  [126,] "Bacteria" "Bacteroidetes"       "Bacteroidia"                
##  [127,] "Bacteria" "Firmicutes"          "Negativicutes"              
##  [128,] "Bacteria" "Proteobacteria"      "Gammaproteobacteria"        
##  [129,] "Bacteria" "Proteobacteria"      "Gammaproteobacteria"        
##  [130,] "Bacteria" "Proteobacteria"      "Gammaproteobacteria"        
##  [131,] "Bacteria" "Proteobacteria"      "Gammaproteobacteria"        
##  [132,] "Bacteria" "Bacteroidetes"       "Bacteroidia"                
##  [133,] "Bacteria" "Patescibacteria"     "Saccharimonadia"            
##  [134,] "Bacteria" "Bacteroidetes"       "Bacteroidia"                
##  [135,] "Bacteria" "Firmicutes"          "Negativicutes"              
##  [136,] "Bacteria" "Firmicutes"          "Negativicutes"              
##  [137,] "Bacteria" "Bacteroidetes"       "Bacteroidia"                
##  [138,] "Bacteria" "Bacteroidetes"       "Bacteroidia"                
##  [139,] "Bacteria" "Bacteroidetes"       "Bacteroidia"                
##  [140,] "Bacteria" "Firmicutes"          "Bacilli"                    
##  [141,] "Bacteria" "Firmicutes"          "Negativicutes"              
##  [142,] "Bacteria" "Firmicutes"          "Negativicutes"              
##  [143,] "Bacteria" "Fusobacteria"        "Fusobacteriia"              
##  [144,] "Bacteria" "Firmicutes"          "Negativicutes"              
##  [145,] "Bacteria" "Firmicutes"          "Bacilli"                    
##  [146,] "Bacteria" "Bacteroidetes"       "Bacteroidia"                
##  [147,] "Bacteria" "Proteobacteria"      "Gammaproteobacteria"        
##  [148,] "Bacteria" "Firmicutes"          "Negativicutes"              
##  [149,] "Bacteria" "Firmicutes"          "Negativicutes"              
##  [150,] "Bacteria" "Firmicutes"          "Negativicutes"              
##  [151,] "Bacteria" "Proteobacteria"      "Gammaproteobacteria"        
##  [152,] "Bacteria" "Bacteroidetes"       "Bacteroidia"                
##  [153,] "Bacteria" "Firmicutes"          "Negativicutes"              
##  [154,] "Bacteria" "Bacteroidetes"       "Bacteroidia"                
##  [155,] "Bacteria" "Firmicutes"          "Clostridia"                 
##  [156,] "Bacteria" "Proteobacteria"      "Gammaproteobacteria"        
##  [157,] "Bacteria" "Firmicutes"          "Negativicutes"              
##  [158,] "Bacteria" "Bacteroidetes"       "Bacteroidia"                
##  [159,] "Bacteria" "Firmicutes"          "Negativicutes"              
##  [160,] "Bacteria" "Firmicutes"          "Clostridia"                 
##  [161,] "Bacteria" "Firmicutes"          "Bacilli"                    
##  [162,] "Bacteria" "Actinobacteria"      "Actinobacteria"             
##  [163,] "Bacteria" "Bacteroidetes"       "Bacteroidia"                
##  [164,] "Bacteria" "Firmicutes"          "Negativicutes"              
##  [165,] "Bacteria" "Epsilonbacteraeota"  "Campylobacteria"            
##  [166,] "Bacteria" "Bacteroidetes"       "Bacteroidia"                
##  [167,] "Bacteria" "Bacteroidetes"       "Bacteroidia"                
##  [168,] "Bacteria" "Firmicutes"          "Bacilli"                    
##  [169,] "Bacteria" "Bacteroidetes"       "Bacteroidia"                
##  [170,] "Bacteria" "Proteobacteria"      "Gammaproteobacteria"        
##  [171,] "Bacteria" "Firmicutes"          "Bacilli"                    
##  [172,] "Bacteria" "Firmicutes"          "Bacilli"                    
##  [173,] "Bacteria" "Patescibacteria"     "Saccharimonadia"            
##  [174,] "Bacteria" "Bacteroidetes"       "Bacteroidia"                
##  [175,] "Bacteria" "Bacteroidetes"       "Bacteroidia"                
##  [176,] "Bacteria" "Firmicutes"          "Clostridia"                 
##  [177,] "Bacteria" "Firmicutes"          "Negativicutes"              
##  [178,] "Bacteria" "Firmicutes"          "Negativicutes"              
##  [179,] "Bacteria" "Bacteroidetes"       "Bacteroidia"                
##  [180,] "Bacteria" "Proteobacteria"      "Gammaproteobacteria"        
##  [181,] "Bacteria" "Patescibacteria"     "Gracilibacteria"            
##  [182,] "Bacteria" "Firmicutes"          "Negativicutes"              
##  [183,] "Bacteria" "Proteobacteria"      "Gammaproteobacteria"        
##  [184,] "Bacteria" "Firmicutes"          "Negativicutes"              
##  [185,] "Bacteria" "Proteobacteria"      "Gammaproteobacteria"        
##  [186,] "Bacteria" "Firmicutes"          "Negativicutes"              
##  [187,] "Bacteria" "Firmicutes"          "Negativicutes"              
##  [188,] "Bacteria" "Bacteroidetes"       "Bacteroidia"                
##  [189,] "Bacteria" "Bacteroidetes"       "Bacteroidia"                
##  [190,] "Bacteria" "Proteobacteria"      "Gammaproteobacteria"        
##  [191,] "Bacteria" "Bacteroidetes"       "Bacteroidia"                
##  [192,] "Bacteria" "Bacteroidetes"       "Bacteroidia"                
##  [193,] "Bacteria" "Proteobacteria"      "Gammaproteobacteria"        
##  [194,] "Bacteria" "Bacteroidetes"       "Bacteroidia"                
##  [195,] "Bacteria" "Firmicutes"          "Negativicutes"              
##  [196,] "Bacteria" "Firmicutes"          "Negativicutes"              
##  [197,] "Bacteria" "Fusobacteria"        "Fusobacteriia"              
##  [198,] "Bacteria" "Firmicutes"          "Negativicutes"              
##  [199,] "Bacteria" "Bacteroidetes"       "Bacteroidia"                
##  [200,] "Bacteria" "Firmicutes"          "Negativicutes"              
##  [201,] "Bacteria" "Bacteroidetes"       "Bacteroidia"                
##  [202,] "Bacteria" "Firmicutes"          "Clostridia"                 
##  [203,] "Bacteria" "Epsilonbacteraeota"  "Campylobacteria"            
##  [204,] "Bacteria" "Bacteroidetes"       "Bacteroidia"                
##  [205,] "Bacteria" "Firmicutes"          "Negativicutes"              
##  [206,] "Bacteria" "Firmicutes"          "Negativicutes"              
##  [207,] "Bacteria" "Bacteroidetes"       "Bacteroidia"                
##  [208,] "Bacteria" "Proteobacteria"      "Gammaproteobacteria"        
##  [209,] "Bacteria" "Proteobacteria"      "Gammaproteobacteria"        
##  [210,] "Bacteria" "Firmicutes"          "Bacilli"                    
##  [211,] "Bacteria" "Firmicutes"          "Clostridia"                 
##  [212,] "Bacteria" "Firmicutes"          "Negativicutes"              
##  [213,] "Bacteria" "Proteobacteria"      "Gammaproteobacteria"        
##  [214,] "Bacteria" "Proteobacteria"      "Gammaproteobacteria"        
##  [215,] "Bacteria" "Firmicutes"          "Negativicutes"              
##  [216,] "Bacteria" "Bacteroidetes"       "Bacteroidia"                
##  [217,] "Bacteria" "Firmicutes"          "Bacilli"                    
##  [218,] "Bacteria" "Proteobacteria"      "Gammaproteobacteria"        
##  [219,] "Bacteria" "Firmicutes"          "Negativicutes"              
##  [220,] "Bacteria" "Firmicutes"          "Bacilli"                    
##  [221,] "Bacteria" "Firmicutes"          "Clostridia"                 
##  [222,] "Bacteria" "Firmicutes"          "Bacilli"                    
##  [223,] "Bacteria" "Firmicutes"          "Clostridia"                 
##  [224,] "Bacteria" "Bacteroidetes"       "Bacteroidia"                
##  [225,] "Bacteria" "Fusobacteria"        "Fusobacteriia"              
##  [226,] "Bacteria" "Firmicutes"          "Negativicutes"              
##  [227,] "Bacteria" "Bacteroidetes"       "Bacteroidia"                
##  [228,] "Bacteria" "Firmicutes"          "Bacilli"                    
##  [229,] "Bacteria" "Bacteroidetes"       "Bacteroidia"                
##  [230,] "Bacteria" "Bacteroidetes"       "Bacteroidia"                
##  [231,] "Bacteria" "Fusobacteria"        "Fusobacteriia"              
##  [232,] "Bacteria" "Fusobacteria"        "Fusobacteriia"              
##  [233,] "Bacteria" "Proteobacteria"      "Gammaproteobacteria"        
##  [234,] "Bacteria" "Fusobacteria"        "Fusobacteriia"              
##  [235,] "Bacteria" "Firmicutes"          "Negativicutes"              
##  [236,] "Bacteria" "Patescibacteria"     "Saccharimonadia"            
##  [237,] "Bacteria" "Bacteroidetes"       "Bacteroidia"                
##  [238,] "Bacteria" "Firmicutes"          "Negativicutes"              
##  [239,] "Bacteria" "Bacteroidetes"       "Bacteroidia"                
##  [240,] "Bacteria" "Firmicutes"          "Negativicutes"              
##  [241,] "Bacteria" "Fusobacteria"        "Fusobacteriia"              
##  [242,] "Bacteria" "Bacteroidetes"       "Bacteroidia"                
##  [243,] "Bacteria" "Firmicutes"          "Negativicutes"              
##  [244,] "Bacteria" "Cyanobacteria"       "Oxyphotobacteria"           
##  [245,] "Bacteria" "Bacteroidetes"       "Bacteroidia"                
##  [246,] "Bacteria" "Proteobacteria"      "Gammaproteobacteria"        
##  [247,] "Bacteria" "Proteobacteria"      "Gammaproteobacteria"        
##  [248,] "Bacteria" "Cyanobacteria"       "Oxyphotobacteria"           
##  [249,] "Bacteria" "Patescibacteria"     "Gracilibacteria"            
##  [250,] "Bacteria" "Bacteroidetes"       "Bacteroidia"                
##  [251,] "Bacteria" "Bacteroidetes"       "Bacteroidia"                
##  [252,] "Bacteria" "Bacteroidetes"       "Bacteroidia"                
##  [253,] "Bacteria" "Firmicutes"          "Clostridia"                 
##  [254,] "Bacteria" "Proteobacteria"      "Gammaproteobacteria"        
##  [255,] "Bacteria" "Proteobacteria"      "Gammaproteobacteria"        
##  [256,] "Bacteria" "Firmicutes"          "Negativicutes"              
##  [257,] "Bacteria" "Firmicutes"          "Negativicutes"              
##  [258,] "Bacteria" "Proteobacteria"      "Gammaproteobacteria"        
##  [259,] "Bacteria" "Patescibacteria"     "Saccharimonadia"            
##  [260,] "Bacteria" "Actinobacteria"      "Actinobacteria"             
##  [261,] "Bacteria" "Firmicutes"          "Clostridia"                 
##  [262,] "Bacteria" "Bacteroidetes"       "Bacteroidia"                
##  [263,] "Bacteria" "Bacteroidetes"       "Bacteroidia"                
##  [264,] "Bacteria" "Proteobacteria"      "Gammaproteobacteria"        
##  [265,] "Bacteria" "Patescibacteria"     "Saccharimonadia"            
##  [266,] "Bacteria" "Proteobacteria"      "Gammaproteobacteria"        
##  [267,] "Bacteria" "Firmicutes"          "Erysipelotrichia"           
##  [268,] "Bacteria" "Bacteroidetes"       "Bacteroidia"                
##  [269,] "Bacteria" "Firmicutes"          "Clostridia"                 
##  [270,] "Bacteria" "Proteobacteria"      "Gammaproteobacteria"        
##  [271,] "Bacteria" "Firmicutes"          "Negativicutes"              
##  [272,] "Bacteria" "Proteobacteria"      "Gammaproteobacteria"        
##  [273,] "Bacteria" "Proteobacteria"      "Gammaproteobacteria"        
##  [274,] "Bacteria" "Firmicutes"          "Negativicutes"              
##  [275,] "Bacteria" "Firmicutes"          "Bacilli"                    
##  [276,] "Bacteria" "Firmicutes"          "Negativicutes"              
##  [277,] "Bacteria" "Bacteroidetes"       "Bacteroidia"                
##  [278,] "Bacteria" "Bacteroidetes"       "Bacteroidia"                
##  [279,] "Bacteria" "Proteobacteria"      "Gammaproteobacteria"        
##  [280,] "Bacteria" "Bacteroidetes"       "Bacteroidia"                
##  [281,] "Bacteria" "Fusobacteria"        "Fusobacteriia"              
##  [282,] "Bacteria" "Firmicutes"          NA                           
##  [283,] "Bacteria" "Fusobacteria"        "Fusobacteriia"              
##  [284,] "Bacteria" "Bacteroidetes"       "Bacteroidia"                
##  [285,] "Bacteria" "Bacteroidetes"       "Bacteroidia"                
##  [286,] "Bacteria" "Patescibacteria"     "Gracilibacteria"            
##  [287,] "Bacteria" "Bacteroidetes"       "Bacteroidia"                
##  [288,] "Bacteria" "Firmicutes"          "Negativicutes"              
##  [289,] "Bacteria" "Bacteroidetes"       "Bacteroidia"                
##  [290,] "Bacteria" "Firmicutes"          "Clostridia"                 
##  [291,] "Bacteria" "Bacteroidetes"       "Bacteroidia"                
##  [292,] "Bacteria" "Proteobacteria"      "Gammaproteobacteria"        
##  [293,] "Bacteria" "Firmicutes"          "Negativicutes"              
##  [294,] "Bacteria" "Patescibacteria"     "Saccharimonadia"            
##  [295,] "Bacteria" "Firmicutes"          "Bacilli"                    
##  [296,] "Bacteria" "Firmicutes"          "Negativicutes"              
##  [297,] "Bacteria" "Firmicutes"          NA                           
##  [298,] "Bacteria" "Firmicutes"          "Negativicutes"              
##  [299,] "Bacteria" "Proteobacteria"      "Gammaproteobacteria"        
##  [300,] "Bacteria" "Bacteroidetes"       "Bacteroidia"                
##  [301,] "Bacteria" "Bacteroidetes"       "Bacteroidia"                
##  [302,] "Bacteria" "Fusobacteria"        "Fusobacteriia"              
##  [303,] "Bacteria" "Bacteroidetes"       "Bacteroidia"                
##  [304,] "Bacteria" "Fusobacteria"        "Fusobacteriia"              
##  [305,] "Bacteria" "Fusobacteria"        "Fusobacteriia"              
##  [306,] "Bacteria" "Bacteroidetes"       "Bacteroidia"                
##  [307,] "Bacteria" "Firmicutes"          "Negativicutes"              
##  [308,] "Bacteria" "Fusobacteria"        "Fusobacteriia"              
##  [309,] "Bacteria" "Firmicutes"          "Clostridia"                 
##  [310,] "Bacteria" "Proteobacteria"      "Gammaproteobacteria"        
##  [311,] "Bacteria" "Firmicutes"          "Negativicutes"              
##  [312,] "Bacteria" "Fusobacteria"        "Fusobacteriia"              
##  [313,] "Bacteria" "Patescibacteria"     "Saccharimonadia"            
##  [314,] "Bacteria" "Epsilonbacteraeota"  "Campylobacteria"            
##  [315,] "Bacteria" "Firmicutes"          "Negativicutes"              
##  [316,] "Bacteria" "Bacteroidetes"       "Bacteroidia"                
##  [317,] "Bacteria" "Firmicutes"          "Negativicutes"              
##  [318,] "Bacteria" "Firmicutes"          "Clostridia"                 
##  [319,] "Bacteria" "Firmicutes"          "Clostridia"                 
##  [320,] "Bacteria" "Bacteroidetes"       "Bacteroidia"                
##  [321,] "Bacteria" "Actinobacteria"      "Coriobacteriia"             
##  [322,] "Bacteria" "Firmicutes"          "Bacilli"                    
##  [323,] "Bacteria" "Patescibacteria"     "Saccharimonadia"            
##  [324,] "Bacteria" "Firmicutes"          "Negativicutes"              
##  [325,] "Bacteria" "Proteobacteria"      "Gammaproteobacteria"        
##  [326,] "Bacteria" "Patescibacteria"     "Saccharimonadia"            
##  [327,] "Bacteria" "Bacteroidetes"       "Bacteroidia"                
##  [328,] "Bacteria" "Actinobacteria"      "Coriobacteriia"             
##  [329,] "Bacteria" "Firmicutes"          "Negativicutes"              
##  [330,] "Bacteria" "Bacteroidetes"       "Bacteroidia"                
##  [331,] "Bacteria" "Bacteroidetes"       "Bacteroidia"                
##  [332,] "Bacteria" "Patescibacteria"     "Saccharimonadia"            
##  [333,] "Bacteria" "Firmicutes"          "Clostridia"                 
##  [334,] "Bacteria" "Firmicutes"          "Negativicutes"              
##  [335,] "Bacteria" "Firmicutes"          "Clostridia"                 
##  [336,] "Bacteria" "Fusobacteria"        "Fusobacteriia"              
##  [337,] "Bacteria" "Firmicutes"          "Bacilli"                    
##  [338,] "Bacteria" "Fusobacteria"        "Fusobacteriia"              
##  [339,] "Bacteria" "Actinobacteria"      "Actinobacteria"             
##  [340,] "Bacteria" "Fusobacteria"        "Fusobacteriia"              
##  [341,] "Bacteria" "Fusobacteria"        "Fusobacteriia"              
##  [342,] "Bacteria" "Bacteroidetes"       "Bacteroidia"                
##  [343,] "Bacteria" "Proteobacteria"      "Gammaproteobacteria"        
##  [344,] "Bacteria" "Bacteroidetes"       "Bacteroidia"                
##  [345,] "Bacteria" "Firmicutes"          "Clostridia"                 
##  [346,] "Bacteria" "Proteobacteria"      "Gammaproteobacteria"        
##  [347,] "Bacteria" "Firmicutes"          NA                           
##  [348,] "Bacteria" "Proteobacteria"      "Gammaproteobacteria"        
##  [349,] "Bacteria" "Proteobacteria"      "Gammaproteobacteria"        
##  [350,] "Bacteria" "Bacteroidetes"       "Bacteroidia"                
##  [351,] "Bacteria" "Firmicutes"          "Clostridia"                 
##  [352,] "Bacteria" "Firmicutes"          "Negativicutes"              
##  [353,] "Bacteria" "Bacteroidetes"       "Bacteroidia"                
##  [354,] "Bacteria" "Proteobacteria"      "Gammaproteobacteria"        
##  [355,] "Bacteria" "Fusobacteria"        "Fusobacteriia"              
##  [356,] "Bacteria" "Firmicutes"          "Negativicutes"              
##  [357,] "Bacteria" "Fusobacteria"        "Fusobacteriia"              
##  [358,] "Bacteria" "Bacteroidetes"       "Bacteroidia"                
##  [359,] "Bacteria" "Firmicutes"          "Negativicutes"              
##  [360,] "Bacteria" "Firmicutes"          "Clostridia"                 
##  [361,] "Bacteria" "Firmicutes"          "Clostridia"                 
##  [362,] "Bacteria" "Proteobacteria"      "Gammaproteobacteria"        
##  [363,] "Bacteria" "Bacteroidetes"       "Bacteroidia"                
##  [364,] "Bacteria" "Firmicutes"          "Clostridia"                 
##  [365,] "Bacteria" "Fusobacteria"        "Fusobacteriia"              
##  [366,] "Bacteria" "Patescibacteria"     "Saccharimonadia"            
##  [367,] "Bacteria" "Firmicutes"          "Clostridia"                 
##  [368,] "Bacteria" "Firmicutes"          "Clostridia"                 
##  [369,] "Bacteria" "Actinobacteria"      "Actinobacteria"             
##  [370,] "Bacteria" "Bacteroidetes"       "Bacteroidia"                
##  [371,] "Bacteria" "Patescibacteria"     "Saccharimonadia"            
##  [372,] "Bacteria" "Firmicutes"          "Bacilli"                    
##  [373,] "Bacteria" "Firmicutes"          "Bacilli"                    
##  [374,] "Bacteria" "Firmicutes"          "Negativicutes"              
##  [375,] "Bacteria" "Patescibacteria"     "Saccharimonadia"            
##  [376,] "Bacteria" "Firmicutes"          "Negativicutes"              
##  [377,] "Bacteria" "Fusobacteria"        "Fusobacteriia"              
##  [378,] "Bacteria" "Firmicutes"          "Bacilli"                    
##  [379,] "Bacteria" "Firmicutes"          "Negativicutes"              
##  [380,] "Bacteria" "Fusobacteria"        "Fusobacteriia"              
##  [381,] "Bacteria" "Fusobacteria"        "Fusobacteriia"              
##  [382,] "Bacteria" "Firmicutes"          "Clostridia"                 
##  [383,] "Bacteria" "Bacteroidetes"       "Bacteroidia"                
##  [384,] "Bacteria" "Bacteroidetes"       "Bacteroidia"                
##  [385,] "Bacteria" "Firmicutes"          "Bacilli"                    
##  [386,] "Bacteria" "Proteobacteria"      "Gammaproteobacteria"        
##  [387,] "Bacteria" "Fusobacteria"        "Fusobacteriia"              
##  [388,] "Bacteria" "Actinobacteria"      "Actinobacteria"             
##  [389,] "Bacteria" "Fusobacteria"        "Fusobacteriia"              
##  [390,] "Bacteria" "Bacteroidetes"       "Bacteroidia"                
##  [391,] "Bacteria" "Actinobacteria"      "Actinobacteria"             
##  [392,] "Bacteria" "Fusobacteria"        "Fusobacteriia"              
##  [393,] "Bacteria" "Proteobacteria"      "Gammaproteobacteria"        
##  [394,] "Bacteria" "Proteobacteria"      "Gammaproteobacteria"        
##  [395,] "Bacteria" "Firmicutes"          "Negativicutes"              
##  [396,] "Bacteria" "Bacteroidetes"       "Bacteroidia"                
##  [397,] "Bacteria" "Bacteroidetes"       "Bacteroidia"                
##  [398,] "Bacteria" "Proteobacteria"      "Gammaproteobacteria"        
##  [399,] "Bacteria" "Fusobacteria"        "Fusobacteriia"              
##  [400,] "Bacteria" "Proteobacteria"      "Gammaproteobacteria"        
##  [401,] "Bacteria" "Firmicutes"          "Bacilli"                    
##  [402,] "Bacteria" "Bacteroidetes"       "Bacteroidia"                
##  [403,] "Bacteria" "Bacteroidetes"       "Bacteroidia"                
##  [404,] "Bacteria" "Bacteroidetes"       "Bacteroidia"                
##  [405,] "Bacteria" "Firmicutes"          "Clostridia"                 
##  [406,] "Bacteria" "Firmicutes"          "Negativicutes"              
##  [407,] "Bacteria" "Bacteroidetes"       "Bacteroidia"                
##  [408,] "Bacteria" "Firmicutes"          "Bacilli"                    
##  [409,] "Bacteria" "Firmicutes"          "Negativicutes"              
##  [410,] "Bacteria" "Bacteroidetes"       "Bacteroidia"                
##  [411,] "Bacteria" "Bacteroidetes"       "Bacteroidia"                
##  [412,] "Bacteria" "Bacteroidetes"       "Bacteroidia"                
##  [413,] "Bacteria" "Proteobacteria"      "Gammaproteobacteria"        
##  [414,] "Bacteria" "Firmicutes"          "Clostridia"                 
##  [415,] "Bacteria" "Bacteroidetes"       "Bacteroidia"                
##  [416,] "Bacteria" "Proteobacteria"      "Gammaproteobacteria"        
##  [417,] "Bacteria" "Bacteroidetes"       "Bacteroidia"                
##  [418,] "Bacteria" "Bacteroidetes"       "Bacteroidia"                
##  [419,] "Bacteria" "Fusobacteria"        "Fusobacteriia"              
##  [420,] "Bacteria" "Bacteroidetes"       "Bacteroidia"                
##  [421,] "Bacteria" "Firmicutes"          "Bacilli"                    
##  [422,] "Bacteria" "Firmicutes"          "Negativicutes"              
##  [423,] "Bacteria" "Firmicutes"          "Negativicutes"              
##  [424,] "Bacteria" "Firmicutes"          "Negativicutes"              
##  [425,] "Bacteria" "Proteobacteria"      "Gammaproteobacteria"        
##  [426,] "Bacteria" "Bacteroidetes"       "Bacteroidia"                
##  [427,] "Bacteria" "Bacteroidetes"       "Bacteroidia"                
##  [428,] "Bacteria" "Bacteroidetes"       "Bacteroidia"                
##  [429,] "Bacteria" "Firmicutes"          "Negativicutes"              
##  [430,] "Bacteria" "Firmicutes"          "Negativicutes"              
##  [431,] "Bacteria" "Firmicutes"          "Negativicutes"              
##  [432,] "Bacteria" "Bacteroidetes"       "Bacteroidia"                
##  [433,] "Bacteria" "Firmicutes"          "Negativicutes"              
##  [434,] "Bacteria" "Bacteroidetes"       "Bacteroidia"                
##  [435,] "Bacteria" "Firmicutes"          "Clostridia"                 
##  [436,] "Bacteria" "Firmicutes"          "Negativicutes"              
##  [437,] "Bacteria" "Bacteroidetes"       "Bacteroidia"                
##  [438,] "Bacteria" "Firmicutes"          "Bacilli"                    
##  [439,] "Bacteria" "Bacteroidetes"       "Bacteroidia"                
##  [440,] "Bacteria" "Proteobacteria"      "Gammaproteobacteria"        
##  [441,] "Bacteria" "Bacteroidetes"       "Bacteroidia"                
##  [442,] "Bacteria" "Fusobacteria"        "Fusobacteriia"              
##  [443,] "Bacteria" "Firmicutes"          "Negativicutes"              
##  [444,] "Bacteria" "Firmicutes"          "Negativicutes"              
##  [445,] "Bacteria" "Firmicutes"          "Negativicutes"              
##  [446,] "Bacteria" "Firmicutes"          "Negativicutes"              
##  [447,] "Bacteria" "Firmicutes"          "Bacilli"                    
##  [448,] "Bacteria" "Fusobacteria"        "Fusobacteriia"              
##  [449,] "Bacteria" "Bacteroidetes"       "Bacteroidia"                
##  [450,] "Bacteria" "Fusobacteria"        "Fusobacteriia"              
##  [451,] "Bacteria" "Bacteroidetes"       "Bacteroidia"                
##  [452,] "Bacteria" "Epsilonbacteraeota"  "Campylobacteria"            
##  [453,] "Bacteria" "Firmicutes"          "Clostridia"                 
##  [454,] "Bacteria" "Bacteroidetes"       "Bacteroidia"                
##  [455,] "Bacteria" "Firmicutes"          "Negativicutes"              
##  [456,] "Bacteria" "Firmicutes"          "Clostridia"                 
##  [457,] "Bacteria" "Proteobacteria"      "Gammaproteobacteria"        
##  [458,] "Bacteria" "Actinobacteria"      "Actinobacteria"             
##  [459,] "Bacteria" "Fusobacteria"        "Fusobacteriia"              
##  [460,] "Bacteria" "Firmicutes"          "Negativicutes"              
##  [461,] "Bacteria" "Patescibacteria"     "Saccharimonadia"            
##  [462,] "Bacteria" "Fusobacteria"        "Fusobacteriia"              
##  [463,] "Bacteria" "Bacteroidetes"       "Bacteroidia"                
##  [464,] "Bacteria" "Firmicutes"          "Bacilli"                    
##  [465,] "Bacteria" "Bacteroidetes"       "Bacteroidia"                
##  [466,] "Bacteria" "Firmicutes"          "Negativicutes"              
##  [467,] "Bacteria" "Bacteroidetes"       "Bacteroidia"                
##  [468,] "Bacteria" "Proteobacteria"      "Gammaproteobacteria"        
##  [469,] "Bacteria" "Patescibacteria"     "Saccharimonadia"            
##  [470,] "Bacteria" "Proteobacteria"      "Gammaproteobacteria"        
##  [471,] "Bacteria" "Proteobacteria"      "Gammaproteobacteria"        
##  [472,] "Bacteria" "Bacteroidetes"       "Bacteroidia"                
##  [473,] "Bacteria" "Bacteroidetes"       "Bacteroidia"                
##  [474,] "Bacteria" "Firmicutes"          "Negativicutes"              
##  [475,] "Bacteria" "Bacteroidetes"       "Bacteroidia"                
##  [476,] "Bacteria" "Proteobacteria"      "Gammaproteobacteria"        
##  [477,] "Bacteria" "Bacteroidetes"       "Bacteroidia"                
##  [478,] "Bacteria" "Proteobacteria"      "Gammaproteobacteria"        
##  [479,] "Bacteria" "Proteobacteria"      "Gammaproteobacteria"        
##  [480,] "Bacteria" "Fusobacteria"        "Fusobacteriia"              
##  [481,] "Bacteria" "Bacteroidetes"       "Bacteroidia"                
##  [482,] "Bacteria" "Bacteroidetes"       "Bacteroidia"                
##  [483,] "Bacteria" "Fusobacteria"        "Fusobacteriia"              
##  [484,] "Bacteria" "Firmicutes"          "Negativicutes"              
##  [485,] "Bacteria" "Firmicutes"          "Bacilli"                    
##  [486,] "Bacteria" "Fusobacteria"        "Fusobacteriia"              
##  [487,] "Bacteria" "Bacteroidetes"       "Bacteroidia"                
##  [488,] "Bacteria" "Fusobacteria"        "Fusobacteriia"              
##  [489,] "Bacteria" "Bacteroidetes"       "Bacteroidia"                
##  [490,] "Bacteria" "Epsilonbacteraeota"  "Campylobacteria"            
##  [491,] "Bacteria" "Firmicutes"          "Negativicutes"              
##  [492,] "Bacteria" "Epsilonbacteraeota"  "Campylobacteria"            
##  [493,] "Bacteria" "Bacteroidetes"       "Bacteroidia"                
##  [494,] "Bacteria" "Fusobacteria"        "Fusobacteriia"              
##  [495,] "Bacteria" "Bacteroidetes"       "Bacteroidia"                
##  [496,] "Bacteria" "Firmicutes"          "Negativicutes"              
##  [497,] "Bacteria" "Proteobacteria"      "Gammaproteobacteria"        
##  [498,] "Bacteria" "Bacteroidetes"       "Bacteroidia"                
##  [499,] "Bacteria" "Firmicutes"          "Clostridia"                 
##  [500,] "Bacteria" "Firmicutes"          "Negativicutes"              
##  [501,] "Bacteria" "Firmicutes"          "Negativicutes"              
##  [502,] "Bacteria" "Actinobacteria"      "Actinobacteria"             
##  [503,] "Bacteria" "Proteobacteria"      "Gammaproteobacteria"        
##  [504,] "Bacteria" "Firmicutes"          "Negativicutes"              
##  [505,] "Bacteria" "Proteobacteria"      "Gammaproteobacteria"        
##  [506,] "Bacteria" "Fusobacteria"        "Fusobacteriia"              
##  [507,] "Bacteria" "Actinobacteria"      "Actinobacteria"             
##  [508,] "Bacteria" "Bacteroidetes"       "Bacteroidia"                
##  [509,] "Bacteria" "Bacteroidetes"       "Bacteroidia"                
##  [510,] "Bacteria" "Proteobacteria"      "Gammaproteobacteria"        
##  [511,] "Bacteria" "Firmicutes"          "Negativicutes"              
##  [512,] "Bacteria" "Firmicutes"          "Negativicutes"              
##  [513,] "Bacteria" "Proteobacteria"      "Gammaproteobacteria"        
##  [514,] "Bacteria" "Firmicutes"          "Negativicutes"              
##  [515,] "Bacteria" "Firmicutes"          "Negativicutes"              
##  [516,] "Bacteria" "Firmicutes"          "Negativicutes"              
##  [517,] "Bacteria" "Bacteroidetes"       "Bacteroidia"                
##  [518,] "Bacteria" "Proteobacteria"      "Gammaproteobacteria"        
##  [519,] "Bacteria" "Firmicutes"          "Negativicutes"              
##  [520,] "Bacteria" "Actinobacteria"      "Actinobacteria"             
##  [521,] "Bacteria" "Firmicutes"          "Bacilli"                    
##  [522,] "Bacteria" "Actinobacteria"      "Actinobacteria"             
##  [523,] "Bacteria" "Firmicutes"          "Clostridia"                 
##  [524,] "Bacteria" "Firmicutes"          "Negativicutes"              
##  [525,] "Bacteria" "Fusobacteria"        "Fusobacteriia"              
##  [526,] "Bacteria" "Bacteroidetes"       "Bacteroidia"                
##  [527,] "Bacteria" "Bacteroidetes"       "Bacteroidia"                
##  [528,] "Bacteria" "Proteobacteria"      "Gammaproteobacteria"        
##  [529,] "Bacteria" "Epsilonbacteraeota"  "Campylobacteria"            
##  [530,] "Bacteria" "Firmicutes"          "Bacilli"                    
##  [531,] "Bacteria" "Proteobacteria"      "Gammaproteobacteria"        
##  [532,] "Bacteria" "Bacteroidetes"       "Bacteroidia"                
##  [533,] "Bacteria" "Patescibacteria"     "Saccharimonadia"            
##  [534,] "Bacteria" "Firmicutes"          "Negativicutes"              
##  [535,] "Bacteria" "Bacteroidetes"       "Bacteroidia"                
##  [536,] "Bacteria" "Bacteroidetes"       "Bacteroidia"                
##  [537,] "Bacteria" "Firmicutes"          "Negativicutes"              
##  [538,] "Bacteria" "Proteobacteria"      "Gammaproteobacteria"        
##  [539,] "Bacteria" "Bacteroidetes"       "Bacteroidia"                
##  [540,] "Bacteria" "Proteobacteria"      "Gammaproteobacteria"        
##  [541,] "Bacteria" "Fusobacteria"        "Fusobacteriia"              
##  [542,] "Bacteria" "Firmicutes"          "Bacilli"                    
##  [543,] "Bacteria" "Bacteroidetes"       "Bacteroidia"                
##  [544,] "Bacteria" "Bacteroidetes"       "Bacteroidia"                
##  [545,] "Bacteria" "Firmicutes"          "Clostridia"                 
##  [546,] "Bacteria" "Bacteroidetes"       "Bacteroidia"                
##  [547,] "Bacteria" "Proteobacteria"      "Gammaproteobacteria"        
##  [548,] "Bacteria" NA                    NA                           
##  [549,] "Bacteria" "Firmicutes"          "Negativicutes"              
##  [550,] "Bacteria" "Bacteroidetes"       "Bacteroidia"                
##  [551,] "Bacteria" "Firmicutes"          "Bacilli"                    
##  [552,] "Bacteria" "Proteobacteria"      "Gammaproteobacteria"        
##  [553,] "Bacteria" "Bacteroidetes"       "Bacteroidia"                
##  [554,] "Bacteria" "Bacteroidetes"       "Bacteroidia"                
##  [555,] "Bacteria" "Bacteroidetes"       "Bacteroidia"                
##  [556,] "Bacteria" "Firmicutes"          "Negativicutes"              
##  [557,] "Bacteria" "Firmicutes"          "Negativicutes"              
##  [558,] "Bacteria" "Firmicutes"          "Bacilli"                    
##  [559,] "Bacteria" "Bacteroidetes"       "Bacteroidia"                
##  [560,] "Bacteria" "Proteobacteria"      "Gammaproteobacteria"        
##  [561,] "Bacteria" "Bacteroidetes"       "Bacteroidia"                
##  [562,] "Bacteria" "Patescibacteria"     "Saccharimonadia"            
##  [563,] "Bacteria" "Proteobacteria"      "Gammaproteobacteria"        
##  [564,] "Bacteria" "Firmicutes"          "Bacilli"                    
##  [565,] "Bacteria" "Firmicutes"          "Negativicutes"              
##  [566,] "Bacteria" "Bacteroidetes"       "Bacteroidia"                
##  [567,] "Bacteria" "Firmicutes"          "Bacilli"                    
##  [568,] "Bacteria" "Bacteroidetes"       "Bacteroidia"                
##  [569,] "Bacteria" "Bacteroidetes"       "Bacteroidia"                
##  [570,] "Bacteria" "Fusobacteria"        "Fusobacteriia"              
##  [571,] "Bacteria" "Bacteroidetes"       "Bacteroidia"                
##  [572,] "Bacteria" "Bacteroidetes"       "Bacteroidia"                
##  [573,] "Bacteria" "Proteobacteria"      "Gammaproteobacteria"        
##  [574,] "Bacteria" "Bacteroidetes"       "Bacteroidia"                
##  [575,] "Bacteria" "Bacteroidetes"       "Bacteroidia"                
##  [576,] "Bacteria" "Bacteroidetes"       "Bacteroidia"                
##  [577,] "Bacteria" "Patescibacteria"     "Saccharimonadia"            
##  [578,] "Bacteria" "Firmicutes"          "Negativicutes"              
##  [579,] "Bacteria" "Firmicutes"          "Negativicutes"              
##  [580,] "Bacteria" "Firmicutes"          "Negativicutes"              
##  [581,] "Bacteria" "Fusobacteria"        "Fusobacteriia"              
##  [582,] "Bacteria" "Proteobacteria"      "Gammaproteobacteria"        
##  [583,] "Bacteria" "Fusobacteria"        "Fusobacteriia"              
##  [584,] "Bacteria" "Bacteroidetes"       "Bacteroidia"                
##  [585,] "Bacteria" "Proteobacteria"      "Gammaproteobacteria"        
##  [586,] "Bacteria" "Fusobacteria"        "Fusobacteriia"              
##  [587,] "Bacteria" "Fusobacteria"        "Fusobacteriia"              
##  [588,] "Bacteria" "Bacteroidetes"       "Bacteroidia"                
##  [589,] "Bacteria" "Fusobacteria"        "Fusobacteriia"              
##  [590,] "Bacteria" "Bacteroidetes"       "Bacteroidia"                
##  [591,] "Bacteria" "Proteobacteria"      "Gammaproteobacteria"        
##  [592,] "Bacteria" "Bacteroidetes"       "Bacteroidia"                
##  [593,] "Bacteria" "Bacteroidetes"       "Bacteroidia"                
##  [594,] "Bacteria" "Bacteroidetes"       "Bacteroidia"                
##  [595,] "Bacteria" "Proteobacteria"      "Gammaproteobacteria"        
##  [596,] "Bacteria" "Bacteroidetes"       "Bacteroidia"                
##  [597,] "Bacteria" "Firmicutes"          "Negativicutes"              
##  [598,] "Bacteria" "Proteobacteria"      "Gammaproteobacteria"        
##  [599,] "Bacteria" "Firmicutes"          "Clostridia"                 
##  [600,] "Bacteria" "Firmicutes"          "Negativicutes"              
##  [601,] "Bacteria" "Proteobacteria"      "Gammaproteobacteria"        
##  [602,] "Bacteria" "Bacteroidetes"       "Bacteroidia"                
##  [603,] "Bacteria" "Bacteroidetes"       "Bacteroidia"                
##  [604,] "Bacteria" "Firmicutes"          "Negativicutes"              
##  [605,] "Bacteria" "Proteobacteria"      "Gammaproteobacteria"        
##  [606,] "Bacteria" "Fusobacteria"        "Fusobacteriia"              
##  [607,] "Bacteria" "Bacteroidetes"       "Bacteroidia"                
##  [608,] "Bacteria" "Patescibacteria"     "Saccharimonadia"            
##  [609,] "Bacteria" "Bacteroidetes"       "Bacteroidia"                
##  [610,] "Bacteria" "Patescibacteria"     "Saccharimonadia"            
##  [611,] "Bacteria" "Proteobacteria"      "Gammaproteobacteria"        
##  [612,] "Bacteria" "Firmicutes"          "Negativicutes"              
##  [613,] "Bacteria" "Fusobacteria"        "Fusobacteriia"              
##  [614,] "Bacteria" "Fusobacteria"        "Fusobacteriia"              
##  [615,] "Bacteria" "Bacteroidetes"       "Bacteroidia"                
##  [616,] "Bacteria" "Firmicutes"          "Negativicutes"              
##  [617,] "Bacteria" "Bacteroidetes"       "Bacteroidia"                
##  [618,] "Bacteria" "Firmicutes"          "Negativicutes"              
##  [619,] "Bacteria" "Firmicutes"          "Bacilli"                    
##  [620,] "Bacteria" "Bacteroidetes"       "Bacteroidia"                
##  [621,] "Bacteria" "Firmicutes"          "Negativicutes"              
##  [622,] "Bacteria" "Bacteroidetes"       "Bacteroidia"                
##  [623,] "Bacteria" "Firmicutes"          "Negativicutes"              
##  [624,] "Bacteria" "Bacteroidetes"       "Bacteroidia"                
##  [625,] "Bacteria" "Bacteroidetes"       "Bacteroidia"                
##  [626,] "Bacteria" "Bacteroidetes"       "Bacteroidia"                
##  [627,] "Bacteria" "Bacteroidetes"       "Bacteroidia"                
##  [628,] "Bacteria" "Fusobacteria"        "Fusobacteriia"              
##  [629,] "Bacteria" "Bacteroidetes"       "Bacteroidia"                
##  [630,] "Bacteria" "Actinobacteria"      "Actinobacteria"             
##  [631,] "Bacteria" "Firmicutes"          "Negativicutes"              
##  [632,] "Bacteria" "Proteobacteria"      "Gammaproteobacteria"        
##  [633,] "Bacteria" "Actinobacteria"      "Coriobacteriia"             
##  [634,] "Bacteria" "Firmicutes"          "Negativicutes"              
##  [635,] "Bacteria" "Patescibacteria"     "Saccharimonadia"            
##  [636,] "Bacteria" "Actinobacteria"      "Actinobacteria"             
##  [637,] "Bacteria" "Bacteroidetes"       "Bacteroidia"                
##  [638,] "Bacteria" "Proteobacteria"      "Gammaproteobacteria"        
##  [639,] "Bacteria" "Firmicutes"          "Bacilli"                    
##  [640,] "Bacteria" "Patescibacteria"     "Gracilibacteria"            
##  [641,] "Bacteria" "Firmicutes"          "Negativicutes"              
##  [642,] "Bacteria" "Actinobacteria"      "Actinobacteria"             
##  [643,] "Bacteria" "Proteobacteria"      "Gammaproteobacteria"        
##  [644,] "Bacteria" "Bacteroidetes"       "Bacteroidia"                
##  [645,] "Bacteria" "Bacteroidetes"       "Bacteroidia"                
##  [646,] "Bacteria" "Bacteroidetes"       "Bacteroidia"                
##  [647,] "Bacteria" "Actinobacteria"      "Actinobacteria"             
##  [648,] "Bacteria" "Firmicutes"          "Bacilli"                    
##  [649,] "Bacteria" "Bacteroidetes"       "Bacteroidia"                
##  [650,] "Bacteria" "Firmicutes"          "Clostridia"                 
##  [651,] "Bacteria" "Firmicutes"          "Negativicutes"              
##  [652,] "Bacteria" "Bacteroidetes"       "Bacteroidia"                
##  [653,] "Bacteria" "Firmicutes"          "Negativicutes"              
##  [654,] "Bacteria" "Patescibacteria"     "Saccharimonadia"            
##  [655,] "Bacteria" "Bacteroidetes"       "Bacteroidia"                
##  [656,] "Bacteria" "Firmicutes"          "Negativicutes"              
##  [657,] "Bacteria" "Fusobacteria"        "Fusobacteriia"              
##  [658,] "Bacteria" "Bacteroidetes"       "Bacteroidia"                
##  [659,] "Bacteria" "Firmicutes"          "Negativicutes"              
##  [660,] "Bacteria" "Fusobacteria"        "Fusobacteriia"              
##  [661,] "Bacteria" "Actinobacteria"      "Actinobacteria"             
##  [662,] "Bacteria" "Fusobacteria"        "Fusobacteriia"              
##  [663,] "Bacteria" "Fusobacteria"        "Fusobacteriia"              
##  [664,] "Bacteria" "Proteobacteria"      "Gammaproteobacteria"        
##  [665,] "Bacteria" "Fusobacteria"        "Fusobacteriia"              
##  [666,] "Bacteria" "Firmicutes"          "Negativicutes"              
##  [667,] "Bacteria" "Cyanobacteria"       "Oxyphotobacteria"           
##  [668,] "Bacteria" "Firmicutes"          "Bacilli"                    
##  [669,] "Bacteria" "Bacteroidetes"       "Bacteroidia"                
##  [670,] "Bacteria" "Bacteroidetes"       "Bacteroidia"                
##  [671,] "Bacteria" "Actinobacteria"      "Actinobacteria"             
##  [672,] "Bacteria" "Actinobacteria"      "Actinobacteria"             
##  [673,] "Bacteria" "Patescibacteria"     "Saccharimonadia"            
##  [674,] "Bacteria" "Bacteroidetes"       "Bacteroidia"                
##  [675,] "Bacteria" "Firmicutes"          "Bacilli"                    
##  [676,] "Bacteria" "Proteobacteria"      "Gammaproteobacteria"        
##  [677,] "Bacteria" "Bacteroidetes"       "Bacteroidia"                
##  [678,] "Bacteria" "Firmicutes"          "Negativicutes"              
##  [679,] "Bacteria" "Bacteroidetes"       "Bacteroidia"                
##  [680,] "Bacteria" "Firmicutes"          NA                           
##  [681,] "Bacteria" "Bacteroidetes"       "Bacteroidia"                
##  [682,] "Bacteria" "Proteobacteria"      "Gammaproteobacteria"        
##  [683,] "Bacteria" "Firmicutes"          "Negativicutes"              
##  [684,] "Bacteria" "Bacteroidetes"       "Bacteroidia"                
##  [685,] "Bacteria" "Bacteroidetes"       "Bacteroidia"                
##  [686,] "Bacteria" "Bacteroidetes"       "Bacteroidia"                
##  [687,] "Bacteria" "Patescibacteria"     "Saccharimonadia"            
##  [688,] "Bacteria" "Firmicutes"          "Bacilli"                    
##  [689,] "Bacteria" "Firmicutes"          "Negativicutes"              
##  [690,] "Bacteria" "Firmicutes"          "Negativicutes"              
##  [691,] "Bacteria" "Bacteroidetes"       "Bacteroidia"                
##  [692,] "Bacteria" "Patescibacteria"     "Saccharimonadia"            
##  [693,] "Bacteria" "Patescibacteria"     "Gracilibacteria"            
##  [694,] "Bacteria" "Bacteroidetes"       "Bacteroidia"                
##  [695,] "Bacteria" "Firmicutes"          "Negativicutes"              
##  [696,] "Bacteria" "Firmicutes"          "Bacilli"                    
##  [697,] "Bacteria" "Firmicutes"          "Negativicutes"              
##  [698,] "Bacteria" "Bacteroidetes"       "Bacteroidia"                
##  [699,] "Bacteria" "Bacteroidetes"       "Bacteroidia"                
##  [700,] "Bacteria" "Spirochaetes"        "Spirochaetia"               
##  [701,] "Bacteria" "Bacteroidetes"       "Bacteroidia"                
##  [702,] "Bacteria" "Firmicutes"          "Negativicutes"              
##  [703,] "Bacteria" "Fusobacteria"        "Fusobacteriia"              
##  [704,] "Bacteria" "Epsilonbacteraeota"  "Campylobacteria"            
##  [705,] "Bacteria" "Bacteroidetes"       "Bacteroidia"                
##  [706,] "Bacteria" "Patescibacteria"     "Saccharimonadia"            
##  [707,] "Bacteria" "Actinobacteria"      "Actinobacteria"             
##  [708,] "Bacteria" "Bacteroidetes"       "Bacteroidia"                
##  [709,] "Bacteria" "Bacteroidetes"       "Bacteroidia"                
##  [710,] "Bacteria" "Proteobacteria"      "Gammaproteobacteria"        
##  [711,] "Bacteria" "Bacteroidetes"       "Bacteroidia"                
##  [712,] "Bacteria" "Firmicutes"          "Negativicutes"              
##  [713,] "Bacteria" "Bacteroidetes"       "Bacteroidia"                
##  [714,] "Bacteria" "Bacteroidetes"       "Bacteroidia"                
##  [715,] "Bacteria" "Proteobacteria"      "Gammaproteobacteria"        
##  [716,] "Bacteria" "Patescibacteria"     "Saccharimonadia"            
##  [717,] "Bacteria" "Firmicutes"          "Bacilli"                    
##  [718,] "Bacteria" "Spirochaetes"        "Spirochaetia"               
##  [719,] "Bacteria" "Bacteroidetes"       "Bacteroidia"                
##  [720,] "Bacteria" "Proteobacteria"      "Gammaproteobacteria"        
##  [721,] "Bacteria" "Bacteroidetes"       "Bacteroidia"                
##  [722,] "Bacteria" "Bacteroidetes"       "Bacteroidia"                
##  [723,] "Bacteria" "Bacteroidetes"       "Bacteroidia"                
##  [724,] "Bacteria" "Proteobacteria"      "Gammaproteobacteria"        
##  [725,] "Bacteria" "Proteobacteria"      "Gammaproteobacteria"        
##  [726,] "Bacteria" "Proteobacteria"      "Gammaproteobacteria"        
##  [727,] "Bacteria" "Proteobacteria"      "Gammaproteobacteria"        
##  [728,] "Bacteria" "Firmicutes"          "Negativicutes"              
##  [729,] "Bacteria" "Firmicutes"          "Negativicutes"              
##  [730,] "Bacteria" "Bacteroidetes"       "Bacteroidia"                
##  [731,] "Bacteria" "Proteobacteria"      "Gammaproteobacteria"        
##  [732,] "Bacteria" "Fusobacteria"        "Fusobacteriia"              
##  [733,] "Bacteria" "Proteobacteria"      "Gammaproteobacteria"        
##  [734,] "Bacteria" "Firmicutes"          "Negativicutes"              
##  [735,] "Bacteria" "Proteobacteria"      "Gammaproteobacteria"        
##  [736,] "Bacteria" "Bacteroidetes"       "Bacteroidia"                
##  [737,] "Bacteria" "Proteobacteria"      "Gammaproteobacteria"        
##  [738,] "Bacteria" "Actinobacteria"      "Actinobacteria"             
##  [739,] "Bacteria" "Proteobacteria"      "Gammaproteobacteria"        
##  [740,] "Bacteria" "Bacteroidetes"       "Bacteroidia"                
##  [741,] "Bacteria" "Fusobacteria"        "Fusobacteriia"              
##  [742,] "Bacteria" "Firmicutes"          "Bacilli"                    
##  [743,] "Bacteria" "Actinobacteria"      "Actinobacteria"             
##  [744,] "Bacteria" "Firmicutes"          "Bacilli"                    
##  [745,] "Bacteria" "Firmicutes"          "Negativicutes"              
##  [746,] "Bacteria" "Firmicutes"          "Negativicutes"              
##  [747,] "Bacteria" "Bacteroidetes"       "Bacteroidia"                
##  [748,] "Bacteria" "Patescibacteria"     "Saccharimonadia"            
##  [749,] "Bacteria" "Fusobacteria"        "Fusobacteriia"              
##  [750,] "Bacteria" "Proteobacteria"      "Gammaproteobacteria"        
##  [751,] "Bacteria" "Bacteroidetes"       "Bacteroidia"                
##  [752,] "Bacteria" "Bacteroidetes"       "Bacteroidia"                
##  [753,] "Bacteria" "Bacteroidetes"       "Bacteroidia"                
##  [754,] "Bacteria" "Bacteroidetes"       "Bacteroidia"                
##  [755,] "Bacteria" "Firmicutes"          "Negativicutes"              
##  [756,] "Bacteria" "Firmicutes"          "Negativicutes"              
##  [757,] "Bacteria" "Bacteroidetes"       "Bacteroidia"                
##  [758,] "Bacteria" "Fusobacteria"        "Fusobacteriia"              
##  [759,] "Bacteria" "Firmicutes"          "Negativicutes"              
##  [760,] "Bacteria" "Bacteroidetes"       "Bacteroidia"                
##  [761,] "Bacteria" "Firmicutes"          "Negativicutes"              
##  [762,] "Bacteria" "Bacteroidetes"       "Bacteroidia"                
##  [763,] "Bacteria" "Bacteroidetes"       "Bacteroidia"                
##  [764,] "Bacteria" "Firmicutes"          "Negativicutes"              
##  [765,] "Bacteria" "Fusobacteria"        "Fusobacteriia"              
##  [766,] "Bacteria" "Firmicutes"          "Negativicutes"              
##  [767,] "Bacteria" "Fusobacteria"        "Fusobacteriia"              
##  [768,] "Bacteria" "Fusobacteria"        "Fusobacteriia"              
##  [769,] "Bacteria" "Bacteroidetes"       "Bacteroidia"                
##  [770,] "Bacteria" "Bacteroidetes"       "Bacteroidia"                
##  [771,] "Bacteria" "Firmicutes"          "Bacilli"                    
##  [772,] "Bacteria" "Fusobacteria"        "Fusobacteriia"              
##  [773,] "Bacteria" "Bacteroidetes"       "Bacteroidia"                
##  [774,] "Bacteria" "Firmicutes"          "Bacilli"                    
##  [775,] "Bacteria" "Actinobacteria"      "Actinobacteria"             
##  [776,] "Bacteria" "Firmicutes"          "Negativicutes"              
##  [777,] "Bacteria" "Firmicutes"          "Negativicutes"              
##  [778,] "Bacteria" "Bacteroidetes"       "Bacteroidia"                
##  [779,] "Bacteria" "Proteobacteria"      "Gammaproteobacteria"        
##  [780,] "Bacteria" "Firmicutes"          "Clostridia"                 
##  [781,] "Bacteria" "Bacteroidetes"       "Bacteroidia"                
##  [782,] "Bacteria" "Proteobacteria"      "Gammaproteobacteria"        
##  [783,] "Bacteria" "Bacteroidetes"       "Bacteroidia"                
##  [784,] "Bacteria" "Proteobacteria"      "Gammaproteobacteria"        
##  [785,] "Bacteria" "Firmicutes"          "Clostridia"                 
##  [786,] "Bacteria" "Firmicutes"          "Bacilli"                    
##  [787,] "Bacteria" "Bacteroidetes"       "Bacteroidia"                
##  [788,] "Bacteria" "Bacteroidetes"       "Bacteroidia"                
##  [789,] "Bacteria" "Proteobacteria"      "Gammaproteobacteria"        
##  [790,] "Bacteria" "Firmicutes"          "Clostridia"                 
##  [791,] "Bacteria" "Firmicutes"          "Negativicutes"              
##  [792,] "Bacteria" "Firmicutes"          "Clostridia"                 
##  [793,] "Bacteria" "Firmicutes"          "Negativicutes"              
##  [794,] "Bacteria" "Fusobacteria"        "Fusobacteriia"              
##  [795,] "Bacteria" "Epsilonbacteraeota"  "Campylobacteria"            
##  [796,] "Bacteria" "Bacteroidetes"       "Bacteroidia"                
##  [797,] "Bacteria" "Firmicutes"          NA                           
##  [798,] "Bacteria" "Firmicutes"          "Negativicutes"              
##  [799,] "Bacteria" "Firmicutes"          "Negativicutes"              
##  [800,] "Bacteria" "Proteobacteria"      "Gammaproteobacteria"        
##  [801,] "Bacteria" "Bacteroidetes"       "Bacteroidia"                
##  [802,] "Bacteria" "Fusobacteria"        "Fusobacteriia"              
##  [803,] "Bacteria" "Bacteroidetes"       "Bacteroidia"                
##  [804,] "Bacteria" "Bacteroidetes"       "Bacteroidia"                
##  [805,] "Bacteria" "Actinobacteria"      "Actinobacteria"             
##  [806,] "Bacteria" "Firmicutes"          "Clostridia"                 
##  [807,] "Bacteria" "Proteobacteria"      "Gammaproteobacteria"        
##  [808,] "Bacteria" "Firmicutes"          "Bacilli"                    
##  [809,] "Bacteria" "Bacteroidetes"       "Bacteroidia"                
##  [810,] "Bacteria" "Patescibacteria"     "Saccharimonadia"            
##  [811,] "Bacteria" "Bacteroidetes"       "Bacteroidia"                
##  [812,] "Bacteria" "Actinobacteria"      "Actinobacteria"             
##  [813,] "Bacteria" "Fusobacteria"        "Fusobacteriia"              
##  [814,] "Bacteria" "Firmicutes"          "Negativicutes"              
##  [815,] "Bacteria" "Bacteroidetes"       "Bacteroidia"                
##  [816,] "Bacteria" "Proteobacteria"      "Gammaproteobacteria"        
##  [817,] "Bacteria" "Bacteroidetes"       "Bacteroidia"                
##  [818,] "Bacteria" "Firmicutes"          "Negativicutes"              
##  [819,] "Bacteria" "Bacteroidetes"       "Bacteroidia"                
##  [820,] "Bacteria" "Patescibacteria"     "Saccharimonadia"            
##  [821,] "Bacteria" "Proteobacteria"      "Gammaproteobacteria"        
##  [822,] "Bacteria" "Fusobacteria"        "Fusobacteriia"              
##  [823,] "Bacteria" "Firmicutes"          "Bacilli"                    
##  [824,] "Bacteria" "Bacteroidetes"       "Bacteroidia"                
##  [825,] "Bacteria" "Fusobacteria"        "Fusobacteriia"              
##  [826,] "Bacteria" "Bacteroidetes"       "Bacteroidia"                
##  [827,] "Bacteria" "Bacteroidetes"       "Bacteroidia"                
##  [828,] "Bacteria" "Proteobacteria"      "Gammaproteobacteria"        
##  [829,] "Bacteria" "Bacteroidetes"       "Bacteroidia"                
##  [830,] "Bacteria" "Proteobacteria"      "Gammaproteobacteria"        
##  [831,] "Bacteria" "Firmicutes"          "Negativicutes"              
##  [832,] "Bacteria" "Bacteroidetes"       "Bacteroidia"                
##  [833,] "Bacteria" "Firmicutes"          "Clostridia"                 
##  [834,] "Bacteria" "Actinobacteria"      "Actinobacteria"             
##  [835,] "Bacteria" "Firmicutes"          "Negativicutes"              
##  [836,] "Bacteria" "Firmicutes"          "Negativicutes"              
##  [837,] "Bacteria" "Firmicutes"          "Bacilli"                    
##  [838,] "Bacteria" "Bacteroidetes"       "Bacteroidia"                
##  [839,] "Bacteria" "Epsilonbacteraeota"  "Campylobacteria"            
##  [840,] "Bacteria" "Patescibacteria"     "Saccharimonadia"            
##  [841,] "Bacteria" "Firmicutes"          "Negativicutes"              
##  [842,] "Bacteria" "Bacteroidetes"       "Bacteroidia"                
##  [843,] "Bacteria" "Firmicutes"          "Clostridia"                 
##  [844,] "Bacteria" "Bacteroidetes"       "Bacteroidia"                
##  [845,] "Bacteria" "Firmicutes"          "Bacilli"                    
##  [846,] "Bacteria" "Actinobacteria"      "Actinobacteria"             
##  [847,] "Bacteria" "Firmicutes"          "Negativicutes"              
##  [848,] "Bacteria" "Actinobacteria"      "Actinobacteria"             
##  [849,] "Bacteria" "Bacteroidetes"       "Bacteroidia"                
##  [850,] "Bacteria" "Proteobacteria"      "Gammaproteobacteria"        
##  [851,] "Bacteria" "Bacteroidetes"       "Bacteroidia"                
##  [852,] "Bacteria" "Firmicutes"          "Bacilli"                    
##  [853,] "Bacteria" "Bacteroidetes"       "Bacteroidia"                
##  [854,] "Bacteria" "Firmicutes"          "Negativicutes"              
##  [855,] "Bacteria" "Bacteroidetes"       "Bacteroidia"                
##  [856,] "Bacteria" "Firmicutes"          "Clostridia"                 
##  [857,] "Bacteria" "Bacteroidetes"       "Bacteroidia"                
##  [858,] "Bacteria" "Firmicutes"          "Bacilli"                    
##  [859,] "Bacteria" "Bacteroidetes"       "Bacteroidia"                
##  [860,] "Bacteria" "Patescibacteria"     "Saccharimonadia"            
##  [861,] "Bacteria" "Firmicutes"          "Clostridia"                 
##  [862,] "Bacteria" "Firmicutes"          "Clostridia"                 
##  [863,] "Bacteria" "Bacteroidetes"       "Bacteroidia"                
##  [864,] "Bacteria" "Bacteroidetes"       "Bacteroidia"                
##  [865,] "Bacteria" "Bacteroidetes"       "Bacteroidia"                
##  [866,] "Bacteria" "Bacteroidetes"       "Bacteroidia"                
##  [867,] "Bacteria" "Actinobacteria"      "Actinobacteria"             
##  [868,] "Bacteria" "Bacteroidetes"       "Bacteroidia"                
##  [869,] "Bacteria" "Firmicutes"          "Negativicutes"              
##  [870,] "Bacteria" "Actinobacteria"      "Actinobacteria"             
##  [871,] "Bacteria" "Actinobacteria"      "Actinobacteria"             
##  [872,] "Bacteria" "Firmicutes"          "Negativicutes"              
##  [873,] "Bacteria" "Bacteroidetes"       "Bacteroidia"                
##  [874,] "Bacteria" "Bacteroidetes"       "Bacteroidia"                
##  [875,] "Bacteria" "Actinobacteria"      "Actinobacteria"             
##  [876,] "Bacteria" "Fusobacteria"        "Fusobacteriia"              
##  [877,] "Bacteria" "Patescibacteria"     "Saccharimonadia"            
##  [878,] "Bacteria" "Firmicutes"          "Negativicutes"              
##  [879,] "Bacteria" "Actinobacteria"      "Actinobacteria"             
##  [880,] "Bacteria" "Firmicutes"          "Negativicutes"              
##  [881,] "Bacteria" "Proteobacteria"      "Gammaproteobacteria"        
##  [882,] "Bacteria" "Firmicutes"          "Clostridia"                 
##  [883,] "Bacteria" "Firmicutes"          "Negativicutes"              
##  [884,] "Bacteria" "Bacteroidetes"       "Bacteroidia"                
##  [885,] "Bacteria" "Bacteroidetes"       "Bacteroidia"                
##  [886,] "Bacteria" "Bacteroidetes"       "Bacteroidia"                
##  [887,] "Bacteria" "Firmicutes"          "Negativicutes"              
##  [888,] "Bacteria" "Actinobacteria"      "Actinobacteria"             
##  [889,] "Bacteria" "Actinobacteria"      "Actinobacteria"             
##  [890,] "Bacteria" "Firmicutes"          "Clostridia"                 
##  [891,] "Bacteria" "Proteobacteria"      "Gammaproteobacteria"        
##  [892,] "Bacteria" "Firmicutes"          "Bacilli"                    
##  [893,] "Bacteria" "Patescibacteria"     "Saccharimonadia"            
##  [894,] "Bacteria" "Bacteroidetes"       "Bacteroidia"                
##  [895,] "Bacteria" "Actinobacteria"      "Actinobacteria"             
##  [896,] "Bacteria" "Bacteroidetes"       "Bacteroidia"                
##  [897,] "Bacteria" "Bacteroidetes"       "Bacteroidia"                
##  [898,] "Bacteria" "Bacteroidetes"       "Bacteroidia"                
##  [899,] "Bacteria" "Patescibacteria"     "Saccharimonadia"            
##  [900,] "Bacteria" "Firmicutes"          NA                           
##  [901,] "Bacteria" "Fusobacteria"        "Fusobacteriia"              
##  [902,] "Bacteria" "Proteobacteria"      "Gammaproteobacteria"        
##  [903,] "Bacteria" "Firmicutes"          "Bacilli"                    
##  [904,] "Bacteria" "Firmicutes"          "Clostridia"                 
##  [905,] "Bacteria" "Actinobacteria"      "Actinobacteria"             
##  [906,] "Bacteria" "Bacteroidetes"       "Bacteroidia"                
##  [907,] "Bacteria" "Firmicutes"          "Bacilli"                    
##  [908,] "Bacteria" "Proteobacteria"      "Gammaproteobacteria"        
##  [909,] "Bacteria" "Firmicutes"          "Negativicutes"              
##  [910,] "Bacteria" "Proteobacteria"      "Gammaproteobacteria"        
##  [911,] "Bacteria" "Firmicutes"          "Negativicutes"              
##  [912,] "Bacteria" "Patescibacteria"     "Saccharimonadia"            
##  [913,] "Bacteria" "Proteobacteria"      "Gammaproteobacteria"        
##  [914,] "Bacteria" "Proteobacteria"      "Gammaproteobacteria"        
##  [915,] "Bacteria" "Actinobacteria"      "Actinobacteria"             
##  [916,] "Bacteria" "Firmicutes"          "Negativicutes"              
##  [917,] "Bacteria" "Bacteroidetes"       "Bacteroidia"                
##  [918,] "Bacteria" "Firmicutes"          "Clostridia"                 
##  [919,] "Bacteria" "Actinobacteria"      "Actinobacteria"             
##  [920,] "Bacteria" "Proteobacteria"      "Gammaproteobacteria"        
##  [921,] "Bacteria" "Epsilonbacteraeota"  "Campylobacteria"            
##  [922,] "Bacteria" "Firmicutes"          "Negativicutes"              
##  [923,] "Bacteria" "Firmicutes"          "Negativicutes"              
##  [924,] "Bacteria" "Actinobacteria"      "Actinobacteria"             
##  [925,] "Bacteria" "Fusobacteria"        "Fusobacteriia"              
##  [926,] "Bacteria" "Proteobacteria"      "Gammaproteobacteria"        
##  [927,] "Bacteria" "Fusobacteria"        "Fusobacteriia"              
##  [928,] "Bacteria" "Fusobacteria"        "Fusobacteriia"              
##  [929,] "Bacteria" "Firmicutes"          "Negativicutes"              
##  [930,] "Bacteria" "Bacteroidetes"       "Bacteroidia"                
##  [931,] "Bacteria" "Fusobacteria"        "Fusobacteriia"              
##  [932,] "Bacteria" "Firmicutes"          "Bacilli"                    
##  [933,] "Bacteria" "Firmicutes"          NA                           
##  [934,] "Bacteria" "Firmicutes"          "Negativicutes"              
##  [935,] "Bacteria" "Bacteroidetes"       "Bacteroidia"                
##  [936,] "Bacteria" "Firmicutes"          "Clostridia"                 
##  [937,] "Bacteria" "Fusobacteria"        "Fusobacteriia"              
##  [938,] "Bacteria" "Firmicutes"          "Erysipelotrichia"           
##  [939,] "Bacteria" "Firmicutes"          "Bacilli"                    
##  [940,] "Bacteria" "Bacteroidetes"       "Bacteroidia"                
##  [941,] "Bacteria" "Firmicutes"          "Bacilli"                    
##  [942,] "Bacteria" "Actinobacteria"      "Actinobacteria"             
##  [943,] "Bacteria" "Epsilonbacteraeota"  "Campylobacteria"            
##  [944,] "Bacteria" "Spirochaetes"        "Spirochaetia"               
##  [945,] "Bacteria" "Firmicutes"          "Bacilli"                    
##  [946,] "Bacteria" "Actinobacteria"      "Actinobacteria"             
##  [947,] "Bacteria" "Firmicutes"          "Bacilli"                    
##  [948,] "Bacteria" "Fusobacteria"        "Fusobacteriia"              
##  [949,] "Bacteria" "Fusobacteria"        "Fusobacteriia"              
##  [950,] "Bacteria" "Firmicutes"          "Clostridia"                 
##  [951,] "Bacteria" "Fusobacteria"        "Fusobacteriia"              
##  [952,] "Bacteria" "Firmicutes"          "Negativicutes"              
##  [953,] "Bacteria" "Firmicutes"          NA                           
##  [954,] "Bacteria" "Spirochaetes"        "Spirochaetia"               
##  [955,] "Bacteria" "Bacteroidetes"       "Bacteroidia"                
##  [956,] "Bacteria" "Actinobacteria"      "Actinobacteria"             
##  [957,] "Bacteria" "Firmicutes"          "Negativicutes"              
##  [958,] "Bacteria" "Patescibacteria"     "Saccharimonadia"            
##  [959,] "Bacteria" "Firmicutes"          "Bacilli"                    
##  [960,] "Bacteria" "Bacteroidetes"       "Bacteroidia"                
##  [961,] "Bacteria" "Firmicutes"          "Negativicutes"              
##  [962,] "Bacteria" "Firmicutes"          "Bacilli"                    
##  [963,] "Bacteria" "Firmicutes"          "Negativicutes"              
##  [964,] "Bacteria" "Proteobacteria"      "Gammaproteobacteria"        
##  [965,] "Bacteria" "Fusobacteria"        "Fusobacteriia"              
##  [966,] "Bacteria" "Firmicutes"          "Clostridia"                 
##  [967,] "Bacteria" "Bacteroidetes"       "Bacteroidia"                
##  [968,] "Bacteria" "Fusobacteria"        "Fusobacteriia"              
##  [969,] "Bacteria" "Firmicutes"          "Clostridia"                 
##  [970,] "Bacteria" "Bacteroidetes"       "Bacteroidia"                
##  [971,] "Bacteria" "Fusobacteria"        "Fusobacteriia"              
##  [972,] "Bacteria" "Firmicutes"          "Negativicutes"              
##  [973,] "Bacteria" "Patescibacteria"     "Saccharimonadia"            
##  [974,] "Bacteria" "Firmicutes"          "Negativicutes"              
##  [975,] "Bacteria" "Bacteroidetes"       "Bacteroidia"                
##  [976,] "Bacteria" "Firmicutes"          "Clostridia"                 
##  [977,] "Bacteria" "Firmicutes"          "Clostridia"                 
##  [978,] "Bacteria" "Firmicutes"          "Negativicutes"              
##  [979,] "Bacteria" "Actinobacteria"      "Actinobacteria"             
##  [980,] "Bacteria" "Actinobacteria"      "Actinobacteria"             
##  [981,] "Bacteria" "Fusobacteria"        "Fusobacteriia"              
##  [982,] "Bacteria" "Bacteroidetes"       "Bacteroidia"                
##  [983,] "Bacteria" "Fusobacteria"        "Fusobacteriia"              
##  [984,] "Bacteria" "Actinobacteria"      "Actinobacteria"             
##  [985,] "Bacteria" "Epsilonbacteraeota"  "Campylobacteria"            
##  [986,] "Bacteria" "Firmicutes"          "Negativicutes"              
##  [987,] "Bacteria" "Actinobacteria"      "Actinobacteria"             
##  [988,] "Bacteria" "Proteobacteria"      "Gammaproteobacteria"        
##  [989,] "Bacteria" "Proteobacteria"      "Gammaproteobacteria"        
##  [990,] "Bacteria" "Firmicutes"          "Negativicutes"              
##  [991,] "Bacteria" "Bacteroidetes"       "Bacteroidia"                
##  [992,] "Bacteria" "Actinobacteria"      "Actinobacteria"             
##  [993,] "Bacteria" "Actinobacteria"      "Actinobacteria"             
##  [994,] "Bacteria" "Firmicutes"          "Negativicutes"              
##  [995,] "Bacteria" "Firmicutes"          "Negativicutes"              
##  [996,] "Bacteria" "Fusobacteria"        "Fusobacteriia"              
##  [997,] "Bacteria" "Fusobacteria"        "Fusobacteriia"              
##  [998,] "Bacteria" "Patescibacteria"     "Saccharimonadia"            
##  [999,] "Bacteria" "Bacteroidetes"       "Bacteroidia"                
## [1000,] "Bacteria" "Firmicutes"          "Negativicutes"              
## [1001,] "Bacteria" "Firmicutes"          "Negativicutes"              
## [1002,] "Bacteria" "Firmicutes"          "Negativicutes"              
## [1003,] "Bacteria" "Proteobacteria"      "Gammaproteobacteria"        
## [1004,] "Bacteria" "Bacteroidetes"       "Bacteroidia"                
## [1005,] "Bacteria" "Bacteroidetes"       "Bacteroidia"                
## [1006,] "Bacteria" "Bacteroidetes"       "Bacteroidia"                
## [1007,] "Bacteria" "Firmicutes"          "Clostridia"                 
## [1008,] "Bacteria" "Bacteroidetes"       "Bacteroidia"                
## [1009,] "Bacteria" "Bacteroidetes"       "Bacteroidia"                
## [1010,] "Bacteria" "Firmicutes"          "Clostridia"                 
## [1011,] "Bacteria" "Firmicutes"          "Negativicutes"              
## [1012,] "Bacteria" "Patescibacteria"     "Saccharimonadia"            
## [1013,] "Bacteria" "Proteobacteria"      "Gammaproteobacteria"        
## [1014,] "Bacteria" "Patescibacteria"     "Saccharimonadia"            
## [1015,] "Bacteria" "Firmicutes"          "Clostridia"                 
## [1016,] "Bacteria" "Proteobacteria"      "Gammaproteobacteria"        
## [1017,] "Bacteria" "Patescibacteria"     "Saccharimonadia"            
## [1018,] "Bacteria" "Bacteroidetes"       "Bacteroidia"                
## [1019,] "Bacteria" "Fusobacteria"        "Fusobacteriia"              
## [1020,] "Bacteria" "Proteobacteria"      "Gammaproteobacteria"        
## [1021,] "Bacteria" "Bacteroidetes"       "Bacteroidia"                
## [1022,] "Bacteria" "Fusobacteria"        "Fusobacteriia"              
## [1023,] "Bacteria" "Spirochaetes"        "Spirochaetia"               
## [1024,] "Bacteria" "Firmicutes"          "Bacilli"                    
## [1025,] "Bacteria" "Firmicutes"          "Bacilli"                    
## [1026,] "Bacteria" "Actinobacteria"      "Actinobacteria"             
## [1027,] "Bacteria" "Fusobacteria"        "Fusobacteriia"              
## [1028,] "Bacteria" "Bacteroidetes"       "Bacteroidia"                
## [1029,] "Bacteria" "Bacteroidetes"       "Bacteroidia"                
## [1030,] "Bacteria" "Proteobacteria"      "Gammaproteobacteria"        
## [1031,] "Bacteria" "Fusobacteria"        "Fusobacteriia"              
## [1032,] "Bacteria" "Fusobacteria"        "Fusobacteriia"              
## [1033,] "Bacteria" "Fusobacteria"        "Fusobacteriia"              
## [1034,] "Bacteria" "Firmicutes"          "Negativicutes"              
## [1035,] "Bacteria" "Bacteroidetes"       "Bacteroidia"                
## [1036,] "Bacteria" "Firmicutes"          "Negativicutes"              
## [1037,] "Bacteria" "Firmicutes"          "Negativicutes"              
## [1038,] "Bacteria" "Firmicutes"          "Clostridia"                 
## [1039,] "Bacteria" "Bacteroidetes"       "Bacteroidia"                
## [1040,] "Bacteria" "Firmicutes"          "Negativicutes"              
## [1041,] "Bacteria" "Bacteroidetes"       "Bacteroidia"                
## [1042,] "Bacteria" "Firmicutes"          "Clostridia"                 
## [1043,] "Bacteria" "Fusobacteria"        "Fusobacteriia"              
## [1044,] "Bacteria" "Firmicutes"          "Negativicutes"              
## [1045,] "Bacteria" "Actinobacteria"      "Actinobacteria"             
## [1046,] "Bacteria" "Bacteroidetes"       "Bacteroidia"                
## [1047,] "Bacteria" "Firmicutes"          "Negativicutes"              
## [1048,] "Bacteria" "Bacteroidetes"       "Bacteroidia"                
## [1049,] "Bacteria" "Firmicutes"          "Negativicutes"              
## [1050,] "Bacteria" "Fusobacteria"        "Fusobacteriia"              
## [1051,] "Bacteria" "Firmicutes"          "Clostridia"                 
## [1052,] "Bacteria" "Firmicutes"          "Bacilli"                    
## [1053,] "Bacteria" "Bacteroidetes"       "Bacteroidia"                
## [1054,] "Bacteria" "Proteobacteria"      "Gammaproteobacteria"        
## [1055,] "Bacteria" "Bacteroidetes"       "Bacteroidia"                
## [1056,] "Bacteria" "Firmicutes"          "Bacilli"                    
## [1057,] "Bacteria" "Proteobacteria"      "Gammaproteobacteria"        
## [1058,] "Bacteria" "Bacteroidetes"       "Bacteroidia"                
## [1059,] "Bacteria" "Firmicutes"          "Clostridia"                 
## [1060,] "Bacteria" "Bacteroidetes"       "Bacteroidia"                
## [1061,] "Bacteria" "Firmicutes"          "Negativicutes"              
## [1062,] "Bacteria" "Firmicutes"          "Bacilli"                    
## [1063,] "Bacteria" "Bacteroidetes"       "Bacteroidia"                
## [1064,] "Bacteria" "Fusobacteria"        "Fusobacteriia"              
## [1065,] "Bacteria" "Firmicutes"          "Negativicutes"              
## [1066,] "Bacteria" "Fusobacteria"        "Fusobacteriia"              
## [1067,] "Bacteria" "Firmicutes"          "Negativicutes"              
## [1068,] "Bacteria" "Proteobacteria"      "Gammaproteobacteria"        
## [1069,] "Bacteria" "Fusobacteria"        "Fusobacteriia"              
## [1070,] "Bacteria" "Firmicutes"          "Negativicutes"              
## [1071,] "Bacteria" "Proteobacteria"      "Gammaproteobacteria"        
## [1072,] "Bacteria" "Bacteroidetes"       "Bacteroidia"                
## [1073,] "Bacteria" "Bacteroidetes"       "Bacteroidia"                
## [1074,] "Bacteria" "Actinobacteria"      "Coriobacteriia"             
## [1075,] "Bacteria" "Bacteroidetes"       "Bacteroidia"                
## [1076,] "Bacteria" "Proteobacteria"      "Gammaproteobacteria"        
## [1077,] "Bacteria" NA                    NA                           
## [1078,] "Bacteria" "Bacteroidetes"       "Bacteroidia"                
## [1079,] "Bacteria" "Firmicutes"          "Negativicutes"              
## [1080,] "Bacteria" "Firmicutes"          "Clostridia"                 
## [1081,] "Bacteria" "Bacteroidetes"       "Bacteroidia"                
## [1082,] "Bacteria" "Patescibacteria"     "Saccharimonadia"            
## [1083,] "Bacteria" "Bacteroidetes"       "Bacteroidia"                
## [1084,] "Bacteria" "Firmicutes"          "Negativicutes"              
## [1085,] "Bacteria" "Firmicutes"          "Negativicutes"              
## [1086,] "Bacteria" "Firmicutes"          "Bacilli"                    
## [1087,] "Bacteria" "Firmicutes"          "Bacilli"                    
## [1088,] "Bacteria" "Bacteroidetes"       "Bacteroidia"                
## [1089,] "Bacteria" "Firmicutes"          "Negativicutes"              
## [1090,] "Bacteria" "Bacteroidetes"       "Bacteroidia"                
## [1091,] "Bacteria" "Actinobacteria"      "Coriobacteriia"             
## [1092,] "Bacteria" "Bacteroidetes"       "Bacteroidia"                
## [1093,] "Bacteria" "Bacteroidetes"       "Bacteroidia"                
## [1094,] "Bacteria" "Bacteroidetes"       "Bacteroidia"                
## [1095,] "Bacteria" "Bacteroidetes"       "Bacteroidia"                
## [1096,] "Bacteria" "Firmicutes"          "Negativicutes"              
## [1097,] "Bacteria" "Bacteroidetes"       "Bacteroidia"                
## [1098,] "Bacteria" "Firmicutes"          "Bacilli"                    
## [1099,] "Bacteria" "Actinobacteria"      "Actinobacteria"             
## [1100,] "Bacteria" "Bacteroidetes"       "Bacteroidia"                
## [1101,] "Bacteria" "Patescibacteria"     "Saccharimonadia"            
## [1102,] "Bacteria" "Firmicutes"          "Negativicutes"              
## [1103,] "Bacteria" "Firmicutes"          "Negativicutes"              
## [1104,] "Bacteria" "Bacteroidetes"       "Bacteroidia"                
## [1105,] "Bacteria" "Bacteroidetes"       "Bacteroidia"                
## [1106,] "Bacteria" "Firmicutes"          "Negativicutes"              
## [1107,] "Bacteria" "Bacteroidetes"       "Bacteroidia"                
## [1108,] "Bacteria" "Fusobacteria"        "Fusobacteriia"              
## [1109,] "Bacteria" "Bacteroidetes"       "Bacteroidia"                
## [1110,] "Bacteria" "Firmicutes"          "Negativicutes"              
## [1111,] "Bacteria" "Bacteroidetes"       "Bacteroidia"                
## [1112,] "Bacteria" "Fusobacteria"        "Fusobacteriia"              
## [1113,] "Bacteria" "Bacteroidetes"       "Bacteroidia"                
## [1114,] "Bacteria" "Firmicutes"          "Clostridia"                 
## [1115,] "Bacteria" "Patescibacteria"     "Saccharimonadia"            
## [1116,] "Bacteria" "Patescibacteria"     "Saccharimonadia"            
## [1117,] "Bacteria" "Bacteroidetes"       "Bacteroidia"                
## [1118,] "Bacteria" "Bacteroidetes"       "Bacteroidia"                
## [1119,] "Bacteria" "Actinobacteria"      "Actinobacteria"             
## [1120,] "Bacteria" "Patescibacteria"     "Saccharimonadia"            
## [1121,] "Bacteria" "Firmicutes"          "Negativicutes"              
## [1122,] "Bacteria" "Firmicutes"          "Negativicutes"              
## [1123,] "Bacteria" "Bacteroidetes"       "Bacteroidia"                
## [1124,] "Bacteria" "Epsilonbacteraeota"  "Campylobacteria"            
## [1125,] "Bacteria" "Actinobacteria"      "Actinobacteria"             
## [1126,] "Bacteria" "Cyanobacteria"       "Oxyphotobacteria"           
## [1127,] "Bacteria" "Bacteroidetes"       "Bacteroidia"                
## [1128,] "Bacteria" "Firmicutes"          NA                           
## [1129,] "Bacteria" "Fusobacteria"        "Fusobacteriia"              
## [1130,] "Bacteria" "Proteobacteria"      "Gammaproteobacteria"        
## [1131,] "Bacteria" "Firmicutes"          "Clostridia"                 
## [1132,] "Bacteria" "Bacteroidetes"       "Bacteroidia"                
## [1133,] "Bacteria" "Bacteroidetes"       "Bacteroidia"                
## [1134,] "Bacteria" "Firmicutes"          "Negativicutes"              
## [1135,] "Bacteria" "Proteobacteria"      "Gammaproteobacteria"        
## [1136,] "Bacteria" "Proteobacteria"      "Gammaproteobacteria"        
## [1137,] "Bacteria" "Bacteroidetes"       "Bacteroidia"                
## [1138,] "Bacteria" "Bacteroidetes"       "Bacteroidia"                
## [1139,] "Bacteria" "Proteobacteria"      "Gammaproteobacteria"        
## [1140,] "Bacteria" "Firmicutes"          "Bacilli"                    
## [1141,] "Bacteria" "Proteobacteria"      "Gammaproteobacteria"        
## [1142,] "Bacteria" "Bacteroidetes"       "Bacteroidia"                
## [1143,] "Bacteria" "Firmicutes"          "Clostridia"                 
## [1144,] "Bacteria" "Firmicutes"          "Bacilli"                    
## [1145,] "Bacteria" "Actinobacteria"      "Actinobacteria"             
## [1146,] "Bacteria" "Firmicutes"          "Bacilli"                    
## [1147,] "Bacteria" "Bacteroidetes"       "Bacteroidia"                
## [1148,] "Bacteria" "Firmicutes"          "Negativicutes"              
## [1149,] "Bacteria" "Firmicutes"          "Clostridia"                 
## [1150,] "Bacteria" "Actinobacteria"      "Actinobacteria"             
## [1151,] "Bacteria" "Fusobacteria"        "Fusobacteriia"              
## [1152,] "Bacteria" "Proteobacteria"      "Gammaproteobacteria"        
## [1153,] "Bacteria" "Firmicutes"          "Bacilli"                    
## [1154,] "Bacteria" "Proteobacteria"      "Gammaproteobacteria"        
## [1155,] "Bacteria" "Bacteroidetes"       "Bacteroidia"                
## [1156,] "Bacteria" "Proteobacteria"      "Gammaproteobacteria"        
## [1157,] "Bacteria" "Firmicutes"          "Bacilli"                    
## [1158,] "Bacteria" "Fusobacteria"        "Fusobacteriia"              
## [1159,] "Bacteria" "Firmicutes"          "Clostridia"                 
## [1160,] "Bacteria" "Bacteroidetes"       "Bacteroidia"                
## [1161,] "Bacteria" "Bacteroidetes"       "Bacteroidia"                
## [1162,] "Bacteria" "Firmicutes"          "Bacilli"                    
## [1163,] "Bacteria" "Tenericutes"         "Mollicutes"                 
## [1164,] "Bacteria" "Proteobacteria"      "Gammaproteobacteria"        
## [1165,] "Bacteria" "Patescibacteria"     "Gracilibacteria"            
## [1166,] "Bacteria" "Firmicutes"          "Negativicutes"              
## [1167,] "Bacteria" "Proteobacteria"      "Gammaproteobacteria"        
## [1168,] "Bacteria" "Firmicutes"          "Negativicutes"              
## [1169,] "Bacteria" "Firmicutes"          "Negativicutes"              
## [1170,] "Bacteria" "Firmicutes"          "Bacilli"                    
## [1171,] "Bacteria" "Bacteroidetes"       "Bacteroidia"                
## [1172,] "Bacteria" "Patescibacteria"     "Saccharimonadia"            
## [1173,] "Bacteria" "Bacteroidetes"       "Bacteroidia"                
## [1174,] "Bacteria" "Firmicutes"          "Negativicutes"              
## [1175,] "Bacteria" "Firmicutes"          "Negativicutes"              
## [1176,] "Bacteria" "Firmicutes"          "Negativicutes"              
## [1177,] "Bacteria" "Bacteroidetes"       "Bacteroidia"                
## [1178,] "Bacteria" "Fusobacteria"        "Fusobacteriia"              
## [1179,] "Bacteria" "Proteobacteria"      "Gammaproteobacteria"        
## [1180,] "Bacteria" "Bacteroidetes"       "Bacteroidia"                
## [1181,] "Bacteria" "Firmicutes"          "Clostridia"                 
## [1182,] "Bacteria" "Firmicutes"          "Clostridia"                 
## [1183,] "Bacteria" "Firmicutes"          "Bacilli"                    
## [1184,] "Bacteria" "Bacteroidetes"       "Bacteroidia"                
## [1185,] "Bacteria" "Proteobacteria"      "Gammaproteobacteria"        
## [1186,] "Bacteria" "Proteobacteria"      "Gammaproteobacteria"        
## [1187,] "Bacteria" "Bacteroidetes"       "Bacteroidia"                
## [1188,] "Bacteria" "Firmicutes"          "Negativicutes"              
## [1189,] "Bacteria" "Bacteroidetes"       "Bacteroidia"                
## [1190,] "Bacteria" "Bacteroidetes"       "Bacteroidia"                
## [1191,] "Bacteria" "Actinobacteria"      "Actinobacteria"             
## [1192,] "Bacteria" "Fusobacteria"        "Fusobacteriia"              
## [1193,] "Bacteria" "Actinobacteria"      "Actinobacteria"             
## [1194,] "Bacteria" "Proteobacteria"      "Gammaproteobacteria"        
## [1195,] "Bacteria" "Bacteroidetes"       "Bacteroidia"                
## [1196,] "Bacteria" "Bacteroidetes"       "Bacteroidia"                
## [1197,] "Bacteria" "Firmicutes"          "Negativicutes"              
## [1198,] "Bacteria" "Firmicutes"          "Clostridia"                 
## [1199,] "Bacteria" "Firmicutes"          "Bacilli"                    
## [1200,] "Bacteria" "Bacteroidetes"       "Bacteroidia"                
## [1201,] "Bacteria" "Firmicutes"          "Negativicutes"              
## [1202,] "Bacteria" "Fusobacteria"        "Fusobacteriia"              
## [1203,] "Bacteria" "Actinobacteria"      "Actinobacteria"             
## [1204,] "Bacteria" "Bacteroidetes"       "Bacteroidia"                
## [1205,] "Bacteria" "Synergistetes"       "Synergistia"                
## [1206,] "Bacteria" "Epsilonbacteraeota"  "Campylobacteria"            
## [1207,] "Bacteria" "Proteobacteria"      "Gammaproteobacteria"        
## [1208,] "Bacteria" "Firmicutes"          "Negativicutes"              
## [1209,] "Bacteria" "Proteobacteria"      "Alphaproteobacteria"        
## [1210,] "Bacteria" "Firmicutes"          "Negativicutes"              
## [1211,] "Bacteria" "Fusobacteria"        "Fusobacteriia"              
## [1212,] "Bacteria" "Patescibacteria"     "Saccharimonadia"            
## [1213,] "Bacteria" "Firmicutes"          "Negativicutes"              
## [1214,] "Bacteria" "Fusobacteria"        "Fusobacteriia"              
## [1215,] "Bacteria" "Patescibacteria"     "Saccharimonadia"            
## [1216,] "Bacteria" "Firmicutes"          "Clostridia"                 
## [1217,] "Bacteria" "Spirochaetes"        "Spirochaetia"               
## [1218,] "Bacteria" "Bacteroidetes"       "Bacteroidia"                
## [1219,] "Bacteria" "Firmicutes"          "Clostridia"                 
## [1220,] "Bacteria" "Firmicutes"          "Clostridia"                 
## [1221,] "Bacteria" "Firmicutes"          "Negativicutes"              
## [1222,] "Bacteria" "Actinobacteria"      "Actinobacteria"             
## [1223,] "Bacteria" "Bacteroidetes"       "Bacteroidia"                
## [1224,] "Bacteria" "Firmicutes"          NA                           
## [1225,] "Bacteria" "Firmicutes"          "Negativicutes"              
## [1226,] "Bacteria" "Firmicutes"          NA                           
## [1227,] "Bacteria" "Firmicutes"          "Negativicutes"              
## [1228,] "Bacteria" "Proteobacteria"      "Gammaproteobacteria"        
## [1229,] "Bacteria" "Bacteroidetes"       "Bacteroidia"                
## [1230,] "Bacteria" "Firmicutes"          "Negativicutes"              
## [1231,] "Bacteria" "Bacteroidetes"       "Bacteroidia"                
## [1232,] "Bacteria" "Bacteroidetes"       "Bacteroidia"                
## [1233,] "Bacteria" "Bacteroidetes"       "Bacteroidia"                
## [1234,] "Bacteria" "Firmicutes"          "Negativicutes"              
## [1235,] "Bacteria" "Bacteroidetes"       "Bacteroidia"                
## [1236,] "Bacteria" "Firmicutes"          "Clostridia"                 
## [1237,] "Bacteria" "Fusobacteria"        "Fusobacteriia"              
## [1238,] "Bacteria" "Actinobacteria"      "Actinobacteria"             
## [1239,] "Bacteria" "Firmicutes"          "Negativicutes"              
## [1240,] "Bacteria" "Patescibacteria"     "Saccharimonadia"            
## [1241,] "Bacteria" "Bacteroidetes"       "Bacteroidia"                
## [1242,] "Bacteria" "Bacteroidetes"       "Bacteroidia"                
## [1243,] "Bacteria" "Firmicutes"          "Clostridia"                 
## [1244,] "Bacteria" "Firmicutes"          "Negativicutes"              
## [1245,] "Bacteria" "Firmicutes"          "Negativicutes"              
## [1246,] "Bacteria" "Fusobacteria"        "Fusobacteriia"              
## [1247,] "Bacteria" "Firmicutes"          NA                           
## [1248,] "Bacteria" "Bacteroidetes"       "Bacteroidia"                
## [1249,] "Bacteria" "Firmicutes"          "Negativicutes"              
## [1250,] "Bacteria" "Actinobacteria"      "Actinobacteria"             
## [1251,] "Bacteria" "Fusobacteria"        "Fusobacteriia"              
## [1252,] "Bacteria" "Bacteroidetes"       "Bacteroidia"                
## [1253,] "Bacteria" "Bacteroidetes"       "Bacteroidia"                
## [1254,] "Bacteria" "Actinobacteria"      "Actinobacteria"             
## [1255,] "Bacteria" "Bacteroidetes"       "Bacteroidia"                
## [1256,] "Bacteria" "Bacteroidetes"       "Bacteroidia"                
## [1257,] "Bacteria" "Firmicutes"          "Negativicutes"              
## [1258,] "Bacteria" "Bacteroidetes"       "Bacteroidia"                
## [1259,] "Bacteria" "Firmicutes"          "Bacilli"                    
## [1260,] "Bacteria" "Bacteroidetes"       "Bacteroidia"                
## [1261,] "Bacteria" "Proteobacteria"      "Gammaproteobacteria"        
## [1262,] "Bacteria" "Firmicutes"          "Negativicutes"              
## [1263,] "Bacteria" "Patescibacteria"     "Saccharimonadia"            
## [1264,] "Bacteria" "Actinobacteria"      "Actinobacteria"             
## [1265,] "Bacteria" "Actinobacteria"      "Actinobacteria"             
## [1266,] "Bacteria" "Firmicutes"          "Negativicutes"              
## [1267,] "Bacteria" "Actinobacteria"      "Coriobacteriia"             
## [1268,] "Bacteria" "Bacteroidetes"       "Bacteroidia"                
## [1269,] "Bacteria" "Firmicutes"          "Clostridia"                 
## [1270,] "Bacteria" "Bacteroidetes"       "Bacteroidia"                
## [1271,] "Bacteria" "Firmicutes"          "Negativicutes"              
## [1272,] "Bacteria" "Firmicutes"          "Negativicutes"              
## [1273,] "Bacteria" "Fusobacteria"        "Fusobacteriia"              
## [1274,] "Bacteria" "Bacteroidetes"       "Bacteroidia"                
## [1275,] "Bacteria" "Spirochaetes"        "Spirochaetia"               
## [1276,] "Bacteria" "Bacteroidetes"       "Bacteroidia"                
## [1277,] "Bacteria" "Firmicutes"          "Negativicutes"              
## [1278,] "Bacteria" "Firmicutes"          "Negativicutes"              
## [1279,] "Bacteria" "Firmicutes"          "Negativicutes"              
## [1280,] "Bacteria" "Proteobacteria"      "Gammaproteobacteria"        
## [1281,] "Bacteria" "Proteobacteria"      "Gammaproteobacteria"        
## [1282,] "Bacteria" "Bacteroidetes"       "Bacteroidia"                
## [1283,] "Bacteria" "Patescibacteria"     "Saccharimonadia"            
## [1284,] "Bacteria" "Firmicutes"          "Negativicutes"              
## [1285,] "Bacteria" "Fusobacteria"        "Fusobacteriia"              
## [1286,] "Bacteria" "Actinobacteria"      "Actinobacteria"             
## [1287,] "Bacteria" "Firmicutes"          "Negativicutes"              
## [1288,] "Bacteria" "Firmicutes"          "Clostridia"                 
## [1289,] "Bacteria" "Tenericutes"         "Mollicutes"                 
## [1290,] "Bacteria" "Fusobacteria"        "Fusobacteriia"              
## [1291,] "Bacteria" "Bacteroidetes"       "Bacteroidia"                
## [1292,] "Bacteria" "Firmicutes"          "Clostridia"                 
## [1293,] "Bacteria" "Firmicutes"          "Clostridia"                 
## [1294,] "Bacteria" "Fusobacteria"        "Fusobacteriia"              
## [1295,] "Bacteria" "Proteobacteria"      "Gammaproteobacteria"        
## [1296,] "Bacteria" "Bacteroidetes"       "Bacteroidia"                
## [1297,] "Bacteria" "Firmicutes"          "Clostridia"                 
## [1298,] "Bacteria" "Bacteroidetes"       "Bacteroidia"                
## [1299,] "Bacteria" "Bacteroidetes"       "Bacteroidia"                
## [1300,] "Bacteria" "Bacteroidetes"       "Bacteroidia"                
## [1301,] "Bacteria" "Firmicutes"          "Clostridia"                 
## [1302,] "Bacteria" "Bacteroidetes"       "Bacteroidia"                
## [1303,] "Bacteria" "Firmicutes"          "Negativicutes"              
## [1304,] "Bacteria" "Bacteroidetes"       "Bacteroidia"                
## [1305,] "Bacteria" "Actinobacteria"      "Actinobacteria"             
## [1306,] "Bacteria" "Firmicutes"          "Bacilli"                    
## [1307,] "Bacteria" "Proteobacteria"      "Gammaproteobacteria"        
## [1308,] "Bacteria" "Proteobacteria"      "Gammaproteobacteria"        
## [1309,] "Bacteria" "Spirochaetes"        "Spirochaetia"               
## [1310,] "Bacteria" "Proteobacteria"      "Gammaproteobacteria"        
## [1311,] "Bacteria" "Proteobacteria"      "Gammaproteobacteria"        
## [1312,] "Bacteria" "Firmicutes"          "Clostridia"                 
## [1313,] "Bacteria" "Bacteroidetes"       "Bacteroidia"                
## [1314,] "Bacteria" "Firmicutes"          "Clostridia"                 
## [1315,] "Bacteria" "Bacteroidetes"       "Bacteroidia"                
## [1316,] "Bacteria" "Bacteroidetes"       "Bacteroidia"                
## [1317,] "Bacteria" "Epsilonbacteraeota"  "Campylobacteria"            
## [1318,] "Bacteria" "Firmicutes"          "Negativicutes"              
## [1319,] "Bacteria" "Firmicutes"          "Negativicutes"              
## [1320,] "Bacteria" "Firmicutes"          "Bacilli"                    
## [1321,] "Bacteria" "Firmicutes"          "Negativicutes"              
## [1322,] "Bacteria" "Epsilonbacteraeota"  "Campylobacteria"            
## [1323,] "Bacteria" "Firmicutes"          "Negativicutes"              
## [1324,] "Bacteria" "Firmicutes"          "Bacilli"                    
## [1325,] "Bacteria" "Bacteroidetes"       "Bacteroidia"                
## [1326,] "Bacteria" "Proteobacteria"      "Gammaproteobacteria"        
## [1327,] "Bacteria" "Firmicutes"          "Bacilli"                    
## [1328,] "Bacteria" "Firmicutes"          "Negativicutes"              
## [1329,] "Bacteria" "Bacteroidetes"       "Bacteroidia"                
## [1330,] "Bacteria" "Bacteroidetes"       "Bacteroidia"                
## [1331,] "Bacteria" "Firmicutes"          "Clostridia"                 
## [1332,] "Bacteria" "Bacteroidetes"       "Bacteroidia"                
## [1333,] "Bacteria" "Firmicutes"          "Negativicutes"              
## [1334,] "Bacteria" "Bacteroidetes"       "Bacteroidia"                
## [1335,] "Bacteria" "Fusobacteria"        "Fusobacteriia"              
## [1336,] "Bacteria" "Fusobacteria"        "Fusobacteriia"              
## [1337,] "Bacteria" "Firmicutes"          "Negativicutes"              
## [1338,] "Bacteria" "Firmicutes"          "Clostridia"                 
## [1339,] "Bacteria" "Proteobacteria"      "Gammaproteobacteria"        
## [1340,] "Bacteria" "Firmicutes"          "Negativicutes"              
## [1341,] "Bacteria" "Firmicutes"          NA                           
## [1342,] "Bacteria" "Fusobacteria"        "Fusobacteriia"              
## [1343,] "Bacteria" "Bacteroidetes"       "Bacteroidia"                
## [1344,] "Bacteria" "Patescibacteria"     "Saccharimonadia"            
## [1345,] "Bacteria" "Bacteroidetes"       "Bacteroidia"                
## [1346,] "Bacteria" "Patescibacteria"     "Saccharimonadia"            
## [1347,] "Bacteria" "Bacteroidetes"       "Bacteroidia"                
## [1348,] "Bacteria" "Bacteroidetes"       "Bacteroidia"                
## [1349,] "Bacteria" "Proteobacteria"      "Gammaproteobacteria"        
## [1350,] "Bacteria" "Actinobacteria"      "Actinobacteria"             
## [1351,] "Bacteria" "Actinobacteria"      "Actinobacteria"             
## [1352,] "Bacteria" "Actinobacteria"      "Actinobacteria"             
## [1353,] "Bacteria" "Bacteroidetes"       "Bacteroidia"                
## [1354,] "Bacteria" "Firmicutes"          "Negativicutes"              
## [1355,] "Bacteria" "Epsilonbacteraeota"  "Campylobacteria"            
## [1356,] "Bacteria" "Firmicutes"          NA                           
## [1357,] "Bacteria" "Bacteroidetes"       "Bacteroidia"                
## [1358,] "Bacteria" "Tenericutes"         "Mollicutes"                 
## [1359,] "Bacteria" "Firmicutes"          "Negativicutes"              
## [1360,] "Bacteria" "Bacteroidetes"       "Bacteroidia"                
## [1361,] "Bacteria" "Proteobacteria"      "Gammaproteobacteria"        
## [1362,] "Bacteria" "Firmicutes"          NA                           
## [1363,] "Bacteria" "Firmicutes"          NA                           
## [1364,] "Bacteria" "Actinobacteria"      "Actinobacteria"             
## [1365,] "Bacteria" "Bacteroidetes"       "Bacteroidia"                
## [1366,] "Bacteria" "Bacteroidetes"       "Bacteroidia"                
## [1367,] "Bacteria" "Bacteroidetes"       "Bacteroidia"                
## [1368,] "Bacteria" "Proteobacteria"      "Gammaproteobacteria"        
## [1369,] "Bacteria" "Fusobacteria"        "Fusobacteriia"              
## [1370,] "Bacteria" "Firmicutes"          "Clostridia"                 
## [1371,] "Bacteria" "Bacteroidetes"       "Bacteroidia"                
## [1372,] "Bacteria" "Fusobacteria"        "Fusobacteriia"              
## [1373,] "Bacteria" "Firmicutes"          "Bacilli"                    
## [1374,] "Bacteria" "Proteobacteria"      "Gammaproteobacteria"        
## [1375,] "Bacteria" "Fusobacteria"        "Fusobacteriia"              
## [1376,] "Bacteria" "Proteobacteria"      "Gammaproteobacteria"        
## [1377,] "Bacteria" "Firmicutes"          "Negativicutes"              
## [1378,] "Bacteria" "Bacteroidetes"       "Bacteroidia"                
## [1379,] "Bacteria" "Bacteroidetes"       "Bacteroidia"                
## [1380,] "Bacteria" "Patescibacteria"     "Gracilibacteria"            
## [1381,] "Bacteria" "Firmicutes"          "Clostridia"                 
## [1382,] "Bacteria" "Bacteroidetes"       "Bacteroidia"                
## [1383,] "Bacteria" "Firmicutes"          "Negativicutes"              
## [1384,] "Bacteria" "Firmicutes"          "Bacilli"                    
## [1385,] "Bacteria" "Firmicutes"          "Negativicutes"              
## [1386,] "Bacteria" "Firmicutes"          "Bacilli"                    
## [1387,] "Bacteria" "Fusobacteria"        "Fusobacteriia"              
## [1388,] "Bacteria" "Bacteroidetes"       "Bacteroidia"                
## [1389,] "Bacteria" "Firmicutes"          "Negativicutes"              
## [1390,] "Bacteria" "Bacteroidetes"       "Bacteroidia"                
## [1391,] "Bacteria" "Patescibacteria"     "Saccharimonadia"            
## [1392,] "Bacteria" "Bacteroidetes"       "Bacteroidia"                
## [1393,] "Bacteria" "Fusobacteria"        "Fusobacteriia"              
## [1394,] "Bacteria" "Bacteroidetes"       "Bacteroidia"                
## [1395,] "Bacteria" "Fusobacteria"        "Fusobacteriia"              
## [1396,] "Bacteria" "Firmicutes"          NA                           
## [1397,] "Bacteria" "Bacteroidetes"       "Bacteroidia"                
## [1398,] "Bacteria" "Bacteroidetes"       "Bacteroidia"                
## [1399,] "Bacteria" "Firmicutes"          "Bacilli"                    
## [1400,] "Bacteria" "Bacteroidetes"       "Bacteroidia"                
## [1401,] "Bacteria" "Fusobacteria"        "Fusobacteriia"              
## [1402,] "Bacteria" "Bacteroidetes"       "Bacteroidia"                
## [1403,] "Bacteria" "Fusobacteria"        "Fusobacteriia"              
## [1404,] "Bacteria" "Bacteroidetes"       "Bacteroidia"                
## [1405,] "Bacteria" "Epsilonbacteraeota"  "Campylobacteria"            
## [1406,] "Bacteria" "Proteobacteria"      "Gammaproteobacteria"        
## [1407,] "Bacteria" "Fusobacteria"        "Fusobacteriia"              
## [1408,] "Bacteria" "Firmicutes"          "Negativicutes"              
## [1409,] "Bacteria" "Proteobacteria"      "Gammaproteobacteria"        
## [1410,] "Bacteria" "Fusobacteria"        "Fusobacteriia"              
## [1411,] "Bacteria" "Firmicutes"          "Clostridia"                 
## [1412,] "Bacteria" "Firmicutes"          "Bacilli"                    
## [1413,] "Bacteria" "Proteobacteria"      "Gammaproteobacteria"        
## [1414,] "Bacteria" "Proteobacteria"      "Gammaproteobacteria"        
## [1415,] "Bacteria" "Bacteroidetes"       "Bacteroidia"                
## [1416,] "Bacteria" "Actinobacteria"      "Actinobacteria"             
## [1417,] "Bacteria" "Spirochaetes"        "Spirochaetia"               
## [1418,] "Bacteria" "Firmicutes"          "Negativicutes"              
## [1419,] "Bacteria" "Fusobacteria"        "Fusobacteriia"              
## [1420,] "Bacteria" "Fusobacteria"        "Fusobacteriia"              
## [1421,] "Bacteria" "Epsilonbacteraeota"  "Campylobacteria"            
## [1422,] "Bacteria" "Firmicutes"          "Negativicutes"              
## [1423,] "Bacteria" "Fusobacteria"        "Fusobacteriia"              
## [1424,] "Bacteria" "Bacteroidetes"       "Bacteroidia"                
## [1425,] "Bacteria" "Firmicutes"          "Negativicutes"              
## [1426,] "Bacteria" "Epsilonbacteraeota"  "Campylobacteria"            
## [1427,] "Bacteria" "Bacteroidetes"       "Bacteroidia"                
## [1428,] "Bacteria" "Actinobacteria"      "Actinobacteria"             
## [1429,] "Bacteria" "Bacteroidetes"       "Bacteroidia"                
## [1430,] "Bacteria" "Bacteroidetes"       "Bacteroidia"                
## [1431,] "Bacteria" "Firmicutes"          "Negativicutes"              
## [1432,] "Bacteria" "Spirochaetes"        "Spirochaetia"               
## [1433,] "Bacteria" "Bacteroidetes"       "Bacteroidia"                
## [1434,] "Bacteria" "Firmicutes"          "Bacilli"                    
## [1435,] "Bacteria" "Proteobacteria"      "Gammaproteobacteria"        
## [1436,] "Bacteria" "Patescibacteria"     "Saccharimonadia"            
## [1437,] "Bacteria" "Firmicutes"          "Clostridia"                 
## [1438,] "Bacteria" "Firmicutes"          "Negativicutes"              
## [1439,] "Bacteria" "Proteobacteria"      "Gammaproteobacteria"        
## [1440,] "Bacteria" "Actinobacteria"      "Actinobacteria"             
## [1441,] "Bacteria" "Firmicutes"          "Negativicutes"              
## [1442,] "Bacteria" "Firmicutes"          "Negativicutes"              
## [1443,] "Bacteria" "Bacteroidetes"       "Bacteroidia"                
## [1444,] "Bacteria" "Bacteroidetes"       "Bacteroidia"                
## [1445,] "Bacteria" "Firmicutes"          "Clostridia"                 
## [1446,] "Bacteria" "Proteobacteria"      "Gammaproteobacteria"        
## [1447,] "Bacteria" NA                    NA                           
## [1448,] "Bacteria" "Proteobacteria"      "Gammaproteobacteria"        
## [1449,] "Bacteria" "Bacteroidetes"       "Bacteroidia"                
## [1450,] "Bacteria" "Firmicutes"          "Bacilli"                    
## [1451,] "Bacteria" "Firmicutes"          "Bacilli"                    
## [1452,] "Bacteria" "Actinobacteria"      "Actinobacteria"             
## [1453,] "Bacteria" "Bacteroidetes"       "Bacteroidia"                
## [1454,] "Bacteria" "Firmicutes"          "Negativicutes"              
## [1455,] "Bacteria" "Firmicutes"          "Negativicutes"              
## [1456,] "Bacteria" "Proteobacteria"      "Gammaproteobacteria"        
## [1457,] "Bacteria" "Bacteroidetes"       "Bacteroidia"                
## [1458,] "Bacteria" "Patescibacteria"     "Saccharimonadia"            
## [1459,] "Bacteria" "Firmicutes"          "Negativicutes"              
## [1460,] "Bacteria" "Fusobacteria"        "Fusobacteriia"              
## [1461,] "Bacteria" "Bacteroidetes"       "Bacteroidia"                
## [1462,] "Bacteria" "Bacteroidetes"       "Bacteroidia"                
## [1463,] "Bacteria" "Spirochaetes"        "Spirochaetia"               
## [1464,] "Bacteria" "Firmicutes"          "Clostridia"                 
## [1465,] "Bacteria" "Fusobacteria"        "Fusobacteriia"              
## [1466,] "Bacteria" "Actinobacteria"      "Actinobacteria"             
## [1467,] "Bacteria" "Bacteroidetes"       "Bacteroidia"                
## [1468,] "Bacteria" "Bacteroidetes"       "Bacteroidia"                
## [1469,] "Bacteria" "Firmicutes"          "Bacilli"                    
## [1470,] "Bacteria" "Bacteroidetes"       "Bacteroidia"                
## [1471,] "Bacteria" NA                    NA                           
## [1472,] "Bacteria" "Fusobacteria"        "Fusobacteriia"              
## [1473,] "Bacteria" "Bacteroidetes"       "Bacteroidia"                
## [1474,] "Bacteria" "Patescibacteria"     "Saccharimonadia"            
## [1475,] "Bacteria" "Bacteroidetes"       "Bacteroidia"                
## [1476,] "Bacteria" "Firmicutes"          "Negativicutes"              
## [1477,] "Bacteria" "Fusobacteria"        "Fusobacteriia"              
## [1478,] "Bacteria" "Firmicutes"          "Clostridia"                 
## [1479,] "Bacteria" "Proteobacteria"      "Gammaproteobacteria"        
## [1480,] "Bacteria" "Firmicutes"          "Negativicutes"              
## [1481,] "Bacteria" "Bacteroidetes"       "Bacteroidia"                
## [1482,] "Bacteria" "Firmicutes"          "Negativicutes"              
## [1483,] "Bacteria" "Bacteroidetes"       "Bacteroidia"                
## [1484,] "Bacteria" "Bacteroidetes"       "Bacteroidia"                
## [1485,] "Bacteria" "Bacteroidetes"       "Bacteroidia"                
## [1486,] "Bacteria" "Firmicutes"          "Negativicutes"              
## [1487,] "Bacteria" "Bacteroidetes"       "Bacteroidia"                
## [1488,] "Bacteria" "Firmicutes"          "Negativicutes"              
## [1489,] "Bacteria" "Proteobacteria"      "Gammaproteobacteria"        
## [1490,] "Bacteria" "Actinobacteria"      "Actinobacteria"             
## [1491,] "Bacteria" "Fusobacteria"        "Fusobacteriia"              
## [1492,] "Bacteria" "Firmicutes"          "Clostridia"                 
## [1493,] "Bacteria" "Firmicutes"          "Clostridia"                 
## [1494,] "Bacteria" "Firmicutes"          "Negativicutes"              
## [1495,] "Bacteria" "Proteobacteria"      "Gammaproteobacteria"        
## [1496,] "Bacteria" "Proteobacteria"      "Gammaproteobacteria"        
## [1497,] "Bacteria" "Bacteroidetes"       "Bacteroidia"                
## [1498,] "Bacteria" "Proteobacteria"      "Gammaproteobacteria"        
## [1499,] "Bacteria" "Patescibacteria"     "Saccharimonadia"            
## [1500,] "Bacteria" "Firmicutes"          "Bacilli"                    
## [1501,] "Bacteria" "Actinobacteria"      "Actinobacteria"             
## [1502,] "Bacteria" "Epsilonbacteraeota"  "Campylobacteria"            
## [1503,] "Bacteria" "Bacteroidetes"       "Bacteroidia"                
## [1504,] "Bacteria" "Firmicutes"          "Negativicutes"              
## [1505,] "Bacteria" "Proteobacteria"      "Gammaproteobacteria"        
## [1506,] "Bacteria" "Patescibacteria"     "Saccharimonadia"            
## [1507,] "Bacteria" "Bacteroidetes"       "Bacteroidia"                
## [1508,] "Bacteria" "Bacteroidetes"       "Bacteroidia"                
## [1509,] "Bacteria" "Proteobacteria"      "Gammaproteobacteria"        
## [1510,] "Bacteria" "Fusobacteria"        "Fusobacteriia"              
## [1511,] "Bacteria" "Proteobacteria"      "Gammaproteobacteria"        
## [1512,] "Bacteria" "Bacteroidetes"       "Bacteroidia"                
## [1513,] "Bacteria" "Bacteroidetes"       "Bacteroidia"                
## [1514,] "Bacteria" "Firmicutes"          "Bacilli"                    
## [1515,] "Bacteria" "Actinobacteria"      "Actinobacteria"             
## [1516,] "Bacteria" "Bacteroidetes"       "Bacteroidia"                
## [1517,] "Bacteria" "Proteobacteria"      "Gammaproteobacteria"        
## [1518,] "Bacteria" "Firmicutes"          "Clostridia"                 
## [1519,] "Bacteria" "Firmicutes"          "Bacilli"                    
## [1520,] "Bacteria" "Bacteroidetes"       "Bacteroidia"                
## [1521,] "Bacteria" "Fusobacteria"        "Fusobacteriia"              
## [1522,] "Bacteria" "Bacteroidetes"       "Bacteroidia"                
## [1523,] "Bacteria" "Fusobacteria"        "Fusobacteriia"              
## [1524,] "Bacteria" "Proteobacteria"      "Gammaproteobacteria"        
## [1525,] "Bacteria" "Actinobacteria"      "Coriobacteriia"             
## [1526,] "Bacteria" "Firmicutes"          "Negativicutes"              
## [1527,] "Bacteria" "Bacteroidetes"       "Bacteroidia"                
## [1528,] "Bacteria" "Firmicutes"          NA                           
## [1529,] "Bacteria" "Bacteroidetes"       "Bacteroidia"                
## [1530,] "Bacteria" "Firmicutes"          "Clostridia"                 
## [1531,] "Bacteria" "Epsilonbacteraeota"  "Campylobacteria"            
## [1532,] "Bacteria" "Patescibacteria"     "Saccharimonadia"            
## [1533,] "Bacteria" "Fusobacteria"        "Fusobacteriia"              
## [1534,] "Bacteria" "Firmicutes"          "Clostridia"                 
## [1535,] "Bacteria" "Firmicutes"          NA                           
## [1536,] "Bacteria" "Bacteroidetes"       "Bacteroidia"                
## [1537,] "Bacteria" "Bacteroidetes"       "Bacteroidia"                
## [1538,] "Bacteria" "Patescibacteria"     "Gracilibacteria"            
## [1539,] "Bacteria" "Actinobacteria"      "Actinobacteria"             
## [1540,] "Bacteria" "Firmicutes"          "Negativicutes"              
## [1541,] "Bacteria" "Firmicutes"          "Negativicutes"              
## [1542,] "Bacteria" "Bacteroidetes"       "Bacteroidia"                
## [1543,] "Bacteria" "Firmicutes"          "Bacilli"                    
## [1544,] "Bacteria" "Spirochaetes"        "Spirochaetia"               
## [1545,] "Bacteria" "Firmicutes"          "Clostridia"                 
## [1546,] "Bacteria" "Firmicutes"          "Negativicutes"              
## [1547,] "Bacteria" "Bacteroidetes"       "Bacteroidia"                
## [1548,] "Bacteria" "Actinobacteria"      "Actinobacteria"             
## [1549,] "Bacteria" "Fusobacteria"        "Fusobacteriia"              
## [1550,] "Bacteria" "Firmicutes"          "Negativicutes"              
## [1551,] "Bacteria" "Firmicutes"          "Clostridia"                 
## [1552,] "Bacteria" "Patescibacteria"     "Gracilibacteria"            
## [1553,] "Bacteria" "Fusobacteria"        "Fusobacteriia"              
## [1554,] "Bacteria" "Firmicutes"          "Negativicutes"              
## [1555,] "Bacteria" "Bacteroidetes"       "Bacteroidia"                
## [1556,] "Bacteria" "Actinobacteria"      "Actinobacteria"             
## [1557,] "Bacteria" "Bacteroidetes"       "Bacteroidia"                
## [1558,] "Bacteria" "Firmicutes"          "Negativicutes"              
## [1559,] "Bacteria" "Fusobacteria"        "Fusobacteriia"              
## [1560,] "Bacteria" "Cyanobacteria"       "Oxyphotobacteria"           
## [1561,] "Bacteria" "Bacteroidetes"       "Bacteroidia"                
## [1562,] "Bacteria" "Bacteroidetes"       "Bacteroidia"                
## [1563,] "Bacteria" "Firmicutes"          "Negativicutes"              
## [1564,] "Bacteria" "Fusobacteria"        "Fusobacteriia"              
## [1565,] "Bacteria" "Bacteroidetes"       "Bacteroidia"                
## [1566,] "Bacteria" "Firmicutes"          "Clostridia"                 
## [1567,] "Bacteria" "Firmicutes"          "Negativicutes"              
## [1568,] "Bacteria" "Firmicutes"          "Bacilli"                    
## [1569,] "Bacteria" "Epsilonbacteraeota"  "Campylobacteria"            
## [1570,] "Bacteria" "Bacteroidetes"       "Bacteroidia"                
## [1571,] "Bacteria" "Bacteroidetes"       "Bacteroidia"                
## [1572,] "Bacteria" "Bacteroidetes"       "Bacteroidia"                
## [1573,] "Bacteria" "Firmicutes"          "Negativicutes"              
## [1574,] "Bacteria" "Fusobacteria"        "Fusobacteriia"              
## [1575,] "Bacteria" "Bacteroidetes"       "Bacteroidia"                
## [1576,] "Bacteria" "Bacteroidetes"       "Bacteroidia"                
## [1577,] "Bacteria" "Fusobacteria"        "Fusobacteriia"              
## [1578,] "Bacteria" "Bacteroidetes"       "Bacteroidia"                
## [1579,] "Bacteria" "Bacteroidetes"       "Bacteroidia"                
## [1580,] "Bacteria" "Bacteroidetes"       "Bacteroidia"                
## [1581,] "Bacteria" "Bacteroidetes"       "Bacteroidia"                
## [1582,] "Bacteria" "Firmicutes"          "Negativicutes"              
## [1583,] "Bacteria" "Firmicutes"          "Bacilli"                    
## [1584,] "Bacteria" "Spirochaetes"        "Spirochaetia"               
## [1585,] "Bacteria" "Epsilonbacteraeota"  "Campylobacteria"            
## [1586,] "Bacteria" "Bacteroidetes"       "Bacteroidia"                
## [1587,] "Bacteria" "Firmicutes"          "Negativicutes"              
## [1588,] "Bacteria" "Firmicutes"          "Negativicutes"              
## [1589,] "Bacteria" "Bacteroidetes"       "Bacteroidia"                
## [1590,] "Bacteria" "Epsilonbacteraeota"  "Campylobacteria"            
## [1591,] "Bacteria" "Bacteroidetes"       "Bacteroidia"                
## [1592,] "Bacteria" "Firmicutes"          "Bacilli"                    
## [1593,] "Bacteria" "Proteobacteria"      "Gammaproteobacteria"        
## [1594,] "Bacteria" "Bacteroidetes"       "Bacteroidia"                
## [1595,] "Bacteria" "Proteobacteria"      "Gammaproteobacteria"        
## [1596,] "Bacteria" "Firmicutes"          "Negativicutes"              
## [1597,] "Bacteria" "Bacteroidetes"       "Bacteroidia"                
## [1598,] "Bacteria" "Bacteroidetes"       "Bacteroidia"                
## [1599,] "Bacteria" "Bacteroidetes"       "Bacteroidia"                
## [1600,] "Bacteria" "Bacteroidetes"       "Bacteroidia"                
## [1601,] "Bacteria" "Firmicutes"          "Negativicutes"              
## [1602,] "Bacteria" "Bacteroidetes"       "Bacteroidia"                
## [1603,] "Bacteria" "Bacteroidetes"       "Bacteroidia"                
## [1604,] "Bacteria" "Patescibacteria"     "Saccharimonadia"            
## [1605,] "Bacteria" "Patescibacteria"     "Saccharimonadia"            
## [1606,] "Bacteria" "Firmicutes"          "Bacilli"                    
## [1607,] "Bacteria" "Bacteroidetes"       "Bacteroidia"                
## [1608,] "Bacteria" "Firmicutes"          "Negativicutes"              
## [1609,] "Bacteria" "Proteobacteria"      "Gammaproteobacteria"        
## [1610,] "Bacteria" "Bacteroidetes"       "Bacteroidia"                
## [1611,] "Bacteria" "Fusobacteria"        "Fusobacteriia"              
## [1612,] "Bacteria" "Fusobacteria"        "Fusobacteriia"              
## [1613,] "Bacteria" "Proteobacteria"      "Gammaproteobacteria"        
## [1614,] "Bacteria" "Bacteroidetes"       "Bacteroidia"                
## [1615,] "Bacteria" "Proteobacteria"      "Gammaproteobacteria"        
## [1616,] "Bacteria" "Cyanobacteria"       "Oxyphotobacteria"           
## [1617,] "Bacteria" "Actinobacteria"      "Actinobacteria"             
## [1618,] "Bacteria" "Firmicutes"          "Negativicutes"              
## [1619,] "Bacteria" "Firmicutes"          "Negativicutes"              
## [1620,] "Bacteria" "Fusobacteria"        "Fusobacteriia"              
## [1621,] "Bacteria" "Actinobacteria"      "Actinobacteria"             
## [1622,] "Bacteria" "Proteobacteria"      "Gammaproteobacteria"        
## [1623,] "Bacteria" "Fusobacteria"        "Fusobacteriia"              
## [1624,] "Bacteria" "Bacteroidetes"       "Bacteroidia"                
## [1625,] "Bacteria" "Fusobacteria"        "Fusobacteriia"              
## [1626,] "Bacteria" "Firmicutes"          "Bacilli"                    
## [1627,] "Bacteria" "Bacteroidetes"       "Bacteroidia"                
## [1628,] "Bacteria" "Firmicutes"          "Clostridia"                 
## [1629,] "Bacteria" "Firmicutes"          "Clostridia"                 
## [1630,] "Bacteria" "Bacteroidetes"       "Bacteroidia"                
## [1631,] "Bacteria" "Bacteroidetes"       "Bacteroidia"                
## [1632,] "Bacteria" "Firmicutes"          "Negativicutes"              
## [1633,] "Bacteria" "Fusobacteria"        "Fusobacteriia"              
## [1634,] "Bacteria" "Firmicutes"          "Negativicutes"              
## [1635,] "Bacteria" "Bacteroidetes"       "Bacteroidia"                
## [1636,] "Bacteria" "Proteobacteria"      "Gammaproteobacteria"        
## [1637,] "Bacteria" "Firmicutes"          "Negativicutes"              
## [1638,] "Bacteria" "Bacteroidetes"       "Bacteroidia"                
## [1639,] "Bacteria" "Bacteroidetes"       "Bacteroidia"                
## [1640,] "Bacteria" "Bacteroidetes"       "Bacteroidia"                
## [1641,] "Bacteria" "Actinobacteria"      "Actinobacteria"             
## [1642,] "Bacteria" "Bacteroidetes"       "Bacteroidia"                
## [1643,] "Bacteria" "Firmicutes"          "Clostridia"                 
## [1644,] "Bacteria" "Patescibacteria"     "Gracilibacteria"            
## [1645,] "Bacteria" "Bacteroidetes"       "Bacteroidia"                
## [1646,] "Bacteria" "Fusobacteria"        "Fusobacteriia"              
## [1647,] "Bacteria" "Firmicutes"          "Clostridia"                 
## [1648,] "Bacteria" "Firmicutes"          "Clostridia"                 
## [1649,] "Bacteria" "Fusobacteria"        "Fusobacteriia"              
## [1650,] "Bacteria" "Fusobacteria"        "Fusobacteriia"              
## [1651,] "Bacteria" "Firmicutes"          "Clostridia"                 
## [1652,] "Bacteria" "Bacteroidetes"       "Bacteroidia"                
## [1653,] "Bacteria" "Firmicutes"          "Negativicutes"              
## [1654,] "Bacteria" "Epsilonbacteraeota"  "Campylobacteria"            
## [1655,] "Bacteria" "Patescibacteria"     "Saccharimonadia"            
## [1656,] "Bacteria" "Actinobacteria"      "Coriobacteriia"             
## [1657,] "Bacteria" "Bacteroidetes"       "Bacteroidia"                
## [1658,] "Bacteria" "Spirochaetes"        "Spirochaetia"               
## [1659,] "Bacteria" "Proteobacteria"      "Gammaproteobacteria"        
## [1660,] "Bacteria" "Proteobacteria"      "Gammaproteobacteria"        
## [1661,] "Bacteria" "Bacteroidetes"       "Bacteroidia"                
## [1662,] "Bacteria" "Bacteroidetes"       "Bacteroidia"                
## [1663,] "Bacteria" "Firmicutes"          "Negativicutes"              
## [1664,] "Bacteria" "Bacteroidetes"       "Bacteroidia"                
## [1665,] "Bacteria" "Bacteroidetes"       "Bacteroidia"                
## [1666,] "Bacteria" "Fusobacteria"        "Fusobacteriia"              
## [1667,] "Bacteria" "Fusobacteria"        "Fusobacteriia"              
## [1668,] "Bacteria" "Firmicutes"          "Clostridia"                 
## [1669,] "Bacteria" "Spirochaetes"        "Spirochaetia"               
## [1670,] "Bacteria" "Fusobacteria"        "Fusobacteriia"              
## [1671,] "Bacteria" "Firmicutes"          NA                           
## [1672,] "Bacteria" "Bacteroidetes"       "Bacteroidia"                
## [1673,] "Bacteria" "Proteobacteria"      "Gammaproteobacteria"        
## [1674,] "Bacteria" "Proteobacteria"      "Gammaproteobacteria"        
## [1675,] "Bacteria" "Bacteroidetes"       "Bacteroidia"                
## [1676,] "Bacteria" "Firmicutes"          "Clostridia"                 
## [1677,] "Bacteria" "Firmicutes"          "Bacilli"                    
## [1678,] "Bacteria" "Firmicutes"          NA                           
## [1679,] "Bacteria" "Proteobacteria"      "Gammaproteobacteria"        
## [1680,] "Bacteria" "Patescibacteria"     "Saccharimonadia"            
## [1681,] "Bacteria" "Bacteroidetes"       "Bacteroidia"                
## [1682,] "Bacteria" "Bacteroidetes"       "Bacteroidia"                
## [1683,] "Bacteria" "Firmicutes"          "Bacilli"                    
## [1684,] "Bacteria" "Proteobacteria"      "Gammaproteobacteria"        
## [1685,] "Bacteria" "Firmicutes"          "Negativicutes"              
## [1686,] "Bacteria" "Firmicutes"          "Bacilli"                    
## [1687,] "Bacteria" "Bacteroidetes"       "Bacteroidia"                
## [1688,] "Bacteria" "Firmicutes"          "Negativicutes"              
## [1689,] "Bacteria" "Fusobacteria"        "Fusobacteriia"              
## [1690,] "Bacteria" "Proteobacteria"      "Gammaproteobacteria"        
## [1691,] "Bacteria" "Fusobacteria"        "Fusobacteriia"              
## [1692,] "Bacteria" "Bacteroidetes"       "Bacteroidia"                
## [1693,] "Bacteria" "Fusobacteria"        "Fusobacteriia"              
## [1694,] "Bacteria" "Bacteroidetes"       "Bacteroidia"                
## [1695,] "Bacteria" "Bacteroidetes"       "Bacteroidia"                
## [1696,] "Bacteria" "Actinobacteria"      "Actinobacteria"             
## [1697,] "Bacteria" "Bacteroidetes"       "Bacteroidia"                
## [1698,] "Bacteria" "Epsilonbacteraeota"  "Campylobacteria"            
## [1699,] "Bacteria" "Proteobacteria"      "Gammaproteobacteria"        
## [1700,] "Bacteria" "Actinobacteria"      "Actinobacteria"             
## [1701,] "Bacteria" "Patescibacteria"     "Saccharimonadia"            
## [1702,] "Bacteria" "Fusobacteria"        "Fusobacteriia"              
## [1703,] "Bacteria" "Firmicutes"          "Bacilli"                    
## [1704,] "Bacteria" "Proteobacteria"      "Gammaproteobacteria"        
## [1705,] "Bacteria" "Bacteroidetes"       "Bacteroidia"                
## [1706,] "Bacteria" "Proteobacteria"      "Gammaproteobacteria"        
## [1707,] "Bacteria" "Firmicutes"          "Bacilli"                    
## [1708,] "Bacteria" "Firmicutes"          "Negativicutes"              
## [1709,] "Bacteria" "Firmicutes"          "Bacilli"                    
## [1710,] "Bacteria" "Bacteroidetes"       "Bacteroidia"                
## [1711,] "Bacteria" "Firmicutes"          "Negativicutes"              
## [1712,] "Bacteria" "Firmicutes"          "Negativicutes"              
## [1713,] "Bacteria" "Firmicutes"          "Clostridia"                 
## [1714,] "Bacteria" "Firmicutes"          "Negativicutes"              
## [1715,] "Bacteria" "Bacteroidetes"       "Bacteroidia"                
## [1716,] "Bacteria" "Fusobacteria"        "Fusobacteriia"              
## [1717,] "Bacteria" "Proteobacteria"      "Gammaproteobacteria"        
## [1718,] "Bacteria" "Bacteroidetes"       "Bacteroidia"                
## [1719,] "Bacteria" "Firmicutes"          "Clostridia"                 
## [1720,] "Bacteria" "Proteobacteria"      "Gammaproteobacteria"        
## [1721,] "Bacteria" "Bacteroidetes"       "Bacteroidia"                
## [1722,] "Bacteria" "Bacteroidetes"       "Bacteroidia"                
## [1723,] "Bacteria" "Firmicutes"          "Negativicutes"              
## [1724,] "Bacteria" "Bacteroidetes"       "Bacteroidia"                
## [1725,] "Bacteria" "Proteobacteria"      "Gammaproteobacteria"        
## [1726,] "Bacteria" "Bacteroidetes"       "Bacteroidia"                
## [1727,] "Bacteria" "Actinobacteria"      "Coriobacteriia"             
## [1728,] "Bacteria" "Bacteroidetes"       "Bacteroidia"                
## [1729,] "Bacteria" "Bacteroidetes"       "Bacteroidia"                
## [1730,] "Bacteria" "Fusobacteria"        "Fusobacteriia"              
## [1731,] "Bacteria" "Firmicutes"          "Negativicutes"              
## [1732,] "Bacteria" "Actinobacteria"      "Actinobacteria"             
## [1733,] "Bacteria" "Bacteroidetes"       "Bacteroidia"                
## [1734,] "Bacteria" "Bacteroidetes"       "Bacteroidia"                
## [1735,] "Bacteria" "Bacteroidetes"       "Bacteroidia"                
## [1736,] "Bacteria" "Bacteroidetes"       "Bacteroidia"                
## [1737,] "Bacteria" "Firmicutes"          "Clostridia"                 
## [1738,] "Bacteria" "Bacteroidetes"       "Bacteroidia"                
## [1739,] "Bacteria" "Bacteroidetes"       "Bacteroidia"                
## [1740,] "Bacteria" "Bacteroidetes"       "Bacteroidia"                
## [1741,] "Bacteria" "Firmicutes"          "Negativicutes"              
## [1742,] "Bacteria" "Bacteroidetes"       "Bacteroidia"                
## [1743,] "Bacteria" "Firmicutes"          "Bacilli"                    
## [1744,] "Bacteria" "Bacteroidetes"       "Bacteroidia"                
## [1745,] "Bacteria" "Proteobacteria"      "Gammaproteobacteria"        
## [1746,] "Bacteria" "Bacteroidetes"       "Bacteroidia"                
## [1747,] "Bacteria" "Bacteroidetes"       "Bacteroidia"                
## [1748,] "Bacteria" "Fusobacteria"        "Fusobacteriia"              
## [1749,] "Bacteria" "Bacteroidetes"       "Bacteroidia"                
## [1750,] "Bacteria" "Bacteroidetes"       "Bacteroidia"                
## [1751,] "Bacteria" "Firmicutes"          "Bacilli"                    
## [1752,] "Bacteria" "Firmicutes"          "Negativicutes"              
## [1753,] "Bacteria" "Bacteroidetes"       "Bacteroidia"                
## [1754,] "Bacteria" "Bacteroidetes"       "Bacteroidia"                
## [1755,] "Bacteria" "Firmicutes"          "Negativicutes"              
## [1756,] "Bacteria" "Firmicutes"          "Bacilli"                    
## [1757,] "Bacteria" "Actinobacteria"      "Actinobacteria"             
## [1758,] "Bacteria" "Bacteroidetes"       "Bacteroidia"                
## [1759,] "Bacteria" "Bacteroidetes"       "Bacteroidia"                
## [1760,] "Bacteria" "Patescibacteria"     "Saccharimonadia"            
## [1761,] "Bacteria" "Bacteroidetes"       "Bacteroidia"                
## [1762,] "Bacteria" "Firmicutes"          "Bacilli"                    
## [1763,] "Bacteria" "Firmicutes"          "Bacilli"                    
## [1764,] "Bacteria" "Firmicutes"          "Negativicutes"              
## [1765,] "Bacteria" "Firmicutes"          "Clostridia"                 
## [1766,] "Bacteria" "Bacteroidetes"       "Bacteroidia"                
## [1767,] "Bacteria" "Firmicutes"          "Clostridia"                 
## [1768,] "Bacteria" "Bacteroidetes"       "Bacteroidia"                
## [1769,] "Bacteria" "Firmicutes"          "Negativicutes"              
## [1770,] "Bacteria" "Bacteroidetes"       "Bacteroidia"                
## [1771,] "Bacteria" "Firmicutes"          "Negativicutes"              
## [1772,] "Bacteria" "Proteobacteria"      "Gammaproteobacteria"        
## [1773,] "Bacteria" "Firmicutes"          "Clostridia"                 
## [1774,] "Bacteria" "Actinobacteria"      "Actinobacteria"             
## [1775,] "Bacteria" "Firmicutes"          "Clostridia"                 
## [1776,] "Bacteria" "Firmicutes"          "Bacilli"                    
## [1777,] "Bacteria" "Proteobacteria"      "Gammaproteobacteria"        
## [1778,] "Bacteria" "Proteobacteria"      "Gammaproteobacteria"        
## [1779,] "Bacteria" "Fusobacteria"        "Fusobacteriia"              
## [1780,] "Bacteria" "Bacteroidetes"       "Bacteroidia"                
## [1781,] "Bacteria" "Firmicutes"          "Clostridia"                 
## [1782,] "Bacteria" "Proteobacteria"      "Gammaproteobacteria"        
## [1783,] "Bacteria" "Actinobacteria"      "Actinobacteria"             
## [1784,] "Bacteria" "Bacteroidetes"       "Bacteroidia"                
## [1785,] "Bacteria" "Bacteroidetes"       "Bacteroidia"                
## [1786,] "Bacteria" "Bacteroidetes"       "Bacteroidia"                
## [1787,] "Bacteria" "Firmicutes"          "Negativicutes"              
## [1788,] "Bacteria" "Bacteroidetes"       "Bacteroidia"                
## [1789,] "Bacteria" "Proteobacteria"      "Gammaproteobacteria"        
## [1790,] "Bacteria" "Fusobacteria"        "Fusobacteriia"              
## [1791,] "Bacteria" "Proteobacteria"      "Gammaproteobacteria"        
## [1792,] "Bacteria" "Firmicutes"          NA                           
## [1793,] "Bacteria" "Proteobacteria"      "Gammaproteobacteria"        
## [1794,] "Bacteria" "Firmicutes"          "Clostridia"                 
## [1795,] "Bacteria" "Firmicutes"          NA                           
## [1796,] "Bacteria" "Bacteroidetes"       "Bacteroidia"                
## [1797,] "Bacteria" "Bacteroidetes"       "Bacteroidia"                
## [1798,] "Bacteria" "Firmicutes"          "Clostridia"                 
## [1799,] "Bacteria" "Firmicutes"          "Bacilli"                    
## [1800,] "Bacteria" "Firmicutes"          "Clostridia"                 
## [1801,] "Bacteria" "Firmicutes"          "Clostridia"                 
## [1802,] "Bacteria" "Firmicutes"          "Negativicutes"              
## [1803,] "Bacteria" "Fusobacteria"        "Fusobacteriia"              
## [1804,] "Bacteria" "Firmicutes"          "Clostridia"                 
## [1805,] "Bacteria" "Fusobacteria"        "Fusobacteriia"              
## [1806,] "Bacteria" "Bacteroidetes"       "Bacteroidia"                
## [1807,] "Bacteria" "Fusobacteria"        "Fusobacteriia"              
## [1808,] "Bacteria" "Firmicutes"          "Bacilli"                    
## [1809,] "Bacteria" "Actinobacteria"      "Actinobacteria"             
## [1810,] "Bacteria" "Actinobacteria"      "Actinobacteria"             
## [1811,] "Bacteria" "Firmicutes"          "Negativicutes"              
## [1812,] "Bacteria" "Firmicutes"          "Negativicutes"              
## [1813,] "Bacteria" "Firmicutes"          "Negativicutes"              
## [1814,] "Bacteria" "Fusobacteria"        "Fusobacteriia"              
## [1815,] "Bacteria" "Proteobacteria"      "Gammaproteobacteria"        
## [1816,] "Bacteria" "Proteobacteria"      "Gammaproteobacteria"        
## [1817,] "Bacteria" "Proteobacteria"      "Gammaproteobacteria"        
## [1818,] "Bacteria" "Bacteroidetes"       "Bacteroidia"                
## [1819,] "Bacteria" "Bacteroidetes"       "Bacteroidia"                
## [1820,] "Bacteria" "Bacteroidetes"       "Bacteroidia"                
## [1821,] "Bacteria" "Proteobacteria"      "Gammaproteobacteria"        
## [1822,] "Bacteria" "Bacteroidetes"       "Bacteroidia"                
## [1823,] "Bacteria" "Firmicutes"          "Negativicutes"              
## [1824,] "Bacteria" "Firmicutes"          "Negativicutes"              
## [1825,] "Bacteria" "Firmicutes"          "Bacilli"                    
## [1826,] "Bacteria" "Firmicutes"          "Negativicutes"              
## [1827,] "Bacteria" "Bacteroidetes"       "Bacteroidia"                
## [1828,] "Bacteria" "Proteobacteria"      "Gammaproteobacteria"        
## [1829,] "Bacteria" "Firmicutes"          "Clostridia"                 
## [1830,] "Bacteria" "Fusobacteria"        "Fusobacteriia"              
## [1831,] "Bacteria" "Fusobacteria"        "Fusobacteriia"              
## [1832,] "Bacteria" "Bacteroidetes"       "Bacteroidia"                
## [1833,] "Bacteria" "Actinobacteria"      "Actinobacteria"             
## [1834,] "Bacteria" "Patescibacteria"     "Saccharimonadia"            
## [1835,] "Bacteria" "Bacteroidetes"       "Bacteroidia"                
## [1836,] "Bacteria" "Spirochaetes"        "Spirochaetia"               
## [1837,] "Bacteria" NA                    NA                           
## [1838,] "Bacteria" "Bacteroidetes"       "Bacteroidia"                
## [1839,] "Bacteria" "Fusobacteria"        "Fusobacteriia"              
## [1840,] "Bacteria" "Spirochaetes"        "Spirochaetia"               
## [1841,] "Bacteria" "Firmicutes"          "Negativicutes"              
## [1842,] "Bacteria" "Firmicutes"          "Clostridia"                 
## [1843,] "Bacteria" "Bacteroidetes"       "Bacteroidia"                
## [1844,] "Bacteria" "Bacteroidetes"       "Bacteroidia"                
## [1845,] "Bacteria" "Firmicutes"          "Clostridia"                 
## [1846,] "Bacteria" "Proteobacteria"      "Gammaproteobacteria"        
## [1847,] "Bacteria" "Bacteroidetes"       "Bacteroidia"                
## [1848,] "Bacteria" "Bacteroidetes"       "Bacteroidia"                
## [1849,] "Bacteria" "Bacteroidetes"       "Bacteroidia"                
## [1850,] "Bacteria" "Bacteroidetes"       "Bacteroidia"                
## [1851,] "Bacteria" "Synergistetes"       "Synergistia"                
## [1852,] "Bacteria" "Bacteroidetes"       "Bacteroidia"                
## [1853,] "Bacteria" "Bacteroidetes"       "Bacteroidia"                
## [1854,] "Bacteria" "Proteobacteria"      "Gammaproteobacteria"        
## [1855,] "Bacteria" "Bacteroidetes"       "Bacteroidia"                
## [1856,] "Bacteria" "Proteobacteria"      "Gammaproteobacteria"        
## [1857,] "Bacteria" "Bacteroidetes"       "Bacteroidia"                
## [1858,] "Bacteria" "Bacteroidetes"       "Bacteroidia"                
## [1859,] "Bacteria" "Proteobacteria"      "Gammaproteobacteria"        
## [1860,] "Bacteria" "Firmicutes"          "Clostridia"                 
## [1861,] "Bacteria" "Proteobacteria"      "Gammaproteobacteria"        
## [1862,] "Bacteria" "Bacteroidetes"       "Bacteroidia"                
## [1863,] "Bacteria" "Proteobacteria"      "Gammaproteobacteria"        
## [1864,] "Bacteria" "Fusobacteria"        "Fusobacteriia"              
## [1865,] "Bacteria" "Proteobacteria"      "Gammaproteobacteria"        
## [1866,] "Bacteria" "Bacteroidetes"       "Bacteroidia"                
## [1867,] "Bacteria" "Fusobacteria"        "Fusobacteriia"              
## [1868,] "Bacteria" "Firmicutes"          "Clostridia"                 
## [1869,] "Bacteria" "Proteobacteria"      "Gammaproteobacteria"        
## [1870,] "Bacteria" "Firmicutes"          "Negativicutes"              
## [1871,] "Bacteria" "Proteobacteria"      "Gammaproteobacteria"        
## [1872,] "Bacteria" "Firmicutes"          "Bacilli"                    
## [1873,] "Bacteria" "Proteobacteria"      "Gammaproteobacteria"        
## [1874,] "Bacteria" "Firmicutes"          "Negativicutes"              
## [1875,] "Bacteria" "Fusobacteria"        "Fusobacteriia"              
## [1876,] "Bacteria" "Firmicutes"          "Negativicutes"              
## [1877,] "Bacteria" "Actinobacteria"      "Actinobacteria"             
## [1878,] "Bacteria" "Bacteroidetes"       "Bacteroidia"                
## [1879,] "Bacteria" "Proteobacteria"      "Gammaproteobacteria"        
## [1880,] "Bacteria" "Firmicutes"          "Clostridia"                 
## [1881,] "Bacteria" "Bacteroidetes"       "Bacteroidia"                
## [1882,] "Bacteria" "Firmicutes"          "Negativicutes"              
## [1883,] "Bacteria" "Fusobacteria"        "Fusobacteriia"              
## [1884,] "Bacteria" "Bacteroidetes"       "Bacteroidia"                
## [1885,] "Bacteria" "Actinobacteria"      "Actinobacteria"             
## [1886,] "Bacteria" "Bacteroidetes"       "Bacteroidia"                
## [1887,] "Bacteria" "Firmicutes"          "Negativicutes"              
## [1888,] "Bacteria" "Actinobacteria"      "Actinobacteria"             
## [1889,] "Bacteria" "Bacteroidetes"       "Bacteroidia"                
## [1890,] "Bacteria" "Bacteroidetes"       "Bacteroidia"                
## [1891,] "Bacteria" "Proteobacteria"      "Gammaproteobacteria"        
## [1892,] "Bacteria" "Proteobacteria"      "Gammaproteobacteria"        
## [1893,] "Bacteria" "Firmicutes"          "Negativicutes"              
## [1894,] "Bacteria" "Fusobacteria"        "Fusobacteriia"              
## [1895,] "Bacteria" "Firmicutes"          "Negativicutes"              
## [1896,] "Bacteria" "Firmicutes"          "Clostridia"                 
## [1897,] "Bacteria" "Bacteroidetes"       "Bacteroidia"                
## [1898,] "Bacteria" "Firmicutes"          "Bacilli"                    
## [1899,] "Bacteria" "Proteobacteria"      "Gammaproteobacteria"        
## [1900,] "Bacteria" "Firmicutes"          "Negativicutes"              
## [1901,] "Bacteria" "Firmicutes"          "Clostridia"                 
## [1902,] "Bacteria" "Firmicutes"          "Negativicutes"              
## [1903,] "Bacteria" "Bacteroidetes"       "Bacteroidia"                
## [1904,] "Bacteria" "Bacteroidetes"       "Bacteroidia"                
## [1905,] "Bacteria" "Fusobacteria"        "Fusobacteriia"              
## [1906,] "Bacteria" "Patescibacteria"     "Saccharimonadia"            
## [1907,] "Bacteria" "Firmicutes"          "Clostridia"                 
## [1908,] "Bacteria" "Bacteroidetes"       "Bacteroidia"                
## [1909,] "Bacteria" "Firmicutes"          "Clostridia"                 
## [1910,] "Bacteria" "Fusobacteria"        "Fusobacteriia"              
## [1911,] "Bacteria" "Proteobacteria"      "Gammaproteobacteria"        
## [1912,] "Bacteria" "Bacteroidetes"       "Bacteroidia"                
## [1913,] "Bacteria" "Firmicutes"          "Negativicutes"              
## [1914,] "Bacteria" "Bacteroidetes"       "Bacteroidia"                
## [1915,] "Bacteria" "Bacteroidetes"       "Bacteroidia"                
## [1916,] "Bacteria" "Spirochaetes"        "Spirochaetia"               
## [1917,] "Bacteria" "Bacteroidetes"       "Bacteroidia"                
## [1918,] "Bacteria" "Bacteroidetes"       "Bacteroidia"                
## [1919,] "Bacteria" "Actinobacteria"      "Actinobacteria"             
## [1920,] "Bacteria" "Firmicutes"          "Clostridia"                 
## [1921,] "Bacteria" "Bacteroidetes"       "Bacteroidia"                
## [1922,] "Bacteria" "Firmicutes"          "Clostridia"                 
## [1923,] "Bacteria" "Proteobacteria"      "Gammaproteobacteria"        
## [1924,] "Bacteria" "Proteobacteria"      "Gammaproteobacteria"        
## [1925,] "Bacteria" "Bacteroidetes"       "Bacteroidia"                
## [1926,] "Bacteria" "Bacteroidetes"       "Bacteroidia"                
## [1927,] "Bacteria" "Epsilonbacteraeota"  "Campylobacteria"            
## [1928,] "Bacteria" "Proteobacteria"      "Gammaproteobacteria"        
## [1929,] "Bacteria" "Firmicutes"          "Negativicutes"              
## [1930,] "Bacteria" "Bacteroidetes"       "Bacteroidia"                
## [1931,] "Bacteria" "Fusobacteria"        "Fusobacteriia"              
## [1932,] "Bacteria" "Bacteroidetes"       "Bacteroidia"                
## [1933,] "Bacteria" "Patescibacteria"     "Saccharimonadia"            
## [1934,] "Bacteria" "Bacteroidetes"       "Bacteroidia"                
## [1935,] "Bacteria" "Actinobacteria"      "Actinobacteria"             
## [1936,] "Bacteria" "Actinobacteria"      "Actinobacteria"             
## [1937,] "Bacteria" "Proteobacteria"      "Gammaproteobacteria"        
## [1938,] "Bacteria" "Bacteroidetes"       "Bacteroidia"                
## [1939,] "Bacteria" "Bacteroidetes"       "Bacteroidia"                
## [1940,] "Bacteria" "Fusobacteria"        "Fusobacteriia"              
## [1941,] "Bacteria" "Firmicutes"          "Bacilli"                    
## [1942,] "Bacteria" "Actinobacteria"      "Actinobacteria"             
## [1943,] "Bacteria" "Bacteroidetes"       "Bacteroidia"                
## [1944,] "Bacteria" "Proteobacteria"      "Gammaproteobacteria"        
## [1945,] "Bacteria" "Firmicutes"          "Clostridia"                 
## [1946,] "Bacteria" "Spirochaetes"        "Spirochaetia"               
## [1947,] "Bacteria" "Proteobacteria"      "Gammaproteobacteria"        
## [1948,] "Bacteria" "Fusobacteria"        "Fusobacteriia"              
## [1949,] "Bacteria" "Bacteroidetes"       "Bacteroidia"                
## [1950,] "Bacteria" "Firmicutes"          "Negativicutes"              
## [1951,] "Bacteria" "Bacteroidetes"       "Bacteroidia"                
## [1952,] "Bacteria" "Firmicutes"          "Clostridia"                 
## [1953,] "Bacteria" "Firmicutes"          "Negativicutes"              
## [1954,] "Bacteria" "Proteobacteria"      "Gammaproteobacteria"        
## [1955,] "Bacteria" "Bacteroidetes"       "Bacteroidia"                
## [1956,] "Bacteria" "Bacteroidetes"       "Bacteroidia"                
## [1957,] "Bacteria" "Firmicutes"          "Clostridia"                 
## [1958,] "Bacteria" "Bacteroidetes"       "Bacteroidia"                
## [1959,] "Bacteria" "Bacteroidetes"       "Bacteroidia"                
## [1960,] "Bacteria" "Bacteroidetes"       "Bacteroidia"                
## [1961,] "Bacteria" "Bacteroidetes"       "Bacteroidia"                
## [1962,] "Bacteria" "Actinobacteria"      "Coriobacteriia"             
## [1963,] "Bacteria" "Bacteroidetes"       "Bacteroidia"                
## [1964,] "Bacteria" "Actinobacteria"      "Actinobacteria"             
## [1965,] "Bacteria" "Fusobacteria"        "Fusobacteriia"              
## [1966,] "Bacteria" "Firmicutes"          "Negativicutes"              
## [1967,] "Bacteria" "Bacteroidetes"       "Bacteroidia"                
## [1968,] "Bacteria" "Firmicutes"          "Negativicutes"              
## [1969,] "Bacteria" "Fusobacteria"        "Fusobacteriia"              
## [1970,] "Bacteria" "Proteobacteria"      "Gammaproteobacteria"        
## [1971,] "Bacteria" "Fusobacteria"        "Fusobacteriia"              
## [1972,] "Bacteria" "Firmicutes"          "Clostridia"                 
## [1973,] "Bacteria" "Bacteroidetes"       "Bacteroidia"                
## [1974,] "Bacteria" "Firmicutes"          "Clostridia"                 
## [1975,] "Bacteria" "Firmicutes"          "Clostridia"                 
## [1976,] "Bacteria" "Bacteroidetes"       "Bacteroidia"                
## [1977,] "Bacteria" "Firmicutes"          "Clostridia"                 
## [1978,] "Bacteria" "Bacteroidetes"       "Bacteroidia"                
## [1979,] "Bacteria" "Bacteroidetes"       "Bacteroidia"                
## [1980,] "Bacteria" "Bacteroidetes"       "Bacteroidia"                
## [1981,] "Bacteria" "Bacteroidetes"       "Bacteroidia"                
## [1982,] "Bacteria" "Fusobacteria"        "Fusobacteriia"              
## [1983,] "Bacteria" "Firmicutes"          "Negativicutes"              
## [1984,] "Bacteria" "Proteobacteria"      "Gammaproteobacteria"        
## [1985,] "Bacteria" "Bacteroidetes"       "Bacteroidia"                
## [1986,] "Bacteria" "Firmicutes"          "Negativicutes"              
## [1987,] "Bacteria" "Proteobacteria"      "Gammaproteobacteria"        
## [1988,] "Bacteria" "Actinobacteria"      "Actinobacteria"             
## [1989,] "Bacteria" "Fusobacteria"        "Fusobacteriia"              
## [1990,] "Bacteria" "Firmicutes"          "Bacilli"                    
## [1991,] "Bacteria" "Bacteroidetes"       "Bacteroidia"                
## [1992,] "Bacteria" "Bacteroidetes"       "Bacteroidia"                
## [1993,] "Bacteria" "Bacteroidetes"       "Bacteroidia"                
## [1994,] "Bacteria" "Proteobacteria"      "Gammaproteobacteria"        
## [1995,] "Bacteria" "Firmicutes"          "Negativicutes"              
## [1996,] "Bacteria" "Firmicutes"          "Bacilli"                    
## [1997,] "Bacteria" "Bacteroidetes"       "Bacteroidia"                
## [1998,] "Bacteria" "Firmicutes"          "Negativicutes"              
## [1999,] "Bacteria" "Firmicutes"          "Clostridia"                 
## [2000,] "Bacteria" "Proteobacteria"      "Gammaproteobacteria"        
## [2001,] "Bacteria" "Fusobacteria"        "Fusobacteriia"              
## [2002,] "Bacteria" "Firmicutes"          "Clostridia"                 
## [2003,] "Bacteria" "Fusobacteria"        "Fusobacteriia"              
## [2004,] "Bacteria" "Bacteroidetes"       "Bacteroidia"                
## [2005,] "Bacteria" "Bacteroidetes"       "Bacteroidia"                
## [2006,] "Bacteria" "Firmicutes"          "Clostridia"                 
## [2007,] "Bacteria" "Bacteroidetes"       "Bacteroidia"                
## [2008,] "Bacteria" "Firmicutes"          "Negativicutes"              
## [2009,] "Bacteria" "Bacteroidetes"       "Bacteroidia"                
## [2010,] "Bacteria" "Proteobacteria"      "Gammaproteobacteria"        
## [2011,] "Bacteria" "Bacteroidetes"       "Bacteroidia"                
## [2012,] "Bacteria" "Bacteroidetes"       "Bacteroidia"                
## [2013,] "Bacteria" "Patescibacteria"     "Saccharimonadia"            
## [2014,] "Bacteria" "Bacteroidetes"       "Bacteroidia"                
## [2015,] "Bacteria" "Bacteroidetes"       "Bacteroidia"                
## [2016,] "Bacteria" "Firmicutes"          "Negativicutes"              
## [2017,] "Bacteria" "Bacteroidetes"       "Bacteroidia"                
## [2018,] "Bacteria" "Patescibacteria"     "Saccharimonadia"            
## [2019,] "Bacteria" "Firmicutes"          "Negativicutes"              
## [2020,] "Bacteria" "Fusobacteria"        "Fusobacteriia"              
## [2021,] "Bacteria" "Bacteroidetes"       "Bacteroidia"                
## [2022,] "Bacteria" "Actinobacteria"      "Actinobacteria"             
## [2023,] "Bacteria" "Proteobacteria"      "Gammaproteobacteria"        
## [2024,] "Bacteria" "Proteobacteria"      "Gammaproteobacteria"        
## [2025,] "Bacteria" "Proteobacteria"      "Gammaproteobacteria"        
## [2026,] "Bacteria" "Bacteroidetes"       "Bacteroidia"                
## [2027,] "Bacteria" NA                    NA                           
## [2028,] "Bacteria" "Bacteroidetes"       "Bacteroidia"                
## [2029,] "Bacteria" "Fusobacteria"        "Fusobacteriia"              
## [2030,] "Bacteria" "Fusobacteria"        "Fusobacteriia"              
## [2031,] "Bacteria" "Fusobacteria"        "Fusobacteriia"              
## [2032,] "Bacteria" "Patescibacteria"     "Saccharimonadia"            
## [2033,] "Bacteria" "Fusobacteria"        "Fusobacteriia"              
## [2034,] "Bacteria" "Firmicutes"          "Bacilli"                    
## [2035,] "Bacteria" "Firmicutes"          "Bacilli"                    
## [2036,] "Bacteria" "Firmicutes"          "Bacilli"                    
## [2037,] "Bacteria" "Synergistetes"       "Synergistia"                
## [2038,] "Bacteria" "Bacteroidetes"       "Bacteroidia"                
## [2039,] "Bacteria" "Fusobacteria"        "Fusobacteriia"              
## [2040,] "Bacteria" "Firmicutes"          "Negativicutes"              
## [2041,] "Bacteria" "Firmicutes"          "Negativicutes"              
## [2042,] "Bacteria" "Bacteroidetes"       "Bacteroidia"                
## [2043,] "Bacteria" "Proteobacteria"      "Gammaproteobacteria"        
## [2044,] "Bacteria" "Proteobacteria"      "Gammaproteobacteria"        
## [2045,] "Bacteria" "Patescibacteria"     "Gracilibacteria"            
## [2046,] "Bacteria" "Fusobacteria"        "Fusobacteriia"              
## [2047,] "Bacteria" "Firmicutes"          "Clostridia"                 
## [2048,] "Bacteria" "Firmicutes"          "Negativicutes"              
## [2049,] "Bacteria" "Epsilonbacteraeota"  "Campylobacteria"            
## [2050,] "Bacteria" "Bacteroidetes"       "Bacteroidia"                
## [2051,] "Bacteria" "Bacteroidetes"       "Bacteroidia"                
## [2052,] "Bacteria" "Firmicutes"          "Negativicutes"              
## [2053,] "Bacteria" "Bacteroidetes"       "Bacteroidia"                
## [2054,] "Bacteria" "Actinobacteria"      "Actinobacteria"             
## [2055,] "Bacteria" "Bacteroidetes"       "Bacteroidia"                
## [2056,] "Bacteria" "Firmicutes"          "Negativicutes"              
## [2057,] "Bacteria" "Patescibacteria"     "Saccharimonadia"            
## [2058,] "Bacteria" "Firmicutes"          "Negativicutes"              
## [2059,] "Bacteria" "Bacteroidetes"       "Bacteroidia"                
## [2060,] "Bacteria" "Firmicutes"          "Clostridia"                 
## [2061,] "Bacteria" "Bacteroidetes"       "Bacteroidia"                
## [2062,] "Bacteria" "Firmicutes"          "Negativicutes"              
## [2063,] "Bacteria" "Cyanobacteria"       "Oxyphotobacteria"           
## [2064,] "Bacteria" "Bacteroidetes"       "Bacteroidia"                
## [2065,] "Bacteria" "Firmicutes"          "Negativicutes"              
## [2066,] "Bacteria" "Proteobacteria"      "Gammaproteobacteria"        
## [2067,] "Bacteria" "Bacteroidetes"       "Bacteroidia"                
## [2068,] "Bacteria" "Bacteroidetes"       "Bacteroidia"                
## [2069,] "Bacteria" "Fusobacteria"        "Fusobacteriia"              
## [2070,] "Bacteria" "Patescibacteria"     "Saccharimonadia"            
## [2071,] "Bacteria" "Bacteroidetes"       "Bacteroidia"                
## [2072,] "Bacteria" "Epsilonbacteraeota"  "Campylobacteria"            
## [2073,] "Bacteria" "Firmicutes"          "Clostridia"                 
## [2074,] "Bacteria" "Proteobacteria"      "Gammaproteobacteria"        
## [2075,] "Bacteria" "Fusobacteria"        "Fusobacteriia"              
## [2076,] "Bacteria" "Firmicutes"          "Negativicutes"              
## [2077,] "Bacteria" "Spirochaetes"        "Spirochaetia"               
## [2078,] "Bacteria" "Fusobacteria"        "Fusobacteriia"              
## [2079,] "Bacteria" "Patescibacteria"     "Saccharimonadia"            
## [2080,] "Bacteria" "Bacteroidetes"       "Bacteroidia"                
## [2081,] "Bacteria" "Bacteroidetes"       "Bacteroidia"                
## [2082,] "Bacteria" "Bacteroidetes"       "Bacteroidia"                
## [2083,] "Bacteria" "Fusobacteria"        "Fusobacteriia"              
## [2084,] "Bacteria" NA                    NA                           
## [2085,] "Bacteria" "Proteobacteria"      "Gammaproteobacteria"        
## [2086,] "Bacteria" "Fusobacteria"        "Fusobacteriia"              
## [2087,] "Bacteria" "Firmicutes"          "Clostridia"                 
## [2088,] "Bacteria" "Bacteroidetes"       "Bacteroidia"                
## [2089,] "Bacteria" "Firmicutes"          "Negativicutes"              
## [2090,] "Bacteria" "Bacteroidetes"       "Bacteroidia"                
## [2091,] "Bacteria" "Spirochaetes"        "Spirochaetia"               
## [2092,] "Bacteria" "Actinobacteria"      "Actinobacteria"             
## [2093,] "Bacteria" "Actinobacteria"      "Actinobacteria"             
## [2094,] "Bacteria" "Bacteroidetes"       "Bacteroidia"                
## [2095,] "Bacteria" "Bacteroidetes"       "Bacteroidia"                
## [2096,] "Bacteria" "Fusobacteria"        "Fusobacteriia"              
## [2097,] "Bacteria" "Bacteroidetes"       "Bacteroidia"                
## [2098,] "Bacteria" "Spirochaetes"        "Spirochaetia"               
## [2099,] "Bacteria" "Fusobacteria"        "Fusobacteriia"              
## [2100,] "Bacteria" "Bacteroidetes"       "Bacteroidia"                
## [2101,] "Bacteria" "Bacteroidetes"       "Bacteroidia"                
## [2102,] "Bacteria" "Fusobacteria"        "Fusobacteriia"              
## [2103,] "Bacteria" "Firmicutes"          "Clostridia"                 
## [2104,] "Bacteria" "Bacteroidetes"       "Bacteroidia"                
## [2105,] "Bacteria" "Fusobacteria"        "Fusobacteriia"              
## [2106,] "Bacteria" "Proteobacteria"      "Gammaproteobacteria"        
## [2107,] "Bacteria" "Bacteroidetes"       "Bacteroidia"                
## [2108,] "Bacteria" "Firmicutes"          "Negativicutes"              
## [2109,] "Bacteria" "Bacteroidetes"       "Bacteroidia"                
## [2110,] "Bacteria" "Bacteroidetes"       "Bacteroidia"                
## [2111,] "Bacteria" "Bacteroidetes"       "Bacteroidia"                
## [2112,] "Bacteria" "Firmicutes"          "Clostridia"                 
## [2113,] "Bacteria" "Firmicutes"          "Negativicutes"              
## [2114,] "Bacteria" "Proteobacteria"      "Gammaproteobacteria"        
## [2115,] "Bacteria" "Bacteroidetes"       "Bacteroidia"                
## [2116,] "Bacteria" "Firmicutes"          NA                           
## [2117,] "Bacteria" "Firmicutes"          "Bacilli"                    
## [2118,] "Bacteria" "Bacteroidetes"       "Bacteroidia"                
## [2119,] "Bacteria" "Firmicutes"          "Negativicutes"              
## [2120,] "Bacteria" "Proteobacteria"      "Gammaproteobacteria"        
## [2121,] "Bacteria" "Bacteroidetes"       "Bacteroidia"                
## [2122,] "Bacteria" "Bacteroidetes"       "Bacteroidia"                
## [2123,] "Bacteria" "Patescibacteria"     "Gracilibacteria"            
## [2124,] "Bacteria" "Firmicutes"          "Negativicutes"              
## [2125,] "Bacteria" "Bacteroidetes"       "Bacteroidia"                
## [2126,] "Bacteria" "Firmicutes"          "Bacilli"                    
## [2127,] "Bacteria" "Bacteroidetes"       "Bacteroidia"                
## [2128,] "Bacteria" "Firmicutes"          "Negativicutes"              
## [2129,] "Bacteria" "Firmicutes"          "Bacilli"                    
## [2130,] "Bacteria" "Proteobacteria"      "Gammaproteobacteria"        
## [2131,] "Bacteria" "Bacteroidetes"       "Bacteroidia"                
## [2132,] "Bacteria" "Epsilonbacteraeota"  "Campylobacteria"            
## [2133,] "Bacteria" "Actinobacteria"      "Actinobacteria"             
## [2134,] "Bacteria" "Fusobacteria"        "Fusobacteriia"              
## [2135,] "Bacteria" "Bacteroidetes"       "Bacteroidia"                
## [2136,] "Bacteria" "Firmicutes"          "Negativicutes"              
## [2137,] "Bacteria" "Actinobacteria"      "Actinobacteria"             
## [2138,] "Bacteria" "Bacteroidetes"       "Bacteroidia"                
## [2139,] "Bacteria" "Bacteroidetes"       "Bacteroidia"                
## [2140,] "Bacteria" "Firmicutes"          "Clostridia"                 
## [2141,] "Bacteria" "Spirochaetes"        "Spirochaetia"               
## [2142,] "Bacteria" "Epsilonbacteraeota"  "Campylobacteria"            
## [2143,] "Bacteria" "Proteobacteria"      "Gammaproteobacteria"        
## [2144,] "Bacteria" "Firmicutes"          "Negativicutes"              
## [2145,] "Bacteria" "Bacteroidetes"       "Bacteroidia"                
## [2146,] "Bacteria" "Bacteroidetes"       "Bacteroidia"                
## [2147,] "Bacteria" "Bacteroidetes"       "Bacteroidia"                
## [2148,] "Bacteria" "Bacteroidetes"       "Bacteroidia"                
## [2149,] "Bacteria" "Fusobacteria"        "Fusobacteriia"              
## [2150,] "Bacteria" "Proteobacteria"      "Gammaproteobacteria"        
## [2151,] "Bacteria" "Patescibacteria"     "Saccharimonadia"            
## [2152,] "Bacteria" "Proteobacteria"      "Gammaproteobacteria"        
## [2153,] "Bacteria" "Epsilonbacteraeota"  "Campylobacteria"            
## [2154,] "Bacteria" "Bacteroidetes"       "Bacteroidia"                
## [2155,] "Bacteria" "Bacteroidetes"       "Bacteroidia"                
## [2156,] "Bacteria" "Bacteroidetes"       "Bacteroidia"                
## [2157,] "Bacteria" "Proteobacteria"      "Gammaproteobacteria"        
## [2158,] "Bacteria" NA                    NA                           
## [2159,] "Bacteria" "Bacteroidetes"       "Bacteroidia"                
## [2160,] "Bacteria" "Firmicutes"          "Bacilli"                    
## [2161,] "Bacteria" "Proteobacteria"      "Gammaproteobacteria"        
## [2162,] "Bacteria" "Actinobacteria"      "Actinobacteria"             
## [2163,] "Bacteria" "Fusobacteria"        "Fusobacteriia"              
## [2164,] "Bacteria" "Bacteroidetes"       "Bacteroidia"                
## [2165,] "Bacteria" "Firmicutes"          "Negativicutes"              
## [2166,] "Bacteria" "Bacteroidetes"       "Bacteroidia"                
## [2167,] "Bacteria" "Bacteroidetes"       "Bacteroidia"                
## [2168,] "Bacteria" "Spirochaetes"        "Spirochaetia"               
## [2169,] "Bacteria" "Bacteroidetes"       "Bacteroidia"                
## [2170,] "Bacteria" "Firmicutes"          "Clostridia"                 
## [2171,] "Bacteria" "Fusobacteria"        "Fusobacteriia"              
## [2172,] "Bacteria" "Firmicutes"          "Clostridia"                 
## [2173,] "Bacteria" "Actinobacteria"      "Actinobacteria"             
## [2174,] "Bacteria" "Bacteroidetes"       "Bacteroidia"                
## [2175,] "Bacteria" "Bacteroidetes"       "Bacteroidia"                
## [2176,] "Bacteria" "Fusobacteria"        "Fusobacteriia"              
## [2177,] "Bacteria" "Bacteroidetes"       "Bacteroidia"                
## [2178,] "Bacteria" "Firmicutes"          "Clostridia"                 
## [2179,] "Bacteria" "Firmicutes"          "Negativicutes"              
## [2180,] "Bacteria" "Firmicutes"          "Negativicutes"              
## [2181,] "Bacteria" "Fusobacteria"        "Fusobacteriia"              
## [2182,] "Bacteria" "Firmicutes"          "Clostridia"                 
## [2183,] "Bacteria" "Bacteroidetes"       "Bacteroidia"                
## [2184,] "Bacteria" "Bacteroidetes"       "Bacteroidia"                
## [2185,] "Bacteria" "Bacteroidetes"       "Bacteroidia"                
## [2186,] "Bacteria" "Fusobacteria"        "Fusobacteriia"              
## [2187,] "Bacteria" "Firmicutes"          "Negativicutes"              
## [2188,] "Bacteria" "Firmicutes"          "Negativicutes"              
## [2189,] "Bacteria" "Fusobacteria"        "Fusobacteriia"              
## [2190,] "Bacteria" "Firmicutes"          "Negativicutes"              
## [2191,] "Bacteria" "Bacteroidetes"       "Bacteroidia"                
## [2192,] "Bacteria" "Proteobacteria"      "Gammaproteobacteria"        
## [2193,] "Bacteria" "Bacteroidetes"       "Bacteroidia"                
## [2194,] "Bacteria" "Firmicutes"          "Negativicutes"              
## [2195,] "Bacteria" "Proteobacteria"      "Gammaproteobacteria"        
## [2196,] "Bacteria" "Patescibacteria"     "Saccharimonadia"            
## [2197,] "Bacteria" "Bacteroidetes"       "Bacteroidia"                
## [2198,] "Bacteria" "Fusobacteria"        "Fusobacteriia"              
## [2199,] "Bacteria" "Bacteroidetes"       "Bacteroidia"                
## [2200,] "Bacteria" "Firmicutes"          "Negativicutes"              
## [2201,] "Bacteria" "Actinobacteria"      "Actinobacteria"             
## [2202,] "Bacteria" "Bacteroidetes"       "Bacteroidia"                
## [2203,] "Bacteria" "Firmicutes"          "Negativicutes"              
## [2204,] "Bacteria" "Fusobacteria"        "Fusobacteriia"              
## [2205,] "Bacteria" "Firmicutes"          "Negativicutes"              
## [2206,] "Bacteria" "Proteobacteria"      "Gammaproteobacteria"        
## [2207,] "Bacteria" "Bacteroidetes"       "Bacteroidia"                
## [2208,] "Bacteria" "Proteobacteria"      "Gammaproteobacteria"        
## [2209,] "Bacteria" "Patescibacteria"     "Saccharimonadia"            
## [2210,] "Bacteria" "Spirochaetes"        "Spirochaetia"               
## [2211,] "Bacteria" "Firmicutes"          "Clostridia"                 
## [2212,] "Bacteria" "Bacteroidetes"       "Bacteroidia"                
## [2213,] "Bacteria" "Bacteroidetes"       "Bacteroidia"                
## [2214,] "Bacteria" "Firmicutes"          "Negativicutes"              
## [2215,] "Bacteria" "Bacteroidetes"       "Bacteroidia"                
## [2216,] "Bacteria" "Bacteroidetes"       "Bacteroidia"                
## [2217,] "Bacteria" "Bacteroidetes"       "Bacteroidia"                
## [2218,] "Bacteria" "Firmicutes"          "Negativicutes"              
## [2219,] "Bacteria" "Bacteroidetes"       "Bacteroidia"                
## [2220,] "Bacteria" "Bacteroidetes"       "Bacteroidia"                
## [2221,] "Bacteria" "Patescibacteria"     "Saccharimonadia"            
## [2222,] "Bacteria" "Bacteroidetes"       "Bacteroidia"                
## [2223,] "Bacteria" "Cyanobacteria"       "Oxyphotobacteria"           
## [2224,] "Bacteria" "Bacteroidetes"       "Bacteroidia"                
## [2225,] "Bacteria" "Firmicutes"          "Negativicutes"              
## [2226,] "Bacteria" "Patescibacteria"     "Saccharimonadia"            
## [2227,] "Bacteria" "Firmicutes"          "Bacilli"                    
## [2228,] "Bacteria" "Bacteroidetes"       "Bacteroidia"                
## [2229,] "Bacteria" "Firmicutes"          "Clostridia"                 
## [2230,] "Bacteria" NA                    NA                           
## [2231,] "Bacteria" "Actinobacteria"      "Actinobacteria"             
## [2232,] "Bacteria" "Firmicutes"          "Negativicutes"              
## [2233,] "Bacteria" "Bacteroidetes"       "Bacteroidia"                
## [2234,] "Bacteria" "Firmicutes"          "Negativicutes"              
## [2235,] "Bacteria" "Firmicutes"          "Clostridia"                 
## [2236,] "Bacteria" "Firmicutes"          "Negativicutes"              
## [2237,] "Bacteria" "Bacteroidetes"       "Bacteroidia"                
## [2238,] "Bacteria" "Fusobacteria"        "Fusobacteriia"              
## [2239,] "Bacteria" "Firmicutes"          "Negativicutes"              
## [2240,] "Bacteria" "Actinobacteria"      "Actinobacteria"             
## [2241,] "Bacteria" "Bacteroidetes"       "Bacteroidia"                
## [2242,] "Bacteria" "Cyanobacteria"       "Oxyphotobacteria"           
## [2243,] "Bacteria" "Bacteroidetes"       "Bacteroidia"                
## [2244,] "Bacteria" "Bacteroidetes"       "Bacteroidia"                
## [2245,] "Bacteria" "Proteobacteria"      "Gammaproteobacteria"        
## [2246,] "Bacteria" "Bacteroidetes"       "Bacteroidia"                
## [2247,] "Bacteria" "Bacteroidetes"       "Bacteroidia"                
## [2248,] "Bacteria" "Bacteroidetes"       "Bacteroidia"                
## [2249,] "Bacteria" NA                    NA                           
## [2250,] "Bacteria" "Patescibacteria"     "Saccharimonadia"            
## [2251,] "Bacteria" "Actinobacteria"      "Coriobacteriia"             
## [2252,] "Bacteria" "Bacteroidetes"       "Bacteroidia"                
## [2253,] "Bacteria" "Firmicutes"          "Negativicutes"              
## [2254,] "Bacteria" "Firmicutes"          "Negativicutes"              
## [2255,] "Bacteria" "Synergistetes"       "Synergistia"                
## [2256,] "Bacteria" "Bacteroidetes"       "Bacteroidia"                
## [2257,] "Bacteria" "Bacteroidetes"       "Bacteroidia"                
## [2258,] "Bacteria" "Firmicutes"          "Clostridia"                 
## [2259,] "Bacteria" "Actinobacteria"      "Actinobacteria"             
## [2260,] "Bacteria" "Firmicutes"          "Bacilli"                    
## [2261,] "Bacteria" "Firmicutes"          "Negativicutes"              
## [2262,] "Bacteria" "Proteobacteria"      "Gammaproteobacteria"        
## [2263,] "Bacteria" "Proteobacteria"      "Gammaproteobacteria"        
## [2264,] "Bacteria" "Fusobacteria"        "Fusobacteriia"              
## [2265,] "Bacteria" "Bacteroidetes"       "Bacteroidia"                
## [2266,] "Bacteria" "Fusobacteria"        "Fusobacteriia"              
## [2267,] "Bacteria" "Firmicutes"          "Negativicutes"              
## [2268,] "Bacteria" "Proteobacteria"      "Gammaproteobacteria"        
## [2269,] "Bacteria" "Bacteroidetes"       "Bacteroidia"                
## [2270,] "Bacteria" "Bacteroidetes"       "Bacteroidia"                
## [2271,] "Bacteria" "Fusobacteria"        "Fusobacteriia"              
## [2272,] "Bacteria" "Patescibacteria"     "Gracilibacteria"            
## [2273,] "Bacteria" "Fusobacteria"        "Fusobacteriia"              
## [2274,] "Bacteria" "Firmicutes"          "Clostridia"                 
## [2275,] "Bacteria" "Bacteroidetes"       "Bacteroidia"                
## [2276,] "Bacteria" "Bacteroidetes"       "Bacteroidia"                
## [2277,] "Bacteria" "Firmicutes"          "Clostridia"                 
## [2278,] "Bacteria" "Firmicutes"          "Negativicutes"              
## [2279,] "Bacteria" "Proteobacteria"      "Gammaproteobacteria"        
## [2280,] "Bacteria" "Firmicutes"          NA                           
## [2281,] "Bacteria" "Proteobacteria"      "Gammaproteobacteria"        
## [2282,] "Bacteria" "Firmicutes"          "Clostridia"                 
## [2283,] "Bacteria" "Bacteroidetes"       "Bacteroidia"                
## [2284,] "Bacteria" "Bacteroidetes"       "Bacteroidia"                
## [2285,] "Bacteria" "Bacteroidetes"       "Bacteroidia"                
## [2286,] "Bacteria" "Firmicutes"          "Clostridia"                 
## [2287,] "Bacteria" "Bacteroidetes"       "Bacteroidia"                
## [2288,] "Bacteria" "Bacteroidetes"       "Bacteroidia"                
## [2289,] "Bacteria" "Patescibacteria"     "Gracilibacteria"            
## [2290,] "Bacteria" "Fusobacteria"        "Fusobacteriia"              
## [2291,] "Bacteria" "Firmicutes"          "Negativicutes"              
## [2292,] "Bacteria" "Proteobacteria"      "Gammaproteobacteria"        
## [2293,] "Bacteria" "Bacteroidetes"       "Bacteroidia"                
## [2294,] "Bacteria" "Bacteroidetes"       "Bacteroidia"                
## [2295,] "Bacteria" "Bacteroidetes"       "Bacteroidia"                
## [2296,] "Bacteria" "Proteobacteria"      "Gammaproteobacteria"        
## [2297,] "Bacteria" "Bacteroidetes"       "Bacteroidia"                
## [2298,] "Bacteria" "Bacteroidetes"       "Bacteroidia"                
## [2299,] "Bacteria" "Bacteroidetes"       "Bacteroidia"                
## [2300,] "Bacteria" "Bacteroidetes"       "Bacteroidia"                
## [2301,] "Bacteria" "Firmicutes"          "Negativicutes"              
## [2302,] "Bacteria" "Firmicutes"          "Clostridia"                 
## [2303,] "Bacteria" "Actinobacteria"      "Actinobacteria"             
## [2304,] "Bacteria" "Firmicutes"          "Clostridia"                 
## [2305,] "Bacteria" "Spirochaetes"        "Spirochaetia"               
## [2306,] "Bacteria" "Bacteroidetes"       "Bacteroidia"                
## [2307,] "Bacteria" "Patescibacteria"     "Saccharimonadia"            
## [2308,] "Bacteria" "Bacteroidetes"       "Bacteroidia"                
## [2309,] "Bacteria" "Firmicutes"          "Clostridia"                 
## [2310,] "Bacteria" "Bacteroidetes"       "Bacteroidia"                
## [2311,] "Bacteria" "Firmicutes"          "Clostridia"                 
## [2312,] "Bacteria" "Firmicutes"          "Negativicutes"              
## [2313,] "Bacteria" "Actinobacteria"      "Actinobacteria"             
## [2314,] "Bacteria" "Bacteroidetes"       "Bacteroidia"                
## [2315,] "Bacteria" "Fusobacteria"        "Fusobacteriia"              
## [2316,] "Bacteria" "Actinobacteria"      "Actinobacteria"             
## [2317,] "Bacteria" "Firmicutes"          NA                           
## [2318,] "Bacteria" "Bacteroidetes"       "Bacteroidia"                
## [2319,] "Bacteria" "Firmicutes"          "Clostridia"                 
## [2320,] "Bacteria" "Patescibacteria"     "Saccharimonadia"            
## [2321,] "Bacteria" "Bacteroidetes"       "Bacteroidia"                
## [2322,] "Bacteria" "Bacteroidetes"       "Bacteroidia"                
## [2323,] "Bacteria" "Bacteroidetes"       "Bacteroidia"                
## [2324,] "Bacteria" "Bacteroidetes"       "Bacteroidia"                
## [2325,] "Bacteria" "Bacteroidetes"       "Bacteroidia"                
## [2326,] "Bacteria" "Firmicutes"          NA                           
## [2327,] "Bacteria" "Fusobacteria"        "Fusobacteriia"              
## [2328,] "Bacteria" "Bacteroidetes"       "Bacteroidia"                
## [2329,] "Bacteria" "Firmicutes"          "Clostridia"                 
## [2330,] "Bacteria" "Bacteroidetes"       "Bacteroidia"                
## [2331,] "Bacteria" "Bacteroidetes"       "Bacteroidia"                
## [2332,] "Bacteria" "Bacteroidetes"       "Bacteroidia"                
## [2333,] "Bacteria" "Bacteroidetes"       "Bacteroidia"                
## [2334,] "Bacteria" "Bacteroidetes"       "Bacteroidia"                
## [2335,] "Bacteria" "Proteobacteria"      "Gammaproteobacteria"        
## [2336,] "Bacteria" "Proteobacteria"      "Gammaproteobacteria"        
## [2337,] "Bacteria" "Bacteroidetes"       "Bacteroidia"                
## [2338,] "Bacteria" "Bacteroidetes"       "Bacteroidia"                
## [2339,] "Bacteria" "Firmicutes"          "Bacilli"                    
## [2340,] "Bacteria" "Bacteroidetes"       "Bacteroidia"                
## [2341,] "Bacteria" "Patescibacteria"     "Saccharimonadia"            
## [2342,] "Bacteria" "Firmicutes"          "Clostridia"                 
## [2343,] "Bacteria" "Proteobacteria"      "Gammaproteobacteria"        
## [2344,] "Bacteria" "Bacteroidetes"       "Bacteroidia"                
## [2345,] "Bacteria" "Bacteroidetes"       "Bacteroidia"                
## [2346,] "Bacteria" "Actinobacteria"      "Actinobacteria"             
## [2347,] "Bacteria" "Firmicutes"          "Negativicutes"              
## [2348,] "Bacteria" "Actinobacteria"      "Actinobacteria"             
## [2349,] "Bacteria" "Synergistetes"       "Synergistia"                
## [2350,] "Bacteria" "Spirochaetes"        "Spirochaetia"               
## [2351,] "Bacteria" "Bacteroidetes"       "Bacteroidia"                
## [2352,] "Bacteria" "Bacteroidetes"       "Bacteroidia"                
## [2353,] "Bacteria" "Actinobacteria"      "Actinobacteria"             
## [2354,] "Bacteria" "Actinobacteria"      "Actinobacteria"             
## [2355,] "Bacteria" "Firmicutes"          "Clostridia"                 
## [2356,] "Bacteria" "Bacteroidetes"       "Bacteroidia"                
## [2357,] "Bacteria" "Firmicutes"          "Negativicutes"              
## [2358,] "Bacteria" "Bacteroidetes"       "Bacteroidia"                
## [2359,] "Bacteria" "Bacteroidetes"       "Bacteroidia"                
## [2360,] "Bacteria" "Firmicutes"          "Negativicutes"              
## [2361,] "Bacteria" "Proteobacteria"      "Gammaproteobacteria"        
## [2362,] "Bacteria" "Firmicutes"          "Negativicutes"              
## [2363,] "Bacteria" "Bacteroidetes"       "Bacteroidia"                
## [2364,] "Bacteria" "Firmicutes"          "Negativicutes"              
## [2365,] "Bacteria" "Firmicutes"          "Negativicutes"              
## [2366,] "Bacteria" "Fusobacteria"        "Fusobacteriia"              
## [2367,] "Bacteria" "Actinobacteria"      "Actinobacteria"             
## [2368,] "Bacteria" "Bacteroidetes"       "Bacteroidia"                
## [2369,] "Bacteria" "Proteobacteria"      "Gammaproteobacteria"        
## [2370,] "Bacteria" "Fusobacteria"        "Fusobacteriia"              
## [2371,] "Bacteria" "Bacteroidetes"       "Bacteroidia"                
## [2372,] "Bacteria" "Firmicutes"          "Negativicutes"              
## [2373,] "Bacteria" "Bacteroidetes"       "Bacteroidia"                
## [2374,] "Bacteria" "Bacteroidetes"       "Bacteroidia"                
## [2375,] "Bacteria" "Firmicutes"          "Negativicutes"              
## [2376,] "Bacteria" "Firmicutes"          "Clostridia"                 
## [2377,] "Bacteria" "Proteobacteria"      "Gammaproteobacteria"        
## [2378,] "Bacteria" "Spirochaetes"        "Spirochaetia"               
## [2379,] "Bacteria" "Bacteroidetes"       "Bacteroidia"                
## [2380,] "Bacteria" "Fusobacteria"        "Fusobacteriia"              
## [2381,] "Bacteria" "Bacteroidetes"       "Bacteroidia"                
## [2382,] "Bacteria" "Fusobacteria"        "Fusobacteriia"              
## [2383,] "Bacteria" "Firmicutes"          "Clostridia"                 
## [2384,] "Bacteria" "Bacteroidetes"       "Bacteroidia"                
## [2385,] "Bacteria" "Proteobacteria"      "Gammaproteobacteria"        
## [2386,] "Bacteria" "Bacteroidetes"       "Bacteroidia"                
## [2387,] "Bacteria" "Firmicutes"          "Negativicutes"              
## [2388,] "Bacteria" "Actinobacteria"      "Actinobacteria"             
## [2389,] "Bacteria" "Bacteroidetes"       "Bacteroidia"                
## [2390,] "Bacteria" "Firmicutes"          "Negativicutes"              
## [2391,] "Bacteria" "Fusobacteria"        "Fusobacteriia"              
## [2392,] "Bacteria" "Firmicutes"          "Negativicutes"              
## [2393,] "Bacteria" "Fusobacteria"        "Fusobacteriia"              
## [2394,] "Bacteria" "Patescibacteria"     "Saccharimonadia"            
## [2395,] "Bacteria" "Firmicutes"          "Negativicutes"              
## [2396,] "Bacteria" "Firmicutes"          "Negativicutes"              
## [2397,] "Bacteria" "Firmicutes"          "Negativicutes"              
## [2398,] "Bacteria" "Actinobacteria"      "Actinobacteria"             
## [2399,] "Bacteria" "Actinobacteria"      "Actinobacteria"             
## [2400,] "Bacteria" "Bacteroidetes"       "Bacteroidia"                
## [2401,] "Bacteria" "Actinobacteria"      "Coriobacteriia"             
## [2402,] "Bacteria" "Spirochaetes"        "Spirochaetia"               
## [2403,] "Bacteria" "Bacteroidetes"       "Bacteroidia"                
## [2404,] "Bacteria" "Fusobacteria"        "Fusobacteriia"              
## [2405,] "Bacteria" "Firmicutes"          "Clostridia"                 
## [2406,] "Bacteria" "Firmicutes"          "Negativicutes"              
## [2407,] "Bacteria" "Bacteroidetes"       "Bacteroidia"                
## [2408,] "Bacteria" "Bacteroidetes"       "Bacteroidia"                
## [2409,] "Bacteria" "Fusobacteria"        "Fusobacteriia"              
## [2410,] "Bacteria" "Bacteroidetes"       "Bacteroidia"                
## [2411,] "Bacteria" "Firmicutes"          "Negativicutes"              
## [2412,] "Bacteria" "Firmicutes"          NA                           
## [2413,] "Bacteria" "Patescibacteria"     "Saccharimonadia"            
## [2414,] "Bacteria" "Bacteroidetes"       "Bacteroidia"                
## [2415,] "Bacteria" "Firmicutes"          "Negativicutes"              
## [2416,] "Bacteria" "Firmicutes"          "Negativicutes"              
## [2417,] "Bacteria" "Firmicutes"          "Negativicutes"              
## [2418,] "Bacteria" "Bacteroidetes"       "Bacteroidia"                
## [2419,] "Bacteria" "Bacteroidetes"       "Bacteroidia"                
## [2420,] "Bacteria" "Firmicutes"          "Clostridia"                 
## [2421,] "Bacteria" "Proteobacteria"      "Gammaproteobacteria"        
## [2422,] "Bacteria" "Actinobacteria"      "Actinobacteria"             
## [2423,] "Bacteria" "Actinobacteria"      "Actinobacteria"             
## [2424,] "Bacteria" "Firmicutes"          NA                           
## [2425,] "Bacteria" "Bacteroidetes"       "Bacteroidia"                
## [2426,] "Bacteria" "Cyanobacteria"       "Oxyphotobacteria"           
## [2427,] "Bacteria" "Bacteroidetes"       "Bacteroidia"                
## [2428,] "Bacteria" "Bacteroidetes"       "Bacteroidia"                
## [2429,] "Bacteria" "Bacteroidetes"       "Bacteroidia"                
## [2430,] "Bacteria" "Bacteroidetes"       "Bacteroidia"                
## [2431,] "Bacteria" "Bacteroidetes"       "Bacteroidia"                
## [2432,] "Bacteria" "Firmicutes"          "Clostridia"                 
## [2433,] "Bacteria" "Firmicutes"          "Negativicutes"              
## [2434,] "Bacteria" "Bacteroidetes"       "Bacteroidia"                
## [2435,] "Bacteria" "Actinobacteria"      "Actinobacteria"             
## [2436,] "Bacteria" "Bacteroidetes"       "Bacteroidia"                
## [2437,] "Bacteria" "Spirochaetes"        "Spirochaetia"               
## [2438,] "Bacteria" "Firmicutes"          NA                           
## [2439,] "Bacteria" "Bacteroidetes"       "Bacteroidia"                
## [2440,] "Bacteria" "Cyanobacteria"       "Oxyphotobacteria"           
## [2441,] "Bacteria" "Actinobacteria"      "Actinobacteria"             
## [2442,] "Bacteria" "Bacteroidetes"       "Bacteroidia"                
## [2443,] "Bacteria" "Proteobacteria"      "Gammaproteobacteria"        
## [2444,] "Bacteria" "Proteobacteria"      "Gammaproteobacteria"        
## [2445,] "Bacteria" "Firmicutes"          "Clostridia"                 
## [2446,] "Bacteria" "Bacteroidetes"       "Bacteroidia"                
## [2447,] "Bacteria" "Bacteroidetes"       "Bacteroidia"                
## [2448,] "Bacteria" "Firmicutes"          "Bacilli"                    
## [2449,] "Bacteria" "Patescibacteria"     "Saccharimonadia"            
## [2450,] "Bacteria" "Bacteroidetes"       "Bacteroidia"                
## [2451,] "Bacteria" "Firmicutes"          NA                           
## [2452,] "Bacteria" "Bacteroidetes"       "Bacteroidia"                
## [2453,] "Bacteria" "Proteobacteria"      "Gammaproteobacteria"        
## [2454,] "Bacteria" "Bacteroidetes"       "Bacteroidia"                
## [2455,] "Bacteria" "Firmicutes"          "Clostridia"                 
## [2456,] "Bacteria" "Actinobacteria"      "Actinobacteria"             
## [2457,] "Bacteria" "Proteobacteria"      "Gammaproteobacteria"        
## [2458,] "Bacteria" "Bacteroidetes"       "Bacteroidia"                
## [2459,] "Bacteria" "Firmicutes"          "Clostridia"                 
## [2460,] "Bacteria" "Spirochaetes"        "Spirochaetia"               
## [2461,] "Bacteria" "Firmicutes"          "Clostridia"                 
## [2462,] "Bacteria" "Firmicutes"          "Negativicutes"              
## [2463,] "Bacteria" "Bacteroidetes"       "Bacteroidia"                
## [2464,] "Bacteria" "Firmicutes"          "Clostridia"                 
## [2465,] "Bacteria" "Bacteroidetes"       "Bacteroidia"                
## [2466,] "Bacteria" "Bacteroidetes"       "Bacteroidia"                
## [2467,] "Bacteria" "Proteobacteria"      "Gammaproteobacteria"        
## [2468,] "Bacteria" "Bacteroidetes"       "Bacteroidia"                
## [2469,] "Bacteria" "Bacteroidetes"       "Bacteroidia"                
## [2470,] "Bacteria" "Bacteroidetes"       "Bacteroidia"                
## [2471,] "Bacteria" "Proteobacteria"      "Gammaproteobacteria"        
## [2472,] "Bacteria" "Proteobacteria"      "Gammaproteobacteria"        
## [2473,] "Bacteria" "Firmicutes"          "Negativicutes"              
## [2474,] "Bacteria" "Firmicutes"          "Clostridia"                 
## [2475,] "Bacteria" "Bacteroidetes"       "Bacteroidia"                
## [2476,] "Bacteria" "Bacteroidetes"       "Bacteroidia"                
## [2477,] "Bacteria" "Patescibacteria"     "Saccharimonadia"            
## [2478,] "Bacteria" "Firmicutes"          "Negativicutes"              
## [2479,] "Bacteria" "Firmicutes"          "Negativicutes"              
## [2480,] "Bacteria" "Actinobacteria"      "Actinobacteria"             
## [2481,] "Bacteria" "Actinobacteria"      "Actinobacteria"             
## [2482,] "Bacteria" "Firmicutes"          "Clostridia"                 
## [2483,] "Bacteria" "Bacteroidetes"       "Bacteroidia"                
## [2484,] "Bacteria" "Spirochaetes"        "Spirochaetia"               
## [2485,] "Bacteria" "Proteobacteria"      "Gammaproteobacteria"        
## [2486,] "Bacteria" "Firmicutes"          "Clostridia"                 
## [2487,] "Bacteria" "Bacteroidetes"       "Bacteroidia"                
## [2488,] "Bacteria" "Bacteroidetes"       "Bacteroidia"                
## [2489,] "Bacteria" "Proteobacteria"      "Gammaproteobacteria"        
## [2490,] "Bacteria" "Bacteroidetes"       "Bacteroidia"                
## [2491,] "Bacteria" "Fusobacteria"        "Fusobacteriia"              
## [2492,] "Bacteria" "Firmicutes"          "Negativicutes"              
## [2493,] "Bacteria" "Bacteroidetes"       "Bacteroidia"                
## [2494,] "Bacteria" "Bacteroidetes"       "Bacteroidia"                
## [2495,] "Bacteria" "Bacteroidetes"       "Bacteroidia"                
## [2496,] "Bacteria" "Fusobacteria"        "Fusobacteriia"              
## [2497,] "Bacteria" "Actinobacteria"      "Actinobacteria"             
## [2498,] "Bacteria" "Proteobacteria"      "Gammaproteobacteria"        
## [2499,] "Bacteria" "Bacteroidetes"       "Bacteroidia"                
## [2500,] "Bacteria" "Actinobacteria"      "Actinobacteria"             
## [2501,] "Bacteria" "Firmicutes"          "Negativicutes"              
## [2502,] "Bacteria" "Bacteroidetes"       "Bacteroidia"                
## [2503,] "Bacteria" "Bacteroidetes"       "Bacteroidia"                
## [2504,] "Bacteria" "Proteobacteria"      "Gammaproteobacteria"        
## [2505,] "Bacteria" "Fusobacteria"        "Fusobacteriia"              
## [2506,] "Bacteria" "Bacteroidetes"       "Bacteroidia"                
## [2507,] "Bacteria" "Bacteroidetes"       "Bacteroidia"                
## [2508,] "Bacteria" "Bacteroidetes"       "Bacteroidia"                
## [2509,] "Bacteria" "Firmicutes"          "Bacilli"                    
## [2510,] "Bacteria" "Patescibacteria"     "Saccharimonadia"            
## [2511,] "Bacteria" "Bacteroidetes"       "Bacteroidia"                
## [2512,] "Bacteria" "Proteobacteria"      "Gammaproteobacteria"        
## [2513,] "Bacteria" "Firmicutes"          "Clostridia"                 
## [2514,] "Bacteria" "Firmicutes"          "Clostridia"                 
## [2515,] "Bacteria" "Bacteroidetes"       "Bacteroidia"                
## [2516,] "Bacteria" "Proteobacteria"      "Gammaproteobacteria"        
## [2517,] "Bacteria" "Actinobacteria"      "Actinobacteria"             
## [2518,] "Bacteria" "Fusobacteria"        "Fusobacteriia"              
## [2519,] "Bacteria" "Actinobacteria"      "Actinobacteria"             
## [2520,] "Bacteria" "Actinobacteria"      "Actinobacteria"             
## [2521,] "Bacteria" "Proteobacteria"      "Gammaproteobacteria"        
## [2522,] "Bacteria" "Fusobacteria"        "Fusobacteriia"              
## [2523,] "Bacteria" "Bacteroidetes"       "Bacteroidia"                
## [2524,] "Bacteria" "Firmicutes"          "Negativicutes"              
## [2525,] "Bacteria" "Proteobacteria"      "Gammaproteobacteria"        
## [2526,] "Bacteria" "Actinobacteria"      "Actinobacteria"             
## [2527,] "Bacteria" "Bacteroidetes"       "Bacteroidia"                
## [2528,] "Bacteria" "Firmicutes"          "Bacilli"                    
## [2529,] "Bacteria" "Fusobacteria"        "Fusobacteriia"              
## [2530,] "Bacteria" "Bacteroidetes"       "Bacteroidia"                
## [2531,] "Bacteria" "Firmicutes"          "Negativicutes"              
## [2532,] "Bacteria" "Firmicutes"          "Negativicutes"              
## [2533,] "Bacteria" "Bacteroidetes"       "Bacteroidia"                
## [2534,] "Bacteria" "Bacteroidetes"       "Bacteroidia"                
## [2535,] "Bacteria" "Bacteroidetes"       "Bacteroidia"                
## [2536,] "Bacteria" "Proteobacteria"      "Gammaproteobacteria"        
## [2537,] "Bacteria" "Firmicutes"          "Bacilli"                    
## [2538,] "Bacteria" "Bacteroidetes"       "Bacteroidia"                
## [2539,] "Bacteria" "Actinobacteria"      "Actinobacteria"             
## [2540,] "Bacteria" "Proteobacteria"      "Gammaproteobacteria"        
## [2541,] "Bacteria" "Bacteroidetes"       "Bacteroidia"                
## [2542,] "Bacteria" "Firmicutes"          "Clostridia"                 
## [2543,] "Bacteria" "Fusobacteria"        "Fusobacteriia"              
## [2544,] "Bacteria" "Bacteroidetes"       "Bacteroidia"                
## [2545,] "Bacteria" "Tenericutes"         "Mollicutes"                 
## [2546,] "Bacteria" "Fusobacteria"        "Fusobacteriia"              
## [2547,] "Bacteria" "Fusobacteria"        "Fusobacteriia"              
## [2548,] "Bacteria" "Actinobacteria"      "Actinobacteria"             
## [2549,] "Bacteria" "Firmicutes"          "Negativicutes"              
## [2550,] "Bacteria" "Firmicutes"          "Clostridia"                 
## [2551,] "Bacteria" "Firmicutes"          "Negativicutes"              
## [2552,] "Bacteria" "Bacteroidetes"       "Bacteroidia"                
## [2553,] "Bacteria" "Fusobacteria"        "Fusobacteriia"              
## [2554,] "Bacteria" "Fusobacteria"        "Fusobacteriia"              
## [2555,] "Bacteria" "Fusobacteria"        "Fusobacteriia"              
## [2556,] "Bacteria" "Bacteroidetes"       "Bacteroidia"                
## [2557,] "Bacteria" "Proteobacteria"      "Gammaproteobacteria"        
## [2558,] "Bacteria" "Spirochaetes"        "Spirochaetia"               
## [2559,] "Bacteria" "Bacteroidetes"       "Bacteroidia"                
## [2560,] "Bacteria" "Firmicutes"          "Negativicutes"              
## [2561,] "Bacteria" "Firmicutes"          "Negativicutes"              
## [2562,] "Bacteria" "Spirochaetes"        "Spirochaetia"               
## [2563,] "Bacteria" "Spirochaetes"        "Spirochaetia"               
## [2564,] "Bacteria" "Proteobacteria"      "Gammaproteobacteria"        
## [2565,] "Bacteria" "Firmicutes"          "Clostridia"                 
## [2566,] "Bacteria" "Firmicutes"          "Bacilli"                    
## [2567,] "Bacteria" "Proteobacteria"      "Gammaproteobacteria"        
## [2568,] "Bacteria" "Firmicutes"          "Bacilli"                    
## [2569,] "Bacteria" "Bacteroidetes"       "Bacteroidia"                
## [2570,] "Bacteria" "Firmicutes"          "Negativicutes"              
## [2571,] "Bacteria" "Bacteroidetes"       "Bacteroidia"                
## [2572,] "Bacteria" "Bacteroidetes"       "Bacteroidia"                
## [2573,] "Bacteria" "Proteobacteria"      "Gammaproteobacteria"        
## [2574,] "Bacteria" "Fusobacteria"        "Fusobacteriia"              
## [2575,] "Bacteria" "Patescibacteria"     "Saccharimonadia"            
## [2576,] "Bacteria" "Bacteroidetes"       "Bacteroidia"                
## [2577,] "Bacteria" "Actinobacteria"      "Actinobacteria"             
## [2578,] "Bacteria" "Fusobacteria"        "Fusobacteriia"              
## [2579,] "Bacteria" "Bacteroidetes"       "Bacteroidia"                
## [2580,] "Bacteria" "Bacteroidetes"       "Bacteroidia"                
## [2581,] "Bacteria" "Bacteroidetes"       "Bacteroidia"                
## [2582,] "Bacteria" "Fusobacteria"        "Fusobacteriia"              
## [2583,] "Bacteria" "Bacteroidetes"       "Bacteroidia"                
## [2584,] "Bacteria" "Firmicutes"          "Negativicutes"              
## [2585,] "Bacteria" "Bacteroidetes"       "Bacteroidia"                
## [2586,] "Bacteria" "Patescibacteria"     "Saccharimonadia"            
## [2587,] "Bacteria" "Bacteroidetes"       "Bacteroidia"                
## [2588,] "Bacteria" "Firmicutes"          "Clostridia"                 
## [2589,] "Bacteria" "Firmicutes"          "Clostridia"                 
## [2590,] "Bacteria" "Bacteroidetes"       "Bacteroidia"                
## [2591,] "Bacteria" "Epsilonbacteraeota"  "Campylobacteria"            
## [2592,] "Bacteria" "Firmicutes"          "Negativicutes"              
## [2593,] "Bacteria" "Firmicutes"          "Bacilli"                    
## [2594,] "Bacteria" "Proteobacteria"      "Gammaproteobacteria"        
## [2595,] "Bacteria" "Bacteroidetes"       "Bacteroidia"                
## [2596,] "Bacteria" "Firmicutes"          "Bacilli"                    
## [2597,] "Bacteria" "Bacteroidetes"       "Bacteroidia"                
## [2598,] "Bacteria" "Firmicutes"          NA                           
## [2599,] "Bacteria" "Bacteroidetes"       "Bacteroidia"                
## [2600,] "Bacteria" "Firmicutes"          "Negativicutes"              
## [2601,] "Bacteria" "Firmicutes"          "Negativicutes"              
## [2602,] "Bacteria" "Proteobacteria"      "Gammaproteobacteria"        
## [2603,] "Bacteria" "Bacteroidetes"       "Bacteroidia"                
## [2604,] "Bacteria" "Firmicutes"          "Negativicutes"              
## [2605,] "Bacteria" "Bacteroidetes"       "Bacteroidia"                
## [2606,] "Bacteria" "Proteobacteria"      "Gammaproteobacteria"        
## [2607,] "Bacteria" "Actinobacteria"      "Actinobacteria"             
## [2608,] "Bacteria" NA                    NA                           
## [2609,] "Bacteria" "Firmicutes"          NA                           
## [2610,] "Bacteria" "Actinobacteria"      "Actinobacteria"             
## [2611,] "Bacteria" "Firmicutes"          "Clostridia"                 
## [2612,] "Bacteria" "Bacteroidetes"       "Bacteroidia"                
## [2613,] "Bacteria" "Actinobacteria"      "Actinobacteria"             
## [2614,] "Bacteria" "Bacteroidetes"       "Bacteroidia"                
## [2615,] "Bacteria" "Firmicutes"          NA                           
## [2616,] "Bacteria" "Bacteroidetes"       "Bacteroidia"                
## [2617,] "Bacteria" "Proteobacteria"      "Gammaproteobacteria"        
## [2618,] "Bacteria" "Firmicutes"          "Negativicutes"              
## [2619,] "Bacteria" "Proteobacteria"      "Gammaproteobacteria"        
## [2620,] "Bacteria" "Fusobacteria"        "Fusobacteriia"              
## [2621,] "Bacteria" "Fusobacteria"        "Fusobacteriia"              
## [2622,] "Bacteria" "Bacteroidetes"       "Bacteroidia"                
## [2623,] "Bacteria" "Bacteroidetes"       "Bacteroidia"                
## [2624,] "Bacteria" "Proteobacteria"      "Gammaproteobacteria"        
## [2625,] "Bacteria" "Firmicutes"          "Clostridia"                 
## [2626,] "Bacteria" "Patescibacteria"     "Gracilibacteria"            
## [2627,] "Bacteria" "Firmicutes"          "Negativicutes"              
## [2628,] "Bacteria" "Firmicutes"          "Bacilli"                    
## [2629,] "Bacteria" "Actinobacteria"      "Actinobacteria"             
## [2630,] "Bacteria" "Firmicutes"          "Clostridia"                 
## [2631,] "Bacteria" "Firmicutes"          NA                           
## [2632,] "Bacteria" "Spirochaetes"        "Spirochaetia"               
## [2633,] "Bacteria" "Firmicutes"          "Bacilli"                    
## [2634,] "Bacteria" "Bacteroidetes"       "Bacteroidia"                
## [2635,] "Bacteria" "Epsilonbacteraeota"  "Campylobacteria"            
## [2636,] "Bacteria" "Firmicutes"          "Negativicutes"              
## [2637,] "Bacteria" "Epsilonbacteraeota"  "Campylobacteria"            
## [2638,] "Bacteria" "Epsilonbacteraeota"  "Campylobacteria"            
## [2639,] "Bacteria" "Bacteroidetes"       "Bacteroidia"                
## [2640,] "Bacteria" "Fusobacteria"        "Fusobacteriia"              
## [2641,] "Bacteria" "Epsilonbacteraeota"  "Campylobacteria"            
## [2642,] "Bacteria" "Firmicutes"          "Negativicutes"              
## [2643,] "Bacteria" "Bacteroidetes"       "Bacteroidia"                
## [2644,] "Bacteria" "Firmicutes"          "Bacilli"                    
## [2645,] "Bacteria" "Proteobacteria"      "Gammaproteobacteria"        
## [2646,] "Bacteria" "Firmicutes"          "Clostridia"                 
## [2647,] "Bacteria" "Bacteroidetes"       "Bacteroidia"                
## [2648,] "Bacteria" "Firmicutes"          "Clostridia"                 
## [2649,] "Bacteria" "Bacteroidetes"       "Bacteroidia"                
## [2650,] "Bacteria" "Firmicutes"          "Bacilli"                    
## [2651,] "Bacteria" "Spirochaetes"        "Spirochaetia"               
## [2652,] "Bacteria" "Proteobacteria"      "Gammaproteobacteria"        
## [2653,] "Bacteria" "Fusobacteria"        "Fusobacteriia"              
## [2654,] "Bacteria" "Bacteroidetes"       "Bacteroidia"                
## [2655,] "Bacteria" "Firmicutes"          "Negativicutes"              
## [2656,] "Bacteria" "Bacteroidetes"       "Bacteroidia"                
## [2657,] "Bacteria" "Bacteroidetes"       "Bacteroidia"                
## [2658,] "Bacteria" "Firmicutes"          "Clostridia"                 
## [2659,] "Bacteria" "Bacteroidetes"       "Bacteroidia"                
## [2660,] "Bacteria" "Bacteroidetes"       "Bacteroidia"                
## [2661,] "Bacteria" "Bacteroidetes"       "Bacteroidia"                
## [2662,] "Bacteria" "Firmicutes"          "Negativicutes"              
## [2663,] "Bacteria" "Actinobacteria"      "Actinobacteria"             
## [2664,] "Bacteria" "Actinobacteria"      "Actinobacteria"             
## [2665,] "Bacteria" "Fusobacteria"        "Fusobacteriia"              
## [2666,] "Bacteria" "Firmicutes"          "Negativicutes"              
## [2667,] "Bacteria" NA                    NA                           
## [2668,] "Bacteria" "Firmicutes"          "Negativicutes"              
## [2669,] "Bacteria" "Firmicutes"          "Clostridia"                 
## [2670,] "Bacteria" "Bacteroidetes"       "Bacteroidia"                
## [2671,] "Bacteria" "Firmicutes"          "Negativicutes"              
## [2672,] "Bacteria" "Proteobacteria"      "Gammaproteobacteria"        
## [2673,] "Bacteria" "Bacteroidetes"       "Bacteroidia"                
## [2674,] "Bacteria" "Bacteroidetes"       "Bacteroidia"                
## [2675,] "Bacteria" "Spirochaetes"        "Spirochaetia"               
## [2676,] "Bacteria" "Bacteroidetes"       "Bacteroidia"                
## [2677,] "Bacteria" "Actinobacteria"      "Actinobacteria"             
## [2678,] "Bacteria" "Tenericutes"         "Mollicutes"                 
## [2679,] "Bacteria" "Proteobacteria"      "Gammaproteobacteria"        
## [2680,] "Bacteria" "Bacteroidetes"       "Bacteroidia"                
## [2681,] "Bacteria" "Proteobacteria"      "Gammaproteobacteria"        
## [2682,] "Bacteria" "Bacteroidetes"       "Bacteroidia"                
## [2683,] "Bacteria" "Actinobacteria"      "Actinobacteria"             
## [2684,] "Bacteria" "Bacteroidetes"       "Bacteroidia"                
## [2685,] "Bacteria" "Bacteroidetes"       "Bacteroidia"                
## [2686,] "Bacteria" "Fusobacteria"        "Fusobacteriia"              
## [2687,] "Bacteria" "Actinobacteria"      "Coriobacteriia"             
## [2688,] "Bacteria" "Patescibacteria"     "Saccharimonadia"            
## [2689,] "Bacteria" "Actinobacteria"      "Actinobacteria"             
## [2690,] "Bacteria" NA                    NA                           
## [2691,] "Bacteria" "Firmicutes"          "Clostridia"                 
## [2692,] "Bacteria" "Fusobacteria"        "Fusobacteriia"              
## [2693,] "Bacteria" "Firmicutes"          "Clostridia"                 
## [2694,] "Bacteria" "Bacteroidetes"       "Bacteroidia"                
## [2695,] "Bacteria" "Bacteroidetes"       "Bacteroidia"                
## [2696,] "Bacteria" "Firmicutes"          NA                           
## [2697,] "Bacteria" "Firmicutes"          "Negativicutes"              
## [2698,] "Bacteria" "Firmicutes"          "Negativicutes"              
## [2699,] "Bacteria" "Firmicutes"          "Negativicutes"              
## [2700,] "Bacteria" "Bacteroidetes"       "Bacteroidia"                
## [2701,] "Bacteria" NA                    NA                           
## [2702,] "Bacteria" "Firmicutes"          "Negativicutes"              
## [2703,] "Bacteria" "Bacteroidetes"       "Bacteroidia"                
## [2704,] "Bacteria" "Bacteroidetes"       "Bacteroidia"                
## [2705,] "Bacteria" "Proteobacteria"      "Gammaproteobacteria"        
## [2706,] "Bacteria" "Fusobacteria"        "Fusobacteriia"              
## [2707,] "Bacteria" "Firmicutes"          "Negativicutes"              
## [2708,] "Bacteria" "Fusobacteria"        "Fusobacteriia"              
## [2709,] "Bacteria" "Firmicutes"          "Negativicutes"              
## [2710,] "Bacteria" "Bacteroidetes"       "Bacteroidia"                
## [2711,] "Bacteria" "Bacteroidetes"       "Bacteroidia"                
## [2712,] "Bacteria" "Fusobacteria"        "Fusobacteriia"              
## [2713,] "Bacteria" "Firmicutes"          "Negativicutes"              
## [2714,] "Bacteria" "Fusobacteria"        "Fusobacteriia"              
## [2715,] "Bacteria" "Bacteroidetes"       "Bacteroidia"                
## [2716,] "Bacteria" "Fusobacteria"        "Fusobacteriia"              
## [2717,] "Bacteria" "Fusobacteria"        "Fusobacteriia"              
## [2718,] "Bacteria" "Bacteroidetes"       "Bacteroidia"                
## [2719,] "Bacteria" "Proteobacteria"      "Gammaproteobacteria"        
## [2720,] "Bacteria" "Firmicutes"          "Clostridia"                 
## [2721,] "Bacteria" "Spirochaetes"        "Spirochaetia"               
## [2722,] "Bacteria" "Fusobacteria"        "Fusobacteriia"              
## [2723,] "Bacteria" "Firmicutes"          "Clostridia"                 
## [2724,] "Bacteria" "Bacteroidetes"       "Bacteroidia"                
## [2725,] "Bacteria" "Bacteroidetes"       "Bacteroidia"                
## [2726,] "Bacteria" "Proteobacteria"      "Gammaproteobacteria"        
## [2727,] "Bacteria" "Bacteroidetes"       "Bacteroidia"                
## [2728,] "Bacteria" "Actinobacteria"      "Actinobacteria"             
## [2729,] "Bacteria" "Bacteroidetes"       "Bacteroidia"                
## [2730,] "Bacteria" "Epsilonbacteraeota"  "Campylobacteria"            
## [2731,] "Bacteria" "Firmicutes"          "Clostridia"                 
## [2732,] "Bacteria" "Proteobacteria"      "Gammaproteobacteria"        
## [2733,] "Bacteria" "Bacteroidetes"       "Bacteroidia"                
## [2734,] "Bacteria" "Fusobacteria"        "Fusobacteriia"              
## [2735,] "Bacteria" "Fusobacteria"        "Fusobacteriia"              
## [2736,] "Bacteria" "Fusobacteria"        "Fusobacteriia"              
## [2737,] "Bacteria" "Proteobacteria"      "Gammaproteobacteria"        
## [2738,] "Bacteria" "Bacteroidetes"       "Bacteroidia"                
## [2739,] "Bacteria" "Actinobacteria"      "Actinobacteria"             
## [2740,] "Bacteria" NA                    NA                           
## [2741,] "Bacteria" "Firmicutes"          NA                           
## [2742,] "Bacteria" "Bacteroidetes"       "Bacteroidia"                
## [2743,] "Bacteria" "Firmicutes"          "Erysipelotrichia"           
## [2744,] "Bacteria" "Firmicutes"          "Negativicutes"              
## [2745,] "Bacteria" "Proteobacteria"      "Gammaproteobacteria"        
## [2746,] "Bacteria" "Proteobacteria"      "Gammaproteobacteria"        
## [2747,] "Bacteria" "Firmicutes"          "Negativicutes"              
## [2748,] "Bacteria" "Bacteroidetes"       "Bacteroidia"                
## [2749,] "Bacteria" "Bacteroidetes"       "Bacteroidia"                
## [2750,] "Bacteria" "Bacteroidetes"       "Bacteroidia"                
## [2751,] "Bacteria" "Patescibacteria"     "Saccharimonadia"            
## [2752,] "Bacteria" "Bacteroidetes"       "Bacteroidia"                
## [2753,] "Bacteria" "Bacteroidetes"       "Bacteroidia"                
## [2754,] "Bacteria" "Bacteroidetes"       "Bacteroidia"                
## [2755,] "Bacteria" "Proteobacteria"      "Gammaproteobacteria"        
## [2756,] "Bacteria" NA                    NA                           
## [2757,] "Bacteria" "Bacteroidetes"       "Bacteroidia"                
## [2758,] "Bacteria" "Fusobacteria"        "Fusobacteriia"              
## [2759,] "Bacteria" "Bacteroidetes"       "Bacteroidia"                
## [2760,] "Bacteria" "Actinobacteria"      "Coriobacteriia"             
## [2761,] "Bacteria" "Bacteroidetes"       "Bacteroidia"                
## [2762,] "Bacteria" "Bacteroidetes"       "Bacteroidia"                
## [2763,] "Bacteria" "Bacteroidetes"       "Bacteroidia"                
## [2764,] "Bacteria" "Actinobacteria"      "Actinobacteria"             
## [2765,] "Bacteria" "Actinobacteria"      "Actinobacteria"             
## [2766,] "Bacteria" "Bacteroidetes"       "Bacteroidia"                
## [2767,] "Bacteria" "Firmicutes"          "Bacilli"                    
## [2768,] "Bacteria" "Firmicutes"          "Bacilli"                    
## [2769,] "Bacteria" "Firmicutes"          "Clostridia"                 
## [2770,] "Bacteria" "Firmicutes"          "Negativicutes"              
## [2771,] "Bacteria" "Patescibacteria"     "Saccharimonadia"            
## [2772,] "Bacteria" "Firmicutes"          "Bacilli"                    
## [2773,] "Bacteria" "Bacteroidetes"       "Bacteroidia"                
## [2774,] "Bacteria" "Firmicutes"          "Negativicutes"              
## [2775,] "Bacteria" "Epsilonbacteraeota"  "Campylobacteria"            
## [2776,] "Bacteria" "Actinobacteria"      "Actinobacteria"             
## [2777,] "Bacteria" "Proteobacteria"      "Gammaproteobacteria"        
## [2778,] "Bacteria" "Actinobacteria"      "Actinobacteria"             
## [2779,] "Bacteria" "Bacteroidetes"       "Bacteroidia"                
## [2780,] "Bacteria" "Bacteroidetes"       "Bacteroidia"                
## [2781,] "Bacteria" "Firmicutes"          NA                           
## [2782,] "Bacteria" "Bacteroidetes"       "Bacteroidia"                
## [2783,] "Bacteria" "Firmicutes"          "Clostridia"                 
## [2784,] "Bacteria" "Firmicutes"          "Clostridia"                 
## [2785,] "Bacteria" "Actinobacteria"      "Actinobacteria"             
## [2786,] "Bacteria" "Actinobacteria"      "Actinobacteria"             
## [2787,] "Bacteria" "Bacteroidetes"       "Bacteroidia"                
## [2788,] "Bacteria" "Firmicutes"          "Negativicutes"              
## [2789,] "Bacteria" NA                    NA                           
## [2790,] "Bacteria" "Bacteroidetes"       "Bacteroidia"                
## [2791,] "Bacteria" "Bacteroidetes"       "Bacteroidia"                
## [2792,] "Bacteria" "Bacteroidetes"       "Bacteroidia"                
## [2793,] "Bacteria" "Firmicutes"          "Negativicutes"              
## [2794,] "Bacteria" "Firmicutes"          "Clostridia"                 
## [2795,] "Bacteria" "Proteobacteria"      "Gammaproteobacteria"        
## [2796,] "Bacteria" "Bacteroidetes"       "Bacteroidia"                
## [2797,] "Bacteria" "Bacteroidetes"       "Bacteroidia"                
## [2798,] "Bacteria" "Bacteroidetes"       "Bacteroidia"                
## [2799,] "Bacteria" "Bacteroidetes"       "Bacteroidia"                
## [2800,] "Bacteria" "Proteobacteria"      "Gammaproteobacteria"        
## [2801,] "Bacteria" "Proteobacteria"      "Gammaproteobacteria"        
## [2802,] "Bacteria" "Bacteroidetes"       "Bacteroidia"                
## [2803,] "Bacteria" "Bacteroidetes"       "Bacteroidia"                
## [2804,] "Bacteria" "Firmicutes"          "Negativicutes"              
## [2805,] "Bacteria" "Firmicutes"          "Clostridia"                 
## [2806,] "Bacteria" "Bacteroidetes"       "Bacteroidia"                
## [2807,] "Bacteria" "Bacteroidetes"       "Bacteroidia"                
## [2808,] "Bacteria" "Firmicutes"          "Negativicutes"              
## [2809,] "Bacteria" "Fusobacteria"        "Fusobacteriia"              
## [2810,] "Bacteria" "Bacteroidetes"       "Bacteroidia"                
## [2811,] "Bacteria" "Bacteroidetes"       "Bacteroidia"                
## [2812,] "Bacteria" "Fusobacteria"        "Fusobacteriia"              
## [2813,] "Bacteria" "Proteobacteria"      "Gammaproteobacteria"        
## [2814,] "Bacteria" "Fusobacteria"        "Fusobacteriia"              
## [2815,] "Bacteria" "Bacteroidetes"       "Bacteroidia"                
## [2816,] "Bacteria" "Bacteroidetes"       "Bacteroidia"                
## [2817,] "Bacteria" "Bacteroidetes"       "Bacteroidia"                
## [2818,] "Bacteria" "Firmicutes"          "Negativicutes"              
## [2819,] "Bacteria" "Firmicutes"          "Clostridia"                 
## [2820,] "Bacteria" "Bacteroidetes"       "Bacteroidia"                
## [2821,] "Bacteria" "Firmicutes"          "Negativicutes"              
## [2822,] "Bacteria" "Bacteroidetes"       "Bacteroidia"                
## [2823,] "Bacteria" "Bacteroidetes"       "Bacteroidia"                
## [2824,] "Bacteria" "Bacteroidetes"       "Bacteroidia"                
## [2825,] "Bacteria" "Bacteroidetes"       "Bacteroidia"                
## [2826,] "Bacteria" "Bacteroidetes"       "Bacteroidia"                
## [2827,] "Bacteria" "Firmicutes"          "Negativicutes"              
## [2828,] "Bacteria" "Firmicutes"          "Negativicutes"              
## [2829,] "Bacteria" "Patescibacteria"     "Saccharimonadia"            
## [2830,] "Bacteria" "Firmicutes"          "Clostridia"                 
## [2831,] "Bacteria" "Patescibacteria"     "Saccharimonadia"            
## [2832,] "Bacteria" "Proteobacteria"      "Gammaproteobacteria"        
## [2833,] "Bacteria" "Proteobacteria"      "Gammaproteobacteria"        
## [2834,] "Bacteria" "Spirochaetes"        "Spirochaetia"               
## [2835,] "Bacteria" "Firmicutes"          "Negativicutes"              
## [2836,] "Bacteria" "Bacteroidetes"       "Bacteroidia"                
## [2837,] "Bacteria" "Firmicutes"          "Negativicutes"              
## [2838,] "Bacteria" "Fusobacteria"        "Fusobacteriia"              
## [2839,] "Bacteria" "Bacteroidetes"       "Bacteroidia"                
## [2840,] "Bacteria" "Firmicutes"          "Clostridia"                 
## [2841,] "Bacteria" "Firmicutes"          "Clostridia"                 
## [2842,] "Bacteria" "Firmicutes"          "Negativicutes"              
## [2843,] "Bacteria" "Proteobacteria"      "Gammaproteobacteria"        
## [2844,] "Bacteria" "Firmicutes"          "Negativicutes"              
## [2845,] "Bacteria" "Bacteroidetes"       "Bacteroidia"                
## [2846,] "Bacteria" "Firmicutes"          "Negativicutes"              
## [2847,] "Bacteria" "Proteobacteria"      "Gammaproteobacteria"        
## [2848,] "Bacteria" "Firmicutes"          "Bacilli"                    
## [2849,] "Bacteria" "Firmicutes"          "Negativicutes"              
## [2850,] "Bacteria" "Bacteroidetes"       "Bacteroidia"                
## [2851,] "Bacteria" "Proteobacteria"      "Gammaproteobacteria"        
## [2852,] "Bacteria" "Fusobacteria"        "Fusobacteriia"              
## [2853,] "Bacteria" "Firmicutes"          "Negativicutes"              
## [2854,] "Bacteria" "Bacteroidetes"       "Bacteroidia"                
## [2855,] "Bacteria" "Proteobacteria"      "Gammaproteobacteria"        
## [2856,] "Bacteria" "Actinobacteria"      "Actinobacteria"             
## [2857,] "Bacteria" "Proteobacteria"      "Gammaproteobacteria"        
## [2858,] "Bacteria" "Bacteroidetes"       "Bacteroidia"                
## [2859,] "Bacteria" "Bacteroidetes"       "Bacteroidia"                
## [2860,] "Bacteria" "Proteobacteria"      "Gammaproteobacteria"        
## [2861,] "Bacteria" "Firmicutes"          "Negativicutes"              
## [2862,] "Bacteria" "Firmicutes"          "Negativicutes"              
## [2863,] "Bacteria" "Bacteroidetes"       "Bacteroidia"                
## [2864,] "Bacteria" "Actinobacteria"      "Actinobacteria"             
## [2865,] "Bacteria" "Bacteroidetes"       "Bacteroidia"                
## [2866,] "Bacteria" "Tenericutes"         "Mollicutes"                 
## [2867,] "Bacteria" "Bacteroidetes"       "Bacteroidia"                
## [2868,] "Bacteria" "Firmicutes"          "Clostridia"                 
## [2869,] "Bacteria" "Actinobacteria"      "Actinobacteria"             
## [2870,] "Bacteria" "Firmicutes"          "Clostridia"                 
## [2871,] "Bacteria" "Bacteroidetes"       "Bacteroidia"                
## [2872,] "Bacteria" "Firmicutes"          NA                           
## [2873,] "Bacteria" "Proteobacteria"      "Gammaproteobacteria"        
## [2874,] "Bacteria" "Bacteroidetes"       "Bacteroidia"                
## [2875,] "Bacteria" "Bacteroidetes"       "Bacteroidia"                
## [2876,] "Bacteria" "Bacteroidetes"       "Bacteroidia"                
## [2877,] "Bacteria" "Proteobacteria"      "Gammaproteobacteria"        
## [2878,] "Bacteria" "Bacteroidetes"       "Bacteroidia"                
## [2879,] "Bacteria" "Firmicutes"          "Bacilli"                    
## [2880,] "Bacteria" "Firmicutes"          NA                           
## [2881,] "Bacteria" "Bacteroidetes"       "Bacteroidia"                
## [2882,] "Bacteria" "Proteobacteria"      "Gammaproteobacteria"        
## [2883,] "Bacteria" "Firmicutes"          "Negativicutes"              
## [2884,] "Bacteria" "Firmicutes"          "Clostridia"                 
## [2885,] "Bacteria" "Bacteroidetes"       "Bacteroidia"                
## [2886,] "Bacteria" "Firmicutes"          "Clostridia"                 
## [2887,] "Bacteria" "Firmicutes"          "Clostridia"                 
## [2888,] "Bacteria" "Fusobacteria"        "Fusobacteriia"              
## [2889,] "Bacteria" "Bacteroidetes"       "Bacteroidia"                
## [2890,] "Bacteria" "Bacteroidetes"       "Bacteroidia"                
## [2891,] "Bacteria" "Proteobacteria"      "Gammaproteobacteria"        
## [2892,] "Bacteria" "Bacteroidetes"       "Bacteroidia"                
## [2893,] "Bacteria" "Bacteroidetes"       "Bacteroidia"                
## [2894,] "Bacteria" "Proteobacteria"      "Gammaproteobacteria"        
## [2895,] "Bacteria" "Firmicutes"          NA                           
## [2896,] "Bacteria" "Fusobacteria"        "Fusobacteriia"              
## [2897,] "Bacteria" "Bacteroidetes"       "Bacteroidia"                
## [2898,] "Bacteria" "Bacteroidetes"       "Bacteroidia"                
## [2899,] "Bacteria" "Patescibacteria"     "Saccharimonadia"            
## [2900,] "Bacteria" "Fusobacteria"        "Fusobacteriia"              
## [2901,] "Bacteria" "Bacteroidetes"       "Bacteroidia"                
## [2902,] "Bacteria" "Firmicutes"          "Negativicutes"              
## [2903,] "Bacteria" "Actinobacteria"      "Actinobacteria"             
## [2904,] "Bacteria" "Bacteroidetes"       "Bacteroidia"                
## [2905,] "Bacteria" "Spirochaetes"        "Spirochaetia"               
## [2906,] "Bacteria" "Proteobacteria"      "Gammaproteobacteria"        
## [2907,] "Bacteria" "Spirochaetes"        "Spirochaetia"               
## [2908,] "Bacteria" "Epsilonbacteraeota"  "Campylobacteria"            
## [2909,] "Bacteria" "Firmicutes"          "Clostridia"                 
## [2910,] "Bacteria" "Proteobacteria"      "Gammaproteobacteria"        
## [2911,] "Bacteria" "Firmicutes"          "Negativicutes"              
## [2912,] "Bacteria" "Tenericutes"         "Mollicutes"                 
## [2913,] "Bacteria" "Bacteroidetes"       "Bacteroidia"                
## [2914,] "Bacteria" NA                    NA                           
## [2915,] "Bacteria" "Patescibacteria"     "Saccharimonadia"            
## [2916,] "Bacteria" "Bacteroidetes"       "Bacteroidia"                
## [2917,] "Bacteria" "Patescibacteria"     "Saccharimonadia"            
## [2918,] "Bacteria" "Firmicutes"          "Negativicutes"              
## [2919,] "Bacteria" "Fusobacteria"        "Fusobacteriia"              
## [2920,] "Bacteria" "Firmicutes"          "Negativicutes"              
## [2921,] "Bacteria" "Bacteroidetes"       "Bacteroidia"                
## [2922,] "Bacteria" "Actinobacteria"      "Actinobacteria"             
## [2923,] "Bacteria" "Proteobacteria"      "Gammaproteobacteria"        
## [2924,] "Bacteria" "Firmicutes"          "Bacilli"                    
## [2925,] "Bacteria" "Firmicutes"          "Clostridia"                 
## [2926,] "Bacteria" "Fusobacteria"        "Fusobacteriia"              
## [2927,] "Bacteria" "Proteobacteria"      "Gammaproteobacteria"        
## [2928,] "Bacteria" "Firmicutes"          "Bacilli"                    
## [2929,] "Bacteria" "Actinobacteria"      "Actinobacteria"             
## [2930,] "Bacteria" "Actinobacteria"      "Actinobacteria"             
## [2931,] "Bacteria" "Firmicutes"          "Clostridia"                 
## [2932,] "Bacteria" "Proteobacteria"      "Gammaproteobacteria"        
## [2933,] "Bacteria" "Firmicutes"          "Bacilli"                    
## [2934,] "Bacteria" "Actinobacteria"      "Coriobacteriia"             
## [2935,] "Bacteria" "Actinobacteria"      "Actinobacteria"             
## [2936,] "Bacteria" "Firmicutes"          "Negativicutes"              
## [2937,] "Bacteria" "Firmicutes"          NA                           
## [2938,] "Bacteria" "Fusobacteria"        "Fusobacteriia"              
## [2939,] "Bacteria" "Bacteroidetes"       "Bacteroidia"                
## [2940,] "Bacteria" "Bacteroidetes"       "Bacteroidia"                
## [2941,] "Bacteria" "Actinobacteria"      "Actinobacteria"             
## [2942,] "Bacteria" "Patescibacteria"     "Saccharimonadia"            
## [2943,] "Bacteria" "Bacteroidetes"       "Bacteroidia"                
## [2944,] "Bacteria" "Firmicutes"          "Bacilli"                    
## [2945,] "Bacteria" "Bacteroidetes"       "Bacteroidia"                
## [2946,] "Bacteria" "Bacteroidetes"       "Bacteroidia"                
## [2947,] "Bacteria" "Bacteroidetes"       "Bacteroidia"                
## [2948,] "Bacteria" "Firmicutes"          NA                           
## [2949,] "Bacteria" "Firmicutes"          "Clostridia"                 
## [2950,] "Bacteria" "Firmicutes"          "Negativicutes"              
## [2951,] "Bacteria" "Firmicutes"          "Negativicutes"              
## [2952,] "Bacteria" "Bacteroidetes"       "Bacteroidia"                
## [2953,] "Bacteria" "Bacteroidetes"       "Bacteroidia"                
## [2954,] "Bacteria" "Bacteroidetes"       "Bacteroidia"                
## [2955,] "Bacteria" "Firmicutes"          "Clostridia"                 
## [2956,] "Bacteria" "Cyanobacteria"       "Oxyphotobacteria"           
## [2957,] "Bacteria" "Proteobacteria"      "Gammaproteobacteria"        
## [2958,] "Bacteria" "Fusobacteria"        "Fusobacteriia"              
## [2959,] "Bacteria" "Firmicutes"          "Negativicutes"              
## [2960,] "Bacteria" "Firmicutes"          "Negativicutes"              
## [2961,] "Bacteria" "Bacteroidetes"       "Bacteroidia"                
## [2962,] "Bacteria" "Proteobacteria"      "Gammaproteobacteria"        
## [2963,] "Bacteria" "Firmicutes"          "Negativicutes"              
## [2964,] "Bacteria" "Firmicutes"          "Negativicutes"              
## [2965,] "Bacteria" "Proteobacteria"      "Gammaproteobacteria"        
## [2966,] "Bacteria" "Firmicutes"          "Negativicutes"              
## [2967,] "Bacteria" "Firmicutes"          "Negativicutes"              
## [2968,] "Bacteria" "Firmicutes"          NA                           
## [2969,] "Bacteria" "Bacteroidetes"       "Bacteroidia"                
## [2970,] "Bacteria" "Fusobacteria"        "Fusobacteriia"              
## [2971,] "Bacteria" "Proteobacteria"      "Gammaproteobacteria"        
## [2972,] "Bacteria" "Patescibacteria"     "Gracilibacteria"            
## [2973,] "Bacteria" "Bacteroidetes"       "Bacteroidia"                
## [2974,] "Bacteria" "Bacteroidetes"       "Bacteroidia"                
## [2975,] "Bacteria" "Firmicutes"          "Bacilli"                    
## [2976,] "Bacteria" "Spirochaetes"        "Spirochaetia"               
## [2977,] "Bacteria" "Firmicutes"          "Clostridia"                 
## [2978,] "Bacteria" "Tenericutes"         "Mollicutes"                 
## [2979,] "Bacteria" "Firmicutes"          "Negativicutes"              
## [2980,] "Bacteria" "Bacteroidetes"       "Bacteroidia"                
## [2981,] "Bacteria" "Proteobacteria"      "Gammaproteobacteria"        
## [2982,] "Bacteria" "Spirochaetes"        "Spirochaetia"               
## [2983,] "Bacteria" "Epsilonbacteraeota"  "Campylobacteria"            
## [2984,] "Bacteria" "Bacteroidetes"       "Bacteroidia"                
## [2985,] "Bacteria" "Firmicutes"          "Clostridia"                 
## [2986,] "Bacteria" "Proteobacteria"      "Gammaproteobacteria"        
## [2987,] "Bacteria" "Actinobacteria"      "Actinobacteria"             
## [2988,] "Bacteria" "Bacteroidetes"       "Bacteroidia"                
## [2989,] "Bacteria" "Bacteroidetes"       "Bacteroidia"                
## [2990,] "Bacteria" "Proteobacteria"      "Gammaproteobacteria"        
## [2991,] "Bacteria" "Firmicutes"          "Negativicutes"              
## [2992,] "Bacteria" "Synergistetes"       "Synergistia"                
## [2993,] "Bacteria" "Firmicutes"          "Bacilli"                    
## [2994,] "Bacteria" "Actinobacteria"      "Actinobacteria"             
## [2995,] "Bacteria" "Bacteroidetes"       "Bacteroidia"                
## [2996,] "Bacteria" "Firmicutes"          "Clostridia"                 
## [2997,] "Bacteria" "Firmicutes"          "Negativicutes"              
## [2998,] "Bacteria" "Patescibacteria"     "Saccharimonadia"            
## [2999,] "Bacteria" "Proteobacteria"      "Gammaproteobacteria"        
## [3000,] "Bacteria" "Firmicutes"          "Negativicutes"              
## [3001,] "Bacteria" "Bacteroidetes"       "Bacteroidia"                
## [3002,] "Bacteria" "Fusobacteria"        "Fusobacteriia"              
## [3003,] "Bacteria" "Firmicutes"          NA                           
## [3004,] "Bacteria" "Bacteroidetes"       "Bacteroidia"                
## [3005,] "Bacteria" "Proteobacteria"      "Gammaproteobacteria"        
## [3006,] "Bacteria" "Proteobacteria"      "Gammaproteobacteria"        
## [3007,] "Bacteria" "Firmicutes"          "Negativicutes"              
## [3008,] "Bacteria" "Actinobacteria"      "Actinobacteria"             
## [3009,] "Bacteria" "Firmicutes"          "Clostridia"                 
## [3010,] "Bacteria" "Firmicutes"          "Negativicutes"              
## [3011,] "Bacteria" "Fusobacteria"        "Fusobacteriia"              
## [3012,] "Bacteria" "Firmicutes"          "Negativicutes"              
## [3013,] "Bacteria" "Firmicutes"          NA                           
## [3014,] "Bacteria" "Bacteroidetes"       "Bacteroidia"                
## [3015,] "Bacteria" "Spirochaetes"        "Spirochaetia"               
## [3016,] "Bacteria" NA                    NA                           
## [3017,] "Bacteria" "Proteobacteria"      "Gammaproteobacteria"        
## [3018,] "Bacteria" "Firmicutes"          "Negativicutes"              
## [3019,] "Bacteria" "Bacteroidetes"       "Bacteroidia"                
## [3020,] "Bacteria" NA                    NA                           
## [3021,] "Bacteria" "Bacteroidetes"       "Bacteroidia"                
## [3022,] "Bacteria" "Firmicutes"          "Clostridia"                 
## [3023,] "Bacteria" "Bacteroidetes"       "Bacteroidia"                
## [3024,] "Bacteria" "Bacteroidetes"       "Bacteroidia"                
## [3025,] "Bacteria" "Bacteroidetes"       "Bacteroidia"                
## [3026,] "Bacteria" "Patescibacteria"     "Gracilibacteria"            
## [3027,] "Bacteria" "Bacteroidetes"       "Bacteroidia"                
## [3028,] "Bacteria" "Patescibacteria"     "Gracilibacteria"            
## [3029,] "Bacteria" "Fusobacteria"        "Fusobacteriia"              
## [3030,] "Bacteria" "Bacteroidetes"       "Bacteroidia"                
## [3031,] "Bacteria" "Bacteroidetes"       "Bacteroidia"                
## [3032,] "Bacteria" "Proteobacteria"      "Gammaproteobacteria"        
## [3033,] "Bacteria" "Bacteroidetes"       "Bacteroidia"                
## [3034,] "Bacteria" NA                    NA                           
## [3035,] "Bacteria" "Firmicutes"          "Bacilli"                    
## [3036,] "Bacteria" "Firmicutes"          "Clostridia"                 
## [3037,] "Bacteria" "Firmicutes"          "Clostridia"                 
## [3038,] "Bacteria" "Bacteroidetes"       "Bacteroidia"                
## [3039,] "Bacteria" "Actinobacteria"      "Actinobacteria"             
## [3040,] "Bacteria" "Firmicutes"          "Bacilli"                    
## [3041,] "Bacteria" "Epsilonbacteraeota"  "Campylobacteria"            
## [3042,] "Bacteria" "Bacteroidetes"       "Bacteroidia"                
## [3043,] "Bacteria" "Firmicutes"          "Negativicutes"              
## [3044,] "Bacteria" "Firmicutes"          "Negativicutes"              
## [3045,] "Bacteria" "Bacteroidetes"       "Bacteroidia"                
## [3046,] "Bacteria" "Proteobacteria"      "Gammaproteobacteria"        
## [3047,] "Bacteria" "Actinobacteria"      "Actinobacteria"             
## [3048,] "Bacteria" "Proteobacteria"      "Gammaproteobacteria"        
## [3049,] "Bacteria" "Patescibacteria"     "Gracilibacteria"            
## [3050,] "Bacteria" "Firmicutes"          "Clostridia"                 
## [3051,] "Bacteria" "Bacteroidetes"       "Bacteroidia"                
## [3052,] "Bacteria" "Proteobacteria"      "Gammaproteobacteria"        
## [3053,] "Bacteria" "Fusobacteria"        "Fusobacteriia"              
## [3054,] "Bacteria" "Fusobacteria"        "Fusobacteriia"              
## [3055,] "Bacteria" "Firmicutes"          "Clostridia"                 
## [3056,] "Bacteria" "Bacteroidetes"       "Bacteroidia"                
## [3057,] "Bacteria" "Bacteroidetes"       "Bacteroidia"                
## [3058,] "Bacteria" "Fusobacteria"        "Fusobacteriia"              
## [3059,] "Bacteria" "Bacteroidetes"       "Bacteroidia"                
## [3060,] "Bacteria" "Bacteroidetes"       "Bacteroidia"                
## [3061,] "Bacteria" "Bacteroidetes"       "Bacteroidia"                
## [3062,] "Bacteria" "Fusobacteria"        "Fusobacteriia"              
## [3063,] "Bacteria" "Firmicutes"          "Negativicutes"              
## [3064,] "Bacteria" "Firmicutes"          "Clostridia"                 
## [3065,] "Bacteria" "Actinobacteria"      "Actinobacteria"             
## [3066,] "Bacteria" "Firmicutes"          "Clostridia"                 
## [3067,] "Bacteria" "Firmicutes"          "Negativicutes"              
## [3068,] "Bacteria" "Firmicutes"          "Negativicutes"              
## [3069,] "Bacteria" "Patescibacteria"     "Saccharimonadia"            
## [3070,] "Bacteria" "Actinobacteria"      "Actinobacteria"             
## [3071,] "Bacteria" "Firmicutes"          "Clostridia"                 
## [3072,] "Bacteria" "Firmicutes"          "Negativicutes"              
## [3073,] "Bacteria" "Bacteroidetes"       "Bacteroidia"                
## [3074,] "Bacteria" "Fusobacteria"        "Fusobacteriia"              
## [3075,] "Bacteria" "Bacteroidetes"       "Bacteroidia"                
## [3076,] "Bacteria" "Proteobacteria"      "Gammaproteobacteria"        
## [3077,] "Bacteria" "Firmicutes"          NA                           
## [3078,] "Bacteria" "Fusobacteria"        "Fusobacteriia"              
## [3079,] "Bacteria" "Proteobacteria"      "Gammaproteobacteria"        
## [3080,] "Bacteria" "Bacteroidetes"       "Bacteroidia"                
## [3081,] "Bacteria" "Spirochaetes"        "Spirochaetia"               
## [3082,] "Bacteria" "Actinobacteria"      "Actinobacteria"             
## [3083,] "Bacteria" "Actinobacteria"      "Actinobacteria"             
## [3084,] "Bacteria" "Bacteroidetes"       "Bacteroidia"                
## [3085,] "Bacteria" "Fusobacteria"        "Fusobacteriia"              
## [3086,] "Bacteria" "Firmicutes"          NA                           
## [3087,] "Bacteria" "Bacteroidetes"       "Bacteroidia"                
## [3088,] "Bacteria" "Proteobacteria"      "Gammaproteobacteria"        
## [3089,] "Bacteria" "Patescibacteria"     "Saccharimonadia"            
## [3090,] "Bacteria" "Firmicutes"          "Negativicutes"              
## [3091,] "Bacteria" "Firmicutes"          "Negativicutes"              
## [3092,] "Bacteria" "Actinobacteria"      "Actinobacteria"             
## [3093,] "Bacteria" "Fusobacteria"        "Fusobacteriia"              
## [3094,] "Bacteria" "Actinobacteria"      "Actinobacteria"             
## [3095,] "Bacteria" "Proteobacteria"      "Gammaproteobacteria"        
## [3096,] "Bacteria" "Bacteroidetes"       "Bacteroidia"                
## [3097,] "Bacteria" "Firmicutes"          "Negativicutes"              
## [3098,] "Bacteria" "Firmicutes"          "Negativicutes"              
## [3099,] "Bacteria" "Firmicutes"          NA                           
## [3100,] "Bacteria" "Bacteroidetes"       "Bacteroidia"                
## [3101,] "Bacteria" "Bacteroidetes"       "Bacteroidia"                
## [3102,] "Bacteria" "Spirochaetes"        "Spirochaetia"               
## [3103,] "Bacteria" NA                    NA                           
## [3104,] "Bacteria" "Bacteroidetes"       "Bacteroidia"                
## [3105,] "Bacteria" "Firmicutes"          NA                           
## [3106,] "Bacteria" "Actinobacteria"      "Actinobacteria"             
## [3107,] "Bacteria" "Bacteroidetes"       "Bacteroidia"                
## [3108,] "Bacteria" "Bacteroidetes"       "Bacteroidia"                
## [3109,] "Bacteria" "Proteobacteria"      "Gammaproteobacteria"        
## [3110,] "Bacteria" "Actinobacteria"      "Actinobacteria"             
## [3111,] "Bacteria" "Firmicutes"          NA                           
## [3112,] "Bacteria" "Fusobacteria"        "Fusobacteriia"              
## [3113,] "Bacteria" "Firmicutes"          "Negativicutes"              
## [3114,] "Bacteria" "Firmicutes"          "Clostridia"                 
## [3115,] "Bacteria" "Proteobacteria"      "Gammaproteobacteria"        
## [3116,] "Bacteria" "Firmicutes"          "Clostridia"                 
## [3117,] "Bacteria" "Firmicutes"          "Bacilli"                    
## [3118,] "Bacteria" "Bacteroidetes"       "Bacteroidia"                
## [3119,] "Bacteria" "Fusobacteria"        "Fusobacteriia"              
## [3120,] "Bacteria" "Bacteroidetes"       "Bacteroidia"                
## [3121,] "Bacteria" "Bacteroidetes"       "Bacteroidia"                
## [3122,] "Bacteria" "Actinobacteria"      "Actinobacteria"             
## [3123,] "Bacteria" "Actinobacteria"      "Actinobacteria"             
## [3124,] "Bacteria" "Bacteroidetes"       "Bacteroidia"                
## [3125,] "Bacteria" "Bacteroidetes"       "Bacteroidia"                
## [3126,] "Bacteria" "Firmicutes"          "Clostridia"                 
## [3127,] "Bacteria" "Firmicutes"          "Bacilli"                    
## [3128,] "Bacteria" "Bacteroidetes"       "Bacteroidia"                
## [3129,] "Bacteria" "Actinobacteria"      "Actinobacteria"             
## [3130,] "Bacteria" "Actinobacteria"      "Actinobacteria"             
## [3131,] "Bacteria" "Firmicutes"          "Negativicutes"              
## [3132,] "Bacteria" "Bacteroidetes"       "Bacteroidia"                
## [3133,] "Bacteria" "Bacteroidetes"       "Bacteroidia"                
## [3134,] "Bacteria" "Proteobacteria"      "Gammaproteobacteria"        
## [3135,] "Bacteria" "Bacteroidetes"       "Bacteroidia"                
## [3136,] "Bacteria" "Actinobacteria"      "Actinobacteria"             
## [3137,] "Bacteria" "Epsilonbacteraeota"  "Campylobacteria"            
## [3138,] "Bacteria" "Bacteroidetes"       "Bacteroidia"                
## [3139,] "Bacteria" "Bacteroidetes"       "Bacteroidia"                
## [3140,] "Bacteria" "Bacteroidetes"       "Bacteroidia"                
## [3141,] "Bacteria" "Bacteroidetes"       "Bacteroidia"                
## [3142,] "Bacteria" "Bacteroidetes"       "Bacteroidia"                
## [3143,] "Bacteria" "Bacteroidetes"       "Bacteroidia"                
## [3144,] "Bacteria" "Proteobacteria"      "Gammaproteobacteria"        
## [3145,] "Bacteria" "Patescibacteria"     "Saccharimonadia"            
## [3146,] "Bacteria" "Fusobacteria"        "Fusobacteriia"              
## [3147,] "Bacteria" "Fusobacteria"        "Fusobacteriia"              
## [3148,] "Bacteria" "Actinobacteria"      "Actinobacteria"             
## [3149,] "Bacteria" "Firmicutes"          NA                           
## [3150,] "Bacteria" "Spirochaetes"        "Spirochaetia"               
## [3151,] "Bacteria" "Firmicutes"          "Clostridia"                 
## [3152,] "Bacteria" "Proteobacteria"      "Alphaproteobacteria"        
## [3153,] "Bacteria" "Spirochaetes"        "Spirochaetia"               
## [3154,] "Bacteria" "Bacteroidetes"       "Bacteroidia"                
## [3155,] "Bacteria" "Bacteroidetes"       "Bacteroidia"                
## [3156,] "Bacteria" "Bacteroidetes"       "Bacteroidia"                
## [3157,] "Bacteria" "Firmicutes"          "Negativicutes"              
## [3158,] "Bacteria" "Firmicutes"          "Clostridia"                 
## [3159,] "Bacteria" "Proteobacteria"      "Gammaproteobacteria"        
## [3160,] "Bacteria" "Bacteroidetes"       "Bacteroidia"                
## [3161,] "Bacteria" "Epsilonbacteraeota"  "Campylobacteria"            
## [3162,] "Bacteria" "Proteobacteria"      "Gammaproteobacteria"        
## [3163,] "Bacteria" "Bacteroidetes"       "Bacteroidia"                
## [3164,] "Bacteria" "Bacteroidetes"       "Bacteroidia"                
## [3165,] "Bacteria" "Firmicutes"          "Clostridia"                 
## [3166,] "Bacteria" "Proteobacteria"      "Gammaproteobacteria"        
## [3167,] "Bacteria" "Bacteroidetes"       "Bacteroidia"                
## [3168,] "Bacteria" "Spirochaetes"        "Spirochaetia"               
## [3169,] "Bacteria" "Spirochaetes"        "Spirochaetia"               
## [3170,] "Bacteria" "Fusobacteria"        "Fusobacteriia"              
## [3171,] "Bacteria" "Epsilonbacteraeota"  "Campylobacteria"            
## [3172,] "Bacteria" "Tenericutes"         "Mollicutes"                 
## [3173,] "Bacteria" "Actinobacteria"      "Actinobacteria"             
## [3174,] "Bacteria" "Fusobacteria"        "Fusobacteriia"              
## [3175,] "Bacteria" "Cyanobacteria"       "Oxyphotobacteria"           
## [3176,] "Bacteria" "Epsilonbacteraeota"  "Campylobacteria"            
## [3177,] "Bacteria" "Bacteroidetes"       "Bacteroidia"                
## [3178,] "Bacteria" "Actinobacteria"      "Actinobacteria"             
## [3179,] "Bacteria" "Fusobacteria"        "Fusobacteriia"              
## [3180,] "Bacteria" "Proteobacteria"      "Gammaproteobacteria"        
## [3181,] "Bacteria" "Bacteroidetes"       "Bacteroidia"                
## [3182,] "Bacteria" "Spirochaetes"        "Spirochaetia"               
## [3183,] "Bacteria" "Fusobacteria"        "Fusobacteriia"              
## [3184,] "Bacteria" "Bacteroidetes"       "Bacteroidia"                
## [3185,] "Bacteria" "Fusobacteria"        "Fusobacteriia"              
## [3186,] "Bacteria" "Bacteroidetes"       "Bacteroidia"                
## [3187,] "Bacteria" "Proteobacteria"      "Gammaproteobacteria"        
## [3188,] "Bacteria" "Patescibacteria"     "Saccharimonadia"            
## [3189,] "Bacteria" "Bacteroidetes"       "Bacteroidia"                
## [3190,] "Bacteria" "Bacteroidetes"       "Bacteroidia"                
## [3191,] "Bacteria" "Spirochaetes"        "Spirochaetia"               
## [3192,] "Bacteria" "Proteobacteria"      "Gammaproteobacteria"        
## [3193,] "Bacteria" "Bacteroidetes"       "Bacteroidia"                
## [3194,] "Bacteria" "Fusobacteria"        "Fusobacteriia"              
## [3195,] "Bacteria" "Proteobacteria"      "Gammaproteobacteria"        
## [3196,] "Bacteria" "Fusobacteria"        "Fusobacteriia"              
## [3197,] "Bacteria" "Bacteroidetes"       "Bacteroidia"                
## [3198,] "Bacteria" "Bacteroidetes"       "Bacteroidia"                
## [3199,] "Bacteria" "Bacteroidetes"       "Bacteroidia"                
## [3200,] "Bacteria" "Epsilonbacteraeota"  "Campylobacteria"            
## [3201,] "Bacteria" "Proteobacteria"      "Gammaproteobacteria"        
## [3202,] "Bacteria" "Firmicutes"          "Clostridia"                 
## [3203,] "Bacteria" "Epsilonbacteraeota"  "Campylobacteria"            
## [3204,] "Bacteria" "Patescibacteria"     "Saccharimonadia"            
## [3205,] "Bacteria" "Actinobacteria"      "Actinobacteria"             
## [3206,] "Bacteria" "Bacteroidetes"       "Bacteroidia"                
## [3207,] "Bacteria" "Bacteroidetes"       "Bacteroidia"                
## [3208,] "Bacteria" "Bacteroidetes"       "Bacteroidia"                
## [3209,] "Bacteria" NA                    NA                           
## [3210,] "Bacteria" "Bacteroidetes"       "Bacteroidia"                
## [3211,] "Bacteria" "Actinobacteria"      "Actinobacteria"             
## [3212,] "Bacteria" "Bacteroidetes"       "Bacteroidia"                
## [3213,] "Bacteria" "Firmicutes"          "Clostridia"                 
## [3214,] "Bacteria" "Firmicutes"          "Clostridia"                 
## [3215,] "Bacteria" "Bacteroidetes"       "Bacteroidia"                
## [3216,] "Bacteria" "Bacteroidetes"       "Bacteroidia"                
## [3217,] "Bacteria" "Fusobacteria"        "Fusobacteriia"              
## [3218,] "Bacteria" "Bacteroidetes"       "Bacteroidia"                
## [3219,] "Bacteria" "Bacteroidetes"       "Bacteroidia"                
## [3220,] "Bacteria" "Firmicutes"          "Negativicutes"              
## [3221,] "Bacteria" "Firmicutes"          "Clostridia"                 
## [3222,] "Bacteria" "Firmicutes"          "Clostridia"                 
## [3223,] "Bacteria" "Firmicutes"          "Bacilli"                    
## [3224,] "Bacteria" "Firmicutes"          "Clostridia"                 
## [3225,] "Bacteria" "Firmicutes"          "Clostridia"                 
## [3226,] "Bacteria" "Actinobacteria"      "Actinobacteria"             
## [3227,] "Bacteria" "Fusobacteria"        "Fusobacteriia"              
## [3228,] "Bacteria" "Fusobacteria"        "Fusobacteriia"              
## [3229,] "Bacteria" "Firmicutes"          "Negativicutes"              
## [3230,] "Bacteria" "Fusobacteria"        "Fusobacteriia"              
## [3231,] "Bacteria" "Firmicutes"          NA                           
## [3232,] "Bacteria" "Bacteroidetes"       "Bacteroidia"                
## [3233,] "Bacteria" "Fusobacteria"        "Fusobacteriia"              
## [3234,] "Bacteria" "Proteobacteria"      "Gammaproteobacteria"        
## [3235,] "Bacteria" "Bacteroidetes"       "Bacteroidia"                
## [3236,] "Bacteria" NA                    NA                           
## [3237,] "Bacteria" "Cyanobacteria"       "Oxyphotobacteria"           
## [3238,] "Bacteria" "Firmicutes"          "Negativicutes"              
## [3239,] "Bacteria" "Firmicutes"          "Bacilli"                    
## [3240,] "Bacteria" "Bacteroidetes"       "Bacteroidia"                
## [3241,] "Bacteria" "Bacteroidetes"       "Bacteroidia"                
## [3242,] "Bacteria" "Firmicutes"          "Negativicutes"              
## [3243,] "Bacteria" "Bacteroidetes"       "Bacteroidia"                
## [3244,] "Bacteria" "Firmicutes"          "Clostridia"                 
## [3245,] "Bacteria" "Actinobacteria"      "Actinobacteria"             
## [3246,] "Bacteria" "Firmicutes"          "Clostridia"                 
## [3247,] "Bacteria" "Epsilonbacteraeota"  "Campylobacteria"            
## [3248,] "Bacteria" "Actinobacteria"      "Coriobacteriia"             
## [3249,] "Bacteria" "Bacteroidetes"       "Bacteroidia"                
## [3250,] "Bacteria" "Patescibacteria"     "Saccharimonadia"            
## [3251,] "Bacteria" "Actinobacteria"      "Coriobacteriia"             
## [3252,] "Bacteria" "Actinobacteria"      "Actinobacteria"             
## [3253,] "Bacteria" "Bacteroidetes"       "Bacteroidia"                
## [3254,] "Bacteria" "Bacteroidetes"       "Bacteroidia"                
## [3255,] "Bacteria" "Proteobacteria"      "Gammaproteobacteria"        
## [3256,] "Bacteria" "Bacteroidetes"       "Bacteroidia"                
## [3257,] "Bacteria" "Patescibacteria"     "Saccharimonadia"            
## [3258,] "Bacteria" "Fusobacteria"        "Fusobacteriia"              
## [3259,] "Bacteria" "Firmicutes"          "Bacilli"                    
## [3260,] "Bacteria" "Epsilonbacteraeota"  "Campylobacteria"            
## [3261,] "Bacteria" "Epsilonbacteraeota"  "Campylobacteria"            
## [3262,] "Bacteria" "Actinobacteria"      "Actinobacteria"             
## [3263,] "Bacteria" "Patescibacteria"     "Saccharimonadia"            
## [3264,] "Bacteria" "Fusobacteria"        "Fusobacteriia"              
## [3265,] "Bacteria" "Actinobacteria"      "Actinobacteria"             
## [3266,] "Bacteria" "Bacteroidetes"       "Bacteroidia"                
## [3267,] "Bacteria" "Bacteroidetes"       "Bacteroidia"                
## [3268,] "Bacteria" "Firmicutes"          "Bacilli"                    
## [3269,] "Bacteria" "Bacteroidetes"       "Bacteroidia"                
## [3270,] "Bacteria" "Fusobacteria"        "Fusobacteriia"              
## [3271,] "Bacteria" "Bacteroidetes"       "Bacteroidia"                
## [3272,] "Bacteria" "Bacteroidetes"       "Bacteroidia"                
## [3273,] "Bacteria" "Bacteroidetes"       "Bacteroidia"                
## [3274,] "Bacteria" "Patescibacteria"     "Saccharimonadia"            
## [3275,] "Bacteria" "Bacteroidetes"       "Bacteroidia"                
## [3276,] "Bacteria" "Actinobacteria"      "Actinobacteria"             
## [3277,] "Bacteria" NA                    NA                           
## [3278,] "Bacteria" "Firmicutes"          "Bacilli"                    
## [3279,] "Bacteria" "Spirochaetes"        "Spirochaetia"               
## [3280,] "Bacteria" "Bacteroidetes"       "Bacteroidia"                
## [3281,] "Bacteria" "Firmicutes"          "Negativicutes"              
## [3282,] "Bacteria" "Proteobacteria"      "Gammaproteobacteria"        
## [3283,] "Bacteria" "Bacteroidetes"       "Bacteroidia"                
## [3284,] "Bacteria" "Bacteroidetes"       "Bacteroidia"                
## [3285,] "Bacteria" "Proteobacteria"      "Gammaproteobacteria"        
## [3286,] "Bacteria" "Proteobacteria"      "Gammaproteobacteria"        
## [3287,] "Bacteria" "Proteobacteria"      "Gammaproteobacteria"        
## [3288,] "Bacteria" "Spirochaetes"        "Spirochaetia"               
## [3289,] "Bacteria" "Firmicutes"          "Clostridia"                 
## [3290,] "Bacteria" "Spirochaetes"        "Spirochaetia"               
## [3291,] "Bacteria" "Bacteroidetes"       "Bacteroidia"                
## [3292,] "Bacteria" "Proteobacteria"      "Gammaproteobacteria"        
## [3293,] "Bacteria" "Firmicutes"          "Negativicutes"              
## [3294,] "Bacteria" "Bacteroidetes"       "Bacteroidia"                
## [3295,] "Bacteria" "Bacteroidetes"       "Bacteroidia"                
## [3296,] "Bacteria" "Bacteroidetes"       "Bacteroidia"                
## [3297,] "Bacteria" "Actinobacteria"      "Actinobacteria"             
## [3298,] "Bacteria" "Bacteroidetes"       "Bacteroidia"                
## [3299,] "Bacteria" "Firmicutes"          "Bacilli"                    
## [3300,] "Bacteria" "Bacteroidetes"       "Bacteroidia"                
## [3301,] "Bacteria" "Bacteroidetes"       "Bacteroidia"                
## [3302,] "Bacteria" "Bacteroidetes"       "Bacteroidia"                
## [3303,] "Bacteria" "Bacteroidetes"       "Bacteroidia"                
## [3304,] "Bacteria" "Proteobacteria"      "Gammaproteobacteria"        
## [3305,] "Bacteria" "Proteobacteria"      "Gammaproteobacteria"        
## [3306,] "Bacteria" "Bacteroidetes"       "Bacteroidia"                
## [3307,] "Bacteria" "Bacteroidetes"       "Bacteroidia"                
## [3308,] "Bacteria" "Patescibacteria"     "Saccharimonadia"            
## [3309,] "Bacteria" "Proteobacteria"      "Gammaproteobacteria"        
## [3310,] "Bacteria" "Bacteroidetes"       "Bacteroidia"                
## [3311,] "Bacteria" "Bacteroidetes"       "Bacteroidia"                
## [3312,] "Bacteria" "Fusobacteria"        "Fusobacteriia"              
## [3313,] "Bacteria" "Bacteroidetes"       "Bacteroidia"                
## [3314,] "Bacteria" "Spirochaetes"        "Spirochaetia"               
## [3315,] "Bacteria" "Bacteroidetes"       "Bacteroidia"                
## [3316,] "Bacteria" "Patescibacteria"     "Saccharimonadia"            
## [3317,] "Bacteria" "Firmicutes"          "Clostridia"                 
## [3318,] "Bacteria" "Fusobacteria"        "Fusobacteriia"              
## [3319,] "Bacteria" "Proteobacteria"      "Gammaproteobacteria"        
## [3320,] "Bacteria" "Bacteroidetes"       "Bacteroidia"                
## [3321,] "Bacteria" "Bacteroidetes"       "Bacteroidia"                
## [3322,] "Bacteria" "Fusobacteria"        "Fusobacteriia"              
## [3323,] "Bacteria" "Bacteroidetes"       "Bacteroidia"                
## [3324,] "Bacteria" "Bacteroidetes"       "Bacteroidia"                
## [3325,] "Bacteria" "Bacteroidetes"       "Bacteroidia"                
## [3326,] "Bacteria" NA                    NA                           
## [3327,] "Bacteria" "Actinobacteria"      "Actinobacteria"             
## [3328,] "Bacteria" "Bacteroidetes"       "Bacteroidia"                
## [3329,] "Bacteria" "Bacteroidetes"       "Bacteroidia"                
## [3330,] "Bacteria" "Spirochaetes"        "Spirochaetia"               
## [3331,] "Bacteria" "Bacteroidetes"       "Bacteroidia"                
## [3332,] "Bacteria" "Bacteroidetes"       "Bacteroidia"                
## [3333,] "Bacteria" "Firmicutes"          "Negativicutes"              
## [3334,] "Bacteria" "Actinobacteria"      "Actinobacteria"             
## [3335,] "Bacteria" "Firmicutes"          "Bacilli"                    
## [3336,] "Bacteria" "Proteobacteria"      "Gammaproteobacteria"        
## [3337,] "Bacteria" "Bacteroidetes"       "Bacteroidia"                
## [3338,] "Bacteria" "Fusobacteria"        "Fusobacteriia"              
## [3339,] "Bacteria" "Patescibacteria"     "Saccharimonadia"            
## [3340,] "Bacteria" "Patescibacteria"     "Saccharimonadia"            
## [3341,] "Bacteria" "Firmicutes"          "Clostridia"                 
## [3342,] "Bacteria" "Fusobacteria"        "Fusobacteriia"              
## [3343,] "Bacteria" "Firmicutes"          NA                           
## [3344,] "Bacteria" "Firmicutes"          "Clostridia"                 
## [3345,] "Bacteria" "Patescibacteria"     "Saccharimonadia"            
## [3346,] "Bacteria" "Epsilonbacteraeota"  "Campylobacteria"            
## [3347,] "Bacteria" "Bacteroidetes"       "Bacteroidia"                
## [3348,] "Bacteria" "Proteobacteria"      "Gammaproteobacteria"        
## [3349,] "Bacteria" "Bacteroidetes"       "Bacteroidia"                
## [3350,] "Bacteria" "Fusobacteria"        "Fusobacteriia"              
## [3351,] "Bacteria" "Proteobacteria"      "Gammaproteobacteria"        
## [3352,] "Bacteria" "Proteobacteria"      "Gammaproteobacteria"        
## [3353,] "Bacteria" "Bacteroidetes"       "Bacteroidia"                
## [3354,] "Bacteria" "Proteobacteria"      "Gammaproteobacteria"        
## [3355,] "Bacteria" "Proteobacteria"      "Gammaproteobacteria"        
## [3356,] "Bacteria" "Firmicutes"          "Negativicutes"              
## [3357,] "Bacteria" "Actinobacteria"      "Coriobacteriia"             
## [3358,] "Bacteria" "Fusobacteria"        "Fusobacteriia"              
## [3359,] "Bacteria" "Spirochaetes"        "Spirochaetia"               
## [3360,] "Bacteria" "Firmicutes"          "Negativicutes"              
## [3361,] "Bacteria" "Actinobacteria"      "Actinobacteria"             
## [3362,] "Bacteria" "Bacteroidetes"       "Bacteroidia"                
## [3363,] "Bacteria" "Firmicutes"          "Negativicutes"              
## [3364,] "Bacteria" "Firmicutes"          "Clostridia"                 
## [3365,] "Bacteria" "Bacteroidetes"       "Bacteroidia"                
## [3366,] "Bacteria" "Bacteroidetes"       "Bacteroidia"                
## [3367,] "Bacteria" "Proteobacteria"      "Gammaproteobacteria"        
## [3368,] "Bacteria" "Actinobacteria"      "Actinobacteria"             
## [3369,] "Bacteria" "Bacteroidetes"       "Bacteroidia"                
## [3370,] "Bacteria" "Firmicutes"          "Negativicutes"              
## [3371,] "Bacteria" "Bacteroidetes"       "Bacteroidia"                
## [3372,] "Bacteria" "Fusobacteria"        "Fusobacteriia"              
## [3373,] "Bacteria" "Fusobacteria"        "Fusobacteriia"              
## [3374,] "Bacteria" "Spirochaetes"        "Spirochaetia"               
## [3375,] "Bacteria" "Bacteroidetes"       "Bacteroidia"                
## [3376,] "Bacteria" "Bacteroidetes"       "Bacteroidia"                
## [3377,] "Bacteria" "Bacteroidetes"       "Bacteroidia"                
## [3378,] "Bacteria" "Fusobacteria"        "Fusobacteriia"              
## [3379,] "Bacteria" "Bacteroidetes"       "Bacteroidia"                
## [3380,] "Bacteria" "Bacteroidetes"       "Bacteroidia"                
## [3381,] "Bacteria" "Bacteroidetes"       "Bacteroidia"                
## [3382,] "Bacteria" "Bacteroidetes"       "Bacteroidia"                
## [3383,] "Bacteria" "Firmicutes"          "Clostridia"                 
## [3384,] "Bacteria" "Bacteroidetes"       "Bacteroidia"                
## [3385,] "Bacteria" "Spirochaetes"        "Spirochaetia"               
## [3386,] "Bacteria" "Bacteroidetes"       "Bacteroidia"                
## [3387,] "Bacteria" "Firmicutes"          "Bacilli"                    
## [3388,] "Bacteria" "Firmicutes"          "Clostridia"                 
## [3389,] "Bacteria" "Bacteroidetes"       "Bacteroidia"                
## [3390,] "Bacteria" "Bacteroidetes"       "Bacteroidia"                
## [3391,] "Bacteria" "Firmicutes"          "Bacilli"                    
## [3392,] "Bacteria" "Bacteroidetes"       "Bacteroidia"                
## [3393,] "Bacteria" "Patescibacteria"     "Saccharimonadia"            
## [3394,] "Bacteria" NA                    NA                           
## [3395,] "Bacteria" "Fusobacteria"        "Fusobacteriia"              
## [3396,] "Bacteria" "Actinobacteria"      "Actinobacteria"             
## [3397,] "Bacteria" "Bacteroidetes"       "Bacteroidia"                
## [3398,] "Bacteria" "Actinobacteria"      "Actinobacteria"             
## [3399,] "Bacteria" "Firmicutes"          "Clostridia"                 
## [3400,] "Bacteria" "Bacteroidetes"       "Bacteroidia"                
## [3401,] "Bacteria" "Proteobacteria"      "Gammaproteobacteria"        
## [3402,] "Bacteria" "Proteobacteria"      "Gammaproteobacteria"        
## [3403,] "Bacteria" "Firmicutes"          "Clostridia"                 
## [3404,] "Bacteria" "Bacteroidetes"       "Bacteroidia"                
## [3405,] "Bacteria" "Bacteroidetes"       "Bacteroidia"                
## [3406,] "Bacteria" "Bacteroidetes"       "Bacteroidia"                
## [3407,] "Bacteria" "Bacteroidetes"       "Bacteroidia"                
## [3408,] "Bacteria" "Firmicutes"          "Bacilli"                    
## [3409,] "Bacteria" "Bacteroidetes"       "Bacteroidia"                
## [3410,] "Bacteria" "Proteobacteria"      "Gammaproteobacteria"        
## [3411,] "Bacteria" "Actinobacteria"      "Coriobacteriia"             
## [3412,] "Bacteria" "Bacteroidetes"       "Bacteroidia"                
## [3413,] "Bacteria" "Spirochaetes"        "Spirochaetia"               
## [3414,] "Bacteria" "Bacteroidetes"       "Bacteroidia"                
## [3415,] "Bacteria" "Bacteroidetes"       "Bacteroidia"                
## [3416,] "Bacteria" "Bacteroidetes"       "Bacteroidia"                
## [3417,] "Bacteria" "Fusobacteria"        "Fusobacteriia"              
## [3418,] "Bacteria" "Bacteroidetes"       "Bacteroidia"                
## [3419,] "Bacteria" "Firmicutes"          "Negativicutes"              
## [3420,] "Bacteria" "Firmicutes"          NA                           
## [3421,] "Bacteria" "Bacteroidetes"       "Bacteroidia"                
## [3422,] "Bacteria" "Firmicutes"          "Clostridia"                 
## [3423,] "Bacteria" "Proteobacteria"      "Gammaproteobacteria"        
## [3424,] "Bacteria" "Bacteroidetes"       "Bacteroidia"                
## [3425,] "Bacteria" "Proteobacteria"      "Gammaproteobacteria"        
## [3426,] "Bacteria" "Bacteroidetes"       "Bacteroidia"                
## [3427,] "Bacteria" "Bacteroidetes"       "Bacteroidia"                
## [3428,] "Bacteria" "Bacteroidetes"       "Bacteroidia"                
## [3429,] "Bacteria" "Proteobacteria"      "Gammaproteobacteria"        
## [3430,] "Bacteria" "Firmicutes"          "Bacilli"                    
## [3431,] "Bacteria" "Bacteroidetes"       "Bacteroidia"                
## [3432,] "Bacteria" "Bacteroidetes"       "Bacteroidia"                
## [3433,] "Bacteria" "Bacteroidetes"       "Bacteroidia"                
## [3434,] "Bacteria" "Bacteroidetes"       "Bacteroidia"                
## [3435,] "Bacteria" "Firmicutes"          NA                           
## [3436,] "Bacteria" "Proteobacteria"      "Gammaproteobacteria"        
## [3437,] "Bacteria" "Proteobacteria"      "Gammaproteobacteria"        
## [3438,] "Bacteria" "Firmicutes"          "Negativicutes"              
## [3439,] "Bacteria" "Proteobacteria"      "Gammaproteobacteria"        
## [3440,] "Bacteria" "Firmicutes"          "Bacilli"                    
## [3441,] "Bacteria" "Bacteroidetes"       "Bacteroidia"                
## [3442,] "Bacteria" "Actinobacteria"      "Coriobacteriia"             
## [3443,] "Bacteria" "Firmicutes"          "Clostridia"                 
## [3444,] "Bacteria" "Firmicutes"          "Negativicutes"              
## [3445,] "Bacteria" "Bacteroidetes"       "Bacteroidia"                
## [3446,] "Bacteria" "Firmicutes"          "Clostridia"                 
## [3447,] "Bacteria" "Epsilonbacteraeota"  "Campylobacteria"            
## [3448,] "Bacteria" "Bacteroidetes"       "Bacteroidia"                
## [3449,] "Bacteria" "Bacteroidetes"       "Bacteroidia"                
## [3450,] "Bacteria" "Fusobacteria"        "Fusobacteriia"              
## [3451,] "Bacteria" "Bacteroidetes"       "Bacteroidia"                
## [3452,] "Bacteria" "Fusobacteria"        "Fusobacteriia"              
## [3453,] "Bacteria" "Bacteroidetes"       "Bacteroidia"                
## [3454,] "Bacteria" "Bacteroidetes"       "Bacteroidia"                
## [3455,] "Bacteria" "Firmicutes"          "Negativicutes"              
## [3456,] "Bacteria" "Bacteroidetes"       "Bacteroidia"                
## [3457,] "Bacteria" "Fusobacteria"        "Fusobacteriia"              
## [3458,] "Bacteria" "Firmicutes"          "Negativicutes"              
## [3459,] "Bacteria" "Proteobacteria"      "Gammaproteobacteria"        
## [3460,] "Bacteria" "Actinobacteria"      "Actinobacteria"             
## [3461,] "Bacteria" "Firmicutes"          "Negativicutes"              
## [3462,] "Bacteria" "Proteobacteria"      "Gammaproteobacteria"        
## [3463,] "Bacteria" "Firmicutes"          "Clostridia"                 
## [3464,] "Bacteria" "Bacteroidetes"       "Bacteroidia"                
## [3465,] "Bacteria" "Bacteroidetes"       "Bacteroidia"                
## [3466,] "Bacteria" "Bacteroidetes"       "Bacteroidia"                
## [3467,] "Bacteria" "Firmicutes"          "Negativicutes"              
## [3468,] "Bacteria" "Firmicutes"          "Bacilli"                    
## [3469,] "Bacteria" "Bacteroidetes"       "Bacteroidia"                
## [3470,] "Bacteria" "Bacteroidetes"       "Bacteroidia"                
## [3471,] "Bacteria" NA                    NA                           
## [3472,] "Bacteria" "Proteobacteria"      "Gammaproteobacteria"        
## [3473,] "Bacteria" "Firmicutes"          "Clostridia"                 
## [3474,] "Bacteria" "Actinobacteria"      "Actinobacteria"             
## [3475,] "Bacteria" "Bacteroidetes"       "Bacteroidia"                
## [3476,] "Bacteria" "Bacteroidetes"       "Bacteroidia"                
## [3477,] "Bacteria" "Bacteroidetes"       "Bacteroidia"                
## [3478,] "Bacteria" "Proteobacteria"      "Gammaproteobacteria"        
## [3479,] "Bacteria" "Bacteroidetes"       "Bacteroidia"                
## [3480,] "Bacteria" "Firmicutes"          "Negativicutes"              
## [3481,] "Bacteria" "Proteobacteria"      "Gammaproteobacteria"        
## [3482,] "Bacteria" "Bacteroidetes"       "Bacteroidia"                
## [3483,] "Bacteria" "Firmicutes"          "Bacilli"                    
## [3484,] "Bacteria" "Firmicutes"          "Clostridia"                 
## [3485,] "Bacteria" "Bacteroidetes"       "Bacteroidia"                
## [3486,] "Bacteria" "Bacteroidetes"       "Bacteroidia"                
## [3487,] "Bacteria" "Bacteroidetes"       "Bacteroidia"                
## [3488,] "Bacteria" "Firmicutes"          "Negativicutes"              
## [3489,] "Bacteria" "Firmicutes"          "Negativicutes"              
## [3490,] "Bacteria" "Patescibacteria"     "Saccharimonadia"            
## [3491,] "Bacteria" "Bacteroidetes"       "Bacteroidia"                
## [3492,] "Bacteria" "Firmicutes"          "Negativicutes"              
## [3493,] "Bacteria" "Firmicutes"          "Negativicutes"              
## [3494,] "Bacteria" "Actinobacteria"      "Actinobacteria"             
## [3495,] "Bacteria" "Firmicutes"          NA                           
## [3496,] "Bacteria" "Proteobacteria"      "Gammaproteobacteria"        
## [3497,] "Bacteria" "Bacteroidetes"       "Bacteroidia"                
## [3498,] "Bacteria" "Cyanobacteria"       "Oxyphotobacteria"           
## [3499,] "Bacteria" "Firmicutes"          "Bacilli"                    
## [3500,] "Bacteria" "Bacteroidetes"       "Bacteroidia"                
## [3501,] "Bacteria" "Bacteroidetes"       "Bacteroidia"                
## [3502,] "Bacteria" "Firmicutes"          "Bacilli"                    
## [3503,] "Bacteria" "Proteobacteria"      "Gammaproteobacteria"        
## [3504,] "Bacteria" "Actinobacteria"      "Actinobacteria"             
## [3505,] "Bacteria" "Bacteroidetes"       "Bacteroidia"                
## [3506,] "Bacteria" "Bacteroidetes"       "Bacteroidia"                
## [3507,] "Bacteria" "Fusobacteria"        "Fusobacteriia"              
## [3508,] "Bacteria" "Fusobacteria"        "Fusobacteriia"              
## [3509,] "Bacteria" "Fusobacteria"        "Fusobacteriia"              
## [3510,] "Bacteria" "Patescibacteria"     "Saccharimonadia"            
## [3511,] "Bacteria" "Bacteroidetes"       "Bacteroidia"                
## [3512,] "Bacteria" "Proteobacteria"      "Gammaproteobacteria"        
## [3513,] "Bacteria" "Firmicutes"          "Clostridia"                 
## [3514,] "Bacteria" "Bacteroidetes"       "Bacteroidia"                
## [3515,] "Bacteria" "Firmicutes"          "Clostridia"                 
## [3516,] "Bacteria" "Firmicutes"          "Negativicutes"              
## [3517,] "Bacteria" "Firmicutes"          "Negativicutes"              
## [3518,] "Bacteria" "Proteobacteria"      "Alphaproteobacteria"        
## [3519,] "Bacteria" "Firmicutes"          "Negativicutes"              
## [3520,] "Bacteria" "Proteobacteria"      "Gammaproteobacteria"        
## [3521,] "Bacteria" "Proteobacteria"      "Gammaproteobacteria"        
## [3522,] "Bacteria" "Bacteroidetes"       "Bacteroidia"                
## [3523,] "Bacteria" "Firmicutes"          "Negativicutes"              
## [3524,] "Bacteria" "Proteobacteria"      "Gammaproteobacteria"        
## [3525,] "Bacteria" "Bacteroidetes"       "Bacteroidia"                
## [3526,] "Bacteria" "Firmicutes"          "Bacilli"                    
## [3527,] "Bacteria" "Proteobacteria"      "Gammaproteobacteria"        
## [3528,] "Bacteria" "Proteobacteria"      "Gammaproteobacteria"        
## [3529,] "Bacteria" "Bacteroidetes"       "Bacteroidia"                
## [3530,] "Bacteria" "Firmicutes"          "Bacilli"                    
## [3531,] "Bacteria" "Bacteroidetes"       "Bacteroidia"                
## [3532,] "Bacteria" "Bacteroidetes"       "Bacteroidia"                
## [3533,] "Bacteria" NA                    NA                           
## [3534,] "Bacteria" "Fusobacteria"        "Fusobacteriia"              
## [3535,] "Bacteria" "Firmicutes"          NA                           
## [3536,] "Bacteria" "Patescibacteria"     "Gracilibacteria"            
## [3537,] "Bacteria" "Patescibacteria"     "Saccharimonadia"            
## [3538,] "Bacteria" "Firmicutes"          "Negativicutes"              
## [3539,] "Bacteria" "Fusobacteria"        "Fusobacteriia"              
## [3540,] "Bacteria" "Bacteroidetes"       "Bacteroidia"                
## [3541,] "Bacteria" "Bacteroidetes"       "Bacteroidia"                
## [3542,] "Bacteria" "Fusobacteria"        "Fusobacteriia"              
## [3543,] "Bacteria" "Fusobacteria"        "Fusobacteriia"              
## [3544,] "Bacteria" "Proteobacteria"      "Gammaproteobacteria"        
## [3545,] "Bacteria" "Fusobacteria"        "Fusobacteriia"              
## [3546,] "Bacteria" "Firmicutes"          "Clostridia"                 
## [3547,] "Bacteria" "Proteobacteria"      "Gammaproteobacteria"        
## [3548,] "Bacteria" "Bacteroidetes"       "Bacteroidia"                
## [3549,] "Bacteria" "Actinobacteria"      "Actinobacteria"             
## [3550,] "Bacteria" NA                    NA                           
## [3551,] "Bacteria" "Firmicutes"          "Clostridia"                 
## [3552,] "Bacteria" "Firmicutes"          "Clostridia"                 
## [3553,] "Bacteria" "Proteobacteria"      "Gammaproteobacteria"        
## [3554,] "Bacteria" "Proteobacteria"      "Gammaproteobacteria"        
## [3555,] "Bacteria" "Spirochaetes"        "Spirochaetia"               
## [3556,] "Bacteria" "Firmicutes"          "Bacilli"                    
## [3557,] "Bacteria" "Firmicutes"          "Bacilli"                    
## [3558,] "Bacteria" "Actinobacteria"      "Actinobacteria"             
## [3559,] "Bacteria" "Actinobacteria"      "Actinobacteria"             
## [3560,] "Bacteria" "Firmicutes"          "Bacilli"                    
## [3561,] "Bacteria" "Spirochaetes"        "Spirochaetia"               
## [3562,] "Bacteria" "Firmicutes"          "Clostridia"                 
## [3563,] "Bacteria" "Epsilonbacteraeota"  "Campylobacteria"            
## [3564,] "Bacteria" "Firmicutes"          "Clostridia"                 
## [3565,] "Bacteria" "Bacteroidetes"       "Bacteroidia"                
## [3566,] "Bacteria" "Firmicutes"          "Negativicutes"              
## [3567,] "Bacteria" "Proteobacteria"      "Gammaproteobacteria"        
## [3568,] "Bacteria" "Firmicutes"          "Bacilli"                    
## [3569,] "Bacteria" "Firmicutes"          "Negativicutes"              
## [3570,] "Bacteria" "Bacteroidetes"       "Bacteroidia"                
## [3571,] "Bacteria" "Bacteroidetes"       "Bacteroidia"                
## [3572,] "Bacteria" "Epsilonbacteraeota"  "Campylobacteria"            
## [3573,] "Bacteria" "Fusobacteria"        "Fusobacteriia"              
## [3574,] "Bacteria" "Bacteroidetes"       "Bacteroidia"                
## [3575,] "Bacteria" "Firmicutes"          "Negativicutes"              
## [3576,] "Bacteria" "Firmicutes"          "Clostridia"                 
## [3577,] "Bacteria" "Firmicutes"          "Bacilli"                    
## [3578,] "Bacteria" "Firmicutes"          "Clostridia"                 
## [3579,] "Bacteria" "Bacteroidetes"       "Bacteroidia"                
## [3580,] "Bacteria" "Firmicutes"          "Clostridia"                 
## [3581,] "Bacteria" "Bacteroidetes"       "Bacteroidia"                
## [3582,] "Bacteria" "Spirochaetes"        "Spirochaetia"               
## [3583,] "Bacteria" "Firmicutes"          "Clostridia"                 
## [3584,] "Bacteria" "Actinobacteria"      "Actinobacteria"             
## [3585,] "Bacteria" "Bacteroidetes"       "Bacteroidia"                
## [3586,] "Bacteria" "Proteobacteria"      "Gammaproteobacteria"        
## [3587,] "Bacteria" "Bacteroidetes"       "Bacteroidia"                
## [3588,] "Bacteria" "Bacteroidetes"       "Bacteroidia"                
## [3589,] "Bacteria" "Proteobacteria"      NA                           
## [3590,] "Bacteria" "Epsilonbacteraeota"  "Campylobacteria"            
## [3591,] "Bacteria" "Firmicutes"          "Bacilli"                    
## [3592,] "Bacteria" "Bacteroidetes"       "Bacteroidia"                
## [3593,] "Bacteria" "Firmicutes"          "Bacilli"                    
## [3594,] "Bacteria" "Bacteroidetes"       "Bacteroidia"                
## [3595,] "Bacteria" NA                    NA                           
## [3596,] "Bacteria" "Fusobacteria"        "Fusobacteriia"              
## [3597,] "Bacteria" "Bacteroidetes"       "Bacteroidia"                
## [3598,] "Bacteria" "Bacteroidetes"       "Bacteroidia"                
## [3599,] "Bacteria" NA                    NA                           
## [3600,] "Bacteria" "Epsilonbacteraeota"  "Campylobacteria"            
## [3601,] "Bacteria" "Bacteroidetes"       "Bacteroidia"                
## [3602,] "Bacteria" "Proteobacteria"      "Gammaproteobacteria"        
## [3603,] "Bacteria" "Firmicutes"          "Negativicutes"              
## [3604,] "Bacteria" "Proteobacteria"      "Gammaproteobacteria"        
## [3605,] "Bacteria" "Actinobacteria"      "Actinobacteria"             
## [3606,] "Bacteria" "Bacteroidetes"       "Bacteroidia"                
## [3607,] "Bacteria" "Firmicutes"          "Bacilli"                    
## [3608,] "Bacteria" "Proteobacteria"      "Gammaproteobacteria"        
## [3609,] "Bacteria" "Fusobacteria"        "Fusobacteriia"              
## [3610,] "Bacteria" "Actinobacteria"      "Actinobacteria"             
## [3611,] "Bacteria" "Bacteroidetes"       "Bacteroidia"                
## [3612,] "Bacteria" "Firmicutes"          "Clostridia"                 
## [3613,] "Bacteria" "Fusobacteria"        "Fusobacteriia"              
## [3614,] "Bacteria" "Firmicutes"          "Negativicutes"              
## [3615,] "Bacteria" "Fusobacteria"        "Fusobacteriia"              
## [3616,] "Bacteria" "Firmicutes"          "Clostridia"                 
## [3617,] "Bacteria" "Fusobacteria"        "Fusobacteriia"              
## [3618,] "Bacteria" "Firmicutes"          "Clostridia"                 
## [3619,] "Bacteria" "Proteobacteria"      "Gammaproteobacteria"        
## [3620,] "Bacteria" "Actinobacteria"      "Coriobacteriia"             
## [3621,] "Bacteria" "Actinobacteria"      "Actinobacteria"             
## [3622,] "Bacteria" "Bacteroidetes"       "Bacteroidia"                
## [3623,] "Bacteria" "Proteobacteria"      "Gammaproteobacteria"        
## [3624,] "Bacteria" "Bacteroidetes"       "Bacteroidia"                
## [3625,] "Bacteria" "Bacteroidetes"       "Bacteroidia"                
## [3626,] "Bacteria" "Bacteroidetes"       "Bacteroidia"                
## [3627,] "Bacteria" "Proteobacteria"      "Gammaproteobacteria"        
## [3628,] "Bacteria" "Firmicutes"          "Clostridia"                 
## [3629,] "Bacteria" "Bacteroidetes"       "Bacteroidia"                
## [3630,] "Bacteria" "Patescibacteria"     "Saccharimonadia"            
## [3631,] "Bacteria" "Bacteroidetes"       "Bacteroidia"                
## [3632,] "Bacteria" "Bacteroidetes"       "Bacteroidia"                
## [3633,] "Bacteria" "Proteobacteria"      "Gammaproteobacteria"        
## [3634,] "Bacteria" "Firmicutes"          "Bacilli"                    
## [3635,] "Bacteria" "Bacteroidetes"       "Bacteroidia"                
## [3636,] "Bacteria" "Patescibacteria"     "Saccharimonadia"            
## [3637,] "Bacteria" "Bacteroidetes"       "Bacteroidia"                
## [3638,] "Bacteria" "Firmicutes"          "Clostridia"                 
## [3639,] "Bacteria" "Firmicutes"          "Negativicutes"              
## [3640,] "Bacteria" "Firmicutes"          "Negativicutes"              
## [3641,] "Bacteria" "Firmicutes"          "Negativicutes"              
## [3642,] "Bacteria" "Proteobacteria"      "Alphaproteobacteria"        
## [3643,] "Bacteria" "Proteobacteria"      "Gammaproteobacteria"        
## [3644,] "Bacteria" "Bacteroidetes"       "Bacteroidia"                
## [3645,] "Bacteria" "Proteobacteria"      "Gammaproteobacteria"        
## [3646,] "Bacteria" "Fusobacteria"        "Fusobacteriia"              
## [3647,] "Bacteria" "Actinobacteria"      "Actinobacteria"             
## [3648,] "Bacteria" "Firmicutes"          NA                           
## [3649,] "Bacteria" "Firmicutes"          "Clostridia"                 
## [3650,] "Bacteria" "Spirochaetes"        "Spirochaetia"               
## [3651,] "Bacteria" "Firmicutes"          "Clostridia"                 
## [3652,] "Bacteria" "Bacteroidetes"       "Bacteroidia"                
## [3653,] "Bacteria" NA                    NA                           
## [3654,] "Bacteria" "Bacteroidetes"       "Bacteroidia"                
## [3655,] "Bacteria" "Bacteroidetes"       "Bacteroidia"                
## [3656,] "Bacteria" "Proteobacteria"      "Gammaproteobacteria"        
## [3657,] "Bacteria" NA                    NA                           
## [3658,] "Bacteria" "Bacteroidetes"       "Bacteroidia"                
## [3659,] "Bacteria" "Firmicutes"          "Negativicutes"              
## [3660,] "Bacteria" "Epsilonbacteraeota"  "Campylobacteria"            
## [3661,] "Bacteria" "Firmicutes"          "Clostridia"                 
## [3662,] "Bacteria" "Actinobacteria"      "Actinobacteria"             
## [3663,] "Bacteria" "Bacteroidetes"       "Bacteroidia"                
## [3664,] "Bacteria" "Bacteroidetes"       "Bacteroidia"                
## [3665,] "Bacteria" "Bacteroidetes"       "Bacteroidia"                
## [3666,] "Bacteria" "Firmicutes"          "Negativicutes"              
## [3667,] "Bacteria" "Proteobacteria"      "Gammaproteobacteria"        
## [3668,] "Bacteria" "Proteobacteria"      "Gammaproteobacteria"        
## [3669,] "Bacteria" NA                    NA                           
## [3670,] "Bacteria" "Actinobacteria"      "Coriobacteriia"             
## [3671,] "Bacteria" "Firmicutes"          NA                           
## [3672,] "Bacteria" "Firmicutes"          NA                           
## [3673,] "Bacteria" "Bacteroidetes"       "Bacteroidia"                
## [3674,] "Bacteria" "Bacteroidetes"       "Bacteroidia"                
## [3675,] "Bacteria" "Firmicutes"          "Clostridia"                 
## [3676,] "Bacteria" "Bacteroidetes"       "Bacteroidia"                
## [3677,] "Bacteria" "Spirochaetes"        "Spirochaetia"               
## [3678,] "Bacteria" "Bacteroidetes"       "Bacteroidia"                
## [3679,] "Bacteria" "Bacteroidetes"       "Bacteroidia"                
## [3680,] "Bacteria" "Bacteroidetes"       "Bacteroidia"                
## [3681,] "Bacteria" "Firmicutes"          "Negativicutes"              
## [3682,] "Bacteria" "Bacteroidetes"       "Bacteroidia"                
## [3683,] "Bacteria" "Proteobacteria"      "Gammaproteobacteria"        
## [3684,] "Bacteria" "Proteobacteria"      "Gammaproteobacteria"        
## [3685,] "Bacteria" "Firmicutes"          "Bacilli"                    
## [3686,] "Bacteria" "Firmicutes"          "Clostridia"                 
## [3687,] "Bacteria" "Firmicutes"          NA                           
## [3688,] "Bacteria" "Bacteroidetes"       "Bacteroidia"                
## [3689,] "Bacteria" "Bacteroidetes"       "Bacteroidia"                
## [3690,] "Bacteria" "Firmicutes"          "Clostridia"                 
## [3691,] "Bacteria" "Fusobacteria"        "Fusobacteriia"              
## [3692,] "Bacteria" "Fusobacteria"        "Fusobacteriia"              
## [3693,] "Bacteria" "Bacteroidetes"       "Bacteroidia"                
## [3694,] "Bacteria" "Spirochaetes"        "Spirochaetia"               
## [3695,] "Bacteria" "Firmicutes"          "Clostridia"                 
## [3696,] "Bacteria" "Firmicutes"          "Negativicutes"              
## [3697,] "Bacteria" "Firmicutes"          "Clostridia"                 
## [3698,] "Bacteria" "Proteobacteria"      "Gammaproteobacteria"        
## [3699,] "Bacteria" "Fusobacteria"        "Fusobacteriia"              
## [3700,] "Bacteria" "Bacteroidetes"       "Bacteroidia"                
## [3701,] "Bacteria" "Firmicutes"          "Negativicutes"              
## [3702,] "Bacteria" "Epsilonbacteraeota"  "Campylobacteria"            
## [3703,] "Bacteria" NA                    NA                           
## [3704,] "Bacteria" "Fusobacteria"        "Fusobacteriia"              
## [3705,] "Bacteria" "Proteobacteria"      "Gammaproteobacteria"        
## [3706,] "Bacteria" "Firmicutes"          NA                           
## [3707,] "Bacteria" "Bacteroidetes"       "Bacteroidia"                
## [3708,] "Bacteria" "Proteobacteria"      "Gammaproteobacteria"        
## [3709,] "Bacteria" "Bacteroidetes"       "Bacteroidia"                
## [3710,] "Bacteria" "Bacteroidetes"       "Bacteroidia"                
## [3711,] "Bacteria" "Firmicutes"          "Clostridia"                 
## [3712,] "Bacteria" "Actinobacteria"      "Coriobacteriia"             
## [3713,] "Bacteria" "Firmicutes"          "Bacilli"                    
## [3714,] "Bacteria" "Firmicutes"          "Clostridia"                 
## [3715,] "Bacteria" "Firmicutes"          "Clostridia"                 
## [3716,] "Bacteria" "Proteobacteria"      "Gammaproteobacteria"        
## [3717,] "Bacteria" "Firmicutes"          "Bacilli"                    
## [3718,] "Bacteria" "Fusobacteria"        "Fusobacteriia"              
## [3719,] "Bacteria" "Patescibacteria"     "Saccharimonadia"            
## [3720,] "Bacteria" "Firmicutes"          "Bacilli"                    
## [3721,] "Bacteria" "Firmicutes"          "Negativicutes"              
## [3722,] "Bacteria" "Firmicutes"          "Bacilli"                    
## [3723,] "Bacteria" "Proteobacteria"      "Gammaproteobacteria"        
## [3724,] "Bacteria" "Firmicutes"          "Clostridia"                 
## [3725,] "Bacteria" "Firmicutes"          "Clostridia"                 
## [3726,] "Bacteria" "Epsilonbacteraeota"  "Campylobacteria"            
## [3727,] "Bacteria" "Epsilonbacteraeota"  "Campylobacteria"            
## [3728,] "Bacteria" "Firmicutes"          "Clostridia"                 
## [3729,] "Bacteria" "Firmicutes"          "Clostridia"                 
## [3730,] "Bacteria" "Firmicutes"          "Negativicutes"              
## [3731,] "Bacteria" "Firmicutes"          "Bacilli"                    
## [3732,] "Bacteria" "Spirochaetes"        "Spirochaetia"               
## [3733,] "Bacteria" "Firmicutes"          "Negativicutes"              
## [3734,] "Bacteria" "Bacteroidetes"       "Bacteroidia"                
## [3735,] "Bacteria" "Fusobacteria"        "Fusobacteriia"              
## [3736,] "Bacteria" "Firmicutes"          "Bacilli"                    
## [3737,] "Bacteria" "Bacteroidetes"       "Bacteroidia"                
## [3738,] "Bacteria" "Firmicutes"          "Clostridia"                 
## [3739,] "Bacteria" "Bacteroidetes"       "Bacteroidia"                
## [3740,] "Bacteria" "Bacteroidetes"       "Bacteroidia"                
## [3741,] "Bacteria" "Bacteroidetes"       "Bacteroidia"                
## [3742,] "Bacteria" "Proteobacteria"      "Gammaproteobacteria"        
## [3743,] "Bacteria" "Epsilonbacteraeota"  "Campylobacteria"            
## [3744,] "Bacteria" "Firmicutes"          "Bacilli"                    
## [3745,] "Bacteria" "Patescibacteria"     "Saccharimonadia"            
## [3746,] "Bacteria" "Bacteroidetes"       "Bacteroidia"                
## [3747,] "Bacteria" "Bacteroidetes"       "Bacteroidia"                
## [3748,] "Bacteria" "Bacteroidetes"       "Bacteroidia"                
## [3749,] "Bacteria" "Bacteroidetes"       "Bacteroidia"                
## [3750,] "Bacteria" "Epsilonbacteraeota"  "Campylobacteria"            
## [3751,] "Bacteria" "Firmicutes"          "Clostridia"                 
## [3752,] "Bacteria" "Firmicutes"          "Bacilli"                    
## [3753,] "Bacteria" "Firmicutes"          "Negativicutes"              
## [3754,] "Bacteria" "Bacteroidetes"       "Bacteroidia"                
## [3755,] "Bacteria" "Firmicutes"          "Clostridia"                 
## [3756,] "Bacteria" "Firmicutes"          "Clostridia"                 
## [3757,] "Bacteria" "Bacteroidetes"       "Bacteroidia"                
## [3758,] "Bacteria" "Firmicutes"          "Bacilli"                    
## [3759,] "Bacteria" "Firmicutes"          "Negativicutes"              
## [3760,] "Bacteria" "Patescibacteria"     "Saccharimonadia"            
## [3761,] "Bacteria" "Bacteroidetes"       "Bacteroidia"                
## [3762,] "Bacteria" "Firmicutes"          "Bacilli"                    
## [3763,] "Bacteria" "Spirochaetes"        "Spirochaetia"               
## [3764,] "Bacteria" "Bacteroidetes"       "Bacteroidia"                
## [3765,] "Bacteria" "Bacteroidetes"       "Bacteroidia"                
## [3766,] "Bacteria" "Bacteroidetes"       "Bacteroidia"                
## [3767,] "Bacteria" "Proteobacteria"      "Gammaproteobacteria"        
## [3768,] "Bacteria" "Firmicutes"          "Negativicutes"              
## [3769,] "Bacteria" "Actinobacteria"      "Actinobacteria"             
## [3770,] "Bacteria" "Proteobacteria"      "Gammaproteobacteria"        
## [3771,] "Bacteria" "Bacteroidetes"       "Bacteroidia"                
## [3772,] "Bacteria" "Patescibacteria"     "Saccharimonadia"            
## [3773,] "Bacteria" "Firmicutes"          "Clostridia"                 
## [3774,] "Bacteria" "Firmicutes"          "Clostridia"                 
## [3775,] "Bacteria" "Firmicutes"          "Negativicutes"              
## [3776,] "Bacteria" "Bacteroidetes"       "Bacteroidia"                
## [3777,] "Bacteria" "Bacteroidetes"       "Bacteroidia"                
## [3778,] "Bacteria" "Bacteroidetes"       "Bacteroidia"                
## [3779,] "Bacteria" "Bacteroidetes"       "Bacteroidia"                
## [3780,] "Bacteria" "Firmicutes"          "Clostridia"                 
## [3781,] "Bacteria" "Bacteroidetes"       "Bacteroidia"                
## [3782,] "Bacteria" "Bacteroidetes"       "Bacteroidia"                
## [3783,] "Bacteria" "Proteobacteria"      "Gammaproteobacteria"        
## [3784,] "Bacteria" "Bacteroidetes"       "Bacteroidia"                
## [3785,] "Bacteria" "Fusobacteria"        "Fusobacteriia"              
## [3786,] "Bacteria" "Bacteroidetes"       "Bacteroidia"                
## [3787,] "Bacteria" NA                    NA                           
## [3788,] "Bacteria" "Patescibacteria"     "Saccharimonadia"            
## [3789,] "Bacteria" "Proteobacteria"      "Gammaproteobacteria"        
## [3790,] "Bacteria" "Firmicutes"          "Negativicutes"              
## [3791,] "Bacteria" "Spirochaetes"        "Spirochaetia"               
## [3792,] "Bacteria" "Patescibacteria"     "Saccharimonadia"            
## [3793,] "Bacteria" "Bacteroidetes"       "Bacteroidia"                
## [3794,] "Bacteria" "Bacteroidetes"       "Bacteroidia"                
## [3795,] "Bacteria" "Firmicutes"          "Negativicutes"              
## [3796,] "Bacteria" "Patescibacteria"     "Saccharimonadia"            
## [3797,] "Bacteria" "Patescibacteria"     "Gracilibacteria"            
## [3798,] "Bacteria" "Firmicutes"          "Bacilli"                    
## [3799,] "Bacteria" "Bacteroidetes"       "Bacteroidia"                
## [3800,] "Bacteria" "Proteobacteria"      "Gammaproteobacteria"        
## [3801,] "Bacteria" "Firmicutes"          "Negativicutes"              
## [3802,] "Bacteria" "Bacteroidetes"       "Bacteroidia"                
## [3803,] "Bacteria" "Proteobacteria"      "Gammaproteobacteria"        
## [3804,] "Bacteria" "Firmicutes"          "Clostridia"                 
## [3805,] "Bacteria" "Firmicutes"          "Bacilli"                    
## [3806,] "Bacteria" "Patescibacteria"     "Saccharimonadia"            
## [3807,] "Bacteria" "Firmicutes"          "Clostridia"                 
## [3808,] "Bacteria" "Bacteroidetes"       "Bacteroidia"                
## [3809,] "Bacteria" "Actinobacteria"      "Actinobacteria"             
## [3810,] "Bacteria" "Actinobacteria"      "Actinobacteria"             
## [3811,] "Bacteria" "Firmicutes"          "Negativicutes"              
## [3812,] "Bacteria" "Firmicutes"          "Negativicutes"              
## [3813,] "Bacteria" "Firmicutes"          "Bacilli"                    
## [3814,] "Bacteria" "Firmicutes"          "Bacilli"                    
## [3815,] "Bacteria" "Bacteroidetes"       "Bacteroidia"                
## [3816,] "Bacteria" "Bacteroidetes"       "Bacteroidia"                
## [3817,] "Bacteria" "Firmicutes"          "Negativicutes"              
## [3818,] "Bacteria" "Firmicutes"          "Negativicutes"              
## [3819,] "Bacteria" "Bacteroidetes"       "Bacteroidia"                
## [3820,] "Bacteria" "Bacteroidetes"       "Bacteroidia"                
## [3821,] "Bacteria" "Patescibacteria"     "Gracilibacteria"            
## [3822,] "Bacteria" "Spirochaetes"        "Spirochaetia"               
## [3823,] "Bacteria" "Bacteroidetes"       "Bacteroidia"                
## [3824,] "Bacteria" "Firmicutes"          "Negativicutes"              
## [3825,] "Bacteria" "Spirochaetes"        "Spirochaetia"               
## [3826,] "Bacteria" "Firmicutes"          "Clostridia"                 
## [3827,] "Bacteria" "Patescibacteria"     "Saccharimonadia"            
## [3828,] "Bacteria" "Proteobacteria"      "Gammaproteobacteria"        
## [3829,] "Bacteria" "Firmicutes"          "Bacilli"                    
## [3830,] "Bacteria" "Patescibacteria"     "Saccharimonadia"            
## [3831,] "Bacteria" "Bacteroidetes"       "Bacteroidia"                
## [3832,] "Bacteria" "Proteobacteria"      "Gammaproteobacteria"        
## [3833,] "Bacteria" "Fusobacteria"        "Fusobacteriia"              
## [3834,] "Bacteria" "Actinobacteria"      "Actinobacteria"             
## [3835,] "Bacteria" "Bacteroidetes"       "Bacteroidia"                
## [3836,] "Bacteria" "Bacteroidetes"       "Bacteroidia"                
## [3837,] "Bacteria" "Firmicutes"          "Bacilli"                    
## [3838,] "Bacteria" "Bacteroidetes"       "Bacteroidia"                
## [3839,] "Bacteria" "Bacteroidetes"       "Bacteroidia"                
## [3840,] "Bacteria" "Firmicutes"          "Clostridia"                 
## [3841,] "Bacteria" "Bacteroidetes"       "Bacteroidia"                
## [3842,] "Bacteria" "Bacteroidetes"       "Bacteroidia"                
## [3843,] "Bacteria" "Bacteroidetes"       "Bacteroidia"                
## [3844,] "Bacteria" "Bacteroidetes"       "Bacteroidia"                
## [3845,] "Bacteria" "Firmicutes"          NA                           
## [3846,] "Bacteria" "Epsilonbacteraeota"  "Campylobacteria"            
## [3847,] "Bacteria" "Bacteroidetes"       "Bacteroidia"                
## [3848,] "Bacteria" "Firmicutes"          "Clostridia"                 
## [3849,] "Bacteria" "Spirochaetes"        "Spirochaetia"               
## [3850,] "Bacteria" "Firmicutes"          "Clostridia"                 
## [3851,] "Bacteria" "Proteobacteria"      "Gammaproteobacteria"        
## [3852,] "Bacteria" "Proteobacteria"      "Gammaproteobacteria"        
## [3853,] "Bacteria" "Bacteroidetes"       "Bacteroidia"                
## [3854,] "Bacteria" "Firmicutes"          "Bacilli"                    
## [3855,] "Bacteria" "Epsilonbacteraeota"  "Campylobacteria"            
## [3856,] "Bacteria" "Firmicutes"          "Bacilli"                    
## [3857,] "Bacteria" "Fusobacteria"        "Fusobacteriia"              
## [3858,] "Bacteria" "Bacteroidetes"       "Bacteroidia"                
## [3859,] "Bacteria" "Bacteroidetes"       "Bacteroidia"                
## [3860,] "Bacteria" "Bacteroidetes"       "Bacteroidia"                
## [3861,] "Bacteria" "Firmicutes"          "Bacilli"                    
## [3862,] "Bacteria" "Firmicutes"          "Clostridia"                 
## [3863,] "Bacteria" "Firmicutes"          "Clostridia"                 
## [3864,] "Bacteria" "Bacteroidetes"       "Bacteroidia"                
## [3865,] "Bacteria" "Fusobacteria"        "Fusobacteriia"              
## [3866,] "Bacteria" "Firmicutes"          "Clostridia"                 
## [3867,] "Bacteria" "Fusobacteria"        "Fusobacteriia"              
## [3868,] "Bacteria" "Bacteroidetes"       "Bacteroidia"                
## [3869,] "Bacteria" "Patescibacteria"     "Saccharimonadia"            
## [3870,] "Bacteria" "Bacteroidetes"       "Bacteroidia"                
## [3871,] "Bacteria" "Proteobacteria"      "Gammaproteobacteria"        
## [3872,] "Bacteria" "Proteobacteria"      "Gammaproteobacteria"        
## [3873,] "Bacteria" "Firmicutes"          "Bacilli"                    
## [3874,] "Bacteria" "Bacteroidetes"       "Bacteroidia"                
## [3875,] "Bacteria" "Firmicutes"          NA                           
## [3876,] "Bacteria" "Proteobacteria"      "Gammaproteobacteria"        
## [3877,] "Bacteria" "Bacteroidetes"       "Bacteroidia"                
## [3878,] "Bacteria" "Epsilonbacteraeota"  "Campylobacteria"            
## [3879,] "Bacteria" "Bacteroidetes"       "Bacteroidia"                
## [3880,] "Bacteria" "Firmicutes"          "Clostridia"                 
## [3881,] "Bacteria" "Spirochaetes"        "Spirochaetia"               
## [3882,] "Bacteria" "Bacteroidetes"       "Bacteroidia"                
## [3883,] "Bacteria" "Firmicutes"          "Bacilli"                    
## [3884,] "Bacteria" "Firmicutes"          "Negativicutes"              
## [3885,] "Bacteria" "Proteobacteria"      "Gammaproteobacteria"        
## [3886,] "Bacteria" "Firmicutes"          "Clostridia"                 
## [3887,] "Bacteria" "Firmicutes"          "Clostridia"                 
## [3888,] "Bacteria" "Firmicutes"          "Clostridia"                 
## [3889,] "Bacteria" "Firmicutes"          "Bacilli"                    
## [3890,] "Bacteria" "Firmicutes"          "Clostridia"                 
## [3891,] "Bacteria" "Patescibacteria"     "Saccharimonadia"            
## [3892,] "Bacteria" "Firmicutes"          NA                           
## [3893,] "Bacteria" "Firmicutes"          "Clostridia"                 
## [3894,] "Bacteria" "Firmicutes"          "Clostridia"                 
## [3895,] "Bacteria" "Actinobacteria"      "Actinobacteria"             
## [3896,] "Bacteria" "Bacteroidetes"       "Bacteroidia"                
## [3897,] "Bacteria" "Epsilonbacteraeota"  "Campylobacteria"            
## [3898,] "Bacteria" "Proteobacteria"      "Gammaproteobacteria"        
## [3899,] "Bacteria" "Proteobacteria"      "Gammaproteobacteria"        
## [3900,] "Bacteria" "Firmicutes"          "Bacilli"                    
## [3901,] "Bacteria" "Firmicutes"          "Bacilli"                    
## [3902,] "Bacteria" "Firmicutes"          "Negativicutes"              
## [3903,] "Bacteria" "Bacteroidetes"       "Bacteroidia"                
## [3904,] "Bacteria" "Bacteroidetes"       "Bacteroidia"                
## [3905,] "Bacteria" "Spirochaetes"        "Spirochaetia"               
## [3906,] "Bacteria" "Firmicutes"          "Clostridia"                 
## [3907,] "Bacteria" "Bacteroidetes"       "Bacteroidia"                
## [3908,] "Bacteria" "Proteobacteria"      "Gammaproteobacteria"        
## [3909,] "Bacteria" "Patescibacteria"     "Gracilibacteria"            
## [3910,] "Bacteria" "Bacteroidetes"       "Bacteroidia"                
## [3911,] "Bacteria" "Bacteroidetes"       "Bacteroidia"                
## [3912,] "Bacteria" "Epsilonbacteraeota"  "Campylobacteria"            
## [3913,] "Bacteria" "Firmicutes"          "Negativicutes"              
## [3914,] "Bacteria" "Firmicutes"          "Erysipelotrichia"           
## [3915,] "Bacteria" "Firmicutes"          "Bacilli"                    
## [3916,] "Bacteria" "Firmicutes"          "Bacilli"                    
## [3917,] "Bacteria" "Bacteroidetes"       "Bacteroidia"                
## [3918,] "Bacteria" "Fusobacteria"        "Fusobacteriia"              
## [3919,] "Bacteria" "Proteobacteria"      "Gammaproteobacteria"        
## [3920,] "Bacteria" "Firmicutes"          "Negativicutes"              
## [3921,] "Bacteria" "Actinobacteria"      "Actinobacteria"             
## [3922,] "Bacteria" "Bacteroidetes"       "Bacteroidia"                
## [3923,] "Bacteria" "Firmicutes"          NA                           
## [3924,] "Bacteria" "Proteobacteria"      "Gammaproteobacteria"        
## [3925,] "Bacteria" "Actinobacteria"      "Actinobacteria"             
## [3926,] "Bacteria" "Firmicutes"          "Negativicutes"              
## [3927,] "Bacteria" "Fusobacteria"        "Fusobacteriia"              
## [3928,] "Bacteria" "Proteobacteria"      "Gammaproteobacteria"        
## [3929,] "Bacteria" "Deinococcus-Thermus" "Deinococci"                 
## [3930,] "Bacteria" "Epsilonbacteraeota"  "Campylobacteria"            
## [3931,] "Bacteria" "Firmicutes"          "Clostridia"                 
## [3932,] "Bacteria" "Firmicutes"          "Clostridia"                 
## [3933,] "Bacteria" NA                    NA                           
## [3934,] "Bacteria" "Firmicutes"          "Clostridia"                 
## [3935,] "Bacteria" "Proteobacteria"      "Gammaproteobacteria"        
## [3936,] "Bacteria" "Spirochaetes"        "Spirochaetia"               
## [3937,] "Bacteria" "Firmicutes"          "Bacilli"                    
## [3938,] "Bacteria" "Proteobacteria"      "Gammaproteobacteria"        
## [3939,] "Bacteria" "Firmicutes"          "Bacilli"                    
## [3940,] "Bacteria" "Firmicutes"          "Clostridia"                 
## [3941,] "Bacteria" "Bacteroidetes"       "Bacteroidia"                
## [3942,] "Bacteria" "Bacteroidetes"       "Bacteroidia"                
## [3943,] "Bacteria" "Proteobacteria"      "Gammaproteobacteria"        
## [3944,] "Bacteria" "Firmicutes"          "Clostridia"                 
## [3945,] "Bacteria" "Bacteroidetes"       "Bacteroidia"                
## [3946,] "Bacteria" "Firmicutes"          "Bacilli"                    
## [3947,] "Bacteria" "Firmicutes"          "Negativicutes"              
## [3948,] "Bacteria" "Firmicutes"          "Bacilli"                    
## [3949,] "Bacteria" "Bacteroidetes"       "Bacteroidia"                
## [3950,] "Bacteria" "Firmicutes"          "Bacilli"                    
## [3951,] "Bacteria" "Firmicutes"          "Bacilli"                    
## [3952,] "Bacteria" "Bacteroidetes"       "Bacteroidia"                
## [3953,] "Bacteria" "Fusobacteria"        "Fusobacteriia"              
## [3954,] "Bacteria" "Proteobacteria"      "Gammaproteobacteria"        
## [3955,] "Bacteria" "Epsilonbacteraeota"  "Campylobacteria"            
## [3956,] "Bacteria" NA                    NA                           
## [3957,] "Bacteria" "Epsilonbacteraeota"  "Campylobacteria"            
## [3958,] "Bacteria" "Patescibacteria"     "Saccharimonadia"            
## [3959,] "Bacteria" "Bacteroidetes"       "Bacteroidia"                
## [3960,] "Bacteria" "Firmicutes"          "Negativicutes"              
## [3961,] "Bacteria" "Fusobacteria"        "Fusobacteriia"              
## [3962,] "Bacteria" "Patescibacteria"     "Saccharimonadia"            
## [3963,] "Bacteria" "Bacteroidetes"       "Bacteroidia"                
## [3964,] "Bacteria" "Bacteroidetes"       "Bacteroidia"                
## [3965,] "Bacteria" "Bacteroidetes"       "Bacteroidia"                
## [3966,] "Bacteria" "Firmicutes"          "Bacilli"                    
## [3967,] "Bacteria" NA                    NA                           
## [3968,] "Bacteria" "Fusobacteria"        "Fusobacteriia"              
## [3969,] "Bacteria" "Fusobacteria"        "Fusobacteriia"              
## [3970,] "Bacteria" "Epsilonbacteraeota"  "Campylobacteria"            
## [3971,] "Bacteria" "Firmicutes"          NA                           
## [3972,] "Bacteria" "Tenericutes"         "Mollicutes"                 
## [3973,] "Bacteria" "Tenericutes"         "Mollicutes"                 
## [3974,] "Bacteria" "Bacteroidetes"       "Bacteroidia"                
## [3975,] "Bacteria" "Proteobacteria"      "Deltaproteobacteria"        
## [3976,] "Bacteria" "Patescibacteria"     "Gracilibacteria"            
## [3977,] "Bacteria" "Bacteroidetes"       "Bacteroidia"                
## [3978,] "Bacteria" "Bacteroidetes"       "Bacteroidia"                
## [3979,] "Bacteria" "Firmicutes"          "Bacilli"                    
## [3980,] "Bacteria" "Fusobacteria"        "Fusobacteriia"              
## [3981,] "Bacteria" "Firmicutes"          "Clostridia"                 
## [3982,] "Bacteria" "Firmicutes"          "Bacilli"                    
## [3983,] "Bacteria" "Bacteroidetes"       "Bacteroidia"                
## [3984,] "Bacteria" "Firmicutes"          "Negativicutes"              
## [3985,] "Bacteria" "Firmicutes"          "Clostridia"                 
## [3986,] "Bacteria" "Bacteroidetes"       "Bacteroidia"                
## [3987,] "Bacteria" "Bacteroidetes"       "Bacteroidia"                
## [3988,] "Bacteria" "Bacteroidetes"       "Bacteroidia"                
## [3989,] "Bacteria" "Actinobacteria"      "Coriobacteriia"             
## [3990,] "Bacteria" "Firmicutes"          "Negativicutes"              
## [3991,] "Bacteria" "Fusobacteria"        "Fusobacteriia"              
## [3992,] "Bacteria" "Epsilonbacteraeota"  "Campylobacteria"            
## [3993,] "Bacteria" "Bacteroidetes"       "Bacteroidia"                
## [3994,] "Bacteria" "Firmicutes"          "Bacilli"                    
## [3995,] "Bacteria" "Spirochaetes"        "Spirochaetia"               
## [3996,] "Bacteria" "Bacteroidetes"       "Bacteroidia"                
## [3997,] "Bacteria" "Firmicutes"          "Clostridia"                 
## [3998,] "Bacteria" "Bacteroidetes"       "Bacteroidia"                
## [3999,] "Bacteria" "Patescibacteria"     "Saccharimonadia"            
## [4000,] "Bacteria" "Firmicutes"          NA                           
## [4001,] "Bacteria" "Firmicutes"          "Bacilli"                    
## [4002,] "Bacteria" "Spirochaetes"        "Spirochaetia"               
## [4003,] "Bacteria" "Firmicutes"          "Bacilli"                    
## [4004,] "Bacteria" "Bacteroidetes"       "Bacteroidia"                
## [4005,] "Bacteria" "Fusobacteria"        "Fusobacteriia"              
## [4006,] "Bacteria" "Bacteroidetes"       "Bacteroidia"                
## [4007,] "Bacteria" "Firmicutes"          "Negativicutes"              
## [4008,] "Bacteria" "Bacteroidetes"       "Bacteroidia"                
## [4009,] "Bacteria" "Bacteroidetes"       "Bacteroidia"                
## [4010,] "Bacteria" "Proteobacteria"      "Gammaproteobacteria"        
## [4011,] "Bacteria" "Firmicutes"          "Negativicutes"              
## [4012,] "Bacteria" "Bacteroidetes"       "Bacteroidia"                
## [4013,] "Bacteria" "Firmicutes"          "Clostridia"                 
## [4014,] "Bacteria" "Firmicutes"          "Negativicutes"              
## [4015,] "Bacteria" "Actinobacteria"      "Actinobacteria"             
## [4016,] "Bacteria" "Firmicutes"          "Bacilli"                    
## [4017,] "Bacteria" "Bacteroidetes"       "Bacteroidia"                
## [4018,] "Bacteria" "Firmicutes"          "Clostridia"                 
## [4019,] "Bacteria" "Firmicutes"          "Clostridia"                 
## [4020,] "Bacteria" "Actinobacteria"      "Actinobacteria"             
## [4021,] "Bacteria" "Bacteroidetes"       "Bacteroidia"                
## [4022,] "Bacteria" "Actinobacteria"      "Actinobacteria"             
## [4023,] "Bacteria" "Firmicutes"          "Negativicutes"              
## [4024,] "Bacteria" "Patescibacteria"     "Saccharimonadia"            
## [4025,] "Bacteria" "Bacteroidetes"       "Bacteroidia"                
## [4026,] "Bacteria" "Bacteroidetes"       "Bacteroidia"                
## [4027,] "Bacteria" "Firmicutes"          "Bacilli"                    
## [4028,] "Bacteria" "Firmicutes"          "Negativicutes"              
## [4029,] "Bacteria" "Firmicutes"          "Bacilli"                    
## [4030,] "Bacteria" "Patescibacteria"     "Saccharimonadia"            
## [4031,] "Bacteria" "Proteobacteria"      "Gammaproteobacteria"        
## [4032,] "Bacteria" "Actinobacteria"      "Actinobacteria"             
## [4033,] "Bacteria" "Spirochaetes"        "Spirochaetia"               
## [4034,] "Bacteria" "Firmicutes"          "Bacilli"                    
## [4035,] "Bacteria" "Fusobacteria"        "Fusobacteriia"              
## [4036,] "Bacteria" "Firmicutes"          "Clostridia"                 
## [4037,] "Bacteria" "Patescibacteria"     "Gracilibacteria"            
## [4038,] "Bacteria" "Firmicutes"          "Negativicutes"              
## [4039,] "Bacteria" "Proteobacteria"      "Gammaproteobacteria"        
## [4040,] "Bacteria" "Firmicutes"          "Bacilli"                    
## [4041,] "Bacteria" "Bacteroidetes"       "Bacteroidia"                
## [4042,] "Bacteria" "Fusobacteria"        "Fusobacteriia"              
## [4043,] "Bacteria" "Bacteroidetes"       "Bacteroidia"                
## [4044,] "Bacteria" "Bacteroidetes"       "Bacteroidia"                
## [4045,] "Bacteria" "Firmicutes"          "Clostridia"                 
## [4046,] "Bacteria" "Patescibacteria"     "Saccharimonadia"            
## [4047,] "Bacteria" "Bacteroidetes"       "Bacteroidia"                
## [4048,] "Bacteria" NA                    NA                           
## [4049,] "Bacteria" "Fusobacteria"        "Fusobacteriia"              
## [4050,] "Bacteria" "Bacteroidetes"       "Bacteroidia"                
## [4051,] "Bacteria" "Bacteroidetes"       "Bacteroidia"                
## [4052,] "Bacteria" "Bacteroidetes"       "Bacteroidia"                
## [4053,] "Bacteria" "Firmicutes"          "Bacilli"                    
## [4054,] "Bacteria" "Bacteroidetes"       "Bacteroidia"                
## [4055,] "Bacteria" "Actinobacteria"      "Actinobacteria"             
## [4056,] "Bacteria" "Fusobacteria"        "Fusobacteriia"              
## [4057,] "Bacteria" "Proteobacteria"      "Gammaproteobacteria"        
## [4058,] "Bacteria" "Bacteroidetes"       "Bacteroidia"                
## [4059,] "Bacteria" "Firmicutes"          "Bacilli"                    
## [4060,] "Bacteria" "Fusobacteria"        "Fusobacteriia"              
## [4061,] "Bacteria" "Cyanobacteria"       "Oxyphotobacteria"           
## [4062,] "Bacteria" "Actinobacteria"      "Actinobacteria"             
## [4063,] "Bacteria" "Patescibacteria"     "Saccharimonadia"            
## [4064,] "Bacteria" "Fusobacteria"        "Fusobacteriia"              
## [4065,] "Bacteria" "Firmicutes"          NA                           
## [4066,] "Bacteria" "Bacteroidetes"       "Bacteroidia"                
## [4067,] "Bacteria" "Firmicutes"          "Clostridia"                 
## [4068,] "Bacteria" "Firmicutes"          "Clostridia"                 
## [4069,] "Bacteria" "Bacteroidetes"       "Bacteroidia"                
## [4070,] "Bacteria" "Actinobacteria"      "Actinobacteria"             
## [4071,] "Bacteria" "Firmicutes"          "Bacilli"                    
## [4072,] "Bacteria" "Patescibacteria"     "Gracilibacteria"            
## [4073,] "Bacteria" "Firmicutes"          "Negativicutes"              
## [4074,] "Bacteria" "Firmicutes"          "Negativicutes"              
## [4075,] "Bacteria" "Firmicutes"          "Negativicutes"              
## [4076,] "Bacteria" "Firmicutes"          "Clostridia"                 
## [4077,] "Bacteria" "Bacteroidetes"       "Bacteroidia"                
## [4078,] "Bacteria" "Proteobacteria"      "Gammaproteobacteria"        
## [4079,] "Bacteria" "Bacteroidetes"       "Bacteroidia"                
## [4080,] "Bacteria" "Bacteroidetes"       "Bacteroidia"                
## [4081,] "Bacteria" "Firmicutes"          "Negativicutes"              
## [4082,] "Bacteria" "Firmicutes"          "Negativicutes"              
## [4083,] "Bacteria" "Firmicutes"          "Clostridia"                 
## [4084,] "Bacteria" "Epsilonbacteraeota"  "Campylobacteria"            
## [4085,] "Bacteria" "Cyanobacteria"       "Oxyphotobacteria"           
## [4086,] "Bacteria" "Firmicutes"          "Bacilli"                    
## [4087,] "Bacteria" "Bacteroidetes"       "Bacteroidia"                
## [4088,] "Bacteria" "Proteobacteria"      "Gammaproteobacteria"        
## [4089,] "Bacteria" "Actinobacteria"      "Actinobacteria"             
## [4090,] "Bacteria" NA                    NA                           
## [4091,] "Bacteria" "Actinobacteria"      "Actinobacteria"             
## [4092,] "Bacteria" "Firmicutes"          "Bacilli"                    
## [4093,] "Bacteria" "Firmicutes"          "Clostridia"                 
## [4094,] "Bacteria" "Firmicutes"          "Clostridia"                 
## [4095,] "Bacteria" "Proteobacteria"      "Gammaproteobacteria"        
## [4096,] "Bacteria" "Firmicutes"          "Clostridia"                 
## [4097,] "Bacteria" "Patescibacteria"     "Saccharimonadia"            
## [4098,] "Bacteria" "Proteobacteria"      "Gammaproteobacteria"        
## [4099,] "Bacteria" "Bacteroidetes"       "Bacteroidia"                
## [4100,] "Bacteria" "Firmicutes"          "Clostridia"                 
## [4101,] "Bacteria" "Actinobacteria"      "Actinobacteria"             
## [4102,] "Bacteria" "Bacteroidetes"       "Bacteroidia"                
## [4103,] "Bacteria" "Firmicutes"          "Clostridia"                 
## [4104,] "Bacteria" "Firmicutes"          "Clostridia"                 
## [4105,] "Bacteria" "Bacteroidetes"       "Bacteroidia"                
## [4106,] "Bacteria" "Bacteroidetes"       "Bacteroidia"                
## [4107,] "Bacteria" "Patescibacteria"     "Saccharimonadia"            
## [4108,] "Bacteria" "Patescibacteria"     "Saccharimonadia"            
## [4109,] "Bacteria" "Bacteroidetes"       "Bacteroidia"                
## [4110,] "Bacteria" "Firmicutes"          "Negativicutes"              
## [4111,] "Bacteria" "Firmicutes"          "Bacilli"                    
## [4112,] "Bacteria" "Firmicutes"          "Negativicutes"              
## [4113,] "Bacteria" "Fusobacteria"        "Fusobacteriia"              
## [4114,] "Bacteria" "Actinobacteria"      "Actinobacteria"             
## [4115,] "Bacteria" "Actinobacteria"      "Actinobacteria"             
## [4116,] "Bacteria" "Firmicutes"          "Clostridia"                 
## [4117,] "Bacteria" "Actinobacteria"      "Coriobacteriia"             
## [4118,] "Bacteria" "Bacteroidetes"       "Bacteroidia"                
## [4119,] "Bacteria" "Actinobacteria"      "Coriobacteriia"             
## [4120,] "Bacteria" "Bacteroidetes"       "Bacteroidia"                
## [4121,] "Bacteria" "Elusimicrobia"       "Elusimicrobia"              
## [4122,] "Bacteria" "Epsilonbacteraeota"  "Campylobacteria"            
## [4123,] "Bacteria" "Proteobacteria"      NA                           
## [4124,] "Bacteria" "Epsilonbacteraeota"  "Campylobacteria"            
## [4125,] "Bacteria" "Bacteroidetes"       "Bacteroidia"                
## [4126,] "Bacteria" "Patescibacteria"     "Saccharimonadia"            
## [4127,] "Bacteria" "Firmicutes"          "Bacilli"                    
## [4128,] "Bacteria" "Proteobacteria"      "Gammaproteobacteria"        
## [4129,] "Bacteria" "Bacteroidetes"       "Bacteroidia"                
## [4130,] "Bacteria" "Firmicutes"          "Clostridia"                 
## [4131,] "Bacteria" "Firmicutes"          "Bacilli"                    
## [4132,] "Bacteria" "Firmicutes"          "Clostridia"                 
## [4133,] "Bacteria" "Bacteroidetes"       "Bacteroidia"                
## [4134,] "Bacteria" "Bacteroidetes"       "Bacteroidia"                
## [4135,] "Bacteria" "Patescibacteria"     "Saccharimonadia"            
## [4136,] "Bacteria" "Patescibacteria"     "Saccharimonadia"            
## [4137,] "Bacteria" "Proteobacteria"      "Gammaproteobacteria"        
## [4138,] "Bacteria" "Patescibacteria"     "Saccharimonadia"            
## [4139,] "Bacteria" "Proteobacteria"      "Gammaproteobacteria"        
## [4140,] "Bacteria" "Firmicutes"          "Clostridia"                 
## [4141,] "Bacteria" "Proteobacteria"      "Gammaproteobacteria"        
## [4142,] "Bacteria" "Firmicutes"          "Clostridia"                 
## [4143,] "Bacteria" "Firmicutes"          "Clostridia"                 
## [4144,] "Bacteria" "Bacteroidetes"       "Bacteroidia"                
## [4145,] "Bacteria" "Firmicutes"          "Bacilli"                    
## [4146,] "Bacteria" "Bacteroidetes"       "Bacteroidia"                
## [4147,] "Bacteria" "Fusobacteria"        "Fusobacteriia"              
## [4148,] "Bacteria" "Firmicutes"          "Negativicutes"              
## [4149,] "Bacteria" "Bacteroidetes"       "Bacteroidia"                
## [4150,] "Bacteria" "Firmicutes"          "Clostridia"                 
## [4151,] "Bacteria" "Firmicutes"          "Clostridia"                 
## [4152,] "Bacteria" "Spirochaetes"        "Spirochaetia"               
## [4153,] "Bacteria" "Proteobacteria"      "Gammaproteobacteria"        
## [4154,] "Bacteria" "Firmicutes"          "Negativicutes"              
## [4155,] "Bacteria" "Bacteroidetes"       "Bacteroidia"                
## [4156,] "Bacteria" "Actinobacteria"      "Actinobacteria"             
## [4157,] "Bacteria" "Firmicutes"          "Negativicutes"              
## [4158,] "Bacteria" "Firmicutes"          "Bacilli"                    
## [4159,] "Bacteria" "Proteobacteria"      "Gammaproteobacteria"        
## [4160,] "Bacteria" "Bacteroidetes"       "Bacteroidia"                
## [4161,] "Bacteria" "Firmicutes"          "Clostridia"                 
## [4162,] "Bacteria" "Firmicutes"          "Negativicutes"              
## [4163,] "Bacteria" "Firmicutes"          "Clostridia"                 
## [4164,] "Bacteria" "Patescibacteria"     "Saccharimonadia"            
## [4165,] "Bacteria" "Fusobacteria"        "Fusobacteriia"              
## [4166,] "Bacteria" "Proteobacteria"      "Gammaproteobacteria"        
## [4167,] "Bacteria" "Firmicutes"          "Bacilli"                    
## [4168,] "Bacteria" "Fusobacteria"        "Fusobacteriia"              
## [4169,] "Bacteria" "Tenericutes"         "Mollicutes"                 
## [4170,] "Bacteria" "Firmicutes"          "Negativicutes"              
## [4171,] "Bacteria" "Bacteroidetes"       "Bacteroidia"                
## [4172,] "Bacteria" "Cyanobacteria"       "Oxyphotobacteria"           
## [4173,] "Bacteria" "Bacteroidetes"       "Bacteroidia"                
## [4174,] "Bacteria" "Firmicutes"          "Clostridia"                 
## [4175,] "Bacteria" "Spirochaetes"        "Spirochaetia"               
## [4176,] "Bacteria" "Proteobacteria"      "Gammaproteobacteria"        
## [4177,] "Bacteria" "Firmicutes"          "Clostridia"                 
## [4178,] "Bacteria" "Proteobacteria"      "Gammaproteobacteria"        
## [4179,] "Bacteria" "Bacteroidetes"       "Bacteroidia"                
## [4180,] "Bacteria" "Patescibacteria"     "Saccharimonadia"            
## [4181,] "Bacteria" "Firmicutes"          "Bacilli"                    
## [4182,] "Bacteria" "Proteobacteria"      "Gammaproteobacteria"        
## [4183,] "Bacteria" "Tenericutes"         "Mollicutes"                 
## [4184,] "Bacteria" "Bacteroidetes"       "Bacteroidia"                
## [4185,] "Bacteria" "Patescibacteria"     "Gracilibacteria"            
## [4186,] "Bacteria" "Firmicutes"          "Clostridia"                 
## [4187,] "Bacteria" "Firmicutes"          NA                           
## [4188,] "Bacteria" "Firmicutes"          "Negativicutes"              
## [4189,] "Bacteria" NA                    NA                           
## [4190,] "Bacteria" "Spirochaetes"        "Spirochaetia"               
## [4191,] "Bacteria" "Proteobacteria"      "Gammaproteobacteria"        
## [4192,] "Bacteria" "Proteobacteria"      "Gammaproteobacteria"        
## [4193,] "Bacteria" "Firmicutes"          "Bacilli"                    
## [4194,] "Bacteria" "Epsilonbacteraeota"  "Campylobacteria"            
## [4195,] "Bacteria" "Bacteroidetes"       "Bacteroidia"                
## [4196,] "Bacteria" "Firmicutes"          "Clostridia"                 
## [4197,] "Bacteria" "Firmicutes"          "Clostridia"                 
## [4198,] "Bacteria" "Firmicutes"          "Negativicutes"              
## [4199,] "Bacteria" "Actinobacteria"      "Actinobacteria"             
## [4200,] "Bacteria" "Proteobacteria"      "Gammaproteobacteria"        
## [4201,] "Bacteria" "Firmicutes"          "Bacilli"                    
## [4202,] "Bacteria" "Fusobacteria"        "Fusobacteriia"              
## [4203,] "Bacteria" "Firmicutes"          "Clostridia"                 
## [4204,] "Bacteria" "Bacteroidetes"       "Bacteroidia"                
## [4205,] "Bacteria" "Firmicutes"          "Negativicutes"              
## [4206,] "Bacteria" "Bacteroidetes"       "Bacteroidia"                
## [4207,] "Bacteria" "Actinobacteria"      "Actinobacteria"             
## [4208,] "Bacteria" "Firmicutes"          "Clostridia"                 
## [4209,] "Bacteria" "Fusobacteria"        "Fusobacteriia"              
## [4210,] "Bacteria" "Spirochaetes"        "Spirochaetia"               
## [4211,] "Bacteria" "Patescibacteria"     "Saccharimonadia"            
## [4212,] "Bacteria" "Firmicutes"          NA                           
## [4213,] "Bacteria" "Epsilonbacteraeota"  "Campylobacteria"            
## [4214,] "Bacteria" "Bacteroidetes"       "Bacteroidia"                
## [4215,] "Bacteria" "Bacteroidetes"       "Bacteroidia"                
## [4216,] "Bacteria" "Firmicutes"          NA                           
## [4217,] "Bacteria" "Patescibacteria"     "Saccharimonadia"            
## [4218,] "Bacteria" "Bacteroidetes"       "Bacteroidia"                
## [4219,] "Bacteria" "Firmicutes"          "Negativicutes"              
## [4220,] "Bacteria" "Epsilonbacteraeota"  "Campylobacteria"            
## [4221,] "Bacteria" "Proteobacteria"      "Gammaproteobacteria"        
## [4222,] "Bacteria" "Fusobacteria"        "Fusobacteriia"              
## [4223,] "Bacteria" "Bacteroidetes"       "Bacteroidia"                
## [4224,] "Bacteria" "Bacteroidetes"       "Bacteroidia"                
## [4225,] "Bacteria" "Firmicutes"          "Clostridia"                 
## [4226,] "Bacteria" "Fusobacteria"        "Fusobacteriia"              
## [4227,] "Bacteria" "Bacteroidetes"       "Bacteroidia"                
## [4228,] "Bacteria" "Firmicutes"          "Negativicutes"              
## [4229,] "Bacteria" "Proteobacteria"      "Gammaproteobacteria"        
## [4230,] "Bacteria" "Firmicutes"          "Bacilli"                    
## [4231,] "Bacteria" "Bacteroidetes"       "Bacteroidia"                
## [4232,] "Bacteria" "Proteobacteria"      "Gammaproteobacteria"        
## [4233,] "Bacteria" "Patescibacteria"     "Gracilibacteria"            
## [4234,] "Bacteria" "Bacteroidetes"       "Bacteroidia"                
## [4235,] "Bacteria" "Firmicutes"          "Bacilli"                    
## [4236,] "Bacteria" "Patescibacteria"     "Saccharimonadia"            
## [4237,] "Bacteria" "Firmicutes"          "Bacilli"                    
## [4238,] "Bacteria" "Spirochaetes"        "Spirochaetia"               
## [4239,] "Bacteria" "Actinobacteria"      "Actinobacteria"             
## [4240,] "Bacteria" "Proteobacteria"      "Gammaproteobacteria"        
## [4241,] "Bacteria" "Epsilonbacteraeota"  "Campylobacteria"            
## [4242,] "Bacteria" "Cyanobacteria"       "Oxyphotobacteria"           
## [4243,] "Bacteria" "Patescibacteria"     "Saccharimonadia"            
## [4244,] "Bacteria" "Firmicutes"          "Bacilli"                    
## [4245,] "Bacteria" "Bacteroidetes"       "Bacteroidia"                
## [4246,] "Bacteria" "Actinobacteria"      "Actinobacteria"             
## [4247,] "Bacteria" "Spirochaetes"        "Spirochaetia"               
## [4248,] "Bacteria" "Firmicutes"          "Bacilli"                    
## [4249,] "Bacteria" "Bacteroidetes"       "Bacteroidia"                
## [4250,] "Bacteria" "Spirochaetes"        "Spirochaetia"               
## [4251,] "Bacteria" "Firmicutes"          "Clostridia"                 
## [4252,] "Bacteria" "Firmicutes"          NA                           
## [4253,] "Bacteria" "Proteobacteria"      "Deltaproteobacteria"        
## [4254,] "Bacteria" "Tenericutes"         "Mollicutes"                 
## [4255,] "Bacteria" "Firmicutes"          "Bacilli"                    
## [4256,] "Bacteria" "Bacteroidetes"       "Bacteroidia"                
## [4257,] "Bacteria" "Actinobacteria"      "Actinobacteria"             
## [4258,] "Bacteria" "Epsilonbacteraeota"  "Campylobacteria"            
## [4259,] "Bacteria" "Bacteroidetes"       "Bacteroidia"                
## [4260,] "Bacteria" "Bacteroidetes"       "Bacteroidia"                
## [4261,] "Bacteria" "Patescibacteria"     "Saccharimonadia"            
## [4262,] "Bacteria" "Firmicutes"          "Bacilli"                    
## [4263,] "Bacteria" NA                    NA                           
## [4264,] "Bacteria" "Bacteroidetes"       "Bacteroidia"                
## [4265,] "Bacteria" "Bacteroidetes"       "Bacteroidia"                
## [4266,] "Bacteria" "Patescibacteria"     "Saccharimonadia"            
## [4267,] "Bacteria" "Actinobacteria"      "Actinobacteria"             
## [4268,] "Bacteria" "Actinobacteria"      "Coriobacteriia"             
## [4269,] "Bacteria" "Bacteroidetes"       "Bacteroidia"                
## [4270,] "Bacteria" "Firmicutes"          "Bacilli"                    
## [4271,] "Bacteria" "Firmicutes"          "Clostridia"                 
## [4272,] "Bacteria" "Firmicutes"          "Clostridia"                 
## [4273,] "Bacteria" "Actinobacteria"      "Actinobacteria"             
## [4274,] "Bacteria" "Proteobacteria"      "Gammaproteobacteria"        
## [4275,] "Bacteria" "Bacteroidetes"       "Bacteroidia"                
## [4276,] "Bacteria" "Firmicutes"          "Clostridia"                 
## [4277,] "Bacteria" "Firmicutes"          "Clostridia"                 
## [4278,] "Bacteria" "Firmicutes"          "Bacilli"                    
## [4279,] "Bacteria" "Firmicutes"          "Bacilli"                    
## [4280,] "Bacteria" "Bacteroidetes"       "Bacteroidia"                
## [4281,] "Bacteria" "Proteobacteria"      "Gammaproteobacteria"        
## [4282,] "Bacteria" "Fusobacteria"        "Fusobacteriia"              
## [4283,] "Bacteria" "Fusobacteria"        "Fusobacteriia"              
## [4284,] "Bacteria" "Proteobacteria"      "Gammaproteobacteria"        
## [4285,] "Bacteria" "Firmicutes"          "Erysipelotrichia"           
## [4286,] "Bacteria" "Firmicutes"          "Clostridia"                 
## [4287,] "Bacteria" "Bacteroidetes"       "Bacteroidia"                
## [4288,] "Bacteria" "Firmicutes"          "Bacilli"                    
## [4289,] "Bacteria" "Firmicutes"          NA                           
## [4290,] "Bacteria" "Firmicutes"          "Clostridia"                 
## [4291,] "Bacteria" "Firmicutes"          "Negativicutes"              
## [4292,] "Bacteria" "Bacteroidetes"       "Bacteroidia"                
## [4293,] "Bacteria" "Proteobacteria"      "Gammaproteobacteria"        
## [4294,] "Bacteria" "Bacteroidetes"       "Bacteroidia"                
## [4295,] "Bacteria" "Actinobacteria"      "Coriobacteriia"             
## [4296,] "Bacteria" "Bacteroidetes"       "Bacteroidia"                
## [4297,] "Bacteria" "Actinobacteria"      "Actinobacteria"             
## [4298,] "Bacteria" NA                    NA                           
## [4299,] "Bacteria" "Proteobacteria"      "Gammaproteobacteria"        
## [4300,] "Bacteria" "Firmicutes"          "Bacilli"                    
## [4301,] "Bacteria" "Firmicutes"          "Bacilli"                    
## [4302,] "Bacteria" "Spirochaetes"        "Spirochaetia"               
## [4303,] "Bacteria" "Bacteroidetes"       "Bacteroidia"                
## [4304,] "Bacteria" "Firmicutes"          "Negativicutes"              
## [4305,] "Bacteria" "Proteobacteria"      "Alphaproteobacteria"        
## [4306,] "Bacteria" "Firmicutes"          "Clostridia"                 
## [4307,] "Bacteria" "Spirochaetes"        "Spirochaetia"               
## [4308,] "Bacteria" "Firmicutes"          "Bacilli"                    
## [4309,] "Bacteria" "Proteobacteria"      "Gammaproteobacteria"        
## [4310,] "Bacteria" "Firmicutes"          "Negativicutes"              
## [4311,] "Bacteria" "Proteobacteria"      "Gammaproteobacteria"        
## [4312,] "Bacteria" "Firmicutes"          "Clostridia"                 
## [4313,] "Bacteria" "Patescibacteria"     "Saccharimonadia"            
## [4314,] "Bacteria" "Proteobacteria"      "Gammaproteobacteria"        
## [4315,] "Bacteria" "Bacteroidetes"       "Bacteroidia"                
## [4316,] "Bacteria" "Firmicutes"          "Clostridia"                 
## [4317,] "Bacteria" "Firmicutes"          NA                           
## [4318,] "Bacteria" "Actinobacteria"      "Actinobacteria"             
## [4319,] "Bacteria" "Spirochaetes"        "Spirochaetia"               
## [4320,] "Bacteria" "Firmicutes"          "Clostridia"                 
## [4321,] "Bacteria" "Firmicutes"          "Clostridia"                 
## [4322,] "Bacteria" "Proteobacteria"      "Gammaproteobacteria"        
## [4323,] "Bacteria" "Firmicutes"          "Clostridia"                 
## [4324,] "Bacteria" "Epsilonbacteraeota"  "Campylobacteria"            
## [4325,] "Bacteria" "Patescibacteria"     "Saccharimonadia"            
## [4326,] "Bacteria" "Proteobacteria"      "Gammaproteobacteria"        
## [4327,] "Bacteria" "Bacteroidetes"       "Bacteroidia"                
## [4328,] "Bacteria" "Firmicutes"          "Erysipelotrichia"           
## [4329,] "Bacteria" "Actinobacteria"      "Actinobacteria"             
## [4330,] "Bacteria" "Bacteroidetes"       "Bacteroidia"                
## [4331,] "Bacteria" "Firmicutes"          "Negativicutes"              
## [4332,] "Bacteria" "Proteobacteria"      "Gammaproteobacteria"        
## [4333,] "Bacteria" "Proteobacteria"      "Gammaproteobacteria"        
## [4334,] "Bacteria" "Bacteroidetes"       "Bacteroidia"                
## [4335,] "Bacteria" "Firmicutes"          "Bacilli"                    
## [4336,] "Bacteria" "Epsilonbacteraeota"  "Campylobacteria"            
## [4337,] "Bacteria" "Spirochaetes"        "Spirochaetia"               
## [4338,] "Bacteria" "Firmicutes"          "Clostridia"                 
## [4339,] "Bacteria" "Proteobacteria"      "Alphaproteobacteria"        
## [4340,] "Bacteria" "Proteobacteria"      "Gammaproteobacteria"        
## [4341,] "Bacteria" "Proteobacteria"      "Gammaproteobacteria"        
## [4342,] "Bacteria" "Firmicutes"          "Bacilli"                    
## [4343,] "Bacteria" "Bacteroidetes"       "Bacteroidia"                
## [4344,] "Bacteria" "Fusobacteria"        "Fusobacteriia"              
## [4345,] "Bacteria" "Bacteroidetes"       "Bacteroidia"                
## [4346,] "Bacteria" "Bacteroidetes"       "Bacteroidia"                
## [4347,] "Bacteria" "Firmicutes"          "Clostridia"                 
## [4348,] "Bacteria" "Bacteroidetes"       "Bacteroidia"                
## [4349,] "Bacteria" "Proteobacteria"      "Gammaproteobacteria"        
## [4350,] "Bacteria" "Bacteroidetes"       "Bacteroidia"                
## [4351,] "Bacteria" "Proteobacteria"      "Alphaproteobacteria"        
## [4352,] "Bacteria" "Firmicutes"          "Clostridia"                 
## [4353,] "Bacteria" "Bacteroidetes"       "Bacteroidia"                
## [4354,] "Bacteria" "Firmicutes"          "Bacilli"                    
## [4355,] "Bacteria" "Bacteroidetes"       "Bacteroidia"                
## [4356,] "Bacteria" "Firmicutes"          "Bacilli"                    
## [4357,] "Bacteria" "Proteobacteria"      "Gammaproteobacteria"        
## [4358,] "Bacteria" "Fusobacteria"        "Fusobacteriia"              
## [4359,] "Bacteria" "Proteobacteria"      "Gammaproteobacteria"        
## [4360,] "Bacteria" "Firmicutes"          "Clostridia"                 
## [4361,] "Bacteria" "Bacteroidetes"       "Bacteroidia"                
## [4362,] "Bacteria" "Fusobacteria"        "Fusobacteriia"              
## [4363,] "Bacteria" "Bacteroidetes"       "Bacteroidia"                
## [4364,] "Bacteria" "Firmicutes"          "Clostridia"                 
## [4365,] "Bacteria" "Proteobacteria"      "Gammaproteobacteria"        
## [4366,] "Bacteria" "Proteobacteria"      "Gammaproteobacteria"        
## [4367,] "Bacteria" "Firmicutes"          NA                           
## [4368,] "Bacteria" "Epsilonbacteraeota"  "Campylobacteria"            
## [4369,] "Bacteria" "Firmicutes"          "Clostridia"                 
## [4370,] "Bacteria" "Fusobacteria"        "Fusobacteriia"              
## [4371,] "Bacteria" "Proteobacteria"      "Gammaproteobacteria"        
## [4372,] "Bacteria" "Proteobacteria"      "Gammaproteobacteria"        
## [4373,] "Bacteria" "Firmicutes"          "Negativicutes"              
## [4374,] "Bacteria" "Fusobacteria"        "Fusobacteriia"              
## [4375,] "Bacteria" "Firmicutes"          "Negativicutes"              
## [4376,] "Bacteria" "Spirochaetes"        "Spirochaetia"               
## [4377,] "Bacteria" "Bacteroidetes"       "Bacteroidia"                
## [4378,] "Bacteria" "Firmicutes"          "Bacilli"                    
## [4379,] "Bacteria" "Firmicutes"          "Clostridia"                 
## [4380,] "Bacteria" "Proteobacteria"      "Gammaproteobacteria"        
## [4381,] "Bacteria" "Firmicutes"          NA                           
## [4382,] "Bacteria" "Bacteroidetes"       "Bacteroidia"                
## [4383,] "Bacteria" "Firmicutes"          "Bacilli"                    
## [4384,] "Bacteria" "Actinobacteria"      "Coriobacteriia"             
## [4385,] "Bacteria" "Firmicutes"          "Clostridia"                 
## [4386,] "Bacteria" "Bacteroidetes"       "Bacteroidia"                
## [4387,] "Bacteria" "Firmicutes"          "Negativicutes"              
## [4388,] "Bacteria" "Firmicutes"          "Clostridia"                 
## [4389,] "Bacteria" "Actinobacteria"      "Actinobacteria"             
## [4390,] "Bacteria" "Firmicutes"          "Negativicutes"              
## [4391,] "Bacteria" "Epsilonbacteraeota"  "Campylobacteria"            
## [4392,] "Bacteria" "Epsilonbacteraeota"  "Campylobacteria"            
## [4393,] "Bacteria" "Fusobacteria"        "Fusobacteriia"              
## [4394,] "Bacteria" "Firmicutes"          "Negativicutes"              
## [4395,] "Bacteria" "Firmicutes"          "Erysipelotrichia"           
## [4396,] "Bacteria" "Firmicutes"          "Negativicutes"              
## [4397,] "Bacteria" "Firmicutes"          "Negativicutes"              
## [4398,] "Bacteria" "Firmicutes"          "Negativicutes"              
## [4399,] "Bacteria" "Firmicutes"          "Bacilli"                    
## [4400,] "Bacteria" "Patescibacteria"     "Saccharimonadia"            
## [4401,] "Bacteria" "Firmicutes"          "Negativicutes"              
## [4402,] "Bacteria" "Fusobacteria"        "Fusobacteriia"              
## [4403,] "Bacteria" "Proteobacteria"      "Gammaproteobacteria"        
## [4404,] "Bacteria" "Patescibacteria"     "Saccharimonadia"            
## [4405,] "Bacteria" "Actinobacteria"      "Coriobacteriia"             
## [4406,] "Bacteria" "Bacteroidetes"       "Bacteroidia"                
## [4407,] "Bacteria" "Bacteroidetes"       "Bacteroidia"                
## [4408,] "Bacteria" "Proteobacteria"      "Alphaproteobacteria"        
## [4409,] "Bacteria" "Firmicutes"          "Bacilli"                    
## [4410,] "Bacteria" "Bacteroidetes"       "Bacteroidia"                
## [4411,] "Bacteria" "Firmicutes"          "Clostridia"                 
## [4412,] "Bacteria" "Firmicutes"          "Bacilli"                    
## [4413,] "Bacteria" "Actinobacteria"      "Coriobacteriia"             
## [4414,] "Bacteria" "Spirochaetes"        "Spirochaetia"               
## [4415,] "Bacteria" "Bacteroidetes"       "Bacteroidia"                
## [4416,] "Bacteria" "Epsilonbacteraeota"  "Campylobacteria"            
## [4417,] "Bacteria" "Synergistetes"       "Synergistia"                
## [4418,] "Bacteria" "Actinobacteria"      "Actinobacteria"             
## [4419,] "Bacteria" "Actinobacteria"      "Actinobacteria"             
## [4420,] "Bacteria" "Actinobacteria"      "Coriobacteriia"             
## [4421,] "Bacteria" "Firmicutes"          "Clostridia"                 
## [4422,] "Bacteria" "Tenericutes"         "Mollicutes"                 
## [4423,] "Bacteria" "Proteobacteria"      "Gammaproteobacteria"        
## [4424,] "Bacteria" "Firmicutes"          "Bacilli"                    
## [4425,] "Bacteria" "Epsilonbacteraeota"  "Campylobacteria"            
## [4426,] "Bacteria" "Actinobacteria"      "Actinobacteria"             
## [4427,] "Bacteria" "Firmicutes"          "Clostridia"                 
## [4428,] "Bacteria" "Firmicutes"          "Negativicutes"              
## [4429,] "Bacteria" "Bacteroidetes"       "Bacteroidia"                
## [4430,] "Bacteria" "Actinobacteria"      "Coriobacteriia"             
## [4431,] "Bacteria" "Bacteroidetes"       "Bacteroidia"                
## [4432,] "Bacteria" "Bacteroidetes"       "Bacteroidia"                
## [4433,] "Bacteria" "Patescibacteria"     "Saccharimonadia"            
## [4434,] "Bacteria" "Firmicutes"          "Clostridia"                 
## [4435,] "Bacteria" NA                    NA                           
## [4436,] "Bacteria" "Actinobacteria"      "Actinobacteria"             
## [4437,] "Bacteria" "Firmicutes"          NA                           
## [4438,] "Bacteria" "Patescibacteria"     "Saccharimonadia"            
## [4439,] "Bacteria" "Proteobacteria"      "Gammaproteobacteria"        
## [4440,] "Bacteria" "Firmicutes"          "Negativicutes"              
## [4441,] "Bacteria" "Firmicutes"          "Negativicutes"              
## [4442,] "Bacteria" "Proteobacteria"      "Gammaproteobacteria"        
## [4443,] "Bacteria" "Firmicutes"          "Erysipelotrichia"           
## [4444,] "Bacteria" "Proteobacteria"      "Gammaproteobacteria"        
## [4445,] "Bacteria" "Bacteroidetes"       "Bacteroidia"                
## [4446,] "Bacteria" "Spirochaetes"        "Spirochaetia"               
## [4447,] "Bacteria" "Bacteroidetes"       "Bacteroidia"                
## [4448,] "Bacteria" "Firmicutes"          "Bacilli"                    
## [4449,] "Bacteria" "Bacteroidetes"       "Bacteroidia"                
## [4450,] "Bacteria" "Bacteroidetes"       "Bacteroidia"                
## [4451,] "Bacteria" NA                    NA                           
## [4452,] "Bacteria" "Bacteroidetes"       "Bacteroidia"                
## [4453,] "Bacteria" "Actinobacteria"      "Actinobacteria"             
## [4454,] "Bacteria" "Proteobacteria"      "Gammaproteobacteria"        
## [4455,] "Bacteria" "Bacteroidetes"       "Bacteroidia"                
## [4456,] "Bacteria" "Fusobacteria"        "Fusobacteriia"              
## [4457,] "Bacteria" "Proteobacteria"      "Gammaproteobacteria"        
## [4458,] "Bacteria" "Firmicutes"          "Erysipelotrichia"           
## [4459,] "Bacteria" "Bacteroidetes"       "Bacteroidia"                
## [4460,] "Bacteria" "Proteobacteria"      "Gammaproteobacteria"        
## [4461,] "Bacteria" "Epsilonbacteraeota"  "Campylobacteria"            
## [4462,] "Bacteria" "Fusobacteria"        "Fusobacteriia"              
## [4463,] "Bacteria" "Firmicutes"          "Bacilli"                    
## [4464,] "Bacteria" "Bacteroidetes"       "Bacteroidia"                
## [4465,] "Bacteria" "Proteobacteria"      "Gammaproteobacteria"        
## [4466,] "Bacteria" "Actinobacteria"      "Coriobacteriia"             
## [4467,] "Bacteria" "Proteobacteria"      "Gammaproteobacteria"        
## [4468,] "Bacteria" "Firmicutes"          "Bacilli"                    
## [4469,] "Bacteria" NA                    NA                           
## [4470,] "Bacteria" "Bacteroidetes"       "Bacteroidia"                
## [4471,] "Bacteria" "Firmicutes"          "Clostridia"                 
## [4472,] "Bacteria" "Firmicutes"          "Clostridia"                 
## [4473,] "Bacteria" "Bacteroidetes"       "Bacteroidia"                
## [4474,] "Bacteria" "Bacteroidetes"       "Bacteroidia"                
## [4475,] "Bacteria" "Firmicutes"          "Clostridia"                 
## [4476,] "Bacteria" "Bacteroidetes"       "Bacteroidia"                
## [4477,] "Bacteria" "Firmicutes"          "Negativicutes"              
## [4478,] "Bacteria" "Bacteroidetes"       "Bacteroidia"                
## [4479,] "Bacteria" "Bacteroidetes"       "Bacteroidia"                
## [4480,] "Bacteria" "Firmicutes"          "Bacilli"                    
## [4481,] "Bacteria" "Proteobacteria"      "Gammaproteobacteria"        
## [4482,] "Bacteria" "Firmicutes"          "Clostridia"                 
## [4483,] "Bacteria" "Firmicutes"          "Bacilli"                    
## [4484,] "Bacteria" "Bacteroidetes"       "Bacteroidia"                
## [4485,] "Bacteria" "Firmicutes"          "Negativicutes"              
## [4486,] "Bacteria" "Firmicutes"          "Erysipelotrichia"           
## [4487,] "Bacteria" "Patescibacteria"     "Saccharimonadia"            
## [4488,] "Bacteria" "Patescibacteria"     "Saccharimonadia"            
## [4489,] "Bacteria" "Spirochaetes"        "Spirochaetia"               
## [4490,] "Bacteria" "Firmicutes"          "Clostridia"                 
## [4491,] "Bacteria" "Firmicutes"          "Clostridia"                 
## [4492,] "Bacteria" "Proteobacteria"      "Gammaproteobacteria"        
## [4493,] "Bacteria" "Firmicutes"          "Clostridia"                 
## [4494,] "Bacteria" "Proteobacteria"      "Gammaproteobacteria"        
## [4495,] "Bacteria" "Bacteroidetes"       "Bacteroidia"                
## [4496,] "Bacteria" "Proteobacteria"      "Gammaproteobacteria"        
## [4497,] "Bacteria" "Proteobacteria"      "Alphaproteobacteria"        
## [4498,] "Bacteria" "Firmicutes"          "Bacilli"                    
## [4499,] "Bacteria" "Bacteroidetes"       "Bacteroidia"                
## [4500,] "Bacteria" "Firmicutes"          "Negativicutes"              
## [4501,] "Bacteria" "Acidobacteria"       "Blastocatellia_(Subgroup_4)"
## [4502,] "Bacteria" "Firmicutes"          "Bacilli"                    
## [4503,] "Bacteria" "Proteobacteria"      "Gammaproteobacteria"        
## [4504,] "Bacteria" "Firmicutes"          "Clostridia"                 
## [4505,] "Bacteria" "Fusobacteria"        "Fusobacteriia"              
## [4506,] "Bacteria" "Bacteroidetes"       "Bacteroidia"                
## [4507,] "Bacteria" "Bacteroidetes"       "Bacteroidia"                
## [4508,] "Bacteria" "Firmicutes"          "Clostridia"                 
## [4509,] "Bacteria" "Patescibacteria"     "Saccharimonadia"            
## [4510,] "Bacteria" "Proteobacteria"      "Gammaproteobacteria"        
## [4511,] "Bacteria" "Bacteroidetes"       "Bacteroidia"                
## [4512,] "Bacteria" "Bacteroidetes"       "Bacteroidia"                
## [4513,] "Bacteria" "Actinobacteria"      "Actinobacteria"             
## [4514,] "Bacteria" "Firmicutes"          "Bacilli"                    
## [4515,] "Bacteria" "Fusobacteria"        "Fusobacteriia"              
## [4516,] "Bacteria" "Patescibacteria"     "Saccharimonadia"            
## [4517,] "Bacteria" "Fusobacteria"        "Fusobacteriia"              
## [4518,] "Bacteria" "Firmicutes"          "Negativicutes"              
## [4519,] "Bacteria" "Bacteroidetes"       "Bacteroidia"                
## [4520,] "Bacteria" "Firmicutes"          "Bacilli"                    
## [4521,] "Bacteria" "Proteobacteria"      "Gammaproteobacteria"        
## [4522,] "Bacteria" "Bacteroidetes"       "Bacteroidia"                
## [4523,] "Bacteria" "Bacteroidetes"       "Bacteroidia"                
## [4524,] "Bacteria" "Proteobacteria"      "Gammaproteobacteria"        
## [4525,] "Bacteria" "Firmicutes"          "Clostridia"                 
## [4526,] "Bacteria" "Bacteroidetes"       "Bacteroidia"                
## [4527,] "Bacteria" "Patescibacteria"     "Saccharimonadia"            
## [4528,] "Bacteria" "Bacteroidetes"       "Bacteroidia"                
## [4529,] "Bacteria" "Firmicutes"          "Negativicutes"              
## [4530,] "Bacteria" "Bacteroidetes"       "Bacteroidia"                
## [4531,] "Bacteria" "Bacteroidetes"       "Bacteroidia"                
## [4532,] "Bacteria" "Bacteroidetes"       "Bacteroidia"                
## [4533,] "Bacteria" NA                    NA                           
## [4534,] "Bacteria" "Proteobacteria"      "Gammaproteobacteria"        
## [4535,] "Bacteria" "Bacteroidetes"       "Bacteroidia"                
## [4536,] "Bacteria" "Actinobacteria"      "Actinobacteria"             
## [4537,] "Bacteria" "Actinobacteria"      "Coriobacteriia"             
## [4538,] "Bacteria" "Firmicutes"          "Bacilli"                    
## [4539,] "Bacteria" "Bacteroidetes"       "Bacteroidia"                
## [4540,] "Bacteria" "Bacteroidetes"       "Bacteroidia"                
## [4541,] "Bacteria" "Firmicutes"          "Clostridia"                 
## [4542,] "Bacteria" "Firmicutes"          "Clostridia"                 
## [4543,] "Bacteria" NA                    NA                           
## [4544,] "Bacteria" "Bacteroidetes"       "Bacteroidia"                
## [4545,] "Bacteria" "Epsilonbacteraeota"  "Campylobacteria"            
## [4546,] "Bacteria" "Spirochaetes"        "Spirochaetia"               
## [4547,] "Bacteria" "Proteobacteria"      "Gammaproteobacteria"        
## [4548,] "Bacteria" "Bacteroidetes"       "Bacteroidia"                
## [4549,] "Bacteria" "Firmicutes"          "Negativicutes"              
## [4550,] "Bacteria" "Firmicutes"          "Bacilli"                    
## [4551,] "Bacteria" "Fusobacteria"        "Fusobacteriia"              
## [4552,] "Bacteria" "Firmicutes"          "Bacilli"                    
## [4553,] "Bacteria" "Spirochaetes"        "Spirochaetia"               
## [4554,] "Bacteria" "Bacteroidetes"       "Bacteroidia"                
## [4555,] "Bacteria" "Firmicutes"          "Clostridia"                 
##         [,4]                          [,5]                           
##    [1,] "Selenomonadales"             "Veillonellaceae"              
##    [2,] "Lactobacillales"             "Streptococcaceae"             
##    [3,] "Bacteroidales"               "Prevotellaceae"               
##    [4,] "Pasteurellales"              "Pasteurellaceae"              
##    [5,] "Selenomonadales"             "Veillonellaceae"              
##    [6,] "Selenomonadales"             "Veillonellaceae"              
##    [7,] "Selenomonadales"             "Veillonellaceae"              
##    [8,] "Saccharimonadales"           "Saccharimonadaceae"           
##    [9,] "Betaproteobacteriales"       "Neisseriaceae"                
##   [10,] "Selenomonadales"             "Veillonellaceae"              
##   [11,] "Betaproteobacteriales"       "Neisseriaceae"                
##   [12,] "Selenomonadales"             "Veillonellaceae"              
##   [13,] "Fusobacteriales"             "Fusobacteriaceae"             
##   [14,] "Selenomonadales"             "Veillonellaceae"              
##   [15,] "Bacteroidales"               "Prevotellaceae"               
##   [16,] "Campylobacterales"           "Campylobacteraceae"           
##   [17,] "Pasteurellales"              "Pasteurellaceae"              
##   [18,] "Clostridiales"               "Lachnospiraceae"              
##   [19,] "Selenomonadales"             "Veillonellaceae"              
##   [20,] "Bacteroidales"               "Prevotellaceae"               
##   [21,] "Lactobacillales"             "Carnobacteriaceae"            
##   [22,] "Bacteroidales"               "Prevotellaceae"               
##   [23,] "Bacteroidales"               "Prevotellaceae"               
##   [24,] "Selenomonadales"             "Veillonellaceae"              
##   [25,] "Pasteurellales"              "Pasteurellaceae"              
##   [26,] "Bacteroidales"               "Prevotellaceae"               
##   [27,] "Bacteroidales"               "Porphyromonadaceae"           
##   [28,] "Selenomonadales"             "Veillonellaceae"              
##   [29,] "Lactobacillales"             "Streptococcaceae"             
##   [30,] "Lactobacillales"             "Streptococcaceae"             
##   [31,] "Bacteroidales"               "Porphyromonadaceae"           
##   [32,] "Lactobacillales"             "Streptococcaceae"             
##   [33,] "Bacteroidales"               "Prevotellaceae"               
##   [34,] "Bacteroidales"               "Prevotellaceae"               
##   [35,] "Clostridiales"               "Peptostreptococcaceae"        
##   [36,] "Selenomonadales"             "Veillonellaceae"              
##   [37,] "Bacteroidales"               "Porphyromonadaceae"           
##   [38,] "Selenomonadales"             "Veillonellaceae"              
##   [39,] "Clostridiales"               "Lachnospiraceae"              
##   [40,] "Lactobacillales"             "Streptococcaceae"             
##   [41,] "Bacteroidales"               "Prevotellaceae"               
##   [42,] "Lactobacillales"             "Streptococcaceae"             
##   [43,] "Lactobacillales"             "Streptococcaceae"             
##   [44,] "Bacteroidales"               "Prevotellaceae"               
##   [45,] "Betaproteobacteriales"       "Neisseriaceae"                
##   [46,] "Bacteroidales"               "Prevotellaceae"               
##   [47,] "Selenomonadales"             "Veillonellaceae"              
##   [48,] "Selenomonadales"             "Veillonellaceae"              
##   [49,] "Clostridiales"               "Lachnospiraceae"              
##   [50,] "Fusobacteriales"             "Leptotrichiaceae"             
##   [51,] "Lactobacillales"             "Streptococcaceae"             
##   [52,] "Pasteurellales"              "Pasteurellaceae"              
##   [53,] "Lactobacillales"             "Streptococcaceae"             
##   [54,] "Fusobacteriales"             "Leptotrichiaceae"             
##   [55,] "Bacteroidales"               "Prevotellaceae"               
##   [56,] "Bacteroidales"               "Prevotellaceae"               
##   [57,] "Selenomonadales"             "Veillonellaceae"              
##   [58,] "Clostridiales"               "Lachnospiraceae"              
##   [59,] "Coriobacteriales"            "Atopobiaceae"                 
##   [60,] "Pasteurellales"              "Pasteurellaceae"              
##   [61,] "Pasteurellales"              "Pasteurellaceae"              
##   [62,] "Bacteroidales"               "Prevotellaceae"               
##   [63,] "Clostridiales"               "Lachnospiraceae"              
##   [64,] "Selenomonadales"             "Veillonellaceae"              
##   [65,] "Pasteurellales"              "Pasteurellaceae"              
##   [66,] "Selenomonadales"             "Veillonellaceae"              
##   [67,] "Saccharimonadales"           "Saccharimonadaceae"           
##   [68,] "Micrococcales"               "Micrococcaceae"               
##   [69,] "Betaproteobacteriales"       "Neisseriaceae"                
##   [70,] "Clostridiales"               "Family_XI"                    
##   [71,] "Bacteroidales"               "Prevotellaceae"               
##   [72,] "Micrococcales"               "Micrococcaceae"               
##   [73,] "Clostridiales"               "Lachnospiraceae"              
##   [74,] "Selenomonadales"             "Veillonellaceae"              
##   [75,] "Micrococcales"               "Micrococcaceae"               
##   [76,] "Betaproteobacteriales"       "Neisseriaceae"                
##   [77,] "Clostridiales"               "Family_XIII"                  
##   [78,] "Bacteroidales"               "Porphyromonadaceae"           
##   [79,] "Selenomonadales"             "Veillonellaceae"              
##   [80,] "Clostridiales"               "Lachnospiraceae"              
##   [81,] "Pasteurellales"              "Pasteurellaceae"              
##   [82,] "Selenomonadales"             "Veillonellaceae"              
##   [83,] "Bacteroidales"               "Prevotellaceae"               
##   [84,] "Bacteroidales"               "Prevotellaceae"               
##   [85,] "Selenomonadales"             "Veillonellaceae"              
##   [86,] "Bacteroidales"               "Prevotellaceae"               
##   [87,] "Fusobacteriales"             "Leptotrichiaceae"             
##   [88,] "Clostridiales"               "Ruminococcaceae"              
##   [89,] "Bacteroidales"               "Prevotellaceae"               
##   [90,] "Clostridiales"               "Ruminococcaceae"              
##   [91,] "Micrococcales"               "Micrococcaceae"               
##   [92,] "Betaproteobacteriales"       "Neisseriaceae"                
##   [93,] "Micrococcales"               "Micrococcaceae"               
##   [94,] "Bacillales"                  "Family_XI"                    
##   [95,] "Bacteroidales"               "Prevotellaceae"               
##   [96,] "Betaproteobacteriales"       "Neisseriaceae"                
##   [97,] "Fusobacteriales"             "Fusobacteriaceae"             
##   [98,] "Selenomonadales"             "Veillonellaceae"              
##   [99,] "Fusobacteriales"             "Fusobacteriaceae"             
##  [100,] "Bacteroidales"               "Prevotellaceae"               
##  [101,] "Bacteroidales"               "Prevotellaceae"               
##  [102,] "Pasteurellales"              "Pasteurellaceae"              
##  [103,] "Pasteurellales"              "Pasteurellaceae"              
##  [104,] "Bacteroidales"               "Prevotellaceae"               
##  [105,] "Lactobacillales"             "Carnobacteriaceae"            
##  [106,] "Bacillales"                  "Family_XI"                    
##  [107,] "Selenomonadales"             "Veillonellaceae"              
##  [108,] "Selenomonadales"             "Veillonellaceae"              
##  [109,] "Micrococcales"               "Micrococcaceae"               
##  [110,] "Pasteurellales"              "Pasteurellaceae"              
##  [111,] "Campylobacterales"           "Campylobacteraceae"           
##  [112,] "Bacteroidales"               "Prevotellaceae"               
##  [113,] "Betaproteobacteriales"       "Neisseriaceae"                
##  [114,] "Betaproteobacteriales"       "Neisseriaceae"                
##  [115,] "Saccharimonadales"           NA                             
##  [116,] "Bacteroidales"               "Prevotellaceae"               
##  [117,] "Betaproteobacteriales"       "Neisseriaceae"                
##  [118,] "Pasteurellales"              "Pasteurellaceae"              
##  [119,] "Pasteurellales"              "Pasteurellaceae"              
##  [120,] "Pasteurellales"              "Pasteurellaceae"              
##  [121,] "Pasteurellales"              "Pasteurellaceae"              
##  [122,] "Bacteroidales"               "Prevotellaceae"               
##  [123,] "Lactobacillales"             "Streptococcaceae"             
##  [124,] "Bacteroidales"               "Prevotellaceae"               
##  [125,] "Betaproteobacteriales"       "Neisseriaceae"                
##  [126,] "Bacteroidales"               "Prevotellaceae"               
##  [127,] "Selenomonadales"             "Veillonellaceae"              
##  [128,] "Pasteurellales"              "Pasteurellaceae"              
##  [129,] "Pasteurellales"              "Pasteurellaceae"              
##  [130,] "Pasteurellales"              "Pasteurellaceae"              
##  [131,] "Betaproteobacteriales"       "Burkholderiaceae"             
##  [132,] "Bacteroidales"               "Prevotellaceae"               
##  [133,] "Saccharimonadales"           "Saccharimonadaceae"           
##  [134,] "Bacteroidales"               "Prevotellaceae"               
##  [135,] "Selenomonadales"             "Veillonellaceae"              
##  [136,] "Selenomonadales"             "Veillonellaceae"              
##  [137,] "Bacteroidales"               "Prevotellaceae"               
##  [138,] "Flavobacteriales"            "Flavobacteriaceae"            
##  [139,] "Bacteroidales"               "Prevotellaceae"               
##  [140,] "Lactobacillales"             "Streptococcaceae"             
##  [141,] "Selenomonadales"             "Veillonellaceae"              
##  [142,] "Selenomonadales"             "Veillonellaceae"              
##  [143,] "Fusobacteriales"             "Leptotrichiaceae"             
##  [144,] "Selenomonadales"             "Veillonellaceae"              
##  [145,] "Lactobacillales"             "Streptococcaceae"             
##  [146,] "Flavobacteriales"            "Weeksellaceae"                
##  [147,] "Betaproteobacteriales"       "Neisseriaceae"                
##  [148,] "Selenomonadales"             "Veillonellaceae"              
##  [149,] "Selenomonadales"             "Veillonellaceae"              
##  [150,] "Selenomonadales"             "Veillonellaceae"              
##  [151,] "Betaproteobacteriales"       "Neisseriaceae"                
##  [152,] "Bacteroidales"               "Prevotellaceae"               
##  [153,] "Selenomonadales"             "Veillonellaceae"              
##  [154,] "Bacteroidales"               "Prevotellaceae"               
##  [155,] "Clostridiales"               "Lachnospiraceae"              
##  [156,] "Enterobacteriales"           "Enterobacteriaceae"           
##  [157,] "Selenomonadales"             "Veillonellaceae"              
##  [158,] "Bacteroidales"               "Prevotellaceae"               
##  [159,] "Selenomonadales"             "Veillonellaceae"              
##  [160,] "Clostridiales"               "Lachnospiraceae"              
##  [161,] "Lactobacillales"             "Streptococcaceae"             
##  [162,] "Micrococcales"               "Micrococcaceae"               
##  [163,] "Bacteroidales"               "Prevotellaceae"               
##  [164,] "Selenomonadales"             "Veillonellaceae"              
##  [165,] "Campylobacterales"           "Campylobacteraceae"           
##  [166,] "Bacteroidales"               "Prevotellaceae"               
##  [167,] "Bacteroidales"               "Porphyromonadaceae"           
##  [168,] "Lactobacillales"             "Streptococcaceae"             
##  [169,] "Bacteroidales"               "Prevotellaceae"               
##  [170,] "Betaproteobacteriales"       "Neisseriaceae"                
##  [171,] "Lactobacillales"             "Streptococcaceae"             
##  [172,] "Lactobacillales"             "Streptococcaceae"             
##  [173,] "Saccharimonadales"           "Saccharimonadaceae"           
##  [174,] "Bacteroidales"               "Prevotellaceae"               
##  [175,] "Bacteroidales"               "Prevotellaceae"               
##  [176,] "Clostridiales"               "Lachnospiraceae"              
##  [177,] "Selenomonadales"             "Veillonellaceae"              
##  [178,] "Selenomonadales"             "Veillonellaceae"              
##  [179,] "Bacteroidales"               "Prevotellaceae"               
##  [180,] "Betaproteobacteriales"       "Neisseriaceae"                
##  [181,] "Absconditabacteriales_(SR1)" NA                             
##  [182,] "Selenomonadales"             "Veillonellaceae"              
##  [183,] "Betaproteobacteriales"       "Neisseriaceae"                
##  [184,] "Selenomonadales"             "Veillonellaceae"              
##  [185,] "Pasteurellales"              "Pasteurellaceae"              
##  [186,] "Selenomonadales"             "Veillonellaceae"              
##  [187,] "Selenomonadales"             "Veillonellaceae"              
##  [188,] "Bacteroidales"               "Prevotellaceae"               
##  [189,] "Bacteroidales"               "Prevotellaceae"               
##  [190,] "Pasteurellales"              "Pasteurellaceae"              
##  [191,] "Bacteroidales"               "Prevotellaceae"               
##  [192,] "Bacteroidales"               "Porphyromonadaceae"           
##  [193,] "Betaproteobacteriales"       "Neisseriaceae"                
##  [194,] "Bacteroidales"               "Prevotellaceae"               
##  [195,] "Selenomonadales"             "Veillonellaceae"              
##  [196,] "Selenomonadales"             "Veillonellaceae"              
##  [197,] "Fusobacteriales"             "Fusobacteriaceae"             
##  [198,] "Selenomonadales"             "Veillonellaceae"              
##  [199,] "Bacteroidales"               "Prevotellaceae"               
##  [200,] "Selenomonadales"             "Veillonellaceae"              
##  [201,] "Bacteroidales"               "Prevotellaceae"               
##  [202,] "Clostridiales"               "Lachnospiraceae"              
##  [203,] "Campylobacterales"           "Campylobacteraceae"           
##  [204,] "Bacteroidales"               "Prevotellaceae"               
##  [205,] "Selenomonadales"             "Veillonellaceae"              
##  [206,] "Selenomonadales"             "Veillonellaceae"              
##  [207,] "Flavobacteriales"            "Flavobacteriaceae"            
##  [208,] "Betaproteobacteriales"       "Neisseriaceae"                
##  [209,] "Betaproteobacteriales"       "Neisseriaceae"                
##  [210,] "Lactobacillales"             "Streptococcaceae"             
##  [211,] "Clostridiales"               "Family_XIII"                  
##  [212,] "Selenomonadales"             "Veillonellaceae"              
##  [213,] "Pasteurellales"              "Pasteurellaceae"              
##  [214,] "Betaproteobacteriales"       "Neisseriaceae"                
##  [215,] "Selenomonadales"             "Veillonellaceae"              
##  [216,] "Bacteroidales"               "Prevotellaceae"               
##  [217,] "Lactobacillales"             "Streptococcaceae"             
##  [218,] "Betaproteobacteriales"       "Neisseriaceae"                
##  [219,] "Selenomonadales"             "Veillonellaceae"              
##  [220,] "Lactobacillales"             "Streptococcaceae"             
##  [221,] "Clostridiales"               "Lachnospiraceae"              
##  [222,] "Lactobacillales"             "Streptococcaceae"             
##  [223,] "Clostridiales"               "Lachnospiraceae"              
##  [224,] "Bacteroidales"               "Prevotellaceae"               
##  [225,] "Fusobacteriales"             "Leptotrichiaceae"             
##  [226,] "Selenomonadales"             "Veillonellaceae"              
##  [227,] "Bacteroidales"               "Prevotellaceae"               
##  [228,] "Lactobacillales"             "Streptococcaceae"             
##  [229,] "Flavobacteriales"            "Flavobacteriaceae"            
##  [230,] "Flavobacteriales"            "Flavobacteriaceae"            
##  [231,] "Fusobacteriales"             "Leptotrichiaceae"             
##  [232,] "Fusobacteriales"             "Fusobacteriaceae"             
##  [233,] "Pasteurellales"              "Pasteurellaceae"              
##  [234,] "Fusobacteriales"             "Fusobacteriaceae"             
##  [235,] "Selenomonadales"             "Veillonellaceae"              
##  [236,] "Saccharimonadales"           NA                             
##  [237,] "Bacteroidales"               "Prevotellaceae"               
##  [238,] "Selenomonadales"             "Veillonellaceae"              
##  [239,] "Bacteroidales"               "Prevotellaceae"               
##  [240,] "Selenomonadales"             "Veillonellaceae"              
##  [241,] "Fusobacteriales"             "Fusobacteriaceae"             
##  [242,] "Bacteroidales"               "Porphyromonadaceae"           
##  [243,] "Selenomonadales"             "Veillonellaceae"              
##  [244,] "Chloroplast"                 NA                             
##  [245,] "Bacteroidales"               "Prevotellaceae"               
##  [246,] "Pasteurellales"              "Pasteurellaceae"              
##  [247,] "Betaproteobacteriales"       "Neisseriaceae"                
##  [248,] "Chloroplast"                 NA                             
##  [249,] "Absconditabacteriales_(SR1)" NA                             
##  [250,] "Bacteroidales"               "Prevotellaceae"               
##  [251,] "Bacteroidales"               "Prevotellaceae"               
##  [252,] "Bacteroidales"               "Prevotellaceae"               
##  [253,] "Clostridiales"               "Lachnospiraceae"              
##  [254,] "Pasteurellales"              "Pasteurellaceae"              
##  [255,] "Enterobacteriales"           "Enterobacteriaceae"           
##  [256,] "Selenomonadales"             "Veillonellaceae"              
##  [257,] "Selenomonadales"             "Veillonellaceae"              
##  [258,] "Pasteurellales"              "Pasteurellaceae"              
##  [259,] "Saccharimonadales"           "Saccharimonadaceae"           
##  [260,] "Actinomycetales"             "Actinomycetaceae"             
##  [261,] "Clostridiales"               "Lachnospiraceae"              
##  [262,] "Bacteroidales"               "Prevotellaceae"               
##  [263,] "Bacteroidales"               "Prevotellaceae"               
##  [264,] "Betaproteobacteriales"       "Neisseriaceae"                
##  [265,] "Saccharimonadales"           NA                             
##  [266,] "Betaproteobacteriales"       "Neisseriaceae"                
##  [267,] "Erysipelotrichales"          "Erysipelotrichaceae"          
##  [268,] "Bacteroidales"               "Prevotellaceae"               
##  [269,] "Clostridiales"               "Family_XI"                    
##  [270,] "Betaproteobacteriales"       "Neisseriaceae"                
##  [271,] "Selenomonadales"             "Veillonellaceae"              
##  [272,] "Betaproteobacteriales"       "Neisseriaceae"                
##  [273,] "Betaproteobacteriales"       "Neisseriaceae"                
##  [274,] "Selenomonadales"             "Veillonellaceae"              
##  [275,] "Lactobacillales"             "Streptococcaceae"             
##  [276,] "Selenomonadales"             "Veillonellaceae"              
##  [277,] "Bacteroidales"               "Prevotellaceae"               
##  [278,] "Flavobacteriales"            "Flavobacteriaceae"            
##  [279,] "Pasteurellales"              "Pasteurellaceae"              
##  [280,] "Bacteroidales"               "Prevotellaceae"               
##  [281,] "Fusobacteriales"             "Fusobacteriaceae"             
##  [282,] NA                            NA                             
##  [283,] "Fusobacteriales"             "Fusobacteriaceae"             
##  [284,] "Bacteroidales"               "Porphyromonadaceae"           
##  [285,] "Bacteroidales"               "Prevotellaceae"               
##  [286,] "Absconditabacteriales_(SR1)" NA                             
##  [287,] "Bacteroidales"               "Prevotellaceae"               
##  [288,] "Selenomonadales"             "Veillonellaceae"              
##  [289,] "Bacteroidales"               "Prevotellaceae"               
##  [290,] "Clostridiales"               "Lachnospiraceae"              
##  [291,] "Bacteroidales"               "Prevotellaceae"               
##  [292,] "Pasteurellales"              "Pasteurellaceae"              
##  [293,] "Selenomonadales"             "Veillonellaceae"              
##  [294,] "Saccharimonadales"           "Saccharimonadaceae"           
##  [295,] "Lactobacillales"             "Streptococcaceae"             
##  [296,] "Selenomonadales"             "Veillonellaceae"              
##  [297,] NA                            NA                             
##  [298,] "Selenomonadales"             "Veillonellaceae"              
##  [299,] "Betaproteobacteriales"       "Neisseriaceae"                
##  [300,] "Bacteroidales"               "Prevotellaceae"               
##  [301,] "Bacteroidales"               "Prevotellaceae"               
##  [302,] "Fusobacteriales"             "Leptotrichiaceae"             
##  [303,] "Bacteroidales"               "Prevotellaceae"               
##  [304,] "Fusobacteriales"             "Fusobacteriaceae"             
##  [305,] "Fusobacteriales"             "Fusobacteriaceae"             
##  [306,] "Bacteroidales"               "Prevotellaceae"               
##  [307,] "Selenomonadales"             "Veillonellaceae"              
##  [308,] "Fusobacteriales"             "Fusobacteriaceae"             
##  [309,] "Clostridiales"               "Peptostreptococcaceae"        
##  [310,] "Pasteurellales"              "Pasteurellaceae"              
##  [311,] "Selenomonadales"             "Veillonellaceae"              
##  [312,] "Fusobacteriales"             "Fusobacteriaceae"             
##  [313,] "Saccharimonadales"           NA                             
##  [314,] "Campylobacterales"           "Campylobacteraceae"           
##  [315,] "Selenomonadales"             "Veillonellaceae"              
##  [316,] "Bacteroidales"               "Prevotellaceae"               
##  [317,] "Selenomonadales"             "Veillonellaceae"              
##  [318,] "Clostridiales"               "Lachnospiraceae"              
##  [319,] "Clostridiales"               "Family_XIII"                  
##  [320,] "Bacteroidales"               "Prevotellaceae"               
##  [321,] "Coriobacteriales"            "Atopobiaceae"                 
##  [322,] "Lactobacillales"             "Streptococcaceae"             
##  [323,] "Saccharimonadales"           NA                             
##  [324,] "Selenomonadales"             "Veillonellaceae"              
##  [325,] "Pasteurellales"              "Pasteurellaceae"              
##  [326,] "Saccharimonadales"           "Saccharimonadaceae"           
##  [327,] "Bacteroidales"               "Porphyromonadaceae"           
##  [328,] "Coriobacteriales"            "Atopobiaceae"                 
##  [329,] "Selenomonadales"             "Veillonellaceae"              
##  [330,] "Bacteroidales"               "Prevotellaceae"               
##  [331,] "Bacteroidales"               "Prevotellaceae"               
##  [332,] "Saccharimonadales"           NA                             
##  [333,] "Clostridiales"               "Lachnospiraceae"              
##  [334,] "Selenomonadales"             "Veillonellaceae"              
##  [335,] "Clostridiales"               "Lachnospiraceae"              
##  [336,] "Fusobacteriales"             "Leptotrichiaceae"             
##  [337,] "Lactobacillales"             "Aerococcaceae"                
##  [338,] "Fusobacteriales"             "Leptotrichiaceae"             
##  [339,] "Micrococcales"               "Micrococcaceae"               
##  [340,] "Fusobacteriales"             "Fusobacteriaceae"             
##  [341,] "Fusobacteriales"             "Fusobacteriaceae"             
##  [342,] "Bacteroidales"               "Prevotellaceae"               
##  [343,] "Betaproteobacteriales"       "Neisseriaceae"                
##  [344,] "Bacteroidales"               "Prevotellaceae"               
##  [345,] "Clostridiales"               "Peptostreptococcaceae"        
##  [346,] "Betaproteobacteriales"       "Neisseriaceae"                
##  [347,] NA                            NA                             
##  [348,] "Betaproteobacteriales"       "Neisseriaceae"                
##  [349,] "Pasteurellales"              "Pasteurellaceae"              
##  [350,] "Bacteroidales"               "Prevotellaceae"               
##  [351,] "Clostridiales"               "Family_XIII"                  
##  [352,] "Selenomonadales"             "Veillonellaceae"              
##  [353,] "Bacteroidales"               "Prevotellaceae"               
##  [354,] "Pasteurellales"              "Pasteurellaceae"              
##  [355,] "Fusobacteriales"             "Fusobacteriaceae"             
##  [356,] "Selenomonadales"             "Veillonellaceae"              
##  [357,] "Fusobacteriales"             "Fusobacteriaceae"             
##  [358,] "Bacteroidales"               "Prevotellaceae"               
##  [359,] "Selenomonadales"             "Veillonellaceae"              
##  [360,] "Clostridiales"               "Peptococcaceae"               
##  [361,] "Clostridiales"               "Family_XIII"                  
##  [362,] "Pasteurellales"              "Pasteurellaceae"              
##  [363,] "Bacteroidales"               "Prevotellaceae"               
##  [364,] "Clostridiales"               "Peptostreptococcaceae"        
##  [365,] "Fusobacteriales"             NA                             
##  [366,] "Saccharimonadales"           NA                             
##  [367,] "Clostridiales"               "Lachnospiraceae"              
##  [368,] "Clostridiales"               "Lachnospiraceae"              
##  [369,] "Actinomycetales"             "Actinomycetaceae"             
##  [370,] "Bacteroidales"               "Prevotellaceae"               
##  [371,] "Saccharimonadales"           NA                             
##  [372,] "Bacillales"                  "Family_XI"                    
##  [373,] "Lactobacillales"             "Streptococcaceae"             
##  [374,] "Selenomonadales"             "Veillonellaceae"              
##  [375,] "Saccharimonadales"           NA                             
##  [376,] "Selenomonadales"             "Veillonellaceae"              
##  [377,] "Fusobacteriales"             "Fusobacteriaceae"             
##  [378,] "Lactobacillales"             "Streptococcaceae"             
##  [379,] "Selenomonadales"             "Veillonellaceae"              
##  [380,] "Fusobacteriales"             "Leptotrichiaceae"             
##  [381,] "Fusobacteriales"             "Fusobacteriaceae"             
##  [382,] "Clostridiales"               "Family_XI"                    
##  [383,] "Bacteroidales"               "Prevotellaceae"               
##  [384,] "Bacteroidales"               "Prevotellaceae"               
##  [385,] "Lactobacillales"             "Streptococcaceae"             
##  [386,] "Enterobacteriales"           "Enterobacteriaceae"           
##  [387,] "Fusobacteriales"             "Leptotrichiaceae"             
##  [388,] "Actinomycetales"             "Actinomycetaceae"             
##  [389,] "Fusobacteriales"             "Leptotrichiaceae"             
##  [390,] "Bacteroidales"               "Prevotellaceae"               
##  [391,] "Actinomycetales"             "Actinomycetaceae"             
##  [392,] "Fusobacteriales"             "Fusobacteriaceae"             
##  [393,] "Pasteurellales"              "Pasteurellaceae"              
##  [394,] "Enterobacteriales"           "Enterobacteriaceae"           
##  [395,] "Selenomonadales"             "Veillonellaceae"              
##  [396,] "Bacteroidales"               "Prevotellaceae"               
##  [397,] "Bacteroidales"               "Prevotellaceae"               
##  [398,] "Pasteurellales"              "Pasteurellaceae"              
##  [399,] "Fusobacteriales"             "Leptotrichiaceae"             
##  [400,] "Pasteurellales"              "Pasteurellaceae"              
##  [401,] "Lactobacillales"             "Streptococcaceae"             
##  [402,] "Flavobacteriales"            "Flavobacteriaceae"            
##  [403,] "Bacteroidales"               "Prevotellaceae"               
##  [404,] "Bacteroidales"               "Prevotellaceae"               
##  [405,] "Clostridiales"               "Lachnospiraceae"              
##  [406,] "Selenomonadales"             "Veillonellaceae"              
##  [407,] "Bacteroidales"               "Prevotellaceae"               
##  [408,] "Bacillales"                  "Family_XI"                    
##  [409,] "Selenomonadales"             "Veillonellaceae"              
##  [410,] "Bacteroidales"               "Porphyromonadaceae"           
##  [411,] "Bacteroidales"               "Prevotellaceae"               
##  [412,] "Bacteroidales"               "Prevotellaceae"               
##  [413,] "Betaproteobacteriales"       "Neisseriaceae"                
##  [414,] "Clostridiales"               "Lachnospiraceae"              
##  [415,] "Bacteroidales"               "Prevotellaceae"               
##  [416,] "Pseudomonadales"             "Moraxellaceae"                
##  [417,] "Bacteroidales"               "Prevotellaceae"               
##  [418,] "Bacteroidales"               "Prevotellaceae"               
##  [419,] "Fusobacteriales"             "Leptotrichiaceae"             
##  [420,] "Bacteroidales"               "Prevotellaceae"               
##  [421,] "Bacillales"                  "Family_XI"                    
##  [422,] "Selenomonadales"             "Veillonellaceae"              
##  [423,] "Selenomonadales"             "Veillonellaceae"              
##  [424,] "Selenomonadales"             "Veillonellaceae"              
##  [425,] "Pasteurellales"              "Pasteurellaceae"              
##  [426,] "Bacteroidales"               "Prevotellaceae"               
##  [427,] "Bacteroidales"               "Prevotellaceae"               
##  [428,] "Bacteroidales"               "Prevotellaceae"               
##  [429,] "Selenomonadales"             "Veillonellaceae"              
##  [430,] "Selenomonadales"             "Veillonellaceae"              
##  [431,] "Selenomonadales"             "Veillonellaceae"              
##  [432,] "Bacteroidales"               "Prevotellaceae"               
##  [433,] "Selenomonadales"             "Veillonellaceae"              
##  [434,] "Bacteroidales"               "Prevotellaceae"               
##  [435,] "Clostridiales"               "Lachnospiraceae"              
##  [436,] "Selenomonadales"             "Veillonellaceae"              
##  [437,] "Bacteroidales"               "Prevotellaceae"               
##  [438,] "Lactobacillales"             "Streptococcaceae"             
##  [439,] "Bacteroidales"               "Prevotellaceae"               
##  [440,] "Betaproteobacteriales"       "Neisseriaceae"                
##  [441,] "Bacteroidales"               "Porphyromonadaceae"           
##  [442,] "Fusobacteriales"             "Fusobacteriaceae"             
##  [443,] "Selenomonadales"             "Veillonellaceae"              
##  [444,] "Selenomonadales"             "Veillonellaceae"              
##  [445,] "Selenomonadales"             "Veillonellaceae"              
##  [446,] "Selenomonadales"             "Veillonellaceae"              
##  [447,] "Lactobacillales"             "Streptococcaceae"             
##  [448,] "Fusobacteriales"             "Leptotrichiaceae"             
##  [449,] "Flavobacteriales"            "Flavobacteriaceae"            
##  [450,] "Fusobacteriales"             "Leptotrichiaceae"             
##  [451,] "Bacteroidales"               "Prevotellaceae"               
##  [452,] "Campylobacterales"           NA                             
##  [453,] "Clostridiales"               "Family_XIII"                  
##  [454,] "Flavobacteriales"            "Flavobacteriaceae"            
##  [455,] "Selenomonadales"             "Veillonellaceae"              
##  [456,] "Clostridiales"               "Ruminococcaceae"              
##  [457,] "Betaproteobacteriales"       "Neisseriaceae"                
##  [458,] "Actinomycetales"             "Actinomycetaceae"             
##  [459,] "Fusobacteriales"             "Leptotrichiaceae"             
##  [460,] "Selenomonadales"             "Veillonellaceae"              
##  [461,] "Saccharimonadales"           "Saccharimonadaceae"           
##  [462,] "Fusobacteriales"             "Leptotrichiaceae"             
##  [463,] "Bacteroidales"               "Prevotellaceae"               
##  [464,] "Lactobacillales"             "Streptococcaceae"             
##  [465,] "Bacteroidales"               "Prevotellaceae"               
##  [466,] "Selenomonadales"             "Veillonellaceae"              
##  [467,] "Bacteroidales"               "Prevotellaceae"               
##  [468,] "Pasteurellales"              "Pasteurellaceae"              
##  [469,] "Saccharimonadales"           NA                             
##  [470,] "Betaproteobacteriales"       "Neisseriaceae"                
##  [471,] "Pasteurellales"              "Pasteurellaceae"              
##  [472,] "Bacteroidales"               "Prevotellaceae"               
##  [473,] "Bacteroidales"               "Prevotellaceae"               
##  [474,] "Selenomonadales"             "Veillonellaceae"              
##  [475,] "Bacteroidales"               "Prevotellaceae"               
##  [476,] "Pasteurellales"              "Pasteurellaceae"              
##  [477,] "Bacteroidales"               "Prevotellaceae"               
##  [478,] "Pasteurellales"              "Pasteurellaceae"              
##  [479,] "Enterobacteriales"           "Enterobacteriaceae"           
##  [480,] "Fusobacteriales"             "Leptotrichiaceae"             
##  [481,] "Bacteroidales"               "Prevotellaceae"               
##  [482,] "Bacteroidales"               "Prevotellaceae"               
##  [483,] "Fusobacteriales"             "Fusobacteriaceae"             
##  [484,] "Selenomonadales"             "Veillonellaceae"              
##  [485,] "Lactobacillales"             "Streptococcaceae"             
##  [486,] "Fusobacteriales"             NA                             
##  [487,] "Bacteroidales"               "Prevotellaceae"               
##  [488,] "Fusobacteriales"             "Leptotrichiaceae"             
##  [489,] "Bacteroidales"               "Porphyromonadaceae"           
##  [490,] "Campylobacterales"           "Campylobacteraceae"           
##  [491,] "Selenomonadales"             "Veillonellaceae"              
##  [492,] "Campylobacterales"           NA                             
##  [493,] "Bacteroidales"               "Prevotellaceae"               
##  [494,] "Fusobacteriales"             "Fusobacteriaceae"             
##  [495,] "Bacteroidales"               "Prevotellaceae"               
##  [496,] "Selenomonadales"             "Veillonellaceae"              
##  [497,] "Enterobacteriales"           "Enterobacteriaceae"           
##  [498,] "Bacteroidales"               "Prevotellaceae"               
##  [499,] "Clostridiales"               "Peptostreptococcaceae"        
##  [500,] "Selenomonadales"             "Veillonellaceae"              
##  [501,] "Selenomonadales"             "Veillonellaceae"              
##  [502,] "Actinomycetales"             "Actinomycetaceae"             
##  [503,] "Enterobacteriales"           "Enterobacteriaceae"           
##  [504,] "Selenomonadales"             "Veillonellaceae"              
##  [505,] "Betaproteobacteriales"       "Neisseriaceae"                
##  [506,] "Fusobacteriales"             "Leptotrichiaceae"             
##  [507,] "Actinomycetales"             "Actinomycetaceae"             
##  [508,] "Bacteroidales"               "Prevotellaceae"               
##  [509,] "Bacteroidales"               "Prevotellaceae"               
##  [510,] "Pasteurellales"              "Pasteurellaceae"              
##  [511,] "Selenomonadales"             "Veillonellaceae"              
##  [512,] "Selenomonadales"             "Veillonellaceae"              
##  [513,] "Pasteurellales"              "Pasteurellaceae"              
##  [514,] "Selenomonadales"             "Veillonellaceae"              
##  [515,] "Selenomonadales"             "Veillonellaceae"              
##  [516,] "Selenomonadales"             "Veillonellaceae"              
##  [517,] "Bacteroidales"               "Prevotellaceae"               
##  [518,] "Pasteurellales"              "Pasteurellaceae"              
##  [519,] "Selenomonadales"             "Veillonellaceae"              
##  [520,] "Actinomycetales"             "Actinomycetaceae"             
##  [521,] "Lactobacillales"             "Streptococcaceae"             
##  [522,] "Actinomycetales"             "Actinomycetaceae"             
##  [523,] "Clostridiales"               "Family_XI"                    
##  [524,] "Selenomonadales"             "Veillonellaceae"              
##  [525,] "Fusobacteriales"             "Fusobacteriaceae"             
##  [526,] "Bacteroidales"               "Prevotellaceae"               
##  [527,] "Bacteroidales"               "Prevotellaceae"               
##  [528,] "Betaproteobacteriales"       "Neisseriaceae"                
##  [529,] "Campylobacterales"           "Campylobacteraceae"           
##  [530,] "Lactobacillales"             "Streptococcaceae"             
##  [531,] "Cardiobacteriales"           "Cardiobacteriaceae"           
##  [532,] "Bacteroidales"               "Prevotellaceae"               
##  [533,] "Saccharimonadales"           NA                             
##  [534,] "Selenomonadales"             "Veillonellaceae"              
##  [535,] "Flavobacteriales"            "Flavobacteriaceae"            
##  [536,] "Bacteroidales"               "Prevotellaceae"               
##  [537,] "Selenomonadales"             "Veillonellaceae"              
##  [538,] "Pasteurellales"              "Pasteurellaceae"              
##  [539,] "Bacteroidales"               "Paludibacteraceae"            
##  [540,] "Enterobacteriales"           "Enterobacteriaceae"           
##  [541,] "Fusobacteriales"             "Leptotrichiaceae"             
##  [542,] "Lactobacillales"             "Streptococcaceae"             
##  [543,] "Bacteroidales"               "Porphyromonadaceae"           
##  [544,] "Bacteroidales"               "Prevotellaceae"               
##  [545,] "Clostridiales"               "Family_XIII"                  
##  [546,] "Bacteroidales"               "Prevotellaceae"               
##  [547,] "Betaproteobacteriales"       "Neisseriaceae"                
##  [548,] NA                            NA                             
##  [549,] "Selenomonadales"             "Veillonellaceae"              
##  [550,] "Bacteroidales"               "Prevotellaceae"               
##  [551,] "Bacillales"                  "Family_XI"                    
##  [552,] "Betaproteobacteriales"       "Neisseriaceae"                
##  [553,] "Bacteroidales"               "Prevotellaceae"               
##  [554,] "Flavobacteriales"            "Flavobacteriaceae"            
##  [555,] "Bacteroidales"               "Prevotellaceae"               
##  [556,] "Selenomonadales"             "Veillonellaceae"              
##  [557,] "Selenomonadales"             "Veillonellaceae"              
##  [558,] "Lactobacillales"             "Streptococcaceae"             
##  [559,] "Bacteroidales"               "Prevotellaceae"               
##  [560,] "Pasteurellales"              "Pasteurellaceae"              
##  [561,] "Bacteroidales"               "Porphyromonadaceae"           
##  [562,] "Saccharimonadales"           "Saccharimonadaceae"           
##  [563,] "Pasteurellales"              "Pasteurellaceae"              
##  [564,] "Lactobacillales"             "Streptococcaceae"             
##  [565,] "Selenomonadales"             "Veillonellaceae"              
##  [566,] "Bacteroidales"               "Prevotellaceae"               
##  [567,] "Lactobacillales"             "Streptococcaceae"             
##  [568,] "Bacteroidales"               "Prevotellaceae"               
##  [569,] "Bacteroidales"               "Prevotellaceae"               
##  [570,] "Fusobacteriales"             "Leptotrichiaceae"             
##  [571,] "Bacteroidales"               "Prevotellaceae"               
##  [572,] "Bacteroidales"               "Tannerellaceae"               
##  [573,] "Betaproteobacteriales"       "Neisseriaceae"                
##  [574,] "Bacteroidales"               "Prevotellaceae"               
##  [575,] "Flavobacteriales"            "Flavobacteriaceae"            
##  [576,] "Bacteroidales"               "Prevotellaceae"               
##  [577,] "Saccharimonadales"           "Saccharimonadaceae"           
##  [578,] "Selenomonadales"             "Veillonellaceae"              
##  [579,] "Selenomonadales"             "Veillonellaceae"              
##  [580,] "Selenomonadales"             "Veillonellaceae"              
##  [581,] "Fusobacteriales"             "Leptotrichiaceae"             
##  [582,] "Pasteurellales"              "Pasteurellaceae"              
##  [583,] "Fusobacteriales"             "Fusobacteriaceae"             
##  [584,] "Bacteroidales"               "Porphyromonadaceae"           
##  [585,] "Betaproteobacteriales"       "Neisseriaceae"                
##  [586,] "Fusobacteriales"             "Fusobacteriaceae"             
##  [587,] "Fusobacteriales"             "Fusobacteriaceae"             
##  [588,] "Bacteroidales"               "Tannerellaceae"               
##  [589,] "Fusobacteriales"             "Fusobacteriaceae"             
##  [590,] "Bacteroidales"               "Prevotellaceae"               
##  [591,] "Betaproteobacteriales"       "Neisseriaceae"                
##  [592,] "Bacteroidales"               "Prevotellaceae"               
##  [593,] "Bacteroidales"               "Prevotellaceae"               
##  [594,] "Bacteroidales"               "Prevotellaceae"               
##  [595,] "Pasteurellales"              "Pasteurellaceae"              
##  [596,] "Bacteroidales"               "Prevotellaceae"               
##  [597,] "Selenomonadales"             "Veillonellaceae"              
##  [598,] "Betaproteobacteriales"       "Neisseriaceae"                
##  [599,] "Clostridiales"               "Peptostreptococcaceae"        
##  [600,] "Selenomonadales"             "Veillonellaceae"              
##  [601,] "Pasteurellales"              "Pasteurellaceae"              
##  [602,] "Bacteroidales"               "Prevotellaceae"               
##  [603,] "Bacteroidales"               "Prevotellaceae"               
##  [604,] "Selenomonadales"             "Veillonellaceae"              
##  [605,] "Betaproteobacteriales"       "Neisseriaceae"                
##  [606,] "Fusobacteriales"             "Fusobacteriaceae"             
##  [607,] "Bacteroidales"               "Prevotellaceae"               
##  [608,] "Saccharimonadales"           "Saccharimonadaceae"           
##  [609,] "Bacteroidales"               "Prevotellaceae"               
##  [610,] "Saccharimonadales"           "Saccharimonadaceae"           
##  [611,] "Pasteurellales"              "Pasteurellaceae"              
##  [612,] "Selenomonadales"             "Veillonellaceae"              
##  [613,] "Fusobacteriales"             "Fusobacteriaceae"             
##  [614,] "Fusobacteriales"             "Leptotrichiaceae"             
##  [615,] "Bacteroidales"               "Prevotellaceae"               
##  [616,] "Selenomonadales"             "Veillonellaceae"              
##  [617,] "Bacteroidales"               "Prevotellaceae"               
##  [618,] "Selenomonadales"             "Veillonellaceae"              
##  [619,] "Bacillales"                  "Family_XI"                    
##  [620,] "Bacteroidales"               "Porphyromonadaceae"           
##  [621,] "Selenomonadales"             "Veillonellaceae"              
##  [622,] "Bacteroidales"               "Paludibacteraceae"            
##  [623,] "Selenomonadales"             "Veillonellaceae"              
##  [624,] "Bacteroidales"               "Prevotellaceae"               
##  [625,] "Bacteroidales"               "Prevotellaceae"               
##  [626,] "Bacteroidales"               "Prevotellaceae"               
##  [627,] "Bacteroidales"               "Prevotellaceae"               
##  [628,] "Fusobacteriales"             "Fusobacteriaceae"             
##  [629,] "Bacteroidales"               "Prevotellaceae"               
##  [630,] "Micrococcales"               "Micrococcaceae"               
##  [631,] "Selenomonadales"             "Veillonellaceae"              
##  [632,] "Pasteurellales"              "Pasteurellaceae"              
##  [633,] "Coriobacteriales"            "Atopobiaceae"                 
##  [634,] "Selenomonadales"             "Veillonellaceae"              
##  [635,] "Saccharimonadales"           "Saccharimonadaceae"           
##  [636,] "Micrococcales"               NA                             
##  [637,] "Bacteroidales"               "Porphyromonadaceae"           
##  [638,] "Pasteurellales"              "Pasteurellaceae"              
##  [639,] "Lactobacillales"             "Streptococcaceae"             
##  [640,] "Absconditabacteriales_(SR1)" NA                             
##  [641,] "Selenomonadales"             "Veillonellaceae"              
##  [642,] NA                            NA                             
##  [643,] "Enterobacteriales"           "Enterobacteriaceae"           
##  [644,] "Bacteroidales"               "Prevotellaceae"               
##  [645,] "Bacteroidales"               "Prevotellaceae"               
##  [646,] "Bacteroidales"               "Prevotellaceae"               
##  [647,] "Micrococcales"               "Micrococcaceae"               
##  [648,] "Lactobacillales"             "Streptococcaceae"             
##  [649,] "Bacteroidales"               "Prevotellaceae"               
##  [650,] "Clostridiales"               "Peptostreptococcaceae"        
##  [651,] "Selenomonadales"             "Veillonellaceae"              
##  [652,] "Bacteroidales"               "Prevotellaceae"               
##  [653,] "Selenomonadales"             "Veillonellaceae"              
##  [654,] "Saccharimonadales"           NA                             
##  [655,] "Bacteroidales"               "Prevotellaceae"               
##  [656,] "Selenomonadales"             "Veillonellaceae"              
##  [657,] "Fusobacteriales"             "Leptotrichiaceae"             
##  [658,] "Bacteroidales"               "Porphyromonadaceae"           
##  [659,] "Selenomonadales"             "Veillonellaceae"              
##  [660,] "Fusobacteriales"             "Leptotrichiaceae"             
##  [661,] "Micrococcales"               NA                             
##  [662,] "Fusobacteriales"             "Fusobacteriaceae"             
##  [663,] "Fusobacteriales"             "Fusobacteriaceae"             
##  [664,] "Pasteurellales"              "Pasteurellaceae"              
##  [665,] "Fusobacteriales"             "Leptotrichiaceae"             
##  [666,] "Selenomonadales"             "Veillonellaceae"              
##  [667,] "Chloroplast"                 NA                             
##  [668,] "Lactobacillales"             NA                             
##  [669,] "Sphingobacteriales"          "Lentimicrobiaceae"            
##  [670,] "Bacteroidales"               "Tannerellaceae"               
##  [671,] "Actinomycetales"             "Actinomycetaceae"             
##  [672,] "Micrococcales"               "Micrococcaceae"               
##  [673,] "Saccharimonadales"           NA                             
##  [674,] "Bacteroidales"               "Prevotellaceae"               
##  [675,] "Lactobacillales"             "Streptococcaceae"             
##  [676,] "Pasteurellales"              "Pasteurellaceae"              
##  [677,] "Bacteroidales"               "Prevotellaceae"               
##  [678,] "Selenomonadales"             "Veillonellaceae"              
##  [679,] "Bacteroidales"               "Prevotellaceae"               
##  [680,] NA                            NA                             
##  [681,] "Flavobacteriales"            "Flavobacteriaceae"            
##  [682,] "Pasteurellales"              "Pasteurellaceae"              
##  [683,] "Selenomonadales"             "Veillonellaceae"              
##  [684,] "Bacteroidales"               "Porphyromonadaceae"           
##  [685,] "Bacteroidales"               "Prevotellaceae"               
##  [686,] "Bacteroidales"               "Prevotellaceae"               
##  [687,] "Saccharimonadales"           "Saccharimonadaceae"           
##  [688,] "Lactobacillales"             "Streptococcaceae"             
##  [689,] "Selenomonadales"             "Veillonellaceae"              
##  [690,] "Selenomonadales"             "Veillonellaceae"              
##  [691,] "Bacteroidales"               "Porphyromonadaceae"           
##  [692,] "Saccharimonadales"           "Saccharimonadaceae"           
##  [693,] NA                            NA                             
##  [694,] "Bacteroidales"               "Prevotellaceae"               
##  [695,] "Selenomonadales"             "Veillonellaceae"              
##  [696,] "Lactobacillales"             "Streptococcaceae"             
##  [697,] "Selenomonadales"             "Veillonellaceae"              
##  [698,] "Bacteroidales"               "Prevotellaceae"               
##  [699,] "Bacteroidales"               "Prevotellaceae"               
##  [700,] "Spirochaetales"              "Spirochaetaceae"              
##  [701,] "Bacteroidales"               "Prevotellaceae"               
##  [702,] "Selenomonadales"             "Veillonellaceae"              
##  [703,] "Fusobacteriales"             "Leptotrichiaceae"             
##  [704,] "Campylobacterales"           "P6-b51"                       
##  [705,] "Bacteroidales"               "Prevotellaceae"               
##  [706,] "Saccharimonadales"           "Saccharimonadaceae"           
##  [707,] "Actinomycetales"             "Actinomycetaceae"             
##  [708,] "Bacteroidales"               "Prevotellaceae"               
##  [709,] "Bacteroidales"               "Prevotellaceae"               
##  [710,] "Pasteurellales"              "Pasteurellaceae"              
##  [711,] "Bacteroidales"               "Prevotellaceae"               
##  [712,] "Selenomonadales"             "Veillonellaceae"              
##  [713,] "Bacteroidales"               "Porphyromonadaceae"           
##  [714,] "Bacteroidales"               "Prevotellaceae"               
##  [715,] "Betaproteobacteriales"       "Neisseriaceae"                
##  [716,] "Saccharimonadales"           NA                             
##  [717,] "Lactobacillales"             "Streptococcaceae"             
##  [718,] "Spirochaetales"              "Spirochaetaceae"              
##  [719,] "Bacteroidales"               "Prevotellaceae"               
##  [720,] "Pasteurellales"              "Pasteurellaceae"              
##  [721,] "Bacteroidales"               "Prevotellaceae"               
##  [722,] "Bacteroidales"               "Prevotellaceae"               
##  [723,] "Flavobacteriales"            "Flavobacteriaceae"            
##  [724,] "Betaproteobacteriales"       "Burkholderiaceae"             
##  [725,] "Pasteurellales"              "Pasteurellaceae"              
##  [726,] "Betaproteobacteriales"       "Neisseriaceae"                
##  [727,] "Pasteurellales"              "Pasteurellaceae"              
##  [728,] "Selenomonadales"             "Veillonellaceae"              
##  [729,] "Selenomonadales"             "Veillonellaceae"              
##  [730,] "Flavobacteriales"            "Flavobacteriaceae"            
##  [731,] "Betaproteobacteriales"       "Neisseriaceae"                
##  [732,] "Fusobacteriales"             "Fusobacteriaceae"             
##  [733,] "Pasteurellales"              "Pasteurellaceae"              
##  [734,] "Selenomonadales"             "Veillonellaceae"              
##  [735,] "Betaproteobacteriales"       "Neisseriaceae"                
##  [736,] "Flavobacteriales"            "Flavobacteriaceae"            
##  [737,] "Betaproteobacteriales"       "Neisseriaceae"                
##  [738,] "Actinomycetales"             "Actinomycetaceae"             
##  [739,] "Betaproteobacteriales"       "Neisseriaceae"                
##  [740,] "Flavobacteriales"            "Weeksellaceae"                
##  [741,] "Fusobacteriales"             "Leptotrichiaceae"             
##  [742,] "Lactobacillales"             "Streptococcaceae"             
##  [743,] "Micrococcales"               "Micrococcaceae"               
##  [744,] "Lactobacillales"             "Streptococcaceae"             
##  [745,] "Selenomonadales"             "Veillonellaceae"              
##  [746,] "Selenomonadales"             "Veillonellaceae"              
##  [747,] "Bacteroidales"               "Prevotellaceae"               
##  [748,] "Saccharimonadales"           NA                             
##  [749,] "Fusobacteriales"             "Fusobacteriaceae"             
##  [750,] "Pasteurellales"              "Pasteurellaceae"              
##  [751,] "Bacteroidales"               "Prevotellaceae"               
##  [752,] "Bacteroidales"               "Prevotellaceae"               
##  [753,] "Bacteroidales"               "Prevotellaceae"               
##  [754,] "Flavobacteriales"            "Flavobacteriaceae"            
##  [755,] "Selenomonadales"             "Veillonellaceae"              
##  [756,] "Selenomonadales"             "Veillonellaceae"              
##  [757,] "Bacteroidales"               "Prevotellaceae"               
##  [758,] "Fusobacteriales"             "Leptotrichiaceae"             
##  [759,] "Selenomonadales"             "Veillonellaceae"              
##  [760,] "Bacteroidales"               "Porphyromonadaceae"           
##  [761,] "Selenomonadales"             "Veillonellaceae"              
##  [762,] "Bacteroidales"               "Porphyromonadaceae"           
##  [763,] "Bacteroidales"               "Prevotellaceae"               
##  [764,] "Selenomonadales"             "Veillonellaceae"              
##  [765,] "Fusobacteriales"             "Fusobacteriaceae"             
##  [766,] "Selenomonadales"             "Veillonellaceae"              
##  [767,] "Fusobacteriales"             "Fusobacteriaceae"             
##  [768,] "Fusobacteriales"             "Fusobacteriaceae"             
##  [769,] "Bacteroidales"               "Prevotellaceae"               
##  [770,] "Bacteroidales"               "Prevotellaceae"               
##  [771,] "Lactobacillales"             "Streptococcaceae"             
##  [772,] "Fusobacteriales"             "Leptotrichiaceae"             
##  [773,] "Bacteroidales"               "Prevotellaceae"               
##  [774,] "Lactobacillales"             "Streptococcaceae"             
##  [775,] "Actinomycetales"             "Actinomycetaceae"             
##  [776,] "Selenomonadales"             "Veillonellaceae"              
##  [777,] "Selenomonadales"             "Veillonellaceae"              
##  [778,] "Bacteroidales"               "Prevotellaceae"               
##  [779,] "Betaproteobacteriales"       "Neisseriaceae"                
##  [780,] "Clostridiales"               "Lachnospiraceae"              
##  [781,] "Bacteroidales"               "Prevotellaceae"               
##  [782,] "Pasteurellales"              "Pasteurellaceae"              
##  [783,] "Bacteroidales"               "Prevotellaceae"               
##  [784,] "Pasteurellales"              "Pasteurellaceae"              
##  [785,] "Clostridiales"               "Family_XI"                    
##  [786,] "Lactobacillales"             "Aerococcaceae"                
##  [787,] "Bacteroidales"               "Prevotellaceae"               
##  [788,] "Bacteroidales"               "Prevotellaceae"               
##  [789,] "Pasteurellales"              "Pasteurellaceae"              
##  [790,] "Clostridiales"               "Lachnospiraceae"              
##  [791,] "Selenomonadales"             "Veillonellaceae"              
##  [792,] "Clostridiales"               "Lachnospiraceae"              
##  [793,] "Selenomonadales"             "Veillonellaceae"              
##  [794,] "Fusobacteriales"             "Leptotrichiaceae"             
##  [795,] "Campylobacterales"           "Campylobacteraceae"           
##  [796,] "Bacteroidales"               "Prevotellaceae"               
##  [797,] NA                            NA                             
##  [798,] "Selenomonadales"             "Veillonellaceae"              
##  [799,] "Selenomonadales"             "Veillonellaceae"              
##  [800,] "Betaproteobacteriales"       "Neisseriaceae"                
##  [801,] "Bacteroidales"               "Prevotellaceae"               
##  [802,] "Fusobacteriales"             "Leptotrichiaceae"             
##  [803,] "Bacteroidales"               "Porphyromonadaceae"           
##  [804,] "Bacteroidales"               "Prevotellaceae"               
##  [805,] "Micrococcales"               NA                             
##  [806,] "Clostridiales"               "Lachnospiraceae"              
##  [807,] "Pasteurellales"              "Pasteurellaceae"              
##  [808,] "Lactobacillales"             "Streptococcaceae"             
##  [809,] "Bacteroidales"               "Prevotellaceae"               
##  [810,] "Saccharimonadales"           "Saccharimonadaceae"           
##  [811,] "Bacteroidales"               "Prevotellaceae"               
##  [812,] "Actinomycetales"             "Actinomycetaceae"             
##  [813,] "Fusobacteriales"             "Leptotrichiaceae"             
##  [814,] "Selenomonadales"             "Veillonellaceae"              
##  [815,] "Bacteroidales"               "Prevotellaceae"               
##  [816,] "Pasteurellales"              "Pasteurellaceae"              
##  [817,] "Bacteroidales"               "Porphyromonadaceae"           
##  [818,] "Selenomonadales"             "Veillonellaceae"              
##  [819,] "Bacteroidales"               "Prevotellaceae"               
##  [820,] "Saccharimonadales"           "Saccharimonadaceae"           
##  [821,] "Betaproteobacteriales"       "Neisseriaceae"                
##  [822,] "Fusobacteriales"             "Leptotrichiaceae"             
##  [823,] "Lactobacillales"             "Carnobacteriaceae"            
##  [824,] "Bacteroidales"               "Prevotellaceae"               
##  [825,] "Fusobacteriales"             "Leptotrichiaceae"             
##  [826,] "Bacteroidales"               "Prevotellaceae"               
##  [827,] "Bacteroidales"               "Prevotellaceae"               
##  [828,] "Betaproteobacteriales"       NA                             
##  [829,] "Bacteroidales"               "Prevotellaceae"               
##  [830,] "Pasteurellales"              "Pasteurellaceae"              
##  [831,] "Selenomonadales"             "Veillonellaceae"              
##  [832,] "Bacteroidales"               "Prevotellaceae"               
##  [833,] "Clostridiales"               "Peptostreptococcaceae"        
##  [834,] "Micrococcales"               NA                             
##  [835,] "Selenomonadales"             "Veillonellaceae"              
##  [836,] "Selenomonadales"             "Veillonellaceae"              
##  [837,] "Lactobacillales"             "Carnobacteriaceae"            
##  [838,] "Bacteroidales"               "Prevotellaceae"               
##  [839,] "Campylobacterales"           "Campylobacteraceae"           
##  [840,] "Saccharimonadales"           "Saccharimonadaceae"           
##  [841,] "Selenomonadales"             "Veillonellaceae"              
##  [842,] "Bacteroidales"               "Porphyromonadaceae"           
##  [843,] "Clostridiales"               "Family_XI"                    
##  [844,] "Bacteroidales"               "Porphyromonadaceae"           
##  [845,] NA                            NA                             
##  [846,] "Actinomycetales"             "Actinomycetaceae"             
##  [847,] "Selenomonadales"             "Veillonellaceae"              
##  [848,] "Micrococcales"               "Micrococcaceae"               
##  [849,] "Bacteroidales"               "Prevotellaceae"               
##  [850,] "Pasteurellales"              "Pasteurellaceae"              
##  [851,] "Bacteroidales"               "Prevotellaceae"               
##  [852,] "Lactobacillales"             "Streptococcaceae"             
##  [853,] "Bacteroidales"               "Prevotellaceae"               
##  [854,] "Selenomonadales"             "Veillonellaceae"              
##  [855,] "Bacteroidales"               "Prevotellaceae"               
##  [856,] "Clostridiales"               "Peptostreptococcaceae"        
##  [857,] "Bacteroidales"               "Prevotellaceae"               
##  [858,] "Lactobacillales"             "Streptococcaceae"             
##  [859,] "Bacteroidales"               "Prevotellaceae"               
##  [860,] "Saccharimonadales"           "Saccharimonadaceae"           
##  [861,] "Clostridiales"               "Defluviitaleaceae"            
##  [862,] "Clostridiales"               "Lachnospiraceae"              
##  [863,] "Bacteroidales"               "Prevotellaceae"               
##  [864,] "Bacteroidales"               "Prevotellaceae"               
##  [865,] "Bacteroidales"               "Prevotellaceae"               
##  [866,] "Bacteroidales"               "Porphyromonadaceae"           
##  [867,] "Actinomycetales"             "Actinomycetaceae"             
##  [868,] "Bacteroidales"               "Prevotellaceae"               
##  [869,] "Selenomonadales"             "Veillonellaceae"              
##  [870,] "Actinomycetales"             "Actinomycetaceae"             
##  [871,] "Actinomycetales"             "Actinomycetaceae"             
##  [872,] "Selenomonadales"             "Veillonellaceae"              
##  [873,] "Bacteroidales"               "Porphyromonadaceae"           
##  [874,] "Bacteroidales"               "Porphyromonadaceae"           
##  [875,] "Actinomycetales"             "Actinomycetaceae"             
##  [876,] "Fusobacteriales"             "Leptotrichiaceae"             
##  [877,] "Saccharimonadales"           "Saccharimonadaceae"           
##  [878,] "Selenomonadales"             "Veillonellaceae"              
##  [879,] "Actinomycetales"             "Actinomycetaceae"             
##  [880,] "Selenomonadales"             "Veillonellaceae"              
##  [881,] "Pasteurellales"              "Pasteurellaceae"              
##  [882,] "Clostridiales"               "Lachnospiraceae"              
##  [883,] "Selenomonadales"             "Veillonellaceae"              
##  [884,] "Bacteroidales"               "Prevotellaceae"               
##  [885,] "Bacteroidales"               "Prevotellaceae"               
##  [886,] "Bacteroidales"               "Porphyromonadaceae"           
##  [887,] "Selenomonadales"             "Veillonellaceae"              
##  [888,] "Actinomycetales"             "Actinomycetaceae"             
##  [889,] "Micrococcales"               NA                             
##  [890,] "Clostridiales"               "Lachnospiraceae"              
##  [891,] "Pasteurellales"              "Pasteurellaceae"              
##  [892,] "Lactobacillales"             "Streptococcaceae"             
##  [893,] "Saccharimonadales"           "Saccharimonadaceae"           
##  [894,] "Flavobacteriales"            "Weeksellaceae"                
##  [895,] "Actinomycetales"             "Actinomycetaceae"             
##  [896,] "Bacteroidales"               "Prevotellaceae"               
##  [897,] "Bacteroidales"               "Prevotellaceae"               
##  [898,] "Bacteroidales"               "Prevotellaceae"               
##  [899,] "Saccharimonadales"           "Saccharimonadaceae"           
##  [900,] NA                            NA                             
##  [901,] "Fusobacteriales"             "Fusobacteriaceae"             
##  [902,] "Pasteurellales"              "Pasteurellaceae"              
##  [903,] "Lactobacillales"             "Streptococcaceae"             
##  [904,] "Clostridiales"               "Ruminococcaceae"              
##  [905,] "Micrococcales"               NA                             
##  [906,] "Bacteroidales"               "Prevotellaceae"               
##  [907,] "Lactobacillales"             "Streptococcaceae"             
##  [908,] "Pasteurellales"              "Pasteurellaceae"              
##  [909,] "Selenomonadales"             "Veillonellaceae"              
##  [910,] "Pasteurellales"              "Pasteurellaceae"              
##  [911,] "Selenomonadales"             "Veillonellaceae"              
##  [912,] "Saccharimonadales"           "Saccharimonadaceae"           
##  [913,] "Betaproteobacteriales"       "Neisseriaceae"                
##  [914,] "Pasteurellales"              "Pasteurellaceae"              
##  [915,] "Bifidobacteriales"           "Bifidobacteriaceae"           
##  [916,] "Selenomonadales"             "Veillonellaceae"              
##  [917,] "Bacteroidales"               "Prevotellaceae"               
##  [918,] "Clostridiales"               "Lachnospiraceae"              
##  [919,] "Actinomycetales"             "Actinomycetaceae"             
##  [920,] "Betaproteobacteriales"       "Neisseriaceae"                
##  [921,] "Campylobacterales"           "Campylobacteraceae"           
##  [922,] "Selenomonadales"             "Veillonellaceae"              
##  [923,] "Selenomonadales"             "Veillonellaceae"              
##  [924,] NA                            NA                             
##  [925,] "Fusobacteriales"             "Leptotrichiaceae"             
##  [926,] "Betaproteobacteriales"       "Neisseriaceae"                
##  [927,] "Fusobacteriales"             "Fusobacteriaceae"             
##  [928,] "Fusobacteriales"             "Fusobacteriaceae"             
##  [929,] "Selenomonadales"             "Veillonellaceae"              
##  [930,] "Bacteroidales"               "Prevotellaceae"               
##  [931,] "Fusobacteriales"             "Leptotrichiaceae"             
##  [932,] "Lactobacillales"             "Streptococcaceae"             
##  [933,] NA                            NA                             
##  [934,] "Selenomonadales"             "Veillonellaceae"              
##  [935,] "Bacteroidales"               "Prevotellaceae"               
##  [936,] "Clostridiales"               "Family_XIII"                  
##  [937,] "Fusobacteriales"             "Leptotrichiaceae"             
##  [938,] "Erysipelotrichales"          "Erysipelotrichaceae"          
##  [939,] "Lactobacillales"             "Streptococcaceae"             
##  [940,] "Bacteroidales"               "Prevotellaceae"               
##  [941,] "Lactobacillales"             "Streptococcaceae"             
##  [942,] "Actinomycetales"             "Actinomycetaceae"             
##  [943,] "Campylobacterales"           NA                             
##  [944,] "Spirochaetales"              "Spirochaetaceae"              
##  [945,] "Lactobacillales"             "Streptococcaceae"             
##  [946,] "Actinomycetales"             "Actinomycetaceae"             
##  [947,] "Lactobacillales"             "Streptococcaceae"             
##  [948,] "Fusobacteriales"             "Leptotrichiaceae"             
##  [949,] "Fusobacteriales"             "Leptotrichiaceae"             
##  [950,] "Clostridiales"               "Lachnospiraceae"              
##  [951,] "Fusobacteriales"             "Fusobacteriaceae"             
##  [952,] "Selenomonadales"             "Veillonellaceae"              
##  [953,] NA                            NA                             
##  [954,] "Spirochaetales"              "Spirochaetaceae"              
##  [955,] "Bacteroidales"               "Prevotellaceae"               
##  [956,] "Micrococcales"               "Micrococcaceae"               
##  [957,] "Selenomonadales"             "Veillonellaceae"              
##  [958,] "Saccharimonadales"           "Saccharimonadaceae"           
##  [959,] "Lactobacillales"             "Streptococcaceae"             
##  [960,] "Bacteroidales"               "Porphyromonadaceae"           
##  [961,] "Selenomonadales"             "Veillonellaceae"              
##  [962,] "Lactobacillales"             "Streptococcaceae"             
##  [963,] "Selenomonadales"             "Veillonellaceae"              
##  [964,] "Enterobacteriales"           "Enterobacteriaceae"           
##  [965,] "Fusobacteriales"             "Fusobacteriaceae"             
##  [966,] "Clostridiales"               "Lachnospiraceae"              
##  [967,] "Bacteroidales"               "Prevotellaceae"               
##  [968,] "Fusobacteriales"             "Fusobacteriaceae"             
##  [969,] "Clostridiales"               "Ruminococcaceae"              
##  [970,] "Bacteroidales"               "Prevotellaceae"               
##  [971,] "Fusobacteriales"             "Fusobacteriaceae"             
##  [972,] "Selenomonadales"             "Veillonellaceae"              
##  [973,] "Saccharimonadales"           NA                             
##  [974,] "Selenomonadales"             "Veillonellaceae"              
##  [975,] "Bacteroidales"               "Prevotellaceae"               
##  [976,] "Clostridiales"               "Lachnospiraceae"              
##  [977,] "Clostridiales"               "Lachnospiraceae"              
##  [978,] "Selenomonadales"             "Veillonellaceae"              
##  [979,] "Actinomycetales"             "Actinomycetaceae"             
##  [980,] "Actinomycetales"             "Actinomycetaceae"             
##  [981,] "Fusobacteriales"             "Fusobacteriaceae"             
##  [982,] "Bacteroidales"               "Prevotellaceae"               
##  [983,] "Fusobacteriales"             "Leptotrichiaceae"             
##  [984,] "Actinomycetales"             "Actinomycetaceae"             
##  [985,] "Campylobacterales"           NA                             
##  [986,] "Selenomonadales"             "Veillonellaceae"              
##  [987,] "Actinomycetales"             "Actinomycetaceae"             
##  [988,] "Pasteurellales"              "Pasteurellaceae"              
##  [989,] "Pasteurellales"              "Pasteurellaceae"              
##  [990,] "Selenomonadales"             "Veillonellaceae"              
##  [991,] "Bacteroidales"               "Prevotellaceae"               
##  [992,] "Micrococcales"               NA                             
##  [993,] "Actinomycetales"             "Actinomycetaceae"             
##  [994,] "Selenomonadales"             "Veillonellaceae"              
##  [995,] "Selenomonadales"             "Veillonellaceae"              
##  [996,] "Fusobacteriales"             "Leptotrichiaceae"             
##  [997,] "Fusobacteriales"             "Leptotrichiaceae"             
##  [998,] "Saccharimonadales"           "Saccharimonadaceae"           
##  [999,] "Bacteroidales"               "Prevotellaceae"               
## [1000,] "Selenomonadales"             "Veillonellaceae"              
## [1001,] "Selenomonadales"             "Veillonellaceae"              
## [1002,] "Selenomonadales"             "Veillonellaceae"              
## [1003,] "Betaproteobacteriales"       "Neisseriaceae"                
## [1004,] "Bacteroidales"               "Porphyromonadaceae"           
## [1005,] "Bacteroidales"               "Prevotellaceae"               
## [1006,] "Bacteroidales"               "Prevotellaceae"               
## [1007,] "Clostridiales"               "Lachnospiraceae"              
## [1008,] "Bacteroidales"               "Prevotellaceae"               
## [1009,] "Bacteroidales"               "Prevotellaceae"               
## [1010,] "Clostridiales"               "Family_XI"                    
## [1011,] "Selenomonadales"             "Veillonellaceae"              
## [1012,] "Saccharimonadales"           "Saccharimonadaceae"           
## [1013,] "Betaproteobacteriales"       "Neisseriaceae"                
## [1014,] "Saccharimonadales"           NA                             
## [1015,] "Clostridiales"               "Lachnospiraceae"              
## [1016,] "Betaproteobacteriales"       "Neisseriaceae"                
## [1017,] "Saccharimonadales"           "Saccharimonadaceae"           
## [1018,] "Bacteroidales"               "Prevotellaceae"               
## [1019,] "Fusobacteriales"             "Fusobacteriaceae"             
## [1020,] "Enterobacteriales"           "Enterobacteriaceae"           
## [1021,] "Bacteroidales"               "Prevotellaceae"               
## [1022,] "Fusobacteriales"             "Leptotrichiaceae"             
## [1023,] "Spirochaetales"              "Spirochaetaceae"              
## [1024,] "Lactobacillales"             "Carnobacteriaceae"            
## [1025,] "Lactobacillales"             "Carnobacteriaceae"            
## [1026,] "Corynebacteriales"           "Corynebacteriaceae"           
## [1027,] "Fusobacteriales"             "Leptotrichiaceae"             
## [1028,] "Bacteroidales"               "Prevotellaceae"               
## [1029,] "Bacteroidales"               "Prevotellaceae"               
## [1030,] "Betaproteobacteriales"       "Neisseriaceae"                
## [1031,] "Fusobacteriales"             "Fusobacteriaceae"             
## [1032,] "Fusobacteriales"             "Leptotrichiaceae"             
## [1033,] "Fusobacteriales"             "Fusobacteriaceae"             
## [1034,] "Selenomonadales"             "Veillonellaceae"              
## [1035,] "Bacteroidales"               "Prevotellaceae"               
## [1036,] "Selenomonadales"             "Veillonellaceae"              
## [1037,] "Selenomonadales"             "Veillonellaceae"              
## [1038,] "Clostridiales"               "Lachnospiraceae"              
## [1039,] "Bacteroidales"               "Prevotellaceae"               
## [1040,] "Selenomonadales"             "Veillonellaceae"              
## [1041,] "Bacteroidales"               "Prevotellaceae"               
## [1042,] "Clostridiales"               "Lachnospiraceae"              
## [1043,] "Fusobacteriales"             "Leptotrichiaceae"             
## [1044,] "Selenomonadales"             "Veillonellaceae"              
## [1045,] "Actinomycetales"             "Actinomycetaceae"             
## [1046,] "Bacteroidales"               "Prevotellaceae"               
## [1047,] "Selenomonadales"             "Veillonellaceae"              
## [1048,] "Bacteroidales"               "Prevotellaceae"               
## [1049,] "Selenomonadales"             "Veillonellaceae"              
## [1050,] "Fusobacteriales"             NA                             
## [1051,] "Clostridiales"               "Lachnospiraceae"              
## [1052,] "Lactobacillales"             "Carnobacteriaceae"            
## [1053,] "Bacteroidales"               "Prevotellaceae"               
## [1054,] "Pasteurellales"              "Pasteurellaceae"              
## [1055,] "Bacteroidales"               "Prevotellaceae"               
## [1056,] "Lactobacillales"             "Streptococcaceae"             
## [1057,] "Pasteurellales"              "Pasteurellaceae"              
## [1058,] "Bacteroidales"               "Prevotellaceae"               
## [1059,] "Clostridiales"               "Lachnospiraceae"              
## [1060,] "Bacteroidales"               "Prevotellaceae"               
## [1061,] "Selenomonadales"             "Veillonellaceae"              
## [1062,] "Lactobacillales"             "Streptococcaceae"             
## [1063,] "Bacteroidales"               "Prevotellaceae"               
## [1064,] "Fusobacteriales"             "Fusobacteriaceae"             
## [1065,] "Selenomonadales"             "Veillonellaceae"              
## [1066,] "Fusobacteriales"             "Leptotrichiaceae"             
## [1067,] "Selenomonadales"             "Veillonellaceae"              
## [1068,] "Pasteurellales"              "Pasteurellaceae"              
## [1069,] "Fusobacteriales"             "Leptotrichiaceae"             
## [1070,] "Selenomonadales"             "Veillonellaceae"              
## [1071,] "Pasteurellales"              "Pasteurellaceae"              
## [1072,] "Bacteroidales"               "Porphyromonadaceae"           
## [1073,] "Bacteroidales"               "Prevotellaceae"               
## [1074,] "Coriobacteriales"            "Atopobiaceae"                 
## [1075,] "Flavobacteriales"            "Flavobacteriaceae"            
## [1076,] "Betaproteobacteriales"       "Neisseriaceae"                
## [1077,] NA                            NA                             
## [1078,] "Bacteroidales"               "Prevotellaceae"               
## [1079,] "Selenomonadales"             "Veillonellaceae"              
## [1080,] "Clostridiales"               "Lachnospiraceae"              
## [1081,] "Bacteroidales"               "Prevotellaceae"               
## [1082,] "Saccharimonadales"           "Saccharimonadaceae"           
## [1083,] "Flavobacteriales"            "Flavobacteriaceae"            
## [1084,] "Selenomonadales"             "Veillonellaceae"              
## [1085,] "Selenomonadales"             "Veillonellaceae"              
## [1086,] NA                            NA                             
## [1087,] "Lactobacillales"             "Streptococcaceae"             
## [1088,] "Bacteroidales"               "Prevotellaceae"               
## [1089,] "Selenomonadales"             "Veillonellaceae"              
## [1090,] "Bacteroidales"               "Prevotellaceae"               
## [1091,] "Coriobacteriales"            "Atopobiaceae"                 
## [1092,] "Bacteroidales"               "Prevotellaceae"               
## [1093,] "Bacteroidales"               "Prevotellaceae"               
## [1094,] "Bacteroidales"               "Prevotellaceae"               
## [1095,] "Flavobacteriales"            "Flavobacteriaceae"            
## [1096,] "Selenomonadales"             "Veillonellaceae"              
## [1097,] "Bacteroidales"               "Porphyromonadaceae"           
## [1098,] "Lactobacillales"             "Streptococcaceae"             
## [1099,] "Actinomycetales"             "Actinomycetaceae"             
## [1100,] "Bacteroidales"               "Prevotellaceae"               
## [1101,] "Saccharimonadales"           "Saccharimonadaceae"           
## [1102,] "Selenomonadales"             "Veillonellaceae"              
## [1103,] "Selenomonadales"             "Veillonellaceae"              
## [1104,] "Bacteroidales"               "Prevotellaceae"               
## [1105,] "Bacteroidales"               "Prevotellaceae"               
## [1106,] "Selenomonadales"             "Veillonellaceae"              
## [1107,] "Bacteroidales"               "Prevotellaceae"               
## [1108,] "Fusobacteriales"             "Fusobacteriaceae"             
## [1109,] "Flavobacteriales"            "Flavobacteriaceae"            
## [1110,] "Selenomonadales"             "Veillonellaceae"              
## [1111,] "Bacteroidales"               "Prevotellaceae"               
## [1112,] "Fusobacteriales"             "Leptotrichiaceae"             
## [1113,] "Bacteroidales"               "Prevotellaceae"               
## [1114,] "Clostridiales"               "Peptostreptococcaceae"        
## [1115,] "Saccharimonadales"           "Saccharimonadaceae"           
## [1116,] "Saccharimonadales"           "Saccharimonadaceae"           
## [1117,] "Flavobacteriales"            "Flavobacteriaceae"            
## [1118,] "Bacteroidales"               "Prevotellaceae"               
## [1119,] "Micrococcales"               "Micrococcaceae"               
## [1120,] "Saccharimonadales"           "Saccharimonadaceae"           
## [1121,] "Selenomonadales"             "Veillonellaceae"              
## [1122,] "Selenomonadales"             "Veillonellaceae"              
## [1123,] "Bacteroidales"               "Prevotellaceae"               
## [1124,] "Campylobacterales"           "Campylobacteraceae"           
## [1125,] "Actinomycetales"             "Actinomycetaceae"             
## [1126,] "Chloroplast"                 NA                             
## [1127,] "Bacteroidales"               "Porphyromonadaceae"           
## [1128,] NA                            NA                             
## [1129,] "Fusobacteriales"             "Fusobacteriaceae"             
## [1130,] "Pasteurellales"              "Pasteurellaceae"              
## [1131,] "Clostridiales"               "Lachnospiraceae"              
## [1132,] "Bacteroidales"               "Prevotellaceae"               
## [1133,] "Bacteroidales"               "Prevotellaceae"               
## [1134,] "Selenomonadales"             "Veillonellaceae"              
## [1135,] "Pasteurellales"              "Pasteurellaceae"              
## [1136,] "Pasteurellales"              "Pasteurellaceae"              
## [1137,] "Bacteroidales"               "Prevotellaceae"               
## [1138,] "Bacteroidales"               "Prevotellaceae"               
## [1139,] "Betaproteobacteriales"       "Neisseriaceae"                
## [1140,] "Lactobacillales"             NA                             
## [1141,] "Pseudomonadales"             "Moraxellaceae"                
## [1142,] "Bacteroidales"               "Prevotellaceae"               
## [1143,] "Clostridiales"               "Lachnospiraceae"              
## [1144,] "Lactobacillales"             "Streptococcaceae"             
## [1145,] "Actinomycetales"             "Actinomycetaceae"             
## [1146,] "Lactobacillales"             "Streptococcaceae"             
## [1147,] "Bacteroidales"               "Prevotellaceae"               
## [1148,] "Selenomonadales"             "Veillonellaceae"              
## [1149,] "Clostridiales"               "Lachnospiraceae"              
## [1150,] "Micrococcales"               NA                             
## [1151,] "Fusobacteriales"             "Leptotrichiaceae"             
## [1152,] "Pasteurellales"              "Pasteurellaceae"              
## [1153,] "Lactobacillales"             "Streptococcaceae"             
## [1154,] "Betaproteobacteriales"       "Neisseriaceae"                
## [1155,] "Bacteroidales"               "Prevotellaceae"               
## [1156,] "Pasteurellales"              "Pasteurellaceae"              
## [1157,] "Lactobacillales"             "Streptococcaceae"             
## [1158,] "Fusobacteriales"             "Fusobacteriaceae"             
## [1159,] "Clostridiales"               "Lachnospiraceae"              
## [1160,] "Bacteroidales"               "Prevotellaceae"               
## [1161,] "Bacteroidales"               "Prevotellaceae"               
## [1162,] "Lactobacillales"             "Streptococcaceae"             
## [1163,] "Mycoplasmatales"             "Mycoplasmataceae"             
## [1164,] "Pasteurellales"              "Pasteurellaceae"              
## [1165,] "Absconditabacteriales_(SR1)" NA                             
## [1166,] "Selenomonadales"             "Veillonellaceae"              
## [1167,] "Betaproteobacteriales"       "Neisseriaceae"                
## [1168,] "Selenomonadales"             "Veillonellaceae"              
## [1169,] "Selenomonadales"             "Veillonellaceae"              
## [1170,] "Lactobacillales"             "Streptococcaceae"             
## [1171,] "Bacteroidales"               "Prevotellaceae"               
## [1172,] "Saccharimonadales"           "Saccharimonadaceae"           
## [1173,] "Bacteroidales"               "Prevotellaceae"               
## [1174,] "Selenomonadales"             "Veillonellaceae"              
## [1175,] "Selenomonadales"             "Veillonellaceae"              
## [1176,] "Selenomonadales"             "Veillonellaceae"              
## [1177,] "Bacteroidales"               "Porphyromonadaceae"           
## [1178,] "Fusobacteriales"             "Leptotrichiaceae"             
## [1179,] "Betaproteobacteriales"       "Neisseriaceae"                
## [1180,] "Bacteroidales"               "Prevotellaceae"               
## [1181,] "Clostridiales"               "Lachnospiraceae"              
## [1182,] "Clostridiales"               "Peptostreptococcaceae"        
## [1183,] "Lactobacillales"             "Streptococcaceae"             
## [1184,] "Bacteroidales"               "Prevotellaceae"               
## [1185,] "Pasteurellales"              "Pasteurellaceae"              
## [1186,] "Pasteurellales"              "Pasteurellaceae"              
## [1187,] "Bacteroidales"               "Prevotellaceae"               
## [1188,] "Selenomonadales"             "Veillonellaceae"              
## [1189,] "Bacteroidales"               "Prevotellaceae"               
## [1190,] "Bacteroidales"               "Prevotellaceae"               
## [1191,] "Actinomycetales"             "Actinomycetaceae"             
## [1192,] "Fusobacteriales"             "Leptotrichiaceae"             
## [1193,] "Actinomycetales"             "Actinomycetaceae"             
## [1194,] "Pasteurellales"              "Pasteurellaceae"              
## [1195,] "Bacteroidales"               "Porphyromonadaceae"           
## [1196,] "Flavobacteriales"            "Flavobacteriaceae"            
## [1197,] "Selenomonadales"             "Veillonellaceae"              
## [1198,] "Clostridiales"               "Lachnospiraceae"              
## [1199,] "Lactobacillales"             "Streptococcaceae"             
## [1200,] "Bacteroidales"               "Prevotellaceae"               
## [1201,] "Selenomonadales"             "Veillonellaceae"              
## [1202,] "Fusobacteriales"             "Fusobacteriaceae"             
## [1203,] "Actinomycetales"             "Actinomycetaceae"             
## [1204,] "Bacteroidales"               "Prevotellaceae"               
## [1205,] "Synergistales"               "Synergistaceae"               
## [1206,] "Campylobacterales"           NA                             
## [1207,] "Betaproteobacteriales"       "Burkholderiaceae"             
## [1208,] "Selenomonadales"             "Veillonellaceae"              
## [1209,] "Rhizobiales"                 "Rhizobiaceae"                 
## [1210,] "Selenomonadales"             "Veillonellaceae"              
## [1211,] "Fusobacteriales"             "Fusobacteriaceae"             
## [1212,] "Saccharimonadales"           "Saccharimonadaceae"           
## [1213,] "Selenomonadales"             "Veillonellaceae"              
## [1214,] "Fusobacteriales"             "Fusobacteriaceae"             
## [1215,] "Saccharimonadales"           "Saccharimonadaceae"           
## [1216,] "Clostridiales"               "Lachnospiraceae"              
## [1217,] "Spirochaetales"              "Spirochaetaceae"              
## [1218,] "Bacteroidales"               "Prevotellaceae"               
## [1219,] "Clostridiales"               "Peptostreptococcaceae"        
## [1220,] "Clostridiales"               "Peptostreptococcaceae"        
## [1221,] "Selenomonadales"             "Veillonellaceae"              
## [1222,] "Actinomycetales"             "Actinomycetaceae"             
## [1223,] "Bacteroidales"               "Prevotellaceae"               
## [1224,] NA                            NA                             
## [1225,] "Selenomonadales"             "Veillonellaceae"              
## [1226,] NA                            NA                             
## [1227,] "Selenomonadales"             "Veillonellaceae"              
## [1228,] "Betaproteobacteriales"       "Burkholderiaceae"             
## [1229,] "Bacteroidales"               "Prevotellaceae"               
## [1230,] "Selenomonadales"             "Veillonellaceae"              
## [1231,] "Bacteroidales"               "Porphyromonadaceae"           
## [1232,] "Bacteroidales"               "Prevotellaceae"               
## [1233,] "Flavobacteriales"            "Weeksellaceae"                
## [1234,] "Selenomonadales"             "Veillonellaceae"              
## [1235,] "Bacteroidales"               "Porphyromonadaceae"           
## [1236,] "Clostridiales"               "Lachnospiraceae"              
## [1237,] "Fusobacteriales"             "Fusobacteriaceae"             
## [1238,] "Micrococcales"               "Micrococcaceae"               
## [1239,] "Selenomonadales"             "Veillonellaceae"              
## [1240,] "Saccharimonadales"           "Saccharimonadaceae"           
## [1241,] "Bacteroidales"               "Porphyromonadaceae"           
## [1242,] "Bacteroidales"               "Porphyromonadaceae"           
## [1243,] "Clostridiales"               "Peptostreptococcaceae"        
## [1244,] "Selenomonadales"             "Veillonellaceae"              
## [1245,] "Selenomonadales"             "Veillonellaceae"              
## [1246,] "Fusobacteriales"             "Leptotrichiaceae"             
## [1247,] NA                            NA                             
## [1248,] "Bacteroidales"               "Prevotellaceae"               
## [1249,] "Selenomonadales"             "Veillonellaceae"              
## [1250,] "Micrococcales"               NA                             
## [1251,] "Fusobacteriales"             NA                             
## [1252,] "Bacteroidales"               "Prevotellaceae"               
## [1253,] "Bacteroidales"               "Prevotellaceae"               
## [1254,] "Micrococcales"               NA                             
## [1255,] "Bacteroidales"               "Prevotellaceae"               
## [1256,] "Bacteroidales"               "Paludibacteraceae"            
## [1257,] "Selenomonadales"             "Veillonellaceae"              
## [1258,] "Bacteroidales"               "Prevotellaceae"               
## [1259,] "Lactobacillales"             "Streptococcaceae"             
## [1260,] "Bacteroidales"               "Prevotellaceae"               
## [1261,] "Betaproteobacteriales"       "Neisseriaceae"                
## [1262,] "Selenomonadales"             "Veillonellaceae"              
## [1263,] "Saccharimonadales"           "Saccharimonadaceae"           
## [1264,] NA                            NA                             
## [1265,] "Actinomycetales"             "Actinomycetaceae"             
## [1266,] "Selenomonadales"             "Veillonellaceae"              
## [1267,] "Coriobacteriales"            "Atopobiaceae"                 
## [1268,] "Bacteroidales"               "Prevotellaceae"               
## [1269,] "Clostridiales"               "Family_XIII"                  
## [1270,] "Bacteroidales"               "Prevotellaceae"               
## [1271,] "Selenomonadales"             "Veillonellaceae"              
## [1272,] "Selenomonadales"             "Veillonellaceae"              
## [1273,] "Fusobacteriales"             "Fusobacteriaceae"             
## [1274,] "Bacteroidales"               "Prevotellaceae"               
## [1275,] "Spirochaetales"              "Spirochaetaceae"              
## [1276,] "Bacteroidales"               "Prevotellaceae"               
## [1277,] "Selenomonadales"             "Veillonellaceae"              
## [1278,] "Selenomonadales"             "Veillonellaceae"              
## [1279,] "Selenomonadales"             "Veillonellaceae"              
## [1280,] "Pasteurellales"              "Pasteurellaceae"              
## [1281,] "Pasteurellales"              "Pasteurellaceae"              
## [1282,] "Bacteroidales"               "Prevotellaceae"               
## [1283,] "Saccharimonadales"           "Saccharimonadaceae"           
## [1284,] "Selenomonadales"             "Veillonellaceae"              
## [1285,] "Fusobacteriales"             "Fusobacteriaceae"             
## [1286,] "Corynebacteriales"           "Corynebacteriaceae"           
## [1287,] "Selenomonadales"             "Veillonellaceae"              
## [1288,] "Clostridiales"               "Lachnospiraceae"              
## [1289,] "Mycoplasmatales"             "Mycoplasmataceae"             
## [1290,] "Fusobacteriales"             "Fusobacteriaceae"             
## [1291,] "Bacteroidales"               "Prevotellaceae"               
## [1292,] "Clostridiales"               "Ruminococcaceae"              
## [1293,] "Clostridiales"               "Lachnospiraceae"              
## [1294,] "Fusobacteriales"             "Fusobacteriaceae"             
## [1295,] "Betaproteobacteriales"       "Burkholderiaceae"             
## [1296,] "Bacteroidales"               "Prevotellaceae"               
## [1297,] "Clostridiales"               "Lachnospiraceae"              
## [1298,] "Flavobacteriales"            "Weeksellaceae"                
## [1299,] "Bacteroidales"               "Prevotellaceae"               
## [1300,] "Bacteroidales"               "Porphyromonadaceae"           
## [1301,] "Clostridiales"               "Lachnospiraceae"              
## [1302,] "Bacteroidales"               "Prevotellaceae"               
## [1303,] "Selenomonadales"             "Veillonellaceae"              
## [1304,] "Bacteroidales"               "Prevotellaceae"               
## [1305,] "Actinomycetales"             "Actinomycetaceae"             
## [1306,] "Lactobacillales"             "Streptococcaceae"             
## [1307,] "Betaproteobacteriales"       "Neisseriaceae"                
## [1308,] "Betaproteobacteriales"       "Neisseriaceae"                
## [1309,] "Spirochaetales"              "Spirochaetaceae"              
## [1310,] "Pasteurellales"              "Pasteurellaceae"              
## [1311,] "Pasteurellales"              "Pasteurellaceae"              
## [1312,] "Clostridiales"               "Family_XIII"                  
## [1313,] "Bacteroidales"               "Prevotellaceae"               
## [1314,] "Clostridiales"               "Lachnospiraceae"              
## [1315,] "Bacteroidales"               "Prevotellaceae"               
## [1316,] "Sphingobacteriales"          "Lentimicrobiaceae"            
## [1317,] "Campylobacterales"           NA                             
## [1318,] "Selenomonadales"             "Veillonellaceae"              
## [1319,] "Selenomonadales"             "Veillonellaceae"              
## [1320,] "Lactobacillales"             "Streptococcaceae"             
## [1321,] "Selenomonadales"             "Veillonellaceae"              
## [1322,] "Campylobacterales"           "Campylobacteraceae"           
## [1323,] "Selenomonadales"             "Veillonellaceae"              
## [1324,] "Lactobacillales"             "Streptococcaceae"             
## [1325,] "Bacteroidales"               "Prevotellaceae"               
## [1326,] "Pasteurellales"              "Pasteurellaceae"              
## [1327,] "Lactobacillales"             "Streptococcaceae"             
## [1328,] "Selenomonadales"             "Veillonellaceae"              
## [1329,] "Bacteroidales"               "Prevotellaceae"               
## [1330,] "Bacteroidales"               "Porphyromonadaceae"           
## [1331,] "Clostridiales"               "Lachnospiraceae"              
## [1332,] "Bacteroidales"               "Prevotellaceae"               
## [1333,] "Selenomonadales"             "Veillonellaceae"              
## [1334,] "Bacteroidales"               "Prevotellaceae"               
## [1335,] "Fusobacteriales"             "Fusobacteriaceae"             
## [1336,] "Fusobacteriales"             "Leptotrichiaceae"             
## [1337,] "Selenomonadales"             "Veillonellaceae"              
## [1338,] "Clostridiales"               "Lachnospiraceae"              
## [1339,] "Pasteurellales"              "Pasteurellaceae"              
## [1340,] "Selenomonadales"             "Veillonellaceae"              
## [1341,] NA                            NA                             
## [1342,] "Fusobacteriales"             "Leptotrichiaceae"             
## [1343,] "Bacteroidales"               "Prevotellaceae"               
## [1344,] "Saccharimonadales"           "Saccharimonadaceae"           
## [1345,] "Flavobacteriales"            "Flavobacteriaceae"            
## [1346,] "Saccharimonadales"           NA                             
## [1347,] "Bacteroidales"               "Prevotellaceae"               
## [1348,] "Bacteroidales"               "Prevotellaceae"               
## [1349,] "Betaproteobacteriales"       "Neisseriaceae"                
## [1350,] "Micrococcales"               NA                             
## [1351,] "Actinomycetales"             "Actinomycetaceae"             
## [1352,] "Actinomycetales"             "Actinomycetaceae"             
## [1353,] "Bacteroidales"               "Porphyromonadaceae"           
## [1354,] "Selenomonadales"             "Veillonellaceae"              
## [1355,] "Campylobacterales"           NA                             
## [1356,] NA                            NA                             
## [1357,] "Bacteroidales"               "Prevotellaceae"               
## [1358,] "Acholeplasmatales"           "Acholeplasmataceae"           
## [1359,] "Selenomonadales"             "Veillonellaceae"              
## [1360,] "Bacteroidales"               "Prevotellaceae"               
## [1361,] "Betaproteobacteriales"       "Neisseriaceae"                
## [1362,] NA                            NA                             
## [1363,] NA                            NA                             
## [1364,] "Micrococcales"               NA                             
## [1365,] "Bacteroidales"               "Prevotellaceae"               
## [1366,] "Bacteroidales"               "Prevotellaceae"               
## [1367,] "Bacteroidales"               "Porphyromonadaceae"           
## [1368,] "Pasteurellales"              "Pasteurellaceae"              
## [1369,] "Fusobacteriales"             "Fusobacteriaceae"             
## [1370,] "Clostridiales"               "Ruminococcaceae"              
## [1371,] "Bacteroidales"               "Prevotellaceae"               
## [1372,] "Fusobacteriales"             "Leptotrichiaceae"             
## [1373,] "Bacillales"                  "Family_XI"                    
## [1374,] "Betaproteobacteriales"       "Neisseriaceae"                
## [1375,] "Fusobacteriales"             "Fusobacteriaceae"             
## [1376,] "Betaproteobacteriales"       "Neisseriaceae"                
## [1377,] "Selenomonadales"             "Veillonellaceae"              
## [1378,] "Bacteroidales"               "Prevotellaceae"               
## [1379,] "Bacteroidales"               "Prevotellaceae"               
## [1380,] NA                            NA                             
## [1381,] "Clostridiales"               "Lachnospiraceae"              
## [1382,] "Bacteroidales"               "Prevotellaceae"               
## [1383,] "Selenomonadales"             "Veillonellaceae"              
## [1384,] "Lactobacillales"             "Streptococcaceae"             
## [1385,] "Selenomonadales"             "Veillonellaceae"              
## [1386,] "Lactobacillales"             "Streptococcaceae"             
## [1387,] "Fusobacteriales"             "Fusobacteriaceae"             
## [1388,] "Bacteroidales"               "Prevotellaceae"               
## [1389,] "Selenomonadales"             "Veillonellaceae"              
## [1390,] "Bacteroidales"               "Prevotellaceae"               
## [1391,] "Saccharimonadales"           "Saccharimonadaceae"           
## [1392,] "Bacteroidales"               "Prevotellaceae"               
## [1393,] "Fusobacteriales"             "Fusobacteriaceae"             
## [1394,] "Bacteroidales"               "Prevotellaceae"               
## [1395,] "Fusobacteriales"             "Leptotrichiaceae"             
## [1396,] NA                            NA                             
## [1397,] "Bacteroidales"               "Prevotellaceae"               
## [1398,] "Bacteroidales"               "Prevotellaceae"               
## [1399,] "Lactobacillales"             "Carnobacteriaceae"            
## [1400,] "Flavobacteriales"            "Weeksellaceae"                
## [1401,] "Fusobacteriales"             "Leptotrichiaceae"             
## [1402,] "Bacteroidales"               "Prevotellaceae"               
## [1403,] "Fusobacteriales"             "Fusobacteriaceae"             
## [1404,] "Bacteroidales"               "Prevotellaceae"               
## [1405,] "Campylobacterales"           "Campylobacteraceae"           
## [1406,] "Pasteurellales"              "Pasteurellaceae"              
## [1407,] "Fusobacteriales"             "Leptotrichiaceae"             
## [1408,] "Selenomonadales"             "Veillonellaceae"              
## [1409,] "Pasteurellales"              "Pasteurellaceae"              
## [1410,] "Fusobacteriales"             "Fusobacteriaceae"             
## [1411,] "Clostridiales"               "Ruminococcaceae"              
## [1412,] "Bacillales"                  "Staphylococcaceae"            
## [1413,] "Betaproteobacteriales"       NA                             
## [1414,] "Betaproteobacteriales"       "Neisseriaceae"                
## [1415,] "Bacteroidales"               "Prevotellaceae"               
## [1416,] "Micrococcales"               NA                             
## [1417,] "Spirochaetales"              "Spirochaetaceae"              
## [1418,] "Selenomonadales"             "Veillonellaceae"              
## [1419,] "Fusobacteriales"             "Leptotrichiaceae"             
## [1420,] "Fusobacteriales"             "Leptotrichiaceae"             
## [1421,] "Campylobacterales"           "P6-b51"                       
## [1422,] "Selenomonadales"             "Veillonellaceae"              
## [1423,] "Fusobacteriales"             "Fusobacteriaceae"             
## [1424,] "Bacteroidales"               "Prevotellaceae"               
## [1425,] "Selenomonadales"             "Veillonellaceae"              
## [1426,] "Campylobacterales"           "Campylobacteraceae"           
## [1427,] "Bacteroidales"               "Prevotellaceae"               
## [1428,] "Actinomycetales"             "Actinomycetaceae"             
## [1429,] "Bacteroidales"               "Prevotellaceae"               
## [1430,] "Bacteroidales"               "Prevotellaceae"               
## [1431,] "Selenomonadales"             "Veillonellaceae"              
## [1432,] "Spirochaetales"              "Spirochaetaceae"              
## [1433,] "Bacteroidales"               "Prevotellaceae"               
## [1434,] "Lactobacillales"             "Streptococcaceae"             
## [1435,] "Pasteurellales"              "Pasteurellaceae"              
## [1436,] "Saccharimonadales"           "Saccharimonadaceae"           
## [1437,] "Clostridiales"               "Family_XI"                    
## [1438,] "Selenomonadales"             "Veillonellaceae"              
## [1439,] "Pasteurellales"              "Pasteurellaceae"              
## [1440,] "Corynebacteriales"           "Corynebacteriaceae"           
## [1441,] "Selenomonadales"             "Veillonellaceae"              
## [1442,] "Selenomonadales"             "Veillonellaceae"              
## [1443,] "Bacteroidales"               "Prevotellaceae"               
## [1444,] "Flavobacteriales"            "Weeksellaceae"                
## [1445,] "Clostridiales"               "Lachnospiraceae"              
## [1446,] "Pasteurellales"              "Pasteurellaceae"              
## [1447,] NA                            NA                             
## [1448,] "Pasteurellales"              "Pasteurellaceae"              
## [1449,] "Bacteroidales"               "Prevotellaceae"               
## [1450,] "Lactobacillales"             "Streptococcaceae"             
## [1451,] "Lactobacillales"             "Streptococcaceae"             
## [1452,] "Actinomycetales"             "Actinomycetaceae"             
## [1453,] "Bacteroidales"               "Prevotellaceae"               
## [1454,] "Selenomonadales"             "Veillonellaceae"              
## [1455,] "Selenomonadales"             "Veillonellaceae"              
## [1456,] "Enterobacteriales"           "Enterobacteriaceae"           
## [1457,] "Bacteroidales"               "Prevotellaceae"               
## [1458,] "Saccharimonadales"           "Saccharimonadaceae"           
## [1459,] "Selenomonadales"             "Veillonellaceae"              
## [1460,] "Fusobacteriales"             "Fusobacteriaceae"             
## [1461,] "Bacteroidales"               "Porphyromonadaceae"           
## [1462,] "Bacteroidales"               "Prevotellaceae"               
## [1463,] "Spirochaetales"              "Spirochaetaceae"              
## [1464,] "Clostridiales"               "Lachnospiraceae"              
## [1465,] "Fusobacteriales"             "Leptotrichiaceae"             
## [1466,] "Micrococcales"               NA                             
## [1467,] "Bacteroidales"               "Rikenellaceae"                
## [1468,] "Bacteroidales"               NA                             
## [1469,] "Lactobacillales"             "Streptococcaceae"             
## [1470,] "Bacteroidales"               "Prevotellaceae"               
## [1471,] NA                            NA                             
## [1472,] "Fusobacteriales"             "Fusobacteriaceae"             
## [1473,] "Bacteroidales"               "Prevotellaceae"               
## [1474,] "Saccharimonadales"           NA                             
## [1475,] "Bacteroidales"               "Prevotellaceae"               
## [1476,] "Selenomonadales"             "Veillonellaceae"              
## [1477,] "Fusobacteriales"             "Leptotrichiaceae"             
## [1478,] "Clostridiales"               "Lachnospiraceae"              
## [1479,] "Pasteurellales"              "Pasteurellaceae"              
## [1480,] "Selenomonadales"             "Veillonellaceae"              
## [1481,] "Bacteroidales"               "Prevotellaceae"               
## [1482,] "Selenomonadales"             "Veillonellaceae"              
## [1483,] "Bacteroidales"               "Prevotellaceae"               
## [1484,] "Bacteroidales"               "Prevotellaceae"               
## [1485,] "Bacteroidales"               "Prevotellaceae"               
## [1486,] "Selenomonadales"             "Veillonellaceae"              
## [1487,] "Bacteroidales"               "Prevotellaceae"               
## [1488,] "Selenomonadales"             "Veillonellaceae"              
## [1489,] "Betaproteobacteriales"       "Neisseriaceae"                
## [1490,] "Actinomycetales"             "Actinomycetaceae"             
## [1491,] "Fusobacteriales"             "Leptotrichiaceae"             
## [1492,] "Clostridiales"               "Lachnospiraceae"              
## [1493,] "Clostridiales"               "Lachnospiraceae"              
## [1494,] "Selenomonadales"             "Veillonellaceae"              
## [1495,] "Pseudomonadales"             "Moraxellaceae"                
## [1496,] "Pasteurellales"              "Pasteurellaceae"              
## [1497,] "Bacteroidales"               "Prevotellaceae"               
## [1498,] "Pasteurellales"              "Pasteurellaceae"              
## [1499,] "Saccharimonadales"           "Saccharimonadaceae"           
## [1500,] "Lactobacillales"             "Streptococcaceae"             
## [1501,] "Corynebacteriales"           "Corynebacteriaceae"           
## [1502,] "Campylobacterales"           "Campylobacteraceae"           
## [1503,] "Flavobacteriales"            "Weeksellaceae"                
## [1504,] "Selenomonadales"             "Veillonellaceae"              
## [1505,] "Betaproteobacteriales"       "Neisseriaceae"                
## [1506,] "Saccharimonadales"           "Saccharimonadaceae"           
## [1507,] "Bacteroidales"               "Prevotellaceae"               
## [1508,] "Bacteroidales"               "Prevotellaceae"               
## [1509,] "Pasteurellales"              "Pasteurellaceae"              
## [1510,] "Fusobacteriales"             "Fusobacteriaceae"             
## [1511,] "Betaproteobacteriales"       "Neisseriaceae"                
## [1512,] "Bacteroidales"               "Prevotellaceae"               
## [1513,] "Flavobacteriales"            "Flavobacteriaceae"            
## [1514,] "Lactobacillales"             "Carnobacteriaceae"            
## [1515,] "Micrococcales"               NA                             
## [1516,] "Bacteroidales"               "Prevotellaceae"               
## [1517,] "Pasteurellales"              "Pasteurellaceae"              
## [1518,] "Clostridiales"               "Lachnospiraceae"              
## [1519,] "Lactobacillales"             "Carnobacteriaceae"            
## [1520,] "Bacteroidales"               "Prevotellaceae"               
## [1521,] "Fusobacteriales"             "Leptotrichiaceae"             
## [1522,] "Bacteroidales"               "Prevotellaceae"               
## [1523,] "Fusobacteriales"             "Fusobacteriaceae"             
## [1524,] "Betaproteobacteriales"       "Neisseriaceae"                
## [1525,] "Coriobacteriales"            "Atopobiaceae"                 
## [1526,] "Selenomonadales"             "Veillonellaceae"              
## [1527,] "Bacteroidales"               "Prevotellaceae"               
## [1528,] NA                            NA                             
## [1529,] "Bacteroidales"               "Prevotellaceae"               
## [1530,] "Clostridiales"               "Lachnospiraceae"              
## [1531,] "Campylobacterales"           "Campylobacteraceae"           
## [1532,] "Saccharimonadales"           "Saccharimonadaceae"           
## [1533,] "Fusobacteriales"             "Fusobacteriaceae"             
## [1534,] "Clostridiales"               "Lachnospiraceae"              
## [1535,] NA                            NA                             
## [1536,] "Bacteroidales"               "Prevotellaceae"               
## [1537,] "Bacteroidales"               "Prevotellaceae"               
## [1538,] "Absconditabacteriales_(SR1)" NA                             
## [1539,] "Actinomycetales"             "Actinomycetaceae"             
## [1540,] "Selenomonadales"             "Veillonellaceae"              
## [1541,] "Selenomonadales"             "Veillonellaceae"              
## [1542,] "Bacteroidales"               "Prevotellaceae"               
## [1543,] "Lactobacillales"             "Streptococcaceae"             
## [1544,] "Spirochaetales"              "Spirochaetaceae"              
## [1545,] "Clostridiales"               "Ruminococcaceae"              
## [1546,] "Selenomonadales"             "Veillonellaceae"              
## [1547,] "Bacteroidales"               "Prevotellaceae"               
## [1548,] "Micrococcales"               NA                             
## [1549,] "Fusobacteriales"             "Fusobacteriaceae"             
## [1550,] "Selenomonadales"             "Veillonellaceae"              
## [1551,] "Clostridiales"               "Peptostreptococcaceae"        
## [1552,] "Absconditabacteriales_(SR1)" NA                             
## [1553,] "Fusobacteriales"             "Leptotrichiaceae"             
## [1554,] "Selenomonadales"             "Veillonellaceae"              
## [1555,] "Bacteroidales"               "Prevotellaceae"               
## [1556,] "Actinomycetales"             "Actinomycetaceae"             
## [1557,] "Bacteroidales"               "Prevotellaceae"               
## [1558,] "Selenomonadales"             "Veillonellaceae"              
## [1559,] "Fusobacteriales"             "Fusobacteriaceae"             
## [1560,] "Chloroplast"                 NA                             
## [1561,] "Bacteroidales"               "Prevotellaceae"               
## [1562,] "Bacteroidales"               "Prevotellaceae"               
## [1563,] "Selenomonadales"             "Veillonellaceae"              
## [1564,] "Fusobacteriales"             "Leptotrichiaceae"             
## [1565,] "Bacteroidales"               "Prevotellaceae"               
## [1566,] "Clostridiales"               "Peptococcaceae"               
## [1567,] "Selenomonadales"             "Veillonellaceae"              
## [1568,] "Lactobacillales"             "Carnobacteriaceae"            
## [1569,] "Campylobacterales"           NA                             
## [1570,] "Bacteroidales"               "Rikenellaceae"                
## [1571,] "Bacteroidales"               "Prevotellaceae"               
## [1572,] "Bacteroidales"               "Prevotellaceae"               
## [1573,] "Selenomonadales"             "Veillonellaceae"              
## [1574,] "Fusobacteriales"             "Fusobacteriaceae"             
## [1575,] "Bacteroidales"               "Prevotellaceae"               
## [1576,] "Bacteroidales"               "Porphyromonadaceae"           
## [1577,] "Fusobacteriales"             "Fusobacteriaceae"             
## [1578,] "Bacteroidales"               "Prevotellaceae"               
## [1579,] "Bacteroidales"               "Prevotellaceae"               
## [1580,] "Bacteroidales"               "Prevotellaceae"               
## [1581,] "Bacteroidales"               "Prevotellaceae"               
## [1582,] "Selenomonadales"             "Veillonellaceae"              
## [1583,] "Lactobacillales"             "Streptococcaceae"             
## [1584,] "Spirochaetales"              "Spirochaetaceae"              
## [1585,] "Campylobacterales"           NA                             
## [1586,] "Bacteroidales"               "Prevotellaceae"               
## [1587,] "Selenomonadales"             "Veillonellaceae"              
## [1588,] "Selenomonadales"             "Veillonellaceae"              
## [1589,] "Sphingobacteriales"          "Lentimicrobiaceae"            
## [1590,] "Campylobacterales"           NA                             
## [1591,] "Bacteroidales"               "Prevotellaceae"               
## [1592,] "Lactobacillales"             "Carnobacteriaceae"            
## [1593,] "Pasteurellales"              "Pasteurellaceae"              
## [1594,] "Flavobacteriales"            "Flavobacteriaceae"            
## [1595,] "Pasteurellales"              "Pasteurellaceae"              
## [1596,] "Selenomonadales"             "Veillonellaceae"              
## [1597,] "Bacteroidales"               "Prevotellaceae"               
## [1598,] "Bacteroidales"               "Prevotellaceae"               
## [1599,] "Bacteroidales"               "Prevotellaceae"               
## [1600,] "Bacteroidales"               "Prevotellaceae"               
## [1601,] "Selenomonadales"             "Veillonellaceae"              
## [1602,] "Bacteroidales"               "Porphyromonadaceae"           
## [1603,] "Bacteroidales"               "Prevotellaceae"               
## [1604,] "Saccharimonadales"           "Saccharimonadaceae"           
## [1605,] "Saccharimonadales"           "Saccharimonadaceae"           
## [1606,] "Lactobacillales"             "Streptococcaceae"             
## [1607,] "Bacteroidales"               "Prevotellaceae"               
## [1608,] "Selenomonadales"             "Veillonellaceae"              
## [1609,] "Betaproteobacteriales"       "Neisseriaceae"                
## [1610,] "Bacteroidales"               "Prevotellaceae"               
## [1611,] "Fusobacteriales"             "Fusobacteriaceae"             
## [1612,] "Fusobacteriales"             "Leptotrichiaceae"             
## [1613,] "Pasteurellales"              "Pasteurellaceae"              
## [1614,] "Bacteroidales"               "Prevotellaceae"               
## [1615,] "Betaproteobacteriales"       "Neisseriaceae"                
## [1616,] "Chloroplast"                 NA                             
## [1617,] "Actinomycetales"             "Actinomycetaceae"             
## [1618,] "Selenomonadales"             "Veillonellaceae"              
## [1619,] "Selenomonadales"             "Veillonellaceae"              
## [1620,] "Fusobacteriales"             "Leptotrichiaceae"             
## [1621,] "Actinomycetales"             "Actinomycetaceae"             
## [1622,] "Pasteurellales"              "Pasteurellaceae"              
## [1623,] "Fusobacteriales"             "Fusobacteriaceae"             
## [1624,] "Bacteroidales"               "Porphyromonadaceae"           
## [1625,] "Fusobacteriales"             "Leptotrichiaceae"             
## [1626,] "Lactobacillales"             "Streptococcaceae"             
## [1627,] "Bacteroidales"               "Prevotellaceae"               
## [1628,] "Clostridiales"               "Lachnospiraceae"              
## [1629,] "Clostridiales"               "Lachnospiraceae"              
## [1630,] "Bacteroidales"               "Prevotellaceae"               
## [1631,] "Bacteroidales"               "Prevotellaceae"               
## [1632,] "Selenomonadales"             "Veillonellaceae"              
## [1633,] "Fusobacteriales"             "Leptotrichiaceae"             
## [1634,] "Selenomonadales"             "Veillonellaceae"              
## [1635,] "Bacteroidales"               "Prevotellaceae"               
## [1636,] "Pasteurellales"              "Pasteurellaceae"              
## [1637,] "Selenomonadales"             "Veillonellaceae"              
## [1638,] "Bacteroidales"               "Prevotellaceae"               
## [1639,] "Bacteroidales"               "Prevotellaceae"               
## [1640,] "Bacteroidales"               "Porphyromonadaceae"           
## [1641,] "Actinomycetales"             "Actinomycetaceae"             
## [1642,] "Bacteroidales"               "Prevotellaceae"               
## [1643,] "Clostridiales"               "Family_XIII"                  
## [1644,] NA                            NA                             
## [1645,] "Bacteroidales"               "Prevotellaceae"               
## [1646,] "Fusobacteriales"             NA                             
## [1647,] "Clostridiales"               "Lachnospiraceae"              
## [1648,] "Clostridiales"               "Ruminococcaceae"              
## [1649,] "Fusobacteriales"             "Leptotrichiaceae"             
## [1650,] "Fusobacteriales"             "Leptotrichiaceae"             
## [1651,] "Clostridiales"               "Lachnospiraceae"              
## [1652,] "Bacteroidales"               "Prevotellaceae"               
## [1653,] "Selenomonadales"             "Veillonellaceae"              
## [1654,] "Campylobacterales"           "Campylobacteraceae"           
## [1655,] "Saccharimonadales"           "Saccharimonadaceae"           
## [1656,] "Coriobacteriales"            "Eggerthellaceae"              
## [1657,] "Bacteroidales"               "Prevotellaceae"               
## [1658,] "Spirochaetales"              "Spirochaetaceae"              
## [1659,] "Pasteurellales"              "Pasteurellaceae"              
## [1660,] "Enterobacteriales"           "Enterobacteriaceae"           
## [1661,] "Bacteroidales"               "Prevotellaceae"               
## [1662,] "Bacteroidales"               "Prevotellaceae"               
## [1663,] "Selenomonadales"             "Veillonellaceae"              
## [1664,] "Bacteroidales"               "Porphyromonadaceae"           
## [1665,] "Bacteroidales"               "Prevotellaceae"               
## [1666,] "Fusobacteriales"             "Leptotrichiaceae"             
## [1667,] "Fusobacteriales"             "Leptotrichiaceae"             
## [1668,] "Clostridiales"               "Lachnospiraceae"              
## [1669,] "Spirochaetales"              "Spirochaetaceae"              
## [1670,] "Fusobacteriales"             "Leptotrichiaceae"             
## [1671,] NA                            NA                             
## [1672,] "Sphingobacteriales"          "Lentimicrobiaceae"            
## [1673,] "Pseudomonadales"             "Moraxellaceae"                
## [1674,] "Pasteurellales"              "Pasteurellaceae"              
## [1675,] "Bacteroidales"               "Prevotellaceae"               
## [1676,] "Clostridiales"               "Lachnospiraceae"              
## [1677,] "Lactobacillales"             "Streptococcaceae"             
## [1678,] NA                            NA                             
## [1679,] "Pasteurellales"              "Pasteurellaceae"              
## [1680,] "Saccharimonadales"           "Saccharimonadaceae"           
## [1681,] "Bacteroidales"               "Prevotellaceae"               
## [1682,] "Bacteroidales"               "Porphyromonadaceae"           
## [1683,] "Lactobacillales"             NA                             
## [1684,] "Pasteurellales"              "Pasteurellaceae"              
## [1685,] "Selenomonadales"             "Veillonellaceae"              
## [1686,] "Lactobacillales"             "Streptococcaceae"             
## [1687,] "Bacteroidales"               "Prevotellaceae"               
## [1688,] "Selenomonadales"             "Veillonellaceae"              
## [1689,] "Fusobacteriales"             "Fusobacteriaceae"             
## [1690,] "Pasteurellales"              "Pasteurellaceae"              
## [1691,] "Fusobacteriales"             "Fusobacteriaceae"             
## [1692,] "Bacteroidales"               "Prevotellaceae"               
## [1693,] "Fusobacteriales"             "Leptotrichiaceae"             
## [1694,] "Bacteroidales"               "Prevotellaceae"               
## [1695,] "Bacteroidales"               "Prevotellaceae"               
## [1696,] "Actinomycetales"             "Actinomycetaceae"             
## [1697,] "Bacteroidales"               "Prevotellaceae"               
## [1698,] "Campylobacterales"           "Campylobacteraceae"           
## [1699,] "Pasteurellales"              "Pasteurellaceae"              
## [1700,] "Actinomycetales"             "Actinomycetaceae"             
## [1701,] "Saccharimonadales"           NA                             
## [1702,] "Fusobacteriales"             "Fusobacteriaceae"             
## [1703,] "Lactobacillales"             "Streptococcaceae"             
## [1704,] "Pasteurellales"              "Pasteurellaceae"              
## [1705,] "Bacteroidales"               "Prevotellaceae"               
## [1706,] "Betaproteobacteriales"       "Neisseriaceae"                
## [1707,] "Lactobacillales"             "Streptococcaceae"             
## [1708,] "Selenomonadales"             "Veillonellaceae"              
## [1709,] "Lactobacillales"             "Streptococcaceae"             
## [1710,] "Bacteroidales"               "Prevotellaceae"               
## [1711,] "Selenomonadales"             "Veillonellaceae"              
## [1712,] "Selenomonadales"             "Veillonellaceae"              
## [1713,] "Clostridiales"               "Lachnospiraceae"              
## [1714,] "Selenomonadales"             "Veillonellaceae"              
## [1715,] "Bacteroidales"               "Prevotellaceae"               
## [1716,] "Fusobacteriales"             "Leptotrichiaceae"             
## [1717,] "Pasteurellales"              "Pasteurellaceae"              
## [1718,] "Flavobacteriales"            "Flavobacteriaceae"            
## [1719,] "Clostridiales"               "Lachnospiraceae"              
## [1720,] "Betaproteobacteriales"       "Neisseriaceae"                
## [1721,] "Bacteroidales"               "Prevotellaceae"               
## [1722,] "Flavobacteriales"            "Weeksellaceae"                
## [1723,] "Selenomonadales"             "Veillonellaceae"              
## [1724,] "Bacteroidales"               "Prevotellaceae"               
## [1725,] "Pasteurellales"              "Pasteurellaceae"              
## [1726,] "Bacteroidales"               "Prevotellaceae"               
## [1727,] "Coriobacteriales"            "Atopobiaceae"                 
## [1728,] "Bacteroidales"               "Prevotellaceae"               
## [1729,] "Bacteroidales"               "Prevotellaceae"               
## [1730,] "Fusobacteriales"             "Fusobacteriaceae"             
## [1731,] "Selenomonadales"             "Veillonellaceae"              
## [1732,] "Corynebacteriales"           "Corynebacteriaceae"           
## [1733,] "Bacteroidales"               "Prevotellaceae"               
## [1734,] "Sphingobacteriales"          "Lentimicrobiaceae"            
## [1735,] "Bacteroidales"               "Prevotellaceae"               
## [1736,] "Bacteroidales"               "Prevotellaceae"               
## [1737,] "Clostridiales"               "Defluviitaleaceae"            
## [1738,] "Bacteroidales"               "Prevotellaceae"               
## [1739,] "Bacteroidales"               "Prevotellaceae"               
## [1740,] "Bacteroidales"               "Prevotellaceae"               
## [1741,] "Selenomonadales"             "Veillonellaceae"              
## [1742,] "Bacteroidales"               "Prevotellaceae"               
## [1743,] "Lactobacillales"             "Streptococcaceae"             
## [1744,] "Bacteroidales"               "Prevotellaceae"               
## [1745,] "Betaproteobacteriales"       "Neisseriaceae"                
## [1746,] "Bacteroidales"               "Porphyromonadaceae"           
## [1747,] "Bacteroidales"               "Porphyromonadaceae"           
## [1748,] "Fusobacteriales"             "Fusobacteriaceae"             
## [1749,] "Bacteroidales"               "Prevotellaceae"               
## [1750,] "Bacteroidales"               "Prevotellaceae"               
## [1751,] "Bacillales"                  "Family_XI"                    
## [1752,] "Selenomonadales"             "Veillonellaceae"              
## [1753,] "Bacteroidales"               "Prevotellaceae"               
## [1754,] "Bacteroidales"               "Prevotellaceae"               
## [1755,] "Selenomonadales"             "Veillonellaceae"              
## [1756,] "Lactobacillales"             "Carnobacteriaceae"            
## [1757,] "Micrococcales"               NA                             
## [1758,] "Bacteroidales"               "Prevotellaceae"               
## [1759,] "Bacteroidales"               "Prevotellaceae"               
## [1760,] "Saccharimonadales"           "Saccharimonadaceae"           
## [1761,] "Bacteroidales"               "Porphyromonadaceae"           
## [1762,] "Lactobacillales"             "Carnobacteriaceae"            
## [1763,] "Lactobacillales"             "Streptococcaceae"             
## [1764,] "Selenomonadales"             "Veillonellaceae"              
## [1765,] "Clostridiales"               "Lachnospiraceae"              
## [1766,] "Bacteroidales"               "Prevotellaceae"               
## [1767,] "Clostridiales"               "Ruminococcaceae"              
## [1768,] "Bacteroidales"               "Prevotellaceae"               
## [1769,] "Selenomonadales"             "Veillonellaceae"              
## [1770,] "Bacteroidales"               "Porphyromonadaceae"           
## [1771,] "Selenomonadales"             "Veillonellaceae"              
## [1772,] "Cardiobacteriales"           "Cardiobacteriaceae"           
## [1773,] "Clostridiales"               "Lachnospiraceae"              
## [1774,] "Actinomycetales"             "Actinomycetaceae"             
## [1775,] "Clostridiales"               "Ruminococcaceae"              
## [1776,] "Bacillales"                  "Family_XI"                    
## [1777,] "Pasteurellales"              "Pasteurellaceae"              
## [1778,] "Betaproteobacteriales"       "Neisseriaceae"                
## [1779,] "Fusobacteriales"             "Fusobacteriaceae"             
## [1780,] "Bacteroidales"               "Prevotellaceae"               
## [1781,] "Clostridiales"               "Lachnospiraceae"              
## [1782,] "Pasteurellales"              "Pasteurellaceae"              
## [1783,] "Micrococcales"               NA                             
## [1784,] "Bacteroidales"               "Prevotellaceae"               
## [1785,] "Bacteroidales"               "Prevotellaceae"               
## [1786,] "Bacteroidales"               "Prevotellaceae"               
## [1787,] "Selenomonadales"             "Veillonellaceae"              
## [1788,] "Bacteroidales"               "Prevotellaceae"               
## [1789,] "Pasteurellales"              "Pasteurellaceae"              
## [1790,] "Fusobacteriales"             "Leptotrichiaceae"             
## [1791,] "Pasteurellales"              "Pasteurellaceae"              
## [1792,] NA                            NA                             
## [1793,] NA                            NA                             
## [1794,] "Clostridiales"               "Peptostreptococcaceae"        
## [1795,] NA                            NA                             
## [1796,] "Bacteroidales"               "Porphyromonadaceae"           
## [1797,] "Bacteroidales"               "Prevotellaceae"               
## [1798,] "Clostridiales"               "Lachnospiraceae"              
## [1799,] "Lactobacillales"             "Streptococcaceae"             
## [1800,] "Clostridiales"               "Family_XIII"                  
## [1801,] "Clostridiales"               "Lachnospiraceae"              
## [1802,] "Selenomonadales"             "Veillonellaceae"              
## [1803,] "Fusobacteriales"             "Fusobacteriaceae"             
## [1804,] "Clostridiales"               "Lachnospiraceae"              
## [1805,] "Fusobacteriales"             "Fusobacteriaceae"             
## [1806,] "Flavobacteriales"            "Flavobacteriaceae"            
## [1807,] "Fusobacteriales"             "Fusobacteriaceae"             
## [1808,] "Lactobacillales"             "Carnobacteriaceae"            
## [1809,] "Micrococcales"               "Micrococcaceae"               
## [1810,] "Actinomycetales"             "Actinomycetaceae"             
## [1811,] "Selenomonadales"             "Veillonellaceae"              
## [1812,] "Selenomonadales"             "Veillonellaceae"              
## [1813,] "Selenomonadales"             "Veillonellaceae"              
## [1814,] "Fusobacteriales"             "Fusobacteriaceae"             
## [1815,] "Betaproteobacteriales"       "Neisseriaceae"                
## [1816,] "Pasteurellales"              "Pasteurellaceae"              
## [1817,] "Enterobacteriales"           "Enterobacteriaceae"           
## [1818,] "Bacteroidales"               "Prevotellaceae"               
## [1819,] "Bacteroidales"               "Porphyromonadaceae"           
## [1820,] "Bacteroidales"               "Prevotellaceae"               
## [1821,] "Betaproteobacteriales"       "Neisseriaceae"                
## [1822,] "Bacteroidales"               "Prevotellaceae"               
## [1823,] "Selenomonadales"             "Veillonellaceae"              
## [1824,] "Selenomonadales"             "Veillonellaceae"              
## [1825,] "Lactobacillales"             "Streptococcaceae"             
## [1826,] "Selenomonadales"             "Veillonellaceae"              
## [1827,] "Bacteroidales"               "Porphyromonadaceae"           
## [1828,] "Pasteurellales"              "Pasteurellaceae"              
## [1829,] "Clostridiales"               "Lachnospiraceae"              
## [1830,] "Fusobacteriales"             "Leptotrichiaceae"             
## [1831,] "Fusobacteriales"             "Leptotrichiaceae"             
## [1832,] "Bacteroidales"               "Prevotellaceae"               
## [1833,] "Corynebacteriales"           "Corynebacteriaceae"           
## [1834,] "Saccharimonadales"           "Saccharimonadaceae"           
## [1835,] "Flavobacteriales"            "Weeksellaceae"                
## [1836,] "Spirochaetales"              "Spirochaetaceae"              
## [1837,] NA                            NA                             
## [1838,] "Bacteroidales"               "Prevotellaceae"               
## [1839,] "Fusobacteriales"             "Leptotrichiaceae"             
## [1840,] "Spirochaetales"              "Spirochaetaceae"              
## [1841,] "Selenomonadales"             "Veillonellaceae"              
## [1842,] "Clostridiales"               "Family_XI"                    
## [1843,] "Bacteroidales"               "Prevotellaceae"               
## [1844,] "Bacteroidales"               "Porphyromonadaceae"           
## [1845,] "Clostridiales"               "Peptostreptococcaceae"        
## [1846,] "Betaproteobacteriales"       "Burkholderiaceae"             
## [1847,] "Bacteroidales"               "Prevotellaceae"               
## [1848,] "Bacteroidales"               "Prevotellaceae"               
## [1849,] "Bacteroidales"               "Prevotellaceae"               
## [1850,] "Bacteroidales"               "Prevotellaceae"               
## [1851,] "Synergistales"               "Synergistaceae"               
## [1852,] "Bacteroidales"               "Prevotellaceae"               
## [1853,] "Bacteroidales"               "Prevotellaceae"               
## [1854,] "Pasteurellales"              "Pasteurellaceae"              
## [1855,] "Bacteroidales"               "Prevotellaceae"               
## [1856,] "Pasteurellales"              "Pasteurellaceae"              
## [1857,] "Bacteroidales"               "Prevotellaceae"               
## [1858,] "Bacteroidales"               "Porphyromonadaceae"           
## [1859,] "Betaproteobacteriales"       "Neisseriaceae"                
## [1860,] "Clostridiales"               "Lachnospiraceae"              
## [1861,] "Pasteurellales"              "Pasteurellaceae"              
## [1862,] "Flavobacteriales"            "Flavobacteriaceae"            
## [1863,] "Betaproteobacteriales"       "Neisseriaceae"                
## [1864,] "Fusobacteriales"             "Fusobacteriaceae"             
## [1865,] "Pasteurellales"              "Pasteurellaceae"              
## [1866,] "Bacteroidales"               "Prevotellaceae"               
## [1867,] "Fusobacteriales"             "Leptotrichiaceae"             
## [1868,] "Clostridiales"               "Lachnospiraceae"              
## [1869,] "Pasteurellales"              "Pasteurellaceae"              
## [1870,] "Selenomonadales"             "Veillonellaceae"              
## [1871,] "Pasteurellales"              "Pasteurellaceae"              
## [1872,] "Lactobacillales"             "Streptococcaceae"             
## [1873,] "Pasteurellales"              "Pasteurellaceae"              
## [1874,] "Selenomonadales"             "Veillonellaceae"              
## [1875,] "Fusobacteriales"             "Fusobacteriaceae"             
## [1876,] "Selenomonadales"             "Veillonellaceae"              
## [1877,] NA                            NA                             
## [1878,] "Flavobacteriales"            "Flavobacteriaceae"            
## [1879,] "Enterobacteriales"           "Enterobacteriaceae"           
## [1880,] "Clostridiales"               "Ruminococcaceae"              
## [1881,] "Bacteroidales"               "Prevotellaceae"               
## [1882,] "Selenomonadales"             "Veillonellaceae"              
## [1883,] "Fusobacteriales"             "Fusobacteriaceae"             
## [1884,] "Bacteroidales"               "Prevotellaceae"               
## [1885,] "Actinomycetales"             "Actinomycetaceae"             
## [1886,] NA                            NA                             
## [1887,] "Selenomonadales"             "Veillonellaceae"              
## [1888,] "Micrococcales"               "Micrococcaceae"               
## [1889,] "Bacteroidales"               "Prevotellaceae"               
## [1890,] "Bacteroidales"               "Prevotellaceae"               
## [1891,] "Betaproteobacteriales"       "Neisseriaceae"                
## [1892,] "Betaproteobacteriales"       "Neisseriaceae"                
## [1893,] "Selenomonadales"             "Veillonellaceae"              
## [1894,] "Fusobacteriales"             "Fusobacteriaceae"             
## [1895,] "Selenomonadales"             "Veillonellaceae"              
## [1896,] "Clostridiales"               "Lachnospiraceae"              
## [1897,] "Bacteroidales"               "Prevotellaceae"               
## [1898,] "Lactobacillales"             "Streptococcaceae"             
## [1899,] "Pasteurellales"              "Pasteurellaceae"              
## [1900,] "Selenomonadales"             "Veillonellaceae"              
## [1901,] "Clostridiales"               "Lachnospiraceae"              
## [1902,] "Selenomonadales"             "Veillonellaceae"              
## [1903,] "Bacteroidales"               "Prevotellaceae"               
## [1904,] "Bacteroidales"               "Prevotellaceae"               
## [1905,] "Fusobacteriales"             "Fusobacteriaceae"             
## [1906,] "Saccharimonadales"           "Saccharimonadaceae"           
## [1907,] "Clostridiales"               "Peptostreptococcaceae"        
## [1908,] "Bacteroidales"               "Prevotellaceae"               
## [1909,] "Clostridiales"               "Lachnospiraceae"              
## [1910,] "Fusobacteriales"             "Fusobacteriaceae"             
## [1911,] "Pasteurellales"              "Pasteurellaceae"              
## [1912,] "Bacteroidales"               "Prevotellaceae"               
## [1913,] "Selenomonadales"             "Veillonellaceae"              
## [1914,] "Flavobacteriales"            "Flavobacteriaceae"            
## [1915,] "Bacteroidales"               "Prevotellaceae"               
## [1916,] "Spirochaetales"              "Spirochaetaceae"              
## [1917,] "Flavobacteriales"            "Flavobacteriaceae"            
## [1918,] "Flavobacteriales"            "Flavobacteriaceae"            
## [1919,] "Micrococcales"               NA                             
## [1920,] "Clostridiales"               "Lachnospiraceae"              
## [1921,] "Bacteroidales"               "Prevotellaceae"               
## [1922,] "Clostridiales"               "Lachnospiraceae"              
## [1923,] "Pseudomonadales"             "Pseudomonadaceae"             
## [1924,] "Enterobacteriales"           "Enterobacteriaceae"           
## [1925,] "Bacteroidales"               "Prevotellaceae"               
## [1926,] "Bacteroidales"               "Prevotellaceae"               
## [1927,] "Campylobacterales"           "Campylobacteraceae"           
## [1928,] "Pasteurellales"              "Pasteurellaceae"              
## [1929,] "Selenomonadales"             "Veillonellaceae"              
## [1930,] "Bacteroidales"               "Prevotellaceae"               
## [1931,] "Fusobacteriales"             "Leptotrichiaceae"             
## [1932,] "Bacteroidales"               "Prevotellaceae"               
## [1933,] "Saccharimonadales"           "Saccharimonadaceae"           
## [1934,] "Bacteroidales"               "Prevotellaceae"               
## [1935,] "Micrococcales"               NA                             
## [1936,] "Micrococcales"               NA                             
## [1937,] "Betaproteobacteriales"       "Neisseriaceae"                
## [1938,] "Bacteroidales"               "Prevotellaceae"               
## [1939,] "Bacteroidales"               "Prevotellaceae"               
## [1940,] "Fusobacteriales"             "Fusobacteriaceae"             
## [1941,] "Bacillales"                  "Family_XI"                    
## [1942,] "Micrococcales"               NA                             
## [1943,] "Bacteroidales"               "Prevotellaceae"               
## [1944,] "Betaproteobacteriales"       "Neisseriaceae"                
## [1945,] "Clostridiales"               "Lachnospiraceae"              
## [1946,] "Spirochaetales"              "Spirochaetaceae"              
## [1947,] "Betaproteobacteriales"       "Neisseriaceae"                
## [1948,] "Fusobacteriales"             "Fusobacteriaceae"             
## [1949,] "Flavobacteriales"            "Flavobacteriaceae"            
## [1950,] "Selenomonadales"             "Veillonellaceae"              
## [1951,] "Bacteroidales"               "Prevotellaceae"               
## [1952,] "Clostridiales"               "Lachnospiraceae"              
## [1953,] "Selenomonadales"             "Veillonellaceae"              
## [1954,] "Pasteurellales"              "Pasteurellaceae"              
## [1955,] "Bacteroidales"               "Prevotellaceae"               
## [1956,] "Bacteroidales"               "Prevotellaceae"               
## [1957,] "Clostridiales"               "Lachnospiraceae"              
## [1958,] "Bacteroidales"               "Porphyromonadaceae"           
## [1959,] "Bacteroidales"               "Prevotellaceae"               
## [1960,] "Bacteroidales"               "Prevotellaceae"               
## [1961,] "Flavobacteriales"            "Flavobacteriaceae"            
## [1962,] "Coriobacteriales"            "Atopobiaceae"                 
## [1963,] "Bacteroidales"               "Prevotellaceae"               
## [1964,] "Actinomycetales"             "Actinomycetaceae"             
## [1965,] "Fusobacteriales"             "Leptotrichiaceae"             
## [1966,] "Selenomonadales"             "Veillonellaceae"              
## [1967,] "Bacteroidales"               "Prevotellaceae"               
## [1968,] "Selenomonadales"             "Veillonellaceae"              
## [1969,] "Fusobacteriales"             "Fusobacteriaceae"             
## [1970,] "Betaproteobacteriales"       "Neisseriaceae"                
## [1971,] "Fusobacteriales"             NA                             
## [1972,] "Clostridiales"               "Peptostreptococcaceae"        
## [1973,] "Bacteroidales"               "Prevotellaceae"               
## [1974,] "Clostridiales"               "Lachnospiraceae"              
## [1975,] "Clostridiales"               "Family_XI"                    
## [1976,] "Bacteroidales"               "Prevotellaceae"               
## [1977,] "Clostridiales"               "Peptostreptococcaceae"        
## [1978,] "Bacteroidales"               "Prevotellaceae"               
## [1979,] "Bacteroidales"               "Porphyromonadaceae"           
## [1980,] "Bacteroidales"               "Prevotellaceae"               
## [1981,] "Bacteroidales"               "Prevotellaceae"               
## [1982,] "Fusobacteriales"             "Fusobacteriaceae"             
## [1983,] "Selenomonadales"             "Veillonellaceae"              
## [1984,] "Betaproteobacteriales"       "Neisseriaceae"                
## [1985,] "Bacteroidales"               "Porphyromonadaceae"           
## [1986,] "Selenomonadales"             "Veillonellaceae"              
## [1987,] "Pasteurellales"              "Pasteurellaceae"              
## [1988,] "Micrococcales"               NA                             
## [1989,] "Fusobacteriales"             "Leptotrichiaceae"             
## [1990,] "Lactobacillales"             "Streptococcaceae"             
## [1991,] "Bacteroidales"               "Prevotellaceae"               
## [1992,] "Bacteroidales"               "Rikenellaceae"                
## [1993,] "Flavobacteriales"            "Flavobacteriaceae"            
## [1994,] "Betaproteobacteriales"       "Neisseriaceae"                
## [1995,] "Selenomonadales"             "Veillonellaceae"              
## [1996,] "Bacillales"                  "Family_XI"                    
## [1997,] "Bacteroidales"               "Tannerellaceae"               
## [1998,] "Selenomonadales"             "Veillonellaceae"              
## [1999,] "Clostridiales"               "Lachnospiraceae"              
## [2000,] "Betaproteobacteriales"       "Neisseriaceae"                
## [2001,] "Fusobacteriales"             "Leptotrichiaceae"             
## [2002,] "Clostridiales"               "Peptostreptococcaceae"        
## [2003,] "Fusobacteriales"             "Fusobacteriaceae"             
## [2004,] "Bacteroidales"               "Paludibacteraceae"            
## [2005,] "Bacteroidales"               "Prevotellaceae"               
## [2006,] "Clostridiales"               "Lachnospiraceae"              
## [2007,] "Bacteroidales"               "Prevotellaceae"               
## [2008,] "Selenomonadales"             "Veillonellaceae"              
## [2009,] "Bacteroidales"               "Prevotellaceae"               
## [2010,] "Pasteurellales"              "Pasteurellaceae"              
## [2011,] "Bacteroidales"               "Prevotellaceae"               
## [2012,] "Bacteroidales"               "Porphyromonadaceae"           
## [2013,] "Saccharimonadales"           NA                             
## [2014,] "Bacteroidales"               "Prevotellaceae"               
## [2015,] "Bacteroidales"               "Prevotellaceae"               
## [2016,] "Selenomonadales"             "Veillonellaceae"              
## [2017,] "Flavobacteriales"            "Flavobacteriaceae"            
## [2018,] "Saccharimonadales"           "Saccharimonadaceae"           
## [2019,] "Selenomonadales"             "Veillonellaceae"              
## [2020,] "Fusobacteriales"             "Fusobacteriaceae"             
## [2021,] "Bacteroidales"               "Porphyromonadaceae"           
## [2022,] "Actinomycetales"             "Actinomycetaceae"             
## [2023,] "Pasteurellales"              "Pasteurellaceae"              
## [2024,] "Pasteurellales"              "Pasteurellaceae"              
## [2025,] "Betaproteobacteriales"       "Neisseriaceae"                
## [2026,] "Bacteroidales"               "Prevotellaceae"               
## [2027,] NA                            NA                             
## [2028,] "Bacteroidales"               "Prevotellaceae"               
## [2029,] "Fusobacteriales"             "Leptotrichiaceae"             
## [2030,] "Fusobacteriales"             NA                             
## [2031,] "Fusobacteriales"             "Leptotrichiaceae"             
## [2032,] "Saccharimonadales"           "Saccharimonadaceae"           
## [2033,] "Fusobacteriales"             "Fusobacteriaceae"             
## [2034,] "Lactobacillales"             "Streptococcaceae"             
## [2035,] "Lactobacillales"             "Streptococcaceae"             
## [2036,] "Lactobacillales"             "Carnobacteriaceae"            
## [2037,] "Synergistales"               "Synergistaceae"               
## [2038,] "Bacteroidales"               "Prevotellaceae"               
## [2039,] "Fusobacteriales"             "Fusobacteriaceae"             
## [2040,] "Selenomonadales"             "Veillonellaceae"              
## [2041,] "Selenomonadales"             "Veillonellaceae"              
## [2042,] "Bacteroidales"               "Prevotellaceae"               
## [2043,] "Pasteurellales"              "Pasteurellaceae"              
## [2044,] "Betaproteobacteriales"       "Neisseriaceae"                
## [2045,] "Absconditabacteriales_(SR1)" NA                             
## [2046,] "Fusobacteriales"             "Leptotrichiaceae"             
## [2047,] "Clostridiales"               "Ruminococcaceae"              
## [2048,] "Selenomonadales"             "Veillonellaceae"              
## [2049,] "Campylobacterales"           "Campylobacteraceae"           
## [2050,] "Bacteroidales"               "Prevotellaceae"               
## [2051,] "Bacteroidales"               "Prevotellaceae"               
## [2052,] "Selenomonadales"             "Veillonellaceae"              
## [2053,] "Bacteroidales"               "Prevotellaceae"               
## [2054,] "Actinomycetales"             "Actinomycetaceae"             
## [2055,] "Bacteroidales"               "Prevotellaceae"               
## [2056,] "Selenomonadales"             "Veillonellaceae"              
## [2057,] "Saccharimonadales"           "Saccharimonadaceae"           
## [2058,] "Selenomonadales"             "Veillonellaceae"              
## [2059,] "Bacteroidales"               "Prevotellaceae"               
## [2060,] "Clostridiales"               "Lachnospiraceae"              
## [2061,] "Bacteroidales"               "Prevotellaceae"               
## [2062,] "Selenomonadales"             "Veillonellaceae"              
## [2063,] "Chloroplast"                 NA                             
## [2064,] "Bacteroidales"               "Porphyromonadaceae"           
## [2065,] "Selenomonadales"             "Veillonellaceae"              
## [2066,] "Betaproteobacteriales"       "Neisseriaceae"                
## [2067,] "Bacteroidales"               "Prevotellaceae"               
## [2068,] "Bacteroidales"               "Prevotellaceae"               
## [2069,] "Fusobacteriales"             "Fusobacteriaceae"             
## [2070,] "Saccharimonadales"           "Saccharimonadaceae"           
## [2071,] "Bacteroidales"               "Prevotellaceae"               
## [2072,] "Campylobacterales"           NA                             
## [2073,] "Clostridiales"               "Lachnospiraceae"              
## [2074,] "Pasteurellales"              "Pasteurellaceae"              
## [2075,] "Fusobacteriales"             NA                             
## [2076,] "Selenomonadales"             "Veillonellaceae"              
## [2077,] "Spirochaetales"              "Spirochaetaceae"              
## [2078,] "Fusobacteriales"             "Fusobacteriaceae"             
## [2079,] "Saccharimonadales"           "Saccharimonadaceae"           
## [2080,] "Bacteroidales"               "Porphyromonadaceae"           
## [2081,] "Flavobacteriales"            "Flavobacteriaceae"            
## [2082,] "Bacteroidales"               "Prevotellaceae"               
## [2083,] "Fusobacteriales"             "Fusobacteriaceae"             
## [2084,] NA                            NA                             
## [2085,] "Betaproteobacteriales"       "Neisseriaceae"                
## [2086,] "Fusobacteriales"             "Leptotrichiaceae"             
## [2087,] "Clostridiales"               "Lachnospiraceae"              
## [2088,] "Bacteroidales"               "Prevotellaceae"               
## [2089,] "Selenomonadales"             "Veillonellaceae"              
## [2090,] "Bacteroidales"               "Prevotellaceae"               
## [2091,] "Spirochaetales"              "Spirochaetaceae"              
## [2092,] "Micrococcales"               NA                             
## [2093,] "Micrococcales"               NA                             
## [2094,] "Bacteroidales"               "Prevotellaceae"               
## [2095,] "Bacteroidales"               "Porphyromonadaceae"           
## [2096,] "Fusobacteriales"             "Fusobacteriaceae"             
## [2097,] "Bacteroidales"               "Porphyromonadaceae"           
## [2098,] "Spirochaetales"              "Spirochaetaceae"              
## [2099,] "Fusobacteriales"             "Leptotrichiaceae"             
## [2100,] "Bacteroidales"               "Prevotellaceae"               
## [2101,] "Bacteroidales"               "Prevotellaceae"               
## [2102,] "Fusobacteriales"             "Leptotrichiaceae"             
## [2103,] "Clostridiales"               "Lachnospiraceae"              
## [2104,] "Bacteroidales"               "Prevotellaceae"               
## [2105,] "Fusobacteriales"             "Leptotrichiaceae"             
## [2106,] "Pasteurellales"              "Pasteurellaceae"              
## [2107,] "Bacteroidales"               "Prevotellaceae"               
## [2108,] "Selenomonadales"             "Veillonellaceae"              
## [2109,] "Bacteroidales"               "Prevotellaceae"               
## [2110,] "Bacteroidales"               "Prevotellaceae"               
## [2111,] "Bacteroidales"               "Prevotellaceae"               
## [2112,] "Clostridiales"               "Lachnospiraceae"              
## [2113,] "Selenomonadales"             "Veillonellaceae"              
## [2114,] "Betaproteobacteriales"       "Neisseriaceae"                
## [2115,] "Bacteroidales"               "Prevotellaceae"               
## [2116,] NA                            NA                             
## [2117,] "Lactobacillales"             "Carnobacteriaceae"            
## [2118,] "Bacteroidales"               "Prevotellaceae"               
## [2119,] "Selenomonadales"             "Veillonellaceae"              
## [2120,] "Pasteurellales"              "Pasteurellaceae"              
## [2121,] "Flavobacteriales"            "Flavobacteriaceae"            
## [2122,] "Bacteroidales"               "Prevotellaceae"               
## [2123,] "JGI_0000069-P22"             NA                             
## [2124,] "Selenomonadales"             "Veillonellaceae"              
## [2125,] "Bacteroidales"               "Prevotellaceae"               
## [2126,] "Lactobacillales"             "Streptococcaceae"             
## [2127,] "Bacteroidales"               "Prevotellaceae"               
## [2128,] "Selenomonadales"             "Veillonellaceae"              
## [2129,] "Lactobacillales"             "Streptococcaceae"             
## [2130,] "Pasteurellales"              "Pasteurellaceae"              
## [2131,] "Bacteroidales"               "Prevotellaceae"               
## [2132,] "Campylobacterales"           NA                             
## [2133,] "Actinomycetales"             "Actinomycetaceae"             
## [2134,] "Fusobacteriales"             "Fusobacteriaceae"             
## [2135,] "Bacteroidales"               "Prevotellaceae"               
## [2136,] "Selenomonadales"             "Veillonellaceae"              
## [2137,] "Micrococcales"               "Micrococcaceae"               
## [2138,] "Bacteroidales"               "Prevotellaceae"               
## [2139,] "Bacteroidales"               "Prevotellaceae"               
## [2140,] "Clostridiales"               "Family_XIII"                  
## [2141,] "Spirochaetales"              "Spirochaetaceae"              
## [2142,] "Campylobacterales"           NA                             
## [2143,] "Betaproteobacteriales"       "Neisseriaceae"                
## [2144,] "Selenomonadales"             "Veillonellaceae"              
## [2145,] "Bacteroidales"               "Prevotellaceae"               
## [2146,] "Bacteroidales"               "Prevotellaceae"               
## [2147,] "Bacteroidales"               "Prevotellaceae"               
## [2148,] "Bacteroidales"               "Porphyromonadaceae"           
## [2149,] "Fusobacteriales"             "Leptotrichiaceae"             
## [2150,] "Betaproteobacteriales"       "Neisseriaceae"                
## [2151,] "Saccharimonadales"           "Saccharimonadaceae"           
## [2152,] "Cardiobacteriales"           "Cardiobacteriaceae"           
## [2153,] "Campylobacterales"           "Campylobacteraceae"           
## [2154,] "Bacteroidales"               "Prevotellaceae"               
## [2155,] "Bacteroidales"               "Prevotellaceae"               
## [2156,] "Bacteroidales"               "Prevotellaceae"               
## [2157,] "Betaproteobacteriales"       "Neisseriaceae"                
## [2158,] NA                            NA                             
## [2159,] "Bacteroidales"               "Porphyromonadaceae"           
## [2160,] "Lactobacillales"             "Carnobacteriaceae"            
## [2161,] "Pasteurellales"              "Pasteurellaceae"              
## [2162,] "Actinomycetales"             "Actinomycetaceae"             
## [2163,] "Fusobacteriales"             "Fusobacteriaceae"             
## [2164,] "Flavobacteriales"            "Flavobacteriaceae"            
## [2165,] "Selenomonadales"             "Veillonellaceae"              
## [2166,] "Bacteroidales"               "Porphyromonadaceae"           
## [2167,] "Bacteroidales"               "Prevotellaceae"               
## [2168,] "Spirochaetales"              "Spirochaetaceae"              
## [2169,] "Bacteroidales"               "Prevotellaceae"               
## [2170,] "Clostridiales"               "Lachnospiraceae"              
## [2171,] "Fusobacteriales"             "Fusobacteriaceae"             
## [2172,] "Clostridiales"               "Lachnospiraceae"              
## [2173,] "Actinomycetales"             "Actinomycetaceae"             
## [2174,] "Bacteroidales"               "Prevotellaceae"               
## [2175,] "Bacteroidales"               "Prevotellaceae"               
## [2176,] "Fusobacteriales"             "Leptotrichiaceae"             
## [2177,] "Flavobacteriales"            "Flavobacteriaceae"            
## [2178,] "Clostridiales"               "Lachnospiraceae"              
## [2179,] "Selenomonadales"             "Veillonellaceae"              
## [2180,] "Selenomonadales"             "Veillonellaceae"              
## [2181,] "Fusobacteriales"             "Fusobacteriaceae"             
## [2182,] "Clostridiales"               "Peptostreptococcaceae"        
## [2183,] "Bacteroidales"               "Porphyromonadaceae"           
## [2184,] "Bacteroidales"               "Prevotellaceae"               
## [2185,] "Bacteroidales"               "Prevotellaceae"               
## [2186,] "Fusobacteriales"             "Fusobacteriaceae"             
## [2187,] "Selenomonadales"             "Veillonellaceae"              
## [2188,] "Selenomonadales"             "Veillonellaceae"              
## [2189,] "Fusobacteriales"             "Fusobacteriaceae"             
## [2190,] "Selenomonadales"             "Veillonellaceae"              
## [2191,] "Bacteroidales"               "Prevotellaceae"               
## [2192,] "Betaproteobacteriales"       "Neisseriaceae"                
## [2193,] "Flavobacteriales"            "Weeksellaceae"                
## [2194,] "Selenomonadales"             "Veillonellaceae"              
## [2195,] "Pasteurellales"              "Pasteurellaceae"              
## [2196,] "Saccharimonadales"           NA                             
## [2197,] "Bacteroidales"               "Porphyromonadaceae"           
## [2198,] "Fusobacteriales"             "Fusobacteriaceae"             
## [2199,] "Bacteroidales"               "Prevotellaceae"               
## [2200,] "Selenomonadales"             "Veillonellaceae"              
## [2201,] "Actinomycetales"             "Actinomycetaceae"             
## [2202,] NA                            NA                             
## [2203,] "Selenomonadales"             "Veillonellaceae"              
## [2204,] "Fusobacteriales"             "Fusobacteriaceae"             
## [2205,] "Selenomonadales"             "Veillonellaceae"              
## [2206,] "Pasteurellales"              "Pasteurellaceae"              
## [2207,] "Bacteroidales"               "Prevotellaceae"               
## [2208,] "Pasteurellales"              "Pasteurellaceae"              
## [2209,] "Saccharimonadales"           "Saccharimonadaceae"           
## [2210,] "Spirochaetales"              "Spirochaetaceae"              
## [2211,] "Clostridiales"               "Peptostreptococcaceae"        
## [2212,] "Bacteroidales"               "Prevotellaceae"               
## [2213,] "Bacteroidales"               "Porphyromonadaceae"           
## [2214,] "Selenomonadales"             "Veillonellaceae"              
## [2215,] "Bacteroidales"               "Prevotellaceae"               
## [2216,] "Bacteroidales"               "Prevotellaceae"               
## [2217,] "Flavobacteriales"            "Flavobacteriaceae"            
## [2218,] "Selenomonadales"             "Veillonellaceae"              
## [2219,] "Bacteroidales"               "Prevotellaceae"               
## [2220,] "Bacteroidales"               "Prevotellaceae"               
## [2221,] "Saccharimonadales"           NA                             
## [2222,] "Flavobacteriales"            "Flavobacteriaceae"            
## [2223,] "Chloroplast"                 NA                             
## [2224,] "Flavobacteriales"            "Flavobacteriaceae"            
## [2225,] "Selenomonadales"             "Veillonellaceae"              
## [2226,] "Saccharimonadales"           "Saccharimonadaceae"           
## [2227,] "Lactobacillales"             "Streptococcaceae"             
## [2228,] "Bacteroidales"               "Prevotellaceae"               
## [2229,] "Clostridiales"               "Lachnospiraceae"              
## [2230,] NA                            NA                             
## [2231,] "Actinomycetales"             "Actinomycetaceae"             
## [2232,] "Selenomonadales"             "Veillonellaceae"              
## [2233,] "Bacteroidales"               "Prevotellaceae"               
## [2234,] "Selenomonadales"             "Veillonellaceae"              
## [2235,] "Clostridiales"               "Lachnospiraceae"              
## [2236,] "Selenomonadales"             "Veillonellaceae"              
## [2237,] "Bacteroidales"               "Prevotellaceae"               
## [2238,] "Fusobacteriales"             "Fusobacteriaceae"             
## [2239,] "Selenomonadales"             "Veillonellaceae"              
## [2240,] "Micrococcales"               NA                             
## [2241,] "Bacteroidales"               "Prevotellaceae"               
## [2242,] "Chloroplast"                 NA                             
## [2243,] "Bacteroidales"               "Porphyromonadaceae"           
## [2244,] "Bacteroidales"               "Prevotellaceae"               
## [2245,] "Pasteurellales"              "Pasteurellaceae"              
## [2246,] "Bacteroidales"               "Porphyromonadaceae"           
## [2247,] "Bacteroidales"               "Prevotellaceae"               
## [2248,] "Bacteroidales"               "Porphyromonadaceae"           
## [2249,] NA                            NA                             
## [2250,] "Saccharimonadales"           "Saccharimonadaceae"           
## [2251,] "Coriobacteriales"            "Atopobiaceae"                 
## [2252,] "Bacteroidales"               "Prevotellaceae"               
## [2253,] "Selenomonadales"             "Veillonellaceae"              
## [2254,] "Selenomonadales"             "Veillonellaceae"              
## [2255,] "Synergistales"               "Synergistaceae"               
## [2256,] "Bacteroidales"               "Prevotellaceae"               
## [2257,] "Bacteroidales"               "Prevotellaceae"               
## [2258,] "Clostridiales"               "Peptostreptococcaceae"        
## [2259,] "Corynebacteriales"           "Corynebacteriaceae"           
## [2260,] "Bacillales"                  "Family_XI"                    
## [2261,] "Selenomonadales"             "Veillonellaceae"              
## [2262,] "Pasteurellales"              "Pasteurellaceae"              
## [2263,] "Betaproteobacteriales"       "Neisseriaceae"                
## [2264,] "Fusobacteriales"             "Leptotrichiaceae"             
## [2265,] "Bacteroidales"               "Prevotellaceae"               
## [2266,] "Fusobacteriales"             "Leptotrichiaceae"             
## [2267,] "Selenomonadales"             "Veillonellaceae"              
## [2268,] "Betaproteobacteriales"       "Neisseriaceae"                
## [2269,] "Flavobacteriales"            "Flavobacteriaceae"            
## [2270,] "Bacteroidales"               "Prevotellaceae"               
## [2271,] "Fusobacteriales"             "Leptotrichiaceae"             
## [2272,] "Absconditabacteriales_(SR1)" NA                             
## [2273,] "Fusobacteriales"             NA                             
## [2274,] "Clostridiales"               "Lachnospiraceae"              
## [2275,] "Bacteroidales"               "Porphyromonadaceae"           
## [2276,] "Flavobacteriales"            "Flavobacteriaceae"            
## [2277,] "Clostridiales"               "Lachnospiraceae"              
## [2278,] "Selenomonadales"             "Veillonellaceae"              
## [2279,] "Pasteurellales"              "Pasteurellaceae"              
## [2280,] NA                            NA                             
## [2281,] "Betaproteobacteriales"       "Neisseriaceae"                
## [2282,] "Clostridiales"               "Lachnospiraceae"              
## [2283,] "Bacteroidales"               "Prevotellaceae"               
## [2284,] "Bacteroidales"               "Prevotellaceae"               
## [2285,] "Bacteroidales"               "Prevotellaceae"               
## [2286,] "Clostridiales"               "Lachnospiraceae"              
## [2287,] "Bacteroidales"               "Prevotellaceae"               
## [2288,] "Sphingobacteriales"          "Lentimicrobiaceae"            
## [2289,] "Absconditabacteriales_(SR1)" NA                             
## [2290,] "Fusobacteriales"             "Leptotrichiaceae"             
## [2291,] "Selenomonadales"             "Veillonellaceae"              
## [2292,] "Pasteurellales"              "Pasteurellaceae"              
## [2293,] "Bacteroidales"               "Prevotellaceae"               
## [2294,] "Bacteroidales"               "Prevotellaceae"               
## [2295,] "Bacteroidales"               "Porphyromonadaceae"           
## [2296,] "Betaproteobacteriales"       "Neisseriaceae"                
## [2297,] "Bacteroidales"               "Prevotellaceae"               
## [2298,] "Bacteroidales"               "Prevotellaceae"               
## [2299,] "Bacteroidales"               "Prevotellaceae"               
## [2300,] "Bacteroidales"               "Prevotellaceae"               
## [2301,] "Selenomonadales"             "Veillonellaceae"              
## [2302,] "Clostridiales"               "Peptostreptococcaceae"        
## [2303,] "Bifidobacteriales"           "Bifidobacteriaceae"           
## [2304,] "Clostridiales"               "Peptostreptococcaceae"        
## [2305,] "Spirochaetales"              "Spirochaetaceae"              
## [2306,] "Flavobacteriales"            "Weeksellaceae"                
## [2307,] "Saccharimonadales"           "Saccharimonadaceae"           
## [2308,] "Flavobacteriales"            "Flavobacteriaceae"            
## [2309,] "Clostridiales"               "Peptostreptococcaceae"        
## [2310,] "Bacteroidales"               "Prevotellaceae"               
## [2311,] "Clostridiales"               "Lachnospiraceae"              
## [2312,] "Selenomonadales"             "Veillonellaceae"              
## [2313,] "Micrococcales"               NA                             
## [2314,] "Bacteroidales"               "Prevotellaceae"               
## [2315,] "Fusobacteriales"             NA                             
## [2316,] "Micrococcales"               NA                             
## [2317,] NA                            NA                             
## [2318,] "Bacteroidales"               "Porphyromonadaceae"           
## [2319,] "Clostridiales"               "Lachnospiraceae"              
## [2320,] "Saccharimonadales"           NA                             
## [2321,] "Bacteroidales"               "Prevotellaceae"               
## [2322,] "Bacteroidales"               "Prevotellaceae"               
## [2323,] "Bacteroidales"               "Prevotellaceae"               
## [2324,] "Bacteroidales"               "Prevotellaceae"               
## [2325,] "Bacteroidales"               "Prevotellaceae"               
## [2326,] NA                            NA                             
## [2327,] "Fusobacteriales"             "Leptotrichiaceae"             
## [2328,] "Bacteroidales"               "Prevotellaceae"               
## [2329,] "Clostridiales"               "Ruminococcaceae"              
## [2330,] "Bacteroidales"               "Prevotellaceae"               
## [2331,] "Bacteroidales"               "Porphyromonadaceae"           
## [2332,] NA                            NA                             
## [2333,] "Bacteroidales"               "Prevotellaceae"               
## [2334,] "Bacteroidales"               "Prevotellaceae"               
## [2335,] "Betaproteobacteriales"       "Neisseriaceae"                
## [2336,] "Betaproteobacteriales"       "Neisseriaceae"                
## [2337,] "Bacteroidales"               "Prevotellaceae"               
## [2338,] "Bacteroidales"               "Prevotellaceae"               
## [2339,] "Lactobacillales"             "Streptococcaceae"             
## [2340,] "Bacteroidales"               "Prevotellaceae"               
## [2341,] "Saccharimonadales"           "Saccharimonadaceae"           
## [2342,] "Clostridiales"               "Lachnospiraceae"              
## [2343,] "Pasteurellales"              "Pasteurellaceae"              
## [2344,] "Bacteroidales"               "Porphyromonadaceae"           
## [2345,] "Bacteroidales"               "Porphyromonadaceae"           
## [2346,] "Micrococcales"               NA                             
## [2347,] "Selenomonadales"             "Veillonellaceae"              
## [2348,] "Corynebacteriales"           "Corynebacteriaceae"           
## [2349,] "Synergistales"               "Synergistaceae"               
## [2350,] "Spirochaetales"              "Spirochaetaceae"              
## [2351,] "Bacteroidales"               "Prevotellaceae"               
## [2352,] "Bacteroidales"               "Prevotellaceae"               
## [2353,] "Actinomycetales"             "Actinomycetaceae"             
## [2354,] "Micrococcales"               NA                             
## [2355,] "Clostridiales"               "Lachnospiraceae"              
## [2356,] "Bacteroidales"               "Prevotellaceae"               
## [2357,] "Selenomonadales"             "Veillonellaceae"              
## [2358,] "Bacteroidales"               "Prevotellaceae"               
## [2359,] "Bacteroidales"               "Prevotellaceae"               
## [2360,] "Selenomonadales"             "Veillonellaceae"              
## [2361,] "Enterobacteriales"           "Enterobacteriaceae"           
## [2362,] "Selenomonadales"             "Veillonellaceae"              
## [2363,] "Bacteroidales"               "Prevotellaceae"               
## [2364,] "Selenomonadales"             "Veillonellaceae"              
## [2365,] "Selenomonadales"             "Veillonellaceae"              
## [2366,] "Fusobacteriales"             "Fusobacteriaceae"             
## [2367,] "Actinomycetales"             "Actinomycetaceae"             
## [2368,] "Bacteroidales"               "Prevotellaceae"               
## [2369,] "Betaproteobacteriales"       "Neisseriaceae"                
## [2370,] "Fusobacteriales"             "Leptotrichiaceae"             
## [2371,] "Bacteroidales"               "Porphyromonadaceae"           
## [2372,] "Selenomonadales"             "Veillonellaceae"              
## [2373,] "Bacteroidales"               "Prevotellaceae"               
## [2374,] "Bacteroidales"               "Prevotellaceae"               
## [2375,] "Selenomonadales"             "Veillonellaceae"              
## [2376,] "Clostridiales"               "Peptostreptococcaceae"        
## [2377,] "Betaproteobacteriales"       "Neisseriaceae"                
## [2378,] "Spirochaetales"              "Spirochaetaceae"              
## [2379,] "Bacteroidales"               "Prevotellaceae"               
## [2380,] "Fusobacteriales"             "Fusobacteriaceae"             
## [2381,] "Bacteroidales"               "Prevotellaceae"               
## [2382,] "Fusobacteriales"             "Leptotrichiaceae"             
## [2383,] "Clostridiales"               "Family_XIII"                  
## [2384,] "Bacteroidales"               "Paludibacteraceae"            
## [2385,] "Pasteurellales"              "Pasteurellaceae"              
## [2386,] "Bacteroidales"               "Prevotellaceae"               
## [2387,] "Selenomonadales"             "Veillonellaceae"              
## [2388,] "Actinomycetales"             "Actinomycetaceae"             
## [2389,] "Bacteroidales"               "Prevotellaceae"               
## [2390,] "Selenomonadales"             "Veillonellaceae"              
## [2391,] "Fusobacteriales"             "Leptotrichiaceae"             
## [2392,] "Selenomonadales"             "Veillonellaceae"              
## [2393,] "Fusobacteriales"             "Leptotrichiaceae"             
## [2394,] "Saccharimonadales"           "Saccharimonadaceae"           
## [2395,] "Selenomonadales"             "Veillonellaceae"              
## [2396,] "Selenomonadales"             "Veillonellaceae"              
## [2397,] "Selenomonadales"             "Veillonellaceae"              
## [2398,] "Corynebacteriales"           "Corynebacteriaceae"           
## [2399,] "Actinomycetales"             "Actinomycetaceae"             
## [2400,] "Bacteroidales"               "Prevotellaceae"               
## [2401,] "Coriobacteriales"            "Atopobiaceae"                 
## [2402,] "Spirochaetales"              "Spirochaetaceae"              
## [2403,] "Bacteroidales"               "Prevotellaceae"               
## [2404,] "Fusobacteriales"             "Fusobacteriaceae"             
## [2405,] "Clostridiales"               "Lachnospiraceae"              
## [2406,] "Selenomonadales"             "Veillonellaceae"              
## [2407,] "Bacteroidales"               "Prevotellaceae"               
## [2408,] "Bacteroidales"               "Prevotellaceae"               
## [2409,] "Fusobacteriales"             "Leptotrichiaceae"             
## [2410,] "Bacteroidales"               "Prevotellaceae"               
## [2411,] "Selenomonadales"             "Veillonellaceae"              
## [2412,] NA                            NA                             
## [2413,] "Saccharimonadales"           "Saccharimonadaceae"           
## [2414,] "Bacteroidales"               "Prevotellaceae"               
## [2415,] "Selenomonadales"             "Veillonellaceae"              
## [2416,] "Selenomonadales"             "Veillonellaceae"              
## [2417,] "Selenomonadales"             "Veillonellaceae"              
## [2418,] "Bacteroidales"               "Prevotellaceae"               
## [2419,] "Bacteroidales"               "Prevotellaceae"               
## [2420,] "Clostridiales"               "Lachnospiraceae"              
## [2421,] "Pasteurellales"              "Pasteurellaceae"              
## [2422,] "Micrococcales"               NA                             
## [2423,] "Micrococcales"               NA                             
## [2424,] NA                            NA                             
## [2425,] "Bacteroidales"               "Prevotellaceae"               
## [2426,] "Chloroplast"                 NA                             
## [2427,] "Bacteroidales"               "Prevotellaceae"               
## [2428,] "Bacteroidales"               "Porphyromonadaceae"           
## [2429,] "Flavobacteriales"            "Flavobacteriaceae"            
## [2430,] "Bacteroidales"               "Prevotellaceae"               
## [2431,] "Flavobacteriales"            "Flavobacteriaceae"            
## [2432,] "Clostridiales"               "Lachnospiraceae"              
## [2433,] "Selenomonadales"             "Veillonellaceae"              
## [2434,] "Bacteroidales"               "Prevotellaceae"               
## [2435,] NA                            NA                             
## [2436,] "Bacteroidales"               "Prevotellaceae"               
## [2437,] "Spirochaetales"              "Spirochaetaceae"              
## [2438,] NA                            NA                             
## [2439,] "Bacteroidales"               "Prevotellaceae"               
## [2440,] "Chloroplast"                 NA                             
## [2441,] "Actinomycetales"             "Actinomycetaceae"             
## [2442,] "Bacteroidales"               "Porphyromonadaceae"           
## [2443,] "Betaproteobacteriales"       "Neisseriaceae"                
## [2444,] "Pasteurellales"              "Pasteurellaceae"              
## [2445,] "Clostridiales"               "Ruminococcaceae"              
## [2446,] "Bacteroidales"               "Prevotellaceae"               
## [2447,] "Bacteroidales"               "Prevotellaceae"               
## [2448,] "Lactobacillales"             "Streptococcaceae"             
## [2449,] "Saccharimonadales"           "Saccharimonadaceae"           
## [2450,] "Bacteroidales"               "Prevotellaceae"               
## [2451,] NA                            NA                             
## [2452,] "Bacteroidales"               "Prevotellaceae"               
## [2453,] "Betaproteobacteriales"       "Neisseriaceae"                
## [2454,] "Bacteroidales"               "Prevotellaceae"               
## [2455,] "Clostridiales"               "Lachnospiraceae"              
## [2456,] "Actinomycetales"             "Actinomycetaceae"             
## [2457,] "Betaproteobacteriales"       "Neisseriaceae"                
## [2458,] "Bacteroidales"               "Prevotellaceae"               
## [2459,] "Clostridiales"               "Lachnospiraceae"              
## [2460,] "Spirochaetales"              "Spirochaetaceae"              
## [2461,] "Clostridiales"               "Lachnospiraceae"              
## [2462,] "Selenomonadales"             "Veillonellaceae"              
## [2463,] "Bacteroidales"               "Prevotellaceae"               
## [2464,] "Clostridiales"               "Peptostreptococcaceae"        
## [2465,] "Bacteroidales"               "Prevotellaceae"               
## [2466,] "Bacteroidales"               "Prevotellaceae"               
## [2467,] "Betaproteobacteriales"       "Neisseriaceae"                
## [2468,] "Flavobacteriales"            "Weeksellaceae"                
## [2469,] "Bacteroidales"               "Prevotellaceae"               
## [2470,] "Bacteroidales"               "Prevotellaceae"               
## [2471,] "Betaproteobacteriales"       "Neisseriaceae"                
## [2472,] "Pasteurellales"              "Pasteurellaceae"              
## [2473,] "Selenomonadales"             "Veillonellaceae"              
## [2474,] "Clostridiales"               "Peptococcaceae"               
## [2475,] "Bacteroidales"               "Prevotellaceae"               
## [2476,] "Bacteroidales"               "Prevotellaceae"               
## [2477,] "Saccharimonadales"           NA                             
## [2478,] "Selenomonadales"             "Veillonellaceae"              
## [2479,] "Selenomonadales"             "Veillonellaceae"              
## [2480,] "Micrococcales"               NA                             
## [2481,] "Micrococcales"               NA                             
## [2482,] "Clostridiales"               "Lachnospiraceae"              
## [2483,] "Bacteroidales"               "Prevotellaceae"               
## [2484,] "Spirochaetales"              "Spirochaetaceae"              
## [2485,] "Betaproteobacteriales"       "Neisseriaceae"                
## [2486,] "Clostridiales"               "Family_XIII"                  
## [2487,] "Bacteroidales"               "Prevotellaceae"               
## [2488,] "Bacteroidales"               "Prevotellaceae"               
## [2489,] "Pasteurellales"              "Pasteurellaceae"              
## [2490,] "Bacteroidales"               "Prevotellaceae"               
## [2491,] "Fusobacteriales"             "Fusobacteriaceae"             
## [2492,] "Selenomonadales"             "Veillonellaceae"              
## [2493,] "Sphingobacteriales"          "Lentimicrobiaceae"            
## [2494,] "Bacteroidales"               "Prevotellaceae"               
## [2495,] "Flavobacteriales"            "Flavobacteriaceae"            
## [2496,] "Fusobacteriales"             "Fusobacteriaceae"             
## [2497,] "Actinomycetales"             "Actinomycetaceae"             
## [2498,] "Pasteurellales"              "Pasteurellaceae"              
## [2499,] "Bacteroidales"               NA                             
## [2500,] "Corynebacteriales"           "Corynebacteriaceae"           
## [2501,] "Selenomonadales"             "Veillonellaceae"              
## [2502,] "Bacteroidales"               "Prevotellaceae"               
## [2503,] "Bacteroidales"               "Prevotellaceae"               
## [2504,] "Enterobacteriales"           "Enterobacteriaceae"           
## [2505,] "Fusobacteriales"             "Fusobacteriaceae"             
## [2506,] "Bacteroidales"               "Prevotellaceae"               
## [2507,] "Flavobacteriales"            "Flavobacteriaceae"            
## [2508,] "Bacteroidales"               "Prevotellaceae"               
## [2509,] "Lactobacillales"             "Streptococcaceae"             
## [2510,] "Saccharimonadales"           "Saccharimonadaceae"           
## [2511,] "Flavobacteriales"            "Flavobacteriaceae"            
## [2512,] "Betaproteobacteriales"       "Neisseriaceae"                
## [2513,] "Clostridiales"               "Lachnospiraceae"              
## [2514,] "Clostridiales"               "Peptostreptococcaceae"        
## [2515,] "Bacteroidales"               "Prevotellaceae"               
## [2516,] "Betaproteobacteriales"       "Neisseriaceae"                
## [2517,] "Micrococcales"               "Micrococcaceae"               
## [2518,] "Fusobacteriales"             "Leptotrichiaceae"             
## [2519,] "Actinomycetales"             "Actinomycetaceae"             
## [2520,] "Actinomycetales"             "Actinomycetaceae"             
## [2521,] "Betaproteobacteriales"       "Neisseriaceae"                
## [2522,] "Fusobacteriales"             "Fusobacteriaceae"             
## [2523,] "Bacteroidales"               "Prevotellaceae"               
## [2524,] "Selenomonadales"             "Veillonellaceae"              
## [2525,] "Betaproteobacteriales"       "Neisseriaceae"                
## [2526,] "Micrococcales"               NA                             
## [2527,] "Bacteroidales"               "Prevotellaceae"               
## [2528,] "Lactobacillales"             "Streptococcaceae"             
## [2529,] "Fusobacteriales"             NA                             
## [2530,] "Bacteroidales"               "Tannerellaceae"               
## [2531,] "Selenomonadales"             "Veillonellaceae"              
## [2532,] "Selenomonadales"             "Veillonellaceae"              
## [2533,] "Bacteroidales"               "Prevotellaceae"               
## [2534,] "Flavobacteriales"            "Flavobacteriaceae"            
## [2535,] "Bacteroidales"               "Prevotellaceae"               
## [2536,] "Pasteurellales"              "Pasteurellaceae"              
## [2537,] "Lactobacillales"             "Streptococcaceae"             
## [2538,] "Sphingobacteriales"          "Lentimicrobiaceae"            
## [2539,] "Micrococcales"               NA                             
## [2540,] "Pasteurellales"              "Pasteurellaceae"              
## [2541,] "Bacteroidales"               "Prevotellaceae"               
## [2542,] "Clostridiales"               "Ruminococcaceae"              
## [2543,] "Fusobacteriales"             "Fusobacteriaceae"             
## [2544,] "Bacteroidales"               "Prevotellaceae"               
## [2545,] NA                            NA                             
## [2546,] "Fusobacteriales"             "Fusobacteriaceae"             
## [2547,] "Fusobacteriales"             "Leptotrichiaceae"             
## [2548,] "Micrococcales"               NA                             
## [2549,] "Selenomonadales"             "Veillonellaceae"              
## [2550,] "Clostridiales"               "Peptostreptococcaceae"        
## [2551,] "Selenomonadales"             "Veillonellaceae"              
## [2552,] "Bacteroidales"               "Prevotellaceae"               
## [2553,] "Fusobacteriales"             "Leptotrichiaceae"             
## [2554,] "Fusobacteriales"             "Leptotrichiaceae"             
## [2555,] "Fusobacteriales"             "Leptotrichiaceae"             
## [2556,] "Flavobacteriales"            "Flavobacteriaceae"            
## [2557,] "Pasteurellales"              "Pasteurellaceae"              
## [2558,] "Spirochaetales"              "Spirochaetaceae"              
## [2559,] "Bacteroidales"               NA                             
## [2560,] "Selenomonadales"             "Veillonellaceae"              
## [2561,] "Selenomonadales"             "Veillonellaceae"              
## [2562,] "Spirochaetales"              "Spirochaetaceae"              
## [2563,] "Spirochaetales"              "Spirochaetaceae"              
## [2564,] "Betaproteobacteriales"       "Neisseriaceae"                
## [2565,] "Clostridiales"               "Family_XI"                    
## [2566,] "Lactobacillales"             "Carnobacteriaceae"            
## [2567,] "Pasteurellales"              "Pasteurellaceae"              
## [2568,] "Lactobacillales"             "Streptococcaceae"             
## [2569,] "Bacteroidales"               "Prevotellaceae"               
## [2570,] "Selenomonadales"             "Veillonellaceae"              
## [2571,] "Bacteroidales"               "Prevotellaceae"               
## [2572,] "Bacteroidales"               "Prevotellaceae"               
## [2573,] "Pasteurellales"              "Pasteurellaceae"              
## [2574,] "Fusobacteriales"             "Leptotrichiaceae"             
## [2575,] "Saccharimonadales"           "Saccharimonadaceae"           
## [2576,] "Bacteroidales"               "Prevotellaceae"               
## [2577,] "Actinomycetales"             "Actinomycetaceae"             
## [2578,] "Fusobacteriales"             "Leptotrichiaceae"             
## [2579,] "Flavobacteriales"            "Flavobacteriaceae"            
## [2580,] "Bacteroidales"               "Porphyromonadaceae"           
## [2581,] "Bacteroidales"               "Porphyromonadaceae"           
## [2582,] "Fusobacteriales"             "Fusobacteriaceae"             
## [2583,] "Bacteroidales"               "Prevotellaceae"               
## [2584,] "Selenomonadales"             "Veillonellaceae"              
## [2585,] "Flavobacteriales"            "Flavobacteriaceae"            
## [2586,] "Saccharimonadales"           NA                             
## [2587,] "Bacteroidales"               "Rikenellaceae"                
## [2588,] "Clostridiales"               "Lachnospiraceae"              
## [2589,] "Clostridiales"               "Peptostreptococcaceae"        
## [2590,] "Bacteroidales"               "Prevotellaceae"               
## [2591,] "Campylobacterales"           NA                             
## [2592,] "Selenomonadales"             "Veillonellaceae"              
## [2593,] "Lactobacillales"             "Lactobacillaceae"             
## [2594,] "Betaproteobacteriales"       "Neisseriaceae"                
## [2595,] "Bacteroidales"               "Prevotellaceae"               
## [2596,] "Lactobacillales"             "Streptococcaceae"             
## [2597,] "Bacteroidales"               "Prevotellaceae"               
## [2598,] NA                            NA                             
## [2599,] "Sphingobacteriales"          "Lentimicrobiaceae"            
## [2600,] "Selenomonadales"             "Veillonellaceae"              
## [2601,] "Selenomonadales"             "Veillonellaceae"              
## [2602,] "Cardiobacteriales"           "Cardiobacteriaceae"           
## [2603,] "Bacteroidales"               "Prevotellaceae"               
## [2604,] "Selenomonadales"             "Veillonellaceae"              
## [2605,] "Bacteroidales"               "Prevotellaceae"               
## [2606,] "Betaproteobacteriales"       "Neisseriaceae"                
## [2607,] "Micrococcales"               NA                             
## [2608,] NA                            NA                             
## [2609,] NA                            NA                             
## [2610,] "Micrococcales"               NA                             
## [2611,] "Clostridiales"               "Peptostreptococcaceae"        
## [2612,] "Bacteroidales"               "Prevotellaceae"               
## [2613,] "Micrococcales"               NA                             
## [2614,] "Bacteroidales"               "Prevotellaceae"               
## [2615,] NA                            NA                             
## [2616,] "Bacteroidales"               "Prevotellaceae"               
## [2617,] "Pasteurellales"              "Pasteurellaceae"              
## [2618,] "Selenomonadales"             "Veillonellaceae"              
## [2619,] "Pasteurellales"              "Pasteurellaceae"              
## [2620,] "Fusobacteriales"             "Leptotrichiaceae"             
## [2621,] "Fusobacteriales"             "Fusobacteriaceae"             
## [2622,] "Bacteroidales"               "Prevotellaceae"               
## [2623,] "Flavobacteriales"            "Flavobacteriaceae"            
## [2624,] "Enterobacteriales"           "Enterobacteriaceae"           
## [2625,] "Clostridiales"               "Lachnospiraceae"              
## [2626,] "Absconditabacteriales_(SR1)" NA                             
## [2627,] "Selenomonadales"             "Veillonellaceae"              
## [2628,] "Lactobacillales"             "Carnobacteriaceae"            
## [2629,] "Corynebacteriales"           "Corynebacteriaceae"           
## [2630,] "Clostridiales"               "Lachnospiraceae"              
## [2631,] NA                            NA                             
## [2632,] "Spirochaetales"              "Spirochaetaceae"              
## [2633,] "Bacillales"                  "Family_XI"                    
## [2634,] "Bacteroidales"               "Prevotellaceae"               
## [2635,] "Campylobacterales"           NA                             
## [2636,] "Selenomonadales"             "Veillonellaceae"              
## [2637,] "Campylobacterales"           NA                             
## [2638,] "Campylobacterales"           NA                             
## [2639,] "Bacteroidales"               "Prevotellaceae"               
## [2640,] "Fusobacteriales"             "Leptotrichiaceae"             
## [2641,] "Campylobacterales"           "Campylobacteraceae"           
## [2642,] "Selenomonadales"             "Veillonellaceae"              
## [2643,] "Bacteroidales"               "Prevotellaceae"               
## [2644,] "Lactobacillales"             "Streptococcaceae"             
## [2645,] "Pasteurellales"              "Pasteurellaceae"              
## [2646,] "Clostridiales"               "Family_XI"                    
## [2647,] "Bacteroidales"               "Prevotellaceae"               
## [2648,] "Clostridiales"               "Ruminococcaceae"              
## [2649,] "Flavobacteriales"            "Flavobacteriaceae"            
## [2650,] "Lactobacillales"             "Streptococcaceae"             
## [2651,] "Spirochaetales"              "Spirochaetaceae"              
## [2652,] "Pasteurellales"              "Pasteurellaceae"              
## [2653,] "Fusobacteriales"             "Leptotrichiaceae"             
## [2654,] "Bacteroidales"               "Prevotellaceae"               
## [2655,] "Selenomonadales"             "Veillonellaceae"              
## [2656,] "Bacteroidales"               "Tannerellaceae"               
## [2657,] "Bacteroidales"               "Prevotellaceae"               
## [2658,] "Clostridiales"               "Peptostreptococcaceae"        
## [2659,] "Bacteroidales"               "Prevotellaceae"               
## [2660,] "Flavobacteriales"            "Flavobacteriaceae"            
## [2661,] "Flavobacteriales"            "Flavobacteriaceae"            
## [2662,] "Selenomonadales"             "Veillonellaceae"              
## [2663,] "Actinomycetales"             "Actinomycetaceae"             
## [2664,] "Actinomycetales"             "Actinomycetaceae"             
## [2665,] "Fusobacteriales"             "Leptotrichiaceae"             
## [2666,] "Selenomonadales"             "Veillonellaceae"              
## [2667,] NA                            NA                             
## [2668,] "Selenomonadales"             "Veillonellaceae"              
## [2669,] "Clostridiales"               "Family_XI"                    
## [2670,] "Flavobacteriales"            "Flavobacteriaceae"            
## [2671,] "Selenomonadales"             "Veillonellaceae"              
## [2672,] "Pasteurellales"              "Pasteurellaceae"              
## [2673,] "Bacteroidales"               "Prevotellaceae"               
## [2674,] "Bacteroidales"               "Prevotellaceae"               
## [2675,] "Spirochaetales"              "Spirochaetaceae"              
## [2676,] "Bacteroidales"               "Prevotellaceae"               
## [2677,] "Actinomycetales"             "Actinomycetaceae"             
## [2678,] "Mollicutes_RF39"             NA                             
## [2679,] "Enterobacteriales"           "Enterobacteriaceae"           
## [2680,] "Bacteroidales"               "Prevotellaceae"               
## [2681,] "Betaproteobacteriales"       "Burkholderiaceae"             
## [2682,] "Bacteroidales"               "Prevotellaceae"               
## [2683,] "Actinomycetales"             "Actinomycetaceae"             
## [2684,] "Flavobacteriales"            "Flavobacteriaceae"            
## [2685,] "Bacteroidales"               "Prevotellaceae"               
## [2686,] "Fusobacteriales"             "Leptotrichiaceae"             
## [2687,] "Coriobacteriales"            "Atopobiaceae"                 
## [2688,] "Saccharimonadales"           "Saccharimonadaceae"           
## [2689,] "Micrococcales"               NA                             
## [2690,] NA                            NA                             
## [2691,] "Clostridiales"               "Lachnospiraceae"              
## [2692,] "Fusobacteriales"             "Fusobacteriaceae"             
## [2693,] "Clostridiales"               "Clostridiales_vadinBB60_group"
## [2694,] "Bacteroidales"               "Prevotellaceae"               
## [2695,] "Bacteroidales"               "Prevotellaceae"               
## [2696,] NA                            NA                             
## [2697,] "Selenomonadales"             "Veillonellaceae"              
## [2698,] "Selenomonadales"             "Veillonellaceae"              
## [2699,] "Selenomonadales"             "Veillonellaceae"              
## [2700,] "Bacteroidales"               "Prevotellaceae"               
## [2701,] NA                            NA                             
## [2702,] "Selenomonadales"             "Veillonellaceae"              
## [2703,] "Sphingobacteriales"          "Lentimicrobiaceae"            
## [2704,] "Bacteroidales"               "Prevotellaceae"               
## [2705,] "Pasteurellales"              "Pasteurellaceae"              
## [2706,] "Fusobacteriales"             "Fusobacteriaceae"             
## [2707,] "Selenomonadales"             "Veillonellaceae"              
## [2708,] "Fusobacteriales"             "Fusobacteriaceae"             
## [2709,] "Selenomonadales"             "Veillonellaceae"              
## [2710,] "Bacteroidales"               "Prevotellaceae"               
## [2711,] "Flavobacteriales"            "Flavobacteriaceae"            
## [2712,] "Fusobacteriales"             "Fusobacteriaceae"             
## [2713,] "Selenomonadales"             "Veillonellaceae"              
## [2714,] "Fusobacteriales"             "Fusobacteriaceae"             
## [2715,] "Bacteroidales"               "Prevotellaceae"               
## [2716,] "Fusobacteriales"             "Leptotrichiaceae"             
## [2717,] "Fusobacteriales"             "Fusobacteriaceae"             
## [2718,] "Bacteroidales"               "Prevotellaceae"               
## [2719,] "Enterobacteriales"           "Enterobacteriaceae"           
## [2720,] "Clostridiales"               "Lachnospiraceae"              
## [2721,] "Spirochaetales"              "Spirochaetaceae"              
## [2722,] "Fusobacteriales"             "Leptotrichiaceae"             
## [2723,] "Clostridiales"               "Lachnospiraceae"              
## [2724,] "Bacteroidales"               "Prevotellaceae"               
## [2725,] "Bacteroidales"               "Prevotellaceae"               
## [2726,] "Betaproteobacteriales"       "Neisseriaceae"                
## [2727,] "Bacteroidales"               "Prevotellaceae"               
## [2728,] "Actinomycetales"             "Actinomycetaceae"             
## [2729,] "Bacteroidales"               "Prevotellaceae"               
## [2730,] "Campylobacterales"           NA                             
## [2731,] "Clostridiales"               "Lachnospiraceae"              
## [2732,] "Betaproteobacteriales"       "Neisseriaceae"                
## [2733,] "Bacteroidales"               "Prevotellaceae"               
## [2734,] "Fusobacteriales"             "Leptotrichiaceae"             
## [2735,] "Fusobacteriales"             "Fusobacteriaceae"             
## [2736,] "Fusobacteriales"             "Leptotrichiaceae"             
## [2737,] "Pasteurellales"              "Pasteurellaceae"              
## [2738,] "Bacteroidales"               "Prevotellaceae"               
## [2739,] "Micrococcales"               NA                             
## [2740,] NA                            NA                             
## [2741,] NA                            NA                             
## [2742,] "Bacteroidales"               "Prevotellaceae"               
## [2743,] "Erysipelotrichales"          "Erysipelotrichaceae"          
## [2744,] "Selenomonadales"             "Veillonellaceae"              
## [2745,] "Betaproteobacteriales"       "Neisseriaceae"                
## [2746,] "Betaproteobacteriales"       "Neisseriaceae"                
## [2747,] "Selenomonadales"             "Veillonellaceae"              
## [2748,] "Bacteroidales"               "Prevotellaceae"               
## [2749,] "Bacteroidales"               "Prevotellaceae"               
## [2750,] "Bacteroidales"               "Prevotellaceae"               
## [2751,] "Saccharimonadales"           "Saccharimonadaceae"           
## [2752,] "Bacteroidales"               "Prevotellaceae"               
## [2753,] "Bacteroidales"               "Tannerellaceae"               
## [2754,] "Bacteroidales"               "Prevotellaceae"               
## [2755,] "Betaproteobacteriales"       "Neisseriaceae"                
## [2756,] NA                            NA                             
## [2757,] "Flavobacteriales"            "Flavobacteriaceae"            
## [2758,] "Fusobacteriales"             "Fusobacteriaceae"             
## [2759,] "Bacteroidales"               "Prevotellaceae"               
## [2760,] "Coriobacteriales"            "Atopobiaceae"                 
## [2761,] "Bacteroidales"               "Prevotellaceae"               
## [2762,] "Bacteroidales"               "Porphyromonadaceae"           
## [2763,] "Bacteroidales"               "Prevotellaceae"               
## [2764,] "Micrococcales"               NA                             
## [2765,] NA                            NA                             
## [2766,] "Bacteroidales"               "Prevotellaceae"               
## [2767,] "Lactobacillales"             "Streptococcaceae"             
## [2768,] "Lactobacillales"             NA                             
## [2769,] "Clostridiales"               NA                             
## [2770,] "Selenomonadales"             "Veillonellaceae"              
## [2771,] "Saccharimonadales"           "Saccharimonadaceae"           
## [2772,] "Lactobacillales"             "Carnobacteriaceae"            
## [2773,] "Bacteroidales"               "Prevotellaceae"               
## [2774,] "Selenomonadales"             "Veillonellaceae"              
## [2775,] "Campylobacterales"           NA                             
## [2776,] "Corynebacteriales"           "Corynebacteriaceae"           
## [2777,] "Betaproteobacteriales"       "Neisseriaceae"                
## [2778,] "Corynebacteriales"           "Corynebacteriaceae"           
## [2779,] "Bacteroidales"               "Tannerellaceae"               
## [2780,] "Flavobacteriales"            "Weeksellaceae"                
## [2781,] NA                            NA                             
## [2782,] "Flavobacteriales"            "Flavobacteriaceae"            
## [2783,] "Clostridiales"               "Clostridiales_vadinBB60_group"
## [2784,] "Clostridiales"               "Family_XIII"                  
## [2785,] "Corynebacteriales"           "Corynebacteriaceae"           
## [2786,] "Actinomycetales"             "Actinomycetaceae"             
## [2787,] "Bacteroidales"               "Prevotellaceae"               
## [2788,] "Selenomonadales"             "Veillonellaceae"              
## [2789,] NA                            NA                             
## [2790,] "Bacteroidales"               "Prevotellaceae"               
## [2791,] "Bacteroidales"               "Prevotellaceae"               
## [2792,] "Flavobacteriales"            "Weeksellaceae"                
## [2793,] "Selenomonadales"             "Veillonellaceae"              
## [2794,] "Clostridiales"               NA                             
## [2795,] "Betaproteobacteriales"       "Neisseriaceae"                
## [2796,] "Bacteroidales"               "Prevotellaceae"               
## [2797,] "Bacteroidales"               "Prevotellaceae"               
## [2798,] "Flavobacteriales"            "Flavobacteriaceae"            
## [2799,] "Bacteroidales"               "Prevotellaceae"               
## [2800,] "Pasteurellales"              "Pasteurellaceae"              
## [2801,] "Pasteurellales"              "Pasteurellaceae"              
## [2802,] "Flavobacteriales"            "Flavobacteriaceae"            
## [2803,] "Bacteroidales"               "Prevotellaceae"               
## [2804,] "Selenomonadales"             "Veillonellaceae"              
## [2805,] "Clostridiales"               "Family_XIII"                  
## [2806,] "Bacteroidales"               "Prevotellaceae"               
## [2807,] "Bacteroidales"               "Prevotellaceae"               
## [2808,] "Selenomonadales"             "Veillonellaceae"              
## [2809,] "Fusobacteriales"             "Leptotrichiaceae"             
## [2810,] "Bacteroidales"               "Prevotellaceae"               
## [2811,] "Bacteroidales"               "Prevotellaceae"               
## [2812,] "Fusobacteriales"             "Fusobacteriaceae"             
## [2813,] "Pasteurellales"              "Pasteurellaceae"              
## [2814,] "Fusobacteriales"             "Leptotrichiaceae"             
## [2815,] "Bacteroidales"               "Prevotellaceae"               
## [2816,] "Bacteroidales"               "Prevotellaceae"               
## [2817,] "Bacteroidales"               "Prevotellaceae"               
## [2818,] "Selenomonadales"             "Veillonellaceae"              
## [2819,] "Clostridiales"               "Lachnospiraceae"              
## [2820,] "Bacteroidales"               "Prevotellaceae"               
## [2821,] "Selenomonadales"             "Veillonellaceae"              
## [2822,] "Bacteroidales"               "Prevotellaceae"               
## [2823,] "Bacteroidales"               "Prevotellaceae"               
## [2824,] "Bacteroidales"               "Prevotellaceae"               
## [2825,] "Flavobacteriales"            "Weeksellaceae"                
## [2826,] "Bacteroidales"               NA                             
## [2827,] "Selenomonadales"             "Veillonellaceae"              
## [2828,] "Selenomonadales"             "Veillonellaceae"              
## [2829,] "Saccharimonadales"           NA                             
## [2830,] "Clostridiales"               "Peptostreptococcaceae"        
## [2831,] "Saccharimonadales"           "Saccharimonadaceae"           
## [2832,] "Pasteurellales"              "Pasteurellaceae"              
## [2833,] "Pasteurellales"              "Pasteurellaceae"              
## [2834,] "Spirochaetales"              "Spirochaetaceae"              
## [2835,] "Selenomonadales"             "Veillonellaceae"              
## [2836,] "Bacteroidales"               "Prevotellaceae"               
## [2837,] "Selenomonadales"             "Veillonellaceae"              
## [2838,] "Fusobacteriales"             "Fusobacteriaceae"             
## [2839,] "Bacteroidales"               "Prevotellaceae"               
## [2840,] "Clostridiales"               "Family_XI"                    
## [2841,] "Clostridiales"               "Lachnospiraceae"              
## [2842,] "Selenomonadales"             "Veillonellaceae"              
## [2843,] "Betaproteobacteriales"       "Burkholderiaceae"             
## [2844,] "Selenomonadales"             "Veillonellaceae"              
## [2845,] "Bacteroidales"               "Prevotellaceae"               
## [2846,] "Selenomonadales"             "Veillonellaceae"              
## [2847,] "Pasteurellales"              "Pasteurellaceae"              
## [2848,] "Lactobacillales"             "Carnobacteriaceae"            
## [2849,] "Selenomonadales"             "Veillonellaceae"              
## [2850,] "Bacteroidales"               "Prevotellaceae"               
## [2851,] "Betaproteobacteriales"       "Neisseriaceae"                
## [2852,] "Fusobacteriales"             "Fusobacteriaceae"             
## [2853,] "Selenomonadales"             "Veillonellaceae"              
## [2854,] "Bacteroidales"               "Prevotellaceae"               
## [2855,] "Betaproteobacteriales"       "Neisseriaceae"                
## [2856,] "Actinomycetales"             "Actinomycetaceae"             
## [2857,] "Betaproteobacteriales"       "Neisseriaceae"                
## [2858,] "Bacteroidales"               "Porphyromonadaceae"           
## [2859,] "Bacteroidales"               "Prevotellaceae"               
## [2860,] "Betaproteobacteriales"       "Neisseriaceae"                
## [2861,] "Selenomonadales"             "Veillonellaceae"              
## [2862,] "Selenomonadales"             "Veillonellaceae"              
## [2863,] "Flavobacteriales"            "Weeksellaceae"                
## [2864,] "Micrococcales"               NA                             
## [2865,] "Bacteroidales"               "Tannerellaceae"               
## [2866,] "Mollicutes_RF39"             NA                             
## [2867,] "Sphingobacteriales"          "Lentimicrobiaceae"            
## [2868,] "Clostridiales"               "Lachnospiraceae"              
## [2869,] "Micrococcales"               NA                             
## [2870,] "Clostridiales"               "Lachnospiraceae"              
## [2871,] "Bacteroidales"               "Prevotellaceae"               
## [2872,] NA                            NA                             
## [2873,] "Pasteurellales"              "Pasteurellaceae"              
## [2874,] "Bacteroidales"               "Prevotellaceae"               
## [2875,] "Flavobacteriales"            "Flavobacteriaceae"            
## [2876,] "Bacteroidales"               "Prevotellaceae"               
## [2877,] "Enterobacteriales"           "Enterobacteriaceae"           
## [2878,] "Bacteroidales"               "Prevotellaceae"               
## [2879,] "Lactobacillales"             "Streptococcaceae"             
## [2880,] NA                            NA                             
## [2881,] "Sphingobacteriales"          "Lentimicrobiaceae"            
## [2882,] "Cardiobacteriales"           "Cardiobacteriaceae"           
## [2883,] "Selenomonadales"             "Veillonellaceae"              
## [2884,] "Clostridiales"               "Lachnospiraceae"              
## [2885,] "Bacteroidales"               "Prevotellaceae"               
## [2886,] "Clostridiales"               "Peptostreptococcaceae"        
## [2887,] "Clostridiales"               "Lachnospiraceae"              
## [2888,] "Fusobacteriales"             NA                             
## [2889,] "Flavobacteriales"            "Flavobacteriaceae"            
## [2890,] "Bacteroidales"               "Prevotellaceae"               
## [2891,] "Betaproteobacteriales"       "Neisseriaceae"                
## [2892,] "Bacteroidales"               "Prevotellaceae"               
## [2893,] "Bacteroidales"               "Prevotellaceae"               
## [2894,] "Pasteurellales"              "Pasteurellaceae"              
## [2895,] NA                            NA                             
## [2896,] "Fusobacteriales"             "Leptotrichiaceae"             
## [2897,] "Bacteroidales"               "Prevotellaceae"               
## [2898,] "Bacteroidales"               "Prevotellaceae"               
## [2899,] "Saccharimonadales"           NA                             
## [2900,] "Fusobacteriales"             "Fusobacteriaceae"             
## [2901,] "Bacteroidales"               "Prevotellaceae"               
## [2902,] "Selenomonadales"             "Veillonellaceae"              
## [2903,] "Actinomycetales"             "Actinomycetaceae"             
## [2904,] "Bacteroidales"               "Prevotellaceae"               
## [2905,] "Spirochaetales"              "Spirochaetaceae"              
## [2906,] "Pasteurellales"              "Pasteurellaceae"              
## [2907,] "Spirochaetales"              "Spirochaetaceae"              
## [2908,] "Campylobacterales"           "Campylobacteraceae"           
## [2909,] "Clostridiales"               "Lachnospiraceae"              
## [2910,] "Oceanospirillales"           "Halomonadaceae"               
## [2911,] "Selenomonadales"             "Veillonellaceae"              
## [2912,] "Mycoplasmatales"             "Mycoplasmataceae"             
## [2913,] "Bacteroidales"               "Prevotellaceae"               
## [2914,] NA                            NA                             
## [2915,] "Saccharimonadales"           "Saccharimonadaceae"           
## [2916,] "Flavobacteriales"            "Flavobacteriaceae"            
## [2917,] "Saccharimonadales"           "Saccharimonadaceae"           
## [2918,] "Selenomonadales"             "Veillonellaceae"              
## [2919,] "Fusobacteriales"             "Fusobacteriaceae"             
## [2920,] "Selenomonadales"             "Veillonellaceae"              
## [2921,] "Bacteroidales"               "Porphyromonadaceae"           
## [2922,] "Actinomycetales"             "Actinomycetaceae"             
## [2923,] "Betaproteobacteriales"       NA                             
## [2924,] "Lactobacillales"             "Streptococcaceae"             
## [2925,] "Clostridiales"               "Lachnospiraceae"              
## [2926,] "Fusobacteriales"             "Leptotrichiaceae"             
## [2927,] "Betaproteobacteriales"       "Neisseriaceae"                
## [2928,] "Lactobacillales"             "Streptococcaceae"             
## [2929,] "Actinomycetales"             "Actinomycetaceae"             
## [2930,] "Micrococcales"               NA                             
## [2931,] "Clostridiales"               "Ruminococcaceae"              
## [2932,] "Pasteurellales"              "Pasteurellaceae"              
## [2933,] "Lactobacillales"             "Carnobacteriaceae"            
## [2934,] "Coriobacteriales"            "Atopobiaceae"                 
## [2935,] "Actinomycetales"             "Actinomycetaceae"             
## [2936,] "Selenomonadales"             "Veillonellaceae"              
## [2937,] NA                            NA                             
## [2938,] "Fusobacteriales"             "Fusobacteriaceae"             
## [2939,] "Bacteroidales"               "Prevotellaceae"               
## [2940,] "Bacteroidales"               "Prevotellaceae"               
## [2941,] "Micrococcales"               NA                             
## [2942,] "Saccharimonadales"           "Saccharimonadaceae"           
## [2943,] "Bacteroidales"               "Prevotellaceae"               
## [2944,] "Bacillales"                  "Staphylococcaceae"            
## [2945,] "Flavobacteriales"            "Flavobacteriaceae"            
## [2946,] "Sphingobacteriales"          "Lentimicrobiaceae"            
## [2947,] "Bacteroidales"               "Tannerellaceae"               
## [2948,] NA                            NA                             
## [2949,] "Clostridiales"               "Lachnospiraceae"              
## [2950,] "Selenomonadales"             "Veillonellaceae"              
## [2951,] "Selenomonadales"             "Veillonellaceae"              
## [2952,] "Bacteroidales"               "Prevotellaceae"               
## [2953,] "Bacteroidales"               "Prevotellaceae"               
## [2954,] "Flavobacteriales"            "Flavobacteriaceae"            
## [2955,] "Clostridiales"               "Lachnospiraceae"              
## [2956,] "Chloroplast"                 NA                             
## [2957,] "Pasteurellales"              "Pasteurellaceae"              
## [2958,] "Fusobacteriales"             "Leptotrichiaceae"             
## [2959,] "Selenomonadales"             "Veillonellaceae"              
## [2960,] "Selenomonadales"             "Veillonellaceae"              
## [2961,] "Bacteroidales"               "Prevotellaceae"               
## [2962,] "Pasteurellales"              "Pasteurellaceae"              
## [2963,] "Selenomonadales"             "Veillonellaceae"              
## [2964,] "Selenomonadales"             "Veillonellaceae"              
## [2965,] "Betaproteobacteriales"       "Neisseriaceae"                
## [2966,] "Selenomonadales"             "Veillonellaceae"              
## [2967,] "Selenomonadales"             "Veillonellaceae"              
## [2968,] NA                            NA                             
## [2969,] "Bacteroidales"               "Prevotellaceae"               
## [2970,] "Fusobacteriales"             "Leptotrichiaceae"             
## [2971,] "Betaproteobacteriales"       "Neisseriaceae"                
## [2972,] "JGI_0000069-P22"             NA                             
## [2973,] "Bacteroidales"               "Prevotellaceae"               
## [2974,] "Bacteroidales"               "Rikenellaceae"                
## [2975,] "Bacillales"                  "Family_XI"                    
## [2976,] "Spirochaetales"              "Spirochaetaceae"              
## [2977,] "Clostridiales"               "Lachnospiraceae"              
## [2978,] "Mycoplasmatales"             "Mycoplasmataceae"             
## [2979,] "Selenomonadales"             "Veillonellaceae"              
## [2980,] "Bacteroidales"               "Prevotellaceae"               
## [2981,] "Enterobacteriales"           "Enterobacteriaceae"           
## [2982,] "Spirochaetales"              "Spirochaetaceae"              
## [2983,] "Campylobacterales"           "P6-b51"                       
## [2984,] "Bacteroidales"               "Porphyromonadaceae"           
## [2985,] "Clostridiales"               "Lachnospiraceae"              
## [2986,] "Pasteurellales"              "Pasteurellaceae"              
## [2987,] "Actinomycetales"             "Actinomycetaceae"             
## [2988,] "Bacteroidales"               "Prevotellaceae"               
## [2989,] "Flavobacteriales"            "Flavobacteriaceae"            
## [2990,] "Pasteurellales"              "Pasteurellaceae"              
## [2991,] "Selenomonadales"             "Veillonellaceae"              
## [2992,] "Synergistales"               "Synergistaceae"               
## [2993,] "Lactobacillales"             "Streptococcaceae"             
## [2994,] "Bifidobacteriales"           "Bifidobacteriaceae"           
## [2995,] "Bacteroidales"               "Prevotellaceae"               
## [2996,] "Clostridiales"               "Lachnospiraceae"              
## [2997,] "Selenomonadales"             "Veillonellaceae"              
## [2998,] "Saccharimonadales"           NA                             
## [2999,] "Enterobacteriales"           "Enterobacteriaceae"           
## [3000,] "Selenomonadales"             "Veillonellaceae"              
## [3001,] "Flavobacteriales"            "Flavobacteriaceae"            
## [3002,] "Fusobacteriales"             "Leptotrichiaceae"             
## [3003,] NA                            NA                             
## [3004,] "Bacteroidales"               "Prevotellaceae"               
## [3005,] "Betaproteobacteriales"       "Neisseriaceae"                
## [3006,] "Pasteurellales"              "Pasteurellaceae"              
## [3007,] "Selenomonadales"             "Veillonellaceae"              
## [3008,] "Corynebacteriales"           "Corynebacteriaceae"           
## [3009,] "Clostridiales"               "Lachnospiraceae"              
## [3010,] "Selenomonadales"             "Veillonellaceae"              
## [3011,] "Fusobacteriales"             "Fusobacteriaceae"             
## [3012,] "Selenomonadales"             "Veillonellaceae"              
## [3013,] NA                            NA                             
## [3014,] "Bacteroidales"               "Prevotellaceae"               
## [3015,] "Spirochaetales"              "Spirochaetaceae"              
## [3016,] NA                            NA                             
## [3017,] "Pasteurellales"              "Pasteurellaceae"              
## [3018,] "Selenomonadales"             "Veillonellaceae"              
## [3019,] "Bacteroidales"               "Porphyromonadaceae"           
## [3020,] NA                            NA                             
## [3021,] "Bacteroidales"               "Prevotellaceae"               
## [3022,] "Clostridiales"               "Peptostreptococcaceae"        
## [3023,] "Bacteroidales"               "Prevotellaceae"               
## [3024,] "Bacteroidales"               "Prevotellaceae"               
## [3025,] "Bacteroidales"               "Prevotellaceae"               
## [3026,] "Absconditabacteriales_(SR1)" NA                             
## [3027,] "Bacteroidales"               "Prevotellaceae"               
## [3028,] "Absconditabacteriales_(SR1)" NA                             
## [3029,] "Fusobacteriales"             "Fusobacteriaceae"             
## [3030,] "Flavobacteriales"            "Flavobacteriaceae"            
## [3031,] "Bacteroidales"               "Prevotellaceae"               
## [3032,] "Betaproteobacteriales"       "Neisseriaceae"                
## [3033,] "Bacteroidales"               "Prevotellaceae"               
## [3034,] NA                            NA                             
## [3035,] "Lactobacillales"             "Carnobacteriaceae"            
## [3036,] "Clostridiales"               "Lachnospiraceae"              
## [3037,] "Clostridiales"               "Lachnospiraceae"              
## [3038,] "Bacteroidales"               "Prevotellaceae"               
## [3039,] "Actinomycetales"             "Actinomycetaceae"             
## [3040,] "Lactobacillales"             "Streptococcaceae"             
## [3041,] "Campylobacterales"           NA                             
## [3042,] "Bacteroidales"               "Prevotellaceae"               
## [3043,] "Selenomonadales"             "Veillonellaceae"              
## [3044,] "Selenomonadales"             "Veillonellaceae"              
## [3045,] "Flavobacteriales"            "Flavobacteriaceae"            
## [3046,] "Betaproteobacteriales"       NA                             
## [3047,] "Actinomycetales"             "Actinomycetaceae"             
## [3048,] "Pasteurellales"              "Pasteurellaceae"              
## [3049,] "Absconditabacteriales_(SR1)" NA                             
## [3050,] "Clostridiales"               "Lachnospiraceae"              
## [3051,] "Bacteroidales"               "Prevotellaceae"               
## [3052,] "Pseudomonadales"             "Moraxellaceae"                
## [3053,] "Fusobacteriales"             "Fusobacteriaceae"             
## [3054,] "Fusobacteriales"             "Fusobacteriaceae"             
## [3055,] "Clostridiales"               "Peptostreptococcaceae"        
## [3056,] "Bacteroidales"               "Porphyromonadaceae"           
## [3057,] "Bacteroidales"               "Prevotellaceae"               
## [3058,] "Fusobacteriales"             "Leptotrichiaceae"             
## [3059,] "Bacteroidales"               "Prevotellaceae"               
## [3060,] "Flavobacteriales"            "Flavobacteriaceae"            
## [3061,] "Bacteroidales"               "Porphyromonadaceae"           
## [3062,] "Fusobacteriales"             "Leptotrichiaceae"             
## [3063,] "Selenomonadales"             "Veillonellaceae"              
## [3064,] "Clostridiales"               "Lachnospiraceae"              
## [3065,] "Bifidobacteriales"           "Bifidobacteriaceae"           
## [3066,] "Clostridiales"               "Lachnospiraceae"              
## [3067,] "Selenomonadales"             "Veillonellaceae"              
## [3068,] "Selenomonadales"             "Veillonellaceae"              
## [3069,] "Saccharimonadales"           "Saccharimonadaceae"           
## [3070,] "Micrococcales"               NA                             
## [3071,] "Clostridiales"               "Family_XIII"                  
## [3072,] "Selenomonadales"             "Veillonellaceae"              
## [3073,] "Flavobacteriales"            "Flavobacteriaceae"            
## [3074,] "Fusobacteriales"             "Leptotrichiaceae"             
## [3075,] "Bacteroidales"               "Prevotellaceae"               
## [3076,] "Betaproteobacteriales"       "Burkholderiaceae"             
## [3077,] NA                            NA                             
## [3078,] "Fusobacteriales"             "Leptotrichiaceae"             
## [3079,] "Betaproteobacteriales"       "Neisseriaceae"                
## [3080,] "Bacteroidales"               "Porphyromonadaceae"           
## [3081,] "Spirochaetales"              "Spirochaetaceae"              
## [3082,] "Actinomycetales"             "Actinomycetaceae"             
## [3083,] "Micrococcales"               "Micrococcaceae"               
## [3084,] "Bacteroidales"               "Prevotellaceae"               
## [3085,] "Fusobacteriales"             "Fusobacteriaceae"             
## [3086,] NA                            NA                             
## [3087,] "Bacteroidales"               "Prevotellaceae"               
## [3088,] "Betaproteobacteriales"       "Neisseriaceae"                
## [3089,] "Saccharimonadales"           "Saccharimonadaceae"           
## [3090,] "Selenomonadales"             "Veillonellaceae"              
## [3091,] "Selenomonadales"             "Veillonellaceae"              
## [3092,] "Corynebacteriales"           "Corynebacteriaceae"           
## [3093,] "Fusobacteriales"             "Fusobacteriaceae"             
## [3094,] "Actinomycetales"             "Actinomycetaceae"             
## [3095,] "Pasteurellales"              "Pasteurellaceae"              
## [3096,] "Bacteroidales"               "Prevotellaceae"               
## [3097,] "Selenomonadales"             "Veillonellaceae"              
## [3098,] "Selenomonadales"             "Veillonellaceae"              
## [3099,] NA                            NA                             
## [3100,] "Bacteroidales"               "Prevotellaceae"               
## [3101,] "Bacteroidales"               "Prevotellaceae"               
## [3102,] "Spirochaetales"              "Spirochaetaceae"              
## [3103,] NA                            NA                             
## [3104,] "Sphingobacteriales"          "Lentimicrobiaceae"            
## [3105,] NA                            NA                             
## [3106,] "Actinomycetales"             "Actinomycetaceae"             
## [3107,] "Bacteroidales"               "Prevotellaceae"               
## [3108,] "Bacteroidales"               "Prevotellaceae"               
## [3109,] "Pasteurellales"              "Pasteurellaceae"              
## [3110,] "Micrococcales"               NA                             
## [3111,] NA                            NA                             
## [3112,] "Fusobacteriales"             "Leptotrichiaceae"             
## [3113,] "Selenomonadales"             "Veillonellaceae"              
## [3114,] "Clostridiales"               "Lachnospiraceae"              
## [3115,] "Pasteurellales"              "Pasteurellaceae"              
## [3116,] "Clostridiales"               "Family_XIII"                  
## [3117,] "Lactobacillales"             "Streptococcaceae"             
## [3118,] "Flavobacteriales"            "Weeksellaceae"                
## [3119,] "Fusobacteriales"             "Leptotrichiaceae"             
## [3120,] "Bacteroidales"               "Prevotellaceae"               
## [3121,] "Bacteroidales"               "Prevotellaceae"               
## [3122,] "Micrococcales"               NA                             
## [3123,] "Actinomycetales"             "Actinomycetaceae"             
## [3124,] "Bacteroidales"               "Prevotellaceae"               
## [3125,] "Bacteroidales"               "Prevotellaceae"               
## [3126,] "Clostridiales"               "Defluviitaleaceae"            
## [3127,] "Lactobacillales"             "Carnobacteriaceae"            
## [3128,] "Bacteroidales"               "Prevotellaceae"               
## [3129,] "Micrococcales"               NA                             
## [3130,] "Actinomycetales"             "Actinomycetaceae"             
## [3131,] "Selenomonadales"             "Veillonellaceae"              
## [3132,] "Bacteroidales"               "Prevotellaceae"               
## [3133,] "Flavobacteriales"            "Flavobacteriaceae"            
## [3134,] "Pasteurellales"              "Pasteurellaceae"              
## [3135,] "Bacteroidales"               "Prevotellaceae"               
## [3136,] NA                            NA                             
## [3137,] "Campylobacterales"           NA                             
## [3138,] "Flavobacteriales"            "Flavobacteriaceae"            
## [3139,] "Flavobacteriales"            "Flavobacteriaceae"            
## [3140,] "Bacteroidales"               "Porphyromonadaceae"           
## [3141,] "Bacteroidales"               "Prevotellaceae"               
## [3142,] "Bacteroidales"               "Prevotellaceae"               
## [3143,] "Bacteroidales"               "Prevotellaceae"               
## [3144,] "Pasteurellales"              "Pasteurellaceae"              
## [3145,] "Saccharimonadales"           "Saccharimonadaceae"           
## [3146,] "Fusobacteriales"             "Leptotrichiaceae"             
## [3147,] "Fusobacteriales"             "Leptotrichiaceae"             
## [3148,] "Micrococcales"               NA                             
## [3149,] NA                            NA                             
## [3150,] "Spirochaetales"              "Spirochaetaceae"              
## [3151,] "Clostridiales"               "Ruminococcaceae"              
## [3152,] NA                            NA                             
## [3153,] "Spirochaetales"              "Spirochaetaceae"              
## [3154,] "Bacteroidales"               "Prevotellaceae"               
## [3155,] "Bacteroidales"               "Prevotellaceae"               
## [3156,] "Bacteroidales"               "Prevotellaceae"               
## [3157,] "Selenomonadales"             "Veillonellaceae"              
## [3158,] "Clostridiales"               "Lachnospiraceae"              
## [3159,] "Betaproteobacteriales"       "Neisseriaceae"                
## [3160,] "Bacteroidales"               "Tannerellaceae"               
## [3161,] "Campylobacterales"           NA                             
## [3162,] "Cardiobacteriales"           "Cardiobacteriaceae"           
## [3163,] "Bacteroidales"               "Porphyromonadaceae"           
## [3164,] "Bacteroidales"               "Prevotellaceae"               
## [3165,] "Clostridiales"               "Lachnospiraceae"              
## [3166,] "Pasteurellales"              "Pasteurellaceae"              
## [3167,] "Bacteroidales"               "Prevotellaceae"               
## [3168,] "Spirochaetales"              "Spirochaetaceae"              
## [3169,] "Spirochaetales"              "Spirochaetaceae"              
## [3170,] "Fusobacteriales"             "Leptotrichiaceae"             
## [3171,] "Campylobacterales"           "Campylobacteraceae"           
## [3172,] "Mollicutes_RF39"             NA                             
## [3173,] "Actinomycetales"             "Actinomycetaceae"             
## [3174,] "Fusobacteriales"             "Fusobacteriaceae"             
## [3175,] "Chloroplast"                 NA                             
## [3176,] "Campylobacterales"           NA                             
## [3177,] "Bacteroidales"               "Prevotellaceae"               
## [3178,] NA                            NA                             
## [3179,] "Fusobacteriales"             "Leptotrichiaceae"             
## [3180,] "Betaproteobacteriales"       "Neisseriaceae"                
## [3181,] "Flavobacteriales"            "Flavobacteriaceae"            
## [3182,] "Spirochaetales"              "Spirochaetaceae"              
## [3183,] "Fusobacteriales"             "Leptotrichiaceae"             
## [3184,] "Bacteroidales"               "Prevotellaceae"               
## [3185,] "Fusobacteriales"             "Fusobacteriaceae"             
## [3186,] "Bacteroidales"               "Rikenellaceae"                
## [3187,] "Betaproteobacteriales"       "Neisseriaceae"                
## [3188,] "Saccharimonadales"           NA                             
## [3189,] "Flavobacteriales"            "Flavobacteriaceae"            
## [3190,] "Bacteroidales"               "Prevotellaceae"               
## [3191,] "Spirochaetales"              "Spirochaetaceae"              
## [3192,] "Cardiobacteriales"           "Cardiobacteriaceae"           
## [3193,] "Bacteroidales"               "Prevotellaceae"               
## [3194,] "Fusobacteriales"             "Leptotrichiaceae"             
## [3195,] "Pasteurellales"              "Pasteurellaceae"              
## [3196,] "Fusobacteriales"             "Leptotrichiaceae"             
## [3197,] "Bacteroidales"               "Prevotellaceae"               
## [3198,] "Bacteroidales"               "Prevotellaceae"               
## [3199,] "Bacteroidales"               "Prevotellaceae"               
## [3200,] "Campylobacterales"           NA                             
## [3201,] "Pasteurellales"              "Pasteurellaceae"              
## [3202,] "Clostridiales"               "Family_XI"                    
## [3203,] "Campylobacterales"           "P6-b51"                       
## [3204,] "Saccharimonadales"           NA                             
## [3205,] "Micrococcales"               NA                             
## [3206,] "Bacteroidales"               "Prevotellaceae"               
## [3207,] "Flavobacteriales"            "Flavobacteriaceae"            
## [3208,] "Bacteroidales"               "Prevotellaceae"               
## [3209,] NA                            NA                             
## [3210,] "Bacteroidales"               "Prevotellaceae"               
## [3211,] "Corynebacteriales"           "Corynebacteriaceae"           
## [3212,] "Bacteroidales"               "Prevotellaceae"               
## [3213,] "Clostridiales"               "Lachnospiraceae"              
## [3214,] "Clostridiales"               "Peptostreptococcaceae"        
## [3215,] "Bacteroidales"               "Prevotellaceae"               
## [3216,] "Bacteroidales"               "Prevotellaceae"               
## [3217,] "Fusobacteriales"             "Leptotrichiaceae"             
## [3218,] "Bacteroidales"               "Prevotellaceae"               
## [3219,] "Bacteroidales"               "Bacteroidaceae"               
## [3220,] "Selenomonadales"             "Veillonellaceae"              
## [3221,] "Clostridiales"               "Ruminococcaceae"              
## [3222,] "Clostridiales"               "Lachnospiraceae"              
## [3223,] "Lactobacillales"             "Carnobacteriaceae"            
## [3224,] "Clostridiales"               "Lachnospiraceae"              
## [3225,] "Clostridiales"               "Ruminococcaceae"              
## [3226,] "Actinomycetales"             "Actinomycetaceae"             
## [3227,] "Fusobacteriales"             "Fusobacteriaceae"             
## [3228,] "Fusobacteriales"             "Leptotrichiaceae"             
## [3229,] "Selenomonadales"             "Veillonellaceae"              
## [3230,] "Fusobacteriales"             "Leptotrichiaceae"             
## [3231,] NA                            NA                             
## [3232,] "Bacteroidales"               "Prevotellaceae"               
## [3233,] "Fusobacteriales"             "Leptotrichiaceae"             
## [3234,] "Pasteurellales"              "Pasteurellaceae"              
## [3235,] "Bacteroidales"               "Prevotellaceae"               
## [3236,] NA                            NA                             
## [3237,] "Chloroplast"                 NA                             
## [3238,] "Selenomonadales"             "Veillonellaceae"              
## [3239,] "Lactobacillales"             "Streptococcaceae"             
## [3240,] "Bacteroidales"               "Prevotellaceae"               
## [3241,] "Bacteroidales"               "Prevotellaceae"               
## [3242,] "Selenomonadales"             "Veillonellaceae"              
## [3243,] "Bacteroidales"               NA                             
## [3244,] "Clostridiales"               "Lachnospiraceae"              
## [3245,] "Micrococcales"               "Micrococcaceae"               
## [3246,] "Clostridiales"               "Lachnospiraceae"              
## [3247,] "Campylobacterales"           NA                             
## [3248,] "Coriobacteriales"            "Coriobacteriaceae"            
## [3249,] "Bacteroidales"               "Prevotellaceae"               
## [3250,] "Saccharimonadales"           "Saccharimonadaceae"           
## [3251,] "Coriobacteriales"            "Atopobiaceae"                 
## [3252,] "Micrococcales"               NA                             
## [3253,] "Bacteroidales"               "Prevotellaceae"               
## [3254,] "Sphingobacteriales"          "Lentimicrobiaceae"            
## [3255,] "Betaproteobacteriales"       "Burkholderiaceae"             
## [3256,] "Bacteroidales"               "Porphyromonadaceae"           
## [3257,] "Saccharimonadales"           "Saccharimonadaceae"           
## [3258,] "Fusobacteriales"             "Leptotrichiaceae"             
## [3259,] "Lactobacillales"             "Streptococcaceae"             
## [3260,] "Campylobacterales"           "Campylobacteraceae"           
## [3261,] "Campylobacterales"           "Campylobacteraceae"           
## [3262,] "Micrococcales"               NA                             
## [3263,] "Saccharimonadales"           "Saccharimonadaceae"           
## [3264,] "Fusobacteriales"             "Leptotrichiaceae"             
## [3265,] "Micrococcales"               NA                             
## [3266,] "Bacteroidales"               "Prevotellaceae"               
## [3267,] "Bacteroidales"               "Prevotellaceae"               
## [3268,] "Lactobacillales"             "Carnobacteriaceae"            
## [3269,] "Bacteroidales"               "Prevotellaceae"               
## [3270,] "Fusobacteriales"             NA                             
## [3271,] "Bacteroidales"               "Prevotellaceae"               
## [3272,] "Bacteroidales"               "Prevotellaceae"               
## [3273,] "Bacteroidales"               "Prevotellaceae"               
## [3274,] "Saccharimonadales"           "Saccharimonadaceae"           
## [3275,] "Bacteroidales"               "Prevotellaceae"               
## [3276,] "Actinomycetales"             "Actinomycetaceae"             
## [3277,] NA                            NA                             
## [3278,] "Lactobacillales"             "Carnobacteriaceae"            
## [3279,] "Spirochaetales"              "Spirochaetaceae"              
## [3280,] "Bacteroidales"               "Prevotellaceae"               
## [3281,] "Selenomonadales"             "Veillonellaceae"              
## [3282,] "Cardiobacteriales"           "Cardiobacteriaceae"           
## [3283,] "Bacteroidales"               "Prevotellaceae"               
## [3284,] "Bacteroidales"               "Prevotellaceae"               
## [3285,] "Pasteurellales"              "Pasteurellaceae"              
## [3286,] "Pasteurellales"              "Pasteurellaceae"              
## [3287,] "Pasteurellales"              "Pasteurellaceae"              
## [3288,] "Spirochaetales"              "Spirochaetaceae"              
## [3289,] "Clostridiales"               "Lachnospiraceae"              
## [3290,] "Spirochaetales"              "Spirochaetaceae"              
## [3291,] "Bacteroidales"               "Prevotellaceae"               
## [3292,] "Pasteurellales"              "Pasteurellaceae"              
## [3293,] "Selenomonadales"             "Veillonellaceae"              
## [3294,] "Bacteroidales"               "Prevotellaceae"               
## [3295,] "Bacteroidales"               "Prevotellaceae"               
## [3296,] "Bacteroidales"               "Prevotellaceae"               
## [3297,] "Actinomycetales"             "Actinomycetaceae"             
## [3298,] "Bacteroidales"               "Prevotellaceae"               
## [3299,] "Lactobacillales"             "Streptococcaceae"             
## [3300,] "Bacteroidales"               "Prevotellaceae"               
## [3301,] "Bacteroidales"               "Prevotellaceae"               
## [3302,] "Flavobacteriales"            "Weeksellaceae"                
## [3303,] "Bacteroidales"               "Tannerellaceae"               
## [3304,] "Betaproteobacteriales"       "Neisseriaceae"                
## [3305,] "Betaproteobacteriales"       "Neisseriaceae"                
## [3306,] "Bacteroidales"               "Prevotellaceae"               
## [3307,] "Bacteroidales"               "Prevotellaceae"               
## [3308,] "Saccharimonadales"           "Saccharimonadaceae"           
## [3309,] "Pasteurellales"              "Pasteurellaceae"              
## [3310,] "Bacteroidales"               "Prevotellaceae"               
## [3311,] "Bacteroidales"               "Prevotellaceae"               
## [3312,] "Fusobacteriales"             "Leptotrichiaceae"             
## [3313,] "Flavobacteriales"            "Flavobacteriaceae"            
## [3314,] "Spirochaetales"              "Spirochaetaceae"              
## [3315,] "Bacteroidales"               "Prevotellaceae"               
## [3316,] "Saccharimonadales"           "Saccharimonadaceae"           
## [3317,] "Clostridiales"               "Lachnospiraceae"              
## [3318,] "Fusobacteriales"             "Leptotrichiaceae"             
## [3319,] "Betaproteobacteriales"       "Neisseriaceae"                
## [3320,] "Bacteroidales"               "Prevotellaceae"               
## [3321,] "Bacteroidales"               "Prevotellaceae"               
## [3322,] "Fusobacteriales"             "Leptotrichiaceae"             
## [3323,] "Bacteroidales"               "Prevotellaceae"               
## [3324,] "Bacteroidales"               "Prevotellaceae"               
## [3325,] "Bacteroidales"               "Prevotellaceae"               
## [3326,] NA                            NA                             
## [3327,] "Corynebacteriales"           "Corynebacteriaceae"           
## [3328,] "Flavobacteriales"            "Weeksellaceae"                
## [3329,] "Bacteroidales"               "Prevotellaceae"               
## [3330,] "Spirochaetales"              "Spirochaetaceae"              
## [3331,] "Bacteroidales"               "Porphyromonadaceae"           
## [3332,] "Bacteroidales"               "Prevotellaceae"               
## [3333,] "Selenomonadales"             "Veillonellaceae"              
## [3334,] "Micrococcales"               NA                             
## [3335,] "Bacillales"                  "Family_XI"                    
## [3336,] "Cardiobacteriales"           "Cardiobacteriaceae"           
## [3337,] "Bacteroidales"               "Prevotellaceae"               
## [3338,] "Fusobacteriales"             "Leptotrichiaceae"             
## [3339,] "Saccharimonadales"           "Saccharimonadaceae"           
## [3340,] "Saccharimonadales"           "Saccharimonadaceae"           
## [3341,] "Clostridiales"               "Family_XIII"                  
## [3342,] "Fusobacteriales"             "Fusobacteriaceae"             
## [3343,] NA                            NA                             
## [3344,] "Clostridiales"               "Lachnospiraceae"              
## [3345,] "Saccharimonadales"           "Saccharimonadaceae"           
## [3346,] "Campylobacterales"           NA                             
## [3347,] "Bacteroidales"               "Porphyromonadaceae"           
## [3348,] "Betaproteobacteriales"       "Neisseriaceae"                
## [3349,] "Flavobacteriales"            "Weeksellaceae"                
## [3350,] "Fusobacteriales"             "Leptotrichiaceae"             
## [3351,] "Pasteurellales"              "Pasteurellaceae"              
## [3352,] "Betaproteobacteriales"       "Neisseriaceae"                
## [3353,] "Bacteroidales"               "Prevotellaceae"               
## [3354,] "Betaproteobacteriales"       "Neisseriaceae"                
## [3355,] "Betaproteobacteriales"       "Neisseriaceae"                
## [3356,] "Selenomonadales"             "Veillonellaceae"              
## [3357,] "Coriobacteriales"            "Atopobiaceae"                 
## [3358,] "Fusobacteriales"             "Fusobacteriaceae"             
## [3359,] "Spirochaetales"              "Spirochaetaceae"              
## [3360,] "Selenomonadales"             "Veillonellaceae"              
## [3361,] NA                            NA                             
## [3362,] "Bacteroidales"               "Prevotellaceae"               
## [3363,] "Selenomonadales"             "Veillonellaceae"              
## [3364,] "Clostridiales"               "Family_XI"                    
## [3365,] "Bacteroidales"               "Prevotellaceae"               
## [3366,] "Bacteroidales"               "Prevotellaceae"               
## [3367,] "Pasteurellales"              "Pasteurellaceae"              
## [3368,] "Actinomycetales"             "Actinomycetaceae"             
## [3369,] "Flavobacteriales"            "Flavobacteriaceae"            
## [3370,] "Selenomonadales"             "Veillonellaceae"              
## [3371,] "Flavobacteriales"            "Flavobacteriaceae"            
## [3372,] "Fusobacteriales"             "Fusobacteriaceae"             
## [3373,] "Fusobacteriales"             "Leptotrichiaceae"             
## [3374,] "Spirochaetales"              "Spirochaetaceae"              
## [3375,] "Bacteroidales"               "Prevotellaceae"               
## [3376,] "Flavobacteriales"            "Flavobacteriaceae"            
## [3377,] "Bacteroidales"               "Prevotellaceae"               
## [3378,] "Fusobacteriales"             "Leptotrichiaceae"             
## [3379,] "Bacteroidales"               "Rikenellaceae"                
## [3380,] "Flavobacteriales"            "Flavobacteriaceae"            
## [3381,] "Flavobacteriales"            "Flavobacteriaceae"            
## [3382,] "Bacteroidales"               "Prevotellaceae"               
## [3383,] "Clostridiales"               "Lachnospiraceae"              
## [3384,] "Bacteroidales"               "Prevotellaceae"               
## [3385,] "Spirochaetales"              "Spirochaetaceae"              
## [3386,] "Bacteroidales"               "Porphyromonadaceae"           
## [3387,] "Lactobacillales"             "Streptococcaceae"             
## [3388,] "Clostridiales"               "Peptococcaceae"               
## [3389,] "Bacteroidales"               "Prevotellaceae"               
## [3390,] "Bacteroidales"               "Prevotellaceae"               
## [3391,] "Lactobacillales"             "Carnobacteriaceae"            
## [3392,] "Flavobacteriales"            "Flavobacteriaceae"            
## [3393,] "Saccharimonadales"           "Saccharimonadaceae"           
## [3394,] NA                            NA                             
## [3395,] "Fusobacteriales"             "Leptotrichiaceae"             
## [3396,] "Micrococcales"               "Micrococcaceae"               
## [3397,] "Bacteroidales"               "Porphyromonadaceae"           
## [3398,] "Micrococcales"               NA                             
## [3399,] "Clostridiales"               "Lachnospiraceae"              
## [3400,] "Bacteroidales"               "Porphyromonadaceae"           
## [3401,] "Enterobacteriales"           "Enterobacteriaceae"           
## [3402,] "Betaproteobacteriales"       "Neisseriaceae"                
## [3403,] "Clostridiales"               "Family_XI"                    
## [3404,] "Flavobacteriales"            "Flavobacteriaceae"            
## [3405,] "Bacteroidales"               "Prevotellaceae"               
## [3406,] "Bacteroidales"               "Prevotellaceae"               
## [3407,] "Bacteroidales"               "Prevotellaceae"               
## [3408,] "Lactobacillales"             "Aerococcaceae"                
## [3409,] "Flavobacteriales"            "Flavobacteriaceae"            
## [3410,] "Pasteurellales"              "Pasteurellaceae"              
## [3411,] "Coriobacteriales"            "Atopobiaceae"                 
## [3412,] "Bacteroidales"               "Prevotellaceae"               
## [3413,] "Spirochaetales"              "Spirochaetaceae"              
## [3414,] "Bacteroidales"               "Prevotellaceae"               
## [3415,] "Bacteroidales"               "Prevotellaceae"               
## [3416,] "Bacteroidales"               "Prevotellaceae"               
## [3417,] "Fusobacteriales"             "Leptotrichiaceae"             
## [3418,] "Bacteroidales"               "Porphyromonadaceae"           
## [3419,] "Selenomonadales"             "Veillonellaceae"              
## [3420,] NA                            NA                             
## [3421,] "Bacteroidales"               "Prevotellaceae"               
## [3422,] "Clostridiales"               "Lachnospiraceae"              
## [3423,] NA                            NA                             
## [3424,] "Bacteroidales"               "Prevotellaceae"               
## [3425,] "Pasteurellales"              "Pasteurellaceae"              
## [3426,] "Bacteroidales"               "Prevotellaceae"               
## [3427,] "Bacteroidales"               "Prevotellaceae"               
## [3428,] "Flavobacteriales"            "Flavobacteriaceae"            
## [3429,] "Betaproteobacteriales"       NA                             
## [3430,] NA                            NA                             
## [3431,] "Bacteroidales"               "Prevotellaceae"               
## [3432,] "Bacteroidales"               "Prevotellaceae"               
## [3433,] "Flavobacteriales"            "Weeksellaceae"                
## [3434,] "Bacteroidales"               "Prevotellaceae"               
## [3435,] NA                            NA                             
## [3436,] "Enterobacteriales"           "Enterobacteriaceae"           
## [3437,] "Pasteurellales"              "Pasteurellaceae"              
## [3438,] "Selenomonadales"             "Veillonellaceae"              
## [3439,] "Pseudomonadales"             "Pseudomonadaceae"             
## [3440,] "Lactobacillales"             "Aerococcaceae"                
## [3441,] "Flavobacteriales"            "Flavobacteriaceae"            
## [3442,] "Coriobacteriales"            "Atopobiaceae"                 
## [3443,] "Clostridiales"               "Peptostreptococcaceae"        
## [3444,] "Selenomonadales"             "Veillonellaceae"              
## [3445,] "Bacteroidales"               "Prevotellaceae"               
## [3446,] "Clostridiales"               "Lachnospiraceae"              
## [3447,] "Campylobacterales"           NA                             
## [3448,] "Bacteroidales"               "Prevotellaceae"               
## [3449,] "Bacteroidales"               "Prevotellaceae"               
## [3450,] "Fusobacteriales"             "Fusobacteriaceae"             
## [3451,] "Bacteroidales"               "Prevotellaceae"               
## [3452,] "Fusobacteriales"             "Leptotrichiaceae"             
## [3453,] "Bacteroidales"               "Prevotellaceae"               
## [3454,] "Bacteroidales"               "Prevotellaceae"               
## [3455,] "Selenomonadales"             "Veillonellaceae"              
## [3456,] "Bacteroidales"               "Porphyromonadaceae"           
## [3457,] "Fusobacteriales"             NA                             
## [3458,] "Selenomonadales"             "Veillonellaceae"              
## [3459,] "Pasteurellales"              "Pasteurellaceae"              
## [3460,] "Micrococcales"               NA                             
## [3461,] "Selenomonadales"             "Veillonellaceae"              
## [3462,] "Betaproteobacteriales"       "Neisseriaceae"                
## [3463,] "Clostridiales"               "Lachnospiraceae"              
## [3464,] "Bacteroidales"               "Prevotellaceae"               
## [3465,] "Bacteroidales"               "Porphyromonadaceae"           
## [3466,] "Bacteroidales"               NA                             
## [3467,] "Selenomonadales"             "Veillonellaceae"              
## [3468,] NA                            NA                             
## [3469,] "Bacteroidales"               "Prevotellaceae"               
## [3470,] "Bacteroidales"               "Prevotellaceae"               
## [3471,] NA                            NA                             
## [3472,] "Betaproteobacteriales"       NA                             
## [3473,] "Clostridiales"               "Lachnospiraceae"              
## [3474,] "Actinomycetales"             "Actinomycetaceae"             
## [3475,] "Bacteroidales"               "Porphyromonadaceae"           
## [3476,] "Bacteroidales"               "Prevotellaceae"               
## [3477,] "Bacteroidales"               "Porphyromonadaceae"           
## [3478,] "Betaproteobacteriales"       NA                             
## [3479,] "Bacteroidales"               NA                             
## [3480,] "Selenomonadales"             "Veillonellaceae"              
## [3481,] "Pasteurellales"              "Pasteurellaceae"              
## [3482,] "Flavobacteriales"            "Flavobacteriaceae"            
## [3483,] "Lactobacillales"             "Carnobacteriaceae"            
## [3484,] "Clostridiales"               "Lachnospiraceae"              
## [3485,] "Bacteroidales"               "Prevotellaceae"               
## [3486,] "Bacteroidales"               "Prevotellaceae"               
## [3487,] "Flavobacteriales"            "Flavobacteriaceae"            
## [3488,] "Selenomonadales"             "Veillonellaceae"              
## [3489,] "Selenomonadales"             "Veillonellaceae"              
## [3490,] "Saccharimonadales"           NA                             
## [3491,] "Bacteroidales"               "Prevotellaceae"               
## [3492,] "Selenomonadales"             "Veillonellaceae"              
## [3493,] "Selenomonadales"             "Veillonellaceae"              
## [3494,] "Micrococcales"               NA                             
## [3495,] NA                            NA                             
## [3496,] "Pasteurellales"              "Pasteurellaceae"              
## [3497,] "Bacteroidales"               "Prevotellaceae"               
## [3498,] "Chloroplast"                 NA                             
## [3499,] "Lactobacillales"             "Streptococcaceae"             
## [3500,] "Bacteroidales"               "Prevotellaceae"               
## [3501,] NA                            NA                             
## [3502,] "Lactobacillales"             "Streptococcaceae"             
## [3503,] "Pasteurellales"              "Pasteurellaceae"              
## [3504,] "Micrococcales"               NA                             
## [3505,] "Bacteroidales"               "Paludibacteraceae"            
## [3506,] "Bacteroidales"               "Prevotellaceae"               
## [3507,] "Fusobacteriales"             "Leptotrichiaceae"             
## [3508,] "Fusobacteriales"             "Fusobacteriaceae"             
## [3509,] "Fusobacteriales"             "Leptotrichiaceae"             
## [3510,] "Saccharimonadales"           "Saccharimonadaceae"           
## [3511,] "Flavobacteriales"            "Flavobacteriaceae"            
## [3512,] "Pasteurellales"              "Pasteurellaceae"              
## [3513,] "Clostridiales"               "Peptococcaceae"               
## [3514,] "Bacteroidales"               "Prevotellaceae"               
## [3515,] "Clostridiales"               "Lachnospiraceae"              
## [3516,] "Selenomonadales"             "Veillonellaceae"              
## [3517,] "Selenomonadales"             "Veillonellaceae"              
## [3518,] "Rickettsiales"               "Mitochondria"                 
## [3519,] "Selenomonadales"             "Veillonellaceae"              
## [3520,] "Pseudomonadales"             "Pseudomonadaceae"             
## [3521,] "Pasteurellales"              "Pasteurellaceae"              
## [3522,] "Bacteroidales"               "Prevotellaceae"               
## [3523,] "Selenomonadales"             "Veillonellaceae"              
## [3524,] "Pasteurellales"              "Pasteurellaceae"              
## [3525,] "Bacteroidales"               "Prevotellaceae"               
## [3526,] "Lactobacillales"             "Carnobacteriaceae"            
## [3527,] "Betaproteobacteriales"       "Neisseriaceae"                
## [3528,] "Betaproteobacteriales"       "Neisseriaceae"                
## [3529,] "Bacteroidales"               "Porphyromonadaceae"           
## [3530,] "Lactobacillales"             "Streptococcaceae"             
## [3531,] "Bacteroidales"               "Prevotellaceae"               
## [3532,] "Bacteroidales"               "Prevotellaceae"               
## [3533,] NA                            NA                             
## [3534,] "Fusobacteriales"             "Fusobacteriaceae"             
## [3535,] NA                            NA                             
## [3536,] NA                            NA                             
## [3537,] "Saccharimonadales"           "Saccharimonadaceae"           
## [3538,] "Selenomonadales"             "Veillonellaceae"              
## [3539,] "Fusobacteriales"             "Leptotrichiaceae"             
## [3540,] "Bacteroidales"               "Prevotellaceae"               
## [3541,] "Bacteroidales"               "Prevotellaceae"               
## [3542,] "Fusobacteriales"             "Leptotrichiaceae"             
## [3543,] "Fusobacteriales"             "Leptotrichiaceae"             
## [3544,] "Betaproteobacteriales"       "Neisseriaceae"                
## [3545,] "Fusobacteriales"             "Leptotrichiaceae"             
## [3546,] "Clostridiales"               "Family_XI"                    
## [3547,] NA                            NA                             
## [3548,] "Bacteroidales"               "Tannerellaceae"               
## [3549,] "Micrococcales"               "Micrococcaceae"               
## [3550,] NA                            NA                             
## [3551,] "Clostridiales"               "Lachnospiraceae"              
## [3552,] "Clostridiales"               "Lachnospiraceae"              
## [3553,] "Betaproteobacteriales"       "Neisseriaceae"                
## [3554,] "Pasteurellales"              "Pasteurellaceae"              
## [3555,] "Spirochaetales"              "Spirochaetaceae"              
## [3556,] "Lactobacillales"             "Streptococcaceae"             
## [3557,] "Lactobacillales"             "Streptococcaceae"             
## [3558,] "Micrococcales"               NA                             
## [3559,] "Micrococcales"               NA                             
## [3560,] "Lactobacillales"             "Lactobacillaceae"             
## [3561,] "Spirochaetales"              "Spirochaetaceae"              
## [3562,] "Clostridiales"               "Lachnospiraceae"              
## [3563,] "Campylobacterales"           NA                             
## [3564,] "Clostridiales"               "Peptostreptococcaceae"        
## [3565,] "Bacteroidales"               "Prevotellaceae"               
## [3566,] "Selenomonadales"             "Veillonellaceae"              
## [3567,] "Betaproteobacteriales"       "Neisseriaceae"                
## [3568,] "Lactobacillales"             "Streptococcaceae"             
## [3569,] "Selenomonadales"             "Veillonellaceae"              
## [3570,] "Bacteroidales"               "Prevotellaceae"               
## [3571,] "Flavobacteriales"            "Flavobacteriaceae"            
## [3572,] "Campylobacterales"           NA                             
## [3573,] "Fusobacteriales"             "Fusobacteriaceae"             
## [3574,] "Bacteroidales"               "Porphyromonadaceae"           
## [3575,] "Selenomonadales"             "Veillonellaceae"              
## [3576,] "Clostridiales"               "Lachnospiraceae"              
## [3577,] "Lactobacillales"             "Streptococcaceae"             
## [3578,] "Clostridiales"               "Lachnospiraceae"              
## [3579,] "Bacteroidales"               NA                             
## [3580,] "Clostridiales"               "Lachnospiraceae"              
## [3581,] "Bacteroidales"               "Prevotellaceae"               
## [3582,] "Spirochaetales"              "Spirochaetaceae"              
## [3583,] "Clostridiales"               "Peptococcaceae"               
## [3584,] "Micrococcales"               NA                             
## [3585,] "Bacteroidales"               "Prevotellaceae"               
## [3586,] "Betaproteobacteriales"       "Neisseriaceae"                
## [3587,] "Bacteroidales"               "Prevotellaceae"               
## [3588,] "Bacteroidales"               "Prevotellaceae"               
## [3589,] NA                            NA                             
## [3590,] "Campylobacterales"           NA                             
## [3591,] "Lactobacillales"             "Carnobacteriaceae"            
## [3592,] "Bacteroidales"               "Prevotellaceae"               
## [3593,] "Lactobacillales"             "Carnobacteriaceae"            
## [3594,] "Bacteroidales"               "Porphyromonadaceae"           
## [3595,] NA                            NA                             
## [3596,] "Fusobacteriales"             "Leptotrichiaceae"             
## [3597,] "Bacteroidales"               "Prevotellaceae"               
## [3598,] "Flavobacteriales"            "Flavobacteriaceae"            
## [3599,] NA                            NA                             
## [3600,] "Campylobacterales"           NA                             
## [3601,] "Bacteroidales"               "Prevotellaceae"               
## [3602,] "Betaproteobacteriales"       NA                             
## [3603,] "Selenomonadales"             "Veillonellaceae"              
## [3604,] "Betaproteobacteriales"       "Neisseriaceae"                
## [3605,] "Micrococcales"               NA                             
## [3606,] "Flavobacteriales"            "Weeksellaceae"                
## [3607,] "Lactobacillales"             "Streptococcaceae"             
## [3608,] "Betaproteobacteriales"       "Burkholderiaceae"             
## [3609,] "Fusobacteriales"             "Leptotrichiaceae"             
## [3610,] "Corynebacteriales"           "Corynebacteriaceae"           
## [3611,] "Bacteroidales"               "Porphyromonadaceae"           
## [3612,] "Clostridiales"               "Lachnospiraceae"              
## [3613,] "Fusobacteriales"             "Leptotrichiaceae"             
## [3614,] "Selenomonadales"             "Veillonellaceae"              
## [3615,] "Fusobacteriales"             "Fusobacteriaceae"             
## [3616,] "Clostridiales"               "Lachnospiraceae"              
## [3617,] "Fusobacteriales"             "Leptotrichiaceae"             
## [3618,] "Clostridiales"               "Peptostreptococcaceae"        
## [3619,] "Betaproteobacteriales"       "Neisseriaceae"                
## [3620,] "Coriobacteriales"            "Atopobiaceae"                 
## [3621,] "Micrococcales"               "Micrococcaceae"               
## [3622,] "Bacteroidales"               "Tannerellaceae"               
## [3623,] "Betaproteobacteriales"       "Neisseriaceae"                
## [3624,] "Bacteroidales"               "Prevotellaceae"               
## [3625,] "Bacteroidales"               "Rikenellaceae"                
## [3626,] "Bacteroidales"               "Prevotellaceae"               
## [3627,] "Pasteurellales"              "Pasteurellaceae"              
## [3628,] "Clostridiales"               "Lachnospiraceae"              
## [3629,] "Bacteroidales"               "Tannerellaceae"               
## [3630,] "Saccharimonadales"           NA                             
## [3631,] "Bacteroidales"               "Porphyromonadaceae"           
## [3632,] "Bacteroidales"               "Prevotellaceae"               
## [3633,] "Betaproteobacteriales"       "Neisseriaceae"                
## [3634,] "Lactobacillales"             NA                             
## [3635,] "Sphingobacteriales"          "Lentimicrobiaceae"            
## [3636,] "Saccharimonadales"           NA                             
## [3637,] "Flavobacteriales"            "Weeksellaceae"                
## [3638,] "Clostridiales"               "Ruminococcaceae"              
## [3639,] "Selenomonadales"             "Veillonellaceae"              
## [3640,] "Selenomonadales"             "Veillonellaceae"              
## [3641,] "Selenomonadales"             "Veillonellaceae"              
## [3642,] "Rhodospirillales"            NA                             
## [3643,] "Cardiobacteriales"           "Cardiobacteriaceae"           
## [3644,] "Bacteroidales"               NA                             
## [3645,] "Betaproteobacteriales"       NA                             
## [3646,] "Fusobacteriales"             "Fusobacteriaceae"             
## [3647,] NA                            NA                             
## [3648,] NA                            NA                             
## [3649,] "Clostridiales"               "Lachnospiraceae"              
## [3650,] "Spirochaetales"              "Spirochaetaceae"              
## [3651,] "Clostridiales"               "Ruminococcaceae"              
## [3652,] "Flavobacteriales"            "Flavobacteriaceae"            
## [3653,] NA                            NA                             
## [3654,] "Flavobacteriales"            "Flavobacteriaceae"            
## [3655,] "Bacteroidales"               "Prevotellaceae"               
## [3656,] "Pasteurellales"              "Pasteurellaceae"              
## [3657,] NA                            NA                             
## [3658,] NA                            NA                             
## [3659,] "Selenomonadales"             "Veillonellaceae"              
## [3660,] "Campylobacterales"           NA                             
## [3661,] "Clostridiales"               "Peptostreptococcaceae"        
## [3662,] "Micrococcales"               NA                             
## [3663,] "Bacteroidales"               "Prevotellaceae"               
## [3664,] "Bacteroidales"               "Prevotellaceae"               
## [3665,] "Bacteroidales"               "Prevotellaceae"               
## [3666,] "Selenomonadales"             "Veillonellaceae"              
## [3667,] "Pasteurellales"              "Pasteurellaceae"              
## [3668,] "Pasteurellales"              "Pasteurellaceae"              
## [3669,] NA                            NA                             
## [3670,] "Coriobacteriales"            "Atopobiaceae"                 
## [3671,] NA                            NA                             
## [3672,] NA                            NA                             
## [3673,] "Flavobacteriales"            "Flavobacteriaceae"            
## [3674,] "Bacteroidales"               "Prevotellaceae"               
## [3675,] "Clostridiales"               "Lachnospiraceae"              
## [3676,] "Bacteroidales"               "Prevotellaceae"               
## [3677,] "Spirochaetales"              "Spirochaetaceae"              
## [3678,] "Bacteroidales"               "Prevotellaceae"               
## [3679,] "Flavobacteriales"            "Weeksellaceae"                
## [3680,] "Bacteroidales"               "Prevotellaceae"               
## [3681,] "Selenomonadales"             "Veillonellaceae"              
## [3682,] "Bacteroidales"               "Prevotellaceae"               
## [3683,] "Betaproteobacteriales"       "Neisseriaceae"                
## [3684,] "Pasteurellales"              "Pasteurellaceae"              
## [3685,] "Lactobacillales"             "Aerococcaceae"                
## [3686,] "Clostridiales"               "Lachnospiraceae"              
## [3687,] NA                            NA                             
## [3688,] "Bacteroidales"               "Prevotellaceae"               
## [3689,] "Bacteroidales"               "Prevotellaceae"               
## [3690,] "Clostridiales"               "Lachnospiraceae"              
## [3691,] "Fusobacteriales"             "Leptotrichiaceae"             
## [3692,] "Fusobacteriales"             "Fusobacteriaceae"             
## [3693,] "Sphingobacteriales"          "Lentimicrobiaceae"            
## [3694,] "Spirochaetales"              "Spirochaetaceae"              
## [3695,] "Clostridiales"               "Lachnospiraceae"              
## [3696,] "Selenomonadales"             "Veillonellaceae"              
## [3697,] "Clostridiales"               "Ruminococcaceae"              
## [3698,] "Cardiobacteriales"           "Cardiobacteriaceae"           
## [3699,] "Fusobacteriales"             "Leptotrichiaceae"             
## [3700,] "Bacteroidales"               "Prevotellaceae"               
## [3701,] "Selenomonadales"             "Veillonellaceae"              
## [3702,] "Campylobacterales"           "Campylobacteraceae"           
## [3703,] NA                            NA                             
## [3704,] "Fusobacteriales"             "Leptotrichiaceae"             
## [3705,] "Betaproteobacteriales"       "Neisseriaceae"                
## [3706,] NA                            NA                             
## [3707,] "Bacteroidales"               "Prevotellaceae"               
## [3708,] "Betaproteobacteriales"       "Neisseriaceae"                
## [3709,] "Bacteroidales"               "Porphyromonadaceae"           
## [3710,] "Bacteroidales"               "Prevotellaceae"               
## [3711,] "Clostridiales"               "Lachnospiraceae"              
## [3712,] "Coriobacteriales"            "Atopobiaceae"                 
## [3713,] "Lactobacillales"             "Streptococcaceae"             
## [3714,] "Clostridiales"               "Family_XIII"                  
## [3715,] "Clostridiales"               "Lachnospiraceae"              
## [3716,] "Betaproteobacteriales"       "Burkholderiaceae"             
## [3717,] "Lactobacillales"             "Aerococcaceae"                
## [3718,] "Fusobacteriales"             "Fusobacteriaceae"             
## [3719,] "Saccharimonadales"           "Saccharimonadaceae"           
## [3720,] "Lactobacillales"             "Carnobacteriaceae"            
## [3721,] "Selenomonadales"             "Veillonellaceae"              
## [3722,] "Lactobacillales"             NA                             
## [3723,] "Pasteurellales"              "Pasteurellaceae"              
## [3724,] "Clostridiales"               "Lachnospiraceae"              
## [3725,] "Clostridiales"               "Lachnospiraceae"              
## [3726,] "Campylobacterales"           NA                             
## [3727,] "Campylobacterales"           NA                             
## [3728,] "Clostridiales"               "Peptostreptococcaceae"        
## [3729,] "Clostridiales"               "Lachnospiraceae"              
## [3730,] "Selenomonadales"             "Veillonellaceae"              
## [3731,] "Lactobacillales"             "Lactobacillaceae"             
## [3732,] "Spirochaetales"              "Spirochaetaceae"              
## [3733,] "Selenomonadales"             "Veillonellaceae"              
## [3734,] "Sphingobacteriales"          "Lentimicrobiaceae"            
## [3735,] "Fusobacteriales"             "Fusobacteriaceae"             
## [3736,] "Bacillales"                  "Family_XI"                    
## [3737,] "Bacteroidales"               "Prevotellaceae"               
## [3738,] "Clostridiales"               NA                             
## [3739,] "Bacteroidales"               "Porphyromonadaceae"           
## [3740,] "Bacteroidales"               "Prevotellaceae"               
## [3741,] "Bacteroidales"               "Prevotellaceae"               
## [3742,] "Betaproteobacteriales"       "Neisseriaceae"                
## [3743,] "Campylobacterales"           "Campylobacteraceae"           
## [3744,] "Lactobacillales"             "Streptococcaceae"             
## [3745,] "Saccharimonadales"           NA                             
## [3746,] "Flavobacteriales"            "Flavobacteriaceae"            
## [3747,] "Bacteroidales"               "Prevotellaceae"               
## [3748,] "Bacteroidales"               "Prevotellaceae"               
## [3749,] "Bacteroidales"               "Rikenellaceae"                
## [3750,] "Campylobacterales"           NA                             
## [3751,] "Clostridiales"               "Lachnospiraceae"              
## [3752,] "Lactobacillales"             "Streptococcaceae"             
## [3753,] "Selenomonadales"             "Veillonellaceae"              
## [3754,] "Bacteroidales"               "Prevotellaceae"               
## [3755,] "Clostridiales"               "Lachnospiraceae"              
## [3756,] "Clostridiales"               "Lachnospiraceae"              
## [3757,] "Bacteroidales"               "Prevotellaceae"               
## [3758,] "Lactobacillales"             "Streptococcaceae"             
## [3759,] "Selenomonadales"             "Veillonellaceae"              
## [3760,] "Saccharimonadales"           NA                             
## [3761,] "Bacteroidales"               "Prevotellaceae"               
## [3762,] "Lactobacillales"             "Streptococcaceae"             
## [3763,] "Spirochaetales"              "Spirochaetaceae"              
## [3764,] "Flavobacteriales"            "Weeksellaceae"                
## [3765,] "Bacteroidales"               "Prevotellaceae"               
## [3766,] "Bacteroidales"               "Prevotellaceae"               
## [3767,] "Betaproteobacteriales"       "Burkholderiaceae"             
## [3768,] "Selenomonadales"             "Veillonellaceae"              
## [3769,] "Corynebacteriales"           "Corynebacteriaceae"           
## [3770,] "Pasteurellales"              "Pasteurellaceae"              
## [3771,] "Bacteroidales"               "Prevotellaceae"               
## [3772,] "Saccharimonadales"           "Saccharimonadaceae"           
## [3773,] "Clostridiales"               "Peptostreptococcaceae"        
## [3774,] "Clostridiales"               "Lachnospiraceae"              
## [3775,] "Selenomonadales"             "Veillonellaceae"              
## [3776,] "Bacteroidales"               "Prevotellaceae"               
## [3777,] "Bacteroidales"               "Prevotellaceae"               
## [3778,] "Bacteroidales"               "Prevotellaceae"               
## [3779,] "Bacteroidales"               "Prevotellaceae"               
## [3780,] "Clostridiales"               "Family_XIII"                  
## [3781,] "Bacteroidales"               "Rikenellaceae"                
## [3782,] "Bacteroidales"               "Prevotellaceae"               
## [3783,] "Betaproteobacteriales"       "Neisseriaceae"                
## [3784,] "Bacteroidales"               "Prevotellaceae"               
## [3785,] "Fusobacteriales"             NA                             
## [3786,] "Bacteroidales"               "Tannerellaceae"               
## [3787,] NA                            NA                             
## [3788,] "Saccharimonadales"           "Saccharimonadaceae"           
## [3789,] "Betaproteobacteriales"       "Neisseriaceae"                
## [3790,] "Selenomonadales"             "Veillonellaceae"              
## [3791,] "Spirochaetales"              "Spirochaetaceae"              
## [3792,] "Saccharimonadales"           "Saccharimonadaceae"           
## [3793,] "Flavobacteriales"            "Flavobacteriaceae"            
## [3794,] "Bacteroidales"               "Prevotellaceae"               
## [3795,] "Selenomonadales"             "Veillonellaceae"              
## [3796,] "Saccharimonadales"           NA                             
## [3797,] "Absconditabacteriales_(SR1)" NA                             
## [3798,] "Lactobacillales"             "Carnobacteriaceae"            
## [3799,] "Bacteroidales"               "Prevotellaceae"               
## [3800,] "Pasteurellales"              "Pasteurellaceae"              
## [3801,] "Selenomonadales"             "Veillonellaceae"              
## [3802,] "Bacteroidales"               "Prevotellaceae"               
## [3803,] "Betaproteobacteriales"       "Neisseriaceae"                
## [3804,] "Clostridiales"               "Lachnospiraceae"              
## [3805,] "Lactobacillales"             "Streptococcaceae"             
## [3806,] "Saccharimonadales"           "Saccharimonadaceae"           
## [3807,] "Clostridiales"               "Lachnospiraceae"              
## [3808,] "Bacteroidales"               "Prevotellaceae"               
## [3809,] "Micrococcales"               NA                             
## [3810,] "Micrococcales"               NA                             
## [3811,] "Selenomonadales"             "Veillonellaceae"              
## [3812,] "Selenomonadales"             "Veillonellaceae"              
## [3813,] "Lactobacillales"             "Streptococcaceae"             
## [3814,] "Lactobacillales"             "Streptococcaceae"             
## [3815,] "Bacteroidales"               "Prevotellaceae"               
## [3816,] "Flavobacteriales"            "Weeksellaceae"                
## [3817,] "Selenomonadales"             "Veillonellaceae"              
## [3818,] "Selenomonadales"             "Veillonellaceae"              
## [3819,] "Flavobacteriales"            "Flavobacteriaceae"            
## [3820,] "Bacteroidales"               "Prevotellaceae"               
## [3821,] "Absconditabacteriales_(SR1)" NA                             
## [3822,] "Spirochaetales"              "Spirochaetaceae"              
## [3823,] "Bacteroidales"               "Prevotellaceae"               
## [3824,] "Selenomonadales"             "Veillonellaceae"              
## [3825,] "Spirochaetales"              "Spirochaetaceae"              
## [3826,] "Clostridiales"               "Lachnospiraceae"              
## [3827,] "Saccharimonadales"           NA                             
## [3828,] "Pasteurellales"              "Pasteurellaceae"              
## [3829,] "Lactobacillales"             "Lactobacillaceae"             
## [3830,] "Saccharimonadales"           "Saccharimonadaceae"           
## [3831,] "Bacteroidales"               "Prevotellaceae"               
## [3832,] "Betaproteobacteriales"       "Neisseriaceae"                
## [3833,] "Fusobacteriales"             "Leptotrichiaceae"             
## [3834,] "Micrococcales"               "Micrococcaceae"               
## [3835,] "Bacteroidales"               "Porphyromonadaceae"           
## [3836,] "Bacteroidales"               "Prevotellaceae"               
## [3837,] NA                            NA                             
## [3838,] "Bacteroidales"               "Prevotellaceae"               
## [3839,] "Bacteroidales"               "Prevotellaceae"               
## [3840,] "Clostridiales"               "Lachnospiraceae"              
## [3841,] "Bacteroidales"               "Prevotellaceae"               
## [3842,] "Flavobacteriales"            "Flavobacteriaceae"            
## [3843,] "Sphingobacteriales"          NA                             
## [3844,] "Bacteroidales"               "Bacteroidales_Incertae_Sedis" 
## [3845,] NA                            NA                             
## [3846,] "Campylobacterales"           NA                             
## [3847,] "Bacteroidales"               "Prevotellaceae"               
## [3848,] "Clostridiales"               "Lachnospiraceae"              
## [3849,] "Spirochaetales"              "Spirochaetaceae"              
## [3850,] "Clostridiales"               "Lachnospiraceae"              
## [3851,] "Betaproteobacteriales"       "Neisseriaceae"                
## [3852,] "Betaproteobacteriales"       "Neisseriaceae"                
## [3853,] "Bacteroidales"               NA                             
## [3854,] "Lactobacillales"             "Carnobacteriaceae"            
## [3855,] "Campylobacterales"           "Campylobacteraceae"           
## [3856,] "Lactobacillales"             NA                             
## [3857,] "Fusobacteriales"             "Leptotrichiaceae"             
## [3858,] "Bacteroidales"               "Prevotellaceae"               
## [3859,] "Bacteroidales"               "Prevotellaceae"               
## [3860,] "Flavobacteriales"            "Flavobacteriaceae"            
## [3861,] "Lactobacillales"             "Carnobacteriaceae"            
## [3862,] "Clostridiales"               "Lachnospiraceae"              
## [3863,] "Clostridiales"               "Lachnospiraceae"              
## [3864,] "Bacteroidales"               NA                             
## [3865,] "Fusobacteriales"             "Leptotrichiaceae"             
## [3866,] "Clostridiales"               "Lachnospiraceae"              
## [3867,] "Fusobacteriales"             NA                             
## [3868,] "Bacteroidales"               "Prevotellaceae"               
## [3869,] "Saccharimonadales"           "Saccharimonadaceae"           
## [3870,] "Bacteroidales"               "Porphyromonadaceae"           
## [3871,] "Pasteurellales"              "Pasteurellaceae"              
## [3872,] "Betaproteobacteriales"       "Burkholderiaceae"             
## [3873,] "Bacillales"                  NA                             
## [3874,] "Bacteroidales"               "Prevotellaceae"               
## [3875,] NA                            NA                             
## [3876,] "Pasteurellales"              "Pasteurellaceae"              
## [3877,] "Bacteroidales"               "Prevotellaceae"               
## [3878,] "Campylobacterales"           "Campylobacteraceae"           
## [3879,] "Flavobacteriales"            "Weeksellaceae"                
## [3880,] "Clostridiales"               "Family_XIII"                  
## [3881,] "Spirochaetales"              "Spirochaetaceae"              
## [3882,] "Bacteroidales"               "Prevotellaceae"               
## [3883,] "Lactobacillales"             "Carnobacteriaceae"            
## [3884,] "Selenomonadales"             "Veillonellaceae"              
## [3885,] "Pasteurellales"              "Pasteurellaceae"              
## [3886,] "Clostridiales"               "Lachnospiraceae"              
## [3887,] "Clostridiales"               "Family_XI"                    
## [3888,] "Clostridiales"               "Ruminococcaceae"              
## [3889,] "Lactobacillales"             "Streptococcaceae"             
## [3890,] "Clostridiales"               "Family_XIII"                  
## [3891,] "Saccharimonadales"           NA                             
## [3892,] NA                            NA                             
## [3893,] "Clostridiales"               "Lachnospiraceae"              
## [3894,] "Clostridiales"               "Lachnospiraceae"              
## [3895,] "Actinomycetales"             "Actinomycetaceae"             
## [3896,] "Bacteroidales"               "Prevotellaceae"               
## [3897,] "Campylobacterales"           NA                             
## [3898,] "Betaproteobacteriales"       NA                             
## [3899,] "Cardiobacteriales"           "Cardiobacteriaceae"           
## [3900,] "Lactobacillales"             "Carnobacteriaceae"            
## [3901,] "Bacillales"                  "Family_XI"                    
## [3902,] "Selenomonadales"             "Veillonellaceae"              
## [3903,] "Bacteroidales"               "Prevotellaceae"               
## [3904,] "Bacteroidales"               "Porphyromonadaceae"           
## [3905,] "Spirochaetales"              "Spirochaetaceae"              
## [3906,] "Clostridiales"               "Lachnospiraceae"              
## [3907,] "Bacteroidales"               "Prevotellaceae"               
## [3908,] "Pasteurellales"              "Pasteurellaceae"              
## [3909,] "JGI_0000069-P22"             NA                             
## [3910,] "Bacteroidales"               "Prevotellaceae"               
## [3911,] "Bacteroidales"               "Prevotellaceae"               
## [3912,] "Campylobacterales"           NA                             
## [3913,] "Selenomonadales"             "Veillonellaceae"              
## [3914,] "Erysipelotrichales"          "Erysipelotrichaceae"          
## [3915,] "Lactobacillales"             NA                             
## [3916,] "Bacillales"                  "Family_XI"                    
## [3917,] "Bacteroidales"               "Prevotellaceae"               
## [3918,] "Fusobacteriales"             "Fusobacteriaceae"             
## [3919,] "Pasteurellales"              "Pasteurellaceae"              
## [3920,] "Selenomonadales"             "Veillonellaceae"              
## [3921,] "Micrococcales"               NA                             
## [3922,] "Bacteroidales"               "Prevotellaceae"               
## [3923,] NA                            NA                             
## [3924,] "Betaproteobacteriales"       "Burkholderiaceae"             
## [3925,] "Micrococcales"               NA                             
## [3926,] "Selenomonadales"             "Veillonellaceae"              
## [3927,] "Fusobacteriales"             "Leptotrichiaceae"             
## [3928,] "Betaproteobacteriales"       "Burkholderiaceae"             
## [3929,] "Thermales"                   "Thermaceae"                   
## [3930,] "Campylobacterales"           NA                             
## [3931,] "Clostridiales"               "Lachnospiraceae"              
## [3932,] "Clostridiales"               "Family_XI"                    
## [3933,] NA                            NA                             
## [3934,] "Clostridiales"               NA                             
## [3935,] "Betaproteobacteriales"       "Burkholderiaceae"             
## [3936,] "Spirochaetales"              "Spirochaetaceae"              
## [3937,] "Lactobacillales"             "Lactobacillaceae"             
## [3938,] "Pseudomonadales"             "Moraxellaceae"                
## [3939,] "Lactobacillales"             "Lactobacillaceae"             
## [3940,] "Clostridiales"               "Family_XI"                    
## [3941,] "Bacteroidales"               "Prevotellaceae"               
## [3942,] "Bacteroidales"               "Prevotellaceae"               
## [3943,] "Pasteurellales"              "Pasteurellaceae"              
## [3944,] "Clostridiales"               "Lachnospiraceae"              
## [3945,] "Bacteroidales"               "Prevotellaceae"               
## [3946,] "Lactobacillales"             "Carnobacteriaceae"            
## [3947,] "Selenomonadales"             "Veillonellaceae"              
## [3948,] "Lactobacillales"             "Carnobacteriaceae"            
## [3949,] "Bacteroidales"               "Porphyromonadaceae"           
## [3950,] "Lactobacillales"             "Streptococcaceae"             
## [3951,] "Lactobacillales"             "Streptococcaceae"             
## [3952,] "Bacteroidales"               "Prevotellaceae"               
## [3953,] "Fusobacteriales"             "Leptotrichiaceae"             
## [3954,] "Pasteurellales"              "Pasteurellaceae"              
## [3955,] "Campylobacterales"           "Campylobacteraceae"           
## [3956,] NA                            NA                             
## [3957,] "Campylobacterales"           "P6-b51"                       
## [3958,] "Saccharimonadales"           "Saccharimonadaceae"           
## [3959,] "Bacteroidales"               "Prevotellaceae"               
## [3960,] "Selenomonadales"             "Veillonellaceae"              
## [3961,] "Fusobacteriales"             "Fusobacteriaceae"             
## [3962,] "Saccharimonadales"           NA                             
## [3963,] "Bacteroidales"               "Prevotellaceae"               
## [3964,] NA                            NA                             
## [3965,] "Bacteroidales"               "Prevotellaceae"               
## [3966,] "Lactobacillales"             "Carnobacteriaceae"            
## [3967,] NA                            NA                             
## [3968,] "Fusobacteriales"             "Leptotrichiaceae"             
## [3969,] "Fusobacteriales"             "Fusobacteriaceae"             
## [3970,] "Campylobacterales"           "Campylobacteraceae"           
## [3971,] NA                            NA                             
## [3972,] "Acholeplasmatales"           "Acholeplasmataceae"           
## [3973,] "Mycoplasmatales"             "Mycoplasmataceae"             
## [3974,] "Flavobacteriales"            "Weeksellaceae"                
## [3975,] "Desulfovibrionales"          "Desulfovibrionaceae"          
## [3976,] NA                            NA                             
## [3977,] "Bacteroidales"               "Prevotellaceae"               
## [3978,] "Bacteroidales"               "Porphyromonadaceae"           
## [3979,] "Lactobacillales"             "Streptococcaceae"             
## [3980,] "Fusobacteriales"             "Leptotrichiaceae"             
## [3981,] "Clostridiales"               "Lachnospiraceae"              
## [3982,] "Lactobacillales"             "Streptococcaceae"             
## [3983,] "Bacteroidales"               "Prevotellaceae"               
## [3984,] "Selenomonadales"             "Veillonellaceae"              
## [3985,] "Clostridiales"               "Lachnospiraceae"              
## [3986,] "Bacteroidales"               "Prevotellaceae"               
## [3987,] "Bacteroidales"               "Prevotellaceae"               
## [3988,] "Flavobacteriales"            "Flavobacteriaceae"            
## [3989,] "Coriobacteriales"            "Eggerthellaceae"              
## [3990,] "Selenomonadales"             "Veillonellaceae"              
## [3991,] "Fusobacteriales"             "Leptotrichiaceae"             
## [3992,] "Campylobacterales"           NA                             
## [3993,] "Bacteroidales"               "Prevotellaceae"               
## [3994,] "Lactobacillales"             "Streptococcaceae"             
## [3995,] "Spirochaetales"              "Spirochaetaceae"              
## [3996,] "Flavobacteriales"            "Flavobacteriaceae"            
## [3997,] "Clostridiales"               "Peptostreptococcaceae"        
## [3998,] "Sphingobacteriales"          "Lentimicrobiaceae"            
## [3999,] "Saccharimonadales"           NA                             
## [4000,] NA                            NA                             
## [4001,] "Lactobacillales"             "Streptococcaceae"             
## [4002,] "Spirochaetales"              "Spirochaetaceae"              
## [4003,] "Bacillales"                  "Family_XI"                    
## [4004,] "Flavobacteriales"            "Flavobacteriaceae"            
## [4005,] "Fusobacteriales"             "Leptotrichiaceae"             
## [4006,] "Bacteroidales"               "Prevotellaceae"               
## [4007,] "Selenomonadales"             "Veillonellaceae"              
## [4008,] "Bacteroidales"               "Prevotellaceae"               
## [4009,] "Flavobacteriales"            "Weeksellaceae"                
## [4010,] "Pasteurellales"              "Pasteurellaceae"              
## [4011,] "Selenomonadales"             "Veillonellaceae"              
## [4012,] "Bacteroidales"               "Tannerellaceae"               
## [4013,] "Clostridiales"               "Peptostreptococcaceae"        
## [4014,] "Selenomonadales"             "Veillonellaceae"              
## [4015,] "Actinomycetales"             "Actinomycetaceae"             
## [4016,] "Lactobacillales"             "Carnobacteriaceae"            
## [4017,] "Bacteroidales"               "Prevotellaceae"               
## [4018,] "Clostridiales"               "Clostridiales_vadinBB60_group"
## [4019,] "Clostridiales"               "Peptococcaceae"               
## [4020,] "Micrococcales"               "Micrococcaceae"               
## [4021,] "Bacteroidales"               "Prevotellaceae"               
## [4022,] "Actinomycetales"             "Actinomycetaceae"             
## [4023,] "Selenomonadales"             "Veillonellaceae"              
## [4024,] "Saccharimonadales"           NA                             
## [4025,] "Flavobacteriales"            "Flavobacteriaceae"            
## [4026,] "Bacteroidales"               "Prevotellaceae"               
## [4027,] "Lactobacillales"             "Streptococcaceae"             
## [4028,] "Selenomonadales"             "Veillonellaceae"              
## [4029,] "Lactobacillales"             NA                             
## [4030,] "Saccharimonadales"           NA                             
## [4031,] "Betaproteobacteriales"       "Neisseriaceae"                
## [4032,] "Actinomycetales"             "Actinomycetaceae"             
## [4033,] "Spirochaetales"              "Spirochaetaceae"              
## [4034,] "Lactobacillales"             "Carnobacteriaceae"            
## [4035,] "Fusobacteriales"             NA                             
## [4036,] "Clostridiales"               "Eubacteriaceae"               
## [4037,] "Absconditabacteriales_(SR1)" NA                             
## [4038,] "Selenomonadales"             "Veillonellaceae"              
## [4039,] "Betaproteobacteriales"       "Neisseriaceae"                
## [4040,] "Bacillales"                  NA                             
## [4041,] "Bacteroidales"               "Prevotellaceae"               
## [4042,] "Fusobacteriales"             "Fusobacteriaceae"             
## [4043,] "Bacteroidales"               "Prevotellaceae"               
## [4044,] "Flavobacteriales"            "Flavobacteriaceae"            
## [4045,] "Clostridiales"               "Lachnospiraceae"              
## [4046,] "Saccharimonadales"           "Saccharimonadaceae"           
## [4047,] "Bacteroidales"               "Prevotellaceae"               
## [4048,] NA                            NA                             
## [4049,] "Fusobacteriales"             NA                             
## [4050,] "Flavobacteriales"            "Weeksellaceae"                
## [4051,] "Bacteroidales"               "Prevotellaceae"               
## [4052,] "Bacteroidales"               "Prevotellaceae"               
## [4053,] "Lactobacillales"             NA                             
## [4054,] NA                            NA                             
## [4055,] NA                            NA                             
## [4056,] "Fusobacteriales"             "Fusobacteriaceae"             
## [4057,] NA                            NA                             
## [4058,] "Bacteroidales"               NA                             
## [4059,] "Lactobacillales"             "Streptococcaceae"             
## [4060,] "Fusobacteriales"             NA                             
## [4061,] "Chloroplast"                 NA                             
## [4062,] "Actinomycetales"             "Actinomycetaceae"             
## [4063,] "Saccharimonadales"           "Saccharimonadaceae"           
## [4064,] "Fusobacteriales"             "Leptotrichiaceae"             
## [4065,] NA                            NA                             
## [4066,] "Bacteroidales"               "Tannerellaceae"               
## [4067,] "Clostridiales"               "Lachnospiraceae"              
## [4068,] "Clostridiales"               "Lachnospiraceae"              
## [4069,] "Bacteroidales"               NA                             
## [4070,] "Corynebacteriales"           "Corynebacteriaceae"           
## [4071,] "Lactobacillales"             "Streptococcaceae"             
## [4072,] "Absconditabacteriales_(SR1)" NA                             
## [4073,] "Selenomonadales"             "Veillonellaceae"              
## [4074,] "Selenomonadales"             "Veillonellaceae"              
## [4075,] "Selenomonadales"             "Veillonellaceae"              
## [4076,] "Clostridiales"               "Lachnospiraceae"              
## [4077,] "Flavobacteriales"            "Flavobacteriaceae"            
## [4078,] "Pasteurellales"              "Pasteurellaceae"              
## [4079,] "Bacteroidales"               "Prevotellaceae"               
## [4080,] "Flavobacteriales"            "Flavobacteriaceae"            
## [4081,] "Selenomonadales"             "Veillonellaceae"              
## [4082,] "Selenomonadales"             "Veillonellaceae"              
## [4083,] "Clostridiales"               "Lachnospiraceae"              
## [4084,] "Campylobacterales"           NA                             
## [4085,] "Chloroplast"                 NA                             
## [4086,] "Lactobacillales"             NA                             
## [4087,] "Bacteroidales"               NA                             
## [4088,] "Betaproteobacteriales"       "Burkholderiaceae"             
## [4089,] "Corynebacteriales"           "Corynebacteriaceae"           
## [4090,] NA                            NA                             
## [4091,] NA                            NA                             
## [4092,] "Lactobacillales"             NA                             
## [4093,] "Clostridiales"               "Peptostreptococcaceae"        
## [4094,] "Clostridiales"               "Lachnospiraceae"              
## [4095,] "Betaproteobacteriales"       "Neisseriaceae"                
## [4096,] "Clostridiales"               "Ruminococcaceae"              
## [4097,] "Saccharimonadales"           "Saccharimonadaceae"           
## [4098,] "Pasteurellales"              "Pasteurellaceae"              
## [4099,] "Bacteroidales"               "Prevotellaceae"               
## [4100,] "Clostridiales"               "Peptococcaceae"               
## [4101,] "Micrococcales"               NA                             
## [4102,] "Flavobacteriales"            "Weeksellaceae"                
## [4103,] "Clostridiales"               "Family_XIII"                  
## [4104,] "Clostridiales"               "Family_XIII"                  
## [4105,] "Bacteroidales"               "Prevotellaceae"               
## [4106,] "Bacteroidales"               "Prevotellaceae"               
## [4107,] "Saccharimonadales"           NA                             
## [4108,] "Saccharimonadales"           "Saccharimonadaceae"           
## [4109,] "Bacteroidales"               "Porphyromonadaceae"           
## [4110,] "Selenomonadales"             "Veillonellaceae"              
## [4111,] "Lactobacillales"             "Carnobacteriaceae"            
## [4112,] "Selenomonadales"             "Veillonellaceae"              
## [4113,] "Fusobacteriales"             "Leptotrichiaceae"             
## [4114,] "Micrococcales"               NA                             
## [4115,] "Micrococcales"               NA                             
## [4116,] "Clostridiales"               "Peptostreptococcaceae"        
## [4117,] "Coriobacteriales"            "Atopobiaceae"                 
## [4118,] "Bacteroidales"               "Rikenellaceae"                
## [4119,] "Coriobacteriales"            "Atopobiaceae"                 
## [4120,] NA                            NA                             
## [4121,] "Lineage_IV"                  NA                             
## [4122,] "Campylobacterales"           "Rs-M59_termite_group"         
## [4123,] NA                            NA                             
## [4124,] "Campylobacterales"           NA                             
## [4125,] NA                            NA                             
## [4126,] "Saccharimonadales"           NA                             
## [4127,] "Lactobacillales"             "Streptococcaceae"             
## [4128,] "Pasteurellales"              "Pasteurellaceae"              
## [4129,] "Bacteroidales"               "Prevotellaceae"               
## [4130,] "Clostridiales"               "Lachnospiraceae"              
## [4131,] "Lactobacillales"             "Carnobacteriaceae"            
## [4132,] "Clostridiales"               "Peptostreptococcaceae"        
## [4133,] "Flavobacteriales"            "Flavobacteriaceae"            
## [4134,] "Bacteroidales"               "Prevotellaceae"               
## [4135,] "Saccharimonadales"           "Saccharimonadaceae"           
## [4136,] "Saccharimonadales"           "Saccharimonadaceae"           
## [4137,] "Betaproteobacteriales"       NA                             
## [4138,] "Saccharimonadales"           NA                             
## [4139,] "Cardiobacteriales"           "Cardiobacteriaceae"           
## [4140,] "Clostridiales"               "Ruminococcaceae"              
## [4141,] "Betaproteobacteriales"       "Neisseriaceae"                
## [4142,] "Clostridiales"               "Family_XI"                    
## [4143,] "Clostridiales"               "Lachnospiraceae"              
## [4144,] "Bacteroidales"               "Prevotellaceae"               
## [4145,] "Lactobacillales"             "Carnobacteriaceae"            
## [4146,] "Bacteroidales"               "Prevotellaceae"               
## [4147,] "Fusobacteriales"             "Leptotrichiaceae"             
## [4148,] "Selenomonadales"             "Veillonellaceae"              
## [4149,] "Bacteroidales"               "Prevotellaceae"               
## [4150,] "Clostridiales"               "Peptostreptococcaceae"        
## [4151,] "Clostridiales"               "Lachnospiraceae"              
## [4152,] "Spirochaetales"              "Spirochaetaceae"              
## [4153,] "Pasteurellales"              "Pasteurellaceae"              
## [4154,] "Selenomonadales"             "Veillonellaceae"              
## [4155,] "Bacteroidales"               "Prevotellaceae"               
## [4156,] "Corynebacteriales"           "Corynebacteriaceae"           
## [4157,] "Selenomonadales"             "Veillonellaceae"              
## [4158,] NA                            NA                             
## [4159,] "Betaproteobacteriales"       NA                             
## [4160,] "Bacteroidales"               "Prevotellaceae"               
## [4161,] "Clostridiales"               "Lachnospiraceae"              
## [4162,] "Selenomonadales"             "Veillonellaceae"              
## [4163,] "Clostridiales"               "Lachnospiraceae"              
## [4164,] "Saccharimonadales"           NA                             
## [4165,] "Fusobacteriales"             "Fusobacteriaceae"             
## [4166,] "Pasteurellales"              "Pasteurellaceae"              
## [4167,] "Bacillales"                  "Family_XI"                    
## [4168,] "Fusobacteriales"             "Fusobacteriaceae"             
## [4169,] NA                            NA                             
## [4170,] "Selenomonadales"             "Veillonellaceae"              
## [4171,] "Bacteroidales"               "Prevotellaceae"               
## [4172,] "Chloroplast"                 NA                             
## [4173,] "Bacteroidales"               "Prevotellaceae"               
## [4174,] "Clostridiales"               "Lachnospiraceae"              
## [4175,] "Spirochaetales"              "Spirochaetaceae"              
## [4176,] "Betaproteobacteriales"       "Burkholderiaceae"             
## [4177,] "Clostridiales"               "Family_XIII"                  
## [4178,] "Betaproteobacteriales"       "Neisseriaceae"                
## [4179,] "Bacteroidales"               "Prevotellaceae"               
## [4180,] "Saccharimonadales"           NA                             
## [4181,] "Lactobacillales"             "Carnobacteriaceae"            
## [4182,] "Betaproteobacteriales"       "Neisseriaceae"                
## [4183,] "Mycoplasmatales"             "Mycoplasmataceae"             
## [4184,] "Bacteroidales"               "Bacteroidaceae"               
## [4185,] "JGI_0000069-P22"             NA                             
## [4186,] "Clostridiales"               "Lachnospiraceae"              
## [4187,] NA                            NA                             
## [4188,] "Selenomonadales"             "Veillonellaceae"              
## [4189,] NA                            NA                             
## [4190,] "Spirochaetales"              "Spirochaetaceae"              
## [4191,] "Pasteurellales"              "Pasteurellaceae"              
## [4192,] "Betaproteobacteriales"       "Neisseriaceae"                
## [4193,] "Bacillales"                  "Family_XI"                    
## [4194,] "Campylobacterales"           NA                             
## [4195,] "Flavobacteriales"            "Flavobacteriaceae"            
## [4196,] "Clostridiales"               "Family_XIII"                  
## [4197,] "Clostridiales"               "Family_XI"                    
## [4198,] "Selenomonadales"             "Veillonellaceae"              
## [4199,] "Micrococcales"               NA                             
## [4200,] "Pseudomonadales"             "Moraxellaceae"                
## [4201,] "Lactobacillales"             "Streptococcaceae"             
## [4202,] "Fusobacteriales"             "Fusobacteriaceae"             
## [4203,] "Clostridiales"               "Lachnospiraceae"              
## [4204,] "Bacteroidales"               "Prevotellaceae"               
## [4205,] "Selenomonadales"             "Veillonellaceae"              
## [4206,] "Bacteroidales"               NA                             
## [4207,] "Corynebacteriales"           "Corynebacteriaceae"           
## [4208,] "Clostridiales"               "Lachnospiraceae"              
## [4209,] "Fusobacteriales"             "Leptotrichiaceae"             
## [4210,] "Spirochaetales"              "Spirochaetaceae"              
## [4211,] "Saccharimonadales"           "Saccharimonadaceae"           
## [4212,] NA                            NA                             
## [4213,] "Campylobacterales"           NA                             
## [4214,] "Bacteroidales"               "Prevotellaceae"               
## [4215,] "Bacteroidales"               "Prevotellaceae"               
## [4216,] NA                            NA                             
## [4217,] "Saccharimonadales"           "Saccharimonadaceae"           
## [4218,] "Bacteroidales"               "Prevotellaceae"               
## [4219,] "Selenomonadales"             "Veillonellaceae"              
## [4220,] "Campylobacterales"           NA                             
## [4221,] "Betaproteobacteriales"       NA                             
## [4222,] "Fusobacteriales"             NA                             
## [4223,] "Flavobacteriales"            "Weeksellaceae"                
## [4224,] "Flavobacteriales"            "Flavobacteriaceae"            
## [4225,] "Clostridiales"               "Lachnospiraceae"              
## [4226,] "Fusobacteriales"             "Leptotrichiaceae"             
## [4227,] "Bacteroidales"               "Prevotellaceae"               
## [4228,] "Selenomonadales"             "Veillonellaceae"              
## [4229,] "Pasteurellales"              "Pasteurellaceae"              
## [4230,] "Lactobacillales"             "Streptococcaceae"             
## [4231,] "Bacteroidales"               "Prevotellaceae"               
## [4232,] "Pasteurellales"              "Pasteurellaceae"              
## [4233,] "JGI_0000069-P22"             NA                             
## [4234,] "Bacteroidales"               "Prevotellaceae"               
## [4235,] "Lactobacillales"             "Carnobacteriaceae"            
## [4236,] "Saccharimonadales"           "Saccharimonadaceae"           
## [4237,] "Lactobacillales"             "Carnobacteriaceae"            
## [4238,] "Spirochaetales"              "Spirochaetaceae"              
## [4239,] "Corynebacteriales"           "Corynebacteriaceae"           
## [4240,] "Betaproteobacteriales"       "Neisseriaceae"                
## [4241,] "Campylobacterales"           NA                             
## [4242,] "Chloroplast"                 NA                             
## [4243,] "Saccharimonadales"           NA                             
## [4244,] "Lactobacillales"             "Carnobacteriaceae"            
## [4245,] "Bacteroidales"               "Porphyromonadaceae"           
## [4246,] "Actinomycetales"             "Actinomycetaceae"             
## [4247,] "Spirochaetales"              "Spirochaetaceae"              
## [4248,] "Bacillales"                  "Family_XI"                    
## [4249,] "Bacteroidales"               NA                             
## [4250,] "Spirochaetales"              "Spirochaetaceae"              
## [4251,] "Clostridiales"               "Family_XI"                    
## [4252,] NA                            NA                             
## [4253,] "Desulfobacterales"           "Desulfobulbaceae"             
## [4254,] "Izimaplasmatales"            NA                             
## [4255,] "Lactobacillales"             "Streptococcaceae"             
## [4256,] "Bacteroidales"               "Prevotellaceae"               
## [4257,] "Micrococcales"               NA                             
## [4258,] "Campylobacterales"           NA                             
## [4259,] "Flavobacteriales"            "Flavobacteriaceae"            
## [4260,] "Flavobacteriales"            "Weeksellaceae"                
## [4261,] "Saccharimonadales"           "Saccharimonadaceae"           
## [4262,] "Lactobacillales"             NA                             
## [4263,] NA                            NA                             
## [4264,] "Bacteroidales"               "Prevotellaceae"               
## [4265,] "Bacteroidales"               NA                             
## [4266,] "Saccharimonadales"           NA                             
## [4267,] "Micrococcales"               NA                             
## [4268,] "Coriobacteriales"            "Atopobiaceae"                 
## [4269,] "Bacteroidales"               "Prevotellaceae"               
## [4270,] "Lactobacillales"             "Streptococcaceae"             
## [4271,] "Clostridiales"               "Peptostreptococcaceae"        
## [4272,] "Clostridiales"               "Lachnospiraceae"              
## [4273,] NA                            NA                             
## [4274,] "Pasteurellales"              "Pasteurellaceae"              
## [4275,] "Bacteroidales"               "Prevotellaceae"               
## [4276,] "Clostridiales"               "Lachnospiraceae"              
## [4277,] "Clostridiales"               "Family_XIII"                  
## [4278,] "Lactobacillales"             "Streptococcaceae"             
## [4279,] "Lactobacillales"             "Streptococcaceae"             
## [4280,] "Bacteroidales"               "Porphyromonadaceae"           
## [4281,] "Betaproteobacteriales"       NA                             
## [4282,] "Fusobacteriales"             "Leptotrichiaceae"             
## [4283,] "Fusobacteriales"             "Fusobacteriaceae"             
## [4284,] "Betaproteobacteriales"       "Neisseriaceae"                
## [4285,] "Erysipelotrichales"          "Erysipelotrichaceae"          
## [4286,] "Clostridiales"               "Lachnospiraceae"              
## [4287,] "Bacteroidales"               "Bacteroidaceae"               
## [4288,] "Lactobacillales"             "Carnobacteriaceae"            
## [4289,] NA                            NA                             
## [4290,] "Clostridiales"               "Peptostreptococcaceae"        
## [4291,] "Selenomonadales"             "Veillonellaceae"              
## [4292,] "Bacteroidales"               NA                             
## [4293,] "Pasteurellales"              "Pasteurellaceae"              
## [4294,] NA                            NA                             
## [4295,] "Coriobacteriales"            "Atopobiaceae"                 
## [4296,] "Bacteroidales"               "Rikenellaceae"                
## [4297,] "Micrococcales"               "Micrococcaceae"               
## [4298,] NA                            NA                             
## [4299,] "Pasteurellales"              "Pasteurellaceae"              
## [4300,] "Lactobacillales"             "Carnobacteriaceae"            
## [4301,] "Lactobacillales"             NA                             
## [4302,] "Spirochaetales"              "Spirochaetaceae"              
## [4303,] "Flavobacteriales"            "Weeksellaceae"                
## [4304,] "Selenomonadales"             "Veillonellaceae"              
## [4305,] "Rickettsiales"               "Mitochondria"                 
## [4306,] "Clostridiales"               "Lachnospiraceae"              
## [4307,] "Spirochaetales"              "Spirochaetaceae"              
## [4308,] "Lactobacillales"             "Carnobacteriaceae"            
## [4309,] "Pasteurellales"              "Pasteurellaceae"              
## [4310,] "Selenomonadales"             "Veillonellaceae"              
## [4311,] "Betaproteobacteriales"       "Neisseriaceae"                
## [4312,] "Clostridiales"               "Peptostreptococcaceae"        
## [4313,] "Saccharimonadales"           NA                             
## [4314,] "Enterobacteriales"           "Enterobacteriaceae"           
## [4315,] "Bacteroidales"               NA                             
## [4316,] "Clostridiales"               "Lachnospiraceae"              
## [4317,] NA                            NA                             
## [4318,] "Micrococcales"               NA                             
## [4319,] "Spirochaetales"              "Spirochaetaceae"              
## [4320,] "Clostridiales"               "Peptostreptococcaceae"        
## [4321,] "Clostridiales"               "Ruminococcaceae"              
## [4322,] "Pasteurellales"              "Pasteurellaceae"              
## [4323,] "Clostridiales"               "Ruminococcaceae"              
## [4324,] "Campylobacterales"           NA                             
## [4325,] "Saccharimonadales"           "Saccharimonadaceae"           
## [4326,] "Cardiobacteriales"           "Cardiobacteriaceae"           
## [4327,] "Bacteroidales"               "Prevotellaceae"               
## [4328,] "Erysipelotrichales"          "Erysipelotrichaceae"          
## [4329,] "Bifidobacteriales"           "Bifidobacteriaceae"           
## [4330,] "Bacteroidales"               NA                             
## [4331,] "Selenomonadales"             "Veillonellaceae"              
## [4332,] "Betaproteobacteriales"       NA                             
## [4333,] "Betaproteobacteriales"       "Neisseriaceae"                
## [4334,] "Bacteroidales"               NA                             
## [4335,] "Lactobacillales"             "Streptococcaceae"             
## [4336,] "Campylobacterales"           "Helicobacteraceae"            
## [4337,] "Spirochaetales"              "Spirochaetaceae"              
## [4338,] "Clostridiales"               "Lachnospiraceae"              
## [4339,] "Sphingomonadales"            "Sphingomonadaceae"            
## [4340,] "Xanthomonadales"             "Xanthomonadaceae"             
## [4341,] "Betaproteobacteriales"       NA                             
## [4342,] "Bacillales"                  "Family_XI"                    
## [4343,] "Bacteroidales"               "Porphyromonadaceae"           
## [4344,] "Fusobacteriales"             "Leptotrichiaceae"             
## [4345,] "Bacteroidales"               "Rikenellaceae"                
## [4346,] "Bacteroidales"               NA                             
## [4347,] "Clostridiales"               "Family_XI"                    
## [4348,] "Bacteroidales"               "Prevotellaceae"               
## [4349,] "Betaproteobacteriales"       NA                             
## [4350,] "Bacteroidales"               "Tannerellaceae"               
## [4351,] "Rickettsiales"               "Mitochondria"                 
## [4352,] "Clostridiales"               "Defluviitaleaceae"            
## [4353,] "Bacteroidales"               "Prevotellaceae"               
## [4354,] "Bacillales"                  "Family_XI"                    
## [4355,] "Bacteroidales"               "Prevotellaceae"               
## [4356,] "Lactobacillales"             "Streptococcaceae"             
## [4357,] "Betaproteobacteriales"       "Burkholderiaceae"             
## [4358,] "Fusobacteriales"             "Fusobacteriaceae"             
## [4359,] "Pasteurellales"              "Pasteurellaceae"              
## [4360,] "Clostridiales"               "Lachnospiraceae"              
## [4361,] "Bacteroidales"               "Marinilabiliaceae"            
## [4362,] "Fusobacteriales"             "Leptotrichiaceae"             
## [4363,] "Flavobacteriales"            "Weeksellaceae"                
## [4364,] "Clostridiales"               "Peptostreptococcaceae"        
## [4365,] "Betaproteobacteriales"       NA                             
## [4366,] "Betaproteobacteriales"       NA                             
## [4367,] NA                            NA                             
## [4368,] "Campylobacterales"           NA                             
## [4369,] "Clostridiales"               "Family_XI"                    
## [4370,] "Fusobacteriales"             "Fusobacteriaceae"             
## [4371,] "Enterobacteriales"           "Enterobacteriaceae"           
## [4372,] "Cardiobacteriales"           "Cardiobacteriaceae"           
## [4373,] "Selenomonadales"             "Veillonellaceae"              
## [4374,] "Fusobacteriales"             "Leptotrichiaceae"             
## [4375,] "Selenomonadales"             "Veillonellaceae"              
## [4376,] "Spirochaetales"              "Spirochaetaceae"              
## [4377,] "Bacteroidales"               "Prevotellaceae"               
## [4378,] "Lactobacillales"             "Carnobacteriaceae"            
## [4379,] "Clostridiales"               "Lachnospiraceae"              
## [4380,] "Pasteurellales"              "Pasteurellaceae"              
## [4381,] NA                            NA                             
## [4382,] "Bacteroidales"               "Prevotellaceae"               
## [4383,] "Lactobacillales"             "Streptococcaceae"             
## [4384,] "Coriobacteriales"            "Eggerthellaceae"              
## [4385,] "Clostridiales"               "Lachnospiraceae"              
## [4386,] "Bacteroidales"               "Rikenellaceae"                
## [4387,] "Selenomonadales"             "Veillonellaceae"              
## [4388,] "Clostridiales"               "Lachnospiraceae"              
## [4389,] "Micrococcales"               NA                             
## [4390,] "Selenomonadales"             "Veillonellaceae"              
## [4391,] "Campylobacterales"           "P6-b51"                       
## [4392,] "Campylobacterales"           NA                             
## [4393,] "Fusobacteriales"             NA                             
## [4394,] "Selenomonadales"             "Veillonellaceae"              
## [4395,] "Erysipelotrichales"          "Erysipelotrichaceae"          
## [4396,] "Selenomonadales"             "Veillonellaceae"              
## [4397,] "Selenomonadales"             "Veillonellaceae"              
## [4398,] "Selenomonadales"             "Veillonellaceae"              
## [4399,] NA                            NA                             
## [4400,] "Saccharimonadales"           NA                             
## [4401,] "Selenomonadales"             "Veillonellaceae"              
## [4402,] "Fusobacteriales"             "Fusobacteriaceae"             
## [4403,] "Betaproteobacteriales"       "Neisseriaceae"                
## [4404,] "Saccharimonadales"           "Saccharimonadaceae"           
## [4405,] "Coriobacteriales"            "Atopobiaceae"                 
## [4406,] "Bacteroidales"               "Prevotellaceae"               
## [4407,] "Bacteroidales"               NA                             
## [4408,] "Rhizobiales"                 "Beijerinckiaceae"             
## [4409,] "Lactobacillales"             "Carnobacteriaceae"            
## [4410,] NA                            NA                             
## [4411,] "Clostridiales"               "Family_XIII"                  
## [4412,] "Lactobacillales"             "Lactobacillaceae"             
## [4413,] "Coriobacteriales"            "Atopobiaceae"                 
## [4414,] "Spirochaetales"              "Spirochaetaceae"              
## [4415,] "Flavobacteriales"            "Flavobacteriaceae"            
## [4416,] "Campylobacterales"           NA                             
## [4417,] "Synergistales"               "Synergistaceae"               
## [4418,] "Micrococcales"               NA                             
## [4419,] "Micrococcales"               NA                             
## [4420,] "Coriobacteriales"            "Atopobiaceae"                 
## [4421,] "Clostridiales"               "Lachnospiraceae"              
## [4422,] "Mycoplasmatales"             "Mycoplasmataceae"             
## [4423,] "Cardiobacteriales"           "Cardiobacteriaceae"           
## [4424,] "Bacillales"                  "Staphylococcaceae"            
## [4425,] "Campylobacterales"           NA                             
## [4426,] "Micrococcales"               NA                             
## [4427,] "Clostridiales"               "Lachnospiraceae"              
## [4428,] "Selenomonadales"             "Veillonellaceae"              
## [4429,] "Bacteroidales"               "Marinilabiliaceae"            
## [4430,] "Coriobacteriales"            "Atopobiaceae"                 
## [4431,] "Bacteroidales"               "Porphyromonadaceae"           
## [4432,] "Bacteroidales"               "Prevotellaceae"               
## [4433,] "Saccharimonadales"           NA                             
## [4434,] "Clostridiales"               "Peptococcaceae"               
## [4435,] NA                            NA                             
## [4436,] "Micrococcales"               NA                             
## [4437,] NA                            NA                             
## [4438,] "Saccharimonadales"           "Saccharimonadaceae"           
## [4439,] "Pasteurellales"              "Pasteurellaceae"              
## [4440,] "Selenomonadales"             "Veillonellaceae"              
## [4441,] "Selenomonadales"             "Veillonellaceae"              
## [4442,] "Pasteurellales"              "Pasteurellaceae"              
## [4443,] "Erysipelotrichales"          "Erysipelotrichaceae"          
## [4444,] "Cardiobacteriales"           "Cardiobacteriaceae"           
## [4445,] "Flavobacteriales"            "Flavobacteriaceae"            
## [4446,] "Spirochaetales"              "Spirochaetaceae"              
## [4447,] "Flavobacteriales"            "Flavobacteriaceae"            
## [4448,] "Bacillales"                  "Family_XI"                    
## [4449,] "Bacteroidales"               "Porphyromonadaceae"           
## [4450,] "Bacteroidales"               "Prevotellaceae"               
## [4451,] NA                            NA                             
## [4452,] "Bacteroidales"               "Prevotellaceae"               
## [4453,] "Actinomycetales"             "Actinomycetaceae"             
## [4454,] "Pasteurellales"              "Pasteurellaceae"              
## [4455,] "Flavobacteriales"            "Flavobacteriaceae"            
## [4456,] "Fusobacteriales"             "Fusobacteriaceae"             
## [4457,] "Betaproteobacteriales"       "Neisseriaceae"                
## [4458,] "Erysipelotrichales"          "Erysipelotrichaceae"          
## [4459,] "Bacteroidales"               "Prevotellaceae"               
## [4460,] "Pasteurellales"              "Pasteurellaceae"              
## [4461,] "Campylobacterales"           NA                             
## [4462,] "Fusobacteriales"             "Leptotrichiaceae"             
## [4463,] "Lactobacillales"             NA                             
## [4464,] "Bacteroidales"               NA                             
## [4465,] "Betaproteobacteriales"       "Neisseriaceae"                
## [4466,] "Coriobacteriales"            "Atopobiaceae"                 
## [4467,] "Betaproteobacteriales"       "Burkholderiaceae"             
## [4468,] "Bacillales"                  NA                             
## [4469,] NA                            NA                             
## [4470,] "Bacteroidales"               "Prevotellaceae"               
## [4471,] "Clostridiales"               "Lachnospiraceae"              
## [4472,] "Clostridiales"               "Lachnospiraceae"              
## [4473,] "Bacteroidales"               "Prevotellaceae"               
## [4474,] "Bacteroidales"               "Prevotellaceae"               
## [4475,] "Clostridiales"               "Lachnospiraceae"              
## [4476,] "Bacteroidales"               "Porphyromonadaceae"           
## [4477,] "Selenomonadales"             "Veillonellaceae"              
## [4478,] "Bacteroidales"               "Prevotellaceae"               
## [4479,] "Bacteroidales"               "Tannerellaceae"               
## [4480,] "Lactobacillales"             "Streptococcaceae"             
## [4481,] "Betaproteobacteriales"       NA                             
## [4482,] "Clostridiales"               "Peptococcaceae"               
## [4483,] "Lactobacillales"             "Streptococcaceae"             
## [4484,] "Bacteroidales"               "Prevotellaceae"               
## [4485,] "Selenomonadales"             "Veillonellaceae"              
## [4486,] "Erysipelotrichales"          "Erysipelotrichaceae"          
## [4487,] "Saccharimonadales"           NA                             
## [4488,] "Saccharimonadales"           NA                             
## [4489,] "Spirochaetales"              "Spirochaetaceae"              
## [4490,] "Clostridiales"               "Lachnospiraceae"              
## [4491,] "Clostridiales"               "Peptostreptococcaceae"        
## [4492,] "Betaproteobacteriales"       "Burkholderiaceae"             
## [4493,] "Clostridiales"               "Lachnospiraceae"              
## [4494,] "Pasteurellales"              "Pasteurellaceae"              
## [4495,] "Bacteroidales"               "Prevotellaceae"               
## [4496,] "Pseudomonadales"             "Pseudomonadaceae"             
## [4497,] "Rhizobiales"                 "Xanthobacteraceae"            
## [4498,] "Bacillales"                  "Family_XI"                    
## [4499,] "Bacteroidales"               "Prevotellaceae"               
## [4500,] "Selenomonadales"             "Veillonellaceae"              
## [4501,] "Blastocatellales"            "Blastocatellaceae"            
## [4502,] "Lactobacillales"             "Streptococcaceae"             
## [4503,] "Betaproteobacteriales"       "Neisseriaceae"                
## [4504,] "Clostridiales"               "Lachnospiraceae"              
## [4505,] "Fusobacteriales"             "Leptotrichiaceae"             
## [4506,] "Bacteroidales"               "Prevotellaceae"               
## [4507,] "Bacteroidales"               "Prevotellaceae"               
## [4508,] "Clostridiales"               "Lachnospiraceae"              
## [4509,] "Saccharimonadales"           "Saccharimonadaceae"           
## [4510,] "Pasteurellales"              "Pasteurellaceae"              
## [4511,] "Bacteroidales"               "Prevotellaceae"               
## [4512,] NA                            NA                             
## [4513,] "Micrococcales"               NA                             
## [4514,] "Lactobacillales"             "Carnobacteriaceae"            
## [4515,] "Fusobacteriales"             "Fusobacteriaceae"             
## [4516,] "Saccharimonadales"           "Saccharimonadaceae"           
## [4517,] "Fusobacteriales"             "Leptotrichiaceae"             
## [4518,] "Selenomonadales"             "Veillonellaceae"              
## [4519,] "Bacteroidales"               "Prevotellaceae"               
## [4520,] "Lactobacillales"             "Streptococcaceae"             
## [4521,] "Betaproteobacteriales"       "Neisseriaceae"                
## [4522,] "Bacteroidales"               "Prevotellaceae"               
## [4523,] "Bacteroidales"               "Prevotellaceae"               
## [4524,] "Betaproteobacteriales"       "Burkholderiaceae"             
## [4525,] "Clostridiales"               "Family_XI"                    
## [4526,] "Flavobacteriales"            "Flavobacteriaceae"            
## [4527,] "Saccharimonadales"           "Saccharimonadaceae"           
## [4528,] "Bacteroidales"               "Prevotellaceae"               
## [4529,] "Selenomonadales"             "Veillonellaceae"              
## [4530,] NA                            NA                             
## [4531,] "Bacteroidales"               "Porphyromonadaceae"           
## [4532,] "Bacteroidales"               "Porphyromonadaceae"           
## [4533,] NA                            NA                             
## [4534,] "Pasteurellales"              "Pasteurellaceae"              
## [4535,] "Bacteroidales"               "Porphyromonadaceae"           
## [4536,] "Micrococcales"               NA                             
## [4537,] "Coriobacteriales"            NA                             
## [4538,] "Lactobacillales"             "Streptococcaceae"             
## [4539,] "Bacteroidales"               "Bacteroidaceae"               
## [4540,] "Bacteroidales"               NA                             
## [4541,] "Clostridiales"               "Lachnospiraceae"              
## [4542,] "Clostridiales"               "Lachnospiraceae"              
## [4543,] NA                            NA                             
## [4544,] "Bacteroidales"               "Prevotellaceae"               
## [4545,] "Campylobacterales"           NA                             
## [4546,] "Spirochaetales"              "Spirochaetaceae"              
## [4547,] NA                            NA                             
## [4548,] "Bacteroidales"               "Rikenellaceae"                
## [4549,] "Selenomonadales"             "Veillonellaceae"              
## [4550,] "Lactobacillales"             "Streptococcaceae"             
## [4551,] "Fusobacteriales"             "Leptotrichiaceae"             
## [4552,] "Lactobacillales"             "Carnobacteriaceae"            
## [4553,] "Spirochaetales"              "Spirochaetaceae"              
## [4554,] "Bacteroidales"               "Prevotellaceae"               
## [4555,] "Clostridiales"               "Clostridiales_vadinBB60_group"
##         [,6]                              [,7]                   
##    [1,] "Veillonella"                     "dispar"               
##    [2,] "Streptococcus"                   NA                     
##    [3,] "Prevotella_7"                    "melaninogenica"       
##    [4,] "Haemophilus"                     NA                     
##    [5,] "Veillonella"                     "rogosae"              
##    [6,] "Veillonella"                     "atypica"              
##    [7,] "Veillonella"                     "atypica"              
##    [8,] NA                                NA                     
##    [9,] "Neisseria"                       NA                     
##   [10,] "Veillonella"                     "tobetsuensis"         
##   [11,] "Neisseria"                       NA                     
##   [12,] "Veillonella"                     NA                     
##   [13,] "Fusobacterium"                   "periodonticum"        
##   [14,] "Veillonella"                     NA                     
##   [15,] "Prevotella_7"                    NA                     
##   [16,] "Campylobacter"                   "concisus"             
##   [17,] "Haemophilus"                     "parainfluenzae"       
##   [18,] "Oribacterium"                    "sinus"                
##   [19,] "Veillonella"                     NA                     
##   [20,] "Alloprevotella"                  NA                     
##   [21,] "Granulicatella"                  NA                     
##   [22,] "Prevotella_7"                    "histicola"            
##   [23,] "Prevotella_7"                    "melaninogenica"       
##   [24,] "Veillonella"                     NA                     
##   [25,] "Haemophilus"                     NA                     
##   [26,] "Prevotella_7"                    "histicola"            
##   [27,] "Porphyromonas"                   NA                     
##   [28,] "Veillonella"                     NA                     
##   [29,] "Streptococcus"                   NA                     
##   [30,] "Streptococcus"                   NA                     
##   [31,] "Porphyromonas"                   "pasteri"              
##   [32,] "Streptococcus"                   NA                     
##   [33,] "Prevotella_6"                    "salivae"              
##   [34,] "Prevotella"                      "pallens"              
##   [35,] "Peptostreptococcus"              "stomatis"             
##   [36,] "Megasphaera"                     "micronuciformis"      
##   [37,] "Porphyromonas"                   NA                     
##   [38,] "Megasphaera"                     NA                     
##   [39,] "Lachnoanaerobaculum"             NA                     
##   [40,] "Streptococcus"                   NA                     
##   [41,] "Prevotella_7"                    NA                     
##   [42,] "Streptococcus"                   NA                     
##   [43,] "Streptococcus"                   "cristatus"            
##   [44,] "Prevotella"                      NA                     
##   [45,] "Neisseria"                       NA                     
##   [46,] "Prevotella_7"                    NA                     
##   [47,] "Megasphaera"                     "micronuciformis"      
##   [48,] "Veillonella"                     NA                     
##   [49,] "Lachnoanaerobaculum"             "cf."                  
##   [50,] "Leptotrichia"                    NA                     
##   [51,] "Streptococcus"                   NA                     
##   [52,] "Haemophilus"                     "parainfluenzae"       
##   [53,] "Streptococcus"                   NA                     
##   [54,] "Leptotrichia"                    NA                     
##   [55,] "Alloprevotella"                  NA                     
##   [56,] "Prevotella_7"                    NA                     
##   [57,] "Veillonella"                     "atypica"              
##   [58,] "Oribacterium"                    "parvum"               
##   [59,] "Atopobium"                       "parvulum"             
##   [60,] "Haemophilus"                     "parainfluenzae"       
##   [61,] "Haemophilus"                     NA                     
##   [62,] "Prevotella_6"                    NA                     
##   [63,] "Oribacterium"                    "asaccharolyticum"     
##   [64,] "Veillonella"                     NA                     
##   [65,] "Haemophilus"                     NA                     
##   [66,] "Selenomonas_3"                   NA                     
##   [67,] "Candidatus_Saccharimonas"        NA                     
##   [68,] "Rothia"                          "mucilaginosa"         
##   [69,] "Neisseria"                       NA                     
##   [70,] "Parvimonas"                      NA                     
##   [71,] "Alloprevotella"                  NA                     
##   [72,] "Rothia"                          "mucilaginosa"         
##   [73,] "Catonella"                       NA                     
##   [74,] "Veillonella"                     NA                     
##   [75,] "Rothia"                          NA                     
##   [76,] "Neisseria"                       "lactamica"            
##   [77,] NA                                NA                     
##   [78,] "Porphyromonas"                   NA                     
##   [79,] "Veillonella"                     "parvula"              
##   [80,] "Stomatobaculum"                  NA                     
##   [81,] "Haemophilus"                     NA                     
##   [82,] "Veillonella"                     "parvula"              
##   [83,] "Prevotella_2"                    "shahii"               
##   [84,] "Prevotella"                      NA                     
##   [85,] "Megasphaera"                     NA                     
##   [86,] "Prevotella_7"                    NA                     
##   [87,] "Leptotrichia"                    NA                     
##   [88,] "Ruminococcaceae_UCG-014"         NA                     
##   [89,] "Alloprevotella"                  NA                     
##   [90,] "Ruminococcaceae_UCG-014"         NA                     
##   [91,] "Rothia"                          "dentocariosa"         
##   [92,] "Neisseria"                       NA                     
##   [93,] "Rothia"                          NA                     
##   [94,] "Gemella"                         "sanguinis"            
##   [95,] "Prevotella_7"                    NA                     
##   [96,] "Neisseria"                       NA                     
##   [97,] "Fusobacterium"                   NA                     
##   [98,] "Veillonella"                     NA                     
##   [99,] "Fusobacterium"                   NA                     
##  [100,] "Prevotella_7"                    NA                     
##  [101,] "Prevotella_7"                    NA                     
##  [102,] "Haemophilus"                     "parainfluenzae"       
##  [103,] "Haemophilus"                     NA                     
##  [104,] "Prevotella"                      "nanceiensis"          
##  [105,] "Granulicatella"                  "elegans"              
##  [106,] "Gemella"                         NA                     
##  [107,] "Veillonella"                     NA                     
##  [108,] "Veillonella"                     NA                     
##  [109,] "Rothia"                          "mucilaginosa"         
##  [110,] "Aggregatibacter"                 "segnis"               
##  [111,] "Campylobacter"                   "concisus"             
##  [112,] "Prevotella"                      "nanceiensis"          
##  [113,] "Neisseria"                       NA                     
##  [114,] "Neisseria"                       NA                     
##  [115,] NA                                NA                     
##  [116,] "Prevotella"                      "nanceiensis"          
##  [117,] "Neisseria"                       NA                     
##  [118,] "Actinobacillus"                  NA                     
##  [119,] "Haemophilus"                     NA                     
##  [120,] "Haemophilus"                     NA                     
##  [121,] "Haemophilus"                     "sputorum"             
##  [122,] "Prevotella"                      NA                     
##  [123,] "Streptococcus"                   NA                     
##  [124,] "Alloprevotella"                  NA                     
##  [125,] "Neisseria"                       NA                     
##  [126,] "Prevotella"                      "nigrescens"           
##  [127,] "Veillonella"                     NA                     
##  [128,] "Haemophilus"                     "sputorum"             
##  [129,] "Aggregatibacter"                 NA                     
##  [130,] "Haemophilus"                     NA                     
##  [131,] "Lautropia"                       "mirabilis"            
##  [132,] "Prevotella"                      "aurantiaca"           
##  [133,] "Candidatus_Saccharimonas"        NA                     
##  [134,] "Prevotella_7"                    NA                     
##  [135,] "Veillonella"                     NA                     
##  [136,] "Veillonella"                     NA                     
##  [137,] "Prevotella_7"                    NA                     
##  [138,] "Capnocytophaga"                  "sputigena"            
##  [139,] "Prevotella_7"                    NA                     
##  [140,] "Streptococcus"                   NA                     
##  [141,] "Veillonella"                     NA                     
##  [142,] "Megasphaera"                     NA                     
##  [143,] "Leptotrichia"                    NA                     
##  [144,] "Veillonella"                     NA                     
##  [145,] "Streptococcus"                   "pseudopneumoniae"     
##  [146,] "Bergeyella"                      NA                     
##  [147,] "Neisseria"                       "oralis"               
##  [148,] "Selenomonas_3"                   NA                     
##  [149,] "Veillonella"                     NA                     
##  [150,] "Veillonella"                     NA                     
##  [151,] "Neisseria"                       NA                     
##  [152,] "Prevotella_7"                    NA                     
##  [153,] "Veillonella"                     NA                     
##  [154,] "Alloprevotella"                  NA                     
##  [155,] "Butyrivibrio_2"                  NA                     
##  [156,] "Klebsiella"                      NA                     
##  [157,] NA                                NA                     
##  [158,] "Prevotella_7"                    NA                     
##  [159,] "Veillonella"                     NA                     
##  [160,] "Oribacterium"                    NA                     
##  [161,] "Streptococcus"                   NA                     
##  [162,] "Rothia"                          NA                     
##  [163,] "Alloprevotella"                  NA                     
##  [164,] "Veillonella"                     NA                     
##  [165,] "Campylobacter"                   NA                     
##  [166,] "Prevotella_7"                    "scopos"               
##  [167,] "Porphyromonas"                   NA                     
##  [168,] "Streptococcus"                   "parasanguinis"        
##  [169,] "Alloprevotella"                  NA                     
##  [170,] "Neisseria"                       NA                     
##  [171,] "Streptococcus"                   NA                     
##  [172,] "Streptococcus"                   NA                     
##  [173,] "Candidatus_Saccharimonas"        NA                     
##  [174,] "Prevotella"                      NA                     
##  [175,] "Prevotella_7"                    NA                     
##  [176,] "Stomatobaculum"                  "longum"               
##  [177,] "Veillonella"                     NA                     
##  [178,] "Veillonella"                     "parvula"              
##  [179,] "Prevotella_7"                    NA                     
##  [180,] "Neisseria"                       "flavescens"           
##  [181,] NA                                NA                     
##  [182,] "Veillonella"                     NA                     
##  [183,] "Neisseria"                       "mucosa"               
##  [184,] "Veillonella"                     NA                     
##  [185,] "Aggregatibacter"                 "aphrophilus"          
##  [186,] "Veillonella"                     "parvula"              
##  [187,] "Veillonella"                     NA                     
##  [188,] "Prevotella"                      NA                     
##  [189,] "Prevotella_7"                    NA                     
##  [190,] "Aggregatibacter"                 NA                     
##  [191,] "Alloprevotella"                  NA                     
##  [192,] "Porphyromonas"                   NA                     
##  [193,] "Neisseria"                       "subflava"             
##  [194,] "Prevotella_7"                    NA                     
##  [195,] "Veillonella"                     NA                     
##  [196,] "Veillonella"                     NA                     
##  [197,] "Fusobacterium"                   "nucleatum"            
##  [198,] "Dialister"                       "invisus"              
##  [199,] "Prevotella"                      NA                     
##  [200,] "Selenomonas_3"                   NA                     
##  [201,] "Alloprevotella"                  NA                     
##  [202,] "Oribacterium"                    NA                     
##  [203,] "Campylobacter"                   NA                     
##  [204,] "Alloprevotella"                  NA                     
##  [205,] "Veillonella"                     "atypica"              
##  [206,] "Veillonella"                     NA                     
##  [207,] "Capnocytophaga"                  NA                     
##  [208,] "Neisseria"                       "flavescens"           
##  [209,] "Kingella"                        "oralis"               
##  [210,] "Streptococcus"                   NA                     
##  [211,] NA                                NA                     
##  [212,] "Veillonella"                     NA                     
##  [213,] "Haemophilus"                     NA                     
##  [214,] "Neisseria"                       NA                     
##  [215,] "Megasphaera"                     NA                     
##  [216,] "Prevotella_7"                    "veroralis"            
##  [217,] "Streptococcus"                   NA                     
##  [218,] "Neisseria"                       NA                     
##  [219,] "Megasphaera"                     NA                     
##  [220,] "Streptococcus"                   NA                     
##  [221,] "Oribacterium"                    NA                     
##  [222,] "Streptococcus"                   NA                     
##  [223,] "Johnsonella"                     NA                     
##  [224,] "Alloprevotella"                  "rava"                 
##  [225,] "Leptotrichia"                    NA                     
##  [226,] "Veillonella"                     NA                     
##  [227,] "Alloprevotella"                  NA                     
##  [228,] "Streptococcus"                   NA                     
##  [229,] "Capnocytophaga"                  NA                     
##  [230,] "Capnocytophaga"                  NA                     
##  [231,] "Leptotrichia"                    NA                     
##  [232,] "Fusobacterium"                   "necrophorum"          
##  [233,] "Haemophilus"                     "pittmaniae"           
##  [234,] "Fusobacterium"                   "nucleatum"            
##  [235,] "Veillonella"                     NA                     
##  [236,] NA                                NA                     
##  [237,] "Alloprevotella"                  NA                     
##  [238,] "Veillonella"                     NA                     
##  [239,] "Prevotella_7"                    NA                     
##  [240,] "Veillonella"                     NA                     
##  [241,] "Fusobacterium"                   NA                     
##  [242,] "Porphyromonas"                   "endodontalis"         
##  [243,] NA                                NA                     
##  [244,] NA                                NA                     
##  [245,] "Prevotella_7"                    NA                     
##  [246,] "Aggregatibacter"                 NA                     
##  [247,] "Neisseria"                       NA                     
##  [248,] NA                                NA                     
##  [249,] NA                                NA                     
##  [250,] "Prevotella_7"                    NA                     
##  [251,] "Alloprevotella"                  NA                     
##  [252,] "Prevotella_7"                    NA                     
##  [253,] "Johnsonella"                     NA                     
##  [254,] "Haemophilus"                     NA                     
##  [255,] "Enterobacter"                    NA                     
##  [256,] "Veillonella"                     NA                     
##  [257,] "Veillonella"                     NA                     
##  [258,] "Actinobacillus"                  NA                     
##  [259,] NA                                NA                     
##  [260,] "Actinomyces"                     "odontolyticus"        
##  [261,] "Lachnoanaerobaculum"             NA                     
##  [262,] "Prevotella_7"                    NA                     
##  [263,] "Alloprevotella"                  NA                     
##  [264,] "Neisseria"                       NA                     
##  [265,] NA                                NA                     
##  [266,] "Bergeriella"                     NA                     
##  [267,] "Solobacterium"                   NA                     
##  [268,] "Prevotella"                      NA                     
##  [269,] "Parvimonas"                      "micra"                
##  [270,] NA                                NA                     
##  [271,] "Veillonella"                     NA                     
##  [272,] "Neisseria"                       NA                     
##  [273,] "Neisseria"                       NA                     
##  [274,] "Veillonella"                     NA                     
##  [275,] "Streptococcus"                   "lactarius"            
##  [276,] "Megasphaera"                     NA                     
##  [277,] "Prevotella"                      NA                     
##  [278,] "Capnocytophaga"                  "leadbetteri"          
##  [279,] "Haemophilus"                     "haemolyticus"         
##  [280,] "Alloprevotella"                  NA                     
##  [281,] "Fusobacterium"                   "nucleatum"            
##  [282,] NA                                NA                     
##  [283,] "Fusobacterium"                   NA                     
##  [284,] "Porphyromonas"                   NA                     
##  [285,] "Prevotella"                      NA                     
##  [286,] NA                                NA                     
##  [287,] "Alloprevotella"                  "tannerae"             
##  [288,] "Veillonella"                     NA                     
##  [289,] "Alloprevotella"                  NA                     
##  [290,] "Oribacterium"                    NA                     
##  [291,] "Alloprevotella"                  NA                     
##  [292,] "Haemophilus"                     NA                     
##  [293,] "Veillonella"                     NA                     
##  [294,] "Candidatus_Saccharimonas"        NA                     
##  [295,] "Anthococcus"                     NA                     
##  [296,] "Veillonella"                     NA                     
##  [297,] NA                                NA                     
##  [298,] "Selenomonas_3"                   NA                     
##  [299,] "Neisseria"                       NA                     
##  [300,] "Alloprevotella"                  "tannerae"             
##  [301,] "Prevotella_7"                    NA                     
##  [302,] "Leptotrichia"                    "buccalis"             
##  [303,] "Prevotella"                      "nigrescens"           
##  [304,] NA                                NA                     
##  [305,] "Fusobacterium"                   NA                     
##  [306,] "Prevotella_7"                    NA                     
##  [307,] "Veillonella"                     NA                     
##  [308,] NA                                NA                     
##  [309,] NA                                NA                     
##  [310,] "Haemophilus"                     NA                     
##  [311,] "Veillonella"                     NA                     
##  [312,] "Fusobacterium"                   "necrophorum"          
##  [313,] NA                                NA                     
##  [314,] "Campylobacter"                   NA                     
##  [315,] "Veillonella"                     NA                     
##  [316,] "Prevotella_7"                    NA                     
##  [317,] "Veillonella"                     NA                     
##  [318,] "Catonella"                       NA                     
##  [319,] NA                                NA                     
##  [320,] "Prevotella"                      NA                     
##  [321,] "Atopobium"                       NA                     
##  [322,] "Streptococcus"                   NA                     
##  [323,] NA                                NA                     
##  [324,] "Veillonella"                     NA                     
##  [325,] "Haemophilus"                     NA                     
##  [326,] NA                                NA                     
##  [327,] "Porphyromonas"                   NA                     
##  [328,] "Atopobium"                       "parvulum"             
##  [329,] "Veillonella"                     NA                     
##  [330,] "Prevotella"                      NA                     
##  [331,] "Alloprevotella"                  "rava"                 
##  [332,] NA                                NA                     
##  [333,] "Lachnoanaerobaculum"             NA                     
##  [334,] "Veillonella"                     NA                     
##  [335,] "Catonella"                       NA                     
##  [336,] "Leptotrichia"                    NA                     
##  [337,] "Abiotrophia"                     "defectiva"            
##  [338,] "Leptotrichia"                    NA                     
##  [339,] "Rothia"                          NA                     
##  [340,] "Fusobacterium"                   NA                     
##  [341,] "Fusobacterium"                   NA                     
##  [342,] "Prevotella_6"                    NA                     
##  [343,] "Neisseria"                       NA                     
##  [344,] "Prevotella"                      NA                     
##  [345,] "Peptostreptococcus"              NA                     
##  [346,] "Kingella"                        NA                     
##  [347,] NA                                NA                     
##  [348,] "Neisseria"                       NA                     
##  [349,] "Haemophilus"                     "parainfluenzae"       
##  [350,] "Prevotella"                      NA                     
##  [351,] NA                                NA                     
##  [352,] "Veillonella"                     NA                     
##  [353,] "Alloprevotella"                  NA                     
##  [354,] "Actinobacillus"                  NA                     
##  [355,] "Fusobacterium"                   "nucleatum"            
##  [356,] "Veillonella"                     NA                     
##  [357,] "Fusobacterium"                   "periodonticum"        
##  [358,] "Prevotella_7"                    NA                     
##  [359,] "Veillonella"                     NA                     
##  [360,] "Peptococcus"                     NA                     
##  [361,] NA                                NA                     
##  [362,] NA                                NA                     
##  [363,] "Alloprevotella"                  NA                     
##  [364,] "Peptoanaerobacter"               NA                     
##  [365,] NA                                NA                     
##  [366,] NA                                NA                     
##  [367,] "Lachnoanaerobaculum"             NA                     
##  [368,] "Lachnoanaerobaculum"             NA                     
##  [369,] "Actinomyces"                     NA                     
##  [370,] "Prevotella_2"                    NA                     
##  [371,] NA                                NA                     
##  [372,] "Gemella"                         NA                     
##  [373,] "Streptococcus"                   NA                     
##  [374,] "Veillonella"                     NA                     
##  [375,] NA                                NA                     
##  [376,] "Veillonella"                     NA                     
##  [377,] "Fusobacterium"                   "nucleatum"            
##  [378,] NA                                NA                     
##  [379,] "Selenomonas"                     "sputigena"            
##  [380,] "Leptotrichia"                    NA                     
##  [381,] "Fusobacterium"                   "nucleatum"            
##  [382,] "Parvimonas"                      NA                     
##  [383,] "Alloprevotella"                  NA                     
##  [384,] "Prevotella"                      NA                     
##  [385,] "Streptococcus"                   NA                     
##  [386,] "Enterobacter"                    NA                     
##  [387,] "Leptotrichia"                    NA                     
##  [388,] "Actinomyces"                     NA                     
##  [389,] "Leptotrichia"                    NA                     
##  [390,] "Prevotella_6"                    NA                     
##  [391,] "Actinomyces"                     "lingnae"              
##  [392,] "Fusobacterium"                   NA                     
##  [393,] NA                                NA                     
##  [394,] "Enterobacter"                    NA                     
##  [395,] "Veillonella"                     "parvula"              
##  [396,] "Prevotella"                      NA                     
##  [397,] "Prevotella"                      NA                     
##  [398,] "Haemophilus"                     "haemolyticus"         
##  [399,] "Leptotrichia"                    NA                     
##  [400,] "Haemophilus"                     NA                     
##  [401,] "Streptococcus"                   NA                     
##  [402,] "Capnocytophaga"                  "granulosa"            
##  [403,] "Prevotella"                      NA                     
##  [404,] "Prevotella_7"                    "denticola"            
##  [405,] "Catonella"                       "morbi"                
##  [406,] "Veillonella"                     NA                     
##  [407,] "Alloprevotella"                  NA                     
##  [408,] "Gemella"                         "morbillorum"          
##  [409,] "Veillonella"                     NA                     
##  [410,] "Porphyromonas"                   NA                     
##  [411,] "Prevotella_7"                    NA                     
##  [412,] "Prevotella_7"                    NA                     
##  [413,] "Neisseria"                       NA                     
##  [414,] "Stomatobaculum"                  NA                     
##  [415,] "Prevotella_7"                    "denticola"            
##  [416,] "Moraxella"                       NA                     
##  [417,] "Alloprevotella"                  NA                     
##  [418,] "Prevotella_7"                    NA                     
##  [419,] "Leptotrichia"                    NA                     
##  [420,] "Prevotella_7"                    NA                     
##  [421,] "Gemella"                         "morbillorum"          
##  [422,] "Veillonella"                     NA                     
##  [423,] "Veillonella"                     NA                     
##  [424,] "Veillonella"                     NA                     
##  [425,] "Haemophilus"                     NA                     
##  [426,] "Alloprevotella"                  NA                     
##  [427,] "Prevotella"                      "oris"                 
##  [428,] "Prevotella"                      NA                     
##  [429,] "Veillonella"                     NA                     
##  [430,] "Veillonella"                     "denticariosi"         
##  [431,] "Selenomonas_3"                   NA                     
##  [432,] "Prevotella_7"                    NA                     
##  [433,] "Veillonella"                     NA                     
##  [434,] "Prevotella_7"                    NA                     
##  [435,] "Oribacterium"                    NA                     
##  [436,] "Veillonella"                     NA                     
##  [437,] "Prevotella_7"                    NA                     
##  [438,] "Streptococcus"                   "mitis"                
##  [439,] "Prevotella_7"                    NA                     
##  [440,] NA                                NA                     
##  [441,] "Porphyromonas"                   NA                     
##  [442,] "Fusobacterium"                   "nucleatum"            
##  [443,] "Veillonella"                     NA                     
##  [444,] "Veillonella"                     NA                     
##  [445,] "Veillonella"                     NA                     
##  [446,] NA                                NA                     
##  [447,] "Streptococcus"                   NA                     
##  [448,] "Leptotrichia"                    NA                     
##  [449,] "Capnocytophaga"                  "gingivalis"           
##  [450,] "Leptotrichia"                    "hongkongensis"        
##  [451,] "Prevotella"                      NA                     
##  [452,] NA                                NA                     
##  [453,] "Mogibacterium"                   NA                     
##  [454,] "Capnocytophaga"                  NA                     
##  [455,] "Veillonella"                     NA                     
##  [456,] "Ruminococcaceae_UCG-014"         NA                     
##  [457,] NA                                NA                     
##  [458,] "Actinomyces"                     NA                     
##  [459,] "Leptotrichia"                    "hofstadii"            
##  [460,] "Veillonella"                     NA                     
##  [461,] NA                                NA                     
##  [462,] "Leptotrichia"                    NA                     
##  [463,] "Prevotella"                      NA                     
##  [464,] "Streptococcus"                   "oralis"               
##  [465,] "Prevotella_7"                    NA                     
##  [466,] "Dialister"                       "pneumosintes"         
##  [467,] "Prevotella_7"                    NA                     
##  [468,] "Haemophilus"                     NA                     
##  [469,] NA                                NA                     
##  [470,] "Neisseria"                       NA                     
##  [471,] "Haemophilus"                     NA                     
##  [472,] "Alloprevotella"                  NA                     
##  [473,] "Prevotella_7"                    "pleuritidis"          
##  [474,] "Veillonella"                     NA                     
##  [475,] "Prevotella_7"                    "veroralis"            
##  [476,] "Haemophilus"                     NA                     
##  [477,] "Prevotella"                      NA                     
##  [478,] "Aggregatibacter"                 "aphrophilus"          
##  [479,] "Klebsiella"                      NA                     
##  [480,] "Leptotrichia"                    NA                     
##  [481,] "Alloprevotella"                  NA                     
##  [482,] "Prevotella_7"                    NA                     
##  [483,] "Fusobacterium"                   NA                     
##  [484,] "Veillonella"                     NA                     
##  [485,] "Streptococcus"                   NA                     
##  [486,] NA                                NA                     
##  [487,] "Prevotella_7"                    NA                     
##  [488,] "Leptotrichia"                    NA                     
##  [489,] "Porphyromonas"                   NA                     
##  [490,] "Campylobacter"                   "concisus"             
##  [491,] "Veillonella"                     NA                     
##  [492,] NA                                NA                     
##  [493,] "Prevotella_7"                    NA                     
##  [494,] NA                                NA                     
##  [495,] "Prevotella_7"                    NA                     
##  [496,] "Veillonella"                     NA                     
##  [497,] "Klebsiella"                      "michiganensis"        
##  [498,] "Prevotella_7"                    NA                     
##  [499,] "Filifactor"                      "alocis"               
##  [500,] "Veillonella"                     NA                     
##  [501,] "Veillonella"                     NA                     
##  [502,] "Actinomyces"                     NA                     
##  [503,] "Serratia"                        NA                     
##  [504,] "Veillonella"                     NA                     
##  [505,] "Neisseria"                       "elongata"             
##  [506,] "Leptotrichia"                    NA                     
##  [507,] "Actinomyces"                     NA                     
##  [508,] "Alloprevotella"                  NA                     
##  [509,] "Prevotella_7"                    NA                     
##  [510,] "Actinobacillus"                  NA                     
##  [511,] "Veillonella"                     NA                     
##  [512,] "Veillonella"                     NA                     
##  [513,] "Aggregatibacter"                 NA                     
##  [514,] "Veillonella"                     NA                     
##  [515,] "Veillonella"                     NA                     
##  [516,] "Megasphaera"                     NA                     
##  [517,] "Prevotella_7"                    "melaninogenica"       
##  [518,] "Haemophilus"                     NA                     
##  [519,] "Veillonella"                     NA                     
##  [520,] "Actinomyces"                     NA                     
##  [521,] "Anthococcus"                     NA                     
##  [522,] "Actinomyces"                     NA                     
##  [523,] "Parvimonas"                      "micra"                
##  [524,] "Veillonella"                     NA                     
##  [525,] "Fusobacterium"                   NA                     
##  [526,] "Prevotella_7"                    NA                     
##  [527,] "Prevotella_7"                    NA                     
##  [528,] "Neisseria"                       NA                     
##  [529,] "Campylobacter"                   "gracilis"             
##  [530,] "Streptococcus"                   NA                     
##  [531,] "Cardiobacterium"                 "hominis"              
##  [532,] "Prevotella"                      "intermedia"           
##  [533,] NA                                NA                     
##  [534,] "Veillonella"                     NA                     
##  [535,] "Capnocytophaga"                  NA                     
##  [536,] "Prevotella"                      NA                     
##  [537,] "Veillonella"                     NA                     
##  [538,] "Actinobacillus"                  NA                     
##  [539,] "F0058"                           NA                     
##  [540,] "Enterobacter"                    NA                     
##  [541,] "Leptotrichia"                    NA                     
##  [542,] NA                                NA                     
##  [543,] "Porphyromonas"                   NA                     
##  [544,] "Alloprevotella"                  NA                     
##  [545,] "Family_XIII_UCG-001"             NA                     
##  [546,] "Prevotella_7"                    NA                     
##  [547,] "Neisseria"                       "flavescens"           
##  [548,] NA                                NA                     
##  [549,] "Veillonella"                     NA                     
##  [550,] "Alloprevotella"                  NA                     
##  [551,] "Gemella"                         NA                     
##  [552,] "Neisseria"                       NA                     
##  [553,] "Prevotella"                      NA                     
##  [554,] "Capnocytophaga"                  NA                     
##  [555,] "Prevotella"                      NA                     
##  [556,] "Veillonella"                     NA                     
##  [557,] NA                                NA                     
##  [558,] "Streptococcus"                   NA                     
##  [559,] "Prevotella"                      NA                     
##  [560,] "Cricetibacter"                   NA                     
##  [561,] "Porphyromonas"                   "catoniae"             
##  [562,] NA                                NA                     
##  [563,] "Haemophilus"                     NA                     
##  [564,] "Lactococcus"                     NA                     
##  [565,] "Megasphaera"                     NA                     
##  [566,] "Prevotella_7"                    NA                     
##  [567,] "Streptococcus"                   NA                     
##  [568,] "Prevotella_7"                    NA                     
##  [569,] "Alloprevotella"                  NA                     
##  [570,] "Leptotrichia"                    NA                     
##  [571,] "Prevotella_7"                    NA                     
##  [572,] "Tannerella"                      NA                     
##  [573,] "Neisseria"                       NA                     
##  [574,] "Prevotella_7"                    NA                     
##  [575,] "Capnocytophaga"                  NA                     
##  [576,] "Prevotella"                      NA                     
##  [577,] NA                                NA                     
##  [578,] NA                                NA                     
##  [579,] "Veillonella"                     NA                     
##  [580,] "Veillonella"                     NA                     
##  [581,] "Leptotrichia"                    NA                     
##  [582,] "Haemophilus"                     NA                     
##  [583,] "Fusobacterium"                   NA                     
##  [584,] "Porphyromonas"                   NA                     
##  [585,] NA                                NA                     
##  [586,] "Fusobacterium"                   NA                     
##  [587,] "Fusobacterium"                   NA                     
##  [588,] "Tannerella"                      "forsythia"            
##  [589,] "Fusobacterium"                   NA                     
##  [590,] "Prevotella"                      NA                     
##  [591,] "Neisseria"                       NA                     
##  [592,] "Prevotella_7"                    NA                     
##  [593,] "Prevotella_7"                    NA                     
##  [594,] "Prevotella_7"                    NA                     
##  [595,] "Actinobacillus"                  NA                     
##  [596,] "Prevotella_7"                    NA                     
##  [597,] "Veillonella"                     NA                     
##  [598,] "Neisseria"                       NA                     
##  [599,] "Peptostreptococcus"              NA                     
##  [600,] "Veillonella"                     NA                     
##  [601,] "Haemophilus"                     NA                     
##  [602,] "Prevotella_7"                    NA                     
##  [603,] "Prevotella_7"                    NA                     
##  [604,] "Selenomonas_3"                   "infelix"              
##  [605,] "Neisseria"                       NA                     
##  [606,] "Fusobacterium"                   NA                     
##  [607,] "Prevotella_7"                    NA                     
##  [608,] NA                                NA                     
##  [609,] "Alloprevotella"                  NA                     
##  [610,] NA                                NA                     
##  [611,] "Haemophilus"                     NA                     
##  [612,] "Veillonella"                     NA                     
##  [613,] "Fusobacterium"                   "nucleatum"            
##  [614,] "Leptotrichia"                    "trevisanii"           
##  [615,] "Prevotella"                      NA                     
##  [616,] "Veillonella"                     NA                     
##  [617,] "Alloprevotella"                  NA                     
##  [618,] "Veillonella"                     NA                     
##  [619,] "Gemella"                         "morbillorum"          
##  [620,] "Porphyromonas"                   "gingivalis"           
##  [621,] "Selenomonas_3"                   "noxia"                
##  [622,] "F0058"                           NA                     
##  [623,] "Veillonella"                     NA                     
##  [624,] "Prevotella_7"                    NA                     
##  [625,] "Prevotella_6"                    NA                     
##  [626,] "Prevotella"                      NA                     
##  [627,] "Prevotella"                      NA                     
##  [628,] "Fusobacterium"                   NA                     
##  [629,] "Prevotella"                      "intermedia"           
##  [630,] "Rothia"                          "mucilaginosa"         
##  [631,] "Megasphaera"                     NA                     
##  [632,] NA                                NA                     
##  [633,] "Atopobium"                       NA                     
##  [634,] "Selenomonas"                     NA                     
##  [635,] NA                                NA                     
##  [636,] NA                                NA                     
##  [637,] "Porphyromonas"                   "gingivalis"           
##  [638,] "Haemophilus"                     "haemolyticus"         
##  [639,] "Streptococcus"                   NA                     
##  [640,] NA                                NA                     
##  [641,] "Veillonella"                     NA                     
##  [642,] NA                                NA                     
##  [643,] "Serratia"                        NA                     
##  [644,] "Prevotella_7"                    NA                     
##  [645,] "Prevotella"                      "oris"                 
##  [646,] "Prevotella_7"                    NA                     
##  [647,] "Rothia"                          "mucilaginosa"         
##  [648,] "Streptococcus"                   NA                     
##  [649,] "Alloprevotella"                  NA                     
##  [650,] "Filifactor"                      NA                     
##  [651,] "Veillonella"                     NA                     
##  [652,] "Prevotella"                      NA                     
##  [653,] "Veillonella"                     NA                     
##  [654,] NA                                NA                     
##  [655,] "Alloprevotella"                  NA                     
##  [656,] "Veillonella"                     NA                     
##  [657,] "Leptotrichia"                    NA                     
##  [658,] "Porphyromonas"                   NA                     
##  [659,] "Veillonella"                     NA                     
##  [660,] "Leptotrichia"                    NA                     
##  [661,] NA                                NA                     
##  [662,] "Fusobacterium"                   NA                     
##  [663,] NA                                NA                     
##  [664,] "Haemophilus"                     "haemolyticus"         
##  [665,] "Leptotrichia"                    NA                     
##  [666,] "Veillonella"                     NA                     
##  [667,] NA                                NA                     
##  [668,] NA                                NA                     
##  [669,] NA                                NA                     
##  [670,] "Tannerella"                      NA                     
##  [671,] "Actinomyces"                     "odontolyticus"        
##  [672,] "Rothia"                          NA                     
##  [673,] NA                                NA                     
##  [674,] "Alloprevotella"                  NA                     
##  [675,] NA                                NA                     
##  [676,] "Haemophilus"                     NA                     
##  [677,] "Prevotella_7"                    NA                     
##  [678,] "Veillonella"                     NA                     
##  [679,] "Prevotella_7"                    NA                     
##  [680,] NA                                NA                     
##  [681,] "Capnocytophaga"                  "gingivalis"           
##  [682,] "Haemophilus"                     NA                     
##  [683,] "Selenomonas"                     NA                     
##  [684,] "Porphyromonas"                   NA                     
##  [685,] "Prevotella_7"                    NA                     
##  [686,] NA                                NA                     
##  [687,] NA                                NA                     
##  [688,] "Streptococcus"                   NA                     
##  [689,] "Megasphaera"                     NA                     
##  [690,] "Veillonella"                     NA                     
##  [691,] "Porphyromonas"                   NA                     
##  [692,] "Candidatus_Saccharimonas"        NA                     
##  [693,] NA                                NA                     
##  [694,] "Prevotella_7"                    NA                     
##  [695,] "Veillonella"                     NA                     
##  [696,] "Streptococcus"                   NA                     
##  [697,] "Megasphaera"                     NA                     
##  [698,] "Alloprevotella"                  NA                     
##  [699,] "Prevotellaceae_YAB2003_group"    NA                     
##  [700,] "Treponema_2"                     "vincentii"            
##  [701,] "Prevotella_7"                    NA                     
##  [702,] "Veillonella"                     NA                     
##  [703,] "Leptotrichia"                    NA                     
##  [704,] NA                                NA                     
##  [705,] "Prevotella"                      NA                     
##  [706,] NA                                NA                     
##  [707,] "Actinomyces"                     "graevenitzii"         
##  [708,] "Prevotella_7"                    NA                     
##  [709,] "Alloprevotella"                  NA                     
##  [710,] "Haemophilus"                     NA                     
##  [711,] "Prevotella_7"                    NA                     
##  [712,] "Veillonella"                     NA                     
##  [713,] "Porphyromonas"                   NA                     
##  [714,] "Prevotella"                      NA                     
##  [715,] "Neisseria"                       NA                     
##  [716,] NA                                NA                     
##  [717,] NA                                NA                     
##  [718,] "Treponema_2"                     NA                     
##  [719,] "Prevotella_2"                    "conceptionensis"      
##  [720,] NA                                NA                     
##  [721,] "Prevotella"                      "nigrescens"           
##  [722,] "Prevotella"                      NA                     
##  [723,] "Capnocytophaga"                  NA                     
##  [724,] "Lautropia"                       NA                     
##  [725,] "Actinobacillus"                  NA                     
##  [726,] "Neisseria"                       NA                     
##  [727,] "Haemophilus"                     "haemolyticus"         
##  [728,] "Veillonella"                     NA                     
##  [729,] "Veillonella"                     NA                     
##  [730,] "Capnocytophaga"                  NA                     
##  [731,] "Eikenella"                       NA                     
##  [732,] "Fusobacterium"                   NA                     
##  [733,] "Aggregatibacter"                 NA                     
##  [734,] "Selenomonas_3"                   NA                     
##  [735,] NA                                NA                     
##  [736,] "Capnocytophaga"                  NA                     
##  [737,] "Eikenella"                       "corrodens"            
##  [738,] "Actinomyces"                     NA                     
##  [739,] "Neisseria"                       NA                     
##  [740,] "Bergeyella"                      NA                     
##  [741,] "Leptotrichia"                    NA                     
##  [742,] "Streptococcus"                   "salivarius"           
##  [743,] "Rothia"                          NA                     
##  [744,] "Streptococcus"                   NA                     
##  [745,] "Veillonella"                     NA                     
##  [746,] "Veillonella"                     NA                     
##  [747,] "Prevotella"                      NA                     
##  [748,] NA                                NA                     
##  [749,] "Fusobacterium"                   NA                     
##  [750,] NA                                NA                     
##  [751,] "Prevotella"                      NA                     
##  [752,] "Alloprevotella"                  NA                     
##  [753,] "Prevotellaceae_YAB2003_group"    NA                     
##  [754,] "Capnocytophaga"                  NA                     
##  [755,] NA                                NA                     
##  [756,] "Veillonella"                     NA                     
##  [757,] "Alloprevotella"                  NA                     
##  [758,] "Leptotrichia"                    NA                     
##  [759,] "Selenomonas"                     "sputigena"            
##  [760,] "Porphyromonas"                   NA                     
##  [761,] "Veillonella"                     NA                     
##  [762,] "Porphyromonas"                   NA                     
##  [763,] "Prevotella"                      "oulorum"              
##  [764,] "Selenomonas"                     NA                     
##  [765,] "Fusobacterium"                   NA                     
##  [766,] "Veillonella"                     NA                     
##  [767,] "Fusobacterium"                   "nucleatum"            
##  [768,] "Fusobacterium"                   NA                     
##  [769,] "Alloprevotella"                  NA                     
##  [770,] "Alloprevotella"                  NA                     
##  [771,] "Streptococcus"                   NA                     
##  [772,] "Leptotrichia"                    "hofstadii"            
##  [773,] "Alloprevotella"                  NA                     
##  [774,] "Streptococcus"                   NA                     
##  [775,] "Actinomyces"                     NA                     
##  [776,] "Veillonella"                     NA                     
##  [777,] "Veillonella"                     NA                     
##  [778,] "Alloprevotella"                  NA                     
##  [779,] "Neisseria"                       NA                     
##  [780,] "Oribacterium"                    NA                     
##  [781,] "Alloprevotella"                  NA                     
##  [782,] "Haemophilus"                     "haemolyticus"         
##  [783,] "Prevotella_7"                    NA                     
##  [784,] "Aggregatibacter"                 NA                     
##  [785,] "Parvimonas"                      NA                     
##  [786,] "Abiotrophia"                     "defectiva"            
##  [787,] "Prevotella_2"                    NA                     
##  [788,] "Alloprevotella"                  NA                     
##  [789,] "Aggregatibacter"                 "aphrophilus"          
##  [790,] "Lachnoanaerobaculum"             NA                     
##  [791,] "Megasphaera"                     NA                     
##  [792,] "Oribacterium"                    NA                     
##  [793,] "Veillonella"                     NA                     
##  [794,] "Leptotrichia"                    NA                     
##  [795,] "Campylobacter"                   NA                     
##  [796,] "Prevotella"                      NA                     
##  [797,] NA                                NA                     
##  [798,] "Veillonella"                     NA                     
##  [799,] "Veillonella"                     NA                     
##  [800,] "Neisseria"                       NA                     
##  [801,] "Prevotella_7"                    NA                     
##  [802,] "Leptotrichia"                    NA                     
##  [803,] "Porphyromonas"                   NA                     
##  [804,] "Prevotellaceae_YAB2003_group"    NA                     
##  [805,] NA                                NA                     
##  [806,] NA                                NA                     
##  [807,] "Haemophilus"                     NA                     
##  [808,] NA                                NA                     
##  [809,] "Prevotella_7"                    NA                     
##  [810,] NA                                NA                     
##  [811,] "Prevotella_7"                    NA                     
##  [812,] "Actinomyces"                     NA                     
##  [813,] "Leptotrichia"                    NA                     
##  [814,] "Veillonella"                     NA                     
##  [815,] "Prevotella"                      NA                     
##  [816,] "Haemophilus"                     NA                     
##  [817,] "Porphyromonas"                   NA                     
##  [818,] "Selenomonas_3"                   "noxia"                
##  [819,] "Prevotella"                      NA                     
##  [820,] NA                                NA                     
##  [821,] "Bergeriella"                     NA                     
##  [822,] "Leptotrichia"                    NA                     
##  [823,] "Granulicatella"                  NA                     
##  [824,] "Alloprevotella"                  NA                     
##  [825,] "Leptotrichia"                    NA                     
##  [826,] "Prevotellaceae_YAB2003_group"    NA                     
##  [827,] "Alloprevotella"                  NA                     
##  [828,] NA                                NA                     
##  [829,] "Prevotella_7"                    NA                     
##  [830,] "Haemophilus"                     NA                     
##  [831,] "Veillonella"                     NA                     
##  [832,] "Prevotella_7"                    NA                     
##  [833,] NA                                NA                     
##  [834,] NA                                NA                     
##  [835,] "Megasphaera"                     NA                     
##  [836,] "Selenomonas_3"                   NA                     
##  [837,] "Granulicatella"                  NA                     
##  [838,] "Alloprevotella"                  NA                     
##  [839,] "Campylobacter"                   "gracilis"             
##  [840,] NA                                NA                     
##  [841,] "Veillonella"                     NA                     
##  [842,] "Porphyromonas"                   NA                     
##  [843,] "Parvimonas"                      NA                     
##  [844,] "Porphyromonas"                   NA                     
##  [845,] NA                                NA                     
##  [846,] "Actinomyces"                     "graevenitzii"         
##  [847,] "Veillonella"                     NA                     
##  [848,] "Rothia"                          NA                     
##  [849,] "Alloprevotella"                  NA                     
##  [850,] "Haemophilus"                     "influenzae"           
##  [851,] "Prevotella_7"                    NA                     
##  [852,] "Streptococcus"                   NA                     
##  [853,] "Prevotella_7"                    NA                     
##  [854,] "Megasphaera"                     NA                     
##  [855,] "Prevotella_7"                    "denticola"            
##  [856,] "Peptostreptococcus"              NA                     
##  [857,] "Alloprevotella"                  NA                     
##  [858,] "Streptococcus"                   NA                     
##  [859,] "Alloprevotella"                  NA                     
##  [860,] NA                                NA                     
##  [861,] "Defluviitaleaceae_UCG-011"       NA                     
##  [862,] "Lachnoanaerobaculum"             NA                     
##  [863,] "Prevotella_7"                    NA                     
##  [864,] "Alloprevotella"                  NA                     
##  [865,] "Prevotella"                      NA                     
##  [866,] "Porphyromonas"                   NA                     
##  [867,] "Actinomyces"                     NA                     
##  [868,] "Prevotella_7"                    NA                     
##  [869,] "Selenomonas"                     NA                     
##  [870,] "Actinomyces"                     NA                     
##  [871,] "Actinomyces"                     NA                     
##  [872,] "Selenomonas_3"                   NA                     
##  [873,] "Porphyromonas"                   NA                     
##  [874,] "Porphyromonas"                   NA                     
##  [875,] "Actinomyces"                     NA                     
##  [876,] "Leptotrichia"                    NA                     
##  [877,] NA                                NA                     
##  [878,] NA                                NA                     
##  [879,] "Actinomyces"                     NA                     
##  [880,] "Veillonella"                     NA                     
##  [881,] "Aggregatibacter"                 NA                     
##  [882,] "Oribacterium"                    NA                     
##  [883,] "Selenomonas_3"                   NA                     
##  [884,] "Prevotella"                      "intermedia"           
##  [885,] "Alloprevotella"                  NA                     
##  [886,] "Porphyromonas"                   NA                     
##  [887,] NA                                NA                     
##  [888,] "Actinomyces"                     "odontolyticus"        
##  [889,] NA                                NA                     
##  [890,] NA                                NA                     
##  [891,] "Haemophilus"                     NA                     
##  [892,] "Streptococcus"                   NA                     
##  [893,] NA                                NA                     
##  [894,] "Bergeyella"                      NA                     
##  [895,] "Actinomyces"                     NA                     
##  [896,] "Prevotella_7"                    NA                     
##  [897,] "Prevotella_2"                    NA                     
##  [898,] "Prevotella_2"                    NA                     
##  [899,] NA                                NA                     
##  [900,] NA                                NA                     
##  [901,] "Fusobacterium"                   NA                     
##  [902,] "Aggregatibacter"                 NA                     
##  [903,] "Streptococcus"                   NA                     
##  [904,] "Ruminococcaceae_UCG-014"         NA                     
##  [905,] NA                                NA                     
##  [906,] "Prevotella_2"                    NA                     
##  [907,] "Streptococcus"                   NA                     
##  [908,] "Aggregatibacter"                 NA                     
##  [909,] "Veillonella"                     NA                     
##  [910,] "Haemophilus"                     NA                     
##  [911,] "Veillonella"                     "parvula"              
##  [912,] NA                                NA                     
##  [913,] "Neisseria"                       NA                     
##  [914,] "Aggregatibacter"                 NA                     
##  [915,] "Alloscardovia"                   "omnicolens"           
##  [916,] "Veillonella"                     NA                     
##  [917,] "Prevotella_6"                    NA                     
##  [918,] "Lachnoanaerobaculum"             NA                     
##  [919,] "Actinomyces"                     NA                     
##  [920,] "Neisseria"                       NA                     
##  [921,] "Campylobacter"                   "showae"               
##  [922,] NA                                NA                     
##  [923,] "Veillonella"                     NA                     
##  [924,] NA                                NA                     
##  [925,] NA                                NA                     
##  [926,] "Neisseria"                       NA                     
##  [927,] NA                                NA                     
##  [928,] "Fusobacterium"                   NA                     
##  [929,] "Anaeroglobus"                    NA                     
##  [930,] "Alloprevotella"                  NA                     
##  [931,] "Leptotrichia"                    NA                     
##  [932,] "Streptococcus"                   NA                     
##  [933,] NA                                NA                     
##  [934,] "Veillonella"                     NA                     
##  [935,] "Alloprevotella"                  NA                     
##  [936,] NA                                NA                     
##  [937,] "Leptotrichia"                    NA                     
##  [938,] "Solobacterium"                   NA                     
##  [939,] "Streptococcus"                   NA                     
##  [940,] "Prevotella_7"                    NA                     
##  [941,] "Streptococcus"                   NA                     
##  [942,] "Actinomyces"                     NA                     
##  [943,] NA                                NA                     
##  [944,] "Treponema_2"                     "denticola"            
##  [945,] "Streptococcus"                   NA                     
##  [946,] "Actinomyces"                     NA                     
##  [947,] "Streptococcus"                   NA                     
##  [948,] "Sneathia"                        "sanguinegens"         
##  [949,] "Leptotrichia"                    NA                     
##  [950,] "Lachnoanaerobaculum"             NA                     
##  [951,] NA                                NA                     
##  [952,] "Veillonella"                     NA                     
##  [953,] NA                                NA                     
##  [954,] "Treponema_2"                     NA                     
##  [955,] "Alloprevotella"                  NA                     
##  [956,] "Rothia"                          NA                     
##  [957,] "Megasphaera"                     NA                     
##  [958,] "Candidatus_Saccharimonas"        NA                     
##  [959,] "Streptococcus"                   NA                     
##  [960,] "Porphyromonas"                   NA                     
##  [961,] NA                                NA                     
##  [962,] "Streptococcus"                   NA                     
##  [963,] "Veillonella"                     NA                     
##  [964,] "Serratia"                        NA                     
##  [965,] "Fusobacterium"                   NA                     
##  [966,] "Lachnoanaerobaculum"             NA                     
##  [967,] "Prevotella"                      NA                     
##  [968,] "Fusobacterium"                   NA                     
##  [969,] "Ruminococcaceae_UCG-014"         NA                     
##  [970,] "Alloprevotella"                  NA                     
##  [971,] "Fusobacterium"                   NA                     
##  [972,] "Dialister"                       NA                     
##  [973,] NA                                NA                     
##  [974,] "Selenomonas_3"                   NA                     
##  [975,] "Prevotella_7"                    NA                     
##  [976,] "Lachnoanaerobaculum"             NA                     
##  [977,] "Oribacterium"                    NA                     
##  [978,] "Selenomonas_3"                   NA                     
##  [979,] "Actinomyces"                     NA                     
##  [980,] "Actinomyces"                     NA                     
##  [981,] "Fusobacterium"                   NA                     
##  [982,] "Prevotella"                      "intermedia"           
##  [983,] "Leptotrichia"                    NA                     
##  [984,] "Actinomyces"                     NA                     
##  [985,] NA                                NA                     
##  [986,] "Veillonella"                     NA                     
##  [987,] "Actinomyces"                     NA                     
##  [988,] "Aggregatibacter"                 NA                     
##  [989,] "Haemophilus"                     NA                     
##  [990,] "Veillonella"                     NA                     
##  [991,] "Prevotella"                      NA                     
##  [992,] NA                                NA                     
##  [993,] "Mobiluncus"                      NA                     
##  [994,] "Veillonella"                     NA                     
##  [995,] "Veillonella"                     NA                     
##  [996,] "Leptotrichia"                    NA                     
##  [997,] "Leptotrichia"                    NA                     
##  [998,] NA                                NA                     
##  [999,] "Alloprevotella"                  NA                     
## [1000,] "Veillonella"                     NA                     
## [1001,] "Veillonella"                     NA                     
## [1002,] "Veillonella"                     NA                     
## [1003,] "Simonsiella"                     NA                     
## [1004,] "Porphyromonas"                   NA                     
## [1005,] "Prevotella"                      "intermedia"           
## [1006,] "Alloprevotella"                  NA                     
## [1007,] "Lachnoanaerobaculum"             "umeaense"             
## [1008,] "Prevotella_7"                    NA                     
## [1009,] "Prevotella"                      NA                     
## [1010,] "Parvimonas"                      "micra"                
## [1011,] "Veillonella"                     NA                     
## [1012,] NA                                NA                     
## [1013,] "Alysiella"                       NA                     
## [1014,] NA                                NA                     
## [1015,] "Oribacterium"                    NA                     
## [1016,] NA                                NA                     
## [1017,] NA                                NA                     
## [1018,] "Prevotella_2"                    NA                     
## [1019,] "Fusobacterium"                   NA                     
## [1020,] "Serratia"                        NA                     
## [1021,] "Prevotella_7"                    NA                     
## [1022,] "Leptotrichia"                    NA                     
## [1023,] "Treponema_2"                     NA                     
## [1024,] "Granulicatella"                  NA                     
## [1025,] "Granulicatella"                  NA                     
## [1026,] "Corynebacterium"                 "durum"                
## [1027,] "Streptobacillus"                 NA                     
## [1028,] "Alloprevotella"                  NA                     
## [1029,] "Prevotella"                      NA                     
## [1030,] NA                                NA                     
## [1031,] NA                                NA                     
## [1032,] "Leptotrichia"                    "wadei"                
## [1033,] "Fusobacterium"                   NA                     
## [1034,] "Veillonella"                     NA                     
## [1035,] "Prevotella_7"                    NA                     
## [1036,] "Veillonella"                     NA                     
## [1037,] "Veillonella"                     NA                     
## [1038,] "Catonella"                       NA                     
## [1039,] "Alloprevotella"                  NA                     
## [1040,] "Veillonella"                     NA                     
## [1041,] "Prevotella_7"                    NA                     
## [1042,] "Johnsonella"                     NA                     
## [1043,] NA                                NA                     
## [1044,] "Veillonella"                     NA                     
## [1045,] "Actinomyces"                     "odontolyticus"        
## [1046,] "Prevotella"                      NA                     
## [1047,] "Veillonella"                     NA                     
## [1048,] "Prevotella_7"                    NA                     
## [1049,] "Veillonella"                     NA                     
## [1050,] NA                                NA                     
## [1051,] NA                                NA                     
## [1052,] "Granulicatella"                  NA                     
## [1053,] "Prevotellaceae_YAB2003_group"    NA                     
## [1054,] "Aggregatibacter"                 NA                     
## [1055,] "Prevotella_7"                    NA                     
## [1056,] "Streptococcus"                   NA                     
## [1057,] "Haemophilus"                     NA                     
## [1058,] "Prevotella"                      NA                     
## [1059,] "Catonella"                       "morbi"                
## [1060,] "Alloprevotella"                  NA                     
## [1061,] "Veillonella"                     NA                     
## [1062,] "Streptococcus"                   NA                     
## [1063,] "Prevotella"                      NA                     
## [1064,] "Fusobacterium"                   NA                     
## [1065,] "Selenomonas"                     NA                     
## [1066,] "Leptotrichia"                    NA                     
## [1067,] "Veillonella"                     NA                     
## [1068,] "Haemophilus"                     NA                     
## [1069,] "Leptotrichia"                    NA                     
## [1070,] "Veillonella"                     NA                     
## [1071,] "Haemophilus"                     NA                     
## [1072,] "Porphyromonas"                   NA                     
## [1073,] "Alloprevotella"                  NA                     
## [1074,] "Atopobium"                       "rimae"                
## [1075,] "Capnocytophaga"                  NA                     
## [1076,] NA                                NA                     
## [1077,] NA                                NA                     
## [1078,] "Alloprevotella"                  NA                     
## [1079,] "Veillonella"                     NA                     
## [1080,] "Oribacterium"                    NA                     
## [1081,] "Prevotella_7"                    NA                     
## [1082,] NA                                NA                     
## [1083,] "Capnocytophaga"                  NA                     
## [1084,] NA                                NA                     
## [1085,] "Veillonella"                     NA                     
## [1086,] NA                                NA                     
## [1087,] "Streptococcus"                   "salivarius"           
## [1088,] "Prevotella_2"                    NA                     
## [1089,] "Selenomonas"                     NA                     
## [1090,] "Prevotella_7"                    NA                     
## [1091,] "Atopobium"                       NA                     
## [1092,] "Prevotella"                      NA                     
## [1093,] "Prevotella"                      NA                     
## [1094,] "Prevotella_6"                    NA                     
## [1095,] "Capnocytophaga"                  NA                     
## [1096,] "Veillonella"                     NA                     
## [1097,] "Porphyromonas"                   NA                     
## [1098,] NA                                NA                     
## [1099,] "Actinomyces"                     NA                     
## [1100,] "Prevotella"                      NA                     
## [1101,] NA                                NA                     
## [1102,] "Veillonella"                     NA                     
## [1103,] "Veillonella"                     NA                     
## [1104,] "Prevotella"                      NA                     
## [1105,] "Prevotella"                      NA                     
## [1106,] "Selenomonas_3"                   NA                     
## [1107,] "Prevotella_2"                    NA                     
## [1108,] "Fusobacterium"                   "massiliense"          
## [1109,] "Capnocytophaga"                  NA                     
## [1110,] "Veillonella"                     NA                     
## [1111,] "Alloprevotella"                  NA                     
## [1112,] "Leptotrichia"                    NA                     
## [1113,] "Prevotella_2"                    NA                     
## [1114,] "Peptoanaerobacter"               NA                     
## [1115,] "Candidatus_Saccharimonas"        NA                     
## [1116,] NA                                NA                     
## [1117,] "Capnocytophaga"                  NA                     
## [1118,] "Prevotella_7"                    NA                     
## [1119,] "Rothia"                          NA                     
## [1120,] NA                                NA                     
## [1121,] "Veillonella"                     NA                     
## [1122,] NA                                NA                     
## [1123,] "Prevotella_7"                    NA                     
## [1124,] "Campylobacter"                   NA                     
## [1125,] "Actinomyces"                     NA                     
## [1126,] NA                                NA                     
## [1127,] "Porphyromonas"                   NA                     
## [1128,] NA                                NA                     
## [1129,] "Fusobacterium"                   NA                     
## [1130,] "Aggregatibacter"                 NA                     
## [1131,] "Catonella"                       NA                     
## [1132,] "Prevotella_7"                    NA                     
## [1133,] "Prevotella_6"                    NA                     
## [1134,] "Anaeroglobus"                    "geminatus"            
## [1135,] "Haemophilus"                     NA                     
## [1136,] "Haemophilus"                     NA                     
## [1137,] "Prevotella"                      NA                     
## [1138,] "Prevotella_7"                    NA                     
## [1139,] "Kingella"                        NA                     
## [1140,] NA                                NA                     
## [1141,] "Moraxella"                       NA                     
## [1142,] "Prevotella"                      NA                     
## [1143,] "Johnsonella"                     NA                     
## [1144,] "Streptococcus"                   NA                     
## [1145,] "Actinomyces"                     NA                     
## [1146,] NA                                NA                     
## [1147,] "Prevotella_2"                    NA                     
## [1148,] "Veillonella"                     NA                     
## [1149,] "Lachnoanaerobaculum"             NA                     
## [1150,] NA                                NA                     
## [1151,] "Leptotrichia"                    NA                     
## [1152,] "Aggregatibacter"                 NA                     
## [1153,] "Streptococcus"                   NA                     
## [1154,] "Neisseria"                       NA                     
## [1155,] "Prevotella_7"                    NA                     
## [1156,] "Haemophilus"                     NA                     
## [1157,] "Streptococcus"                   NA                     
## [1158,] "Fusobacterium"                   NA                     
## [1159,] NA                                NA                     
## [1160,] "Prevotella_7"                    NA                     
## [1161,] "Prevotella"                      NA                     
## [1162,] "Streptococcus"                   NA                     
## [1163,] "Mycoplasma"                      "salivarium"           
## [1164,] NA                                NA                     
## [1165,] NA                                NA                     
## [1166,] "Veillonella"                     NA                     
## [1167,] "Kingella"                        NA                     
## [1168,] "Megasphaera"                     NA                     
## [1169,] "Veillonella"                     NA                     
## [1170,] NA                                NA                     
## [1171,] "Prevotella"                      NA                     
## [1172,] NA                                NA                     
## [1173,] "Prevotella"                      NA                     
## [1174,] "Selenomonas"                     NA                     
## [1175,] "Veillonella"                     NA                     
## [1176,] "Veillonella"                     NA                     
## [1177,] "Porphyromonas"                   NA                     
## [1178,] "Leptotrichia"                    NA                     
## [1179,] "Eikenella"                       NA                     
## [1180,] "Alloprevotella"                  NA                     
## [1181,] "Stomatobaculum"                  NA                     
## [1182,] "Peptostreptococcus"              NA                     
## [1183,] "Streptococcus"                   "cristatus"            
## [1184,] "Prevotella_7"                    NA                     
## [1185,] NA                                NA                     
## [1186,] "Cricetibacter"                   NA                     
## [1187,] "Prevotella_2"                    NA                     
## [1188,] "Veillonella"                     NA                     
## [1189,] "Prevotella_7"                    NA                     
## [1190,] "Prevotella_6"                    NA                     
## [1191,] "Actinomyces"                     NA                     
## [1192,] "Leptotrichia"                    NA                     
## [1193,] "Actinomyces"                     NA                     
## [1194,] "Haemophilus"                     "haemolyticus"         
## [1195,] "Porphyromonas"                   NA                     
## [1196,] "Capnocytophaga"                  "granulosa"            
## [1197,] "Veillonella"                     NA                     
## [1198,] "Lachnoanaerobaculum"             NA                     
## [1199,] "Streptococcus"                   "sinensis"             
## [1200,] "Prevotella_7"                    NA                     
## [1201,] NA                                NA                     
## [1202,] "Fusobacterium"                   NA                     
## [1203,] "Actinomyces"                     NA                     
## [1204,] "Prevotella_7"                    NA                     
## [1205,] "Fretibacterium"                  NA                     
## [1206,] NA                                NA                     
## [1207,] "Lautropia"                       NA                     
## [1208,] "Selenomonas_3"                   "noxia"                
## [1209,] "Candidatus_Liberibacter"         NA                     
## [1210,] "Dialister"                       NA                     
## [1211,] "Fusobacterium"                   NA                     
## [1212,] NA                                NA                     
## [1213,] NA                                NA                     
## [1214,] "Fusobacterium"                   NA                     
## [1215,] NA                                NA                     
## [1216,] "Lachnospiraceae_AC2044_group"    NA                     
## [1217,] "Treponema_2"                     NA                     
## [1218,] "Alloprevotella"                  "tannerae"             
## [1219,] "Peptostreptococcus"              NA                     
## [1220,] "Peptostreptococcus"              NA                     
## [1221,] "Selenomonas_3"                   NA                     
## [1222,] "Actinomyces"                     NA                     
## [1223,] "Alloprevotella"                  NA                     
## [1224,] NA                                NA                     
## [1225,] "Veillonella"                     NA                     
## [1226,] NA                                NA                     
## [1227,] "Selenomonas_3"                   NA                     
## [1228,] "Lautropia"                       NA                     
## [1229,] "Prevotella"                      NA                     
## [1230,] "Megasphaera"                     NA                     
## [1231,] "Porphyromonas"                   NA                     
## [1232,] "Prevotella_2"                    NA                     
## [1233,] "Bergeyella"                      NA                     
## [1234,] "Veillonella"                     NA                     
## [1235,] "Porphyromonas"                   NA                     
## [1236,] "Stomatobaculum"                  NA                     
## [1237,] "Fusobacterium"                   NA                     
## [1238,] "Rothia"                          NA                     
## [1239,] "Veillonella"                     NA                     
## [1240,] NA                                NA                     
## [1241,] "Porphyromonas"                   NA                     
## [1242,] "Porphyromonas"                   NA                     
## [1243,] NA                                NA                     
## [1244,] NA                                NA                     
## [1245,] "Veillonella"                     NA                     
## [1246,] "Leptotrichia"                    NA                     
## [1247,] NA                                NA                     
## [1248,] "Prevotella_7"                    NA                     
## [1249,] "Selenomonas_3"                   NA                     
## [1250,] NA                                NA                     
## [1251,] NA                                NA                     
## [1252,] "Prevotella"                      NA                     
## [1253,] "Prevotella_7"                    NA                     
## [1254,] NA                                NA                     
## [1255,] "Alloprevotella"                  NA                     
## [1256,] "F0058"                           NA                     
## [1257,] "Selenomonas_3"                   NA                     
## [1258,] "Alloprevotella"                  NA                     
## [1259,] "Anthococcus"                     NA                     
## [1260,] "Prevotella_7"                    NA                     
## [1261,] "Simonsiella"                     NA                     
## [1262,] "Veillonella"                     NA                     
## [1263,] NA                                NA                     
## [1264,] NA                                NA                     
## [1265,] "Actinomyces"                     NA                     
## [1266,] "Veillonella"                     NA                     
## [1267,] "Atopobium"                       NA                     
## [1268,] "Prevotella_7"                    NA                     
## [1269,] NA                                NA                     
## [1270,] "Prevotella_2"                    NA                     
## [1271,] NA                                NA                     
## [1272,] "Veillonella"                     NA                     
## [1273,] NA                                NA                     
## [1274,] "Prevotella_2"                    NA                     
## [1275,] "Treponema_2"                     NA                     
## [1276,] "Alloprevotella"                  NA                     
## [1277,] "Selenomonas_3"                   NA                     
## [1278,] "Veillonella"                     NA                     
## [1279,] "Veillonella"                     NA                     
## [1280,] "Aggregatibacter"                 NA                     
## [1281,] "Actinobacillus"                  NA                     
## [1282,] "Prevotella"                      "maculosa"             
## [1283,] "Candidatus_Saccharimonas"        NA                     
## [1284,] "Selenomonas_3"                   NA                     
## [1285,] "Fusobacterium"                   NA                     
## [1286,] "Corynebacterium"                 "durum"                
## [1287,] "Megasphaera"                     NA                     
## [1288,] "Lachnoanaerobaculum"             NA                     
## [1289,] "Mycoplasma"                      "faucium"              
## [1290,] "Fusobacterium"                   "nucleatum"            
## [1291,] "Prevotella_7"                    "melaninogenica"       
## [1292,] "Ruminococcaceae_UCG-014"         NA                     
## [1293,] "Lachnoanaerobaculum"             NA                     
## [1294,] "Fusobacterium"                   NA                     
## [1295,] "Lautropia"                       NA                     
## [1296,] "Prevotellaceae_YAB2003_group"    NA                     
## [1297,] "Lachnoanaerobaculum"             NA                     
## [1298,] "Bergeyella"                      NA                     
## [1299,] "Prevotella_7"                    NA                     
## [1300,] "Porphyromonas"                   NA                     
## [1301,] "Stomatobaculum"                  NA                     
## [1302,] "Prevotella"                      NA                     
## [1303,] "Veillonella"                     NA                     
## [1304,] "Prevotella"                      "micans"               
## [1305,] "Actinomyces"                     NA                     
## [1306,] "Streptococcus"                   NA                     
## [1307,] "Kingella"                        NA                     
## [1308,] "Simonsiella"                     NA                     
## [1309,] "Treponema_2"                     NA                     
## [1310,] "Actinobacillus"                  NA                     
## [1311,] NA                                NA                     
## [1312,] NA                                NA                     
## [1313,] "Prevotella_7"                    NA                     
## [1314,] "Catonella"                       NA                     
## [1315,] NA                                NA                     
## [1316,] NA                                NA                     
## [1317,] NA                                NA                     
## [1318,] "Veillonella"                     NA                     
## [1319,] "Selenomonas"                     NA                     
## [1320,] "Streptococcus"                   "mutans"               
## [1321,] NA                                NA                     
## [1322,] "Campylobacter"                   NA                     
## [1323,] "Veillonella"                     NA                     
## [1324,] "Streptococcus"                   NA                     
## [1325,] "Prevotella_7"                    NA                     
## [1326,] "Haemophilus"                     NA                     
## [1327,] NA                                NA                     
## [1328,] "Selenomonas_3"                   NA                     
## [1329,] "Prevotella_7"                    NA                     
## [1330,] "Porphyromonas"                   NA                     
## [1331,] "Catonella"                       NA                     
## [1332,] "Prevotella_2"                    NA                     
## [1333,] "Veillonella"                     NA                     
## [1334,] "Prevotella_7"                    NA                     
## [1335,] NA                                NA                     
## [1336,] NA                                NA                     
## [1337,] "Veillonella"                     NA                     
## [1338,] "Johnsonella"                     NA                     
## [1339,] NA                                NA                     
## [1340,] "Veillonella"                     NA                     
## [1341,] NA                                NA                     
## [1342,] "Leptotrichia"                    NA                     
## [1343,] "Alloprevotella"                  NA                     
## [1344,] "Candidatus_Saccharimonas"        NA                     
## [1345,] "Capnocytophaga"                  NA                     
## [1346,] NA                                NA                     
## [1347,] "Alloprevotella"                  NA                     
## [1348,] "Prevotella"                      NA                     
## [1349,] "Alysiella"                       NA                     
## [1350,] NA                                NA                     
## [1351,] "Actinomyces"                     NA                     
## [1352,] "Actinomyces"                     NA                     
## [1353,] "Porphyromonas"                   NA                     
## [1354,] "Selenomonas"                     NA                     
## [1355,] NA                                NA                     
## [1356,] NA                                NA                     
## [1357,] "Prevotella"                      NA                     
## [1358,] "Acholeplasma"                    NA                     
## [1359,] "Veillonella"                     NA                     
## [1360,] "Alloprevotella"                  NA                     
## [1361,] "Kingella"                        NA                     
## [1362,] NA                                NA                     
## [1363,] NA                                NA                     
## [1364,] NA                                NA                     
## [1365,] "Prevotella_7"                    NA                     
## [1366,] "Prevotella_7"                    NA                     
## [1367,] "Porphyromonas"                   NA                     
## [1368,] "Haemophilus"                     NA                     
## [1369,] "Fusobacterium"                   NA                     
## [1370,] "Ruminococcaceae_UCG-014"         NA                     
## [1371,] "Alloprevotella"                  NA                     
## [1372,] "Leptotrichia"                    NA                     
## [1373,] "Gemella"                         "parahaemolysans"      
## [1374,] "Neisseria"                       NA                     
## [1375,] NA                                NA                     
## [1376,] "Neisseria"                       NA                     
## [1377,] "Veillonella"                     NA                     
## [1378,] "Prevotella_7"                    NA                     
## [1379,] "Prevotella_7"                    NA                     
## [1380,] NA                                NA                     
## [1381,] "Butyrivibrio_2"                  NA                     
## [1382,] "Prevotella_7"                    NA                     
## [1383,] "Veillonella"                     NA                     
## [1384,] "Streptococcus"                   NA                     
## [1385,] NA                                NA                     
## [1386,] "Streptococcus"                   NA                     
## [1387,] "Fusobacterium"                   NA                     
## [1388,] "Prevotella_7"                    NA                     
## [1389,] NA                                NA                     
## [1390,] "Alloprevotella"                  NA                     
## [1391,] NA                                NA                     
## [1392,] "Alloprevotella"                  NA                     
## [1393,] "Fusobacterium"                   NA                     
## [1394,] "Prevotella_7"                    NA                     
## [1395,] "Leptotrichia"                    NA                     
## [1396,] NA                                NA                     
## [1397,] "Prevotella_7"                    "fusca"                
## [1398,] "Prevotella_7"                    NA                     
## [1399,] NA                                NA                     
## [1400,] "Bergeyella"                      NA                     
## [1401,] "Streptobacillus"                 NA                     
## [1402,] "Alloprevotella"                  NA                     
## [1403,] "Fusobacterium"                   NA                     
## [1404,] "Alloprevotella"                  NA                     
## [1405,] "Campylobacter"                   NA                     
## [1406,] "Haemophilus"                     NA                     
## [1407,] "Leptotrichia"                    NA                     
## [1408,] "Veillonella"                     NA                     
## [1409,] "Haemophilus"                     NA                     
## [1410,] "Fusobacterium"                   NA                     
## [1411,] "Ruminococcaceae_UCG-014"         NA                     
## [1412,] "Staphylococcus"                  NA                     
## [1413,] NA                                NA                     
## [1414,] "Eikenella"                       "corrodens"            
## [1415,] "Prevotella_2"                    NA                     
## [1416,] NA                                NA                     
## [1417,] "Treponema_2"                     NA                     
## [1418,] "Selenomonas"                     NA                     
## [1419,] NA                                NA                     
## [1420,] "Leptotrichia"                    "wadei"                
## [1421,] NA                                NA                     
## [1422,] "Megasphaera"                     NA                     
## [1423,] "Fusobacterium"                   NA                     
## [1424,] "Prevotella_7"                    NA                     
## [1425,] "Veillonella"                     NA                     
## [1426,] "Campylobacter"                   NA                     
## [1427,] "Prevotellaceae_YAB2003_group"    NA                     
## [1428,] "Actinomyces"                     "odontolyticus"        
## [1429,] "Prevotella"                      "micans"               
## [1430,] "Prevotella"                      "intermedia"           
## [1431,] "Selenomonas"                     NA                     
## [1432,] "Treponema_2"                     NA                     
## [1433,] "Prevotella_2"                    NA                     
## [1434,] "Anthococcus"                     NA                     
## [1435,] "Actinobacillus"                  NA                     
## [1436,] NA                                NA                     
## [1437,] "Parvimonas"                      NA                     
## [1438,] "Veillonella"                     NA                     
## [1439,] "Haemophilus"                     NA                     
## [1440,] "Corynebacterium"                 NA                     
## [1441,] NA                                NA                     
## [1442,] "Veillonella"                     NA                     
## [1443,] "Alloprevotella"                  NA                     
## [1444,] "Bergeyella"                      NA                     
## [1445,] "Shuttleworthia"                  "satelles"             
## [1446,] "Actinobacillus"                  NA                     
## [1447,] NA                                NA                     
## [1448,] "Chelonobacter"                   NA                     
## [1449,] "Prevotella"                      NA                     
## [1450,] "Streptococcus"                   NA                     
## [1451,] NA                                NA                     
## [1452,] "Actinomyces"                     NA                     
## [1453,] "Prevotella"                      "intermedia"           
## [1454,] "Veillonella"                     NA                     
## [1455,] "Megasphaera"                     NA                     
## [1456,] "Serratia"                        NA                     
## [1457,] "Prevotella"                      NA                     
## [1458,] NA                                NA                     
## [1459,] "Veillonella"                     NA                     
## [1460,] "Fusobacterium"                   NA                     
## [1461,] "Porphyromonas"                   NA                     
## [1462,] "Alloprevotella"                  NA                     
## [1463,] "Treponema_2"                     "lecithinolyticum"     
## [1464,] "Johnsonella"                     NA                     
## [1465,] "Leptotrichia"                    NA                     
## [1466,] NA                                NA                     
## [1467,] "Rikenellaceae_RC9_gut_group"     NA                     
## [1468,] NA                                NA                     
## [1469,] NA                                NA                     
## [1470,] "Prevotella"                      NA                     
## [1471,] NA                                NA                     
## [1472,] NA                                NA                     
## [1473,] "Prevotella"                      NA                     
## [1474,] NA                                NA                     
## [1475,] "Prevotella"                      NA                     
## [1476,] "Selenomonas_3"                   NA                     
## [1477,] "Leptotrichia"                    NA                     
## [1478,] "Lachnoanaerobaculum"             "saburreum"            
## [1479,] "Aggregatibacter"                 NA                     
## [1480,] "Selenomonas"                     "sputigena"            
## [1481,] "Prevotella"                      "intermedia"           
## [1482,] "Selenomonas_3"                   NA                     
## [1483,] "Prevotella_6"                    NA                     
## [1484,] "Alloprevotella"                  NA                     
## [1485,] "Prevotella"                      NA                     
## [1486,] "Selenomonas_3"                   "artemidis"            
## [1487,] "Prevotella"                      NA                     
## [1488,] "Veillonella"                     NA                     
## [1489,] "Neisseria"                       NA                     
## [1490,] "Actinomyces"                     NA                     
## [1491,] "Leptotrichia"                    NA                     
## [1492,] "Stomatobaculum"                  NA                     
## [1493,] "Lachnoanaerobaculum"             NA                     
## [1494,] "Veillonella"                     NA                     
## [1495,] "Moraxella"                       NA                     
## [1496,] "Haemophilus"                     NA                     
## [1497,] "Prevotella"                      NA                     
## [1498,] "Actinobacillus"                  NA                     
## [1499,] NA                                NA                     
## [1500,] "Streptococcus"                   NA                     
## [1501,] "Corynebacterium"                 NA                     
## [1502,] "Campylobacter"                   NA                     
## [1503,] "Bergeyella"                      NA                     
## [1504,] "Selenomonas_3"                   NA                     
## [1505,] "Neisseria"                       NA                     
## [1506,] NA                                NA                     
## [1507,] "Prevotella_7"                    NA                     
## [1508,] "Alloprevotella"                  NA                     
## [1509,] "Aggregatibacter"                 NA                     
## [1510,] "Fusobacterium"                   NA                     
## [1511,] "Neisseria"                       NA                     
## [1512,] "Prevotella"                      NA                     
## [1513,] "Capnocytophaga"                  NA                     
## [1514,] "Granulicatella"                  NA                     
## [1515,] NA                                NA                     
## [1516,] "Alloprevotella"                  NA                     
## [1517,] NA                                NA                     
## [1518,] "Catonella"                       NA                     
## [1519,] "Granulicatella"                  NA                     
## [1520,] "Prevotella"                      "intermedia"           
## [1521,] "Leptotrichia"                    NA                     
## [1522,] "Prevotella_6"                    NA                     
## [1523,] "Fusobacterium"                   NA                     
## [1524,] "Simonsiella"                     "muelleri"             
## [1525,] "Atopobium"                       NA                     
## [1526,] "Veillonella"                     NA                     
## [1527,] "Prevotella_7"                    NA                     
## [1528,] NA                                NA                     
## [1529,] "Prevotella"                      NA                     
## [1530,] "Stomatobaculum"                  NA                     
## [1531,] "Campylobacter"                   "concisus"             
## [1532,] NA                                NA                     
## [1533,] NA                                NA                     
## [1534,] "Catonella"                       NA                     
## [1535,] NA                                NA                     
## [1536,] "Prevotella_7"                    NA                     
## [1537,] "Prevotella_2"                    NA                     
## [1538,] NA                                NA                     
## [1539,] "Actinomyces"                     NA                     
## [1540,] NA                                NA                     
## [1541,] "Selenomonas_3"                   NA                     
## [1542,] "Prevotella_6"                    NA                     
## [1543,] "Streptococcus"                   NA                     
## [1544,] "Treponema_2"                     NA                     
## [1545,] "Ruminococcaceae_UCG-014"         NA                     
## [1546,] "Selenomonas_3"                   NA                     
## [1547,] "Alloprevotella"                  NA                     
## [1548,] NA                                NA                     
## [1549,] "Fusobacterium"                   NA                     
## [1550,] "Selenomonas_3"                   NA                     
## [1551,] NA                                NA                     
## [1552,] NA                                NA                     
## [1553,] "Leptotrichia"                    NA                     
## [1554,] "Veillonella"                     NA                     
## [1555,] "Alloprevotella"                  NA                     
## [1556,] "Actinomyces"                     NA                     
## [1557,] "Prevotella_2"                    NA                     
## [1558,] "Megasphaera"                     NA                     
## [1559,] "Fusobacterium"                   NA                     
## [1560,] NA                                NA                     
## [1561,] "Prevotella_7"                    NA                     
## [1562,] "Prevotella_2"                    NA                     
## [1563,] "Selenomonas_3"                   NA                     
## [1564,] "Leptotrichia"                    NA                     
## [1565,] "Prevotella_7"                    NA                     
## [1566,] "Peptococcus"                     NA                     
## [1567,] "Megasphaera"                     NA                     
## [1568,] NA                                NA                     
## [1569,] NA                                NA                     
## [1570,] "Rikenellaceae_RC9_gut_group"     NA                     
## [1571,] "Prevotella_2"                    "loescheii"            
## [1572,] "Prevotella_2"                    NA                     
## [1573,] "Veillonella"                     NA                     
## [1574,] "Fusobacterium"                   NA                     
## [1575,] "Prevotella"                      NA                     
## [1576,] "Porphyromonas"                   NA                     
## [1577,] "Fusobacterium"                   NA                     
## [1578,] "Prevotella_2"                    NA                     
## [1579,] "Alloprevotella"                  NA                     
## [1580,] "Prevotella"                      NA                     
## [1581,] "Prevotellaceae_YAB2003_group"    NA                     
## [1582,] "Veillonella"                     NA                     
## [1583,] "Streptococcus"                   NA                     
## [1584,] "Treponema_2"                     NA                     
## [1585,] NA                                NA                     
## [1586,] "Alloprevotella"                  NA                     
## [1587,] "Veillonella"                     NA                     
## [1588,] "Veillonella"                     NA                     
## [1589,] NA                                NA                     
## [1590,] NA                                NA                     
## [1591,] "Prevotella"                      NA                     
## [1592,] "Granulicatella"                  NA                     
## [1593,] "Haemophilus"                     NA                     
## [1594,] "Capnocytophaga"                  NA                     
## [1595,] "Actinobacillus"                  NA                     
## [1596,] "Veillonella"                     NA                     
## [1597,] "Prevotella_2"                    NA                     
## [1598,] "Prevotella_2"                    NA                     
## [1599,] "Prevotella_7"                    NA                     
## [1600,] "Prevotellaceae_YAB2003_group"    NA                     
## [1601,] "Veillonella"                     NA                     
## [1602,] "Porphyromonas"                   NA                     
## [1603,] "Prevotella_6"                    NA                     
## [1604,] NA                                NA                     
## [1605,] NA                                NA                     
## [1606,] "Streptococcus"                   NA                     
## [1607,] "Prevotella"                      NA                     
## [1608,] "Megasphaera"                     NA                     
## [1609,] "Neisseria"                       NA                     
## [1610,] "Prevotella_2"                    NA                     
## [1611,] "Fusobacterium"                   NA                     
## [1612,] "Leptotrichia"                    NA                     
## [1613,] "Haemophilus"                     NA                     
## [1614,] "Prevotella"                      NA                     
## [1615,] NA                                NA                     
## [1616,] NA                                NA                     
## [1617,] "Actinomyces"                     NA                     
## [1618,] "Veillonella"                     NA                     
## [1619,] "Selenomonas"                     NA                     
## [1620,] "Leptotrichia"                    NA                     
## [1621,] "Actinomyces"                     NA                     
## [1622,] "Haemophilus"                     NA                     
## [1623,] "Fusobacterium"                   "nucleatum"            
## [1624,] "Porphyromonas"                   NA                     
## [1625,] "Leptotrichia"                    NA                     
## [1626,] "Streptococcus"                   NA                     
## [1627,] "Prevotella_7"                    NA                     
## [1628,] "Lachnoanaerobaculum"             NA                     
## [1629,] "Oribacterium"                    NA                     
## [1630,] "Prevotella_7"                    NA                     
## [1631,] "Prevotella"                      NA                     
## [1632,] "Selenomonas"                     NA                     
## [1633,] "Leptotrichia"                    NA                     
## [1634,] NA                                NA                     
## [1635,] "Alloprevotella"                  NA                     
## [1636,] "Haemophilus"                     NA                     
## [1637,] "Selenomonas_3"                   NA                     
## [1638,] "Prevotella"                      NA                     
## [1639,] "Alloprevotella"                  NA                     
## [1640,] "Porphyromonas"                   NA                     
## [1641,] "Actinomyces"                     NA                     
## [1642,] "Prevotella"                      "intermedia"           
## [1643,] NA                                NA                     
## [1644,] NA                                NA                     
## [1645,] "Prevotella_2"                    NA                     
## [1646,] NA                                NA                     
## [1647,] "Lachnoanaerobaculum"             NA                     
## [1648,] "Ruminococcaceae_UCG-014"         NA                     
## [1649,] "Leptotrichia"                    NA                     
## [1650,] "Leptotrichia"                    NA                     
## [1651,] "Stomatobaculum"                  NA                     
## [1652,] "Alloprevotella"                  NA                     
## [1653,] "Selenomonas"                     "sputigena"            
## [1654,] "Campylobacter"                   "showae"               
## [1655,] "Candidatus_Saccharimonas"        NA                     
## [1656,] "Cryptobacterium"                 "curtum"               
## [1657,] "Prevotella_7"                    NA                     
## [1658,] "Treponema_2"                     NA                     
## [1659,] "Haemophilus"                     NA                     
## [1660,] "Rahnella"                        NA                     
## [1661,] "Alloprevotella"                  NA                     
## [1662,] "Alloprevotella"                  NA                     
## [1663,] "Veillonella"                     NA                     
## [1664,] "Porphyromonas"                   NA                     
## [1665,] "Alloprevotella"                  NA                     
## [1666,] "Leptotrichia"                    NA                     
## [1667,] "Leptotrichia"                    NA                     
## [1668,] "Lachnoanaerobaculum"             NA                     
## [1669,] "Treponema_2"                     NA                     
## [1670,] "Leptotrichia"                    NA                     
## [1671,] NA                                NA                     
## [1672,] NA                                NA                     
## [1673,] "Moraxella"                       NA                     
## [1674,] "Haemophilus"                     NA                     
## [1675,] "Prevotella_7"                    NA                     
## [1676,] "Lachnoanaerobaculum"             NA                     
## [1677,] "Anthococcus"                     NA                     
## [1678,] NA                                NA                     
## [1679,] "Cricetibacter"                   NA                     
## [1680,] NA                                NA                     
## [1681,] "Prevotella"                      NA                     
## [1682,] NA                                NA                     
## [1683,] NA                                NA                     
## [1684,] "Haemophilus"                     "parainfluenzae"       
## [1685,] "Selenomonas_3"                   NA                     
## [1686,] "Streptococcus"                   NA                     
## [1687,] "Prevotella"                      NA                     
## [1688,] "Veillonella"                     NA                     
## [1689,] "Fusobacterium"                   NA                     
## [1690,] "Actinobacillus"                  NA                     
## [1691,] "Fusobacterium"                   NA                     
## [1692,] "Prevotellaceae_YAB2003_group"    NA                     
## [1693,] "Streptobacillus"                 NA                     
## [1694,] "Alloprevotella"                  NA                     
## [1695,] "Prevotella_2"                    "saccharolytica"       
## [1696,] "Actinomyces"                     NA                     
## [1697,] "Prevotellaceae_YAB2003_group"    NA                     
## [1698,] "Campylobacter"                   NA                     
## [1699,] "Haemophilus"                     NA                     
## [1700,] "Actinomyces"                     NA                     
## [1701,] NA                                NA                     
## [1702,] NA                                NA                     
## [1703,] "Streptococcus"                   NA                     
## [1704,] "Haemophilus"                     NA                     
## [1705,] "Prevotella"                      NA                     
## [1706,] "Kingella"                        NA                     
## [1707,] "Streptococcus"                   NA                     
## [1708,] "Selenomonas"                     NA                     
## [1709,] "Streptococcus"                   NA                     
## [1710,] "Prevotella_7"                    NA                     
## [1711,] "Selenomonas_3"                   "artemidis"            
## [1712,] NA                                NA                     
## [1713,] "Lachnoanaerobaculum"             NA                     
## [1714,] "Veillonella"                     NA                     
## [1715,] "Alloprevotella"                  NA                     
## [1716,] "Leptotrichia"                    NA                     
## [1717,] "Haemophilus"                     NA                     
## [1718,] "Capnocytophaga"                  "ochracea"             
## [1719,] "Oribacterium"                    NA                     
## [1720,] "Neisseria"                       "oralis"               
## [1721,] "Prevotella"                      NA                     
## [1722,] "Bergeyella"                      NA                     
## [1723,] "Veillonella"                     NA                     
## [1724,] "Prevotella_7"                    NA                     
## [1725,] "Aggregatibacter"                 NA                     
## [1726,] "Prevotella"                      NA                     
## [1727,] "Atopobium"                       NA                     
## [1728,] "Prevotella_7"                    NA                     
## [1729,] "Prevotella_6"                    NA                     
## [1730,] "Fusobacterium"                   NA                     
## [1731,] "Veillonella"                     NA                     
## [1732,] "Corynebacterium"                 "matruchotii"          
## [1733,] "Prevotella"                      NA                     
## [1734,] NA                                NA                     
## [1735,] "Alloprevotella"                  NA                     
## [1736,] "Prevotella"                      NA                     
## [1737,] "Defluviitaleaceae_UCG-011"       NA                     
## [1738,] "Prevotella"                      NA                     
## [1739,] "Prevotella_7"                    NA                     
## [1740,] "Prevotella"                      NA                     
## [1741,] NA                                NA                     
## [1742,] "Alloprevotella"                  NA                     
## [1743,] "Streptococcus"                   NA                     
## [1744,] "Prevotella"                      NA                     
## [1745,] "Neisseria"                       NA                     
## [1746,] "Porphyromonas"                   NA                     
## [1747,] "Porphyromonas"                   NA                     
## [1748,] "Fusobacterium"                   NA                     
## [1749,] "Prevotella_7"                    NA                     
## [1750,] "Alloprevotella"                  NA                     
## [1751,] "Gemella"                         NA                     
## [1752,] "Veillonella"                     NA                     
## [1753,] "Prevotella_7"                    NA                     
## [1754,] "Prevotella_2"                    NA                     
## [1755,] NA                                NA                     
## [1756,] "Granulicatella"                  NA                     
## [1757,] NA                                NA                     
## [1758,] "Alloprevotella"                  NA                     
## [1759,] "Prevotella"                      NA                     
## [1760,] "Candidatus_Saccharimonas"        NA                     
## [1761,] "Porphyromonas"                   NA                     
## [1762,] "Granulicatella"                  NA                     
## [1763,] "Streptococcus"                   NA                     
## [1764,] "Selenomonas"                     NA                     
## [1765,] NA                                NA                     
## [1766,] "Prevotella_6"                    NA                     
## [1767,] "Ruminococcaceae_UCG-014"         NA                     
## [1768,] "Prevotella_7"                    NA                     
## [1769,] "Selenomonas_3"                   NA                     
## [1770,] "Porphyromonas"                   NA                     
## [1771,] "Selenomonas"                     NA                     
## [1772,] "Cardiobacterium"                 NA                     
## [1773,] "Stomatobaculum"                  NA                     
## [1774,] "Actinomyces"                     NA                     
## [1775,] "Ruminococcaceae_UCG-014"         NA                     
## [1776,] "Gemella"                         NA                     
## [1777,] "Haemophilus"                     NA                     
## [1778,] NA                                NA                     
## [1779,] "Fusobacterium"                   NA                     
## [1780,] "Prevotella_7"                    NA                     
## [1781,] NA                                NA                     
## [1782,] "Necropsobacter"                  NA                     
## [1783,] NA                                NA                     
## [1784,] "Alloprevotella"                  NA                     
## [1785,] "Prevotella_7"                    NA                     
## [1786,] "Alloprevotella"                  NA                     
## [1787,] "Centipeda"                       NA                     
## [1788,] "Prevotella"                      NA                     
## [1789,] NA                                NA                     
## [1790,] "Leptotrichia"                    NA                     
## [1791,] "Haemophilus"                     NA                     
## [1792,] NA                                NA                     
## [1793,] NA                                NA                     
## [1794,] "Peptostreptococcus"              NA                     
## [1795,] NA                                NA                     
## [1796,] NA                                NA                     
## [1797,] "Alloprevotella"                  NA                     
## [1798,] "Catonella"                       NA                     
## [1799,] "Streptococcus"                   NA                     
## [1800,] NA                                NA                     
## [1801,] "Oribacterium"                    NA                     
## [1802,] "Veillonella"                     NA                     
## [1803,] "Fusobacterium"                   NA                     
## [1804,] "Johnsonella"                     NA                     
## [1805,] "Fusobacterium"                   NA                     
## [1806,] "Capnocytophaga"                  NA                     
## [1807,] NA                                NA                     
## [1808,] "Granulicatella"                  NA                     
## [1809,] "Rothia"                          NA                     
## [1810,] "Actinomyces"                     NA                     
## [1811,] "Veillonella"                     NA                     
## [1812,] "Veillonella"                     NA                     
## [1813,] "Veillonella"                     NA                     
## [1814,] "Fusobacterium"                   NA                     
## [1815,] NA                                NA                     
## [1816,] NA                                NA                     
## [1817,] "Atlantibacter"                   NA                     
## [1818,] "Prevotella_7"                    NA                     
## [1819,] "Porphyromonas"                   NA                     
## [1820,] "Alloprevotella"                  NA                     
## [1821,] "Alysiella"                       NA                     
## [1822,] "Prevotella"                      NA                     
## [1823,] "Selenomonas_4"                   NA                     
## [1824,] "Veillonella"                     NA                     
## [1825,] "Anthococcus"                     NA                     
## [1826,] "Anaeroglobus"                    NA                     
## [1827,] "Porphyromonas"                   NA                     
## [1828,] "Aggregatibacter"                 NA                     
## [1829,] "Oribacterium"                    NA                     
## [1830,] "Leptotrichia"                    NA                     
## [1831,] NA                                NA                     
## [1832,] "Prevotella_2"                    NA                     
## [1833,] "Corynebacterium"                 NA                     
## [1834,] NA                                NA                     
## [1835,] "Bergeyella"                      NA                     
## [1836,] "Treponema_2"                     NA                     
## [1837,] NA                                NA                     
## [1838,] "Prevotellaceae_YAB2003_group"    NA                     
## [1839,] "Leptotrichia"                    NA                     
## [1840,] "Treponema_2"                     NA                     
## [1841,] "Veillonella"                     NA                     
## [1842,] "Ezakiella"                       NA                     
## [1843,] "Prevotella_7"                    NA                     
## [1844,] "Porphyromonas"                   NA                     
## [1845,] "Peptostreptococcus"              NA                     
## [1846,] "Lautropia"                       NA                     
## [1847,] "Alloprevotella"                  NA                     
## [1848,] "Prevotella_7"                    NA                     
## [1849,] "Prevotella_2"                    NA                     
## [1850,] "Prevotella_2"                    NA                     
## [1851,] "Fretibacterium"                  "fastidiosum"          
## [1852,] "Prevotella_7"                    NA                     
## [1853,] "Alloprevotella"                  NA                     
## [1854,] "Haemophilus"                     NA                     
## [1855,] "Prevotella_7"                    NA                     
## [1856,] "Haemophilus"                     NA                     
## [1857,] "Prevotella_7"                    NA                     
## [1858,] "Porphyromonas"                   NA                     
## [1859,] NA                                NA                     
## [1860,] "Lachnoanaerobaculum"             NA                     
## [1861,] "Actinobacillus"                  NA                     
## [1862,] "Capnocytophaga"                  NA                     
## [1863,] "Alysiella"                       NA                     
## [1864,] "Fusobacterium"                   NA                     
## [1865,] "Aggregatibacter"                 NA                     
## [1866,] "Prevotella_2"                    NA                     
## [1867,] NA                                NA                     
## [1868,] "Oribacterium"                    NA                     
## [1869,] "Haemophilus"                     NA                     
## [1870,] NA                                NA                     
## [1871,] "Haemophilus"                     NA                     
## [1872,] "Streptococcus"                   NA                     
## [1873,] "Haemophilus"                     "haemolyticus"         
## [1874,] "Selenomonas_4"                   NA                     
## [1875,] "Fusobacterium"                   "nucleatum"            
## [1876,] NA                                NA                     
## [1877,] NA                                NA                     
## [1878,] "Capnocytophaga"                  NA                     
## [1879,] NA                                NA                     
## [1880,] "Ruminococcaceae_UCG-014"         NA                     
## [1881,] "Alloprevotella"                  NA                     
## [1882,] "Selenomonas_3"                   NA                     
## [1883,] "Fusobacterium"                   NA                     
## [1884,] "Prevotella_7"                    NA                     
## [1885,] "Actinomyces"                     "odontolyticus"        
## [1886,] NA                                NA                     
## [1887,] NA                                NA                     
## [1888,] "Rothia"                          NA                     
## [1889,] "Alloprevotella"                  NA                     
## [1890,] "Prevotella"                      NA                     
## [1891,] "Neisseria"                       NA                     
## [1892,] "Kingella"                        NA                     
## [1893,] "Veillonella"                     NA                     
## [1894,] NA                                NA                     
## [1895,] "Veillonella"                     NA                     
## [1896,] "Oribacterium"                    NA                     
## [1897,] "Prevotella_7"                    NA                     
## [1898,] "Streptococcus"                   NA                     
## [1899,] "Aggregatibacter"                 NA                     
## [1900,] "Veillonella"                     NA                     
## [1901,] "Johnsonella"                     NA                     
## [1902,] "Veillonella"                     NA                     
## [1903,] "Alloprevotella"                  NA                     
## [1904,] "Alloprevotella"                  NA                     
## [1905,] "Fusobacterium"                   NA                     
## [1906,] NA                                NA                     
## [1907,] NA                                NA                     
## [1908,] "Alloprevotella"                  NA                     
## [1909,] "Catonella"                       NA                     
## [1910,] "Fusobacterium"                   NA                     
## [1911,] "Aggregatibacter"                 NA                     
## [1912,] "Alloprevotella"                  NA                     
## [1913,] "Veillonella"                     NA                     
## [1914,] "Capnocytophaga"                  "gingivalis"           
## [1915,] "Prevotella_7"                    NA                     
## [1916,] "Treponema_2"                     NA                     
## [1917,] "Capnocytophaga"                  NA                     
## [1918,] "Capnocytophaga"                  NA                     
## [1919,] NA                                NA                     
## [1920,] "Lachnoanaerobaculum"             NA                     
## [1921,] "Prevotella_7"                    NA                     
## [1922,] "Stomatobaculum"                  NA                     
## [1923,] "Pseudomonas"                     NA                     
## [1924,] "Rahnella"                        NA                     
## [1925,] "Prevotella_6"                    NA                     
## [1926,] "Prevotella"                      NA                     
## [1927,] "Campylobacter"                   NA                     
## [1928,] "Actinobacillus"                  NA                     
## [1929,] NA                                NA                     
## [1930,] "Alloprevotella"                  NA                     
## [1931,] "Leptotrichia"                    NA                     
## [1932,] "Prevotella"                      NA                     
## [1933,] NA                                NA                     
## [1934,] "Prevotella"                      NA                     
## [1935,] NA                                NA                     
## [1936,] NA                                NA                     
## [1937,] NA                                NA                     
## [1938,] "Prevotella_7"                    NA                     
## [1939,] "Prevotella_2"                    NA                     
## [1940,] NA                                NA                     
## [1941,] "Gemella"                         NA                     
## [1942,] NA                                NA                     
## [1943,] "Prevotella"                      NA                     
## [1944,] "Bergeriella"                     NA                     
## [1945,] "Johnsonella"                     NA                     
## [1946,] "Treponema_2"                     NA                     
## [1947,] "Kingella"                        NA                     
## [1948,] NA                                NA                     
## [1949,] "Capnocytophaga"                  NA                     
## [1950,] NA                                NA                     
## [1951,] "Prevotella_7"                    NA                     
## [1952,] "Lachnoanaerobaculum"             NA                     
## [1953,] "Veillonella"                     NA                     
## [1954,] NA                                NA                     
## [1955,] NA                                NA                     
## [1956,] "Prevotella_2"                    NA                     
## [1957,] "Lachnoanaerobaculum"             NA                     
## [1958,] "Porphyromonas"                   NA                     
## [1959,] "Prevotella_7"                    NA                     
## [1960,] "Prevotella"                      NA                     
## [1961,] "Capnocytophaga"                  NA                     
## [1962,] "Atopobium"                       NA                     
## [1963,] "Alloprevotella"                  NA                     
## [1964,] "Actinomyces"                     NA                     
## [1965,] "Leptotrichia"                    NA                     
## [1966,] NA                                NA                     
## [1967,] "Prevotella_7"                    NA                     
## [1968,] "Veillonella"                     NA                     
## [1969,] "Fusobacterium"                   NA                     
## [1970,] "Neisseria"                       NA                     
## [1971,] NA                                NA                     
## [1972,] "Peptoanaerobacter"               "stomatis"             
## [1973,] "Alloprevotella"                  NA                     
## [1974,] "Johnsonella"                     "ignava"               
## [1975,] "Peptoniphilus"                   "lacrimalis"           
## [1976,] "Alloprevotella"                  NA                     
## [1977,] "Peptostreptococcus"              NA                     
## [1978,] "Alloprevotella"                  NA                     
## [1979,] "Porphyromonas"                   NA                     
## [1980,] "Prevotella_7"                    NA                     
## [1981,] "Prevotella_7"                    NA                     
## [1982,] "Fusobacterium"                   NA                     
## [1983,] "Veillonella"                     NA                     
## [1984,] "Neisseria"                       NA                     
## [1985,] "Porphyromonas"                   NA                     
## [1986,] "Veillonella"                     NA                     
## [1987,] "Haemophilus"                     NA                     
## [1988,] NA                                NA                     
## [1989,] "Leptotrichia"                    NA                     
## [1990,] NA                                NA                     
## [1991,] "Alloprevotella"                  NA                     
## [1992,] "Rikenellaceae_RC9_gut_group"     NA                     
## [1993,] "Capnocytophaga"                  NA                     
## [1994,] NA                                NA                     
## [1995,] "Centipeda"                       NA                     
## [1996,] "Gemella"                         NA                     
## [1997,] "Tannerella"                      NA                     
## [1998,] "Veillonella"                     NA                     
## [1999,] NA                                NA                     
## [2000,] "Neisseria"                       NA                     
## [2001,] "Leptotrichia"                    NA                     
## [2002,] "Peptostreptococcus"              NA                     
## [2003,] "Fusobacterium"                   NA                     
## [2004,] "F0058"                           NA                     
## [2005,] "Prevotella_7"                    NA                     
## [2006,] "Lachnoanaerobaculum"             NA                     
## [2007,] "Prevotella"                      NA                     
## [2008,] "Veillonella"                     NA                     
## [2009,] "Prevotella"                      NA                     
## [2010,] "Haemophilus"                     NA                     
## [2011,] "Prevotella"                      NA                     
## [2012,] "Porphyromonas"                   NA                     
## [2013,] NA                                NA                     
## [2014,] "Prevotella_7"                    NA                     
## [2015,] "Prevotella"                      NA                     
## [2016,] "Veillonella"                     NA                     
## [2017,] "Capnocytophaga"                  NA                     
## [2018,] NA                                NA                     
## [2019,] "Veillonella"                     NA                     
## [2020,] "Fusobacterium"                   "nucleatum"            
## [2021,] "Porphyromonas"                   NA                     
## [2022,] "Actinomyces"                     NA                     
## [2023,] "Actinobacillus"                  NA                     
## [2024,] "Haemophilus"                     "haemolyticus"         
## [2025,] "Neisseria"                       NA                     
## [2026,] "Prevotella_7"                    NA                     
## [2027,] NA                                NA                     
## [2028,] "Prevotella"                      NA                     
## [2029,] "Leptotrichia"                    NA                     
## [2030,] NA                                NA                     
## [2031,] "Leptotrichia"                    NA                     
## [2032,] "Candidatus_Saccharimonas"        NA                     
## [2033,] NA                                NA                     
## [2034,] "Anthococcus"                     NA                     
## [2035,] "Anthococcus"                     NA                     
## [2036,] "Granulicatella"                  NA                     
## [2037,] "Fretibacterium"                  NA                     
## [2038,] "Prevotella_7"                    NA                     
## [2039,] "Fusobacterium"                   NA                     
## [2040,] "Selenomonas"                     NA                     
## [2041,] "Veillonella"                     NA                     
## [2042,] "Prevotella_7"                    NA                     
## [2043,] "Haemophilus"                     NA                     
## [2044,] "Bergeriella"                     NA                     
## [2045,] NA                                NA                     
## [2046,] "Leptotrichia"                    NA                     
## [2047,] "Ruminococcaceae_UCG-014"         NA                     
## [2048,] "Veillonella"                     NA                     
## [2049,] "Campylobacter"                   NA                     
## [2050,] "Prevotella_6"                    NA                     
## [2051,] "Prevotella_7"                    NA                     
## [2052,] "Dialister"                       NA                     
## [2053,] "Prevotella_7"                    NA                     
## [2054,] "Actinomyces"                     NA                     
## [2055,] "Prevotella_7"                    NA                     
## [2056,] "Selenomonas"                     NA                     
## [2057,] "Candidatus_Saccharimonas"        NA                     
## [2058,] NA                                NA                     
## [2059,] "Prevotella"                      NA                     
## [2060,] "Catonella"                       NA                     
## [2061,] "Alloprevotella"                  NA                     
## [2062,] "Veillonella"                     NA                     
## [2063,] NA                                NA                     
## [2064,] "Porphyromonas"                   NA                     
## [2065,] "Veillonella"                     NA                     
## [2066,] "Neisseria"                       NA                     
## [2067,] "Prevotella_7"                    NA                     
## [2068,] "Prevotella"                      NA                     
## [2069,] "Fusobacterium"                   NA                     
## [2070,] "Candidatus_Saccharimonas"        NA                     
## [2071,] "Prevotellaceae_YAB2003_group"    NA                     
## [2072,] NA                                NA                     
## [2073,] "Oribacterium"                    NA                     
## [2074,] "Haemophilus"                     NA                     
## [2075,] NA                                NA                     
## [2076,] "Veillonella"                     NA                     
## [2077,] "Treponema_2"                     NA                     
## [2078,] "Fusobacterium"                   NA                     
## [2079,] "Candidatus_Saccharimonas"        NA                     
## [2080,] "Porphyromonas"                   NA                     
## [2081,] "Capnocytophaga"                  NA                     
## [2082,] "Prevotella_7"                    NA                     
## [2083,] NA                                NA                     
## [2084,] NA                                NA                     
## [2085,] "Neisseria"                       NA                     
## [2086,] "Leptotrichia"                    NA                     
## [2087,] "Lachnoanaerobaculum"             NA                     
## [2088,] "Prevotella"                      NA                     
## [2089,] "Dialister"                       NA                     
## [2090,] "Prevotella_2"                    NA                     
## [2091,] "Treponema_2"                     NA                     
## [2092,] NA                                NA                     
## [2093,] NA                                NA                     
## [2094,] "Prevotella"                      NA                     
## [2095,] "Porphyromonas"                   NA                     
## [2096,] "Fusobacterium"                   NA                     
## [2097,] NA                                NA                     
## [2098,] "Treponema_2"                     "socranskii"           
## [2099,] "Leptotrichia"                    NA                     
## [2100,] "Prevotella"                      NA                     
## [2101,] "Prevotella_7"                    NA                     
## [2102,] "Leptotrichia"                    NA                     
## [2103,] "Catonella"                       NA                     
## [2104,] "Prevotella"                      NA                     
## [2105,] "Leptotrichia"                    NA                     
## [2106,] "Aggregatibacter"                 NA                     
## [2107,] "Prevotella_7"                    NA                     
## [2108,] "Veillonella"                     NA                     
## [2109,] "Prevotella_2"                    NA                     
## [2110,] "Prevotella_2"                    "saccharolytica"       
## [2111,] "Alloprevotella"                  NA                     
## [2112,] NA                                NA                     
## [2113,] "Selenomonas_3"                   NA                     
## [2114,] "Neisseria"                       NA                     
## [2115,] "Prevotellaceae_YAB2003_group"    NA                     
## [2116,] NA                                NA                     
## [2117,] "Granulicatella"                  NA                     
## [2118,] "Prevotella_2"                    NA                     
## [2119,] "Selenomonas_3"                   NA                     
## [2120,] "Haemophilus"                     NA                     
## [2121,] "Capnocytophaga"                  NA                     
## [2122,] "Prevotellaceae_YAB2003_group"    NA                     
## [2123,] NA                                NA                     
## [2124,] "Selenomonas"                     NA                     
## [2125,] "Prevotella_7"                    NA                     
## [2126,] "Streptococcus"                   NA                     
## [2127,] "Alloprevotella"                  NA                     
## [2128,] "Veillonella"                     NA                     
## [2129,] "Streptococcus"                   NA                     
## [2130,] "Haemophilus"                     NA                     
## [2131,] "Alloprevotella"                  NA                     
## [2132,] NA                                NA                     
## [2133,] "Actinomyces"                     NA                     
## [2134,] "Fusobacterium"                   NA                     
## [2135,] "Prevotella_6"                    NA                     
## [2136,] "Veillonella"                     NA                     
## [2137,] "Rothia"                          NA                     
## [2138,] "Alloprevotella"                  NA                     
## [2139,] "Prevotella_7"                    NA                     
## [2140,] NA                                NA                     
## [2141,] "Treponema_2"                     NA                     
## [2142,] NA                                NA                     
## [2143,] NA                                NA                     
## [2144,] "Veillonella"                     NA                     
## [2145,] "Prevotella_7"                    NA                     
## [2146,] "Alloprevotella"                  NA                     
## [2147,] "Alloprevotella"                  NA                     
## [2148,] "Porphyromonas"                   "gingivalis"           
## [2149,] "Leptotrichia"                    NA                     
## [2150,] NA                                NA                     
## [2151,] NA                                NA                     
## [2152,] "Cardiobacterium"                 "valvarum"             
## [2153,] "Campylobacter"                   NA                     
## [2154,] "Prevotella_2"                    NA                     
## [2155,] "Prevotellaceae_YAB2003_group"    NA                     
## [2156,] "Prevotella_7"                    NA                     
## [2157,] "Neisseria"                       NA                     
## [2158,] NA                                NA                     
## [2159,] "Porphyromonas"                   NA                     
## [2160,] "Granulicatella"                  NA                     
## [2161,] "Haemophilus"                     NA                     
## [2162,] "Actinomyces"                     NA                     
## [2163,] NA                                NA                     
## [2164,] "Capnocytophaga"                  NA                     
## [2165,] "Selenomonas_3"                   NA                     
## [2166,] NA                                NA                     
## [2167,] "Prevotellaceae_YAB2003_group"    NA                     
## [2168,] "Treponema_2"                     NA                     
## [2169,] "Prevotella_2"                    NA                     
## [2170,] "Butyrivibrio_2"                  NA                     
## [2171,] "Fusobacterium"                   NA                     
## [2172,] "Oribacterium"                    NA                     
## [2173,] "Actinomyces"                     NA                     
## [2174,] "Alloprevotella"                  NA                     
## [2175,] "Alloprevotella"                  NA                     
## [2176,] "Leptotrichia"                    NA                     
## [2177,] "Capnocytophaga"                  NA                     
## [2178,] NA                                NA                     
## [2179,] "Veillonella"                     NA                     
## [2180,] NA                                NA                     
## [2181,] NA                                NA                     
## [2182,] "Peptostreptococcus"              NA                     
## [2183,] "Porphyromonas"                   NA                     
## [2184,] "Prevotella"                      NA                     
## [2185,] "Prevotella"                      NA                     
## [2186,] "Fusobacterium"                   NA                     
## [2187,] "Veillonella"                     NA                     
## [2188,] "Selenomonas_3"                   NA                     
## [2189,] "Fusobacterium"                   "nucleatum"            
## [2190,] NA                                NA                     
## [2191,] "Prevotella"                      NA                     
## [2192,] "Simonsiella"                     "muelleri"             
## [2193,] "Bergeyella"                      NA                     
## [2194,] "Veillonella"                     NA                     
## [2195,] "Haemophilus"                     NA                     
## [2196,] NA                                NA                     
## [2197,] "Porphyromonas"                   NA                     
## [2198,] NA                                NA                     
## [2199,] "Alloprevotella"                  NA                     
## [2200,] NA                                NA                     
## [2201,] "F0332"                           NA                     
## [2202,] NA                                NA                     
## [2203,] "Selenomonas_3"                   NA                     
## [2204,] NA                                NA                     
## [2205,] NA                                NA                     
## [2206,] "Haemophilus"                     NA                     
## [2207,] "Prevotella_7"                    NA                     
## [2208,] "Aggregatibacter"                 NA                     
## [2209,] "Candidatus_Saccharimonas"        NA                     
## [2210,] "Treponema_2"                     "medium"               
## [2211,] NA                                NA                     
## [2212,] "Prevotella_7"                    NA                     
## [2213,] "Porphyromonas"                   NA                     
## [2214,] "Selenomonas_3"                   NA                     
## [2215,] NA                                NA                     
## [2216,] "Prevotella"                      NA                     
## [2217,] "Capnocytophaga"                  NA                     
## [2218,] "Selenomonas_3"                   NA                     
## [2219,] "Prevotella_7"                    NA                     
## [2220,] NA                                NA                     
## [2221,] NA                                NA                     
## [2222,] "Capnocytophaga"                  NA                     
## [2223,] NA                                NA                     
## [2224,] "Capnocytophaga"                  NA                     
## [2225,] "Veillonella"                     NA                     
## [2226,] NA                                NA                     
## [2227,] "Streptococcus"                   NA                     
## [2228,] "Prevotella_7"                    NA                     
## [2229,] "Lachnoanaerobaculum"             NA                     
## [2230,] NA                                NA                     
## [2231,] "Actinomyces"                     NA                     
## [2232,] "Selenomonas_3"                   NA                     
## [2233,] "Prevotella_2"                    NA                     
## [2234,] "Selenomonas"                     NA                     
## [2235,] "Lachnoanaerobaculum"             NA                     
## [2236,] "Veillonella"                     NA                     
## [2237,] "Prevotella_7"                    NA                     
## [2238,] NA                                NA                     
## [2239,] "Veillonella"                     NA                     
## [2240,] NA                                NA                     
## [2241,] "Prevotella_7"                    NA                     
## [2242,] NA                                NA                     
## [2243,] "Porphyromonas"                   NA                     
## [2244,] "Prevotellaceae_YAB2003_group"    NA                     
## [2245,] "Haemophilus"                     NA                     
## [2246,] NA                                NA                     
## [2247,] "Prevotellaceae_YAB2003_group"    NA                     
## [2248,] "Porphyromonas"                   NA                     
## [2249,] NA                                NA                     
## [2250,] "Candidatus_Saccharimonas"        NA                     
## [2251,] "Atopobium"                       NA                     
## [2252,] "Prevotella_2"                    NA                     
## [2253,] "Selenomonas"                     NA                     
## [2254,] "Dialister"                       NA                     
## [2255,] "Fretibacterium"                  NA                     
## [2256,] "Prevotella"                      "oulorum"              
## [2257,] "Prevotella_7"                    NA                     
## [2258,] "Peptoanaerobacter"               NA                     
## [2259,] "Corynebacterium"                 NA                     
## [2260,] "Gemella"                         NA                     
## [2261,] "Veillonella"                     NA                     
## [2262,] "Necropsobacter"                  NA                     
## [2263,] NA                                NA                     
## [2264,] "Leptotrichia"                    NA                     
## [2265,] "Prevotella_7"                    NA                     
## [2266,] "Leptotrichia"                    NA                     
## [2267,] "Anaeroglobus"                    NA                     
## [2268,] NA                                NA                     
## [2269,] "Capnocytophaga"                  NA                     
## [2270,] "Prevotella_2"                    NA                     
## [2271,] "Leptotrichia"                    NA                     
## [2272,] NA                                NA                     
## [2273,] NA                                NA                     
## [2274,] NA                                NA                     
## [2275,] "Porphyromonas"                   NA                     
## [2276,] "Capnocytophaga"                  NA                     
## [2277,] "Stomatobaculum"                  NA                     
## [2278,] "Selenomonas"                     NA                     
## [2279,] "Actinobacillus"                  NA                     
## [2280,] NA                                NA                     
## [2281,] NA                                NA                     
## [2282,] NA                                NA                     
## [2283,] "Prevotella_7"                    NA                     
## [2284,] "Prevotella_2"                    NA                     
## [2285,] "Prevotella"                      NA                     
## [2286,] "Lachnoanaerobaculum"             NA                     
## [2287,] "Alloprevotella"                  NA                     
## [2288,] NA                                NA                     
## [2289,] NA                                NA                     
## [2290,] "Leptotrichia"                    NA                     
## [2291,] "Veillonella"                     NA                     
## [2292,] "Aggregatibacter"                 NA                     
## [2293,] "Prevotella"                      NA                     
## [2294,] "Alloprevotella"                  NA                     
## [2295,] "Porphyromonas"                   NA                     
## [2296,] "Neisseria"                       NA                     
## [2297,] "Prevotella_7"                    NA                     
## [2298,] "Prevotella"                      NA                     
## [2299,] "Prevotella_7"                    NA                     
## [2300,] "Prevotella_7"                    NA                     
## [2301,] NA                                NA                     
## [2302,] NA                                NA                     
## [2303,] "Scardovia"                       "wiggsiae"             
## [2304,] NA                                NA                     
## [2305,] "Treponema_2"                     NA                     
## [2306,] "Bergeyella"                      NA                     
## [2307,] NA                                NA                     
## [2308,] "Capnocytophaga"                  NA                     
## [2309,] "Peptostreptococcus"              NA                     
## [2310,] "Prevotella_7"                    NA                     
## [2311,] NA                                NA                     
## [2312,] "Veillonella"                     NA                     
## [2313,] NA                                NA                     
## [2314,] "Prevotella"                      NA                     
## [2315,] NA                                NA                     
## [2316,] NA                                NA                     
## [2317,] NA                                NA                     
## [2318,] "Porphyromonas"                   NA                     
## [2319,] "Stomatobaculum"                  NA                     
## [2320,] NA                                NA                     
## [2321,] "Alloprevotella"                  NA                     
## [2322,] "Prevotella_7"                    NA                     
## [2323,] "Prevotella_6"                    NA                     
## [2324,] "Alloprevotella"                  NA                     
## [2325,] "Prevotella_6"                    NA                     
## [2326,] NA                                NA                     
## [2327,] NA                                NA                     
## [2328,] "Prevotella_7"                    NA                     
## [2329,] "Ruminococcaceae_UCG-014"         NA                     
## [2330,] "Alloprevotella"                  NA                     
## [2331,] "Porphyromonas"                   NA                     
## [2332,] NA                                NA                     
## [2333,] "Prevotella"                      NA                     
## [2334,] "Prevotellaceae_YAB2003_group"    NA                     
## [2335,] "Neisseria"                       "elongata"             
## [2336,] "Kingella"                        NA                     
## [2337,] "Prevotella"                      NA                     
## [2338,] "Prevotella"                      NA                     
## [2339,] "Streptococcus"                   NA                     
## [2340,] "Alloprevotella"                  NA                     
## [2341,] NA                                NA                     
## [2342,] "Lachnoanaerobaculum"             NA                     
## [2343,] "Haemophilus"                     "parainfluenzae"       
## [2344,] "Porphyromonas"                   NA                     
## [2345,] "Porphyromonas"                   NA                     
## [2346,] NA                                NA                     
## [2347,] NA                                NA                     
## [2348,] "Corynebacterium"                 NA                     
## [2349,] "Fretibacterium"                  NA                     
## [2350,] "Treponema_2"                     NA                     
## [2351,] NA                                NA                     
## [2352,] "Prevotella"                      NA                     
## [2353,] "Actinomyces"                     NA                     
## [2354,] NA                                NA                     
## [2355,] "Lachnoanaerobaculum"             NA                     
## [2356,] "Prevotella"                      NA                     
## [2357,] "Veillonella"                     NA                     
## [2358,] "Prevotella"                      NA                     
## [2359,] "Prevotella_7"                    "dentalis"             
## [2360,] "Selenomonas_3"                   "infelix"              
## [2361,] NA                                NA                     
## [2362,] "Selenomonas_3"                   NA                     
## [2363,] "Prevotella_7"                    NA                     
## [2364,] NA                                NA                     
## [2365,] "Selenomonas"                     NA                     
## [2366,] NA                                NA                     
## [2367,] "Actinomyces"                     NA                     
## [2368,] "Prevotella"                      NA                     
## [2369,] "Eikenella"                       NA                     
## [2370,] "Oceanivirga"                     NA                     
## [2371,] "Porphyromonas"                   NA                     
## [2372,] "Veillonella"                     NA                     
## [2373,] "Alloprevotella"                  NA                     
## [2374,] "Alloprevotella"                  NA                     
## [2375,] "Selenomonas_3"                   NA                     
## [2376,] "Peptoanaerobacter"               "stomatis"             
## [2377,] "Alysiella"                       NA                     
## [2378,] "Treponema_2"                     NA                     
## [2379,] "Prevotella"                      NA                     
## [2380,] "Fusobacterium"                   NA                     
## [2381,] "Prevotella_7"                    NA                     
## [2382,] "Leptotrichia"                    NA                     
## [2383,] NA                                NA                     
## [2384,] "F0058"                           NA                     
## [2385,] "Haemophilus"                     NA                     
## [2386,] "Alloprevotella"                  NA                     
## [2387,] "Veillonella"                     NA                     
## [2388,] "Actinomyces"                     NA                     
## [2389,] "Alloprevotella"                  NA                     
## [2390,] "Veillonella"                     NA                     
## [2391,] "Leptotrichia"                    NA                     
## [2392,] "Selenomonas_3"                   NA                     
## [2393,] "Leptotrichia"                    NA                     
## [2394,] NA                                NA                     
## [2395,] "Selenomonas"                     NA                     
## [2396,] "Veillonella"                     NA                     
## [2397,] "Veillonella"                     NA                     
## [2398,] "Corynebacterium"                 NA                     
## [2399,] "Actinomyces"                     NA                     
## [2400,] "Prevotella_7"                    NA                     
## [2401,] "Atopobium"                       NA                     
## [2402,] "Treponema_2"                     NA                     
## [2403,] "Prevotella_7"                    NA                     
## [2404,] NA                                NA                     
## [2405,] "Lachnoanaerobaculum"             NA                     
## [2406,] "Veillonella"                     NA                     
## [2407,] "Alloprevotella"                  NA                     
## [2408,] "Prevotella_7"                    NA                     
## [2409,] "Leptotrichia"                    NA                     
## [2410,] "Prevotella_7"                    NA                     
## [2411,] "Veillonella"                     NA                     
## [2412,] NA                                NA                     
## [2413,] "Candidatus_Saccharimonas"        NA                     
## [2414,] "Prevotellaceae_YAB2003_group"    NA                     
## [2415,] "Selenomonas_3"                   NA                     
## [2416,] "Selenomonas_3"                   NA                     
## [2417,] "Veillonella"                     NA                     
## [2418,] "Alloprevotella"                  NA                     
## [2419,] "Prevotella_7"                    NA                     
## [2420,] "Johnsonella"                     NA                     
## [2421,] "Haemophilus"                     NA                     
## [2422,] NA                                NA                     
## [2423,] NA                                NA                     
## [2424,] NA                                NA                     
## [2425,] "Alloprevotella"                  NA                     
## [2426,] NA                                NA                     
## [2427,] "Prevotella_6"                    NA                     
## [2428,] "Porphyromonas"                   NA                     
## [2429,] "Capnocytophaga"                  NA                     
## [2430,] "Alloprevotella"                  NA                     
## [2431,] "Capnocytophaga"                  NA                     
## [2432,] "Johnsonella"                     NA                     
## [2433,] "Selenomonas_3"                   NA                     
## [2434,] "Prevotella_2"                    NA                     
## [2435,] NA                                NA                     
## [2436,] "Prevotella_7"                    NA                     
## [2437,] "Treponema_2"                     NA                     
## [2438,] NA                                NA                     
## [2439,] "Prevotella_7"                    NA                     
## [2440,] NA                                NA                     
## [2441,] "Actinomyces"                     NA                     
## [2442,] "Porphyromonas"                   NA                     
## [2443,] NA                                NA                     
## [2444,] "Haemophilus"                     NA                     
## [2445,] "Ruminococcaceae_UCG-014"         NA                     
## [2446,] "Alloprevotella"                  NA                     
## [2447,] "Prevotella_6"                    NA                     
## [2448,] "Anthococcus"                     NA                     
## [2449,] NA                                NA                     
## [2450,] NA                                NA                     
## [2451,] NA                                NA                     
## [2452,] "Prevotella_7"                    NA                     
## [2453,] "Bergeriella"                     NA                     
## [2454,] "Prevotellaceae_YAB2003_group"    NA                     
## [2455,] "Johnsonella"                     NA                     
## [2456,] "Actinomyces"                     NA                     
## [2457,] "Kingella"                        NA                     
## [2458,] "Prevotella_2"                    NA                     
## [2459,] "Lachnoanaerobaculum"             NA                     
## [2460,] "Treponema_2"                     NA                     
## [2461,] "Stomatobaculum"                  NA                     
## [2462,] "Dialister"                       NA                     
## [2463,] "Alloprevotella"                  NA                     
## [2464,] "Peptostreptococcus"              NA                     
## [2465,] "Prevotella"                      NA                     
## [2466,] "Prevotella_6"                    NA                     
## [2467,] NA                                NA                     
## [2468,] "Bergeyella"                      NA                     
## [2469,] "Prevotella"                      NA                     
## [2470,] "Prevotella_2"                    NA                     
## [2471,] "Bergeriella"                     NA                     
## [2472,] "Aggregatibacter"                 NA                     
## [2473,] NA                                NA                     
## [2474,] "Peptococcus"                     NA                     
## [2475,] "Prevotellaceae_YAB2003_group"    NA                     
## [2476,] NA                                NA                     
## [2477,] NA                                NA                     
## [2478,] "Selenomonas_3"                   NA                     
## [2479,] "Centipeda"                       NA                     
## [2480,] NA                                NA                     
## [2481,] NA                                NA                     
## [2482,] "Lachnoanaerobaculum"             NA                     
## [2483,] "Prevotella_7"                    NA                     
## [2484,] "Treponema_2"                     NA                     
## [2485,] "Neisseria"                       NA                     
## [2486,] "Family_XIII_UCG-001"             NA                     
## [2487,] "Prevotella_7"                    NA                     
## [2488,] "Prevotella_7"                    NA                     
## [2489,] "Haemophilus"                     NA                     
## [2490,] "Prevotella"                      NA                     
## [2491,] NA                                NA                     
## [2492,] "Veillonella"                     NA                     
## [2493,] NA                                NA                     
## [2494,] "Prevotella"                      NA                     
## [2495,] "Capnocytophaga"                  NA                     
## [2496,] "Fusobacterium"                   NA                     
## [2497,] "Actinomyces"                     NA                     
## [2498,] NA                                NA                     
## [2499,] NA                                NA                     
## [2500,] "Corynebacterium"                 NA                     
## [2501,] "Selenomonas_3"                   NA                     
## [2502,] "Prevotella"                      NA                     
## [2503,] "Alloprevotella"                  NA                     
## [2504,] NA                                NA                     
## [2505,] "Fusobacterium"                   "simiae"               
## [2506,] "Alloprevotella"                  NA                     
## [2507,] "Capnocytophaga"                  "ochracea"             
## [2508,] "Prevotella"                      NA                     
## [2509,] "Streptococcus"                   NA                     
## [2510,] NA                                NA                     
## [2511,] "Capnocytophaga"                  NA                     
## [2512,] "Neisseria"                       "macacae"              
## [2513,] "Lachnoanaerobaculum"             NA                     
## [2514,] NA                                NA                     
## [2515,] "Alloprevotella"                  NA                     
## [2516,] "Kingella"                        NA                     
## [2517,] "Rothia"                          "mucilaginosa"         
## [2518,] NA                                NA                     
## [2519,] "Actinomyces"                     NA                     
## [2520,] "Actinomyces"                     NA                     
## [2521,] "Bergeriella"                     NA                     
## [2522,] NA                                NA                     
## [2523,] "Alloprevotella"                  NA                     
## [2524,] "Selenomonas"                     NA                     
## [2525,] "Neisseria"                       NA                     
## [2526,] NA                                NA                     
## [2527,] "Prevotella"                      NA                     
## [2528,] NA                                NA                     
## [2529,] NA                                NA                     
## [2530,] "Tannerella"                      NA                     
## [2531,] "Selenomonas_3"                   NA                     
## [2532,] "Dialister"                       NA                     
## [2533,] "Alloprevotella"                  NA                     
## [2534,] "Capnocytophaga"                  NA                     
## [2535,] "Prevotella_7"                    NA                     
## [2536,] NA                                NA                     
## [2537,] "Anthococcus"                     NA                     
## [2538,] NA                                NA                     
## [2539,] NA                                NA                     
## [2540,] "Haemophilus"                     NA                     
## [2541,] "Prevotellaceae_YAB2003_group"    NA                     
## [2542,] "Ruminococcaceae_UCG-014"         NA                     
## [2543,] NA                                NA                     
## [2544,] "Prevotella_6"                    NA                     
## [2545,] NA                                NA                     
## [2546,] NA                                NA                     
## [2547,] "Leptotrichia"                    "wadei"                
## [2548,] NA                                NA                     
## [2549,] "Veillonella"                     NA                     
## [2550,] NA                                NA                     
## [2551,] "Megasphaera"                     NA                     
## [2552,] "Prevotella_7"                    NA                     
## [2553,] "Leptotrichia"                    NA                     
## [2554,] "Leptotrichia"                    NA                     
## [2555,] "Streptobacillus"                 NA                     
## [2556,] "Capnocytophaga"                  NA                     
## [2557,] "Haemophilus"                     NA                     
## [2558,] "Treponema_2"                     NA                     
## [2559,] NA                                NA                     
## [2560,] "Dialister"                       NA                     
## [2561,] "Veillonella"                     NA                     
## [2562,] "Treponema_2"                     NA                     
## [2563,] "Treponema_2"                     NA                     
## [2564,] NA                                NA                     
## [2565,] "Parvimonas"                      NA                     
## [2566,] NA                                NA                     
## [2567,] "Haemophilus"                     NA                     
## [2568,] "Anthococcus"                     NA                     
## [2569,] "Prevotella_6"                    NA                     
## [2570,] "Veillonella"                     NA                     
## [2571,] "Prevotella_6"                    NA                     
## [2572,] "Prevotella_7"                    NA                     
## [2573,] "Aggregatibacter"                 NA                     
## [2574,] "Leptotrichia"                    NA                     
## [2575,] NA                                NA                     
## [2576,] "Prevotella_7"                    NA                     
## [2577,] "Actinomyces"                     NA                     
## [2578,] "Leptotrichia"                    NA                     
## [2579,] "Capnocytophaga"                  NA                     
## [2580,] "Porphyromonas"                   NA                     
## [2581,] "Porphyromonas"                   NA                     
## [2582,] "Fusobacterium"                   NA                     
## [2583,] "Prevotella_7"                    NA                     
## [2584,] "Selenomonas_3"                   NA                     
## [2585,] "Capnocytophaga"                  NA                     
## [2586,] NA                                NA                     
## [2587,] "Rikenellaceae_RC9_gut_group"     NA                     
## [2588,] NA                                NA                     
## [2589,] "Peptostreptococcus"              NA                     
## [2590,] "Prevotella_2"                    "marshii"              
## [2591,] NA                                NA                     
## [2592,] "Veillonella"                     NA                     
## [2593,] "Lactobacillus"                   "delbrueckii"          
## [2594,] "Neisseria"                       NA                     
## [2595,] "Prevotella"                      NA                     
## [2596,] "Streptococcus"                   NA                     
## [2597,] "Prevotella_7"                    NA                     
## [2598,] NA                                NA                     
## [2599,] NA                                NA                     
## [2600,] "Selenomonas_3"                   NA                     
## [2601,] "Selenomonas_3"                   NA                     
## [2602,] "Cardiobacterium"                 NA                     
## [2603,] "Prevotella_7"                    NA                     
## [2604,] "Veillonella"                     NA                     
## [2605,] "Prevotella"                      NA                     
## [2606,] NA                                NA                     
## [2607,] NA                                NA                     
## [2608,] NA                                NA                     
## [2609,] NA                                NA                     
## [2610,] NA                                NA                     
## [2611,] "Filifactor"                      NA                     
## [2612,] "Prevotella_7"                    NA                     
## [2613,] NA                                NA                     
## [2614,] "Prevotella"                      NA                     
## [2615,] NA                                NA                     
## [2616,] "Prevotella"                      NA                     
## [2617,] "Haemophilus"                     NA                     
## [2618,] "Veillonella"                     NA                     
## [2619,] "Aggregatibacter"                 NA                     
## [2620,] "Leptotrichia"                    NA                     
## [2621,] "Fusobacterium"                   NA                     
## [2622,] "Prevotella"                      NA                     
## [2623,] "Capnocytophaga"                  NA                     
## [2624,] "Atlantibacter"                   NA                     
## [2625,] "Lachnoanaerobaculum"             NA                     
## [2626,] NA                                NA                     
## [2627,] "Veillonella"                     NA                     
## [2628,] NA                                NA                     
## [2629,] "Corynebacterium"                 NA                     
## [2630,] "Oribacterium"                    NA                     
## [2631,] NA                                NA                     
## [2632,] "Treponema_2"                     NA                     
## [2633,] "Gemella"                         NA                     
## [2634,] "Prevotella_7"                    NA                     
## [2635,] NA                                NA                     
## [2636,] "Dialister"                       NA                     
## [2637,] NA                                NA                     
## [2638,] NA                                NA                     
## [2639,] "Alloprevotella"                  NA                     
## [2640,] NA                                NA                     
## [2641,] "Campylobacter"                   "curvus"               
## [2642,] "Veillonella"                     NA                     
## [2643,] "Prevotellaceae_YAB2003_group"    NA                     
## [2644,] "Streptococcus"                   NA                     
## [2645,] NA                                NA                     
## [2646,] "Parvimonas"                      NA                     
## [2647,] "Prevotella"                      NA                     
## [2648,] "Ruminococcaceae_UCG-014"         NA                     
## [2649,] "Capnocytophaga"                  NA                     
## [2650,] "Streptococcus"                   NA                     
## [2651,] "Treponema_2"                     NA                     
## [2652,] "Actinobacillus"                  NA                     
## [2653,] "Leptotrichia"                    NA                     
## [2654,] "Prevotella_7"                    "baroniae"             
## [2655,] "Selenomonas_3"                   NA                     
## [2656,] "Tannerella"                      NA                     
## [2657,] "Alloprevotella"                  NA                     
## [2658,] "Peptoanaerobacter"               NA                     
## [2659,] "Prevotellaceae_YAB2003_group"    NA                     
## [2660,] "Capnocytophaga"                  NA                     
## [2661,] "Capnocytophaga"                  NA                     
## [2662,] "Selenomonas_3"                   NA                     
## [2663,] "Actinomyces"                     NA                     
## [2664,] "Actinomyces"                     NA                     
## [2665,] "Leptotrichia"                    NA                     
## [2666,] "Selenomonas_3"                   NA                     
## [2667,] NA                                NA                     
## [2668,] "Selenomonas_3"                   NA                     
## [2669,] "Parvimonas"                      NA                     
## [2670,] "Capnocytophaga"                  NA                     
## [2671,] "Veillonella"                     NA                     
## [2672,] "Necropsobacter"                  NA                     
## [2673,] "Prevotella"                      "intermedia"           
## [2674,] "Prevotella"                      NA                     
## [2675,] "Treponema_2"                     "maltophilum"          
## [2676,] "Prevotella"                      NA                     
## [2677,] "Actinomyces"                     NA                     
## [2678,] NA                                NA                     
## [2679,] "Raoultella"                      NA                     
## [2680,] NA                                NA                     
## [2681,] "Lautropia"                       NA                     
## [2682,] "Prevotella_7"                    NA                     
## [2683,] "Actinomyces"                     NA                     
## [2684,] "Capnocytophaga"                  NA                     
## [2685,] "Prevotella_7"                    NA                     
## [2686,] NA                                NA                     
## [2687,] "Atopobium"                       NA                     
## [2688,] NA                                NA                     
## [2689,] NA                                NA                     
## [2690,] NA                                NA                     
## [2691,] NA                                NA                     
## [2692,] "Fusobacterium"                   NA                     
## [2693,] NA                                NA                     
## [2694,] "Prevotella"                      NA                     
## [2695,] "Prevotella_2"                    NA                     
## [2696,] NA                                NA                     
## [2697,] "Veillonella"                     NA                     
## [2698,] "Veillonella"                     NA                     
## [2699,] "Veillonella"                     NA                     
## [2700,] "Prevotella_7"                    NA                     
## [2701,] NA                                NA                     
## [2702,] "Veillonella"                     NA                     
## [2703,] NA                                NA                     
## [2704,] "Prevotella"                      NA                     
## [2705,] "Chelonobacter"                   NA                     
## [2706,] "Fusobacterium"                   NA                     
## [2707,] "Selenomonas_3"                   NA                     
## [2708,] "Fusobacterium"                   NA                     
## [2709,] "Veillonella"                     NA                     
## [2710,] "Prevotella_7"                    NA                     
## [2711,] "Capnocytophaga"                  NA                     
## [2712,] NA                                NA                     
## [2713,] "Veillonella"                     NA                     
## [2714,] "Fusobacterium"                   NA                     
## [2715,] "Prevotella_7"                    NA                     
## [2716,] "Leptotrichia"                    NA                     
## [2717,] NA                                NA                     
## [2718,] "Prevotella_2"                    NA                     
## [2719,] "Atlantibacter"                   NA                     
## [2720,] NA                                NA                     
## [2721,] "Treponema_2"                     "maltophilum"          
## [2722,] NA                                NA                     
## [2723,] "Oribacterium"                    NA                     
## [2724,] "Prevotella_7"                    NA                     
## [2725,] "Prevotella"                      NA                     
## [2726,] "Kingella"                        NA                     
## [2727,] "Prevotella_7"                    NA                     
## [2728,] "Actinomyces"                     NA                     
## [2729,] "Prevotella_7"                    NA                     
## [2730,] NA                                NA                     
## [2731,] NA                                NA                     
## [2732,] "Kingella"                        NA                     
## [2733,] "Prevotella_2"                    NA                     
## [2734,] "Leptotrichia"                    NA                     
## [2735,] "Fusobacterium"                   NA                     
## [2736,] "Leptotrichia"                    NA                     
## [2737,] "Aggregatibacter"                 "actinomycetemcomitans"
## [2738,] "Prevotella_7"                    NA                     
## [2739,] NA                                NA                     
## [2740,] NA                                NA                     
## [2741,] NA                                NA                     
## [2742,] NA                                NA                     
## [2743,] "Solobacterium"                   "moorei"               
## [2744,] "Selenomonas_4"                   NA                     
## [2745,] "Neisseria"                       NA                     
## [2746,] "Kingella"                        NA                     
## [2747,] "Centipeda"                       NA                     
## [2748,] "Prevotella_2"                    NA                     
## [2749,] "Prevotella_6"                    NA                     
## [2750,] "Prevotella_6"                    NA                     
## [2751,] NA                                NA                     
## [2752,] "Prevotella_7"                    NA                     
## [2753,] "Tannerella"                      NA                     
## [2754,] "Alloprevotella"                  NA                     
## [2755,] "Eikenella"                       NA                     
## [2756,] NA                                NA                     
## [2757,] "Capnocytophaga"                  NA                     
## [2758,] "Fusobacterium"                   NA                     
## [2759,] "Prevotella_7"                    NA                     
## [2760,] "Atopobium"                       NA                     
## [2761,] "Prevotella_7"                    NA                     
## [2762,] "Porphyromonas"                   NA                     
## [2763,] "Prevotella_2"                    NA                     
## [2764,] NA                                NA                     
## [2765,] NA                                NA                     
## [2766,] "Alloprevotella"                  NA                     
## [2767,] "Streptococcus"                   NA                     
## [2768,] NA                                NA                     
## [2769,] NA                                NA                     
## [2770,] "Selenomonas"                     NA                     
## [2771,] NA                                NA                     
## [2772,] NA                                NA                     
## [2773,] "Prevotellaceae_YAB2003_group"    NA                     
## [2774,] "Selenomonas"                     NA                     
## [2775,] NA                                NA                     
## [2776,] "Corynebacterium"                 NA                     
## [2777,] "Eikenella"                       NA                     
## [2778,] "Corynebacterium"                 "argentoratense"       
## [2779,] "Tannerella"                      NA                     
## [2780,] "Bergeyella"                      NA                     
## [2781,] NA                                NA                     
## [2782,] "Capnocytophaga"                  NA                     
## [2783,] NA                                NA                     
## [2784,] NA                                NA                     
## [2785,] "Corynebacterium"                 NA                     
## [2786,] "Actinomyces"                     NA                     
## [2787,] "Prevotella_7"                    NA                     
## [2788,] "Veillonella"                     NA                     
## [2789,] NA                                NA                     
## [2790,] "Alloprevotella"                  NA                     
## [2791,] "Prevotella"                      NA                     
## [2792,] "Bergeyella"                      NA                     
## [2793,] NA                                NA                     
## [2794,] NA                                NA                     
## [2795,] "Neisseria"                       NA                     
## [2796,] "Prevotellaceae_YAB2003_group"    NA                     
## [2797,] "Prevotella_7"                    NA                     
## [2798,] "Capnocytophaga"                  NA                     
## [2799,] "Prevotella"                      NA                     
## [2800,] NA                                NA                     
## [2801,] "Actinobacillus"                  NA                     
## [2802,] "Capnocytophaga"                  NA                     
## [2803,] "Prevotella_7"                    NA                     
## [2804,] "Veillonella"                     NA                     
## [2805,] NA                                NA                     
## [2806,] "Prevotella_7"                    NA                     
## [2807,] "Alloprevotella"                  NA                     
## [2808,] NA                                NA                     
## [2809,] "Leptotrichia"                    NA                     
## [2810,] "Prevotella_7"                    NA                     
## [2811,] "Prevotella_7"                    NA                     
## [2812,] "Fusobacterium"                   NA                     
## [2813,] "Haemophilus"                     NA                     
## [2814,] NA                                NA                     
## [2815,] "Prevotella"                      NA                     
## [2816,] "Prevotella_7"                    NA                     
## [2817,] "Prevotella_7"                    NA                     
## [2818,] "Selenomonas_3"                   NA                     
## [2819,] "Stomatobaculum"                  NA                     
## [2820,] "Alloprevotella"                  NA                     
## [2821,] "Selenomonas_3"                   NA                     
## [2822,] "Prevotella"                      NA                     
## [2823,] "Alloprevotella"                  NA                     
## [2824,] "Alloprevotella"                  NA                     
## [2825,] "Bergeyella"                      NA                     
## [2826,] NA                                NA                     
## [2827,] "Megasphaera"                     NA                     
## [2828,] "Selenomonas"                     NA                     
## [2829,] NA                                NA                     
## [2830,] "Peptoanaerobacter"               NA                     
## [2831,] "Candidatus_Saccharimonas"        NA                     
## [2832,] "Aggregatibacter"                 NA                     
## [2833,] "Haemophilus"                     NA                     
## [2834,] "Treponema_2"                     NA                     
## [2835,] NA                                NA                     
## [2836,] "Alloprevotella"                  NA                     
## [2837,] "Dialister"                       NA                     
## [2838,] NA                                NA                     
## [2839,] "Prevotella_7"                    NA                     
## [2840,] "Parvimonas"                      NA                     
## [2841,] "Lachnoanaerobaculum"             NA                     
## [2842,] "Selenomonas_3"                   NA                     
## [2843,] "Lautropia"                       NA                     
## [2844,] "Selenomonas_3"                   NA                     
## [2845,] "Prevotellaceae_YAB2003_group"    NA                     
## [2846,] "Selenomonas"                     NA                     
## [2847,] "Aggregatibacter"                 NA                     
## [2848,] NA                                NA                     
## [2849,] "Selenomonas_3"                   NA                     
## [2850,] "Prevotella_7"                    NA                     
## [2851,] "Kingella"                        NA                     
## [2852,] "Fusobacterium"                   NA                     
## [2853,] "Selenomonas_3"                   NA                     
## [2854,] "Prevotella_7"                    NA                     
## [2855,] "Neisseria"                       "meningitidis"         
## [2856,] "Actinomyces"                     "oris"                 
## [2857,] "Simonsiella"                     NA                     
## [2858,] "Porphyromonas"                   NA                     
## [2859,] "Prevotella_2"                    NA                     
## [2860,] "Bergeriella"                     NA                     
## [2861,] "Selenomonas_3"                   NA                     
## [2862,] NA                                NA                     
## [2863,] "Bergeyella"                      NA                     
## [2864,] NA                                NA                     
## [2865,] "Tannerella"                      NA                     
## [2866,] NA                                NA                     
## [2867,] NA                                NA                     
## [2868,] NA                                NA                     
## [2869,] NA                                NA                     
## [2870,] "Lachnospiraceae_AC2044_group"    NA                     
## [2871,] "Prevotella"                      NA                     
## [2872,] NA                                NA                     
## [2873,] "Haemophilus"                     NA                     
## [2874,] "Prevotella_7"                    NA                     
## [2875,] "Capnocytophaga"                  NA                     
## [2876,] "Prevotella_7"                    "baroniae"             
## [2877,] NA                                NA                     
## [2878,] "Prevotella_7"                    NA                     
## [2879,] "Anthococcus"                     NA                     
## [2880,] NA                                NA                     
## [2881,] NA                                NA                     
## [2882,] "Cardiobacterium"                 NA                     
## [2883,] "Selenomonas"                     NA                     
## [2884,] NA                                NA                     
## [2885,] "Prevotella"                      NA                     
## [2886,] NA                                NA                     
## [2887,] "Stomatobaculum"                  NA                     
## [2888,] NA                                NA                     
## [2889,] "Capnocytophaga"                  NA                     
## [2890,] "Prevotella_7"                    NA                     
## [2891,] "Neisseria"                       NA                     
## [2892,] "Alloprevotella"                  NA                     
## [2893,] "Prevotella_7"                    NA                     
## [2894,] NA                                NA                     
## [2895,] NA                                NA                     
## [2896,] "Leptotrichia"                    NA                     
## [2897,] "Alloprevotella"                  NA                     
## [2898,] NA                                NA                     
## [2899,] NA                                NA                     
## [2900,] "Fusobacterium"                   NA                     
## [2901,] "Prevotella"                      NA                     
## [2902,] "Dialister"                       NA                     
## [2903,] "Actinomyces"                     NA                     
## [2904,] "Prevotella"                      NA                     
## [2905,] "Treponema_2"                     NA                     
## [2906,] "Haemophilus"                     NA                     
## [2907,] "Treponema_2"                     NA                     
## [2908,] "Campylobacter"                   NA                     
## [2909,] "Lachnoanaerobaculum"             NA                     
## [2910,] "Cobetia"                         NA                     
## [2911,] "Veillonella"                     NA                     
## [2912,] "Mycoplasma"                      "orale"                
## [2913,] "Alloprevotella"                  NA                     
## [2914,] NA                                NA                     
## [2915,] NA                                NA                     
## [2916,] "Capnocytophaga"                  "ochracea"             
## [2917,] "Candidatus_Saccharimonas"        NA                     
## [2918,] NA                                NA                     
## [2919,] "Fusobacterium"                   NA                     
## [2920,] "Veillonella"                     NA                     
## [2921,] "Porphyromonas"                   NA                     
## [2922,] "Actinomyces"                     NA                     
## [2923,] NA                                NA                     
## [2924,] "Anthococcus"                     NA                     
## [2925,] "Catonella"                       NA                     
## [2926,] "Leptotrichia"                    NA                     
## [2927,] NA                                NA                     
## [2928,] "Streptococcus"                   NA                     
## [2929,] "Arcanobacterium"                 "haemolyticum"         
## [2930,] NA                                NA                     
## [2931,] "Faecalibacterium"                NA                     
## [2932,] "Necropsobacter"                  NA                     
## [2933,] NA                                NA                     
## [2934,] "Atopobium"                       NA                     
## [2935,] "Actinomyces"                     "naeslundii"           
## [2936,] "Veillonella"                     NA                     
## [2937,] NA                                NA                     
## [2938,] NA                                NA                     
## [2939,] "Prevotella_7"                    NA                     
## [2940,] "Prevotella"                      NA                     
## [2941,] NA                                NA                     
## [2942,] NA                                NA                     
## [2943,] "Alloprevotella"                  NA                     
## [2944,] "Staphylococcus"                  NA                     
## [2945,] "Capnocytophaga"                  NA                     
## [2946,] NA                                NA                     
## [2947,] "Tannerella"                      NA                     
## [2948,] NA                                NA                     
## [2949,] "Catonella"                       NA                     
## [2950,] "Selenomonas"                     NA                     
## [2951,] NA                                NA                     
## [2952,] "Prevotella_7"                    NA                     
## [2953,] "Prevotella_7"                    NA                     
## [2954,] "Capnocytophaga"                  NA                     
## [2955,] "Oribacterium"                    NA                     
## [2956,] NA                                NA                     
## [2957,] "Haemophilus"                     "haemolyticus"         
## [2958,] "Leptotrichia"                    NA                     
## [2959,] "Veillonella"                     NA                     
## [2960,] "Veillonella"                     NA                     
## [2961,] "Alloprevotella"                  NA                     
## [2962,] "Aggregatibacter"                 NA                     
## [2963,] "Veillonella"                     NA                     
## [2964,] "Veillonella"                     NA                     
## [2965,] "Eikenella"                       NA                     
## [2966,] "Selenomonas_3"                   NA                     
## [2967,] "Veillonella"                     NA                     
## [2968,] NA                                NA                     
## [2969,] NA                                NA                     
## [2970,] "Leptotrichia"                    NA                     
## [2971,] "Neisseria"                       NA                     
## [2972,] NA                                NA                     
## [2973,] "Prevotella_7"                    NA                     
## [2974,] "Rikenellaceae_RC9_gut_group"     NA                     
## [2975,] "Gemella"                         NA                     
## [2976,] "Treponema_2"                     NA                     
## [2977,] "Lachnoanaerobaculum"             NA                     
## [2978,] "Mycoplasma"                      NA                     
## [2979,] "Veillonella"                     NA                     
## [2980,] "Prevotella"                      NA                     
## [2981,] "Citrobacter"                     NA                     
## [2982,] "Treponema_2"                     NA                     
## [2983,] NA                                NA                     
## [2984,] "Porphyromonas"                   NA                     
## [2985,] "Johnsonella"                     NA                     
## [2986,] NA                                NA                     
## [2987,] "F0332"                           NA                     
## [2988,] "Prevotella_2"                    NA                     
## [2989,] "Capnocytophaga"                  NA                     
## [2990,] NA                                NA                     
## [2991,] "Dialister"                       NA                     
## [2992,] "Fretibacterium"                  NA                     
## [2993,] "Streptococcus"                   NA                     
## [2994,] "Alloscardovia"                   NA                     
## [2995,] "Prevotella_6"                    NA                     
## [2996,] "Catonella"                       NA                     
## [2997,] "Dialister"                       NA                     
## [2998,] NA                                NA                     
## [2999,] "Rahnella"                        "aquatilis"            
## [3000,] "Selenomonas_3"                   NA                     
## [3001,] "Capnocytophaga"                  NA                     
## [3002,] "Leptotrichia"                    NA                     
## [3003,] NA                                NA                     
## [3004,] "Prevotella_7"                    NA                     
## [3005,] NA                                NA                     
## [3006,] "Cricetibacter"                   NA                     
## [3007,] "Veillonella"                     NA                     
## [3008,] "Lawsonella"                      NA                     
## [3009,] "Johnsonella"                     NA                     
## [3010,] "Selenomonas"                     NA                     
## [3011,] NA                                NA                     
## [3012,] "Selenomonas_3"                   NA                     
## [3013,] NA                                NA                     
## [3014,] "Prevotella"                      NA                     
## [3015,] "Treponema_2"                     NA                     
## [3016,] NA                                NA                     
## [3017,] NA                                NA                     
## [3018,] NA                                NA                     
## [3019,] "Porphyromonas"                   NA                     
## [3020,] NA                                NA                     
## [3021,] "Alloprevotella"                  NA                     
## [3022,] "Peptoanaerobacter"               NA                     
## [3023,] "Prevotella_7"                    NA                     
## [3024,] "Prevotella"                      NA                     
## [3025,] "Prevotella_7"                    NA                     
## [3026,] NA                                NA                     
## [3027,] "Prevotella_7"                    NA                     
## [3028,] NA                                NA                     
## [3029,] "Fusobacterium"                   NA                     
## [3030,] "Capnocytophaga"                  NA                     
## [3031,] "Prevotella"                      NA                     
## [3032,] NA                                NA                     
## [3033,] "Prevotella"                      NA                     
## [3034,] NA                                NA                     
## [3035,] NA                                NA                     
## [3036,] "Stomatobaculum"                  NA                     
## [3037,] "Lachnoanaerobaculum"             NA                     
## [3038,] "Alloprevotella"                  NA                     
## [3039,] "Actinomyces"                     NA                     
## [3040,] "Anthococcus"                     NA                     
## [3041,] NA                                NA                     
## [3042,] "Alloprevotella"                  NA                     
## [3043,] "Selenomonas_3"                   NA                     
## [3044,] "Selenomonas"                     NA                     
## [3045,] "Capnocytophaga"                  NA                     
## [3046,] NA                                NA                     
## [3047,] "Actinomyces"                     "naeslundii"           
## [3048,] "Cricetibacter"                   NA                     
## [3049,] NA                                NA                     
## [3050,] NA                                NA                     
## [3051,] NA                                NA                     
## [3052,] "Moraxella"                       NA                     
## [3053,] NA                                NA                     
## [3054,] NA                                NA                     
## [3055,] "Filifactor"                      NA                     
## [3056,] "Porphyromonas"                   NA                     
## [3057,] "Prevotella"                      NA                     
## [3058,] "Oceanivirga"                     NA                     
## [3059,] "Prevotella_7"                    NA                     
## [3060,] "Capnocytophaga"                  NA                     
## [3061,] "Porphyromonas"                   NA                     
## [3062,] "Leptotrichia"                    "buccalis"             
## [3063,] "Veillonella"                     NA                     
## [3064,] "Lachnoanaerobaculum"             NA                     
## [3065,] "Alloscardovia"                   NA                     
## [3066,] "Lachnoanaerobaculum"             NA                     
## [3067,] "Selenomonas_3"                   NA                     
## [3068,] "Selenomonas_3"                   NA                     
## [3069,] "Candidatus_Saccharimonas"        NA                     
## [3070,] NA                                NA                     
## [3071,] NA                                NA                     
## [3072,] "Selenomonas_3"                   NA                     
## [3073,] "Capnocytophaga"                  NA                     
## [3074,] "Streptobacillus"                 NA                     
## [3075,] "Alloprevotella"                  NA                     
## [3076,] "Lautropia"                       NA                     
## [3077,] NA                                NA                     
## [3078,] "Leptotrichia"                    NA                     
## [3079,] "Simonsiella"                     NA                     
## [3080,] "Porphyromonas"                   NA                     
## [3081,] "Treponema_2"                     NA                     
## [3082,] "Actinomyces"                     NA                     
## [3083,] "Rothia"                          NA                     
## [3084,] "Prevotella_7"                    NA                     
## [3085,] "Fusobacterium"                   NA                     
## [3086,] NA                                NA                     
## [3087,] "Prevotellaceae_YAB2003_group"    NA                     
## [3088,] NA                                NA                     
## [3089,] NA                                NA                     
## [3090,] "Selenomonas_3"                   NA                     
## [3091,] "Selenomonas_3"                   NA                     
## [3092,] "Corynebacterium"                 NA                     
## [3093,] NA                                NA                     
## [3094,] "Actinomyces"                     NA                     
## [3095,] NA                                NA                     
## [3096,] NA                                NA                     
## [3097,] "Centipeda"                       NA                     
## [3098,] "Veillonella"                     NA                     
## [3099,] NA                                NA                     
## [3100,] "Prevotella_7"                    NA                     
## [3101,] "Prevotella"                      NA                     
## [3102,] "Treponema_2"                     NA                     
## [3103,] NA                                NA                     
## [3104,] NA                                NA                     
## [3105,] NA                                NA                     
## [3106,] "Actinomyces"                     NA                     
## [3107,] "Prevotella"                      NA                     
## [3108,] "Prevotella_7"                    NA                     
## [3109,] "Haemophilus"                     "parainfluenzae"       
## [3110,] NA                                NA                     
## [3111,] NA                                NA                     
## [3112,] NA                                NA                     
## [3113,] NA                                NA                     
## [3114,] "Lachnoanaerobaculum"             NA                     
## [3115,] NA                                NA                     
## [3116,] NA                                NA                     
## [3117,] "Anthococcus"                     NA                     
## [3118,] "Bergeyella"                      NA                     
## [3119,] NA                                NA                     
## [3120,] "Prevotella_2"                    NA                     
## [3121,] "Alloprevotella"                  NA                     
## [3122,] NA                                NA                     
## [3123,] "Actinomyces"                     NA                     
## [3124,] "Prevotellaceae_YAB2003_group"    NA                     
## [3125,] "Prevotella_7"                    NA                     
## [3126,] "Defluviitaleaceae_UCG-011"       NA                     
## [3127,] NA                                NA                     
## [3128,] "Prevotella_2"                    NA                     
## [3129,] NA                                NA                     
## [3130,] "Actinomyces"                     "odontolyticus"        
## [3131,] "Selenomonas_3"                   NA                     
## [3132,] "Prevotella_7"                    NA                     
## [3133,] "Capnocytophaga"                  NA                     
## [3134,] NA                                NA                     
## [3135,] "Prevotella_7"                    NA                     
## [3136,] NA                                NA                     
## [3137,] NA                                NA                     
## [3138,] "Capnocytophaga"                  NA                     
## [3139,] "Capnocytophaga"                  NA                     
## [3140,] "Porphyromonas"                   NA                     
## [3141,] "Prevotella_7"                    NA                     
## [3142,] "Prevotella"                      NA                     
## [3143,] "Prevotella"                      NA                     
## [3144,] "Aggregatibacter"                 NA                     
## [3145,] "Candidatus_Saccharimonas"        NA                     
## [3146,] "Leptotrichia"                    NA                     
## [3147,] "Leptotrichia"                    NA                     
## [3148,] NA                                NA                     
## [3149,] NA                                NA                     
## [3150,] "Treponema_2"                     NA                     
## [3151,] "Ruminococcaceae_UCG-014"         NA                     
## [3152,] NA                                NA                     
## [3153,] "Treponema_2"                     NA                     
## [3154,] "Prevotella"                      NA                     
## [3155,] "Prevotella"                      NA                     
## [3156,] "Prevotella_7"                    NA                     
## [3157,] "Veillonella"                     NA                     
## [3158,] "Johnsonella"                     NA                     
## [3159,] "Neisseria"                       NA                     
## [3160,] "Tannerella"                      NA                     
## [3161,] NA                                NA                     
## [3162,] "Cardiobacterium"                 "valvarum"             
## [3163,] "Porphyromonas"                   NA                     
## [3164,] "Prevotella_7"                    NA                     
## [3165,] "Stomatobaculum"                  NA                     
## [3166,] "Aggregatibacter"                 NA                     
## [3167,] "Alloprevotella"                  NA                     
## [3168,] "Treponema_2"                     NA                     
## [3169,] "Treponema_2"                     "maltophilum"          
## [3170,] "Leptotrichia"                    NA                     
## [3171,] "Campylobacter"                   NA                     
## [3172,] NA                                NA                     
## [3173,] "Actinomyces"                     NA                     
## [3174,] NA                                NA                     
## [3175,] NA                                NA                     
## [3176,] NA                                NA                     
## [3177,] "Alloprevotella"                  NA                     
## [3178,] NA                                NA                     
## [3179,] NA                                NA                     
## [3180,] NA                                NA                     
## [3181,] "Capnocytophaga"                  "ochracea"             
## [3182,] "Treponema_2"                     NA                     
## [3183,] "Leptotrichia"                    NA                     
## [3184,] "Prevotella_2"                    NA                     
## [3185,] "Fusobacterium"                   NA                     
## [3186,] "Blvii28_wastewater-sludge_group" NA                     
## [3187,] "Neisseria"                       NA                     
## [3188,] NA                                NA                     
## [3189,] "Capnocytophaga"                  NA                     
## [3190,] "Prevotella_7"                    NA                     
## [3191,] "Treponema_2"                     NA                     
## [3192,] "Cardiobacterium"                 NA                     
## [3193,] "Prevotella_7"                    NA                     
## [3194,] NA                                NA                     
## [3195,] NA                                NA                     
## [3196,] NA                                NA                     
## [3197,] "Prevotella_7"                    NA                     
## [3198,] "Prevotella_7"                    NA                     
## [3199,] "Prevotella"                      NA                     
## [3200,] NA                                NA                     
## [3201,] NA                                NA                     
## [3202,] "Anaerococcus"                    NA                     
## [3203,] NA                                NA                     
## [3204,] NA                                NA                     
## [3205,] NA                                NA                     
## [3206,] "Prevotella"                      NA                     
## [3207,] "Capnocytophaga"                  NA                     
## [3208,] "Prevotellaceae_YAB2003_group"    NA                     
## [3209,] NA                                NA                     
## [3210,] "Alloprevotella"                  NA                     
## [3211,] "Corynebacterium"                 NA                     
## [3212,] "Prevotella"                      NA                     
## [3213,] "Butyrivibrio_2"                  NA                     
## [3214,] NA                                NA                     
## [3215,] "Prevotella_7"                    NA                     
## [3216,] "Prevotella_7"                    NA                     
## [3217,] "Leptotrichia"                    NA                     
## [3218,] "Prevotellaceae_YAB2003_group"    NA                     
## [3219,] "Bacteroides"                     NA                     
## [3220,] "Anaeroglobus"                    NA                     
## [3221,] NA                                NA                     
## [3222,] "Johnsonella"                     NA                     
## [3223,] "Granulicatella"                  NA                     
## [3224,] NA                                NA                     
## [3225,] "Faecalibacterium"                NA                     
## [3226,] "Actinomyces"                     NA                     
## [3227,] NA                                NA                     
## [3228,] NA                                NA                     
## [3229,] "Selenomonas_3"                   NA                     
## [3230,] "Leptotrichia"                    NA                     
## [3231,] NA                                NA                     
## [3232,] "Prevotella_7"                    NA                     
## [3233,] "Leptotrichia"                    NA                     
## [3234,] "Necropsobacter"                  NA                     
## [3235,] "Prevotella_2"                    NA                     
## [3236,] NA                                NA                     
## [3237,] NA                                NA                     
## [3238,] "Selenomonas"                     NA                     
## [3239,] NA                                NA                     
## [3240,] "Prevotella"                      NA                     
## [3241,] "Prevotella_7"                    NA                     
## [3242,] "Selenomonas"                     NA                     
## [3243,] NA                                NA                     
## [3244,] "Lachnoanaerobaculum"             NA                     
## [3245,] "Rothia"                          NA                     
## [3246,] "Shuttleworthia"                  "satelles"             
## [3247,] NA                                NA                     
## [3248,] "Collinsella"                     "aerofaciens"          
## [3249,] "Prevotella_7"                    NA                     
## [3250,] NA                                NA                     
## [3251,] "Atopobium"                       NA                     
## [3252,] NA                                NA                     
## [3253,] "Prevotella"                      NA                     
## [3254,] NA                                NA                     
## [3255,] "Lautropia"                       NA                     
## [3256,] "Porphyromonas"                   NA                     
## [3257,] NA                                NA                     
## [3258,] "Leptotrichia"                    NA                     
## [3259,] "Anthococcus"                     NA                     
## [3260,] "Campylobacter"                   NA                     
## [3261,] "Campylobacter"                   "ureolyticus"          
## [3262,] NA                                NA                     
## [3263,] "Candidatus_Saccharimonas"        NA                     
## [3264,] NA                                NA                     
## [3265,] NA                                NA                     
## [3266,] NA                                NA                     
## [3267,] "Alloprevotella"                  NA                     
## [3268,] NA                                NA                     
## [3269,] "Alloprevotella"                  NA                     
## [3270,] NA                                NA                     
## [3271,] "Alloprevotella"                  NA                     
## [3272,] "Alloprevotella"                  NA                     
## [3273,] "Prevotella"                      NA                     
## [3274,] NA                                NA                     
## [3275,] "Prevotella"                      NA                     
## [3276,] "Actinomyces"                     NA                     
## [3277,] NA                                NA                     
## [3278,] NA                                NA                     
## [3279,] "Treponema_2"                     NA                     
## [3280,] "Prevotella_7"                    "buccae"               
## [3281,] "Selenomonas_3"                   NA                     
## [3282,] "Cardiobacterium"                 NA                     
## [3283,] "Alloprevotella"                  NA                     
## [3284,] "Prevotella_2"                    NA                     
## [3285,] NA                                NA                     
## [3286,] "Necropsobacter"                  NA                     
## [3287,] "Cricetibacter"                   NA                     
## [3288,] "Treponema_2"                     "socranskii"           
## [3289,] "Lachnoanaerobaculum"             NA                     
## [3290,] "Treponema_2"                     NA                     
## [3291,] "Prevotella_6"                    NA                     
## [3292,] "Necropsobacter"                  NA                     
## [3293,] NA                                NA                     
## [3294,] "Prevotella_7"                    NA                     
## [3295,] "Prevotella"                      NA                     
## [3296,] "Prevotella"                      NA                     
## [3297,] "Actinomyces"                     "odontolyticus"        
## [3298,] "Prevotella"                      NA                     
## [3299,] NA                                NA                     
## [3300,] "Prevotella_7"                    NA                     
## [3301,] "Alloprevotella"                  NA                     
## [3302,] "Bergeyella"                      NA                     
## [3303,] "Tannerella"                      NA                     
## [3304,] "Kingella"                        NA                     
## [3305,] "Kingella"                        NA                     
## [3306,] "Prevotella_2"                    NA                     
## [3307,] "Prevotella_7"                    NA                     
## [3308,] NA                                NA                     
## [3309,] NA                                NA                     
## [3310,] "Prevotellaceae_YAB2003_group"    NA                     
## [3311,] "Prevotella"                      NA                     
## [3312,] "Leptotrichia"                    NA                     
## [3313,] "Capnocytophaga"                  NA                     
## [3314,] "Treponema_2"                     "vincentii"            
## [3315,] NA                                NA                     
## [3316,] NA                                NA                     
## [3317,] "Catonella"                       NA                     
## [3318,] "Leptotrichia"                    NA                     
## [3319,] "Bergeriella"                     NA                     
## [3320,] "Alloprevotella"                  NA                     
## [3321,] "Prevotella"                      NA                     
## [3322,] "Leptotrichia"                    NA                     
## [3323,] "Prevotella_7"                    NA                     
## [3324,] "Prevotella"                      NA                     
## [3325,] "Prevotella"                      NA                     
## [3326,] NA                                NA                     
## [3327,] "Corynebacterium"                 NA                     
## [3328,] "Bergeyella"                      NA                     
## [3329,] "Prevotellaceae_YAB2003_group"    NA                     
## [3330,] "Treponema_2"                     NA                     
## [3331,] "Porphyromonas"                   NA                     
## [3332,] "Prevotella"                      NA                     
## [3333,] NA                                NA                     
## [3334,] NA                                NA                     
## [3335,] "Gemella"                         NA                     
## [3336,] "Cardiobacterium"                 NA                     
## [3337,] "Prevotella_7"                    NA                     
## [3338,] "Leptotrichia"                    NA                     
## [3339,] "Candidatus_Saccharimonas"        NA                     
## [3340,] NA                                NA                     
## [3341,] NA                                NA                     
## [3342,] NA                                NA                     
## [3343,] NA                                NA                     
## [3344,] NA                                NA                     
## [3345,] NA                                NA                     
## [3346,] NA                                NA                     
## [3347,] NA                                NA                     
## [3348,] "Bergeriella"                     NA                     
## [3349,] NA                                NA                     
## [3350,] "Leptotrichia"                    NA                     
## [3351,] NA                                NA                     
## [3352,] "Bergeriella"                     NA                     
## [3353,] "Prevotella"                      NA                     
## [3354,] NA                                NA                     
## [3355,] NA                                NA                     
## [3356,] "Veillonella"                     NA                     
## [3357,] "Atopobium"                       "rimae"                
## [3358,] NA                                NA                     
## [3359,] "Treponema_2"                     NA                     
## [3360,] "Veillonella"                     NA                     
## [3361,] NA                                NA                     
## [3362,] "Prevotella_2"                    NA                     
## [3363,] NA                                NA                     
## [3364,] "Peptoniphilus"                   NA                     
## [3365,] "Prevotella"                      NA                     
## [3366,] "Prevotella_7"                    NA                     
## [3367,] "Necropsobacter"                  NA                     
## [3368,] "Actinomyces"                     NA                     
## [3369,] "Capnocytophaga"                  NA                     
## [3370,] "Veillonella"                     NA                     
## [3371,] "Capnocytophaga"                  NA                     
## [3372,] "Fusobacterium"                   NA                     
## [3373,] "Leptotrichia"                    NA                     
## [3374,] "Treponema_2"                     NA                     
## [3375,] "Prevotella"                      NA                     
## [3376,] "Capnocytophaga"                  NA                     
## [3377,] "Prevotella_7"                    NA                     
## [3378,] "Leptotrichia"                    NA                     
## [3379,] "Rikenellaceae_RC9_gut_group"     NA                     
## [3380,] "Capnocytophaga"                  NA                     
## [3381,] "Capnocytophaga"                  NA                     
## [3382,] "Prevotella_7"                    NA                     
## [3383,] "Lachnoanaerobaculum"             NA                     
## [3384,] "Prevotella_7"                    NA                     
## [3385,] "Treponema_2"                     NA                     
## [3386,] "Porphyromonas"                   NA                     
## [3387,] "Streptococcus"                   NA                     
## [3388,] "Peptococcus"                     NA                     
## [3389,] "Prevotella_7"                    NA                     
## [3390,] "Prevotella_2"                    NA                     
## [3391,] "Granulicatella"                  NA                     
## [3392,] "Capnocytophaga"                  NA                     
## [3393,] NA                                NA                     
## [3394,] NA                                NA                     
## [3395,] "Leptotrichia"                    NA                     
## [3396,] "Rothia"                          NA                     
## [3397,] "Porphyromonas"                   NA                     
## [3398,] NA                                NA                     
## [3399,] NA                                NA                     
## [3400,] "Porphyromonas"                   NA                     
## [3401,] "Enterobacter"                    NA                     
## [3402,] NA                                NA                     
## [3403,] "Parvimonas"                      NA                     
## [3404,] "Capnocytophaga"                  NA                     
## [3405,] "Prevotella_7"                    NA                     
## [3406,] "Alloprevotella"                  NA                     
## [3407,] "Prevotella"                      NA                     
## [3408,] "Abiotrophia"                     "defectiva"            
## [3409,] "Capnocytophaga"                  NA                     
## [3410,] "Cricetibacter"                   NA                     
## [3411,] "Atopobium"                       NA                     
## [3412,] "Alloprevotella"                  NA                     
## [3413,] "Treponema_2"                     NA                     
## [3414,] "Prevotella"                      NA                     
## [3415,] "Prevotella"                      NA                     
## [3416,] "Prevotella_7"                    NA                     
## [3417,] "Leptotrichia"                    NA                     
## [3418,] "Porphyromonas"                   NA                     
## [3419,] "Veillonella"                     NA                     
## [3420,] NA                                NA                     
## [3421,] NA                                NA                     
## [3422,] "Oribacterium"                    NA                     
## [3423,] NA                                NA                     
## [3424,] "Prevotella"                      NA                     
## [3425,] "Necropsobacter"                  NA                     
## [3426,] "Prevotellaceae_YAB2003_group"    NA                     
## [3427,] "Alloprevotella"                  NA                     
## [3428,] "Capnocytophaga"                  NA                     
## [3429,] NA                                NA                     
## [3430,] NA                                NA                     
## [3431,] NA                                NA                     
## [3432,] "Prevotella"                      NA                     
## [3433,] "Soonwooa"                        NA                     
## [3434,] "Prevotellaceae_YAB2003_group"    NA                     
## [3435,] NA                                NA                     
## [3436,] "Pantoea"                         NA                     
## [3437,] NA                                NA                     
## [3438,] NA                                NA                     
## [3439,] "Pseudomonas"                     NA                     
## [3440,] "Abiotrophia"                     NA                     
## [3441,] "Capnocytophaga"                  NA                     
## [3442,] NA                                NA                     
## [3443,] "Peptostreptococcus"              NA                     
## [3444,] NA                                NA                     
## [3445,] "Prevotella"                      NA                     
## [3446,] "Stomatobaculum"                  NA                     
## [3447,] NA                                NA                     
## [3448,] "Prevotella"                      NA                     
## [3449,] "Prevotella_2"                    NA                     
## [3450,] NA                                NA                     
## [3451,] "Prevotella_7"                    "multiformis"          
## [3452,] "Leptotrichia"                    NA                     
## [3453,] "Prevotella_7"                    NA                     
## [3454,] "Alloprevotella"                  NA                     
## [3455,] "Selenomonas_3"                   NA                     
## [3456,] "Porphyromonas"                   NA                     
## [3457,] NA                                NA                     
## [3458,] NA                                NA                     
## [3459,] "Aggregatibacter"                 NA                     
## [3460,] NA                                NA                     
## [3461,] "Selenomonas_4"                   NA                     
## [3462,] "Simonsiella"                     NA                     
## [3463,] "Johnsonella"                     NA                     
## [3464,] "Prevotella_2"                    NA                     
## [3465,] "Porphyromonas"                   NA                     
## [3466,] NA                                NA                     
## [3467,] "Centipeda"                       NA                     
## [3468,] NA                                NA                     
## [3469,] "Prevotella"                      NA                     
## [3470,] "Alloprevotella"                  NA                     
## [3471,] NA                                NA                     
## [3472,] NA                                NA                     
## [3473,] NA                                NA                     
## [3474,] "Actinomyces"                     NA                     
## [3475,] NA                                NA                     
## [3476,] "Prevotellaceae_YAB2003_group"    NA                     
## [3477,] "Porphyromonas"                   NA                     
## [3478,] NA                                NA                     
## [3479,] NA                                NA                     
## [3480,] "Selenomonas_3"                   NA                     
## [3481,] "Necropsobacter"                  NA                     
## [3482,] "Capnocytophaga"                  NA                     
## [3483,] NA                                NA                     
## [3484,] "Howardella"                      NA                     
## [3485,] "Prevotella_7"                    NA                     
## [3486,] "Alloprevotella"                  NA                     
## [3487,] "Capnocytophaga"                  NA                     
## [3488,] "Selenomonas_3"                   NA                     
## [3489,] NA                                NA                     
## [3490,] NA                                NA                     
## [3491,] "Prevotella_7"                    NA                     
## [3492,] "Selenomonas_3"                   NA                     
## [3493,] "Selenomonas_4"                   NA                     
## [3494,] NA                                NA                     
## [3495,] NA                                NA                     
## [3496,] NA                                NA                     
## [3497,] "Prevotella_7"                    NA                     
## [3498,] NA                                NA                     
## [3499,] "Anthococcus"                     NA                     
## [3500,] "Prevotella_6"                    NA                     
## [3501,] NA                                NA                     
## [3502,] NA                                NA                     
## [3503,] "Aggregatibacter"                 NA                     
## [3504,] NA                                NA                     
## [3505,] "F0058"                           NA                     
## [3506,] "Alloprevotella"                  NA                     
## [3507,] NA                                NA                     
## [3508,] NA                                NA                     
## [3509,] "Leptotrichia"                    NA                     
## [3510,] NA                                NA                     
## [3511,] "Capnocytophaga"                  NA                     
## [3512,] "Actinobacillus"                  NA                     
## [3513,] "Peptococcus"                     NA                     
## [3514,] "Prevotella_6"                    NA                     
## [3515,] NA                                NA                     
## [3516,] "Selenomonas_3"                   NA                     
## [3517,] NA                                NA                     
## [3518,] NA                                NA                     
## [3519,] "Selenomonas_3"                   NA                     
## [3520,] "Pseudomonas"                     NA                     
## [3521,] "Necropsobacter"                  NA                     
## [3522,] NA                                NA                     
## [3523,] NA                                NA                     
## [3524,] NA                                NA                     
## [3525,] "Prevotella"                      NA                     
## [3526,] NA                                NA                     
## [3527,] NA                                NA                     
## [3528,] "Neisseria"                       NA                     
## [3529,] "Porphyromonas"                   NA                     
## [3530,] "Anthococcus"                     NA                     
## [3531,] "Prevotella"                      NA                     
## [3532,] "Prevotella"                      NA                     
## [3533,] NA                                NA                     
## [3534,] NA                                NA                     
## [3535,] NA                                NA                     
## [3536,] NA                                NA                     
## [3537,] NA                                NA                     
## [3538,] "Selenomonas"                     NA                     
## [3539,] "Leptotrichia"                    NA                     
## [3540,] NA                                NA                     
## [3541,] "Prevotella_7"                    NA                     
## [3542,] "Streptobacillus"                 NA                     
## [3543,] "Leptotrichia"                    NA                     
## [3544,] "Neisseria"                       NA                     
## [3545,] "Leptotrichia"                    NA                     
## [3546,] "Parvimonas"                      NA                     
## [3547,] NA                                NA                     
## [3548,] "Tannerella"                      NA                     
## [3549,] NA                                NA                     
## [3550,] NA                                NA                     
## [3551,] NA                                NA                     
## [3552,] NA                                NA                     
## [3553,] "Neisseria"                       NA                     
## [3554,] "Cricetibacter"                   NA                     
## [3555,] NA                                NA                     
## [3556,] NA                                NA                     
## [3557,] "Anthococcus"                     NA                     
## [3558,] NA                                NA                     
## [3559,] NA                                NA                     
## [3560,] "Lactobacillus"                   NA                     
## [3561,] "Treponema_2"                     NA                     
## [3562,] "Johnsonella"                     NA                     
## [3563,] NA                                NA                     
## [3564,] "Peptostreptococcus"              NA                     
## [3565,] "Prevotella"                      NA                     
## [3566,] "Selenomonas_3"                   NA                     
## [3567,] "Neisseria"                       NA                     
## [3568,] NA                                NA                     
## [3569,] "Veillonella"                     NA                     
## [3570,] "Alloprevotella"                  NA                     
## [3571,] "Capnocytophaga"                  "ochracea"             
## [3572,] NA                                NA                     
## [3573,] NA                                NA                     
## [3574,] NA                                NA                     
## [3575,] NA                                NA                     
## [3576,] "Oribacterium"                    NA                     
## [3577,] NA                                NA                     
## [3578,] "Lachnoanaerobaculum"             NA                     
## [3579,] NA                                NA                     
## [3580,] "Howardella"                      NA                     
## [3581,] "Prevotella_7"                    NA                     
## [3582,] "Treponema_2"                     NA                     
## [3583,] "Peptococcus"                     NA                     
## [3584,] NA                                NA                     
## [3585,] "Prevotella_7"                    NA                     
## [3586,] "Bergeriella"                     NA                     
## [3587,] "Prevotellaceae_YAB2003_group"    NA                     
## [3588,] "Prevotella_2"                    NA                     
## [3589,] NA                                NA                     
## [3590,] NA                                NA                     
## [3591,] NA                                NA                     
## [3592,] "Alloprevotella"                  NA                     
## [3593,] NA                                NA                     
## [3594,] "Porphyromonas"                   NA                     
## [3595,] NA                                NA                     
## [3596,] NA                                NA                     
## [3597,] "Alloprevotella"                  NA                     
## [3598,] "Capnocytophaga"                  NA                     
## [3599,] NA                                NA                     
## [3600,] NA                                NA                     
## [3601,] "Prevotella"                      NA                     
## [3602,] NA                                NA                     
## [3603,] NA                                NA                     
## [3604,] "Bergeriella"                     NA                     
## [3605,] NA                                NA                     
## [3606,] "Bergeyella"                      NA                     
## [3607,] NA                                NA                     
## [3608,] "Comamonas"                       NA                     
## [3609,] "Leptotrichia"                    NA                     
## [3610,] "Corynebacterium"                 NA                     
## [3611,] "Porphyromonas"                   NA                     
## [3612,] "Johnsonella"                     NA                     
## [3613,] "Leptotrichia"                    NA                     
## [3614,] "Dialister"                       NA                     
## [3615,] NA                                NA                     
## [3616,] NA                                NA                     
## [3617,] "Leptotrichia"                    NA                     
## [3618,] "Peptostreptococcus"              NA                     
## [3619,] "Kingella"                        NA                     
## [3620,] "Atopobium"                       NA                     
## [3621,] "Rothia"                          NA                     
## [3622,] "Tannerella"                      "forsythia"            
## [3623,] "Kingella"                        NA                     
## [3624,] "Prevotella"                      NA                     
## [3625,] "Rikenellaceae_RC9_gut_group"     NA                     
## [3626,] "Prevotella_7"                    NA                     
## [3627,] NA                                NA                     
## [3628,] NA                                NA                     
## [3629,] "Tannerella"                      NA                     
## [3630,] NA                                NA                     
## [3631,] "Porphyromonas"                   NA                     
## [3632,] "Prevotella"                      NA                     
## [3633,] "Bergeriella"                     NA                     
## [3634,] NA                                NA                     
## [3635,] NA                                NA                     
## [3636,] NA                                NA                     
## [3637,] "Bergeyella"                      NA                     
## [3638,] "Ruminococcaceae_UCG-014"         NA                     
## [3639,] "Veillonella"                     NA                     
## [3640,] "Veillonella"                     NA                     
## [3641,] "Veillonella"                     NA                     
## [3642,] NA                                NA                     
## [3643,] "Cardiobacterium"                 NA                     
## [3644,] NA                                NA                     
## [3645,] NA                                NA                     
## [3646,] NA                                NA                     
## [3647,] NA                                NA                     
## [3648,] NA                                NA                     
## [3649,] "Shuttleworthia"                  NA                     
## [3650,] "Treponema_2"                     NA                     
## [3651,] "Ruminococcaceae_UCG-014"         NA                     
## [3652,] "Capnocytophaga"                  NA                     
## [3653,] NA                                NA                     
## [3654,] "Capnocytophaga"                  NA                     
## [3655,] "Alloprevotella"                  NA                     
## [3656,] "Necropsobacter"                  NA                     
## [3657,] NA                                NA                     
## [3658,] NA                                NA                     
## [3659,] NA                                NA                     
## [3660,] NA                                NA                     
## [3661,] "Peptostreptococcus"              NA                     
## [3662,] NA                                NA                     
## [3663,] "Prevotella_7"                    NA                     
## [3664,] "Prevotella"                      NA                     
## [3665,] "Prevotella_7"                    NA                     
## [3666,] "Veillonella"                     NA                     
## [3667,] "Rodentibacter"                   NA                     
## [3668,] NA                                NA                     
## [3669,] NA                                NA                     
## [3670,] "Atopobium"                       NA                     
## [3671,] NA                                NA                     
## [3672,] NA                                NA                     
## [3673,] "Capnocytophaga"                  NA                     
## [3674,] "Prevotella"                      NA                     
## [3675,] NA                                NA                     
## [3676,] "Prevotella_7"                    NA                     
## [3677,] "Treponema_2"                     "maltophilum"          
## [3678,] "Prevotella"                      NA                     
## [3679,] "Bergeyella"                      NA                     
## [3680,] "Prevotellaceae_YAB2003_group"    NA                     
## [3681,] "Selenomonas_3"                   NA                     
## [3682,] "Alloprevotella"                  NA                     
## [3683,] "Kingella"                        NA                     
## [3684,] "Haemophilus"                     NA                     
## [3685,] "Abiotrophia"                     NA                     
## [3686,] NA                                NA                     
## [3687,] NA                                NA                     
## [3688,] "Prevotella_2"                    NA                     
## [3689,] "Prevotella_7"                    "buccae"               
## [3690,] "Agathobacter"                    NA                     
## [3691,] NA                                NA                     
## [3692,] NA                                NA                     
## [3693,] NA                                NA                     
## [3694,] "Treponema_2"                     NA                     
## [3695,] "Butyrivibrio_2"                  NA                     
## [3696,] NA                                NA                     
## [3697,] "Faecalibacterium"                NA                     
## [3698,] "Cardiobacterium"                 "valvarum"             
## [3699,] "Leptotrichia"                    NA                     
## [3700,] "Prevotella_7"                    NA                     
## [3701,] "Selenomonas_3"                   NA                     
## [3702,] "Campylobacter"                   NA                     
## [3703,] NA                                NA                     
## [3704,] "Leptotrichia"                    NA                     
## [3705,] "Kingella"                        NA                     
## [3706,] NA                                NA                     
## [3707,] "Prevotella"                      NA                     
## [3708,] "Kingella"                        NA                     
## [3709,] NA                                NA                     
## [3710,] "Prevotella_7"                    NA                     
## [3711,] "Lachnoanaerobaculum"             NA                     
## [3712,] "Atopobium"                       NA                     
## [3713,] NA                                NA                     
## [3714,] NA                                NA                     
## [3715,] NA                                NA                     
## [3716,] "Comamonas"                       NA                     
## [3717,] "Abiotrophia"                     NA                     
## [3718,] "Fusobacterium"                   NA                     
## [3719,] NA                                NA                     
## [3720,] NA                                NA                     
## [3721,] "Selenomonas_3"                   NA                     
## [3722,] NA                                NA                     
## [3723,] "Aggregatibacter"                 "actinomycetemcomitans"
## [3724,] "Lachnoanaerobaculum"             NA                     
## [3725,] "Shuttleworthia"                  NA                     
## [3726,] NA                                NA                     
## [3727,] NA                                NA                     
## [3728,] "Filifactor"                      NA                     
## [3729,] "Lachnoanaerobaculum"             NA                     
## [3730,] NA                                NA                     
## [3731,] "Lactobacillus"                   "sanfranciscensis"     
## [3732,] "Treponema_2"                     NA                     
## [3733,] NA                                NA                     
## [3734,] NA                                NA                     
## [3735,] NA                                NA                     
## [3736,] "Gemella"                         NA                     
## [3737,] "Prevotella"                      NA                     
## [3738,] NA                                NA                     
## [3739,] "Porphyromonas"                   NA                     
## [3740,] "Prevotellaceae_YAB2003_group"    NA                     
## [3741,] "Prevotella_6"                    NA                     
## [3742,] "Simonsiella"                     NA                     
## [3743,] "Campylobacter"                   NA                     
## [3744,] NA                                NA                     
## [3745,] NA                                NA                     
## [3746,] "Capnocytophaga"                  NA                     
## [3747,] "Prevotella_7"                    NA                     
## [3748,] "Prevotella_7"                    NA                     
## [3749,] "Rikenellaceae_RC9_gut_group"     NA                     
## [3750,] NA                                NA                     
## [3751,] "Stomatobaculum"                  NA                     
## [3752,] NA                                NA                     
## [3753,] NA                                NA                     
## [3754,] "Prevotella_7"                    NA                     
## [3755,] "Stomatobaculum"                  NA                     
## [3756,] "Lachnoanaerobaculum"             NA                     
## [3757,] "Prevotellaceae_YAB2003_group"    NA                     
## [3758,] NA                                NA                     
## [3759,] "Selenomonas_3"                   NA                     
## [3760,] NA                                NA                     
## [3761,] "Alloprevotella"                  NA                     
## [3762,] NA                                NA                     
## [3763,] "Treponema_2"                     NA                     
## [3764,] "Bergeyella"                      NA                     
## [3765,] "Prevotella_7"                    NA                     
## [3766,] "Prevotella_7"                    NA                     
## [3767,] "Lautropia"                       NA                     
## [3768,] "Dialister"                       NA                     
## [3769,] "Corynebacterium"                 NA                     
## [3770,] "Chelonobacter"                   NA                     
## [3771,] "Prevotellaceae_YAB2003_group"    NA                     
## [3772,] NA                                NA                     
## [3773,] "Peptostreptococcus"              NA                     
## [3774,] "Lachnoanaerobaculum"             NA                     
## [3775,] "Selenomonas_3"                   NA                     
## [3776,] "Alloprevotella"                  NA                     
## [3777,] "Prevotella_7"                    NA                     
## [3778,] "Alloprevotella"                  NA                     
## [3779,] "Prevotella_7"                    NA                     
## [3780,] NA                                NA                     
## [3781,] "Rikenellaceae_RC9_gut_group"     NA                     
## [3782,] "Prevotellaceae_YAB2003_group"    NA                     
## [3783,] "Simonsiella"                     NA                     
## [3784,] "Alloprevotella"                  NA                     
## [3785,] NA                                NA                     
## [3786,] "Tannerella"                      NA                     
## [3787,] NA                                NA                     
## [3788,] NA                                NA                     
## [3789,] NA                                NA                     
## [3790,] "Selenomonas"                     NA                     
## [3791,] "Treponema_2"                     "socranskii"           
## [3792,] NA                                NA                     
## [3793,] "Capnocytophaga"                  NA                     
## [3794,] "Alloprevotella"                  NA                     
## [3795,] "Selenomonas"                     NA                     
## [3796,] NA                                NA                     
## [3797,] NA                                NA                     
## [3798,] NA                                NA                     
## [3799,] "Prevotella"                      NA                     
## [3800,] NA                                NA                     
## [3801,] NA                                NA                     
## [3802,] "Prevotellaceae_YAB2003_group"    NA                     
## [3803,] "Simonsiella"                     NA                     
## [3804,] "Lachnoanaerobaculum"             NA                     
## [3805,] "Anthococcus"                     NA                     
## [3806,] "Candidatus_Saccharimonas"        NA                     
## [3807,] "Johnsonella"                     NA                     
## [3808,] "Prevotella"                      NA                     
## [3809,] NA                                NA                     
## [3810,] NA                                NA                     
## [3811,] "Megasphaera"                     NA                     
## [3812,] "Veillonella"                     NA                     
## [3813,] "Anthococcus"                     NA                     
## [3814,] NA                                NA                     
## [3815,] "Prevotellaceae_YAB2003_group"    NA                     
## [3816,] "Bergeyella"                      NA                     
## [3817,] "Selenomonas_3"                   NA                     
## [3818,] "Selenomonas_3"                   NA                     
## [3819,] "Capnocytophaga"                  "ochracea"             
## [3820,] "Prevotella"                      NA                     
## [3821,] NA                                NA                     
## [3822,] "Treponema_2"                     NA                     
## [3823,] "Prevotella_7"                    NA                     
## [3824,] "Veillonella"                     NA                     
## [3825,] "Treponema_2"                     NA                     
## [3826,] "Lachnoanaerobaculum"             NA                     
## [3827,] NA                                NA                     
## [3828,] NA                                NA                     
## [3829,] "Lactobacillus"                   NA                     
## [3830,] NA                                NA                     
## [3831,] "Prevotella_7"                    NA                     
## [3832,] "Simonsiella"                     NA                     
## [3833,] NA                                NA                     
## [3834,] "Rothia"                          NA                     
## [3835,] "Porphyromonas"                   NA                     
## [3836,] "Alloprevotella"                  NA                     
## [3837,] NA                                NA                     
## [3838,] "Prevotella_2"                    NA                     
## [3839,] "Prevotella_7"                    NA                     
## [3840,] "Catonella"                       NA                     
## [3841,] NA                                NA                     
## [3842,] "Capnocytophaga"                  NA                     
## [3843,] NA                                NA                     
## [3844,] "Phocaeicola"                     "abscessus"            
## [3845,] NA                                NA                     
## [3846,] NA                                NA                     
## [3847,] "Prevotella_7"                    NA                     
## [3848,] "Cuneatibacter"                   NA                     
## [3849,] "Treponema_2"                     NA                     
## [3850,] "Lachnoanaerobaculum"             NA                     
## [3851,] "Bergeriella"                     NA                     
## [3852,] "Eikenella"                       NA                     
## [3853,] NA                                NA                     
## [3854,] NA                                NA                     
## [3855,] "Campylobacter"                   NA                     
## [3856,] NA                                NA                     
## [3857,] "Leptotrichia"                    NA                     
## [3858,] "Prevotella_7"                    NA                     
## [3859,] "Alloprevotella"                  NA                     
## [3860,] "Capnocytophaga"                  NA                     
## [3861,] NA                                NA                     
## [3862,] "Lachnoanaerobaculum"             NA                     
## [3863,] "Johnsonella"                     NA                     
## [3864,] NA                                NA                     
## [3865,] NA                                NA                     
## [3866,] "Lachnoanaerobaculum"             NA                     
## [3867,] NA                                NA                     
## [3868,] "Prevotella"                      NA                     
## [3869,] NA                                NA                     
## [3870,] "Porphyromonas"                   NA                     
## [3871,] "Haemophilus"                     "haemolyticus"         
## [3872,] "Comamonas"                       NA                     
## [3873,] NA                                NA                     
## [3874,] "Prevotellaceae_YAB2003_group"    NA                     
## [3875,] NA                                NA                     
## [3876,] "Necropsobacter"                  NA                     
## [3877,] "Prevotella_7"                    NA                     
## [3878,] "Campylobacter"                   NA                     
## [3879,] "Bergeyella"                      NA                     
## [3880,] NA                                NA                     
## [3881,] "Treponema_2"                     NA                     
## [3882,] "Prevotella_7"                    "buccae"               
## [3883,] NA                                NA                     
## [3884,] "Dialister"                       NA                     
## [3885,] NA                                NA                     
## [3886,] NA                                NA                     
## [3887,] "Ezakiella"                       NA                     
## [3888,] "Ruminococcaceae_UCG-014"         NA                     
## [3889,] NA                                NA                     
## [3890,] "Mogibacterium"                   NA                     
## [3891,] NA                                NA                     
## [3892,] NA                                NA                     
## [3893,] "Lachnoanaerobaculum"             NA                     
## [3894,] NA                                NA                     
## [3895,] "Actinomyces"                     NA                     
## [3896,] "Prevotella"                      NA                     
## [3897,] NA                                NA                     
## [3898,] NA                                NA                     
## [3899,] "Cardiobacterium"                 NA                     
## [3900,] NA                                NA                     
## [3901,] "Gemella"                         NA                     
## [3902,] "Selenomonas_4"                   NA                     
## [3903,] "Prevotella_2"                    NA                     
## [3904,] "Porphyromonas"                   NA                     
## [3905,] "Treponema_2"                     "maltophilum"          
## [3906,] "Catonella"                       NA                     
## [3907,] "Prevotella"                      NA                     
## [3908,] "Necropsobacter"                  NA                     
## [3909,] NA                                NA                     
## [3910,] "Prevotella_7"                    NA                     
## [3911,] "Prevotella_7"                    NA                     
## [3912,] NA                                NA                     
## [3913,] "Selenomonas_3"                   NA                     
## [3914,] "Solobacterium"                   NA                     
## [3915,] NA                                NA                     
## [3916,] "Gemella"                         NA                     
## [3917,] "Prevotella_7"                    NA                     
## [3918,] NA                                NA                     
## [3919,] "Aggregatibacter"                 NA                     
## [3920,] NA                                NA                     
## [3921,] NA                                NA                     
## [3922,] "Prevotella"                      NA                     
## [3923,] NA                                NA                     
## [3924,] "Duganella"                       NA                     
## [3925,] NA                                NA                     
## [3926,] "Anaeroglobus"                    NA                     
## [3927,] NA                                NA                     
## [3928,] NA                                NA                     
## [3929,] "Thermus"                         NA                     
## [3930,] NA                                NA                     
## [3931,] NA                                NA                     
## [3932,] NA                                NA                     
## [3933,] NA                                NA                     
## [3934,] NA                                NA                     
## [3935,] "Comamonas"                       NA                     
## [3936,] "Treponema_2"                     NA                     
## [3937,] "Lactobacillus"                   NA                     
## [3938,] "Enhydrobacter"                   "aerosaccus"           
## [3939,] "Lactobacillus"                   "fuchuensis"           
## [3940,] "Anaerococcus"                    NA                     
## [3941,] "Prevotella_7"                    NA                     
## [3942,] "Alloprevotella"                  NA                     
## [3943,] NA                                NA                     
## [3944,] "Stomatobaculum"                  NA                     
## [3945,] "Prevotella_2"                    NA                     
## [3946,] NA                                NA                     
## [3947,] NA                                NA                     
## [3948,] NA                                NA                     
## [3949,] "Porphyromonas"                   NA                     
## [3950,] NA                                NA                     
## [3951,] "Anthococcus"                     NA                     
## [3952,] "Prevotella"                      NA                     
## [3953,] NA                                NA                     
## [3954,] NA                                NA                     
## [3955,] "Campylobacter"                   "curvus"               
## [3956,] NA                                NA                     
## [3957,] NA                                NA                     
## [3958,] NA                                NA                     
## [3959,] "Prevotella"                      NA                     
## [3960,] "Veillonella"                     NA                     
## [3961,] NA                                NA                     
## [3962,] NA                                NA                     
## [3963,] "Prevotella_6"                    NA                     
## [3964,] NA                                NA                     
## [3965,] "Prevotella_7"                    NA                     
## [3966,] NA                                NA                     
## [3967,] NA                                NA                     
## [3968,] "Leptotrichia"                    NA                     
## [3969,] NA                                NA                     
## [3970,] "Campylobacter"                   NA                     
## [3971,] NA                                NA                     
## [3972,] "Acholeplasma"                    NA                     
## [3973,] "Mycoplasma"                      "hominis"              
## [3974,] "Bergeyella"                      NA                     
## [3975,] "Desulfovibrio"                   NA                     
## [3976,] NA                                NA                     
## [3977,] "Prevotella"                      NA                     
## [3978,] NA                                NA                     
## [3979,] "Anthococcus"                     NA                     
## [3980,] NA                                NA                     
## [3981,] NA                                NA                     
## [3982,] "Anthococcus"                     NA                     
## [3983,] "Prevotella"                      NA                     
## [3984,] "Centipeda"                       NA                     
## [3985,] "Stomatobaculum"                  NA                     
## [3986,] "Alloprevotella"                  NA                     
## [3987,] "Prevotella"                      NA                     
## [3988,] "Capnocytophaga"                  "ochracea"             
## [3989,] "Cryptobacterium"                 NA                     
## [3990,] "Dialister"                       NA                     
## [3991,] "Leptotrichia"                    NA                     
## [3992,] NA                                NA                     
## [3993,] "Alloprevotella"                  NA                     
## [3994,] "Streptococcus"                   "constellatus"         
## [3995,] "Treponema_2"                     "socranskii"           
## [3996,] "Capnocytophaga"                  NA                     
## [3997,] "Peptoanaerobacter"               NA                     
## [3998,] NA                                NA                     
## [3999,] NA                                NA                     
## [4000,] NA                                NA                     
## [4001,] "Anthococcus"                     NA                     
## [4002,] "Treponema_2"                     NA                     
## [4003,] "Gemella"                         NA                     
## [4004,] "Capnocytophaga"                  NA                     
## [4005,] "Leptotrichia"                    NA                     
## [4006,] "Prevotella_7"                    NA                     
## [4007,] "Selenomonas_3"                   NA                     
## [4008,] "Alloprevotella"                  NA                     
## [4009,] "Bergeyella"                      NA                     
## [4010,] "Aggregatibacter"                 NA                     
## [4011,] NA                                NA                     
## [4012,] "Tannerella"                      NA                     
## [4013,] "Peptostreptococcus"              NA                     
## [4014,] "Selenomonas_3"                   NA                     
## [4015,] "Actinomyces"                     NA                     
## [4016,] NA                                NA                     
## [4017,] "Prevotella"                      NA                     
## [4018,] NA                                NA                     
## [4019,] "Peptococcus"                     NA                     
## [4020,] "Rothia"                          NA                     
## [4021,] "Prevotella_7"                    NA                     
## [4022,] "Actinomyces"                     NA                     
## [4023,] "Veillonella"                     NA                     
## [4024,] NA                                NA                     
## [4025,] "Capnocytophaga"                  "ochracea"             
## [4026,] "Prevotella"                      NA                     
## [4027,] "Anthococcus"                     NA                     
## [4028,] "Selenomonas_3"                   NA                     
## [4029,] NA                                NA                     
## [4030,] NA                                NA                     
## [4031,] "Neisseria"                       NA                     
## [4032,] "Actinomyces"                     NA                     
## [4033,] "Treponema_2"                     NA                     
## [4034,] "Isobaculum"                      NA                     
## [4035,] NA                                NA                     
## [4036,] "Pseudoramibacter"                "alactolyticus"        
## [4037,] NA                                NA                     
## [4038,] "Veillonella"                     NA                     
## [4039,] "Neisseria"                       NA                     
## [4040,] NA                                NA                     
## [4041,] "Alloprevotella"                  NA                     
## [4042,] NA                                NA                     
## [4043,] "Prevotella_2"                    NA                     
## [4044,] "Capnocytophaga"                  NA                     
## [4045,] "Lachnoanaerobaculum"             NA                     
## [4046,] NA                                NA                     
## [4047,] "Prevotella_7"                    NA                     
## [4048,] NA                                NA                     
## [4049,] NA                                NA                     
## [4050,] "Bergeyella"                      "cardium"              
## [4051,] "Prevotella_7"                    NA                     
## [4052,] "Prevotella_7"                    NA                     
## [4053,] NA                                NA                     
## [4054,] NA                                NA                     
## [4055,] NA                                NA                     
## [4056,] NA                                NA                     
## [4057,] NA                                NA                     
## [4058,] NA                                NA                     
## [4059,] "Streptococcus"                   NA                     
## [4060,] NA                                NA                     
## [4061,] NA                                NA                     
## [4062,] "Actinomyces"                     "gerencseriae"         
## [4063,] "Candidatus_Saccharimonas"        NA                     
## [4064,] NA                                NA                     
## [4065,] NA                                NA                     
## [4066,] "Tannerella"                      NA                     
## [4067,] "Stomatobaculum"                  NA                     
## [4068,] NA                                NA                     
## [4069,] NA                                NA                     
## [4070,] "Corynebacterium"                 NA                     
## [4071,] "Anthococcus"                     NA                     
## [4072,] NA                                NA                     
## [4073,] "Selenomonas_4"                   NA                     
## [4074,] "Selenomonas_3"                   NA                     
## [4075,] NA                                NA                     
## [4076,] "Lachnoanaerobaculum"             NA                     
## [4077,] "Capnocytophaga"                  NA                     
## [4078,] NA                                NA                     
## [4079,] "Alloprevotella"                  NA                     
## [4080,] "Capnocytophaga"                  NA                     
## [4081,] NA                                NA                     
## [4082,] "Anaeroglobus"                    NA                     
## [4083,] NA                                NA                     
## [4084,] NA                                NA                     
## [4085,] NA                                NA                     
## [4086,] NA                                NA                     
## [4087,] NA                                NA                     
## [4088,] "Lautropia"                       NA                     
## [4089,] "Corynebacterium"                 NA                     
## [4090,] NA                                NA                     
## [4091,] NA                                NA                     
## [4092,] NA                                NA                     
## [4093,] "Peptostreptococcus"              NA                     
## [4094,] "Stomatobaculum"                  NA                     
## [4095,] NA                                NA                     
## [4096,] "Ruminococcaceae_UCG-014"         NA                     
## [4097,] "Candidatus_Saccharimonas"        NA                     
## [4098,] NA                                NA                     
## [4099,] "Alloprevotella"                  NA                     
## [4100,] "Peptococcus"                     NA                     
## [4101,] NA                                NA                     
## [4102,] "Chryseobacterium"                NA                     
## [4103,] NA                                NA                     
## [4104,] "Mogibacterium"                   NA                     
## [4105,] "Alloprevotella"                  NA                     
## [4106,] "Prevotella_7"                    NA                     
## [4107,] NA                                NA                     
## [4108,] NA                                NA                     
## [4109,] "Porphyromonas"                   NA                     
## [4110,] "Anaeroglobus"                    NA                     
## [4111,] "Granulicatella"                  NA                     
## [4112,] "Veillonella"                     NA                     
## [4113,] NA                                NA                     
## [4114,] NA                                NA                     
## [4115,] NA                                NA                     
## [4116,] "Filifactor"                      NA                     
## [4117,] "Atopobium"                       NA                     
## [4118,] "Rikenellaceae_RC9_gut_group"     NA                     
## [4119,] "Atopobium"                       NA                     
## [4120,] NA                                NA                     
## [4121,] NA                                NA                     
## [4122,] NA                                NA                     
## [4123,] NA                                NA                     
## [4124,] NA                                NA                     
## [4125,] NA                                NA                     
## [4126,] NA                                NA                     
## [4127,] "Anthococcus"                     NA                     
## [4128,] "Necropsobacter"                  NA                     
## [4129,] "Prevotella"                      NA                     
## [4130,] NA                                NA                     
## [4131,] NA                                NA                     
## [4132,] NA                                NA                     
## [4133,] "Capnocytophaga"                  NA                     
## [4134,] "Prevotella"                      NA                     
## [4135,] NA                                NA                     
## [4136,] NA                                NA                     
## [4137,] NA                                NA                     
## [4138,] NA                                NA                     
## [4139,] "Cardiobacterium"                 NA                     
## [4140,] "Ruminococcaceae_UCG-014"         NA                     
## [4141,] "Bergeriella"                     NA                     
## [4142,] "Parvimonas"                      NA                     
## [4143,] "Butyrivibrio_2"                  NA                     
## [4144,] "Prevotella_7"                    NA                     
## [4145,] NA                                NA                     
## [4146,] "Prevotella_2"                    NA                     
## [4147,] NA                                NA                     
## [4148,] "Veillonella"                     NA                     
## [4149,] "Prevotella"                      NA                     
## [4150,] "Filifactor"                      NA                     
## [4151,] NA                                NA                     
## [4152,] "Treponema_2"                     NA                     
## [4153,] NA                                NA                     
## [4154,] "Dialister"                       NA                     
## [4155,] "Prevotella_2"                    NA                     
## [4156,] "Corynebacterium"                 NA                     
## [4157,] "Veillonella"                     NA                     
## [4158,] NA                                NA                     
## [4159,] NA                                NA                     
## [4160,] "Prevotella_7"                    NA                     
## [4161,] NA                                NA                     
## [4162,] NA                                NA                     
## [4163,] NA                                NA                     
## [4164,] NA                                NA                     
## [4165,] NA                                NA                     
## [4166,] "Necropsobacter"                  NA                     
## [4167,] "Gemella"                         NA                     
## [4168,] NA                                NA                     
## [4169,] NA                                NA                     
## [4170,] "Anaeroglobus"                    NA                     
## [4171,] "Prevotella_2"                    NA                     
## [4172,] NA                                NA                     
## [4173,] "Alloprevotella"                  NA                     
## [4174,] "Johnsonella"                     NA                     
## [4175,] "Treponema_2"                     NA                     
## [4176,] "Comamonas"                       NA                     
## [4177,] NA                                NA                     
## [4178,] "Neisseria"                       NA                     
## [4179,] "Prevotellaceae_YAB2003_group"    NA                     
## [4180,] NA                                NA                     
## [4181,] NA                                NA                     
## [4182,] "Bergeriella"                     NA                     
## [4183,] "Mycoplasma"                      "spermatophilum"       
## [4184,] "Bacteroides"                     NA                     
## [4185,] NA                                NA                     
## [4186,] NA                                NA                     
## [4187,] NA                                NA                     
## [4188,] "Selenomonas"                     "sputigena"            
## [4189,] NA                                NA                     
## [4190,] "Treponema_2"                     NA                     
## [4191,] NA                                NA                     
## [4192,] "Bergeriella"                     NA                     
## [4193,] "Gemella"                         NA                     
## [4194,] NA                                NA                     
## [4195,] "Capnocytophaga"                  NA                     
## [4196,] NA                                NA                     
## [4197,] "Parvimonas"                      NA                     
## [4198,] "Veillonella"                     NA                     
## [4199,] NA                                NA                     
## [4200,] "Acinetobacter"                   "proteolyticus"        
## [4201,] "Anthococcus"                     NA                     
## [4202,] "Fusobacterium"                   NA                     
## [4203,] "Stomatobaculum"                  NA                     
## [4204,] "Prevotella_7"                    NA                     
## [4205,] "Veillonella"                     NA                     
## [4206,] NA                                NA                     
## [4207,] "Lawsonella"                      NA                     
## [4208,] "Lachnoanaerobaculum"             NA                     
## [4209,] "Streptobacillus"                 NA                     
## [4210,] "Treponema_2"                     "denticola"            
## [4211,] NA                                NA                     
## [4212,] NA                                NA                     
## [4213,] NA                                NA                     
## [4214,] "Prevotella_7"                    NA                     
## [4215,] "Prevotella_2"                    NA                     
## [4216,] NA                                NA                     
## [4217,] "Candidatus_Saccharimonas"        NA                     
## [4218,] "Prevotella_7"                    NA                     
## [4219,] NA                                NA                     
## [4220,] NA                                NA                     
## [4221,] NA                                NA                     
## [4222,] NA                                NA                     
## [4223,] "Soonwooa"                        NA                     
## [4224,] "Capnocytophaga"                  NA                     
## [4225,] NA                                NA                     
## [4226,] "Leptotrichia"                    NA                     
## [4227,] "Prevotella_7"                    NA                     
## [4228,] NA                                NA                     
## [4229,] "Necropsobacter"                  NA                     
## [4230,] NA                                NA                     
## [4231,] "Prevotella_7"                    NA                     
## [4232,] "Actinobacillus"                  NA                     
## [4233,] NA                                NA                     
## [4234,] "Prevotella_7"                    NA                     
## [4235,] NA                                NA                     
## [4236,] "Candidatus_Saccharimonas"        NA                     
## [4237,] NA                                NA                     
## [4238,] "Treponema_2"                     NA                     
## [4239,] "Corynebacterium"                 NA                     
## [4240,] "Bergeriella"                     NA                     
## [4241,] NA                                NA                     
## [4242,] NA                                NA                     
## [4243,] NA                                NA                     
## [4244,] NA                                NA                     
## [4245,] "Porphyromonas"                   NA                     
## [4246,] "Actinomyces"                     NA                     
## [4247,] "Treponema_2"                     NA                     
## [4248,] "Gemella"                         NA                     
## [4249,] NA                                NA                     
## [4250,] "Treponema_2"                     NA                     
## [4251,] "Finegoldia"                      "magna"                
## [4252,] NA                                NA                     
## [4253,] "Desulfobulbus"                   NA                     
## [4254,] NA                                NA                     
## [4255,] NA                                NA                     
## [4256,] "Prevotella_7"                    NA                     
## [4257,] NA                                NA                     
## [4258,] NA                                NA                     
## [4259,] "Capnocytophaga"                  NA                     
## [4260,] "Bergeyella"                      NA                     
## [4261,] NA                                NA                     
## [4262,] NA                                NA                     
## [4263,] NA                                NA                     
## [4264,] "Prevotella"                      NA                     
## [4265,] NA                                NA                     
## [4266,] NA                                NA                     
## [4267,] NA                                NA                     
## [4268,] "Atopobium"                       NA                     
## [4269,] "Prevotella"                      NA                     
## [4270,] "Anthococcus"                     NA                     
## [4271,] "Peptostreptococcus"              NA                     
## [4272,] "Lachnoanaerobaculum"             NA                     
## [4273,] NA                                NA                     
## [4274,] NA                                NA                     
## [4275,] "Prevotella_2"                    NA                     
## [4276,] "Catonella"                       NA                     
## [4277,] NA                                NA                     
## [4278,] "Anthococcus"                     NA                     
## [4279,] "Anthococcus"                     NA                     
## [4280,] "Porphyromonas"                   NA                     
## [4281,] NA                                NA                     
## [4282,] NA                                NA                     
## [4283,] NA                                NA                     
## [4284,] NA                                NA                     
## [4285,] "Solobacterium"                   NA                     
## [4286,] NA                                NA                     
## [4287,] "Bacteroides"                     NA                     
## [4288,] NA                                NA                     
## [4289,] NA                                NA                     
## [4290,] "Peptostreptococcus"              NA                     
## [4291,] NA                                NA                     
## [4292,] NA                                NA                     
## [4293,] "Cricetibacter"                   NA                     
## [4294,] NA                                NA                     
## [4295,] "Atopobium"                       NA                     
## [4296,] "Rikenellaceae_RC9_gut_group"     NA                     
## [4297,] NA                                NA                     
## [4298,] NA                                NA                     
## [4299,] NA                                NA                     
## [4300,] NA                                NA                     
## [4301,] NA                                NA                     
## [4302,] "Treponema_2"                     NA                     
## [4303,] "Chryseobacterium"                NA                     
## [4304,] "Selenomonas_3"                   NA                     
## [4305,] NA                                NA                     
## [4306,] NA                                NA                     
## [4307,] "Treponema_2"                     "parvum"               
## [4308,] NA                                NA                     
## [4309,] NA                                NA                     
## [4310,] "Veillonella"                     NA                     
## [4311,] "Bergeriella"                     NA                     
## [4312,] "Peptostreptococcus"              NA                     
## [4313,] NA                                NA                     
## [4314,] NA                                NA                     
## [4315,] NA                                NA                     
## [4316,] NA                                NA                     
## [4317,] NA                                NA                     
## [4318,] NA                                NA                     
## [4319,] "Treponema_2"                     NA                     
## [4320,] "Filifactor"                      NA                     
## [4321,] "Ruminococcaceae_UCG-014"         NA                     
## [4322,] NA                                NA                     
## [4323,] "Subdoligranulum"                 NA                     
## [4324,] NA                                NA                     
## [4325,] NA                                NA                     
## [4326,] "Cardiobacterium"                 NA                     
## [4327,] NA                                NA                     
## [4328,] "Erysipelotrichaceae_UCG-006"     NA                     
## [4329,] "Bifidobacterium"                 "longum"               
## [4330,] NA                                NA                     
## [4331,] "Veillonella"                     NA                     
## [4332,] NA                                NA                     
## [4333,] "Bergeriella"                     NA                     
## [4334,] NA                                NA                     
## [4335,] "Anthococcus"                     NA                     
## [4336,] "Wolinella"                       NA                     
## [4337,] "Treponema_2"                     NA                     
## [4338,] NA                                NA                     
## [4339,] "Sphingobium"                     "yanoikuyae"           
## [4340,] "Xanthomonas"                     NA                     
## [4341,] NA                                NA                     
## [4342,] "Gemella"                         NA                     
## [4343,] NA                                NA                     
## [4344,] "Leptotrichia"                    NA                     
## [4345,] "Rikenellaceae_RC9_gut_group"     NA                     
## [4346,] NA                                NA                     
## [4347,] "Parvimonas"                      NA                     
## [4348,] "Prevotella"                      NA                     
## [4349,] NA                                NA                     
## [4350,] "Tannerella"                      NA                     
## [4351,] NA                                NA                     
## [4352,] "Defluviitaleaceae_UCG-011"       NA                     
## [4353,] "Prevotella_2"                    NA                     
## [4354,] "Gemella"                         NA                     
## [4355,] "Alloprevotella"                  NA                     
## [4356,] NA                                NA                     
## [4357,] NA                                NA                     
## [4358,] "Fusobacterium"                   NA                     
## [4359,] NA                                NA                     
## [4360,] NA                                NA                     
## [4361,] NA                                NA                     
## [4362,] "Leptotrichia"                    NA                     
## [4363,] "Bergeyella"                      NA                     
## [4364,] "Peptostreptococcus"              NA                     
## [4365,] NA                                NA                     
## [4366,] NA                                NA                     
## [4367,] NA                                NA                     
## [4368,] NA                                NA                     
## [4369,] "Parvimonas"                      NA                     
## [4370,] NA                                NA                     
## [4371,] "Lelliottia"                      NA                     
## [4372,] "Cardiobacterium"                 "valvarum"             
## [4373,] NA                                NA                     
## [4374,] "Leptotrichia"                    NA                     
## [4375,] "Dialister"                       NA                     
## [4376,] "Treponema_2"                     NA                     
## [4377,] "Prevotella"                      NA                     
## [4378,] NA                                NA                     
## [4379,] "Lachnoanaerobaculum"             NA                     
## [4380,] NA                                NA                     
## [4381,] NA                                NA                     
## [4382,] "Prevotella"                      NA                     
## [4383,] NA                                NA                     
## [4384,] "Cryptobacterium"                 NA                     
## [4385,] NA                                NA                     
## [4386,] "Rikenellaceae_RC9_gut_group"     NA                     
## [4387,] "Veillonella"                     NA                     
## [4388,] "Lachnoanaerobaculum"             NA                     
## [4389,] NA                                NA                     
## [4390,] "Selenomonas_4"                   NA                     
## [4391,] NA                                NA                     
## [4392,] NA                                NA                     
## [4393,] NA                                NA                     
## [4394,] "Selenomonas_4"                   NA                     
## [4395,] "Solobacterium"                   NA                     
## [4396,] "Centipeda"                       NA                     
## [4397,] "Centipeda"                       NA                     
## [4398,] NA                                NA                     
## [4399,] NA                                NA                     
## [4400,] NA                                NA                     
## [4401,] "Anaeroglobus"                    NA                     
## [4402,] NA                                NA                     
## [4403,] "Kingella"                        NA                     
## [4404,] "Candidatus_Saccharimonas"        NA                     
## [4405,] "Atopobium"                       NA                     
## [4406,] "Prevotella"                      NA                     
## [4407,] NA                                NA                     
## [4408,] "Methylobacterium"                NA                     
## [4409,] NA                                NA                     
## [4410,] NA                                NA                     
## [4411,] "Family_XIII_UCG-001"             NA                     
## [4412,] "Lactobacillus"                   NA                     
## [4413,] "Atopobium"                       NA                     
## [4414,] "Treponema_2"                     NA                     
## [4415,] NA                                NA                     
## [4416,] NA                                NA                     
## [4417,] "Fretibacterium"                  NA                     
## [4418,] NA                                NA                     
## [4419,] NA                                NA                     
## [4420,] "Atopobium"                       NA                     
## [4421,] NA                                NA                     
## [4422,] "Mycoplasma"                      NA                     
## [4423,] "Cardiobacterium"                 NA                     
## [4424,] "Staphylococcus"                  NA                     
## [4425,] NA                                NA                     
## [4426,] NA                                NA                     
## [4427,] "Johnsonella"                     NA                     
## [4428,] NA                                NA                     
## [4429,] NA                                NA                     
## [4430,] "Atopobium"                       NA                     
## [4431,] NA                                NA                     
## [4432,] "Prevotellaceae_YAB2003_group"    NA                     
## [4433,] NA                                NA                     
## [4434,] "Peptococcus"                     NA                     
## [4435,] NA                                NA                     
## [4436,] NA                                NA                     
## [4437,] NA                                NA                     
## [4438,] "Candidatus_Saccharimonas"        NA                     
## [4439,] NA                                NA                     
## [4440,] NA                                NA                     
## [4441,] "Megasphaera"                     NA                     
## [4442,] "Actinobacillus"                  NA                     
## [4443,] "Bulleidia"                       "extructa"             
## [4444,] "Suttonella"                      NA                     
## [4445,] "Capnocytophaga"                  NA                     
## [4446,] "Treponema_2"                     NA                     
## [4447,] "Capnocytophaga"                  NA                     
## [4448,] "Gemella"                         NA                     
## [4449,] "Porphyromonas"                   NA                     
## [4450,] "Prevotella"                      NA                     
## [4451,] NA                                NA                     
## [4452,] "Prevotella"                      NA                     
## [4453,] "Actinomyces"                     NA                     
## [4454,] "Chelonobacter"                   NA                     
## [4455,] NA                                NA                     
## [4456,] NA                                NA                     
## [4457,] "Bergeriella"                     NA                     
## [4458,] "Solobacterium"                   NA                     
## [4459,] "Alloprevotella"                  NA                     
## [4460,] NA                                NA                     
## [4461,] NA                                NA                     
## [4462,] NA                                NA                     
## [4463,] NA                                NA                     
## [4464,] NA                                NA                     
## [4465,] NA                                NA                     
## [4466,] "Atopobium"                       NA                     
## [4467,] "Lautropia"                       NA                     
## [4468,] NA                                NA                     
## [4469,] NA                                NA                     
## [4470,] "Prevotella_7"                    NA                     
## [4471,] "Anaerostipes"                    NA                     
## [4472,] "Stomatobaculum"                  NA                     
## [4473,] "Prevotella"                      NA                     
## [4474,] "Alloprevotella"                  NA                     
## [4475,] "Johnsonella"                     NA                     
## [4476,] "Porphyromonas"                   NA                     
## [4477,] "Selenomonas_4"                   NA                     
## [4478,] "Prevotella_7"                    NA                     
## [4479,] "Tannerella"                      NA                     
## [4480,] "Anthococcus"                     NA                     
## [4481,] NA                                NA                     
## [4482,] "Peptococcus"                     NA                     
## [4483,] NA                                NA                     
## [4484,] "Alloprevotella"                  NA                     
## [4485,] "Selenomonas"                     NA                     
## [4486,] "Erysipelotrichaceae_UCG-008"     NA                     
## [4487,] NA                                NA                     
## [4488,] NA                                NA                     
## [4489,] "Treponema_2"                     NA                     
## [4490,] "Catonella"                       NA                     
## [4491,] "Peptoanaerobacter"               NA                     
## [4492,] "Delftia"                         NA                     
## [4493,] NA                                NA                     
## [4494,] NA                                NA                     
## [4495,] "Prevotella"                      NA                     
## [4496,] "Pseudomonas"                     NA                     
## [4497,] "Afipia"                          NA                     
## [4498,] "Gemella"                         NA                     
## [4499,] "Prevotella_7"                    NA                     
## [4500,] NA                                NA                     
## [4501,] NA                                NA                     
## [4502,] "Anthococcus"                     NA                     
## [4503,] "Bergeriella"                     NA                     
## [4504,] "Johnsonella"                     NA                     
## [4505,] NA                                NA                     
## [4506,] "Prevotella_7"                    "denticola"            
## [4507,] "Prevotella"                      NA                     
## [4508,] "Lachnoanaerobaculum"             NA                     
## [4509,] NA                                NA                     
## [4510,] "Necropsobacter"                  NA                     
## [4511,] "Prevotella_7"                    NA                     
## [4512,] NA                                NA                     
## [4513,] NA                                NA                     
## [4514,] NA                                NA                     
## [4515,] NA                                NA                     
## [4516,] "Candidatus_Saccharimonas"        NA                     
## [4517,] "Leptotrichia"                    NA                     
## [4518,] "Dialister"                       NA                     
## [4519,] NA                                NA                     
## [4520,] NA                                NA                     
## [4521,] "Bergeriella"                     NA                     
## [4522,] "Prevotella_7"                    NA                     
## [4523,] "Prevotella_7"                    NA                     
## [4524,] "Brachymonas"                     NA                     
## [4525,] "W5053"                           NA                     
## [4526,] "Capnocytophaga"                  "haemolytica"          
## [4527,] "Candidatus_Saccharimonas"        NA                     
## [4528,] "Alloprevotella"                  NA                     
## [4529,] "Centipeda"                       NA                     
## [4530,] NA                                NA                     
## [4531,] "Porphyromonas"                   NA                     
## [4532,] NA                                NA                     
## [4533,] NA                                NA                     
## [4534,] NA                                NA                     
## [4535,] "Porphyromonas"                   NA                     
## [4536,] NA                                NA                     
## [4537,] NA                                NA                     
## [4538,] "Anthococcus"                     NA                     
## [4539,] "Bacteroides"                     NA                     
## [4540,] NA                                NA                     
## [4541,] NA                                NA                     
## [4542,] "Stomatobaculum"                  NA                     
## [4543,] NA                                NA                     
## [4544,] NA                                NA                     
## [4545,] NA                                NA                     
## [4546,] "Treponema_2"                     NA                     
## [4547,] NA                                NA                     
## [4548,] "Rikenellaceae_RC9_gut_group"     NA                     
## [4549,] "Centipeda"                       NA                     
## [4550,] "Streptococcus"                   NA                     
## [4551,] "Leptotrichia"                    NA                     
## [4552,] NA                                NA                     
## [4553,] "Treponema_2"                     NA                     
## [4554,] "Prevotella_2"                    NA                     
## [4555,] NA                                NA
```

```
#inspect the taxonomic assignments after adding species
taxa.print <- taxa
rownames(taxa.print) <- NULL
head(taxa.print)
```

```
##      Kingdom    Phylum           Class                 Order            
## [1,] "Bacteria" "Firmicutes"     "Negativicutes"       "Selenomonadales"
## [2,] "Bacteria" "Firmicutes"     "Bacilli"             "Lactobacillales"
## [3,] "Bacteria" "Bacteroidetes"  "Bacteroidia"         "Bacteroidales"  
## [4,] "Bacteria" "Proteobacteria" "Gammaproteobacteria" "Pasteurellales" 
## [5,] "Bacteria" "Firmicutes"     "Negativicutes"       "Selenomonadales"
## [6,] "Bacteria" "Firmicutes"     "Negativicutes"       "Selenomonadales"
##      Family             Genus           Species         
## [1,] "Veillonellaceae"  "Veillonella"   "dispar"        
## [2,] "Streptococcaceae" "Streptococcus" NA              
## [3,] "Prevotellaceae"   "Prevotella_7"  "melaninogenica"
## [4,] "Pasteurellaceae"  "Haemophilus"   NA              
## [5,] "Veillonellaceae"  "Veillonella"   "rogosae"       
## [6,] "Veillonellaceae"  "Veillonella"   "atypica"
```

```
table(nchar(getSequences(seqtab)))
```

```
## 
##  395 
## 4555
```

```
# Write to disk
saveRDS(seqtab, "/Users/luizroesch/Desktop/Saliva/seqs/dada2/fastq/seqtab_final.rds") 
saveRDS(tax, "/Users/luizroesch/Desktop/Saliva/seqs/dada2/fastq/tax_final.rds")
saveRDS(taxa, "/Users/luizroesch/Desktop/Saliva/seqs/dada2/fastq/taxa_sp_final.rds")
```
