## Supplementary material for "PIME: a package for discovery of novel differences among microbial communities": Detailed and reproducible description of PIME data analysis.

### Pime Validation Saliva dataset

*Roesch et al.*

*4/11/2019*

**Likelihood of introducing bias while building prevalence-filtered datasets** **7**

##### Load objects generated by dada2

##### Make a phyloseq object

```
library(phyloseq)
library(ggplot2)
library(microbiome)
library(vegan)
library(knitr)
library(ggpubr)
library(pime)
seqtab = readRDS("/Users/luizroesch/Desktop/Saliva/seqs/dada2/fastq/seqtab_final.rds")
tax= readRDS("/Users/luizroesch/Desktop/Saliva/seqs/dada2/fastq/tax_final.rds")
taxa = readRDS("/Users/luizroesch/Desktop/Saliva/seqs/dada2/fastq/taxa_sp_final.rds")
map <- "/Users/luizroesch/Desktop/Saliva/seqs/dada2/map.txt"
ps <- phyloseq(otu_table(seqtab, taxa_are_rows=FALSE),
               tax_table(taxa))
sample_metadata = import_qiime_sample_data(map)
saliva =merge_phyloseq(ps, sample_metadata)
saliva
```

```
## phyloseq-class experiment-level object
## otu_table()   OTU Table:         [ 4555 taxa and 125 samples ]
## sample_data() Sample Data:       [ 125 samples by 21 sample variables ]
## tax_table()   Taxonomy Table:    [ 4555 taxa by 7 taxonomic ranks ]
```

```
summarize_phyloseq(saliva)
```

```
## Compositional = NO
## 1] Min. number of reads = 24956
```

```
## 2] Max. number of reads = 223138
## 3] Total number of reads = 5532280
## 4] Average number of reads = 44258.24
## 5] Median number of reads = 39432
## 7] Sparsity = 0.95352096597146
## 6] Any OTU sum to 1 or less? NO
## 8] Number of singletons = 0
## 9] Percent of OTUs that are singletons 0
## 10] Number of sample variables are: 21
## X.NAME
## KIDMED_index
## Gender
## Age
## Fruit.daily.
## Second.fruit.daily.
## Vegetable.daily.
## Second.vegetable.daily.
## Fish.regularly.
## Pulses..1.weekly
## Pasta.rice..5.weekly
## Cereals.grains.breakfast.
## Nut.2.3.weekly
## Olive.oil.used.
## Dairy.at.breakfast
## yogurt.cheese.3.weekly.
## Fast.food..1.weekly
## Skip.breakfast..3.weekly.
## Baked.goods.pastires.breakfast.
## Sweets.3.daily.
## Total
```

```
rank_names(saliva)
```

```
## [1] "Kingdom" "Phylum"  "Class"   "Order"   "Family"  "Genus"   "Species"
```

```
set.seed(2125)
salivaR = rarefy_even_depth(saliva, sample.size = 24900)
```

### Finding microbiota differences among diet without PIME filtering

#### Randon Forest approach

```
library(randomForest)
train= otu_table(salivaR)
# Make one column for our outcome/response variable
KIDMED_index <- as.factor(sample_data(salivaR)$KIDMED_index)
# Combine them into 1 data frame
training.set <- data.frame(KIDMED_index, train)
train.model = randomForest(KIDMED_index ~ ., data = training.set, importance = TRUE)
print(train.model)
```

```
##
## Call:
```

```
##  randomForest(formula = KIDMED_index ~ ., data = training.set,      importance = TRUE)
##                Type of random forest: classification
##                      Number of trees: 500
## No. of variables tried at each split: 66
##
##         OOB estimate of  error rate: 47.2%
## Confusion matrix:
##        High Low Medium class.error
## High      0   1     23   1.0000000
## Low       0   4     33   0.8918919
## Medium    0   2     62   0.0312500
```

The OOB error rate is 47.2% This estimate of error suggests that when the resulting model is applied to new observations, the answers will be in error 47.2% of the time. That is, it is 52.8% accurate, which is not a reasonable model. We still can calculate the variable importance but the results will be inaccurate. The table lists each ASV and then measures of importance for each ASV. Higher values indicate that the variable is relatively more important.

```
imp= train.model$importance
ftable= cbind.data.frame(imp, (tax_table(salivaR)))
rownames(ftable) <- NULL
library(knitr)
my_kable = function(x, max.rows=30, ...) {
  kable(x[1:max.rows, ], ...)
}
my_kable(ftable[ , c(1:4, 11:12)], caption = "Importance of ASVs to differentiate diet
         - dataset not filtered by PIME",
         full_width = F)
```

Table 1: Importance of ASVs to differentiate diet - dataset not filtered by PIME

| High | Low | Medium | MeanDecreaseAccuracy | Genus | Species |
|---:|---:|---:|---:|---|---|
| 0.0001412 | 0.0002420 | -0.0003781 | -0.0002383 | Veillonella | dispar |
| 0.0005316 | -0.0000757 | 0.0003621 | 0.0002460 | Streptococcus | NA |
| -0.0007087 | -0.0002381 | -0.0009387 | -0.0006611 | Prevotella_7 | melaninogenica |
| 0.0005970 | -0.0013382 | 0.0004795 | -0.0000990 | Haemophilus | NA |
| -0.0002831 | -0.0007718 | -0.0007693 | -0.0006733 | Veillonella | rogosae |
| 0.0010591 | -0.0017230 | 0.0017728 | 0.0006763 | Veillonella | atypica |
| -0.0003485 | -0.0003950 | -0.0005560 | -0.0004844 | Veillonella | atypica |
| -0.0017937 | -0.0016305 | 0.0006960 | -0.0004635 | NA | NA |
| 0.0001222 | -0.0006316 | -0.0008806 | -0.0005597 | Neisseria | NA |
| 0.0003855 | -0.0007201 | -0.0002264 | -0.0000468 | Veillonella | tobetsuensis |
| 0.0088414 | 0.0002376 | 0.0041391 | 0.0037162 | Neisseria | NA |
| 0.0003636 | 0.0001971 | -0.0004207 | -0.0000435 | Veillonella | NA |
| 0.0003571 | -0.0005756 | 0.0002678 | -0.0000314 | Fusobacterium | periodonticum |
| -0.0002325 | 0.0018595 | 0.0021934 | 0.0015672 | Veillonella | NA |
| -0.0014024 | 0.0003352 | -0.0005835 | -0.0003285 | Prevotella_7 | NA |
| -0.0000500 | -0.0002927 | 0.0011552 | 0.0004818 | Campylobacter | concisus |
| -0.0004937 | 0.0005244 | -0.0000235 | 0.0001619 | Haemophilus | parainfluenzae |
| -0.0000808 | -0.0000694 | 0.0005813 | 0.0004070 | Oribacterium | sinus |
| 0.0010420 | -0.0002127 | 0.0004703 | 0.0003438 | Veillonella | NA |
| -0.0016994 | -0.0005211 | -0.0013471 | -0.0011476 | Alloprevotella | NA |
| -0.0006175 | -0.0002555 | 0.0001594 | 0.0000610 | Granulicatella | NA |
| -0.0005944 | 0.0000784 | -0.0004873 | -0.0003954 | Prevotella_7 | histicola |

| High | Low | Medium | MeanDecreaseAccuracy | Genus | Species |
|---|---|---|---|---|---|
| -0.0014318 | 0.0003491 | 0.0002360 | -0.0000664 | Prevotella_7 | melaninogenica |
| 0.0014731 | -0.0011255 | 0.0010072 | 0.0004806 | Veillonella | NA |
| 0.0024232 | -0.0000084 | -0.0007168 | 0.0000653 | Haemophilus | NA |
| 0.0012032 | -0.0000730 | 0.0010958 | 0.0007395 | Prevotella_7 | histicola |
| 0.0003660 | 0.0021627 | 0.0014040 | 0.0014680 | Porphyromonas | NA |
| 0.0003190 | -0.0002518 | 0.0004577 | 0.0002383 | Veillonella | NA |
| 0.0005707 | -0.0006580 | 0.0000373 | -0.0000560 | Streptococcus | NA |
| -0.0006699 | 0.0000757 | -0.0014930 | -0.0008448 | Streptococcus | NA |

Negative values of Mean Decrease Accuracy are a clear warning sign that your model might be overfitting noise.

### Running PIME

#### Prediction using random forests on full dataset

```
OOB_error_full=pime.oob.error(salivaR, "KIDMED_index")
OOB_error_full
```

```
## [1] 0.48
```

The OOB error rate <=0.1, indicated the dataset present large differences, and pime might not remove much of the noise. Higher OOB error rate indicates that the next functions should be run to find the best prevalence interval for the dataset.

The best prevalence interval is that one with OOB error rate close to zero. Here we will test the dataset at prevalence interval of 65% but other intervals of prevalence can be tested. For instance, the prevalence interval of 25% has OOB error of 7.2%. This indicates that the model is 92.8% accurate, which is a reasonably good model.

##### To get the table with ASV importance of any chosen prevalence interval.

```r
imp65=best.prev$`Importance`$`Prevalence 65`
library(knitr)
kable(imp65[ ,c(2:5,12:13)], caption = "Importance of ASVs to differentiate diet treatments")
```

Table 2: Importance of ASVs to differentiate diet treatments

| High | Low | Medium | MeanDecreaseAccuracy | Genus | Species |
|---:|---:|---:|---:|---|---|
| 0.0199890 | 0.0286621 | 0.0882889 | 0.0572971 | Veillonella | atypica |
| 0.0490419 | 0.0060162 | 0.0776384 | 0.0504103 | Neisseria | NA |
| 0.0166776 | 0.0153442 | 0.0566553 | 0.0363870 | Neisseria | NA |
| 0.0317372 | 0.0454642 | 0.0317731 | 0.0349324 | Prevotella_7 | NA |
| 0.0278527 | 0.0344717 | 0.0260247 | 0.0286506 | Alloprevotella | NA |
| 0.0236309 | 0.0311944 | 0.0277640 | 0.0276086 | Prevotella_7 | melaninogenica |
| 0.0087427 | 0.0045976 | 0.0482760 | 0.0273222 | Haemophilus | NA |
| 0.0200383 | 0.0652593 | 0.0026502 | 0.0239046 | Porphyromonas | NA |
| 0.0145192 | 0.0250054 | 0.0250142 | 0.0227588 | Megasphaera | micronuciformis |
| 0.0188147 | 0.0205621 | 0.0233007 | 0.0216439 | Selenomonas_3 | NA |
| 0.0278639 | 0.0261529 | 0.0167201 | 0.0213493 | Haemophilus | NA |
| 0.0457836 | 0.0107400 | 0.0155470 | 0.0197877 | Lautropia | mirabilis |
| 0.0500097 | 0.0074543 | 0.0123015 | 0.0177629 | Capnocytophaga | sputigena |
| 0.0304463 | 0.0273772 | 0.0090561 | 0.0175554 | Campylobacter | NA |

| High | Low | Medium | MeanDecreaseAccuracy | Genus | Species |
|---:|---:|---:|---:|---|---|
| 0.0382307 | 0.0063770 | 0.0156118 | 0.0165545 | Prevotella | pallens |
| 0.0240077 | 0.0149386 | 0.0113565 | 0.0148534 | Porphyromonas | NA |
| 0.0031292 | 0.0354766 | 0.0031614 | 0.0127990 | Megasphaera | NA |
| 0.0234088 | 0.0166500 | 0.0072665 | 0.0126463 | Fusobacterium | NA |
| 0.0456022 | 0.0048811 | 0.0039165 | 0.0122248 | Haemophilus | NA |
| 0.0547647 | 0.0050642 | 0.0011161 | 0.0118354 | Granulicatella | elegans |
| 0.0034961 | 0.0332455 | 0.0015585 | 0.0109399 | Gemella | NA |
| 0.0196707 | 0.0140168 | 0.0058010 | 0.0108141 | Rothia | NA |
| 0.0174090 | 0.0112367 | 0.0081562 | 0.0105538 | Dialister | invisus |
| 0.0423435 | 0.0008468 | 0.0011259 | 0.0084171 | Solobacterium | NA |
| 0.0373067 | 0.0043891 | 0.0001041 | 0.0082167 | Prevotella_7 | histicola |
| 0.0052723 | 0.0223817 | 0.0000747 | 0.0074152 | Prevotella_7 | histicola |
| 0.0292156 | 0.0005130 | 0.0012151 | 0.0059559 | Prevotella | nigrescens |
| -0.0000150 | -0.0003763 | 0.0019169 | 0.0008718 | Butyrivibrio_2 | NA |
| 0.0001048 | 0.0005321 | 0.0012789 | 0.0008716 | NA | NA |
| -0.0007024 | 0.0007604 | 0.0013591 | 0.0007723 | Kingella | oralis |

#### Likelihood of introducing bias while building prevalence-filtered datasets

```
randomized=pime.error.prediction(salivaR, "KIDMED_index", bootstrap = 100, parallel = TRUE, max.prev = [illegible]
randomized$Plot
```

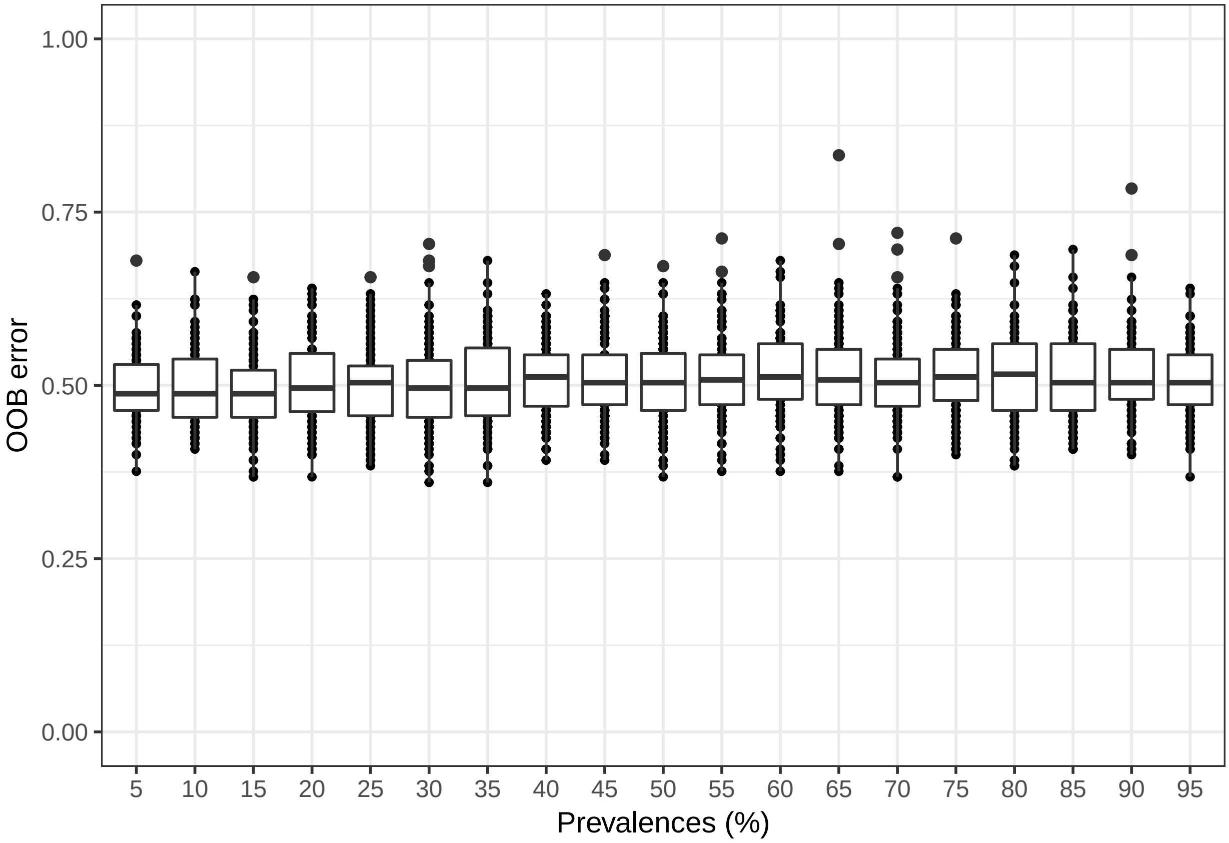

The first function randomizes the samples labels into arbitrary groupings using 100 random permutations. For each randomized prevalence filtered dataset, the OOB error rate is calculated to determine whether differences in the original groups occur by chance.

```
replicated.oob.error= pime.oob.replicate(prevalences, "KIDMED_index", bootstrap = 100, parallel = TRUE)
replicated.oob.error$Plot
```

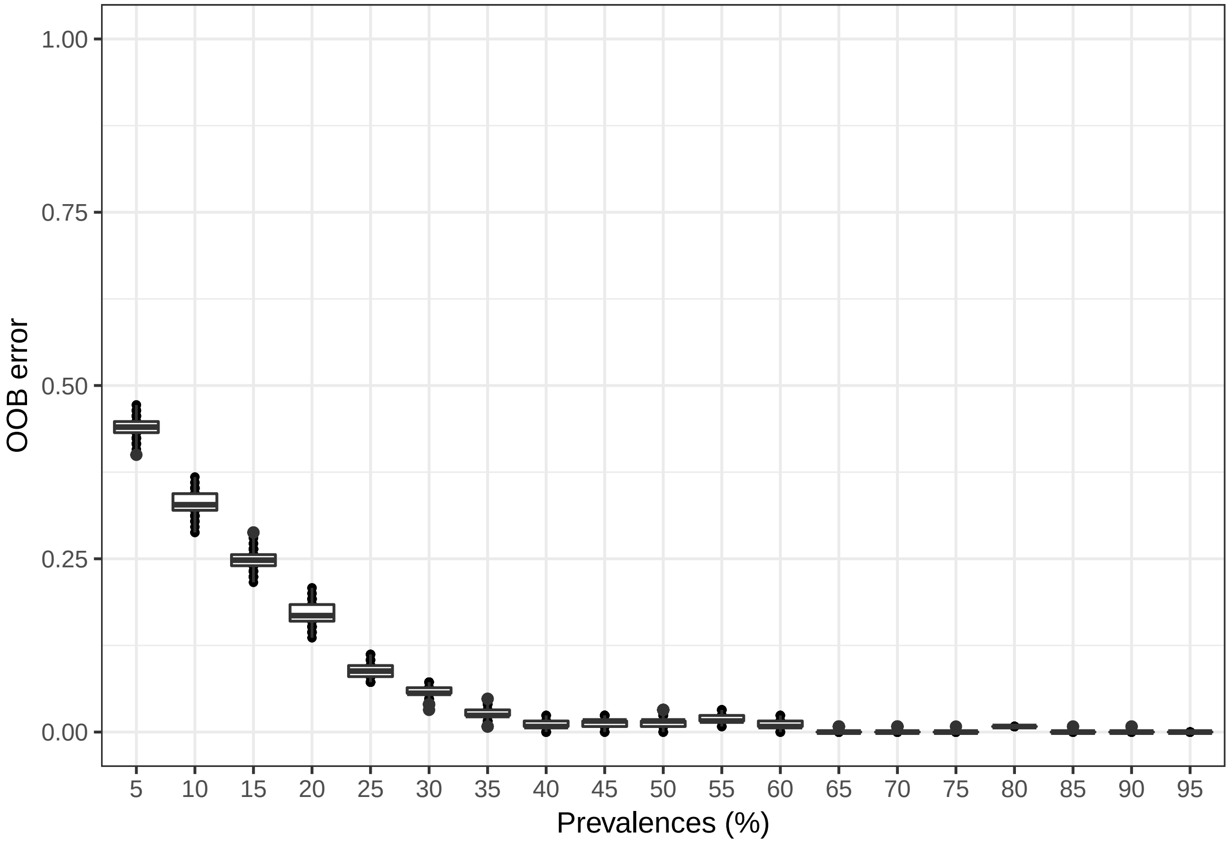

The second function performs the Random Forest analyses and computes the OOB error for 100 replications in each prevalence interval without randomizing the sample labels. The biological difference among samples is expected to be greater than the differences generated randomly. Thus, the greatest fraction of randomizations should generate high error rates. On the other hand, no improvement in accuracy is expected within the randomized dataset.

### Pime Validation 2nd dataset

*Roesch et al.*

*4/11/2019*

**Likelihood of introducing bias while building prevalence-filtered datasets** **7**

#### Running the dataset from Davis-Richardson et al. (2014)

##### Load objects generated by dada2

##### Make a phyloseq object

```
library(phyloseq)
library(ggplot2)
library(microbiome)
library(vegan)
library(knitr)
library(ggpubr)
library(pime)
seqtab.nochim = readRDS("/Users/luizroesch/Desktop/D3C/dada2/seqtab_nonchim.rds")
taxa = readRDS("/Users/luizroesch/Desktop/D3C/dada2/taxa.rds")
map <- "/Users/luizroesch/Desktop/D3C/dada2/map.txt"
ps <- phyloseq(otu_table(seqtab.nochim, taxa_are_rows=FALSE),
             tax_table(taxa))
sample_metadata = import_qiime_sample_data(map)
dipp1 =merge_phyloseq(ps, sample_metadata)
dipp1
```

```
## phyloseq-class experiment-level object
## otu_table()   OTU Table:         [ 10726 taxa and 931 samples ]
## sample_data() Sample Data:       [ 931 samples by 33 sample variables ]
## tax_table()   Taxonomy Table:    [ 10726 taxa by 6 taxonomic ranks ]
```

```r
rank_names(dipp1)
```

```
## [1] "Kingdom" "Phylum"  "Class"   "Order"   "Family"  "Genus"
```

```r
dipp = subset_samples(dipp1, current_status=="control" | current_status=="pre_sero")
Turku = subset_samples(dipp, site=="Turku")
Turku.1 = subset_samples(Turku, TechReps=="FALSE")
summarize_phyloseq(Turku.1)
```

```
## Compositional = NO
## 1] Min. number of reads = 18094
## 2] Max. number of reads = 271818
## 3] Total number of reads = 36576086
## 4] Average number of reads = 102454.022408964
## 5] Median number of reads = 93757
## 7] Sparsity = 0.977152300412986
## 6] Any OTU sum to 1 or less? YES
## 8] Number of singletons = 4468
## 9] Percent of OTUs that are singletons 1251.5406162465
## 10] Number of sample variables are: 33
## X.SampleID
## illumina_id
## mask_id
## sample_id
## site
## TechRep_type
## TechReps
## age_at_sampling
## Age_Months
## pair
## seroconverted
## current_status
## age_first_sc
## aa_count
## aa_ICA
## aa_IAA
## aa_GADA
## aa_IA2A
## age_GADA
## age_IA2A
## age_IAA
## age_ICA
## has_T1D
## age_T1D
## DQB1_cat
## Gender
## MoD_simp
## Breast_feeding_any
## Duration_Breast_Feeding_months
## Duration_exclusive_breast_feeding_weeks
## total_reads
## antibiotics
## antibiotic_courses
```

```r
set.seed(2125)
inputR = rarefy_even_depth(Turku.1, sample.size = 9450)
```

### Finding microbiota differences between autoimmune and control subjects without PIME filtering

#### Randon Forest approach

```r
library(randomForest)
train= otu_table(inputR)
# Make one column for our outcome/response variable
seroconverted <- as.factor(sample_data(inputR)$seroconverted)
# Combine them into 1 data frame
training.set <- data.frame(seroconverted, train)
train.model = randomForest(seroconverted ~ ., data = training.set, importance = TRUE)
print(train.model)
```

```
##
## Call:
##  randomForest(formula = seroconverted ~ ., data = training.set,      importance = TRUE)
##                Type of random forest: classification
##                      Number of trees: 500
## No. of variables tried at each split: 71
##
##         OOB estimate of  error rate: 19.05%
## Confusion matrix:
##       FALSE TRUE class.error
## FALSE   210   14    0.062500
## TRUE     54   79    0.406015
```

The OOB error rate is 19.05% This estimate of error suggests that when the resulting model is applied to new observations, the answers will be in error 47.2% of the time. That is, it is 80.95% accurate, which is a reasonable good model.

We can calculate the variable importance and expect a relatively accurate model. The table lists each ASV and then measures of importance for each ASV. Higher values indicate that the variable is relatively more important.

```r
imp= train.model$importance
ftable= cbind.data.frame(imp, (tax_table(inputR)))
rownames(ftable) <- NULL
library(knitr)
my_kable = function(x, max.rows=30, ...) {
  kable(x[1:max.rows, ], ...)
}
my_kable(ftable[ , c(1:3, 10)], caption = "Importance of ASVs to differentiate autoimmune vs. control s
         - dataset not filtered by PIME",
         full_width = F)
```

Table 1: Importance of ASVs to differentiate autoimmune vs. control subjects - dataset not filtered by PIME

| FALSE | TRUE | MeanDecreaseAccuracy | Genus |
|---|---|---|---|
| 0.0027737 | 0.0061260 | 0.0040229 | Bacteroides |
| 0.0030593 | 0.0032532 | 0.0030675 | Bacteroides |
| 0.0011049 | 0.0036452 | 0.0020475 | Bacteroides |
| 0.0036919 | 0.0017167 | 0.0030263 | Bacteroides |
| 0.0005608 | 0.0012160 | 0.0007776 | Bacteroides |
| 0.0011502 | 0.0003429 | 0.0008414 | Bacteroides |
| 0.0024688 | 0.0020223 | 0.0023129 | Bacteroides |
| 0.0021085 | 0.0014692 | 0.0018860 | Bacteroides |
| 0.0028044 | 0.0047162 | 0.0034930 | Bacteroides |
| 0.0005086 | 0.0008318 | 0.0006262 | Bifidobacterium |
| 0.0029725 | 0.0019331 | 0.0026164 | Bacteroides |
| 0.0009011 | 0.0007443 | 0.0007863 | NA |
| 0.0002948 | 0.0008422 | 0.0005063 | Bifidobacterium |
| 0.0030590 | 0.0035091 | 0.0031696 | Bacteroides |
| 0.0006980 | 0.0015966 | 0.0010235 | NA |
| 0.0031825 | 0.0044689 | 0.0036732 | Bacteroides |
| 0.0025244 | 0.0040647 | 0.0031011 | Bacteroides |
| 0.0042504 | 0.0042783 | 0.0042528 | Bacteroides |
| 0.0001046 | 0.0003151 | 0.0002130 | Blautia |
| 0.0006497 | 0.0007506 | 0.0006734 | Bacteroides |
| 0.0010946 | 0.0007470 | 0.0010054 | Bacteroides |
| 0.0007692 | 0.0011580 | 0.0009358 | NA |
| 0.0006069 | 0.0016966 | 0.0010199 | Bacteroides |
| 0.0009788 | 0.0000343 | 0.0006665 | Bacteroides |
| 0.0004795 | -0.0000395 | 0.0003009 | Bifidobacterium |
| 0.0005750 | 0.0015003 | 0.0009386 | Bacteroides |
| 0.0006706 | 0.0010255 | 0.0008338 | Bifidobacterium |
| 0.0008356 | 0.0009516 | 0.0009099 | NA |
| 0.0004466 | 0.0003679 | 0.0004381 | Bacteroides |
| 0.0002399 | 0.0013466 | 0.0006526 | Bifidobacterium |

The first nine ASVs were classified as important to differentiate autoimune cases vs. controls. Those ASVs were close related to Bacteroidetes. Previously, Davis-Richardson et al. (2014) discovered that the relative abundance of Bacteroides was significantly higher in autoimmune vs. control subjects.

To validate our workflow we expect to confirm the results found by Davis-Richardson et al. (2014) by using PIME.

### Running PIME

#### Prediction using random forests on full dataset

```
OOB_error_full=pime.oob.error(inputR, "seroconverted")
OOB_error_full
```

```
## [1] 0.1960784
```

The OOB error rate <=0.1, indicated the dataset present large differences, and pime might not remove much of the noise. Higher OOB error rate indicates that the next functions should be run to find the best prevalence interval for the dataset.

##### Split the dataset by predictor variable

```r
per_variable_obj= pime.split.by.variable(inputR, "seroconverted")
per_variable_obj
```

```
## $`FALSE`
## phyloseq-class experiment-level object
## otu_table()   OTU Table:         [ 4167 taxa and 224 samples ]
## sample_data() Sample Data:       [ 224 samples by 33 sample variables ]
## tax_table()   Taxonomy Table:    [ 4167 taxa by 6 taxonomic ranks ]
##
## $`TRUE`
## phyloseq-class experiment-level object
## otu_table()   OTU Table:         [ 2904 taxa and 133 samples ]
## sample_data() Sample Data:       [ 133 samples by 33 sample variables ]
## tax_table()   Taxonomy Table:    [ 2904 taxa by 6 taxonomic ranks ]
```

##### Calculate the highest possible prevalence intervals

This function calculates prevalence for different intervals by increments of 5. The input file is the output from the pime.split.by.variable(per__variable__obj)

```r
prevalences=pime.prevalence(per_variable_obj)
head(prevalences)
```

```
## $`5`
## phyloseq-class experiment-level object
## otu_table()   OTU Table:         [ 979 taxa and 357 samples ]
## sample_data() Sample Data:       [ 357 samples by 33 sample variables ]
## tax_table()   Taxonomy Table:    [ 979 taxa by 6 taxonomic ranks ]
##
## $`10`
## phyloseq-class experiment-level object
## otu_table()   OTU Table:         [ 556 taxa and 357 samples ]
## sample_data() Sample Data:       [ 357 samples by 33 sample variables ]
## tax_table()   Taxonomy Table:    [ 556 taxa by 6 taxonomic ranks ]
##
## $`15`
## phyloseq-class experiment-level object
## otu_table()   OTU Table:         [ 431 taxa and 357 samples ]
## sample_data() Sample Data:       [ 357 samples by 33 sample variables ]
## tax_table()   Taxonomy Table:    [ 431 taxa by 6 taxonomic ranks ]
##
## $`20`
## phyloseq-class experiment-level object
## otu_table()   OTU Table:         [ 335 taxa and 357 samples ]
## sample_data() Sample Data:       [ 357 samples by 33 sample variables ]
## tax_table()   Taxonomy Table:    [ 335 taxa by 6 taxonomic ranks ]
##
```

```
## $`25`
## phyloseq-class experiment-level object
## otu_table()   OTU Table:         [ 258 taxa and 357 samples ]
## sample_data() Sample Data:       [ 357 samples by 33 sample variables ]
## tax_table()   Taxonomy Table:    [ 258 taxa by 6 taxonomic ranks ]
##
## $`30`
## phyloseq-class experiment-level object
## otu_table()   OTU Table:         [ 219 taxa and 357 samples ]
## sample_data() Sample Data:       [ 357 samples by 33 sample variables ]
## tax_table()   Taxonomy Table:    [ 219 taxa by 6 taxonomic ranks ]
```

#### Calculate the best prevalence interval for the dataset

This function will return a table with Out of Bag error from random forests for each prevalence interval. The number of taxa and the number of remaining sequences for each prevalence interval are also computed. The best prevalence interval value provides the clearest separation of communities while still including a majority of the taxa in the analysis. If true differences are present. It will be represented by the first interval in which the OOB error rate is zero or close to zero. The input file is the list of prevalences generated by the pime.prevalence (prevalences) and the predictor variable ("seroconverted").

```r
set.seed(2125)
best.prev=pime.best.prevalence(prevalences, "seroconverted")
```

```
##        Interval OOB error rate (%) OTUs   Nseqs
##   Prevalence 5%              13.17  979 3152779
##  Prevalence 10%              10.64  556 2975527
##  Prevalence 15%               6.16  431 2891465
##  Prevalence 20%               3.64  335 2773432
##  Prevalence 25%               3.36  258 2556564
##  Prevalence 30%                2.8  219 2457867
##  Prevalence 35%               2.24  175 2339966
##  Prevalence 40%               2.24  143 2119062
##  Prevalence 45%                1.4  115 1922701
##  Prevalence 50%               2.52   99 1733801
##  Prevalence 55%               1.12   74 1357557
##  Prevalence 60%                  0   62 1165304
##  Prevalence 65%               0.28   49 1026879
##  Prevalence 70%               0.56   39  897367
```

The best prevalence interval is that one with OOB error rate close to zero. Here we will test the dataset at prevalence interval of 60% but other intervals of prevalence can be tested. For instance, the prevalence interval of 15% has OOB error of 10.64%. This indicates that the model is about 90% accurate, which is a reasonably good model.

#### Get the table with ASV importance of any chosen prevalence interval.

```r
imp60=best.prev$`Importance`$`Prevalence 60`
library(knitr)
kable(imp60[ ,c(2:4,11)], caption = "Importance of ASVs to differentiate autoimmune vs. control subjects
```

Table 2: Importance of ASVs to differentiate autoimmune vs. control subjects

| FALSE. | TRUE. | MeanDecreaseAccuracy | Genus |
|---|---|---|---|
| 0.0588016 | 0.0147796 | 0.0421928 | Bacteroides |
| 0.0470208 | 0.0118621 | 0.0338832 | Bacteroides |
| 0.0386230 | 0.0118253 | 0.0284351 | Bifidobacterium |
| 0.0371892 | 0.0072908 | 0.0260364 | Bifidobacterium |
| 0.0374713 | 0.0062792 | 0.0257281 | Bacteroides |
| 0.0365809 | 0.0070485 | 0.0255316 | Bacteroides |
| 0.0355649 | 0.0073829 | 0.0251251 | Bacteroides |
| 0.0371600 | 0.0044714 | 0.0251073 | Bacteroides |
| 0.0365959 | 0.0057748 | 0.0250061 | Bacteroides |
| 0.0345973 | 0.0075772 | 0.0245141 | Bacteroides |
| 0.0338302 | 0.0062864 | 0.0234888 | Bacteroides |
| 0.0345660 | 0.0036705 | 0.0230925 | Bacteroides |
| 0.0320649 | 0.0060689 | 0.0222983 | Bacteroides |
| 0.0302848 | 0.0073825 | 0.0216310 | Bacteroides |
| 0.0277696 | 0.0081509 | 0.0203184 | Bacteroides |
| 0.0285186 | 0.0050400 | 0.0196142 | Blautia |
| 0.0256354 | 0.0056571 | 0.0181410 | Blautia |
| 0.0261868 | 0.0046818 | 0.0178867 | Blautia |
| 0.0251795 | 0.0056913 | 0.0176057 | Veillonella |
| 0.0262274 | 0.0021835 | 0.0172836 | Veillonella |
| 0.0242705 | 0.0025109 | 0.0161146 | NA |
| 0.0233260 | 0.0018888 | 0.0152578 | Blautia |
| 0.0217452 | 0.0019501 | 0.0141460 | Veillonella |
| 0.0199318 | 0.0025108 | 0.0133215 | Veillonella |
| 0.0170497 | 0.0034517 | 0.0118528 | Veillonella |
| 0.0129373 | 0.0009900 | 0.0084477 | Veillonella |
| 0.0009143 | 0.0000755 | 0.0005954 | Veillonella |
| 0.0001202 | 0.0008933 | 0.0003859 | NA |
| 0.0000427 | 0.0007290 | 0.0003016 | Intestinibacter |
| 0.0003467 | 0.0001654 | 0.0002801 | Flavonifractor |

PIME still show the importance of Bacteroidetes to classify TD1 subjects pre-seroconversion

### Likelihood of introducing bias while building prevalence-filtered datasets

```
randomized=pime.error.prediction(inputR, "seroconverted", bootstrap = 100, parallel = TRUE, max.prev = 9
randomized$Plot
```

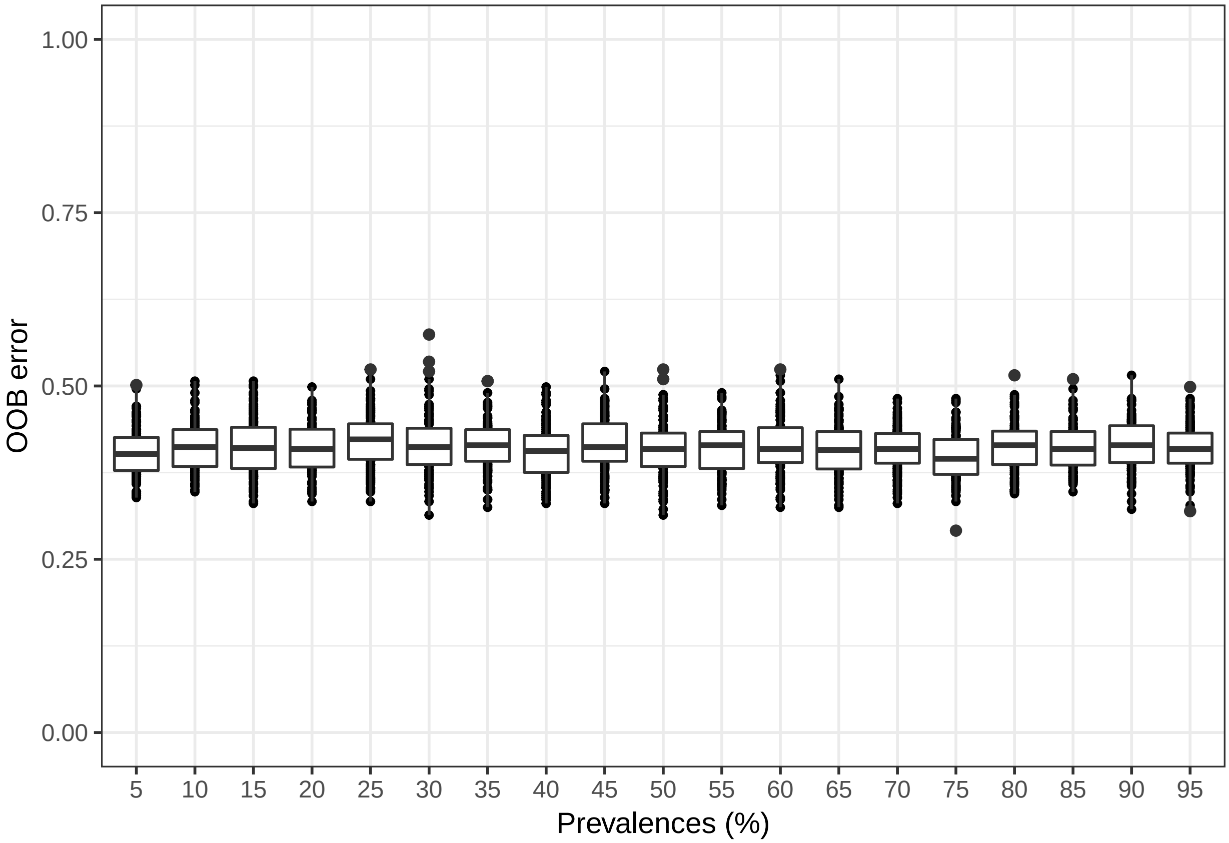

The first function randomizes the samples labels into arbitrary groupings using 100 random permutations. For each randomized prevalence filtered dataset, the OOB error rate is calculated to determine whether differences in the original groups occur by chance.

```
replicated.oob.error= pime.oob.replicate(prevalences, "seroconverted", bootstrap = 100, parallel = TRUE)
replicated.oob.error$Plot
```

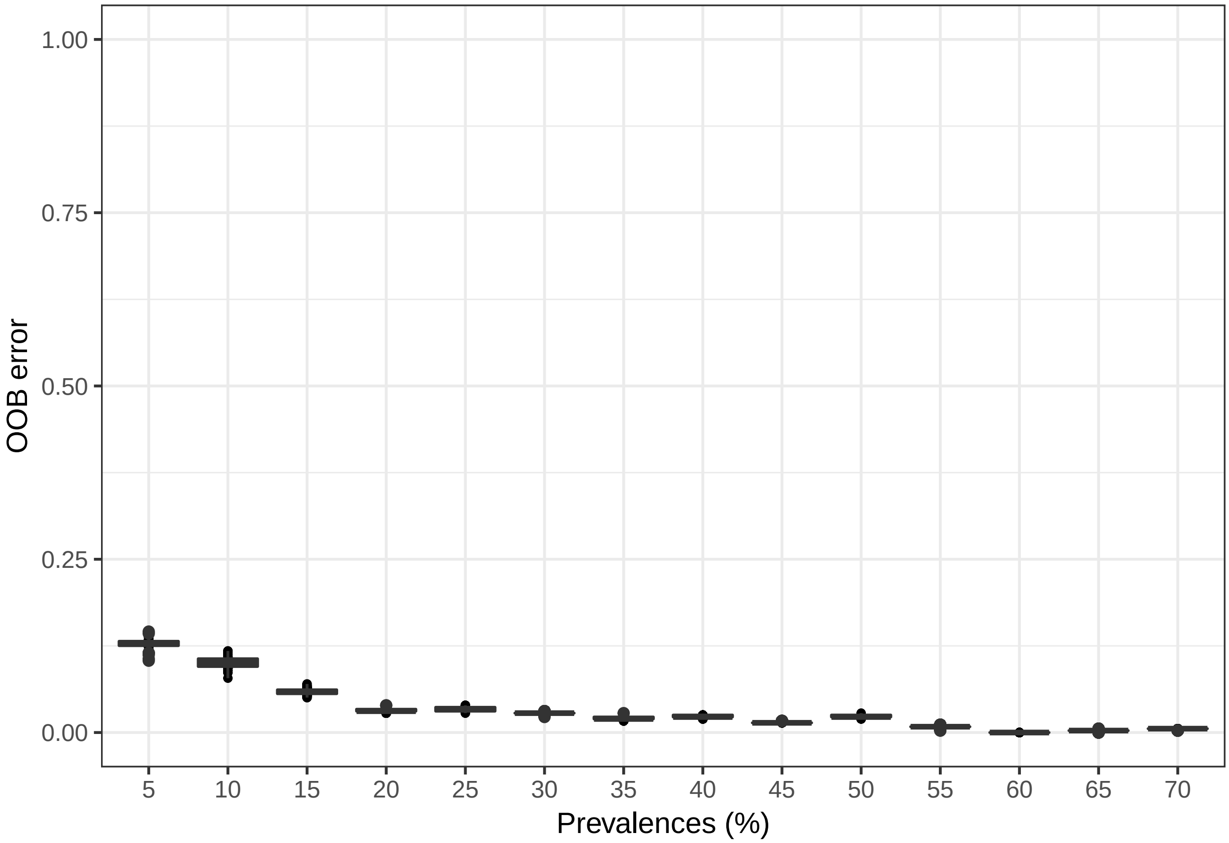

The second function performs the Random Forest analyses and computes the OOB error for 100 replications in each prevalence interval without randomizing the sample labels. The biological difference among samples is expected to be greater than the differences generated randomly. Thus, the greatest fraction of randomizations should generate high error rates. On the other hand, no improvement in accuracy is expected within the randomized dataset.

### Pime Validation 3rd dataset

*Roesch et al.*

*4/11/2019*

#### Contents

**Running the dataset from Avershina et al., 2017** **1**
Load files and make a phyloseq object . . . . . 1

**Finding differences between the vaginal microbiome in pregnant women randomized to receive milk with or without probiotic bacterial strains without PIME filtering** **2**

**Likelihood of introducing bias while building prevalence-filtered datasets** **6**

### Running the dataset from Avershina et al., 2017

#### Load files and make a phyloseq object

```
library(phyloseq)
library(ggplot2)
library(microbiome)
library(vegan)
library(knitr)
library(ggpubr)
library(pime)
jsonbiomfile = "/Users/luizroesch/Desktop/Saliva/seqs/dada2/Core/vaginal_microbiota/otu_table_tax.biom"
mapfile1 = "/Users/luizroesch/Desktop/Saliva/seqs/dada2/Core/vaginal_microbiota/map.txt"
biom1 = import_biom(jsonbiomfile, mapfile1, parseFunction=parse_taxonomy_default)
map1 = import_qiime_sample_data(mapfile1)
inputR = merge_phyloseq(biom1,map1)
inputR
```

```
## phyloseq-class experiment-level object
## otu_table()   OTU Table:         [ 463 taxa and 335 samples ]
## sample_data() Sample Data:       [ 335 samples by 4 sample variables ]
## tax_table()   Taxonomy Table:    [ 463 taxa by 7 taxonomic ranks ]
```

```
summarize_phyloseq(inputR)
```

```
## Compositional = NO
## 1] Min. number of reads = 3000
## 2] Max. number of reads = 3000
## 3] Total number of reads = 1005000
```

```
## 4] Average number of reads = 3000
## 5] Median number of reads = 3000
## 7] Sparsity = 0.948467167402727
## 6] Any OTU sum to 1 or less? YES
## 8] Number of singletons = 21
## 9] Percent of OTUs that are singletons 4.53563714902808
## 10] Number of sample variables are: 4
## ID
## Group
## Probiotic
## Blood
```

```
rank_names(inputR)
```

```
## [1] "Rank1" "Rank2" "Rank3" "Rank4" "Rank5" "Rank6" "Rank7"
```

```
colnames(tax_table(inputR)) = c( "Kingdom", "Phylum", "Class", "Order", "Family", "Genus", "Species")
rank_names(inputR)
```

```
## [1] "Kingdom" "Phylum"  "Class"   "Order"   "Family"  "Genus"   "Species"
```

### Finding differences between the vaginal microbiome in pregnant women randomized to receive milk with or without probiotic bacterial strains without PIME filtering

#### Randon Forest approach

```
library(randomForest)
train= t(otu_table(inputR))
# Make one column for our outcome/response variable
Probiotic <- as.factor(sample_data(inputR)$Probiotic)
# Combine them into 1 data frame
training.set <- data.frame(Probiotic, train)
train.model = randomForest(Probiotic ~ ., data = training.set, importance = TRUE)
print(train.model)
```

```
##
## Call:
##  randomForest(formula = Probiotic ~ ., data = training.set, importance = TRUE)
##                Type of random forest: classification
##                      Number of trees: 500
## No. of variables tried at each split: 21
##
##         OOB estimate of  error rate: 48.06%
## Confusion matrix:
##     no yes class.error
## no  99  72   0.4210526
## yes 89  75   0.5426829
```

The OOB error rate is 45.37% This estimate of error suggests that when the resulting model is applied to new observations, the answers will be in error 45.37% of the time. That is, it is 54.63% accurate, which is not a reasonable model.

We still can calculate the variable importance but the results will be inaccurate. The table lists each ASV and then measures of importance for each ASV. Higher values indicate that the variable is relatively more important.

```
importance = importance(train.model)
ftable= cbind.data.frame(importance, (tax_table(inputR)))
rownames(ftable) <- NULL
library(knitr)
my_kable = function(x, max.rows=30, ...) {
  kable(x[1:max.rows, ], ...)
}
my_kable(ftable[ , c(1:3, 10)], caption = "Importance of ASVs to differentiate microbiome in pregnant w
         - dataset not filtered by PIME",
         full_width = F)
```

Table 1: Importance of ASVs to differentiate microbiome in pregnant women randomized to receive milk with or without probiotic bacterial strains - dataset not filtered by PIME

| no | yes | MeanDecreaseAccuracy | Genus |
|---:|---:|---:|:---|
| -1.1041002 | 0.0936667 | -0.8556814 | g__Lactobacillus |
| 2.1393620 | 4.0529250 | 4.3208797 | g__Lactobacillus |
| 8.3510078 | -1.0430953 | 5.1463426 | g__Lactobacillus |
| 0.5912786 | 4.0968771 | 3.1715829 | g__Lactobacillus |
| 0.4331607 | 0.4226862 | 0.5868810 | g__Lactobacillus |
| 1.2545575 | -2.2089412 | -0.5384735 | NA |
| 1.7529480 | -1.0210985 | 0.5948827 | g__ |
| -0.0827764 | -1.4061197 | -1.0838549 | g__ |
| 0.0000000 | 0.0000000 | 0.0000000 | g__Oscillospira |
| 0.7471672 | -0.5426303 | 0.1265587 | g__Pseudoalteromonas |
| 2.6670291 | -1.9235154 | 0.4335792 | g__Ralstonia |
| 0.6881364 | -1.5345408 | -0.4403926 | g__Lactobacillus |
| -2.6435365 | -0.6995249 | -2.3865977 | g__ |
| -1.3451455 | -0.0903209 | -1.0013189 | g__Sneathia |
| 0.2379641 | -0.5435183 | -0.1878372 | g__ |
| 1.2990199 | 0.1528160 | 1.0756434 | g__Pseudomonas |
| -0.5905026 | -0.5153560 | -0.9560497 | g__Megasphaera |
| 0.4037907 | 1.0010015 | 0.6202631 | g__ |
| 2.6131553 | 1.2708747 | 2.7345901 | g__Sphingomonas |
| 1.8465427 | -1.4643195 | 0.4409543 | g__Staphylococcus |
| 0.7391351 | 0.1303608 | 0.5268786 | g__ |
| -1.0672804 | -0.5459137 | -1.1887389 | g__Peptoniphilus |
| -1.3585546 | 1.9023533 | 0.1568671 | g__ |
| 3.2982886 | -0.9967514 | 1.3027347 | g__Atopobium |
| 0.7056267 | 2.6545632 | 2.0978635 | g__Ureaplasma |
| -0.4094100 | -1.1532525 | -1.1419941 | g__Pseudomonas |
| -0.4898722 | -1.1129731 | -1.1713058 | g__Methylobacterium |
| 1.3774403 | -1.4745811 | -0.4039159 | g__Aerococcus |
| 0.5437540 | -1.2290182 | -0.5614829 | g__Streptococcus |
| -1.7152791 | -0.4626517 | -1.6394801 | g__Parvimonas |

Negative values of Mean Decrease Accuracy are a clear warning sign that your model might be overfitting noise.

### Running PIME

#### Prediction using random forests on full dataset

```r
OOB_error_full=pime.oob.error(inputR, "Probiotic")
OOB_error_full
```

```
## [1] 0.4507463
```

The OOB error rate <=0.1, indicated the dataset present large differences, and pime might not remove much of the noise. Higher OOB error rate indicates that the next functions should be run to find the best prevalence interval for the dataset.

#### Split the dataset by predictor variable

```r
per_variable_obj= pime.split.by.variable(inputR, "Probiotic")
per_variable_obj
```

```
## $no
## phyloseq-class experiment-level object
## otu_table()   OTU Table:         [ 377 taxa and 171 samples ]
## sample_data() Sample Data:       [ 171 samples by 4 sample variables ]
## tax_table()   Taxonomy Table:    [ 377 taxa by 7 taxonomic ranks ]
##
## $yes
## phyloseq-class experiment-level object
## otu_table()   OTU Table:         [ 413 taxa and 164 samples ]
## sample_data() Sample Data:       [ 164 samples by 4 sample variables ]
## tax_table()   Taxonomy Table:    [ 413 taxa by 7 taxonomic ranks ]
```

#### Calculate the highest possible prevalence intervals

This function calculates prevalence for different intervals by increments of 5. The input file is the output from the pime.split.by.variable(per__variable__obj)

```r
prevalences=pime.prevalence(per_variable_obj)
head(prevalences)
```

```
## $`5`
## phyloseq-class experiment-level object
## otu_table()   OTU Table:         [ 123 taxa and 335 samples ]
## sample_data() Sample Data:       [ 335 samples by 4 sample variables ]
## tax_table()   Taxonomy Table:    [ 123 taxa by 7 taxonomic ranks ]
##
## $`10`
## phyloseq-class experiment-level object
## otu_table()   OTU Table:         [ 68 taxa and 335 samples ]
## sample_data() Sample Data:       [ 335 samples by 4 sample variables ]
## tax_table()   Taxonomy Table:    [ 68 taxa by 7 taxonomic ranks ]
##
## $`15`
## phyloseq-class experiment-level object
```

```
## otu_table()   OTU Table:         [ 40 taxa and 335 samples ]
## sample_data() Sample Data:       [ 335 samples by 4 sample variables ]
## tax_table()   Taxonomy Table:    [ 40 taxa by 7 taxonomic ranks ]
##
## $`20`
## phyloseq-class experiment-level object
## otu_table()   OTU Table:         [ 26 taxa and 335 samples ]
## sample_data() Sample Data:       [ 335 samples by 4 sample variables ]
## tax_table()   Taxonomy Table:    [ 26 taxa by 7 taxonomic ranks ]
##
## $`25`
## phyloseq-class experiment-level object
## otu_table()   OTU Table:         [ 22 taxa and 335 samples ]
## sample_data() Sample Data:       [ 335 samples by 4 sample variables ]
## tax_table()   Taxonomy Table:    [ 22 taxa by 7 taxonomic ranks ]
##
## $`30`
## phyloseq-class experiment-level object
## otu_table()   OTU Table:         [ 17 taxa and 335 samples ]
## sample_data() Sample Data:       [ 335 samples by 4 sample variables ]
## tax_table()   Taxonomy Table:    [ 17 taxa by 7 taxonomic ranks ]
```

#### Calculate the best prevalence interval for the dataset

This function will return a table with Out of Bag error from random forests for each prevalence interval. The number of taxa and the number of remaining sequences for each prevalence interval are also computed. The best prevalence interval value provides the clearest separation of communities while still including a majority of the taxa in the analysis. If true differences are present. It will be represented by the first interval in which the OOB error rate is zero or close to zero. The input file is the list of prevalences generated by the pime.prevalence (prevalences) and the predictor variable ("Probiotic").

```
set.seed(2125)
best.prev=pime.best.prevalence(prevalences, "Probiotic")
```

```
##          Interval OOB error rate (%) OTUs  Nseqs
##    Prevalence 5%               28.06  123 988309
##   Prevalence 10%               14.93   68 967461
##   Prevalence 15%                20.9   40 954379
##   Prevalence 20%               21.19   26 931177
##   Prevalence 25%               27.46   22 929563
##   Prevalence 30%               25.67   17 903330
##   Prevalence 35%               33.73   14 899992
##   Prevalence 40%               17.91   14 898656
##   Prevalence 45%                  20   10 889244
##   Prevalence 50%               42.69    7 881104
##   Prevalence 55%               14.63    7 853874
##   Prevalence 60%               17.01    6 853217
##   Prevalence 65%               18.21    5 797394
##   Prevalence 70%               53.73    3 745832
##   Prevalence 75%               47.16    2 663347
##   Prevalence 80%               46.57    2 663347
##   Prevalence 85%               46.87    2 663347
##   Prevalence 90%               47.16    2 663347
```

The best prevalence interval is that one with OOB error rate close to zero. Here no prevalence interval was close to zero. That means PIME was unable to fit a reasonable model for the dataset. In other words, PIME did not find differences between treatments confirming the previews results from Avershina et al., (2017) demonstrating our approach is not prone to type I errors and do not generate false positive differences.

For this dataset, the OOB error obtained for each prevalence interval indicate that the calculations of ASV importance will not generate accurate results.

#### Likelihood of introducing bias while building prevalence-filtered datasets

```
randomized=pime.error.prediction(inputR, "Probiotic", bootstrap = 100, parallel = TRUE, max.prev = 95)
randomized$Plot
```

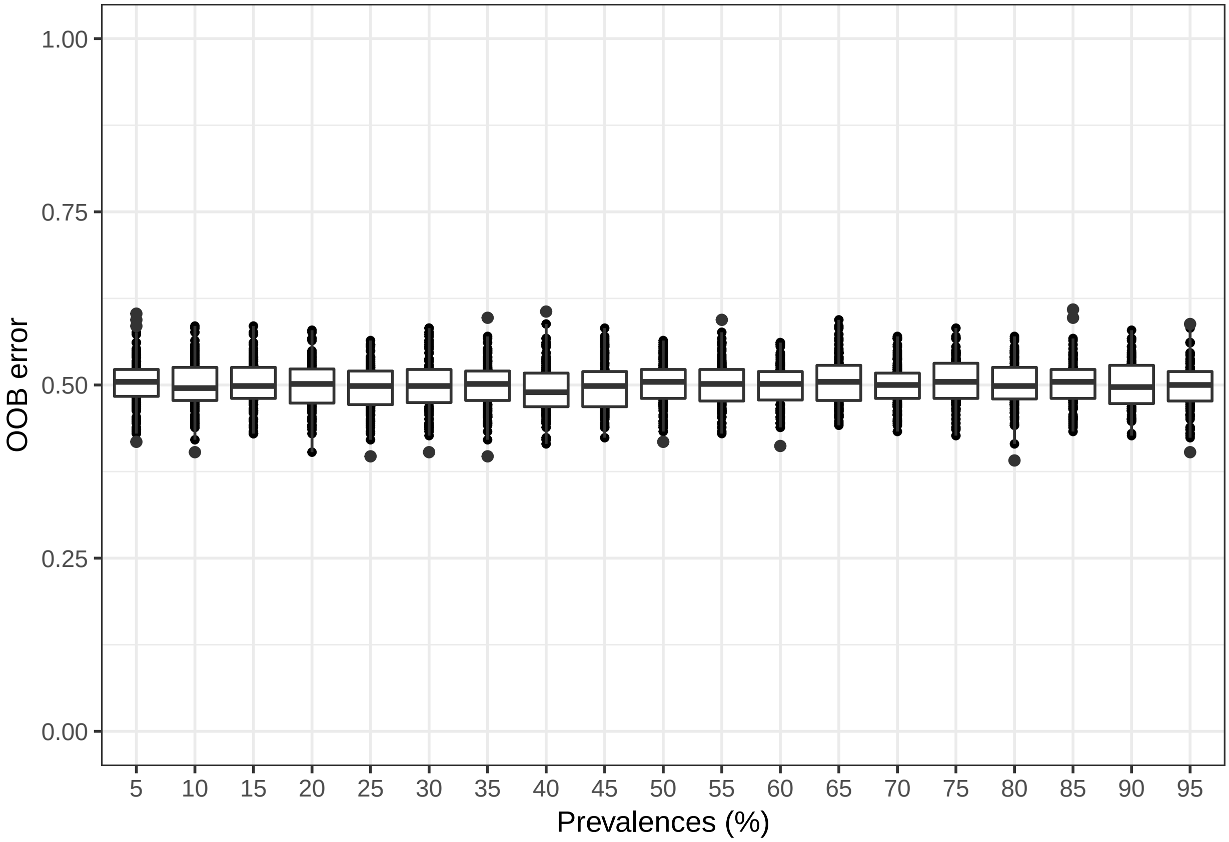

The first function randomizes the samples labels into arbitrary groupings using 100 random permutations. For each randomized prevalence filtered dataset, the OOB error rate is calculated to determine whether differences in the original groups occur by chance.

```
replicated.oob.error= pime.oob.replicate(prevalences, "Probiotic", bootstrap = 100, parallel = TRUE)
replicated.oob.error$Plot
```

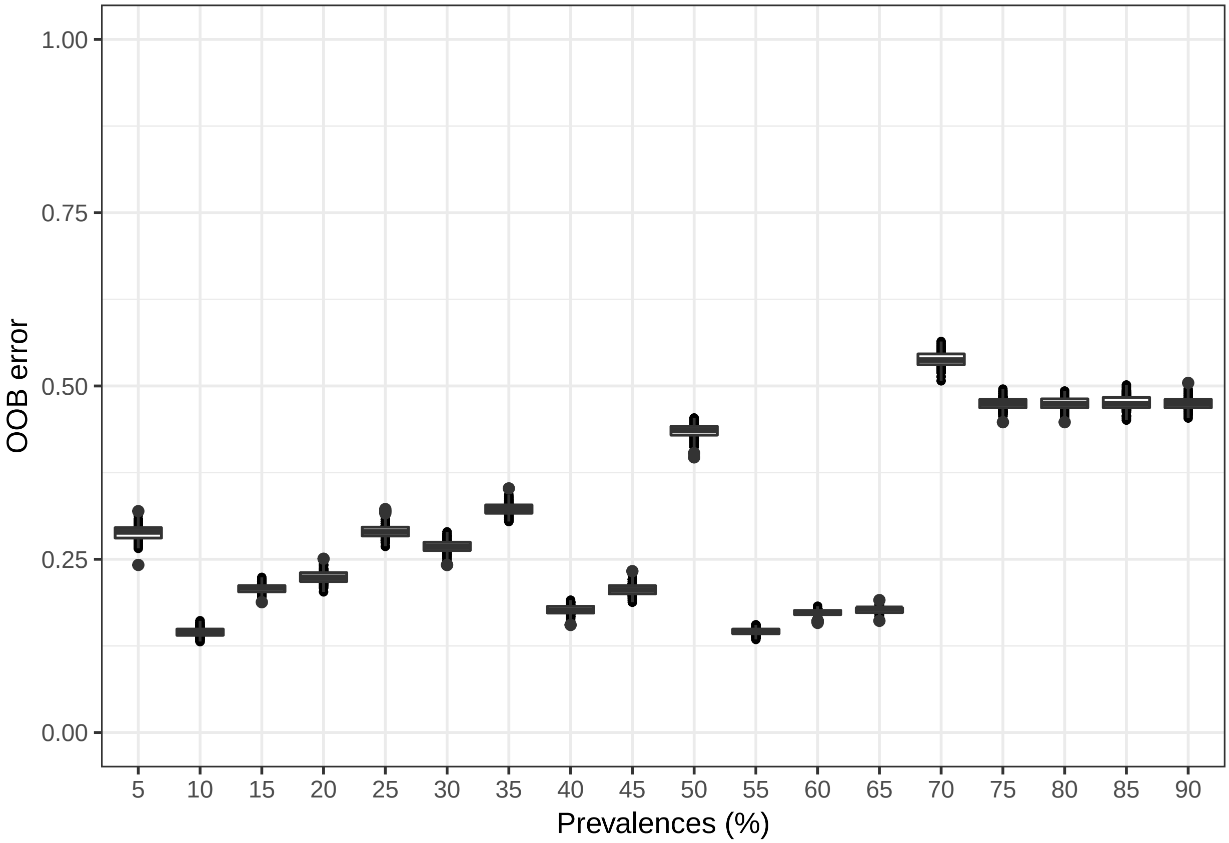

The second function performs the Random Forest analyses and computes the OOB error for 100 replications in each prevalence interval without randomizing the sample labels. The biological difference among samples is expected to be greater than the differences generated randomly. Thus, the greatest fraction of randomizations should generate high error rates. On the other hand, no improvement in accuracy is expected within the randomized dataset.

### Pime Validation 4th dataset

*Roesch et al.*

*4/11/2019*

**Likelihood of introducing bias while building prevalence-filtered datasets** **5**

### Running a dataset from the Human Microbiome Project - association between saliva microbiome and the left antecubital fossa

#### Load files and make a phyloseq object

```
library(phyloseq)
library(ggplot2)
library(microbiome)
library(vegan)
library(knitr)
library(ggpubr)
library(pime)
jsonbiomfile = "/Users/luizroesch/Desktop/Saliva/seqs/dada2/Core/HMP/otu_table_fix.biom"
mapfile = "/Users/luizroesch/Desktop/Saliva/seqs/dada2/Core/HMP/v35_map_uniquebyPSN.txt"
biom = import_biom(jsonbiomfile, mapfile, parseFunction=parse_taxonomy_default)
map = import_qiime_sample_data(mapfile)
input = merge_phyloseq(biom,map)
rank_names(input)
```

```
## [1] "Rank1" "Rank2" "Rank3" "Rank4" "Rank5" "Rank6"
```

```
colnames(tax_table(input)) = c( "Kingdom", "Phylum", "Class", "Order", "Family", "Genus")
filtered = filter_taxa(input, function(x) sum(x) >1, TRUE)
HMP = filtered %>% subset_samples(HMPbodysubsite=="Saliva" | HMPbodysubsite=="Left_Antecubital_fossa")
HMP= prune_taxa(taxa_sums(HMP) > 0, HMP)
set.seed(2125)
```

```r
inputR = rarefy_even_depth(HMP, sample.size = 2000)
inputR
```

```
## phyloseq-class experiment-level object
## otu_table()   OTU Table:         [ 17153 taxa and 172 samples ]
## sample_data() Sample Data:       [ 172 samples by 9 sample variables ]
## tax_table()   Taxonomy Table:    [ 17153 taxa by 6 taxonomic ranks ]
```

#### Finding association between saliva microbiome and the left antecubital fossa without PIME filtering

##### Randon Forest approach

```r
library(randomForest)
filtered = filter_taxa(inputR, function(x) sum(x > 1) > (0.1*length(x)), TRUE)
train= t(otu_table(filtered))
# Make one column for our outcome/response variable
HMPbodysubsite <- as.factor(sample_data(filtered)$HMPbodysubsite)
# Combine them into 1 data frame
training.set <- data.frame(HMPbodysubsite, train)
train.model = randomForest(HMPbodysubsite ~ ., data = training.set, importance = TRUE)
print(train.model)
```

```
##
## Call:
##  randomForest(formula = HMPbodysubsite ~ ., data = training.set,      importance = TRUE)
##                Type of random forest: classification
##                      Number of trees: 500
## No. of variables tried at each split: 22
##
##         OOB estimate of  error rate: 0.58%
## Confusion matrix:
##                       Left_Antecubital_fossa Saliva class.error
## Left_Antecubital_fossa                     59      0 0.000000000
## Saliva                                      1    112 0.008849558
```

The OOB error rate is 0.58% This estimate of error suggests that when the resulting model is applied to new observations, the answers will be in error 0.58% of the time. That is, it is 99.42% accurate, which is a very good model.

We can calculate the variable importance even without running PIME because no filtering step will icrease the accuracy of the prediction (which is highly accurate already). The table lists each ASV and then measures of importance for each ASV. Higher values indicate that the variable is relatively more important.

```r
importance = importance(train.model)
ftable= cbind.data.frame(importance, (tax_table(filtered)))
rownames(ftable) <- NULL
library(knitr)
my_kable = function(x, max.rows=30, ...) {
  kable(x[1:max.rows, ], ...)
}
my_kable(ftable[ , c(1:3, 10)], caption = "Importance of ASVs to differentiate saliva microbiome and th
```

```
- dataset not filtered by PIME",
full_width = F)
```

Table 1: Importance of ASVs to differentiate saliva microbiome and the left antecubital fossa - dataset not filtered by PIME

| Left_Antecubital_fossa | Saliva | MeanDecreaseAccuracy | Genus |
|---:|---:|---:|:---|
| 0.000000 | 1.0010015 | 1.0010015 | g___Veillonella |
| 1.389425 | 0.0000000 | 1.3777712 | NA |
| 0.000000 | 0.0000000 | 0.0000000 | g___Peptostreptococcus |
| 0.000000 | 0.0000000 | 0.0000000 | g___ |
| 0.000000 | 0.4093271 | 0.4877615 | g___ |
| 0.000000 | 0.0000000 | 0.0000000 | g___Catonella |
| 0.000000 | -1.0010015 | -1.0010015 | g___Bulleidia |
| 1.001002 | 0.0000000 | 1.0010015 | g___Fusobacterium |
| 2.090924 | 0.8153591 | 2.0605531 | g___Veillonella |
| 0.000000 | 0.0000000 | 0.0000000 | g___Haemophilus |
| 0.000000 | 0.0000000 | 0.0000000 | g___Streptococcus |
| 2.126639 | 1.4170505 | 2.0472485 | g___ |
| 0.000000 | 0.0000000 | 0.0000000 | g___Prevotella |
| 1.001002 | 0.0000000 | 1.0010015 | g___Selenomonas |
| 0.000000 | 0.0000000 | 0.0000000 | g___Veillonella |
| 0.000000 | 0.0000000 | 0.0000000 | g___ |
| 0.000000 | -1.0010015 | -1.0010015 | g___Prevotella |
| 0.000000 | 0.0000000 | 0.0000000 | g___Veillonella |
| 1.001002 | 0.0000000 | 1.0010015 | g___Prevotella |
| 1.784745 | 0.0000000 | 1.7861912 | g___Oribacterium |
| 1.636275 | 1.0010015 | 1.5415402 | g___Streptococcus |
| 0.000000 | 0.0000000 | 0.0000000 | g___Porphyromonas |
| 1.001002 | 0.0000000 | 1.0010015 | g___Streptococcus |
| 1.463971 | 1.4126375 | 1.4670551 | g___Mogibacterium |
| 0.000000 | 0.0000000 | 0.0000000 | g___Veillonella |
| 0.000000 | 0.0000000 | 0.0000000 | g___ |
| 1.001002 | 0.0000000 | 1.0010015 | g___Neisseria |
| 1.001002 | -1.0010015 | 0.0000000 | g___Prevotella |
| 0.000000 | 0.0000000 | 0.0000000 | g___ |
| 0.000000 | 0.0000000 | 0.0000000 | NA |

### Running PIME

#### Prediction using random forests on full dataset

```
OOB_error_full=pime.oob.error(filtered, "HMPbodysubsite")
```

```
## [1] "OOB error rate is zero. Your data presents large differences.\n                    Prevalen
```

```
OOB_error_full
```

```
## [1] "OOB error rate is zero. Your data presents large differences.\n                    Prevalen
```

The OOB error rate zero, indicated the dataset present large differences with little or no noise to be removed by PIME. Running PIME will not improve the accuracy of the model. We will run PIME but no decrease into the OOB error is expected.

#### Split the dataset by predictor variable

```
per_variable_obj= pime.split.by.variable(filtered, "HMPbodysubsite")
per_variable_obj
```

```
## $Left_Antecubital_fossa
## phyloseq-class experiment-level object
## otu_table()   OTU Table:         [ 383 taxa and 59 samples ]
## sample_data() Sample Data:       [ 59 samples by 9 sample variables ]
## tax_table()   Taxonomy Table:    [ 383 taxa by 6 taxonomic ranks ]
##
## $Saliva
## phyloseq-class experiment-level object
## otu_table()   OTU Table:         [ 475 taxa and 113 samples ]
## sample_data() Sample Data:       [ 113 samples by 9 sample variables ]
## tax_table()   Taxonomy Table:    [ 475 taxa by 6 taxonomic ranks ]
```

#### Calculate the highest possible prevalence intervals

This function calculates prevalence for different intervals by increments of 5. The input file is the output from the pime.split.by.variable(per_variable_obj)

```
prevalences=pime.prevalence(per_variable_obj)
head(prevalences)
```

```
## $`5`
## phyloseq-class experiment-level object
## otu_table()   OTU Table:         [ 509 taxa and 172 samples ]
## sample_data() Sample Data:       [ 172 samples by 9 sample variables ]
## tax_table()   Taxonomy Table:    [ 509 taxa by 6 taxonomic ranks ]
##
## $`10`
## phyloseq-class experiment-level object
## otu_table()   OTU Table:         [ 509 taxa and 172 samples ]
## sample_data() Sample Data:       [ 172 samples by 9 sample variables ]
## tax_table()   Taxonomy Table:    [ 509 taxa by 6 taxonomic ranks ]
##
## $`15`
## phyloseq-class experiment-level object
## otu_table()   OTU Table:         [ 509 taxa and 172 samples ]
## sample_data() Sample Data:       [ 172 samples by 9 sample variables ]
## tax_table()   Taxonomy Table:    [ 509 taxa by 6 taxonomic ranks ]
##
## $`20`
## phyloseq-class experiment-level object
## otu_table()   OTU Table:         [ 506 taxa and 172 samples ]
## sample_data() Sample Data:       [ 172 samples by 9 sample variables ]
## tax_table()   Taxonomy Table:    [ 506 taxa by 6 taxonomic ranks ]
##
## $`25`
## phyloseq-class experiment-level object
```

```
## otu_table()   OTU Table:         [ 473 taxa and 172 samples ]
## sample_data() Sample Data:       [ 172 samples by 9 sample variables ]
## tax_table()   Taxonomy Table:    [ 473 taxa by 6 taxonomic ranks ]
##
## $`30`
## phyloseq-class experiment-level object
## otu_table()   OTU Table:         [ 421 taxa and 172 samples ]
## sample_data() Sample Data:       [ 172 samples by 9 sample variables ]
## tax_table()   Taxonomy Table:    [ 421 taxa by 6 taxonomic ranks ]
```

##### Calculate the best prevalence interval for the dataset

This function will return a table with Out of Bag error from random forests for each prevalence interval. The number of taxa and the number of remaining sequences for each prevalence interval are also computed. The best prevalence interval value provides the clearest separation of communities while still including a majority of the taxa in the analysis. If true differences are present. It will be represented by the first interval in which the OOB error rate is zero or close to zero. The input file is the list of prevalences generated by the pime.prevalence (prevalences) and the predictor variable ("HMPbodysubsite").

```
set.seed(2125)
best.prev=pime.best.prevalence(prevalences, "HMPbodysubsite")
```

```
##        Interval OOB error rate (%) OTUs  Nseqs
##   Prevalence 5%                  0  509 190455
##  Prevalence 10%                  0  509 189296
##  Prevalence 15%                  0  509 188555
##  Prevalence 20%                  0  506 188189
##  Prevalence 25%                  0  473 183574
##  Prevalence 30%                  0  421 176217
##  Prevalence 35%                  0  328 161346
##  Prevalence 40%                  0  274 151525
##  Prevalence 45%                  0  234 142112
##  Prevalence 50%                  0  185 129369
##  Prevalence 55%                  0  143 112978
##  Prevalence 60%                  0  118 104474
##  Prevalence 65%                  0   74  82052
##  Prevalence 70%                  0   48  62817
##  Prevalence 75%                  0   34  56457
##  Prevalence 80%                  0   23  46475
##  Prevalence 85%                  0   13  34459
```

The best prevalence interval is that one with OOB error rate close to zero. Here all prevalence intervals were zero. That means PIME did not find greater differences in the dataset.

#### Likelihood of introducing bias while building prevalence-filtered datasets

```
randomized=pime.error.prediction(filtered, "HMPbodysubsite", bootstrap = 100, parallel = TRUE, max.prev
randomized$Plot
```

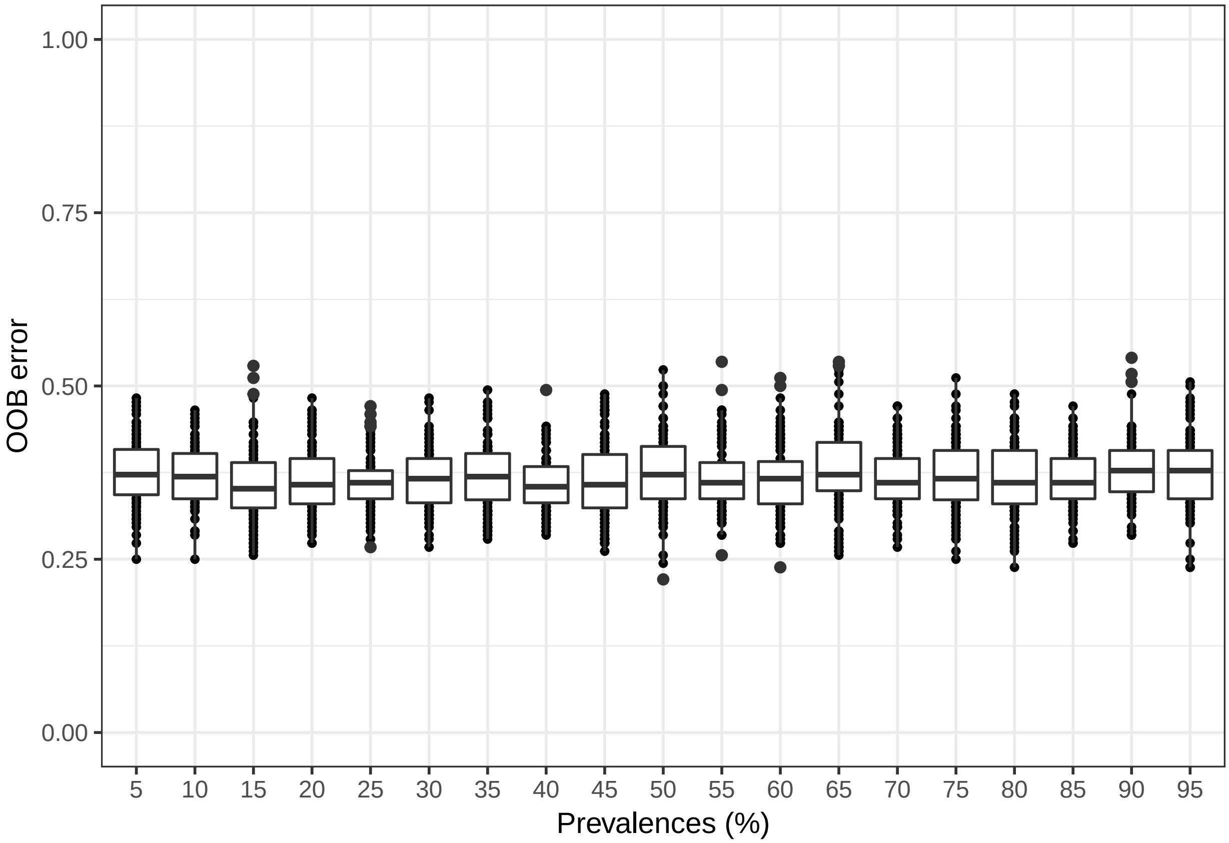

The first function randomizes the samples labels into arbitrary groupings using 100 random permutations. For each randomized prevalence filtered dataset, the OOB error rate is calculated to determine whether differences in the original groups occur by chance.

```
replicated.oob.error= pime.oob.replicate(prevalences, "HMPbodysubsite", bootstrap = 100, parallel = TRU[illegible]
replicated.oob.error$Plot
```

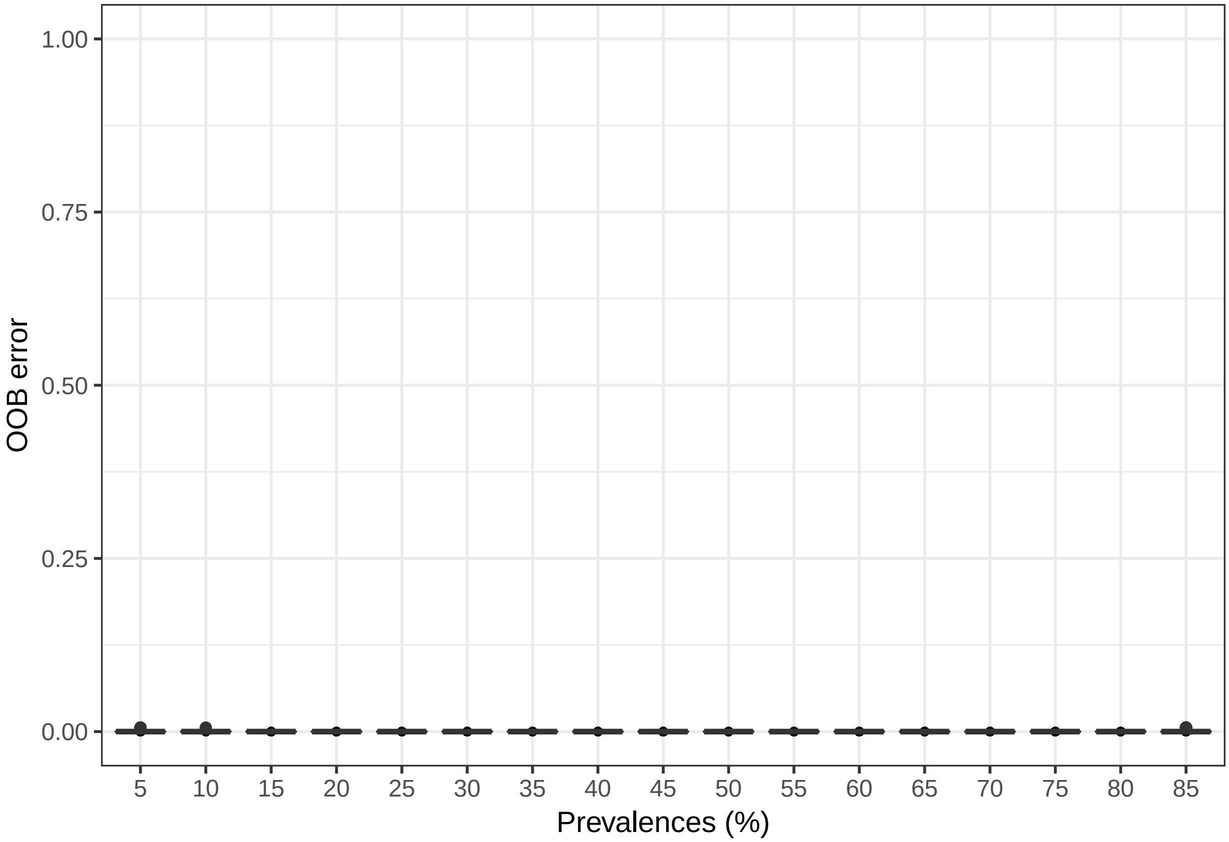

The second function performs the Random Forest analyses and computes the OOB error for 100 replications in each prevalence interval without randomizing the sample labels. The biological difference among samples is expected to be greater than the differences generated randomly. Thus, the greatest fraction of randomizations should generate high error rates. On the other hand, no improvement in accuracy is expected within the randomized dataset.
